# HOXB8 counteracts MAPK/ERK oncogenic signaling in a chicken embryo model of neoplasia

**DOI:** 10.1101/2021.06.11.448019

**Authors:** Axelle Wilmerding, Lauranne Bouteille, Lucrezia Rinaldi, Nathalie Caruso, Yacine Graba, Marie-Claire Delfini

## Abstract

HOX transcription factors are members of an evolutionarily conserved family of proteins required for the establishment of the anteroposterior body axis during bilaterian development. Although they are often deregulated in cancers, the molecular mechanisms by which they act as oncogenes or tumor suppressor genes are only partially understood. Since the MAPK/ERK signaling pathway is deregulated in most cancers, we aimed at apprehending if and how Hox proteins interact with ERK oncogenicity.

Using an *in vivo* neoplasia model in the chicken embryo that we have developed, consisting in the overactivation of the ERK1/2 kinases in the trunk neural tube, we analyzed the consequences of HOXB8 gain of function at morphological and transcriptional level in this model. We found that HOXB8 acts as a tumor suppressor, counteracting ERK-induced neoplasia. HOXB8 tumor suppressor function in this model relies on a large reversion of the oncogenic transcriptome induced by ERK. In addition to showing that HOXB8 protein controls the transcriptional responsiveness to ERK oncogenic signaling, our study identified new downstream targets of ERK oncogenic activation in an *in vivo* context that could provide clues for therapeutic strategies.

## INTRODUCTION

The HOX family is an evolutionary conserved set of proteins that regulate developmental processes such as anteroposterior body axis patterning, organ morphogenesis and cell fate. As transcription factors, their mainly act by target gene transcriptional activation or repression (Krumlauf, 2018; Rezsohazy et al., 2015). There are 39 HOX genes in vertebrates, arranged in a contiguous manner in 4 groups of 9 to 11 genes, A, B, C and D, located in Human on 4 different chromosomes, at 7p15, 17p21, 12q13, and 2q31, respectively (Grier et al., 2005). HOX genes are numbered 1 through 13, such that each HOX has up to 4 paralogs on each of the 4 chromosomal loci. Their role in embryonic development during which their spatial and temporal expression is tightly governed is well established (Rezsohazy et al., 2015). HOX genes remain expressed in adult tissues and participate, among other, in tissue-specific stem cell differentiation contributing to the maintenance and function of various organs and tissues (Seifert, 2015).

Aberrant expression of HOX genes is associated to many Human pathologies including cancers (Bhatlekar et al., 2018; Shah, 2010). The roles of HOX transcription factors as oncogenes or tumor suppressor genes, depending on the cellular context, are well established, and implies modulation of angiogenesis, differentiation, apoptosis, proliferation, epithelial to mesenchymal transition and DNA repair (Grier et al., 2005). Highlighting the precise context-dependent and versatile role of HOX proteins, the same HOX protein can act either as a tumor suppressor or oncogene (Jonkers et al., 2020). For example, HOXA9 drives leukemia (Collins and Hess, 2016), but suppresses metastasis and growth in breast cancer (Sun et al., 2013). Similarly, HOXB8 acts as an oncogene in several cancers including colorectal cancer (Ying et al., 2020), but is a favorable prognostic marker in renal cancer (The Human Protein Atlas) (Uhlen et al., 2017). HOXB8 was also recently shown to positively controls the transcriptional expression of the tumor suppressor *LZTS1* (Wilmerding et al., 2021a). A better understanding of the molecular mechanisms that promote either the oncogenic or tumor suppressor activity of HOX proteins remains unsolved.

The mitogen-activated protein kinase (MAPK) cascades are critical pathways for human cancer cell survival, dissemination, and resistance to drug therapy (Braicu et al., 2019; De Luca et al., 2012). In particular, the MAPK/extracellular signal-regulated kinase (ERK1/2) pathway is a signaling node that receives input from numerous stimuli, including internal metabolic stress and DNA damage pathways, as well as through signaling from external growth factors, cell-matrix interactions, and communication from other cells (Burotto et al., 2014). Through sequential activation of Ras-like GTPase, RAF, MEK, and ERK1/2, the ERK1/2 pathway transduces the signal of growth factors across the cell membrane to control cell cycle progression, proliferation, survival and differentiation. This pathway is finely regulated by feedback loops and nearly strictly converges towards the phosphorylation and activation of ERK1 and ERK2 kinases, which interact with many cytosolic and nuclear substrates (> 300 described) (Samatar and Poulikakos, 2014; Yang et al., 2019a). That overactivation of the RTK/RAS/RAF/MEK/ERK signal is a key oncogenic event is now known for more than two decades (Hilger et al., 2002) and has been recently confirmed by pan-cancer genomic analyses which have indeed shown that it is the signaling pathway with the highest median frequency of alterations (46% of samples) across all cancer types (Sanchez-Vega et al., 2018). The clinical and therapeutic success of RAF and MEK1/2 inhibitors has revolutionized the existing treatment schemes for previously incurable cancers. However, the overall therapeutic efficacies are still largely compromised by side effects and emerging drug resistance mechanisms (Samatar and Poulikakos, 2014). It is therefore crucial to increase our understanding of the molecular mechanisms underlying the oncogenic activity of the ERK pathway to develop more effective therapies.

Study on zebrafish embryo suggested that posterior Hox genes control the responsiveness to RTK/FGF signal (which has ERK as cellular effector in this tissue) (Lunn et al., 2007) during early development (Shimizu et al., 2006). In addition, recent studies showed that HOXB7, HOXA3 and HOXC6 act as oncogenes by activating ERK pathway in pancreas, colon and glioblastoma cancers respectively (Tsuboi et al., 2017; Yang et al., 2019b; Zhang et al., 2018). To gain insights in the molecular mechanisms by which HOX proteins control ERK oncogenic pathways, we developed a chicken embryo neoplasia model obtained by ERK1/2 overactivation in the neural tube (Wilmerding et al., 2021b) and investigated the effects of HOXB8 expression in this model. We found that HOXB8 largely counteracts ERK oncogenic activity in this neoplasia model. Transcriptomic analysis showed that HOXB8 tumor suppressor function relies on a large reversion of the oncogenic transcriptome induced by ERK overactivation. HOXB8 effect is independent of ERK phosphorylation and nuclear translocation and does not rely on the transcriptional control of the early response genes of the EGR and ETV. This study also identified new transcriptional downstream targets of ERK controlled by HOXB8 that could provide clues for therapeutic applications. We found that one of the genes most upregulated by ERK overactivation and whose expression is reversed by HOXB8 is the gene encoding CHST15, whose expression is unfavorable in most human cancers, underlining the relevance of this chicken *in vivo* model to identify new targets pertinent in human pathology.

## RESULTS

### HOXB8 transcription factor inhibits neoplasia induced by ERK overactivation in the chicken embryo neural tube

To test the effect of HOXB8 on ERK oncogenic activity in an *in vivo* context, we took advantage of a model of ERK-induced neoplasia we developed in the chicken embryo neural tube (Wilmerding et al., 2021b). The model consists in the expression of a constitutively active form of the kinase MEK1, acting upstream of ERK1/2 (MEK1ca) (Delfini et al., 2005; Mansour et al., 1994). Transfection is obtained by electroporation in the trunk neural tube of 2-days old chicken embryos. While MEK1ca expression (using the pCIG-MEK1ca expressing vector co-expressing nuclear GFP) induces a strong neoplasic phenotype three days post-electroporation ((Wilmerding et al., 2021b), Figure 1A-B), co-electroporation of pCIG-MEK1ca with a HOXB8 expressing vector (also co-expressing GFP) leads to a milder phenotype (Figure 1A-B). Immunofluorescences on transverse sections with a GFP antibody to stain transfected cells (Figure 1B), co-stained with phalloidin (Supplementary Figure 1) allowed to highlight that while the neural tube is still morphologically affected (compare to control/pCIG condition and to contralateral/non-electroporated part of the neural tube), the extent of neoplasia is much weaker in all the embryos analyzed when MEK1ca is co-transfected with the HOXB8 expressing vector (>10 embryos analyzed for each condition). This morphological analysis suggested that in this context, HOXB8 gain of function works as a tumor suppressor.

**Figure 1:**
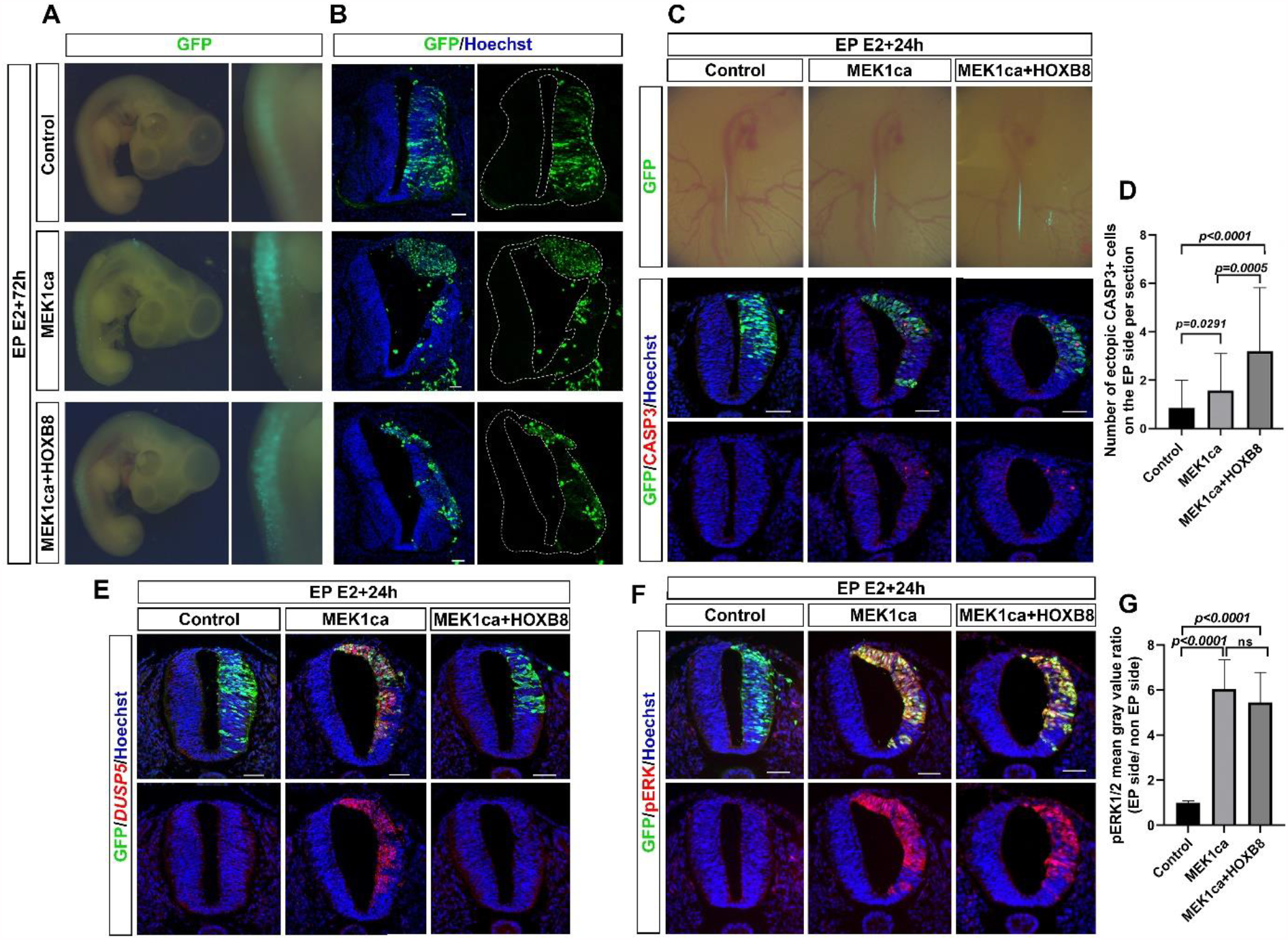
HOXB8 suppresses MEK1ca-induced neoplasia in the trunk neural tube. **A-**Dorsal view of electroporated embryos three days post-electroporation with the control pCIG, MEK1ca, or MEK1ca+HOXB8 expressing vectors. Transfected cells are identified by GFP. **B-**Immunofluorescences on transverse sections of chicken embryo with anti-GFP antibody three days post-electroporation with pCIG, MEK1ca, or MEK1ca+HOXB8 expressing vectors. **C-**(Top) Dorsal view of electroporated embryos one day post-electroporation with the control pCIG, MEK1ca, or MEK1ca+HOXB8 expressing vectors. Transfected cells are identified by GFP. (Bottom) Immunofluorescences on transverse sections of chicken embryo with anti-GFP and anti-cleaved caspase 3 (CASP3) antibodies. **D-**Number of ectopic CASP3+ cells on the electroporated side one day after electroporation in the three conditions (n=3 animals/18 sections). The quantifications show a significant increase of Casp3 cells in neural tube after MEK1ca and MEK1ca+HOXB8 conditions compared to control, with more Casp3 cells in the MEK1ca+HOXB8 condition than in MEK1ca alone (two-tailed Mann–Whitney test, error bars represent s.d.). **E-**Fluorescent *in situ* hybridizations with DUSP5 probe and immunofluorescences with anti-GFP antibody on trunk transverse sections of chicken embryo one day post-electroporation in the three conditions. **F**-Immunofluorescences on transverse sections with anti-pERK1/2 and anti-GFP antibodies one day post-electroporation with PCIG, MEK1ca or MEK1ca+HOXB8 expressing vectors. **G**-pERK1/2 mean gray value ratio (electroporated/contralateral side) one day after electroporation in the three conditions (n=3 animals/18 sections). The quantifications show no significant difference between MEK1ca and MEK1ca+HOXB8 conditions (two-tailed Mann–Whitney test, error bars represent s.d.). Blue is Hoechst staining. Scale bar: 50µm.

We next decided to investigate the molecular mechanisms by which HOXB8 inhibits ERK1/2 overactivation induced neoplasia. HOXB8 is endogenously expressed in the trunk neural tube of chicken embryo from E3, controlling neuronal delamination (Wilmerding et al., 2021a). Since HOXB8 overexpression in the neural tube leads to an increase of apoptosis (Wilmerding et al., 2021a), HOXB8 tumor suppressor effect may be due to cell death induction in MEK1ca expressing cells. We thus performed immunofluorescences on transverse sections of chicken embryos one day after electroporation and quantified the cleaved caspase3 (CASP3) apoptotic marker in the neural tube after transfection of control pCIG, MEK1ca, or MEK1ca+HOXB8. We found increased apoptosis in the MEK1ca condition (as already described in (Wilmerding et al., 2021b) which is further enhanced in MEK1ca+HOXB8 condition (Figure C-D), in agreement with the hypothesis that HOXB8-induced cell death contribute to reduce MEK1ca-induced neoplasia. Since in a developmental context in zebrafish, Hox genes have been shown to control the responsiveness of FGF/ERK signaling (Shimizu et al., 2006), next weprobed if in addition to increase cell death in MEK1ca expressing cells, HOXB8 gain of function might also counteracts MEK1ca-induced oncogenic transcriptome. To test this hypothesis, we performed fluorescent *in situ* hybridizations on transverse sections with probes against target genes of the ERK pathway, DUSP5, GREB1 and IL17RD genes, which transcripts are strongly upregulated after MEK1ca expression in the chicken neural tube (Wilmerding et al., 2021b). We found that when HOXB8 is co-expressed, MEK1ca does not induce the expression of any of these three genes (Figure 1E, Supplementary Figures 2 and 3). We next asked if the suppressing effect of HOXB8 on MEK1ca-induced target genes observed occurs upstream, by modifying ERK1/2 activation, or downstream of ERK1/2 by modifying the transcriptional response to ERK1/2 signaling. We performed immunofluorescences on transverse sections with the phosphoThr202/Tyr204-ERK1/2 (pERK) antibody which reflects ERK1/2 kinases activity. We found that HOXB8 co-expression does not change the level of MEK1ca-induced ERK activation, and that pERK staining was still localized both in the cytoplasm and nucleus, as is the case following MEK1ca induction (Figure 1F-G). Altogether, these results show that HOXB8 functions as a tumor suppressor in this neoplasic context, by increasing cell death and preventing MEK1ca-induced target gene activation, acting downstream of ERK phosphorylation and nuclear translocation.

### HOXB8 largely reverses the ERK-induced oncogenic transcriptome

To identify the global transcriptional changes underlying HOXB8-mediated suppression of MEK1ca-induced neoplasia, we performed RNA-seq on GFP-sorted positive neural tube cells (Figure 2A). E2 neural tubes were bilaterally electroporated with either the control vector pCIG (control condition), pCIG-MEK1ca (Wilmerding et al., 2021b), pCIG-HOXB8 (Wilmerding et al., 2021a), or pCIG-MEK1ca plus pCIG-HOXB8 (Figure 2A). The regions of the neural tube expressing the GFP were dissected 18 hours after electroporation and dissociated (Figure 2A). GFP-expressing cells were isolated by FACS with the use of a dead cell exclusion (DCE)/discrimination dye (DAPI) to eliminate dying cells. Two independent RNAs samples by condition were extracted, reverse transcribed, and cDNAs were amplified using a linear amplification system and used for sequencing library building. Qualitative analysis of RNA-seq data after alignment to the Galgal4 genome assembly (17108 genes) from the 2 biological replicates for the 4 conditions shows a high Pearson correlation score (>0,99) indicative of experimental reproducibility, confirmed by principal component analysis (Supplementary Figure. 4).

**Figure 2:**
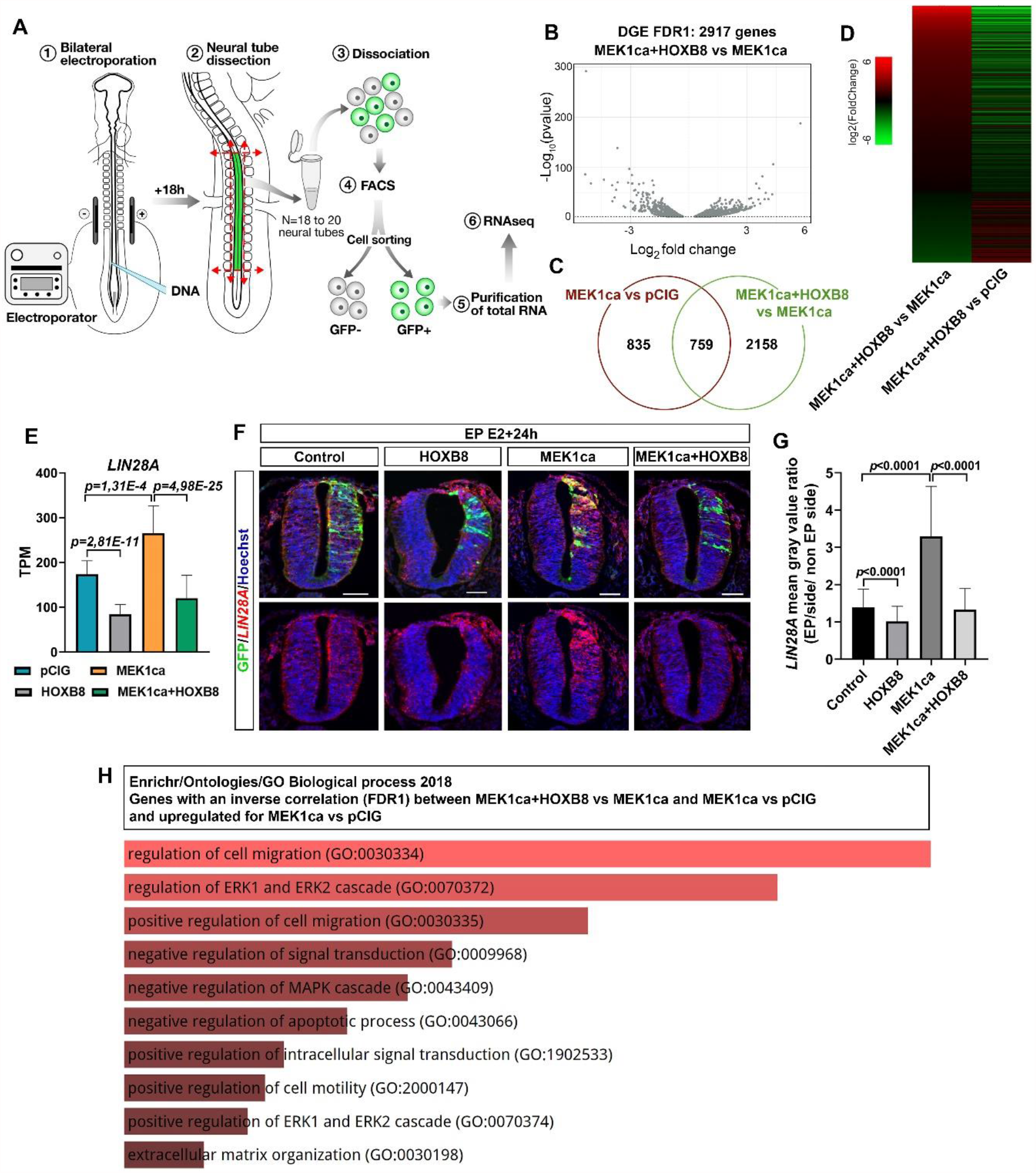
Transcriptomic analysis of MEK1ca-induced neoplasia suppression by HOXB8. **A-**Experimental design. 18 h after bilateral electroporation of trunk neural tube at the stage HH12, the electroporated region of the neural tube was dissected (18-20 embryos per condition in duplicate) and the GFP positive cells were sorted by FACS. **B-**Volcano plot of differential gene expression (DGE, FDR = 1%) for the MEK1ca+HOXB8 versus MEK1ca conditions (2917 genes). Generated using VolaNoseR **C-**Venn diagram showing gene overlaps between MEK1ca versus pCIG and MEK1ca+HOXB8 versus MEK1ca conditions (FDR = 1%) **D-**Heat map showing the expression of the 759 genes upregulated by MEK1ca and reversed by HOXB8 in MEK1ca versus pCIG (line 1), MEK1ca+HOXB8 versus MEK1ca (line 2) and MEK1ca+HOXB8 versus MEK1ca (line 3), highlighting the global reversion of MEK1ca deregulated genes by HOXB8. **E**-Mean expression of LIN28A TPM (transcripts per kilobase million), obtained for the two replicates of the control (pCIG1, pCIG2), HOXB8 (HOXB8-1 and HOXB8-2), MEK1ca (MEK1ca-1 and MEK1ca-2) and MEK1ca+HoxB8 (MEK1ca+HOXB8-1 and MEK1ca+HOXB8-2) expressing samples, with the corresponding p value according to the DGE (FDR= 1%) of each pair. **F-**Fluorescent *in situ* hybridizations with *LIN28A* probe and immunofluorescences with anti-GFP antibody on trunk transverse sections of chicken embryo one day post-electroporation in the pCIG, HOXB8, MEK1ca and MEK1ca+HOXB8 conditions. Blue is Hoechst staining. Scale bar: 50µm. **G-***LIN28A* expression ratio (electroporated/contralateral side) one day after electroporation in the 4 conditions (n=3 animals/6 sections min). The quantifications show that HOXB8 reverses MEK1ca-induced expression of LIN28A (two-tailed Mann–Whitney test, error bars represent s.d.). **H**-Gene ontology enrichment analysis using the EnrichR analytical tool for biological processes (GO biological processes 2018), for the upregulated genes in the MEK1ca versus pCIG condition among the 759 commonly deregulated genes (FDR = 1%) (between “MEK1ca+HOXB8 versus MEK1ca” and “MEK1ca versus pCIG”), with inverse correlation (518 genes).

Comparison of the differential gene expression (DGE) for a FDR1 (False Discovery Rate = 1%) between “MEK1ca+HOXB8 versus MEK1ca” (Figure 2B, Supplementary Table 1) and “MEK1ca versus pCIG” (Supplementary Table 2 for FDR1, and FDR5 is available in Wilmerding et al., 2021b) identifies 759 genes commonly deregulated (Figure 2C). Surprisingly, more than 90% (689 genes) of the commonly deregulated genes are inversely correlated, as illustrated by the expression heatmap (Figure 2D). In other words, among the commonly deregulated genes, most of the genes deregulated in the MEK1ca condition display expression levels nearly normal when HOXB8 is co-transfected (Figure 2D). Among these genes are DUSP5, IL17RD and GREB1 for which the *in situ* hybridization experiments showed a reversion of the MEK1ca-induced phenotype in presence of HOXB8 (Figure 1E and Supplementary Figures 2 and 3). The RNA-seq data thus identifies a HOXB8-mediated reversion of the MEK1ca-induced transcriptome. Of note, 19 of the 20 genes that are most upregulated by MEK1ca, display reduced expression (for some of them nearly completely) when HOXB8 is co-expressed (Supplementary Figure 5).

Using *in situ* hybridization on tissue sections, we confirmed the RNA-seq data, including genes for which MEK1ca has a modest effect, as illustrated for *LIN28A* (Figure 2E, F): *LIN28A* expression is upregulated by MEK1ca (MEK1ca versus pCIG) (Wilmerding et al., 2021b), and downregulated by HOXB8 alone (HOXB8 versus pCIG) (Wilmerding et al., 2021a) or by the combination of both (MEK1ca+HOXB8 versus pCIG) (Figure 2E). Chicken *LIN28A* is expressed in E15/E16 embryos (Yokoyama et al., 2008), the stage at which the RNA-seq was performed. We found that in the neural tube, *LIN28A* is expressed in a gradient along the rostro-caudal axis, consistent with a regulation by the FGF/ERK signal produced as a gradient from the most caudal part of the embryo (Supplementary Figure 6) (Wilmerding et al., 2021b). This posterior gradient is more pronounced at E2, the stage at which we performed electroporations. *In situ* hybridization on transverse trunk sections showed that MEK1ca upregulates *LIN28A*, while HOXB8 downregulates *LIN28A*, including when co-expressed with MEK1ca (Figure 2F-G).

*LIN28A* encodes an RNA binding protein highly conserved that represses microRNAs including *let-7* and influences mRNA translation, thus regulating the self-renewal of embryonic stem cells (Shyh-Chang and Daley, 2013). *LIN28A* is also important for body growth and metabolism, tissue development and somatic reprogramming. Furthermore, its role as oncogenes has been widely demonstrated (Balzeau et al., 2017). High levels of LIN28A (or LIN28B, the second Lin protein) are associated with many human cancers such as glioblastoma, ovarian, gastric, prostate and breast cancer, as well as in pediatric cancers (Carmel-Gross et al., 2016). In mice the ectopic expression of Lin28a is sufficient to induce and/or accelerate tumorigenesis by a let-7-dependent mechanism (Balzeau et al., 2017). In addition to confirm the RNA-seq data, the study of LIN28A regulation by MEKIca and HOXB8 together with LIN28A oncogenic activity suggests that HOXB8 reversion of MEK1ca neoplasia at least partly relies on transcriptional downregulation of LIN28A.

In order to identify other mechanisms that might account for HOXB8 tumor suppressor function in the MEK1ca context, we performed a gene ontology enrichment analysis using the EnrichR” analytical tool (Chen et al., 2013; Kuleshov et al., 2016; Xie et al., 2021) for biological processes (GO biological processes 2018), for the upregulated genes in the MEK1ca versus pCIG condition among the 759 commonly deregulated genes (FDR = 1%) (between “MEK1ca+HOXB8 versus MEK1ca” and “MEK1ca versus pCIG”), with inverse correlation (518 genes) (Figure 2H, Supplementary Table 3). The hits are coherent with the tumor suppressor role of HOXB8 in this context. Indeed, genes of this list have as biological processes regulation of cell migration and cell motility, regulation of ERK1/2 (intracellular signal transduction, negative regulation of apoptotic process and extracellular matrix organization (Figure 2H). In conclusion, the RNAseq results, validated by in situ hybridization, are consistent with gene ontology analysis with the phenotypic tumor suppressor effect of HOXB8 gain of function in the MEK1ca-neoplasia induced context observed morphologically.

### Clustering analysis of the RNA-seq suggests mechanisms of HOXB8-mediated reversion of MEK1ca-induced neoplasia

To further the analysis of the RNA-seq to identify molecular mechanisms implicated in HOXB8 tumor suppressor function, and to identify new oncogenes and tumor suppressors, we performed a clustering analysis of the RNA-seq data starting with the four experimental conditions and using “kmeans” on the list of the 2316 genes deregulated upon MEK1ca expression (MEK1ca versus pCIG) (FDR = 5%) (Wilmerding et al., 2021b) (Figure 3, Supplementary Figure 7, Supplementary Table 4 and 5). Twelve is the number of clusters that we found best describes the dynamics of gene expression in the different conditions, although some of the clusters are similar to each other.

**Figure 3:**
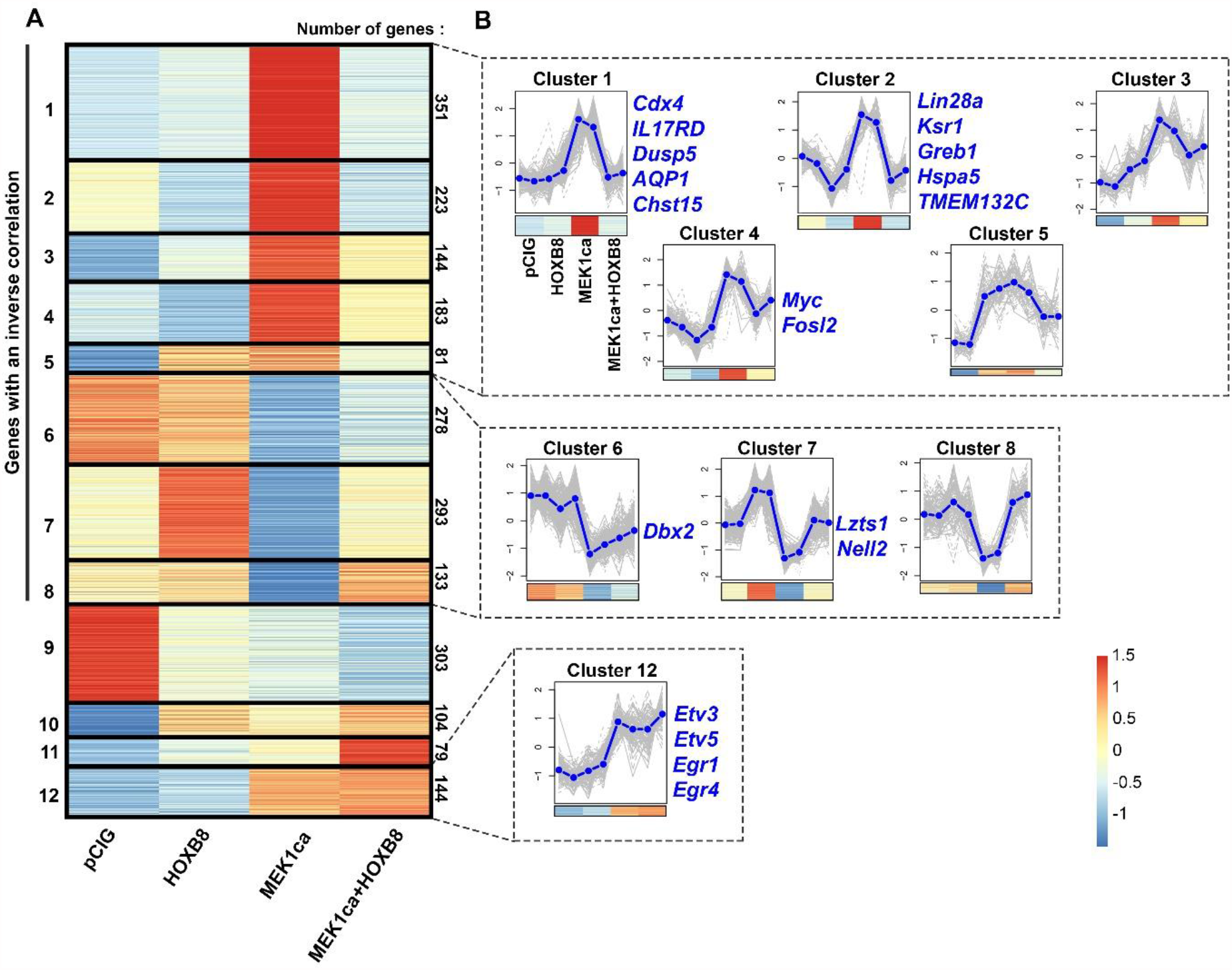
Clustering of genes deregulated by MEK1ca. **A-**Heat map of the 2316 genes deregulated in the “MEK1ca versus PCIG” condition (FDR= 5%) clustered using “kmeans” with k=12, for pCIG, HOXB8, MEK1ca, and MEK1ca+HOXB8 conditions. 1686 genes (72,8%, clusters 1 to 8) display an inverse correlation between MEK1ca versus pCIG and MEK1ca+HOXB8 versus MEK1ca conditions. Only 144 genes (6.2%, cluster 12) behave the same in MEK1ca and MEK1ca+HOXB8 condition (i.e., are not reversed by HOXB8). **B-**Plots of clusters 1 to 8 and 12 with the two replicates of each condition (control (pCIG1, pCIG2), HOXB8 (HOXB8-1 and HOXB8-2), MEK1ca (MEK1ca-1 and MEK1ca-2) and MEK1ca+HOXB8 (MEK1ca+HOXB8-1 and MEK1ca+HOXB8-2), with the mean (in blue), and with the corresponding heat map on the bottom. Some representative genes of clusters are on the right side of the corresponding cluster.

42,3% (980/2316) of the genes are in clusters where expression is increased by MEK1ca and reversed by HOXB8 (clusters 1-5) (Figure 3A-B). Among them, cluster 1 (351 genes) regroups genes that are not regulated by HOXB8 alone (Figure 3B). These genes are (1) genes which play a physiological role in maintaining cells in an immature state downstream of the FGF/ERK pathway, (including CDX4), (2) genes controlling the FGF/ERK signaling pathway (including DUSP5 and IL17RD), and (3) genes which are endogenously not expressed in the embryo at that stage and thus do not control rostro-caudal maturation of the embryo in physiological conditions (including IL1R1 and AQP1) (Figure 3B and Supplementary Figure 8) (Wilmerding et al., 2021b). Among the clusters where gene expression is increased by MEK1ca and reversed by HOXB8 in the MEK1ca context (MEK1ca+HOXB8), are genes of clusters 2 and 4 (Figure 3B) that behave like LIN28A, (i.e., which are downregulated by HOXB8 alone). These genes might act as oncogenes and because of their negative regulation by HOXB8, they may contribute to reverse the MEK1ca-induced oncogenic phenotype. Among these genes is as example KSR1 gene (Cluster 2, Figure 3B, Supplementary Figure 8). It encodes a molecular scaffold Kinase Suppressor of RAS which plays potent roles in promoting RAS-mediated signaling through the RAF/MEK/ERK kinases cascade (Neilsen et al., 2017). Another gene of this group that might be interesting to consider is TMEM132C (Cluster 2, Figure 3B). Indeed, the molecular functions of TMEM132 genes remain poorly understood and under-investigated despite their mutations associated with cancer (Sanchez-Pulido and Ponting, 2018). Structural analysis of this gene predicts a cellular adhesion function, connecting the extracellular medium with the intracellular ACTIN cytoskeleton (Sanchez-Pulido and Ponting, 2018). For this gene, we described for the first time its expression in vertebrate embryo (Figure 4A) and confirmed the bioinformatics data (Figure 4B) by *in situ* hybridization on tissue section (Figure 4C). Its expression during the development of the spinal cord suggests that it might play physiological function in the differentiation of the motor neurons and in the fate and/or migration of the neural crest cells of the trunk (Figure 4A). Outside of the neural tube, it is also expressed in the dorsal root ganglia and in the myotome, where it might also play key functions (Figure 4A). *In situ* hybridization on tissue section (Figure 4C) confirms the dynamics of *TMEM132C* expression identified from the RNA-seq data (compare Figure 4C and 4B). In conclusion, the list of the genes upregulated by MEK1ca and reversed by HOXB8 (cluster 1 to 5) might thus allow to identify new putative oncogenes.

**Figure 4:**
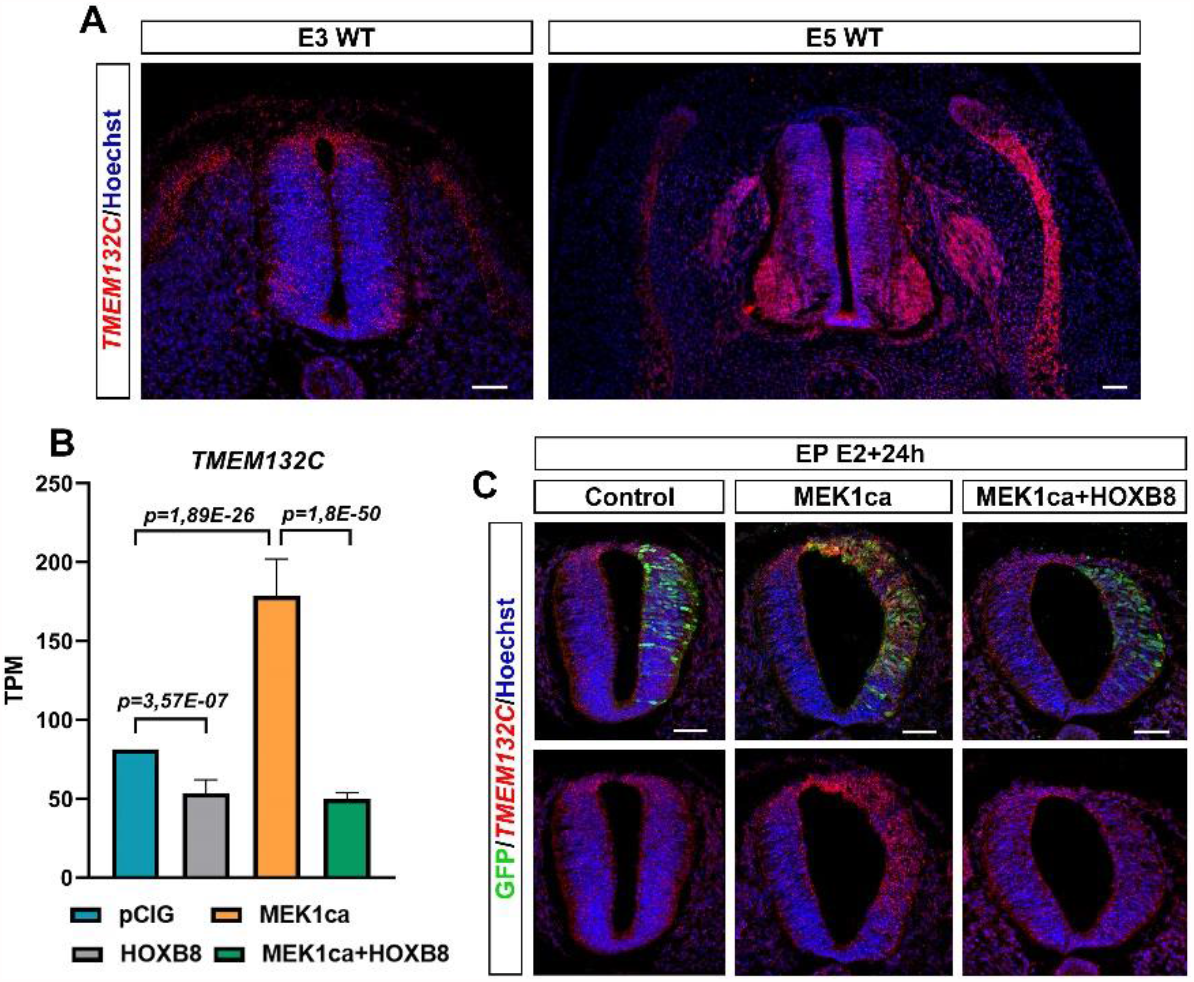
TMEM132C regulation by MEK1ca and HOXB8. **A-**Fluorescent *in situ* hybridizations with TMEM132C probe on trunk transverse sections of wild type chicken embryo at E3 and E4. **B**-Mean expression of *TMEM132C* in TPM (transcripts per kilobase million), obtained for the two replicates of the control (pCIG1, pCIG2), HOXB8 (HOXB8-1 and HOXB8-2), MEK1ca (MEK1ca-1 and MEK1ca-2) and MEK1ca+HOXB8 (MEK1ca+HOXB8-1 and MEK1ca+HOXB8-2) expressing samples, with the corresponding p-value according to the DGE of each pair. **C-**Fluorescent *in situ* hybridizations with TMEM132C probe and immunofluorescences with anti-GFP antibody on trunk transverse sections of chicken embryo one day post-electroporation in the pCIG, HOXB8, MEK1ca and MEK1ca+HOXB8 conditions.

30,4% (704/2316) of genes (clusters 6-8) behave oppositely to genes of clusters 1-5: their expression are downregulated by MEK1ca and reversed by HOXB8. These include genes not regulated alone by HOXB8 alone (cluster 6, Figure 3B), such as DBX2 (Supplementary Figure 8), involved in primary neurogenesis (Ma et al., 2011). They also include genes regulated by HOXB8 alone (clusters 7-8, Figure 3B), such as *LZTS1* (cluster 7, Figure 3B, Supplementary Figure 8), a Leucine zipper tumor suppressor that suppresses colorectal cancer proliferation through inhibition of the AKT pathway (Zhou et al., 2015) and that we recently found to control neuronal delamination in the trunk neural tube of chicken embryo (Wilmerding et al., 2021a). *LZTS1* is transcriptionally activated by HOXB8 and might as other genes from the same cluster provide additional mechanisms by which HOXB8 reverse MEK1ca-induced neoplasia. *NELL2* (Neural EGFL Like 2) for example, expressed in the neural tube and involved in neural development (Nelson et al., 2004), has also been described as a tumor suppressor since enriched in normal nerve cells compared with nervous system tumors (Maeda et al., 2001) and inhibiting cancer cell migration in renal cell carcinoma (Nakamura et al., 2015). In conclusion, the list of the genes downregulated by MEK1ca and reversed by HOXB8 (cluster 6, 7 and 8) might thus identify new putative tumor suppressors.

Clustering analysis of the RNA-seq data also identifies genes not reversed by HOXB8. These genes, grouped in cluster 12, only represents 6% of the genes analyzed (114/2316). Surprisingly, several ERK early response genes upregulated by MEK1ca belong to this group (cluster 12, Figure 3B): EGR1, EGR4, ETV3 and ETV5 (Figure 3B, and Figure 5A). *In situ* hybridization with the EGR1 probe confirms the dynamics of expression identified from the RNA-seq data (compare Figure 5A and 5C). These results highlight the specificity of the mechanisms leading to HOXB8 reversion of MEK1ca-induced neoplasia. Since other early genes are however part of the genes reversed by HOXB8 expression, including c-MYC (cluster 4, Figure 5B, Supplementary Figure 9), the global transcriptional reversion of the MEK1ca phenotype by HOXB8 could rely on the transcriptional control of these genes by HOXB8 which might work as nodes of the global transcriptional regulation of the other genes. The MYC genes, which consists of 3 paralogs encoding transcription factors C-MYC, L-MYC and N-MYC, are one of the most frequently deregulated driver genes in human cancer (Duffy et al., 2021). Since our transcriptomic data shows that HOXB8 inhibits the expression of all three MYC genes in the trunk neural tube (Wilmerding et al., 2021a) (Supplementary Figure 9B and 10), one simple explanation of the global transcriptional reversion of the MEK1ca-induced phenotype by HOXB8 is via MYC transcriptional inhibition. It fits at least for LIN28A gene which is controlled at the transcriptional level via MYC (Chang et al., 2009). However, we do not exclude that other gene such as FOS (FOSL2 gene is in the cluster 4, Figure 3B, Supplementary Figure 6) or other early response genes also act as relay of MEK1ca targets reversed by HOXB8, or that HOXB8 acts more as a global epigenetic factor which controls transcription in direct competition with ERK or one (or few) of its key target genes.

**Figure 5:**
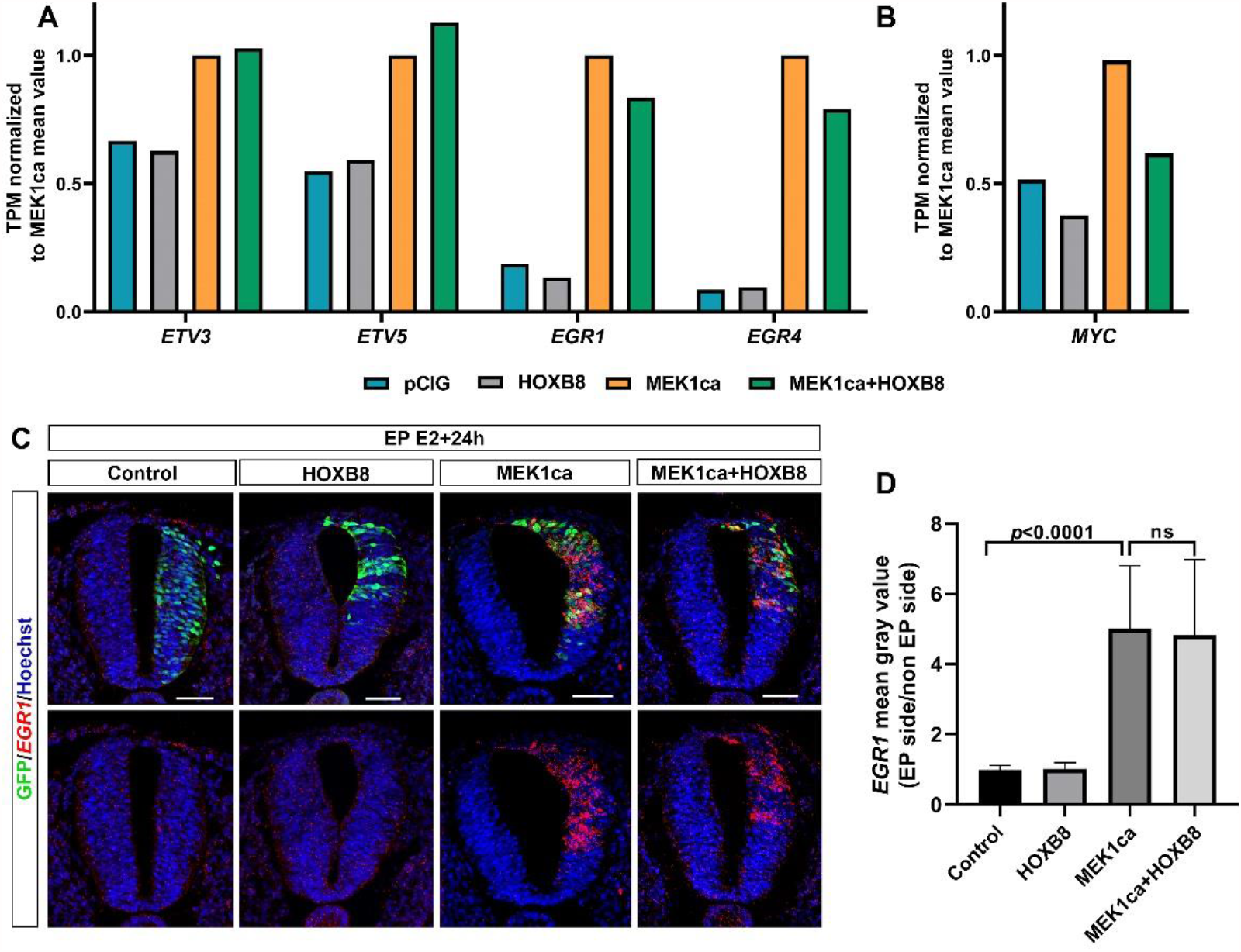
The MEK1ca-induced ETV and EGR gene expression are not reversed by HOXB8. **A and B –** Mean expression in TPM (transcripts per kilobase million) normalized to MEK1ca value for pCIG, HoxB8 and MEK1ca+HOXB8 (mean of the two replicates of each condition) for ETV3, ETB5, EGR1 and EGR1 genes (**A**), and MYC gene (**B**). **C-**Fluorescent *in situ* hybridizations with EGR1 probe and immunofluorescences with anti-GFP antibody on trunk transverse sections of chicken embryo one day post-electroporation in the pCIG, HOXB8, MEK1ca and MEK1ca+HOXB8 conditions. Blue is Hoechst staining. Scale bar: 50µm. **D-***EGR1* expression ratio (electroporated/contralateral side) one day after electroporation in the four conditions (n=3 animals/18 sections). The quantifications show that HOXB8 does not reverse MEK1ca-induced EGR1 expression (two-tailed Mann–Whitney test, error bars represent s.d.).

### CHST15, a target of HOXB8 transcriptional reversion of MEK1ca-induced neoplasia, correlates with poor survival in many human cancers

Among genes of cluster 1 (Figure 3B), the *CHST15* gene caught our attention because it represents one of the most upregulated gene by MEK1ca reversed by HOXB8 (after IL1R1, CDX1 like, CDX4 and DUSP5, and just before AQP1, Supplementary Figure 11). Carbohydrate sulfotransferase 15 (CHST15) is a specific enzyme that biosynthesizes Chondroitin sulfate E (CS-E) (Takakura et al., 2015), a highly sulfated glycosaminoglycan promoting tumor invasion and metastasis. Although CHST15 has been described in the literature as being oncogenic in some cancers including in breast cancer (Liu et al., 2019), its regulation by the ERK pathway or by HOX transcription factors was not described. During embryonic development, *CHST15* is expressed very specifically in the most caudal region of the chicken embryo (Kimura et al., 2011), where FGF8 signaling and pERK are high (Lunn et al., 2007; Wilmerding et al., 2021b) (Figure 6A). We validated the RNA-seq results obtained for *CHST15* (Figure 6B) by *in situ* hybridizations on chicken embryo transverse sections with a *CHST15* probe: MEK1ca increases *CHST15* expression, and this, only in the absence of HOXB8 (Figure 6C-D). Using data from the Human Protein Atlas (TCGA RNA-seq data) (Courtesy of Human Protein Atlas, www.proteinatlas.org) (Uhlen et al., 2017), we highlighted that CHST15, which is already described as an unfavorable prognostic marker for renal cancer, displays a high expression correlated with poor outcome in most human cancers in survival analysis (15/17 cancer types) (Figure 6E-F and Supplementary Figure 12). In conclusion, our finding that CHST15, which seems to be unfavourable for most human cancers, is a positive target of ERK in oncogenic condition, and transcriptionally reversed by HOXB8 transcription factor gain of function, underlines the relevance of the chicken ‘MEK1ca/HOXB8” *in vivo* model to identify new key oncogenes and tumor suppressors, pertinent in human pathologies.

**Figure 6:**
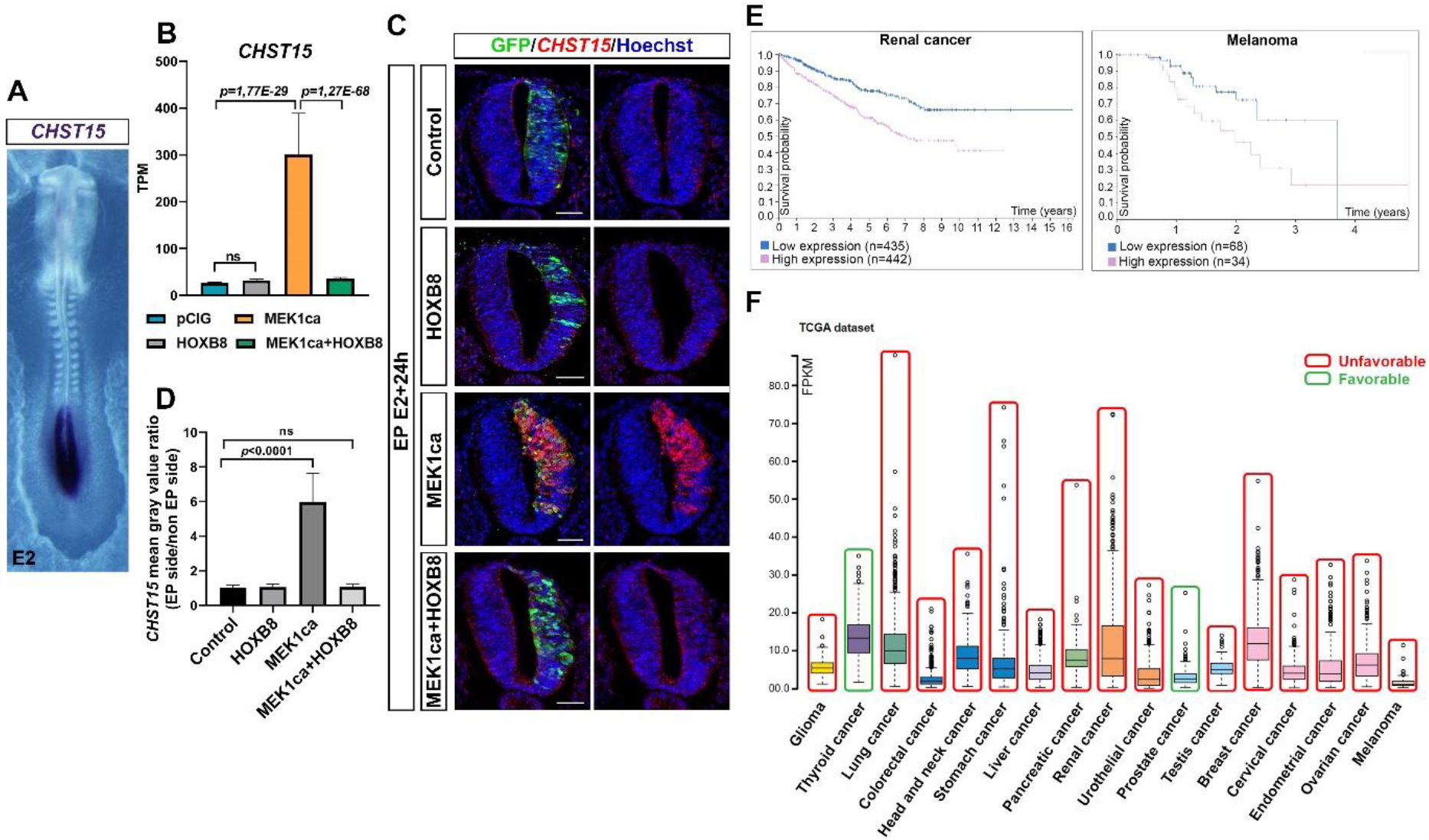
CHST15 upregulated by MEK1ca and reversed by HOXB8 is unfavorable for many cancers. **A-**Dorsal view of a two-days old chicken embryo after a whole mount *in situ* hybridization with a *CHST15* probe highlighting its expression only in the most caudal part of the embryo. **B**-Mean expression of *CHST15* in TPM (transcripts per kilobase million), obtained for the two replicates of the control (pCIG1, pCIG2), HOXB8 (HOXB8-1 and HOXB8-2), MEK1ca (MEK1ca-1 and MEK1ca-2) and MEK1ca+HOXB8 (MEK1ca+HOXB8-1 and MEK1ca+HOXB8-2) expressing samples, with the corresponding p value according to the DGE (FDR5) of each pair. **C-**Fluorescent *in situ* hybridizations with a *Chst15* probe and immunofluorescences with anti-GFP antibody on trunk transverse sections of chicken embryo one day post-electroporation in the pCIG, HOXB8, MEK1ca and MEK1ca+HOXB8 conditions. Blue is Hoechst staining. Scale bar: 50µm. **D-***CHST15* staining mean gray value ratio (electroporated/contralateral side) one day after electroporation in the 4 conditions (n=3 animals/18 sections) (two-tailed Mann–Whitney test, error bars represent s.d. **E-**Survival analysis data from the Human Protein Atlas (Courtesy of Human Protein Atlas, www.proteinatlas.org) highlighting that CHST15 displays a high expression correlated with poor outcome in Renal and Melanoma human cancers. **F**-RNA expression overview (RNA-seq data in 17 cancer types as median FPKM (Fragment Per Kilobase of exon per Million reads), generated by The Cancer Genome Atlas (TCGA) (Courtesy of Human Protein Atlas, www.proteinatlas.org), on which we added a color code (red unfavorable, green favorable) to highlight that most cancers (15/17) have a high expression correlated with poor outcome in the survival analysis data (Human Protein Atlas).

## DISCUSSION

The RAS/RAF/MEK/ERK (MAPK/ERK) pathway is hyperactivated and takes an active part in the malignant transformation in most cancers (Hoshino et al., 1999; Imperial et al., 2019; Maik-Rachline and Seger, 2016), including melanoma (Wellbrock and Arozarena, 2016), and neuroblastoma (Eleveld et al., 2018) cancers (having neural tube cells as embryological origin). Inhibitors of this pathway are used clinically and improve the prognosis of cancer patients for melanomas but are poorly efficient in other cancers (Konieczkowski et al., 2014). In addition, a major problem with the current therapies is that patients often develop resistance (Imperial et al., 2019). Although immunotherapy has recently emerged as an effective therapeutic approach (Hu-Lieskovan et al., 2014), ERK1/2 activity inhibition is still considered a prime target for the treatment of most cancers. As a result, much effort is being made to understand the barriers to current treatments and to discover new therapeutic strategies to counteract ERK hyper-activity in oncology (Imperial et al., 2019).

In this study, we used the chicken embryo neural tube as a platform to explore novel regulatory/interfering paths for ERK oncogenic activity. The model of neoplasia induced by MEK1ca expression we recently developed (Wilmerding et al., 2021b) is experimentally convenient, economic, respects the 3R (Sneddon et al., 2017) and allows to apprehend the epistatic relationship between the ERK pathway and other proteins including transcription factors during oncogenic progression. Furthermore, results obtained identified putative new oncogenes and tumor suppressor genes, that may be particularly relevant for cancers originating from the embryonic neural tube. The data presented also suggest several non-exclusive mechanisms though which HOXB8 acts as tumor suppressor in the neural tube downstream of oncogenic ERK activation. This includes the increase of cell death, the repression of “node” oncogenic genes including MYC (c-MYC, MYC-L and MYC-N) and FOS which control the transcription of a myriad of downstream targets, and the activation of tumor suppressor genes such as *LZTS1*.

One of the transcriptional targets of MYC is *LIN28A* (Chang et al., 2009) which we found upregulated by MEK1ca and reversed by HOXB8 suggesting that HOXB8 might at least counteract MEK1ca oncogenic activity by negatively regulating LIN28A. Recent results obtained in mice suggest that during caudal bud development, the expression of *Lin28* genes are controlled by Hox proteins (Hoxb13 and Hoxc13) (Aires et al., 2019). The transcriptional control of *LIN28* genes by the HOXB8 protein in the chicken neural tube context could thus be a function shared between the HOX proteins and not restricted to this tissue (neural tube) and organism (chicken). We recently highlighted shared or generic HOX functions during the development of the spinal cord in chicken embryo (Wilmerding et al., 2021a), and more generally in other biological contexts and species (Banreti et al., 2014; Saurin et al., 2018). It would thus be interesting to investigate if other HOX proteins behave as HOXB8 in the specific context of MEK1ca neoplasia induction in the embryonic neural tube.

Previous studies also described HOX genes as tumor suppressor. For example, elevated HOXB9 expression predicts a favorable outcome in colon adenocarcinoma patients (Zhan et al., 2014). Also, HOXA5 in colon cancer, is downregulated, and its re-expression induces loss of the cancer stem cell phenotype, preventing tumor progression and metastasis (Ordóñez-Morán et al., 2015). Of note, in renal cancer, HOXB8 is prognostic, with high expression depicted as favorable (Human Protein Atlas resource) (Uhlen et al., 2017)). If these HOX tumor suppressors activities operates by counteracting ERK oncogenic activity already associated to these cancers (Stadler, 2005) is to our knowledge not known.

Finally, our work opens perspectives to explore novel ERK interfering strategies with longer term therapeutic potential. First it identifies a mean to globally block transcriptional changes induced by ERK signaling, downstream ERK phosphorylation and translocation in the nucleus. This defines a biological context and a unique tool to explore ERK interfering approaches in the nucleus. Second, it identifies genes (IL1R1, CDX1-like, CDX4, DUSP5, and AQP1) (Supplementary Figure 11), strongly activated by MEK1ca and nearly fully reversed by HOXB8, which may be used as readouts to screen in an *in vivo* vertebrate context for new molecules able to counteract ERK activity, downstream of pERK nuclear translocation. To render the MEK1ca neoplasia model more convenient and independent of the electroporation step, chicken/quail transgenic animals allowing for in ovo doxycycline induction of MEK1ca could be generated.

## MATERIALS AND METHODS

### Ethics statement

Experiments performed with non-hatched avian embryos in the first twothird of embryonic developmental time are not considered animal experiments according to the directive 2010/63/EU.

### Chicken embryos

Fertilized chicken eggs were obtained from EARL les Bruyeres (Dangers, France) and incubated horizontally at 38°C in a humidified incubator. Embryos were staged according to the developmental table of Hamburger and Hamilton (HH) (Hamburger and Hamilton, 1992) or according to days of incubation (E).

### In ovo electroporation and plasmids

Neural tube *in ovo* electroporations were performed around HH11. Eggs were windowed, and the DNA solution was injected in neural tube lumen. Needle L-shape platinum electrodes (CUY613P5) were placed on both sides of the embryo at trunk level (5 mm apart), with the cathode always at its right. Five 50 ms pulses of 25 volts were given unilateral (or bilateral for RNA-seq experiments) at 50 ms intervals with an electroporator NEPA21 (Nepagene).

The plasmids used for the gain-of-function experiments co-express a cytoplasmic or nuclear GFP (pCAGGS and pCIG respectively, used alone as controls) and the coding sequence (CDS) of the gene of interest. Vectors used are: pCIG-MEK1ca (Delfini et al., 2005), pCIG-HOXB8 (gifted by Doctor Olivier Pourquié). All the plasmids used for electroporation were purified using the Nucleobond Xtra Midi kit (Macherey-Nagel). Final concentration of DNA delivered by embryo for electroporation is between 1 to 2µg/µl.

### Tissue section

Embryos were fixed in 4% buffered formaldehyde in PBS then treated with a sucrose gradient (15% and 30% in PBS), embedded in OCT medium and stored at −80°C. Embryos were sectioned into 16 µm sections with a Leica cryostat and the slides were conserved at – 80°C or directly used for FISH and/or immunofluorescence.

### Immunofluorescence

Slides were rehydrated in PBS then blocked with 10% goat serum, 3% BSA, 0,4% Triton X-100 in PBS for one hour. Primary antibodies were incubated over-night diluted in the same solution at 4°C. The following primary antibodies were used in this study: chicken anti-GFP 1:1000 (1020 AVES), rabbit anti-Phospho-p44/42 MAPK (ERK1/2) (Thr202/Tyr204) Antibody #9101 (Cell Signaling Technology), rabbit anti-SOX2 1:500 (AB5603 Merck Millipore), mouse anti-Tuj1 1:500 (801202 Biolegend), rat anti-pH3 1: 250 (S28, abcam ab10543), rabbit anti-Caspase 3 1:500 (Asp175, CST 9661). The secondary antibodies used were: anti-chicken, anti-rabbit, anti-mouse or anti-rat with fluorochromes (488, 568 or 647) at 1:500 (Alexa Fluor, abcam). They were incubated for one hour in the blocking solution containing Hoechst (1:1000). Slides were washed, mounted (Thermo Scientific Shandon Immu-Mount) and imaged with a Zeiss microscope Z1 equipped with Apotome or a confocal LSM 780.

### In situ hybridization

RNA probes used for *in situ* hybridization were: DUSP5, IL17RD, LIN28A, EGR1 CHST15 (Wilmerding et al., 2021b), TMEM132C (fw: CTGCCTTGAAATGGCCGGT and rev: AGGTGTCTGCACCAGATCGT, GREB1 (fw: TATGCTGGACAGCTCAAGACA, rev: TTGCGCCCATTATCATCTGGA). The probes were produced from PCR product amplified from cDNA neural tube from E3 chicken embryo (WT or transfected by MEK1ca). All the forward primers contain the T7 RNA polymerase promotor sequence: TAATACGACTCACTATAGGGC.

#### Fluorescent in situ hybridization on tissue section

The slides were treated with proteinase K 10 µg/ml (3 minutes at 37°C) in a solution of TrisHCl 50 mM pH 7.5, then in triethanolamine 0.1M and 0.25% acetic anhydride. They were pre-incubated with hybridization buffer (50% formamide, SSC 5X, Denhardts 5X, yeast tRNA 250 µg/ml and herring sperm DNA 500 µg/ml) for 3h at room temperature and incubated in the same buffer with DIG-labelled RNA probes over-night at 55°C in a wet chamber. The slides were then washed twice with 0.2X SSC for 30 minutes at 65°C. After 5 minutes in TNT buffer (100 mM Tris pH7.5, 150mM NaCl and 0.1% Tween-20), they were then blocked for 1h in buffer containing TNT 1X, 1% Blocking reagent (Roche) and 10% goat serum, then incubated in the same buffer for 3h with anti-DIG-POD antibodies (1:500, Roche) and revealed using the kit TSA-Plus Cyanin-3 (Perkin Elmer).

### Whole mount in situ hybridization

Embryos were fixed 2h at RT in 4% formaldehyde in PBS. Embryos were dehydrated with sequential washes in 50% ethanol/ PBS+ 0.1% Tween20 and 100% ethanol and conserved at − 20°c. Embryos were bleached for 45mn in 80% ethanol + 20% of 30% H2O2 and then rehydrated. They were treated X minutes with proteinase K 10µg/ml at RT and refixed with 4% formaldehyde, 0.2% glutaraldehyde. After 1h of blocking in the hybridization buffer (50% formamide, SSC 5x, 50μg/mL Heparine, yeast tRNA 50μg/mL, SDS 1%) hybridization with DIG-labelled RNA probes was performed at 68°C overnight. The next day, embryos were washed (3 times 30 minutes) in hybridization buffer and 1 time in TBS (25 mM Tris, 150 mM NaCl, 2 mM KCl, pH 7.4) +0.1% Tween 20. They were incubated 1h at RT in a blocking buffer (20% Blocking reagent + 20% Goat serum) and then overnight with an anti-DIG-AP antibody (1:2000, Roche) in the blocking buffer. After 3 washes (1 hour) in TBS+0.1% Tween 20, embryos were equilibrated (2 times 10 minutes) in NTMT buffer (NaCl 100mM, TrisHCl 100mM pH9,5, MgCl2 50mM, 2%Tween20) and incubated in NBT/BCIP (Promega) at RT in the dark until color development. Pictures of whole embryos were made using a BinoFluo MZFLIII and a color camera.

### RNA-seq analysis

Electroporations were carried out as described in a previous section but with 5 bilateral pulses, with the protocol described in (Wilmerding et al., 2021a). Results for pCIG (2µg/µl), pCIG-MEK1ca (1 µg/µl + pCIG 1µg/µl) and pCIG-HOXB8 (1 µg/µl + pCIG 1µg/µl) were published in (Wilmerding et al., 2021a) and (Wilmerding et al., 2021b), and compared with pCIG-MEK1ca + pCIG-HOXB8 (1 µg/µl of each) condition. Part of the neural tube expressing the GFP were dissected 18 hours after electroporation and dissociated (Trypsin-EDTA 0,25%). GFP-expressing cells was isolated by FACS with the use of a dead cell exclusion (DCE)/discrimination dye (DAPI) to eliminate dying cells (Wilmerding et al., 2021a). RNA was extracted (RNeasy Mini Kit) and reverse transcribed and cDNA was amplified using a linear amplification system and used for sequencing library building (GATC): Random primed cDNA library, purification of poly-A containing mRNA molecules, mRNA fragmentation, random primed cDNA synthesis, adapter ligation and adapter specific PCR amplification, Illumina technology, 50 000 000 reads paired end with 2 x 50 bp read length. Bioinformatics analyses were done using the galgal4.0 chicken genome. Qualitative analysis of RNA-seq data from the four biological replicates shows a high Pearson Correlation score (>0,99) indicative of the experimental reproducibility (Supplementary Figure 4).

### Quantifications and statistical significance

The number of embryos and number sections analyzed are indicated in the figure legends. A minimum of 3 embryos and 6 sections per embryo were used to quantify. All quantifications were made using the cell counter tool of Fiji software. The results were analyzed and plotted using Prism 8 software (GraphPad software). Statistical analyses were performed using a two-tailed Mann Whitney test and considered significant when p-value < 0,05. All p-values are indicated on the graphs. The error bars represent the standard deviation (SD).

## Acknowledgements

We sincerely thank Dr Andrew Saurin for his help in the RNA-seq analysis. We would like to thank Sophie Gournet (IBPS, CNRS UMR7622) for the drawings of Figures 2, and the Optical Imaging Platform of the IBDM. We sincerely thank Heather Etchevers for her critical reading of the manuscript. We thank Dr Olivier Pourquié for its generous gifts of pCIG-HOXB8 expression vector. We thank Dr Bianca Habermann for her advice on bioinformatics and Laure Lore for their technical support at the beginning of this study. We also would like to thank Emilie Bonzom, Nomenjanahary Solotahiana and Lalla-Kamilya Gaouzi for their implication in this work during their Master internship. FACS experiments were done at the CRCM (Marseille, France) and transcriptome sequencing at the GATC. We would like to warmly thank LA LIGUE CONTRE LE CANCER, the Cancéropôle PACA, the AMIDEX, the ANR and the IBDM for their founding support.

