## SUPPLEMENTARY FIGURES AND TABLES Supplementary for "HOXB8 counteracts MAPK/ERK oncogenic signaling in a chicken embryo model of neoplasia"

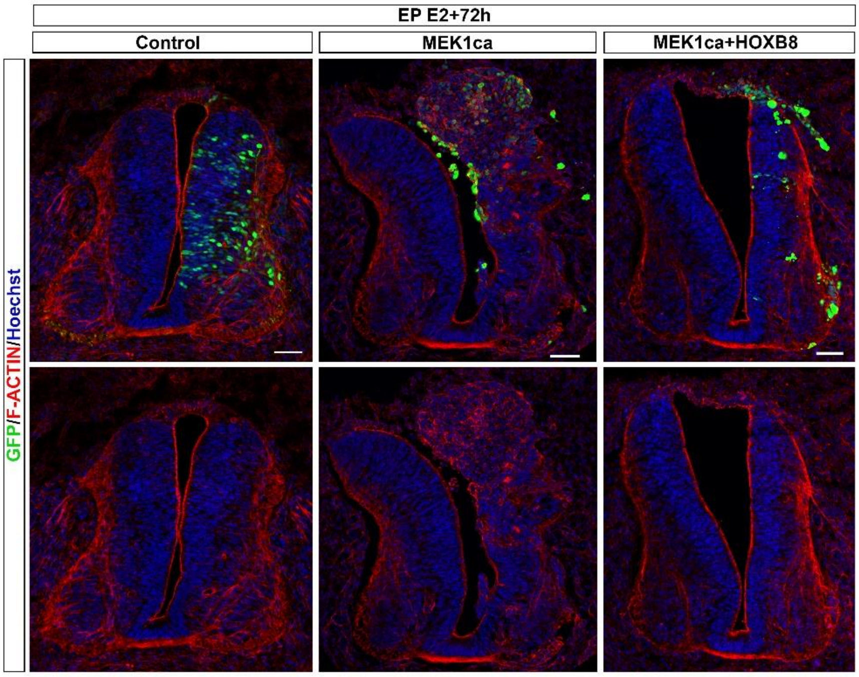

**Supplementary Figure 1: HOXB8 gain of function in the trunk neural tube of the chicken embryo transfected by MEK1ca expressing vector inhibits neoplasia formation**
Immunofluorescences on transverse sections of chicken embryo with anti-GFP antibody with a co-staining against F-actin (phalloidin), 3 days day post-electroporation with PCIG, MEK1ca or MEK1ca+HOXB8 expressing vectors. Blue is Hoechst staining. Scale bar: 50µm.

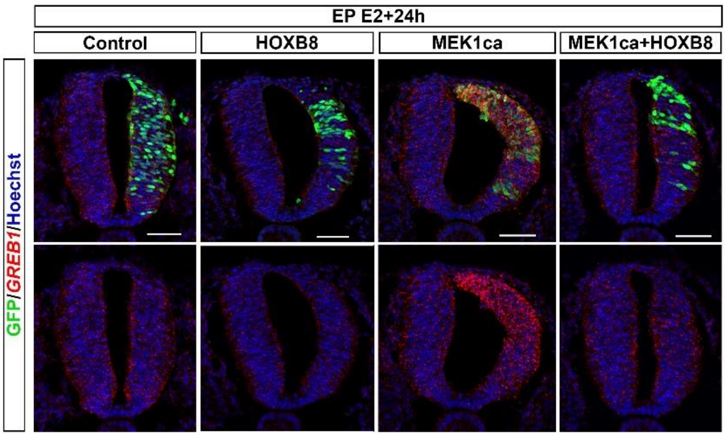

**Supplementary Figure 2: GREB1 is upregulated by MEK1ca in the chicken neural tube and reversed by HOXB8 co-expression**

Fluorescent *in situ* hybridizations with GREB1 probe and immunofluorescences with anti-GFP antibody on trunk transverse sections of chicken embryo one day post-electroporation in the pCIG, HOXB8, MEK1ca and MEK1ca+HOXB8 conditions. Blue is Hoechst staining. Scale bar: 50µm.

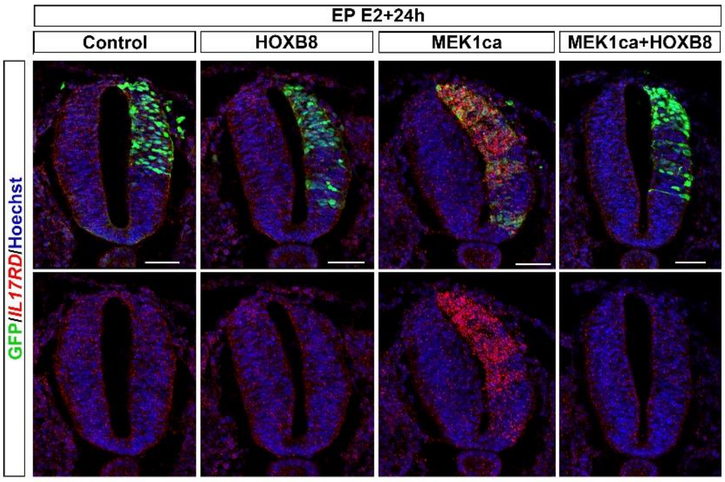

**Supplementary Figure 3: IL17RD is upregulated by MEK1ca in the chicken neural tube and reversed by HOXB8 co-expression**

Fluorescent *in situ* hybridization with IL17RD probe and immunofluorescences with anti-GFP antibody on trunk transverse sections of chicken embryo one day post-electroporation in the pCIG, HOXB8, MEK1ca and MEK1ca+HOXB8 conditions. Blue is Hoechst staining. Scale bar: 50µm.

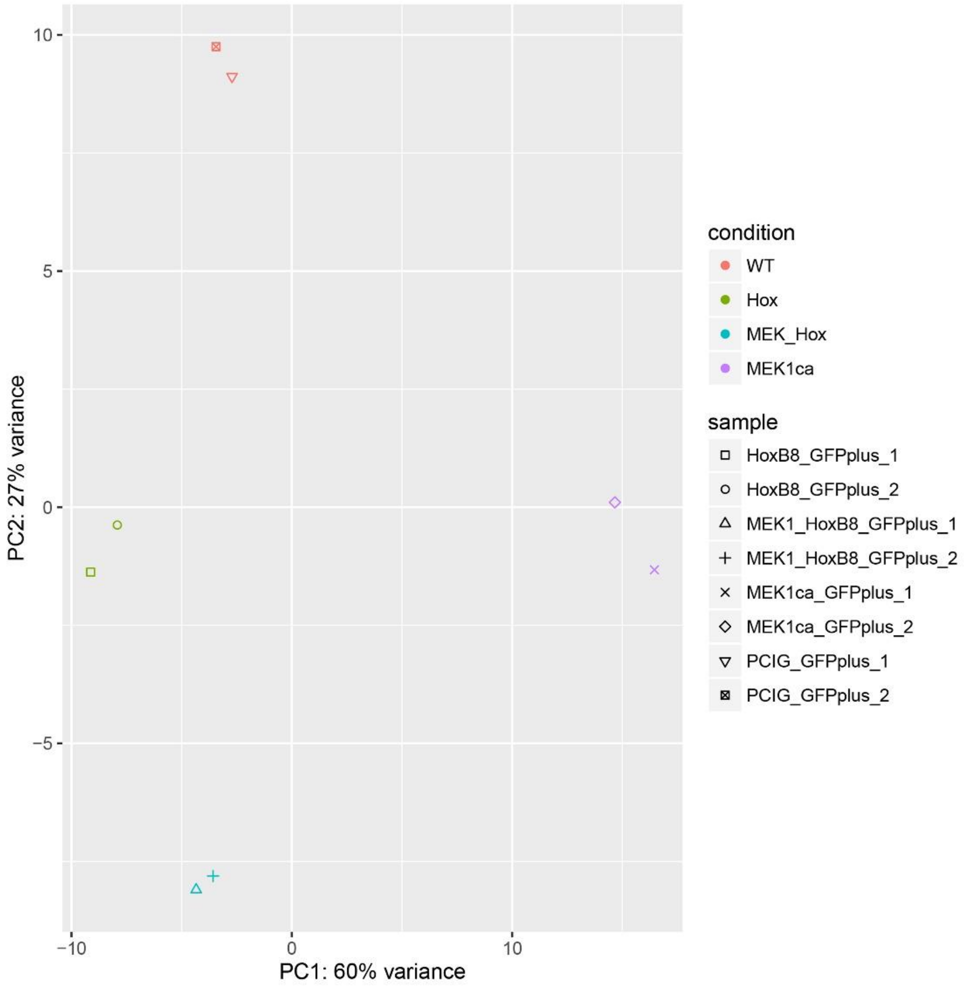

**Supplementary Figure 4:** Principal component analysis (PCA) obtained from RNA-seq counting data using galgal4 assembly for the two replicates of the control (pCIG1, pCIG2), HOXB8 (HOXB8-1 and HOXB8-2), MEK1ca (MEK1ca-1 and MEK1ca-2) and MEK1ca+HOXB8 (MEK1ca+HOXB8-1 and MEK1ca+HOXB8-2) expressing samples, highlighting the quality of the biological replicates.

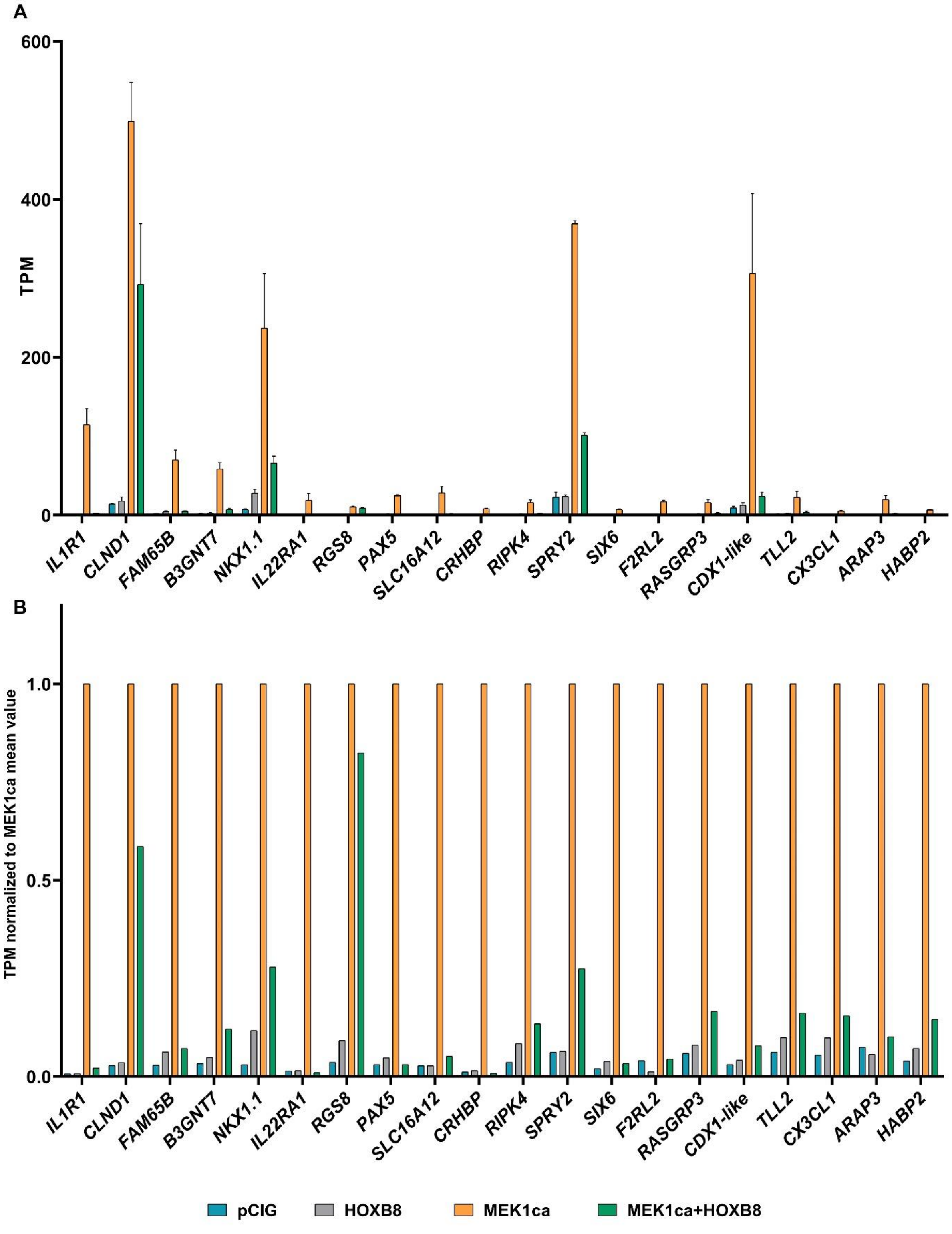

**Supplementary Figure 5**

**Supplementary Figure 5: A and B-** Graph of the mean of the TPM (transcripts per kilobase million) number for these 20 genes, obtained for the two replicates of the control (pCIG1, pCIG2), HOXB8 (HOXB8-1 and HOXB8-2), MEK1ca (MEK1ca-1 and MEK1ca-2) and MEK1ca+HOXB8 (MEK1ca+HOXB8-1 and MEK1ca+HOXB8-2) expressing samples (**A**) and normalized to MEK1ca value (**B**).

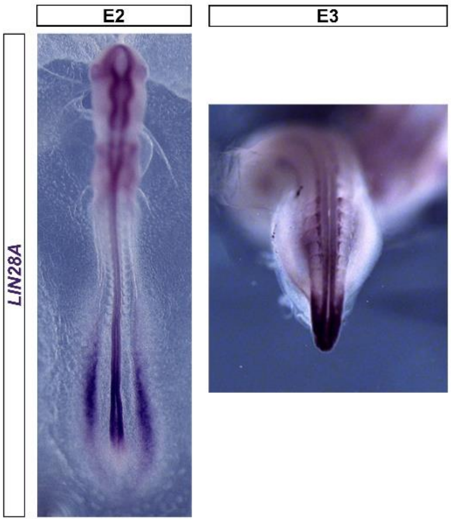

**Supplementary Figure 6: A and B-** Dorsal view of a two-days old (**A**) and three-days old (only the most caudal part) (**B**) chicken embryo after a whole mount *in situ* hybridization with LIN28A probe highlighting that its expression is the strongest in the most caudal part of the embryo.

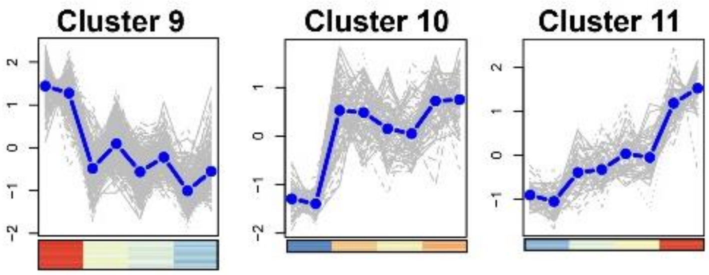

**Supplementary Figure 7:** Plots of clusters 9, 10 and 11 (related to Figure 3) with the two replicates of each condition (control (pCIG1, pCIG2), HOXB8 (HOXB8-1 and HOXB8-2), MEK1ca (MEK1ca-1 and MEK1ca-2) and MEK1ca+HOXB8 (MEK1ca+HOXB8-1 and MEK1ca+HOXB8-2)), with the mean (in blue), and with the corresponding heat map on the bottom.

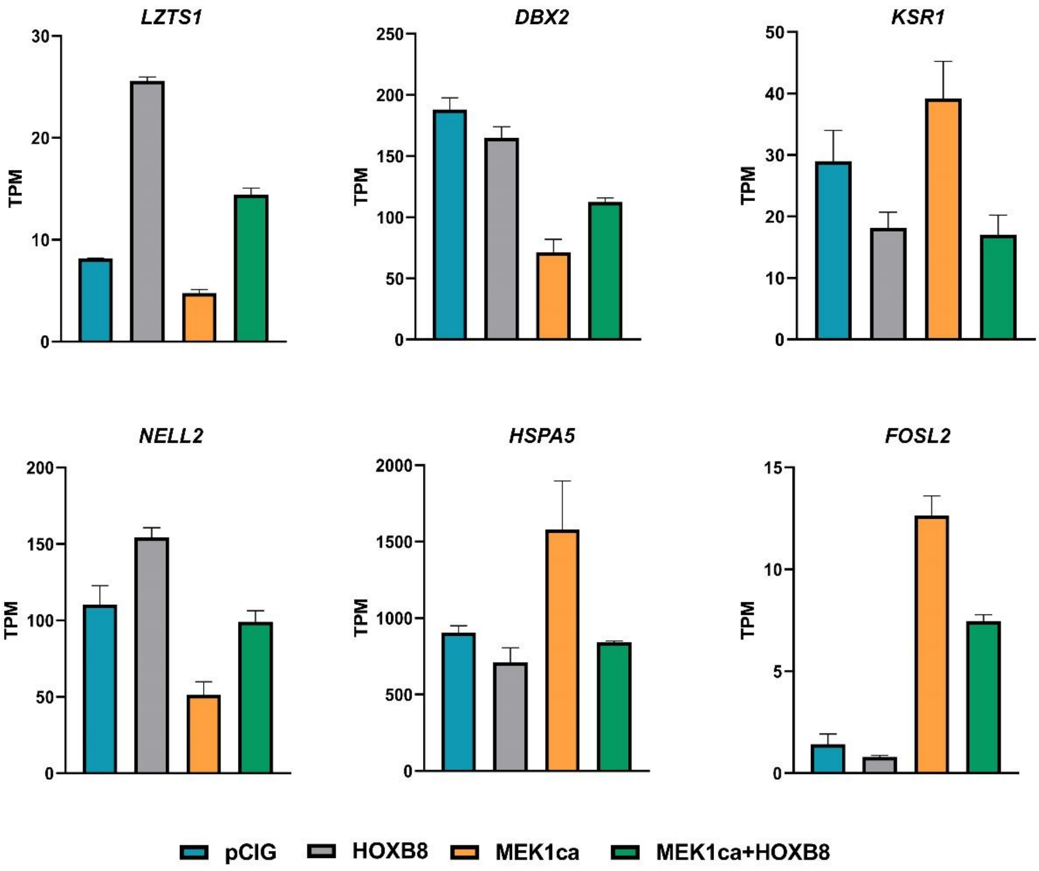

**Supplementary Figure 8:** Graph of the mean of the number of the TPM (transcripts per kilobase million) for LZTS1, KRS1 and NELL2 genes, obtained for the two replicates of the control (pCIG1, pCIG2), HOXB8 (HOXB8-1 and HOXB8-2), MEK1ca (MEK1ca-1 and MEK1ca-2) and MEK1ca+HOXB8 (MEK1ca+HOXB8-1 and MEK1ca+HOXB8-2) expressing samples, with the corresponding p value according to the DGE (FDR= 5%5) of each pair.

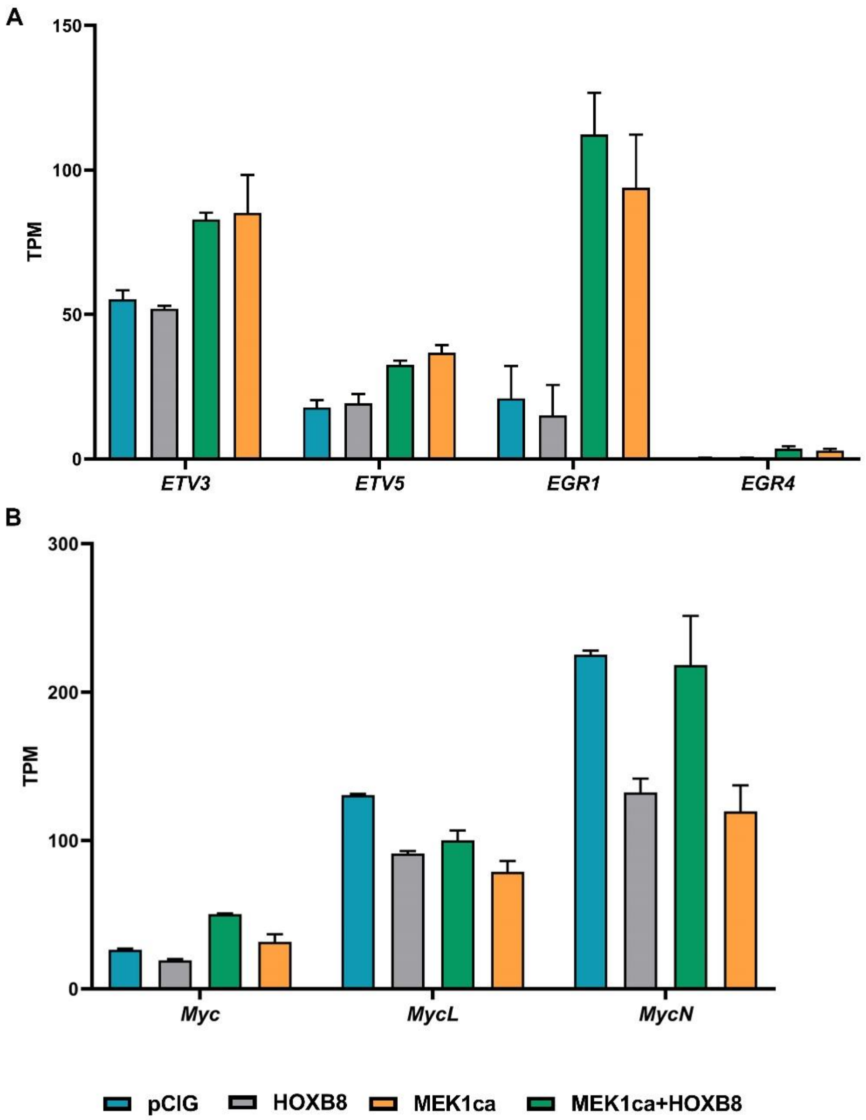

**Supplementary Figure 9: A and B -** Graph of the mean of the number of the TPM (transcripts per kilobase million) for ETV3, ETV, EGR1 and EGR4 genes (**A**), and MYC, MYCL and MYCN (**B**) obtained for the two replicates of the control (pCIG1, pCIG2), HOXB8 (HOXB8-1 and HOXB8-2), MEK1ca (MEK1ca-1 and MEK1ca-2) and MEK1ca+HOXB8 (MEK1ca+HOXB8-1 and MEK1ca+HOXB8-2) expressing samples, with the corresponding p value according to the DGE (FDR= 5%) of each pair.

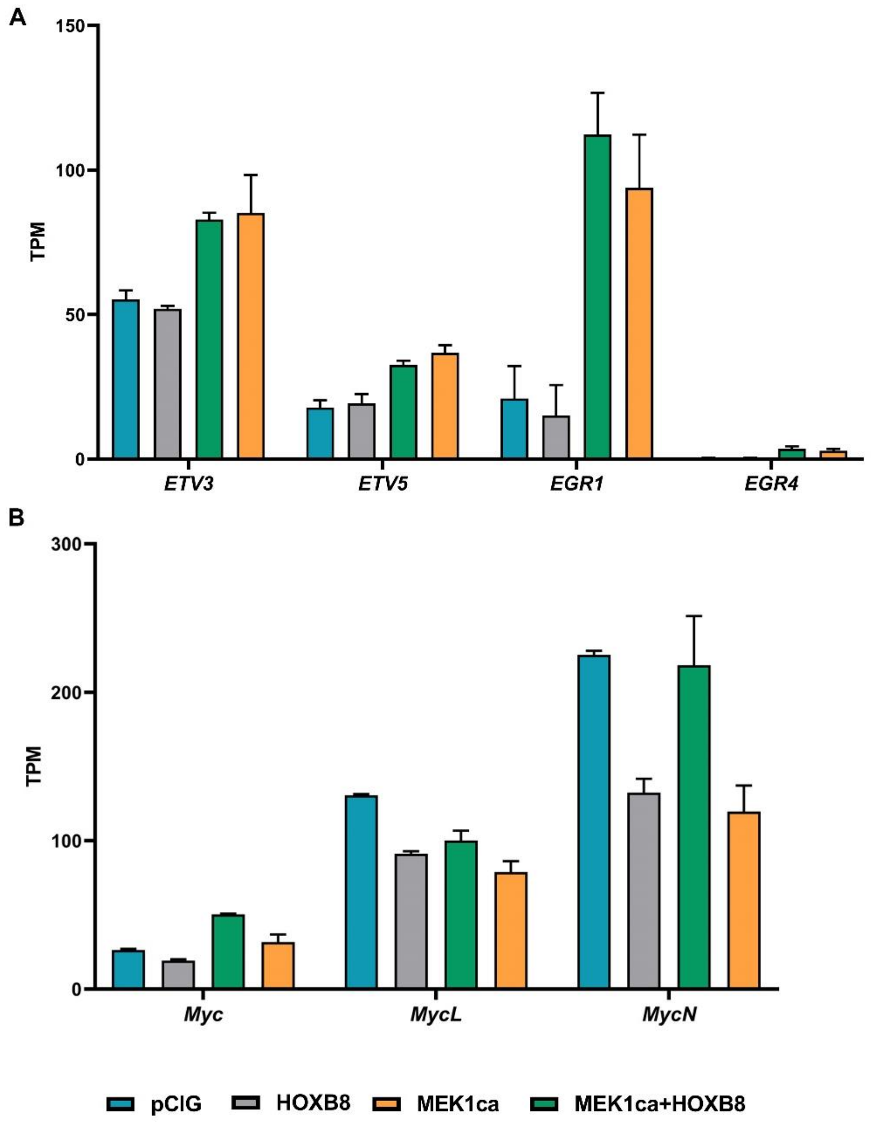

**Supplementary Figure 10:** Graph of TPM (transcripts per kilobase million) normalized on MEK1ca value for pCIG, HOXB8 and MEK1ca+HOXB8 (mean of the two replicates of each condition) for MYC, MYCL and MYCN genes.

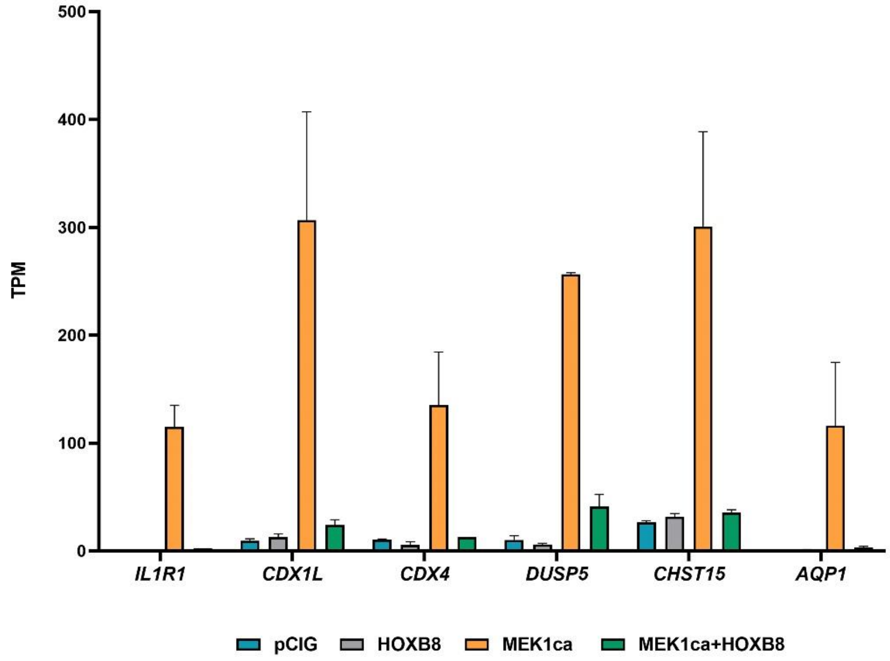

**Supplementary Figure 11:** Graph of the mean of the number of the TPM (transcripts per kilobase million) for IL1R1, CDX1l, CDX4, DUSP5, CHST15 and AQP1 genes obtained for the two replicates of the control (pCIG1, pCIG2), HOXB8 (HOXB8-1 and HOXB8-2), MEK1ca (MEK1ca-1 and MEK1ca-2) and MEK1ca+HOXB8 (MEK1ca+HOXB8-1 and MEK1ca+HOXB8-2) expressing samples.

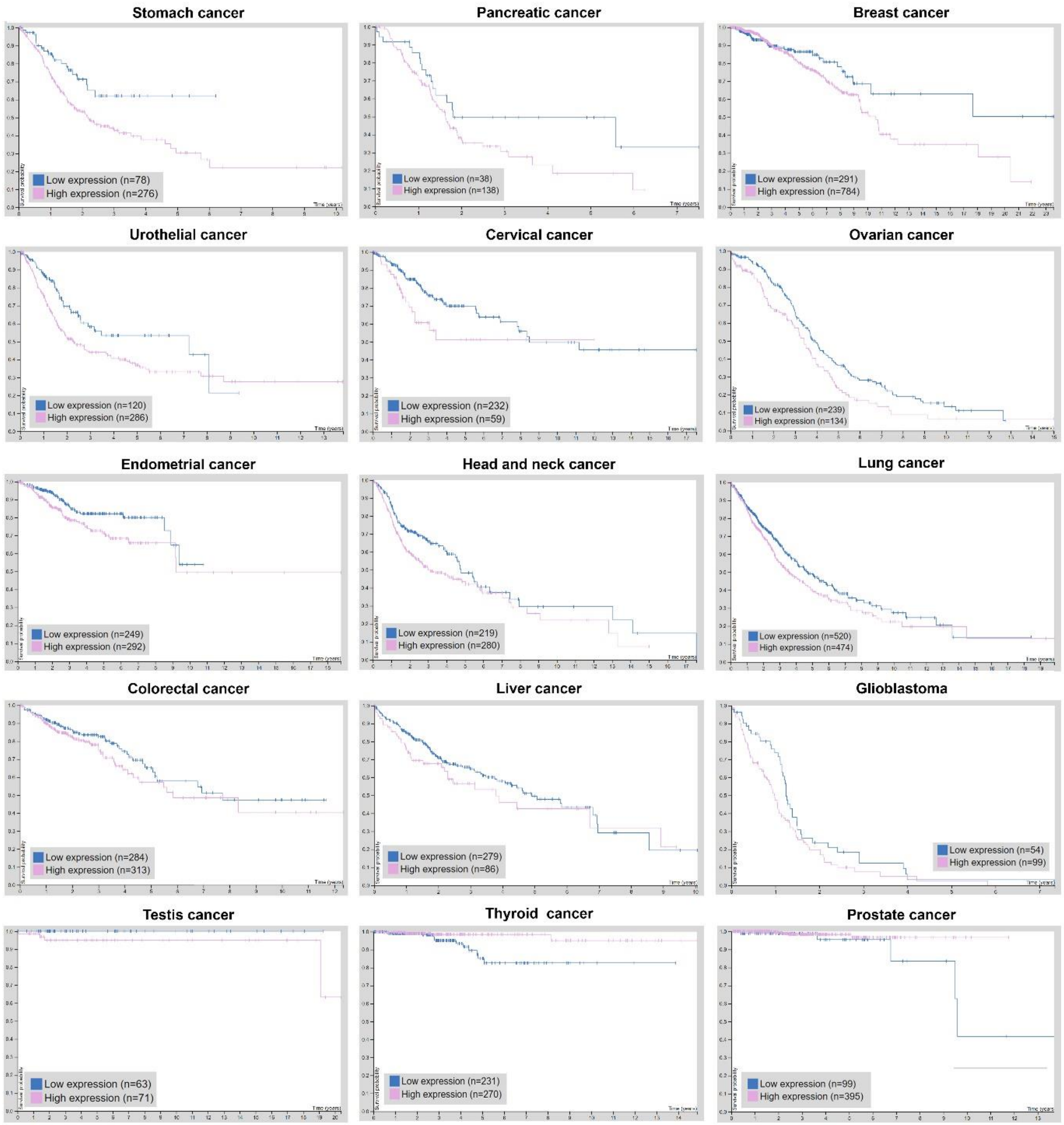

**Supplementary Figure 12:** Survival analysis data from the Human Protein Atlas (Courtesy of Human Protein Atlas, www.proteinatlas.org) for CHST15 displays (for all cancer types except Renal cancer and melanoma display Figure 6).

### SUPPLEMENTARY TABLE LEGENDS

**Supplementary Table 1:** MEK1ca+HOXB8 versus MEK1ca differentially regulated genes (False Discovery Rate (padj) <= 0.0001)

**Supplementary Table 2:** MEK1ca vs pCIG differentially regulated genes (False Discovery Rate (padj) <= 0.0001)

**Supplementary Table 3:** Commonly differentially regulated genes between MEK1+HOXB8 vs MEK1ca and pCIG vs MEK1ca (False Discovery Rate (padj) <= 0.0001). Genes that are inversely regulated are highlighted.

**Supplementary Table 4:** MEK1ca+HOXB8 versus MEK1ca differentially regulated genes (False Discovery Rate (padj) <= 0.005)

**Supplementary Table 5**: Classification of the 2316 genes deregulated in the “MEK1ca versus pCIG” condition (False Discovery Rate (padj) <= 0.005) clustered using “kmeans” with k=12, with the mean TPM (transcripts per kilobase million) of each gene for the two replicates of pCIG, HOXB8, MEK1ca, and MEK1ca+HOXB8 in each condition.

**Supplementary Table 1:** MEK1ca+HOXB8 versus MEK1ca differentially regulated genes (False Discovery Rate (padj) <= 0.0001)

| ID | baseMean | log2FoldChange | lfcSE | stat | pvalue | padj | Symbol |
|---|---|---|---|---|---|---|---|
| ENSGALG00000026853 | 1756,76051 | 5,751530736 | 0,19626668 | 29,3046727 | 9,04E-189 | 6,16E-185 | FGF3 |
| ENSGALG00000011433 | 901,861183 | 4,353428748 | 0,19799365 | 21,9877188 | 3,77E-107 | 1,28E-103 | |
| ENSGALG00000009778 | 267,922713 | 4,29765073 | 0,29733506 | 14,4538982 | 2,37E-47 | 9,77E-45 | |
| ENSGALG00000011582 | 175,170205 | 4,101846269 | 0,31827959 | 12,8875569 | 5,29E-38 | 1,53E-35 | |
| ENSGALG00000027742 | 312,342696 | 3,96517198 | 0,26630875 | 14,8893793 | 3,86E-50 | 1,81E-47 | KLHL33 |
| ENSGALG00000028044 | 194,51466 | 3,90733608 | 0,30410231 | 12,848755 | 8,74E-38 | 2,48E-35 | |
| ENSGALG00000002108 | 813,374631 | 3,799830272 | 0,19658831 | 19,328872 | 3,07E-83 | 5,23E-80 | EVPL |
| ENSGALG00000021395 | 138,406928 | 3,681673719 | 0,32878732 | 11,1977363 | 4,18E-29 | 6,70E-27 | |
| ENSGALG00000001080 | 589,095015 | 3,620989798 | 0,22311063 | 16,2295708 | 3,12E-59 | 2,02E-56 | NR5A1 |
| ENSGALG00000005292 | 233,995114 | 3,593969871 | 0,28192744 | 12,7478541 | 3,20E-37 | 8,90E-35 | KLHL40 |
| ENSGALG00000028376 | 1521,83584 | 3,47510163 | 0,19828778 | 17,525546 | 9,15E-69 | 7,78E-66 | FGF19 |
| ENSGALG00000023843 | 120,841933 | 3,393126537 | 0,33077585 | 10,2580842 | 1,09E-24 | 1,35E-22 | FRMD7 |
| ENSGALG00000006471 | 279,298959 | 3,309144864 | 0,28892507 | 11,4532978 | 2,26E-30 | 4,04E-28 | SLCO2A1 |
| ENSGALG00000028304 | 118,793101 | 3,251722149 | 0,33476041 | 9,71358044 | 2,64E-22 | 2,58E-20 | |
| ENSGALG00000004526 | 421,691059 | 3,217263436 | 0,24373894 | 13,1996283 | 8,82E-40 | 2,93E-37 | TNR |
| ENSGALG00000008069 | 109,411674 | 3,182915984 | 0,34138391 | 9,32356755 | 1,12E-20 | 9,51E-19 | |
| ENSGALG00000004134 | 133,434282 | 3,164728278 | 0,32672711 | 9,6861515 | 3,45E-22 | 3,31E-20 | GLRA1 |
| ENSGALG00000012254 | 87,731989 | 3,162704218 | 0,3459847 | 9,14116788 | 6,18E-20 | 4,98E-18 | KCNJ4 |
| ENSGALG00000006944 | 469,144675 | 3,116432937 | 0,23983811 | 12,9939021 | 1,32E-38 | 4,01E-36 | |
| ENSGALG00000028612 | 103,60478 | 3,070056483 | 0,34649284 | 8,86037489 | 7,97E-19 | 5,81E-17 | |
| ENSGALG00000005285 | 100,237429 | 2,996752626 | 0,34025326 | 8,80741788 | 1,28E-18 | 8,94E-17 | TBX4 |
| ENSGALG00000007335 | 107,298878 | 2,985281892 | 0,34055773 | 8,76586154 | 1,85E-18 | 1,28E-16 | KBTBD13 |
| ENSGALG00000001256 | 266,944995 | 2,963677659 | 0,29793308 | 9,94746081 | 2,59E-23 | 2,71E-21 | NLRC5 |
| ENSGALG00000016685 | 93,4474561 | 2,945393761 | 0,33836846 | 8,70469363 | 3,18E-18 | 2,13E-16 | ASMT |
| ENSGALG00000016721 | 160,3282 | 2,940591405 | 0,30420224 | 9,66656738 | 4,18E-22 | 3,93E-20 | PLCXD1 |
| ENSGALG00000026167 | 171,614092 | 2,925839411 | 0,31778334 | 9,20702576 | 3,35E-20 | 2,78E-18 | PIK3R5 |
| ENSGALG00000002926 | 76,2289496 | 2,900739748 | 0,35004614 | 8,28673542 | 1,16E-16 | 6,86E-15 | |
| ENSGALG00000004700 | 97,8406537 | 2,88173675 | 0,33435932 | 8,61868224 | 6,77E-18 | 4,45E-16 | NCF2 |
| ENSGALG00000016722 | 73,4311203 | 2,829128945 | 0,35273145 | 8,0206314 | 1,05E-15 | 5,52E-14 | |
| ENSGALG00000012830 | 227,950727 | 2,760094911 | 0,2851044 | 9,68099721 | 3,63E-22 | 3,44E-20 | |
| ENSGALG00000027247 | 97,4363295 | 2,75413056 | 0,33274463 | 8,27700976 | 1,26E-16 | 7,41E-15 | EOMES |
| ENSGALG00000002823 | 276,033949 | 2,746001119 | 0,25652175 | 10,7047499 | 9,67E-27 | 1,39E-24 | |
| ENSGALG00000006614 | 540,926615 | 2,721769719 | 0,26010327 | 10,4641889 | 1,26E-25 | 1,70E-23 | |
| ENSGALG00000006744 | 446,950734 | 2,709854582 | 0,26391565 | 10,2678814 | 9,83E-25 | 1,23E-22 | |
| ENSGALG00000020485 | 352,825328 | 2,707785881 | 0,28649565 | 9,45140328 | 3,34E-21 | 2,90E-19 | |
| ENSGALG00000015322 | 401,95748 | 2,694878773 | 0,24687156 | 10,9161165 | 9,65E-28 | 1,43E-25 | IMPG2 |
| ENSGALG00000025886 | 234,932728 | 2,681244585 | 0,27696529 | 9,68079628 | 3,64E-22 | 3,44E-20 | SUSD3 |
| ENSGALG00000013922 | 145,692651 | 2,651462599 | 0,31447645 | 8,43135507 | 3,42E-17 | 2,11E-15 | |
| ENSGALG00000026590 | 256,390276 | 2,649598672 | 0,26552851 | 9,97858437 | 1,89E-23 | 2,01E-21 | |
| ENSGALG00000016325 | 669,814955 | 2,632246576 | 0,2229691 | 11,8054319 | 3,66E-32 | 7,66E-30 | |
| ENSGALG00000026627 | 138,559837 | 2,620280418 | 0,32529098 | 8,0551894 | 7,94E-16 | 4,29E-14 | |
| ENSGALG00000001780 | 98,3971 | 2,613843166 | 0,33586268 | 7,78247584 | 7,11E-15 | 3,32E-13 | PROC |
| ENSGALG00000002987 | 118,456359 | 2,607861451 | 0,32067745 | 8,13235054 | 4,21E-16 | 2,34E-14 | SERPINF2 |
| ENSGALG00000027059 | 200,391038 | 2,606173672 | 0,28900153 | 9,01785421 | 1,92E-19 | 1,48E-17 | |
| ENSGALG00000023821 | 140,669435 | 2,605691323 | 0,31496616 | 8,2729246 | 1,31E-16 | 7,64E-15 | |
| ENSGALG00000006032 | 296,771768 | 2,599165832 | 0,27225162 | 9,54692527 | 1,34E-21 | 1,19E-19 | |
| ENSGALG00000002960 | 103,932494 | 2,574242623 | 0,33553695 | 7,67200941 | 1,69E-14 | 7,76E-13 | HMGCS2 |
| ENSGALG00000010841 | 110,200839 | 2,5587632 | 0,33214766 | 7,70369186 | 1,32E-14 | 6,10E-13 | |
| ENSGALG00000002829 | 649,133449 | 2,553310959 | 0,20495773 | 12,4577442 | 1,27E-35 | 3,20E-33 | |
| ENSGALG00000021198 | 61,5032288 | 2,552636185 | 0,35110067 | 7,27038252 | 3,58E-13 | 1,39E-11 | |
| ENSGALG00000015141 | 58,9626486 | 2,548558749 | 0,35207483 | 7,23868481 | 4,53E-13 | 1,74E-11 | GDA |
| ENSGALG00000005122 | 338,26914 | 2,528381619 | 0,24275499 | 10,415364 | 2,11E-25 | 2,79E-23 | MYO1H |
| ENSGALG00000010145 | 278,247492 | 2,508391386 | 0,30141971 | 8,3219221 | 8,65E-17 | 5,15E-15 | MDGA1 |
| ENSGALG00000014372 | 67,9784132 | 2,495880293 | 0,34928387 | 7,14570731 | 8,95E-13 | 3,29E-11 | SLC34A2 |
| ENSGALG00000029144 | 72,3286127 | 2,46980754 | 0,34882621 | 7,08033811 | 1,44E-12 | 5,17E-11 | |
| ENSGALG00000021399 | 89,3295109 | 2,464358792 | 0,33686101 | 7,31565468 | 2,56E-13 | 1,01E-11 | |
| ENSGALG00000001111 | 752,305162 | 2,451755117 | 0,21130589 | 11,6028718 | 3,98E-31 | 7,64E-29 | |
| ENSGALG00000012985 | 261,612669 | 2,44205884 | 0,26586807 | 9,18522799 | 4,11E-20 | 3,39E-18 | SLC6A13 |
| ENSGALG00000008105 | 47,87761 | 2,404806479 | 0,35395793 | 6,79404614 | 1,09E-11 | 3,44E-10 | |
| ENSGALG00000012812 | 176,061446 | 2,39422427 | 0,29597013 | 8,08941196 | 6,00E-16 | 3,29E-14 | SVOPL |

| | | | | | | | |
|---|---|---|---|---|---|---|---|
| ENSGALG00000002638 | 117,914057 | 2,388124012 | 0,32831388 | 7,27390518 | 3,49E-13 | 1,35E-11 | |
| ENSGALG00000002318 | 63,4096631 | 2,376044719 | 0,3494777 | 6,79884494 | 1,05E-11 | 3,33E-10 | LVRN |
| ENSGALG00000002015 | 214,323208 | 2,374226325 | 0,28640932 | 8,28962651 | 1,14E-16 | 6,72E-15 | |
| ENSGALG00000026256 | 199,837668 | 2,357924834 | 0,33320574 | 7,0764832 | 1,48E-12 | 5,30E-11 | |
| ENSGALG00000007395 | 402,677514 | 2,356587415 | 0,22616715 | 10,4196716 | 2,02E-25 | 2,69E-23 | ABCC2 |
| ENSGALG00000002046 | 463,721693 | 2,341658671 | 0,24228891 | 9,66473716 | 4,26E-22 | 3,97E-20 | |
| ENSGALG00000007340 | 53,630639 | 2,338672246 | 0,3529687 | 6,62572131 | 3,46E-11 | 1,01E-09 | |
| ENSGALG00000023934 | 52,5759497 | 2,337998666 | 0,35383511 | 6,60759382 | 3,91E-11 | 1,12E-09 | |
| ENSGALG00000028341 | 324,387866 | 2,327426751 | 0,23601987 | 9,86114768 | 6,13E-23 | 6,23E-21 | |
| ENSGALG00000002260 | 279,413842 | 2,326075861 | 0,254913 | 9,12497939 | 7,17E-20 | 5,71E-18 | CISH |
| ENSGALG00000006402 | 199,281869 | 2,315345967 | 0,27769375 | 8,33776765 | 7,57E-17 | 4,56E-15 | |
| ENSGALG00000008709 | 98,4372913 | 2,291854243 | 0,32683951 | 7,0121701 | 2,35E-12 | 8,17E-11 | MCOLN3 |
| ENSGALG00000013006 | 357,98483 | 2,275189997 | 0,25794845 | 8,8203283 | 1,14E-18 | 8,05E-17 | ROPN1L |
| ENSGALG00000006992 | 182,223575 | 2,270300547 | 0,29844555 | 7,60708468 | 2,80E-14 | 1,27E-12 | MMP9 |
| ENSGALG00000011113 | 527,000372 | 2,26759493 | 0,2713629 | 8,35631896 | 6,47E-17 | 3,93E-15 | SGIP1 |
| ENSGALG00000023577 | 321,7197 | 2,257832359 | 0,26699218 | 8,45654861 | 2,75E-17 | 1,74E-15 | |
| ENSGALG00000016852 | 56,0309211 | 2,256531595 | 0,35169224 | 6,41621099 | 1,40E-10 | 3,74E-09 | |
| ENSGALG00000027617 | 219,458041 | 2,255993416 | 0,28513386 | 7,91205035 | 2,53E-15 | 1,25E-13 | |
| ENSGALG00000003677 | 113,607816 | 2,250263529 | 0,34495968 | 6,52326543 | 6,88E-11 | 1,92E-09 | NTNG2 |
| ENSGALG00000023742 | 249,604133 | 2,24719921 | 0,28204891 | 7,96740977 | 1,62E-15 | 8,26E-14 | |
| ENSGALG00000001589 | 111,257666 | 2,246033051 | 0,34415728 | 6,52618204 | 6,75E-11 | 1,89E-09 | |
| ENSGALG00000016369 | 952,122279 | 2,242110682 | 0,1770401 | 12,6644225 | 9,31E-37 | 2,54E-34 | |
| ENSGALG00000027703 | 404,868805 | 2,238234147 | 0,25355456 | 8,82742624 | 1,07E-18 | 7,64E-17 | |
| ENSGALG00000003389 | 248,600828 | 2,230534118 | 0,27052364 | 8,24524646 | 1,65E-16 | 9,51E-15 | |
| ENSGALG00000015244 | 123,748282 | 2,228224412 | 0,32046392 | 6,95312109 | 3,57E-12 | 1,20E-10 | TBX19 |
| ENSGALG00000001857 | 255,91037 | 2,203449137 | 0,25950427 | 8,49099387 | 2,05E-17 | 1,31E-15 | FAM132A |
| ENSGALG00000019861 | 47,6042668 | 2,196772492 | 0,35394524 | 6,2065321 | 5,42E-10 | 1,33E-08 | |
| ENSGALG00000008514 | 959,211971 | 2,19329695 | 0,17694652 | 12,3952535 | 2,77E-35 | 6,86E-33 | DLL4 |
| ENSGALG00000021714 | 88,4059279 | 2,185372855 | 0,33433945 | 6,53638944 | 6,30E-11 | 1,78E-09 | |
| ENSGALG00000009499 | 136,985306 | 2,185164296 | 0,31520463 | 6,93252602 | 4,13E-12 | 1,37E-10 | NPY5R |
| ENSGALG00000026467 | 67,1746865 | 2,184133106 | 0,34748728 | 6,28550519 | 3,27E-10 | 8,22E-09 | |
| ENSGALG00000027532 | 181,565157 | 2,181261094 | 0,33533315 | 6,50475832 | 7,78E-11 | 2,16E-09 | |
| ENSGALG00000002817 | 43,7219895 | 2,174112777 | 0,35312942 | 6,15670254 | 7,43E-10 | 1,79E-08 | |
| ENSGALG00000028339 | 572,220786 | 2,143746023 | 0,25356336 | 8,45447874 | 2,80E-17 | 1,76E-15 | |
| ENSGALG00000003548 | 717,979314 | 2,131798574 | 0,1846233 | 11,546747 | 7,67E-31 | 1,43E-28 | BAIAP2L1 |
| ENSGALG00000029168 | 574,122828 | 2,125452005 | 0,26338005 | 8,06990504 | 7,04E-16 | 3,85E-14 | TMEM27 |
| ENSGALG00000005062 | 695,377154 | 2,112402161 | 0,18917938 | 11,1661333 | 5,97E-29 | 9,35E-27 | FoxN4 |
| ENSGALG00000001879 | 730,682744 | 2,106018407 | 0,18389159 | 11,4524996 | 2,28E-30 | 4,04E-28 | |
| ENSGALG00000028551 | 2508,94496 | 2,105237568 | 0,17723341 | 11,8783337 | 1,53E-32 | 3,32E-30 | |
| ENSGALG00000001696 | 267,698482 | 2,102610472 | 0,25640045 | 8,20049448 | 2,39E-16 | 1,36E-14 | SAG |
| ENSGALG00000008275 | 48,5714847 | 2,098791425 | 0,35395495 | 5,92954394 | 3,04E-09 | 6,67E-08 | |
| ENSGALG00000021567 | 284,86115 | 2,081191963 | 0,24661499 | 8,43903259 | 3,20E-17 | 1,99E-15 | EPHA8 |
| ENSGALG00000001874 | 67,2449867 | 2,080511409 | 0,35389195 | 5,87894534 | 4,13E-09 | 8,88E-08 | |
| ENSGALG00000028654 | 92,9286403 | 2,074935686 | 0,34576621 | 6,00097882 | 1,96E-09 | 4,45E-08 | |
| ENSGALG00000006260 | 56,9564609 | 2,062296513 | 0,3527945 | 5,84560272 | 5,05E-09 | 1,06E-07 | KLHL30 |
| ENSGALG00000007570 | 256,243886 | 2,055518081 | 0,25713147 | 7,99403557 | 1,31E-15 | 6,73E-14 | |
| ENSGALG00000019527 | 38,8970283 | 2,052781357 | 0,35219468 | 5,82854162 | 5,59E-09 | 1,16E-07 | SLC6A19 |
| ENSGALG00000000656 | 95,7621403 | 2,036711968 | 0,34147678 | 5,96442297 | 2,46E-09 | 5,51E-08 | |
| ENSGALG00000007104 | 125,617756 | 2,032418742 | 0,32005632 | 6,35019094 | 2,15E-10 | 5,59E-09 | |
| ENSGALG00000026541 | 175,990929 | 2,028653787 | 0,29769667 | 6,81449952 | 9,46E-12 | 3,00E-10 | |
| ENSGALG00000014203 | 138,123283 | 2,025939839 | 0,30058823 | 6,7399175 | 1,58E-11 | 4,87E-10 | PARVG |
| ENSGALG00000020195 | 143,592197 | 2,021370929 | 0,30337177 | 6,66301589 | 2,68E-11 | 7,92E-10 | |
| ENSGALG00000012204 | 102,979277 | 2,019423041 | 0,32821004 | 6,1528374 | 7,61E-10 | 1,83E-08 | |
| ENSGALG00000005797 | 87,7942641 | 2,014484064 | 0,33418994 | 6,02796146 | 1,66E-09 | 3,83E-08 | COL20A1 |
| ENSGALG00000000437 | 37,0932943 | 2,013731373 | 0,35182907 | 5,72360709 | 1,04E-08 | 2,04E-07 | |
| ENSGALG00000002931 | 53,6926993 | 2,013598707 | 0,3534025 | 5,69774883 | 1,21E-08 | 2,37E-07 | |
| ENSGALG00000006273 | 107,39879 | 2,011386223 | 0,32297819 | 6,22762252 | 4,74E-10 | 1,16E-08 | MYLK2 |
| ENSGALG00000000722 | 46,5643306 | 2,006522785 | 0,35375492 | 5,67207026 | 1,41E-08 | 2,72E-07 | |
| ENSGALG00000028147 | 229,824659 | 2,001096858 | 0,26643674 | 7,51058915 | 5,89E-14 | 2,49E-12 | CTSK |
| ENSGALG00000004248 | 56,9012525 | 1,998749091 | 0,35206496 | 5,67721676 | 1,37E-08 | 2,65E-07 | TACR2 |
| ENSGALG00000003075 | 93,0788327 | 1,997443704 | 0,3325728 | 6,0060344 | 1,90E-09 | 4,34E-08 | SLC34A1 |
| ENSGALG00000001571 | 57,1169097 | 1,994083312 | 0,35177003 | 5,66871286 | 1,44E-08 | 2,77E-07 | MYO1F |
| ENSGALG00000010458 | 184,378204 | 1,990178775 | 0,28021881 | 7,10223132 | 1,23E-12 | 4,45E-11 | |
| ENSGALG00000011889 | 230,923015 | 1,988400853 | 0,26765629 | 7,42893376 | 1,09E-13 | 4,48E-12 | PRKCH |

| | | | | | | | |
|---|---|---|---|---|---|---|---|
| ENSGALG00000010924 | 209,968742 | 1,986898996 | 0,28742319 | 6,91279987 | 4,75E-12 | 1,56E-10 | cNFI-A |
| ENSGALG00000015846 | 313,730622 | 1,982793598 | 0,24658548 | 8,04099914 | 8,91E-16 | 4,78E-14 | SNAP91 |
| ENSGALG00000006835 | 112,628775 | 1,974805269 | 0,3171624 | 6,22647971 | 4,77E-10 | 1,17E-08 | TNNC2 |
| ENSGALG00000013253 | 157,862049 | 1,972349113 | 0,31981557 | 6,16714546 | 6,95E-10 | 1,68E-08 | INSRR |
| ENSGALG00000009302 | 367,162682 | 1,971912015 | 0,28798838 | 6,84719294 | 7,53E-12 | 2,42E-10 | EMX2 |
| ENSGALG00000004414 | 101,957841 | 1,959564216 | 0,32123225 | 6,10014788 | 1,06E-09 | 2,50E-08 | |
| ENSGALG00000009489 | 127,740493 | 1,956908773 | 0,32047595 | 6,1062578 | 1,02E-09 | 2,42E-08 | |
| ENSGALG00000005822 | 147,056462 | 1,951526015 | 0,30270273 | 6,44700496 | 1,14E-10 | 3,07E-09 | KCNQ2 |
| ENSGALG00000027147 | 228,058252 | 1,950896881 | 0,28014131 | 6,96397433 | 3,31E-12 | 1,13E-10 | |
| ENSGALG00000021869 | 645,941994 | 1,950468731 | 0,24355446 | 8,00834733 | 1,16E-15 | 6,04E-14 | PODN |
| ENSGALG00000028975 | 532,930635 | 1,945491073 | 0,20584343 | 9,45131484 | 3,35E-21 | 2,90E-19 | |
| ENSGALG00000007666 | 61,9703125 | 1,945055405 | 0,35143193 | 5,53465757 | 3,12E-08 | 5,56E-07 | KCHIP2 |
| ENSGALG00000021656 | 37,5118215 | 1,943986445 | 0,3507973 | 5,5416231 | 3,00E-08 | 5,37E-07 | PTPN22 |
| ENSGALG00000008188 | 164,942808 | 1,939910796 | 0,29070864 | 6,67304152 | 2,51E-11 | 7,46E-10 | TRIM36 |
| ENSGALG00000010362 | 56,1305671 | 1,930559856 | 0,35381555 | 5,45640199 | 4,86E-08 | 8,33E-07 | |
| ENSGALG00000000560 | 58,7581666 | 1,925570368 | 0,35333497 | 5,44970217 | 5,05E-08 | 8,61E-07 | |
| ENSGALG00000015021 | 44,6818444 | 1,923323022 | 0,35395775 | 5,43376434 | 5,52E-08 | 9,36E-07 | TMEM174 |
| ENSGALG00000015551 | 1931,12818 | 1,921428632 | 0,17206371 | 11,1669603 | 5,92E-29 | 9,35E-27 | GRIN3A |
| ENSGALG00000010867 | 80,2669926 | 1,919026004 | 0,34078038 | 5,63126911 | 1,79E-08 | 3,37E-07 | TACSTD2 |
| ENSGALG00000001004 | 271,271449 | 1,914258624 | 0,25960686 | 7,37368273 | 1,66E-13 | 6,69E-12 | |
| ENSGALG00000011347 | 48,911798 | 1,909599526 | 0,35390611 | 5,39578011 | 6,82E-08 | 1,14E-06 | IHH |
| ENSGALG00000006413 | 117,165789 | 1,90767902 | 0,31128952 | 6,12831113 | 8,88E-10 | 2,11E-08 | CEMIP |
| ENSGALG00000003836 | 138,248419 | 1,907173269 | 0,34505436 | 5,52716754 | 3,25E-08 | 5,78E-07 | |
| ENSGALG00000007345 | 111,972561 | 1,905774604 | 0,32456789 | 5,87172866 | 4,31E-09 | 9,20E-08 | |
| ENSGALG00000008701 | 97,1365525 | 1,905747917 | 0,3400514 | 5,60429376 | 2,09E-08 | 3,87E-07 | XDH |
| ENSGALG00000010853 | 484,768782 | 1,904256394 | 0,21726773 | 8,76456169 | 1,88E-18 | 1,29E-16 | C8B |
| ENSGALG00000000221 | 51,6768021 | 1,900065815 | 0,35396003 | 5,3680237 | 7,96E-08 | 1,31E-06 | |
| ENSGALG00000001617 | 1078,32932 | 1,898502237 | 0,21231379 | 8,9419639 | 3,82E-19 | 2,86E-17 | CAMKK1 |
| ENSGALG00000019290 | 78,1238371 | 1,892925943 | 0,34969839 | 5,41302448 | 6,20E-08 | 1,04E-06 | SLC6A12 |
| ENSGALG00000009265 | 88,1743252 | 1,891791729 | 0,34591457 | 5,46895654 | 4,53E-08 | 7,80E-07 | KCNK18 |
| ENSGALG00000003923 | 2335,33874 | 1,889033194 | 0,15037048 | 12,5625268 | 3,39E-36 | 8,88E-34 | COL6A3 |
| ENSGALG00000008995 | 104,546365 | 1,882652598 | 0,3282959 | 5,73462107 | 9,77E-09 | 1,93E-07 | |
| ENSGALG00000000544 | 347,726242 | 1,877518925 | 0,23287565 | 8,06232383 | 7,49E-16 | 4,08E-14 | RAB44 |
| ENSGALG00000007706 | 86,703491 | 1,877381913 | 0,33476816 | 5,60800615 | 2,05E-08 | 3,82E-07 | FGF8 |
| ENSGALG00000015347 | 73,1732063 | 1,876981559 | 0,34452788 | 5,44798159 | 5,09E-08 | 8,66E-07 | ZPLD1 |
| ENSGALG00000009078 | 124,066106 | 1,868595491 | 0,33382944 | 5,59745563 | 2,18E-08 | 4,01E-07 | GALNT14 |
| ENSGALG00000000899 | 303,594278 | 1,867380532 | 0,25292797 | 7,38305256 | 1,55E-13 | 6,29E-12 | |
| ENSGALG00000023459 | 62,5233629 | 1,863475253 | 0,35063414 | 5,31458595 | 1,07E-07 | 1,71E-06 | |
| ENSGALG00000013953 | 153,510812 | 1,853929495 | 0,31615798 | 5,86393381 | 4,52E-09 | 9,60E-08 | |
| ENSGALG00000020316 | 66,9879377 | 1,852228829 | 0,34370538 | 5,38900158 | 7,09E-08 | 1,17E-06 | IL13RA2 |
| ENSGALG00000014770 | 4689,68364 | 1,850573228 | 0,19171766 | 9,65259643 | 4,79E-22 | 4,41E-20 | |
| ENSGALG00000015406 | 32,8242769 | 1,850052547 | 0,34954162 | 5,29279624 | 1,20E-07 | 1,90E-06 | |
| ENSGALG00000007047 | 78,9873557 | 1,844593336 | 0,3380656 | 5,4563178 | 4,86E-08 | 8,33E-07 | GAL |
| ENSGALG00000011421 | 155,787271 | 1,840566258 | 0,29419428 | 6,25629526 | 3,94E-10 | 9,81E-09 | HAL |
| ENSGALG00000003751 | 52,9480261 | 1,835104614 | 0,35380373 | 5,18678703 | 2,14E-07 | 3,22E-06 | |
| ENSGALG00000021693 | 69,3129231 | 1,830329106 | 0,34305164 | 5,33543326 | 9,53E-08 | 1,54E-06 | |
| ENSGALG00000005634 | 92,2321464 | 1,826798031 | 0,32741142 | 5,57951828 | 2,41E-08 | 4,40E-07 | CRYBA4 |
| ENSGALG00000005644 | 54,6398918 | 1,819377424 | 0,3504577 | 5,19143234 | 2,09E-07 | 3,15E-06 | MYO1G |
| ENSGALG00000029006 | 89,8196778 | 1,803532169 | 0,3296621 | 5,47085084 | 4,48E-08 | 7,73E-07 | SYPL1 |
| ENSGALG00000010126 | 116,195874 | 1,799221778 | 0,32988987 | 5,4540074 | 4,92E-08 | 8,42E-07 | |
| ENSGALG00000019361 | 44,228613 | 1,792121861 | 0,35385967 | 5,06449882 | 4,09E-07 | 5,78E-06 | FAM19A2 |
| ENSGALG00000020982 | 53,8722608 | 1,789263062 | 0,35132044 | 5,09296598 | 3,53E-07 | 5,05E-06 | |
| ENSGALG00000004841 | 236,709128 | 1,788099107 | 0,27635881 | 6,47020836 | 9,79E-11 | 2,66E-09 | APLNR |
| ENSGALG00000008845 | 211,090699 | 1,784489653 | 0,26979829 | 6,61416227 | 3,74E-11 | 1,08E-09 | HAO1 |
| ENSGALG00000023354 | 87,3213221 | 1,782015396 | 0,34472234 | 5,16942237 | 2,35E-07 | 3,49E-06 | FSCN2 |
| ENSGALG00000010364 | 75,7123588 | 1,777216343 | 0,34965008 | 5,08284262 | 3,72E-07 | 5,29E-06 | |
| ENSGALG00000027090 | 122,235387 | 1,772101833 | 0,32386305 | 5,47176288 | 4,46E-08 | 7,70E-07 | |
| ENSGALG00000005043 | 168,034499 | 1,771501032 | 0,28869817 | 6,13617001 | 8,45E-10 | 2,02E-08 | ACACB |
| ENSGALG00000005747 | 39,0007429 | 1,767653479 | 0,35360417 | 4,99896104 | 5,76E-07 | 7,89E-06 | |
| ENSGALG00000014252 | 107,135531 | 1,765576737 | 0,34633901 | 5,09782814 | 3,44E-07 | 4,94E-06 | |
| ENSGALG00000006007 | 80,5203103 | 1,763247242 | 0,34316546 | 5,13818391 | 2,77E-07 | 4,06E-06 | SAMD10 |
| ENSGALG00000000630 | 44,9860229 | 1,762139369 | 0,35277196 | 4,99512312 | 5,88E-07 | 8,02E-06 | HAS3 |
| ENSGALG00000013971 | 623,626799 | 1,754983652 | 0,22177863 | 7,91322269 | 2,51E-15 | 1,24E-13 | SGK1 |
| ENSGALG00000025804 | 57,8298229 | 1,753854275 | 0,34868015 | 5,02998031 | 4,91E-07 | 6,84E-06 | CLCN2 |

| | | | | | | | |
|---|---|---|---|---|---|---|---|
| ENSGALG00000016233 | 282,390757 | 1,751740847 | 0,25916092 | 6,75927863 | 1,39E-11 | 4,28E-10 | |
| ENSGALG00000000299 | 146,622524 | 1,750555222 | 0,31737522 | 5,5157274 | 3,47E-08 | 6,13E-07 | PKP1 |
| ENSGALG00000015154 | 107,544575 | 1,749162012 | 0,31926267 | 5,47875512 | 4,28E-08 | 7,43E-07 | TRPM6 |
| ENSGALG00000028801 | 326,04496 | 1,749152851 | 0,23171525 | 7,54871702 | 4,40E-14 | 1,90E-12 | YPEL2 |
| ENSGALG00000003103 | 147,012955 | 1,745350116 | 0,2934725 | 5,94723559 | 2,73E-09 | 6,09E-08 | MST1R |
| ENSGALG00000010408 | 48,6937618 | 1,74244975 | 0,35251743 | 4,94287544 | 7,70E-07 | 1,02E-05 | |
| ENSGALG00000016580 | 156,416853 | 1,740943756 | 0,31375774 | 5,54868778 | 2,88E-08 | 5,20E-07 | SCARA5 |
| ENSGALG00000020779 | 287,434053 | 1,737619419 | 0,24470316 | 7,10092768 | 1,24E-12 | 4,48E-11 | |
| ENSGALG00000015544 | 64,576519 | 1,719200832 | 0,35119647 | 4,89526802 | 9,82E-07 | 1,28E-05 | ALDOB |
| ENSGALG00000015305 | 121,835301 | 1,715527315 | 0,30924605 | 5,547451 | 2,90E-08 | 5,21E-07 | NR2E1 |
| ENSGALG00000008926 | 1594,75407 | 1,713961304 | 0,15633781 | 10,9631913 | 5,74E-28 | 8,69E-26 | CADPS2 |
| ENSGALG00000002957 | 403,400084 | 1,713056332 | 0,23618065 | 7,25316116 | 4,07E-13 | 1,57E-11 | SLC12A3 |
| ENSGALG00000017212 | 76,5473495 | 1,710911672 | 0,34240676 | 4,9967228 | 5,83E-07 | 7,97E-06 | GPR83 |
| ENSGALG00000004827 | 60,764569 | 1,70800582 | 0,34662638 | 4,92751255 | 8,33E-07 | 1,10E-05 | |
| ENSGALG00000003242 | 570,220114 | 1,707949225 | 0,21013146 | 8,12800356 | 4,36E-16 | 2,42E-14 | STAR |
| ENSGALG00000001912 | 146,494381 | 1,699528733 | 0,30128418 | 5,64094911 | 1,69E-08 | 3,21E-07 | |
| ENSGALG00000003003 | 42,6697686 | 1,694427776 | 0,35203153 | 4,81328409 | 1,48E-06 | 1,86E-05 | CCL20 |
| ENSGALG00000016261 | 158,706658 | 1,693178751 | 0,31450243 | 5,38367466 | 7,30E-08 | 1,20E-06 | CYBB |
| ENSGALG00000011007 | 1150,32901 | 1,688733669 | 0,19077647 | 8,85189717 | 8,60E-19 | 6,20E-17 | OSBPL3 |
| ENSGALG00000015117 | 149,514374 | 1,688648354 | 0,29548541 | 5,71482827 | 1,10E-08 | 2,15E-07 | DRD3 |
| ENSGALG00000012429 | 6013,8279 | 1,680823476 | 0,15201043 | 11,0572901 | 2,02E-28 | 3,09E-26 | BMP4 |
| ENSGALG00000000458 | 427,66514 | 1,679795969 | 0,21744886 | 7,72501627 | 1,12E-14 | 5,20E-13 | TMPRSS9 |
| ENSGALG00000011622 | 28,6515601 | 1,67904306 | 0,34603089 | 4,85229247 | 1,22E-06 | 1,56E-05 | |
| ENSGALG00000016572 | 77,2006942 | 1,678816771 | 0,34344316 | 4,88819394 | 1,02E-06 | 1,32E-05 | |
| ENSGALG00000001084 | 353,564869 | 1,677708871 | 0,24186887 | 6,93643975 | 4,02E-12 | 1,34E-10 | ATP8B3 |
| ENSGALG00000026473 | 1522,15108 | 1,674638021 | 0,21127677 | 7,92627606 | 2,26E-15 | 1,13E-13 | |
| ENSGALG00000008178 | 111,711666 | 1,67186128 | 0,32258264 | 5,18273795 | 2,19E-07 | 3,28E-06 | |
| ENSGALG00000005140 | 240,214893 | 1,671709253 | 0,25797704 | 6,48006996 | 9,17E-11 | 2,52E-09 | RRAD |
| ENSGALG00000001234 | 40,8626534 | 1,668621796 | 0,35391846 | 4,7147069 | 2,42E-06 | 2,87E-05 | CETP |
| ENSGALG00000006514 | 1126,00554 | 1,665636549 | 0,17514671 | 9,50995042 | 1,91E-21 | 1,69E-19 | |
| ENSGALG00000004936 | 57,472904 | 1,658043102 | 0,34946885 | 4,74446601 | 2,09E-06 | 2,51E-05 | GLRA4 |
| ENSGALG00000028174 | 599,327274 | 1,656109672 | 0,24081026 | 6,87723897 | 6,10E-12 | 1,98E-10 | |
| ENSGALG00000011882 | 446,567699 | 1,655648124 | 0,21164389 | 7,82280161 | 5,17E-15 | 2,44E-13 | |
| ENSGALG00000020143 | 148,781899 | 1,654526791 | 0,32189845 | 5,13990289 | 2,75E-07 | 4,02E-06 | GABRG1 |
| ENSGALG00000027314 | 27,9768099 | 1,648583002 | 0,34526179 | 4,77487828 | 1,80E-06 | 2,19E-05 | SAXO1 |
| ENSGALG00000019019 | 89,6714541 | 1,647447024 | 0,34390312 | 4,79043927 | 1,66E-06 | 2,05E-05 | NTRK1 |
| ENSGALG00000026161 | 123,209261 | 1,647025641 | 0,3125278 | 5,27001315 | 1,36E-07 | 2,13E-06 | |
| ENSGALG00000024298 | 32,0306965 | 1,646495761 | 0,34885964 | 4,71965109 | 2,36E-06 | 2,81E-05 | ADAMTSL5 |
| ENSGALG00000004394 | 167,659182 | 1,640660466 | 0,28374074 | 5,7822521 | 7,37E-09 | 1,49E-07 | PCBP3 |
| ENSGALG00000016262 | 396,403367 | 1,640510999 | 0,22878691 | 7,17047578 | 7,47E-13 | 2,77E-11 | XK |
| ENSGALG00000015905 | 84,004407 | 1,632452898 | 0,3358609 | 4,86050296 | 1,17E-06 | 1,51E-05 | |
| ENSGALG00000005293 | 115,927539 | 1,631397185 | 0,32721191 | 4,98575121 | 6,17E-07 | 8,36E-06 | |
| ENSGALG00000025946 | 137,584784 | 1,630278266 | 0,31187717 | 5,22730883 | 1,72E-07 | 2,64E-06 | MOB3C |
| ENSGALG00000011806 | 33,3533195 | 1,625032877 | 0,34750572 | 4,67627665 | 2,92E-06 | 3,39E-05 | |
| ENSGALG00000010721 | 516,489319 | 1,61713725 | 0,19825254 | 8,15695606 | 3,44E-16 | 1,92E-14 | |
| ENSGALG00000008133 | 122,891283 | 1,61551597 | 0,32520429 | 4,96769575 | 6,78E-07 | 9,10E-06 | AS3MT |
| ENSGALG00000000820 | 127,586167 | 1,614544959 | 0,31856994 | 5,06810204 | 4,02E-07 | 5,68E-06 | HTR1D |
| ENSGALG00000002832 | 501,681621 | 1,610990532 | 0,21276938 | 7,57153378 | 3,69E-14 | 1,62E-12 | IFI35 |
| ENSGALG00000006606 | 3620,8517 | 1,610976489 | 0,12247757 | 13,1532366 | 1,63E-39 | 5,29E-37 | |
| ENSGALG00000011200 | 907,828768 | 1,609360966 | 0,24666698 | 6,52442818 | 6,83E-11 | 1,91E-09 | THBS2 |
| ENSGALG00000007980 | 258,344877 | 1,607363167 | 0,27462432 | 5,85295284 | 4,83E-09 | 1,02E-07 | KCNG1 |
| ENSGALG00000001062 | 680,579051 | 1,606326071 | 0,19183595 | 8,37343595 | 5,60E-17 | 3,43E-15 | GGT7 |
| ENSGALG00000003106 | 661,981272 | 1,604428011 | 0,22228284 | 7,21795705 | 5,28E-13 | 2,01E-11 | TOX2 |
| ENSGALG00000007815 | 333,865811 | 1,603027021 | 0,27971291 | 5,73097251 | 9,99E-09 | 1,96E-07 | ANKK1 |
| ENSGALG00000015366 | 80,0727703 | 1,602600172 | 0,34326453 | 4,66870313 | 3,03E-06 | 3,52E-05 | |
| ENSGALG00000005725 | 91,1085716 | 1,601523904 | 0,35202702 | 4,54943466 | 5,38E-06 | 5,90E-05 | CSF1R |
| ENSGALG00000022644 | 26,8742275 | 1,595235245 | 0,34379409 | 4,64008918 | 3,48E-06 | 3,98E-05 | |
| ENSGALG00000007759 | 856,866355 | 1,594985498 | 0,19890567 | 8,01880348 | 1,07E-15 | 5,57E-14 | |
| ENSGALG00000008830 | 781,049383 | 1,591806155 | 0,21140245 | 7,52974324 | 5,08E-14 | 2,17E-12 | |
| ENSGALG00000006412 | 53,0587145 | 1,591456503 | 0,35279252 | 4,51102674 | 6,45E-06 | 6,90E-05 | VGLL1 |
| ENSGALG00000029174 | 5348,15015 | 1,589321167 | 0,12909013 | 12,3117174 | 7,83E-35 | 1,87E-32 | |
| ENSGALG00000006245 | 402,970032 | 1,589094556 | 0,22775101 | 6,97733278 | 3,01E-12 | 1,04E-10 | |
| ENSGALG00000026733 | 77,5005998 | 1,588089103 | 0,34066443 | 4,6617403 | 3,14E-06 | 3,62E-05 | FFAR4 |
| ENSGALG00000016138 | 93,2512783 | 1,581494384 | 0,32966262 | 4,79731185 | 1,61E-06 | 2,00E-05 | DSCAM |

| | | | | | | | |
|---|---|---|---|---|---|---|---|
| ENSGALG00000016498 | 94,7917055 | 1,580375213 | 0,32949342 | 4,7963786 | 1,62E-06 | 2,01E-05 | MFSD2B |
| ENSGALG00000011343 | 80,565753 | 1,579403764 | 0,33480013 | 4,71745272 | 2,39E-06 | 2,83E-05 | CNPPD1 |
| ENSGALG00000000733 | 1063,13098 | 1,578595816 | 0,19876941 | 7,94184501 | 1,99E-15 | 1,00E-13 | |
| ENSGALG00000028175 | 72,9348346 | 1,575180638 | 0,35396003 | 4,45016529 | 8,58E-06 | 9,01E-05 | GJA9 |
| ENSGALG00000015293 | 333,004659 | 1,574970296 | 0,24444617 | 6,4430147 | 1,17E-10 | 3,15E-09 | TMEM45A |
| ENSGALG00000012115 | 228,294388 | 1,572479232 | 0,27344667 | 5,75058837 | 8,89E-09 | 1,77E-07 | WT1 |
| ENSGALG00000004692 | 57,025385 | 1,559793633 | 0,35370154 | 4,40991479 | 1,03E-05 | 0,0001064 | |
| ENSGALG00000005958 | 558,635772 | 1,558583818 | 0,23299183 | 6,68943537 | 2,24E-11 | 6,70E-10 | TRIM66 |
| ENSGALG00000000826 | 97,953466 | 1,551558986 | 0,32368653 | 4,79339992 | 1,64E-06 | 2,03E-05 | |
| ENSGALG00000007690 | 2070,9973 | 1,54950396 | 0,17555383 | 8,82637523 | 1,08E-18 | 7,67E-17 | TFAP2C |
| ENSGALG00000006964 | 225,191012 | 1,546629935 | 0,29302707 | 5,27811285 | 1,31E-07 | 2,05E-06 | |
| ENSGALG00000005967 | 735,636675 | 1,545091384 | 0,22167917 | 6,96994379 | 3,17E-12 | 1,09E-10 | |
| ENSGALG00000008440 | 429,095272 | 1,543692993 | 0,21647003 | 7,13120884 | 9,95E-13 | 3,64E-11 | CHST1 |
| ENSGALG00000001882 | 38,7098275 | 1,540579881 | 0,35211396 | 4,37523089 | 1,21E-05 | 0,00012243 | |
| ENSGALG00000015373 | 27,6017686 | 1,538791481 | 0,34451153 | 4,46658922 | 7,95E-06 | 8,37E-05 | PDE6B |
| ENSGALG00000011577 | 399,56293 | 1,531134988 | 0,21728602 | 7,0466337 | 1,83E-12 | 6,48E-11 | CNTNAP5 |
| ENSGALG00000025774 | 5724,96959 | 1,527733684 | 0,12704021 | 12,0255916 | 2,61E-33 | 5,82E-31 | |
| ENSGALG00000005473 | 107,686637 | 1,527501434 | 0,32073027 | 4,76257338 | 1,91E-06 | 2,31E-05 | |
| ENSGALG00000000479 | 244,695616 | 1,527264642 | 0,25520876 | 5,984374 | 2,17E-09 | 4,90E-08 | |
| ENSGALG00000011071 | 944,33324 | 1,527069671 | 0,16947196 | 9,01075121 | 2,05E-19 | 1,57E-17 | |
| ENSGALG00000020459 | 25,5154221 | 1,526419374 | 0,34167333 | 4,46748172 | 7,91E-06 | 8,34E-05 | |
| ENSGALG00000014267 | 99,3352174 | 1,525149591 | 0,31972846 | 4,77014025 | 1,84E-06 | 2,24E-05 | RBM47 |
| ENSGALG00000007803 | 165,807287 | 1,522432144 | 0,29566688 | 5,14914667 | 2,62E-07 | 3,86E-06 | |
| ENSGALG00000001569 | 88,7153734 | 1,518412601 | 0,34222784 | 4,43684706 | 9,13E-06 | 9,51E-05 | RHOC |
| ENSGALG00000020745 | 195,274039 | 1,515524715 | 0,27430933 | 5,52487479 | 3,30E-08 | 5,85E-07 | RHO |
| ENSGALG00000011227 | 778,979814 | 1,515331444 | 0,18180249 | 8,3350423 | 7,75E-17 | 4,65E-15 | CDS1 |
| ENSGALG00000007594 | 126,922974 | 1,512071087 | 0,30201682 | 5,00657906 | 5,54E-07 | 7,63E-06 | ASB12 |
| ENSGALG00000002314 | 85,1314877 | 1,511687395 | 0,33330967 | 4,53538412 | 5,75E-06 | 6,23E-05 | CDHR1 |
| ENSGALG00000002353 | 221,898313 | 1,511078773 | 0,26237238 | 5,75929059 | 8,45E-09 | 1,69E-07 | C1orf216 |
| ENSGALG00000024397 | 46,3770198 | 1,510326295 | 0,3539532 | 4,2670226 | 1,98E-05 | 0,00018598 | |
| ENSGALG00000023866 | 254,336531 | 1,510055693 | 0,28025871 | 5,38807762 | 7,12E-08 | 1,18E-06 | |
| ENSGALG00000026089 | 41,8112969 | 1,509966763 | 0,35044188 | 4,30875091 | 1,64E-05 | 0,00015808 | SLC26A6 |
| ENSGALG00000028386 | 982,725203 | 1,507238343 | 0,16997711 | 8,86730176 | 7,49E-19 | 5,49E-17 | |
| ENSGALG00000006479 | 34,3468527 | 1,50588664 | 0,34819791 | 4,32480087 | 1,53E-05 | 0,00014837 | CCM2L |
| ENSGALG00000005297 | 304,659703 | 1,505251197 | 0,24557254 | 6,12955833 | 8,81E-10 | 2,10E-08 | HHATL |
| ENSGALG00000004448 | 547,114848 | 1,504790559 | 0,23230258 | 6,4777179 | 9,31E-11 | 2,55E-09 | BRINP2 |
| ENSGALG00000026983 | 590,103546 | 1,50209184 | 0,21370636 | 7,02876535 | 2,08E-12 | 7,31E-11 | CHRD |
| ENSGALG00000024264 | 40,6812127 | 1,501303133 | 0,34955904 | 4,2948485 | 1,75E-05 | 0,00016679 | |
| ENSGALG00000028347 | 76,9725475 | 1,501152953 | 0,34262878 | 4,38128098 | 1,18E-05 | 0,00011917 | SDCBP2 |
| ENSGALG00000016556 | 93,5327309 | 1,500831481 | 0,32530567 | 4,6136038 | 3,96E-06 | 4,47E-05 | |
| ENSGALG00000008424 | 31,5559973 | 1,500581439 | 0,34950735 | 4,29341889 | 1,76E-05 | 0,00016751 | CALML3 |
| ENSGALG00000016287 | 253,693238 | 1,497802744 | 0,250661 | 5,9754121 | 2,30E-09 | 5,17E-08 | |
| ENSGALG00000028069 | 276,234024 | 1,497716991 | 0,25564361 | 5,858613 | 4,67E-09 | 9,87E-08 | WNT16 |
| ENSGALG00000015410 | 210,477674 | 1,497100835 | 0,26827858 | 5,58039641 | 2,40E-08 | 4,38E-07 | BVES |
| ENSGALG00000014074 | 676,433259 | 1,49566297 | 0,19193599 | 7,79250911 | 6,57E-15 | 3,08E-13 | |
| ENSGALG00000003713 | 62,0071346 | 1,494159857 | 0,34951056 | 4,2750063 | 1,91E-05 | 0,00018034 | |
| ENSGALG00000007409 | 87,0801504 | 1,493981037 | 0,32899137 | 4,54109494 | 5,60E-06 | 6,09E-05 | SCN4B |
| ENSGALG00000028823 | 172,118142 | 1,491796906 | 0,2989787 | 4,98964273 | 6,05E-07 | 8,21E-06 | |
| ENSGALG00000026585 | 53,5711009 | 1,487448109 | 0,3507492 | 4,24077402 | 2,23E-05 | 0,00020575 | |
| ENSGALG00000027548 | 35,7639328 | 1,48608799 | 0,3490327 | 4,25773286 | 2,07E-05 | 0,00019271 | |
| ENSGALG00000028770 | 169,214437 | 1,483894729 | 0,30979476 | 4,78992848 | 1,67E-06 | 2,06E-05 | INHBB |
| ENSGALG00000015237 | 27,2937688 | 1,481694783 | 0,34447168 | 4,30135446 | 1,70E-05 | 0,00016265 | PPIL6 |
| ENSGALG00000015084 | 1284,76928 | 1,480304896 | 0,18534154 | 7,98690291 | 1,38E-15 | 7,11E-14 | IGSF11 |
| ENSGALG00000027357 | 83,3635346 | 1,478068348 | 0,33426191 | 4,4218869 | 9,78E-06 | 0,000101 | |
| ENSGALG00000007908 | 523,875729 | 1,477721596 | 0,21194377 | 6,97223428 | 3,12E-12 | 1,08E-10 | |
| ENSGALG00000008888 | 216,095633 | 1,475198435 | 0,26634382 | 5,5386996 | 3,05E-08 | 5,45E-07 | |
| ENSGALG00000000984 | 31,631294 | 1,47391651 | 0,3504148 | 4,2062051 | 2,60E-05 | 0,00023462 | |
| ENSGALG00000007862 | 431,761835 | 1,47265082 | 0,23340463 | 6,30943289 | 2,80E-10 | 7,11E-09 | CNTN3 |
| ENSGALG00000016538 | 1671,27665 | 1,472170836 | 0,14445232 | 10,1913963 | 2,17E-24 | 2,59E-22 | RAI2 |
| ENSGALG00000002442 | 183,2009 | 1,464263461 | 0,28517222 | 5,13466379 | 2,83E-07 | 4,12E-06 | ANKS4B |
| ENSGALG00000010107 | 1860,62891 | 1,462273804 | 0,14178721 | 10,3131576 | 6,14E-25 | 7,75E-23 | |
| ENSGALG00000026677 | 510,78911 | 1,461550873 | 0,27216608 | 5,37006991 | 7,87E-08 | 1,29E-06 | F10 |
| ENSGALG00000000703 | 748,407891 | 1,459568013 | 0,22886918 | 6,37730267 | 1,80E-10 | 4,77E-09 | SLC45A3 |
| ENSGALG00000026069 | 33,9757189 | 1,459502097 | 0,34690094 | 4,20725901 | 2,58E-05 | 0,00023392 | |

| | | | | | | | |
|---|---|---|---|---|---|---|---|
| ENSGALG00000001319 | 188,5387 | 1,458388464 | 0,27459113 | 5,31112739 | 1,09E-07 | 1,73E-06 | CAMK1G |
| ENSGALG00000014297 | 511,060783 | 1,457150717 | 0,20851139 | 6,98835066 | 2,78E-12 | 9,64E-11 | IRF7 |
| ENSGALG00000004565 | 123,010215 | 1,456247274 | 0,33891637 | 4,29677471 | 1,73E-05 | 0,00016569 | |
| ENSGALG00000010024 | 105,10355 | 1,453185654 | 0,31957147 | 4,5472947 | 5,43E-06 | 5,94E-05 | |
| ENSGALG00000014941 | 729,891458 | 1,451697765 | 0,1864124 | 7,78756017 | 6,83E-15 | 3,20E-13 | |
| ENSGALG00000018878 | 40,6695287 | 1,44967545 | 0,35380851 | 4,09734482 | 4,18E-05 | 0,00035297 | |
| ENSGALG00000015128 | 44,3465543 | 1,44940013 | 0,35395981 | 4,09481559 | 4,23E-05 | 0,00035587 | AQP4 |
| ENSGALG00000006751 | 459,348637 | 1,449311054 | 0,21417443 | 6,76696594 | 1,32E-11 | 4,09E-10 | THY1 |
| ENSGALG00000017041 | 876,052013 | 1,44806939 | 0,20136843 | 7,19114421 | 6,43E-13 | 2,41E-11 | |
| ENSGALG00000007211 | 323,240319 | 1,44654086 | 0,24592568 | 5,88202449 | 4,05E-09 | 8,74E-08 | CDH22 |
| ENSGALG00000001727 | 728,893661 | 1,445847057 | 0,18026579 | 8,02064026 | 1,05E-15 | 5,52E-14 | LZTS1 |
| ENSGALG00000006239 | 29,3987017 | 1,443346989 | 0,34917352 | 4,13360955 | 3,57E-05 | 0,00030929 | |
| ENSGALG00000023395 | 151,597821 | 1,441830671 | 0,29992743 | 4,80726505 | 1,53E-06 | 1,91E-05 | PLIN1 |
| ENSGALG00000013640 | 91,8229663 | 1,440450208 | 0,32437446 | 4,44070167 | 8,97E-06 | 9,38E-05 | |
| ENSGALG00000014962 | 26,4645058 | 1,440003001 | 0,34319536 | 4,19586969 | 2,72E-05 | 0,00024358 | FAM26F |
| ENSGALG00000008267 | 52,9646856 | 1,439464257 | 0,35338111 | 4,07340464 | 4,63E-05 | 0,0003844 | IL5RA |
| ENSGALG00000011202 | 37,1408249 | 1,438867662 | 0,3522352 | 4,08496273 | 4,41E-05 | 0,00036755 | COLQ |
| ENSGALG00000012652 | 778,180225 | 1,438231617 | 0,2169215 | 6,63019384 | 3,35E-11 | 9,79E-10 | |
| ENSGALG00000001710 | 36,6972134 | 1,437606263 | 0,3472289 | 4,14022641 | 3,47E-05 | 0,00030127 | |
| ENSGALG00000014695 | 237,142454 | 1,436054042 | 0,28464919 | 5,0449961 | 4,54E-07 | 6,35E-06 | CLCN1 |
| ENSGALG00000024368 | 80,858185 | 1,435967902 | 0,33830768 | 4,24456188 | 2,19E-05 | 0,00020286 | |
| ENSGALG00000003678 | 3218,39181 | 1,429238494 | 0,12475534 | 11,4563312 | 2,19E-30 | 3,97E-28 | |
| ENSGALG00000012313 | 84,3836892 | 1,428353126 | 0,34635998 | 4,1238977 | 3,73E-05 | 0,00031938 | |
| ENSGALG00000000914 | 24,8181662 | 1,42659313 | 0,3378824 | 4,22215874 | 2,42E-05 | 0,00022111 | |
| ENSGALG00000006565 | 175,499575 | 1,424795357 | 0,32613752 | 4,36869506 | 1,25E-05 | 0,00012543 | |
| ENSGALG00000011888 | 40,65783 | 1,420709802 | 0,353895 | 4,01449525 | 5,96E-05 | 0,00047852 | |
| ENSGALG00000010213 | 97,7805158 | 1,420151903 | 0,3251042 | 4,36829761 | 1,25E-05 | 0,00012548 | |
| ENSGALG00000008901 | 121,607035 | 1,417468594 | 0,30853867 | 4,59413595 | 4,35E-06 | 4,87E-05 | FP |
| ENSGALG00000009596 | 108,17405 | 1,416089625 | 0,32143686 | 4,40549862 | 1,06E-05 | 0,00010813 | TMEM117 |
| ENSGALG00000014412 | 49,0629688 | 1,415817045 | 0,35185351 | 4,02388215 | 5,72E-05 | 0,00046283 | CSTA |
| ENSGALG00000000589 | 81,385408 | 1,415474788 | 0,33862107 | 4,18011435 | 2,91E-05 | 0,00025877 | |
| ENSGALG00000023834 | 58,0264688 | 1,413727582 | 0,35080504 | 4,02995233 | 5,58E-05 | 0,00045266 | |
| ENSGALG00000018998 | 220,946842 | 1,412120959 | 0,28106361 | 5,02420412 | 5,06E-07 | 7,03E-06 | |
| ENSGALG00000013081 | 81,6196368 | 1,411618895 | 0,34436277 | 4,09922035 | 4,15E-05 | 0,00035078 | VWC2 |
| ENSGALG00000027520 | 176,34242 | 1,409426755 | 0,294554 | 4,78495208 | 1,71E-06 | 2,10E-05 | |
| ENSGALG00000020423 | 91,6730687 | 1,409349068 | 0,34445303 | 4,09155661 | 4,28E-05 | 0,00035977 | |
| ENSGALG00000026957 | 482,71907 | 1,407844107 | 0,20929005 | 6,72676095 | 1,73E-11 | 5,28E-10 | SEMA4G |
| ENSGALG00000010486 | 221,460325 | 1,406315246 | 0,26026568 | 5,40338331 | 6,54E-08 | 1,10E-06 | |
| ENSGALG00000028936 | 318,091175 | 1,405167114 | 0,23410156 | 6,0023824 | 1,94E-09 | 4,43E-08 | |
| ENSGALG00000002637 | 48,9524533 | 1,404353552 | 0,35337987 | 3,97406211 | 7,07E-05 | 0,00055575 | RBP7 |
| ENSGALG00000026948 | 265,898108 | 1,40406892 | 0,28394262 | 4,94490374 | 7,62E-07 | 1,01E-05 | ADD2 |
| ENSGALG00000016782 | 106,880394 | 1,400781577 | 0,32848891 | 4,26431921 | 2,01E-05 | 0,00018802 | |
| ENSGALG00000015918 | 313,695882 | 1,399804901 | 0,23804553 | 5,88040816 | 4,09E-09 | 8,82E-08 | |
| ENSGALG00000011594 | 42,2108128 | 1,39976735 | 0,35390556 | 3,95520025 | 7,65E-05 | 0,00059291 | NR1H4 |
| ENSGALG00000009390 | 538,734312 | 1,398258988 | 0,20555082 | 6,80249786 | 1,03E-11 | 3,26E-10 | |
| ENSGALG00000004252 | 60,1197029 | 1,3977567 | 0,34729606 | 4,02468343 | 5,71E-05 | 0,00046153 | |
| ENSGALG00000019864 | 58,8923664 | 1,392548079 | 0,34877295 | 3,9927067 | 6,53E-05 | 0,00051768 | |
| ENSGALG00000029083 | 179,607144 | 1,390642479 | 0,28146869 | 4,94066488 | 7,79E-07 | 1,03E-05 | NXPH2 |
| ENSGALG00000012875 | 68,4672034 | 1,389711553 | 0,3440597 | 4,03915817 | 5,36E-05 | 0,0004376 | WEE2 |
| ENSGALG00000023689 | 74,9775085 | 1,388126911 | 0,34057145 | 4,07587575 | 4,58E-05 | 0,0003808 | ASS1 |
| ENSGALG00000026536 | 52,5121501 | 1,387507758 | 0,35369019 | 3,92294664 | 8,75E-05 | 0,00067057 | IP6K3 |
| ENSGALG00000014857 | 696,584571 | 1,387229623 | 0,18283909 | 7,58716103 | 3,27E-14 | 1,45E-12 | SELENOP |
| ENSGALG00000002319 | 151,603804 | 1,386034084 | 0,29385541 | 4,71672129 | 2,40E-06 | 2,84E-05 | TEKT2 |
| ENSGALG00000015776 | 28,4892202 | 1,382386109 | 0,34372541 | 4,02177453 | 5,78E-05 | 0,00046644 | GABRR2 |
| ENSGALG00000007636 | 33,4789778 | 1,381489214 | 0,3508105 | 3,93799273 | 8,22E-05 | 0,00063274 | PCK1 |
| ENSGALG00000007014 | 150,696913 | 1,378668928 | 0,28986414 | 4,75625908 | 1,97E-06 | 2,38E-05 | |
| ENSGALG00000027973 | 59,7315947 | 1,378158223 | 0,3525539 | 3,90907098 | 9,27E-05 | 0,0007038 | CATHL1 |
| ENSGALG00000017099 | 99,5011022 | 1,377440865 | 0,3228202 | 4,26689803 | 1,98E-05 | 0,00018598 | RASL11A |
| ENSGALG00000005493 | 91,8625446 | 1,375327372 | 0,3276333 | 4,19776428 | 2,70E-05 | 0,00024241 | ITGB1BP2 |
| ENSGALG00000006522 | 45,3533386 | 1,373172 | 0,35333674 | 3,88629841 | 0,00010178 | 0,00076226 | HCK |
| ENSGALG00000012956 | 1426,50489 | 1,372804533 | 0,18631922 | 7,36802439 | 1,73E-13 | 6,93E-12 | TMTC1 |
| ENSGALG00000002105 | 44,5411838 | 1,371897103 | 0,35367265 | 3,8790025 | 0,00010489 | 0,00078376 | FOXI1 |
| ENSGALG00000016584 | 157,587524 | 1,371284929 | 0,29956385 | 4,57760477 | 4,70E-06 | 5,25E-05 | EGFL6 |
| ENSGALG00000000730 | 181,141324 | 1,369179792 | 0,28016862 | 4,88698482 | 1,02E-06 | 1,33E-05 | |

| | | | | | | | |
|---|---|---|---|---|---|---|---|
| ENSGALG00000009151 | 104,263114 | 1,368816021 | 0,32143562 | 4,25844529 | 2,06E-05 | 0,00019223 | |
| ENSGALG00000027420 | 141,879195 | 1,367365407 | 0,29420706 | 4,64762942 | 3,36E-06 | 3,86E-05 | POPDC3 |
| ENSGALG00000012971 | 180,161283 | 1,365874431 | 0,27502418 | 4,96637936 | 6,82E-07 | 9,15E-06 | CCDC170 |
| ENSGALG00000005852 | 42,716737 | 1,364112477 | 0,35370786 | 3,85660773 | 0,00011497 | 0,00084385 | SRMS |
| ENSGALG00000020458 | 27,1299508 | 1,361629377 | 0,33978429 | 4,00733472 | 6,14E-05 | 0,00049065 | |
| ENSGALG00000013221 | 46,5848114 | 1,36089575 | 0,35262592 | 3,85931853 | 0,0001137 | 0,00083635 | ARHGEF33 |
| ENSGALG00000016155 | 67,5195979 | 1,359832396 | 0,3425225 | 3,97005278 | 7,19E-05 | 0,00056355 | |
| ENSGALG00000004840 | 833,065062 | 1,358453061 | 0,16895662 | 8,04024766 | 8,97E-16 | 4,78E-14 | RPH3A |
| ENSGALG00000014913 | 89,5865526 | 1,358196979 | 0,33321545 | 4,07603243 | 4,58E-05 | 0,00038078 | ROS1 |
| ENSGALG00000028822 | 179,379351 | 1,357789969 | 0,27926636 | 4,86198902 | 1,16E-06 | 1,50E-05 | RNF152 |
| ENSGALG00000001932 | 150,41219 | 1,357009459 | 0,31045596 | 4,37102081 | 1,24E-05 | 0,00012454 | SRRM3 |
| ENSGALG00000006104 | 74,2255943 | 1,356707559 | 0,33688447 | 4,02721902 | 5,64E-05 | 0,00045713 | ADAMTSL3 |
| ENSGALG00000013914 | 741,463509 | 1,355434386 | 0,22319941 | 6,0727508 | 1,26E-09 | 2,95E-08 | |
| ENSGALG00000001863 | 95,5730164 | 1,354146203 | 0,34957981 | 3,87363962 | 0,00010722 | 0,00079659 | |
| ENSGALG00000024064 | 33,3700028 | 1,353791502 | 0,35292071 | 3,83596505 | 0,00012507 | 0,00090819 | |
| ENSGALG00000017036 | 3151,42236 | 1,353726483 | 0,14349833 | 9,43374367 | 3,96E-21 | 3,41E-19 | LHFP |
| ENSGALG00000016342 | 64,6889555 | 1,353678127 | 0,34807358 | 3,88905739 | 0,00010063 | 0,00075407 | MYOM2 |
| ENSGALG00000028162 | 25,784822 | 1,353451729 | 0,34077076 | 3,97173668 | 7,14E-05 | 0,00056055 | |
| ENSGALG00000025857 | 83,4628776 | 1,352908608 | 0,3353198 | 4,03468155 | 5,47E-05 | 0,0004451 | |
| ENSGALG00000005898 | 30,8597554 | 1,351366295 | 0,34832633 | 3,87959844 | 0,00010463 | 0,00078228 | |
| ENSGALG00000026817 | 190,25856 | 1,350039834 | 0,30726986 | 4,39366176 | 1,11E-05 | 0,00011375 | |
| ENSGALG00000011259 | 318,18554 | 1,346953088 | 0,23787295 | 5,66248952 | 1,49E-08 | 2,86E-07 | RPE65 |
| ENSGALG00000015186 | 1129,15289 | 1,345677898 | 0,16699384 | 8,05824872 | 7,74E-16 | 4,20E-14 | |
| ENSGALG00000027568 | 132,010856 | 1,344832291 | 0,31080687 | 4,32690662 | 1,51E-05 | 0,00014748 | |
| ENSGALG00000003671 | 47,5471462 | 1,341551712 | 0,35388746 | 3,79089929 | 0,0001501 | 0,00106718 | |
| ENSGALG00000007106 | 977,161228 | 1,340640592 | 0,20719828 | 6,47032681 | 9,78E-11 | 2,66E-09 | |
| ENSGALG00000002466 | 67,363583 | 1,337320513 | 0,34594234 | 3,86573239 | 0,00011076 | 0,00081776 | SLC2A5 |
| ENSGALG00000008283 | 62,9561587 | 1,336619904 | 0,34809582 | 3,83980453 | 0,00012313 | 0,00089506 | |
| ENSGALG00000009058 | 854,57904 | 1,336327117 | 0,17615198 | 7,58621684 | 3,29E-14 | 1,46E-12 | |
| ENSGALG00000014523 | 712,216802 | 1,335395904 | 0,18088858 | 7,38242249 | 1,55E-13 | 6,30E-12 | |
| ENSGALG00000027624 | 90,7091995 | 1,332865286 | 0,32442701 | 4,10836716 | 3,98E-05 | 0,00033971 | |
| ENSGALG00000020380 | 160,77545 | 1,332253662 | 0,29757918 | 4,47697203 | 7,57E-06 | 8,01E-05 | |
| ENSGALG00000014117 | 36,1408998 | 1,330198207 | 0,35376021 | 3,76016911 | 0,0001698 | 0,00119227 | |
| ENSGALG00000005330 | 2519,77115 | 1,328511787 | 0,13424164 | 9,89642129 | 4,31E-23 | 4,45E-21 | SNCAIP |
| ENSGALG00000013481 | 89,3340529 | 1,327640637 | 0,33153417 | 4,00453641 | 6,21E-05 | 0,00049533 | IKZF3 |
| ENSGALG00000008434 | 453,347477 | 1,327566577 | 0,22347729 | 5,94049892 | 2,84E-09 | 6,30E-08 | SORCS3 |
| ENSGALG00000006134 | 56,0911679 | 1,326807835 | 0,34869604 | 3,80505559 | 0,00014177 | 0,00101484 | |
| ENSGALG00000004903 | 45,997304 | 1,326402261 | 0,35371776 | 3,74988883 | 0,00017691 | 0,00123332 | CHST8 |
| ENSGALG00000015309 | 102,618363 | 1,321942011 | 0,33018568 | 4,00363218 | 6,24E-05 | 0,00049694 | SCML4 |
| ENSGALG00000016551 | 5221,05508 | 1,321564075 | 0,11774551 | 11,2239022 | 3,11E-29 | 5,11E-27 | AP1S2 |
| ENSGALG00000013302 | 789,25879 | 1,321288665 | 0,17573754 | 7,51853389 | 5,54E-14 | 2,35E-12 | |
| ENSGALG00000009315 | 90,4745544 | 1,318764229 | 0,33354287 | 3,9538073 | 7,69E-05 | 0,00059603 | PAPLN |
| ENSGALG00000027575 | 153,600714 | 1,316495304 | 0,2966273 | 4,43821352 | 9,07E-06 | 9,46E-05 | |
| ENSGALG00000003050 | 83,7827729 | 1,314582894 | 0,33514644 | 3,92241347 | 8,77E-05 | 0,00067168 | |
| ENSGALG00000002792 | 26,9329424 | 1,31306542 | 0,34508002 | 3,80510415 | 0,00014174 | 0,00101484 | SPDEF |
| ENSGALG00000016368 | 37,7471182 | 1,308737431 | 0,35393883 | 3,69763729 | 0,00021762 | 0,00147112 | |
| ENSGALG00000002389 | 532,146018 | 1,308458099 | 0,19760492 | 6,62158668 | 3,55E-11 | 1,03E-09 | ITGB4 |
| ENSGALG00000000164 | 213,826705 | 1,307977843 | 0,27864581 | 4,69405176 | 2,68E-06 | 3,15E-05 | MYBPH |
| ENSGALG00000007732 | 613,17499 | 1,306274335 | 0,18526209 | 7,0509534 | 1,78E-12 | 6,32E-11 | RTN4R |
| ENSGALG00000013838 | 384,542651 | 1,305966304 | 0,21582461 | 6,05105365 | 1,44E-09 | 3,34E-08 | |
| ENSGALG00000003316 | 80,9265708 | 1,303670454 | 0,33354405 | 3,90854055 | 9,29E-05 | 0,0007043 | |
| ENSGALG00000005802 | 409,595368 | 1,302158184 | 0,2262217 | 5,75611525 | 8,61E-09 | 1,72E-07 | FLT4 |
| ENSGALG00000016109 | 151,672095 | 1,299400996 | 0,28890898 | 4,49761372 | 6,87E-06 | 7,31E-05 | |
| ENSGALG00000009700 | 477,109619 | 1,299326338 | 0,23266824 | 5,58445936 | 2,34E-08 | 4,28E-07 | PDK4 |
| ENSGALG00000006045 | 136,045181 | 1,297721233 | 0,30150221 | 4,30418474 | 1,68E-05 | 0,0001609 | SH3GL3 |
| ENSGALG00000006165 | 67,1698457 | 1,296804529 | 0,34433853 | 3,76607445 | 0,00016583 | 0,00116563 | WDFY4 |
| ENSGALG00000002234 | 407,001304 | 1,296274176 | 0,21729503 | 5,96550322 | 2,44E-09 | 5,48E-08 | UNC13D |
| ENSGALG00000002051 | 21,0390889 | 1,294841211 | 0,33239734 | 3,89546205 | 9,80E-05 | 0,00073766 | |
| ENSGALG00000008022 | 294,029309 | 1,293915137 | 0,23960701 | 5,40015561 | 6,66E-08 | 1,11E-06 | |
| ENSGALG00000000775 | 136,777503 | 1,291374316 | 0,31998815 | 4,03569419 | 5,44E-05 | 0,00044358 | CTSS |
| ENSGALG00000010540 | 264,704566 | 1,289706596 | 0,24325268 | 5,30192147 | 1,15E-07 | 1,81E-06 | TTC39A |
| ENSGALG00000005790 | 146,077421 | 1,289660896 | 0,31496039 | 4,09467652 | 4,23E-05 | 0,00035587 | |
| ENSGALG00000004859 | 354,90162 | 1,288425333 | 0,24778178 | 5,19983886 | 1,99E-07 | 3,02E-06 | ZNFX1 |
| ENSGALG00000003118 | 754,378386 | 1,286216079 | 0,18775875 | 6,85036558 | 7,37E-12 | 2,38E-10 | ADGRA2 |

| | | | | | | | |
|---|---|---|---|---|---|---|---|
| ENSGALG00000006779 | 194,184542 | 1,285794374 | 0,29046865 | 4,42662009 | 9,57E-06 | 9,92E-05 | PDZD3 |
| ENSGALG00000012362 | 940,671677 | 1,285048698 | 0,17252552 | 7,44845563 | 9,44E-14 | 3,88E-12 | THSD7B |
| ENSGALG00000004044 | 637,181076 | 1,285008651 | 0,19785383 | 6,49473719 | 8,32E-11 | 2,30E-09 | CSRP3 |
| ENSGALG00000003580 | 1991,2953 | 1,284312011 | 0,19422271 | 6,61257377 | 3,78E-11 | 1,09E-09 | MMP2 |
| ENSGALG00000002855 | 1183,64483 | 1,278753967 | 0,16211473 | 7,88795656 | 3,07E-15 | 1,50E-13 | SARDH |
| ENSGALG00000016564 | 80,407544 | 1,275019755 | 0,34133276 | 3,73541576 | 0,00018741 | 0,00129256 | PTK2B |
| ENSGALG00000001028 | 434,619355 | 1,272121671 | 0,22824463 | 5,5735011 | 2,50E-08 | 4,54E-07 | RNF43 |
| ENSGALG00000007078 | 141,279073 | 1,270778013 | 0,30664146 | 4,14418193 | 3,41E-05 | 0,00029631 | |
| ENSGALG00000009689 | 798,333776 | 1,269454305 | 0,17898437 | 7,09254266 | 1,32E-12 | 4,74E-11 | |
| ENSGALG00000012740 | 435,216388 | 1,269186578 | 0,21598232 | 5,8763447 | 4,19E-09 | 8,99E-08 | |
| ENSGALG00000000909 | 237,543246 | 1,268775969 | 0,26254608 | 4,83258398 | 1,35E-06 | 1,70E-05 | GFAP |
| ENSGALG00000016967 | 712,9034 | 1,267335006 | 0,21530236 | 5,88630325 | 3,95E-09 | 8,55E-08 | ENOX1 |
| ENSGALG00000019797 | 403,10583 | 1,266041829 | 0,28583293 | 4,42930704 | 9,45E-06 | 9,81E-05 | |
| ENSGALG00000007562 | 30,6870844 | 1,263491411 | 0,34960809 | 3,61402221 | 0,00030148 | 0,00194997 | FGF4 |
| ENSGALG00000008953 | 507,742329 | 1,263384491 | 0,20357748 | 6,20591478 | 5,44E-10 | 1,33E-08 | AASS |
| ENSGALG00000001795 | 978,865317 | 1,261084777 | 0,18732211 | 6,73217257 | 1,67E-11 | 5,13E-10 | |
| ENSGALG00000012978 | 566,09757 | 1,261073108 | 0,1899643 | 6,63847421 | 3,17E-11 | 9,30E-10 | CDYL2 |
| ENSGALG00000006725 | 183,577155 | 1,259943343 | 0,32489293 | 3,87802635 | 0,00010531 | 0,00078648 | ACAN |
| ENSGALG00000014398 | 578,994538 | 1,259804122 | 0,2168754 | 5,80888439 | 6,29E-09 | 1,29E-07 | |
| ENSGALG00000015598 | 190,820863 | 1,259472248 | 0,27786596 | 4,53266109 | 5,82E-06 | 6,30E-05 | RASGRF2 |
| ENSGALG00000002647 | 518,746939 | 1,258418808 | 0,24512541 | 5,13377553 | 2,84E-07 | 4,13E-06 | PLS1 |
| ENSGALG00000003490 | 147,804607 | 1,25835771 | 0,29722539 | 4,23368174 | 2,30E-05 | 0,0002112 | |
| ENSGALG00000005458 | 78,5157022 | 1,258264066 | 0,3397845 | 3,70312374 | 0,00021296 | 0,00144324 | RPL3L |
| ENSGALG00000005843 | 363,336884 | 1,25767053 | 0,2724846 | 4,61556557 | 3,92E-06 | 4,43E-05 | EEF1A2 |
| ENSGALG00000006809 | 89,4927086 | 1,25736391 | 0,32504634 | 3,86826053 | 0,00010961 | 0,00081065 | POU2AF1 |
| ENSGALG00000014014 | 690,531459 | 1,255177749 | 0,17796048 | 7,05312653 | 1,75E-12 | 6,24E-11 | |
| ENSGALG00000001348 | 34,354329 | 1,253857774 | 0,350669 | 3,57561625 | 0,0003494 | 0,00221571 | TEKT3 |
| ENSGALG00000021855 | 271,814276 | 1,251787198 | 0,25254197 | 4,9567492 | 7,17E-07 | 9,57E-06 | CCDC39 |
| ENSGALG00000007193 | 236,114906 | 1,251769992 | 0,27435374 | 4,56261315 | 5,05E-06 | 5,59E-05 | |
| ENSGALG00000018557 | 36,5653316 | 1,251460959 | 0,35373058 | 3,53789302 | 0,00040333 | 0,00250291 | SOD3 |
| ENSGALG00000025896 | 117,288757 | 1,251281202 | 0,32116119 | 3,89611589 | 9,77E-05 | 0,00073608 | PLPPR4 |
| ENSGALG00000012547 | 219,756071 | 1,25117298 | 0,29086976 | 4,30148871 | 1,70E-05 | 0,00016265 | |
| ENSGALG00000013356 | 85,2230662 | 1,250931573 | 0,33961603 | 3,68337025 | 0,00023017 | 0,00154304 | IKBKE |
| ENSGALG00000003404 | 2332,06775 | 1,24748878 | 0,13904188 | 8,97203598 | 2,91E-19 | 2,21E-17 | |
| ENSGALG00000001206 | 49,0370075 | 1,247009104 | 0,35307655 | 3,53183782 | 0,00041268 | 0,00255627 | |
| ENSGALG00000028496 | 22,1201797 | 1,24376929 | 0,33591389 | 3,70264323 | 0,00021336 | 0,00144525 | |
| ENSGALG00000007065 | 27,8385164 | 1,239888336 | 0,34784058 | 3,5645304 | 0,00036451 | 0,00229546 | GJD4 |
| ENSGALG00000015345 | 537,385222 | 1,237723953 | 0,20004279 | 6,18729599 | 6,12E-10 | 1,49E-08 | |
| ENSGALG00000028448 | 68,098086 | 1,236909029 | 0,34651471 | 3,5695715 | 0,00035757 | 0,00225696 | SH2D1B |
| ENSGALG00000001564 | 906,154794 | 1,234344815 | 0,19698595 | 6,26615653 | 3,70E-10 | 9,23E-09 | |
| ENSGALG00000023036 | 562,73194 | 1,234170444 | 0,20582667 | 5,99616393 | 2,02E-09 | 4,57E-08 | FRMPD1 |
| ENSGALG00000010927 | 22,2125847 | 1,233723332 | 0,33533284 | 3,67910081 | 0,00023406 | 0,00156441 | |
| ENSGALG00000028681 | 168,618009 | 1,232657979 | 0,29055151 | 4,2424766 | 2,21E-05 | 0,00020433 | |
| ENSGALG00000015573 | 43,775019 | 1,232214926 | 0,3535094 | 3,48566378 | 0,00049092 | 0,00295484 | FHL5 |
| ENSGALG00000005298 | 800,818564 | 1,231219861 | 0,17189685 | 7,16255038 | 7,92E-13 | 2,92E-11 | ST6GALNAC6 |
| ENSGALG00000016112 | 539,87209 | 1,228482119 | 0,20939423 | 5,86683837 | 4,44E-09 | 9,46E-08 | |
| ENSGALG00000000107 | 237,729231 | 1,227291307 | 0,27709903 | 4,42907111 | 9,46E-06 | 9,81E-05 | TRIM7 |
| ENSGALG00000002086 | 83,650156 | 1,225708617 | 0,33362046 | 3,67396119 | 0,00023882 | 0,00159076 | RASL10B |
| ENSGALG00000002667 | 699,199625 | 1,225466735 | 0,17683961 | 6,92982057 | 4,21E-12 | 1,40E-10 | OLFML2B |
| ENSGALG00000013431 | 35,4490725 | 1,223926576 | 0,35351054 | 3,46220673 | 0,00053577 | 0,00317564 | |
| ENSGALG00000028031 | 210,360934 | 1,22342747 | 0,2837755 | 4,31125118 | 1,62E-05 | 0,00015642 | C1QL1 |
| ENSGALG00000012019 | 150,440096 | 1,222353012 | 0,29815258 | 4,09975655 | 4,14E-05 | 0,00035053 | |
| ENSGALG00000000048 | 677,127073 | 1,221920851 | 0,19907606 | 6,13795972 | 8,36E-10 | 2,00E-08 | |
| ENSGALG00000028946 | 391,056645 | 1,217039733 | 0,25108871 | 4,84705086 | 1,25E-06 | 1,59E-05 | |
| ENSGALG00000014407 | 118,55781 | 1,216198172 | 0,31990769 | 3,80171597 | 0,0001437 | 0,00102593 | |
| ENSGALG00000012658 | 2563,96357 | 1,214376265 | 0,15238934 | 7,96890584 | 1,60E-15 | 8,19E-14 | TCP11L2 |
| ENSGALG00000028402 | 33,811586 | 1,211737057 | 0,34956854 | 3,46637901 | 0,00052752 | 0,00313769 | |
| ENSGALG00000003345 | 1065,51677 | 1,211546204 | 0,16093702 | 7,52807643 | 5,15E-14 | 2,19E-12 | |
| ENSGALG00000005083 | 47,8682724 | 1,210240985 | 0,35126406 | 3,44538807 | 0,00057024 | 0,00334551 | SLC5A10 |
| ENSGALG00000003803 | 110,857566 | 1,208011558 | 0,33977968 | 3,55527901 | 0,00037758 | 0,0023603 | DAB2 |
| ENSGALG00000026578 | 55,333978 | 1,20748664 | 0,35388721 | 3,41206638 | 0,00064472 | 0,00370533 | |
| ENSGALG00000014126 | 18,9923327 | 1,207239433 | 0,3279747 | 3,6808919 | 0,00023242 | 0,00155583 | |
| ENSGALG00000015764 | 5967,75227 | 1,20416113 | 0,19086925 | 6,30882741 | 2,81E-10 | 7,13E-09 | |
| ENSGALG00000015528 | 170,233046 | 1,204136401 | 0,30064358 | 4,00519575 | 6,20E-05 | 0,00049428 | JAKMIP1 |

| | | | | | | | |
|---|---|---|---|---|---|---|---|
| ENSGALG00000002355 | 234,006388 | 1,203134529 | 0,25489167 | 4,72017987 | 2,36E-06 | 2,80E-05 | |
| ENSGALG00000004641 | 146,280836 | 1,20257315 | 0,29654538 | 4,05527534 | 5,01E-05 | 0,0004122 | |
| ENSGALG00000026903 | 111,361959 | 1,200671289 | 0,32831415 | 3,65708057 | 0,0002551 | 0,00168686 | CALB2 |
| ENSGALG00000004745 | 1802,35745 | 1,199713581 | 0,14187699 | 8,45601213 | 2,77E-17 | 1,74E-15 | NATD1 |
| ENSGALG00000013890 | 33,5607486 | 1,1996045 | 0,35201412 | 3,40783067 | 0,00065482 | 0,00374908 | |
| ENSGALG00000019280 | 990,724576 | 1,199070585 | 0,17339981 | 6,91506298 | 4,68E-12 | 1,54E-10 | |
| ENSGALG00000004066 | 55,6194601 | 1,199060923 | 0,35390165 | 3,38811906 | 0,00070374 | 0,00399391 | HTR6 |
| ENSGALG00000002082 | 57,1027996 | 1,198569946 | 0,34848133 | 3,43940933 | 0,00058299 | 0,00340658 | AMPD1 |
| ENSGALG00000013090 | 44,7148094 | 1,197838739 | 0,35381435 | 3,38550073 | 0,00071049 | 0,00402652 | LOXL4 |
| ENSGALG00000014696 | 222,596022 | 1,197793931 | 0,25975439 | 4,61125584 | 4,00E-06 | 4,51E-05 | |
| ENSGALG00000006922 | 19,3844584 | 1,194740346 | 0,3271281 | 3,65220948 | 0,00025999 | 0,0017142 | |
| ENSGALG00000011489 | 4656,68128 | 1,193784841 | 0,12166568 | 9,8120096 | 1,00E-22 | 9,93E-21 | |
| ENSGALG00000016590 | 18,8906226 | 1,192009768 | 0,32708105 | 3,64438646 | 0,00026803 | 0,00175697 | TLR7 |
| ENSGALG00000002407 | 144,980361 | 1,191296249 | 0,30414176 | 3,9169112 | 8,97E-05 | 0,00068373 | HYDIN |
| ENSGALG00000012890 | 273,965985 | 1,191168201 | 0,26407391 | 4,51073784 | 6,46E-06 | 6,90E-05 | DGKI |
| ENSGALG00000027218 | 239,44089 | 1,190798556 | 0,27577811 | 4,317959 | 1,57E-05 | 0,00015239 | |
| ENSGALG00000017040 | 33,0805894 | 1,190245244 | 0,3523261 | 3,37824885 | 0,00072949 | 0,00411775 | |
| ENSGALG00000008362 | 175,023576 | 1,190011885 | 0,27541961 | 4,32072321 | 1,56E-05 | 0,00015103 | SSTR4 |
| ENSGALG00000004521 | 401,353263 | 1,187877806 | 0,21973208 | 5,40602807 | 6,44E-08 | 1,08E-06 | GPX3 |
| ENSGALG00000006074 | 272,240776 | 1,186152058 | 0,28359005 | 4,18262934 | 2,88E-05 | 0,00025631 | SOX6 |
| ENSGALG00000003842 | 39,7811882 | 1,185854219 | 0,35383776 | 3,35140662 | 0,00080402 | 0,00446624 | GHRH |
| ENSGALG00000021882 | 58,6974367 | 1,185518376 | 0,34682245 | 3,41822852 | 0,0006303 | 0,00363471 | |
| ENSGALG00000002744 | 252,461162 | 1,184999187 | 0,25185371 | 4,70510904 | 2,54E-06 | 3,00E-05 | PCDH15 |
| ENSGALG00000006768 | 37,969199 | 1,18499895 | 0,35365822 | 3,35068965 | 0,00080611 | 0,00447275 | |
| ENSGALG00000014812 | 711,71818 | 1,183271769 | 0,18701629 | 6,32710549 | 2,50E-10 | 6,39E-09 | SIDT1 |
| ENSGALG00000017331 | 630,733948 | 1,182854332 | 0,18865205 | 6,27003173 | 3,61E-10 | 9,05E-09 | |
| ENSGALG00000005547 | 144,897217 | 1,180323703 | 0,29897868 | 3,94785238 | 7,89E-05 | 0,00060932 | PIK3AP1 |
| ENSGALG00000017291 | 662,461807 | 1,180080414 | 0,20904403 | 5,64512838 | 1,65E-08 | 3,14E-07 | FAR2 |
| ENSGALG00000006308 | 28,0784724 | 1,179461013 | 0,33709473 | 3,4989008 | 0,00046718 | 0,00283376 | |
| ENSGALG00000013112 | 320,804038 | 1,179282262 | 0,23305967 | 5,06000147 | 4,19E-07 | 5,90E-06 | RERGL |
| ENSGALG00000007333 | 325,518561 | 1,178440858 | 0,25844246 | 4,55978035 | 5,12E-06 | 5,65E-05 | UBAP1L |
| ENSGALG00000001021 | 191,271341 | 1,17756894 | 0,28381375 | 4,14909056 | 3,34E-05 | 0,00029114 | MYOCD |
| ENSGALG00000003821 | 322,044025 | 1,177115475 | 0,23475754 | 5,01417543 | 5,33E-07 | 7,38E-06 | RAMP1 |
| ENSGALG00000003164 | 243,863169 | 1,17697714 | 0,25907141 | 4,54306075 | 5,54E-06 | 6,04E-05 | |
| ENSGALG00000016651 | 946,900962 | 1,174964335 | 0,18486316 | 6,35585995 | 2,07E-10 | 5,41E-09 | |
| ENSGALG00000002435 | 70,1914893 | 1,174947125 | 0,34690999 | 3,38689333 | 0,00070689 | 0,00400845 | |
| ENSGALG00000001331 | 1782,64917 | 1,174179888 | 0,1437468 | 8,16839003 | 3,13E-16 | 1,75E-14 | ST14 |
| ENSGALG00000009114 | 216,592643 | 1,170782693 | 0,26080261 | 4,48915246 | 7,15E-06 | 7,60E-05 | |
| ENSGALG00000028580 | 59,1862765 | 1,170297276 | 0,34946527 | 3,34882286 | 0,00081156 | 0,00449343 | |
| ENSGALG00000006099 | 3185,42607 | 1,169037408 | 0,1372144 | 8,51978685 | 1,60E-17 | 1,03E-15 | ZFPM1 |
| ENSGALG00000014261 | 23192,4244 | 1,166708788 | 0,14807674 | 7,87908222 | 3,30E-15 | 1,59E-13 | |
| ENSGALG00000006627 | 44,7235286 | 1,164076921 | 0,35292296 | 3,29838819 | 0,00097242 | 0,00521033 | UPB1 |
| ENSGALG00000012668 | 622,845457 | 1,1628752 | 0,24480349 | 4,75023949 | 2,03E-06 | 2,44E-05 | |
| ENSGALG00000011446 | 368,843846 | 1,161895934 | 0,22151576 | 5,24520663 | 1,56E-07 | 2,42E-06 | TNFAIP2 |
| ENSGALG00000005630 | 38,0970872 | 1,161740711 | 0,34283622 | 3,38861721 | 0,00070246 | 0,00398832 | CRYBB1 |
| ENSGALG00000023879 | 439,210587 | 1,160823609 | 0,21167651 | 5,48395108 | 4,16E-08 | 7,24E-07 | |
| ENSGALG00000007596 | 99,2163667 | 1,160341621 | 0,32378836 | 3,58364221 | 0,00033884 | 0,00216077 | |
| ENSGALG00000005721 | 236,127639 | 1,159323873 | 0,25145197 | 4,61051812 | 4,02E-06 | 4,52E-05 | DGKG |
| ENSGALG00000013911 | 546,725872 | 1,159209839 | 0,2024827 | 5,72498212 | 1,03E-08 | 2,03E-07 | |
| ENSGALG00000011418 | 18,2422739 | 1,158885244 | 0,32516535 | 3,56398753 | 0,00036526 | 0,00229703 | CCR6 |
| ENSGALG00000028489 | 20,5369622 | 1,158414063 | 0,33177044 | 3,4916132 | 0,00048011 | 0,00290006 | |
| ENSGALG00000001031 | 327,10314 | 1,156740289 | 0,24403187 | 4,74011969 | 2,14E-06 | 2,56E-05 | HSF5 |
| ENSGALG00000002016 | 953,709981 | 1,156233761 | 0,16494826 | 7,0096752 | 2,39E-12 | 8,30E-11 | PRKCD |
| ENSGALG00000001095 | 18,2554746 | 1,156162616 | 0,32498658 | 3,55757036 | 0,0003743 | 0,00234627 | |
| ENSGALG00000013980 | 288,264915 | 1,155390063 | 0,2983628 | 3,87243334 | 0,00010775 | 0,00079949 | SLC2A12 |
| ENSGALG00000026241 | 639,218946 | 1,154849081 | 0,23028217 | 5,01493056 | 5,31E-07 | 7,36E-06 | GRIN3B |
| ENSGALG00000006208 | 100,690133 | 1,154488936 | 0,32195187 | 3,58590535 | 0,00033591 | 0,00214413 | OTOG |
| ENSGALG00000009641 | 18352,4117 | 1,153845642 | 0,14589909 | 7,90851867 | 2,60E-15 | 1,28E-13 | COL1A2 |
| ENSGALG00000012288 | 52,9340022 | 1,152071153 | 0,35012037 | 3,29050023 | 0,00100009 | 0,00532927 | |
| ENSGALG00000015932 | 523,705524 | 1,152042382 | 0,22404186 | 5,14208532 | 2,72E-07 | 3,98E-06 | |
| ENSGALG00000003789 | 56,0437149 | 1,151405842 | 0,3500328 | 3,28942274 | 0,00100393 | 0,00534762 | FAM78A |
| ENSGALG00000010365 | 313,078692 | 1,151056068 | 0,26208135 | 4,39198002 | 1,12E-05 | 0,0001143 | |
| ENSGALG00000017691 | 74,6333149 | 1,149429932 | 0,34278483 | 3,35321124 | 0,0007988 | 0,00444266 | |
| ENSGALG00000017374 | 478,615413 | 1,149174538 | 0,22225681 | 5,17048077 | 2,33E-07 | 3,48E-06 | |

| | | | | | | | |
|---|---|---|---|---|---|---|---|
| ENSGALG00000028710 | 21,1011731 | 1,148265888 | 0,3321095 | 3,45749187 | 0,00054523 | 0,00322331 | |
| ENSGALG00000012374 | 515,33143 | 1,147327656 | 0,19490406 | 5,88662778 | 3,94E-09 | 8,55E-08 | FRMD6 |
| ENSGALG00000016328 | 5259,3238 | 1,146952103 | 0,14040122 | 8,16910377 | 3,11E-16 | 1,75E-14 | |
| ENSGALG00000025721 | 19,2431462 | 1,146921853 | 0,32414703 | 3,53827662 | 0,00040275 | 0,00250042 | |
| ENSGALG00000012365 | 9658,53942 | 1,146353139 | 0,11013775 | 10,4083585 | 2,27E-25 | 2,97E-23 | PYGL |
| ENSGALG00000005540 | 392,975435 | 1,145405784 | 0,21951203 | 5,21796369 | 1,81E-07 | 2,76E-06 | MICAL2 |
| ENSGALG00000028702 | 48,7217309 | 1,145044679 | 0,35222211 | 3,25091653 | 0,00115034 | 0,0060053 | |
| ENSGALG00000022819 | 93,4187666 | 1,14482978 | 0,33650375 | 3,40213085 | 0,00066863 | 0,00382494 | PCP4 |
| ENSGALG00000001968 | 55,5701756 | 1,14394352 | 0,35227777 | 3,2472771 | 0,00116515 | 0,00606403 | SYT6 |
| ENSGALG00000012505 | 1173,09399 | 1,142096363 | 0,156235 | 7,31011862 | 2,67E-13 | 1,05E-11 | LRFN5 |
| ENSGALG00000015758 | 1198,98398 | 1,141437782 | 0,17077903 | 6,68371174 | 2,33E-11 | 6,96E-10 | |
| ENSGALG00000003876 | 41,5449157 | 1,140755556 | 0,34478607 | 3,30858949 | 0,00093767 | 0,00507207 | |
| ENSGALG00000009791 | 992,573884 | 1,140657464 | 0,21268775 | 5,3630614 | 8,18E-08 | 1,34E-06 | PROX1 |
| ENSGALG00000028130 | 1683,17579 | 1,140619832 | 0,15899173 | 7,17408269 | 7,28E-13 | 2,72E-11 | |
| ENSGALG00000010673 | 22,7777978 | 1,139845847 | 0,34030069 | 3,34952553 | 0,0008095 | 0,00448387 | SLC1A7 |
| ENSGALG00000006436 | 58,7768458 | 1,138634151 | 0,35302664 | 3,22534908 | 0,00125819 | 0,00646426 | |
| ENSGALG00000000620 | 143,16144 | 1,137872957 | 0,29367405 | 3,8746119 | 0,00010679 | 0,00079541 | |
| ENSGALG00000013818 | 524,978654 | 1,136933045 | 0,21089775 | 5,39092064 | 7,01E-08 | 1,17E-06 | CITED2 |
| ENSGALG00000007236 | 26,5853166 | 1,136243459 | 0,34408993 | 3,30217003 | 0,0009594 | 0,00516702 | |
| ENSGALG00000023643 | 36,5223482 | 1,136155947 | 0,35390776 | 3,2103166 | 0,00132589 | 0,00675092 | |
| ENSGALG00000007129 | 549,585731 | 1,134513156 | 0,27598889 | 4,11072037 | 3,94E-05 | 0,00033711 | |
| ENSGALG00000001442 | 382,78952 | 1,131898366 | 0,21564935 | 5,24879104 | 1,53E-07 | 2,37E-06 | OPCML |
| ENSGALG00000000217 | 558,865853 | 1,131400306 | 0,21968434 | 5,15011819 | 2,60E-07 | 3,84E-06 | PPFIA4 |
| ENSGALG00000000911 | 18,8021132 | 1,131377385 | 0,32322345 | 3,50029482 | 0,00046474 | 0,00282099 | IL19 |
| ENSGALG00000003028 | 252,863378 | 1,131167716 | 0,25365391 | 4,45949245 | 8,22E-06 | 8,64E-05 | |
| ENSGALG00000027626 | 26,2635541 | 1,129092221 | 0,34723307 | 3,251684 | 0,00114723 | 0,00599848 | |
| ENSGALG00000008063 | 112,836606 | 1,129089778 | 0,31997995 | 3,52862665 | 0,00041772 | 0,00258161 | NOX5 |
| ENSGALG00000016592 | 75,5093263 | 1,127895523 | 0,34582316 | 3,26148062 | 0,00110832 | 0,00582167 | PRPS2 |
| ENSGALG00000021529 | 183,029289 | 1,127864143 | 0,27306603 | 4,13037145 | 3,62E-05 | 0,00031229 | GP1BB |
| ENSGALG00000006503 | 1322,53017 | 1,127315674 | 0,17398315 | 6,47945327 | 9,21E-11 | 2,52E-09 | VDAC1 |
| ENSGALG00000026333 | 170,514796 | 1,126889713 | 0,28096298 | 4,01081206 | 6,05E-05 | 0,00048433 | |
| ENSGALG00000015348 | 3709,65535 | 1,126202791 | 0,16010935 | 7,03395999 | 2,01E-12 | 7,08E-11 | ALCAM |
| ENSGALG00000027872 | 562,415563 | 1,125834202 | 0,2065498 | 5,45066721 | 5,02E-08 | 8,57E-07 | |
| ENSGALG00000007964 | 712,005282 | 1,125506336 | 0,19390731 | 5,80435241 | 6,46E-09 | 1,32E-07 | GATSL3 |
| ENSGALG00000008940 | 122,073119 | 1,125469701 | 0,30445466 | 3,69667421 | 0,00021844 | 0,00147524 | SPTBN5 |
| ENSGALG00000012252 | 31,0664734 | 1,124245419 | 0,34638552 | 3,24564781 | 0,00117184 | 0,00608953 | |
| ENSGALG00000009237 | 283,646852 | 1,124183808 | 0,25265321 | 4,44951324 | 8,61E-06 | 9,03E-05 | |
| ENSGALG00000008629 | 362,430604 | 1,122527446 | 0,28746439 | 3,90492695 | 9,43E-05 | 0,00071253 | |
| ENSGALG00000004649 | 115,318908 | 1,122312882 | 0,33908753 | 3,30980286 | 0,00093362 | 0,00506019 | TRPV1 |
| ENSGALG00000016821 | 91,586414 | 1,121951764 | 0,32755694 | 3,42521141 | 0,00061432 | 0,00356825 | |
| ENSGALG00000000994 | 36,3375502 | 1,120119947 | 0,35335281 | 3,1699761 | 0,00152451 | 0,00753677 | |
| ENSGALG00000009943 | 1812,23536 | 1,119992507 | 0,14209817 | 7,8818219 | 3,23E-15 | 1,56E-13 | FAM117A |
| ENSGALG00000028795 | 3146,83309 | 1,118039364 | 0,16042604 | 6,96918862 | 3,19E-12 | 1,09E-10 | |
| ENSGALG00000026758 | 74,5546131 | 1,11741437 | 0,33538362 | 3,33174999 | 0,00086302 | 0,0047277 | |
| ENSGALG00000001204 | 1443,12547 | 1,117076992 | 0,14668872 | 7,61528921 | 2,63E-14 | 1,19E-12 | ENKD1 |
| ENSGALG00000012055 | 30,6319383 | 1,115864912 | 0,34954847 | 3,19230382 | 0,00141143 | 0,00709885 | |
| ENSGALG00000015469 | 3626,20782 | 1,115704157 | 0,1321025 | 8,44574565 | 3,02E-17 | 1,89E-15 | |
| ENSGALG00000009963 | 76,7800387 | 1,114244364 | 0,33758559 | 3,30062774 | 0,00096469 | 0,00518524 | LYZ |
| ENSGALG00000011414 | 712,898703 | 1,112714675 | 0,17966211 | 6,19337424 | 5,89E-10 | 1,44E-08 | AMDHD1 |
| ENSGALG00000015454 | 87,0613625 | 1,111985108 | 0,33768276 | 3,29298749 | 0,00099129 | 0,00529478 | |
| ENSGALG00000010233 | 149,38519 | 1,111330421 | 0,289789 | 3,83496413 | 0,00012558 | 0,00091141 | SYNDIG1L |
| ENSGALG00000024411 | 95,5715091 | 1,110994905 | 0,32259096 | 3,44397412 | 0,00057323 | 0,00335823 | |
| ENSGALG00000003016 | 111,975898 | 1,109545556 | 0,31007137 | 3,57835536 | 0,00034576 | 0,00219771 | MASP2 |
| ENSGALG00000006792 | 447,856144 | 1,109047707 | 0,21887998 | 5,06692165 | 4,04E-07 | 5,71E-06 | VSTM2L |
| ENSGALG00000009823 | 207,863068 | 1,108978484 | 0,26701827 | 4,15319326 | 3,28E-05 | 0,00028633 | PTPRE |
| ENSGALG00000027501 | 1178,23042 | 1,108782799 | 0,15555529 | 7,12790156 | 1,02E-12 | 3,72E-11 | ANPEP |
| ENSGALG00000004017 | 335,243952 | 1,108435951 | 0,22759443 | 4,87022448 | 1,11E-06 | 1,44E-05 | TOR1AIP2 |
| ENSGALG00000025912 | 62,6339712 | 1,108425226 | 0,34376932 | 3,22432853 | 0,00126268 | 0,00648 | |
| ENSGALG00000000158 | 209,106145 | 1,107475494 | 0,2643096 | 4,19006906 | 2,79E-05 | 0,00024946 | |
| ENSGALG00000002610 | 604,590631 | 1,105862871 | 0,18967796 | 5,83021282 | 5,54E-09 | 1,15E-07 | |
| ENSGALG00000023621 | 702,550009 | 1,105773477 | 0,19719495 | 5,60751405 | 2,05E-08 | 3,82E-07 | SFXN3 |
| ENSGALG00000009574 | 24,9756977 | 1,105678443 | 0,34380548 | 3,21600002 | 0,00129991 | 0,00664829 | |
| ENSGALG00000006000 | 170,909865 | 1,105166021 | 0,29241804 | 3,77940435 | 0,0001572 | 0,00111071 | |
| ENSGALG00000001011 | 623,472513 | 1,104831537 | 0,2206424 | 5,00733998 | 5,52E-07 | 7,61E-06 | |

| | | | | | | | |
|---|---|---|---|---|---|---|---|
| ENSGALG00000011473 | 347,60043 | 1,104703027 | 0,22754114 | 4,85495953 | 1,20E-06 | 1,54E-05 | RPS6KA2 |
| ENSGALG00000026834 | 233,708518 | 1,10373431 | 0,25983013 | 4,24790728 | 2,16E-05 | 0,00020012 | |
| ENSGALG00000006003 | 58,9954831 | 1,10329195 | 0,3504724 | 3,14801378 | 0,00164384 | 0,00802229 | |
| ENSGALG00000001281 | 270,657774 | 1,101845994 | 0,24895157 | 4,42594511 | 9,60E-06 | 9,94E-05 | |
| ENSGALG00000006407 | 926,885262 | 1,100961832 | 0,17822617 | 6,17732963 | 6,52E-10 | 1,58E-08 | |
| ENSGALG00000011003 | 180,050652 | 1,099169303 | 0,28561474 | 3,84843338 | 0,00011888 | 0,00086689 | SLC35F3 |
| ENSGALG00000000168 | 20,2372176 | 1,099055711 | 0,33636253 | 3,26747371 | 0,00108512 | 0,00571524 | ADORA1 |
| ENSGALG00000014798 | 1444,6552 | 1,098754437 | 0,18532469 | 5,92880749 | 3,05E-09 | 6,69E-08 | SERINC5 |
| ENSGALG00000015076 | 118,818737 | 1,094512369 | 0,30475359 | 3,59146666 | 0,00032882 | 0,00210678 | |
| ENSGALG00000002579 | 438,862484 | 1,09231668 | 0,24079762 | 4,53624361 | 5,73E-06 | 6,21E-05 | RIMBP2 |
| ENSGALG00000027102 | 27,9313615 | 1,090659115 | 0,34855662 | 3,12907298 | 0,00175359 | 0,00846159 | |
| ENSGALG00000007740 | 159,049731 | 1,089704597 | 0,29794304 | 3,6574259 | 0,00025476 | 0,00168541 | |
| ENSGALG00000004972 | 200,694244 | 1,089536909 | 0,30285765 | 3,59752149 | 0,00032126 | 0,00206127 | |
| ENSGALG00000004538 | 131,202356 | 1,088596778 | 0,30030855 | 3,62492773 | 0,00028904 | 0,00188202 | TNN |
| ENSGALG00000005470 | 915,927743 | 1,087631823 | 0,1770332 | 6,14366031 | 8,06E-10 | 1,93E-08 | PLPPR5 |
| ENSGALG00000006189 | 81,0476646 | 1,08750661 | 0,33966232 | 3,20172876 | 0,00136606 | 0,00691664 | CRYGN |
| ENSGALG00000001716 | 219,086022 | 1,084481493 | 0,27662697 | 3,92037508 | 8,84E-05 | 0,00067587 | C15orf59 |
| ENSGALG00000017106 | 609,784044 | 1,082115941 | 0,19278194 | 5,61316028 | 1,99E-08 | 3,72E-07 | ATP8A2 |
| ENSGALG00000014264 | 24,119483 | 1,081789708 | 0,34218632 | 3,16140549 | 0,0015701 | 0,00773684 | NSUN7 |
| ENSGALG00000001470 | 804,498578 | 1,081645785 | 0,18189855 | 5,94642334 | 2,74E-09 | 6,10E-08 | STOML1 |
| ENSGALG00000023636 | 80,78387 | 1,079731402 | 0,33574779 | 3,21590026 | 0,00130036 | 0,00664829 | |
| ENSGALG00000021303 | 70,1790621 | 1,078464744 | 0,33870785 | 3,18405595 | 0,00145227 | 0,00725779 | |
| ENSGALG00000011353 | 167,024074 | 1,077984355 | 0,29480885 | 3,65655358 | 0,00025563 | 0,00168869 | |
| ENSGALG00000026651 | 469,211499 | 1,076820994 | 0,26273591 | 4,09849186 | 4,16E-05 | 0,00035167 | SCARA3 |
| ENSGALG00000005031 | 222,866745 | 1,076697936 | 0,27611031 | 3,89952093 | 9,64E-05 | 0,00072701 | DRP2 |
| ENSGALG00000005076 | 658,308514 | 1,075685444 | 0,20135322 | 5,34228092 | 9,18E-08 | 1,49E-06 | |
| ENSGALG00000013180 | 23,7835918 | 1,075255786 | 0,33793034 | 3,18188592 | 0,00146319 | 0,0072892 | SLC6A18 |
| ENSGALG00000004043 | 1743,12209 | 1,074636101 | 0,18109836 | 5,93399142 | 2,96E-09 | 6,52E-08 | |
| ENSGALG00000000420 | 275,786063 | 1,072751202 | 0,24334062 | 4,40843449 | 1,04E-05 | 0,00010699 | |
| ENSGALG00000003640 | 1881,52679 | 1,071788322 | 0,14728433 | 7,2770018 | 3,41E-13 | 1,33E-11 | |
| ENSGALG00000009355 | 164,050555 | 1,070935457 | 0,2815733 | 3,80339851 | 0,00014272 | 0,00102045 | CLDN11 |
| ENSGALG00000026748 | 19,4954498 | 1,069944148 | 0,32787916 | 3,26322706 | 0,00110151 | 0,00578815 | |
| ENSGALG00000004406 | 128,12145 | 1,066534863 | 0,31159618 | 3,42281114 | 0,00061977 | 0,00358919 | |
| ENSGALG00000006370 | 31,1681716 | 1,066397762 | 0,34558448 | 3,08578021 | 0,00203019 | 0,00956436 | |
| ENSGALG00000012377 | 68,8818144 | 1,065846022 | 0,33849137 | 3,14881303 | 0,00163935 | 0,00801428 | HNMT |
| ENSGALG00000010752 | 133,532542 | 1,063752645 | 0,31701468 | 3,35553118 | 0,00079213 | 0,00441384 | CDCP2 |
| ENSGALG00000021049 | 25,1605315 | 1,063749147 | 0,33499236 | 3,17544297 | 0,00149608 | 0,00741774 | |
| ENSGALG00000006374 | 154,22163 | 1,06372742 | 0,34611406 | 3,07334361 | 0,00211675 | 0,00991383 | TBX6 |
| ENSGALG00000007780 | 1769,76781 | 1,063721106 | 0,16415536 | 6,47996553 | 9,17E-11 | 2,52E-09 | EIF4E3 |
| ENSGALG00000021542 | 348,907918 | 1,061917275 | 0,24470276 | 4,3396212 | 1,43E-05 | 0,00014102 | SNCB |
| ENSGALG00000002055 | 5059,12603 | 1,061032342 | 0,12896244 | 8,22745271 | 1,91E-16 | 1,10E-14 | hairy1 |
| ENSGALG00000028742 | 318,003255 | 1,06018982 | 0,24476244 | 4,33150539 | 1,48E-05 | 0,0001453 | |
| ENSGALG00000001040 | 379,37277 | 1,060173693 | 0,22651608 | 4,68034629 | 2,86E-06 | 3,34E-05 | |
| ENSGALG00000008734 | 1277,14965 | 1,060092794 | 0,17848485 | 5,93939919 | 2,86E-09 | 6,33E-08 | PCSK2 |
| ENSGALG00000025771 | 418,454799 | 1,059868939 | 0,21305921 | 4,97452772 | 6,54E-07 | 8,82E-06 | |
| ENSGALG00000017311 | 141,945952 | 1,059578058 | 0,29313349 | 3,61466049 | 0,00030074 | 0,00194647 | |
| ENSGALG00000002554 | 211,299245 | 1,058725863 | 0,27265415 | 3,88303593 | 0,00010316 | 0,00077214 | TSPAN2 |
| ENSGALG00000011898 | 1941,78412 | 1,056447793 | 0,1400944 | 7,54097066 | 4,66E-14 | 2,00E-12 | FAM109B |
| ENSGALG00000012545 | 20,3956262 | 1,051934631 | 0,33312924 | 3,15773734 | 0,00158999 | 0,00780659 | CYTIP |
| ENSGALG00000010497 | 795,792907 | 1,048579632 | 0,17878894 | 5,86490213 | 4,49E-09 | 9,56E-08 | SLC29A4 |
| ENSGALG00000000503 | 90,5216838 | 1,0483641 | 0,32883068 | 3,18815785 | 0,00143182 | 0,00718021 | |
| ENSGALG00000007959 | 24,9928179 | 1,04699816 | 0,3339903 | 3,13481605 | 0,00171962 | 0,00832534 | SLC26A3 |
| ENSGALG00000012045 | 94,1610251 | 1,046487546 | 0,33468277 | 3,1268044 | 0,00176717 | 0,00850781 | |
| ENSGALG00000009340 | 441,508564 | 1,044750001 | 0,25059513 | 4,16907546 | 3,06E-05 | 0,00027021 | |
| ENSGALG00000007617 | 161,132474 | 1,044420437 | 0,30323964 | 3,44420813 | 0,00057273 | 0,00335677 | |
| ENSGALG00000007837 | 153,44217 | 1,044277791 | 0,31727304 | 3,29141673 | 0,00099684 | 0,0053161 | KAZALD1 |
| ENSGALG00000016020 | 82,6868728 | 1,041815502 | 0,33102199 | 3,14726979 | 0,00164803 | 0,0080365 | |
| ENSGALG00000016608 | 1071,03802 | 1,041630113 | 0,16148809 | 6,45019776 | 1,12E-10 | 3,01E-09 | ADCY3 |
| ENSGALG00000014258 | 1564,45283 | 1,040436704 | 0,17049678 | 6,10238319 | 1,04E-09 | 2,47E-08 | LIMCH1 |
| ENSGALG00000027096 | 1851,9247 | 1,040113834 | 0,13778609 | 7,54875793 | 4,39E-14 | 1,90E-12 | LGALSL |
| ENSGALG00000024369 | 197,407512 | 1,038738473 | 0,26515652 | 3,91745396 | 8,95E-05 | 0,00068257 | |
| ENSGALG00000028234 | 617,735502 | 1,03829388 | 0,19109872 | 5,43328543 | 5,53E-08 | 9,37E-07 | |
| ENSGALG00000007219 | 1265,62807 | 1,038287687 | 0,15486967 | 6,70426727 | 2,02E-11 | 6,12E-10 | |
| ENSGALG00000007253 | 1268,2995 | 1,036667095 | 0,15805846 | 6,55875735 | 5,43E-11 | 1,54E-09 | IL1RAP |

| | | | | | | | |
|---|---|---|---|---|---|---|---|
| ENSGALG00000005460 | 763,147126 | 1,036332157 | 0,18413261 | 5,6281837 | 1,82E-08 | 3,43E-07 | SLCO4A1 |
| ENSGALG00000026714 | 2323,49209 | 1,035206481 | 0,15670753 | 6,60597808 | 3,95E-11 | 1,13E-09 | |
| ENSGALG00000024122 | 36,985836 | 1,03453695 | 0,33443583 | 3,09337955 | 0,00197891 | 0,00935579 | |
| ENSGALG00000023779 | 766,128835 | 1,033222085 | 0,19891303 | 5,19434077 | 2,05E-07 | 3,11E-06 | |
| ENSGALG00000028318 | 1679,83733 | 1,033139386 | 0,18498818 | 5,584894 | 2,34E-08 | 4,28E-07 | CDKN1A |
| ENSGALG00000005114 | 245,746736 | 1,032666035 | 0,25457852 | 4,05637544 | 4,98E-05 | 0,00041076 | |
| ENSGALG00000028628 | 176,658931 | 1,032210406 | 0,27931003 | 3,6955723 | 0,00021939 | 0,00148092 | |
| ENSGALG00000011875 | 520,49686 | 1,032103241 | 0,20932071 | 4,93072682 | 8,19E-07 | 1,08E-05 | TRPC3 |
| ENSGALG00000001304 | 107,91968 | 1,031984172 | 0,31646483 | 3,26097581 | 0,0011103 | 0,0058298 | |
| ENSGALG00000004860 | 237,1229 | 1,03038994 | 0,25186264 | 4,09107893 | 4,29E-05 | 0,00035997 | RASD1 |
| ENSGALG00000010566 | 1230,63642 | 1,029946325 | 0,16433565 | 6,26733359 | 3,67E-10 | 9,18E-09 | |
| ENSGALG00000012791 | 100,570938 | 1,028853172 | 0,32574707 | 3,15844184 | 0,00158615 | 0,00779056 | TBXAS1 |
| ENSGALG00000024024 | 333,882358 | 1,028182094 | 0,2283532 | 4,5025956 | 6,71E-06 | 7,16E-05 | TGFA |
| ENSGALG00000009739 | 1255,18637 | 1,027863615 | 0,18332662 | 5,60673425 | 2,06E-08 | 3,83E-07 | AMIGO2 |
| ENSGALG00000010078 | 138,046016 | 1,027109659 | 0,2951454 | 3,48001248 | 0,00050139 | 0,00300724 | GLP1R |
| ENSGALG00000003349 | 284,814958 | 1,025286613 | 0,23608456 | 4,34287875 | 1,41E-05 | 0,00013925 | PLEKHA2 |
| ENSGALG00000009722 | 2700,68769 | 1,024969081 | 0,12748481 | 8,03993117 | 8,99E-16 | 4,78E-14 | DYNC1I1 |
| ENSGALG00000013200 | 141,95986 | 1,024952745 | 0,30040339 | 3,4119214 | 0,00064507 | 0,00370573 | GABBR2 |
| ENSGALG00000029169 | 143,486281 | 1,024627176 | 0,29797907 | 3,43858777 | 0,00058476 | 0,00341401 | |
| ENSGALG00000025906 | 86,1054184 | 1,019071961 | 0,32995746 | 3,08849502 | 0,00201173 | 0,00949054 | |
| ENSGALG00000012564 | 122,738702 | 1,018929434 | 0,30201581 | 3,37376189 | 0,00074148 | 0,00416817 | |
| ENSGALG00000009686 | 819,704721 | 1,017336386 | 0,20309235 | 5,00923036 | 5,46E-07 | 7,54E-06 | |
| ENSGALG00000001698 | 101,577734 | 1,017252154 | 0,31962329 | 3,18265971 | 0,00145929 | 0,00727774 | GABRA1 |
| ENSGALG00000002991 | 1547,8463 | 1,016425478 | 0,15188757 | 6,69195943 | 2,20E-11 | 6,60E-10 | |
| ENSGALG00000001191 | 726,030438 | 1,014738695 | 0,17607486 | 5,76310952 | 8,26E-09 | 1,66E-07 | MEOX1 |
| ENSGALG00000009066 | 83,1568975 | 1,013658719 | 0,32821816 | 3,08836875 | 0,00201259 | 0,00949129 | |
| ENSGALG00000016223 | 759,19748 | 1,013131958 | 0,18257274 | 5,54919615 | 2,87E-08 | 5,19E-07 | |
| ENSGALG00000010203 | 639,913614 | 1,012100046 | 0,1964193 | 5,15275255 | 2,57E-07 | 3,79E-06 | |
| ENSGALG00000008874 | 19,4465425 | 1,010524458 | 0,32352202 | 3,1235106 | 0,00178707 | 0,00857934 | SLC13A1 |
| ENSGALG00000022718 | 221,713706 | 1,010197798 | 0,26186126 | 3,85775962 | 0,00011443 | 0,00084033 | |
| ENSGALG00000014455 | 15,3755596 | 1,009369671 | 0,31516038 | 3,20271755 | 0,00136137 | 0,0068955 | LPAR5 |
| ENSGALG00000008080 | 251,329122 | 1,008867324 | 0,25119635 | 4,01624997 | 5,91E-05 | 0,00047553 | C1QTNF4 |
| ENSGALG00000001951 | 315,95633 | 1,007959576 | 0,24129054 | 4,17736875 | 2,95E-05 | 0,00026123 | |
| ENSGALG00000027712 | 289,192222 | 1,006836187 | 0,24626896 | 4,08836006 | 4,34E-05 | 0,00036327 | |
| ENSGALG00000028186 | 897,733771 | 1,005437673 | 0,18601887 | 5,40503063 | 6,48E-08 | 1,09E-06 | RERG |
| ENSGALG00000003816 | 91,3783387 | 1,005135037 | 0,32711976 | 3,07268213 | 0,00212144 | 0,009926 | NT5C1A |
| ENSGALG00000007713 | 153,549096 | 1,002069085 | 0,29743797 | 3,36900193 | 0,00075441 | 0,00423331 | |
| ENSGALG00000016251 | 6835,37033 | 1,000647408 | 0,15469938 | 6,46833505 | 9,91E-11 | 2,69E-09 | |
| ENSGALG00000023953 | 637,175356 | 0,998916427 | 0,18661623 | 5,35278435 | 8,66E-08 | 1,41E-06 | |
| ENSGALG00000005887 | 163,632495 | 0,998359123 | 0,30751037 | 3,24658685 | 0,00116798 | 0,00607411 | |
| ENSGALG00000012480 | 541,532617 | 0,99597856 | 0,2064035 | 4,82539577 | 1,40E-06 | 1,76E-05 | |
| ENSGALG00000014681 | 398,451477 | 0,995892967 | 0,23114252 | 4,30856669 | 1,64E-05 | 0,0001581 | PCSK1 |
| ENSGALG00000017316 | 218,980953 | 0,995598244 | 0,30126263 | 3,30475185 | 0,00095061 | 0,0051298 | UCP3 |
| ENSGALG00000001453 | 365,617866 | 0,994068241 | 0,24549624 | 4,04921983 | 5,14E-05 | 0,00042072 | |
| ENSGALG00000002437 | 722,53208 | 0,993124174 | 0,18855357 | 5,26706645 | 1,39E-07 | 2,17E-06 | FAM69B |
| ENSGALG00000012613 | 121,972608 | 0,992227725 | 0,31630972 | 3,13688661 | 0,00170752 | 0,00827328 | FBP1 |
| ENSGALG00000028682 | 130,825911 | 0,990483134 | 0,3063312 | 3,23337336 | 0,00122338 | 0,00631397 | CBLN2 |
| ENSGALG00000013183 | 389,466708 | 0,988585254 | 0,21644464 | 4,56738155 | 4,94E-06 | 5,48E-05 | TERT |
| ENSGALG00000009638 | 342,698465 | 0,988233263 | 0,2260661 | 4,3714351 | 1,23E-05 | 0,00012439 | ANO6 |
| ENSGALG00000016224 | 196,097983 | 0,988198095 | 0,27726141 | 3,56413865 | 0,00036505 | 0,00229676 | MAOA |
| ENSGALG00000009052 | 3269,15476 | 0,987910707 | 0,16241285 | 6,08271266 | 1,18E-09 | 2,78E-08 | LBH |
| ENSGALG00000026042 | 849,913797 | 0,987626868 | 0,23717847 | 4,16406621 | 3,13E-05 | 0,00027514 | |
| ENSGALG00000012536 | 763,863052 | 0,987580114 | 0,21162614 | 4,66662634 | 3,06E-06 | 3,54E-05 | GALNT13 |
| ENSGALG00000016599 | 16,0014025 | 0,987073109 | 0,31315435 | 3,15203387 | 0,00162137 | 0,00793209 | |
| ENSGALG00000009463 | 2838,70473 | 0,986691121 | 0,14031176 | 7,03213428 | 2,03E-12 | 7,16E-11 | |
| ENSGALG00000005030 | 1142,72142 | 0,984706358 | 0,17789423 | 5,53534734 | 3,11E-08 | 5,55E-07 | DOCK10 |
| ENSGALG00000027429 | 108,061421 | 0,983232181 | 0,31233583 | 3,14799676 | 0,00164394 | 0,00802229 | HSD17B1 |
| ENSGALG00000023847 | 205,80734 | 0,981888346 | 0,26827405 | 3,66001979 | 0,0002522 | 0,00166925 | |
| ENSGALG00000012552 | 632,633121 | 0,981819434 | 0,18202425 | 5,39389345 | 6,89E-08 | 1,15E-06 | TOM1 |
| ENSGALG00000027739 | 223,979623 | 0,981670308 | 0,28088257 | 3,49494923 | 0,00047415 | 0,00287042 | |
| ENSGALG00000012715 | 380,224027 | 0,978917656 | 0,24539961 | 3,98907579 | 6,63E-05 | 0,00052445 | CAP2 |
| ENSGALG00000012709 | 399,326921 | 0,978132572 | 0,259146 | 3,77444597 | 0,00016036 | 0,0011301 | ATXN1 |
| ENSGALG00000028301 | 585,856129 | 0,976931086 | 0,20530374 | 4,75846719 | 1,95E-06 | 2,36E-05 | |
| ENSGALG00000006882 | 2766,70659 | 0,976472658 | 0,13081296 | 7,46464785 | 8,35E-14 | 3,47E-12 | ST8SIA2 |

| | | | | | | | |
|---|---|---|---|---|---|---|---|
| ENSGALG00000024059 | 15,053333 | 0,976199065 | 0,31251247 | 3,12371231 | 0,00178585 | 0,00857649 | |
| ENSGALG00000010797 | 412,002343 | 0,975073157 | 0,21003242 | 4,64248879 | 3,44E-06 | 3,95E-05 | |
| ENSGALG00000000902 | 273,817902 | 0,974915117 | 0,24422686 | 3,99184233 | 6,56E-05 | 0,00051867 | KDF1 |
| ENSGALG00000004591 | 574,278704 | 0,974905454 | 0,19696934 | 4,949529 | 7,44E-07 | 9,89E-06 | SERPINC1 |
| ENSGALG00000018875 | 14,7404116 | 0,973272591 | 0,3123576 | 3,11589219 | 0,00183389 | 0,00876086 | |
| ENSGALG00000003886 | 1023,87733 | 0,971801068 | 0,17739435 | 5,47819617 | 4,30E-08 | 7,44E-07 | SGCD |
| ENSGALG00000005844 | 1501,38821 | 0,969445519 | 0,15286931 | 6,34166216 | 2,27E-10 | 5,87E-09 | |
| ENSGALG00000016820 | 787,309833 | 0,968970008 | 0,17832491 | 5,43373323 | 5,52E-08 | 9,36E-07 | GAS6 |
| ENSGALG00000027287 | 736,986058 | 0,968636988 | 0,18931485 | 5,11653998 | 3,11E-07 | 4,50E-06 | |
| ENSGALG00000003170 | 1633,73146 | 0,967045053 | 0,17416807 | 5,55236702 | 2,82E-08 | 5,10E-07 | |
| ENSGALG00000026870 | 2578,97026 | 0,966911139 | 0,16329066 | 5,92141112 | 3,19E-09 | 6,99E-08 | |
| ENSGALG00000027511 | 3914,70702 | 0,966828969 | 0,15258887 | 6,33616983 | 2,36E-10 | 6,06E-09 | |
| ENSGALG00000001209 | 151,276779 | 0,963839438 | 0,28852453 | 3,34058053 | 0,00083603 | 0,00460089 | |
| ENSGALG00000012681 | 687,549661 | 0,96179459 | 0,19337383 | 4,97375784 | 6,57E-07 | 8,84E-06 | MBOAT1 |
| ENSGALG00000026914 | 303,146434 | 0,960831377 | 0,2341299 | 4,10383883 | 4,06E-05 | 0,00034533 | DOK3 |
| ENSGALG00000016597 | 546,580903 | 0,96044418 | 0,2115615 | 4,53978708 | 5,63E-06 | 6,12E-05 | |
| ENSGALG00000028041 | 172,712765 | 0,96010795 | 0,29281701 | 3,27886677 | 0,00104225 | 0,00552416 | DKK2 |
| ENSGALG00000008317 | 155,968503 | 0,958791686 | 0,29825236 | 3,21469944 | 0,00130581 | 0,00666615 | SDSL |
| ENSGALG00000006192 | 588,22456 | 0,958606154 | 0,19869528 | 4,82450397 | 1,40E-06 | 1,77E-05 | |
| ENSGALG00000009896 | 1725,9341 | 0,956710434 | 0,138947 | 6,88543432 | 5,76E-12 | 1,88E-10 | TRERF1 |
| ENSGALG00000008936 | 346,610654 | 0,956545148 | 0,23935184 | 3,99639774 | 6,43E-05 | 0,00051087 | |
| ENSGALG00000012577 | 1716,87461 | 0,956309243 | 0,13789127 | 6,93524147 | 4,06E-12 | 1,35E-10 | FRMD3 |
| ENSGALG00000016945 | 3627,68031 | 0,953551228 | 0,1270066 | 7,5078871 | 6,01E-14 | 2,53E-12 | LECT1 |
| ENSGALG00000014843 | 607,093989 | 0,953307805 | 0,21113346 | 4,51519047 | 6,33E-06 | 6,80E-05 | TPD52L1 |
| ENSGALG00000029150 | 2513,11501 | 0,953288378 | 0,14860816 | 6,41477801 | 1,41E-10 | 3,76E-09 | AK1 |
| ENSGALG00000000295 | 158,361127 | 0,951838119 | 0,2874721 | 3,31106259 | 0,00092942 | 0,00504148 | |
| ENSGALG00000002008 | 770,021515 | 0,951550674 | 0,26369144 | 3,60857625 | 0,00030788 | 0,00198664 | RNF157 |
| ENSGALG00000017926 | 117,345158 | 0,950771836 | 0,30495213 | 3,11777407 | 0,00182222 | 0,00871735 | 7SK |
| ENSGALG00000009123 | 138,335079 | 0,94996493 | 0,29437793 | 3,22702502 | 0,00125085 | 0,00643402 | |
| ENSGALG00000011046 | 918,666805 | 0,946061534 | 0,22454083 | 4,21331632 | 2,52E-05 | 0,00022872 | SNX10 |
| ENSGALG00000004792 | 282,665704 | 0,945718297 | 0,26094655 | 3,62418393 | 0,00028988 | 0,00188564 | |
| ENSGALG00000006988 | 545,963197 | 0,944077449 | 0,23809277 | 3,96516647 | 7,33E-05 | 0,00057357 | |
| ENSGALG00000009002 | 180,825204 | 0,943061124 | 0,27453332 | 3,43514266 | 0,00059224 | 0,00345327 | |
| ENSGALG00000003370 | 6860,89746 | 0,941055859 | 0,11588753 | 8,12042401 | 4,65E-16 | 2,56E-14 | KIAA1191 |
| ENSGALG00000015183 | 541,002513 | 0,940741895 | 0,24692948 | 3,80975932 | 0,0001391 | 0,00099941 | RAB13 |
| ENSGALG00000010977 | 629,762637 | 0,94045823 | 0,18538133 | 5,07310108 | 3,91E-07 | 5,55E-06 | FAM221A |
| ENSGALG00000017169 | 2527,40437 | 0,94013642 | 0,1550564 | 6,06319017 | 1,33E-09 | 3,11E-08 | ELMOD1 |
| ENSGALG00000006351 | 260,931586 | 0,940035694 | 0,25660603 | 3,66334215 | 0,00024895 | 0,00165256 | FAS |
| ENSGALG00000007415 | 599,165563 | 0,938743099 | 0,23705941 | 3,95994876 | 7,50E-05 | 0,0005829 | |
| ENSGALG00000002678 | 150,077896 | 0,938682309 | 0,29361496 | 3,19698392 | 0,00138873 | 0,00700797 | CSPG4 |
| ENSGALG00000003832 | 1422,76359 | 0,938218604 | 0,14754036 | 6,35906414 | 2,03E-10 | 5,31E-09 | TEX264 |
| ENSGALG00000009725 | 763,677984 | 0,938023358 | 0,22425701 | 4,182805 | 2,88E-05 | 0,00025631 | SCG5 |
| ENSGALG00000015967 | 204,249989 | 0,936813576 | 0,2634911 | 3,55538981 | 0,00037742 | 0,0023603 | LRRTM1 |
| ENSGALG00000002473 | 480,317696 | 0,93617222 | 0,20261658 | 4,62041265 | 3,83E-06 | 4,35E-05 | GPR157 |
| ENSGALG00000015926 | 1110,05542 | 0,934566237 | 0,15987762 | 5,84550999 | 5,05E-09 | 1,06E-07 | |
| ENSGALG00000001398 | 801,871514 | 0,933503925 | 0,17565694 | 5,31435844 | 1,07E-07 | 1,71E-06 | |
| ENSGALG00000028292 | 453,150865 | 0,931661788 | 0,25026444 | 3,72270944 | 0,0001971 | 0,00134712 | |
| ENSGALG00000008735 | 158,650282 | 0,931477748 | 0,29513805 | 3,15607472 | 0,00159908 | 0,00784557 | |
| ENSGALG00000010139 | 381,674313 | 0,931031468 | 0,2175137 | 4,28033488 | 1,87E-05 | 0,00017632 | SSTR1 |
| ENSGALG00000002843 | 586,866611 | 0,930974563 | 0,19321147 | 4,81842275 | 1,45E-06 | 1,82E-05 | RILP |
| ENSGALG00000023626 | 1834,26904 | 0,929476271 | 0,16548518 | 5,61667362 | 1,95E-08 | 3,65E-07 | NTN1 |
| ENSGALG00000028091 | 182,303874 | 0,927849672 | 0,28991773 | 3,20038958 | 0,00137242 | 0,00694328 | MYF5 |
| ENSGALG00000023517 | 259,64579 | 0,925400404 | 0,25687314 | 3,60255801 | 0,0003151 | 0,00202459 | AGPAT2 |
| ENSGALG00000005626 | 609,645166 | 0,92496767 | 0,21590969 | 4,2840488 | 1,84E-05 | 0,00017352 | TPST2 |
| ENSGALG00000000311 | 558,408468 | 0,92295118 | 0,21535867 | 4,2856468 | 1,82E-05 | 0,00017251 | |
| ENSGALG00000015207 | 1291,12938 | 0,92039826 | 0,14720064 | 6,25267856 | 4,03E-10 | 1,00E-08 | |
| ENSGALG00000023936 | 805,049393 | 0,918539777 | 0,20126409 | 4,56385321 | 5,02E-06 | 5,56E-05 | |
| ENSGALG00000011709 | 513,362717 | 0,918333355 | 0,23178437 | 3,96201584 | 7,43E-05 | 0,0005792 | |
| ENSGALG00000027622 | 398,876868 | 0,916634803 | 0,26586468 | 3,44774944 | 0,00056528 | 0,00332452 | |
| ENSGALG00000015795 | 539,939541 | 0,91628777 | 0,2153216 | 4,25543833 | 2,09E-05 | 0,00019456 | ADAMTS5 |
| ENSGALG00000005120 | 2531,22001 | 0,916224882 | 0,15496532 | 5,91245108 | 3,37E-09 | 7,37E-08 | |
| ENSGALG00000013071 | 1399,47707 | 0,914850391 | 0,17423001 | 5,25081979 | 1,51E-07 | 2,35E-06 | |
| ENSGALG00000010382 | 346,13588 | 0,914779385 | 0,22801207 | 4,01197789 | 6,02E-05 | 0,00048223 | |
| ENSGALG00000004309 | 938,335478 | 0,913060877 | 0,18553484 | 4,92123686 | 8,60E-07 | 1,13E-05 | GDAP1L1 |

| | | | | | | | |
|---|---|---|---|---|---|---|---|
| ENSGALG00000028622 | 281,675272 | 0,912974481 | 0,25598956 | 3,56645202 | 0,00036185 | 0,00228081 | PEAR1 |
| ENSGALG00000005331 | 677,48087 | 0,912514017 | 0,17989387 | 5,07251308 | 3,93E-07 | 5,56E-06 | GJC2 |
| ENSGALG00000027462 | 1468,67221 | 0,912376278 | 0,14323943 | 6,36958898 | 1,90E-10 | 4,98E-09 | P3H4 |
| ENSGALG00000015789 | 559,929943 | 0,912066241 | 0,21238691 | 4,29436183 | 1,75E-05 | 0,00016704 | |
| ENSGALG00000010551 | 919,316948 | 0,911984879 | 0,18322857 | 4,97730716 | 6,45E-07 | 8,70E-06 | CDC42EP3 |
| ENSGALG00000012298 | 550,04797 | 0,911513817 | 0,22249482 | 4,09678677 | 4,19E-05 | 0,00035361 | SLC39A8 |
| ENSGALG00000003588 | 1398,60382 | 0,911216657 | 0,1584852 | 5,74953779 | 8,95E-09 | 1,78E-07 | |
| ENSGALG00000005814 | 212,002946 | 0,910795359 | 0,27185501 | 3,35029823 | 0,00080725 | 0,00447521 | |
| ENSGALG00000002581 | 512,901786 | 0,910753526 | 0,19519764 | 4,66580184 | 3,07E-06 | 3,56E-05 | HTATIP2 |
| ENSGALG00000004676 | 1678,58309 | 0,910530695 | 0,14330969 | 6,35358792 | 2,10E-10 | 5,48E-09 | |
| ENSGALG00000023195 | 260,809859 | 0,909179266 | 0,24717134 | 3,67833616 | 0,00023476 | 0,00156833 | DLX6 |
| ENSGALG00000015725 | 901,983472 | 0,907979483 | 0,17014105 | 5,33662782 | 9,47E-08 | 1,53E-06 | |
| ENSGALG00000007114 | 305,818525 | 0,907573399 | 0,27679712 | 3,27883977 | 0,00104235 | 0,00552416 | APOA1 |
| ENSGALG00000003018 | 309,190261 | 0,906093174 | 0,24885213 | 3,64109068 | 0,00027149 | 0,00177706 | |
| ENSGALG00000013886 | 841,136602 | 0,904236576 | 0,1804703 | 5,01044535 | 5,43E-07 | 7,51E-06 | LDLRAD4 |
| ENSGALG00000021442 | 473,215804 | 0,903637173 | 0,22582203 | 4,00154571 | 6,29E-05 | 0,00050105 | CARMIL2 |
| ENSGALG00000005014 | 159,559707 | 0,903243848 | 0,28943129 | 3,12075394 | 0,00180389 | 0,00864482 | AIPL1 |
| ENSGALG00000002963 | 189,406665 | 0,902926335 | 0,26991157 | 3,34526731 | 0,00082203 | 0,00453545 | PID1 |
| ENSGALG00000000836 | 482,381548 | 0,902335572 | 0,20852451 | 4,32723978 | 1,51E-05 | 0,00014747 | RFX5 |
| ENSGALG00000011124 | 436,071233 | 0,902068067 | 0,20974945 | 4,30069344 | 1,70E-05 | 0,000163 | ARHGAP24 |
| ENSGALG00000010827 | 4357,48343 | 0,901311467 | 0,11466976 | 7,86006222 | 3,84E-15 | 1,84E-13 | |
| ENSGALG00000008461 | 387,656717 | 0,900341356 | 0,24810826 | 3,62882455 | 0,00028471 | 0,0018574 | |
| ENSGALG00000010119 | 1290,75662 | 0,900097678 | 0,1879287 | 4,78957 | 1,67E-06 | 2,06E-05 | CDKL2 |
| ENSGALG00000020905 | 199,737945 | 0,899806675 | 0,28234109 | 3,18694908 | 0,00143782 | 0,00720498 | KSR2 |
| ENSGALG00000006793 | 399,969639 | 0,898163749 | 0,21225276 | 4,23157623 | 2,32E-05 | 0,0002129 | |
| ENSGALG00000008632 | 774,98804 | 0,897411708 | 0,17277465 | 5,19411687 | 2,06E-07 | 3,11E-06 | ACSS1 |
| ENSGALG00000001896 | 633,555504 | 0,897398544 | 0,18298068 | 4,90433493 | 9,37E-07 | 1,23E-05 | |
| ENSGALG00000006083 | 883,458305 | 0,896818662 | 0,18476425 | 4,85385374 | 1,21E-06 | 1,55E-05 | |
| ENSGALG00000007052 | 507,965349 | 0,895651499 | 0,20347805 | 4,40171068 | 1,07E-05 | 0,00010986 | PXK |
| ENSGALG00000015977 | 788,487319 | 0,89563673 | 0,20705245 | 4,32565141 | 1,52E-05 | 0,00014811 | SMOX |
| ENSGALG00000008262 | 171,789815 | 0,894529697 | 0,28118642 | 3,18126918 | 0,00146631 | 0,0072994 | RASGRF1 |
| ENSGALG00000000374 | 1105,84307 | 0,894239416 | 0,18770457 | 4,76407913 | 1,90E-06 | 2,30E-05 | RNPEP |
| ENSGALG00000017204 | 419,662559 | 0,891856915 | 0,22247021 | 4,00888239 | 6,10E-05 | 0,00048802 | |
| ENSGALG00000002854 | 440,265325 | 0,890941536 | 0,2420509 | 3,68080247 | 0,0002325 | 0,00155583 | |
| ENSGALG00000005203 | 773,684672 | 0,889220424 | 0,2133641 | 4,16761968 | 3,08E-05 | 0,00027177 | OLFM3 |
| ENSGALG00000027603 | 933,535269 | 0,889190298 | 0,16664995 | 5,33567704 | 9,52E-08 | 1,54E-06 | |
| ENSGALG00000003932 | 494,661014 | 0,888085177 | 0,22917231 | 3,87518533 | 0,00010654 | 0,00079397 | SDC4 |
| ENSGALG00000010269 | 1110,19342 | 0,886301871 | 0,16512328 | 5,36751612 | 7,98E-08 | 1,31E-06 | KCNAB1 |
| ENSGALG00000004093 | 635,621253 | 0,885654514 | 0,20975364 | 4,2223558 | 2,42E-05 | 0,00022106 | TBX22 |
| ENSGALG00000015788 | 630,459367 | 0,884593035 | 0,2015792 | 4,38831513 | 1,14E-05 | 0,00011598 | |
| ENSGALG00000004970 | 368,256584 | 0,884420332 | 0,25182556 | 3,51203558 | 0,00044469 | 0,00271743 | |
| ENSGALG00000026152 | 376,28373 | 0,884418189 | 0,23205448 | 3,8112524 | 0,00013826 | 0,00099391 | |
| ENSGALG00000025733 | 660,61374 | 0,882587352 | 0,1955092 | 4,51430076 | 6,35E-06 | 6,83E-05 | |
| ENSGALG00000006008 | 1270,65336 | 0,882382705 | 0,1743852 | 5,05996316 | 4,19E-07 | 5,90E-06 | HOMER2 |
| ENSGALG00000027027 | 1969,08091 | 0,882215122 | 0,16312702 | 5,40814829 | 6,37E-08 | 1,07E-06 | |
| ENSGALG00000003270 | 387,810885 | 0,881994372 | 0,21359889 | 4,12920868 | 3,64E-05 | 0,00031348 | |
| ENSGALG00000008349 | 209,566568 | 0,879571782 | 0,26093775 | 3,37081072 | 0,00074947 | 0,00420787 | LMCD1 |
| ENSGALG00000009601 | 8743,38719 | 0,878730197 | 0,13764581 | 6,38399544 | 1,73E-10 | 4,59E-09 | NELL2 |
| ENSGALG00000027920 | 386,752165 | 0,874011674 | 0,22844437 | 3,82592791 | 0,00013028 | 0,00094249 | |
| ENSGALG00000026059 | 270,416326 | 0,873870501 | 0,23970989 | 3,64553371 | 0,00026684 | 0,00175077 | |
| ENSGALG00000016017 | 590,769047 | 0,873175292 | 0,21544948 | 4,05280753 | 5,06E-05 | 0,00041599 | SLC4A11 |
| ENSGALG00000005974 | 256,914188 | 0,872920311 | 0,25210037 | 3,46259042 | 0,000535 | 0,00317388 | COL6A1 |
| ENSGALG00000001129 | 1785,04064 | 0,871868135 | 0,15304444 | 5,69682996 | 1,22E-08 | 2,38E-07 | |
| ENSGALG00000009867 | 212,190543 | 0,869748432 | 0,26750592 | 3,25132407 | 0,00114869 | 0,00599972 | |
| ENSGALG00000027364 | 6291,5882 | 0,867638443 | 0,1162491 | 7,46361451 | 8,42E-14 | 3,48E-12 | |
| ENSGALG00000011624 | 479,035185 | 0,867273579 | 0,2018671 | 4,29626011 | 1,74E-05 | 0,00016596 | PNLDC1 |
| ENSGALG00000016384 | 268,592265 | 0,867172927 | 0,26856807 | 3,2288758 | 0,00124278 | 0,00640113 | MYT1L |
| ENSGALG00000026172 | 493,174963 | 0,865691508 | 0,21117972 | 4,09931168 | 4,14E-05 | 0,00035078 | |
| ENSGALG00000003895 | 788,021122 | 0,861640305 | 0,25898235 | 3,32702328 | 0,00087779 | 0,00479965 | PRDM12 |
| ENSGALG00000012072 | 201,724233 | 0,861021311 | 0,27532993 | 3,12723472 | 0,00176459 | 0,00849837 | |
| ENSGALG00000000953 | 1344,71246 | 0,860062109 | 0,16163754 | 5,32093044 | 1,03E-07 | 1,66E-06 | |
| ENSGALG00000009902 | 340,496837 | 0,859561208 | 0,24909364 | 3,4507553 | 0,00055902 | 0,00329341 | CFAP57 |
| ENSGALG00000005655 | 299,935919 | 0,858231064 | 0,25571117 | 3,3562518 | 0,00079007 | 0,0044067 | TSPEAR |
| ENSGALG00000017072 | 1099,84084 | 0,857400203 | 0,18100217 | 4,73696081 | 2,17E-06 | 2,59E-05 | N4BP2L1 |

| | | | | | | | |
|---|---|---|---|---|---|---|---|
| ENSGALG00000025919 | 2494,48854 | 0,857090747 | 0,142817 | 6,00132156 | 1,96E-09 | 4,45E-08 | |
| ENSGALG00000015768 | 2710,70193 | 0,855142896 | 0,12974855 | 6,59077029 | 4,38E-11 | 1,25E-09 | |
| ENSGALG00000010664 | 712,177105 | 0,852579615 | 0,22881793 | 3,72601745 | 0,00019453 | 0,00133284 | KIF26B |
| ENSGALG00000001189 | 452,192377 | 0,851541498 | 0,20637637 | 4,12615796 | 3,69E-05 | 0,00031686 | NCF1 |
| ENSGALG00000009536 | 446,627716 | 0,851130737 | 0,22757849 | 3,73994373 | 0,00018406 | 0,00127273 | |
| ENSGALG00000003612 | 268,623077 | 0,850682606 | 0,26697919 | 3,18632551 | 0,00144092 | 0,00721787 | |
| ENSGALG00000023172 | 1255,76162 | 0,849498162 | 0,15375274 | 5,52509283 | 3,29E-08 | 5,85E-07 | |
| ENSGALG00000026671 | 2116,56921 | 0,847898773 | 0,14258676 | 5,94654652 | 2,74E-09 | 6,10E-08 | |
| ENSGALG00000002742 | 2787,33871 | 0,847416181 | 0,14490214 | 5,84819627 | 4,97E-09 | 1,04E-07 | TMEM132B |
| ENSGALG00000005791 | 520,608854 | 0,84592391 | 0,20398465 | 4,14699797 | 3,37E-05 | 0,00029344 | |
| ENSGALG00000009603 | 307,914805 | 0,844955114 | 0,23876567 | 3,53884673 | 0,00040188 | 0,00249616 | |
| ENSGALG00000006379 | 1670,52626 | 0,844378171 | 0,18274684 | 4,62048036 | 3,83E-06 | 4,35E-05 | SHH |
| ENSGALG00000005895 | 481,788237 | 0,844174027 | 0,20498441 | 4,11823536 | 3,82E-05 | 0,00032712 | |
| ENSGALG00000004593 | 2963,70805 | 0,839985137 | 0,15590672 | 5,38774176 | 7,13E-08 | 1,18E-06 | SULF2 |
| ENSGALG00000010857 | 11562,2517 | 0,839106931 | 0,11368089 | 7,38124899 | 1,57E-13 | 6,34E-12 | Dab1 |
| ENSGALG00000026505 | 1579,97582 | 0,837091894 | 0,15268322 | 5,48254005 | 4,19E-08 | 7,29E-07 | |
| ENSGALG00000003354 | 276,830006 | 0,837083394 | 0,24777884 | 3,37834895 | 0,00072922 | 0,00411775 | KCNH4 |
| ENSGALG00000011205 | 1172,74221 | 0,836341019 | 0,20119253 | 4,15691891 | 3,23E-05 | 0,00028261 | SMOC2 |
| ENSGALG00000005161 | 2738,71244 | 0,836184904 | 0,13858143 | 6,03388843 | 1,60E-09 | 3,70E-08 | |
| ENSGALG00000006014 | 961,400691 | 0,836085862 | 0,15943435 | 5,24407593 | 1,57E-07 | 2,43E-06 | |
| ENSGALG00000007770 | 193,517852 | 0,835610241 | 0,26849664 | 3,11218139 | 0,0018571 | 0,00885001 | |
| ENSGALG00000028582 | 1565,14033 | 0,832821984 | 0,15076847 | 5,52384707 | 3,32E-08 | 5,87E-07 | |
| ENSGALG00000005481 | 1783,10991 | 0,83266607 | 0,14481077 | 5,75002865 | 8,92E-09 | 1,77E-07 | ARHGEF3 |
| ENSGALG00000015660 | 2889,81934 | 0,831570906 | 0,12282495 | 6,77037458 | 1,28E-11 | 4,00E-10 | SNX30 |
| ENSGALG00000028125 | 921,395648 | 0,831550629 | 0,18036279 | 4,6104334 | 4,02E-06 | 4,52E-05 | |
| ENSGALG00000025776 | 414,67908 | 0,83014723 | 0,21462899 | 3,86782432 | 0,00010981 | 0,00081166 | |
| ENSGALG00000027903 | 10650,6436 | 0,829170411 | 0,11053296 | 7,50156702 | 6,31E-14 | 2,64E-12 | |
| ENSGALG00000010500 | 3026,41246 | 0,828235743 | 0,13002656 | 6,36974259 | 1,89E-10 | 4,98E-09 | NKX6-2 |
| ENSGALG00000007015 | 336,010164 | 0,825119263 | 0,22964958 | 3,59294921 | 0,00032696 | 0,00209581 | CD40 |
| ENSGALG00000015720 | 662,971603 | 0,824601233 | 0,20777276 | 3,96876479 | 7,22E-05 | 0,00056596 | CHODL |
| ENSGALG00000011744 | 1298,60197 | 0,824149013 | 0,17298432 | 4,76429887 | 1,90E-06 | 2,30E-05 | |
| ENSGALG00000016558 | 512,367292 | 0,822973317 | 0,20099047 | 4,09458879 | 4,23E-05 | 0,00035587 | VEGFD |
| ENSGALG00000001688 | 9554,36542 | 0,822044682 | 0,12651935 | 6,49738304 | 8,17E-11 | 2,27E-09 | sept-05 |
| ENSGALG00000015941 | 330,156486 | 0,819153887 | 0,23360451 | 3,50658414 | 0,0004539 | 0,00276255 | |
| ENSGALG00000019768 | 3223,32241 | 0,819142269 | 0,12221578 | 6,70242617 | 2,05E-11 | 6,19E-10 | ACSF2 |
| ENSGALG00000010698 | 1578,35111 | 0,816916741 | 0,1404823 | 5,81508672 | 6,06E-09 | 1,25E-07 | |
| ENSGALG00000012628 | 1115,10605 | 0,816609796 | 0,18822357 | 4,33850978 | 1,43E-05 | 0,00014153 | HABP4 |
| ENSGALG00000027534 | 844,147038 | 0,815389343 | 0,17402749 | 4,68540539 | 2,79E-06 | 3,27E-05 | |
| ENSGALG00000010397 | 282,533201 | 0,813527008 | 0,25347167 | 3,20953824 | 0,00132948 | 0,0067667 | GRID2 |
| ENSGALG00000027338 | 409,810985 | 0,813282814 | 0,21988916 | 3,6986034 | 0,00021679 | 0,00146626 | |
| ENSGALG00000015297 | 1260,54646 | 0,81224071 | 0,16743713 | 4,8510192 | 1,23E-06 | 1,57E-05 | FOXO3 |
| ENSGALG00000000954 | 405,070556 | 0,812029159 | 0,24204134 | 3,35491929 | 0,00079388 | 0,00442073 | |
| ENSGALG00000006480 | 1820,43736 | 0,810690895 | 0,14795092 | 5,47945836 | 4,27E-08 | 7,41E-07 | TCF7 |
| ENSGALG00000011861 | 430,597577 | 0,810665245 | 0,21257164 | 3,81360956 | 0,00013695 | 0,00098656 | SYT16 |
| ENSGALG00000009680 | 349,521496 | 0,809243098 | 0,22080925 | 3,6648968 | 0,00024744 | 0,00164472 | PAQR7 |
| ENSGALG00000015051 | 777,652051 | 0,807158582 | 0,18251219 | 4,42249142 | 9,76E-06 | 0,00010087 | CDK19 |
| ENSGALG00000013051 | 1092,69729 | 0,806546944 | 0,19222288 | 4,19589469 | 2,72E-05 | 0,00024358 | |
| ENSGALG00000005147 | 486,928416 | 0,80606613 | 0,20058324 | 4,0186116 | 5,85E-05 | 0,00047135 | |
| ENSGALG00000017016 | 265,337815 | 0,804980828 | 0,247719 | 3,24957242 | 0,00115579 | 0,00602451 | FAM124A |
| ENSGALG00000015608 | 746,798973 | 0,802634493 | 0,1733654 | 4,62972702 | 3,66E-06 | 4,17E-05 | ADRA2C |
| ENSGALG00000011394 | 463,382496 | 0,801850887 | 0,20513291 | 3,9089335 | 9,27E-05 | 0,0007038 | |
| ENSGALG00000007565 | 677,702896 | 0,798729286 | 0,22132239 | 3,60889512 | 0,0003075 | 0,00198514 | PPP2R2B |
| ENSGALG00000012073 | 470,076315 | 0,797159786 | 0,20392037 | 3,90917193 | 9,26E-05 | 0,0007038 | |
| ENSGALG00000008408 | 1323,03511 | 0,796075417 | 0,15213548 | 5,23267422 | 1,67E-07 | 2,57E-06 | GAB3 |
| ENSGALG00000014122 | 10545,592 | 0,795943411 | 0,13498611 | 5,89648381 | 3,71E-09 | 8,08E-08 | CYB5R3 |
| ENSGALG00000013489 | 630,058844 | 0,794845244 | 0,18721113 | 4,24571568 | 2,18E-05 | 0,00020195 | |
| ENSGALG00000015233 | 5666,23676 | 0,794464087 | 0,15751312 | 5,0437964 | 4,56E-07 | 6,39E-06 | ATP1B1 |
| ENSGALG00000006507 | 1217,4322 | 0,793097237 | 0,15342058 | 5,16943198 | 2,35E-07 | 3,49E-06 | CLMP |
| ENSGALG00000026605 | 1028,74302 | 0,792369586 | 0,15815822 | 5,00998056 | 5,44E-07 | 7,52E-06 | |
| ENSGALG00000007720 | 244,442979 | 0,792327844 | 0,25557464 | 3,1001818 | 0,00193402 | 0,009178 | AIFM3 |
| ENSGALG00000006319 | 831,267138 | 0,791303644 | 0,17376438 | 4,55388859 | 5,27E-06 | 5,80E-05 | |
| ENSGALG00000028576 | 735,470308 | 0,790565069 | 0,23138694 | 3,4166365 | 0,000634 | 0,00365294 | |
| ENSGALG00000010708 | 2290,46376 | 0,790532368 | 0,15028519 | 5,26021489 | 1,44E-07 | 2,24E-06 | ICA1 |
| ENSGALG00000021589 | 1830,74644 | 0,790216265 | 0,177386 | 4,45478374 | 8,40E-06 | 8,83E-05 | RTN4RL1 |

| | | | | | | | |
|---|---|---|---|---|---|---|---|
| ENSGALG00000027458 | 914,195785 | 0,790135397 | 0,16348734 | 4,83300661 | 1,34E-06 | 1,70E-05 | |
| ENSGALG00000014300 | 310,956459 | 0,790002156 | 0,25045648 | 3,15424928 | 0,00160912 | 0,00788629 | |
| ENSGALG00000013862 | 6295,07557 | 0,789845326 | 0,14390944 | 5,48848856 | 4,05E-08 | 7,08E-07 | OLIG3 |
| ENSGALG00000026899 | 678,260512 | 0,789824953 | 0,18654004 | 4,2340774 | 2,29E-05 | 0,00021097 | |
| ENSGALG00000020522 | 2265,96477 | 0,78974705 | 0,15263795 | 5,17398877 | 2,29E-07 | 3,42E-06 | |
| ENSGALG00000009039 | 1670,74492 | 0,789311851 | 0,14682451 | 5,375886 | 7,62E-08 | 1,25E-06 | |
| ENSGALG00000003933 | 4489,5312 | 0,788744436 | 0,13110215 | 6,01625856 | 1,78E-09 | 4,11E-08 | QSOX1 |
| ENSGALG00000013642 | 1029,71125 | 0,788560165 | 0,16199374 | 4,86784356 | 1,13E-06 | 1,46E-05 | |
| ENSGALG00000006453 | 2069,6286 | 0,788464487 | 0,15333542 | 5,14208958 | 2,72E-07 | 3,98E-06 | |
| ENSGALG00000007623 | 3327,22263 | 0,788173273 | 0,13967263 | 5,64300463 | 1,67E-08 | 3,18E-07 | CACNA1G |
| ENSGALG00000024097 | 498,632385 | 0,787003033 | 0,25367596 | 3,10239499 | 0,00191962 | 0,00911918 | |
| ENSGALG00000005849 | 1033,38166 | 0,784505411 | 0,16373067 | 4,79143821 | 1,66E-06 | 2,05E-05 | PPDPF |
| ENSGALG00000000076 | 922,879923 | 0,7843985 | 0,16221807 | 4,83545708 | 1,33E-06 | 1,68E-05 | |
| ENSGALG00000012116 | 675,676944 | 0,783832372 | 0,18759243 | 4,17837954 | 2,94E-05 | 0,00026058 | |
| ENSGALG00000011635 | 890,924795 | 0,783625313 | 0,17783173 | 4,40655507 | 1,05E-05 | 0,00010776 | TFCP2L1 |
| ENSGALG00000027788 | 2064,81163 | 0,782858625 | 0,18478838 | 4,23651437 | 2,27E-05 | 0,00020926 | CPE |
| ENSGALG00000009866 | 599,485373 | 0,782677583 | 0,20855076 | 3,75293566 | 0,00017478 | 0,00122029 | |
| ENSGALG00000020376 | 2337,04095 | 0,781084057 | 0,12991704 | 6,01217582 | 1,83E-09 | 4,20E-08 | |
| ENSGALG00000007357 | 1727,63044 | 0,780757803 | 0,17882552 | 4,36603111 | 1,27E-05 | 0,00012639 | BCL6 |
| ENSGALG00000009407 | 420,644478 | 0,780277328 | 0,21471092 | 3,63408309 | 0,00027897 | 0,00182342 | |
| ENSGALG00000021620 | 400,193416 | 0,780112352 | 0,21582196 | 3,61461062 | 0,0003008 | 0,00194647 | TMEM59L |
| ENSGALG00000005657 | 1485,50445 | 0,780089243 | 0,20993525 | 3,71585638 | 0,00020252 | 0,00137863 | CRHR2 |
| ENSGALG00000017065 | 605,990734 | 0,779643959 | 0,22129554 | 3,52308929 | 0,00042655 | 0,00263018 | |
| ENSGALG00000015379 | 3926,96098 | 0,779239634 | 0,17207727 | 4,5284285 | 5,94E-06 | 6,41E-05 | TAGLN3 |
| ENSGALG00000005539 | 507,508669 | 0,77704495 | 0,19834408 | 3,9176614 | 8,94E-05 | 0,00068237 | |
| ENSGALG00000009612 | 1284,26197 | 0,776962988 | 0,15672769 | 4,95740715 | 7,14E-07 | 9,55E-06 | |
| ENSGALG00000005421 | 527,433616 | 0,776882668 | 0,19330399 | 4,01896854 | 5,85E-05 | 0,00047091 | |
| ENSGALG00000026326 | 2889,79677 | 0,77526969 | 0,14658762 | 5,28878011 | 1,23E-07 | 1,94E-06 | |
| ENSGALG00000002708 | 2271,23083 | 0,772455755 | 0,1693238 | 4,56200355 | 5,07E-06 | 5,60E-05 | LINGO1 |
| ENSGALG00000013404 | 465,070427 | 0,770980594 | 0,20453334 | 3,76946168 | 0,0001636 | 0,00115171 | |
| ENSGALG00000002283 | 1172,58396 | 0,770174106 | 0,17180246 | 4,482905 | 7,36E-06 | 7,80E-05 | MAPKAPK3 |
| ENSGALG00000017208 | 521,560672 | 0,769802089 | 0,19597474 | 3,92806787 | 8,56E-05 | 0,0006572 | AMOTL1 |
| ENSGALG00000027592 | 1174,35152 | 0,765780883 | 0,16676307 | 4,59202906 | 4,39E-06 | 4,91E-05 | ERN2 |
| ENSGALG00000005779 | 2301,58905 | 0,765244932 | 0,12900388 | 5,93195293 | 2,99E-09 | 6,58E-08 | |
| ENSGALG00000006900 | 414,726982 | 0,764663976 | 0,24206393 | 3,15893399 | 0,00158347 | 0,00778022 | |
| ENSGALG00000004323 | 3702,95548 | 0,7637615 | 0,15205572 | 5,02290526 | 5,09E-07 | 7,07E-06 | RASAL2 |
| ENSGALG00000007744 | 1224,40316 | 0,76351713 | 0,16453915 | 4,64033711 | 3,48E-06 | 3,98E-05 | CASS4 |
| ENSGALG00000002069 | 610,168852 | 0,761037525 | 0,20960313 | 3,63085008 | 0,00028249 | 0,00184376 | |
| ENSGALG00000013948 | 736,872876 | 0,760675086 | 0,17278143 | 4,40252804 | 1,07E-05 | 0,00010953 | RASL11B |
| ENSGALG00000007956 | 2266,7266 | 0,759404394 | 0,14550515 | 5,2190893 | 1,80E-07 | 2,75E-06 | PLXDC2 |
| ENSGALG00000007748 | 2468,69748 | 0,758936347 | 0,12655699 | 5,9967954 | 2,01E-09 | 4,56E-08 | SLC16A2 |
| ENSGALG00000004170 | 385,321336 | 0,75875998 | 0,23438122 | 3,23729002 | 0,00120671 | 0,00624926 | ADA |
| ENSGALG00000015688 | 1188,53051 | 0,758515207 | 0,17302156 | 4,38393456 | 1,17E-05 | 0,00011807 | MXD4 |
| ENSGALG00000003115 | 505,204817 | 0,756684773 | 0,20221429 | 3,74199456 | 0,00018257 | 0,00126624 | |
| ENSGALG00000007889 | 769,261981 | 0,756666335 | 0,18466224 | 4,09756933 | 4,18E-05 | 0,00035285 | |
| ENSGALG00000006346 | 1339,32033 | 0,756405715 | 0,1832878 | 4,12687423 | 3,68E-05 | 0,00031627 | CXCL14 |
| ENSGALG00000026153 | 524,420919 | 0,755983234 | 0,23146908 | 3,26602248 | 0,0010907 | 0,00574239 | |
| ENSGALG00000002958 | 2112,95038 | 0,755313619 | 0,16744802 | 4,51073495 | 6,46E-06 | 6,90E-05 | DNER |
| ENSGALG00000001311 | 1301,90092 | 0,752137925 | 0,15185177 | 4,95310607 | 7,30E-07 | 9,72E-06 | |
| ENSGALG00000027905 | 3422,01429 | 0,751625166 | 0,13298701 | 5,65186918 | 1,59E-08 | 3,03E-07 | CSRNP1 |
| ENSGALG00000009439 | 631,532608 | 0,751125582 | 0,2024985 | 3,70928955 | 0,00020784 | 0,00141276 | GPR85 |
| ENSGALG00000029047 | 339,90745 | 0,749178885 | 0,22666604 | 3,30521004 | 0,00094905 | 0,00512549 | TMEM204 |
| ENSGALG00000015729 | 1439,09217 | 0,747405326 | 0,14976 | 4,99068712 | 6,02E-07 | 8,18E-06 | LPAR1 |
| ENSGALG00000001969 | 1252,39504 | 0,746864271 | 0,15500447 | 4,81834019 | 1,45E-06 | 1,82E-05 | |
| ENSGALG00000004420 | 2554,14464 | 0,745430944 | 0,15350993 | 4,85591338 | 1,20E-06 | 1,54E-05 | SDK1 |
| ENSGALG00000018733 | 341,104932 | 0,744919582 | 0,22358708 | 3,33167538 | 0,00086325 | 0,0047277 | |
| ENSGALG00000005063 | 630,430329 | 0,74466275 | 0,19563887 | 3,8063128 | 0,00014105 | 0,0010113 | ST6GALNAC4 |
| ENSGALG00000026919 | 497,57567 | 0,743491927 | 0,20017411 | 3,71422615 | 0,00020383 | 0,00138686 | |
| ENSGALG00000025937 | 424,74226 | 0,743354667 | 0,21925587 | 3,39035244 | 0,00069803 | 0,00397144 | |
| ENSGALG00000012712 | 2097,8069 | 0,742584862 | 0,14980046 | 4,95716018 | 7,15E-07 | 9,56E-06 | RBM24 |
| ENSGALG00000015994 | 1103,99088 | 0,741778145 | 0,15501143 | 4,78531262 | 1,71E-06 | 2,10E-05 | |
| ENSGALG00000001449 | 3789,5374 | 0,740070536 | 0,14515669 | 5,09842522 | 3,42E-07 | 4,93E-06 | STRA6 |
| ENSGALG00000026449 | 3878,57981 | 0,737442369 | 0,13363533 | 5,51831886 | 3,42E-08 | 6,04E-07 | |
| ENSGALG00000002539 | 465,932804 | 0,737264559 | 0,21127231 | 3,4896412 | 0,00048367 | 0,00291895 | KCNN2 |

| | | | | | | | |
|---|---|---|---|---|---|---|---|
| ENSGALG00000008339 | 615,635733 | 0,736061451 | 0,19489251 | 3,77675604 | 0,00015888 | 0,00112083 | |
| ENSGALG00000013002 | 3217,45336 | 0,734684533 | 0,14696976 | 4,99888241 | 5,77E-07 | 7,89E-06 | DAP1 |
| ENSGALG00000019087 | 2799,71591 | 0,734240753 | 0,12953887 | 5,66811147 | 1,44E-08 | 2,78E-07 | RFXAP |
| ENSGALG00000011059 | 526,882943 | 0,733913318 | 0,20211997 | 3,6310777 | 0,00028224 | 0,00184302 | |
| ENSGALG00000010326 | 286,241823 | 0,733495449 | 0,23722679 | 3,09195881 | 0,0019884 | 0,00938825 | FLVCR2 |
| ENSGALG00000008433 | 2910,77768 | 0,732487 | 0,13031486 | 5,62090169 | 1,90E-08 | 3,57E-07 | |
| ENSGALG00000008076 | 5047,29607 | 0,731953468 | 0,12742676 | 5,74411123 | 9,24E-09 | 1,83E-07 | TMEM164 |
| ENSGALG00000006897 | 1904,48459 | 0,73176707 | 0,13708377 | 5,33810137 | 9,39E-08 | 1,52E-06 | |
| ENSGALG00000025916 | 1511,63504 | 0,730226914 | 0,19711664 | 3,70454229 | 0,00021177 | 0,0014359 | |
| ENSGALG00000002651 | 2712,11427 | 0,730025866 | 0,12886096 | 5,6652212 | 1,47E-08 | 2,82E-07 | ST3GAL2 |
| ENSGALG00000012191 | 7671,58272 | 0,72968259 | 0,12563695 | 5,80786629 | 6,33E-09 | 1,30E-07 | |
| ENSGALG00000002451 | 1225,15716 | 0,728956851 | 0,15337655 | 4,75272686 | 2,01E-06 | 2,41E-05 | |
| ENSGALG00000019317 | 6925,16498 | 0,728083244 | 0,12000428 | 6,06714384 | 1,30E-09 | 3,05E-08 | |
| ENSGALG00000000400 | 352,092636 | 0,727285624 | 0,21940045 | 3,31487749 | 0,00091683 | 0,00498112 | |
| ENSGALG00000014910 | 3784,53082 | 0,72588259 | 0,16231129 | 4,47216332 | 7,74E-06 | 8,18E-05 | COLEC12 |
| ENSGALG00000004875 | 3335,27386 | 0,724796191 | 0,1381226 | 5,24748459 | 1,54E-07 | 2,39E-06 | PEMT |
| ENSGALG00000015123 | 798,450057 | 0,722653298 | 0,17223363 | 4,19577343 | 2,72E-05 | 0,00024358 | ZDHHC23 |
| ENSGALG00000026864 | 531,42097 | 0,721498621 | 0,20303604 | 3,55354952 | 0,00038007 | 0,00237478 | ZCCHC24 |
| ENSGALG00000011426 | 2063,22886 | 0,721230387 | 0,16133795 | 4,4703085 | 7,81E-06 | 8,24E-05 | |
| ENSGALG00000000723 | 8396,32677 | 0,72079059 | 0,14050982 | 5,12982366 | 2,90E-07 | 4,21E-06 | STX12 |
| ENSGALG00000002558 | 389,45186 | 0,720642154 | 0,22908371 | 3,14575908 | 0,00165656 | 0,00806657 | |
| ENSGALG00000001459 | 1037,33334 | 0,719793089 | 0,17879018 | 4,02590944 | 5,68E-05 | 0,00045941 | TNNC1 |
| ENSGALG00000007981 | 2067,61165 | 0,719546729 | 0,14285858 | 5,0367766 | 4,73E-07 | 6,61E-06 | CLN6 |
| ENSGALG00000009042 | 1726,61695 | 0,719468223 | 0,14937565 | 4,81650257 | 1,46E-06 | 1,83E-05 | ISM1 |
| ENSGALG00000003855 | 2094,24658 | 0,719100031 | 0,13684545 | 5,25483329 | 1,48E-07 | 2,31E-06 | |
| ENSGALG00000015403 | 10307,4906 | 0,718054416 | 0,13178716 | 5,44859159 | 5,08E-08 | 8,65E-07 | EPHA3 |
| ENSGALG00000007383 | 515,088556 | 0,717575013 | 0,20175921 | 3,55659118 | 0,0003757 | 0,00235287 | HSPB8 |
| ENSGALG00000012792 | 5661,79807 | 0,717520705 | 0,12510356 | 5,73541388 | 9,73E-09 | 1,92E-07 | HIPK2 |
| ENSGALG00000013583 | 619,651043 | 0,717484093 | 0,2169479 | 3,30717227 | 0,00094243 | 0,00509578 | FAM114A1 |
| ENSGALG00000004812 | 420,513925 | 0,714148656 | 0,22761509 | 3,13752767 | 0,00170379 | 0,00826109 | FAM129A |
| ENSGALG00000003340 | 440,9962 | 0,714011033 | 0,21324869 | 3,34825521 | 0,00081322 | 0,00449533 | GBGT1 |
| ENSGALG00000001233 | 1022,31515 | 0,713516886 | 0,17513042 | 4,07420311 | 4,62E-05 | 0,00038331 | |
| ENSGALG00000007244 | 345,610478 | 0,711922899 | 0,2206692 | 3,22619966 | 0,00125446 | 0,00644774 | SIRT4 |
| ENSGALG00000009628 | 330,245656 | 0,711918795 | 0,2307632 | 3,08506204 | 0,0020351 | 0,00958086 | |
| ENSGALG00000003282 | 5419,41789 | 0,711820753 | 0,11116608 | 6,40321912 | 1,52E-10 | 4,05E-09 | STAT5B |
| ENSGALG00000007196 | 4102,69674 | 0,710478048 | 0,12809281 | 5,54658791 | 2,91E-08 | 5,23E-07 | |
| ENSGALG00000003805 | 6589,80394 | 0,710406685 | 0,12973331 | 5,47590036 | 4,35E-08 | 7,53E-07 | PRKG1 |
| ENSGALG00000001130 | 1241,25137 | 0,709264893 | 0,16240514 | 4,36725635 | 1,26E-05 | 0,00012586 | WBSCR17 |
| ENSGALG00000002038 | 769,11793 | 0,706334316 | 0,17558941 | 4,02264751 | 5,75E-05 | 0,00046499 | DENND2C |
| ENSGALG00000028443 | 5716,4355 | 0,705249503 | 0,16905495 | 4,17171751 | 3,02E-05 | 0,00026727 | |
| ENSGALG00000023738 | 705,328876 | 0,704601132 | 0,2001565 | 3,52025109 | 0,00043114 | 0,00265129 | |
| ENSGALG00000010320 | 1626,02269 | 0,704383823 | 0,13846801 | 5,0869789 | 3,64E-07 | 5,19E-06 | PAWR |
| ENSGALG00000026437 | 5573,48841 | 0,703536871 | 0,11568829 | 6,0813142 | 1,19E-09 | 2,80E-08 | |
| ENSGALG00000004735 | 2180,59814 | 0,703327732 | 0,14882879 | 4,72575054 | 2,29E-06 | 2,73E-05 | |
| ENSGALG00000026973 | 1340,81752 | 0,702510997 | 0,14635529 | 4,80003844 | 1,59E-06 | 1,97E-05 | KIF3C |
| ENSGALG00000026873 | 606,998571 | 0,701267796 | 0,1971895 | 3,55631416 | 0,00037609 | 0,00235426 | |
| ENSGALG00000023266 | 1017,69934 | 0,69953934 | 0,17755104 | 3,93993386 | 8,15E-05 | 0,00062871 | |
| ENSGALG00000027058 | 1118,35276 | 0,699526976 | 0,185719 | 3,76658817 | 0,00016549 | 0,00116401 | EMP2 |
| ENSGALG00000020497 | 375,964711 | 0,698148301 | 0,21726001 | 3,21342288 | 0,00131163 | 0,00669232 | |
| ENSGALG00000004647 | 2581,30282 | 0,697164089 | 0,1609537 | 4,33145725 | 1,48E-05 | 0,0001453 | |
| ENSGALG00000015181 | 1358,59834 | 0,696928823 | 0,1980876 | 3,51828604 | 0,00043434 | 0,00266338 | GAREM1 |
| ENSGALG00000028945 | 1325,88323 | 0,696399862 | 0,19729376 | 3,52976114 | 0,00041594 | 0,0025729 | |
| ENSGALG00000007349 | 388,214699 | 0,696333265 | 0,21422064 | 3,25054234 | 0,00115185 | 0,00601091 | RASL12 |
| ENSGALG00000008455 | 4612,74446 | 0,695894902 | 0,1608753 | 4,32567893 | 1,52E-05 | 0,00014811 | |
| ENSGALG00000010094 | 8338,3892 | 0,695566561 | 0,11023706 | 6,30973457 | 2,80E-10 | 7,11E-09 | B4GALT2 |
| ENSGALG00000010331 | 1143,01041 | 0,693784562 | 0,19974706 | 3,47331558 | 0,00051407 | 0,00306708 | MME |
| ENSGALG00000010406 | 684,710723 | 0,693540735 | 0,17759525 | 3,9051762 | 9,42E-05 | 0,00071219 | TMEM63C |
| ENSGALG00000001889 | 934,570663 | 0,693259058 | 0,17268521 | 4,01458276 | 5,96E-05 | 0,00047852 | CARD9 |
| ENSGALG00000008651 | 2483,20041 | 0,693028969 | 0,15778512 | 4,39223263 | 1,12E-05 | 0,00011425 | APMAP |
| ENSGALG00000005103 | 1726,78257 | 0,692645693 | 0,13835836 | 5,00617158 | 5,55E-07 | 7,63E-06 | GDPD1 |
| ENSGALG00000023188 | 975,838184 | 0,692540652 | 0,16000524 | 4,32823725 | 1,50E-05 | 0,00014706 | NANP |
| ENSGALG00000003636 | 432,590451 | 0,690941222 | 0,20908855 | 3,3045388 | 0,00095133 | 0,00513167 | FBLIM1 |
| ENSGALG00000020575 | 447,180257 | 0,690014438 | 0,22107615 | 3,12116177 | 0,00180139 | 0,00863589 | |
| ENSGALG00000023655 | 726,630068 | 0,689103212 | 0,1781324 | 3,86848893 | 0,00010951 | 0,00081033 | |

| | | | | | | | |
|---|---|---|---|---|---|---|---|
| ENSGALG00000003122 | 2359,00023 | 0,68839827 | 0,16356115 | 4,20881277 | 2,57E-05 | 0,0002327 | ZFAND2A |
| ENSGALG00000002749 | 1561,25313 | 0,687223145 | 0,13989637 | 4,91237283 | 9,00E-07 | 1,18E-05 | |
| ENSGALG00000016844 | 1336,96906 | 0,685906484 | 0,15616188 | 4,39227854 | 1,12E-05 | 0,00011425 | RAB20 |
| ENSGALG00000016123 | 816,293249 | 0,685692574 | 0,20965616 | 3,27055776 | 0,00107336 | 0,00566864 | |
| ENSGALG00000002204 | 670,701965 | 0,684953864 | 0,19602604 | 3,49419842 | 0,00047549 | 0,00287595 | SLC7A4 |
| ENSGALG00000014559 | 1808,90594 | 0,684328525 | 0,1507966 | 4,53808981 | 5,68E-06 | 6,17E-05 | |
| ENSGALG00000009075 | 1175,32124 | 0,683878573 | 0,15401179 | 4,44042999 | 8,98E-06 | 9,38E-05 | CD99L2 |
| ENSGALG00000010770 | 1614,38779 | 0,683740122 | 0,16437692 | 4,15958702 | 3,19E-05 | 0,00028005 | SCIN |
| ENSGALG00000009523 | 834,574184 | 0,681144754 | 0,18607441 | 3,66060408 | 0,00025162 | 0,00166626 | CNTN1 |
| ENSGALG00000007251 | 3608,08828 | 0,681060329 | 0,12144401 | 5,60801928 | 2,05E-08 | 3,82E-07 | PCYT2 |
| ENSGALG00000019702 | 11617,3005 | 0,678512546 | 0,13266103 | 5,11463354 | 3,14E-07 | 4,54E-06 | |
| ENSGALG00000000674 | 418,548479 | 0,67836471 | 0,21301294 | 3,18461735 | 0,00144946 | 0,00725001 | MYO1D |
| ENSGALG00000020688 | 589,72193 | 0,678030575 | 0,21326458 | 3,17929294 | 0,00147635 | 0,00733594 | |
| ENSGALG00000026986 | 1312,77213 | 0,67793682 | 0,16864646 | 4,01986992 | 5,82E-05 | 0,00046939 | |
| ENSGALG00000004240 | 977,710505 | 0,676367039 | 0,17381981 | 3,89119644 | 9,98E-05 | 0,00074827 | ORAI1 |
| ENSGALG00000025764 | 5164,94672 | 0,675250138 | 0,12251808 | 5,51143253 | 3,56E-08 | 6,27E-07 | FKBP10 |
| ENSGALG00000002028 | 1196,67299 | 0,675138088 | 0,19853735 | 3,40055964 | 0,00067248 | 0,00384375 | GPRC5B |
| ENSGALG00000012583 | 448,709518 | 0,674342421 | 0,20797125 | 3,24247904 | 0,00118495 | 0,00615061 | IDNK |
| ENSGALG00000016399 | 1323,25053 | 0,673456474 | 0,16094579 | 4,18436848 | 2,86E-05 | 0,00025497 | |
| ENSGALG00000004472 | 11283,0726 | 0,672299012 | 0,12223168 | 5,50020247 | 3,79E-08 | 6,65E-07 | ASTN1 |
| ENSGALG00000005041 | 1267,24433 | 0,67175632 | 0,1650486 | 4,07005162 | 4,70E-05 | 0,00038926 | TMEM35A |
| ENSGALG00000011896 | 1990,38793 | 0,671754963 | 0,13345151 | 5,03370069 | 4,81E-07 | 6,71E-06 | |
| ENSGALG00000007038 | 2302,52795 | 0,670533206 | 0,13522368 | 4,95869653 | 7,10E-07 | 9,50E-06 | |
| ENSGALG00000005069 | 1045,94718 | 0,669583836 | 0,15806848 | 4,23603637 | 2,27E-05 | 0,00020957 | |
| ENSGALG00000000040 | 692,575173 | 0,669045681 | 0,18677509 | 3,58209261 | 0,00034085 | 0,00217059 | |
| ENSGALG00000000920 | 727,437453 | 0,668851002 | 0,19789956 | 3,37974989 | 0,00072552 | 0,00410213 | CGN |
| ENSGALG00000016415 | 1383,10646 | 0,668618298 | 0,18669663 | 3,58130896 | 0,00034188 | 0,00217443 | MAP7D2 |
| ENSGALG00000016617 | 4902,5535 | 0,667497538 | 0,12877865 | 5,18329341 | 2,18E-07 | 3,27E-06 | NCOA1 |
| ENSGALG00000008970 | 735,022626 | 0,666872911 | 0,18064169 | 3,69168889 | 0,00022277 | 0,00150001 | |
| ENSGALG00000006449 | 910,146444 | 0,666661271 | 0,19615763 | 3,39859966 | 0,00067732 | 0,00386978 | |
| ENSGALG00000003457 | 3064,26122 | 0,662964808 | 0,14313215 | 4,63183715 | 3,62E-06 | 4,14E-05 | CNP |
| ENSGALG00000013766 | 661,354996 | 0,662853472 | 0,19762543 | 3,35409001 | 0,00079626 | 0,00443219 | |
| ENSGALG00000010316 | 7099,3915 | 0,662660688 | 0,11945249 | 5,54748328 | 2,90E-08 | 5,21E-07 | FRAS1 |
| ENSGALG00000002775 | 509,116953 | 0,661278251 | 0,20564659 | 3,21560519 | 0,0013017 | 0,00665263 | FA2H |
| ENSGALG00000012710 | 1279,29723 | 0,660750331 | 0,15866836 | 4,16434841 | 3,12E-05 | 0,00027501 | NCS1 |
| ENSGALG00000016411 | 1960,74491 | 0,657856223 | 0,15713168 | 4,18665555 | 2,83E-05 | 0,00025274 | |
| ENSGALG00000006152 | 1208,2078 | 0,657637784 | 0,1593989 | 4,12573602 | 3,70E-05 | 0,00031724 | |
| ENSGALG00000018967 | 1911,68524 | 0,656123394 | 0,18491847 | 3,54817659 | 0,00038791 | 0,00241489 | |
| ENSGALG00000016543 | 3236,91171 | 0,654951402 | 0,17013318 | 3,84963956 | 0,00011829 | 0,00086402 | NHS |
| ENSGALG00000012023 | 986,869481 | 0,654468757 | 0,16148919 | 4,0527095 | 5,06E-05 | 0,00041599 | DACT1 |
| ENSGALG00000003845 | 2106,1901 | 0,653837192 | 0,13713933 | 4,76768555 | 1,86E-06 | 2,26E-05 | MKNK2 |
| ENSGALG00000003589 | 5654,7984 | 0,652278144 | 0,1207315 | 5,40271723 | 6,56E-08 | 1,10E-06 | VTN |
| ENSGALG00000001426 | 1341,21178 | 0,651807691 | 0,14847266 | 4,39008564 | 1,13E-05 | 0,00011521 | |
| ENSGALG00000002403 | 2862,69893 | 0,651643976 | 0,15300671 | 4,25892429 | 2,05E-05 | 0,00019195 | ULK1 |
| ENSGALG00000001956 | 2538,00503 | 0,650817237 | 0,17613054 | 3,69508455 | 0,00021981 | 0,00148303 | OLFML3 |
| ENSGALG00000008120 | 6095,80519 | 0,649872522 | 0,12621953 | 5,14874782 | 2,62E-07 | 3,86E-06 | TLE3 |
| ENSGALG00000024104 | 1479,60024 | 0,64969951 | 0,15049285 | 4,31714531 | 1,58E-05 | 0,00015284 | |
| ENSGALG00000024488 | 3112,83733 | 0,649250333 | 0,1855727 | 3,49863056 | 0,00046765 | 0,00283487 | CD99 |
| ENSGALG00000008763 | 548,608658 | 0,64905793 | 0,2030908 | 3,19590022 | 0,00139395 | 0,00702134 | SSX2IP |
| ENSGALG00000007848 | 1247,63563 | 0,647083546 | 0,1839624 | 3,51747722 | 0,00043567 | 0,00266711 | PTS |
| ENSGALG00000010422 | 5513,26975 | 0,646628908 | 0,1246783 | 5,1863789 | 2,14E-07 | 3,22E-06 | |
| ENSGALG00000010719 | 1138,21283 | 0,646215963 | 0,15983765 | 4,04295223 | 5,28E-05 | 0,00043187 | NEIL3 |
| ENSGALG00000006036 | 6926,72973 | 0,644947298 | 0,14183609 | 4,54713104 | 5,44E-06 | 5,94E-05 | |
| ENSGALG00000003655 | 1118,25271 | 0,643488267 | 0,18487926 | 3,48058661 | 0,00050032 | 0,00300212 | GAS2 |
| ENSGALG00000000402 | 743,970165 | 0,642116084 | 0,18043203 | 3,55876987 | 0,0003726 | 0,00233882 | LOXL2 |
| ENSGALG00000006947 | 23413,053 | 0,64187784 | 0,11085732 | 5,79012569 | 7,03E-09 | 1,43E-07 | BDH1 |
| ENSGALG00000011335 | 567,87108 | 0,64086738 | 0,205863 | 3,11307701 | 0,00185148 | 0,00882628 | |
| ENSGALG00000026159 | 2624,11695 | 0,639768169 | 0,13669036 | 4,68041921 | 2,86E-06 | 3,34E-05 | PDLIM7 |
| ENSGALG00000026295 | 3751,17129 | 0,638993968 | 0,12561216 | 5,08703909 | 3,64E-07 | 5,19E-06 | CDR2L |
| ENSGALG00000000935 | 1038,82432 | 0,638135684 | 0,19518262 | 3,26942888 | 0,00107765 | 0,0056891 | |
| ENSGALG00000028506 | 1185,60599 | 0,637581477 | 0,16691697 | 3,81975226 | 0,00013359 | 0,00096486 | NHSL2 |
| ENSGALG00000003527 | 1639,11059 | 0,636215476 | 0,14708776 | 4,32541425 | 1,52E-05 | 0,00014816 | ARRDC2 |
| ENSGALG00000028756 | 1081,22564 | 0,636133999 | 0,16633022 | 3,82452444 | 0,00013102 | 0,00094737 | |
| ENSGALG00000004585 | 1609,34206 | 0,634633954 | 0,16521991 | 3,84114698 | 0,00012246 | 0,00089065 | |

| | | | | | | | |
|---|---|---|---|---|---|---|---|
| ENSGALG00000008057 | 2849,96428 | 0,63255389 | 0,14696671 | 4,30406235 | 1,68E-05 | 0,0001609 | LLGL2 |
| ENSGALG00000011053 | 10065,6567 | 0,630484727 | 0,10769501 | 5,85435431 | 4,79E-09 | 1,01E-07 | |
| ENSGALG00000008151 | 5862,81637 | 0,629127593 | 0,11865044 | 5,30236208 | 1,14E-07 | 1,81E-06 | WSB2 |
| ENSGALG00000027658 | 594,253982 | 0,628365542 | 0,19060887 | 3,29662282 | 0,00097855 | 0,005237 | TESK1 |
| ENSGALG00000002664 | 1165,54673 | 0,627944771 | 0,15316524 | 4,09978643 | 4,14E-05 | 0,00035053 | |
| ENSGALG00000029000 | 1142,4251 | 0,627547484 | 0,15842199 | 3,96123969 | 7,46E-05 | 0,00058075 | HYAL3 |
| ENSGALG00000028204 | 8948,35307 | 0,626531339 | 0,12182055 | 5,14306756 | 2,70E-07 | 3,97E-06 | |
| ENSGALG00000001100 | 796,408366 | 0,626475621 | 0,18020964 | 3,47637138 | 0,00050825 | 0,00303934 | |
| ENSGALG00000002313 | 2718,417 | 0,626374122 | 0,13464234 | 4,65213353 | 3,29E-06 | 3,79E-05 | |
| ENSGALG00000026762 | 1809,40153 | 0,625670937 | 0,1367941 | 4,5738152 | 4,79E-06 | 5,34E-05 | |
| ENSGALG00000008728 | 894,547623 | 0,625207283 | 0,17070647 | 3,66246966 | 0,0002498 | 0,00165578 | PTER |
| ENSGALG00000011856 | 1033,65184 | 0,624422163 | 0,15756316 | 3,96299593 | 7,40E-05 | 0,00057782 | FMC1 |
| ENSGALG00000007267 | 4451,86283 | 0,623851268 | 0,11702334 | 5,33099856 | 9,77E-08 | 1,57E-06 | MVD |
| ENSGALG00000014890 | 1413,33309 | 0,620763059 | 0,14626866 | 4,24399216 | 2,20E-05 | 0,00020309 | |
| ENSGALG00000000426 | 554,757236 | 0,620684473 | 0,19913485 | 3,11690528 | 0,0018276 | 0,00873694 | |
| ENSGALG00000006114 | 909,882198 | 0,61972927 | 0,17107416 | 3,62257682 | 0,00029168 | 0,0018965 | |
| ENSGALG00000011900 | 1365,07707 | 0,61861917 | 0,17981254 | 3,44035605 | 0,00058095 | 0,00339761 | NAGA |
| ENSGALG00000007076 | 2564,35145 | 0,617550884 | 0,14376263 | 4,29562882 | 1,74E-05 | 0,00016632 | CCNY |
| ENSGALG00000007381 | 1220,44249 | 0,615500074 | 0,15532009 | 3,96278456 | 7,41E-05 | 0,000578 | SERPING1 |
| ENSGALG00000028543 | 1301,49454 | 0,614319385 | 0,1647996 | 3,72767515 | 0,00019325 | 0,00132563 | FGFR4 |
| ENSGALG00000015734 | 903,95178 | 0,61429478 | 0,19839853 | 3,09626685 | 0,00195974 | 0,00928387 | |
| ENSGALG00000016728 | 5650,40168 | 0,613362339 | 0,11515602 | 5,32635913 | 1,00E-07 | 1,61E-06 | RP2 |
| ENSGALG00000016092 | 954,718038 | 0,611645343 | 0,16439834 | 3,72050802 | 0,00019882 | 0,00135687 | |
| ENSGALG00000011975 | 2830,82544 | 0,61116717 | 0,12275633 | 4,97870188 | 6,40E-07 | 8,65E-06 | |
| ENSGALG00000016143 | 5447,09101 | 0,607480677 | 0,17132527 | 3,54577386 | 0,00039146 | 0,0024359 | |
| ENSGALG00000008512 | 665,497169 | 0,606896507 | 0,19319454 | 3,14137499 | 0,00168157 | 0,00816786 | |
| ENSGALG00000008272 | 1496,82423 | 0,6063144 | 0,15709843 | 3,8594554 | 0,00011364 | 0,00083633 | TMED3 |
| ENSGALG00000015414 | 1285,1197 | 0,605396806 | 0,18042117 | 3,35546434 | 0,00079232 | 0,00441384 | NSUN3 |
| ENSGALG00000002930 | 5292,49241 | 0,603605896 | 0,13766556 | 4,38458178 | 1,16E-05 | 0,00011781 | |
| ENSGALG00000005303 | 729,474387 | 0,603106981 | 0,18273634 | 3,30042173 | 0,0009654 | 0,00518701 | CCDC13 |
| ENSGALG00000006742 | 8579,7495 | 0,60288418 | 0,10750963 | 5,60772266 | 2,05E-08 | 3,82E-07 | |
| ENSGALG00000013964 | 1158,34472 | 0,602864404 | 0,163189 | 3,69427107 | 0,00022052 | 0,00148632 | SPATA18 |
| ENSGALG00000000739 | 566,84692 | 0,602546305 | 0,18860198 | 3,19480372 | 0,00139926 | 0,00704546 | |
| ENSGALG00000009047 | 3574,12193 | 0,601534497 | 0,13925729 | 4,3195906 | 1,56E-05 | 0,0001517 | YPEL5 |
| ENSGALG00000005628 | 1589,65421 | 0,601333554 | 0,18106388 | 3,3211128 | 0,00089659 | 0,00488488 | COL9A3 |
| ENSGALG00000029021 | 4810,26751 | 0,600451025 | 0,1243158 | 4,83004601 | 1,37E-06 | 1,72E-05 | |
| ENSGALG00000023435 | 807,96294 | 0,600270173 | 0,18996812 | 3,15984693 | 0,00157852 | 0,0077671 | GATM |
| ENSGALG00000023876 | 5804,17122 | 0,598475769 | 0,12582855 | 4,75627963 | 1,97E-06 | 2,38E-05 | |
| ENSGALG00000008475 | 1732,98273 | 0,598095279 | 0,145561 | 4,10889791 | 3,98E-05 | 0,00033935 | ARRDC1 |
| ENSGALG00000013167 | 14635,2035 | 0,597899753 | 0,11791074 | 5,07078291 | 3,96E-07 | 5,61E-06 | SDHA |
| ENSGALG00000007801 | 1966,88294 | 0,597201094 | 0,14607757 | 4,08824628 | 4,35E-05 | 0,00036327 | |
| ENSGALG00000027450 | 1958,10116 | 0,596865465 | 0,13602137 | 4,3880272 | 1,14E-05 | 0,00011605 | |
| ENSGALG00000002818 | 12498,9505 | 0,595523293 | 0,12527412 | 4,75376167 | 2,00E-06 | 2,41E-05 | |
| ENSGALG00000008285 | 2205,39659 | 0,594659875 | 0,13357092 | 4,4520161 | 8,51E-06 | 8,94E-05 | NAPEPLD |
| ENSGALG00000026756 | 608,701473 | 0,593058479 | 0,18596131 | 3,1891498 | 0,00142692 | 0,00716353 | |
| ENSGALG00000011145 | 3513,15874 | 0,5922185 | 0,11951587 | 4,95514543 | 7,23E-07 | 9,63E-06 | TRIL |
| ENSGALG00000026180 | 4122,30438 | 0,591162305 | 0,14254212 | 4,14728156 | 3,36E-05 | 0,00029326 | |
| ENSGALG00000005999 | 5083,90869 | 0,591144605 | 0,12187871 | 4,85026945 | 1,23E-06 | 1,57E-05 | ABTB1 |
| ENSGALG00000013436 | 9895,67369 | 0,590008237 | 0,1280495 | 4,60765752 | 4,07E-06 | 4,58E-05 | |
| ENSGALG00000013742 | 6102,59896 | 0,589905166 | 0,14728573 | 4,0051753 | 6,20E-05 | 0,00049428 | EZR |
| ENSGALG00000025745 | 2020,21836 | 0,589609597 | 0,16557075 | 3,56107338 | 0,00036934 | 0,0023216 | |
| ENSGALG00000026081 | 3861,36621 | 0,588777417 | 0,1223416 | 4,81256939 | 1,49E-06 | 1,86E-05 | RBM38 |
| ENSGALG00000000464 | 6973,12042 | 0,587838297 | 0,11000425 | 5,34377783 | 9,10E-08 | 1,48E-06 | |
| ENSGALG00000005102 | 2254,46112 | 0,58732124 | 0,16766068 | 3,50303501 | 0,00045999 | 0,00279338 | CDH4 |
| ENSGALG00000016594 | 1080,56836 | 0,58674351 | 0,17030861 | 3,44517825 | 0,00057068 | 0,00334619 | |
| ENSGALG00000010934 | 813,601759 | 0,586239182 | 0,17064205 | 3,43549068 | 0,00059148 | 0,00345031 | |
| ENSGALG00000025822 | 1186,62395 | 0,586217768 | 0,19001165 | 3,08516751 | 0,00203438 | 0,00958078 | CYP1B1 |
| ENSGALG00000017558 | 835,379673 | 0,585776262 | 0,17286518 | 3,38863076 | 0,00070243 | 0,00398832 | |
| ENSGALG00000002169 | 5307,04919 | 0,58477373 | 0,13930071 | 4,19792355 | 2,69E-05 | 0,0002424 | PLEKHB2 |
| ENSGALG00000025738 | 2313,93547 | 0,584187633 | 0,13782283 | 4,23868547 | 2,25E-05 | 0,00020753 | |
| ENSGALG00000015938 | 2888,98507 | 0,583396297 | 0,14475468 | 4,03024131 | 5,57E-05 | 0,00045237 | |
| ENSGALG00000001574 | 3769,69235 | 0,583342794 | 0,12963845 | 4,49976697 | 6,80E-06 | 7,24E-05 | |
| ENSGALG00000014242 | 1647,30473 | 0,583190799 | 0,15565274 | 3,74674285 | 0,00017915 | 0,0012457 | |
| ENSGALG00000008999 | 3276,84717 | 0,583129446 | 0,15516573 | 3,75810727 | 0,0001712 | 0,00119904 | AFAP1L2 |

| | | | | | | | |
|---|---|---|---|---|---|---|---|
| ENSGALG00000010706 | 6618,96251 | 0,582943162 | 0,11294938 | 5,16110101 | 2,46E-07 | 3,64E-06 | |
| ENSGALG00000004844 | 6201,27995 | 0,581688828 | 0,14767508 | 3,93897768 | 8,18E-05 | 0,00063051 | |
| ENSGALG00000009604 | 5450,24379 | 0,581479374 | 0,13237918 | 4,39252905 | 1,12E-05 | 0,00011425 | DBX2 |
| ENSGALG00000001446 | 6585,16039 | 0,581431635 | 0,15010679 | 3,8734532 | 0,0001073 | 0,00079659 | |
| ENSGALG00000023768 | 4728,02752 | 0,58136874 | 0,11646418 | 4,99182452 | 5,98E-07 | 8,14E-06 | |
| ENSGALG00000002707 | 937,598421 | 0,58133458 | 0,18027574 | 3,22469665 | 0,00126106 | 0,00647656 | CHRNB2 |
| ENSGALG00000029020 | 1577,39916 | 0,581217481 | 0,17198927 | 3,37938213 | 0,00072649 | 0,00410592 | PLPP4 |
| ENSGALG00000028478 | 1030,91271 | 0,580829913 | 0,18242777 | 3,18388976 | 0,0014531 | 0,00725779 | |
| ENSGALG00000007531 | 904,342369 | 0,57752222 | 0,17317755 | 3,33485613 | 0,00085344 | 0,00468152 | ZFYVE27 |
| ENSGALG00000006054 | 759,821789 | 0,576882549 | 0,18206756 | 3,1685082 | 0,00153223 | 0,00756944 | CALCA |
| ENSGALG00000022926 | 4686,66572 | 0,576442566 | 0,12615332 | 4,56938097 | 4,89E-06 | 5,45E-05 | GPR162 |
| ENSGALG00000013185 | 3965,04447 | 0,575558846 | 0,14856351 | 3,87416023 | 0,00010699 | 0,00079645 | |
| ENSGALG00000000869 | 855,333756 | 0,574690811 | 0,17351236 | 3,31210309 | 0,00092597 | 0,00502477 | |
| ENSGALG00000006885 | 6383,54265 | 0,574664846 | 0,12640826 | 4,54610186 | 5,46E-06 | 5,96E-05 | |
| ENSGALG00000015136 | 2417,80657 | 0,573614313 | 0,16092955 | 3,56438149 | 0,00036472 | 0,0022957 | ILDR1 |
| ENSGALG00000015744 | 3529,13316 | 0,573305804 | 0,12554323 | 4,56660089 | 4,96E-06 | 5,50E-05 | |
| ENSGALG00000001489 | 8334,32619 | 0,572451499 | 0,12945603 | 4,42197621 | 9,78E-06 | 0,000101 | SSU72 |
| ENSGALG00000014911 | 1071,97212 | 0,572338387 | 0,15722716 | 3,6402005 | 0,00027243 | 0,00178235 | ARL15 |
| ENSGALG00000015351 | 2719,2778 | 0,572206068 | 0,17042313 | 3,35756104 | 0,00078633 | 0,00439308 | CBLB |
| ENSGALG00000025818 | 8546,90555 | 0,571475138 | 0,10995302 | 5,19744834 | 2,02E-07 | 3,06E-06 | |
| ENSGALG00000005549 | 1034,75448 | 0,569387138 | 0,15615315 | 3,64633795 | 0,000266 | 0,0017479 | CUEDC1 |
| ENSGALG00000024153 | 1062,25024 | 0,56872295 | 0,17097205 | 3,326409 | 0,00087973 | 0,00480831 | |
| ENSGALG00000000186 | 787,093258 | 0,568415238 | 0,17712851 | 3,20905553 | 0,00133172 | 0,00677301 | MTCH1 |
| ENSGALG00000001146 | 1103,97969 | 0,568231367 | 0,16866857 | 3,36892268 | 0,00075463 | 0,00423331 | UBXN6 |
| ENSGALG00000012788 | 1309,52475 | 0,567748724 | 0,16133631 | 3,51903868 | 0,00043311 | 0,00265983 | |
| ENSGALG00000006873 | 2079,27658 | 0,565449117 | 0,13107514 | 4,3139312 | 1,60E-05 | 0,00015464 | PHRF1 |
| ENSGALG00000007093 | 1910,94783 | 0,563647311 | 0,15313163 | 3,68080257 | 0,0002325 | 0,00155583 | CYTH1 |
| ENSGALG00000002778 | 10757,9738 | 0,563240809 | 0,10999952 | 5,12039322 | 3,05E-07 | 4,42E-06 | |
| ENSGALG00000028605 | 2035,83961 | 0,561167945 | 0,15780973 | 3,55597819 | 0,00037658 | 0,00235619 | |
| ENSGALG00000002681 | 1896,59526 | 0,559637304 | 0,14922639 | 3,75025697 | 0,00017665 | 0,00123277 | |
| ENSGALG00000016152 | 7208,33853 | 0,558712272 | 0,12592253 | 4,43695234 | 9,12E-06 | 9,51E-05 | |
| ENSGALG00000010307 | 1309,92001 | 0,558082008 | 0,16022503 | 3,48311375 | 0,00049562 | 0,00297918 | ACYP1 |
| ENSGALG00000010614 | 5022,37444 | 0,557460992 | 0,12556393 | 4,4396588 | 9,01E-06 | 9,41E-05 | SLC25A4 |
| ENSGALG00000001682 | 1516,35079 | 0,557286636 | 0,15010501 | 3,71264502 | 0,0002051 | 0,00139485 | FBXL20 |
| ENSGALG00000001238 | 3616,01439 | 0,556401234 | 0,12857642 | 4,32739711 | 1,51E-05 | 0,00014747 | STX8 |
| ENSGALG00000002286 | 73998,0221 | 0,555324016 | 0,11102707 | 5,00169946 | 5,68E-07 | 7,79E-06 | |
| ENSGALG00000025787 | 2442,89077 | 0,554206894 | 0,13003561 | 4,2619625 | 2,03E-05 | 0,00018949 | |
| ENSGALG00000004849 | 4017,82732 | 0,553445556 | 0,11594023 | 4,77354206 | 1,81E-06 | 2,21E-05 | WSCD2 |
| ENSGALG00000001766 | 1421,20567 | 0,552758547 | 0,17371857 | 3,1819197 | 0,00146302 | 0,0072892 | |
| ENSGALG00000010982 | 4422,55083 | 0,55035484 | 0,13935769 | 3,94922475 | 7,84E-05 | 0,00060618 | |
| ENSGALG00000000316 | 1699,55338 | 0,549437781 | 0,1376171 | 3,99251106 | 6,54E-05 | 0,00051769 | |
| ENSGALG00000027194 | 1293,57158 | 0,548094206 | 0,15171291 | 3,61270652 | 0,00030302 | 0,00195897 | |
| ENSGALG00000016166 | 3894,34548 | 0,547970517 | 0,13938637 | 3,93130636 | 8,45E-05 | 0,0006489 | |
| ENSGALG00000006663 | 9717,81848 | 0,547489923 | 0,11710931 | 4,67503318 | 2,94E-06 | 3,41E-05 | ARHGEF12 |
| ENSGALG00000001966 | 2313,19503 | 0,545673399 | 0,14563945 | 3,74674179 | 0,00017915 | 0,0012457 | |
| ENSGALG00000001864 | 1163,99304 | 0,545343287 | 0,16861317 | 3,23428649 | 0,00121947 | 0,00630098 | C9orf72 |
| ENSGALG00000006598 | 2327,66634 | 0,544504824 | 0,13759012 | 3,95744117 | 7,58E-05 | 0,00058794 | SORL1 |
| ENSGALG00000015107 | 1737,16005 | 0,542439326 | 0,13565302 | 3,99872658 | 6,37E-05 | 0,00050646 | PIP5K1B |
| ENSGALG00000008860 | 2251,91448 | 0,541261109 | 0,1571898 | 3,44336028 | 0,00057453 | 0,00336442 | LONRF3 |
| ENSGALG00000005315 | 1118,21521 | 0,540760838 | 0,15986155 | 3,38268231 | 0,00071782 | 0,00406365 | RPUSD1 |
| ENSGALG00000004033 | 1001,88457 | 0,540669975 | 0,16471038 | 3,28254945 | 0,00102873 | 0,00546475 | GALNT10 |
| ENSGALG00000022713 | 4245,74271 | 0,539793648 | 0,1498998 | 3,60102975 | 0,00031696 | 0,00203557 | |
| ENSGALG00000025800 | 976,781593 | 0,538353026 | 0,16267943 | 3,30928757 | 0,00093534 | 0,00506145 | |
| ENSGALG00000028513 | 5510,29846 | 0,53711532 | 0,11230935 | 4,7824632 | 1,73E-06 | 2,12E-05 | |
| ENSGALG00000015437 | 3953,66098 | 0,536831128 | 0,13281019 | 4,04209288 | 5,30E-05 | 0,00043291 | |
| ENSGALG00000007389 | 2084,84126 | 0,53455468 | 0,13272068 | 4,02766676 | 5,63E-05 | 0,00045653 | ANKRD40 |
| ENSGALG00000016167 | 2827,43941 | 0,530005093 | 0,12787412 | 4,14474083 | 3,40E-05 | 0,00029577 | |
| ENSGALG00000023189 | 809,880728 | 0,529094048 | 0,16742381 | 3,16020792 | 0,00157657 | 0,00776028 | UMAD1 |
| ENSGALG00000009433 | 12972,2996 | 0,528259009 | 0,13672663 | 3,8636145 | 0,00011172 | 0,00082355 | BAG3 |
| ENSGALG00000015372 | 6118,12247 | 0,526646329 | 0,10985979 | 4,79380423 | 1,64E-06 | 2,03E-05 | ATP5I |
| ENSGALG00000024346 | 1302,16175 | 0,525612903 | 0,1486838 | 3,53510544 | 0,00040761 | 0,00252716 | |
| ENSGALG00000015306 | 3107,54178 | 0,524586766 | 0,14678095 | 3,57394315 | 0,00035165 | 0,00222785 | OSTM1 |
| ENSGALG00000026545 | 5534,24488 | 0,524540794 | 0,12628499 | 4,15362728 | 3,27E-05 | 0,00028597 | |
| ENSGALG00000002777 | 12410,374 | 0,524514209 | 0,1049816 | 4,99624903 | 5,85E-07 | 7,98E-06 | |

| | | | | | | | |
|---|---|---|---|---|---|---|---|
| ENSGALG00000015326 | 10708,8753 | 0,524363909 | 0,11790787 | 4,44723436 | 8,70E-06 | 9,12E-05 | CLTA |
| ENSGALG00000014846 | 1408,26305 | 0,524315684 | 0,14757312 | 3,55292141 | 0,00038098 | 0,00237937 | |
| ENSGALG00000011945 | 1101,05327 | 0,524157211 | 0,16753074 | 3,12872254 | 0,00175568 | 0,00846159 | |
| ENSGALG00000006901 | 2767,40121 | 0,523326668 | 0,12428389 | 4,21073609 | 2,55E-05 | 0,00023088 | AATK |
| ENSGALG00000026113 | 4499,07114 | 0,523056553 | 0,14658052 | 3,56839051 | 0,00035918 | 0,0022661 | |
| ENSGALG00000000433 | 145504,42 | 0,523011809 | 0,10274265 | 5,09050316 | 3,57E-07 | 5,11E-06 | |
| ENSGALG00000016886 | 6214,42839 | 0,522545779 | 0,11023413 | 4,7403267 | 2,13E-06 | 2,56E-05 | FARP1 |
| ENSGALG00000008367 | 12605,915 | 0,522250337 | 0,11987187 | 4,35673805 | 1,32E-05 | 0,00013139 | MDK |
| ENSGALG00000008911 | 1167,8871 | 0,520705327 | 0,16373625 | 3,18014692 | 0,001472 | 0,00732237 | RNF208 |
| ENSGALG00000023338 | 5372,42979 | 0,520605081 | 0,12794507 | 4,06897339 | 4,72E-05 | 0,00039083 | CBX2 |
| ENSGALG00000001944 | 1567,50719 | 0,519218499 | 0,15022432 | 3,45628783 | 0,00054767 | 0,00323494 | TBC1D2 |
| ENSGALG00000000059 | 125389,821 | 0,519152064 | 0,12052089 | 4,30756896 | 1,65E-05 | 0,00015871 | TUBB3 |
| ENSGALG00000005346 | 2351,81671 | 0,518505546 | 0,12959656 | 4,00092074 | 6,31E-05 | 0,00050208 | PPIC |
| ENSGALG00000028033 | 2016,60758 | 0,517359519 | 0,13377729 | 3,86731945 | 0,00011004 | 0,0008129 | |
| ENSGALG00000013024 | 1376,29845 | 0,516485359 | 0,15574832 | 3,31615365 | 0,00091266 | 0,00496041 | |
| ENSGALG00000028567 | 2485,76766 | 0,515072598 | 0,12787307 | 4,02799904 | 5,63E-05 | 0,00045616 | MYL9 |
| ENSGALG00000001998 | 1765,57571 | 0,514536405 | 0,13830499 | 3,72030241 | 0,00019898 | 0,00135687 | R3HDM4 |
| ENSGALG00000012089 | 5360,59071 | 0,513995299 | 0,12096528 | 4,24911439 | 2,15E-05 | 0,00019918 | C14orf37 |
| ENSGALG00000013708 | 5397,09415 | 0,513478256 | 0,13150987 | 3,90448458 | 9,44E-05 | 0,00071344 | CYB5A |
| ENSGALG00000029159 | 2974,1732 | 0,51305649 | 0,12916503 | 3,97210068 | 7,12E-05 | 0,00056002 | |
| ENSGALG00000005901 | 2799,23507 | 0,510684045 | 0,13243521 | 3,85610467 | 0,00011521 | 0,00084513 | PCMTD2 |
| ENSGALG00000023786 | 1008,22899 | 0,510469642 | 0,16220256 | 3,14711206 | 0,00164892 | 0,00803796 | |
| ENSGALG00000017053 | 1433,26653 | 0,50974214 | 0,14651068 | 3,47921488 | 0,00050289 | 0,00301355 | SPG20 |
| ENSGALG00000004741 | 5108,39092 | 0,509578858 | 0,14467631 | 3,52219976 | 0,00042798 | 0,00263783 | EPHB2 |
| ENSGALG00000012644 | 1843,51714 | 0,509005651 | 0,13724043 | 3,70886091 | 0,00020819 | 0,00141445 | |
| ENSGALG00000029036 | 2011,06708 | 0,506773529 | 0,16255236 | 3,11760184 | 0,00182329 | 0,00871938 | |
| ENSGALG00000011324 | 77589,574 | 0,50624037 | 0,10393882 | 4,8705612 | 1,11E-06 | 1,44E-05 | |
| ENSGALG00000005757 | 2287,1727 | 0,505955901 | 0,14803896 | 3,41772137 | 0,00063148 | 0,00363995 | IRF8 |
| ENSGALG00000012670 | 2710,27627 | 0,505329169 | 0,13956275 | 3,62080263 | 0,00029369 | 0,00190591 | |
| ENSGALG00000028145 | 4237,96065 | 0,505320547 | 0,15549944 | 3,24966153 | 0,00115542 | 0,00602451 | |
| ENSGALG00000027891 | 10377,4455 | 0,504324315 | 0,10846652 | 4,64958504 | 3,33E-06 | 3,83E-05 | |
| ENSGALG00000024295 | 3217,43815 | 0,504249612 | 0,13242093 | 3,80792978 | 0,00014014 | 0,00100577 | |
| ENSGALG00000008631 | 3955,10092 | 0,503418102 | 0,13183167 | 3,81864307 | 0,00013419 | 0,00096819 | TYRO3 |
| ENSGALG00000005811 | 1644,05046 | 0,503226947 | 0,1577089 | 3,19085955 | 0,0014185 | 0,00712916 | |
| ENSGALG00000025977 | 2572,61094 | 0,502103804 | 0,12961035 | 3,87394827 | 0,00010709 | 0,00079659 | GADD45A |
| ENSGALG00000024398 | 21096,2288 | 0,500531072 | 0,11622841 | 4,30644352 | 1,66E-05 | 0,0001594 | RPS28 |
| ENSGALG00000004851 | 3943,15715 | 0,500191453 | 0,14446169 | 3,4624504 | 0,00053528 | 0,00317415 | |
| ENSGALG00000008341 | 3732,2674 | 0,499807137 | 0,12743693 | 3,92199613 | 8,78E-05 | 0,00067247 | FURIN |
| ENSGALG00000011326 | 2703,92213 | 0,4983963 | 0,1426307 | 3,49431285 | 0,00047528 | 0,00287595 | STK16 |
| ENSGALG00000000842 | 2625,68946 | 0,49824929 | 0,14971545 | 3,3279751 | 0,0008748 | 0,0047852 | |
| ENSGALG00000007370 | 4047,93897 | 0,4969378 | 0,12075859 | 4,11513402 | 3,87E-05 | 0,00033124 | CLP1 |
| ENSGALG00000012947 | 2224,14952 | 0,495800698 | 0,13770491 | 3,60045778 | 0,00031766 | 0,00203909 | |
| ENSGALG00000013067 | 8815,63453 | 0,494753671 | 0,11635485 | 4,25211024 | 2,12E-05 | 0,00019721 | |
| ENSGALG00000000527 | 6471,78085 | 0,494652161 | 0,11959043 | 4,13621848 | 3,53E-05 | 0,00030619 | |
| ENSGALG00000012540 | 1788,92176 | 0,494557337 | 0,13808033 | 3,58166386 | 0,00034141 | 0,00217314 | RBFOX2 |
| ENSGALG00000011778 | 2592,19321 | 0,494085012 | 0,14681864 | 3,36527439 | 0,00076468 | 0,00428615 | |
| ENSGALG00000027824 | 1654,69459 | 0,493225579 | 0,14677677 | 3,36037904 | 0,00077836 | 0,00435387 | |
| ENSGALG00000005897 | 4766,71418 | 0,493064333 | 0,12620964 | 3,90670892 | 9,36E-05 | 0,00070927 | HMCES |
| ENSGALG00000012816 | 2261,40613 | 0,492973169 | 0,15659783 | 3,14802046 | 0,0016438 | 0,00802229 | SLC22A23 |
| ENSGALG00000005071 | 6694,01614 | 0,492854493 | 0,11371723 | 4,33403516 | 1,46E-05 | 0,00014402 | MPP1 |
| ENSGALG00000000336 | 3779,81285 | 0,492268059 | 0,13944285 | 3,53024962 | 0,00041517 | 0,00257049 | PP1M M21 subunit |
| ENSGALG00000004902 | 4281,00793 | 0,491426093 | 0,11754092 | 4,18089353 | 2,90E-05 | 0,00025805 | TOM1L2 |
| ENSGALG00000011894 | 1400,22178 | 0,491394089 | 0,14545816 | 3,37825034 | 0,00072949 | 0,00411775 | |
| ENSGALG00000017076 | 6428,3314 | 0,490620681 | 0,14677761 | 3,34261258 | 0,00082994 | 0,00457043 | B3GLCT |
| ENSGALG00000008409 | 5336,4487 | 0,490103959 | 0,14311325 | 3,42458839 | 0,00061573 | 0,00357339 | |
| ENSGALG00000011531 | 2214,44297 | 0,486736013 | 0,15147264 | 3,2133593 | 0,00131192 | 0,00669232 | APAF1 |
| ENSGALG00000006915 | 1392,13905 | 0,485309324 | 0,15555376 | 3,11988167 | 0,00180924 | 0,00866613 | |
| ENSGALG00000000947 | 2256,10973 | 0,482847185 | 0,14942146 | 3,23144465 | 0,00123166 | 0,00635192 | |
| ENSGALG00000014581 | 2314,87998 | 0,482286067 | 0,13822045 | 3,48925269 | 0,00048437 | 0,0029219 | BORCS8 |
| ENSGALG00000011738 | 6621,78437 | 0,482221251 | 0,11823294 | 4,07856946 | 4,53E-05 | 0,00037711 | ARHGDIB |
| ENSGALG00000027688 | 6057,20574 | 0,481915396 | 0,11816406 | 4,07835855 | 4,54E-05 | 0,00037722 | NDUFB4 |
| ENSGALG00000005713 | 4508,08916 | 0,480930735 | 0,11431181 | 4,2071834 | 2,59E-05 | 0,00023392 | NCKIPSD |
| ENSGALG00000003405 | 5095,98918 | 0,480657067 | 0,11244145 | 4,27473194 | 1,91E-05 | 0,00018043 | BSDC1 |

| | | | | | | | |
|---|---|---|---|---|---|---|---|
| ENSGALG00000002033 | 6624,42072 | 0,480537464 | 0,11110056 | 4,32524788 | 1,52E-05 | 0,00014817 | NDUFB6 |
| ENSGALG00000029135 | 7403,52828 | 0,479980361 | 0,12965053 | 3,70210873 | 0,00021381 | 0,00144758 | GAS41 |
| ENSGALG00000011436 | 3916,77111 | 0,477975314 | 0,1192676 | 4,00758713 | 6,13E-05 | 0,00049041 | TMBIM1 |
| ENSGALG00000003466 | 4818,27592 | 0,477695903 | 0,13420724 | 3,55939007 | 0,00037172 | 0,00233545 | AKR1B1L |
| ENSGALG00000001606 | 1726,36227 | 0,477443775 | 0,14690617 | 3,2499912 | 0,00115409 | 0,00602026 | INPP5B |
| ENSGALG00000000785 | 1523,67138 | 0,476932316 | 0,14689834 | 3,24668282 | 0,00116758 | 0,00607411 | HYAL1 |
| ENSGALG00000014765 | 2914,68173 | 0,476665637 | 0,12457546 | 3,82632052 | 0,00013007 | 0,00094149 | TRIM23 |
| ENSGALG00000000943 | 4846,50588 | 0,473716041 | 0,11278203 | 4,20027936 | 2,67E-05 | 0,00024021 | WDTC1 |
| ENSGALG00000006588 | 1949,86566 | 0,473632802 | 0,13867563 | 3,41540052 | 0,00063688 | 0,00366533 | LRRC75B |
| ENSGALG00000023157 | 3625,87384 | 0,472554642 | 0,14223538 | 3,32234238 | 0,00089265 | 0,00487307 | REEP5 |
| ENSGALG00000006702 | 2572,80575 | 0,471728545 | 0,1314835 | 3,58773935 | 0,00033356 | 0,00213052 | MFGE8 |
| ENSGALG00000003359 | 6429,54572 | 0,469661247 | 0,12003738 | 3,91262507 | 9,13E-05 | 0,00069482 | |
| ENSGALG00000027377 | 1588,86326 | 0,469161575 | 0,14290687 | 3,28298835 | 0,00102713 | 0,00545838 | |
| ENSGALG00000013511 | 6097,6262 | 0,469095656 | 0,12052644 | 3,89205602 | 9,94E-05 | 0,00074604 | ANKRA2 |
| ENSGALG00000003348 | 1706,40697 | 0,468370054 | 0,13812202 | 3,39098761 | 0,00069641 | 0,0039639 | |
| ENSGALG00000028342 | 3684,92319 | 0,4672884 | 0,14052674 | 3,32526324 | 0,00088335 | 0,00482617 | |
| ENSGALG00000004581 | 7502,73386 | 0,466855623 | 0,11693534 | 3,99242555 | 6,54E-05 | 0,00051769 | NDST1 |
| ENSGALG00000007238 | 3294,46471 | 0,466556765 | 0,1396865 | 3,34002762 | 0,0008377 | 0,0046082 | |
| ENSGALG00000001556 | 1997,40951 | 0,466292104 | 0,13625952 | 3,42208821 | 0,00062142 | 0,00359569 | |
| ENSGALG00000009655 | 1343,41122 | 0,465196113 | 0,14630894 | 3,17954671 | 0,00147506 | 0,0073322 | AADAT |
| ENSGALG00000002485 | 4126,42787 | 0,464623616 | 0,12758859 | 3,64157666 | 0,00027097 | 0,00177456 | PHKG1 |
| ENSGALG00000011474 | 2398,5188 | 0,463345487 | 0,1386209 | 3,34253707 | 0,00083016 | 0,00457043 | |
| ENSGALG00000013698 | 1914,38769 | 0,462883571 | 0,13997841 | 3,30682108 | 0,00094361 | 0,00510015 | |
| ENSGALG00000007582 | 9089,3986 | 0,462276159 | 0,11185396 | 4,13285464 | 3,58E-05 | 0,00030983 | DPYSL3 |
| ENSGALG00000003976 | 1632,25763 | 0,461042882 | 0,14423767 | 3,19641107 | 0,00139149 | 0,0070167 | BCORL1 |
| ENSGALG00000024391 | 9415,82069 | 0,460947942 | 0,11472094 | 4,01799317 | 5,87E-05 | 0,00047231 | CACTIN |
| ENSGALG00000024034 | 3571,45047 | 0,460548751 | 0,12312106 | 3,74061704 | 0,00018357 | 0,00127126 | PI4KB |
| ENSGALG00000014226 | 4001,88511 | 0,460133597 | 0,13949225 | 3,29863207 | 0,00097157 | 0,00520949 | |
| ENSGALG00000022980 | 7902,73972 | 0,459059074 | 0,1123607 | 4,08558383 | 4,40E-05 | 0,00036679 | CDKN1B |
| ENSGALG00000014324 | 2616,41443 | 0,458208088 | 0,14844789 | 3,08665942 | 0,00202419 | 0,00953942 | |
| ENSGALG00000001881 | 2571,72261 | 0,456046301 | 0,14749617 | 3,09191952 | 0,00198867 | 0,00938825 | |
| ENSGALG00000000921 | 4858,40124 | 0,454646053 | 0,11600766 | 3,9191038 | 8,89E-05 | 0,00067906 | |
| ENSGALG00000004537 | 5387,82075 | 0,454616982 | 0,11547716 | 3,93685619 | 8,26E-05 | 0,00063539 | |
| ENSGALG00000016870 | 5615,99638 | 0,454001604 | 0,11901986 | 3,81450277 | 0,00013646 | 0,00098404 | TMTC4 |
| ENSGALG00000027963 | 10511,1952 | 0,451771641 | 0,1053151 | 4,28971367 | 1,79E-05 | 0,00016974 | COX7C |
| ENSGALG00000009054 | 1742,52432 | 0,451368563 | 0,14445222 | 3,12469106 | 0,00177992 | 0,00855102 | |
| ENSGALG00000017351 | 2392,614 | 0,45115395 | 0,12979024 | 3,47602367 | 0,00050891 | 0,0030416 | |
| ENSGALG00000007286 | 2211,13478 | 0,450759437 | 0,14338367 | 3,14372922 | 0,0016681 | 0,00811112 | SIRT7 |
| ENSGALG00000001376 | 9938,31883 | 0,45012042 | 0,10543845 | 4,26903467 | 1,96E-05 | 0,00018459 | ARID3B |
| ENSGALG00000017126 | 2105,45067 | 0,449836372 | 0,14456606 | 3,11163202 | 0,00186056 | 0,00886339 | |
| ENSGALG00000004353 | 9460,54723 | 0,448918917 | 0,11266612 | 3,98450692 | 6,76E-05 | 0,00053309 | TBCA |
| ENSGALG00000015780 | 22777,7944 | 0,447991765 | 0,11059391 | 4,05078157 | 5,10E-05 | 0,00041817 | PNRC1 |
| ENSGALG00000012218 | 3810,36969 | 0,44776649 | 0,1283039 | 3,48988984 | 0,00048322 | 0,00291753 | CBY1 |
| ENSGALG00000008274 | 1660,02758 | 0,446767639 | 0,14203363 | 3,14550613 | 0,001658 | 0,00807066 | CRBN |
| ENSGALG00000005608 | 3810,24681 | 0,44662731 | 0,13008707 | 3,43329509 | 0,00059629 | 0,00347391 | SLC9A3R2 |
| ENSGALG00000017329 | 3706,96729 | 0,44555284 | 0,13631333 | 3,26859342 | 0,00108083 | 0,00570371 | FCHSD2 |
| ENSGALG00000015987 | 4807,19259 | 0,445391941 | 0,12455693 | 3,57581017 | 0,00034914 | 0,00221571 | |
| ENSGALG00000000067 | 2534,63139 | 0,444870546 | 0,12860379 | 3,45923359 | 0,00054171 | 0,00320811 | SPR |
| ENSGALG00000011921 | 3797,77855 | 0,444716768 | 0,14148109 | 3,14329478 | 0,00167058 | 0,00812027 | DHRS7 |
| ENSGALG00000027337 | 6395,16083 | 0,441351335 | 0,11482343 | 3,84373949 | 0,00012117 | 0,00088176 | |
| ENSGALG00000004907 | 2377,26928 | 0,441232207 | 0,13371969 | 3,29968027 | 0,00096795 | 0,00519663 | KCTD15 |
| ENSGALG00000017096 | 2040,45459 | 0,440278375 | 0,13379312 | 3,29073997 | 0,00099924 | 0,00532681 | |
| ENSGALG00000012043 | 13548,0358 | 0,439717314 | 0,10426627 | 4,21725362 | 2,47E-05 | 0,00022551 | |
| ENSGALG00000012265 | 3656,88094 | 0,439419657 | 0,1289098 | 3,40873745 | 0,00065264 | 0,00373978 | MICALL1 |
| ENSGALG00000002986 | 2686,67203 | 0,439006087 | 0,12512502 | 3,50853962 | 0,00045057 | 0,00274477 | |
| ENSGALG00000005820 | 8527,7428 | 0,438136271 | 0,11284846 | 3,88251872 | 0,00010338 | 0,00077336 | RNF130 |
| ENSGALG00000026116 | 1987,2825 | 0,437155594 | 0,13561904 | 3,22340863 | 0,00126675 | 0,00649596 | |
| ENSGALG00000013091 | 2461,62995 | 0,436802578 | 0,1334788 | 3,27244902 | 0,0010662 | 0,00564179 | |
| ENSGALG00000008444 | 16562,7434 | 0,436671101 | 0,11266409 | 3,87586767 | 0,00010625 | 0,00079262 | TUBAL3 |
| ENSGALG00000012821 | 60743,3322 | 0,436526282 | 0,11022595 | 3,96028605 | 7,49E-05 | 0,00058241 | |
| ENSGALG00000006399 | 2888,16792 | 0,435019992 | 0,12865618 | 3,38125997 | 0,00072154 | 0,00408283 | MESDC1 |
| ENSGALG00000010134 | 2529,19577 | 0,434710794 | 0,12612196 | 3,44674955 | 0,00056737 | 0,00333397 | ZFAND3 |
| ENSGALG00000016059 | 2071,29674 | 0,434358409 | 0,13768426 | 3,15474273 | 0,0016064 | 0,0078758 | ETS2 |
| ENSGALG00000021365 | 7390,77997 | 0,432965219 | 0,11720328 | 3,69413922 | 0,00022063 | 0,00148635 | DCTN3 |

| | | | | | | | |
|---|---|---|---|---|---|---|---|
| ENSGALG00000003447 | 2587,36522 | 0,432921053 | 0,13816024 | 3,1334707 | 0,00172752 | 0,00835531 | |
| ENSGALG00000004725 | 7054,54942 | 0,431307897 | 0,12989304 | 3,32048515 | 0,00089861 | 0,00489384 | |
| ENSGALG00000000551 | 4770,60214 | 0,428841133 | 0,11757382 | 3,64742036 | 0,00026489 | 0,00174139 | C6orf89 |
| ENSGALG00000023612 | 4284,86865 | 0,427551312 | 0,12223705 | 3,4977227 | 0,00046925 | 0,00284327 | LZTS2 |
| ENSGALG00000010838 | 5580,71709 | 0,427079258 | 0,13104043 | 3,25914125 | 0,0011175 | 0,00585859 | |
| ENSGALG00000015304 | 5360,75819 | 0,425237788 | 0,11685875 | 3,63890423 | 0,0002738 | 0,00179049 | SNX3 |
| ENSGALG00000020528 | 5739,52001 | 0,424819785 | 0,11091716 | 3,83006366 | 0,00012811 | 0,00092926 | INIP |
| ENSGALG00000015603 | 3809,74973 | 0,424497436 | 0,13385318 | 3,171366 | 0,00151724 | 0,00750352 | ZCCHC9 |
| ENSGALG00000026692 | 7703,25537 | 0,424103985 | 0,13618907 | 3,11408229 | 0,00184518 | 0,00880243 | NRAS |
| ENSGALG00000012009 | 3507,12317 | 0,422977808 | 0,12358938 | 3,42244456 | 0,00062061 | 0,0035925 | JKAMP |
| ENSGALG00000027710 | 12308,995 | 0,421872733 | 0,10758636 | 3,92124726 | 8,81E-05 | 0,00067418 | |
| ENSGALG00000000436 | 3616,92165 | 0,421563067 | 0,1280766 | 3,29149164 | 0,00099658 | 0,0053161 | RABIF |
| ENSGALG00000008613 | 2544,37784 | 0,420884082 | 0,1331062 | 3,16201722 | 0,0015668 | 0,00772899 | NDUFA1 |
| ENSGALG00000007054 | 5550,73084 | 0,419262331 | 0,12272606 | 3,41624533 | 0,00063491 | 0,00365664 | TBC1D16 |
| ENSGALG00000017675 | 6606,96745 | 0,417757976 | 0,11206995 | 3,72765371 | 0,00019327 | 0,00132563 | |
| ENSGALG00000008818 | 5178,18834 | 0,417517144 | 0,12931442 | 3,22869741 | 0,00124355 | 0,00640113 | IDH1 |
| ENSGALG00000026394 | 6922,09548 | 0,417110916 | 0,11027253 | 3,78254597 | 0,00015523 | 0,00109906 | |
| ENSGALG00000007220 | 48744,1762 | 0,415111704 | 0,10489641 | 3,95734916 | 7,58E-05 | 0,00058794 | FTH1 |
| ENSGALG00000001816 | 2247,2353 | 0,414935997 | 0,13272784 | 3,12621686 | 0,00177071 | 0,00851282 | MAGI3 |
| ENSGALG00000026802 | 39202,809 | 0,413746284 | 0,11461572 | 3,60985629 | 0,00030637 | 0,00197874 | |
| ENSGALG00000015915 | 2924,59886 | 0,413737551 | 0,13087631 | 3,16128683 | 0,00157074 | 0,00773719 | |
| ENSGALG00000012426 | 3567,37244 | 0,412512543 | 0,12505498 | 3,29864939 | 0,00097151 | 0,00520949 | DDHD1 |
| ENSGALG00000001201 | 7498,13604 | 0,411468635 | 0,10775001 | 3,81873406 | 0,00013414 | 0,00096819 | GFOD2 |
| ENSGALG00000024081 | 3055,10774 | 0,410810718 | 0,12976304 | 3,16585318 | 0,00154629 | 0,0076361 | TGIF2 |
| ENSGALG00000003750 | 7887,37064 | 0,409049332 | 0,1094937 | 3,73582537 | 0,0001871 | 0,00129184 | PLCG1 |
| ENSGALG00000008597 | 7184,86692 | 0,404910294 | 0,11988781 | 3,37740998 | 0,00073172 | 0,0041252 | MRPL2 |
| ENSGALG00000006505 | 3147,8767 | 0,40439474 | 0,12215456 | 3,31051698 | 0,00093124 | 0,00504931 | ZFAND6 |
| ENSGALG00000008143 | 14658,924 | 0,402789538 | 0,10639446 | 3,78581321 | 0,00015321 | 0,00108585 | |
| ENSGALG00000003924 | 3485,26038 | 0,402077112 | 0,12622532 | 3,18539184 | 0,00144558 | 0,00723588 | ACBD6 |
| ENSGALG00000008508 | 3096,67282 | 0,401158456 | 0,12032341 | 3,33400181 | 0,00085606 | 0,00469403 | PYGB |
| ENSGALG00000010704 | 15057,8876 | 0,401049242 | 0,10647647 | 3,76655286 | 0,00016552 | 0,00116401 | |
| ENSGALG00000003371 | 6971,35561 | 0,400787786 | 0,11743488 | 3,4128513 | 0,00064287 | 0,00369623 | |
| ENSGALG00000001096 | 5395,30498 | 0,399476875 | 0,12102428 | 3,30079943 | 0,0009641 | 0,00518412 | |
| ENSGALG00000003412 | 4305,48949 | 0,398169298 | 0,11885899 | 3,34992998 | 0,00080832 | 0,00447915 | |
| ENSGALG00000014970 | 13935,5595 | 0,397421157 | 0,10749714 | 3,69703948 | 0,00021813 | 0,00147386 | FSTL1 |
| ENSGALG00000003422 | 9411,16928 | 0,397187449 | 0,11024485 | 3,60277551 | 0,00031484 | 0,00202385 | |
| ENSGALG00000001572 | 4170,73825 | 0,394480266 | 0,11780929 | 3,34846484 | 0,00081261 | 0,00449376 | COMMD4 |
| ENSGALG00000028470 | 5093,19173 | 0,39443368 | 0,12418131 | 3,17627247 | 0,00149181 | 0,00740195 | |
| ENSGALG00000021047 | 9190,88912 | 0,393972396 | 0,12289699 | 3,20571241 | 0,00134729 | 0,00683431 | KIFAP3 |
| ENSGALG00000010490 | 10248,5665 | 0,389494511 | 0,11178754 | 3,4842389 | 0,00049354 | 0,00296799 | DPYSL4 |
| ENSGALG00000013000 | 7034,84966 | 0,388948819 | 0,1248778 | 3,11463547 | 0,00184172 | 0,00878902 | |
| ENSGALG00000000955 | 5256,56158 | 0,388613492 | 0,11456174 | 3,39217531 | 0,0006934 | 0,00395006 | TMEM222 |
| ENSGALG00000015311 | 2435,46157 | 0,387705603 | 0,12617408 | 3,07278335 | 0,00212072 | 0,009926 | RNF38 |
| ENSGALG00000004011 | 4239,17688 | 0,385892717 | 0,12240581 | 3,15256855 | 0,00161841 | 0,00792043 | |
| ENSGALG00000005337 | 6236,10371 | 0,385644449 | 0,10969262 | 3,51568283 | 0,00043862 | 0,00268278 | SNX2 |
| ENSGALG00000004086 | 6417,78519 | 0,385423377 | 0,12458188 | 3,09373553 | 0,00197654 | 0,00935043 | |
| ENSGALG00000026102 | 19802,423 | 0,385336727 | 0,12172597 | 3,1656082 | 0,00154759 | 0,00763976 | |
| ENSGALG00000016639 | 3515,28691 | 0,384196385 | 0,12279677 | 3,1287174 | 0,00175571 | 0,00846159 | |
| ENSGALG00000026188 | 5641,89638 | 0,384179427 | 0,11525662 | 3,33325247 | 0,00085837 | 0,00470479 | |
| ENSGALG00000008736 | 6339,84871 | 0,384117634 | 0,1174006 | 3,27185401 | 0,00106845 | 0,00564929 | PNPLA7 |
| ENSGALG00000008297 | 4909,6132 | 0,379216929 | 0,11488546 | 3,30082615 | 0,00096401 | 0,00518412 | SEMA4B |
| ENSGALG00000007205 | 14498,1893 | 0,37857906 | 0,10495892 | 3,60692592 | 0,00030985 | 0,00199614 | |
| ENSGALG00000028128 | 3864,66289 | 0,378077792 | 0,11727019 | 3,22398887 | 0,00126418 | 0,00648525 | |
| ENSGALG00000015436 | 20607,2425 | 0,377556464 | 0,11268107 | 3,35066444 | 0,00080618 | 0,00447275 | SDCBP |
| ENSGALG00000002053 | 9601,26321 | 0,376194926 | 0,11159082 | 3,37119972 | 0,00074842 | 0,00420366 | |
| ENSGALG00000004191 | 5196,69877 | 0,374652449 | 0,11676506 | 3,20860067 | 0,00133383 | 0,0067774 | RIC8A |
| ENSGALG00000003717 | 5316,70071 | 0,373543889 | 0,11309691 | 3,30286557 | 0,00095702 | 0,00515652 | MYO9B |
| ENSGALG00000026291 | 5117,15968 | 0,37266418 | 0,11704722 | 3,18387886 | 0,00145316 | 0,00725779 | |
| ENSGALG00000026421 | 20179,8154 | 0,372144283 | 0,11694539 | 3,18220567 | 0,00146158 | 0,00728649 | PSMB2 |
| ENSGALG00000006857 | 3516,26088 | 0,36847671 | 0,11911862 | 3,09335936 | 0,00197904 | 0,00935579 | SPATA20 |
| ENSGALG00000028249 | 8201,39743 | 0,366515679 | 0,10750136 | 3,40940496 | 0,00065105 | 0,00373536 | |
| ENSGALG00000021692 | 6544,92461 | 0,361385734 | 0,11652843 | 3,10126669 | 0,00192695 | 0,00914762 | |
| ENSGALG00000007589 | 8938,41629 | 0,360072922 | 0,10858113 | 3,31616491 | 0,00091262 | 0,00496041 | |
| ENSGALG00000011435 | 8069,4783 | 0,35897775 | 0,11230455 | 3,19646662 | 0,00139122 | 0,0070167 | |

| | | | | | | | |
|---|---|---|---|---|---|---|---|
| ENSGALG00000001330 | 11236,9836 | 0,358718136 | 0,11464643 | 3,12890804 | 0,00175457 | 0,00846159 | ATP5G1 |
| ENSGALG00000026517 | 6681,39607 | 0,355711572 | 0,1112537 | 3,1973011 | 0,0013872 | 0,00700287 | RFC5 |
| ENSGALG00000023073 | 19350,4834 | 0,353608475 | 0,10573559 | 3,34427114 | 0,00082499 | 0,00454598 | |
| ENSGALG00000026565 | 5799,2794 | 0,352044819 | 0,11138453 | 3,16062577 | 0,00157431 | 0,00775196 | |
| ENSGALG00000012229 | 22179,4003 | 0,350504617 | 0,10217867 | 3,43031096 | 0,00060289 | 0,00350934 | RPS29 |
| ENSGALG00000010347 | 11028,0806 | 0,346318448 | 0,1080262 | 3,20587467 | 0,00134653 | 0,0068343 | CCNG2 |
| ENSGALG00000026300 | 11402,689 | 0,345400821 | 0,1081778 | 3,19289927 | 0,00140852 | 0,00708947 | |
| ENSGALG00000023447 | 10483,1653 | 0,344836567 | 0,10464307 | 3,29535984 | 0,00098296 | 0,00525853 | |
| ENSGALG00000013784 | 13631,601 | 0,339967639 | 0,10798288 | 3,14834763 | 0,00164196 | 0,00802129 | CDT1 |
| ENSGALG00000015998 | 9692,12229 | 0,339150714 | 0,10588553 | 3,20299389 | 0,00136007 | 0,00689145 | |
| ENSGALG00000024261 | 6604,44476 | 0,336614329 | 0,10865736 | 3,09794308 | 0,00194869 | 0,00924117 | |
| ENSGALG00000000905 | 28204,5829 | 0,330518761 | 0,10726487 | 3,0813327 | 0,00206076 | 0,00967826 | PSMB4 |
| ENSGALG00000007272 | 8940,93056 | 0,327852784 | 0,10564224 | 3,10342513 | 0,00191295 | 0,00909384 | RNF166 |
| ENSGALG00000008443 | 36760,133 | 0,321038212 | 0,10104157 | 3,17728858 | 0,00148659 | 0,00737876 | ARHGDIA |
| ENSGALG00000015664 | 24898,2726 | 0,319058165 | 0,10032221 | 3,18033435 | 0,00147105 | 0,00732031 | ADD1 |
| ENSGALG00000012546 | 35449,6881 | 0,318249627 | 0,09997702 | 3,18322776 | 0,00145643 | 0,0072688 | MCM5 |
| ENSGALG00000011238 | 46142,4631 | 0,316616268 | 0,10217962 | 3,09862447 | 0,00194421 | 0,00922316 | WLS |
| ENSGALG00000020078 | 34717,1709 | 0,313921783 | 0,09887982 | 3,17478105 | 0,0014995 | 0,00743198 | |
| ENSGALG00000028600 | 142500,088 | 0,313870865 | 0,09855851 | 3,18461456 | 0,00144947 | 0,00725001 | |
| ENSGALG00000010961 | 89733,2273 | -0,310634298 | 0,10095703 | -3,076896 | 0,00209168 | 0,00980319 | IGF2BP3 |
| ENSGALG00000003815 | 34063,9206 | -0,31144656 | 0,10097526 | -3,0843849 | 0,00203974 | 0,00959938 | RPN2 |
| ENSGALG00000004375 | 15903,9039 | -0,317187722 | 0,10291656 | -3,0819891 | 0,00205622 | 0,00966361 | VPS35 |
| ENSGALG00000010748 | 12682,3509 | -0,319116199 | 0,1031995 | -3,0922261 | 0,00198661 | 0,00938507 | EXO1 |
| ENSGALG00000005668 | 17427,2491 | -0,321274618 | 0,10214198 | -3,145373 | 0,00165875 | 0,00807145 | WSB1 |
| ENSGALG00000001817 | 20809,4267 | -0,321828859 | 0,10400477 | -3,0943663 | 0,00197234 | 0,00933381 | CTCF |
| ENSGALG00000000235 | 25074,3258 | -0,321994098 | 0,10263119 | -3,1373902 | 0,00170459 | 0,00826202 | PPP2R2A |
| ENSGALG00000010178 | 15181,2565 | -0,322485727 | 0,10328298 | -3,1223512 | 0,00179413 | 0,00861017 | PUM3 |
| ENSGALG00000005758 | 19288,8311 | -0,323694502 | 0,10277128 | -3,149659 | 0,00163461 | 0,00799398 | |
| ENSGALG00000006477 | 10679,2726 | -0,324083678 | 0,10402567 | -3,1154203 | 0,00183683 | 0,00877181 | |
| ENSGALG00000002222 | 13238,2952 | -0,324594377 | 0,10274095 | -3,1593475 | 0,00158123 | 0,00777748 | NOC2L |
| ENSGALG00000000814 | 43324,2546 | -0,325022823 | 0,09923062 | -3,2754289 | 0,00105502 | 0,00558913 | HNRNPR |
| ENSGALG00000014575 | 10529,4114 | -0,325103377 | 0,10553605 | -3,0804959 | 0,00206656 | 0,00969881 | SEPHS1 |
| ENSGALG00000004780 | 17002,1156 | -0,326184617 | 0,10404378 | -3,1350709 | 0,00171813 | 0,0083217 | |
| ENSGALG00000011091 | 12877,9103 | -0,326829517 | 0,1036675 | -3,152671 | 0,00161784 | 0,00792043 | NUP133 |
| ENSGALG00000001262 | 15946,3219 | -0,32743844 | 0,10659706 | -3,0717398 | 0,00212815 | 0,00994712 | CLIC4 |
| ENSGALG00000004161 | 24484,0776 | -0,327725215 | 0,10319068 | -3,1759188 | 0,00149363 | 0,00740828 | CCNB2 |
| ENSGALG00000014286 | 15937,5586 | -0,329230926 | 0,10635463 | -3,0955955 | 0,00196418 | 0,00930168 | |
| ENSGALG00000002062 | 16195,7787 | -0,329944458 | 0,10510199 | -3,139279 | 0,00169364 | 0,00821945 | SRP68 |
| ENSGALG00000001962 | 54240,9734 | -0,330389801 | 0,10379904 | -3,1829756 | 0,0014577 | 0,00727247 | PTBP1 |
| ENSGALG00000010576 | 13964,8988 | -0,330632554 | 0,10260922 | -3,2222498 | 0,00127188 | 0,00651983 | GTF2A1 |
| ENSGALG00000010481 | 12612,5746 | -0,330642085 | 0,10591126 | -3,1218786 | 0,00179701 | 0,00862096 | CMPK1 |
| ENSGALG00000010341 | 71633,3369 | -0,33081711 | 0,10337066 | -3,2002997 | 0,00137285 | 0,00694328 | XPO5 |
| ENSGALG00000007387 | 14223,8054 | -0,330955786 | 0,10651466 | -3,1071383 | 0,00188908 | 0,00898666 | GTF3C1 |
| ENSGALG00000003547 | 13605,0703 | -0,331189406 | 0,10621784 | -3,1180207 | 0,0018207 | 0,00871312 | SKP2 |
| ENSGALG00000014445 | 19258,1997 | -0,331954776 | 0,10745587 | -3,0892197 | 0,00200683 | 0,00947071 | |
| ENSGALG00000007590 | 10666,5636 | -0,332146464 | 0,10813379 | -3,0716251 | 0,00212897 | 0,00994712 | MTMR8 |
| ENSGALG00000004108 | 32339,0897 | -0,332634184 | 0,10036764 | -3,3141577 | 0,0009192 | 0,00499197 | CCAR1 |
| ENSGALG00000007528 | 28367,4799 | -0,33311917 | 0,10609529 | -3,1398111 | 0,00169057 | 0,00820866 | TCERG1 |
| ENSGALG00000005177 | 21314,8021 | -0,333333735 | 0,1014573 | -3,2854583 | 0,00101817 | 0,00541709 | |
| ENSGALG00000001976 | 9937,86887 | -0,333600989 | 0,10865793 | -3,0701945 | 0,00213919 | 0,00998662 | TRIM33 |
| ENSGALG00000009332 | 24425,5213 | -0,334253024 | 0,10229188 | -3,26764 | 0,00108448 | 0,0057141 | RBM25 |
| ENSGALG00000014298 | 14009,4127 | -0,335955725 | 0,10618376 | -3,1639087 | 0,00155666 | 0,00768172 | RFC1 |
| ENSGALG00000003231 | 28849,9366 | -0,336908133 | 0,1031629 | -3,2657877 | 0,0010916 | 0,00574493 | |
| ENSGALG00000014327 | 7067,9269 | -0,338948318 | 0,10818125 | -3,1331523 | 0,0017294 | 0,00836141 | |
| ENSGALG00000002302 | 11281,3267 | -0,339317581 | 0,10876186 | -3,1198214 | 0,00180961 | 0,00866613 | |
| ENSGALG00000005966 | 13999,7426 | -0,340533213 | 0,11012393 | -3,0922727 | 0,0019863 | 0,00938507 | SEC61A1 |
| ENSGALG00000012202 | 8130,08702 | -0,341191282 | 0,10806076 | -3,1574023 | 0,00159182 | 0,00781275 | |
| ENSGALG00000007989 | 28672,5886 | -0,341827011 | 0,10925868 | -3,1286027 | 0,0017564 | 0,0084619 | ADNP |
| ENSGALG00000005237 | 11277,3975 | -0,342909039 | 0,10563626 | -3,24613 | 0,00116985 | 0,00608154 | FGD3 |
| ENSGALG00000016314 | 22102,7105 | -0,343364339 | 0,10699123 | -3,2092755 | 0,0013307 | 0,00677035 | ELOVL5 |
| ENSGALG00000012239 | 9046,42319 | -0,343596083 | 0,10589827 | -3,2445865 | 0,00117621 | 0,0061076 | METAP1 |
| ENSGALG00000001236 | 5792,05952 | -0,345759524 | 0,11038031 | -3,1324386 | 0,00173361 | 0,00837771 | |
| ENSGALG00000012837 | 7840,94956 | -0,345878883 | 0,10966121 | -3,1540676 | 0,00161012 | 0,00788836 | EXOC2 |
| ENSGALG00000006475 | 6864,19423 | -0,347016477 | 0,10794054 | -3,2148856 | 0,00130497 | 0,00666433 | |

| | | | | | | | |
|---|---|---|---|---|---|---|---|
| ENSGALG00000015553 | 10445,9765 | -0,347410904 | 0,11270003 | -3,082616 | 0,0020519 | 0,00964994 | |
| ENSGALG00000011831 | 9083,64488 | -0,347478626 | 0,11085227 | -3,1346101 | 0,00172083 | 0,00832589 | ANKRD50 |
| ENSGALG00000004269 | 64393,6529 | -0,347526191 | 0,10378498 | -3,348521 | 0,00081244 | 0,00449376 | |
| ENSGALG00000004200 | 11378,582 | -0,347970886 | 0,10751163 | -3,2365883 | 0,00120968 | 0,00626227 | DPP9 |
| ENSGALG00000016887 | 64639,1062 | -0,348222743 | 0,09787488 | -3,5578355 | 0,00037392 | 0,00234499 | |
| ENSGALG00000013649 | 18631,1873 | -0,348311839 | 0,10201524 | -3,4143117 | 0,00063943 | 0,00367803 | CNOT7 |
| ENSGALG00000017139 | 10651,9101 | -0,349554954 | 0,10704704 | -3,265433 | 0,00109297 | 0,00574991 | |
| ENSGALG00000002719 | 17659,4459 | -0,349571816 | 0,10180837 | -3,4336254 | 0,00059557 | 0,00347116 | CSNK1D |
| ENSGALG00000008454 | 36878,682 | -0,351028215 | 0,10381886 | -3,3811602 | 0,0007218 | 0,00408283 | NOP58 |
| ENSGALG00000017026 | 9366,38767 | -0,352330056 | 0,10866839 | -3,2422496 | 0,0011859 | 0,00615322 | VPS36 |
| ENSGALG00000007861 | 11084,8862 | -0,352469383 | 0,10824618 | -3,2561832 | 0,00112921 | 0,0059177 | MAGT1 |
| ENSGALG00000005569 | 11340,4378 | -0,352585966 | 0,10935005 | -3,2243787 | 0,00126246 | 0,00648 | USP47 |
| ENSGALG00000009913 | 62516,8642 | -0,353547892 | 0,09880506 | -3,5782367 | 0,00034592 | 0,00219771 | SMARCA5 |
| ENSGALG00000011620 | 9603,38362 | -0,353615473 | 0,10554783 | -3,3502865 | 0,00080728 | 0,00447521 | AKT1 |
| ENSGALG00000010177 | 12046,3189 | -0,353917428 | 0,10807412 | -3,2747656 | 0,0010575 | 0,00560008 | ZFC3H1 |
| ENSGALG00000015043 | 11145,3654 | -0,35411601 | 0,10996307 | -3,2203176 | 0,00128049 | 0,006559 | |
| ENSGALG00000008036 | 5782,62739 | -0,355406874 | 0,1116416 | -3,1834627 | 0,00145525 | 0,00726557 | TMEM209 |
| ENSGALG00000005125 | 7959,32707 | -0,356020767 | 0,10674059 | -3,3353833 | 0,00085182 | 0,00467642 | LSM14B |
| ENSGALG00000001811 | 14773,9745 | -0,357025162 | 0,11419157 | -3,1265456 | 0,00176873 | 0,00851143 | |
| ENSGALG00000008957 | 22456,5253 | -0,357191503 | 0,11566926 | -3,0880418 | 0,0020148 | 0,00949845 | MSH6 |
| ENSGALG00000009330 | 6639,49466 | -0,358006963 | 0,1142065 | -3,1347337 | 0,0017201 | 0,00832534 | DEGS1 |
| ENSGALG00000010763 | 6453,8983 | -0,358059956 | 0,11268149 | -3,1776289 | 0,00148485 | 0,00737279 | TRIP11 |
| ENSGALG00000008507 | 23518,146 | -0,358201653 | 0,10237686 | -3,4988536 | 0,00046726 | 0,00283376 | THOC2 |
| ENSGALG00000009839 | 30979,5758 | -0,358257804 | 0,10121654 | -3,5395182 | 0,00040086 | 0,00249096 | PPIG |
| ENSGALG00000010349 | 7048,15428 | -0,358393493 | 0,11290442 | -3,1743089 | 0,00150194 | 0,00744137 | POLR1C |
| ENSGALG00000007721 | 11622,16 | -0,358457512 | 0,10981787 | -3,2641091 | 0,00109809 | 0,00577239 | ARHGAP21 |
| ENSGALG00000005694 | 30867,6614 | -0,359162398 | 0,10218229 | -3,5149181 | 0,00043989 | 0,00268931 | GARS |
| ENSGALG00000008254 | 17757,2659 | -0,35964551 | 0,10795091 | -3,3315652 | 0,00086359 | 0,0047277 | |
| ENSGALG00000007683 | 31803,1359 | -0,359759566 | 0,1106419 | -3,251567 | 0,00114771 | 0,00599848 | |
| ENSGALG00000004670 | 7571,3318 | -0,360055666 | 0,10915595 | -3,2985436 | 0,00097188 | 0,00520949 | |
| ENSGALG00000024039 | 12181,8658 | -0,36099515 | 0,10495725 | -3,4394494 | 0,0005829 | 0,00340658 | |
| ENSGALG00000004379 | 20460,5219 | -0,361272684 | 0,10336925 | -3,4949725 | 0,00047411 | 0,00287042 | BCL7A |
| ENSGALG00000012763 | 12297,8834 | -0,362574608 | 0,10840365 | -3,3446715 | 0,0008238 | 0,00454275 | GNPTAB |
| ENSGALG00000001942 | 6053,36315 | -0,362623052 | 0,11586077 | -3,1298174 | 0,00174915 | 0,00844792 | SPDL1 |
| ENSGALG00000012916 | 8111,10575 | -0,362729901 | 0,10919421 | -3,3218786 | 0,00089414 | 0,00487922 | DROSHA |
| ENSGALG00000002139 | 98917,1679 | -0,362887161 | 0,10160459 | -3,5715625 | 0,00035486 | 0,00224229 | CSDE1 |
| ENSGALG00000009325 | 21860,6433 | -0,363294006 | 0,11100817 | -3,2726781 | 0,00106534 | 0,00563941 | |
| ENSGALG00000003826 | 8394,88602 | -0,363869123 | 0,10711719 | -3,3969255 | 0,00068148 | 0,00389027 | ANAPC7 |
| ENSGALG00000001790 | 14734,6451 | -0,363943316 | 0,1104661 | -3,2946155 | 0,00098556 | 0,00526834 | |
| ENSGALG00000004553 | 9745,08279 | -0,363967467 | 0,10494768 | -3,4680848 | 0,00052418 | 0,00312057 | SGPL1 |
| ENSGALG00000001663 | 10259,5734 | -0,364202029 | 0,10990467 | -3,3137995 | 0,00092037 | 0,00499637 | MAK16 |
| ENSGALG00000010900 | 12043,9228 | -0,364291872 | 0,11281721 | -3,2290452 | 0,00124204 | 0,00640061 | SPC25 |
| ENSGALG00000003563 | 13287,7659 | -0,364389392 | 0,11010466 | -3,3094822 | 0,00093469 | 0,00506145 | YARS |
| ENSGALG00000002137 | 15220,8927 | -0,364569255 | 0,10476883 | -3,4797493 | 0,00050188 | 0,00300887 | PGAM5 |
| ENSGALG00000001126 | 13435,9211 | -0,365650894 | 0,10394749 | -3,5176502 | 0,00043539 | 0,00266657 | CHAF1A |
| ENSGALG00000015874 | 13228,0992 | -0,365980992 | 0,10796169 | -3,3899153 | 0,00069914 | 0,00397612 | TTK |
| ENSGALG00000003339 | 10767,0615 | -0,366040447 | 0,11772436 | -3,1093007 | 0,00187531 | 0,00892426 | MTOR |
| ENSGALG00000010400 | 7431,8392 | -0,366106792 | 0,10709024 | -3,4186756 | 0,00062927 | 0,00363171 | |
| ENSGALG00000014867 | 5553,43093 | -0,366420731 | 0,11739665 | -3,1212197 | 0,00180104 | 0,00863589 | PAIP1 |
| ENSGALG00000006193 | 20925,4656 | -0,366551091 | 0,11528348 | -3,179563 | 0,00147497 | 0,0073322 | |
| ENSGALG00000005353 | 4596,56596 | -0,367159893 | 0,11954477 | -3,0713171 | 0,00213117 | 0,00995398 | FAR1 |
| ENSGALG00000015683 | 11433,1929 | -0,367498221 | 0,1101651 | -3,335886 | 0,00085028 | 0,00466985 | GNG10 |
| ENSGALG00000003171 | 20304,0548 | -0,367614474 | 0,10192128 | -3,6068472 | 0,00030994 | 0,00199614 | |
| ENSGALG00000028828 | 22676,6926 | -0,367651199 | 0,10280751 | -3,5761124 | 0,00034874 | 0,00221461 | |
| ENSGALG00000024159 | 24561,8039 | -0,368265263 | 0,10651918 | -3,4572673 | 0,00054568 | 0,0032246 | |
| ENSGALG00000003881 | 9972,69972 | -0,368271023 | 0,11805058 | -3,1196037 | 0,00181095 | 0,00866949 | ERCC6L |
| ENSGALG00000001619 | 9403,98709 | -0,368532524 | 0,1142023 | -3,2270149 | 0,00125089 | 0,00643402 | ATG16L1 |
| ENSGALG00000004115 | 26356,0947 | -0,369026879 | 0,10774308 | -3,4250635 | 0,00061466 | 0,00356867 | DNAJA2 |
| ENSGALG00000012266 | 5845,05186 | -0,369135559 | 0,11234414 | -3,2857571 | 0,00101709 | 0,00541346 | |
| ENSGALG00000008526 | 11139,8409 | -0,369313202 | 0,10883113 | -3,3934519 | 0,00069018 | 0,00393499 | |
| ENSGALG00000001485 | 7897,80461 | -0,369653264 | 0,10794906 | -3,4243306 | 0,00061632 | 0,00357526 | |
| ENSGALG00000012263 | 9464,40856 | -0,36996383 | 0,10644059 | -3,4757776 | 0,00050937 | 0,00304173 | NEMF |
| ENSGALG00000007509 | 10510,1509 | -0,37005709 | 0,10740052 | -3,4455801 | 0,00056983 | 0,00334551 | |
| ENSGALG00000008807 | 7903,608 | -0,370129883 | 0,10704402 | -3,4577352 | 0,00054474 | 0,0032218 | DCUN1D1 |

| | | | | | | | |
|---|---|---|---|---|---|---|---|
| ENSGALG00000014582 | 21606,21 | -0,370326496 | 0,10926121 | -3,3893683 | 0,00070054 | 0,00398239 | CADM1 |
| ENSGALG00000010296 | 6483,94047 | -0,370634079 | 0,11535942 | -3,2128637 | 0,00131419 | 0,00670137 | MRPS18A |
| ENSGALG00000011215 | 4111,7689 | -0,370718656 | 0,11588474 | -3,1990291 | 0,00137891 | 0,00696619 | |
| ENSGALG00000004254 | 3591,44306 | -0,370755667 | 0,11847879 | -3,1292998 | 0,00175223 | 0,00845981 | |
| ENSGALG00000015649 | 13211,5423 | -0,370890117 | 0,10670353 | -3,4758935 | 0,00050915 | 0,00304173 | NOP14 |
| ENSGALG00000008557 | 15377,869 | -0,371010327 | 0,10757228 | -3,4489398 | 0,00056279 | 0,00331419 | ARL5B |
| ENSGALG00000005935 | 6194,93972 | -0,371684146 | 0,110711 | -3,3572469 | 0,00078723 | 0,00439447 | |
| ENSGALG00000004834 | 12710,539 | -0,372460145 | 0,10851246 | -3,4324184 | 0,00059822 | 0,00348367 | COPS3 |
| ENSGALG00000016578 | 7057,34653 | -0,372483757 | 0,11837004 | -3,1467739 | 0,00165083 | 0,00804438 | |
| ENSGALG00000003039 | 8776,91996 | -0,372512284 | 0,10936842 | -3,4060314 | 0,00065915 | 0,00377229 | SRM |
| ENSGALG00000015705 | 13014,2962 | -0,372615903 | 0,12101628 | -3,0790561 | 0,00207658 | 0,00974245 | WHSC1 |
| ENSGALG00000011713 | 7918,94572 | -0,372947939 | 0,11560014 | -3,2261893 | 0,0012545 | 0,00644774 | |
| ENSGALG00000000412 | 5015,41988 | -0,373361237 | 0,1191944 | -3,1323723 | 0,001734 | 0,00837771 | |
| ENSGALG00000003942 | 9873,26393 | -0,373719716 | 0,10787221 | -3,4644669 | 0,00053128 | 0,00315594 | LONP2 |
| ENSGALG00000016236 | 30922,9868 | -0,374432078 | 0,11350948 | -3,2986854 | 0,00097139 | 0,00520949 | |
| ENSGALG00000005618 | 12381,3407 | -0,374714579 | 0,10648923 | -3,5188027 | 0,0004335 | 0,002661 | CTR9 |
| ENSGALG00000010744 | 5078,21857 | -0,375014992 | 0,11759876 | -3,1889367 | 0,00142797 | 0,00716617 | GALNT7 |
| ENSGALG00000009626 | 5852,86848 | -0,375704266 | 0,12017862 | -3,1262155 | 0,00177072 | 0,00851282 | THBS1 |
| ENSGALG00000009677 | 24959,7858 | -0,376298569 | 0,10417999 | -3,6120043 | 0,00030384 | 0,00196335 | UBP |
| ENSGALG00000004867 | 10358,5874 | -0,376546151 | 0,10686348 | -3,5236186 | 0,0004257 | 0,00262613 | |
| ENSGALG00000000085 | 18552,0192 | -0,37666785 | 0,10734971 | -3,5087925 | 0,00045015 | 0,00274339 | |
| ENSGALG00000011281 | 6523,20431 | -0,37690737 | 0,11786547 | -3,1977759 | 0,00138492 | 0,00699394 | |
| ENSGALG00000008247 | 7600,6846 | -0,377692943 | 0,10953801 | -3,4480537 | 0,00056464 | 0,00332221 | ARFGAP2 |
| ENSGALG00000005006 | 11070,7539 | -0,377708377 | 0,10759882 | -3,5103394 | 0,00044754 | 0,00273115 | UBA2 |
| ENSGALG00000000978 | 4782,69426 | -0,377763607 | 0,11791314 | -3,2037448 | 0,00135653 | 0,00687606 | CEP104 |
| ENSGALG00000002985 | 14604,5417 | -0,378678888 | 0,10567575 | -3,5834037 | 0,00033915 | 0,00216174 | UTP18 |
| ENSGALG00000016851 | 8342,75174 | -0,378897954 | 0,12010288 | -3,1547783 | 0,0016062 | 0,0078758 | |
| ENSGALG00000008195 | 5727,11476 | -0,378910892 | 0,11315331 | -3,3486506 | 0,00081206 | 0,00449376 | NPHP1 |
| ENSGALG00000024047 | 6900,30697 | -0,380223912 | 0,11575902 | -3,2846159 | 0,00102121 | 0,00543119 | MYCL |
| ENSGALG00000007036 | 8649,57445 | -0,380541953 | 0,10600071 | -3,5899944 | 0,00033069 | 0,00211673 | PITRM1 |
| ENSGALG00000005573 | 47811,3213 | -0,380592325 | 0,10049182 | -3,7872967 | 0,0001523 | 0,00108107 | |
| ENSGALG00000008187 | 22960,7008 | -0,380821067 | 0,1023642 | -3,7202565 | 0,00019902 | 0,00135687 | EIF3J |
| ENSGALG00000000864 | 4682,06821 | -0,380939683 | 0,11598398 | -3,2844163 | 0,00102194 | 0,00543291 | FBXW2 |
| ENSGALG00000004696 | 6529,78459 | -0,381141883 | 0,10996498 | -3,4660297 | 0,0005282 | 0,0031404 | LEO1 |
| ENSGALG00000011330 | 86092,558 | -0,381207584 | 0,11735831 | -3,2482368 | 0,00116123 | 0,00605055 | DYNC1H1 |
| ENSGALG00000012953 | 7564,61929 | -0,381744469 | 0,11014956 | -3,4656922 | 0,00052887 | 0,00314297 | IPO8 |
| ENSGALG00000007559 | 4279,21636 | -0,381904484 | 0,1214524 | -3,1444788 | 0,00166383 | 0,00809326 | PDSS1 |
| ENSGALG00000008587 | 12026,0733 | -0,382953864 | 0,1040771 | -3,679521 | 0,00023367 | 0,0015626 | NDUFAF1 |
| ENSGALG00000001596 | 12885,5925 | -0,383722949 | 0,10902521 | -3,51958 | 0,00043223 | 0,00265657 | MLLT1 |
| ENSGALG00000004839 | 5677,61086 | -0,383869776 | 0,11257961 | -3,409763 | 0,00065019 | 0,00373203 | GPATCH1 |
| ENSGALG00000002346 | 7982,38435 | -0,383971209 | 0,11217493 | -3,422968 | 0,00061941 | 0,00358865 | CLSPN |
| ENSGALG00000016040 | 7227,33788 | -0,384424184 | 0,11393306 | -3,3741233 | 0,00074051 | 0,00416572 | |
| ENSGALG00000011881 | 17920,0092 | -0,384788496 | 0,10123697 | -3,8008694 | 0,00014419 | 0,0010289 | CCNA2 |
| ENSGALG00000017105 | 3524,63272 | -0,385190009 | 0,11906547 | -3,235111 | 0,00121595 | 0,0062852 | RNF6 |
| ENSGALG00000015010 | 9588,37587 | -0,38521292 | 0,11536985 | -3,3389394 | 0,00084099 | 0,00462442 | |
| ENSGALG00000015040 | 4593,81042 | -0,38608617 | 0,11721149 | -3,2939277 | 0,00098798 | 0,00527917 | SLC16A10 |
| ENSGALG00000013061 | 7957,2214 | -0,38617661 | 0,11279114 | -3,4238206 | 0,00061747 | 0,00358045 | PAPD7 |
| ENSGALG00000011357 | 6554,71543 | -0,387039211 | 0,1110437 | -3,4854674 | 0,00049128 | 0,0029557 | VEZT |
| ENSGALG00000006259 | 19676,996 | -0,387217866 | 0,1021071 | -3,7922717 | 0,00014928 | 0,00106185 | |
| ENSGALG00000027774 | 12094,1246 | -0,387231267 | 0,11361387 | -3,4083098 | 0,00065367 | 0,00374408 | |
| ENSGALG00000008280 | 7969,07752 | -0,387606949 | 0,11109303 | -3,4890303 | 0,00048478 | 0,00292304 | PMPCB |
| ENSGALG00000012724 | 10174,9584 | -0,387874274 | 0,1052864 | -3,6839923 | 0,00022961 | 0,00154105 | PAK1IP1 |
| ENSGALG00000015247 | 7003,54363 | -0,387982807 | 0,10807809 | -3,5898377 | 0,00033088 | 0,002117 | GPR161 |
| ENSGALG00000027262 | 21153,6502 | -0,388074983 | 0,10134615 | -3,8292029 | 0,00012856 | 0,00093152 | CAB39 |
| ENSGALG00000005575 | 18548,7794 | -0,388395592 | 0,10808819 | -3,5933214 | 0,00032649 | 0,00209381 | AKAP1 |
| ENSGALG00000010609 | 3709,28931 | -0,388677075 | 0,12431812 | -3,1264715 | 0,00176918 | 0,00851143 | NDUFAF7 |
| ENSGALG00000005012 | 6065,17104 | -0,388993034 | 0,12158189 | -3,1994324 | 0,00137698 | 0,00695904 | |
| ENSGALG00000005904 | 32126,5427 | -0,389017981 | 0,1104816 | -3,5211109 | 0,00042974 | 0,0026439 | TMED5 |
| ENSGALG00000011103 | 30281,6359 | -0,389199345 | 0,10207015 | -3,8130574 | 0,00013726 | 0,0009872 | |
| ENSGALG00000007788 | 4541,30869 | -0,389226557 | 0,11718497 | -3,3214717 | 0,00089544 | 0,00488438 | ABCB7 |
| ENSGALG00000003166 | 4550,92062 | -0,38962163 | 0,1147713 | -3,3947653 | 0,00068687 | 0,0039178 | |
| ENSGALG00000002129 | 6971,21336 | -0,390332447 | 0,11138482 | -3,5043595 | 0,00045771 | 0,00278449 | WDR47 |
| ENSGALG00000007485 | 13617,2364 | -0,391583137 | 0,1247391 | -3,1392172 | 0,001694 | 0,00821945 | TFRC |
| ENSGALG00000008983 | 6819,43432 | -0,391807233 | 0,11487703 | -3,4106665 | 0,00064804 | 0,00372126 | GNB4 |

| | | | | | | | |
|---|---|---|---|---|---|---|---|
| ENSGALG00000009364 | 15815,3872 | -0,391850394 | 0,10698859 | -3,6625439 | 0,00024972 | 0,00165578 | PRKCI |
| ENSGALG00000028035 | 5338,91852 | -0,391942986 | 0,12309173 | -3,1841536 | 0,00145178 | 0,00725779 | |
| ENSGALG00000017064 | 15165,1186 | -0,392266132 | 0,10750673 | -3,6487589 | 0,00026351 | 0,00173402 | RFC3 |
| ENSGALG00000013410 | 8555,69148 | -0,392331865 | 0,10734883 | -3,6547383 | 0,00025744 | 0,00169986 | UBA3 |
| ENSGALG00000003277 | 6528,68057 | -0,393093094 | 0,11637559 | -3,3777968 | 0,00073069 | 0,00412281 | SUZ12 |
| ENSGALG00000002682 | 5643,05491 | -0,393160876 | 0,12282679 | -3,2009375 | 0,00136981 | 0,00693308 | BRD3 |
| ENSGALG00000004481 | 5167,72741 | -0,39336646 | 0,12164503 | -3,2337242 | 0,00122187 | 0,00630861 | |
| ENSGALG00000015865 | 4560,35143 | -0,393610955 | 0,11603457 | -3,3921868 | 0,00069337 | 0,00395006 | |
| ENSGALG00000009569 | 11508,232 | -0,393918558 | 0,10692863 | -3,6839392 | 0,00022966 | 0,00154105 | TLK1 |
| ENSGALG00000009201 | 11168,2886 | -0,39393497 | 0,11185482 | -3,5218417 | 0,00042856 | 0,00263991 | |
| ENSGALG00000028133 | 7664,26535 | -0,394703045 | 0,11653188 | -3,387082 | 0,0007064 | 0,00400736 | UBE2V2 |
| ENSGALG00000028898 | 2507,09906 | -0,395098922 | 0,12861106 | -3,0720448 | 0,00212598 | 0,00994339 | CPTP |
| ENSGALG00000016467 | 8711,55629 | -0,395365049 | 0,11946994 | -3,3093265 | 0,00093521 | 0,00506145 | SMC6 |
| ENSGALG00000003043 | 3489,75229 | -0,395518744 | 0,12292663 | -3,2175188 | 0,00129305 | 0,00661835 | |
| ENSGALG00000002076 | 11878,1242 | -0,395918706 | 0,11403726 | -3,4718362 | 0,00051691 | 0,00308268 | |
| ENSGALG00000008521 | 16274,1776 | -0,395949661 | 0,12761742 | -3,1026303 | 0,00191809 | 0,00911512 | |
| ENSGALG00000014688 | 9710,78641 | -0,396059354 | 0,12498284 | -3,16891 | 0,00153012 | 0,00756173 | ZYX |
| ENSGALG00000004256 | 8236,22082 | -0,396589984 | 0,11400406 | -3,4787357 | 0,00050379 | 0,00301496 | |
| ENSGALG00000010643 | 4501,85662 | -0,396627508 | 0,11751597 | -3,3750945 | 0,0007379 | 0,00415399 | ZYG11B |
| ENSGALG00000012759 | 9679,72927 | -0,396656849 | 0,1130301 | -3,5093027 | 0,00044928 | 0,00274028 | NUP37 |
| ENSGALG00000004435 | 2804,79298 | -0,396797105 | 0,12438966 | -3,1899524 | 0,00142296 | 0,00714894 | RAB27A |
| ENSGALG00000009608 | 5107,67868 | -0,396825966 | 0,12732928 | -3,1165335 | 0,00182991 | 0,0087449 | |
| ENSGALG00000009838 | 16688,775 | -0,397053502 | 0,10163734 | -3,9065711 | 9,36E-05 | 0,00070928 | AQR |
| ENSGALG00000009533 | 7127,79412 | -0,397081701 | 0,12818092 | -3,0978221 | 0,00194948 | 0,00924172 | NMD3 |
| ENSGALG00000015435 | 3490,23228 | -0,397319834 | 0,12169857 | -3,2647865 | 0,00109547 | 0,00576082 | DCAF6 |
| ENSGALG00000015215 | 6704,29174 | -0,397390834 | 0,12048616 | -3,2982281 | 0,00097297 | 0,00521125 | MAPRE2 |
| ENSGALG00000013776 | 3256,83732 | -0,397455529 | 0,12774216 | -3,1113889 | 0,0018621 | 0,00886759 | CEP135 |
| ENSGALG00000002675 | 3557,32889 | -0,397531672 | 0,12525226 | -3,1738483 | 0,00150432 | 0,00744777 | SCUBE3 |
| ENSGALG00000009322 | 5962,2024 | -0,397973054 | 0,11357624 | -3,5040168 | 0,0004583 | 0,00278683 | NVL |
| ENSGALG00000001576 | 13166,5037 | -0,398019961 | 0,10682503 | -3,7259054 | 0,00019462 | 0,00133284 | |
| ENSGALG00000008889 | 7462,94615 | -0,398667809 | 0,10789505 | -3,6949593 | 0,00021992 | 0,00148303 | TTC14 |
| ENSGALG00000010674 | 9662,45958 | -0,399166032 | 0,11134741 | -3,5848703 | 0,00033725 | 0,00215164 | DCTD |
| ENSGALG00000006483 | 5623,94704 | -0,399267586 | 0,12106372 | -3,2979954 | 0,00097378 | 0,00521352 | |
| ENSGALG00000004368 | 3768,91401 | -0,39990946 | 0,1253746 | -3,1897167 | 0,00142412 | 0,00715213 | |
| ENSGALG00000001377 | 9703,5691 | -0,400093128 | 0,10731064 | -3,7283641 | 0,00019273 | 0,0013239 | POLRMT |
| ENSGALG00000020251 | 2658,23573 | -0,40015479 | 0,12471527 | -3,2085468 | 0,00133408 | 0,0067774 | FNIP2 |
| ENSGALG00000007663 | 14617,8334 | -0,400171791 | 0,11471241 | -3,4884785 | 0,00048578 | 0,00292778 | KDM3B |
| ENSGALG00000008289 | 17537,4534 | -0,401142997 | 0,11170507 | -3,5910904 | 0,0003293 | 0,00210884 | RBM19 |
| ENSGALG00000000332 | 20852,8348 | -0,401294712 | 0,10538837 | -3,8077702 | 0,00014023 | 0,00100589 | |
| ENSGALG00000016747 | 15836,0603 | -0,401466819 | 0,10405 | -3,8584028 | 0,00011413 | 0,00083858 | |
| ENSGALG00000001094 | 14665,6234 | -0,40156915 | 0,12409768 | -3,2359118 | 0,00121255 | 0,00627312 | ADGRD2 |
| ENSGALG00000011493 | 4936,29609 | -0,401631833 | 0,1189649 | -3,3760533 | 0,00073534 | 0,00414216 | TMEM169 |
| ENSGALG00000010304 | 4399,91638 | -0,401743756 | 0,11989741 | -3,3507292 | 0,00080599 | 0,00447275 | MLH3 |
| ENSGALG00000001422 | 12037,3785 | -0,402405365 | 0,1260368 | -3,1927608 | 0,0014092 | 0,00709025 | |
| ENSGALG00000005516 | 8167,22794 | -0,402406965 | 0,10796988 | -3,7270299 | 0,00019375 | 0,00132825 | HEATR6 |
| ENSGALG00000012138 | 29874,2309 | -0,402996238 | 0,10409408 | -3,8714616 | 0,00010818 | 0,00080225 | |
| ENSGALG00000011379 | 3502,29926 | -0,403264726 | 0,12185227 | -3,3094561 | 0,00093477 | 0,00506145 | SLC44A5 |
| ENSGALG00000004486 | 7663,27363 | -0,403537703 | 0,11899283 | -3,3912775 | 0,00069568 | 0,00396137 | ZCCHC8 |
| ENSGALG00000011505 | 19437,7359 | -0,403647787 | 0,1041549 | -3,8754564 | 0,00010643 | 0,00079353 | MARK3 |
| ENSGALG00000007610 | 8753,25449 | -0,403710794 | 0,10808926 | -3,734976 | 0,00018773 | 0,00129417 | CDC23 |
| ENSGALG00000003219 | 2817,88024 | -0,403748967 | 0,12502192 | -3,2294255 | 0,00124039 | 0,00639452 | |
| ENSGALG00000003033 | 4208,03288 | -0,4040823 | 0,12394506 | -3,2601726 | 0,00111344 | 0,00583958 | |
| ENSGALG00000009887 | 7499,04151 | -0,404219345 | 0,11161371 | -3,6215922 | 0,0002928 | 0,00190191 | DIEXF |
| ENSGALG00000016070 | 5235,92577 | -0,404284672 | 0,11391617 | -3,5489664 | 0,00038675 | 0,00240876 | WRB |
| ENSGALG00000005558 | 18603,5366 | -0,404537586 | 0,10210074 | -3,9621417 | 7,43E-05 | 0,0005792 | TBL3 |
| ENSGALG00000003358 | 9790,28902 | -0,404595333 | 0,1105122 | -3,6610919 | 0,00025114 | 0,0016639 | UBIAD1 |
| ENSGALG00000010428 | 7926,19447 | -0,404704607 | 0,10820488 | -3,7401697 | 0,0001839 | 0,00127223 | FAM98A |
| ENSGALG00000012509 | 6088,11777 | -0,40489721 | 0,11727746 | -3,4524724 | 0,00055547 | 0,00327393 | |
| ENSGALG00000016034 | 11941,1216 | -0,4051791 | 0,11639003 | -3,4812184 | 0,00049914 | 0,00299769 | ATRN |
| ENSGALG00000011405 | 5639,0186 | -0,405273633 | 0,11718169 | -3,4585065 | 0,00054318 | 0,00321538 | SLC4A7 |
| ENSGALG00000002042 | 9110,02017 | -0,405517273 | 0,11009443 | -3,6833588 | 0,00023018 | 0,00154304 | |
| ENSGALG00000011118 | 5098,98573 | -0,405615226 | 0,11354185 | -3,5723853 | 0,00035374 | 0,00224011 | MIER1 |
| ENSGALG00000004300 | 11426,3375 | -0,405827457 | 0,10825338 | -3,7488663 | 0,00017764 | 0,00123772 | PTDSS2 |
| ENSGALG00000002107 | 25522,2013 | -0,406162719 | 0,10189792 | -3,9859767 | 6,72E-05 | 0,00053042 | |

| | | | | | | | |
|---|---|---|---|---|---|---|---|
| ENSGALG00000011571 | 3230,47914 | -0,406488263 | 0,12109769 | -3,3566972 | 0,00078879 | 0,00440141 | AGPAT4 |
| ENSGALG00000014404 | 9644,71084 | -0,407214942 | 0,11242179 | -3,6222066 | 0,0002921 | 0,00189831 | |
| ENSGALG00000004001 | 29564,3469 | -0,407260268 | 0,12203749 | -3,3371735 | 0,00084635 | 0,00465015 | |
| ENSGALG00000000275 | 5245,75511 | -0,407767805 | 0,11961336 | -3,409049 | 0,0006519 | 0,00373771 | KCTD9 |
| ENSGALG00000013661 | 4102,81323 | -0,40778269 | 0,12930218 | -3,1537185 | 0,00161205 | 0,00789496 | MICU3 |
| ENSGALG00000009534 | 2852,25345 | -0,408065762 | 0,12826207 | -3,1814998 | 0,00146515 | 0,00729626 | |
| ENSGALG00000003143 | 15625,0529 | -0,408123735 | 0,10675451 | -3,8230117 | 0,00013183 | 0,0009527 | ZC3H7A |
| ENSGALG00000001616 | 13431,8281 | -0,408530697 | 0,10735755 | -3,8053281 | 0,00014162 | 0,00101479 | CDC45 |
| ENSGALG00000003995 | 3806,36291 | -0,408610387 | 0,12214717 | -3,3452299 | 0,00082214 | 0,00453545 | ERAL1 |
| ENSGALG00000011100 | 3743,98602 | -0,408960809 | 0,12437236 | -3,2881968 | 0,00100831 | 0,00536886 | |
| ENSGALG00000008523 | 12531,5109 | -0,409027447 | 0,10500568 | -3,8952888 | 9,81E-05 | 0,00073778 | XPNPEP1 |
| ENSGALG00000012607 | 5903,92354 | -0,409282973 | 0,12766873 | -3,20582 | 0,00134678 | 0,0068343 | ZCCHC6 |
| ENSGALG00000009483 | 9325,77474 | -0,409303762 | 0,10929848 | -3,7448259 | 0,00018052 | 0,0012537 | MARK1 |
| ENSGALG00000013858 | 2841,13225 | -0,410414198 | 0,12330011 | -3,3285793 | 0,0008729 | 0,00477675 | |
| ENSGALG00000002673 | 4173,32355 | -0,41052388 | 0,11457704 | -3,5829508 | 0,00033973 | 0,00216448 | GRPEL2 |
| ENSGALG00000016345 | 4179,81268 | -0,410542245 | 0,1255035 | -3,2711616 | 0,00107107 | 0,00566094 | KLHL15 |
| ENSGALG00000006404 | 23302,9151 | -0,411265326 | 0,10229476 | -4,020395 | 5,81E-05 | 0,00046863 | DDX46 |
| ENSGALG00000011662 | 25762,1153 | -0,412100057 | 0,1171723 | -3,5170434 | 0,00043638 | 0,00267027 | |
| ENSGALG00000006157 | 6346,9334 | -0,412112134 | 0,11052549 | -3,7286613 | 0,0001925 | 0,00132315 | |
| ENSGALG00000000780 | 20148,9101 | -0,413093901 | 0,1091845 | -3,7834483 | 0,00015467 | 0,00109565 | |
| ENSGALG00000010734 | 12330,7124 | -0,413140268 | 0,11515498 | -3,5876892 | 0,00033362 | 0,00213052 | PPP4R3A |
| ENSGALG00000027373 | 11590,4668 | -0,413207621 | 0,12758753 | -3,2386208 | 0,00120109 | 0,00622289 | |
| ENSGALG00000014811 | 9624,08099 | -0,413279775 | 0,10683153 | -3,8685189 | 0,0001095 | 0,00081033 | SMCHD1 |
| ENSGALG00000009300 | 6311,47255 | -0,413699825 | 0,11749092 | -3,5211215 | 0,00042973 | 0,0026439 | NUMB |
| ENSGALG00000014978 | 5258,7361 | -0,41400107 | 0,1181716 | -3,5033889 | 0,00045938 | 0,00279153 | |
| ENSGALG00000004947 | 6549,15547 | -0,414737624 | 0,12292028 | -3,3740375 | 0,00074074 | 0,00416572 | POLR3A |
| ENSGALG00000001613 | 28424,6252 | -0,414922322 | 0,10503517 | -3,9503179 | 7,80E-05 | 0,0006041 | GNL3 |
| ENSGALG00000013943 | 3049,80064 | -0,414996341 | 0,12653671 | -3,2796518 | 0,00103935 | 0,00551258 | |
| ENSGALG00000012241 | 2635,25809 | -0,415005332 | 0,12984246 | -3,1962221 | 0,0013924 | 0,0070187 | KLHDC1 |
| ENSGALG00000027593 | 7745,80903 | -0,415214085 | 0,12333575 | -3,3665347 | 0,00076119 | 0,00426837 | |
| ENSGALG00000000800 | 3898,75043 | -0,415275292 | 0,12002346 | -3,4599509 | 0,00054027 | 0,00320097 | |
| ENSGALG00000009634 | 10679,8347 | -0,415471196 | 0,1072605 | -3,8734781 | 0,00010729 | 0,00079659 | GFM1 |
| ENSGALG00000016972 | 5736,72836 | -0,415815188 | 0,11849243 | -3,5092131 | 0,00044943 | 0,00274028 | NUFIP1 |
| ENSGALG00000026470 | 13532,2225 | -0,41655555 | 0,11205019 | -3,7175802 | 0,00020114 | 0,00136995 | BOD1L1 |
| ENSGALG00000003737 | 3492,31762 | -0,417242803 | 0,13295765 | -3,1381631 | 0,0017001 | 0,00824613 | |
| ENSGALG00000014386 | 4214,12952 | -0,417751127 | 0,12151585 | -3,4378325 | 0,00058639 | 0,00342208 | SEPSECS |
| ENSGALG00000014013 | 11695,5327 | -0,418753457 | 0,10545846 | -3,9707907 | 7,16E-05 | 0,00056246 | |
| ENSGALG00000015132 | 49891,3971 | -0,418923213 | 0,10073836 | -4,1585273 | 3,20E-05 | 0,00028099 | CDH2 |
| ENSGALG00000004121 | 4190,99049 | -0,419252057 | 0,12228908 | -3,4283687 | 0,00060722 | 0,00353303 | LPAR4 |
| ENSGALG00000009961 | 27537,6751 | -0,419402034 | 0,1063815 | -3,9424339 | 8,07E-05 | 0,0006229 | RLI |
| ENSGALG00000010201 | 4830,06159 | -0,41947829 | 0,13602431 | -3,0838479 | 0,00204342 | 0,0096134 | ABHD18 |
| ENSGALG00000019228 | 2351,71501 | -0,419863222 | 0,13563777 | -3,0954741 | 0,00196499 | 0,00930225 | SFT2D2 |
| ENSGALG00000007227 | 17010,7893 | -0,420130207 | 0,12508196 | -3,3588394 | 0,00078271 | 0,0043764 | MSI1 |
| ENSGALG00000000975 | 4091,69565 | -0,420207875 | 0,11994528 | -3,5033297 | 0,00045948 | 0,00279153 | |
| ENSGALG00000003985 | 6692,40436 | -0,42022626 | 0,11526594 | -3,6457107 | 0,00026665 | 0,00175048 | TARSL2 |
| ENSGALG00000007736 | 8209,23878 | -0,420323444 | 0,11023063 | -3,8131273 | 0,00013722 | 0,0009872 | |
| ENSGALG00000008989 | 27837,9247 | -0,420708851 | 0,10920494 | -3,852471 | 0,00011693 | 0,00085547 | |
| ENSGALG00000017119 | 1864,30027 | -0,420708881 | 0,13656883 | -3,0805629 | 0,0020661 | 0,00969881 | TNFRSF19 |
| ENSGALG00000010630 | 14099,6 | -0,420914921 | 0,11268105 | -3,7354544 | 0,00018738 | 0,00129256 | AHCTF1 |
| ENSGALG00000004310 | 3386,86871 | -0,4211147 | 0,1211839 | -3,4750055 | 0,00051084 | 0,00304916 | TTYH3 |
| ENSGALG00000010671 | 83381,23 | -0,421121888 | 0,1031007 | -4,0845686 | 4,42E-05 | 0,00036794 | HNRNPU |
| ENSGALG00000008466 | 2872,73209 | -0,421406429 | 0,13329064 | -3,1615605 | 0,00156926 | 0,00773552 | |
| ENSGALG00000014937 | 3129,4872 | -0,421603891 | 0,12557883 | -3,3572847 | 0,00078712 | 0,00439447 | KPNA5 |
| ENSGALG00000018803 | 3405,63382 | -0,421885923 | 0,12639678 | -3,3377901 | 0,00084448 | 0,00464172 | |
| ENSGALG00000015692 | 1953,56891 | -0,422243248 | 0,13365312 | -3,1592472 | 0,00158177 | 0,00777748 | POLN |
| ENSGALG00000013607 | 7250,98124 | -0,422561681 | 0,11201001 | -3,7725349 | 0,0001616 | 0,0011382 | |
| ENSGALG00000006002 | 2601,77756 | -0,422978689 | 0,1302324 | -3,247876 | 0,0011627 | 0,00605359 | WHAMM |
| ENSGALG00000002763 | 8455,88563 | -0,423193935 | 0,11582995 | -3,6535794 | 0,00025861 | 0,00170672 | FRZB |
| ENSGALG00000000962 | 2645,42242 | -0,423212428 | 0,12644055 | -3,3471258 | 0,00081654 | 0,00451002 | PUS3 |
| ENSGALG00000008708 | 9031,55021 | -0,423296325 | 0,10630235 | -3,9820033 | 6,83E-05 | 0,00053843 | |
| ENSGALG00000014568 | 6675,25211 | -0,423460012 | 0,11373368 | -3,7232595 | 0,00019667 | 0,00134486 | |
| ENSGALG00000001850 | 22801,4935 | -0,423798382 | 0,1070003 | -3,9607213 | 7,47E-05 | 0,00058168 | RARS |
| ENSGALG00000016701 | 2038,86851 | -0,423807866 | 0,1319527 | -3,2118167 | 0,00131899 | 0,00671829 | |
| ENSGALG00000003288 | 22428,6978 | -0,423843105 | 0,12398262 | -3,4185687 | 0,00062951 | 0,00363171 | TARS |

| | | | | | | | |
|---|---|---|---|---|---|---|---|
| ENSGALG00000009177 | 7539,35878 | -0,423896895 | 0,11006703 | -3,8512612 | 0,00011751 | 0,00085924 | FMR1 |
| ENSGALG00000012852 | 33011,4945 | -0,42431457 | 0,11899849 | -3,5657139 | 0,00036287 | 0,00228618 | MKRN1 |
| ENSGALG00000002288 | 13379,2923 | -0,424413539 | 0,10575851 | -4,0130439 | 5,99E-05 | 0,0004809 | LIG3 |
| ENSGALG00000011177 | 6032,23061 | -0,424449663 | 0,11472261 | -3,699791 | 0,00021578 | 0,00146014 | CEP290 |
| ENSGALG00000005945 | 41976,0293 | -0,425016747 | 0,10489477 | -4,0518393 | 5,08E-05 | 0,00041729 | |
| ENSGALG00000002119 | 6930,58517 | -0,425364527 | 0,1209038 | -3,5182064 | 0,00043447 | 0,00266338 | C16orf62 |
| ENSGALG00000000033 | 3918,69493 | -0,425866952 | 0,12660413 | -3,3637683 | 0,00076886 | 0,00430606 | |
| ENSGALG00000004181 | 1761,48747 | -0,425938272 | 0,13751656 | -3,0973599 | 0,00195253 | 0,00925292 | |
| ENSGALG00000016911 | 5511,46419 | -0,426013919 | 0,11550644 | -3,6882267 | 0,00022582 | 0,00151831 | RNF219 |
| ENSGALG00000005205 | 5076,63735 | -0,426364463 | 0,13358149 | -3,191793 | 0,00141393 | 0,00710879 | RNFT1 |
| ENSGALG00000016955 | 4576,70807 | -0,426960619 | 0,13063542 | -3,2683373 | 0,00108181 | 0,00570666 | VWA8 |
| ENSGALG00000008616 | 22133,5216 | -0,427012968 | 0,11420797 | -3,738907 | 0,00018482 | 0,00127734 | |
| ENSGALG00000004104 | 16560,1522 | -0,427862285 | 0,12883118 | -3,3211081 | 0,00089661 | 0,00488488 | INTS1 |
| ENSGALG00000002262 | 28425,4355 | -0,428265663 | 0,10466808 | -4,0916548 | 4,28E-05 | 0,00035977 | ETF1 |
| ENSGALG00000004776 | 7278,29032 | -0,428490556 | 0,1147583 | -3,7338524 | 0,00018857 | 0,00129865 | |
| ENSGALG00000012227 | 10485,4738 | -0,428492425 | 0,10916783 | -3,9250796 | 8,67E-05 | 0,00066503 | TSPAN5 |
| ENSGALG00000017111 | 14958,6545 | -0,428517877 | 0,1112294 | -3,8525595 | 0,00011689 | 0,00085547 | NUP58 |
| ENSGALG00000008844 | 4547,72028 | -0,428880605 | 0,13511839 | -3,1741098 | 0,00150297 | 0,00744376 | SLC31A1 |
| ENSGALG00000023571 | 6639,21372 | -0,428973189 | 0,12324229 | -3,4807305 | 0,00050005 | 0,00300183 | TRIM59 |
| ENSGALG00000014953 | 10323,6261 | -0,429472364 | 0,13438161 | -3,1959161 | 0,00139388 | 0,00702134 | RWDD1 |
| ENSGALG00000016454 | 3850,19078 | -0,429504414 | 0,12203619 | -3,5194838 | 0,00043239 | 0,00265657 | E2F6 |
| ENSGALG00000000346 | 4943,52317 | -0,429865287 | 0,11696755 | -3,6750815 | 0,00023777 | 0,00158535 | |
| ENSGALG00000004133 | 28687,5841 | -0,430003398 | 0,1149324 | -3,7413591 | 0,00018303 | 0,00126815 | SRSF10 |
| ENSGALG00000006730 | 17262,8448 | -0,430049755 | 0,10295568 | -4,1770378 | 2,95E-05 | 0,00026144 | |
| ENSGALG00000027835 | 5051,56121 | -0,430117051 | 0,11513346 | -3,7358127 | 0,00018711 | 0,00129184 | PCBD1 |
| ENSGALG00000001501 | 5273,66434 | -0,430190725 | 0,11395127 | -3,7752164 | 0,00015987 | 0,0011272 | MAPK1 |
| ENSGALG00000006500 | 13286,4752 | -0,430344251 | 0,11868069 | -3,626068 | 0,00028777 | 0,00187553 | |
| ENSGALG00000004986 | 14224,0481 | -0,430376609 | 0,11894558 | -3,6182647 | 0,00029658 | 0,00192378 | CERCAM |
| ENSGALG00000004756 | 2781,91816 | -0,43117271 | 0,12469583 | -3,4577958 | 0,00054461 | 0,0032218 | CEP89 |
| ENSGALG00000015184 | 7792,59624 | -0,431383947 | 0,12288745 | -3,5103988 | 0,00044744 | 0,00273115 | |
| ENSGALG00000007705 | 251805,658 | -0,431935045 | 0,09889184 | -4,3677523 | 1,26E-05 | 0,00012567 | |
| ENSGALG00000011555 | 11516,3317 | -0,431962973 | 0,11422487 | -3,7816893 | 0,00015577 | 0,0011017 | QKI |
| ENSGALG00000014222 | 26624,5153 | -0,431998929 | 0,09984019 | -4,3269043 | 1,51E-05 | 0,00014748 | |
| ENSGALG00000011013 | 3992,5007 | -0,43257266 | 0,11842255 | -3,6527896 | 0,00025941 | 0,00171115 | |
| ENSGALG00000008413 | 12232,7391 | -0,432788919 | 0,11510536 | -3,7599371 | 0,00016996 | 0,00119276 | |
| ENSGALG00000007481 | 6576,27254 | -0,43296248 | 0,10915554 | -3,966473 | 7,29E-05 | 0,00057077 | HACD3 |
| ENSGALG00000006378 | 6236,19115 | -0,43312035 | 0,13117722 | -3,3017955 | 0,00096068 | 0,00517187 | |
| ENSGALG00000013920 | 7232,43287 | -0,433479079 | 0,1210408 | -3,5812642 | 0,00034194 | 0,00217443 | SUV39H2 |
| ENSGALG00000002877 | 3840,76126 | -0,433897954 | 0,13612607 | -3,1874713 | 0,00143523 | 0,00719463 | LRRC45 |
| ENSGALG00000011548 | 3329,63059 | -0,434202614 | 0,12811304 | -3,3892149 | 0,00070093 | 0,00398296 | PPP1R13B |
| ENSGALG00000010976 | 5423,33644 | -0,434636867 | 0,12461543 | -3,4878255 | 0,00048697 | 0,00293365 | CCDC126 |
| ENSGALG00000009930 | 17651,9175 | -0,434701609 | 0,10108014 | -4,300564 | 1,70E-05 | 0,000163 | |
| ENSGALG00000006221 | 21172,6562 | -0,434757063 | 0,10200869 | -4,2619612 | 2,03E-05 | 0,00018949 | KIF1A |
| ENSGALG00000028462 | 7479,58699 | -0,434862527 | 0,12695407 | -3,4253531 | 0,000614 | 0,00356791 | |
| ENSGALG00000008355 | 3052,82011 | -0,434887931 | 0,13359721 | -3,2552171 | 0,00113306 | 0,0059333 | |
| ENSGALG00000011543 | 9465,14039 | -0,435034048 | 0,12352777 | -3,521751 | 0,00042871 | 0,00263991 | SDAD1 |
| ENSGALG00000009519 | 11233,6738 | -0,43521067 | 0,10594254 | -4,1079878 | 3,99E-05 | 0,00034005 | ARG2 |
| ENSGALG00000009598 | 50606,5602 | -0,435652779 | 0,12077169 | -3,6072425 | 0,00030947 | 0,00199538 | EPRS |
| ENSGALG00000006409 | 11343,9424 | -0,436770571 | 0,12587936 | -3,4697554 | 0,00052093 | 0,00310258 | |
| ENSGALG00000001717 | 40955,9308 | -0,437024576 | 0,1153453 | -3,7888373 | 0,00015135 | 0,00107551 | SLC16A1 |
| ENSGALG00000012680 | 2735,45258 | -0,437301833 | 0,12310563 | -3,5522488 | 0,00038195 | 0,00238436 | E2F3 |
| ENSGALG00000011475 | 4517,36266 | -0,437819648 | 0,12412869 | -3,5271432 | 0,00042007 | 0,00259376 | |
| ENSGALG00000014375 | 10299,4805 | -0,437825336 | 0,10798758 | -4,0544047 | 5,03E-05 | 0,00041349 | |
| ENSGALG00000017038 | 6592,44922 | -0,437833021 | 0,13661218 | -3,204934 | 0,00135093 | 0,00685027 | PROSER1 |
| ENSGALG00000012642 | 14397,9617 | -0,437930988 | 0,11769371 | -3,720938 | 0,00019848 | 0,00135573 | |
| ENSGALG00000008796 | 7379,05533 | -0,438293488 | 0,11574409 | -3,7867461 | 0,00015263 | 0,00108291 | |
| ENSGALG00000007999 | 6686,69473 | -0,438325223 | 0,11125069 | -3,9399777 | 8,15E-05 | 0,00062871 | MRPS7 |
| ENSGALG00000009448 | 19270,6318 | -0,43880669 | 0,10703455 | -4,0996732 | 4,14E-05 | 0,00035053 | IFRD1 |
| ENSGALG00000008163 | 17633,4055 | -0,439350797 | 0,12629148 | -3,4788633 | 0,00050355 | 0,00301485 | PSME4 |
| ENSGALG00000004522 | 17438,7774 | -0,439397212 | 0,11296651 | -3,8896238 | 0,0001004 | 0,00075272 | |
| ENSGALG00000005854 | 8969,91159 | -0,439557852 | 0,11347852 | -3,8734895 | 0,00010729 | 0,00079659 | CLUH |
| ENSGALG00000014870 | 12682,5875 | -0,439767118 | 0,11699794 | -3,7587596 | 0,00017076 | 0,00119653 | |
| ENSGALG00000009731 | 11305,3134 | -0,439893222 | 0,10611405 | -4,1454759 | 3,39E-05 | 0,00029501 | |
| ENSGALG00000006427 | 15012,1637 | -0,439960428 | 0,10348 | -4,251647 | 2,12E-05 | 0,00019748 | CARS |

| | | | | | | | |
|---|---|---|---|---|---|---|---|
| ENSGALG00000012497 | 3402,65629 | -0,440007974 | 0,12129309 | -3,6276427 | 0,00028602 | 0,00186503 | |
| ENSGALG00000022802 | 18215,8301 | -0,44018485 | 0,113024 | -3,8946139 | 9,84E-05 | 0,00073869 | |
| ENSGALG00000000430 | 10996,8493 | -0,440297246 | 0,11045127 | -3,9863485 | 6,71E-05 | 0,00052989 | SPPL2B |
| ENSGALG00000007145 | 45254,0897 | -0,440432711 | 0,0991199 | -4,4434336 | 8,85E-06 | 9,27E-05 | ITGB1 |
| ENSGALG00000003071 | 6465,89293 | -0,441229907 | 0,11836541 | -3,7276929 | 0,00019324 | 0,00132563 | |
| ENSGALG00000015049 | 15802,9479 | -0,441268065 | 0,11099473 | -3,9755768 | 7,02E-05 | 0,00055286 | |
| ENSGALG00000008988 | 11231,5561 | -0,442984401 | 0,11261589 | -3,933587 | 8,37E-05 | 0,00064373 | ZZZ3 |
| ENSGALG00000006403 | 20700,7028 | -0,442987653 | 0,10779744 | -4,1094449 | 3,97E-05 | 0,00033876 | MESDC2 |
| ENSGALG00000010957 | 23588,3054 | -0,443064247 | 0,10206261 | -4,3411023 | 1,42E-05 | 0,00014017 | USP1 |
| ENSGALG00000020879 | 1971,7439 | -0,443502872 | 0,13705801 | -3,2358771 | 0,0012127 | 0,00627312 | SLC30A7 |
| ENSGALG00000012696 | 9654,978 | -0,443665071 | 0,1168806 | -3,7958828 | 0,00014712 | 0,00104761 | |
| ENSGALG00000012000 | 7157,91614 | -0,444031788 | 0,10961123 | -4,0509697 | 5,10E-05 | 0,00041817 | METTL14 |
| ENSGALG00000005206 | 7037,81003 | -0,444288079 | 0,11301214 | -3,9313307 | 8,45E-05 | 0,0006489 | ACSL3 |
| ENSGALG00000003861 | 8032,14284 | -0,444434747 | 0,10819238 | -4,1078193 | 3,99E-05 | 0,00034009 | HERC4 |
| ENSGALG00000005241 | 3696,95899 | -0,444802963 | 0,12608477 | -3,5278089 | 0,00041901 | 0,00258842 | |
| ENSGALG00000029011 | 22651,8846 | -0,444941401 | 0,10030346 | -4,4359527 | 9,17E-06 | 9,53E-05 | SLC35F1 |
| ENSGALG00000026971 | 4341,37635 | -0,445130127 | 0,11930931 | -3,7308917 | 0,0001908 | 0,00131268 | RRAGD |
| ENSGALG00000007704 | 19378,5947 | -0,445645488 | 0,1426064 | -3,1250034 | 0,00177803 | 0,00854496 | |
| ENSGALG00000003400 | 9822,18573 | -0,446090625 | 0,10733726 | -4,155972 | 3,24E-05 | 0,0002836 | AIMP2 |
| ENSGALG00000009020 | 9304,45489 | -0,446477137 | 0,11401531 | -3,9159402 | 9,01E-05 | 0,0006861 | JAG1 |
| ENSGALG00000013131 | 1864,58441 | -0,447051346 | 0,13334241 | -3,3526569 | 0,0008004 | 0,00444793 | C18orf21 |
| ENSGALG00000002536 | 10005,1999 | -0,447408092 | 0,11487944 | -3,894588 | 9,84E-05 | 0,00073869 | STX2 |
| ENSGALG00000002863 | 21477,0924 | -0,447764784 | 0,12646631 | -3,5405856 | 0,00039924 | 0,00248203 | ERGIC1 |
| ENSGALG00000007437 | 4980,32711 | -0,447866955 | 0,11297477 | -3,9643095 | 7,36E-05 | 0,00057531 | DPP8 |
| ENSGALG00000010612 | 2716,32593 | -0,44797699 | 0,14444805 | -3,1013018 | 0,00192672 | 0,00914762 | PRKD3 |
| ENSGALG00000016385 | 1906,4631 | -0,448510168 | 0,13792552 | -3,2518287 | 0,00114665 | 0,00599848 | TSSC1 |
| ENSGALG00000011642 | 18424,3446 | -0,448893503 | 0,1458916 | -3,0768974 | 0,00209167 | 0,00980319 | ANKRD17 |
| ENSGALG00000001845 | 10040,8917 | -0,449444654 | 0,10707839 | -4,1973421 | 2,70E-05 | 0,00024254 | PHTF1 |
| ENSGALG00000012568 | 2033,21553 | -0,449796718 | 0,13848137 | -3,2480667 | 0,00116192 | 0,00605185 | |
| ENSGALG00000006324 | 5540,59909 | -0,450353768 | 0,14477251 | -3,1107684 | 0,00186601 | 0,00888313 | |
| ENSGALG00000012598 | 6595,51322 | -0,450640697 | 0,12253012 | -3,6777953 | 0,00023526 | 0,00157089 | NAA35 |
| ENSGALG00000002519 | 3426,12944 | -0,450983038 | 0,12415844 | -3,6323189 | 0,00028089 | 0,00183506 | SLC25A33 |
| ENSGALG00000007652 | 11015,3194 | -0,45121126 | 0,11033188 | -4,0895819 | 4,32E-05 | 0,00036185 | SUCLG2 |
| ENSGALG00000005959 | 3317,97301 | -0,451515508 | 0,12097245 | -3,732383 | 0,00018968 | 0,00130559 | GLMN |
| ENSGALG00000016803 | 4775,60706 | -0,451555414 | 0,13299179 | -3,395363 | 0,00068538 | 0,0039109 | ST6GAL2 |
| ENSGALG00000002663 | 8380,31752 | -0,451591009 | 0,11688065 | -3,8636934 | 0,00011169 | 0,00082355 | ATR |
| ENSGALG00000007075 | 11514,1233 | -0,451627608 | 0,12690022 | -3,5589189 | 0,00037238 | 0,00233857 | ATP13A3 |
| ENSGALG00000004049 | 11640,7727 | -0,45169714 | 0,10780601 | -4,1899069 | 2,79E-05 | 0,00024948 | ITFG1 |
| ENSGALG00000013233 | 9834,12864 | -0,4517759 | 0,11414632 | -3,9578665 | 7,56E-05 | 0,00058733 | |
| ENSGALG00000006336 | 7051,08462 | -0,452052929 | 0,12470158 | -3,6250777 | 0,00028887 | 0,00188183 | |
| ENSGALG00000007659 | 35189,3993 | -0,45246564 | 0,10270533 | -4,4054738 | 1,06E-05 | 0,00010813 | HYOU1 |
| ENSGALG00000008088 | 3813,14193 | -0,452924609 | 0,11705672 | -3,8692748 | 0,00010916 | 0,00080882 | ACSL4 |
| ENSGALG00000008848 | 8858,46011 | -0,453382128 | 0,10839768 | -4,1825813 | 2,88E-05 | 0,00025631 | |
| ENSGALG00000009835 | 14808,4285 | -0,453500739 | 0,11172389 | -4,0591207 | 4,93E-05 | 0,00040645 | XPOT |
| ENSGALG00000015458 | 9174,0129 | -0,453606012 | 0,13072342 | -3,4699674 | 0,00052052 | 0,00310149 | ILDR2 |
| ENSGALG00000005840 | 6869,83939 | -0,454270613 | 0,11585326 | -3,9210861 | 8,82E-05 | 0,00067425 | DENND5A |
| ENSGALG00000006094 | 10694,3183 | -0,454736184 | 0,1116856 | -4,0715741 | 4,67E-05 | 0,00038696 | PHF6 |
| ENSGALG00000004713 | 12581,373 | -0,454819325 | 0,1044249 | -4,355468 | 1,33E-05 | 0,00013206 | |
| ENSGALG00000005160 | 9338,8368 | -0,454934871 | 0,12551054 | -3,6246746 | 0,00028933 | 0,00188297 | VMP1 |
| ENSGALG00000021883 | 1337,96899 | -0,455471836 | 0,14673539 | -3,1040354 | 0,001909 | 0,00907827 | |
| ENSGALG00000005489 | 5406,3342 | -0,455551792 | 0,12948295 | -3,5182378 | 0,00043442 | 0,00266338 | |
| ENSGALG00000016196 | 5946,1092 | -0,455745009 | 0,13219264 | -3,4475824 | 0,00056563 | 0,00332514 | |
| ENSGALG00000011191 | 11563,8832 | -0,455971006 | 0,10723132 | -4,2522187 | 2,12E-05 | 0,00019721 | |
| ENSGALG00000008839 | 9872,74847 | -0,456849267 | 0,12866693 | -3,5506348 | 0,0003843 | 0,00239683 | |
| ENSGALG00000008415 | 6540,80438 | -0,456904989 | 0,11868052 | -3,8498736 | 0,00011818 | 0,00086366 | ALS2 |
| ENSGALG00000000884 | 7194,47798 | -0,456928071 | 0,11254093 | -4,0601057 | 4,91E-05 | 0,00040499 | CXXC5 |
| ENSGALG00000002338 | 29369,7857 | -0,457290151 | 0,14291025 | -3,1998415 | 0,00137503 | 0,00695175 | |
| ENSGALG00000001179 | 4301,53132 | -0,45735712 | 0,11540169 | -3,9631753 | 7,40E-05 | 0,00057772 | |
| ENSGALG00000002954 | 6734,12402 | -0,457794297 | 0,12058072 | -3,7965796 | 0,00014671 | 0,00104576 | MBTD1 |
| ENSGALG00000008497 | 15581,4745 | -0,457962816 | 0,11566741 | -3,9593072 | 7,52E-05 | 0,00058414 | SLC38A2 |
| ENSGALG00000007777 | 3900,17866 | -0,45814234 | 0,12874776 | -3,5584491 | 0,00037305 | 0,0023406 | SLC39A10 |
| ENSGALG00000026322 | 5686,93416 | -0,458344915 | 0,11820551 | -3,8775258 | 0,00010552 | 0,00078767 | |
| ENSGALG00000013647 | 1849,99539 | -0,458404091 | 0,13582178 | -3,3750411 | 0,00073805 | 0,00415399 | TFB1M |
| ENSGALG00000026639 | 1529,36894 | -0,458562471 | 0,13972096 | -3,2819876 | 0,00103078 | 0,00547351 | SMCO4 |

| | | | | | | | |
|---|---|---|---|---|---|---|---|
| ENSGALG00000004531 | 7125,02349 | -0,458962037 | 0,11174171 | -4,1073476 | 4,00E-05 | 0,00034057 | PLOD1 |
| ENSGALG00000007842 | 5019,26998 | -0,459124954 | 0,11745791 | -3,9088467 | 9,27E-05 | 0,0007038 | |
| ENSGALG00000016930 | 13372,7628 | -0,459461549 | 0,10645494 | -4,3160189 | 1,59E-05 | 0,0001534 | DIS3 |
| ENSGALG00000008705 | 14944,0116 | -0,4597037 | 0,10963126 | -4,1931808 | 2,75E-05 | 0,00024622 | |
| ENSGALG00000015773 | 9016,27354 | -0,459782065 | 0,10646265 | -4,318717 | 1,57E-05 | 0,00015208 | |
| ENSGALG00000008482 | 22772,2201 | -0,459855147 | 0,11052584 | -4,1606122 | 3,17E-05 | 0,00027898 | STAG2 |
| ENSGALG00000010411 | 14781,2197 | -0,460452211 | 0,1181671 | -3,8966194 | 9,75E-05 | 0,00073496 | RNF13 |
| ENSGALG00000002501 | 3377,42917 | -0,46100497 | 0,12275816 | -3,7553917 | 0,00017307 | 0,00121087 | |
| ENSGALG00000006281 | 5170,2004 | -0,461106682 | 0,11489632 | -4,0132414 | 5,99E-05 | 0,00048078 | |
| ENSGALG00000006078 | 2849,4853 | -0,461126659 | 0,1296477 | -3,5567671 | 0,00037545 | 0,00235237 | PALB2 |
| ENSGALG00000013893 | 1508,86891 | -0,461829974 | 0,1439203 | -3,2089287 | 0,00133231 | 0,00677346 | MNS1 |
| ENSGALG00000015633 | 9547,40157 | -0,462141421 | 0,10577029 | -4,3692935 | 1,25E-05 | 0,00012525 | |
| ENSGALG00000016980 | 3122,81674 | -0,462357479 | 0,12582273 | -3,6746738 | 0,00023815 | 0,00158711 | COG3 |
| ENSGALG00000013136 | 2283,14809 | -0,462408191 | 0,13185399 | -3,5069714 | 0,00045324 | 0,00275976 | |
| ENSGALG00000000687 | 3686,83705 | -0,462411333 | 0,12645225 | -3,6568058 | 0,00025538 | 0,00168785 | |
| ENSGALG00000016445 | 7095,59053 | -0,463408352 | 0,11403536 | -4,0637251 | 4,83E-05 | 0,000399 | NOL10 |
| ENSGALG00000009899 | 35232,7937 | -0,46345514 | 0,10855368 | -4,2693637 | 1,96E-05 | 0,00018445 | |
| ENSGALG00000011794 | 10368,9097 | -0,463824908 | 0,11474911 | -4,0420785 | 5,30E-05 | 0,00043291 | DDX47 |
| ENSGALG00000007057 | 3861,26178 | -0,46473909 | 0,12088959 | -3,8443267 | 0,00012088 | 0,0008806 | LRRC28 |
| ENSGALG00000012655 | 11727,4649 | -0,4651119 | 0,10973943 | -4,2383297 | 2,25E-05 | 0,00020772 | POLR3B |
| ENSGALG00000015138 | 14137,444 | -0,465157078 | 0,12418565 | -3,7456587 | 0,00017992 | 0,00125045 | |
| ENSGALG00000002942 | 14672,6332 | -0,465828236 | 0,13435921 | -3,4670361 | 0,00052623 | 0,0031314 | JMJD1C |
| ENSGALG00000014217 | 2291,12491 | -0,46602463 | 0,13899423 | -3,3528343 | 0,00079989 | 0,0044469 | |
| ENSGALG00000010054 | 3266,33312 | -0,466429533 | 0,12708508 | -3,6702148 | 0,00024235 | 0,00161268 | |
| ENSGALG00000004158 | 2874,23978 | -0,467330867 | 0,12292142 | -3,8018669 | 0,00014361 | 0,00102584 | |
| ENSGALG00000028849 | 26828,63 | -0,467375912 | 0,11688987 | -3,9984296 | 6,38E-05 | 0,0005068 | SCAF11 |
| ENSGALG00000011309 | 5857,33759 | -0,467663025 | 0,12262631 | -3,8137249 | 0,00013689 | 0,00098656 | PLXNC1 |
| ENSGALG00000013728 | 12921,1551 | -0,467853342 | 0,10806143 | -4,3295129 | 1,49E-05 | 0,00014648 | PPAT |
| ENSGALG00000013765 | 8392,68097 | -0,468026469 | 0,11553655 | -4,0508954 | 5,10E-05 | 0,00041817 | |
| ENSGALG00000002282 | 2493,12249 | -0,468163907 | 0,13046007 | -3,5885608 | 0,00033251 | 0,0021254 | TMEM200A |
| ENSGALG00000000895 | 27062,2913 | -0,468166268 | 0,11013528 | -4,2508292 | 2,13E-05 | 0,00019793 | NUDC |
| ENSGALG00000002655 | 6693,97945 | -0,468798244 | 0,13255859 | -3,5365361 | 0,00040541 | 0,00251466 | ITGAV |
| ENSGALG00000006121 | 11011,1981 | -0,468808823 | 0,11657303 | -4,0215891 | 5,78E-05 | 0,00046653 | PIK3C2A |
| ENSGALG00000011232 | 2855,76217 | -0,469477808 | 0,12392263 | -3,7884753 | 0,00015157 | 0,00107652 | |
| ENSGALG00000006638 | 2375,89839 | -0,469621749 | 0,13927345 | -3,3719401 | 0,00074641 | 0,00419411 | GRIK4 |
| ENSGALG00000009954 | 1375,37198 | -0,469791805 | 0,14501118 | -3,2396936 | 0,00119658 | 0,00620391 | DYNC2LI1 |
| ENSGALG00000007058 | 9479,72971 | -0,469857249 | 0,10779741 | -4,3587065 | 1,31E-05 | 0,00013031 | LSG1 |
| ENSGALG00000011101 | 5741,10347 | -0,47005983 | 0,1114966 | -4,2159117 | 2,49E-05 | 0,00022671 | URB2 |
| ENSGALG00000000843 | 7211,68859 | -0,471124949 | 0,14308635 | -3,2925919 | 0,00099268 | 0,00530016 | DOT1L |
| ENSGALG00000011246 | 5026,82702 | -0,471179653 | 0,11657189 | -4,0419664 | 5,30E-05 | 0,00043291 | |
| ENSGALG00000007564 | 1255,86877 | -0,471313446 | 0,14906226 | -3,1618562 | 0,00156767 | 0,00773047 | |
| ENSGALG00000016836 | 6379,66167 | -0,471449962 | 0,11875041 | -3,9700911 | 7,18E-05 | 0,00056355 | TUBGCP3 |
| ENSGALG00000002439 | 1784,31098 | -0,471657703 | 0,15336793 | -3,0753347 | 0,00210266 | 0,00985127 | |
| ENSGALG00000000721 | 4145,80621 | -0,47173717 | 0,14279654 | -3,3035616 | 0,00095465 | 0,00514755 | SLC41A1 |
| ENSGALG00000008837 | 13730,8475 | -0,471745405 | 0,10804461 | -4,3662094 | 1,26E-05 | 0,00012637 | PELI1 |
| ENSGALG00000026412 | 3599,37504 | -0,471881516 | 0,13799409 | -3,4195778 | 0,00062718 | 0,00362134 | |
| ENSGALG00000016195 | 2664,10393 | -0,472549861 | 0,14052256 | -3,3628042 | 0,00077155 | 0,00431935 | |
| ENSGALG00000004228 | 2475,78192 | -0,472593441 | 0,12611515 | -3,747317 | 0,00017874 | 0,00124475 | USP40 |
| ENSGALG00000012647 | 9993,64528 | -0,472701385 | 0,12593301 | -3,7535939 | 0,00017432 | 0,00121772 | RFX4 |
| ENSGALG00000001627 | 4772,90031 | -0,472962819 | 0,1154681 | -4,0960473 | 4,20E-05 | 0,00035452 | |
| ENSGALG00000026192 | 2515,18021 | -0,473341595 | 0,1470736 | -3,2183994 | 0,00128908 | 0,00660055 | ST8SIA4 |
| ENSGALG00000028267 | 15487,1518 | -0,473386218 | 0,11459396 | -4,1309877 | 3,61E-05 | 0,00031165 | |
| ENSGALG00000006296 | 11149,2649 | -0,47366212 | 0,10934983 | -4,3316217 | 1,48E-05 | 0,0001453 | |
| ENSGALG00000004055 | 3994,96157 | -0,474043343 | 0,11949327 | -3,9671132 | 7,27E-05 | 0,00056956 | C7orf50 |
| ENSGALG00000027440 | 2934,02695 | -0,474361212 | 0,12732099 | -3,7257111 | 0,00019477 | 0,0013332 | |
| ENSGALG00000004889 | 8112,95247 | -0,474641509 | 0,1093142 | -4,3419931 | 1,41E-05 | 0,00013971 | SLC2A1 |
| ENSGALG00000005211 | 6813,79977 | -0,474686708 | 0,11553568 | -4,1085724 | 3,98E-05 | 0,00033962 | |
| ENSGALG00000010182 | 7219,05659 | -0,474781283 | 0,11239386 | -4,2242636 | 2,40E-05 | 0,00021949 | SLC29A1 |
| ENSGALG00000015925 | 3585,11623 | -0,475263897 | 0,12827839 | -3,7049413 | 0,00021144 | 0,00143436 | |
| ENSGALG00000026989 | 16347,2783 | -0,47543634 | 0,12391447 | -3,8368104 | 0,00012464 | 0,00090555 | HOXA7 |
| ENSGALG00000002783 | 1895,83684 | -0,475653855 | 0,15057219 | -3,1589755 | 0,00158325 | 0,00778022 | |
| ENSGALG00000016409 | 21993,092 | -0,475709899 | 0,1119047 | -4,2510269 | 2,13E-05 | 0,00019789 | KIDINS220 |
| ENSGALG00000008686 | 2315,39071 | -0,476496754 | 0,12925913 | -3,6863683 | 0,00022748 | 0,00152868 | TRDMT1 |
| ENSGALG00000006906 | 7509,2168 | -0,477103418 | 0,10793556 | -4,4202615 | 9,86E-06 | 0,0001016 | CHM |

| | | | | | | | |
|---|---|---|---|---|---|---|---|
| ENSGALG00000009841 | 1397,50609 | -0,477667905 | 0,15523553 | -3,0770527 | 0,00209058 | 0,00980319 | FASTKD1 |
| ENSGALG00000011342 | 3208,67383 | -0,478233256 | 0,14250555 | -3,3558921 | 0,00079109 | 0,00441063 | FGD6 |
| ENSGALG00000002045 | 30343,4023 | -0,478318973 | 0,09981877 | -4,7918739 | 1,65E-06 | 2,05E-05 | |
| ENSGALG00000002460 | 17009,9997 | -0,480154381 | 0,10514203 | -4,5667216 | 4,95E-06 | 5,50E-05 | |
| ENSGALG00000002862 | 5434,26011 | -0,480739106 | 0,11106412 | -4,3284827 | 1,50E-05 | 0,00014706 | |
| ENSGALG00000012484 | 32855,9599 | -0,4809575 | 0,12116285 | -3,969513 | 7,20E-05 | 0,00056451 | RIF1 |
| ENSGALG00000010884 | 11252,8159 | -0,481115067 | 0,11488944 | -4,1876351 | 2,82E-05 | 0,00025182 | DDX24 |
| ENSGALG00000015901 | 15729,8858 | -0,481643321 | 0,10542744 | -4,5684816 | 4,91E-06 | 5,46E-05 | SENP6 |
| ENSGALG00000023516 | 18924,6811 | -0,48179177 | 0,1052391 | -4,578068 | 4,69E-06 | 5,25E-05 | |
| ENSGALG00000002294 | 3404,11815 | -0,482168758 | 0,12824623 | -3,7597109 | 0,00017011 | 0,00119322 | SEMA6A |
| ENSGALG00000003078 | 20488,0185 | -0,483195985 | 0,10630098 | -4,5455457 | 5,48E-06 | 5,97E-05 | MMD |
| ENSGALG00000008765 | 5344,75739 | -0,483352119 | 0,12788071 | -3,779711 | 0,00015701 | 0,00110992 | C1D |
| ENSGALG00000005134 | 3683,92085 | -0,483437706 | 0,1457126 | -3,3177481 | 0,00090746 | 0,00493612 | ZSWIM8 |
| ENSGALG00000009967 | 51604,1361 | -0,483675694 | 0,10307537 | -4,6924466 | 2,70E-06 | 3,17E-05 | LRPPRC |
| ENSGALG00000012702 | 17854,6411 | -0,483830731 | 0,11942579 | -4,0513085 | 5,09E-05 | 0,00041799 | JARID2 |
| ENSGALG00000015830 | 24016,1217 | -0,484314141 | 0,10300764 | -4,7017304 | 2,58E-06 | 3,04E-05 | SYNCRIP |
| ENSGALG00000004887 | 17300,5109 | -0,484420703 | 0,10687429 | -4,5326215 | 5,83E-06 | 6,30E-05 | |
| ENSGALG00000002781 | 17608,1718 | -0,484707648 | 0,12726008 | -3,8087958 | 0,00013965 | 0,00100278 | |
| ENSGALG00000001950 | 14346,8445 | -0,484721287 | 0,11288002 | -4,2941281 | 1,75E-05 | 0,0001671 | PMPCA |
| ENSGALG00000001777 | 2307,31017 | -0,484938734 | 0,14695286 | -3,2999612 | 0,00096698 | 0,00519347 | MFSD11 |
| ENSGALG00000002198 | 1337,36066 | -0,485078535 | 0,14899049 | -3,2557685 | 0,00113086 | 0,00592407 | FHOD1 |
| ENSGALG00000004416 | 1683,22849 | -0,485858255 | 0,14199597 | -3,4216342 | 0,00062246 | 0,00359944 | TTC19 |
| ENSGALG00000012258 | 6299,78758 | -0,485912726 | 0,11533315 | -4,2131229 | 2,52E-05 | 0,00022876 | |
| ENSGALG00000013745 | 3956,92601 | -0,486562498 | 0,1295758 | -3,7550415 | 0,00017331 | 0,00121132 | RTTN |
| ENSGALG00000008862 | 12336,5298 | -0,486712741 | 0,10633346 | -4,5772303 | 4,71E-06 | 5,26E-05 | DNAJC10 |
| ENSGALG00000008670 | 2394,2538 | -0,486758564 | 0,13455087 | -3,6176545 | 0,00029728 | 0,00192648 | |
| ENSGALG00000009157 | 10495,9323 | -0,48696748 | 0,11167801 | -4,36046 | 1,30E-05 | 0,00012946 | |
| ENSGALG00000002906 | 23481,0393 | -0,487686151 | 0,1087191 | -4,4857451 | 7,27E-06 | 7,71E-05 | |
| ENSGALG00000005198 | 7882,46709 | -0,488262054 | 0,11634671 | -4,1966123 | 2,71E-05 | 0,00024316 | RPS6KB1 |
| ENSGALG00000004750 | 7577,76657 | -0,488295007 | 0,10928295 | -4,4681721 | 7,89E-06 | 8,32E-05 | NOL8 |
| ENSGALG00000010570 | 52092,719 | -0,488515596 | 0,10051531 | -4,8601111 | 1,17E-06 | 1,51E-05 | BTF3L4 |
| ENSGALG00000016909 | 4324,69907 | -0,488526858 | 0,11625721 | -4,2021211 | 2,64E-05 | 0,00023858 | NDFIP2 |
| ENSGALG00000004638 | 3370,76137 | -0,48870476 | 0,14148157 | -3,4541939 | 0,00055194 | 0,00325875 | |
| ENSGALG00000001036 | 9684,70052 | -0,488799588 | 0,1105545 | -4,4213448 | 9,81E-06 | 0,00010117 | ELAC2 |
| ENSGALG00000009981 | 1405,74497 | -0,49003231 | 0,14652981 | -3,3442499 | 0,00082505 | 0,00454598 | PREPL |
| ENSGALG00000010638 | 13156,3134 | -0,49033693 | 0,12087564 | -4,0565407 | 4,98E-05 | 0,00041072 | CASP3 |
| ENSGALG00000006252 | 14647,7489 | -0,49037706 | 0,13939868 | -3,5178027 | 0,00043514 | 0,00266623 | LMO4 |
| ENSGALG00000000272 | 5267,8402 | -0,490419343 | 0,13544359 | -3,6208384 | 0,00029365 | 0,00190591 | |
| ENSGALG00000008553 | 5356,62728 | -0,490452503 | 0,13463719 | -3,6427714 | 0,00026972 | 0,00176719 | INO80 |
| ENSGALG00000016404 | 5806,16288 | -0,492286998 | 0,11929847 | -4,1265157 | 3,68E-05 | 0,00031657 | |
| ENSGALG00000010335 | 1726,88898 | -0,492481433 | 0,14371477 | -3,4267976 | 0,00061074 | 0,0035505 | |
| ENSGALG00000002141 | 14007,0584 | -0,492600599 | 0,10621413 | -4,6378068 | 3,52E-06 | 4,03E-05 | UGGT1 |
| ENSGALG00000008871 | 16063,5897 | -0,492812674 | 0,13495076 | -3,6517962 | 0,00026041 | 0,00171613 | |
| ENSGALG00000015059 | 7575,55005 | -0,49315511 | 0,1131114 | -4,3599064 | 1,30E-05 | 0,00012969 | |
| ENSGALG00000005652 | 5954,57786 | -0,493244478 | 0,11326504 | -4,3547812 | 1,33E-05 | 0,0001323 | |
| ENSGALG00000023576 | 3299,29916 | -0,493564294 | 0,12078163 | -4,0864186 | 4,38E-05 | 0,0003657 | PIM3 |
| ENSGALG00000008798 | 4574,76494 | -0,493864419 | 0,11829778 | -4,1747565 | 2,98E-05 | 0,0002639 | ZMYND19 |
| ENSGALG00000020742 | 13114,1447 | -0,493871262 | 0,13939629 | -3,5429299 | 0,00039571 | 0,0024612 | PPRC1 |
| ENSGALG00000011962 | 7036,62107 | -0,494326456 | 0,13488463 | -3,6648093 | 0,00024752 | 0,00164472 | |
| ENSGALG00000015478 | 5554,56829 | -0,494614804 | 0,13028443 | -3,796423 | 0,0001468 | 0,00104588 | |
| ENSGALG00000007326 | 7076,35907 | -0,494698301 | 0,1094474 | -4,5199637 | 6,19E-06 | 6,66E-05 | PMM2 |
| ENSGALG00000005683 | 5447,0767 | -0,494902817 | 0,11790098 | -4,1976141 | 2,70E-05 | 0,00024241 | ARHGAP29 |
| ENSGALG00000000795 | 1504,93647 | -0,495778483 | 0,1425043 | -3,4790423 | 0,00050321 | 0,00301416 | MFSD12 |
| ENSGALG00000004273 | 11441,3705 | -0,495936282 | 0,1511882 | -3,2802578 | 0,00103712 | 0,00550504 | RALGPS2 |
| ENSGALG00000014998 | 1987,96101 | -0,495971981 | 0,15420457 | -3,2163248 | 0,00129844 | 0,00664345 | |
| ENSGALG00000015684 | 4408,68885 | -0,496145153 | 0,11398344 | -4,3527826 | 1,34E-05 | 0,00013339 | |
| ENSGALG00000004570 | 5026,79729 | -0,496270556 | 0,13511313 | -3,6730002 | 0,00023972 | 0,00159598 | NCOA3 |
| ENSGALG00000013257 | 123007,221 | -0,496798558 | 0,10262669 | -4,8408318 | 1,29E-06 | 1,64E-05 | |
| ENSGALG00000027635 | 9027,18129 | -0,49702612 | 0,11896336 | -4,1779765 | 2,94E-05 | 0,00026087 | NME3 |
| ENSGALG00000003596 | 25796,9116 | -0,497856616 | 0,162082 | -3,0716343 | 0,0021289 | 0,00994712 | TRRAP |
| ENSGALG00000010624 | 9648,37586 | -0,498764877 | 0,10861449 | -4,5920656 | 4,39E-06 | 4,91E-05 | MEMO1 |
| ENSGALG00000026865 | 1245,99167 | -0,49909443 | 0,16196027 | -3,0815855 | 0,00205901 | 0,00967338 | XRCC2 |
| ENSGALG00000007691 | 3206,91346 | -0,499115113 | 0,12945293 | -3,8555722 | 0,00011546 | 0,00084651 | ARMC9 |
| ENSGALG00000007155 | 2000,50129 | -0,499370494 | 0,14915389 | -3,3480218 | 0,00081391 | 0,00449729 | RMI2 |

| | | | | | | | |
|---|---|---|---|---|---|---|---|
| ENSGALG00000003874 | 8943,2552 | -0,499635075 | 0,10978836 | -4,5508931 | 5,34E-06 | 5,87E-05 | |
| ENSGALG00000005695 | 3266,30309 | -0,500082186 | 0,13148632 | -3,803302 | 0,00014278 | 0,00102045 | CHEK2 |
| ENSGALG00000025792 | 3188,10638 | -0,500292681 | 0,12184936 | -4,1058293 | 4,03E-05 | 0,0003426 | FRRS1L |
| ENSGALG00000011264 | 1925,3077 | -0,500518992 | 0,14795759 | -3,3828544 | 0,00071737 | 0,00406279 | |
| ENSGALG00000016758 | 1465,78964 | -0,5005603 | 0,14656111 | -3,4153691 | 0,00063696 | 0,00366533 | LIPT1 |
| ENSGALG00000010716 | 5501,72957 | -0,50085931 | 0,11885573 | -4,2140106 | 2,51E-05 | 0,00022817 | AGA |
| ENSGALG00000009269 | 3597,54557 | -0,501724842 | 0,13020726 | -3,8532784 | 0,00011655 | 0,00085403 | PTGR2 |
| ENSGALG00000011857 | 24467,2306 | -0,502240381 | 0,10517202 | -4,775418 | 1,79E-06 | 2,19E-05 | |
| ENSGALG00000003719 | 6864,04349 | -0,502933526 | 0,135236 | -3,7189322 | 0,00020007 | 0,00136332 | |
| ENSGALG00000006524 | 12378,2851 | -0,503792354 | 0,10770154 | -4,677671 | 2,90E-06 | 3,38E-05 | |
| ENSGALG00000016781 | 17546,5065 | -0,504088725 | 0,12484349 | -4,0377654 | 5,40E-05 | 0,00043994 | MAP4K4 |
| ENSGALG00000014922 | 15076,9971 | -0,504425665 | 0,11868898 | -4,2499788 | 2,14E-05 | 0,00019855 | ROCK1 |
| ENSGALG00000004690 | 52340,4796 | -0,504461763 | 0,10076346 | -5,006396 | 5,55E-07 | 7,63E-06 | |
| ENSGALG00000002728 | 6868,77576 | -0,504726866 | 0,12336417 | -4,0913571 | 4,29E-05 | 0,00035977 | SLC16A3 |
| ENSGALG00000011877 | 4219,79352 | -0,504776099 | 0,12198763 | -4,1379284 | 3,50E-05 | 0,00030411 | |
| ENSGALG00000005533 | 4875,77262 | -0,504976403 | 0,12223566 | -4,1311708 | 3,61E-05 | 0,0003116 | ZMYM3 |
| ENSGALG00000013209 | 5984,63909 | -0,505418476 | 0,11156134 | -4,5304087 | 5,89E-06 | 6,36E-05 | |
| ENSGALG00000015252 | 4947,94416 | -0,505578034 | 0,14235024 | -3,5516487 | 0,00038283 | 0,00238871 | ST3GAL6 |
| ENSGALG00000007234 | 4564,3675 | -0,505755971 | 0,11795303 | -4,2877742 | 1,80E-05 | 0,00017099 | CLCN5 |
| ENSGALG00000010034 | 2578,09523 | -0,50624459 | 0,13221838 | -3,8288518 | 0,00012874 | 0,00093236 | |
| ENSGALG00000003540 | 2816,51357 | -0,506324474 | 0,14263664 | -3,5497505 | 0,0003856 | 0,0024027 | CEP95 |
| ENSGALG00000005992 | 1391,45334 | -0,506625581 | 0,15329789 | -3,3048438 | 0,00095029 | 0,0051298 | PDE8A |
| ENSGALG00000006935 | 12264,1833 | -0,506724591 | 0,10942458 | -4,6308116 | 3,64E-06 | 4,15E-05 | |
| ENSGALG00000019867 | 6166,97553 | -0,506841982 | 0,11192678 | -4,5283354 | 5,95E-06 | 6,41E-05 | WDCP |
| ENSGALG00000008420 | 3322,24803 | -0,507225725 | 0,1221905 | -4,1511061 | 3,31E-05 | 0,00028877 | CAND2 |
| ENSGALG00000009782 | 45976,8535 | -0,507579083 | 0,10626981 | -4,7763242 | 1,79E-06 | 2,18E-05 | NAA15 |
| ENSGALG00000002424 | 2020,65689 | -0,508419582 | 0,16029303 | -3,1718134 | 0,0015149 | 0,0074947 | RECQL5 |
| ENSGALG00000002652 | 8163,31793 | -0,509052755 | 0,13652522 | -3,7286352 | 0,00019252 | 0,00132315 | |
| ENSGALG00000005509 | 6460,5427 | -0,509428544 | 0,14277078 | -3,5681568 | 0,0003595 | 0,00226707 | DPYD |
| ENSGALG00000004990 | 1688,4348 | -0,509590179 | 0,1471207 | -3,4637558 | 0,00053269 | 0,00316154 | |
| ENSGALG00000010735 | 16013,9368 | -0,510060882 | 0,1046006 | -4,8762709 | 1,08E-06 | 1,40E-05 | |
| ENSGALG00000003628 | 14552,598 | -0,510073233 | 0,14143166 | -3,6064997 | 0,00031036 | 0,00199787 | |
| ENSGALG00000015787 | 3935,81115 | -0,510253054 | 0,14104041 | -3,6177791 | 0,00029714 | 0,00192647 | RNGTT |
| ENSGALG00000008778 | 6210,22008 | -0,51039033 | 0,14919074 | -3,4210591 | 0,00062378 | 0,00360473 | SPRED2 |
| ENSGALG00000012135 | 48183,5814 | -0,511735314 | 0,10797482 | -4,7393951 | 2,14E-06 | 2,56E-05 | ATF4 |
| ENSGALG00000017085 | 4116,94944 | -0,511832688 | 0,12507401 | -4,0922387 | 4,27E-05 | 0,00035928 | SLC7A1 |
| ENSGALG00000002709 | 2938,4248 | -0,511853662 | 0,14024529 | -3,6497031 | 0,00026254 | 0,0017285 | MYO1C |
| ENSGALG00000004621 | 8629,88481 | -0,512109985 | 0,12302965 | -4,1624925 | 3,15E-05 | 0,00027687 | PREX1 |
| ENSGALG00000005278 | 9525,24365 | -0,512823817 | 0,15834712 | -3,2386054 | 0,00120116 | 0,00622289 | |
| ENSGALG00000014791 | 22071,9343 | -0,513191503 | 0,12375943 | -4,1466861 | 3,37E-05 | 0,00029365 | WDR3 |
| ENSGALG00000009825 | 16041,5695 | -0,51401966 | 0,11051134 | -4,6512843 | 3,30E-06 | 3,80E-05 | SRGAP1 |
| ENSGALG00000001441 | 10679,0004 | -0,514329303 | 0,10523513 | -4,8874299 | 1,02E-06 | 1,33E-05 | UBE2G1 |
| ENSGALG00000026468 | 7333,85705 | -0,514962132 | 0,10964748 | -4,6965248 | 2,65E-06 | 3,11E-05 | CHSY1 |
| ENSGALG00000002646 | 11173,6378 | -0,515104787 | 0,14774595 | -3,4864225 | 0,00048953 | 0,00294777 | FAM171B |
| ENSGALG00000005653 | 3018,77554 | -0,515161656 | 0,12312595 | -4,1840218 | 2,86E-05 | 0,00025519 | NFKB2 |
| ENSGALG00000011707 | 2975,95888 | -0,516018775 | 0,13531464 | -3,8134734 | 0,00013703 | 0,00098658 | |
| ENSGALG00000003566 | 2456,14829 | -0,516245691 | 0,13569542 | -3,8044444 | 0,00014212 | 0,00101682 | SMURF2 |
| ENSGALG00000003047 | 12574,6527 | -0,516252465 | 0,13206797 | -3,9089908 | 9,27E-05 | 0,0007038 | SUGP2 |
| ENSGALG00000007527 | 4564,36351 | -0,516973735 | 0,11834809 | -4,3682476 | 1,25E-05 | 0,00012548 | ADARB1 |
| ENSGALG00000012550 | 2418,32562 | -0,517745286 | 0,13904231 | -3,7236527 | 0,00019636 | 0,00134344 | HMOX1 |
| ENSGALG00000009207 | 1636,75887 | -0,518081559 | 0,15117707 | -3,426985 | 0,00061032 | 0,00354957 | TRIM2 |
| ENSGALG00000012207 | 4193,90666 | -0,518404651 | 0,11552573 | -4,4873522 | 7,21E-06 | 7,66E-05 | CCNT2 |
| ENSGALG00000009451 | 10542,6813 | -0,518444547 | 0,1054026 | -4,9187072 | 8,71E-07 | 1,14E-05 | PPID |
| ENSGALG00000010189 | 6061,82346 | -0,518843565 | 0,11081881 | -4,6819089 | 2,84E-06 | 3,32E-05 | |
| ENSGALG00000011596 | 8210,15954 | -0,519655791 | 0,1247256 | -4,1663925 | 3,09E-05 | 0,00027306 | |
| ENSGALG00000006622 | 2389,74327 | -0,520113735 | 0,13317469 | -3,9054999 | 9,40E-05 | 0,00071163 | FRA10AC1 |
| ENSGALG00000014784 | 4045,40608 | -0,520975246 | 0,14051541 | -3,7076021 | 0,00020923 | 0,00142008 | MAST4 |
| ENSGALG00000009992 | 11680,4728 | -0,521017168 | 0,11356065 | -4,588008 | 4,47E-06 | 5,01E-05 | ARHGAP5 |
| ENSGALG00000015889 | 13285,7099 | -0,52134607 | 0,14846895 | -3,5114822 | 0,00044562 | 0,00272187 | PHIP |
| ENSGALG00000007752 | 6790,46143 | -0,521363763 | 0,1083691 | -4,8110001 | 1,50E-06 | 1,88E-05 | COPS7B |
| ENSGALG00000005850 | 15672,1065 | -0,521743975 | 0,10948222 | -4,7655589 | 1,88E-06 | 2,29E-05 | FNBP1L |
| ENSGALG00000012354 | 2395,65855 | -0,521764337 | 0,15908224 | -3,2798402 | 0,00103866 | 0,00551105 | |
| ENSGALG00000015314 | 12934,4444 | -0,521997148 | 0,10929519 | -4,7760302 | 1,79E-06 | 2,19E-05 | BEND3 |
| ENSGALG00000004397 | 3452,74049 | -0,523035736 | 0,1275508 | -4,1006073 | 4,12E-05 | 0,00034977 | MLXIP |

| | | | | | | | |
|---|---|---|---|---|---|---|---|
| ENSGALG00000012592 | 3142,27379 | -0,523397807 | 0,14066624 | -3,7208488 | 0,00019855 | 0,00135573 | RMI1 |
| ENSGALG00000002487 | 10163,3626 | -0,523537895 | 0,1313011 | -3,9873079 | 6,68E-05 | 0,00052806 | SFSWAP |
| ENSGALG00000003260 | 9438,41253 | -0,523953501 | 0,11054819 | -4,7395938 | 2,14E-06 | 2,56E-05 | UTP6 |
| ENSGALG00000012720 | 24474,404 | -0,524679114 | 0,11154136 | -4,7038973 | 2,55E-06 | 3,01E-05 | NUP153 |
| ENSGALG00000005425 | 38874,994 | -0,524696803 | 0,11835069 | -4,4334072 | 9,28E-06 | 9,63E-05 | OGT |
| ENSGALG00000010861 | 2392,2986 | -0,524736464 | 0,13282156 | -3,9506874 | 7,79E-05 | 0,00060351 | OMA1 |
| ENSGALG00000004489 | 1483,19946 | -0,524803254 | 0,14557684 | -3,6049914 | 0,00031216 | 0,00200761 | |
| ENSGALG00000011770 | 14710,9194 | -0,525890593 | 0,11884576 | -4,4249841 | 9,64E-06 | 9,98E-05 | UMPS |
| ENSGALG00000002850 | 6570,40738 | -0,526275555 | 0,13474436 | -3,9057334 | 9,39E-05 | 0,00071134 | RASA2 |
| ENSGALG00000023562 | 1492,39213 | -0,526424205 | 0,1608921 | -3,2719084 | 0,00106824 | 0,00564929 | |
| ENSGALG00000015836 | 3297,85416 | -0,526441345 | 0,13246596 | -3,9741633 | 7,06E-05 | 0,00055575 | CEP162 |
| ENSGALG00000014426 | 14003,0497 | -0,526522361 | 0,12072956 | -4,361172 | 1,29E-05 | 0,00012913 | KPNA1 |
| ENSGALG00000011982 | 5512,8717 | -0,527666168 | 0,14035943 | -3,7593924 | 0,00017033 | 0,00119413 | PDE5 |
| ENSGALG00000011149 | 879,074202 | -0,527681437 | 0,16261977 | -3,2448788 | 0,00117501 | 0,00610367 | PLA2R1 |
| ENSGALG00000008096 | 6070,98948 | -0,528190931 | 0,1247684 | -4,2333711 | 2,30E-05 | 0,00021135 | SUFU |
| ENSGALG00000008847 | 3493,18836 | -0,528756478 | 0,12631689 | -4,1859522 | 2,84E-05 | 0,00025336 | VPS54 |
| ENSGALG00000013177 | 2545,01825 | -0,528877548 | 0,14123176 | -3,7447493 | 0,00018057 | 0,0012537 | BCAT1 |
| ENSGALG00000017392 | 1036,56961 | -0,528892969 | 0,15858794 | -3,3350138 | 0,00085295 | 0,00468075 | SPDYA |
| ENSGALG00000012979 | 10526,2614 | -0,528975634 | 0,14809894 | -3,5717718 | 0,00035457 | 0,00224223 | |
| ENSGALG00000000208 | 3421,17661 | -0,529383362 | 0,13520345 | -3,9154575 | 9,02E-05 | 0,00068709 | MCC |
| ENSGALG00000004474 | 1133,2959 | -0,529777367 | 0,15342984 | -3,4528967 | 0,0005546 | 0,0032709 | PLEKHF1 |
| ENSGALG00000013203 | 7478,18622 | -0,52983334 | 0,11632258 | -4,5548622 | 5,24E-06 | 5,78E-05 | |
| ENSGALG00000011419 | 3053,01947 | -0,529859697 | 0,12366202 | -4,2847409 | 1,83E-05 | 0,0001731 | DUSP4 |
| ENSGALG00000010159 | 4422,59589 | -0,529906457 | 0,12230696 | -4,3325945 | 1,47E-05 | 0,00014486 | |
| ENSGALG00000002974 | 1015,77625 | -0,529999669 | 0,16832703 | -3,1486308 | 0,00164037 | 0,0080164 | SPHKAP |
| ENSGALG00000002624 | 31172,8318 | -0,530323004 | 0,11239841 | -4,7182429 | 2,38E-06 | 2,82E-05 | BMS1 |
| ENSGALG00000006826 | 4092,14666 | -0,530661584 | 0,1454636 | -3,6480713 | 0,00026422 | 0,00173783 | KLHL25 |
| ENSGALG00000002253 | 1425,48713 | -0,530784268 | 0,14560305 | -3,6454199 | 0,00026696 | 0,00175077 | HEMK1 |
| ENSGALG00000011569 | 5815,29092 | -0,530885029 | 0,12042999 | -4,408246 | 1,04E-05 | 0,000107 | SCYL2 |
| ENSGALG00000003944 | 1246,48403 | -0,531232705 | 0,1600895 | -3,3183482 | 0,00090552 | 0,00492749 | FAM189A1 |
| ENSGALG00000004347 | 6495,16049 | -0,533321296 | 0,11036285 | -4,832435 | 1,35E-06 | 1,70E-05 | NEDD4 |
| ENSGALG00000000891 | 3716,06871 | -0,533422492 | 0,12306892 | -4,3343397 | 1,46E-05 | 0,00014396 | NSUN5 |
| ENSGALG00000015818 | 5198,53941 | -0,53353462 | 0,12660774 | -4,2140758 | 2,51E-05 | 0,00022817 | |
| ENSGALG00000015448 | 2486,28528 | -0,533672096 | 0,16395778 | -3,2549362 | 0,00113418 | 0,00593689 | |
| ENSGALG00000008312 | 1844,09907 | -0,534378374 | 0,13847665 | -3,8589783 | 0,00011386 | 0,00083706 | GSAP |
| ENSGALG00000004506 | 11897,6421 | -0,534405871 | 0,11012391 | -4,8527686 | 1,22E-06 | 1,55E-05 | URI1 |
| ENSGALG00000013085 | 2436,82685 | -0,535478159 | 0,13668214 | -3,9176895 | 8,94E-05 | 0,00068237 | PTPRO |
| ENSGALG00000008098 | 5171,85115 | -0,536763264 | 0,11526046 | -4,6569592 | 3,21E-06 | 3,70E-05 | NAMPT |
| ENSGALG00000006856 | 8733,4208 | -0,537151634 | 0,11626332 | -4,6201299 | 3,83E-06 | 4,35E-05 | CD151 |
| ENSGALG00000011024 | 13428,0492 | -0,537183599 | 0,10486491 | -5,122625 | 3,01E-07 | 4,37E-06 | CACHD1 |
| ENSGALG00000010564 | 3014,7883 | -0,537373895 | 0,12339974 | -4,354741 | 1,33E-05 | 0,0001323 | TBCK |
| ENSGALG00000015360 | 9022,70642 | -0,537653684 | 0,10556349 | -5,0931783 | 3,52E-07 | 5,05E-06 | KIAA1524 |
| ENSGALG00000026253 | 1652,9426 | -0,5379573 | 0,15060806 | -3,5719024 | 0,0003544 | 0,00224216 | TMEM144 |
| ENSGALG00000003705 | 3639,3367 | -0,538250704 | 0,12318514 | -4,3694452 | 1,25E-05 | 0,00012525 | VPS13C |
| ENSGALG00000012468 | 13779,9504 | -0,538390713 | 0,13666178 | -3,9395852 | 8,16E-05 | 0,00062927 | |
| ENSGALG00000016440 | 2000,28995 | -0,538612864 | 0,16061769 | -3,3533845 | 0,0007983 | 0,00444169 | KLF11 |
| ENSGALG00000016835 | 6008,91026 | -0,539714594 | 0,14683337 | -3,6756945 | 0,0002372 | 0,00158232 | |
| ENSGALG00000002873 | 15190,7659 | -0,539979366 | 0,10838996 | -4,9818208 | 6,30E-07 | 8,52E-06 | |
| ENSGALG00000015701 | 9337,57881 | -0,540112853 | 0,12997092 | -4,1556437 | 3,24E-05 | 0,00028383 | |
| ENSGALG00000015180 | 15097,1011 | -0,540189844 | 0,11292112 | -4,7837805 | 1,72E-06 | 2,11E-05 | PSAT1 |
| ENSGALG00000001389 | 3052,28108 | -0,540945617 | 0,12602912 | -4,2922273 | 1,77E-05 | 0,00016818 | TTLL11 |
| ENSGALG00000015491 | 4074,13394 | -0,541085652 | 0,12196746 | -4,4363116 | 9,15E-06 | 9,53E-05 | |
| ENSGALG00000012121 | 12053,9223 | -0,541445831 | 0,10454307 | -5,179165 | 2,23E-07 | 3,34E-06 | RCN1 |
| ENSGALG00000013438 | 21891,8795 | -0,541667727 | 0,11204746 | -4,8342706 | 1,34E-06 | 1,69E-05 | TEX10 |
| ENSGALG00000001957 | 8870,63197 | -0,541703645 | 0,13410352 | -4,0394438 | 5,36E-05 | 0,00043733 | |
| ENSGALG00000016460 | 6190,66257 | -0,541890883 | 0,14841996 | -3,6510648 | 0,00026116 | 0,00172019 | |
| ENSGALG00000014573 | 17671,9053 | -0,541989088 | 0,11034447 | -4,9117921 | 9,02E-07 | 1,18E-05 | NAT10 |
| ENSGALG00000025884 | 6669,8373 | -0,543082298 | 0,11342421 | -4,7880633 | 1,68E-06 | 2,07E-05 | NABP1 |
| ENSGALG00000007835 | 1738,05901 | -0,543517994 | 0,14833732 | -3,6640679 | 0,00024824 | 0,00164868 | PUS10 |
| ENSGALG00000006895 | 13372,5266 | -0,543915173 | 0,10588883 | -5,1366627 | 2,80E-07 | 4,09E-06 | CEP131 |
| ENSGALG00000004364 | 7914,97638 | -0,544307466 | 0,11563233 | -4,7072254 | 2,51E-06 | 2,97E-05 | ABR |
| ENSGALG00000026452 | 7763,72006 | -0,545277299 | 0,13192983 | -4,1330858 | 3,58E-05 | 0,0003098 | MURC |
| ENSGALG00000012223 | 1554,61989 | -0,545359079 | 0,13992319 | -3,8975604 | 9,72E-05 | 0,00073252 | ZRANB3 |
| ENSGALG00000003440 | 1101,0238 | -0,546469024 | 0,16030168 | -3,4090037 | 0,00065201 | 0,00373771 | |

| | | | | | | | |
|---|---|---|---|---|---|---|---|
| ENSGALG00000011351 | 157820,465 | -0,547374715 | 0,15800979 | -3,4641822 | 0,00053185 | 0,00315791 | |
| ENSGALG00000016768 | 7527,22871 | -0,54750013 | 0,11986999 | -4,5674497 | 4,94E-06 | 5,48E-05 | LONRF2 |
| ENSGALG00000009503 | 5606,80387 | -0,547842813 | 0,11865192 | -4,6172267 | 3,89E-06 | 4,41E-05 | RAB3GAP2 |
| ENSGALG00000015746 | 3303,51188 | -0,547863301 | 0,14036111 | -3,9032414 | 9,49E-05 | 0,00071632 | JAM2 |
| ENSGALG00000003937 | 13547,5021 | -0,548529686 | 0,13001502 | -4,2189717 | 2,45E-05 | 0,0002241 | GEMIN5 |
| ENSGALG00000005727 | 12274,5514 | -0,548590838 | 0,11540717 | -4,7535249 | 2,00E-06 | 2,41E-05 | |
| ENSGALG00000017083 | 1252,48764 | -0,54907192 | 0,15967948 | -3,4385878 | 0,00058476 | 0,00341401 | |
| ENSGALG00000006971 | 10409,6787 | -0,549657911 | 0,11864179 | -4,6329201 | 3,61E-06 | 4,12E-05 | |
| ENSGALG00000003026 | 1987,5595 | -0,550107166 | 0,16872812 | -3,260317 | 0,00111288 | 0,00583885 | MKL2 |
| ENSGALG00000011400 | 12123,5934 | -0,550294288 | 0,10312199 | -5,3363428 | 9,48E-08 | 1,53E-06 | USP44 |
| ENSGALG00000028501 | 7019,46403 | -0,550742687 | 0,11797631 | -4,6682481 | 3,04E-06 | 3,52E-05 | DCK |
| ENSGALG00000007097 | 2035,76357 | -0,551520351 | 0,16963125 | -3,2512898 | 0,00114883 | 0,00599972 | RPS6KA6 |
| ENSGALG00000017006 | 4626,73679 | -0,551986159 | 0,11796255 | -4,6793338 | 2,88E-06 | 3,35E-05 | SETDB2 |
| ENSGALG00000004071 | 3249,44294 | -0,552437495 | 0,12317196 | -4,4850915 | 7,29E-06 | 7,73E-05 | NETO2 |
| ENSGALG00000000684 | 3181,04311 | -0,552570682 | 0,14354544 | -3,849448 | 0,00011838 | 0,00086424 | KLHDC8A |
| ENSGALG00000003835 | 27823,2079 | -0,552761335 | 0,10615776 | -5,20698 | 1,92E-07 | 2,92E-06 | |
| ENSGALG00000016883 | 3231,09542 | -0,555002491 | 0,12790983 | -4,3390135 | 1,43E-05 | 0,00014131 | DOCK9 |
| ENSGALG00000004813 | 26912,1889 | -0,555221474 | 0,11731903 | -4,732578 | 2,22E-06 | 2,65E-05 | IARS |
| ENSGALG00000005774 | 2418,90539 | -0,555555373 | 0,13336098 | -4,1658014 | 3,10E-05 | 0,00027359 | |
| ENSGALG00000013635 | 10619,1615 | -0,555662229 | 0,12215986 | -4,5486483 | 5,40E-06 | 5,91E-05 | SCAF8 |
| ENSGALG00000016468 | 6793,09113 | -0,555855719 | 0,10894695 | -5,1020767 | 3,36E-07 | 4,84E-06 | |
| ENSGALG00000013430 | 2775,68087 | -0,557212283 | 0,1237629 | -4,5022564 | 6,72E-06 | 7,16E-05 | |
| ENSGALG00000021487 | 6790,2413 | -0,55890103 | 0,12790718 | -4,3695829 | 1,24E-05 | 0,00012525 | |
| ENSGALG00000016231 | 123947,503 | -0,55907395 | 0,10983383 | -5,0901798 | 3,58E-07 | 5,11E-06 | |
| ENSGALG00000019178 | 3990,13341 | -0,559313492 | 0,12498595 | -4,475011 | 7,64E-06 | 8,08E-05 | PRDM15 |
| ENSGALG00000010022 | 5530,32804 | -0,559381952 | 0,11898015 | -4,7014729 | 2,58E-06 | 3,04E-05 | PRMT9 |
| ENSGALG00000011692 | 9047,47741 | -0,559417827 | 0,11608282 | -4,8191269 | 1,44E-06 | 1,81E-05 | CDCA4 |
| ENSGALG00000000447 | 760,920402 | -0,559669779 | 0,17125604 | -3,2680295 | 0,00108299 | 0,00571066 | |
| ENSGALG00000011541 | 6975,19884 | -0,559795067 | 0,14049051 | -3,9845755 | 6,76E-05 | 0,00053309 | BIN1 |
| ENSGALG00000003510 | 5135,23749 | -0,562160497 | 0,117815 | -4,771553 | 1,83E-06 | 2,23E-05 | UCK2 |
| ENSGALG00000012508 | 4886,27256 | -0,562203872 | 0,11484296 | -4,8954143 | 9,81E-07 | 1,28E-05 | |
| ENSGALG00000027231 | 9354,14873 | -0,562988244 | 0,15371607 | -3,6625204 | 0,00024975 | 0,00165578 | NRARP |
| ENSGALG00000016897 | 2920,76606 | -0,563110321 | 0,12347462 | -4,560535 | 5,10E-06 | 5,64E-05 | GPR180 |
| ENSGALG00000010081 | 2535,91839 | -0,563235953 | 0,1656002 | -3,4011791 | 0,00067096 | 0,00383667 | SH3D19 |
| ENSGALG00000003074 | 3831,59863 | -0,563796907 | 0,1286586 | -4,382116 | 1,18E-05 | 0,0001188 | HOMER3 |
| ENSGALG00000012256 | 15406,5047 | -0,563809375 | 0,16328803 | -3,4528519 | 0,00055469 | 0,0032709 | UTRN |
| ENSGALG00000026438 | 2075,6069 | -0,563826313 | 0,13056342 | -4,3184093 | 1,57E-05 | 0,00015218 | |
| ENSGALG00000008132 | 7657,49338 | -0,564518166 | 0,11630408 | -4,8538123 | 1,21E-06 | 1,55E-05 | |
| ENSGALG00000022808 | 24156,7928 | -0,565497585 | 0,10524059 | -5,3733791 | 7,73E-08 | 1,27E-06 | |
| ENSGALG00000011228 | 7018,88755 | -0,565699287 | 0,12484784 | -4,5311101 | 5,87E-06 | 6,35E-05 | WARS |
| ENSGALG00000014940 | 2607,30528 | -0,566659105 | 0,15847617 | -3,575674 | 0,00034933 | 0,00221571 | ZUFSP |
| ENSGALG00000016917 | 1018,50229 | -0,567495374 | 0,16489149 | -3,4416292 | 0,00057822 | 0,00338311 | |
| ENSGALG00000015431 | 2610,45813 | -0,567537408 | 0,12799629 | -4,4340144 | 9,25E-06 | 9,61E-05 | |
| ENSGALG00000009855 | 7273,65346 | -0,567662024 | 0,12230573 | -4,6413362 | 3,46E-06 | 3,97E-05 | TBC1D9 |
| ENSGALG00000017079 | 12990,6555 | -0,56791639 | 0,12482195 | -4,5498117 | 5,37E-06 | 5,89E-05 | USPL1 |
| ENSGALG00000000195 | 1824,65996 | -0,568283448 | 0,16007893 | -3,5500202 | 0,0003852 | 0,00240134 | SVCT2 |
| ENSGALG00000003906 | 12475,0381 | -0,568387401 | 0,10947502 | -5,1919368 | 2,08E-07 | 3,14E-06 | MPHOSPH10 |
| ENSGALG00000016281 | 11421,2288 | -0,569360911 | 0,1081146 | -5,2662723 | 1,39E-07 | 2,17E-06 | DMD |
| ENSGALG00000016391 | 2197,60679 | -0,569812042 | 0,1354173 | -4,2078232 | 2,58E-05 | 0,00023357 | CNKSR2 |
| ENSGALG00000015142 | 3202,10584 | -0,570285644 | 0,13342262 | -4,2742801 | 1,92E-05 | 0,00018068 | DSG2 |
| ENSGALG00000009714 | 1150,55938 | -0,570305895 | 0,15558254 | -3,6656163 | 0,00024674 | 0,00164114 | CBR4 |
| ENSGALG00000014783 | 10636,0999 | -0,572122303 | 0,12610507 | -4,5368701 | 5,71E-06 | 6,20E-05 | |
| ENSGALG00000016910 | 23108,1995 | -0,572230903 | 0,11163907 | -5,1257228 | 2,96E-07 | 4,30E-06 | RBM26 |
| ENSGALG00000012849 | 5113,46952 | -0,57279907 | 0,14885384 | -3,8480636 | 0,00011906 | 0,00086774 | SLC37A3 |
| ENSGALG00000001363 | 21664,3518 | -0,573517858 | 0,10988778 | -5,2191233 | 1,80E-07 | 2,75E-06 | MYBBP1A |
| ENSGALG00000003180 | 15412,6584 | -0,573999724 | 0,10818558 | -5,3056952 | 1,12E-07 | 1,78E-06 | |
| ENSGALG00000013031 | 1094,94036 | -0,574014118 | 0,17360499 | -3,306438 | 0,0009449 | 0,0051051 | CECR1 |
| ENSGALG00000016807 | 10081,6726 | -0,574724846 | 0,11386342 | -5,0474932 | 4,48E-07 | 6,28E-06 | |
| ENSGALG00000016841 | 10674,7371 | -0,576008454 | 0,1276781 | -4,5114115 | 6,44E-06 | 6,90E-05 | |
| ENSGALG00000020057 | 1737,42699 | -0,576621955 | 0,13651224 | -4,2239578 | 2,40E-05 | 0,00021964 | PIGF |
| ENSGALG00000007992 | 6680,6643 | -0,5769061 | 0,10933512 | -5,2764942 | 1,32E-07 | 2,06E-06 | GGA3 |
| ENSGALG00000003591 | 1041,85294 | -0,577610116 | 0,17930224 | -3,2214327 | 0,00127551 | 0,00653599 | |
| ENSGALG00000014939 | 2493,31868 | -0,579384653 | 0,17435114 | -3,3230906 | 0,00089026 | 0,00486197 | GFM2 |
| ENSGALG00000004612 | 6633,12124 | -0,579433401 | 0,10947713 | -5,2927345 | 1,21E-07 | 1,90E-06 | MTHFR |

| | | | | | | | |
|---|---|---|---|---|---|---|---|
| ENSGALG00000007008 | 8124,26296 | -0,579649516 | 0,11753702 | -4,9316337 | 8,15E-07 | 1,08E-05 | POLR3E |
| ENSGALG00000004140 | 1503,56079 | -0,579940212 | 0,16796039 | -3,4528392 | 0,00055472 | 0,0032709 | SH3BP4 |
| ENSGALG00000015629 | 16787,8501 | -0,581629702 | 0,10138489 | -5,736848 | 9,65E-09 | 1,91E-07 | |
| ENSGALG00000010202 | 1508,60268 | -0,581722546 | 0,16884284 | -3,4453492 | 0,00057032 | 0,00334551 | ENTPD5 |
| ENSGALG00000006713 | 2292,89242 | -0,58181715 | 0,14500758 | -4,0123223 | 6,01E-05 | 0,00048209 | |
| ENSGALG00000011464 | 79055,7934 | -0,582311331 | 0,10582388 | -5,5026456 | 3,74E-08 | 6,56E-07 | EIF5 |
| ENSGALG00000017130 | 3473,80286 | -0,582570563 | 0,12916331 | -4,510341 | 6,47E-06 | 6,91E-05 | |
| ENSGALG00000015000 | 15510,2528 | -0,584898632 | 0,10441804 | -5,6015092 | 2,12E-08 | 3,93E-07 | LYAR |
| ENSGALG00000020899 | 644,115695 | -0,585391733 | 0,1791404 | -3,2677818 | 0,00108394 | 0,00571345 | |
| ENSGALG00000027517 | 979,679294 | -0,585667855 | 0,16395158 | -3,5722001 | 0,00035399 | 0,00224065 | gga-mir-3064 |
| ENSGALG00000007677 | 1856,87137 | -0,585818002 | 0,14520033 | -4,03455 | 5,47E-05 | 0,0004451 | THNSL1 |
| ENSGALG00000003426 | 1636,16491 | -0,587203746 | 0,15658355 | -3,7500986 | 0,00017677 | 0,00123292 | RHBDL3 |
| ENSGALG00000015765 | 12998,2541 | -0,587270267 | 0,1455632 | -4,0344695 | 5,47E-05 | 0,0004451 | |
| ENSGALG00000015310 | 4463,74355 | -0,587354563 | 0,1636253 | -3,5896318 | 0,00033115 | 0,00211768 | |
| ENSGALG00000004144 | 27007,8593 | -0,588131949 | 0,10975399 | -5,3586383 | 8,39E-08 | 1,37E-06 | |
| ENSGALG00000012946 | 11580,0518 | -0,588849778 | 0,11469761 | -5,1339326 | 2,84E-07 | 4,13E-06 | FAM60A |
| ENSGALG00000016187 | 8513,83254 | -0,588903063 | 0,11295401 | -5,2136534 | 1,85E-07 | 2,82E-06 | WDR4 |
| ENSGALG00000000284 | 7651,65914 | -0,589684267 | 0,12270008 | -4,8058996 | 1,54E-06 | 1,92E-05 | HOXB4 |
| ENSGALG00000015556 | 4802,19905 | -0,591646833 | 0,12793817 | -4,6244747 | 3,76E-06 | 4,27E-05 | AFAP1 |
| ENSGALG00000012147 | 18627,0418 | -0,591734529 | 0,11258805 | -5,255749 | 1,47E-07 | 2,30E-06 | DDX18 |
| ENSGALG00000008376 | 8424,82548 | -0,592197559 | 0,12144073 | -4,8764326 | 1,08E-06 | 1,40E-05 | EIF2B5 |
| ENSGALG00000005803 | 3271,76516 | -0,592808251 | 0,15820978 | -3,746976 | 0,00017898 | 0,0012457 | |
| ENSGALG00000002326 | 890,069728 | -0,593338649 | 0,18376433 | -3,2288021 | 0,0012431 | 0,00640113 | |
| ENSGALG00000011449 | 1934,52119 | -0,594721435 | 0,14281353 | -4,1643215 | 3,12E-05 | 0,00027501 | |
| ENSGALG00000007536 | 1357,15347 | -0,59551277 | 0,15673099 | -3,7995852 | 0,00014494 | 0,0010337 | |
| ENSGALG00000014213 | 4954,50424 | -0,596053951 | 0,12959904 | -4,5992158 | 4,24E-06 | 4,76E-05 | PRR5 |
| ENSGALG00000009458 | 5058,56826 | -0,596085417 | 0,1247109 | -4,7797377 | 1,76E-06 | 2,15E-05 | |
| ENSGALG00000005171 | 1357,22705 | -0,597406416 | 0,14458 | -4,1320128 | 3,60E-05 | 0,00031066 | AP1S3 |
| ENSGALG00000005243 | 13547,5044 | -0,597738338 | 0,10284356 | -5,8121126 | 6,17E-09 | 1,27E-07 | PPP3CB |
| ENSGALG00000020438 | 1854,20177 | -0,597817303 | 0,14024952 | -4,2625265 | 2,02E-05 | 0,00018927 | |
| ENSGALG00000027159 | 3433,75807 | -0,598718251 | 0,16764923 | -3,5712556 | 0,00035527 | 0,00224353 | CNKSR3 |
| ENSGALG00000014615 | 13865,3358 | -0,598774766 | 0,14500125 | -4,1294456 | 3,64E-05 | 0,00031335 | LAMA1 |
| ENSGALG00000011210 | 4763,25316 | -0,598839459 | 0,11293547 | -5,3024922 | 1,14E-07 | 1,81E-06 | FAM175A |
| ENSGALG00000020003 | 15165,0611 | -0,59891041 | 0,11112336 | -5,3895993 | 7,06E-08 | 1,17E-06 | MAP3K4 |
| ENSGALG00000015409 | 17031,8673 | -0,599227418 | 0,14738119 | -4,0658336 | 4,79E-05 | 0,00039565 | PROS1 |
| ENSGALG00000011710 | 2961,63072 | -0,599591029 | 0,18819478 | -3,186013 | 0,00144248 | 0,00722302 | |
| ENSGALG00000001003 | 4385,75735 | -0,600081977 | 0,12148953 | -4,9393718 | 7,84E-07 | 1,04E-05 | SEPN1 |
| ENSGALG00000006256 | 7733,33351 | -0,600800071 | 0,11264625 | -5,3335117 | 9,63E-08 | 1,55E-06 | |
| ENSGALG00000011280 | 7822,14955 | -0,603497491 | 0,12276846 | -4,9157374 | 8,84E-07 | 1,16E-05 | EEA1 |
| ENSGALG00000010603 | 12309,7524 | -0,604746743 | 0,10826314 | -5,585897 | 2,32E-08 | 4,26E-07 | |
| ENSGALG00000005807 | 96623,5673 | -0,604834873 | 0,10467707 | -5,7781028 | 7,55E-09 | 1,52E-07 | IPO7 |
| ENSGALG00000007030 | 1042,03468 | -0,605544059 | 0,16028433 | -3,7779368 | 0,00015813 | 0,00111611 | |
| ENSGALG00000010623 | 6764,86118 | -0,606701235 | 0,11188195 | -5,4226907 | 5,87E-08 | 9,92E-07 | ORC1 |
| ENSGALG00000006147 | 7732,277 | -0,607224855 | 0,1099367 | -5,5234047 | 3,32E-08 | 5,88E-07 | NUCB2 |
| ENSGALG00000009348 | 14314,7884 | -0,607303929 | 0,15062704 | -4,0318387 | 5,53E-05 | 0,00044957 | |
| ENSGALG00000011522 | 2524,77962 | -0,607850232 | 0,13426355 | -4,5272914 | 5,97E-06 | 6,44E-05 | TRMT61A |
| ENSGALG00000010538 | 11960,1749 | -0,60879058 | 0,11003246 | -5,5328272 | 3,15E-08 | 5,61E-07 | |
| ENSGALG00000011618 | 1230,20769 | -0,609057453 | 0,18958218 | -3,2126303 | 0,00131525 | 0,0067043 | ADSSL1 |
| ENSGALG00000015689 | 13523,3466 | -0,609077567 | 0,14382998 | -4,2347051 | 2,29E-05 | 0,00021053 | KIAA0368 |
| ENSGALG00000005492 | 6703,97246 | -0,609713298 | 0,14208413 | -4,2912131 | 1,78E-05 | 0,00016883 | |
| ENSGALG00000009698 | 6354,41268 | -0,610790424 | 0,15890149 | -3,8438306 | 0,00012113 | 0,00088176 | SLC38A1 |
| ENSGALG00000016302 | 7538,43287 | -0,61104581 | 0,14235289 | -4,2924722 | 1,77E-05 | 0,00016811 | |
| ENSGALG00000000438 | 3178,97206 | -0,61235395 | 0,14457738 | -4,2354755 | 2,28E-05 | 0,00020995 | ERMP1 |
| ENSGALG00000004169 | 13748,9297 | -0,613558741 | 0,10706722 | -5,7305937 | 1,00E-08 | 1,97E-07 | |
| ENSGALG00000013760 | 3088,02148 | -0,614432788 | 0,12107074 | -5,0749899 | 3,88E-07 | 5,50E-06 | |
| ENSGALG00000016574 | 1715,06369 | -0,614768808 | 0,14203902 | -4,3281683 | 1,50E-05 | 0,00014706 | GEMIN8 |
| ENSGALG00000007703 | 1248,99291 | -0,614958252 | 0,15352984 | -4,005464 | 6,19E-05 | 0,00049426 | |
| ENSGALG00000013480 | 4896,2242 | -0,615680556 | 0,12423491 | -4,9557774 | 7,20E-07 | 9,61E-06 | |
| ENSGALG00000016808 | 40422,889 | -0,615970468 | 0,12307828 | -5,0047047 | 5,59E-07 | 7,68E-06 | RANBP2 |
| ENSGALG00000006237 | 12767,8147 | -0,616678741 | 0,12728487 | -4,8448706 | 1,27E-06 | 1,61E-05 | PKN2 |
| ENSGALG00000016951 | 1808,55603 | -0,617422601 | 0,13578215 | -4,5471559 | 5,44E-06 | 5,94E-05 | MTRF1 |
| ENSGALG00000008265 | 23793,5701 | -0,619236444 | 0,10979734 | -5,639813 | 1,70E-08 | 3,22E-07 | |
| ENSGALG00000017388 | 6991,3984 | -0,619582917 | 0,11595165 | -5,3434592 | 9,12E-08 | 1,48E-06 | |
| ENSGALG00000005815 | 11029,7481 | -0,619938317 | 0,10990838 | -5,6405009 | 1,70E-08 | 3,22E-07 | TMEM41B |

| | | | | | | | |
|---|---|---|---|---|---|---|---|
| ENSGALG00000012533 | 37098,6017 | -0,619977689 | 0,14353496 | -4,3193496 | 1,56E-05 | 0,00015175 | |
| ENSGALG00000001192 | 22737,8249 | -0,620466491 | 0,15464988 | -4,0120722 | 6,02E-05 | 0,00048223 | |
| ENSGALG00000016653 | 1523,27891 | -0,621295808 | 0,14742536 | -4,2143076 | 2,51E-05 | 0,00022817 | |
| ENSGALG00000012447 | 5839,1863 | -0,62156312 | 0,12782967 | -4,8624324 | 1,16E-06 | 1,49E-05 | |
| ENSGALG00000008539 | 7630,14576 | -0,622137491 | 0,12115192 | -5,1351847 | 2,82E-07 | 4,11E-06 | ALG12 |
| ENSGALG00000023419 | 9218,5287 | -0,622730836 | 0,13105342 | -4,7517328 | 2,02E-06 | 2,42E-05 | HOXD4 |
| ENSGALG00000000234 | 2924,36008 | -0,623968223 | 0,14846856 | -4,2026959 | 2,64E-05 | 0,00023813 | EPB41L4A |
| ENSGALG00000005438 | 938,768709 | -0,624083264 | 0,1616022 | -3,8618489 | 0,00011253 | 0,00082862 | PARVA |
| ENSGALG00000010975 | 6618,99977 | -0,624280901 | 0,11571015 | -5,3952131 | 6,84E-08 | 1,14E-06 | TMTC2 |
| ENSGALG00000005504 | 1081,65735 | -0,624552568 | 0,17314094 | -3,6071917 | 0,00030953 | 0,00199538 | HNF1B |
| ENSGALG00000006609 | 9375,25541 | -0,624834648 | 0,13736708 | -4,5486491 | 5,40E-06 | 5,91E-05 | EPHB1 |
| ENSGALG00000014860 | 10686,5811 | -0,625239897 | 0,10707403 | -5,8393235 | 5,24E-09 | 1,09E-07 | YES1 |
| ENSGALG00000017159 | 5806,67135 | -0,626252438 | 0,14290102 | -4,3824212 | 1,17E-05 | 0,00011872 | ATM |
| ENSGALG00000004046 | 1117,46957 | -0,627070394 | 0,16993702 | -3,6900164 | 0,00022424 | 0,00150916 | LARP6 |
| ENSGALG00000014855 | 4504,15014 | -0,627700184 | 0,1611556 | -3,8949947 | 9,82E-05 | 0,00073827 | |
| ENSGALG00000010702 | 11951,0445 | -0,628390454 | 0,10835766 | -5,7992249 | 6,66E-09 | 1,36E-07 | ADSS |
| ENSGALG00000015472 | 37144,6363 | -0,629536006 | 0,14753725 | -4,266963 | 1,98E-05 | 0,00018598 | |
| ENSGALG00000012966 | 5358,75757 | -0,62976328 | 0,1160975 | -5,4244346 | 5,81E-08 | 9,83E-07 | ZBTB2 |
| ENSGALG00000005450 | 5500,73037 | -0,632586778 | 0,12083814 | -5,2349926 | 1,65E-07 | 2,54E-06 | CCDC66 |
| ENSGALG00000005256 | 11273,3374 | -0,63393871 | 0,16414924 | -3,8619656 | 0,00011248 | 0,00082862 | EPHA4 |
| ENSGALG00000014337 | 1369,70172 | -0,634337238 | 0,14622858 | -4,3379841 | 1,44E-05 | 0,00014177 | NWD2 |
| ENSGALG00000009850 | 11171,8574 | -0,63459607 | 0,10804099 | -5,8736605 | 4,26E-09 | 9,11E-08 | NOVA1 |
| ENSGALG00000004937 | 10776,4972 | -0,634904509 | 0,10951564 | -5,7973865 | 6,74E-09 | 1,37E-07 | VGLL4 |
| ENSGALG00000017368 | 7609,78676 | -0,635358308 | 0,10950994 | -5,8018322 | 6,56E-09 | 1,34E-07 | TRNT1 |
| ENSGALG00000007392 | 3781,92945 | -0,635704603 | 0,14088967 | -4,5120739 | 6,42E-06 | 6,88E-05 | |
| ENSGALG00000003767 | 4407,98301 | -0,636798593 | 0,11771515 | -5,4096572 | 6,31E-08 | 1,06E-06 | NKD1 |
| ENSGALG00000007710 | 1737,58327 | -0,637223247 | 0,15253964 | -4,1774273 | 2,95E-05 | 0,00026123 | |
| ENSGALG00000012813 | 15846,5259 | -0,637913408 | 0,12201058 | -5,228345 | 1,71E-07 | 2,63E-06 | |
| ENSGALG00000005521 | 7067,27493 | -0,637995859 | 0,13170481 | -4,844135 | 1,27E-06 | 1,62E-05 | PER2 |
| ENSGALG00000022490 | 838,24845 | -0,638026939 | 0,20482582 | -3,1149732 | 0,00183962 | 0,00878204 | |
| ENSGALG00000023451 | 10451,6138 | -0,638494594 | 0,10528216 | -6,0646041 | 1,32E-09 | 3,09E-08 | TRPM7 |
| ENSGALG00000007056 | 3476,50953 | -0,638790551 | 0,18102964 | -3,5286518 | 0,00041768 | 0,00258161 | |
| ENSGALG00000003891 | 23052,983 | -0,638963678 | 0,10013766 | -6,3808527 | 1,76E-10 | 4,67E-09 | ZNF423 |
| ENSGALG00000012662 | 3593,37854 | -0,639523486 | 0,12409213 | -5,1536182 | 2,56E-07 | 3,78E-06 | NUAK1 |
| ENSGALG00000014138 | 4048,40305 | -0,640267154 | 0,14067976 | -4,5512386 | 5,33E-06 | 5,86E-05 | NFXL1 |
| ENSGALG00000011162 | 1836,73645 | -0,640538648 | 0,15156233 | -4,2262391 | 2,38E-05 | 0,00021772 | CYP46A1 |
| ENSGALG00000009240 | 3768,49084 | -0,641416186 | 0,11670261 | -5,4961599 | 3,88E-08 | 6,79E-07 | NFE2L2 |
| ENSGALG00000007352 | 21915,6386 | -0,641667347 | 0,10923519 | -5,8741815 | 4,25E-09 | 9,10E-08 | |
| ENSGALG00000009445 | 3221,29491 | -0,642383978 | 0,12265432 | -5,2373531 | 1,63E-07 | 2,51E-06 | INPP5F |
| ENSGALG00000010642 | 1388,2464 | -0,643267076 | 0,1679801 | -3,8294242 | 0,00012844 | 0,00093118 | IRF2 |
| ENSGALG00000005321 | 14326,6827 | -0,645095033 | 0,14628713 | -4,4097866 | 1,03E-05 | 0,0001064 | |
| ENSGALG00000005024 | 4274,06304 | -0,645541534 | 0,13739879 | -4,6983057 | 2,62E-06 | 3,09E-05 | EFCC1 |
| ENSGALG00000026181 | 1587,87421 | -0,645581251 | 0,15765694 | -4,0948483 | 4,22E-05 | 0,00035587 | |
| ENSGALG00000010536 | 11449,0338 | -0,646034394 | 0,1106982 | -5,8359971 | 5,35E-09 | 1,11E-07 | FAF1 |
| ENSGALG00000016480 | 10063,5827 | -0,647091328 | 0,12228762 | -5,2915523 | 1,21E-07 | 1,91E-06 | SDC1 |
| ENSGALG00000000258 | 23273,6972 | -0,649481888 | 0,12291472 | -5,2840043 | 1,26E-07 | 1,98E-06 | |
| ENSGALG00000000761 | 2678,90106 | -0,649687901 | 0,13200757 | -4,9215959 | 8,58E-07 | 1,13E-05 | TSKU |
| ENSGALG00000014902 | 5870,32391 | -0,650325285 | 0,12934166 | -5,0279647 | 4,96E-07 | 6,90E-06 | |
| ENSGALG00000006864 | 655,136383 | -0,651213169 | 0,19973556 | -3,2603768 | 0,00111264 | 0,00583885 | |
| ENSGALG00000014873 | 10860,5943 | -0,6514166 | 0,10605415 | -6,1423019 | 8,13E-10 | 1,95E-08 | GJA1 |
| ENSGALG00000012232 | 769,021092 | -0,652989979 | 0,19777702 | -3,3016474 | 0,00096119 | 0,00517256 | DNAAF2 |
| ENSGALG00000003154 | 3509,17671 | -0,653496705 | 0,12218795 | -5,3482908 | 8,88E-08 | 1,45E-06 | TXNDC11 |
| ENSGALG00000005845 | 8323,59423 | -0,655210424 | 0,12474397 | -5,2524419 | 1,50E-07 | 2,33E-06 | SLC7A5 |
| ENSGALG00000015241 | 1466,0595 | -0,655397927 | 0,14176198 | -4,6232279 | 3,78E-06 | 4,29E-05 | |
| ENSGALG00000015653 | 733,019478 | -0,655959247 | 0,20273574 | -3,2355382 | 0,00121414 | 0,00627819 | |
| ENSGALG00000016681 | 817,923627 | -0,656368052 | 0,1696389 | -3,8692071 | 0,00010919 | 0,00080882 | DHRSX |
| ENSGALG00000000690 | 5466,00123 | -0,657633862 | 0,14151675 | -4,6470389 | 3,37E-06 | 3,87E-05 | GDPD4 |
| ENSGALG00000016817 | 6729,25411 | -0,658902643 | 0,1108429 | -5,944473 | 2,77E-09 | 6,16E-08 | RASA3 |
| ENSGALG00000006975 | 7158,07962 | -0,659116429 | 0,117355 | -5,6164324 | 1,95E-08 | 3,65E-07 | |
| ENSGALG00000006840 | 2751,68484 | -0,659559733 | 0,17118981 | -3,8527979 | 0,00011678 | 0,00085525 | |
| ENSGALG00000006681 | 619,278693 | -0,660452265 | 0,21494413 | -3,0726695 | 0,00212153 | 0,009926 | BRSK2 |
| ENSGALG00000010204 | 972,360103 | -0,660708753 | 0,16806539 | -3,9312602 | 8,45E-05 | 0,0006489 | KCNC2 |
| ENSGALG00000008551 | 7057,17591 | -0,661341847 | 0,12495655 | -5,2925743 | 1,21E-07 | 1,90E-06 | CRELD2 |
| ENSGALG00000007870 | 12818,5577 | -0,66197661 | 0,11665024 | -5,6748844 | 1,39E-08 | 2,68E-07 | |

| | | | | | | | |
|---|---|---|---|---|---|---|---|
| ENSGALG00000005796 | 13227,6265 | -0,664735985 | 0,10635028 | -6,2504396 | 4,09E-10 | 1,01E-08 | XBP1 |
| ENSGALG00000008293 | 829,335943 | -0,66560113 | 0,20054725 | -3,3189242 | 0,00090365 | 0,00491931 | SH3PXD2A |
| ENSGALG00000009915 | 13723,0991 | -0,66741216 | 0,11515882 | -5,7955799 | 6,81E-09 | 1,39E-07 | EML4 |
| ENSGALG00000005512 | 8645,01109 | -0,667653797 | 0,11848963 | -5,6347026 | 1,75E-08 | 3,31E-07 | |
| ENSGALG00000008418 | 2809,25172 | -0,668553187 | 0,12270622 | -5,4484051 | 5,08E-08 | 8,65E-07 | LARGE2 |
| ENSGALG00000011139 | 4725,82862 | -0,66929694 | 0,13400223 | -4,9946703 | 5,89E-07 | 8,03E-06 | |
| ENSGALG00000011686 | 12938,7332 | -0,670528172 | 0,14703222 | -4,5604165 | 5,11E-06 | 5,64E-05 | SEMA5B |
| ENSGALG00000008240 | 2340,29769 | -0,670953542 | 0,12956854 | -5,1783675 | 2,24E-07 | 3,35E-06 | CFLAR |
| ENSGALG00000006080 | 30589,3035 | -0,672288685 | 0,10183892 | -6,6014906 | 4,07E-11 | 1,17E-09 | GPC4 |
| ENSGALG00000028291 | 15611,0719 | -0,672824764 | 0,13419575 | -5,0137561 | 5,34E-07 | 7,39E-06 | |
| ENSGALG00000004715 | 1867,94138 | -0,672888292 | 0,15250355 | -4,4122795 | 1,02E-05 | 0,00010534 | |
| ENSGALG00000015912 | 3388,05891 | -0,674238434 | 0,14603918 | -4,6168326 | 3,90E-06 | 4,41E-05 | |
| ENSGALG00000015708 | 22628,5619 | -0,674375402 | 0,17848955 | -3,7782347 | 0,00015794 | 0,00111536 | FGFR3 |
| ENSGALG00000008684 | 70609,009 | -0,674536207 | 0,12880685 | -5,236804 | 1,63E-07 | 2,52E-06 | |
| ENSGALG00000010049 | 749,198004 | -0,6748978 | 0,20825089 | -3,2407919 | 0,00119198 | 0,00618241 | PTPRB |
| ENSGALG00000003249 | 7468,82123 | -0,675105103 | 0,11166729 | -6,0456838 | 1,49E-09 | 3,45E-08 | CWF19L1 |
| ENSGALG00000006035 | 6729,77158 | -0,676290578 | 0,10833566 | -6,2425484 | 4,30E-10 | 1,06E-08 | PDE3B |
| ENSGALG00000004494 | 4756,9197 | -0,676838081 | 0,11957995 | -5,66013 | 1,51E-08 | 2,89E-07 | CCNE1 |
| ENSGALG00000016451 | 7945,90073 | -0,679900325 | 0,13943898 | -4,8759701 | 1,08E-06 | 1,40E-05 | ROCK2 |
| ENSGALG00000006437 | 3771,99053 | -0,682364759 | 0,12175344 | -5,6044805 | 2,09E-08 | 3,87E-07 | |
| ENSGALG00000009156 | 5048,98485 | -0,682556058 | 0,16761459 | -4,0721757 | 4,66E-05 | 0,0003862 | CDAN1 |
| ENSGALG00000014274 | 3304,6353 | -0,683648491 | 0,13949356 | -4,9009322 | 9,54E-07 | 1,25E-05 | |
| ENSGALG00000012390 | 2074,46154 | -0,684899295 | 0,14739748 | -4,6466147 | 3,37E-06 | 3,87E-05 | |
| ENSGALG00000009152 | 8451,92517 | -0,685418448 | 0,13463812 | -5,0908202 | 3,57E-07 | 5,10E-06 | ATRNL1 |
| ENSGALG00000004631 | 4926,28212 | -0,686947368 | 0,19666311 | -3,4930159 | 0,0004776 | 0,00288615 | DRAXIN |
| ENSGALG00000010836 | 1260,23388 | -0,688393097 | 0,16687599 | -4,1251777 | 3,70E-05 | 0,00031781 | |
| ENSGALG00000015879 | 11968,4164 | -0,689623134 | 0,14800439 | -4,6594775 | 3,17E-06 | 3,66E-05 | HUNK |
| ENSGALG00000008234 | 5250,56204 | -0,689849994 | 0,12730961 | -5,4186797 | 6,00E-08 | 1,01E-06 | ORC2 |
| ENSGALG00000016308 | 5133,40137 | -0,690314419 | 0,12814377 | -5,3870307 | 7,16E-08 | 1,18E-06 | MYC |
| ENSGALG00000014686 | 1912,60456 | -0,691021625 | 0,17304639 | -3,9932738 | 6,52E-05 | 0,00051675 | FBN2 |
| ENSGALG00000025627 | 543,306297 | -0,691903941 | 0,1957541 | -3,5345567 | 0,00040846 | 0,00253126 | SNORD47 |
| ENSGALG00000008292 | 9417,47651 | -0,69263042 | 0,12226746 | -5,6648794 | 1,47E-08 | 2,83E-07 | |
| ENSGALG00000011679 | 1487,18993 | -0,695574487 | 0,17407076 | -3,9959294 | 6,44E-05 | 0,00051158 | |
| ENSGALG00000007371 | 25835,4514 | -0,695805113 | 0,11656083 | -5,9694591 | 2,38E-09 | 5,36E-08 | IGDCC3 |
| ENSGALG00000017220 | 8610,42217 | -0,695886193 | 0,12577786 | -5,5326606 | 3,15E-08 | 5,61E-07 | |
| ENSGALG00000008758 | 3528,49517 | -0,696392948 | 0,11831828 | -5,8857597 | 3,96E-09 | 8,56E-08 | |
| ENSGALG00000028415 | 2970,11444 | -0,698216592 | 0,20411152 | -3,4207604 | 0,00062446 | 0,00360716 | |
| ENSGALG00000016834 | 1336,22512 | -0,698351437 | 0,16279355 | -4,2897979 | 1,79E-05 | 0,00016974 | MCF2L |
| ENSGALG00000002316 | 6016,72913 | -0,700708149 | 0,10987572 | -6,3772792 | 1,80E-10 | 4,77E-09 | NUBP2 |
| ENSGALG00000002689 | 3193,98974 | -0,701352786 | 0,13477202 | -5,2039941 | 1,95E-07 | 2,96E-06 | |
| ENSGALG00000007501 | 495,825919 | -0,702573548 | 0,22104241 | -3,1784559 | 0,00148062 | 0,00735447 | |
| ENSGALG00000006592 | 563,467935 | -0,703616422 | 0,19089215 | -3,6859369 | 0,00022786 | 0,00153051 | |
| ENSGALG00000021316 | 17378,3782 | -0,704586437 | 0,10231836 | -6,8862169 | 5,73E-12 | 1,88E-10 | |
| ENSGALG00000012153 | 842,909942 | -0,706186244 | 0,17364706 | -4,0667908 | 4,77E-05 | 0,00039426 | METTL15 |
| ENSGALG00000002762 | 6873,84376 | -0,70662137 | 0,1320656 | -5,3505333 | 8,77E-08 | 1,43E-06 | UHRF1BP1 |
| ENSGALG00000002726 | 8639,35675 | -0,708755608 | 0,13953348 | -5,0794664 | 3,78E-07 | 5,38E-06 | KIF1B |
| ENSGALG00000007141 | 2245,3684 | -0,70907976 | 0,1459375 | -4,8587906 | 1,18E-06 | 1,52E-05 | LRRK1 |
| ENSGALG00000011680 | 3554,59737 | -0,709602993 | 0,18365321 | -3,8638203 | 0,00011163 | 0,00082355 | EPHA5 |
| ENSGALG00000012848 | 3745,19397 | -0,711139877 | 0,17381561 | -4,0913464 | 4,29E-05 | 0,00035977 | |
| ENSGALG00000022857 | 997,971589 | -0,711204976 | 0,16756929 | -4,2442441 | 2,19E-05 | 0,000203 | |
| ENSGALG00000015733 | 14565,4266 | -0,711632203 | 0,17400694 | -4,0896772 | 4,32E-05 | 0,00036185 | |
| ENSGALG00000017365 | 2855,56643 | -0,712049377 | 0,15174035 | -4,6925512 | 2,70E-06 | 3,17E-05 | |
| ENSGALG00000009562 | 5126,97314 | -0,714412903 | 0,12876119 | -5,548356 | 2,88E-08 | 5,20E-07 | |
| ENSGALG00000003970 | 25720,8179 | -0,714513793 | 0,12407312 | -5,7588122 | 8,47E-09 | 1,70E-07 | |
| ENSGALG00000003161 | 1569,92422 | -0,715518291 | 0,16581373 | -4,3151933 | 1,59E-05 | 0,00015387 | |
| ENSGALG00000016927 | 1190,18904 | -0,715628819 | 0,17643861 | -4,0559649 | 4,99E-05 | 0,00041123 | |
| ENSGALG00000013615 | 780,849369 | -0,715808204 | 0,18333875 | -3,9042931 | 9,45E-05 | 0,00071361 | |
| ENSGALG00000010036 | 359,240321 | -0,717787007 | 0,2219662 | -3,2337671 | 0,00122169 | 0,00630861 | FOSL2 |
| ENSGALG00000015122 | 2528,57664 | -0,719672556 | 0,13908654 | -5,174279 | 2,29E-07 | 3,42E-06 | TAF4B |
| ENSGALG00000020805 | 5789,80169 | -0,719835862 | 0,11391762 | -6,318916 | 2,63E-10 | 6,72E-09 | LRRC8D |
| ENSGALG00000005430 | 950,090619 | -0,72235333 | 0,20225367 | -3,5715216 | 0,00035491 | 0,00224229 | ALS2CL |
| ENSGALG00000002553 | 2782,54105 | -0,723490416 | 0,16062171 | -4,5043127 | 6,66E-06 | 7,10E-05 | PCDH1 |
| ENSGALG00000007242 | 1612,83074 | -0,724284111 | 0,14362964 | -5,0427204 | 4,59E-07 | 6,42E-06 | SLITRK4 |
| ENSGALG00000008078 | 7758,49805 | -0,725207523 | 0,12521944 | -5,7914932 | 6,98E-09 | 1,42E-07 | |

| | | | | | | | |
|---|---|---|---|---|---|---|---|
| ENSGALG00000006978 | 1693,58452 | -0,725851222 | 0,16965998 | -4,2782701 | 1,88E-05 | 0,00017784 | APOOL |
| ENSGALG00000004004 | 8836,32885 | -0,725989536 | 0,12198937 | -5,9512525 | 2,66E-09 | 5,95E-08 | PHKB |
| ENSGALG00000005246 | 9658,47999 | -0,7272373 | 0,12335529 | -5,8954691 | 3,74E-09 | 8,11E-08 | |
| ENSGALG00000028294 | 3609,41474 | -0,727698928 | 0,13043446 | -5,5790387 | 2,42E-08 | 4,40E-07 | VWA5B2 |
| ENSGALG00000006939 | 26463,2292 | -0,728311968 | 0,1092521 | -6,6663427 | 2,62E-11 | 7,76E-10 | PFKP |
| ENSGALG00000006555 | 1381,69164 | -0,729109536 | 0,15543069 | -4,6908983 | 2,72E-06 | 3,19E-05 | |
| ENSGALG00000025743 | 3141,82765 | -0,729896626 | 0,1215881 | -6,0030268 | 1,94E-09 | 4,42E-08 | CDR2 |
| ENSGALG00000016078 | 860,370977 | -0,73004938 | 0,19386389 | -3,7657832 | 0,00016603 | 0,00116639 | |
| ENSGALG00000014419 | 9837,23866 | -0,730417334 | 0,1131065 | -6,4577842 | 1,06E-10 | 2,87E-09 | |
| ENSGALG00000020479 | 3560,77871 | -0,731996794 | 0,12990873 | -5,6347005 | 1,75E-08 | 3,31E-07 | |
| ENSGALG00000003378 | 7303,91386 | -0,732735156 | 0,12423155 | -5,8981405 | 3,68E-09 | 8,01E-08 | USP42 |
| ENSGALG00000015627 | 1040,22039 | -0,73285779 | 0,21291454 | -3,4420279 | 0,00057737 | 0,00337958 | |
| ENSGALG00000026349 | 29196,624 | -0,736634172 | 0,09993715 | -7,3709745 | 1,69E-13 | 6,80E-12 | |
| ENSGALG00000007069 | 592,208375 | -0,738101541 | 0,23259838 | -3,1732876 | 0,00150723 | 0,00745945 | |
| ENSGALG00000016537 | 11629,8341 | -0,739933749 | 0,1619397 | -4,5691932 | 4,90E-06 | 5,45E-05 | SCML2 |
| ENSGALG00000010448 | 40598,9396 | -0,742255902 | 0,1099244 | -6,7524215 | 1,45E-11 | 4,48E-10 | LTBP1 |
| ENSGALG00000002679 | 1869,60302 | -0,743662338 | 0,15155473 | -4,9068896 | 9,25E-07 | 1,21E-05 | GLT1D1 |
| ENSGALG00000007331 | 18072,4344 | -0,744880346 | 0,10843853 | -6,8691486 | 6,46E-12 | 2,09E-10 | SVIL |
| ENSGALG00000027639 | 27019,6011 | -0,74689172 | 0,10811506 | -6,9083041 | 4,90E-12 | 1,61E-10 | |
| ENSGALG00000010522 | 11132,1004 | -0,747786716 | 0,11992122 | -6,2356496 | 4,50E-10 | 1,11E-08 | |
| ENSGALG00000003975 | 3354,11348 | -0,748226558 | 0,14508752 | -5,1570704 | 2,51E-07 | 3,71E-06 | |
| ENSGALG00000008910 | 360,623018 | -0,749041615 | 0,22125446 | -3,3854305 | 0,00071067 | 0,00402652 | CERKL |
| ENSGALG00000003149 | 1878,4222 | -0,749443102 | 0,15295343 | -4,8998124 | 9,59E-07 | 1,25E-05 | |
| ENSGALG00000007011 | 11491,2627 | -0,750635431 | 0,11252384 | -6,6709014 | 2,54E-11 | 7,56E-10 | |
| ENSGALG00000023464 | 2176,79492 | -0,751478603 | 0,16284203 | -4,6147706 | 3,94E-06 | 4,45E-05 | |
| ENSGALG00000019961 | 8882,89233 | -0,7532835 | 0,12923015 | -5,8290072 | 5,58E-09 | 1,16E-07 | SLC18B1 |
| ENSGALG00000003430 | 9094,42264 | -0,754857044 | 0,10828451 | -6,9710529 | 3,15E-12 | 1,08E-10 | PMS2 |
| ENSGALG00000015862 | 12207,7026 | -0,756350316 | 0,14990125 | -5,0456572 | 4,52E-07 | 6,34E-06 | |
| ENSGALG00000008318 | 18620,5264 | -0,756789619 | 0,11792666 | -6,4174599 | 1,39E-10 | 3,71E-09 | SLK |
| ENSGALG00000010412 | 643,758245 | -0,756929566 | 0,17972006 | -4,2117143 | 2,53E-05 | 0,00023004 | WWTR1 |
| ENSGALG00000004981 | 3521,90259 | -0,757070103 | 0,13293065 | -5,6952262 | 1,23E-08 | 2,40E-07 | TSEN2 |
| ENSGALG00000006981 | 1290,89613 | -0,757579596 | 0,16920706 | -4,477234 | 7,56E-06 | 8,01E-05 | |
| ENSGALG00000007667 | 1024,27903 | -0,757882718 | 0,16652285 | -4,5512235 | 5,33E-06 | 5,86E-05 | GPR158 |
| ENSGALG00000000742 | 7852,97756 | -0,757900463 | 0,13118075 | -5,777528 | 7,58E-09 | 1,53E-07 | |
| ENSGALG00000001419 | 2086,70105 | -0,758492411 | 0,14074025 | -5,389307 | 7,07E-08 | 1,17E-06 | DAB2IP |
| ENSGALG00000016294 | 651,260147 | -0,761040192 | 0,2034016 | -3,7415644 | 0,00018288 | 0,00126776 | HMGCLL1 |
| ENSGALG00000009748 | 12417,6034 | -0,762363604 | 0,12194724 | -6,2515854 | 4,06E-10 | 1,01E-08 | ASNS |
| ENSGALG00000006167 | 4665,12691 | -0,763429455 | 0,16136845 | -4,7309711 | 2,23E-06 | 2,66E-05 | LRRC8B |
| ENSGALG00000017462 | 3834,70932 | -0,763892042 | 0,11771244 | -6,4894761 | 8,61E-11 | 2,38E-09 | FNIP1 |
| ENSGALG00000028254 | 428,450125 | -0,764100303 | 0,21188759 | -3,6061588 | 0,00031076 | 0,00199955 | KIAA1033 |
| ENSGALG00000007300 | 11025,5462 | -0,764508198 | 0,12386685 | -6,1720162 | 6,74E-10 | 1,63E-08 | |
| ENSGALG00000026862 | 22574,7451 | -0,764882325 | 0,16241479 | -4,7094378 | 2,48E-06 | 2,94E-05 | CLDN1 |
| ENSGALG00000008621 | 9873,26199 | -0,76612038 | 0,13164901 | -5,8194163 | 5,91E-09 | 1,22E-07 | NRP2 |
| ENSGALG00000027389 | 3852,81541 | -0,766788012 | 0,13085522 | -5,8598199 | 4,63E-09 | 9,83E-08 | RGS7BP |
| ENSGALG00000013782 | 3281,28093 | -0,769533335 | 0,1760353 | -4,3714718 | 1,23E-05 | 0,00012439 | CDH7 |
| ENSGALG00000026897 | 591,300069 | -0,770711311 | 0,20311525 | -3,7944532 | 0,00014797 | 0,00105311 | |
| ENSGALG00000028737 | 5045,35121 | -0,771568179 | 0,14540814 | -5,3062243 | 1,12E-07 | 1,78E-06 | |
| ENSGALG00000026518 | 766,815675 | -0,773283921 | 0,17841103 | -4,3342831 | 1,46E-05 | 0,00014396 | RUNDC3A |
| ENSGALG00000005610 | 640,252491 | -0,773621314 | 0,24264802 | -3,1882449 | 0,00143139 | 0,00718021 | SLC44A3 |
| ENSGALG00000004594 | 1038,5627 | -0,774792529 | 0,20891066 | -3,7087267 | 0,0002083 | 0,00141449 | CD74 |
| ENSGALG00000013627 | 453,51839 | -0,775913381 | 0,21080288 | -3,6807532 | 0,00023255 | 0,00155583 | SLC7A2 |
| ENSGALG00000015887 | 12708,1034 | -0,776552733 | 0,12924098 | -6,0085643 | 1,87E-09 | 4,28E-08 | URB1 |
| ENSGALG00000015208 | 6752,93068 | -0,778489641 | 0,1611505 | -4,8308236 | 1,36E-06 | 1,72E-05 | ROR2 |
| ENSGALG00000006258 | 9759,66383 | -0,778635815 | 0,12249805 | -6,3563117 | 2,07E-10 | 5,40E-09 | SPOCK1 |
| ENSGALG00000009514 | 5928,6059 | -0,779501664 | 0,11478642 | -6,7908877 | 1,11E-11 | 3,50E-10 | NRCAM |
| ENSGALG00000001020 | 7177,33389 | -0,780112538 | 0,1084839 | -7,1910444 | 6,43E-13 | 2,41E-11 | |
| ENSGALG00000013683 | 18236,9052 | -0,780228434 | 0,15315117 | -5,0944986 | 3,50E-07 | 5,02E-06 | ARID1B |
| ENSGALG00000008961 | 32098,1427 | -0,780468343 | 0,16688611 | -4,6766524 | 2,92E-06 | 3,39E-05 | |
| ENSGALG00000012834 | 693,301303 | -0,78090617 | 0,24310402 | -3,2122306 | 0,00131709 | 0,00671113 | AKR1D1 |
| ENSGALG00000009393 | 806,584777 | -0,782303357 | 0,19134913 | -4,088356 | 4,34E-05 | 0,00036327 | |
| ENSGALG00000012329 | 10571,2971 | -0,782765701 | 0,1633862 | -4,7908925 | 1,66E-06 | 2,05E-05 | |
| ENSGALG00000026970 | 1108,65361 | -0,785311616 | 0,15637208 | -5,0220706 | 5,11E-07 | 7,09E-06 | |
| ENSGALG00000009252 | 7419,33117 | -0,788959505 | 0,12427971 | -6,3482569 | 2,18E-10 | 5,65E-09 | |
| ENSGALG00000012808 | 5957,05996 | -0,789210521 | 0,13604537 | -5,8010834 | 6,59E-09 | 1,35E-07 | CDYL |

| | | | | | | | |
|---|---|---|---|---|---|---|---|
| ENSGALG00000006908 | 14243,3267 | -0,789724754 | 0,14863119 | -5,3133179 | 1,08E-07 | 1,72E-06 | DLG1 |
| ENSGALG00000000639 | 611,516533 | -0,79001541 | 0,22610591 | -3,4940061 | 0,00047583 | 0,00287674 | LRRN2 |
| ENSGALG00000016412 | 2161,43136 | -0,791360693 | 0,13463882 | -5,8776562 | 4,16E-09 | 8,94E-08 | MBOAT2 |
| ENSGALG00000005785 | 2650,42523 | -0,791416097 | 0,18516638 | -4,274081 | 1,92E-05 | 0,00018071 | |
| ENSGALG00000009428 | 6528,61969 | -0,792261942 | 0,14165832 | -5,5927667 | 2,23E-08 | 4,11E-07 | SUSD6 |
| ENSGALG00000004662 | 3631,94871 | -0,792400882 | 0,16649406 | -4,7593343 | 1,94E-06 | 2,35E-05 | DISP3 |
| ENSGALG00000001995 | 9089,14914 | -0,794307619 | 0,11001846 | -7,2197668 | 5,21E-13 | 1,99E-11 | LIMS2 |
| ENSGALG00000012317 | 261,455312 | -0,794799886 | 0,25524866 | -3,1138259 | 0,00184678 | 0,008807 | |
| ENSGALG00000025464 | 234,066267 | -0,795229903 | 0,25312021 | -3,1417085 | 0,00167965 | 0,00816148 | SNORA68 |
| ENSGALG00000004132 | 38489,9584 | -0,797075856 | 0,11456459 | -6,9574364 | 3,47E-12 | 1,17E-10 | |
| ENSGALG00000026486 | 578,46167 | -0,797913674 | 0,2123431 | -3,7576624 | 0,00017151 | 0,00120056 | |
| ENSGALG00000008378 | 25019,9708 | -0,798096236 | 0,12248113 | -6,5160751 | 7,22E-11 | 2,01E-09 | SRGAP3 |
| ENSGALG00000009932 | 6037,71039 | -0,798314026 | 0,13917513 | -5,7360394 | 9,69E-09 | 1,92E-07 | THADA |
| ENSGALG00000011045 | 4900,98782 | -0,799075807 | 0,1382254 | -5,7809621 | 7,43E-09 | 1,50E-07 | |
| ENSGALG00000014907 | 2917,0076 | -0,80019622 | 0,15045066 | -5,3186621 | 1,05E-07 | 1,68E-06 | DCBLD1 |
| ENSGALG00000010910 | 3629,98348 | -0,801671392 | 0,12110467 | -6,619657 | 3,60E-11 | 1,04E-09 | |
| ENSGALG00000012576 | 1359,57449 | -0,801916632 | 0,1553997 | -5,1603485 | 2,46E-07 | 3,65E-06 | RASEF |
| ENSGALG00000000583 | 9340,93564 | -0,802342978 | 0,11958733 | -6,7092639 | 1,96E-11 | 5,94E-10 | SOX13 |
| ENSGALG00000002900 | 2509,77542 | -0,803716995 | 0,14588715 | -5,5091691 | 3,61E-08 | 6,34E-07 | NOTUM |
| ENSGALG00000027521 | 307,657411 | -0,803725098 | 0,23936001 | -3,3578085 | 0,00078563 | 0,00439095 | |
| ENSGALG00000013212 | 21085,1244 | -0,805085082 | 0,11564313 | -6,9618063 | 3,36E-12 | 1,14E-10 | |
| ENSGALG00000001883 | 2479,1556 | -0,805297687 | 0,12694065 | -6,3438913 | 2,24E-10 | 5,80E-09 | ST6GALNAC2 |
| ENSGALG00000003105 | 820,425544 | -0,805784417 | 0,19045019 | -4,2309456 | 2,33E-05 | 0,00021336 | ANKFN1 |
| ENSGALG00000002461 | 9290,02388 | -0,808165073 | 0,1405519 | -5,7499407 | 8,93E-09 | 1,77E-07 | |
| ENSGALG00000002242 | 344,643988 | -0,808545932 | 0,22942797 | -3,5241821 | 0,00042479 | 0,00262173 | |
| ENSGALG00000017103 | 2337,4348 | -0,811501513 | 0,12799906 | -6,3399023 | 2,30E-10 | 5,93E-09 | WASF3 |
| ENSGALG00000014957 | 24709,1325 | -0,81285735 | 0,11248513 | -7,226354 | 4,96E-13 | 1,90E-11 | WDR1 |
| ENSGALG00000003427 | 8550,66088 | -0,813323099 | 0,12341029 | -6,5903992 | 4,39E-11 | 1,25E-09 | USP3 |
| ENSGALG00000011089 | 415,157392 | -0,814043658 | 0,22510112 | -3,6163466 | 0,00029879 | 0,00193531 | |
| ENSGALG00000011643 | 20679,5817 | -0,816325927 | 0,14686948 | -5,5581725 | 2,73E-08 | 4,94E-07 | UTP20 |
| ENSGALG00000010785 | 498,938482 | -0,818155142 | 0,1993699 | -4,1037045 | 4,07E-05 | 0,00034533 | ACOT11 |
| ENSGALG00000001220 | 3875,78907 | -0,818155857 | 0,12990789 | -6,2979691 | 3,02E-10 | 7,62E-09 | HERPUD1 |
| ENSGALG00000002598 | 7963,93283 | -0,82047281 | 0,10859443 | -7,5553857 | 4,18E-14 | 1,82E-12 | TPST1 |
| ENSGALG00000011511 | 101055,806 | -0,820640197 | 0,12654102 | -6,4851715 | 8,86E-11 | 2,44E-09 | CKB |
| ENSGALG00000003160 | 1417,99184 | -0,823423786 | 0,17588287 | -4,68166 | 2,85E-06 | 3,32E-05 | |
| ENSGALG00000006872 | 8998,32455 | -0,823650254 | 0,11488863 | -7,1691189 | 7,55E-13 | 2,79E-11 | PISD |
| ENSGALG00000007039 | 11318,9545 | -0,823801989 | 0,1338606 | -6,1541783 | 7,55E-10 | 1,82E-08 | |
| ENSGALG00000004569 | 2222,44652 | -0,82424563 | 0,16546211 | -4,9814767 | 6,31E-07 | 8,53E-06 | UNC5B |
| ENSGALG00000010507 | 1443,23999 | -0,824303692 | 0,1501991 | -5,4880735 | 4,06E-08 | 7,08E-07 | |
| ENSGALG00000003391 | 10590,4517 | -0,827807721 | 0,10949157 | -7,5604701 | 4,02E-14 | 1,76E-12 | EIF2AK1 |
| ENSGALG00000009546 | 1643,01363 | -0,832251183 | 0,15248225 | -5,4580202 | 4,81E-08 | 8,27E-07 | |
| ENSGALG00000007717 | 242,860186 | -0,834765713 | 0,25520347 | -3,270981 | 0,00107175 | 0,00566236 | GCNT7 |
| ENSGALG00000014954 | 7661,41015 | -0,835213371 | 0,11582931 | -7,2107259 | 5,57E-13 | 2,10E-11 | ZNF518B |
| ENSGALG00000008469 | 4008,42168 | -0,83875822 | 0,14975785 | -5,6007628 | 2,13E-08 | 3,94E-07 | SPINT1 |
| ENSGALG00000002543 | 5887,09033 | -0,840502021 | 0,12400471 | -6,7779847 | 1,22E-11 | 3,81E-10 | CSGALNACT2 |
| ENSGALG00000015980 | 410,785738 | -0,841274637 | 0,25003277 | -3,3646576 | 0,00076639 | 0,00429398 | |
| ENSGALG00000011388 | 12926,4854 | -0,842306194 | 0,12106876 | -6,9572544 | 3,47E-12 | 1,17E-10 | RABGGTB |
| ENSGALG00000015519 | 4471,93343 | -0,844394446 | 0,12450731 | -6,7818863 | 1,19E-11 | 3,71E-10 | ROBO2 |
| ENSGALG00000015656 | 12813,5339 | -0,849003497 | 0,12458324 | -6,814749 | 9,44E-12 | 3,00E-10 | |
| ENSGALG00000012998 | 2617,34593 | -0,849149352 | 0,17683084 | -4,8020434 | 1,57E-06 | 1,96E-05 | WNT5B |
| ENSGALG00000024984 | 247,007719 | -0,849437052 | 0,24634113 | -3,4482145 | 0,00056431 | 0,00332167 | SNORD74 |
| ENSGALG00000023609 | 5841,30372 | -0,855145185 | 0,13510752 | -6,3293679 | 2,46E-10 | 6,31E-09 | |
| ENSGALG00000014178 | 3963,51548 | -0,855544377 | 0,15025645 | -5,6938946 | 1,24E-08 | 2,41E-07 | SCUBE1 |
| ENSGALG00000012394 | 13948,8666 | -0,855781498 | 0,10361657 | -8,259118 | 1,47E-16 | 8,54E-15 | NID2 |
| ENSGALG00000003668 | 1594,44053 | -0,85921128 | 0,14225789 | -6,0398144 | 1,54E-09 | 3,57E-08 | |
| ENSGALG00000007846 | 1019,28883 | -0,863598394 | 0,23089134 | -3,7402805 | 0,00018381 | 0,00127223 | |
| ENSGALG00000010311 | 430,972494 | -0,865329105 | 0,20595707 | -4,2015023 | 2,65E-05 | 0,00023907 | NAV3 |
| ENSGALG00000015681 | 7286,43618 | -0,869123123 | 0,13859875 | -6,270786 | 3,59E-10 | 9,02E-09 | NRIP1 |
| ENSGALG00000010010 | 1250,74925 | -0,869227214 | 0,15347637 | -5,6635898 | 1,48E-08 | 2,84E-07 | RHOQ |
| ENSGALG00000008815 | 1450,48158 | -0,86979829 | 0,16636094 | -5,2283804 | 1,71E-07 | 2,63E-06 | LRRN4 |
| ENSGALG00000011376 | 402,726863 | -0,870171475 | 0,21157345 | -4,1128575 | 3,91E-05 | 0,00033421 | ANKRD9 |
| ENSGALG00000016896 | 2175,65472 | -0,874558686 | 0,16022347 | -5,4583681 | 4,81E-08 | 8,26E-07 | ABCC4 |
| ENSGALG00000023581 | 218,882989 | -0,876769713 | 0,27869933 | -3,145934 | 0,00165557 | 0,00806463 | |
| ENSGALG00000009129 | 1209,82081 | -0,878296092 | 0,17155705 | -5,1195568 | 3,06E-07 | 4,43E-06 | DLX5 |

| | | | | | | | |
|---|---|---|---|---|---|---|---|
| ENSGALG00000004542 | 7447,83595 | -0,879244031 | 0,1445917 | -6,0808747 | 1,20E-09 | 2,81E-08 | ZMYND8 |
| ENSGALG00000016462 | 9552,69875 | -0,880157248 | 0,13995983 | -6,2886421 | 3,20E-10 | 8,07E-09 | MYCN |
| ENSGALG00000028602 | 5443,19542 | -0,881510014 | 0,15763284 | -5,5921725 | 2,24E-08 | 4,12E-07 | |
| ENSGALG00000011120 | 494,969816 | -0,88227849 | 0,23574872 | -3,7424529 | 0,00018223 | 0,00126458 | |
| ENSGALG00000003657 | 764,546995 | -0,88270712 | 0,19877995 | -4,4406246 | 8,97E-06 | 9,38E-05 | |
| ENSGALG00000028320 | 417,903769 | -0,88451445 | 0,27202551 | -3,2515863 | 0,00114763 | 0,00599848 | |
| ENSGALG00000001749 | 4549,2907 | -0,88875485 | 0,12757912 | -6,9663033 | 3,25E-12 | 1,11E-10 | ACSBG2 |
| ENSGALG00000005723 | 2016,64963 | -0,890647194 | 0,1398661 | -6,3678562 | 1,92E-10 | 5,03E-09 | |
| ENSGALG00000009360 | 602,385427 | -0,890830846 | 0,19246128 | -4,6286237 | 3,68E-06 | 4,19E-05 | CAPN2 |
| ENSGALG00000002555 | 1358,14983 | -0,892902927 | 0,21167678 | -4,2182374 | 2,46E-05 | 0,00022468 | RET |
| ENSGALG00000000307 | 8913,08352 | -0,894951246 | 0,11346269 | -7,8876258 | 3,08E-15 | 1,50E-13 | |
| ENSGALG00000012012 | 5383,1455 | -0,895703636 | 0,17255692 | -5,1907721 | 2,09E-07 | 3,15E-06 | |
| ENSGALG00000017870 | 493,118342 | -0,896459218 | 0,20520273 | -4,3686516 | 1,25E-05 | 0,00012543 | SNORD24 |
| ENSGALG00000008547 | 11178,1998 | -0,896850751 | 0,12624913 | -7,1038172 | 1,21E-12 | 4,41E-11 | |
| ENSGALG00000008509 | 336,231933 | -0,897122538 | 0,26693729 | -3,3607988 | 0,00077717 | 0,00434905 | |
| ENSGALG00000012814 | 2520,13193 | -0,898069575 | 0,12792713 | -7,0201654 | 2,22E-12 | 7,76E-11 | PXDC1 |
| ENSGALG00000017838 | 224,751439 | -0,898451215 | 0,26076736 | -3,4454129 | 0,00057019 | 0,00334551 | SNORA62 |
| ENSGALG00000009687 | 2155,9656 | -0,89897016 | 0,15726738 | -5,7161896 | 1,09E-08 | 2,13E-07 | KCNK2 |
| ENSGALG00000014645 | 469,375441 | -0,89901657 | 0,21753348 | -4,1327733 | 3,58E-05 | 0,00030983 | MEF2C |
| ENSGALG00000006498 | 20254,306 | -0,899416586 | 0,16914983 | -5,3172776 | 1,05E-07 | 1,69E-06 | |
| ENSGALG00000026790 | 419,691292 | -0,903869706 | 0,25606475 | -3,5298482 | 0,0004158 | 0,0025729 | DMRT2 |
| ENSGALG00000002704 | 377,643334 | -0,906514113 | 0,2414033 | -3,7551852 | 0,00017321 | 0,00121125 | |
| ENSGALG00000026258 | 289,161398 | -0,909500707 | 0,24047052 | -3,7821713 | 0,00015547 | 0,00110014 | MAF |
| ENSGALG00000004231 | 640,948749 | -0,911405982 | 0,18347202 | -4,9675475 | 6,78E-07 | 9,10E-06 | IFNLR1 |
| ENSGALG00000013861 | 1063,53974 | -0,913164571 | 0,17955855 | -5,0856088 | 3,66E-07 | 5,22E-06 | TNFAIP3 |
| ENSGALG00000020723 | 828,748724 | -0,913543779 | 0,17554467 | -5,204053 | 1,95E-07 | 2,96E-06 | |
| ENSGALG00000005994 | 969,598357 | -0,91862167 | 0,17763077 | -5,1715232 | 2,32E-07 | 3,46E-06 | DERL3 |
| ENSGALG00000010683 | 599,988497 | -0,919156764 | 0,20193219 | -4,551809 | 5,32E-06 | 5,85E-05 | |
| ENSGALG00000001656 | 26148,0576 | -0,922800056 | 0,11046642 | -8,3536705 | 6,62E-17 | 4,00E-15 | |
| ENSGALG00000009540 | 5865,4242 | -0,9229711 | 0,11652612 | -7,9207228 | 2,36E-15 | 1,17E-13 | YAF2 |
| ENSGALG00000025166 | 165,346098 | -0,924288426 | 0,28745972 | -3,2153668 | 0,00130278 | 0,00665566 | SNORD75 |
| ENSGALG00000001000 | 110996,795 | -0,924720639 | 0,13746595 | -6,726907 | 1,73E-11 | 5,28E-10 | |
| ENSGALG00000008305 | 1229,58651 | -0,925580955 | 0,18545643 | -4,9908269 | 6,01E-07 | 8,18E-06 | |
| ENSGALG00000016843 | 7409,4478 | -0,926939677 | 0,12855412 | -7,2105013 | 5,57E-13 | 2,10E-11 | COL4A2 |
| ENSGALG00000012911 | 418,92558 | -0,929872331 | 0,26011606 | -3,5748363 | 0,00035045 | 0,00222129 | SYT10 |
| ENSGALG00000008811 | 6177,80088 | -0,931412058 | 0,14890526 | -6,255065 | 3,97E-10 | 9,87E-09 | SLC1A4 |
| ENSGALG00000007952 | 17655,501 | -0,932963194 | 0,1344416 | -6,9395427 | 3,93E-12 | 1,32E-10 | AMOT |
| ENSGALG00000023241 | 1184,67586 | -0,938819916 | 0,18825868 | -4,9868614 | 6,14E-07 | 8,32E-06 | |
| ENSGALG00000028708 | 6182,01115 | -0,938854547 | 0,12586313 | -7,4593292 | 8,70E-14 | 3,59E-12 | |
| ENSGALG00000009415 | 20765,0987 | -0,939656238 | 0,14021273 | -6,7016471 | 2,06E-11 | 6,21E-10 | SMOC1 |
| ENSGALG00000012726 | 95084,9683 | -0,941672551 | 0,12611408 | -7,4668311 | 8,21E-14 | 3,42E-12 | |
| ENSGALG00000015622 | 657,059398 | -0,944352456 | 0,21737746 | -4,344298 | 1,40E-05 | 0,00013845 | |
| ENSGALG00000015028 | 404,3337 | -0,946739996 | 0,25007245 | -3,7858628 | 0,00015318 | 0,00108585 | |
| ENSGALG00000011258 | 44996,7983 | -0,946927699 | 0,14493435 | -6,5334939 | 6,43E-11 | 1,81E-09 | |
| ENSGALG00000011068 | 1649,85658 | -0,947751489 | 0,14451976 | -6,5579373 | 5,46E-11 | 1,54E-09 | COBLL1 |
| ENSGALG00000005601 | 522,730238 | -0,948098308 | 0,22540074 | -4,2062786 | 2,60E-05 | 0,00023462 | |
| ENSGALG00000006808 | 9236,5969 | -0,948868926 | 0,10714219 | -8,8561653 | 8,28E-19 | 6,00E-17 | TSPAN6 |
| ENSGALG00000013193 | 667,613336 | -0,948945842 | 0,21648961 | -4,3833321 | 1,17E-05 | 0,00011831 | IRX2 |
| ENSGALG00000023608 | 3676,98571 | -0,950348621 | 0,13455102 | -7,0631096 | 1,63E-12 | 5,82E-11 | |
| ENSGALG00000005805 | 4566,75731 | -0,954718819 | 0,14551669 | -6,5608886 | 5,35E-11 | 1,52E-09 | PLCD1 |
| ENSGALG00000023463 | 1598,1146 | -0,956997326 | 0,17129068 | -5,5869784 | 2,31E-08 | 4,24E-07 | |
| ENSGALG00000004393 | 1366,86861 | -0,962441655 | 0,2126961 | -4,5249615 | 6,04E-06 | 6,51E-05 | |
| ENSGALG00000004284 | 19106,5125 | -0,967926178 | 0,15442375 | -6,2679877 | 3,66E-10 | 9,15E-09 | |
| ENSGALG00000027342 | 11330,7864 | -0,968330352 | 0,11546028 | -8,3866967 | 5,00E-17 | 3,08E-15 | |
| ENSGALG00000015908 | 1062,96226 | -0,970529728 | 0,20345544 | -4,7702325 | 1,84E-06 | 2,24E-05 | COL12A1 |
| ENSGALG00000011833 | 13327,6665 | -0,971207285 | 0,11374653 | -8,5383464 | 1,36E-17 | 8,83E-16 | |
| ENSGALG00000009381 | 15338,368 | -0,972226453 | 0,1330298 | -7,3083359 | 2,70E-13 | 1,06E-11 | |
| ENSGALG00000015940 | 2397,4582 | -0,972506334 | 0,14243979 | -6,8274905 | 8,64E-12 | 2,76E-10 | |
| ENSGALG00000007367 | 230,098228 | -0,973082329 | 0,26469761 | -3,6762037 | 0,00023673 | 0,00157994 | WFIKKN2 |
| ENSGALG00000010597 | 2446,53537 | -0,973101915 | 0,12722398 | -7,6487303 | 2,03E-14 | 9,27E-13 | |
| ENSGALG00000027497 | 292,355516 | -0,973179721 | 0,23408863 | -4,157313 | 3,22E-05 | 0,00028231 | |
| ENSGALG00000019397 | 12676,4129 | -0,975439675 | 0,11488543 | -8,4905432 | 2,06E-17 | 1,31E-15 | CERK |
| ENSGALG00000002103 | 289,282544 | -0,979001464 | 0,28612607 | -3,4215737 | 0,0006226 | 0,00359944 | TPPP3 |
| ENSGALG00000015624 | 34457,4291 | -0,97910183 | 0,17589785 | -5,5663091 | 2,60E-08 | 4,72E-07 | VCAN |

| | | | | | | | |
|---|---|---|---|---|---|---|---|
| ENSGALG00000015164 | 3228,28619 | -0,982829392 | 0,14802221 | -6,6397428 | 3,14E-11 | 9,24E-10 | |
| ENSGALG00000020788 | 1411,73564 | -0,986044977 | 0,14888196 | -6,6229983 | 3,52E-11 | 1,02E-09 | |
| ENSGALG00000001371 | 2323,32831 | -0,990350626 | 0,1647412 | -6,0115541 | 1,84E-09 | 4,21E-08 | |
| ENSGALG00000009758 | 793,28474 | -0,991410492 | 0,19270405 | -5,1447309 | 2,68E-07 | 3,94E-06 | SLC7A11 |
| ENSGALG00000015440 | 170,656307 | -0,996123586 | 0,28603433 | -3,4825316 | 0,0004967 | 0,00298434 | |
| ENSGALG00000010692 | 6955,66893 | -0,996294098 | 0,13891979 | -7,1717217 | 7,41E-13 | 2,76E-11 | LRP8 |
| ENSGALG00000026582 | 1194,48697 | -0,997100015 | 0,19077348 | -5,2266176 | 1,73E-07 | 2,64E-06 | |
| ENSGALG00000002532 | 116,305922 | -0,997428466 | 0,31689231 | -3,1475313 | 0,00164655 | 0,00803219 | GJA4 |
| ENSGALG00000010163 | 10928,5236 | -0,998496131 | 0,12690773 | -7,8678908 | 3,61E-15 | 1,74E-13 | LGR5 |
| ENSGALG00000007483 | 18280,1601 | -0,999963194 | 0,14992811 | -6,669618 | 2,56E-11 | 7,61E-10 | LRIG1 |
| ENSGALG00000005483 | 1190,77186 | -1,005472997 | 0,15529553 | -6,4745779 | 9,51E-11 | 2,59E-09 | CDH13 |
| ENSGALG00000011295 | 1389,14492 | -1,007421144 | 0,15807983 | -6,3728633 | 1,86E-10 | 4,90E-09 | SOCS2 |
| ENSGALG00000016627 | 15863,8272 | -1,008110823 | 0,10822868 | -9,3146368 | 1,22E-20 | 1,03E-18 | |
| ENSGALG00000008281 | 355,49596 | -1,009962725 | 0,2355243 | -4,2881467 | 1,80E-05 | 0,00017082 | NEURL1 |
| ENSGALG00000015078 | 5440,75757 | -1,010765666 | 0,18233191 | -5,5435478 | 2,96E-08 | 5,32E-07 | MLLT3 |
| ENSGALG00000015112 | 8576,86268 | -1,013051509 | 0,1332664 | -7,6017026 | 2,92E-14 | 1,31E-12 | |
| ENSGALG00000000949 | 297,083499 | -1,013784064 | 0,23293191 | -4,3522764 | 1,35E-05 | 0,0001336 | HBEGF |
| ENSGALG00000016616 | 1435,92487 | -1,01551489 | 0,1675974 | -6,0592521 | 1,37E-09 | 3,18E-08 | |
| ENSGALG00000009450 | 2394,04421 | -1,016064164 | 0,12964787 | -7,8371065 | 4,61E-15 | 2,19E-13 | DUSP10 |
| ENSGALG00000001009 | 527,817322 | -1,018620159 | 0,24676578 | -4,1278826 | 3,66E-05 | 0,00031509 | |
| ENSGALG00000015896 | 2100,38648 | -1,023057651 | 0,16795043 | -6,0914263 | 1,12E-09 | 2,64E-08 | |
| ENSGALG00000010156 | 274,639072 | -1,025083823 | 0,24796248 | -4,134028 | 3,56E-05 | 0,00030892 | |
| ENSGALG00000013124 | 3974,90345 | -1,026819624 | 0,16230106 | -6,3266353 | 2,51E-10 | 6,40E-09 | FHOD3 |
| ENSGALG00000006138 | 2396,05253 | -1,026936244 | 0,15045199 | -6,825674 | 8,75E-12 | 2,79E-10 | |
| ENSGALG00000010579 | 4573,02342 | -1,028169899 | 0,12570944 | -8,1789393 | 2,86E-16 | 1,62E-14 | STON2 |
| ENSGALG00000012439 | 1314,57587 | -1,028638845 | 0,22792485 | -4,5130614 | 6,39E-06 | 6,86E-05 | |
| ENSGALG00000019077 | 11786,6675 | -1,030078317 | 0,1385218 | -7,436218 | 1,04E-13 | 4,25E-12 | |
| ENSGALG00000000977 | 7324,72056 | -1,032464943 | 0,10973117 | -9,4090394 | 5,01E-21 | 4,29E-19 | MAPKAP1 |
| ENSGALG00000016754 | 1717,85966 | -1,034769759 | 0,17703324 | -5,8450593 | 5,06E-09 | 1,06E-07 | |
| ENSGALG00000012412 | 7272,99248 | -1,036824012 | 0,14525293 | -7,1380593 | 9,47E-13 | 3,47E-11 | MTHFD1L |
| ENSGALG00000027213 | 84,2176986 | -1,037130822 | 0,33646244 | -3,0824565 | 0,002053 | 0,00965178 | ALX1 |
| ENSGALG00000007555 | 13407,2897 | -1,038877738 | 0,11478529 | -9,050617 | 1,42E-19 | 1,11E-17 | CCND1 |
| ENSGALG00000008465 | 8001,13375 | -1,045042375 | 0,12988092 | -8,0461579 | 8,54E-16 | 4,60E-14 | SORCS1 |
| ENSGALG00000022800 | 1489,66503 | -1,045072381 | 0,14566903 | -7,1742935 | 7,27E-13 | 2,72E-11 | |
| ENSGALG00000017714 | 665,00414 | -1,052113718 | 0,20833312 | -5,050151 | 4,41E-07 | 6,20E-06 | |
| ENSGALG00000015975 | 5069,41908 | -1,056406602 | 0,12845648 | -8,2238485 | 1,97E-16 | 1,13E-14 | |
| ENSGALG00000011581 | 10286,8368 | -1,059360643 | 0,33858403 | -3,1287969 | 0,00175524 | 0,00846159 | KIF26A |
| ENSGALG00000015736 | 279,628079 | -1,060236168 | 0,31923414 | -3,3211867 | 0,00089636 | 0,00488488 | |
| ENSGALG00000003295 | 2261,70268 | -1,063705711 | 0,15858273 | -6,7075762 | 1,98E-11 | 6,00E-10 | |
| ENSGALG00000009740 | 783,871387 | -1,063900656 | 0,18158266 | -5,8590432 | 4,66E-09 | 9,86E-08 | RASGRP1 |
| ENSGALG00000013604 | 164,158201 | -1,065244459 | 0,28526693 | -3,7342024 | 0,00018831 | 0,0012975 | VIP |
| ENSGALG00000011236 | 190,29168 | -1,070413941 | 0,31505499 | -3,3975464 | 0,00067993 | 0,00388308 | BEGAIN |
| ENSGALG00000017135 | 1406,42375 | -1,071209509 | 0,16120082 | -6,6451865 | 3,03E-11 | 8,92E-10 | CRYL1 |
| ENSGALG00000002988 | 7561,04394 | -1,072357906 | 0,15209899 | -7,0503948 | 1,78E-12 | 6,33E-11 | PHGDH |
| ENSGALG00000001370 | 1171,23107 | -1,073396904 | 0,19159469 | -5,6024356 | 2,11E-08 | 3,91E-07 | ADAMTS8 |
| ENSGALG00000027757 | 10827,9262 | -1,076478944 | 0,10801659 | -9,9658664 | 2,15E-23 | 2,27E-21 | ARL4A |
| ENSGALG00000010891 | 1573,41259 | -1,079203445 | 0,18453813 | -5,8481327 | 4,97E-09 | 1,04E-07 | ABCB11 |
| ENSGALG00000027707 | 112,772942 | -1,082314729 | 0,31181061 | -3,4710644 | 0,0005184 | 0,0030902 | ESM1 |
| ENSGALG00000016899 | 32,6413126 | -1,08275528 | 0,35267355 | -3,0701346 | 0,00213962 | 0,00998662 | DCT |
| ENSGALG00000000177 | 4145,02133 | -1,083701806 | 0,12196683 | -8,8852173 | 6,38E-19 | 4,70E-17 | |
| ENSGALG00000025770 | 345,961225 | -1,089960517 | 0,25250889 | -4,3165234 | 1,59E-05 | 0,00015316 | |
| ENSGALG00000028046 | 6843,87336 | -1,098357134 | 0,11398468 | -9,6360065 | 5,63E-22 | 5,11E-20 | INSIG2 |
| ENSGALG00000012917 | 4548,57829 | -1,102419391 | 0,19071062 | -5,7805874 | 7,44E-09 | 1,50E-07 | CDH6 |
| ENSGALG00000005842 | 443,936227 | -1,104947048 | 0,2118757 | -5,215072 | 1,84E-07 | 2,80E-06 | GDPD2 |
| ENSGALG00000012095 | 202,109683 | -1,108243378 | 0,27163773 | -4,079858 | 4,51E-05 | 0,00037525 | C14orf105 |
| ENSGALG00000002518 | 128,674403 | -1,109129449 | 0,33692493 | -3,2919186 | 0,00099506 | 0,00531078 | |
| ENSGALG00000009567 | 5248,50516 | -1,113206952 | 0,1654659 | -6,7277124 | 1,72E-11 | 5,27E-10 | TLL1 |
| ENSGALG00000011439 | 1241,39048 | -1,116853678 | 0,16224057 | -6,8839359 | 5,82E-12 | 1,90E-10 | |
| ENSGALG00000013548 | 35,2312467 | -1,118469658 | 0,33863762 | -3,3028512 | 0,00095707 | 0,00515652 | GZMA |
| ENSGALG00000014852 | 15743,6418 | -1,119019515 | 0,11136376 | -10,048327 | 9,34E-24 | 1,05E-21 | |
| ENSGALG00000009179 | 4347,97295 | -1,119360692 | 0,13379231 | -8,3664053 | 5,94E-17 | 3,63E-15 | TNFSF10 |
| ENSGALG00000011172 | 2541,5225 | -1,121518184 | 0,12741815 | -8,8018714 | 1,35E-18 | 9,35E-17 | |
| ENSGALG00000013892 | 1340,40625 | -1,122648268 | 0,1612643 | -6,9615422 | 3,37E-12 | 1,14E-10 | MAP3K5 |
| ENSGALG00000001079 | 117,890946 | -1,122757301 | 0,33560094 | -3,345513 | 0,00082131 | 0,0045345 | WNT3 |

| | | | | | | | |
|---|---|---|---|---|---|---|---|
| ENSGALG00000016680 | 793,768127 | -1,12428894 | 0,23420199 | -4,8005098 | 1,58E-06 | 1,97E-05 | TFAP2B |
| ENSGALG00000008014 | 284,580595 | -1,126068911 | 0,26416635 | -4,2627266 | 2,02E-05 | 0,00018923 | CEBPB |
| ENSGALG00000006337 | 35,9815104 | -1,128489411 | 0,34919354 | -3,2317018 | 0,00123055 | 0,00634861 | |
| ENSGALG00000020151 | 65,1037326 | -1,129234483 | 0,35223689 | -3,205895 | 0,00134643 | 0,0068343 | NMU |
| ENSGALG00000006376 | 107,4182 | -1,138122247 | 0,32739087 | -3,4763409 | 0,00050831 | 0,00303934 | TMC3 |
| ENSGALG00000027210 | 288,324353 | -1,142415301 | 0,23806373 | -4,7987792 | 1,60E-06 | 1,98E-05 | |
| ENSGALG00000001085 | 2988,78652 | -1,143093103 | 0,13119308 | -8,7130596 | 2,96E-18 | 1,98E-16 | |
| ENSGALG00000001276 | 18773,5176 | -1,144775826 | 0,13161296 | -8,6980477 | 3,38E-18 | 2,24E-16 | HOXB9 |
| ENSGALG00000004155 | 218,309793 | -1,147734088 | 0,27623338 | -4,1549435 | 3,25E-05 | 0,00028451 | MYOM3 |
| ENSGALG00000027107 | 1206,95445 | -1,149521147 | 0,15677875 | -7,3321234 | 2,27E-13 | 9,02E-12 | |
| ENSGALG00000014864 | 769,880572 | -1,150010609 | 0,17012701 | -6,7597182 | 1,38E-11 | 4,28E-10 | |
| ENSGALG00000027885 | 31,9835607 | -1,151437329 | 0,34943778 | -3,295114 | 0,00098382 | 0,00526107 | C4orf19 |
| ENSGALG00000010926 | 145,649999 | -1,152009571 | 0,31597679 | -3,6458677 | 0,00026649 | 0,00175025 | |
| ENSGALG00000008875 | 1752,66896 | -1,156831558 | 0,15899216 | -7,2760291 | 3,44E-13 | 1,34E-11 | PLCB1 |
| ENSGALG00000007268 | 3702,32926 | -1,168892699 | 0,14170635 | -8,2486969 | 1,60E-16 | 9,28E-15 | |
| ENSGALG00000006584 | 535,759637 | -1,169466644 | 0,23137239 | -5,054478 | 4,32E-07 | 6,07E-06 | |
| ENSGALG00000016114 | 1343,20194 | -1,174926553 | 0,19801604 | -5,9334919 | 2,97E-09 | 6,53E-08 | |
| ENSGALG00000022871 | 818,020575 | -1,176605905 | 0,18677065 | -6,2997367 | 2,98E-10 | 7,55E-09 | |
| ENSGALG00000026763 | 724,902739 | -1,177313255 | 0,20694833 | -5,6889236 | 1,28E-08 | 2,48E-07 | |
| ENSGALG00000017411 | 73,3262064 | -1,17758282 | 0,34864579 | -3,3775909 | 0,00073124 | 0,00412419 | RXFP3 |
| ENSGALG00000016998 | 449,62073 | -1,177970615 | 0,21440175 | -5,494221 | 3,92E-08 | 6,86E-07 | LPAR6 |
| ENSGALG00000008941 | 29,8088387 | -1,184874519 | 0,3508426 | -3,3772253 | 0,00073221 | 0,00412626 | FEZF1 |
| ENSGALG00000002509 | 14719,4349 | -1,185807324 | 0,13269584 | -8,9362812 | 4,02E-19 | 2,98E-17 | |
| ENSGALG00000012357 | 5240,54414 | -1,187472008 | 0,12252712 | -9,6915034 | 3,28E-22 | 3,16E-20 | CXCR4 |
| ENSGALG00000015665 | 1052,14248 | -1,188036426 | 0,17158557 | -6,9238716 | 4,39E-12 | 1,45E-10 | SH3BP2 |
| ENSGALG00000005619 | 863,255886 | -1,189698364 | 0,2239703 | -5,3118577 | 1,09E-07 | 1,73E-06 | |
| ENSGALG00000005685 | 3302,12612 | -1,189953127 | 0,15706062 | -7,576394 | 3,55E-14 | 1,57E-12 | KSR1 |
| ENSGALG00000016083 | 216,722511 | -1,190269979 | 0,27117041 | -4,38938 | 1,14E-05 | 0,0001155 | |
| ENSGALG00000000936 | 1054,13064 | -1,193444435 | 0,15734156 | -7,5850553 | 3,32E-14 | 1,47E-12 | |
| ENSGALG00000006170 | 3710,82448 | -1,194070483 | 0,14887865 | -8,0204281 | 1,05E-15 | 5,52E-14 | LRRC8C |
| ENSGALG00000015433 | 4115,11204 | -1,199115509 | 0,12542332 | -9,560547 | 1,17E-21 | 1,05E-19 | ABCA1 |
| ENSGALG00000002923 | 1146,0005 | -1,199213505 | 0,15769125 | -7,6048196 | 2,85E-14 | 1,29E-12 | |
| ENSGALG00000011104 | 29,5888843 | -1,203717219 | 0,35160223 | -3,4235198 | 0,00061816 | 0,00358289 | GCG |
| ENSGALG00000012224 | 1928,96823 | -1,20528636 | 0,15139325 | -7,961295 | 1,70E-15 | 8,65E-14 | |
| ENSGALG00000003582 | 6407,42119 | -1,218636528 | 0,17846722 | -6,8283493 | 8,59E-12 | 2,75E-10 | SLC1A3 |
| ENSGALG00000009667 | 660,393085 | -1,219318734 | 0,24952478 | -4,8865638 | 1,03E-06 | 1,33E-05 | MFAP3L |
| ENSGALG00000002632 | 1307,71653 | -1,220611399 | 0,17395739 | -7,0167265 | 2,27E-12 | 7,93E-11 | CALCRL |
| ENSGALG00000009215 | 122,787447 | -1,220936874 | 0,30573548 | -3,9934419 | 6,51E-05 | 0,00051668 | |
| ENSGALG00000000141 | 479,975266 | -1,223136562 | 0,25876918 | -4,7267474 | 2,28E-06 | 2,72E-05 | BLB1 |
| ENSGALG00000006473 | 340,04191 | -1,223226887 | 0,26548467 | -4,6075237 | 4,07E-06 | 4,58E-05 | PLXNA4 |
| ENSGALG00000007660 | 3604,79456 | -1,228207524 | 0,16378808 | -7,4987599 | 6,44E-14 | 2,69E-12 | |
| ENSGALG00000013239 | 67578,5287 | -1,231185989 | 0,13442151 | -9,1591442 | 5,23E-20 | 4,24E-18 | |
| ENSGALG00000008615 | 1056,6531 | -1,233824562 | 0,18606143 | -6,6312754 | 3,33E-11 | 9,74E-10 | |
| ENSGALG00000003739 | 39854,6748 | -1,237172502 | 0,12123912 | -10,2044 | 1,89E-24 | 2,28E-22 | |
| ENSGALG00000029023 | 187,057616 | -1,242039059 | 0,26924919 | -4,6129723 | 3,97E-06 | 4,48E-05 | DMRTB1 |
| ENSGALG00000009532 | 9821,7952 | -1,242374232 | 0,11357599 | -10,938705 | 7,53E-28 | 1,13E-25 | GXYLT1 |
| ENSGALG00000011867 | 2059,31955 | -1,242989678 | 0,14218556 | -8,7420246 | 2,29E-18 | 1,56E-16 | SERHL2 |
| ENSGALG00000005427 | 736,096188 | -1,25622691 | 0,18500374 | -6,7902783 | 1,12E-11 | 3,51E-10 | |
| ENSGALG00000013237 | 693,51796 | -1,260226776 | 0,21528589 | -5,8537361 | 4,81E-09 | 1,01E-07 | |
| ENSGALG00000010858 | 12929,1308 | -1,268608971 | 0,32118282 | -3,9498033 | 7,82E-05 | 0,00060506 | LRP2 |
| ENSGALG00000003353 | 12512,0835 | -1,269046359 | 0,17509801 | -7,2476344 | 4,24E-13 | 1,63E-11 | RAI14 |
| ENSGALG00000019552 | 24,9825451 | -1,269295425 | 0,34407756 | -3,6889806 | 0,00022515 | 0,00151456 | SERPINB2 |
| ENSGALG00000004361 | 87800,4537 | -1,276561341 | 0,12683773 | -10,064524 | 7,93E-24 | 8,99E-22 | |
| ENSGALG00000008352 | 151,362805 | -1,283975751 | 0,29199529 | -4,3972481 | 1,10E-05 | 0,00011206 | |
| ENSGALG00000002397 | 837,302305 | -1,291814757 | 0,2176946 | -5,9340689 | 2,96E-09 | 6,52E-08 | PSPH |
| ENSGALG00000006401 | 4187,88984 | -1,299022844 | 0,12943055 | -10,036447 | 1,05E-23 | 1,16E-21 | OSBPL5 |
| ENSGALG00000014866 | 52452,7894 | -1,312013425 | 0,13134051 | -9,9894042 | 1,70E-23 | 1,83E-21 | |
| ENSGALG00000005442 | 270,987771 | -1,327990169 | 0,28317276 | -4,6896819 | 2,74E-06 | 3,20E-05 | PALMD |
| ENSGALG00000016602 | 1659,68203 | -1,339565689 | 0,16372183 | -8,1819615 | 2,79E-16 | 1,58E-14 | |
| ENSGALG00000021879 | 40,0432745 | -1,34129965 | 0,34849488 | -3,8488361 | 0,00011868 | 0,00086593 | |
| ENSGALG00000002443 | 492,983426 | -1,353430658 | 0,22722109 | -5,9564481 | 2,58E-09 | 5,77E-08 | FBXL16 |
| ENSGALG00000007771 | 2716,44612 | -1,353742442 | 0,15148338 | -8,9365738 | 4,01E-19 | 2,98E-17 | FAM210B |
| ENSGALG00000008491 | 3635,92779 | -1,354708919 | 0,1393335 | -9,7227798 | 2,41E-22 | 2,38E-20 | |
| ENSGALG00000009947 | 4729,08145 | -1,358616298 | 0,11919082 | -11,398666 | 4,25E-30 | 7,32E-28 | PLEKHH2 |

| | | | | | | | |
|---|---|---|---|---|---|---|---|
| ENSGALG00000001346 | 421,970817 | -1,362109597 | 0,2357782 | -5,7770803 | 7,60E-09 | 1,53E-07 | |
| ENSGALG00000028048 | 17805,9742 | -1,362450241 | 0,10389675 | -13,113502 | 2,76E-39 | 8,73E-37 | |
| ENSGALG00000028047 | 613,76602 | -1,369008166 | 0,2271663 | -6,026458 | 1,68E-09 | 3,86E-08 | RHOV |
| ENSGALG00000010148 | 96,0680074 | -1,379685752 | 0,32859473 | -4,1987459 | 2,68E-05 | 0,00024168 | |
| ENSGALG00000003706 | 2880,21855 | -1,390323026 | 0,15220073 | -9,1347989 | 6,55E-20 | 5,25E-18 | |
| ENSGALG00000009405 | 351,467766 | -1,391691484 | 0,2252283 | -6,1790258 | 6,45E-10 | 1,57E-08 | GRIA2 |
| ENSGALG00000014983 | 1427,7077 | -1,39278689 | 0,17525642 | -7,9471375 | 1,91E-15 | 9,66E-14 | F2R |
| ENSGALG00000002993 | 1938,95885 | -1,397301236 | 0,15816908 | -8,834225 | 1,01E-18 | 7,22E-17 | CA10 |
| ENSGALG00000015256 | 9909,97899 | -1,397562325 | 0,12338907 | -11,326468 | 9,70E-30 | 1,61E-27 | PAM |
| ENSGALG00000003114 | 426,741711 | -1,398035927 | 0,21620264 | -6,4663221 | 1,00E-10 | 2,72E-09 | NOG |
| ENSGALG00000011306 | 2909,70938 | -1,403496478 | 0,13240055 | -10,600382 | 2,97E-26 | 4,17E-24 | DES |
| ENSGALG00000027133 | 162,089324 | -1,405319409 | 0,31973 | -4,3953317 | 1,11E-05 | 0,00011297 | |
| ENSGALG00000001768 | 6648,81239 | -1,406520646 | 0,15682598 | -8,9686713 | 3,00E-19 | 2,27E-17 | TENM2 |
| ENSGALG00000005123 | 593,996105 | -1,408198856 | 0,22248629 | -6,3293737 | 2,46E-10 | 6,31E-09 | WNT7A |
| ENSGALG00000000680 | 6447,75978 | -1,411570576 | 0,1125835 | -12,537988 | 4,63E-36 | 1,19E-33 | |
| ENSGALG00000009530 | 697,310671 | -1,411853584 | 0,1982451 | -7,1217577 | 1,07E-12 | 3,88E-11 | PDZRN4 |
| ENSGALG00000014343 | 783,229712 | -1,412639292 | 0,1787261 | -7,9039341 | 2,70E-15 | 1,32E-13 | |
| ENSGALG00000026557 | 168,826381 | -1,412974627 | 0,2911389 | -4,8532664 | 1,21E-06 | 1,55E-05 | PGF |
| ENSGALG00000011319 | 10478,7818 | -1,42017882 | 0,12658897 | -11,218819 | 3,30E-29 | 5,34E-27 | ADGRL3 |
| ENSGALG00000014823 | 40,5162501 | -1,422401118 | 0,34565713 | -4,1150637 | 3,87E-05 | 0,00033124 | |
| ENSGALG00000011468 | 1760,29824 | -1,424650779 | 0,18752858 | -7,5969793 | 3,03E-14 | 1,36E-12 | IGFBP5 |
| ENSGALG00000016297 | 52,4647782 | -1,427639697 | 0,35390644 | -4,0339466 | 5,48E-05 | 0,00044582 | HCRTR2 |
| ENSGALG00000019054 | 529,716619 | -1,427749976 | 0,24150691 | -5,911839 | 3,38E-09 | 7,38E-08 | |
| ENSGALG00000011110 | 296,257233 | -1,431359085 | 0,24880708 | -5,7528873 | 8,77E-09 | 1,75E-07 | |
| ENSGALG00000011207 | 13680,733 | -1,434035643 | 0,1207094 | -11,880066 | 1,50E-32 | 3,30E-30 | DUSP6 |
| ENSGALG00000012018 | 2349,85185 | -1,435600968 | 0,18691445 | -7,6805242 | 1,58E-14 | 7,29E-13 | UGT8 |
| ENSGALG00000013402 | 51,692715 | -1,446510354 | 0,35385338 | -4,0878805 | 4,35E-05 | 0,00036362 | AHRR |
| ENSGALG00000011164 | 2326,41675 | -1,446841804 | 0,20831238 | -6,9455393 | 3,77E-12 | 1,26E-10 | CHN2 |
| ENSGALG00000028821 | 11178,852 | -1,46102831 | 0,14885453 | -9,8151415 | 9,69E-23 | 9,70E-21 | GLIPR2 |
| ENSGALG00000001169 | 20569,632 | -1,469628578 | 0,14537593 | -10,10916 | 5,03E-24 | 5,85E-22 | CRB2 |
| ENSGALG00000006072 | 43,451296 | -1,471939488 | 0,35393821 | -4,1587471 | 3,20E-05 | 0,0002809 | SCN10A |
| ENSGALG00000005855 | 6149,83129 | -1,488520056 | 0,16245344 | -9,1627487 | 5,06E-20 | 4,12E-18 | |
| ENSGALG00000026262 | 5433,56566 | -1,4922838 | 0,15452733 | -9,6570863 | 4,59E-22 | 4,25E-20 | |
| ENSGALG00000001621 | 458,454887 | -1,495017381 | 0,22098058 | -6,7653791 | 1,33E-11 | 4,12E-10 | CLDN5 |
| ENSGALG00000011940 | 614,502923 | -1,496267605 | 0,21912179 | -6,8284747 | 8,58E-12 | 2,75E-10 | |
| ENSGALG00000008978 | 15989,3853 | -1,505709648 | 0,13184171 | -11,420587 | 3,30E-30 | 5,76E-28 | |
| ENSGALG00000025773 | 391,159032 | -1,506718569 | 0,21938214 | -6,8680093 | 6,51E-12 | 2,11E-10 | |
| ENSGALG00000015388 | 3557,58163 | -1,507384513 | 0,13257512 | -11,370041 | 5,90E-30 | 1,00E-27 | |
| ENSGALG00000007171 | 156,082312 | -1,518198388 | 0,29373826 | -5,1685414 | 2,36E-07 | 3,50E-06 | CIITA |
| ENSGALG00000010258 | 1087,70771 | -1,528410178 | 0,23204878 | -6,5865901 | 4,50E-11 | 1,28E-09 | LTBP2 |
| ENSGALG00000006084 | 149,653293 | -1,531133293 | 0,30821845 | -4,9676886 | 6,78E-07 | 9,10E-06 | |
| ENSGALG00000005499 | 52125,7921 | -1,531899327 | 0,15487524 | -9,8911829 | 4,55E-23 | 4,65E-21 | IL17RD |
| ENSGALG00000001383 | 88,9612133 | -1,543511015 | 0,34200221 | -4,5131609 | 6,39E-06 | 6,86E-05 | HTR4 |
| ENSGALG00000017179 | 397,23584 | -1,548130217 | 0,2756174 | -5,6169539 | 1,94E-08 | 3,65E-07 | PDGFD |
| ENSGALG00000000388 | 10384,9656 | -1,555950834 | 0,15057476 | -10,333411 | 4,98E-25 | 6,33E-23 | LIN28A |
| ENSGALG00000016293 | 1486,8847 | -1,561397543 | 0,15239221 | -10,245914 | 1,23E-24 | 1,51E-22 | BMP5 |
| ENSGALG00000014657 | 1240,68069 | -1,561923173 | 0,21313827 | -7,3282155 | 2,33E-13 | 9,26E-12 | |
| ENSGALG00000003957 | 1745,61798 | -1,565814231 | 0,14958689 | -10,46759 | 1,22E-25 | 1,66E-23 | |
| ENSGALG00000017242 | 1988,33091 | -1,579394776 | 0,13593267 | -11,618949 | 3,30E-31 | 6,42E-29 | FZD4 |
| ENSGALG00000009170 | 1955,30232 | -1,581054934 | 0,13559524 | -11,660106 | 2,04E-31 | 4,02E-29 | NCEH1 |
| ENSGALG00000003323 | 100,589751 | -1,591926379 | 0,32090266 | -4,9607765 | 7,02E-07 | 9,41E-06 | OSGIN1 |
| ENSGALG00000001645 | 733,714958 | -1,608468631 | 0,20509442 | -7,8425762 | 4,41E-15 | 2,10E-13 | KCNT1 |
| ENSGALG00000011609 | 3466,23734 | -1,61436812 | 0,3265448 | -4,9437876 | 7,66E-07 | 1,02E-05 | |
| ENSGALG00000003126 | 1209,63641 | -1,615974863 | 0,16174479 | -9,990893 | 1,67E-23 | 1,82E-21 | ERBB4 |
| ENSGALG00000019716 | 86,528621 | -1,618712644 | 0,32858792 | -4,9262694 | 8,38E-07 | 1,10E-05 | KRT20 |
| ENSGALG00000016102 | 256,085293 | -1,630912312 | 0,24814594 | -6,5723916 | 4,95E-11 | 1,41E-09 | |
| ENSGALG00000010468 | 1107,42968 | -1,6309453 | 0,15736024 | -10,364405 | 3,60E-25 | 4,67E-23 | ISM2 |
| ENSGALG00000023521 | 1102,90608 | -1,651708778 | 0,19829188 | -8,3296843 | 8,11E-17 | 4,84E-15 | ADRA2A |
| ENSGALG00000019261 | 1018,73464 | -1,655584971 | 0,22406874 | -7,3887368 | 1,48E-13 | 6,04E-12 | |
| ENSGALG00000026892 | 80,0076125 | -1,6734567 | 0,34940022 | -4,7895124 | 1,67E-06 | 2,06E-05 | |
| ENSGALG00000008353 | 2780,09542 | -1,679225856 | 0,19226914 | -8,7337255 | 2,46E-18 | 1,67E-16 | |
| ENSGALG00000006997 | 745,789097 | -1,684979264 | 0,17360387 | -9,7058852 | 2,85E-22 | 2,77E-20 | BRINP1 |
| ENSGALG00000029175 | 12384,2511 | -1,685259625 | 0,16604046 | -10,149693 | 3,32E-24 | 3,90E-22 | EFNB2 |
| ENSGALG00000008918 | 724,315184 | -1,685987772 | 0,1865069 | -9,0398142 | 1,57E-19 | 1,22E-17 | LAMP5 |

| | | | | | | | |
|---|---|---|---|---|---|---|---|
| ENSGALG00000014930 | 4308,58674 | -1,696897635 | 0,15640418 | -10,849439 | 2,01E-27 | 2,94E-25 | ENC1 |
| ENSGALG00000005498 | 1151,84629 | -1,707286081 | 0,21532932 | -7,928721 | 2,21E-15 | 1,11E-13 | |
| ENSGALG00000014235 | 549,444331 | -1,721709963 | 0,26416786 | -6,5174846 | 7,15E-11 | 1,99E-09 | |
| ENSGALG00000013102 | 1811,9189 | -1,724325032 | 0,13966812 | -12,345874 | 5,13E-35 | 1,25E-32 | COBL |
| ENSGALG00000007218 | 6209,27212 | -1,732428983 | 0,17353793 | -9,9829988 | 1,81E-23 | 1,94E-21 | MB21D2 |
| ENSGALG00000014908 | 2604,03468 | -1,732825245 | 0,16330677 | -10,61086 | 2,65E-26 | 3,76E-24 | FST |
| ENSGALG00000005476 | 213,71162 | -1,752170936 | 0,31806755 | -5,5088013 | 3,61E-08 | 6,35E-07 | PTH1R |
| ENSGALG00000011148 | 2471,49648 | -1,754606037 | 0,1499315 | -11,702718 | 1,23E-31 | 2,51E-29 | |
| ENSGALG00000027874 | 793,364535 | -1,754956403 | 0,23265502 | -7,54317 | 4,59E-14 | 1,97E-12 | CHAC1 |
| ENSGALG00000012635 | 1110,85351 | -1,755097361 | 0,16645491 | -10,543981 | 5,42E-26 | 7,45E-24 | |
| ENSGALG00000006995 | 175,827248 | -1,759714805 | 0,30239363 | -5,8192853 | 5,91E-09 | 1,22E-07 | APOD |
| ENSGALG00000027316 | 126,113792 | -1,764476803 | 0,31211141 | -5,6533557 | 1,57E-08 | 3,00E-07 | |
| ENSGALG00000005519 | 2075,75095 | -1,767947956 | 0,33351026 | -5,3010302 | 1,15E-07 | 1,82E-06 | TLL2 |
| ENSGALG00000009282 | 5108,51903 | -1,769054515 | 0,16415004 | -10,777058 | 4,42E-27 | 6,40E-25 | ELMSAN1 |
| ENSGALG00000007113 | 2619,56625 | -1,77145981 | 0,19094943 | -9,277115 | 1,74E-20 | 1,45E-18 | |
| ENSGALG00000015148 | 101,324198 | -1,772924708 | 0,34687715 | -5,1111027 | 3,20E-07 | 4,62E-06 | ANXA1 |
| ENSGALG00000023406 | 3748,56647 | -1,774894708 | 0,1767754 | -10,040394 | 1,01E-23 | 1,12E-21 | |
| ENSGALG00000004930 | 259,836288 | -1,782009807 | 0,32603449 | -5,4657095 | 4,61E-08 | 7,94E-07 | HRH1 |
| ENSGALG00000009920 | 1810,73161 | -1,80046145 | 0,20594812 | -8,742306 | 2,28E-18 | 1,56E-16 | COCH |
| ENSGALG00000014159 | 18203,792 | -1,81424576 | 0,10616787 | -17,088464 | 1,81E-65 | 1,30E-62 | PACSIN2 |
| ENSGALG00000016875 | 1624,38522 | -1,825016077 | 0,18553233 | -9,836647 | 7,83E-23 | 7,89E-21 | |
| ENSGALG00000004306 | 1596,76443 | -1,830041848 | 0,15669253 | -11,67919 | 1,63E-31 | 3,26E-29 | SYT12 |
| ENSGALG00000011708 | 13494,0054 | -1,833902608 | 0,12124784 | -15,12524 | 1,10E-51 | 6,01E-49 | MYLK |
| ENSGALG00000002714 | 18305,7311 | -1,839377527 | 0,12313583 | -14,937793 | 1,87E-50 | 9,10E-48 | TMEM132C |
| ENSGALG00000007963 | 269,370346 | -1,84239576 | 0,2523089 | -7,3021434 | 2,83E-13 | 1,11E-11 | NFATC2 |
| ENSGALG00000014935 | 24043,958 | -1,864724119 | 0,16406086 | -11,366051 | 6,17E-30 | 1,04E-27 | GREB1L |
| ENSGALG00000015089 | 1244,38805 | -1,867848339 | 0,21666451 | -8,6209242 | 6,64E-18 | 4,39E-16 | GAP43 |
| ENSGALG00000016906 | 16521,3421 | -1,870006943 | 0,10236052 | -18,268829 | 1,47E-74 | 2,00E-71 | SPRY2 |
| ENSGALG00000012392 | 124,624801 | -1,879813042 | 0,30530777 | -6,1571084 | 7,41E-10 | 1,79E-08 | IYD |
| ENSGALG00000007786 | 809,873559 | -1,895122662 | 0,25833391 | -7,3359423 | 2,20E-13 | 8,79E-12 | DOK5 |
| ENSGALG00000003437 | 453,52356 | -1,896224168 | 0,2236049 | -8,4802444 | 2,25E-17 | 1,43E-15 | |
| ENSGALG00000009693 | 162,98636 | -1,908196694 | 0,28491477 | -6,69743 | 2,12E-11 | 6,38E-10 | CPXM2 |
| ENSGALG00000026366 | 659,496965 | -1,956428809 | 0,25593066 | -7,6443705 | 2,10E-14 | 9,56E-13 | |
| ENSGALG00000016377 | 39318,7205 | -1,984049771 | 0,15160151 | -13,087269 | 3,89E-39 | 1,20E-36 | PXDN |
| ENSGALG00000007646 | 1162,43082 | -1,999886692 | 0,2341543 | -8,5408926 | 1,33E-17 | 8,72E-16 | |
| ENSGALG00000014727 | 707,495402 | -2,02826974 | 0,22579434 | -8,982819 | 2,64E-19 | 2,02E-17 | PDE4D |
| ENSGALG00000025941 | 508,274649 | -2,032822244 | 0,21082252 | -9,6423394 | 5,30E-22 | 4,84E-20 | RGS20 |
| ENSGALG00000016455 | 36148,3013 | -2,034647249 | 0,14156144 | -14,372892 | 7,66E-47 | 3,07E-44 | GREB1 |
| ENSGALG00000015332 | 1189,98649 | -2,044938557 | 0,2318737 | -8,8191915 | 1,15E-18 | 8,09E-17 | CPLX1 |
| ENSGALG00000001608 | 473,981577 | -2,081100043 | 0,22971307 | -9,059563 | 1,31E-19 | 1,03E-17 | UNC5D |
| ENSGALG00000006797 | 719,73345 | -2,093089949 | 0,18254835 | -11,465949 | 1,96E-30 | 3,60E-28 | ADARB2 |
| ENSGALG00000005347 | 2547,96798 | -2,102469906 | 0,17281317 | -12,166144 | 4,71E-34 | 1,11E-31 | ADAMTS18 |
| ENSGALG00000005061 | 4294,17572 | -2,114741441 | 0,15753482 | -13,423962 | 4,38E-41 | 1,53E-38 | PPM1E |
| ENSGALG00000006822 | 6099,69686 | -2,117548167 | 0,19086302 | -11,094596 | 1,33E-28 | 2,06E-26 | PCDH19 |
| ENSGALG00000002540 | 1690,5022 | -2,132167559 | 0,1444055 | -14,765141 | 2,46E-49 | 1,08E-46 | |
| ENSGALG00000014872 | 706,849976 | -2,137267919 | 0,21173492 | -10,094074 | 5,87E-24 | 6,77E-22 | FGF10 |
| ENSGALG00000005000 | 10982,0355 | -2,158647655 | 0,15123051 | -14,27389 | 3,18E-46 | 1,24E-43 | COMTD1 |
| ENSGALG00000002816 | 1397,74927 | -2,170634895 | 0,17254974 | -12,579763 | 2,73E-36 | 7,28E-34 | STK10 |
| ENSGALG00000024083 | 2014,38425 | -2,175833689 | 0,16411916 | -13,257646 | 4,07E-40 | 1,39E-37 | C8orf4 |
| ENSGALG00000017244 | 1712,46975 | -2,189436741 | 0,14753276 | -14,840343 | 8,03E-50 | 3,65E-47 | PRSS23 |
| ENSGALG00000012106 | 326,650164 | -2,236831516 | 0,28466041 | -7,8578946 | 3,91E-15 | 1,87E-13 | SCTR |
| ENSGALG00000005689 | 443,468063 | -2,271345118 | 0,23720344 | -9,575515 | 1,01E-21 | 9,14E-20 | Pax-2 |
| ENSGALG00000000600 | 217,113311 | -2,27272638 | 0,28275559 | -8,0377769 | 9,15E-16 | 4,85E-14 | |
| ENSGALG00000019426 | 136,015643 | -2,280006749 | 0,30033938 | -7,5914345 | 3,16E-14 | 1,41E-12 | |
| ENSGALG00000012594 | 465,978763 | -2,283939186 | 0,22484628 | -10,157781 | 3,06E-24 | 3,62E-22 | NTRK2 |
| ENSGALG00000012732 | 358,709185 | -2,298252682 | 0,22771061 | -10,092866 | 5,94E-24 | 6,80E-22 | PHACTR1 |
| ENSGALG00000004376 | 1102,6462 | -2,323750078 | 0,19821753 | -11,723232 | 9,69E-32 | 2,00E-29 | KCNJ2 |
| ENSGALG00000008118 | 2184,0618 | -2,340544818 | 0,13264091 | -17,645724 | 1,10E-69 | 1,15E-66 | |
| ENSGALG00000010435 | 1040,62618 | -2,340611141 | 0,22142593 | -10,570628 | 4,08E-26 | 5,66E-24 | RASGRP3 |
| ENSGALG00000007346 | 9971,3513 | -2,361764382 | 0,15659212 | -15,082268 | 2,12E-51 | 1,11E-48 | KIAA1462 |
| ENSGALG00000017091 | 828,298355 | -2,363903034 | 0,17074536 | -13,844611 | 1,37E-43 | 5,05E-41 | FLT1 |
| ENSGALG00000012278 | 11240,1658 | -2,383840075 | 0,16389202 | -14,545187 | 6,27E-48 | 2,67E-45 | |
| ENSGALG00000016785 | 93,1133081 | -2,405348378 | 0,32963393 | -7,2970291 | 2,94E-13 | 1,15E-11 | IL1RL1 |
| ENSGALG00000013033 | 107,445867 | -2,410715686 | 0,31956021 | -7,5438544 | 4,56E-14 | 1,97E-12 | |

| | | | | | | | |
|---|---|---|---|---|---|---|---|
| ENSGALG00000007657 | 4917,86859 | -2,412172788 | 0,33438082 | -7,2138491 | 5,44E-13 | 2,06E-11 | CDX4 |
| ENSGALG00000012927 | 231,318015 | -2,419967236 | 0,26578914 | -9,1048386 | 8,64E-20 | 6,84E-18 | |
| ENSGALG00000002183 | 431,198329 | -2,438499895 | 0,26607955 | -9,1645522 | 4,98E-20 | 4,08E-18 | PRR16 |
| ENSGALG00000015947 | 963,910906 | -2,438977585 | 0,18916847 | -12,893151 | 4,92E-38 | 1,46E-35 | |
| ENSGALG00000008893 | 301,746209 | -2,444595373 | 0,24691931 | -9,9003817 | 4,15E-23 | 4,31E-21 | |
| ENSGALG00000008890 | 114,859766 | -2,449553325 | 0,32616483 | -7,5101701 | 5,91E-14 | 2,49E-12 | ADGRL4 |
| ENSGALG00000008905 | 317,218524 | -2,456637668 | 0,23731673 | -10,351726 | 4,11E-25 | 5,28E-23 | HABP2 |
| ENSGALG00000026663 | 524,422448 | -2,478656591 | 0,20922664 | -11,846755 | 2,24E-32 | 4,76E-30 | CX3CL1 |
| ENSGALG00000012623 | 104,28468 | -2,482707417 | 0,32178291 | -7,7154733 | 1,21E-14 | 5,58E-13 | |
| ENSGALG00000015826 | 274,530912 | -2,484029553 | 0,31827956 | -7,8045526 | 5,97E-15 | 2,81E-13 | HTR1E |
| ENSGALG00000010224 | 221,988072 | -2,486198557 | 0,29117325 | -8,5385541 | 1,36E-17 | 8,83E-16 | |
| ENSGALG00000007940 | 128,052277 | -2,542363859 | 0,32047998 | -7,9329879 | 2,14E-15 | 1,07E-13 | |
| ENSGALG00000001560 | 195,37323 | -2,555527127 | 0,33829668 | -7,5541005 | 4,22E-14 | 1,83E-12 | |
| ENSGALG00000008581 | 16731,9435 | -2,566439645 | 0,14745274 | -17,405167 | 7,54E-68 | 5,70E-65 | DUSP5 |
| ENSGALG00000016784 | 1039,85153 | -2,607549769 | 0,17414233 | -14,97367 | 1,09E-50 | 5,50E-48 | |
| ENSGALG00000006112 | 988,653043 | -2,692857061 | 0,18990629 | -14,179926 | 1,22E-45 | 4,61E-43 | SCN5A |
| ENSGALG00000016154 | 140,890158 | -2,742713554 | 0,30649601 | -8,948611 | 3,60E-19 | 2,71E-17 | ARC |
| ENSGALG00000016145 | 1265,10116 | -2,754137035 | 0,17459535 | -15,774401 | 4,67E-56 | 2,76E-53 | |
| ENSGALG00000011099 | 581,916765 | -2,760215905 | 0,26996697 | -10,224273 | 1,54E-24 | 1,88E-22 | FAP |
| ENSGALG00000005209 | 4151,24313 | -2,791291062 | 0,34877784 | -8,0030631 | 1,21E-15 | 6,28E-14 | AQP1 |
| ENSGALG00000011562 | 1031,63999 | -2,817896047 | 0,18091223 | -15,576039 | 1,06E-54 | 6,01E-52 | PARK2 |
| ENSGALG00000007656 | 17151,759 | -2,823200316 | 0,30126808 | -9,3710569 | 7,18E-21 | 6,11E-19 | |
| ENSGALG00000012051 | 829,367255 | -2,878544854 | 0,24824835 | -11,595424 | 4,35E-31 | 8,22E-29 | |
| ENSGALG00000028250 | 102,796791 | -2,884651456 | 0,33080912 | -8,7199877 | 2,78E-18 | 1,88E-16 | ANKRD22 |
| ENSGALG00000007701 | 2491,57183 | -2,906313035 | 0,16540669 | -17,57071 | 4,13E-69 | 3,75E-66 | B3GNT7 |
| ENSGALG00000014984 | 5902,58959 | -2,931476354 | 0,14956936 | -19,599445 | 1,56E-85 | 3,04E-82 | F2RL1 |
| ENSGALG00000009704 | 18346,8673 | -2,934403789 | 0,16761639 | -17,506664 | 1,27E-68 | 1,02E-65 | CHST15 |
| ENSGALG00000016140 | 355,674118 | -2,998033733 | 0,24925729 | -12,027868 | 2,54E-33 | 5,76E-31 | FAM3B |
| ENSGALG00000015358 | 3246,18076 | -3,063209848 | 0,14576096 | -21,015297 | 4,75E-98 | 1,29E-94 | MYH15 |
| ENSGALG00000002562 | 2027,1289 | -3,12939254 | 0,17645164 | -17,735129 | 2,25E-70 | 2,55E-67 | ARAP3 |
| ENSGALG00000007538 | 106,892168 | -3,275953369 | 0,34614117 | -9,4642118 | 2,96E-21 | 2,60E-19 | |
| ENSGALG00000022962 | 106,156937 | -3,362304357 | 0,3346563 | -10,047037 | 9,47E-24 | 1,06E-21 | |
| ENSGALG00000002594 | 1003,24058 | -3,661637709 | 0,2050909 | -17,853731 | 2,70E-71 | 3,35E-68 | TFPI |
| ENSGALG00000013624 | 4838,43058 | -3,6799039 | 0,14627576 | -25,157305 | 1,18E-139 | 5,34E-136 | |
| ENSGALG00000006421 | 916,754331 | -3,742853665 | 0,23704674 | -15,789517 | 3,67E-56 | 2,27E-53 | SLC16A12 |
| ENSGALG00000020359 | 246,794372 | -3,821393345 | 0,28450232 | -13,431853 | 3,93E-41 | 1,41E-38 | SIX6 |
| ENSGALG00000003456 | 187,124669 | -3,934712358 | 0,32478027 | -12,114998 | 8,80E-34 | 2,03E-31 | CA12 |
| ENSGALG00000023379 | 453,271455 | -3,949811074 | 0,23138338 | -17,070418 | 2,46E-65 | 1,68E-62 | F2RL2 |
| ENSGALG00000016431 | 670,260054 | -4,37340061 | 0,23581962 | -18,545533 | 8,86E-77 | 1,34E-73 | Pax5B |
| ENSGALG00000014994 | 310,707645 | -5,043766073 | 0,28682666 | -17,584718 | 3,23E-69 | 3,14E-66 | CRHBP |
| ENSGALG00000016783 | 9121,10854 | -5,301446621 | 0,14503512 | -36,552847 | 1,61E-292 | 2,19E-288 | IL1R1 |
| ENSGALG00000004221 | 898,014845 | -5,338632777 | 0,26991661 | -19,778823 | 4,53E-87 | 1,03E-83 | IL22RA1 |

**Supplementary Table 2:** MEK1ca vs pCIG differentially regulated genes (False Discovery Rate (padj) <= 0.0001)

| ID | baseMean | log2FoldChange | lfcSE | stat | pvalue | padj | Symbol |
|---|---|---|---|---|---|---|---|
| ENSGALG00000016783 | 8238,73046 | 6,569509059 | 0,14790027 | 44,4185063 | 0 | 0 | IL1R1 |
| ENSGALG00000026862 | 13555,7342 | 4,954899999 | 0,09919193 | 49,9526499 | 0 | 0 | CLDN1 |
| ENSGALG00000013624 | 4270,69482 | 4,748606334 | 0,13766019 | 34,4951317 | 9,49E-261 | 3,96E-257 | FAM65b |
| ENSGALG00000007701 | 2111,8366 | 4,453082303 | 0,15415453 | 28,8871327 | 1,73E-183 | 5,42E-180 | B3GNT7 |
| ENSGALG00000023406 | 2791,74825 | 4,424329106 | 0,17768451 | 24,899914 | 7,46E-137 | 1,04E-133 | NKX1-2 |
| ENSGALG00000004221 | 826,015749 | 4,334100391 | 0,24063814 | 18,0108626 | 1,60E-72 | 9,10E-70 | IL22RA1 |
| ENSGALG00000026263 | 1163,05249 | 4,280447225 | 0,16395242 | 26,107863 | 2,97E-150 | 5,30E-147 | RGS8 |
| ENSGALG00000016431 | 614,504481 | 4,191515764 | 0,19877752 | 21,0864675 | 1,06E-98 | 1,02E-95 | Pax5 |
| ENSGALG00000006421 | 822,905958 | 4,186231336 | 0,20967318 | 19,9655067 | 1,10E-88 | 8,59E-86 | SLC16A12 |
| ENSGALG00000014994 | 285,756349 | 4,160137851 | 0,2463521 | 16,886959 | 5,61E-64 | 2,51E-61 | CRHBP |
| ENSGALG00000016145 | 1063,31727 | 3,924548833 | 0,20804729 | 18,8637341 | 2,27E-79 | 1,35E-76 | RIPK4 |
| ENSGALG00000016906 | 12704,3465 | 3,743251462 | 0,13733846 | 27,2556682 | 1,42E-163 | 3,56E-160 | SPRY2 |
| ENSGALG00000020359 | 223,508759 | 3,572883143 | 0,25377029 | 14,0792017 | 5,10E-45 | 1,39E-42 | SIX6 |
| ENSGALG00000023379 | 414,317659 | 3,494464883 | 0,23081766 | 15,139504 | 8,89E-52 | 3,09E-49 | F2RL2 |
| ENSGALG00000010435 | 869,92769 | 3,471549614 | 0,19291102 | 17,9956001 | 2,11E-72 | 1,15E-69 | RASGRP3 |
| ENSGALG00000007656 | 15062,1816 | 3,445291545 | 0,25219864 | 13,6610236 | 1,74E-42 | 4,43E-40 | CDX-1 |
| ENSGALG00000005519 | 1750,48505 | 3,391598976 | 0,19583928 | 17,3182775 | 3,42E-67 | 1,65E-64 | TLL2 |
| ENSGALG00000026663 | 441,393045 | 3,38430006 | 0,21308555 | 15,8823535 | 8,40E-57 | 3,18E-54 | CX3CL1 |
| ENSGALG00000002562 | 1820,09922 | 3,3452383 | 0,1690746 | 19,785576 | 3,96E-87 | 2,92E-84 | ARAP3 |
| ENSGALG00000008905 | 264,878628 | 3,342984543 | 0,2406955 | 13,8888536 | 7,40E-44 | 1,93E-41 | HABP2 |
| ENSGALG00000016784 | 893,438424 | 3,229015878 | 0,19216246 | 16,803573 | 2,30E-63 | 9,91E-61 | IL1RL2 |
| ENSGALG00000008581 | 13805,345 | 3,209755496 | 0,25216846 | 12,7286158 | 4,10E-37 | 9,50E-35 | DUSP5 |
| ENSGALG00000011099 | 516,017104 | 3,19842177 | 0,21907483 | 14,5996772 | 2,82E-48 | 8,82E-46 | FAP |
| ENSGALG00000003456 | 170,923754 | 3,130034846 | 0,27207277 | 11,5044031 | 1,25E-30 | 1,98E-28 | CA12 |
| ENSGALG00000015358 | 2958,75231 | 3,121559149 | 0,12071098 | 25,8597779 | 1,89E-147 | 2,95E-144 | MYH15 |
| ENSGALG00000011940 | 461,321244 | 2,983749675 | 0,21913664 | 13,6159322 | 3,22E-42 | 8,06E-40 | |
| ENSGALG00000007963 | 212,091292 | 2,963911322 | 0,25006993 | 11,8523298 | 2,09E-32 | 3,59E-30 | NFATC2 |
| ENSGALG00000023521 | 856,317445 | 2,942428455 | 0,19797705 | 14,8624723 | 5,78E-50 | 1,85E-47 | ADRA2A |
| ENSGALG00000007171 | 114,627355 | 2,91119837 | 0,27426227 | 10,6146514 | 2,55E-26 | 3,32E-24 | CIITA |
| ENSGALG00000005000 | 9346,89113 | 2,843605103 | 0,13435603 | 21,1646995 | 2,02E-99 | 2,11E-96 | COMTD1 |
| ENSGALG00000000600 | 182,137118 | 2,828234018 | 0,26246019 | 10,7758591 | 4,48E-27 | 6,15E-25 | |
| ENSGALG00000012018 | 1787,86628 | 2,812280515 | 0,16831135 | 16,7087992 | 1,13E-62 | 4,72E-60 | UGT8 |
| ENSGALG00000000177 | 2934,9664 | 2,765911854 | 0,13662864 | 20,2440117 | 4,01E-91 | 3,35E-88 | |
| ENSGALG00000007346 | 8739,08087 | 2,763402401 | 0,14295509 | 19,330563 | 2,97E-83 | 1,96E-80 | KIAA1462 |
| ENSGALG00000001621 | 341,359461 | 2,731716978 | 0,25022736 | 10,9169398 | 9,57E-28 | 1,38E-25 | CLDN5 |
| ENSGALG00000010224 | 190,389423 | 2,688243072 | 0,2692821 | 9,9829994 | 1,81E-23 | 2,06E-21 | |
| ENSGALG00000009704 | 16406,1404 | 2,653508455 | 0,23537258 | 11,2736518 | 1,77E-29 | 2,70E-27 | CHST15 |
| ENSGALG00000005209 | 3731,94501 | 2,631256548 | 0,27714025 | 9,49431393 | 2,22E-21 | 2,10E-19 | AQP1 |
| ENSGALG00000014984 | 5578,94086 | 2,599579307 | 0,10540138 | 24,6636185 | 2,63E-134 | 3,29E-131 | F2RL1 |
| ENSGALG00000022962 | 95,9619859 | 2,592879602 | 0,27689482 | 9,3641318 | 7,67E-21 | 7,00E-19 | |
| ENSGALG00000002183 | 379,587253 | 2,571301428 | 0,24362434 | 10,5543701 | 4,85E-26 | 6,25E-24 | PRR16 |
| ENSGALG00000015388 | 2544,04823 | 2,539489467 | 0,27173379 | 9,34550488 | 9,15E-21 | 8,29E-19 | |
| ENSGALG00000016140 | 327,169323 | 2,527534269 | 0,2417696 | 10,4543098 | 1,40E-25 | 1,79E-23 | FAM3B |
| ENSGALG00000013102 | 1492,03268 | 2,517151592 | 0,12944514 | 19,4457017 | 3,17E-84 | 2,20E-81 | COBL |
| ENSGALG00000005855 | 4852,14054 | 2,515154305 | 0,15070039 | 16,6897667 | 1,56E-62 | 6,28E-60 | |
| ENSGALG00000002816 | 1217,9424 | 2,498779812 | 0,17668612 | 14,1424792 | 2,08E-45 | 5,78E-43 | STK10 |
| ENSGALG00000026258 | 189,462612 | 2,487122645 | 0,26495328 | 9,3870233 | 6,17E-21 | 5,68E-19 | MAF |
| ENSGALG00000003126 | 980,359046 | 2,475370264 | 0,15056385 | 16,440668 | 9,78E-61 | 3,82E-58 | ERBB4 |
| ENSGALG00000002540 | 1485,05869 | 2,44573463 | 0,13676571 | 17,8826589 | 1,61E-71 | 8,39E-69 | |
| ENSGALG00000015826 | 238,755372 | 2,399148661 | 0,27605492 | 8,69083812 | 3,60E-18 | 2,53E-16 | HTR1E |
| ENSGALG00000007538 | 97,0573797 | 2,386772965 | 0,27690143 | 8,61957623 | 6,72E-18 | 4,64E-16 | |
| ENSGALG00000019054 | 408,32875 | 2,38528849 | 0,2379735 | 10,0233366 | 1,20E-23 | 1,38E-21 | |
| ENSGALG00000026366 | 561,445489 | 2,348234721 | 0,23749642 | 9,88745309 | 4,72E-23 | 5,13E-21 | |
| ENSGALG00000004306 | 1355,5371 | 2,335846949 | 0,15683954 | 14,893227 | 3,65E-50 | 1,20E-47 | SYT12 |
| ENSGALG00000009393 | 548,007386 | 2,335695464 | 0,18992639 | 12,2978985 | 9,30E-35 | 1,85E-32 | |
| ENSGALG00000007657 | 4493,01812 | 2,332413704 | 0,26632361 | 8,75781796 | 1,99E-18 | 1,44E-16 | CDX4 |
| ENSGALG00000017091 | 755,513643 | 2,329587636 | 0,16022975 | 14,5390453 | 6,85E-48 | 2,09E-45 | FLT1 |
| ENSGALG00000011562 | 980,279996 | 2,318757284 | 0,17351326 | 13,3635736 | 9,87E-41 | 2,42E-38 | PARK2 |
| ENSGALG00000012051 | 782,419586 | 2,272220282 | 0,23077218 | 9,84616192 | 7,12E-23 | 7,61E-21 | |
| ENSGALG00000011207 | 11039,4244 | 2,267210819 | 0,10235954 | 22,1494829 | 1,06E-108 | 1,20E-105 | DUSP6 |

| | | | | | | | |
|---|---|---|---|---|---|---|---|
| ENSGALG00000010036 | 233,218083 | 2,253052375 | 0,24656449 | 9,13778123 | 6,37E-20 | 5,35E-18 | FOSL2 |
| ENSGALG00000000949 | 209,314356 | 2,249487158 | 0,24647672 | 9,12657061 | 7,07E-20 | 5,90E-18 | HBEGF |
| ENSGALG00000004376 | 1011,34575 | 2,243095428 | 0,17712966 | 12,6635789 | 9,41E-37 | 2,14E-34 | KCNJ2 |
| ENSGALG00000007543 | 311,281076 | 2,235076903 | 0,23707019 | 9,42791198 | 4,18E-21 | 3,91E-19 | APCDD1L |
| ENSGALG00000014159 | 15753,2456 | 2,234266769 | 0,08275046 | 27,0000523 | 1,48E-160 | 3,08E-157 | PACSIN2 |
| ENSGALG00000016086 | 151,68942 | 2,227609215 | 0,26302517 | 8,46918638 | 2,47E-17 | 1,63E-15 | EGR4 |
| ENSGALG00000017244 | 1556,01813 | 2,226234853 | 0,13126312 | 16,9600944 | 1,62E-64 | 7,51E-62 | PRSS23 |
| ENSGALG00000007786 | 691,837015 | 2,190066734 | 0,24318575 | 9,00573619 | 2,14E-19 | 1,66E-17 | DOK5 |
| ENSGALG00000017242 | 1658,38116 | 2,189728881 | 0,12383488 | 17,6826506 | 5,70E-70 | 2,85E-67 | FZD4 |
| ENSGALG00000005061 | 3913,87544 | 2,1754071 | 0,11345305 | 19,1745135 | 6,04E-82 | 3,78E-79 | PPM1E |
| ENSGALG00000005176 | 768,829888 | 2,156988496 | 0,17567574 | 12,2782377 | 1,19E-34 | 2,32E-32 | |
| ENSGALG00000003437 | 391,254589 | 2,148196892 | 0,22078047 | 9,73001328 | 2,25E-22 | 2,28E-20 | |
| ENSGALG00000027275 | 448,763372 | 2,082192136 | 0,23055734 | 9,03112478 | 1,70E-19 | 1,34E-17 | |
| ENSGALG00000005499 | 41654,758 | 2,068323361 | 0,23020452 | 8,98472101 | 2,59E-19 | 1,99E-17 | IL17RD |
| ENSGALG00000006995 | 146,518321 | 2,052973523 | 0,26905137 | 7,63041466 | 2,34E-14 | 1,21E-12 | APOD |
| ENSGALG00000004594 | 723,032621 | 2,028493881 | 0,21261457 | 9,54070945 | 1,42E-21 | 1,37E-19 | CD74 |
| ENSGALG00000006112 | 965,473662 | 2,027013529 | 0,17450249 | 11,6159577 | 3,42E-31 | 5,48E-29 | SCN5A |
| ENSGALG00000025941 | 463,056002 | 1,970690155 | 0,19459556 | 10,1271078 | 4,19E-24 | 4,99E-22 | RGS20 |
| ENSGALG00000012732 | 335,645133 | 1,970234077 | 0,21232377 | 9,2793854 | 1,70E-20 | 1,49E-18 | PHACTR1 |
| ENSGALG00000017179 | 331,83407 | 1,969058193 | 0,24335518 | 8,09129369 | 5,90E-16 | 3,53E-14 | PDGFD |
| ENSGALG00000007940 | 119,525715 | 1,939184933 | 0,27051161 | 7,16858298 | 7,58E-13 | 3,26E-11 | |
| ENSGALG00000011833 | 10182,5258 | 1,93191024 | 0,0952049 | 20,2921299 | 1,51E-91 | 1,35E-88 | |
| ENSGALG00000003323 | 80,456143 | 1,918109692 | 0,2769549 | 6,92571127 | 4,34E-12 | 1,69E-10 | OSGIN1 |
| ENSGALG00000012635 | 985,437941 | 1,911001153 | 0,1578461 | 12,1067365 | 9,73E-34 | 1,79E-31 | |
| ENSGALG00000022490 | 573,744074 | 1,907440028 | 0,20504083 | 9,30273285 | 1,37E-20 | 1,21E-18 | |
| ENSGALG00000026154 | 1182,06867 | 1,903399885 | 0,15185043 | 12,534702 | 4,82E-36 | 1,02E-33 | CARD19 |
| ENSGALG00000008461 | 127,705778 | 1,900360149 | 0,2708458 | 7,0163913 | 2,28E-12 | 9,25E-11 | |
| ENSGALG00000007415 | 201,020232 | 1,899486225 | 0,25905889 | 7,33225653 | 2,26E-13 | 1,04E-11 | |
| ENSGALG00000000544 | 59,7220256 | 1,895252672 | 0,27550462 | 6,87920476 | 6,02E-12 | 2,30E-10 | RAB44 |
| ENSGALG00000016785 | 81,8603384 | 1,892879929 | 0,27682278 | 6,83787639 | 8,04E-12 | 3,02E-10 | IL1RL1 |
| ENSGALG00000006447 | 1443,0601 | 1,885131113 | 0,20851752 | 9,04063661 | 1,56E-19 | 1,24E-17 | |
| ENSGALG00000005498 | 1010,21765 | 1,873960213 | 0,20189609 | 9,28180559 | 1,67E-20 | 1,47E-18 | |
| ENSGALG00000008353 | 2472,76845 | 1,848707427 | 0,15762664 | 11,7283947 | 9,12E-32 | 1,50E-29 | |
| ENSGALG00000010005 | 472,644462 | 1,841664237 | 0,22183955 | 8,30178484 | 1,03E-16 | 6,45E-15 | EPAS1 |
| ENSGALG00000010597 | 1857,71605 | 1,8345583 | 0,17099128 | 10,728958 | 7,44E-27 | 1,01E-24 | |
| ENSGALG00000012106 | 297,149921 | 1,828065766 | 0,26618963 | 6,86753188 | 6,53E-12 | 2,48E-10 | SCTR |
| ENSGALG00000000141 | 377,173549 | 1,818659764 | 0,24878968 | 7,31002901 | 2,67E-13 | 1,21E-11 | BLB1 |
| ENSGALG00000026262 | 4692,72087 | 1,788029436 | 0,15007014 | 11,9146249 | 9,93E-33 | 1,73E-30 | |
| ENSGALG00000010468 | 963,638886 | 1,7857319 | 0,19155321 | 9,32238057 | 1,14E-20 | 1,02E-18 | ISM2 |
| ENSGALG00000027365 | 919,719868 | 1,784528584 | 0,14750129 | 12,0983934 | 1,08E-33 | 1,95E-31 | HIVEP3 |
| ENSGALG00000003706 | 2459,1448 | 1,784490953 | 0,12794473 | 13,9473579 | 3,26E-44 | 8,69E-42 | |
| ENSGALG00000011581 | 7771,72761 | 1,763989645 | 0,27657569 | 6,37796348 | 1,79E-10 | 5,80E-09 | KIF26A |
| ENSGALG00000023241 | 912,035255 | 1,749836088 | 0,17358051 | 10,080833 | 6,72E-24 | 7,85E-22 | |
| ENSGALG00000027521 | 220,81723 | 1,741041239 | 0,24159087 | 7,20656876 | 5,74E-13 | 2,50E-11 | |
| ENSGALG00000012623 | 98,1782216 | 1,738546704 | 0,2738672 | 6,3481377 | 2,18E-10 | 6,97E-09 | |
| ENSGALG00000029175 | 11184,2893 | 1,731471147 | 0,15062573 | 11,4951888 | 1,39E-30 | 2,18E-28 | EFNB2 |
| ENSGALG00000027331 | 92,8137096 | 1,731139271 | 0,27587874 | 6,27500053 | 3,50E-10 | 1,10E-08 | |
| ENSGALG00000001560 | 177,821121 | 1,726930021 | 0,27637082 | 6,24859752 | 4,14E-10 | 1,27E-08 | |
| ENSGALG00000002203 | 108,392731 | 1,726038051 | 0,27079159 | 6,37404609 | 1,84E-10 | 5,92E-09 | FGF-18 |
| ENSGALG00000024083 | 1953,45291 | 1,719739442 | 0,15521106 | 11,0800061 | 1,57E-28 | 2,31E-26 | C8orf4 |
| ENSGALG00000005069 | 457,203952 | 1,702946767 | 0,18881513 | 9,01912232 | 1,90E-19 | 1,48E-17 | |
| ENSGALG00000005689 | 438,331804 | 1,702270541 | 0,18963661 | 8,97648701 | 2,80E-19 | 2,13E-17 | Pax-2 |
| ENSGALG00000009693 | 145,589702 | 1,692012083 | 0,26448256 | 6,39744289 | 1,58E-10 | 5,13E-09 | CPXM2 |
| ENSGALG00000015332 | 921,622061 | 1,689640626 | 0,27053693 | 6,2455083 | 4,22E-10 | 1,29E-08 | CPLX1 |
| ENSGALG00000006084 | 123,749559 | 1,684603389 | 0,27421003 | 6,14347832 | 8,07E-10 | 2,35E-08 | |
| ENSGALG00000028250 | 102,045127 | 1,684255106 | 0,27227076 | 6,18595672 | 6,17E-10 | 1,83E-08 | ANKRD22 |
| ENSGALG00000010258 | 947,034185 | 1,674285408 | 0,222721 | 7,5174116 | 5,59E-14 | 2,75E-12 | LTBP2 |
| ENSGALG00000003488 | 483,411427 | 1,668824259 | 0,21466802 | 7,77397695 | 7,61E-15 | 4,14E-13 | |
| ENSGALG00000008281 | 275,533181 | 1,659542176 | 0,23132804 | 7,1739776 | 7,28E-13 | 3,14E-11 | NEURL1 |
| ENSGALG00000005263 | 1264,63713 | 1,654921404 | 0,1861074 | 8,89229227 | 5,99E-19 | 4,46E-17 | SOX8 |
| ENSGALG00000016500 | 476,307952 | 1,652658269 | 0,1912173 | 8,64282823 | 5,48E-18 | 3,81E-16 | FKBP12.6 |
| ENSGALG00000013033 | 98,9033207 | 1,649436663 | 0,27685165 | 5,95783573 | 2,56E-09 | 6,80E-08 | |
| ENSGALG00000009740 | 628,788388 | 1,648366239 | 0,18242751 | 9,03573286 | 1,63E-19 | 1,29E-17 | RASGRP1 |
| ENSGALG00000011076 | 175,435833 | 1,644995721 | 0,24871447 | 6,61399281 | 3,74E-11 | 1,29E-09 | SYNE3 |

| | | | | | | | |
|---|---|---|---|---|---|---|---|
| ENSGALG00000010910 | 2761,33893 | 1,643397512 | 0,13038008 | 12,6046669 | 1,99E-36 | 4,45E-34 | |
| ENSGALG00000001188 | 5705,77087 | 1,635057494 | 0,1139957 | 14,3431509 | 1,18E-46 | 3,42E-44 | |
| ENSGALG00000016602 | 1430,32585 | 1,632391617 | 0,15056739 | 10,8416017 | 2,19E-27 | 3,04E-25 | |
| ENSGALG00000026989 | 11486,0599 | 1,621656649 | 0,10265821 | 15,7966585 | 3,28E-56 | 1,21E-53 | HOXA7 |
| ENSGALG00000016377 | 35584,2442 | 1,612690665 | 0,24226856 | 6,65662393 | 2,80E-11 | 9,79E-10 | PXDN |
| ENSGALG00000027218 | 62,1190209 | 1,609222298 | 0,27648022 | 5,82038856 | 5,87E-09 | 1,49E-07 | |
| ENSGALG00000011058 | 222,479144 | 1,607120374 | 0,23535625 | 6,82845848 | 8,58E-12 | 3,20E-10 | LEPR |
| ENSGALG00000008118 | 2195,31441 | 1,603231134 | 0,14691442 | 10,9126874 | 1,00E-27 | 1,43E-25 | |
| ENSGALG00000009282 | 4772,45996 | 1,600131022 | 0,1549655 | 10,3257244 | 5,39E-25 | 6,61E-23 | ELMSAN1 |
| ENSGALG00000027707 | 84,429158 | 1,597314136 | 0,27648459 | 5,77722663 | 7,59E-09 | 1,89E-07 | ESM1 |
| ENSGALG00000028281 | 147,102379 | 1,585304805 | 0,26452499 | 5,99302471 | 2,06E-09 | 5,54E-08 | |
| ENSGALG00000005347 | 2499,3111 | 1,58404156 | 0,16400145 | 9,65870473 | 4,52E-22 | 4,56E-20 | ADAMTS18 |
| ENSGALG00000001383 | 75,1252537 | 1,582310757 | 0,27707894 | 5,71068578 | 1,13E-08 | 2,76E-07 | HTR4 |
| ENSGALG00000026484 | 105,284801 | 1,578601549 | 0,27180452 | 5,80785612 | 6,33E-09 | 1,60E-07 | |
| ENSGALG00000008890 | 110,363216 | 1,553993643 | 0,27546286 | 5,64139075 | 1,69E-08 | 4,04E-07 | ADGRL4 |
| ENSGALG00000017714 | 531,051528 | 1,553191397 | 0,22067465 | 7,038377 | 1,94E-12 | 8,03E-11 | |
| ENSGALG00000011608 | 1442,66872 | 1,548805995 | 0,27671663 | 5,59708325 | 2,18E-08 | 5,17E-07 | INF2 |
| ENSGALG00000011148 | 2343,39868 | 1,54566024 | 0,12322461 | 12,5434379 | 4,32E-36 | 9,31E-34 | |
| ENSGALG00000004730 | 427,286464 | 1,54214284 | 0,20259329 | 7,61201356 | 2,70E-14 | 1,38E-12 | |
| ENSGALG00000011962 | 5021,1701 | 1,540068054 | 0,12375377 | 12,4446156 | 1,50E-35 | 3,12E-33 | |
| ENSGALG00000008875 | 1473,68406 | 1,539400497 | 0,15531993 | 9,91115904 | 3,72E-23 | 4,09E-21 | PLCB1 |
| ENSGALG00000001189 | 175,140381 | 1,519846307 | 0,24672052 | 6,1601942 | 7,27E-10 | 2,13E-08 | NCF1 |
| ENSGALG00000012847 | 213,214453 | 1,514705906 | 0,24982981 | 6,06295113 | 1,34E-09 | 3,74E-08 | |
| ENSGALG00000016754 | 1417,60122 | 1,508945346 | 0,15769298 | 9,56888107 | 1,08E-21 | 1,06E-19 | |
| ENSGALG00000006997 | 697,291921 | 1,507910301 | 0,16647993 | 9,05761005 | 1,33E-19 | 1,08E-17 | BRINP1 |
| ENSGALG00000019426 | 134,849102 | 1,491625247 | 0,25823534 | 5,77622423 | 7,64E-09 | 1,90E-07 | |
| ENSGALG00000016348 | 4059,56984 | 1,488129704 | 0,14344251 | 10,3743978 | 3,24E-25 | 4,06E-23 | SAT1 |
| ENSGALG00000011236 | 148,702178 | 1,484958654 | 0,273941 | 5,42072431 | 5,94E-08 | 1,31E-06 | BEGAIN |
| ENSGALG00000002594 | 917,561594 | 1,484249601 | 0,27345121 | 5,42784068 | 5,70E-08 | 1,27E-06 | TFPI |
| ENSGALG00000023751 | 646,755681 | 1,484039491 | 0,2418537 | 6,13610416 | 8,46E-10 | 2,45E-08 | |
| ENSGALG00000014930 | 4123,64273 | 1,479606742 | 0,09900477 | 14,9448018 | 1,68E-50 | 5,69E-48 | ENC1 |
| ENSGALG00000010683 | 477,522338 | 1,466981653 | 0,20015393 | 7,32926728 | 2,31E-13 | 1,06E-11 | |
| ENSGALG00000026795 | 85,4265267 | 1,459031178 | 0,27670179 | 5,27293727 | 1,34E-07 | 2,78E-06 | |
| ENSGALG00000007660 | 3126,72301 | 1,456310727 | 0,1563383 | 9,31512467 | 1,22E-20 | 1,09E-18 | |
| ENSGALG00000026251 | 159,900115 | 1,452153197 | 0,27592059 | 5,26293896 | 1,42E-07 | 2,93E-06 | |
| ENSGALG00000028130 | 618,2391 | 1,44989129 | 0,1712474 | 8,46664711 | 2,53E-17 | 1,65E-15 | |
| ENSGALG00000015865 | 3211,05522 | 1,445378264 | 0,11980727 | 12,0641947 | 1,63E-33 | 2,92E-31 | |
| ENSGALG00000009667 | 556,468113 | 1,442248321 | 0,24008414 | 6,00726204 | 1,89E-09 | 5,13E-08 | MFAP3L |
| ENSGALG00000005442 | 233,433914 | 1,437319869 | 0,25444764 | 5,64878444 | 1,62E-08 | 3,89E-07 | PALMD |
| ENSGALG00000007771 | 2433,98465 | 1,43708626 | 0,1405035 | 10,2281172 | 1,48E-24 | 1,78E-22 | FAM210B |
| ENSGALG00000026790 | 326,80396 | 1,433302155 | 0,24589155 | 5,82900134 | 5,58E-09 | 1,42E-07 | DMRT2 |
| ENSGALG00000007530 | 576,29845 | 1,423961501 | 0,20873713 | 6,82179313 | 8,99E-12 | 3,33E-10 | |
| ENSGALG00000027210 | 241,661292 | 1,422887936 | 0,22890179 | 6,2161502 | 5,10E-10 | 1,54E-08 | |
| ENSGALG00000008914 | 65,3744448 | 1,411144988 | 0,27696481 | 5,09503356 | 3,49E-07 | 6,83E-06 | NRAP |
| ENSGALG00000007218 | 5981,23565 | 1,41095016 | 0,1549608 | 9,10520721 | 8,61E-20 | 7,09E-18 | MB21D2 |
| ENSGALG00000012702 | 12990,3378 | 1,407925441 | 0,12617763 | 11,1582809 | 6,52E-29 | 9,72E-27 | JARID2 |
| ENSGALG00000009546 | 1312,83637 | 1,402581457 | 0,15185477 | 9,23633455 | 2,55E-20 | 2,20E-18 | |
| ENSGALG00000006314 | 830,004543 | 1,402020858 | 0,15483246 | 9,05508371 | 1,36E-19 | 1,09E-17 | |
| ENSGALG00000020719 | 239,712039 | 1,399243921 | 0,25114775 | 5,57139732 | 2,53E-08 | 5,93E-07 | |
| ENSGALG00000011468 | 1596,02721 | 1,39801296 | 0,18393319 | 7,60065615 | 2,95E-14 | 1,49E-12 | IGFBP5 |
| ENSGALG00000015665 | 918,715368 | 1,394518864 | 0,15230743 | 9,15594797 | 5,39E-20 | 4,55E-18 | SH3BP2 |
| ENSGALG00000011680 | 2741,59968 | 1,389021273 | 0,17555553 | 7,91214743 | 2,53E-15 | 1,43E-13 | EPHA5 |
| ENSGALG00000010470 | 502,578729 | 1,388958129 | 0,18969021 | 7,32224451 | 2,44E-13 | 1,11E-11 | TCERG1L |
| ENSGALG00000011419 | 2270,32976 | 1,382981881 | 0,11393773 | 12,1380504 | 6,64E-34 | 1,26E-31 | DUSP4 |
| ENSGALG00000014852 | 13662,8714 | 1,380781721 | 0,0883531 | 15,6279945 | 4,69E-55 | 1,68E-52 | |
| ENSGALG00000010049 | 550,482195 | 1,375590195 | 0,23963948 | 5,74024862 | 9,45E-09 | 2,33E-07 | PTPRB |
| ENSGALG00000006401 | 3753,95769 | 1,372021166 | 0,13289053 | 10,3244463 | 5,46E-25 | 6,64E-23 | OSBPL5 |
| ENSGALG00000002326 | 668,309268 | 1,369220123 | 0,17190144 | 7,96514611 | 1,65E-15 | 9,51E-14 | |
| ENSGALG00000007846 | 625,227196 | 1,3613354 | 0,26060484 | 5,22375324 | 1,75E-07 | 3,55E-06 | |
| ENSGALG00000011120 | 395,765388 | 1,356774191 | 0,22365321 | 6,06641944 | 1,31E-09 | 3,68E-08 | |
| ENSGALG00000004209 | 184,595238 | 1,355480435 | 0,24429813 | 5,54846835 | 2,88E-08 | 6,70E-07 | ELFN1 |
| ENSGALG00000014516 | 1387,85858 | 1,351178171 | 0,18357479 | 7,36036878 | 1,83E-13 | 8,47E-12 | CPEB2 |
| ENSGALG00000004569 | 1782,95199 | 1,348974691 | 0,16732365 | 8,06206833 | 7,50E-16 | 4,45E-14 | UNC5B |
| ENSGALG00000012911 | 336,285764 | 1,345449733 | 0,24276482 | 5,54219392 | 2,99E-08 | 6,93E-07 | SYT10 |

| | | | | | | | |
|---|---|---|---|---|---|---|---|
| ENSGALG00000015028 | 333,710492 | 1,342064123 | 0,2166346 | 6,19505895 | 5,83E-10 | 1,74E-08 | |
| ENSGALG00000005476 | 194,019797 | 1,338834069 | 0,27604471 | 4,85006238 | 1,23E-06 | 2,21E-05 | PTH1R |
| ENSGALG00000000107 | 64,6686363 | 1,334266505 | 0,27712884 | 4,81460712 | 1,47E-06 | 2,60E-05 | TRIM7 |
| ENSGALG00000006258 | 7855,034 | 1,330333906 | 0,11339109 | 11,7322618 | 8,71E-32 | 1,45E-29 | SPOCK1 |
| ENSGALG00000015089 | 1227,41331 | 1,327665029 | 0,20978484 | 6,32869871 | 2,47E-10 | 7,85E-09 | GAP43 |
| ENSGALG00000027764 | 201,777427 | 1,324450715 | 0,26757908 | 4,94975438 | 7,43E-07 | 1,38E-05 | |
| ENSGALG00000011110 | 269,499781 | 1,322132253 | 0,23248021 | 5,6870744 | 1,29E-08 | 3,16E-07 | |
| ENSGALG00000008893 | 314,010022 | 1,318238187 | 0,24175367 | 5,45281554 | 4,96E-08 | 1,12E-06 | |
| ENSGALG00000011220 | 1013,85256 | 1,316265785 | 0,15106079 | 8,7134842 | 2,95E-18 | 2,11E-16 | SLC25A29 |
| ENSGALG00000005805 | 3857,09426 | 1,313014625 | 0,12265981 | 10,7045222 | 9,69E-27 | 1,29E-24 | PLCD1 |
| ENSGALG00000022909 | 468,481699 | 1,309497831 | 0,23797027 | 5,50277912 | 3,74E-08 | 8,55E-07 | PTPN3 |
| ENSGALG00000007140 | 2510,20659 | 1,309470163 | 0,26722383 | 4,90027474 | 9,57E-07 | 1,75E-05 | NRP1 |
| ENSGALG00000023347 | 339,64918 | 1,299000587 | 0,20361197 | 6,37978494 | 1,77E-10 | 5,75E-09 | PTCHD3 |
| ENSGALG00000010311 | 349,190305 | 1,29508602 | 0,2066853 | 6,26598025 | 3,70E-10 | 1,15E-08 | NAV3 |
| ENSGALG00000004118 | 1210,82766 | 1,294260658 | 0,15836962 | 8,17240512 | 3,02E-16 | 1,85E-14 | |
| ENSGALG00000002993 | 1805,52188 | 1,293663972 | 0,14239974 | 9,0847354 | 1,04E-19 | 8,44E-18 | CA10 |
| ENSGALG00000009808 | 160,258227 | 1,293465516 | 0,26116788 | 4,952621 | 7,32E-07 | 1,36E-05 | |
| ENSGALG00000015256 | 9263,94872 | 1,284509548 | 0,10864322 | 11,8231908 | 2,96E-32 | 5,01E-30 | PAM |
| ENSGALG00000006729 | 4860,44055 | 1,281458961 | 0,27186573 | 4,713573 | 2,43E-06 | 4,09E-05 | |
| ENSGALG00000015908 | 893,550112 | 1,276423655 | 0,19668679 | 6,48962586 | 8,60E-11 | 2,85E-09 | COL12A1 |
| ENSGALG00000014727 | 724,352277 | 1,275557809 | 0,20623272 | 6,18504092 | 6,21E-10 | 1,84E-08 | PDE4D |
| ENSGALG00000008263 | 465,678921 | 1,273175208 | 0,20291488 | 6,27443008 | 3,51E-10 | 1,10E-08 | CNTN4 |
| ENSGALG00000016817 | 5327,13148 | 1,2720896 | 0,0882221 | 14,4191717 | 3,92E-47 | 1,17E-44 | RASA3 |
| ENSGALG00000007253 | 507,277126 | 1,27114813 | 0,17906175 | 7,09893728 | 1,26E-12 | 5,28E-11 | IL1RAP |
| ENSGALG00000009920 | 1787,73878 | 1,271106358 | 0,20301633 | 6,26110393 | 3,82E-10 | 1,18E-08 | COCH |
| ENSGALG00000002632 | 1174,8881 | 1,269452785 | 0,16354378 | 7,76215854 | 8,35E-15 | 4,52E-13 | CALCRL |
| ENSGALG00000016102 | 216,026386 | 1,263113327 | 0,27712691 | 4,55788769 | 5,17E-06 | 8,21E-05 | |
| ENSGALG00000001608 | 486,701914 | 1,262511696 | 0,21788918 | 5,79428346 | 6,86E-09 | 1,72E-07 | UNC5D |
| ENSGALG00000016998 | 394,580006 | 1,256730923 | 0,21467796 | 5,85402864 | 4,80E-09 | 1,23E-07 | LPAR6 |
| ENSGALG00000013503 | 631,301741 | 1,248703074 | 0,19424591 | 6,42846519 | 1,29E-10 | 4,21E-09 | FAM149A |
| ENSGALG00000011831 | 6588,41321 | 1,24831638 | 0,0954127 | 13,0833358 | 4,10E-39 | 9,86E-37 | ANKRD50 |
| ENSGALG00000000054 | 9908,86346 | 1,246870327 | 0,12340391 | 10,1039772 | 5,30E-24 | 6,26E-22 | |
| ENSGALG00000028047 | 547,475566 | 1,243898501 | 0,24576817 | 5,06126779 | 4,16E-07 | 8,03E-06 | RHOV |
| ENSGALG00000010010 | 1045,78332 | 1,237630354 | 0,14486806 | 8,54315544 | 1,31E-17 | 8,74E-16 | RHOQ |
| ENSGALG00000009947 | 4438,8697 | 1,236965236 | 0,09832671 | 12,5801542 | 2,71E-36 | 5,96E-34 | PLEKHH2 |
| ENSGALG00000014983 | 1347,23546 | 1,23637443 | 0,14656523 | 8,43565994 | 3,29E-17 | 2,12E-15 | F2R |
| ENSGALG00000005619 | 769,181048 | 1,235271948 | 0,20759912 | 5,95027545 | 2,68E-09 | 7,08E-08 | |
| ENSGALG00000015653 | 570,869024 | 1,234972794 | 0,196909 | 6,27179447 | 3,57E-10 | 1,11E-08 | |
| ENSGALG00000003114 | 396,293229 | 1,234804889 | 0,20823968 | 5,92972908 | 3,03E-09 | 7,97E-08 | NOG |
| ENSGALG00000019716 | 79,9556086 | 1,234170903 | 0,27584604 | 4,47412944 | 7,67E-06 | 0,00011733 | KRT20 |
| ENSGALG00000019261 | 997,670343 | 1,230099163 | 0,20297117 | 6,06046252 | 1,36E-09 | 3,79E-08 | |
| ENSGALG00000028821 | 10673,5529 | 1,225133338 | 0,13787775 | 8,88564911 | 6,35E-19 | 4,70E-17 | GLIPR2 |
| ENSGALG00000013902 | 978,003089 | 1,221242084 | 0,22061929 | 5,53551816 | 3,10E-08 | 7,19E-07 | |
| ENSGALG00000008999 | 1657,80989 | 1,220108565 | 0,15349781 | 7,94870323 | 1,88E-15 | 1,07E-13 | AFAP1L2 |
| ENSGALG00000014823 | 35,5546103 | 1,217853522 | 0,25543674 | 4,76773037 | 1,86E-06 | 3,21E-05 | |
| ENSGALG00000000307 | 7560,3549 | 1,215832976 | 0,11045819 | 11,0071785 | 3,53E-28 | 5,13E-26 | |
| ENSGALG00000007268 | 3351,96385 | 1,214609385 | 0,1221963 | 9,93982107 | 2,79E-23 | 3,09E-21 | |
| ENSGALG00000000721 | 3131,94473 | 1,214440274 | 0,12625936 | 9,61861552 | 6,67E-22 | 6,57E-20 | SLC41A1 |
| ENSGALG00000011061 | 859,770333 | 1,212027759 | 0,15686437 | 7,72659688 | 1,10E-14 | 5,93E-13 | |
| ENSGALG00000007669 | 6031,34854 | 1,205315602 | 0,27706455 | 4,35030608 | 1,36E-05 | 0,00019547 | EGR1 |
| ENSGALG00000026557 | 144,536518 | 1,203253046 | 0,27652705 | 4,35130324 | 1,35E-05 | 0,00019503 | PGF |
| ENSGALG00000002555 | 1133,81707 | 1,202502668 | 0,20625382 | 5,83020812 | 5,54E-09 | 1,41E-07 | RET |
| ENSGALG00000027133 | 142,021318 | 1,199491369 | 0,27660527 | 4,33647334 | 1,45E-05 | 0,00020682 | |
| ENSGALG00000024035 | 1486,34939 | 1,198706359 | 0,15977826 | 7,50231181 | 6,27E-14 | 3,08E-12 | |
| ENSGALG00000027070 | 3188,10078 | 1,19854946 | 0,13384164 | 8,95498217 | 3,40E-19 | 2,58E-17 | TIMP2 |
| ENSGALG00000001160 | 934,02292 | 1,198076321 | 0,1603006 | 7,47393516 | 7,78E-14 | 3,73E-12 | |
| ENSGALG00000009360 | 505,831372 | 1,196193521 | 0,18849047 | 6,34617507 | 2,21E-10 | 7,04E-09 | CAPN2 |
| ENSGALG00000014080 | 86,7875432 | 1,191385609 | 0,27640604 | 4,31027336 | 1,63E-05 | 0,00022917 | |
| ENSGALG00000028785 | 165,798859 | 1,191015745 | 0,26472615 | 4,49904838 | 6,83E-06 | 0,0001062 | |
| ENSGALG00000014864 | 693,502451 | 1,190480858 | 0,15988047 | 7,44606817 | 9,62E-14 | 4,59E-12 | |
| ENSGALG00000012927 | 251,354318 | 1,187339616 | 0,23405659 | 5,07287416 | 3,92E-07 | 7,58E-06 | |
| ENSGALG00000007809 | 1064,24338 | 1,186972522 | 0,17157276 | 6,91818756 | 4,57E-12 | 1,77E-10 | TSHZ2 |
| ENSGALG00000016455 | 38193,5361 | 1,186649639 | 0,12872943 | 9,2181693 | 3,02E-20 | 2,59E-18 | GREB1 |
| ENSGALG00000013149 | 115,338492 | 1,183860135 | 0,26574609 | 4,45485435 | 8,40E-06 | 0,00012698 | MOCOS |

| | | | | | | | |
|---|---|---|---|---|---|---|---|
| ENSGALG00000012407 | 252,485775 | 1,181422763 | 0,24199801 | 4,8819524 | 1,05E-06 | 1,90E-05 | LRP1B |
| ENSGALG00000004365 | 588,746185 | 1,179727636 | 0,2121902 | 5,55976489 | 2,70E-08 | 6,32E-07 | WDR66 |
| ENSGALG00000011867 | 1901,86124 | 1,179474298 | 0,13477658 | 8,75132991 | 2,11E-18 | 1,52E-16 | SERHL2 |
| ENSGALG00000008369 | 1041,31301 | 1,177660239 | 0,16764215 | 7,02484557 | 2,14E-12 | 8,79E-11 | ABCC5 |
| ENSGALG00000002294 | 2609,13169 | 1,174006641 | 0,10962142 | 10,7096464 | 9,17E-27 | 1,23E-24 | SEMA6A |
| ENSGALG00000002543 | 4971,15753 | 1,169335598 | 0,11955874 | 9,78042788 | 1,37E-22 | 1,43E-20 | CSGALNACT2 |
| ENSGALG00000007608 | 2014,25657 | 1,169181539 | 0,16816744 | 6,95248465 | 3,59E-12 | 1,41E-10 | ARHGEF9 |
| ENSGALG00000013892 | 1213,76955 | 1,167370699 | 0,13409168 | 8,70576514 | 3,15E-18 | 2,24E-16 | MAP3K5 |
| ENSGALG00000011439 | 1116,38335 | 1,165074777 | 0,15459091 | 7,53650268 | 4,83E-14 | 2,41E-12 | |
| ENSGALG00000009450 | 2113,57054 | 1,162765074 | 0,12246309 | 9,49482074 | 2,21E-21 | 2,10E-19 | DUSP10 |
| ENSGALG00000010778 | 367,923952 | 1,156146033 | 0,2139715 | 5,4032712 | 6,54E-08 | 1,43E-06 | CHRM3 |
| ENSGALG00000023768 | 2459,33149 | 1,155472868 | 0,1314993 | 8,78691237 | 1,54E-18 | 1,12E-16 | |
| ENSGALG00000004083 | 1247,88665 | 1,153422459 | 0,18198777 | 6,33791205 | 2,33E-10 | 7,41E-09 | GRIA1 |
| ENSGALG00000016927 | 964,07015 | 1,145535192 | 0,18286396 | 6,26441204 | 3,74E-10 | 1,16E-08 | |
| ENSGALG00000016258 | 101,512923 | 1,141614114 | 0,26933385 | 4,23865814 | 2,25E-05 | 0,0003031 | |
| ENSGALG00000006988 | 211,344493 | 1,141001037 | 0,25510377 | 4,47269383 | 7,72E-06 | 0,00011783 | |
| ENSGALG00000027554 | 57,7750225 | 1,140917568 | 0,26979126 | 4,22889002 | 2,35E-05 | 0,00031419 | |
| ENSGALG00000010166 | 921,824931 | 1,138308624 | 0,1575227 | 7,22631474 | 4,96E-13 | 2,20E-11 | VLDLR |
| ENSGALG00000022887 | 1108,58418 | 1,136537812 | 0,22465067 | 5,05913389 | 4,21E-07 | 8,11E-06 | KBTBD11 |
| ENSGALG00000009556 | 1809,38722 | 1,135549159 | 0,19219403 | 5,90834769 | 3,46E-09 | 9,02E-08 | PRICKLE1 |
| ENSGALG00000029090 | 11253,5016 | 1,13552965 | 0,10420101 | 10,8974918 | 1,18E-27 | 1,67E-25 | |
| ENSGALG00000012998 | 2212,17969 | 1,132683468 | 0,17363381 | 6,52340397 | 6,87E-11 | 2,28E-09 | WNT5B |
| ENSGALG00000015355 | 7944,79086 | 1,129448636 | 0,09943877 | 11,3582319 | 6,75E-30 | 1,04E-27 | CD47 |
| ENSGALG00000028046 | 6226,49016 | 1,129369719 | 0,09319237 | 12,1186929 | 8,41E-34 | 1,57E-31 | INSIG2 |
| ENSGALG00000011690 | 44,529172 | 1,125300184 | 0,27344667 | 4,11524556 | 3,87E-05 | 0,00049258 | |
| ENSGALG00000014866 | 49930,561 | 1,123162841 | 0,1127082 | 9,96522725 | 2,16E-23 | 2,44E-21 | |
| ENSGALG00000004974 | 170,654697 | 1,122849026 | 0,25433468 | 4,41484832 | 1,01E-05 | 0,00014999 | PPARG |
| ENSGALG00000009567 | 4756,82772 | 1,119009905 | 0,16006243 | 6,99108393 | 2,73E-12 | 1,09E-10 | TLL1 |
| ENSGALG00000027255 | 39,1566631 | 1,118838646 | 0,26596115 | 4,20677466 | 2,59E-05 | 0,00034362 | NRXN3 |
| ENSGALG00000026970 | 924,805259 | 1,11847741 | 0,16590864 | 6,74152603 | 1,57E-11 | 5,65E-10 | |
| ENSGALG00000027778 | 219,349797 | 1,117057663 | 0,24992741 | 4,46952849 | 7,84E-06 | 0,0001193 | |
| ENSGALG00000006376 | 88,1016959 | 1,116147137 | 0,27664315 | 4,03460964 | 5,47E-05 | 0,00067008 | TMC3 |
| ENSGALG00000008469 | 3420,38421 | 1,115125474 | 0,13665113 | 8,16038231 | 3,34E-16 | 2,03E-14 | SPINT1 |
| ENSGALG00000014687 | 236,291624 | 1,113638024 | 0,23763257 | 4,68638633 | 2,78E-06 | 4,61E-05 | EPHA1 |
| ENSGALG00000017044 | 264,610201 | 1,113521859 | 0,21886213 | 5,08777759 | 3,62E-07 | 7,06E-06 | TRPC4 |
| ENSGALG00000010692 | 6211,67753 | 1,111701613 | 0,10700281 | 10,3894623 | 2,77E-25 | 3,50E-23 | LRP8 |
| ENSGALG00000016470 | 648,950225 | 1,110403246 | 0,25736453 | 4,3145154 | 1,60E-05 | 0,00022558 | KCNS3 |
| ENSGALG00000010081 | 1999,17595 | 1,110146926 | 0,15885433 | 6,98845877 | 2,78E-12 | 1,11E-10 | SH3D19 |
| ENSGALG00000006337 | 29,3646416 | 1,109144959 | 0,25231504 | 4,39587332 | 1,10E-05 | 0,00016236 | |
| ENSGALG00000026763 | 669,49741 | 1,104647316 | 0,1839101 | 6,00645275 | 1,90E-09 | 5,15E-08 | |
| ENSGALG00000008182 | 270,211279 | 1,102402002 | 0,23087313 | 4,77492558 | 1,80E-06 | 3,11E-05 | CHAC2 |
| ENSGALG00000009170 | 1968,00063 | 1,101783321 | 0,11424532 | 9,64401251 | 5,21E-22 | 5,21E-20 | NCEH1 |
| ENSGALG00000006797 | 779,712052 | 1,10056236 | 0,16329659 | 6,73965302 | 1,59E-11 | 5,71E-10 | ADARB2 |
| ENSGALG00000015980 | 339,410662 | 1,100313329 | 0,24202176 | 4,54634055 | 5,46E-06 | 8,63E-05 | |
| ENSGALG00000012439 | 1157,51774 | 1,098736639 | 0,22006508 | 4,99278059 | 5,95E-07 | 1,12E-05 | |
| ENSGALG00000026518 | 640,345593 | 1,097099741 | 0,17683365 | 6,20413453 | 5,50E-10 | 1,65E-08 | RUNDC3A |
| ENSGALG00000010268 | 256,63927 | 1,095957762 | 0,25692897 | 4,265606 | 1,99E-05 | 0,00027314 | |
| ENSGALG00000028582 | 726,208031 | 1,093959706 | 0,15924578 | 6,86963091 | 6,44E-12 | 2,45E-10 | |
| ENSGALG00000026379 | 373,049265 | 1,093396741 | 0,20587222 | 5,31104558 | 1,09E-07 | 2,29E-06 | |
| ENSGALG00000002714 | 19270,5199 | 1,089564948 | 0,1023794 | 10,6424241 | 1,89E-26 | 2,49E-24 | TMEM132C |
| ENSGALG00000016875 | 1713,92149 | 1,086190922 | 0,14132808 | 7,685599 | 1,52E-14 | 8,00E-13 | |
| ENSGALG00000001434 | 188,612497 | 1,08597113 | 0,2527644 | 4,29637688 | 1,74E-05 | 0,00024177 | STOM |
| ENSGALG00000005691 | 64,1916299 | 1,085096229 | 0,27707745 | 3,91621988 | 8,99E-05 | 0,00105057 | |
| ENSGALG00000005492 | 5397,76531 | 1,08468259 | 0,14476582 | 7,49267029 | 6,75E-14 | 3,28E-12 | |
| ENSGALG00000007113 | 2485,94591 | 1,078758406 | 0,26642197 | 4,04905949 | 5,14E-05 | 0,00063816 | |
| ENSGALG00000006822 | 6257,91272 | 1,077915026 | 0,25174121 | 4,28183776 | 1,85E-05 | 0,00025592 | PCDH19 |
| ENSGALG00000013060 | 290,382045 | 1,076232694 | 0,23892972 | 4,50439019 | 6,66E-06 | 0,00010395 | ADCY2 |
| ENSGALG00000023581 | 182,271241 | 1,073595233 | 0,25670804 | 4,18216447 | 2,89E-05 | 0,0003802 | |
| ENSGALG00000006705 | 1115,46368 | 1,073202663 | 0,18895057 | 5,67980637 | 1,35E-08 | 3,29E-07 | MYOF |
| ENSGALG00000007056 | 2832,90477 | 1,069890641 | 0,15949999 | 6,70777866 | 1,98E-11 | 7,06E-10 | |
| ENSGALG00000014907 | 2488,06414 | 1,069885451 | 0,14008721 | 7,63728156 | 2,22E-14 | 1,15E-12 | DCBLD1 |
| ENSGALG00000006808 | 8235,96126 | 1,068278734 | 0,08180763 | 13,0584237 | 5,69E-39 | 1,34E-36 | TSPAN6 |
| ENSGALG00000022871 | 763,279385 | 1,067659168 | 0,16564303 | 6,44554255 | 1,15E-10 | 3,78E-09 | |
| ENSGALG00000003011 | 132,477548 | 1,06001077 | 0,26925957 | 3,93676173 | 8,26E-05 | 0,00097279 | TOM1L1 |

| | | | | | | | |
|---|---|---|---|---|---|---|---|
| ENSGALG00000028708 | 5484,0424 | 1,054408367 | 0,12654693 | 8,33215293 | 7,94E-17 | 5,07E-15 | |
| ENSGALG00000025773 | 389,005412 | 1,052013322 | 0,20058054 | 5,24484243 | 1,56E-07 | 3,20E-06 | |
| ENSGALG00000001629 | 48,4307098 | 1,051770302 | 0,27008877 | 3,89416521 | 9,85E-05 | 0,00113395 | |
| ENSGALG00000006958 | 461,662682 | 1,051053707 | 0,19372591 | 5,42546797 | 5,78E-08 | 1,28E-06 | COL27A1 |
| ENSGALG00000026534 | 317,277092 | 1,050613394 | 0,22283472 | 4,71476524 | 2,42E-06 | 4,08E-05 | |
| ENSGALG00000001020 | 6102,64046 | 1,049473012 | 0,13282357 | 7,90125574 | 2,76E-15 | 1,54E-13 | |
| ENSGALG00000008623 | 472,488323 | 1,04510086 | 0,23119186 | 4,52049152 | 6,17E-06 | 9,70E-05 | TOR4A |
| ENSGALG00000028602 | 4340,00904 | 1,04212846 | 0,26450876 | 3,93986374 | 8,15E-05 | 0,00096378 | |
| ENSGALG00000006310 | 433,388773 | 1,04012476 | 0,19660584 | 5,2904063 | 1,22E-07 | 2,55E-06 | |
| ENSGALG00000010035 | 1735,92924 | 1,03790767 | 0,15536856 | 6,68029407 | 2,38E-11 | 8,43E-10 | NR3C2 |
| ENSGALG00000013021 | 8977,79318 | 1,037759509 | 0,12348858 | 8,40368799 | 4,33E-17 | 2,78E-15 | |
| ENSGALG00000014815 | 1413,97594 | 1,036175342 | 0,13546391 | 7,64908786 | 2,02E-14 | 1,05E-12 | |
| ENSGALG00000012594 | 522,648656 | 1,03128421 | 0,20236132 | 5,09625171 | 3,46E-07 | 6,79E-06 | NTRK2 |
| ENSGALG00000006472 | 535,008316 | 1,030705214 | 0,20247632 | 5,09049761 | 3,57E-07 | 6,97E-06 | |
| ENSGALG00000017411 | 67,4262747 | 1,030664685 | 0,27683047 | 3,72308975 | 0,0001968 | 0,00206178 | RXFP3 |
| ENSGALG00000023355 | 60,0806602 | 1,029241933 | 0,27415306 | 3,75426024 | 0,00017385 | 0,00185084 | TMEM154 |
| ENSGALG00000011609 | 3308,43376 | 1,027774555 | 0,27692763 | 3,71134712 | 0,00020616 | 0,00214725 | |
| ENSGALG00000009129 | 1054,989 | 1,026339944 | 0,17522926 | 5,85712656 | 4,71E-09 | 1,21E-07 | DLX5 |
| ENSGALG00000004393 | 1206,90024 | 1,024579421 | 0,20985019 | 4,88243272 | 1,05E-06 | 1,89E-05 | |
| ENSGALG00000007646 | 1017,38466 | 1,023532832 | 0,27175014 | 3,76644827 | 0,00016559 | 0,00177191 | |
| ENSGALG00000016841 | 8687,62321 | 1,021854986 | 0,1195056 | 8,55068686 | 1,22E-17 | 8,23E-16 | |
| ENSGALG00000006252 | 11554,9175 | 1,021389981 | 0,14561324 | 7,01440332 | 2,31E-12 | 9,35E-11 | LMO4 |
| ENSGALG00000023430 | 94,0381071 | 1,019281603 | 0,25211693 | 4,04289225 | 5,28E-05 | 0,00065195 | |
| ENSGALG00000008491 | 3560,03676 | 1,018906247 | 0,13531729 | 7,52975657 | 5,08E-14 | 2,52E-12 | |
| ENSGALG00000008856 | 397,953846 | 1,015823816 | 0,23484575 | 4,32549368 | 1,52E-05 | 0,00021609 | |
| ENSGALG00000006638 | 1878,73274 | 1,014405382 | 0,11908476 | 8,51834775 | 1,62E-17 | 1,07E-15 | GRIK4 |
| ENSGALG00000028294 | 3072,93213 | 1,012130398 | 0,11397222 | 8,88050124 | 6,66E-19 | 4,90E-17 | VWA5B2 |
| ENSGALG00000002532 | 98,8821618 | 1,010182876 | 0,27499589 | 3,67344716 | 0,0002393 | 0,00244759 | GJA4 |
| ENSGALG00000014961 | 54,1316457 | 1,009675076 | 0,26059966 | 3,87442974 | 0,00010687 | 0,00122202 | |
| ENSGALG00000007069 | 496,352261 | 1,009530133 | 0,21308127 | 4,73777041 | 2,16E-06 | 3,67E-05 | |
| ENSGALG00000006908 | 12363,8802 | 1,006453251 | 0,11636849 | 8,6488472 | 5,20E-18 | 3,64E-16 | DLG1 |
| ENSGALG00000015440 | 150,286833 | 1,001448719 | 0,25673749 | 3,90067193 | 9,59E-05 | 0,00110798 | |
| ENSGALG00000001419 | 1796,40392 | 0,999950253 | 0,12013706 | 8,32341193 | 8,55E-17 | 5,43E-15 | DAB2IP |
| ENSGALG00000015208 | 5846,95619 | 0,998078994 | 0,1312595 | 7,60386116 | 2,87E-14 | 1,46E-12 | ROR2 |
| ENSGALG00000011623 | 8193,27422 | 0,997864905 | 0,27138614 | 3,67691917 | 0,00023607 | 0,00241849 | ADAMTS3 |
| ENSGALG00000010588 | 53,1286773 | 0,996815581 | 0,26205985 | 3,80377066 | 0,00014251 | 0,00155339 | |
| ENSGALG00000011963 | 622,99738 | 0,995736113 | 0,16724263 | 5,95384154 | 2,62E-09 | 6,94E-08 | PRDM5 |
| ENSGALG00000008918 | 750,133591 | 0,995278864 | 0,19014125 | 5,23441849 | 1,66E-07 | 3,37E-06 | LAMP5 |
| ENSGALG00000013738 | 4323,62842 | 0,994196056 | 0,10739219 | 9,2576201 | 2,09E-20 | 1,82E-18 | OPTN |
| ENSGALG00000011137 | 93,3711188 | 0,990105855 | 0,27694617 | 3,57508407 | 0,00035012 | 0,00339345 | |
| ENSGALG00000021812 | 73,5735967 | 0,986192682 | 0,27617874 | 3,57084933 | 0,00035583 | 0,00343178 | |
| ENSGALG00000006717 | 1503,28109 | 0,985550396 | 0,24782699 | 3,97676777 | 6,99E-05 | 0,00083864 | |
| ENSGALG00000003149 | 1610,9192 | 0,985359473 | 0,14766495 | 6,67294075 | 2,51E-11 | 8,83E-10 | |
| ENSGALG00000004917 | 473,928512 | 0,985260253 | 0,2036331 | 4,83840905 | 1,31E-06 | 2,33E-05 | |
| ENSGALG00000005990 | 258,942154 | 0,985024441 | 0,26677048 | 3,6924042 | 0,00022214 | 0,00229463 | |
| ENSGALG00000005692 | 405,777143 | 0,983308202 | 0,24246612 | 4,05544582 | 5,00E-05 | 0,00062331 | |
| ENSGALG00000020723 | 739,146919 | 0,981929794 | 0,17104516 | 5,74076334 | 9,43E-09 | 2,33E-07 | |
| ENSGALG00000016537 | 9995,53899 | 0,979050265 | 0,13793185 | 7,09807238 | 1,27E-12 | 5,29E-11 | SCML2 |
| ENSGALG00000027497 | 260,124381 | 0,976598532 | 0,22923317 | 4,26028447 | 2,04E-05 | 0,00027881 | |
| ENSGALG00000004930 | 264,560406 | 0,975284752 | 0,27532773 | 3,54226849 | 0,0003967 | 0,00376505 | HRH1 |
| ENSGALG00000016143 | 2862,24797 | 0,975263193 | 0,16498108 | 5,91136394 | 3,39E-09 | 8,88E-08 | |
| ENSGALG00000010926 | 130,297678 | 0,975212806 | 0,27472538 | 3,54977322 | 0,00038556 | 0,00367048 | |
| ENSGALG00000012808 | 5197,90929 | 0,972939427 | 0,12111181 | 8,03339831 | 9,48E-16 | 5,57E-14 | CDYL |
| ENSGALG00000003739 | 38757,8816 | 0,97006222 | 0,0963384 | 10,0693205 | 7,55E-24 | 8,74E-22 | |
| ENSGALG00000003939 | 112,384247 | 0,968631016 | 0,26679037 | 3,63068202 | 0,00028267 | 0,00282651 | |
| ENSGALG00000007343 | 72,8603799 | 0,967183937 | 0,2759661 | 3,50472014 | 0,00045709 | 0,00421349 | |
| ENSGALG00000016341 | 139,195007 | 0,965199253 | 0,25730295 | 3,75121721 | 0,00017598 | 0,00186552 | |
| ENSGALG00000014343 | 785,569079 | 0,963080504 | 0,16453536 | 5,85333465 | 4,82E-09 | 1,24E-07 | |
| ENSGALG00000002509 | 14144,0875 | 0,962423205 | 0,119871 | 8,0288245 | 9,84E-16 | 5,75E-14 | |
| ENSGALG00000021879 | 34,5191458 | 0,950789217 | 0,2438749 | 3,89867607 | 9,67E-05 | 0,00111509 | |
| ENSGALG00000006614 | 57,8438472 | 0,950175613 | 0,27642148 | 3,437416 | 0,00058729 | 0,00519918 | |
| ENSGALG00000006786 | 1099,77465 | 0,950169855 | 0,14131213 | 6,72390847 | 1,77E-11 | 6,34E-10 | |
| ENSGALG00000016843 | 6732,69779 | 0,946578268 | 0,12267336 | 7,71624989 | 1,20E-14 | 6,38E-13 | COL4A2 |
| ENSGALG00000027805 | 264,08193 | 0,945988714 | 0,26256163 | 3,60292061 | 0,00031466 | 0,00309685 | |

| | | | | | | | |
|---|---|---|---|---|---|---|---|
| ENSGALG00000017135 | 1317,87874 | 0,945725278 | 0,15347839 | 6,16194417 | 7,19E-10 | 2,11E-08 | CRYL1 |
| ENSGALG00000011336 | 41,6337265 | 0,944781785 | 0,2663112 | 3,54766074 | 0,00038867 | 0,00369723 | |
| ENSGALG00000012224 | 1861,1774 | 0,94448356 | 0,15084874 | 6,26112991 | 3,82E-10 | 1,18E-08 | |
| ENSGALG00000008014 | 268,980372 | 0,943508119 | 0,22646821 | 4,16618353 | 3,10E-05 | 0,00040481 | CEBPB |
| ENSGALG00000026892 | 80,5665735 | 0,943046494 | 0,2766565 | 3,40872711 | 0,00065267 | 0,00568143 | |
| ENSGALG00000010707 | 23,462609 | 0,94082246 | 0,24072754 | 3,90824608 | 9,30E-05 | 0,0010798 | GPR68 |
| ENSGALG00000015008 | 34,4613342 | 0,940194042 | 0,26427134 | 3,55768451 | 0,00037414 | 0,00358079 | ZNF366 |
| ENSGALG00000001405 | 278,869771 | 0,939567412 | 0,21873168 | 4,29552511 | 1,74E-05 | 0,00024223 | IRF6 |
| ENSGALG00000010163 | 10137,7585 | 0,938977707 | 0,10980213 | 8,55154343 | 1,21E-17 | 8,21E-16 | LGR5 |
| ENSGALG00000004662 | 3197,84361 | 0,937959127 | 0,14071695 | 6,66557313 | 2,64E-11 | 9,24E-10 | DISP3 |
| ENSGALG00000010836 | 1075,45486 | 0,937907565 | 0,15481833 | 6,05811696 | 1,38E-09 | 3,83E-08 | |
| ENSGALG00000008206 | 194,334171 | 0,934029761 | 0,24433133 | 3,82279978 | 0,00013194 | 0,00145547 | TESC |
| ENSGALG00000002397 | 809,586463 | 0,927215999 | 0,22531285 | 4,11523792 | 3,87E-05 | 0,00049258 | PSPH |
| ENSGALG00000010412 | 554,029025 | 0,925287079 | 0,19363821 | 4,77843223 | 1,77E-06 | 3,07E-05 | WWTR1 |
| ENSGALG00000026370 | 105,807662 | 0,924816521 | 0,26789368 | 3,45217742 | 0,00055608 | 0,00495797 | |
| ENSGALG00000004519 | 193,098236 | 0,924449735 | 0,26575159 | 3,47862353 | 0,000504 | 0,00456847 | TRPV2 |
| ENSGALG00000007039 | 10074,0502 | 0,92399781 | 0,125208 | 7,37970282 | 1,59E-13 | 7,38E-12 | |
| ENSGALG00000008778 | 5061,18074 | 0,923325024 | 0,13889594 | 6,6476025 | 2,98E-11 | 1,04E-09 | SPRED2 |
| ENSGALG00000010785 | 431,614767 | 0,922406238 | 0,22004896 | 4,19182269 | 2,77E-05 | 0,00036514 | ACOT11 |
| ENSGALG00000015164 | 3009,10926 | 0,920081875 | 0,10753234 | 8,55632742 | 1,17E-17 | 7,92E-16 | |
| ENSGALG00000014855 | 3796,22691 | 0,916891226 | 0,15123034 | 6,06287898 | 1,34E-09 | 3,74E-08 | |
| ENSGALG00000016443 | 4759,48054 | 0,912806646 | 0,13882964 | 6,57501249 | 4,86E-11 | 1,65E-09 | HPCAL1 |
| ENSGALG00000023193 | 891,994507 | 0,912803471 | 0,15632952 | 5,83897064 | 5,25E-09 | 1,34E-07 | |
| ENSGALG00000007667 | 897,591606 | 0,910790115 | 0,15122323 | 6,02281894 | 1,71E-09 | 4,71E-08 | GPR158 |
| ENSGALG00000007416 | 138,697657 | 0,91055725 | 0,26535747 | 3,43143628 | 0,00060039 | 0,00530016 | CD3E |
| ENSGALG00000012533 | 31364,4878 | 0,909053879 | 0,13047228 | 6,96741015 | 3,23E-12 | 1,27E-10 | |
| ENSGALG00000011618 | 1026,95651 | 0,908730543 | 0,18188258 | 4,99624825 | 5,85E-07 | 1,10E-05 | ADSSL1 |
| ENSGALG00000015624 | 28928,123 | 0,906952704 | 0,26927951 | 3,36807168 | 0,00075696 | 0,00646335 | VCAN |
| ENSGALG00000001645 | 772,05714 | 0,906502233 | 0,18722125 | 4,84187682 | 1,29E-06 | 2,30E-05 | KCNT1 |
| ENSGALG00000011258 | 41539,9101 | 0,899386948 | 0,13199364 | 6,81386582 | 9,50E-12 | 3,51E-10 | |
| ENSGALG00000003401 | 97,8434771 | 0,897409124 | 0,27523071 | 3,26057042 | 0,00111188 | 0,00890653 | FBXL22 |
| ENSGALG00000017652 | 38,6817197 | 0,894447523 | 0,26978 | 3,31547007 | 0,00091489 | 0,00757906 | TTC9 |
| ENSGALG00000004189 | 665,542257 | 0,89261736 | 0,19559277 | 4,56365205 | 5,03E-06 | 8,01E-05 | MAFK |
| ENSGALG00000025723 | 106,160903 | 0,89142204 | 0,27513226 | 3,23997649 | 0,0011954 | 0,00944015 | |
| ENSGALG00000016308 | 4475,13284 | 0,888326042 | 0,09104509 | 9,75698981 | 1,72E-22 | 1,77E-20 | MYC |
| ENSGALG00000007952 | 16328,0045 | 0,887725324 | 0,11490596 | 7,72566813 | 1,11E-14 | 5,95E-13 | AMOT |
| ENSGALG00000001281 | 97,6098799 | 0,886212495 | 0,27180262 | 3,26050025 | 0,00111216 | 0,00890653 | |
| ENSGALG00000012111 | 193,797146 | 0,886059032 | 0,25134734 | 3,52523742 | 0,0004231 | 0,00396449 | TMEM37 |
| ENSGALG00000010148 | 95,0993319 | 0,885936489 | 0,27329578 | 3,24167639 | 0,00118829 | 0,00940681 | |
| ENSGALG00000001276 | 18245,4455 | 0,885240877 | 0,11268934 | 7,8555867 | 3,98E-15 | 2,20E-13 | HOXB9 |
| ENSGALG00000015896 | 1982,50233 | 0,883690573 | 0,14767564 | 5,98399687 | 2,18E-09 | 5,84E-08 | |
| ENSGALG00000010090 | 526,624476 | 0,882987641 | 0,22809712 | 3,87110386 | 0,00010834 | 0,0012348 | FAM160A1 |
| ENSGALG00000009594 | 606,263065 | 0,882488341 | 0,19693947 | 4,48101319 | 7,43E-06 | 0,00011388 | RARRES1 |
| ENSGALG00000005256 | 9190,22784 | 0,880582454 | 0,23610272 | 3,72965825 | 0,00019174 | 0,00201889 | EPHA4 |
| ENSGALG00000009179 | 4215,94774 | 0,880187457 | 0,10683081 | 8,23907883 | 1,74E-16 | 1,08E-14 | TNFSF10 |
| ENSGALG00000011172 | 2449,32108 | 0,879786034 | 0,13379316 | 6,57571768 | 4,84E-11 | 1,65E-09 | |
| ENSGALG00000027947 | 77,7692928 | 0,877628202 | 0,26244862 | 3,34400006 | 0,0008258 | 0,00697665 | GHSR |
| ENSGALG00000027885 | 26,4620215 | 0,876800098 | 0,23993489 | 3,65432511 | 0,00025786 | 0,00260968 | C4orf19 |
| ENSGALG00000014783 | 8965,42611 | 0,875888949 | 0,11142269 | 7,86095696 | 3,81E-15 | 2,12E-13 | |
| ENSGALG00000009405 | 358,011724 | 0,875608773 | 0,20294284 | 4,31455849 | 1,60E-05 | 0,00022558 | GRIA2 |
| ENSGALG00000010996 | 3029,60967 | 0,875351394 | 0,11601158 | 7,54537966 | 4,51E-14 | 2,26E-12 | IRF2BP2 |
| ENSGALG00000028759 | 391,741267 | 0,87268698 | 0,23549591 | 3,70574156 | 0,00021077 | 0,00219166 | LYPD6B |
| ENSGALG00000008018 | 469,138933 | 0,868526004 | 0,20965704 | 4,14260361 | 3,43E-05 | 0,00044328 | SNAI1 |
| ENSGALG00000006008 | 593,4561 | 0,867963452 | 0,20178146 | 4,30150244 | 1,70E-05 | 0,0002371 | HOMER2 |
| ENSGALG00000029023 | 178,138305 | 0,865902724 | 0,26156953 | 3,3104113 | 0,00093159 | 0,00768178 | DMRTB1 |
| ENSGALG00000016909 | 3574,55237 | 0,864393738 | 0,09752748 | 8,86307897 | 7,78E-19 | 5,69E-17 | NDFIP2 |
| ENSGALG00000026582 | 1114,9679 | 0,863598751 | 0,1902411 | 4,53949614 | 5,64E-06 | 8,89E-05 | |
| ENSGALG00000027684 | 3428,7366 | 0,863457403 | 0,14910293 | 5,79101571 | 7,00E-09 | 1,75E-07 | |
| ENSGALG00000014349 | 1819,66371 | 0,860574566 | 0,11955806 | 7,19796341 | 6,11E-13 | 2,65E-11 | |
| ENSGALG00000011613 | 475,600449 | 0,857977883 | 0,20239708 | 4,23908233 | 2,24E-05 | 0,0003031 | |
| ENSGALG00000001617 | 271,795228 | 0,855845667 | 0,23372685 | 3,6617345 | 0,00025051 | 0,0025415 | CAMKK1 |
| ENSGALG00000001926 | 1199,81634 | 0,853283247 | 0,15483373 | 5,51096475 | 3,57E-08 | 8,22E-07 | HSPB1 |
| ENSGALG00000015180 | 12680,3183 | 0,851097646 | 0,10970107 | 7,75833511 | 8,61E-15 | 4,64E-13 | PSAT1 |
| ENSGALG00000009396 | 1375,97204 | 0,850250642 | 0,16438706 | 5,17224786 | 2,31E-07 | 4,59E-06 | |

| | | | | | | | |
|---|---|---|---|---|---|---|---|
| ENSGALG00000009172 | 460,012151 | 0,849588562 | 0,22136087 | 3,83802504 | 0,00012403 | 0,001384 | OSBPL6 |
| ENSGALG00000016807 | 8544,75416 | 0,848101313 | 0,11909181 | 7,12140736 | 1,07E-12 | 4,55E-11 | |
| ENSGALG00000010291 | 5393,74102 | 0,844497871 | 0,10367256 | 8,14581883 | 3,77E-16 | 2,28E-14 | RBPMS |
| ENSGALG00000000977 | 7029,01558 | 0,842188766 | 0,08563672 | 9,83443558 | 8,00E-23 | 8,48E-21 | MAPKAP1 |
| ENSGALG00000009530 | 719,235997 | 0,841159622 | 0,18836636 | 4,46555117 | 7,99E-06 | 0,0001212 | PDZRN4 |
| ENSGALG00000004155 | 211,984568 | 0,841077447 | 0,24078857 | 3,49301238 | 0,0004776 | 0,00436403 | MYOM3 |
| ENSGALG00000009378 | 1120,38163 | 0,840979613 | 0,13880777 | 6,05859175 | 1,37E-09 | 3,83E-08 | PDGFC |
| ENSGALG00000002335 | 624,130373 | 0,840686726 | 0,18168554 | 4,62715265 | 3,71E-06 | 6,03E-05 | DOCK3 |
| ENSGALG00000002623 | 461,663405 | 0,839503746 | 0,18687975 | 4,49221356 | 7,05E-06 | 0,00010872 | PCOLCE2 |
| ENSGALG00000005410 | 3123,4698 | 0,838361241 | 0,15163748 | 5,52872037 | 3,23E-08 | 7,46E-07 | |
| ENSGALG00000007932 | 1262,37356 | 0,837645303 | 0,16803809 | 4,98485361 | 6,20E-07 | 1,16E-05 | TRAF6 |
| ENSGALG00000015409 | 14646,8005 | 0,837042297 | 0,10986863 | 7,61857419 | 2,56E-14 | 1,32E-12 | PROS1 |
| ENSGALG00000012278 | 12900,8817 | 0,835859144 | 0,23517071 | 3,55426545 | 0,00037904 | 0,00361936 | |
| ENSGALG00000010156 | 256,288262 | 0,833499217 | 0,2396958 | 3,47732084 | 0,00050645 | 0,0045874 | |
| ENSGALG00000001220 | 3532,90233 | 0,831042878 | 0,11299613 | 7,35461342 | 1,91E-13 | 8,81E-12 | HERPUD1 |
| ENSGALG00000013085 | 2051,84727 | 0,830893947 | 0,12691125 | 6,54704716 | 5,87E-11 | 1,98E-09 | PTPRO |
| ENSGALG00000025770 | 327,819551 | 0,830841194 | 0,24237973 | 3,42784926 | 0,00060838 | 0,00535557 | |
| ENSGALG00000012354 | 2002,91372 | 0,829092837 | 0,15052249 | 5,50809953 | 3,63E-08 | 8,31E-07 | |
| ENSGALG00000000639 | 543,002187 | 0,827648848 | 0,21824285 | 3,79232975 | 0,00014924 | 0,00161912 | LRRN2 |
| ENSGALG00000005768 | 4764,60581 | 0,825923962 | 0,11172719 | 7,39232759 | 1,44E-13 | 6,76E-12 | ETV5 |
| ENSGALG00000011232 | 2354,98602 | 0,825691237 | 0,13504902 | 6,11401152 | 9,72E-10 | 2,77E-08 | |
| ENSGALG00000017246 | 1270,40827 | 0,825125037 | 0,15896962 | 5,19045723 | 2,10E-07 | 4,19E-06 | ME3 |
| ENSGALG00000008883 | 4099,49868 | 0,82333394 | 0,11713657 | 7,02883754 | 2,08E-12 | 8,57E-11 | TCF7L2 |
| ENSGALG00000020899 | 521,609979 | 0,823141433 | 0,24736905 | 3,3275846 | 0,00087602 | 0,00731033 | |
| ENSGALG00000003726 | 904,249522 | 0,818923475 | 0,22585036 | 3,625956 | 0,00028789 | 0,00286954 | EGFLAM |
| ENSGALG00000010664 | 324,510001 | 0,817672769 | 0,23994525 | 3,40774723 | 0,00065502 | 0,00569168 | KIF26B |
| ENSGALG00000002054 | 1352,81303 | 0,816030183 | 0,13128078 | 6,21591508 | 5,10E-10 | 1,54E-08 | |
| ENSGALG00000008670 | 1990,92325 | 0,815447938 | 0,13744115 | 5,9330699 | 2,97E-09 | 7,83E-08 | |
| ENSGALG00000013683 | 16575,6485 | 0,810200928 | 0,12018138 | 6,74148471 | 1,57E-11 | 5,65E-10 | ARID1B |
| ENSGALG00000012848 | 3319,46891 | 0,809997189 | 0,15987767 | 5,06635591 | 4,06E-07 | 7,83E-06 | |
| ENSGALG00000014872 | 561,414638 | 0,806209646 | 0,23662254 | 3,40715487 | 0,00065644 | 0,0056984 | FGF10 |
| ENSGALG00000009155 | 822,192391 | 0,800986669 | 0,16316786 | 4,90897338 | 9,16E-07 | 1,68E-05 | |
| ENSGALG00000005430 | 836,618226 | 0,79963845 | 0,20651888 | 3,87198711 | 0,00010795 | 0,00123209 | ALS2CL |
| ENSGALG00000012849 | 4391,56992 | 0,79670503 | 0,13188371 | 6,04096596 | 1,53E-09 | 4,24E-08 | SLC37A3 |
| ENSGALG00000012277 | 3279,13345 | 0,79518636 | 0,11025611 | 7,21217515 | 5,51E-13 | 2,42E-11 | |
| ENSGALG00000015952 | 320,695046 | 0,793177929 | 0,23917365 | 3,31632662 | 0,00091209 | 0,00756087 | |
| ENSGALG00000004599 | 1177,35629 | 0,791092143 | 0,17593216 | 4,49657499 | 6,91E-06 | 0,00010704 | |
| ENSGALG00000026631 | 639,083895 | 0,790472215 | 0,23279997 | 3,39549971 | 0,00068503 | 0,00592197 | HOXA10 |
| ENSGALG00000002443 | 515,246567 | 0,789654091 | 0,18384496 | 4,29521754 | 1,75E-05 | 0,0002423 | FBXL16 |
| ENSGALG00000008445 | 716,590826 | 0,788973502 | 0,17118936 | 4,60877658 | 4,05E-06 | 6,52E-05 | SLC24A3 |
| ENSGALG00000009718 | 365,400186 | 0,788380099 | 0,1998647 | 3,94456899 | 7,99E-05 | 0,00094788 | FAM53B |
| ENSGALG00000009252 | 6799,64958 | 0,7810557 | 0,11275794 | 6,92683575 | 4,30E-12 | 1,68E-10 | |
| ENSGALG00000026034 | 890,686008 | 0,780909637 | 0,16319128 | 4,7852411 | 1,71E-06 | 2,98E-05 | RGS17 |
| ENSGALG00000002598 | 7375,09654 | 0,778822215 | 0,08757756 | 8,89294263 | 5,95E-19 | 4,46E-17 | TPST1 |
| ENSGALG00000005310 | 792,651331 | 0,773499973 | 0,20804946 | 3,71786588 | 0,00020091 | 0,0020996 | |
| ENSGALG00000021238 | 292,286817 | 0,771321549 | 0,21541142 | 3,58069015 | 0,00034269 | 0,00333594 | |
| ENSGALG00000026355 | 316,135871 | 0,769808548 | 0,22820508 | 3,37331911 | 0,00074268 | 0,00636312 | |
| ENSGALG00000004158 | 2408,26029 | 0,769428918 | 0,12659815 | 6,07772646 | 1,22E-09 | 3,45E-08 | |
| ENSGALG00000001346 | 433,934694 | 0,76776324 | 0,23273948 | 3,29880968 | 0,00097096 | 0,00794876 | |
| ENSGALG00000013124 | 3883,6626 | 0,765332021 | 0,13324393 | 5,74384156 | 9,26E-09 | 2,29E-07 | FHOD3 |
| ENSGALG00000025780 | 468,175583 | 0,762758431 | 0,19965686 | 3,82034666 | 0,00013326 | 0,00146485 | ITPRIP |
| ENSGALG00000023886 | 345,558396 | 0,76245624 | 0,20569517 | 3,70672893 | 0,00020995 | 0,00218495 | ZNF804A |
| ENSGALG00000011376 | 374,497295 | 0,758366172 | 0,20404377 | 3,71668381 | 0,00020185 | 0,00210768 | ANKRD9 |
| ENSGALG00000016069 | 4206,1542 | 0,758301426 | 0,09589383 | 7,90771867 | 2,62E-15 | 1,47E-13 | |
| ENSGALG00000001000 | 105882,406 | 0,756534693 | 0,12130242 | 6,23676501 | 4,47E-10 | 1,36E-08 | |
| ENSGALG00000013616 | 416,825405 | 0,749712043 | 0,2111769 | 3,55016123 | 0,000385 | 0,00366786 | OPRM1 |
| ENSGALG00000009145 | 1069,11169 | 0,74529577 | 0,2190544 | 3,40233194 | 0,00066813 | 0,00578387 | |
| ENSGALG00000012012 | 5118,87574 | 0,743046962 | 0,14577197 | 5,09732411 | 3,44E-07 | 6,76E-06 | |
| ENSGALG00000011319 | 11225,2963 | 0,737544915 | 0,09161073 | 8,05085711 | 8,22E-16 | 4,85E-14 | ADGRL3 |
| ENSGALG00000005512 | 7764,5793 | 0,736922168 | 0,10571311 | 6,97096281 | 3,15E-12 | 1,25E-10 | |
| ENSGALG00000000505 | 339,344776 | 0,73643955 | 0,20375113 | 3,61440716 | 0,00030104 | 0,00298625 | KCNH6 |
| ENSGALG00000006138 | 2327,25759 | 0,736205494 | 0,17788661 | 4,13862223 | 3,49E-05 | 0,00045058 | |
| ENSGALG00000003701 | 2093,50715 | 0,734267222 | 0,12693056 | 5,78479482 | 7,26E-09 | 1,81E-07 | SGMS1 |
| ENSGALG00000003043 | 2889,49683 | 0,734167184 | 0,12468763 | 5,88805143 | 3,91E-09 | 1,02E-07 | |

| | | | | | | | |
|---|---|---|---|---|---|---|---|
| ENSGALG00000015250 | 1236,64137 | 0,732130121 | 0,18294116 | 4,00199787 | 6,28E-05 | 0,00075985 | |
| ENSGALG00000028155 | 1607,60298 | 0,730708288 | 0,1348053 | 5,42047159 | 5,94E-08 | 1,31E-06 | |
| ENSGALG00000003774 | 6718,46982 | 0,73043073 | 0,11984382 | 6,09485535 | 1,10E-09 | 3,11E-08 | ATP13A2 |
| ENSGALG00000005653 | 2602,66741 | 0,729875771 | 0,10427934 | 6,99923686 | 2,57E-12 | 1,03E-10 | NFKB2 |
| ENSGALG00000007501 | 443,889264 | 0,729453053 | 0,20909338 | 3,48864733 | 0,00048547 | 0,00442944 | |
| ENSGALG00000025887 | 833,811929 | 0,729103686 | 0,19083714 | 3,8205545 | 0,00013315 | 0,00146485 | ATF3 |
| ENSGALG00000015184 | 6562,38853 | 0,727266294 | 0,09772755 | 7,44177369 | 9,93E-14 | 4,72E-12 | |
| ENSGALG00000002955 | 1085,67322 | 0,725828311 | 0,20628485 | 3,51857296 | 0,00043387 | 0,00404688 | ADAMTSL2 |
| ENSGALG00000028935 | 4407,81779 | 0,725128037 | 0,09141746 | 7,9320517 | 2,16E-15 | 1,22E-13 | BCL10 |
| ENSGALG00000009381 | 14937,4753 | 0,723727203 | 0,11606588 | 6,23548619 | 4,50E-10 | 1,37E-08 | |
| ENSGALG00000014441 | 1267,12755 | 0,722905404 | 0,15192936 | 4,75816803 | 1,95E-06 | 3,35E-05 | |
| ENSGALG00000027352 | 11317,2552 | 0,722863534 | 0,10576918 | 6,83434951 | 8,24E-12 | 3,08E-10 | |
| ENSGALG00000001370 | 1171,81779 | 0,722550699 | 0,15690532 | 4,60501077 | 4,12E-06 | 6,62E-05 | ADAMTS8 |
| ENSGALG00000007206 | 17,4582489 | 0,720355559 | 0,22314617 | 3,22817802 | 0,00124581 | 0,00980119 | |
| ENSGALG00000015947 | 765,779345 | 0,718924866 | 0,22274381 | 3,22758633 | 0,00124839 | 0,00980672 | |
| ENSGALG00000008163 | 14921,5324 | 0,713636421 | 0,10768275 | 6,62721234 | 3,42E-11 | 1,18E-09 | PSME4 |
| ENSGALG00000026727 | 425,717003 | 0,713627256 | 0,19018155 | 3,75234749 | 0,00017519 | 0,00186028 | SLC30A10 |
| ENSGALG00000028279 | 322,93543 | 0,711492754 | 0,21583535 | 3,29646077 | 0,00097911 | 0,00800505 | SNORD14 |
| ENSGALG00000007712 | 5151,87784 | 0,709640176 | 0,09468723 | 7,49457113 | 6,65E-14 | 3,25E-12 | MYO1B |
| ENSGALG00000005423 | 1960,17378 | 0,707902433 | 0,13324908 | 5,31262534 | 1,08E-07 | 2,28E-06 | GRHPR |
| ENSGALG00000015975 | 5061,20264 | 0,705892008 | 0,11070199 | 6,3765067 | 1,81E-10 | 5,84E-09 | |
| ENSGALG00000002892 | 626,326751 | 0,704967282 | 0,1963291 | 3,59074262 | 0,00032974 | 0,00322241 | |
| ENSGALG00000015310 | 3979,38646 | 0,701805821 | 0,09924244 | 7,07163008 | 1,53E-12 | 6,34E-11 | |
| ENSGALG00000000452 | 1371,04529 | 0,701787311 | 0,16834665 | 4,16870367 | 3,06E-05 | 0,00040083 | |
| ENSGALG00000002198 | 1124,31872 | 0,701337674 | 0,20146263 | 3,4812296 | 0,00049912 | 0,00452753 | FHOD1 |
| ENSGALG00000010522 | 10232,0506 | 0,699893806 | 0,1480305 | 4,72803795 | 2,27E-06 | 3,84E-05 | |
| ENSGALG00000027983 | 1055,55115 | 0,699268766 | 0,16725899 | 4,18075434 | 2,91E-05 | 0,00038217 | TNK2 |
| ENSGALG00000010204 | 874,20571 | 0,699072042 | 0,16632037 | 4,20316555 | 2,63E-05 | 0,00034841 | KCNC2 |
| ENSGALG00000004088 | 3580,2986 | 0,695518825 | 0,10391048 | 6,69344229 | 2,18E-11 | 7,75E-10 | FNBP1 |
| ENSGALG00000008680 | 626,282132 | 0,694380448 | 0,18287186 | 3,79708749 | 0,00014641 | 0,00159251 | |
| ENSGALG00000011266 | 760,864805 | 0,693047043 | 0,18382819 | 3,77008024 | 0,0001632 | 0,00174928 | PPM1K |
| ENSGALG00000015721 | 1214,39078 | 0,692751518 | 0,18278823 | 3,78991313 | 0,0001507 | 0,00163213 | SVEP1 |
| ENSGALG00000012956 | 536,621695 | 0,692712504 | 0,21091071 | 3,28438757 | 0,00102204 | 0,00828563 | TMTC1 |
| ENSGALG00000011025 | 4159,539 | 0,69039215 | 0,09483213 | 7,28015057 | 3,33E-13 | 1,50E-11 | |
| ENSGALG00000012211 | 4094,58636 | 0,688791118 | 0,10047528 | 6,85532894 | 7,11E-12 | 2,69E-10 | PDLIM5 |
| ENSGALG00000005065 | 974,960396 | 0,685705936 | 0,15970851 | 4,29348406 | 1,76E-05 | 0,00024393 | PLA2G4A |
| ENSGALG00000016835 | 5266,33357 | 0,685225343 | 0,13174668 | 5,20108259 | 1,98E-07 | 3,97E-06 | |
| ENSGALG00000012696 | 8253,81984 | 0,682603698 | 0,09998162 | 6,82729216 | 8,65E-12 | 3,21E-10 | |
| ENSGALG00000016281 | 10145,7509 | 0,681257671 | 0,08559267 | 7,95929876 | 1,73E-15 | 9,93E-14 | DMD |
| ENSGALG00000023725 | 1053,58774 | 0,679129782 | 0,15467581 | 4,39066583 | 1,13E-05 | 0,00016591 | |
| ENSGALG00000003640 | 865,821934 | 0,678918743 | 0,16063195 | 4,22654863 | 2,37E-05 | 0,00031683 | |
| ENSGALG00000027627 | 7249,84284 | 0,677661132 | 0,0846528 | 8,00518259 | 1,19E-15 | 6,94E-14 | |
| ENSGALG00000027342 | 11212,2371 | 0,672830693 | 0,07880349 | 8,5380821 | 1,36E-17 | 9,08E-16 | |
| ENSGALG00000011550 | 2837,22447 | 0,672556317 | 0,14266182 | 4,71433983 | 2,42E-06 | 4,08E-05 | |
| ENSGALG00000027389 | 3615,54123 | 0,672548469 | 0,11214472 | 5,99714785 | 2,01E-09 | 5,41E-08 | RGS7BP |
| ENSGALG00000002646 | 9732,02947 | 0,671803119 | 0,1449266 | 4,63547136 | 3,56E-06 | 5,80E-05 | FAM171B |
| ENSGALG00000004612 | 5923,23561 | 0,671680213 | 0,09027919 | 7,44003379 | 1,01E-13 | 4,77E-12 | MTHFR |
| ENSGALG00000014106 | 16074,824 | 0,667075995 | 0,09963033 | 6,69551116 | 2,15E-11 | 7,66E-10 | PPFIBP1 |
| ENSGALG00000013342 | 2747,02869 | 0,666913547 | 0,1119687 | 5,95624957 | 2,58E-09 | 6,85E-08 | GBX2 |
| ENSGALG00000007704 | 16606,5878 | 0,666805577 | 0,12692892 | 5,25337789 | 1,49E-07 | 3,07E-06 | |
| ENSGALG00000023876 | 3391,40413 | 0,666027019 | 0,11746061 | 5,67021584 | 1,43E-08 | 3,46E-07 | |
| ENSGALG00000015887 | 11962,9806 | 0,665885713 | 0,11836362 | 5,62576338 | 1,85E-08 | 4,42E-07 | URB1 |
| ENSGALG00000003957 | 1955,41186 | 0,665113393 | 0,12648287 | 5,25852528 | 1,45E-07 | 2,99E-06 | |
| ENSGALG00000006872 | 8598,0072 | 0,664035354 | 0,09325855 | 7,12036955 | 1,08E-12 | 4,56E-11 | PISD |
| ENSGALG00000004437 | 1051,52128 | 0,662144004 | 0,15643348 | 4,23275119 | 2,31E-05 | 0,00031017 | LHFPL2 |
| ENSGALG00000014957 | 23554,8466 | 0,661960443 | 0,09128513 | 7,25156958 | 4,12E-13 | 1,84E-11 | WDR1 |
| ENSGALG00000002655 | 5806,05662 | 0,657486198 | 0,10761985 | 6,10933924 | 1,00E-09 | 2,84E-08 | ITGAV |
| ENSGALG00000004661 | 7333,89219 | 0,654997852 | 0,09584164 | 6,83416796 | 8,25E-12 | 3,08E-10 | |
| ENSGALG00000004231 | 624,271591 | 0,654722209 | 0,17330476 | 3,77786627 | 0,00015818 | 0,00170573 | IFNLR1 |
| ENSGALG00000011770 | 13005,5067 | 0,65450966 | 0,08870671 | 7,37835601 | 1,60E-13 | 7,42E-12 | UMPS |
| ENSGALG00000016803 | 4124,86158 | 0,652622211 | 0,11026512 | 5,91866387 | 3,25E-09 | 8,51E-08 | ST6GAL2 |
| ENSGALG00000016719 | 6369,91266 | 0,652460747 | 0,11253545 | 5,79782427 | 6,72E-09 | 1,69E-07 | TNFRSF21 |
| ENSGALG00000010186 | 4216,82392 | 0,652306301 | 0,0936837 | 6,96285777 | 3,33E-12 | 1,31E-10 | |
| ENSGALG00000014545 | 998,185181 | 0,650360352 | 0,17599459 | 3,69534294 | 0,00021959 | 0,00227389 | |

| | | | | | | | |
|---|---|---|---|---|---|---|---|
| ENSGALG00000005906 | 1908,90153 | 0,646437734 | 0,11565287 | 5,5894654 | 2,28E-08 | 5,39E-07 | |
| ENSGALG00000004042 | 4037,76912 | 0,646319325 | 0,11291506 | 5,72394258 | 1,04E-08 | 2,56E-07 | |
| ENSGALG00000018479 | 474,539705 | 0,645095331 | 0,19673783 | 3,27895931 | 0,00104191 | 0,00843028 | |
| ENSGALG00000012877 | 2598,2138 | 0,643827392 | 0,1203305 | 5,35049194 | 8,77E-08 | 1,87E-06 | CREB3L2 |
| ENSGALG00000013031 | 981,078598 | 0,641018743 | 0,1528676 | 4,19329383 | 2,75E-05 | 0,00036355 | CECR1 |
| ENSGALG00000027835 | 4356,75822 | 0,637590923 | 0,09597243 | 6,64348014 | 3,06E-11 | 1,06E-09 | PCBD1 |
| ENSGALG00000024346 | 783,051886 | 0,637233355 | 0,1537054 | 4,14580971 | 3,39E-05 | 0,00043848 | |
| ENSGALG00000002248 | 5849,29235 | 0,635943683 | 0,08808525 | 7,21963901 | 5,21E-13 | 2,30E-11 | |
| ENSGALG00000026776 | 1287,36825 | 0,634230033 | 0,13457396 | 4,71287338 | 2,44E-06 | 4,10E-05 | TIGAR |
| ENSGALG00000008318 | 17655,4148 | 0,633974687 | 0,08385645 | 7,56023739 | 4,02E-14 | 2,02E-12 | SLK |
| ENSGALG00000015842 | 5393,93615 | 0,632020307 | 0,09677563 | 6,53077971 | 6,54E-11 | 2,19E-09 | TIAM1 |
| ENSGALG00000004181 | 1516,86359 | 0,628709896 | 0,12659877 | 4,96616114 | 6,83E-07 | 1,27E-05 | |
| ENSGALG00000014922 | 13357,9815 | 0,627329029 | 0,08060516 | 7,7827402 | 7,10E-15 | 3,88E-13 | ROCK1 |
| ENSGALG00000002602 | 2698,72514 | 0,625832331 | 0,15063415 | 4,15465119 | 3,26E-05 | 0,00042362 | |
| ENSGALG00000012726 | 94562,7089 | 0,625221924 | 0,1022285 | 6,11592611 | 9,60E-10 | 2,75E-08 | |
| ENSGALG00000027696 | 2435,05731 | 0,622663255 | 0,13427256 | 4,6373083 | 3,53E-06 | 5,76E-05 | |
| ENSGALG00000008222 | 3620,89787 | 0,616961764 | 0,10986009 | 5,61588627 | 1,96E-08 | 4,66E-07 | SLC26A5 |
| ENSGALG00000011435 | 5310,48391 | 0,614694006 | 0,09295677 | 6,61268659 | 3,77E-11 | 1,29E-09 | |
| ENSGALG00000001728 | 1163,49973 | 0,610499448 | 0,14119355 | 4,32384786 | 1,53E-05 | 0,00021746 | NACC2 |
| ENSGALG00000026093 | 847,433551 | 0,608505556 | 0,17097819 | 3,55896599 | 0,00037232 | 0,0035661 | |
| ENSGALG00000019397 | 12768,6069 | 0,608262914 | 0,09652905 | 6,30134598 | 2,95E-10 | 9,30E-09 | CERK |
| ENSGALG00000004820 | 960,081767 | 0,608049873 | 0,1426222 | 4,26336072 | 2,01E-05 | 0,0002756 | FAM57A |
| ENSGALG00000026468 | 6537,04521 | 0,607177306 | 0,09886931 | 6,14121105 | 8,19E-10 | 2,38E-08 | CHSY1 |
| ENSGALG00000000795 | 1331,44868 | 0,605895026 | 0,13162727 | 4,60311177 | 4,16E-06 | 6,68E-05 | MFSD12 |
| ENSGALG00000017199 | 2734,428 | 0,605852032 | 0,15883889 | 3,81425504 | 0,00013659 | 0,00149751 | MAML2 |
| ENSGALG00000008903 | 2249,72466 | 0,604190258 | 0,15751307 | 3,83581031 | 0,00012515 | 0,00139302 | SSFA2 |
| ENSGALG00000011537 | 2507,94825 | 0,603878853 | 0,15538039 | 3,88645484 | 0,00010172 | 0,00116734 | PDE10A |
| ENSGALG00000006939 | 25023,5764 | 0,603844144 | 0,10825219 | 5,57812385 | 2,43E-08 | 5,73E-07 | PFKP |
| ENSGALG00000007434 | 2113,20997 | 0,599865863 | 0,15496377 | 3,87100707 | 0,00010839 | 0,0012348 | RHBDF1 |
| ENSGALG00000011488 | 1168,68478 | 0,598872747 | 0,17053454 | 3,51173862 | 0,00044519 | 0,00412506 | CMTM7 |
| ENSGALG00000016834 | 1257,10709 | 0,598292773 | 0,13769793 | 4,34496565 | 1,39E-05 | 0,00019982 | MCF2L |
| ENSGALG00000009988 | 2544,44093 | 0,593615724 | 0,112456 | 5,27864872 | 1,30E-07 | 2,70E-06 | ELOVL1 |
| ENSGALG00000011548 | 2906,3811 | 0,593504022 | 0,11422522 | 5,19591069 | 2,04E-07 | 4,08E-06 | PPP1R13B |
| ENSGALG00000009755 | 3320,85237 | 0,592776412 | 0,10287504 | 5,76210162 | 8,31E-09 | 2,06E-07 | LRIG3 |
| ENSGALG00000002709 | 2623,11643 | 0,59270728 | 0,12665173 | 4,67981968 | 2,87E-06 | 4,75E-05 | MYO1C |
| ENSGALG00000003642 | 1312,51148 | 0,590129639 | 0,16880923 | 3,49583753 | 0,00047258 | 0,0043244 | PDGF-A |
| ENSGALG00000008615 | 1132,69776 | 0,588449543 | 0,17235789 | 3,41411434 | 0,0006399 | 0,00559753 | |
| ENSGALG00000015096 | 745,308008 | 0,586609595 | 0,15533516 | 3,77641219 | 0,0001591 | 0,00171424 | SH3GL2 |
| ENSGALG00000011295 | 1419,82516 | 0,585889038 | 0,13140148 | 4,45877045 | 8,24E-06 | 0,00012483 | SOCS2 |
| ENSGALG00000014933 | 1201,23972 | 0,584450113 | 0,16167735 | 3,61491648 | 0,00030044 | 0,00298275 | HEXB |
| ENSGALG00000012979 | 9491,84596 | 0,583725131 | 0,11596876 | 5,0334689 | 4,82E-07 | 9,19E-06 | |
| ENSGALG00000001076 | 3535,03862 | 0,580936414 | 0,09643753 | 6,02396613 | 1,70E-09 | 4,70E-08 | LIMK1 |
| ENSGALG00000007242 | 1537,31241 | 0,576491214 | 0,1265805 | 4,55434468 | 5,25E-06 | 8,33E-05 | SLITRK4 |
| ENSGALG00000011045 | 4784,96919 | 0,574987094 | 0,09168796 | 6,27113007 | 3,58E-10 | 1,12E-08 | |
| ENSGALG00000020479 | 3396,9575 | 0,574556644 | 0,12483475 | 4,60253786 | 4,17E-06 | 6,68E-05 | |
| ENSGALG00000016276 | 897,324858 | 0,574168384 | 0,14748691 | 3,89301259 | 9,90E-05 | 0,00113831 | KHDRBS2 |
| ENSGALG00000012429 | 2058,77028 | 0,572869136 | 0,16484364 | 3,47522751 | 0,00051042 | 0,00461848 | BMP4 |
| ENSGALG00000019961 | 8549,80541 | 0,57282511 | 0,10454384 | 5,47928114 | 4,27E-08 | 9,70E-07 | SLC18B1 |
| ENSGALG00000009415 | 20919,8487 | 0,570748764 | 0,13417273 | 4,25383594 | 2,10E-05 | 0,00028634 | SMOC1 |
| ENSGALG00000011241 | 1131,08019 | 0,569997915 | 0,15948275 | 3,57404125 | 0,00035151 | 0,00340331 | RFTN1 |
| ENSGALG00000011708 | 16164,7225 | 0,569570289 | 0,12123193 | 4,69818723 | 2,62E-06 | 4,38E-05 | MYLK |
| ENSGALG00000006340 | 2676,99994 | 0,569237942 | 0,12657684 | 4,49717295 | 6,89E-06 | 0,00010687 | |
| ENSGALG00000009156 | 4770,93246 | 0,56553435 | 0,14428903 | 3,91945491 | 8,87E-05 | 0,00103754 | CDAN1 |
| ENSGALG00000012394 | 13820,7507 | 0,564570466 | 0,07862384 | 7,1806525 | 6,94E-13 | 3,00E-11 | NID2 |
| ENSGALG00000016350 | 2492,97331 | 0,563918458 | 0,11007286 | 5,12313797 | 3,00E-07 | 5,92E-06 | ARHGEF10 |
| ENSGALG00000008293 | 782,926286 | 0,562981441 | 0,15976711 | 3,52376315 | 0,00042546 | 0,00397768 | SH3PXD2A |
| ENSGALG00000006655 | 826,977917 | 0,562031156 | 0,15875615 | 3,54021654 | 0,0003998 | 0,00379131 | MAP3K13 |
| ENSGALG00000001664 | 2551,39665 | 0,561926589 | 0,13283895 | 4,23013411 | 2,34E-05 | 0,00031279 | PTPN9 |
| ENSGALG00000008621 | 9550,43432 | 0,561672993 | 0,11796857 | 4,76120892 | 1,92E-06 | 3,30E-05 | NRP2 |
| ENSGALG00000003122 | 1351,29404 | 0,561038511 | 0,1311515 | 4,27778952 | 1,89E-05 | 0,00026004 | ZFAND2A |
| ENSGALG00000015858 | 3586,9625 | 0,559546334 | 0,10146313 | 5,51477519 | 3,49E-08 | 8,06E-07 | DOPEY1 |
| ENSGALG00000013768 | 1494,29229 | 0,55942652 | 0,16037363 | 3,48827007 | 0,00048616 | 0,00442998 | PHACTR2 |
| ENSGALG00000015862 | 11794,0676 | 0,559415098 | 0,12796473 | 4,37163513 | 1,23E-05 | 0,00017979 | |
| ENSGALG00000027757 | 11293,5634 | 0,559044043 | 0,10878229 | 5,13910919 | 2,76E-07 | 5,46E-06 | ARL4A |

| | | | | | | | |
|---|---|---|---|---|---|---|---|
| ENSGALG00000017462 | 3715,9454 | 0,558530219 | 0,09687426 | 5,76551735 | 8,14E-09 | 2,02E-07 | FNIP1 |
| ENSGALG00000000388 | 11929,068 | 0,558245728 | 0,14596937 | 3,82440333 | 0,00013109 | 0,00144858 | LIN28A |
| ENSGALG00000007238 | 2073,56415 | 0,558183051 | 0,16989919 | 3,28537799 | 0,00101846 | 0,00826191 | |
| ENSGALG00000012852 | 29056,4718 | 0,557720187 | 0,10036391 | 5,55697956 | 2,74E-08 | 6,41E-07 | MKRN1 |
| ENSGALG00000001163 | 1519,07571 | 0,557286192 | 0,12356604 | 4,51002717 | 6,48E-06 | 0,00010148 | NDEL1 |
| ENSGALG00000016480 | 9465,84299 | 0,557041155 | 0,0924977 | 6,02221656 | 1,72E-09 | 4,71E-08 | SDC1 |
| ENSGALG00000028291 | 14773,7601 | 0,555590355 | 0,1099194 | 5,05452513 | 4,31E-07 | 8,29E-06 | |
| ENSGALG00000007845 | 1987,98275 | 0,553575037 | 0,11370336 | 4,86859011 | 1,12E-06 | 2,02E-05 | REL |
| ENSGALG00000014084 | 5460,83517 | 0,552765103 | 0,13748262 | 4,02061798 | 5,80E-05 | 0,00070908 | FGFR1OP2 |
| ENSGALG00000009433 | 8016,67263 | 0,551628222 | 0,14479495 | 3,80972014 | 0,00013912 | 0,00152391 | BAG3 |
| ENSGALG00000000680 | 7215,7939 | 0,550999971 | 0,1233941 | 4,46536719 | 7,99E-06 | 0,0001212 | |
| ENSGALG00000005433 | 1687,94239 | 0,550736363 | 0,11950908 | 4,60832215 | 4,06E-06 | 6,53E-05 | |
| ENSGALG00000002461 | 9112,46886 | 0,549280944 | 0,12891627 | 4,2607573 | 2,04E-05 | 0,00027853 | |
| ENSGALG00000007394 | 992,390556 | 0,546419843 | 0,14202976 | 3,84722077 | 0,00011947 | 0,00134267 | |
| ENSGALG00000015215 | 5874,51018 | 0,543882262 | 0,10412914 | 5,22315125 | 1,76E-07 | 3,55E-06 | MAPRE2 |
| ENSGALG00000010164 | 6967,47145 | 0,542805965 | 0,08759539 | 6,19674117 | 5,76E-10 | 1,73E-08 | SMARCA2 |
| ENSGALG00000017365 | 2743,73739 | 0,541410874 | 0,12416381 | 4,36045628 | 1,30E-05 | 0,00018835 | |
| ENSGALG00000014860 | 10032,8437 | 0,540418516 | 0,08281121 | 6,52591008 | 6,76E-11 | 2,25E-09 | YES1 |
| ENSGALG00000016781 | 15938,709 | 0,539823119 | 0,08390658 | 6,43362076 | 1,25E-10 | 4,08E-09 | MAP4K4 |
| ENSGALG00000016883 | 2972,3538 | 0,537559856 | 0,10976916 | 4,89718485 | 9,72E-07 | 1,77E-05 | DOCK9 |
| ENSGALG00000011449 | 1797,12078 | 0,537091274 | 0,13239866 | 4,05662179 | 4,98E-05 | 0,00062155 | |
| ENSGALG00000009855 | 6709,81331 | 0,536813312 | 0,11427025 | 4,69775213 | 2,63E-06 | 4,38E-05 | TBC1D9 |
| ENSGALG00000017185 | 9572,81105 | 0,536156218 | 0,13194516 | 4,06347779 | 4,83E-05 | 0,00060477 | TMEM123 |
| ENSGALG00000013406 | 658,679579 | 0,533541915 | 0,16232604 | 3,28685351 | 0,00101314 | 0,00822408 | EOGT |
| ENSGALG00000026237 | 8325,49172 | 0,532296374 | 0,09561992 | 5,56679355 | 2,59E-08 | 6,08E-07 | |
| ENSGALG00000000669 | 4577,06628 | 0,531970586 | 0,10145026 | 5,24365914 | 1,57E-07 | 3,21E-06 | ETV3 |
| ENSGALG00000015792 | 1821,33594 | 0,531949092 | 0,12531887 | 4,24476438 | 2,19E-05 | 0,00029688 | |
| ENSGALG00000000891 | 3401,91556 | 0,531658469 | 0,10937715 | 4,86078185 | 1,17E-06 | 2,10E-05 | NSUN5 |
| ENSGALG00000015112 | 8927,72556 | 0,528998124 | 0,10528495 | 5,02444216 | 5,05E-07 | 9,60E-06 | |
| ENSGALG00000002471 | 3937,07792 | 0,528654759 | 0,11018735 | 4,79778103 | 1,60E-06 | 2,82E-05 | |
| ENSGALG00000014834 | 2740,7337 | 0,527383014 | 0,1070106 | 4,92832478 | 8,29E-07 | 1,53E-05 | NCOA7 |
| ENSGALG00000027526 | 2310,64445 | 0,527340701 | 0,12420628 | 4,24568455 | 2,18E-05 | 0,00029599 | COL9A2 |
| ENSGALG00000017414 | 7193,90052 | 0,522128726 | 0,09330101 | 5,59617465 | 2,19E-08 | 5,19E-07 | JUP |
| ENSGALG00000002759 | 2898,01396 | 0,520324255 | 0,1531807 | 3,39680035 | 0,00068179 | 0,00589798 | PEAK1 |
| ENSGALG00000015193 | 2485,09904 | 0,518302858 | 0,13474249 | 3,84661781 | 0,00011976 | 0,00134477 | CCDC80 |
| ENSGALG00000002583 | 881,320118 | 0,51810067 | 0,15491432 | 3,34443367 | 0,00082451 | 0,00697347 | PIK3CD |
| ENSGALG00000006002 | 2311,21963 | 0,517539205 | 0,12381422 | 4,1799657 | 2,92E-05 | 0,00038309 | WHAMM |
| ENSGALG00000011162 | 1743,64653 | 0,516837911 | 0,12166248 | 4,24812899 | 2,16E-05 | 0,00029341 | CYP46A1 |
| ENSGALG00000000761 | 2548,35828 | 0,511636934 | 0,11935769 | 4,28658551 | 1,81E-05 | 0,00025079 | TSKU |
| ENSGALG00000008811 | 6319,41069 | 0,510104856 | 0,14038413 | 3,63363628 | 0,00027945 | 0,00279777 | SLC1A4 |
| ENSGALG00000007016 | 944,373297 | 0,509222424 | 0,14442833 | 3,52577944 | 0,00042224 | 0,00395936 | FAM169B |
| ENSGALG00000003668 | 1606,5427 | 0,508101808 | 0,12448719 | 4,08155899 | 4,47E-05 | 0,00056127 | |
| ENSGALG00000003948 | 4679,11028 | 0,503499536 | 0,09252751 | 5,44161999 | 5,28E-08 | 1,19E-06 | ALAS1 |
| ENSGALG00000015458 | 8275,30923 | 0,50234495 | 0,11154476 | 4,50352792 | 6,68E-06 | 0,00010424 | ILDR2 |
| ENSGALG00000006569 | 8235,42763 | 0,500306434 | 0,1289293 | 3,88047121 | 0,00010425 | 0,00119425 | |
| ENSGALG00000012449 | 8575,6897 | 0,498545559 | 0,13392532 | 3,72256398 | 0,00019721 | 0,00206435 | MBD5 |
| ENSGALG00000004710 | 3763,43768 | 0,498075539 | 0,10011571 | 4,97499881 | 6,52E-07 | 1,22E-05 | TMOD3 |
| ENSGALG00000017388 | 6631,01193 | 0,49802444 | 0,09554361 | 5,21253541 | 1,86E-07 | 3,75E-06 | |
| ENSGALG00000004909 | 2041,96193 | 0,497587671 | 0,13255196 | 3,75390642 | 0,0001741 | 0,00185173 | |
| ENSGALG00000005095 | 2394,73163 | 0,496064325 | 0,13816886 | 3,59027581 | 0,00033033 | 0,00322566 | SLC25A25 |
| ENSGALG00000003770 | 2099,85738 | 0,492649703 | 0,12147281 | 4,05563753 | 5,00E-05 | 0,00062331 | ANXA2 |
| ENSGALG00000001007 | 23689,0314 | 0,492162491 | 0,08170528 | 6,0236317 | 1,71E-09 | 4,70E-08 | |
| ENSGALG00000027925 | 2899,91937 | 0,491701601 | 0,15107329 | 3,25472237 | 0,00113503 | 0,0090607 | |
| ENSGALG00000011054 | 3850,38645 | 0,491324799 | 0,13238005 | 3,71147149 | 0,00020606 | 0,00214725 | |
| ENSGALG00000009503 | 5216,27727 | 0,49017284 | 0,10922838 | 4,48759608 | 7,20E-06 | 0,00011097 | RAB3GAP2 |
| ENSGALG00000000936 | 1155,34155 | 0,490067343 | 0,13552456 | 3,61607777 | 0,0002991 | 0,00297412 | |
| ENSGALG00000011539 | 2399,98235 | 0,490040841 | 0,15085476 | 3,24842803 | 0,00116045 | 0,00921652 | |
| ENSGALG00000016768 | 7017,45831 | 0,489550679 | 0,0912379 | 5,3656504 | 8,07E-08 | 1,73E-06 | LONRF2 |
| ENSGALG00000016839 | 2941,70522 | 0,488322587 | 0,14416141 | 3,38733229 | 0,00070576 | 0,00607593 | |
| ENSGALG00000001962 | 47390,8257 | 0,488180846 | 0,07927072 | 6,15840034 | 7,35E-10 | 2,15E-08 | PTBP1 |
| ENSGALG00000008321 | 2140,81231 | 0,487472572 | 0,11701276 | 4,16597802 | 3,10E-05 | 0,00040481 | RNPEPL1 |
| ENSGALG00000026231 | 4317,60577 | 0,487086101 | 0,13624774 | 3,57500308 | 0,00035022 | 0,00339345 | ARID3A |
| ENSGALG00000002316 | 5849,0452 | 0,486190455 | 0,10164247 | 4,78333983 | 1,72E-06 | 3,00E-05 | NUBP2 |
| ENSGALG00000005727 | 11457,1676 | 0,485528575 | 0,09017413 | 5,38434468 | 7,27E-08 | 1,58E-06 | |

| | | | | | | | |
|---|---|---|---|---|---|---|---|
| ENSGALG00000013661 | 3653,88906 | 0,485373842 | 0,14009384 | 3,4646338 | 0,00053095 | 0,00476108 | MICU3 |
| ENSGALG00000006237 | 12165,443 | 0,48464383 | 0,09005848 | 5,38143497 | 7,39E-08 | 1,60E-06 | PKN2 |
| ENSGALG00000006121 | 10037,2715 | 0,483017989 | 0,10132916 | 4,76682134 | 1,87E-06 | 3,22E-05 | PIK3C2A |
| ENSGALG00000003835 | 26017,6944 | 0,482616746 | 0,0853296 | 5,65591273 | 1,55E-08 | 3,75E-07 | |
| ENSGALG00000002219 | 2154,95524 | 0,481514535 | 0,11726813 | 4,10609869 | 4,02E-05 | 0,00051051 | WBP2 |
| ENSGALG00000014719 | 2318,70089 | 0,480107075 | 0,11701682 | 4,10288958 | 4,08E-05 | 0,00051712 | |
| ENSGALG00000007271 | 2336,70848 | 0,480045702 | 0,10812496 | 4,43973058 | 9,01E-06 | 0,00013558 | |
| ENSGALG00000028128 | 2623,01722 | 0,475145623 | 0,1057506 | 4,49307716 | 7,02E-06 | 0,00010841 | |
| ENSGALG00000002830 | 1989,1104 | 0,475072145 | 0,13713011 | 3,46438965 | 0,00053144 | 0,00476199 | NUDT3 |
| ENSGALG00000016283 | 1439,58733 | 0,474889919 | 0,14244125 | 3,33393539 | 0,00085627 | 0,00717416 | BEND6 |
| ENSGALG00000014953 | 9336,02158 | 0,474792932 | 0,1011178 | 4,69544373 | 2,66E-06 | 4,43E-05 | RWDD1 |
| ENSGALG00000014873 | 10464,2969 | 0,474224225 | 0,08830699 | 5,37017768 | 7,87E-08 | 1,69E-06 | GJA1 |
| ENSGALG00000008181 | 4533,21462 | 0,473531005 | 0,11780023 | 4,01978013 | 5,83E-05 | 0,00071091 | MADD |
| ENSGALG00000009532 | 10930,7149 | 0,469457498 | 0,09307351 | 5,0439434 | 4,56E-07 | 8,72E-06 | GXYLT1 |
| ENSGALG00000009086 | 21565,5654 | 0,468835158 | 0,07859861 | 5,96492909 | 2,45E-09 | 6,53E-08 | EHD3 |
| ENSGALG00000016808 | 38690,2199 | 0,46860233 | 0,08636591 | 5,42577901 | 5,77E-08 | 1,28E-06 | RANBP2 |
| ENSGALG00000012033 | 1874,0337 | 0,467396297 | 0,12741413 | 3,66832386 | 0,00024415 | 0,00249111 | SGSM3 |
| ENSGALG00000005422 | 1476,40233 | 0,465795544 | 0,13673042 | 3,40667103 | 0,0006576 | 0,00570455 | PDLIM1 |
| ENSGALG00000000874 | 1565,53262 | 0,465401247 | 0,12194279 | 3,8165539 | 0,00013533 | 0,00148493 | |
| ENSGALG00000003113 | 3657,12064 | 0,464288641 | 0,09568682 | 4,85216929 | 1,22E-06 | 2,19E-05 | RGS14 |
| ENSGALG00000002726 | 8476,00798 | 0,464112357 | 0,12109197 | 3,83272605 | 0,00012673 | 0,00140434 | KIF1B |
| ENSGALG00000025747 | 1304,00667 | 0,463404585 | 0,13117609 | 3,5326909 | 0,00041135 | 0,00388056 | NQO2 |
| ENSGALG00000015036 | 6732,47434 | 0,461955249 | 0,11674428 | 3,95698403 | 7,59E-05 | 0,00090425 | RIC1 |
| ENSGALG00000016495 | 2231,85951 | 0,461211494 | 0,11447108 | 4,02906563 | 5,60E-05 | 0,00068541 | KLHL29 |
| ENSGALG00000000890 | 12926,8583 | 0,459656521 | 0,08135785 | 5,64981129 | 1,61E-08 | 3,87E-07 | |
| ENSGALG00000004144 | 25688,7242 | 0,459129046 | 0,08737901 | 5,25445453 | 1,48E-07 | 3,05E-06 | |
| ENSGALG00000009850 | 10767,8502 | 0,458569963 | 0,088576 | 5,17713554 | 2,25E-07 | 4,48E-06 | NOVA1 |
| ENSGALG00000014688 | 8715,47092 | 0,458090231 | 0,11541233 | 3,96916206 | 7,21E-05 | 0,00086337 | ZYX |
| ENSGALG00000013480 | 4689,04979 | 0,457972294 | 0,11338344 | 4,03914621 | 5,36E-05 | 0,00066049 | |
| ENSGALG00000015940 | 2526,66559 | 0,457737572 | 0,10654486 | 4,2961959 | 1,74E-05 | 0,00024177 | |
| ENSGALG00000005134 | 3403,87437 | 0,45766228 | 0,10876996 | 4,20761648 | 2,58E-05 | 0,00034271 | ZSWIM8 |
| ENSGALG00000003295 | 2431,84104 | 0,455390197 | 0,14052 | 3,24075001 | 0,00119216 | 0,00942648 | |
| ENSGALG00000011286 | 5006,91718 | 0,454714572 | 0,0910514 | 4,99404243 | 5,91E-07 | 1,12E-05 | UBE2E1 |
| ENSGALG00000010361 | 8260,44964 | 0,453176333 | 0,09954173 | 4,55262667 | 5,30E-06 | 8,39E-05 | |
| ENSGALG00000012380 | 1446,14742 | 0,453021582 | 0,1253689 | 3,61350847 | 0,00030208 | 0,00299425 | SPOPL |
| ENSGALG00000002672 | 3384,73075 | 0,451332604 | 0,11773512 | 3,83345767 | 0,00012635 | 0,00140246 | |
| ENSGALG00000008521 | 14688,0623 | 0,447794364 | 0,09286454 | 4,82201686 | 1,42E-06 | 2,51E-05 | |
| ENSGALG00000014371 | 4193,46245 | 0,447550868 | 0,10744176 | 4,16552078 | 3,11E-05 | 0,0004052 | LGALS8 |
| ENSGALG00000006947 | 14379,0645 | 0,446651103 | 0,08611381 | 5,18675352 | 2,14E-07 | 4,26E-06 | BDH1 |
| ENSGALG00000001596 | 11592,0471 | 0,445911724 | 0,08108031 | 5,49963016 | 3,81E-08 | 8,68E-07 | MLLT1 |
| ENSGALG00000016746 | 1667,79177 | 0,445777139 | 0,13374077 | 3,33314327 | 0,00085871 | 0,0071898 | |
| ENSGALG00000006313 | 1456,17134 | 0,445448611 | 0,12850341 | 3,46643413 | 0,00052741 | 0,0047361 | |
| ENSGALG00000003263 | 4582,86968 | 0,445240965 | 0,0980725 | 4,53991643 | 5,63E-06 | 8,89E-05 | |
| ENSGALG00000009300 | 5731,29585 | 0,444814977 | 0,09141833 | 4,86570901 | 1,14E-06 | 2,05E-05 | NUMB |
| ENSGALG00000003862 | 1841,95984 | 0,443615862 | 0,11893475 | 3,72990962 | 0,00019155 | 0,0020186 | |
| ENSGALG00000005505 | 1365,18076 | 0,442779592 | 0,12653606 | 3,49923634 | 0,00046659 | 0,0042759 | PIK3CB |
| ENSGALG00000005785 | 2692,43001 | 0,442398124 | 0,13426574 | 3,29494421 | 0,00098441 | 0,00803597 | |
| ENSGALG00000004169 | 13236,4427 | 0,441293938 | 0,09199916 | 4,79671711 | 1,61E-06 | 2,83E-05 | |
| ENSGALG00000012462 | 8197,06854 | 0,439936793 | 0,10637709 | 4,13563485 | 3,54E-05 | 0,00045554 | KIF5C |
| ENSGALG00000012093 | 1956,24055 | 0,439445143 | 0,11904646 | 3,69137507 | 0,00022304 | 0,00230204 | |
| ENSGALG00000001957 | 8367,03127 | 0,438219772 | 0,11967372 | 3,66178782 | 0,00025046 | 0,0025415 | |
| ENSGALG00000012799 | 2784,84231 | 0,438081252 | 0,12516354 | 3,5000707 | 0,00046513 | 0,00426567 | RREB1 |
| ENSGALG00000013861 | 1114,81045 | 0,437973226 | 0,13471835 | 3,25102872 | 0,00114988 | 0,00916171 | TNFAIP3 |
| ENSGALG00000014419 | 9771,32739 | 0,437804845 | 0,11330954 | 3,86379513 | 0,00011164 | 0,00126494 | |
| ENSGALG00000026184 | 2048,29159 | 0,437296545 | 0,12629201 | 3,4625829 | 0,00053502 | 0,00478722 | |
| ENSGALG00000010030 | 8265,21081 | 0,435973553 | 0,09014275 | 4,83647923 | 1,32E-06 | 2,35E-05 | |
| ENSGALG00000011464 | 75594,5083 | 0,435819807 | 0,08481234 | 5,13863704 | 2,77E-07 | 5,46E-06 | EIF5 |
| ENSGALG00000011643 | 21083,8275 | 0,435706974 | 0,11836787 | 3,68095649 | 0,00023236 | 0,00238441 | UTP20 |
| ENSGALG00000011016 | 2374,56332 | 0,434866192 | 0,12967523 | 3,35350229 | 0,00079796 | 0,00676722 | PGM1 |
| ENSGALG00000011707 | 2790,34805 | 0,43414341 | 0,11909866 | 3,64524173 | 0,00026714 | 0,00268407 | |
| ENSGALG00000004089 | 7953,41422 | 0,433449007 | 0,08322806 | 5,20796739 | 1,91E-07 | 3,84E-06 | |
| ENSGALG00000014686 | 1896,32829 | 0,43250268 | 0,11603822 | 3,72724338 | 0,00019359 | 0,00203322 | FBN2 |
| ENSGALG00000012414 | 1673,15959 | 0,431565773 | 0,12254766 | 3,52161578 | 0,00042893 | 0,00400704 | GNPNAT1 |
| ENSGALG00000009534 | 2593,70172 | 0,430698068 | 0,10888319 | 3,95559732 | 7,63E-05 | 0,00090864 | |

| | | | | | | | |
|---|---|---|---|---|---|---|---|
| ENSGALG00000001376 | 6656,04949 | 0,43069552 | 0,08629025 | 4,99124172 | 6,00E-07 | 1,13E-05 | ARID3B |
| ENSGALG00000016751 | 17995,0581 | 0,429916343 | 0,11291663 | 3,80737831 | 0,00014045 | 0,00153438 | |
| ENSGALG00000007659 | 32471,1723 | 0,429907921 | 0,0785478 | 5,47320101 | 4,42E-08 | 1,00E-06 | HYOU1 |
| ENSGALG00000013134 | 6778,47765 | 0,429746745 | 0,09606485 | 4,47350681 | 7,69E-06 | 0,00011753 | |
| ENSGALG00000010967 | 19058,4836 | 0,429437085 | 0,07951282 | 5,4008533 | 6,63E-08 | 1,45E-06 | |
| ENSGALG00000000243 | 3939,58343 | 0,429169147 | 0,10346666 | 4,14789781 | 3,36E-05 | 0,0004354 | RDM1 |
| ENSGALG00000011682 | 2443,78682 | 0,428788464 | 0,10784939 | 3,97580809 | 7,01E-05 | 0,00084122 | CNOT4 |
| ENSGALG00000014784 | 3810,58741 | 0,427987842 | 0,11343575 | 3,77295391 | 0,00016133 | 0,00173519 | MAST4 |
| ENSGALG00000004638 | 3143,25207 | 0,427666392 | 0,1185082 | 3,60874933 | 0,00030768 | 0,00304007 | |
| ENSGALG00000008128 | 9640,95815 | 0,426089907 | 0,09019502 | 4,72409566 | 2,31E-06 | 3,91E-05 | |
| ENSGALG00000010975 | 6422,3758 | 0,425801788 | 0,09591443 | 4,43939214 | 9,02E-06 | 0,00013563 | TMTC2 |
| ENSGALG00000004184 | 14964,7344 | 0,42382749 | 0,07954333 | 5,32825936 | 9,92E-08 | 2,10E-06 | SPARC |
| ENSGALG00000009298 | 10491,8514 | 0,42324599 | 0,12277761 | 3,44725699 | 0,00056631 | 0,00503123 | TNIK |
| ENSGALG00000015033 | 5942,94152 | 0,422029877 | 0,11922305 | 3,5398347 | 0,00040038 | 0,00379131 | REV3L |
| ENSGALG00000008009 | 4573,71765 | 0,421760643 | 0,09376655 | 4,49798603 | 6,86E-06 | 0,0001066 | BCAP29 |
| ENSGALG00000001546 | 5321,72397 | 0,420857037 | 0,11791501 | 3,56915579 | 0,00035813 | 0,00345138 | ZZEF1 |
| ENSGALG00000009570 | 4990,17008 | 0,419915848 | 0,09759573 | 4,30260451 | 1,69E-05 | 0,00023646 | ATP6V1D |
| ENSGALG00000011246 | 4677,48231 | 0,419525625 | 0,09763 | 4,29709732 | 1,73E-05 | 0,00024132 | |
| ENSGALG00000015942 | 2654,62871 | 0,419245968 | 0,10972783 | 3,82078074 | 0,00013303 | 0,00146485 | |
| ENSGALG00000004531 | 6608,4459 | 0,419189684 | 0,08983065 | 4,66644405 | 3,06E-06 | 5,03E-05 | PLOD1 |
| ENSGALG00000012251 | 1948,91488 | 0,417627661 | 0,12070121 | 3,46001212 | 0,00054015 | 0,00482625 | |
| ENSGALG00000012739 | 5567,17218 | 0,417000863 | 0,09837079 | 4,23907217 | 2,24E-05 | 0,0003031 | HIVEP1 |
| ENSGALG00000020805 | 5785,80923 | 0,416086722 | 0,09259199 | 4,49376567 | 7,00E-06 | 0,0001082 | LRRC8D |
| ENSGALG00000023806 | 3007,51007 | 0,415743003 | 0,11577931 | 3,59082288 | 0,00032964 | 0,00322241 | |
| ENSGALG00000001443 | 21339,3728 | 0,41409085 | 0,11660287 | 3,5512922 | 0,00038334 | 0,00365771 | |
| ENSGALG00000003181 | 2537,40381 | 0,412959133 | 0,11077844 | 3,72779349 | 0,00019316 | 0,00203049 | SURF2 |
| ENSGALG00000005723 | 2106,65447 | 0,412949734 | 0,1250895 | 3,3012343 | 0,0009626 | 0,00789588 | |
| ENSGALG00000007141 | 2241,98683 | 0,412800257 | 0,11510085 | 3,58642238 | 0,00033525 | 0,00327113 | LRRK1 |
| ENSGALG00000005660 | 9409,08057 | 0,411134823 | 0,12248124 | 3,35671682 | 0,00078874 | 0,00669812 | NF1 |
| ENSGALG00000006863 | 4226,48137 | 0,410961417 | 0,09352764 | 4,39401049 | 1,11E-05 | 0,00016357 | |
| ENSGALG00000005249 | 4537,35945 | 0,410952179 | 0,10151336 | 4,04825713 | 5,16E-05 | 0,00063971 | BICD2 |
| ENSGALG00000004644 | 3446,25125 | 0,409508513 | 0,12409834 | 3,2998709 | 0,00096729 | 0,00792395 | |
| ENSGALG00000001194 | 18631,7262 | 0,408514612 | 0,08737937 | 4,67518369 | 2,94E-06 | 4,84E-05 | |
| ENSGALG00000005962 | 14577,1851 | 0,407910696 | 0,09612843 | 4,24339276 | 2,20E-05 | 0,00029838 | |
| ENSGALG00000028637 | 3638,50777 | 0,407745497 | 0,1057417 | 3,85605195 | 0,00011523 | 0,0013033 | |
| ENSGALG00000005796 | 13049,6305 | 0,407045166 | 0,09023433 | 4,51097882 | 6,45E-06 | 0,00010115 | XBP1 |
| ENSGALG00000011254 | 1903,62474 | 0,406188469 | 0,11859058 | 3,42513271 | 0,0006145 | 0,00539801 | SATB1 |
| ENSGALG00000023451 | 10229,4738 | 0,405630741 | 0,10390284 | 3,90394266 | 9,46E-05 | 0,00109512 | TRPM7 |
| ENSGALG00000002399 | 3646,29569 | 0,405489945 | 0,09701983 | 4,17945444 | 2,92E-05 | 0,00038355 | |
| ENSGALG00000012814 | 2642,59396 | 0,404889777 | 0,11881733 | 3,40766613 | 0,00065521 | 0,00569168 | PXDC1 |
| ENSGALG00000028267 | 14506,7903 | 0,403950335 | 0,07758347 | 5,2066546 | 1,92E-07 | 3,86E-06 | |
| ENSGALG00000017085 | 3891,4285 | 0,403507696 | 0,11323732 | 3,56338096 | 0,00036611 | 0,00352281 | SLC7A1 |
| ENSGALG00000009748 | 12590,9107 | 0,403044512 | 0,10500934 | 3,83817788 | 0,00012395 | 0,001384 | ASNS |
| ENSGALG00000010780 | 2934,00779 | 0,401358719 | 0,11055482 | 3,63040448 | 0,00028298 | 0,00282729 | |
| ENSGALG00000014854 | 14435,0938 | 0,401311952 | 0,08622832 | 4,65406195 | 3,25E-06 | 5,33E-05 | |
| ENSGALG00000011101 | 5371,2295 | 0,40109735 | 0,0891436 | 4,49945224 | 6,81E-06 | 0,00010613 | URB2 |
| ENSGALG00000008855 | 8350,52016 | 0,400902672 | 0,10842426 | 3,69753673 | 0,0002177 | 0,00225977 | UGP2 |
| ENSGALG00000026850 | 2047,38224 | 0,400065314 | 0,11989409 | 3,33682262 | 0,00084742 | 0,00711913 | |
| ENSGALG00000013830 | 3806,14371 | 0,399422 | 0,11396388 | 3,50481227 | 0,00045693 | 0,00421349 | PRELID3A |
| ENSGALG00000015459 | 1707,50104 | 0,399319665 | 0,11963093 | 3,3379299 | 0,00084405 | 0,00709694 | |
| ENSGALG00000005582 | 5939,79295 | 0,398284507 | 0,09131089 | 4,36185098 | 1,29E-05 | 0,00018737 | KLHL18 |
| ENSGALG00000008195 | 5223,77429 | 0,394418985 | 0,08931317 | 4,41613483 | 1,00E-05 | 0,00014928 | NPHP1 |
| ENSGALG00000012404 | 2166,93397 | 0,394285725 | 0,11188253 | 3,52410442 | 0,00042492 | 0,00397553 | ERO1A |
| ENSGALG00000012342 | 4988,06672 | 0,391855224 | 0,11601632 | 3,37758709 | 0,00073125 | 0,00626949 | SASH1 |
| ENSGALG00000011542 | 3470,24914 | 0,391703901 | 0,11096577 | 3,52995262 | 0,00041563 | 0,00391209 | |
| ENSGALG00000028203 | 5589,34557 | 0,39127665 | 0,10019534 | 3,90513811 | 9,42E-05 | 0,00109174 | SPRED1 |
| ENSGALG00000017283 | 2269,10202 | 0,390659364 | 0,1093337 | 3,57309192 | 0,00035279 | 0,00341304 | CCND2 |
| ENSGALG00000008676 | 3130,52664 | 0,390658638 | 0,11592196 | 3,37001398 | 0,00075164 | 0,00643113 | RRBP1 |
| ENSGALG00000002763 | 7822,98963 | 0,38996775 | 0,09784285 | 3,98565421 | 6,73E-05 | 0,00081175 | FRZB |
| ENSGALG00000012280 | 4546,76644 | 0,388208613 | 0,09700097 | 4,0021108 | 6,28E-05 | 0,00075985 | |
| ENSGALG00000014902 | 5801,5163 | 0,387988657 | 0,11314858 | 3,42901912 | 0,00060577 | 0,00534005 | |
| ENSGALG00000016779 | 3110,09587 | 0,387851139 | 0,10566697 | 3,67050494 | 0,00024207 | 0,00247392 | RNF149 |
| ENSGALG00000026822 | 5680,41299 | 0,384087371 | 0,09933111 | 3,86673784 | 0,0001103 | 0,00125318 | WDR81 |
| ENSGALG00000008708 | 8378,00616 | 0,384060223 | 0,08161607 | 4,70569382 | 2,53E-06 | 4,24E-05 | |

| | | | | | | | |
|---|---|---|---|---|---|---|---|
| ENSGALG00000004200 | 10300,1977 | 0,383702067 | 0,10034391 | 3,82386986 | 0,00013137 | 0,00145044 | DPP9 |
| ENSGALG00000019077 | 12866,6527 | 0,383662275 | 0,10944193 | 3,50562403 | 0,00045554 | 0,00420541 | |
| ENSGALG00000008542 | 22218,9525 | 0,383345327 | 0,115816 | 3,3099513 | 0,00093312 | 0,00768428 | ADD3 |
| ENSGALG00000015765 | 12660,3185 | 0,383191974 | 0,11316078 | 3,38626134 | 0,00070852 | 0,0060955 | |
| ENSGALG00000001551 | 13003,1544 | 0,382951597 | 0,10019028 | 3,82224294 | 0,00013224 | 0,00145747 | CCNL2 |
| ENSGALG00000012608 | 2396,75353 | 0,382010791 | 0,11550101 | 3,30742386 | 0,00094158 | 0,00774886 | DAPK1 |
| ENSGALG00000008520 | 50311,2923 | 0,381829873 | 0,07410654 | 5,15244531 | 2,57E-07 | 5,10E-06 | EIF4G1 |
| ENSGALG00000000464 | 4473,94851 | 0,378042232 | 0,09693774 | 3,89984576 | 9,63E-05 | 0,00111074 | |
| ENSGALG00000013628 | 17850,2484 | 0,377744473 | 0,0965373 | 3,91293791 | 9,12E-05 | 0,00106213 | C6orf62 |
| ENSGALG00000004509 | 60307,8172 | 0,377502687 | 0,10213082 | 3,69626592 | 0,00021879 | 0,00226752 | |
| ENSGALG00000000385 | 3155,8316 | 0,377133979 | 0,11668273 | 3,23213188 | 0,0012287 | 0,00967875 | |
| ENSGALG00000008240 | 2335,47856 | 0,376593038 | 0,112924 | 3,33492474 | 0,00085322 | 0,00715348 | CFLAR |
| ENSGALG00000016457 | 20077,635 | 0,37591995 | 0,08487595 | 4,42905171 | 9,46E-06 | 0,00014162 | TRIB2 |
| ENSGALG00000004002 | 12560,3704 | 0,375282513 | 0,09982845 | 3,75927432 | 0,00017041 | 0,00181569 | LYPLA2 |
| ENSGALG00000004384 | 6339,40502 | 0,37272222 | 0,09191076 | 4,05526203 | 5,01E-05 | 0,00062331 | |
| ENSGALG00000014213 | 4846,8151 | 0,372421493 | 0,1124273 | 3,3125538 | 0,00092448 | 0,00763324 | PRR5 |
| ENSGALG00000017335 | 8042,45001 | 0,370836945 | 0,08378084 | 4,42627398 | 9,59E-06 | 0,00014329 | |
| ENSGALG00000013605 | 5142,50116 | 0,369173077 | 0,09448513 | 3,90720816 | 9,34E-05 | 0,00108344 | LRRC16A |
| ENSGALG00000009428 | 6757,25752 | 0,369028406 | 0,11310884 | 3,26259556 | 0,00110397 | 0,00886364 | SUSD6 |
| ENSGALG00000003180 | 15013,9328 | 0,367625634 | 0,08522459 | 4,31360964 | 1,61E-05 | 0,00022599 | |
| ENSGALG00000008414 | 11352,9971 | 0,366746111 | 0,095194 | 3,85261804 | 0,00011686 | 0,00131933 | DVL3 |
| ENSGALG00000003748 | 6555,91118 | 0,36621302 | 0,08463061 | 4,32719328 | 1,51E-05 | 0,00021492 | ANO5 |
| ENSGALG00000006336 | 6628,48994 | 0,365567709 | 0,10726355 | 3,4081263 | 0,00065411 | 0,00569 | |
| ENSGALG00000005845 | 8308,86511 | 0,364474801 | 0,09382855 | 3,88447647 | 0,00010255 | 0,00117581 | SLC7A5 |
| ENSGALG00000011288 | 4443,75245 | 0,36304574 | 0,0964958 | 3,76229586 | 0,00016836 | 0,00179695 | DACT2 |
| ENSGALG00000016736 | 21468,1233 | 0,362152411 | 0,10050997 | 3,60314896 | 0,00031439 | 0,00309657 | CYFIP1 |
| ENSGALG00000004696 | 6019,01081 | 0,360778491 | 0,09380179 | 3,84617901 | 0,00011997 | 0,00134597 | LEO1 |
| ENSGALG00000009562 | 5205,50613 | 0,359101036 | 0,11011889 | 3,26103035 | 0,00111008 | 0,00890653 | |
| ENSGALG00000007145 | 42543,9665 | 0,358001987 | 0,07128079 | 5,02241869 | 5,10E-07 | 9,69E-06 | ITGB1 |
| ENSGALG00000023857 | 2295,50244 | 0,357227793 | 0,10931817 | 3,26778052 | 0,00108394 | 0,00871965 | |
| ENSGALG00000010603 | 12138,5477 | 0,356862546 | 0,08271823 | 4,31419487 | 1,60E-05 | 0,00022565 | |
| ENSGALG00000007514 | 7096,59375 | 0,355063614 | 0,08859236 | 4,00783581 | 6,13E-05 | 0,00074492 | |
| ENSGALG00000003563 | 12215,1985 | 0,354826794 | 0,08437076 | 4,20556587 | 2,60E-05 | 0,0003451 | YARS |
| ENSGALG00000009946 | 16182,3882 | 0,351940163 | 0,09459123 | 3,72064246 | 0,00019872 | 0,00207838 | HECTD1 |
| ENSGALG00000017112 | 5413,39298 | 0,351627272 | 0,10588126 | 3,32095857 | 0,00089709 | 0,00746122 | MTMR6 |
| ENSGALG00000005160 | 8835,93843 | 0,351320099 | 0,09529673 | 3,68659118 | 0,00022728 | 0,00234186 | VMP1 |
| ENSGALG00000002954 | 6375,29438 | 0,350618821 | 0,09690427 | 3,61819794 | 0,00029666 | 0,00295222 | MBTD1 |
| ENSGALG00000008099 | 7103,82231 | 0,350057141 | 0,08535874 | 4,10101087 | 4,11E-05 | 0,00052028 | |
| ENSGALG00000005678 | 19719,4298 | 0,349721909 | 0,0904842 | 3,8650054 | 0,00011109 | 0,00125982 | FLNB |
| ENSGALG00000014795 | 5131,9149 | 0,349663833 | 0,09082989 | 3,84965603 | 0,00011828 | 0,00133083 | TGIF1 |
| ENSGALG00000015013 | 15895,2683 | 0,348772486 | 0,09941333 | 3,50830713 | 0,00045097 | 0,00416629 | MSX1 |
| ENSGALG00000011047 | 7708,25037 | 0,346858574 | 0,09143673 | 3,79342699 | 0,00014858 | 0,00161338 | LEPROT |
| ENSGALG00000001749 | 4850,90171 | 0,34679817 | 0,10645528 | 3,25768863 | 0,00112324 | 0,00897799 | ACSBG2 |
| ENSGALG00000015667 | 5950,72909 | 0,346421134 | 0,09935991 | 3,48652816 | 0,00048933 | 0,00445169 | |
| ENSGALG00000003427 | 8942,13088 | 0,346212827 | 0,10652898 | 3,24994036 | 0,00115429 | 0,00918514 | USP3 |
| ENSGALG00000009540 | 6305,96328 | 0,344720194 | 0,10105941 | 3,41106481 | 0,0006471 | 0,00564079 | YAF2 |
| ENSGALG00000006437 | 3815,5669 | 0,344612756 | 0,0964691 | 3,57226061 | 0,00035391 | 0,00342125 | |
| ENSGALG00000001003 | 4334,6017 | 0,344161191 | 0,10407482 | 3,30686318 | 0,00094347 | 0,00775418 | SEPN1 |
| ENSGALG00000028726 | 5513,31502 | 0,343861226 | 0,09087882 | 3,7837332 | 0,00015449 | 0,00167032 | |
| ENSGALG00000014139 | 7333,64409 | 0,34378877 | 0,09116278 | 3,7711527 | 0,0001625 | 0,00174327 | |
| ENSGALG00000007234 | 4391,88237 | 0,343143822 | 0,09912131 | 3,46185737 | 0,00053646 | 0,00479671 | CLCN5 |
| ENSGALG00000002521 | 2909,63624 | 0,341377701 | 0,10552536 | 3,23502994 | 0,0012163 | 0,00959312 | |
| ENSGALG00000025884 | 6496,58774 | 0,339208538 | 0,0928033 | 3,65513456 | 0,00025705 | 0,00260356 | NABP1 |
| ENSGALG00000000546 | 3505,59123 | 0,338887883 | 0,09640887 | 3,51511115 | 0,00043957 | 0,00408392 | |
| ENSGALG00000013677 | 5281,75302 | 0,33880019 | 0,09391468 | 3,60753163 | 0,00030912 | 0,00305196 | |
| ENSGALG00000026349 | 30008,024 | 0,337656858 | 0,07971202 | 4,23595916 | 2,28E-05 | 0,00030611 | |
| ENSGALG00000012202 | 7460,55254 | 0,337252452 | 0,08638462 | 3,90407991 | 9,46E-05 | 0,00109512 | |
| ENSGALG00000010976 | 5122,33312 | 0,337203087 | 0,10038342 | 3,35915108 | 0,00078182 | 0,00665294 | CCDC126 |
| ENSGALG00000006417 | 9187,43681 | 0,336436699 | 0,08945097 | 3,76112978 | 0,00016915 | 0,00180381 | SEC24A |
| ENSGALG00000006975 | 7222,31417 | 0,33439022 | 0,08487768 | 3,93967199 | 8,16E-05 | 0,00096378 | |
| ENSGALG00000002362 | 7456,91772 | 0,333268709 | 0,08848366 | 3,76644362 | 0,00016559 | 0,00177191 | MANF |
| ENSGALG00000014370 | 4742,98825 | 0,332141176 | 0,09950672 | 3,33787686 | 0,00084421 | 0,00709694 | |
| ENSGALG00000003767 | 4420,56627 | 0,331888441 | 0,0963142 | 3,44589313 | 0,00056918 | 0,00504594 | NKD1 |
| ENSGALG00000006256 | 7680,29257 | 0,33180927 | 0,08270052 | 4,01217858 | 6,02E-05 | 0,00073206 | |

| | | | | | | | |
|---|---|---|---|---|---|---|---|
| ENSGALG00000002548 | 7217,2063 | 0,330998688 | 0,08817899 | 3,75371385 | 0,00017423 | 0,00185173 | TLN1 |
| ENSGALG00000004691 | 5665,22107 | 0,330681288 | 0,10090233 | 3,27724134 | 0,00104827 | 0,00847626 | |
| ENSGALG00000005139 | 48915,1737 | 0,327662098 | 0,07590851 | 4,31653954 | 1,58E-05 | 0,00022428 | CLTC |
| ENSGALG00000011055 | 8298,54873 | 0,326999676 | 0,09831999 | 3,32587164 | 0,00088143 | 0,0073456 | |
| ENSGALG00000016630 | 13012,9547 | 0,325439015 | 0,08809758 | 3,6940745 | 0,00022069 | 0,00228149 | EXTL3 |
| ENSGALG00000011630 | 13213,6043 | 0,324450144 | 0,08177951 | 3,96737719 | 7,27E-05 | 0,00086903 | GLI2 |
| ENSGALG00000015049 | 15025,2928 | 0,323309666 | 0,07842419 | 4,12257561 | 3,75E-05 | 0,00047969 | |
| ENSGALG00000004132 | 40450,1788 | 0,317911072 | 0,09276067 | 3,42721839 | 0,0006098 | 0,00536425 | |
| ENSGALG00000009448 | 18330,278 | 0,31659087 | 0,08894466 | 3,55941395 | 0,00037168 | 0,00356548 | IFRD1 |
| ENSGALG00000000258 | 23571,1755 | 0,314352619 | 0,08646657 | 3,63553944 | 0,0002774 | 0,00278044 | |
| ENSGALG00000004063 | 4760,35876 | 0,313966155 | 0,0919727 | 3,41368847 | 0,0006409 | 0,00560237 | SLC25A14 |
| ENSGALG00000008315 | 6755,45974 | 0,312048416 | 0,08519158 | 3,66290207 | 0,00024937 | 0,00253574 | |
| ENSGALG00000007241 | 5082,09102 | 0,311992362 | 0,09448776 | 3,30193399 | 0,00096021 | 0,00788138 | NUBP1 |
| ENSGALG00000009992 | 11411,8856 | 0,311546697 | 0,0854138 | 3,64749847 | 0,00026481 | 0,00266918 | ARHGAP5 |
| ENSGALG00000008837 | 13220,7097 | 0,311145069 | 0,0811437 | 3,83449427 | 0,00012582 | 0,0013978 | PELI1 |
| ENSGALG00000008265 | 23891,3346 | 0,310830618 | 0,08922654 | 3,48361172 | 0,0004947 | 0,00449394 | |
| ENSGALG00000010084 | 23564,6118 | 0,310711961 | 0,07381728 | 4,20920342 | 2,56E-05 | 0,00034103 | EIF4G3 |
| ENSGALG00000011228 | 6953,04279 | 0,310544896 | 0,0872542 | 3,55908236 | 0,00037215 | 0,0035661 | WARS |
| ENSGALG00000004938 | 5563,15704 | 0,309369487 | 0,09539583 | 3,24300846 | 0,00118275 | 0,00936984 | |
| ENSGALG00000017111 | 14220,273 | 0,309121441 | 0,09000983 | 3,43430744 | 0,00059407 | 0,00525175 | NUP58 |
| ENSGALG00000006971 | 10261,5906 | 0,308524903 | 0,09239987 | 3,33901865 | 0,00084075 | 0,00708211 | |
| ENSGALG00000003039 | 8205,74635 | 0,308092774 | 0,08797415 | 3,50208302 | 0,00046164 | 0,0042398 | SRM |
| ENSGALG00000000346 | 4708,45419 | 0,306247189 | 0,08995671 | 3,40438422 | 0,00066313 | 0,00574456 | |
| ENSGALG00000002863 | 20573,4084 | 0,306238468 | 0,09158034 | 3,34393231 | 0,000826 | 0,00697665 | ERGIC1 |
| ENSGALG00000003830 | 15302,2463 | 0,304702804 | 0,07866157 | 3,87359158 | 0,00010724 | 0,00122512 | |
| ENSGALG00000021041 | 6232,92756 | 0,302815674 | 0,08606635 | 3,51839787 | 0,00043416 | 0,00404688 | |
| ENSGALG00000005397 | 4086,38441 | 0,301881405 | 0,09352409 | 3,22784652 | 0,00124726 | 0,00980639 | GGNBP2 |
| ENSGALG00000010349 | 6576,00357 | 0,300633215 | 0,09122991 | 3,29533608 | 0,00098304 | 0,00803191 | POLR1C |
| ENSGALG00000004295 | 16498,2044 | 0,300034342 | 0,08565811 | 3,50269613 | 0,00046057 | 0,00423316 | CPD |
| ENSGALG00000007387 | 13168,5046 | 0,298075088 | 0,08883093 | 3,35553249 | 0,00079212 | 0,0067223 | GTF3C1 |
| ENSGALG00000011357 | 6177,88173 | 0,296426831 | 0,09147912 | 3,24037681 | 0,00119372 | 0,00943286 | VEZT |
| ENSGALG00000010536 | 11637,7722 | 0,29615271 | 0,08796769 | 3,36660761 | 0,00076099 | 0,00649324 | FAF1 |
| ENSGALG00000011400 | 11997,2666 | 0,295706421 | 0,08251808 | 3,58353479 | 0,00033898 | 0,00330495 | USP44 |
| ENSGALG00000012138 | 28311,992 | 0,295412949 | 0,08299351 | 3,55947062 | 0,0003716 | 0,00356548 | |
| ENSGALG00000002985 | 13749,5981 | 0,293105421 | 0,07993628 | 3,6667381 | 0,00024566 | 0,00250449 | UTP18 |
| ENSGALG00000003391 | 11311,3076 | 0,293097669 | 0,08152671 | 3,59511199 | 0,00032425 | 0,00317625 | EIF2AK1 |
| ENSGALG00000006856 | 8627,6039 | 0,290795459 | 0,08950457 | 3,2489453 | 0,00115834 | 0,00920562 | CD151 |
| ENSGALG00000006895 | 13236,4315 | 0,290027475 | 0,08022918 | 3,61498721 | 0,00030036 | 0,00298275 | CEP131 |
| ENSGALG00000026077 | 76281,091 | 0,289436865 | 0,08082386 | 3,58108204 | 0,00034217 | 0,00333353 | ALDOC |
| ENSGALG00000004625 | 14027,7064 | 0,287002136 | 0,08764175 | 3,27471937 | 0,00105767 | 0,00854125 | ATXN2 |
| ENSGALG00000004098 | 13227,4006 | 0,28601688 | 0,0879168 | 3,25326778 | 0,00114086 | 0,00910141 | STK4 |
| ENSGALG00000005464 | 21720,0724 | 0,284610322 | 0,07882026 | 3,61087753 | 0,00030516 | 0,00301783 | |
| ENSGALG00000016910 | 23116,5508 | 0,283615388 | 0,07814404 | 3,62939229 | 0,00028409 | 0,00283596 | RBM26 |
| ENSGALG00000015995 | 13978,5364 | 0,283340125 | 0,08204665 | 3,45340254 | 0,00055356 | 0,00494255 | PTPRA |
| ENSGALG00000009325 | 20595,5323 | 0,276324484 | 0,08171116 | 3,38172254 | 0,00072033 | 0,00618434 | |
| ENSGALG00000024094 | 16575,1537 | 0,274969008 | 0,08126295 | 3,38369463 | 0,00071517 | 0,00614658 | UBAP2L |
| ENSGALG00000003339 | 10161,4212 | 0,27482717 | 0,08370391 | 3,28332526 | 0,0010259 | 0,00831154 | MTOR |
| ENSGALG00000010481 | 11769,112 | 0,274549325 | 0,08028743 | 3,41958023 | 0,00062718 | 0,00549396 | CMPK1 |
| ENSGALG00000005243 | 13689,9827 | 0,272933583 | 0,07658125 | 3,56397392 | 0,00036528 | 0,00351756 | PPP3CB |
| ENSGALG00000009782 | 45274,5885 | 0,272414043 | 0,08362168 | 3,25769643 | 0,0011232 | 0,00897799 | NAA15 |
| ENSGALG00000011596 | 8125,50614 | 0,272230864 | 0,08398232 | 3,24152578 | 0,00118892 | 0,00940681 | |
| ENSGALG00000006935 | 12087,4137 | 0,270244925 | 0,0815817 | 3,31256806 | 0,00092444 | 0,00763324 | |
| ENSGALG00000006564 | 13301,7317 | 0,268766276 | 0,07743018 | 3,47107908 | 0,00051837 | 0,00467506 | |
| ENSGALG00000008348 | 63887,3158 | 0,267187928 | 0,07700129 | 3,46991508 | 0,00052062 | 0,00469198 | PDIA3 |
| ENSGALG00000003078 | 20092,3466 | 0,264245742 | 0,07691385 | 3,43560664 | 0,00059123 | 0,00523033 | MMD |
| ENSGALG00000012137 | 17323,9143 | 0,256853565 | 0,07795662 | 3,29482705 | 0,00098482 | 0,00803597 | KTN1 |
| ENSGALG00000001364 | 18433,6359 | 0,254871955 | 0,07722743 | 3,30027759 | 0,00096589 | 0,00791766 | CSNK1A1 |
| ENSGALG00000010318 | 32908,0068 | 0,246411634 | 0,0714677 | 3,44787431 | 0,00056502 | 0,00502688 | TMED10 |
| ENSGALG00000002745 | 41084,1287 | 0,246172053 | 0,07133347 | 3,45100341 | 0,00055851 | 0,00497604 | |
| ENSGALG00000005930 | 23930,3439 | -0,243448299 | 0,0744484 | -3,2700274 | 0,00107537 | 0,00867299 | PLS3 |
| ENSGALG00000020813 | 18469,9592 | -0,256523625 | 0,07948115 | -3,2274775 | 0,00124887 | 0,00980672 | |
| ENSGALG00000026300 | 10090,9313 | -0,257041042 | 0,0792187 | -3,2447017 | 0,00117574 | 0,00932021 | |
| ENSGALG00000000812 | 26499,6103 | -0,265852554 | 0,08212241 | -3,2372716 | 0,00120679 | 0,00952409 | OAZ1 |
| ENSGALG00000029037 | 7790,04361 | -0,270845781 | 0,08268109 | -3,2757889 | 0,00105367 | 0,00851447 | |

| | | | | | | | |
|---|---|---|---|---|---|---|---|
| ENSGALG00000014756 | 8148,92025 | -0,271609614 | 0,08352364 | -3,2518892 | 0,00114641 | 0,00913984 | PPWD1 |
| ENSGALG00000002377 | 301877,116 | -0,272471138 | 0,0772308 | -3,5280112 | 0,00041869 | 0,00393497 | ENO1 |
| ENSGALG00000020003 | 18619,6916 | -0,275409017 | 0,07553559 | -3,6460829 | 0,00026627 | 0,0026796 | MAP3K4 |
| ENSGALG00000004048 | 24216,2179 | -0,278746659 | 0,07841685 | -3,554678 | 0,00037844 | 0,00361645 | PTPRS |
| ENSGALG00000027963 | 8992,02102 | -0,279887477 | 0,08105728 | -3,4529592 | 0,00055447 | 0,00494715 | COX7C |
| ENSGALG00000023546 | 8219,08438 | -0,280708865 | 0,08585195 | -3,2696854 | 0,00107667 | 0,00867661 | |
| ENSGALG00000012618 | 8450,7599 | -0,282562019 | 0,08497947 | -3,325062 | 0,00088399 | 0,00736206 | FANCC |
| ENSGALG00000010127 | 15511,214 | -0,2840371 | 0,08325403 | -3,4116921 | 0,00064561 | 0,00563262 | GLO1 |
| ENSGALG00000010829 | 19262,8491 | -0,284700815 | 0,07724408 | -3,68573 | 0,00022805 | 0,00234786 | |
| ENSGALG00000014646 | 9607,01137 | -0,290131032 | 0,08710762 | -3,3307195 | 0,00086622 | 0,007243 | SYT11 |
| ENSGALG00000012185 | 5616,14478 | -0,291440425 | 0,08719363 | -3,342451 | 0,00083042 | 0,00699981 | PLA2G12A |
| ENSGALG00000011276 | 26271,7672 | -0,293405435 | 0,08969033 | -3,2713161 | 0,00107048 | 0,00863913 | BTG1 |
| ENSGALG00000012569 | 7527,80321 | -0,295102597 | 0,09077738 | -3,2508383 | 0,00115065 | 0,00916201 | FBXO7 |
| ENSGALG00000027710 | 10733,9814 | -0,296769112 | 0,08642619 | -3,4337868 | 0,00059521 | 0,00525813 | |
| ENSGALG00000001026 | 8524,74214 | -0,297852025 | 0,08248786 | -3,6108589 | 0,00030518 | 0,00301783 | |
| ENSGALG00000013837 | 12275,1858 | -0,300523253 | 0,08475438 | -3,5458138 | 0,0003914 | 0,00371758 | |
| ENSGALG00000027695 | 6526,32859 | -0,301428682 | 0,09097649 | -3,3132592 | 0,00092215 | 0,00762408 | PCBP4 |
| ENSGALG00000004769 | 29935,7205 | -0,301514296 | 0,08604923 | -3,5039743 | 0,00045837 | 0,0042222 | |
| ENSGALG00000015326 | 8955,0401 | -0,301560094 | 0,08584014 | -3,513043 | 0,00044301 | 0,0041079 | CLTA |
| ENSGALG00000013207 | 7371,39959 | -0,304013471 | 0,09243698 | -3,2888729 | 0,00100589 | 0,00818655 | CTNNAL1 |
| ENSGALG00000006094 | 12757,9566 | -0,306148177 | 0,08654729 | -3,5373513 | 0,00040416 | 0,00382136 | PHF6 |
| ENSGALG00000012121 | 14730,7454 | -0,306219207 | 0,07698902 | -3,9774401 | 6,97E-05 | 0,00083828 | RCN1 |
| ENSGALG00000007127 | 7771,85476 | -0,307935097 | 0,08245774 | -3,7344594 | 0,00018812 | 0,00198413 | |
| ENSGALG00000009559 | 5261,85996 | -0,308352593 | 0,09367103 | -3,2918672 | 0,00099525 | 0,00810516 | |
| ENSGALG00000009474 | 18920,9808 | -0,308530687 | 0,08226101 | -3,7506307 | 0,00017639 | 0,0018683 | |
| ENSGALG00000002040 | 12321,1382 | -0,309590748 | 0,08819991 | -3,5101026 | 0,00044793 | 0,00414132 | |
| ENSGALG00000026981 | 16367,9889 | -0,309740111 | 0,09419663 | -3,2882293 | 0,0010082 | 0,00819996 | NHSL1 |
| ENSGALG00000006277 | 4802,8632 | -0,310824597 | 0,09438854 | -3,2930334 | 0,00099113 | 0,00807697 | |
| ENSGALG00000012546 | 32458,4091 | -0,312592383 | 0,09413799 | -3,3205762 | 0,00089832 | 0,00746649 | MCM5 |
| ENSGALG00000003474 | 3939,39547 | -0,313391117 | 0,09428082 | -3,3240177 | 0,00088731 | 0,00738477 | MED20 |
| ENSGALG00000026437 | 4338,56055 | -0,314002706 | 0,09459663 | -3,3193857 | 0,00090216 | 0,00748844 | |
| ENSGALG00000015712 | 7609,04078 | -0,314599398 | 0,08949622 | -3,5152257 | 0,00043938 | 0,00408392 | SLBP |
| ENSGALG00000007131 | 72027,5204 | -0,31501454 | 0,08157254 | -3,8617718 | 0,00011257 | 0,00127431 | |
| ENSGALG00000025764 | 4071,26079 | -0,316576106 | 0,096448 | -3,28235 | 0,00102946 | 0,00833494 | FKBP10 |
| ENSGALG00000007668 | 10342,6342 | -0,317701794 | 0,08996064 | -3,5315644 | 0,00041311 | 0,00389125 | BMP7 |
| ENSGALG00000001506 | 7912,15144 | -0,318173703 | 0,08732074 | -3,6437356 | 0,00026871 | 0,0026955 | |
| ENSGALG00000006483 | 6631,82379 | -0,318521408 | 0,08620504 | -3,6949278 | 0,00021995 | 0,00227572 | |
| ENSGALG00000023131 | 4044,89831 | -0,321300472 | 0,09495836 | -3,3835932 | 0,00071544 | 0,00614658 | PDXP |
| ENSGALG00000013784 | 12402,0503 | -0,32170776 | 0,08351065 | -3,8522964 | 0,00011702 | 0,00131988 | CDT1 |
| ENSGALG00000021021 | 5260,37371 | -0,322180937 | 0,09166421 | -3,5147954 | 0,00044009 | 0,00408392 | ARL4C |
| ENSGALG00000001475 | 99646,137 | -0,322303202 | 0,09596675 | -3,3584881 | 0,0007837 | 0,00666371 | |
| ENSGALG00000016870 | 4880,45934 | -0,322919481 | 0,09246763 | -3,4922437 | 0,00047898 | 0,00437341 | TMTC4 |
| ENSGALG00000020528 | 5056,48494 | -0,323800436 | 0,09904607 | -3,2691901 | 0,00107856 | 0,00868191 | INIP |
| ENSGALG00000007965 | 19359,9019 | -0,323830546 | 0,09621896 | -3,3655586 | 0,00076389 | 0,00650475 | HN1 |
| ENSGALG00000006780 | 12874,7901 | -0,325989737 | 0,10037051 | -3,2478639 | 0,00116275 | 0,00922895 | |
| ENSGALG00000009362 | 9549,85526 | -0,327914914 | 0,09727937 | -3,3708577 | 0,00074935 | 0,00641585 | |
| ENSGALG00000004621 | 10575,1137 | -0,328680536 | 0,09230808 | -3,5606907 | 0,00036988 | 0,00355364 | PREX1 |
| ENSGALG00000010758 | 4794,60081 | -0,329891138 | 0,08975382 | -3,6755107 | 0,00023737 | 0,00242988 | |
| ENSGALG00000028470 | 4548,02852 | -0,330867645 | 0,09287432 | -3,5625312 | 0,0003673 | 0,00353152 | |
| ENSGALG00000021365 | 6497,94756 | -0,33098251 | 0,0871271 | -3,7988467 | 0,00014537 | 0,00158263 | DCTN3 |
| ENSGALG00000017015 | 5143,28194 | -0,331426117 | 0,09201475 | -3,6018804 | 0,00031592 | 0,00310683 | RNASEH2B |
| ENSGALG00000007254 | 4308,53571 | -0,33313089 | 0,09410502 | -3,5399905 | 0,00040014 | 0,00379131 | ARHGAP12 |
| ENSGALG00000003429 | 14328,0151 | -0,333616807 | 0,0996214 | -3,3488469 | 0,00081149 | 0,00687711 | PBX1 |
| ENSGALG00000025818 | 7103,10421 | -0,334464872 | 0,08352702 | -4,0042717 | 6,22E-05 | 0,00075404 | |
| ENSGALG00000002225 | 3236,66516 | -0,336939904 | 0,09933028 | -3,3921168 | 0,00069355 | 0,00597493 | RFFL |
| ENSGALG00000008677 | 87282,2545 | -0,33704479 | 0,09556181 | -3,5269822 | 0,00042033 | 0,00394733 | VIM |
| ENSGALG00000001567 | 14603,7017 | -0,337139369 | 0,08829903 | -3,8181548 | 0,00013445 | 0,00147663 | |
| ENSGALG00000008438 | 5574,45982 | -0,337474785 | 0,10321767 | -3,2695448 | 0,00107721 | 0,00867661 | SLC35C1 |
| ENSGALG00000010745 | 57730,9692 | -0,33788407 | 0,10090108 | -3,3486665 | 0,00081201 | 0,00687711 | HMGB2 |
| ENSGALG00000005739 | 5611,79649 | -0,338102535 | 0,09729228 | -3,4751219 | 0,00051062 | 0,00461848 | SCD |
| ENSGALG00000014496 | 5642,31406 | -0,339160901 | 0,09495091 | -3,5719606 | 0,00035432 | 0,00342253 | Prom1 |
| ENSGALG00000006539 | 5064,49097 | -0,340143008 | 0,09970044 | -3,41165 | 0,00064571 | 0,00563262 | C21orf2 |
| ENSGALG00000003930 | 5156,97026 | -0,340892203 | 0,10217013 | -3,3365153 | 0,00084836 | 0,00712222 | APBA2 |
| ENSGALG00000004581 | 6531,73175 | -0,341091866 | 0,08790547 | -3,8802119 | 0,00010437 | 0,00119443 | NDST1 |

| | | | | | | | |
|---|---|---|---|---|---|---|---|
| ENSGALG00000008525 | 5352,16855 | -0,34141534 | 0,09154577 | -3,7294496 | 0,0001919 | 0,00201889 | ABHD12 |
| ENSGALG00000010533 | 8438,68556 | -0,341487952 | 0,10389076 | -3,2869907 | 0,00101264 | 0,00822408 | ELAVL4 |
| ENSGALG00000010892 | 4569,599 | -0,341632417 | 0,09812504 | -3,481603 | 0,00049842 | 0,0045245 | |
| ENSGALG00000016417 | 9617,73728 | -0,343209424 | 0,08745497 | -3,9244129 | 8,69E-05 | 0,00101926 | |
| ENSGALG00000000684 | 3976,23685 | -0,343510226 | 0,09330701 | -3,681505 | 0,00023186 | 0,00238124 | KLHDC8A |
| ENSGALG00000010444 | 2882,2778 | -0,345177441 | 0,10587293 | -3,2602994 | 0,00111295 | 0,00890713 | |
| ENSGALG00000000231 | 17219,2246 | -0,345221534 | 0,08265368 | -4,1767231 | 2,96E-05 | 0,00038737 | |
| ENSGALG00000010170 | 3503,64077 | -0,34648621 | 0,09982039 | -3,4710967 | 0,00051834 | 0,00467506 | INTU |
| ENSGALG00000027648 | 7668,3666 | -0,347098945 | 0,08376328 | -4,1438078 | 3,42E-05 | 0,00044142 | |
| ENSGALG00000008758 | 4589,06242 | -0,348320506 | 0,0918692 | -3,791483 | 0,00014975 | 0,00162325 | |
| ENSGALG00000010674 | 11516,7147 | -0,348433039 | 0,07775009 | -4,4814487 | 7,41E-06 | 0,00011379 | DCTD |
| ENSGALG00000020956 | 5772,71418 | -0,348683982 | 0,0891113 | -3,9129044 | 9,12E-05 | 0,00106213 | |
| ENSGALG00000016034 | 14272,6354 | -0,349640251 | 0,08038023 | -4,3498288 | 1,36E-05 | 0,00019567 | ATRN |
| ENSGALG00000005961 | 8289,71197 | -0,349909098 | 0,10219515 | -3,4239306 | 0,00061722 | 0,00541814 | dAK |
| ENSGALG00000001733 | 12421,7814 | -0,350339549 | 0,08564084 | -4,0907999 | 4,30E-05 | 0,00054208 | LRIG2 |
| ENSGALG00000014262 | 6153,36709 | -0,35183883 | 0,09238387 | -3,8084442 | 0,00013984 | 0,00152929 | APBB2 |
| ENSGALG00000014970 | 12546,5116 | -0,352181226 | 0,08624907 | -4,0833047 | 4,44E-05 | 0,00055763 | FSTL1 |
| ENSGALG00000009670 | 3220,19002 | -0,352936897 | 0,10843055 | -3,254958 | 0,00113409 | 0,00905897 | SGCE |
| ENSGALG00000001329 | 7348,8892 | -0,35438055 | 0,08566964 | -4,1365943 | 3,52E-05 | 0,00045411 | EPB41 |
| ENSGALG00000001604 | 6686,31029 | -0,355709643 | 0,09818594 | -3,6228163 | 0,00029141 | 0,00290229 | |
| ENSGALG00000000921 | 4280,88802 | -0,356222635 | 0,09720598 | -3,6646163 | 0,00024771 | 0,00252329 | |
| ENSGALG00000013672 | 6274,94537 | -0,356442405 | 0,09450669 | -3,7716103 | 0,0001622 | 0,00174157 | TSHZ1 |
| ENSGALG00000006634 | 43306,474 | -0,358696121 | 0,07111068 | -5,0441945 | 4,55E-07 | 8,72E-06 | |
| ENSGALG00000022891 | 4638,29282 | -0,359408325 | 0,09123912 | -3,9391911 | 8,18E-05 | 0,00096481 | |
| ENSGALG00000006887 | 5925,3009 | -0,35954959 | 0,08731049 | -4,1180572 | 3,82E-05 | 0,00048789 | FAM174B |
| ENSGALG00000017329 | 3279,3596 | -0,359975454 | 0,1052627 | -3,4197818 | 0,00062671 | 0,00549374 | FCHSD2 |
| ENSGALG00000009650 | 11565,6296 | -0,362142597 | 0,07848435 | -4,6142011 | 3,95E-06 | 6,39E-05 | |
| ENSGALG00000007128 | 21899,2708 | -0,362594465 | 0,07501963 | -4,8333277 | 1,34E-06 | 2,38E-05 | SH3BGRL |
| ENSGALG00000004954 | 30674,7809 | -0,365189206 | 0,0801208 | -4,5579828 | 5,16E-06 | 8,21E-05 | |
| ENSGALG00000012395 | 9821,77442 | -0,367131435 | 0,08321753 | -4,4117079 | 1,03E-05 | 0,00015182 | PLEKHG1 |
| ENSGALG00000014749 | 4798,93922 | -0,36753325 | 0,09967341 | -3,6873752 | 0,00022658 | 0,00233659 | EPHB6 |
| ENSGALG00000011078 | 7107,44613 | -0,368557879 | 0,09522589 | -3,8703537 | 0,00010868 | 0,00123699 | |
| ENSGALG00000021340 | 9694,86728 | -0,368750238 | 0,08310139 | -4,4373533 | 9,11E-06 | 0,0001366 | CA9 |
| ENSGALG00000016390 | 3066,76348 | -0,369515643 | 0,106671 | -3,4640683 | 0,00053207 | 0,00476427 | |
| ENSGALG00000015444 | 4115,05377 | -0,370185453 | 0,10012196 | -3,6973451 | 0,00021787 | 0,00225977 | |
| ENSGALG00000012130 | 2125,48893 | -0,373712161 | 0,11059162 | -3,3792088 | 0,00072695 | 0,00623689 | DNAJC24 |
| ENSGALG00000014033 | 4259,25139 | -0,375904723 | 0,10689108 | -3,516708 | 0,00043693 | 0,00406667 | |
| ENSGALG00000021653 | 5151,69547 | -0,375983807 | 0,10509679 | -3,5775006 | 0,0003469 | 0,00337293 | IFT22 |
| ENSGALG00000002940 | 2775,52872 | -0,376363633 | 0,10520586 | -3,5774017 | 0,00034703 | 0,00337293 | |
| ENSGALG00000006886 | 5794,91901 | -0,376920608 | 0,10869717 | -3,4676213 | 0,00052509 | 0,00471861 | DACH2 |
| ENSGALG00000022603 | 14106,8838 | -0,378152541 | 0,07662552 | -4,9350734 | 8,01E-07 | 1,48E-05 | |
| ENSGALG00000026081 | 3245,15307 | -0,379250879 | 0,10385941 | -3,6515795 | 0,00026063 | 0,00263136 | RBM38 |
| ENSGALG00000015469 | 2385,73691 | -0,382238702 | 0,11262796 | -3,393817 | 0,00068926 | 0,00595026 | |
| ENSGALG00000014094 | 3273,18619 | -0,382585394 | 0,11483546 | -3,3315964 | 0,00086349 | 0,00722505 | STK38L |
| ENSGALG00000006702 | 2276,02955 | -0,38370743 | 0,11242363 | -3,4130498 | 0,0006424 | 0,0056116 | MFGE8 |
| ENSGALG00000027472 | 3563,95993 | -0,383715502 | 0,11493616 | -3,3385097 | 0,00084229 | 0,00709032 | TMEM38B |
| ENSGALG00000026313 | 6897,17922 | -0,38698684 | 0,11169382 | -3,4647111 | 0,0005308 | 0,00476108 | RND3 |
| ENSGALG00000024047 | 8317,97675 | -0,387381439 | 0,09008061 | -4,3003863 | 1,71E-05 | 0,00023803 | MYCL |
| ENSGALG00000002249 | 11964,7275 | -0,388441944 | 0,08418985 | -4,6138809 | 3,95E-06 | 6,39E-05 | AGO1 |
| ENSGALG00000002056 | 4578,49374 | -0,389501294 | 0,10629829 | -3,6642291 | 0,00024808 | 0,00252505 | TANGO2 |
| ENSGALG00000014662 | 2243,29 | -0,390476627 | 0,1184381 | -3,2968836 | 0,00097764 | 0,00799823 | RBP5 |
| ENSGALG00000015663 | 9711,19793 | -0,390574583 | 0,09832214 | -3,9723971 | 7,12E-05 | 0,00085254 | HSDL2 |
| ENSGALG00000016148 | 7859,64044 | -0,391482924 | 0,08589693 | -4,557589 | 5,17E-06 | 8,21E-05 | C2CD2 |
| ENSGALG00000006606 | 1856,635 | -0,392475054 | 0,11686619 | -3,3583285 | 0,00078415 | 0,00666371 | |
| ENSGALG00000002599 | 2253,34485 | -0,392633405 | 0,11309679 | -3,4716581 | 0,00051725 | 0,00467172 | ARHGEF39 |
| ENSGALG00000015702 | 9863,18717 | -0,39306292 | 0,08831382 | -4,4507519 | 8,56E-06 | 0,00012928 | |
| ENSGALG00000005214 | 1715,18636 | -0,394482457 | 0,11892754 | -3,3169983 | 0,0009099 | 0,00754772 | |
| ENSGALG00000003135 | 8376,23704 | -0,395749835 | 0,09573043 | -4,1340026 | 3,56E-05 | 0,00045832 | ANK3 |
| ENSGALG00000014753 | 5047,55497 | -0,396666478 | 0,09075263 | -4,3708537 | 1,24E-05 | 0,00018023 | CENPK |
| ENSGALG00000003457 | 2509,63115 | -0,397746039 | 0,12188792 | -3,2632112 | 0,00110157 | 0,00885009 | CNP |
| ENSGALG00000009354 | 2023,12355 | -0,398534666 | 0,12361626 | -3,2239664 | 0,00126428 | 0,00992152 | |
| ENSGALG00000013067 | 7763,34164 | -0,398595631 | 0,09019442 | -4,4192935 | 9,90E-06 | 0,00014729 | |
| ENSGALG00000026203 | 4347,01953 | -0,401070002 | 0,09158837 | -4,3790496 | 1,19E-05 | 0,00017419 | FAM174A |
| ENSGALG00000008425 | 3562,16922 | -0,404601902 | 0,10623889 | -3,8084161 | 0,00013986 | 0,00152929 | |

| | | | | | | | |
|---|---|---|---|---|---|---|---|
| ENSGALG00000012936 | 10559,4624 | -0,405432395 | 0,11906246 | -3,4052077 | 0,00066114 | 0,00573123 | |
| ENSGALG00000028567 | 2180,82431 | -0,405726705 | 0,11524199 | -3,52065 | 0,00043049 | 0,00401866 | MYL9 |
| ENSGALG00000012796 | 9183,72656 | -0,406079242 | 0,08854404 | -4,5861839 | 4,51E-06 | 7,22E-05 | KIAA1549 |
| ENSGALG00000014648 | 2054,34955 | -0,406366184 | 0,11901606 | -3,4143811 | 0,00063927 | 0,00559597 | CETN3 |
| ENSGALG00000016497 | 4692,83115 | -0,407107351 | 0,09447725 | -4,3090517 | 1,64E-05 | 0,00022992 | UBXN2A |
| ENSGALG00000012361 | 12035,1023 | -0,409904876 | 0,11706425 | -3,5015377 | 0,00046258 | 0,00424536 | NIN |
| ENSGALG00000015109 | 6799,6685 | -0,409909583 | 0,09218338 | -4,4466758 | 8,72E-06 | 0,00013143 | |
| ENSGALG00000015386 | 2068,30084 | -0,41009329 | 0,11802543 | -3,4746181 | 0,00051158 | 0,00462382 | RNF170 |
| ENSGALG00000026870 | 1831,07326 | -0,410373931 | 0,12408835 | -3,3071109 | 0,00094264 | 0,00775242 | |
| ENSGALG00000006158 | 8974,72855 | -0,410915864 | 0,08502916 | -4,8326464 | 1,35E-06 | 2,38E-05 | ACVR2B |
| ENSGALG00000016152 | 6212,20219 | -0,412571324 | 0,0962395 | -4,286923 | 1,81E-05 | 0,00025069 | |
| ENSGALG00000015713 | 7773,18487 | -0,41471478 | 0,09908023 | -4,1856459 | 2,84E-05 | 0,00037482 | FAM53A |
| ENSGALG00000012082 | 1953,58525 | -0,41500134 | 0,1210386 | -3,4286694 | 0,00060655 | 0,00534317 | FAIM |
| ENSGALG00000011778 | 2303,47072 | -0,415058611 | 0,12790911 | -3,2449496 | 0,00117471 | 0,00931801 | |
| ENSGALG00000013370 | 2831,43254 | -0,415383566 | 0,10536475 | -3,942339 | 8,07E-05 | 0,00095493 | |
| ENSGALG00000014004 | 7899,02977 | -0,415462509 | 0,09272364 | -4,4806538 | 7,44E-06 | 0,00011394 | |
| ENSGALG00000007304 | 5003,8857 | -0,415655575 | 0,09074301 | -4,5805797 | 4,64E-06 | 7,41E-05 | |
| ENSGALG00000008152 | 3837,90866 | -0,415829645 | 0,10117889 | -4,1098461 | 3,96E-05 | 0,0005028 | |
| ENSGALG00000002831 | 3577,75168 | -0,41600984 | 0,10574326 | -3,9341499 | 8,35E-05 | 0,0009825 | SH3PXD2B |
| ENSGALG00000011238 | 44015,5054 | -0,417063936 | 0,07670305 | -5,4373842 | 5,41E-08 | 1,21E-06 | WLS |
| ENSGALG00000027897 | 2595,29874 | -0,417252062 | 0,11571548 | -3,6058448 | 0,00031114 | 0,00306943 | DTX1 |
| ENSGALG00000006172 | 1892,0827 | -0,41758713 | 0,11598879 | -3,600237 | 0,00031793 | 0,00312163 | ABCC8 |
| ENSGALG00000011558 | 4354,80067 | -0,418200744 | 0,11988928 | -3,4882246 | 0,00048624 | 0,00442998 | |
| ENSGALG00000021616 | 42268,4535 | -0,419663237 | 0,0886351 | -4,7347298 | 2,19E-06 | 3,72E-05 | |
| ENSGALG00000008873 | 22196,3678 | -0,419829297 | 0,09887727 | -4,2459637 | 2,18E-05 | 0,00029594 | POLE3 |
| ENSGALG00000013002 | 2566,67672 | -0,420522444 | 0,10532109 | -3,9927659 | 6,53E-05 | 0,00078854 | DAP1 |
| ENSGALG00000000713 | 13614,4975 | -0,421676085 | 0,08952353 | -4,7102262 | 2,47E-06 | 4,15E-05 | ZFHX3 |
| ENSGALG00000002484 | 4308,16241 | -0,423017499 | 0,1016587 | -4,1611539 | 3,17E-05 | 0,00041259 | CRYM |
| ENSGALG00000026085 | 4035,50855 | -0,424635504 | 0,0949709 | -4,4712169 | 7,78E-06 | 0,0001185 | |
| ENSGALG00000005237 | 13631,0034 | -0,426583983 | 0,07933189 | -5,3772069 | 7,57E-08 | 1,64E-06 | FGD3 |
| ENSGALG00000000966 | 3098,3696 | -0,42899963 | 0,11454974 | -3,7450949 | 0,00018033 | 0,00190837 | |
| ENSGALG00000003742 | 3986,83364 | -0,429013967 | 0,10607378 | -4,0444865 | 5,24E-05 | 0,00064817 | CPAMD8 |
| ENSGALG00000014903 | 2320,30966 | -0,429814699 | 0,12033773 | -3,5717369 | 0,00035462 | 0,00342281 | ITGA2 |
| ENSGALG00000010346 | 4336,77039 | -0,430819834 | 0,09585982 | -4,4942692 | 6,98E-06 | 0,00010807 | TGFB3 |
| ENSGALG00000003602 | 3515,61032 | -0,430901727 | 0,11141807 | -3,8674312 | 0,00010999 | 0,00125076 | |
| ENSGALG00000005330 | 1520,22623 | -0,431988153 | 0,12387209 | -3,4873727 | 0,00048779 | 0,00444089 | SNCAIP |
| ENSGALG00000010158 | 10233,5261 | -0,433584905 | 0,07981872 | -5,4321205 | 5,57E-08 | 1,25E-06 | KANK1 |
| ENSGALG00000016328 | 3476,01841 | -0,434072431 | 0,12020068 | -3,6112311 | 0,00030475 | 0,00301783 | |
| ENSGALG00000026113 | 3962,92871 | -0,434078932 | 0,09966992 | -4,355165 | 1,33E-05 | 0,00019229 | |
| ENSGALG00000017334 | 2609,2748 | -0,435842486 | 0,12736757 | -3,4219266 | 0,00062179 | 0,00545441 | |
| ENSGALG00000009212 | 1797,44087 | -0,438467006 | 0,1197104 | -3,6627312 | 0,00024954 | 0,00253574 | MND1 |
| ENSGALG00000017321 | 1548,21324 | -0,438859423 | 0,12937108 | -3,3922529 | 0,0006932 | 0,00597493 | PLEKHB1 |
| ENSGALG00000000362 | 6059,70576 | -0,440564998 | 0,08939063 | -4,9285366 | 8,28E-07 | 1,53E-05 | |
| ENSGALG00000015368 | 4121,06739 | -0,441174377 | 0,09460983 | -4,6630922 | 3,11E-06 | 5,11E-05 | TMEM175 |
| ENSGALG00000016200 | 8447,45569 | -0,442373266 | 0,1201603 | -3,6815259 | 0,00023184 | 0,00238124 | |
| ENSGALG00000021285 | 5699,38195 | -0,442446231 | 0,11153099 | -3,967025 | 7,28E-05 | 0,00086948 | |
| ENSGALG00000006950 | 4791,25164 | -0,44318092 | 0,09921158 | -4,467028 | 7,93E-06 | 0,00012055 | LIMK2 |
| ENSGALG00000009516 | 4808,75075 | -0,444515061 | 0,09185634 | -4,8392421 | 1,30E-06 | 2,32E-05 | |
| ENSGALG00000013754 | 4623,32074 | -0,444800112 | 0,10725118 | -4,1472749 | 3,36E-05 | 0,00043614 | |
| ENSGALG00000012158 | 18186,212 | -0,44560696 | 0,08214388 | -5,4247128 | 5,80E-08 | 1,29E-06 | |
| ENSGALG00000008700 | 14557,9866 | -0,447098587 | 0,09405696 | -4,7534873 | 2,00E-06 | 3,42E-05 | PDCD4 |
| ENSGALG00000024120 | 15628,5446 | -0,448054591 | 0,0899222 | -4,9826915 | 6,27E-07 | 1,18E-05 | |
| ENSGALG00000029174 | 2839,91825 | -0,450699904 | 0,12705495 | -3,5472832 | 0,00038923 | 0,00369972 | |
| ENSGALG00000010971 | 1510,70666 | -0,45100601 | 0,13003884 | -3,4682407 | 0,00052388 | 0,00471113 | CSRNP3 |
| ENSGALG00000019987 | 7452,0282 | -0,451061537 | 0,11200004 | -4,0273338 | 5,64E-05 | 0,0006898 | LRP11 |
| ENSGALG00000000970 | 6965,54674 | -0,451525139 | 0,09506462 | -4,7496653 | 2,04E-06 | 3,48E-05 | |
| ENSGALG00000008136 | 1284,16167 | -0,451819148 | 0,13252856 | -3,409221 | 0,00065149 | 0,0056751 | C2orf69 |
| ENSGALG00000007839 | 50053,4791 | -0,453579178 | 0,09430846 | -4,8095279 | 1,51E-06 | 2,66E-05 | NCAM1 |
| ENSGALG00000000628 | 9521,22366 | -0,454003443 | 0,10552649 | -4,3022698 | 1,69E-05 | 0,00023655 | |
| ENSGALG00000025915 | 4915,79448 | -0,454200002 | 0,12936551 | -3,5109822 | 0,00044645 | 0,00413069 | ARRDC3 |
| ENSGALG00000015537 | 3136,68818 | -0,459457432 | 0,12608507 | -3,6440272 | 0,00026841 | 0,00269461 | MAN2B2 |
| ENSGALG00000006036 | 5865,93911 | -0,461900797 | 0,11016505 | -4,1928072 | 2,76E-05 | 0,00036394 | |
| ENSGALG00000010705 | 6215,71804 | -0,462215381 | 0,14311354 | -3,2297111 | 0,00123915 | 0,00975492 | ZBTB18 |
| ENSGALG00000022926 | 4103,57352 | -0,463163963 | 0,11247396 | -4,117966 | 3,82E-05 | 0,00048789 | GPR162 |

| | | | | | | | |
|---|---|---|---|---|---|---|---|
| ENSGALG00000006982 | 2095,88213 | -0,463202789 | 0,13888123 | -3,335244 | 0,00085225 | 0,00715006 | |
| ENSGALG00000009064 | 6911,57971 | -0,463588885 | 0,0867529 | -5,3437857 | 9,10E-08 | 1,93E-06 | FZD1 |
| ENSGALG00000000939 | 13136,3898 | -0,464735788 | 0,11044249 | -4,2079439 | 2,58E-05 | 0,00034257 | |
| ENSGALG00000003595 | 2171,0558 | -0,465858104 | 0,11751231 | -3,9643346 | 7,36E-05 | 0,00087851 | SARM1 |
| ENSGALG00000017119 | 2363,66079 | -0,467363832 | 0,11628448 | -4,0191421 | 5,84E-05 | 0,00071214 | TNFRSF19 |
| ENSGALG00000004598 | 1493,7651 | -0,468100334 | 0,1412582 | -3,3137923 | 0,0009204 | 0,00761459 | CUX2 |
| ENSGALG00000009475 | 10491,2397 | -0,469402079 | 0,09789746 | -4,7948339 | 1,63E-06 | 2,85E-05 | |
| ENSGALG00000006885 | 5604,36422 | -0,469804206 | 0,10737161 | -4,3754975 | 1,21E-05 | 0,00017684 | |
| ENSGALG00000010422 | 4706,76498 | -0,470203966 | 0,12428621 | -3,7832354 | 0,0001548 | 0,00167222 | |
| ENSGALG00000007710 | 2351,67196 | -0,471105348 | 0,10711222 | -4,3982409 | 1,09E-05 | 0,00016079 | |
| ENSGALG00000027373 | 14650,434 | -0,47216803 | 0,10227634 | -4,616591 | 3,90E-06 | 6,33E-05 | |
| ENSGALG00000001774 | 34212,4119 | -0,47258081 | 0,07718637 | -6,1225938 | 9,21E-10 | 2,65E-08 | NEO1 |
| ENSGALG00000015107 | 1553,32125 | -0,4734568 | 0,1315676 | -3,598582 | 0,00031996 | 0,00313663 | PIP5K1B |
| ENSGALG00000012658 | 1659,97017 | -0,475116315 | 0,12664668 | -3,7515101 | 0,00017577 | 0,00186492 | TCP11L2 |
| ENSGALG00000015660 | 2274,63097 | -0,475673447 | 0,12210103 | -3,8957367 | 9,79E-05 | 0,00112766 | SNX30 |
| ENSGALG00000012586 | 1975,98196 | -0,476656889 | 0,11252641 | -4,2359557 | 2,28E-05 | 0,00030611 | GKAP1 |
| ENSGALG00000000316 | 1513,96938 | -0,476826949 | 0,12199447 | -3,9085948 | 9,28E-05 | 0,00107924 | |
| ENSGALG00000024481 | 3771,0681 | -0,479725051 | 0,11484828 | -4,177033 | 2,95E-05 | 0,00038725 | |
| ENSGALG00000003782 | 9195,14164 | -0,482389155 | 0,1220329 | -3,9529435 | 7,72E-05 | 0,00091791 | |
| ENSGALG00000014537 | 1887,30146 | -0,482456357 | 0,13681311 | -3,5263898 | 0,00042127 | 0,00395321 | BMF |
| ENSGALG00000029011 | 28886,6858 | -0,483499319 | 0,07402631 | -6,5314526 | 6,51E-11 | 2,18E-09 | SLC35F1 |
| ENSGALG00000015037 | 2876,08049 | -0,483915865 | 0,10209562 | -4,7398299 | 2,14E-06 | 3,64E-05 | EVC |
| ENSGALG00000010316 | 6055,57077 | -0,484012085 | 0,12348301 | -3,9196654 | 8,87E-05 | 0,00103754 | FRAS1 |
| ENSGALG00000016518 | 2706,67897 | -0,485066531 | 0,11148666 | -4,3508932 | 1,36E-05 | 0,00019517 | PHKA2 |
| ENSGALG00000010818 | 5671,54414 | -0,485799462 | 0,10196839 | -4,7642161 | 1,90E-06 | 3,26E-05 | PLPP3 |
| ENSGALG00000009291 | 1052,37449 | -0,489358793 | 0,13781708 | -3,5507847 | 0,00038408 | 0,00366198 | EPHX1 |
| ENSGALG00000001966 | 2082,4586 | -0,490465312 | 0,14000341 | -3,5032383 | 0,00045964 | 0,00423064 | |
| ENSGALG00000005806 | 2070,07142 | -0,492123395 | 0,11517395 | -4,2728706 | 1,93E-05 | 0,00026497 | KIF24 |
| ENSGALG00000016244 | 1547,30105 | -0,492320823 | 0,12815771 | -3,8415233 | 0,00012227 | 0,00136808 | |
| ENSGALG00000001816 | 2143,75098 | -0,495397442 | 0,135894 | -3,6454696 | 0,0002669 | 0,00268384 | MAGI3 |
| ENSGALG00000020626 | 1193,83839 | -0,496278462 | 0,14167036 | -3,5030507 | 0,00045996 | 0,00423064 | |
| ENSGALG00000013722 | 3288,15489 | -0,496572265 | 0,12655366 | -3,9238082 | 8,72E-05 | 0,00102087 | SEMA3B |
| ENSGALG00000016289 | 12122,6528 | -0,49658952 | 0,10671074 | -4,653604 | 3,26E-06 | 5,33E-05 | DST |
| ENSGALG00000011444 | 5389,01007 | -0,497224463 | 0,11523481 | -4,3148806 | 1,60E-05 | 0,00022558 | MFHAS1 |
| ENSGALG00000009207 | 2177,62925 | -0,499904522 | 0,13219125 | -3,7816762 | 0,00015578 | 0,00168128 | TRIM2 |
| ENSGALG00000012017 | 6540,80506 | -0,499993655 | 0,08604614 | -5,8107619 | 6,22E-09 | 1,57E-07 | DAAM1 |
| ENSGALG00000005680 | 1094,2433 | -0,500648312 | 0,13778586 | -3,6335245 | 0,00027958 | 0,00279777 | |
| ENSGALG00000003099 | 3627,50167 | -0,502434401 | 0,11543971 | -4,3523531 | 1,35E-05 | 0,00019432 | PCTP |
| ENSGALG00000003539 | 4597,6882 | -0,503534691 | 0,0959582 | -5,2474378 | 1,54E-07 | 3,16E-06 | KIAA1522 |
| ENSGALG00000026505 | 1253,87743 | -0,503934058 | 0,14609798 | -3,4492883 | 0,00056207 | 0,00500419 | |
| ENSGALG00000008060 | 1648,89018 | -0,504019022 | 0,12955696 | -3,8903277 | 0,00010011 | 0,00114992 | PRKAR2B |
| ENSGALG00000006859 | 2008,47317 | -0,505092886 | 0,12844032 | -3,9325104 | 8,41E-05 | 0,0009883 | |
| ENSGALG00000010811 | 6795,76845 | -0,506356689 | 0,1227478 | -4,1251794 | 3,70E-05 | 0,00047479 | LGMN |
| ENSGALG00000012124 | 3819,62883 | -0,507526075 | 0,09613084 | -5,2795346 | 1,30E-07 | 2,69E-06 | PELI2 |
| ENSGALG00000015786 | 1271,75753 | -0,508635434 | 0,14394592 | -3,5335176 | 0,00041007 | 0,00387136 | REEP1 |
| ENSGALG00000011083 | 1036,15097 | -0,509062257 | 0,15598828 | -3,2634648 | 0,00110059 | 0,00884785 | |
| ENSGALG00000004338 | 30363,8557 | -0,509560052 | 0,07503149 | -6,791283 | 1,11E-11 | 4,07E-10 | |
| ENSGALG00000002777 | 11306,9083 | -0,509947463 | 0,08235673 | -6,1919342 | 5,94E-10 | 1,77E-08 | |
| ENSGALG00000005107 | 11201,627 | -0,51021951 | 0,10373469 | -4,9185041 | 8,72E-07 | 1,60E-05 | |
| ENSGALG00000015152 | 10783,9679 | -0,512951325 | 0,14885454 | -3,4459905 | 0,00056897 | 0,00504594 | BOC |
| ENSGALG00000025977 | 2371,64375 | -0,513209442 | 0,10745001 | -4,7762626 | 1,79E-06 | 3,10E-05 | GADD45A |
| ENSGALG00000028635 | 2843,21769 | -0,514353516 | 0,11810903 | -4,3549042 | 1,33E-05 | 0,00019229 | |
| ENSGALG00000008728 | 786,945495 | -0,517913117 | 0,15751486 | -3,2880271 | 0,00100892 | 0,00820052 | PTER |
| ENSGALG00000000912 | 1664,29263 | -0,519425471 | 0,11917413 | -4,3585423 | 1,31E-05 | 0,00018956 | |
| ENSGALG00000010835 | 74050,5322 | -0,52138496 | 0,08594348 | -6,0666028 | 1,31E-09 | 3,68E-08 | DTX4 |
| ENSGALG00000022713 | 3872,09413 | -0,525946923 | 0,12443951 | -4,2265269 | 2,37E-05 | 0,00031683 | |
| ENSGALG00000027714 | 1668,20814 | -0,526177638 | 0,12300172 | -4,2778071 | 1,89E-05 | 0,00026004 | ATG10 |
| ENSGALG00000005246 | 13679,395 | -0,527306483 | 0,1075472 | -4,903024 | 9,44E-07 | 1,73E-05 | |
| ENSGALG00000014069 | 15161,3096 | -0,529241353 | 0,09850777 | -5,3725847 | 7,76E-08 | 1,68E-06 | TP53I11 |
| ENSGALG00000015628 | 929,886545 | -0,529404749 | 0,14733434 | -3,5932203 | 0,00032662 | 0,0031969 | MSANTD1 |
| ENSGALG00000013182 | 11579,6316 | -0,530382571 | 0,10696981 | -4,9582454 | 7,11E-07 | 1,32E-05 | |
| ENSGALG00000010982 | 4044,72703 | -0,53129631 | 0,13536613 | -3,9248836 | 8,68E-05 | 0,00101918 | |
| ENSGALG00000015847 | 1982,57576 | -0,531317329 | 0,11374907 | -4,6709596 | 3,00E-06 | 4,93E-05 | PRSS35 |
| ENSGALG00000003601 | 11958,9068 | -0,532351713 | 0,09422206 | -5,6499692 | 1,60E-08 | 3,87E-07 | ATP2B4 |

| | | | | | | | |
|---|---|---|---|---|---|---|---|
| ENSGALG00000006480 | 1476,70794 | -0,534658874 | 0,12638611 | -4,230361 | 2,33E-05 | 0,00031279 | TCF7 |
| ENSGALG00000003185 | 3385,21843 | -0,535546504 | 0,12368984 | -4,3297532 | 1,49E-05 | 0,00021268 | SCPEP1 |
| ENSGALG00000011518 | 813,593359 | -0,535965737 | 0,1518756 | -3,5289785 | 0,00041717 | 0,00392357 | POMGNT2 |
| ENSGALG00000027002 | 29714,3795 | -0,536298125 | 0,10571913 | -5,0728579 | 3,92E-07 | 7,58E-06 | |
| ENSGALG00000003648 | 1181,0691 | -0,536868872 | 0,14374328 | -3,7349146 | 0,00018778 | 0,00198222 | SVIP |
| ENSGALG00000013211 | 1995,32702 | -0,539056737 | 0,14233958 | -3,7871176 | 0,0001524 | 0,00164916 | Sial-T2 |
| ENSGALG00000016576 | 12672,1252 | -0,539357045 | 0,13054957 | -4,1314348 | 3,61E-05 | 0,00046252 | OFD1 |
| ENSGALG00000013155 | 2327,87007 | -0,539440335 | 0,12203608 | -4,4203349 | 9,85E-06 | 0,00014675 | |
| ENSGALG00000028015 | 4297,84833 | -0,539854796 | 0,12735891 | -4,238846 | 2,25E-05 | 0,0003031 | |
| ENSGALG00000000449 | 5699,15535 | -0,541504758 | 0,1320378 | -4,1011343 | 4,11E-05 | 0,00052028 | |
| ENSGALG00000011298 | 3487,55527 | -0,543014986 | 0,11354366 | -4,7824332 | 1,73E-06 | 3,01E-05 | RARB |
| ENSGALG00000004270 | 3394,83221 | -0,544775501 | 0,15590131 | -3,4943614 | 0,0004752 | 0,00434521 | ALDH1A2 |
| ENSGALG00000007269 | 2258,18619 | -0,545360128 | 0,11824854 | -4,6119821 | 3,99E-06 | 6,43E-05 | GABRA3 |
| ENSGALG00000016551 | 3348,17917 | -0,548003277 | 0,12821461 | -4,2741096 | 1,92E-05 | 0,00026408 | AP1S2 |
| ENSGALG00000029102 | 8856,23983 | -0,549378242 | 0,09139118 | -6,0112829 | 1,84E-09 | 5,02E-08 | PXYLP1 |
| ENSGALG00000008433 | 2467,54155 | -0,549541331 | 0,11423742 | -4,8105196 | 1,51E-06 | 2,65E-05 | |
| ENSGALG00000007245 | 695,960091 | -0,550277699 | 0,16045688 | -3,4294429 | 0,00060482 | 0,00533548 | |
| ENSGALG00000013964 | 1040,16713 | -0,55034782 | 0,13837128 | -3,977327 | 6,97E-05 | 0,00083828 | SPATA18 |
| ENSGALG00000002552 | 868,094238 | -0,551384108 | 0,15097485 | -3,6521586 | 0,00026005 | 0,00262813 | |
| ENSGALG00000012219 | 2204,64004 | -0,554221866 | 0,12302026 | -4,5051268 | 6,63E-06 | 0,00010372 | UNC5C |
| ENSGALG00000004323 | 3093,09427 | -0,5545441 | 0,12637337 | -4,3881405 | 1,14E-05 | 0,00016765 | RASAL2 |
| ENSGALG00000015987 | 4635,85249 | -0,554680878 | 0,12141534 | -4,5684578 | 4,91E-06 | 7,84E-05 | |
| ENSGALG00000008871 | 21521,8352 | -0,555142332 | 0,09290105 | -5,9756301 | 2,29E-09 | 6,13E-08 | |
| ENSGALG00000004593 | 2415,84953 | -0,555306466 | 0,163609 | -3,3941071 | 0,00068853 | 0,00594806 | SULF2 |
| ENSGALG00000026757 | 7722,36602 | -0,557063112 | 0,08452448 | -6,5905534 | 4,38E-11 | 1,50E-09 | |
| ENSGALG00000014242 | 1501,10358 | -0,557782152 | 0,14125221 | -3,9488383 | 7,85E-05 | 0,0009329 | |
| ENSGALG00000012710 | 1128,84374 | -0,557841328 | 0,15170545 | -3,6771344 | 0,00023587 | 0,00241843 | NCS1 |
| ENSGALG00000004775 | 914,832175 | -0,558344864 | 0,14503927 | -3,8496118 | 0,00011831 | 0,00133083 | |
| ENSGALG00000015593 | 2880,67783 | -0,558901424 | 0,11770789 | -4,7482072 | 2,05E-06 | 3,50E-05 | EPHA7 |
| ENSGALG00000004537 | 5159,7803 | -0,559387674 | 0,08988409 | -6,2234338 | 4,86E-10 | 1,48E-08 | |
| ENSGALG00000017247 | 4207,7876 | -0,562403241 | 0,09675844 | -5,8124463 | 6,16E-09 | 1,56E-07 | |
| ENSGALG00000000692 | 955,152266 | -0,563167267 | 0,16964073 | -3,3197645 | 0,00090093 | 0,00748326 | CDK18 |
| ENSGALG00000009799 | 18742,1488 | -0,563286437 | 0,12204958 | -4,6152265 | 3,93E-06 | 6,36E-05 | |
| ENSGALG00000002375 | 32718,3264 | -0,563646666 | 0,12800002 | -4,403489 | 1,07E-05 | 0,00015732 | NOTCH1 |
| ENSGALG00000012487 | 1377,92439 | -0,566700383 | 0,13846686 | -4,0926789 | 4,26E-05 | 0,00053825 | |
| ENSGALG00000028795 | 2245,71249 | -0,566961474 | 0,16346113 | -3,468479 | 0,00052341 | 0,00471035 | |
| ENSGALG00000006583 | 5537,14066 | -0,567576615 | 0,11179655 | -5,0768704 | 3,84E-07 | 7,44E-06 | LSP1 |
| ENSGALG00000007007 | 719,343847 | -0,57059825 | 0,16187065 | -3,525026 | 0,00042344 | 0,00396469 | |
| ENSGALG00000004741 | 4792,68775 | -0,573167729 | 0,0909464 | -6,3022583 | 2,93E-10 | 9,27E-09 | EPHB2 |
| ENSGALG00000004256 | 10783,2788 | -0,575998874 | 0,08476767 | -6,7950305 | 1,08E-11 | 3,97E-10 | |
| ENSGALG00000004888 | 7031,61915 | -0,578281859 | 0,0937448 | -6,1686821 | 6,89E-10 | 2,03E-08 | RAI1 |
| ENSGALG00000004400 | 2735,32523 | -0,578988451 | 0,11404307 | -5,0769278 | 3,84E-07 | 7,44E-06 | |
| ENSGALG00000013357 | 3130,31216 | -0,580498969 | 0,15257333 | -3,804721 | 0,00014196 | 0,00154959 | |
| ENSGALG00000027173 | 1590,92759 | -0,581363396 | 0,13148887 | -4,4213887 | 9,81E-06 | 0,00014639 | TMEM159 |
| ENSGALG00000014485 | 4340,50749 | -0,582732951 | 0,10497253 | -5,5512902 | 2,84E-08 | 6,61E-07 | LDB2 |
| ENSGALG00000008034 | 2213,75476 | -0,582849514 | 0,15579351 | -3,7411669 | 0,00018317 | 0,00193681 | CASKIN2 |
| ENSGALG00000005396 | 1307,23993 | -0,582889644 | 0,13145181 | -4,4342459 | 9,24E-06 | 0,00013842 | |
| ENSGALG00000011848 | 4663,28267 | -0,584806197 | 0,10489037 | -5,575404 | 2,47E-08 | 5,81E-07 | |
| ENSGALG00000010940 | 1275,78523 | -0,585237912 | 0,15386088 | -3,8036823 | 0,00014256 | 0,00155339 | ACSS3 |
| ENSGALG00000003404 | 1593,24426 | -0,585273519 | 0,17593109 | -3,3267203 | 0,00087875 | 0,00732815 | |
| ENSGALG00000002707 | 872,059957 | -0,585668905 | 0,17399105 | -3,3660864 | 0,00076243 | 0,00649674 | CHRNB2 |
| ENSGALG00000011657 | 4589,29906 | -0,589843047 | 0,0912792 | -6,4619656 | 1,03E-10 | 3,41E-09 | EAF2 |
| ENSGALG00000012748 | 903,483649 | -0,592402199 | 0,17585259 | -3,368743 | 0,00075512 | 0,00645203 | ELOVL2 |
| ENSGALG00000023279 | 941,141249 | -0,593060121 | 0,14801399 | -4,0067841 | 6,16E-05 | 0,00074752 | EPN3 |
| ENSGALG00000008961 | 48076,4841 | -0,593936394 | 0,13546511 | -4,3844233 | 1,16E-05 | 0,00017034 | |
| ENSGALG00000006409 | 15248,7839 | -0,594856883 | 0,11200977 | -5,310759 | 1,09E-07 | 2,29E-06 | |
| ENSGALG00000001916 | 3626,02385 | -0,595193183 | 0,09718325 | -6,124442 | 9,10E-10 | 2,63E-08 | SLIT3 |
| ENSGALG00000020554 | 873,190798 | -0,595564073 | 0,16838642 | -3,5368889 | 0,00040487 | 0,00382516 | |
| ENSGALG00000015903 | 1387,24274 | -0,597526236 | 0,13866253 | -4,309212 | 1,64E-05 | 0,00022992 | |
| ENSGALG00000002116 | 700,369856 | -0,598531501 | 0,16175305 | -3,7002796 | 0,00021536 | 0,00223751 | TEN1 |
| ENSGALG00000005036 | 1758,38822 | -0,599441683 | 0,16374911 | -3,6607325 | 0,0002515 | 0,0025494 | SH2D3C |
| ENSGALG00000026180 | 3833,45288 | -0,59981586 | 0,14976744 | -4,0049816 | 6,20E-05 | 0,00075251 | |
| ENSGALG00000012346 | 1154,67986 | -0,599840638 | 0,13375751 | -4,4845379 | 7,31E-06 | 0,00011243 | |
| ENSGALG00000000927 | 2190,2949 | -0,600422638 | 0,14772586 | -4,0644382 | 4,81E-05 | 0,00060289 | ANGPTL2 |

| | | | | | | | |
|---|---|---|---|---|---|---|---|
| ENSGALG00000000644 | 7209,00394 | -0,60054163 | 0,12270703 | -4,8941093 | 9,88E-07 | 1,80E-05 | |
| ENSGALG00000011350 | 541,285072 | -0,601290146 | 0,17331192 | -3,4694103 | 0,0005216 | 0,00469742 | NEGR1 |
| ENSGALG00000015661 | 1332,78697 | -0,601504233 | 0,14287821 | -4,2099089 | 2,55E-05 | 0,00034033 | |
| ENSGALG00000010960 | 1489,43056 | -0,603326229 | 0,15158134 | -3,9802144 | 6,89E-05 | 0,00082975 | GALNT3 |
| ENSGALG00000007993 | 1260,65804 | -0,603453342 | 0,13715554 | -4,3997738 | 1,08E-05 | 0,00015985 | DCX |
| ENSGALG00000003973 | 3817,71731 | -0,603622901 | 0,09870926 | -6,1151598 | 9,65E-10 | 2,75E-08 | TLCD1 |
| ENSGALG00000006942 | 9609,38746 | -0,606230537 | 0,07966615 | -7,6096378 | 2,75E-14 | 1,40E-12 | |
| ENSGALG00000012697 | 695,072503 | -0,6064194 | 0,18664528 | -3,2490476 | 0,00115792 | 0,00920562 | SLC41A2 |
| ENSGALG00000026941 | 1374,3273 | -0,606678305 | 0,14040651 | -4,3208702 | 1,55E-05 | 0,00022017 | |
| ENSGALG00000012712 | 1815,03517 | -0,607252424 | 0,12916444 | -4,70139 | 2,58E-06 | 4,32E-05 | RBM24 |
| ENSGALG00000015774 | 1065,02106 | -0,60746392 | 0,15041346 | -4,0386273 | 5,38E-05 | 0,0006613 | CYYR1 |
| ENSGALG00000015542 | 9498,31771 | -0,607580543 | 0,12993075 | -4,6761873 | 2,92E-06 | 4,82E-05 | PLPPR1 |
| ENSGALG00000009537 | 490,58222 | -0,607598218 | 0,18032255 | -3,3695076 | 0,00075303 | 0,00643855 | PPM1L |
| ENSGALG00000000623 | 23029,3222 | -0,609505879 | 0,08219995 | -7,4149182 | 1,22E-13 | 5,72E-12 | |
| ENSGALG00000003375 | 5633,57059 | -0,611230825 | 0,08908432 | -6,861262 | 6,83E-12 | 2,59E-10 | |
| ENSGALG00000014627 | 944,560693 | -0,611235613 | 0,15029669 | -4,0668602 | 4,77E-05 | 0,00059726 | |
| ENSGALG00000016251 | 5263,27603 | -0,611728855 | 0,14715775 | -4,1569598 | 3,23E-05 | 0,0004198 | |
| ENSGALG00000002974 | 1437,50977 | -0,611833543 | 0,15307455 | -3,9969645 | 6,42E-05 | 0,00077544 | SPHKAP |
| ENSGALG00000027240 | 7653,65786 | -0,611869645 | 0,14969349 | -4,0874834 | 4,36E-05 | 0,00054823 | TSPAN13 |
| ENSGALG00000002155 | 2491,16451 | -0,612943427 | 0,15977684 | -3,8362469 | 0,00012493 | 0,00139281 | |
| ENSGALG00000008367 | 12046,292 | -0,614602858 | 0,10352854 | -5,9365551 | 2,91E-09 | 7,68E-08 | MDK |
| ENSGALG00000003163 | 4932,62175 | -0,61460754 | 0,11759786 | -5,2263499 | 1,73E-07 | 3,51E-06 | GNAO1 |
| ENSGALG00000010551 | 740,499781 | -0,614933753 | 0,17383652 | -3,5374256 | 0,00040405 | 0,00382136 | CDC42EP3 |
| ENSGALG00000027183 | 953,592441 | -0,617367034 | 0,15820971 | -3,9022069 | 9,53E-05 | 0,00110199 | |
| ENSGALG00000029070 | 932,579973 | -0,617616574 | 0,16266457 | -3,7968722 | 0,00014653 | 0,00159251 | FAM181A |
| ENSGALG00000011798 | 1064,51312 | -0,617821943 | 0,17164092 | -3,5995026 | 0,00031883 | 0,003128 | HEBP1 |
| ENSGALG00000002652 | 11400,8043 | -0,621511478 | 0,12728765 | -4,8827319 | 1,05E-06 | 1,89E-05 | |
| ENSGALG00000013505 | 3208,27244 | -0,621565495 | 0,14984432 | -4,1480752 | 3,35E-05 | 0,0004354 | |
| ENSGALG00000003582 | 10849,7231 | -0,622442826 | 0,14351033 | -4,3372686 | 1,44E-05 | 0,00020648 | SLC1A3 |
| ENSGALG00000008555 | 1679,68251 | -0,624562442 | 0,15429616 | -4,0478159 | 5,17E-05 | 0,00064028 | MXI1 |
| ENSGALG00000007076 | 2388,80102 | -0,624816157 | 0,15466095 | -4,0399089 | 5,35E-05 | 0,000659 | CCNY |
| ENSGALG00000016391 | 3109,4295 | -0,625110592 | 0,10416619 | -6,0010894 | 1,96E-09 | 5,30E-08 | CNKSR2 |
| ENSGALG00000027377 | 1563,60517 | -0,626128444 | 0,12409736 | -5,0454616 | 4,52E-07 | 8,68E-06 | |
| ENSGALG00000008774 | 10001,1995 | -0,627375234 | 0,09280246 | -6,7603298 | 1,38E-11 | 4,99E-10 | |
| ENSGALG00000000584 | 815,161004 | -0,628246913 | 0,15550344 | -4,0400836 | 5,34E-05 | 0,000659 | |
| ENSGALG00000005403 | 6166,93158 | -0,628470531 | 0,10727764 | -5,8583552 | 4,67E-09 | 1,21E-07 | DHRS11 |
| ENSGALG00000007819 | 1168,66551 | -0,629150395 | 0,15548421 | -4,0463942 | 5,20E-05 | 0,00064355 | PDZRN3 |
| ENSGALG00000015799 | 696,846524 | -0,630241657 | 0,16107768 | -3,9126566 | 9,13E-05 | 0,00106223 | ELAVL2 |
| ENSGALG00000015729 | 1257,59964 | -0,630253804 | 0,13571537 | -4,6439383 | 3,42E-06 | 5,58E-05 | LPAR1 |
| ENSGALG00000007892 | 2832,91388 | -0,631636449 | 0,10556148 | -5,9835882 | 2,18E-09 | 5,85E-08 | SPAG6 |
| ENSGALG00000007748 | 2144,94025 | -0,63248212 | 0,11245141 | -5,6244928 | 1,86E-08 | 4,44E-07 | SLC16A2 |
| ENSGALG00000000040 | 635,110846 | -0,633128633 | 0,18365469 | -3,4473862 | 0,00056604 | 0,00503123 | |
| ENSGALG00000002897 | 1585,0892 | -0,633730718 | 0,13210035 | -4,7973432 | 1,61E-06 | 2,82E-05 | SLC16A14 |
| ENSGALG00000010320 | 1455,65862 | -0,636363776 | 0,13429986 | -4,7383799 | 2,15E-06 | 3,66E-05 | PAWR |
| ENSGALG00000006152 | 1112,15045 | -0,63795304 | 0,16197274 | -3,9386446 | 8,19E-05 | 0,0009661 | |
| ENSGALG00000002437 | 563,695653 | -0,638363537 | 0,17402082 | -3,668317 | 0,00024415 | 0,00249111 | FAM69B |
| ENSGALG00000010269 | 920,840765 | -0,639329258 | 0,17097965 | -3,7392125 | 0,0001846 | 0,00195028 | KCNAB1 |
| ENSGALG00000001153 | 10543,8081 | -0,64025058 | 0,09353164 | -6,8452831 | 7,63E-12 | 2,88E-10 | AUTS2 |
| ENSGALG00000006826 | 5812,76792 | -0,641184602 | 0,13004063 | -4,9306482 | 8,20E-07 | 1,51E-05 | KLHL25 |
| ENSGALG00000028543 | 1233,57546 | -0,64250805 | 0,17971356 | -3,5751785 | 0,00034999 | 0,00339345 | FGFR4 |
| ENSGALG00000026460 | 5792,76927 | -0,643922736 | 0,09174473 | -7,018635 | 2,24E-12 | 9,13E-11 | |
| ENSGALG00000012577 | 1370,05679 | -0,644333952 | 0,12936964 | -4,9805656 | 6,34E-07 | 1,19E-05 | FRMD3 |
| ENSGALG00000001516 | 2392,04392 | -0,648414359 | 0,13283038 | -4,8815216 | 1,05E-06 | 1,90E-05 | |
| ENSGALG00000015241 | 2158,94253 | -0,649652118 | 0,12094277 | -5,3715665 | 7,81E-08 | 1,68E-06 | |
| ENSGALG00000017036 | 2072,67923 | -0,649834759 | 0,14869137 | -4,3703596 | 1,24E-05 | 0,00018042 | LHFP |
| ENSGALG00000000750 | 998,349389 | -0,65019981 | 0,16954656 | -3,8349336 | 0,0001256 | 0,00139654 | |
| ENSGALG00000014508 | 1028,74107 | -0,651699626 | 0,19490886 | -3,3436121 | 0,00082695 | 0,00698 | CD38 |
| ENSGALG00000006490 | 2371,83217 | -0,651963707 | 0,12330582 | -5,2873719 | 1,24E-07 | 2,59E-06 | SCN3B |
| ENSGALG00000002125 | 658,342298 | -0,653074319 | 0,17067212 | -3,826485 | 0,00012999 | 0,00143893 | TMOD1 |
| ENSGALG00000028043 | 7943,94691 | -0,65417089 | 0,08543968 | -7,6565229 | 1,91E-14 | 1,00E-12 | |
| ENSGALG00000002840 | 4314,5965 | -0,659899714 | 0,10957661 | -6,0222678 | 1,72E-09 | 4,71E-08 | GRM4 |
| ENSGALG00000005886 | 3052,74002 | -0,660958059 | 0,12281861 | -5,3815792 | 7,38E-08 | 1,60E-06 | WSCD1 |
| ENSGALG00000029072 | 3177,58522 | -0,662820707 | 0,15123881 | -4,3826099 | 1,17E-05 | 0,00017156 | NTN3 |
| ENSGALG00000009439 | 566,603521 | -0,662995235 | 0,19541313 | -3,3927875 | 0,00069185 | 0,00596854 | GPR85 |

| | | | | | | | |
|---|---|---|---|---|---|---|---|
| ENSGALG00000002712 | 9163,53899 | -0,663245926 | 0,11304776 | -5,8669534 | 4,44E-09 | 1,15E-07 | |
| ENSGALG00000005628 | 1530,5988 | -0,664092027 | 0,18442306 | -3,6009165 | 0,0003171 | 0,00311592 | COL9A3 |
| ENSGALG00000005657 | 1315,51328 | -0,664586696 | 0,19971819 | -3,3276222 | 0,00087591 | 0,00731033 | CRHR2 |
| ENSGALG00000011592 | 769,411267 | -0,666316729 | 0,16592744 | -4,0157116 | 5,93E-05 | 0,00072188 | MRAS |
| ENSGALG00000007732 | 416,813973 | -0,671216647 | 0,19001649 | -3,5324126 | 0,00041179 | 0,00388171 | RTN4R |
| ENSGALG00000011019 | 3781,75966 | -0,672110507 | 0,11796056 | -5,697756 | 1,21E-08 | 2,97E-07 | ROR1 |
| ENSGALG00000016100 | 441,759488 | -0,672712737 | 0,2029277 | -3,3150365 | 0,00091631 | 0,0075858 | |
| ENSGALG00000010775 | 2089,69413 | -0,672804554 | 0,12893781 | -5,2180549 | 1,81E-07 | 3,65E-06 | |
| ENSGALG00000010119 | 1066,14976 | -0,672877793 | 0,15931928 | -4,2234548 | 2,41E-05 | 0,00032084 | CDKL2 |
| ENSGALG00000026055 | 2148,9591 | -0,673106345 | 0,1590473 | -4,2321142 | 2,32E-05 | 0,00031072 | PALM |
| ENSGALG00000004309 | 779,594502 | -0,673847887 | 0,18081676 | -3,7266892 | 0,00019401 | 0,00203427 | GDAP1L1 |
| ENSGALG00000005774 | 3489,02871 | -0,674007323 | 0,11106476 | -6,0685974 | 1,29E-09 | 3,64E-08 | |
| ENSGALG00000017106 | 466,641421 | -0,674065701 | 0,1927802 | -3,4965505 | 0,00047132 | 0,00431602 | ATP8A2 |
| ENSGALG00000016945 | 2967,84555 | -0,674331354 | 0,14913858 | -4,5215085 | 6,14E-06 | 9,66E-05 | LECT1 |
| ENSGALG00000012575 | 8218,93791 | -0,674390391 | 0,10186247 | -6,6205974 | 3,58E-11 | 1,23E-09 | TLE1 |
| ENSGALG00000026704 | 817,227006 | -0,675659204 | 0,17070288 | -3,9581008 | 7,55E-05 | 0,00090089 | GPX8 |
| ENSGALG00000007719 | 1302,68364 | -0,67764292 | 0,20270636 | -3,3429781 | 0,00082884 | 0,00699124 | |
| ENSGALG00000006555 | 2117,21497 | -0,681158051 | 0,13379771 | -5,0909546 | 3,56E-07 | 6,96E-06 | |
| ENSGALG00000016575 | 9168,66303 | -0,686420093 | 0,1297794 | -5,2891299 | 1,23E-07 | 2,57E-06 | GPM6B |
| ENSGALG00000000151 | 1937,7168 | -0,687891713 | 0,11424333 | -6,0212854 | 1,73E-09 | 4,73E-08 | ADAMTS19 |
| ENSGALG00000001727 | 463,318695 | -0,689838866 | 0,18509677 | -3,7269093 | 0,00019384 | 0,0020342 | LZTS1 |
| ENSGALG00000005426 | 877,605348 | -0,690145659 | 0,15508752 | -4,45004 | 8,59E-06 | 0,00012955 | FREM1 |
| ENSGALG00000003550 | 652,802151 | -0,692363086 | 0,17977735 | -3,8512254 | 0,00011753 | 0,00132447 | ARHGAP40 |
| ENSGALG00000023073 | 20456,9956 | -0,692447843 | 0,08514089 | -8,1329649 | 4,19E-16 | 2,52E-14 | |
| ENSGALG00000007364 | 1353,3018 | -0,692880741 | 0,18051262 | -3,8384061 | 0,00012384 | 0,001384 | TOB1 |
| ENSGALG00000009626 | 8049,48001 | -0,694120539 | 0,10537026 | -6,5874428 | 4,47E-11 | 1,53E-09 | THBS1 |
| ENSGALG00000012620 | 28578,6606 | -0,694383357 | 0,081059 | -8,566394 | 1,07E-17 | 7,34E-16 | PTCH1 |
| ENSGALG00000003580 | 1352,19249 | -0,694931491 | 0,15746973 | -4,4131115 | 1,02E-05 | 0,00015102 | MMP2 |
| ENSGALG00000027793 | 607,089392 | -0,696507844 | 0,18113408 | -3,8452612 | 0,00012042 | 0,00134981 | SCN9A |
| ENSGALG00000010903 | 2596,79645 | -0,69779548 | 0,11340379 | -6,1531935 | 7,59E-10 | 2,22E-08 | |
| ENSGALG00000002818 | 12036,7414 | -0,699438203 | 0,11243999 | -6,2205469 | 4,95E-10 | 1,50E-08 | |
| ENSGALG00000003589 | 5291,70331 | -0,699453771 | 0,09356739 | -7,4754011 | 7,70E-14 | 3,70E-12 | VTN |
| ENSGALG00000018624 | 844,593067 | -0,700079474 | 0,18303859 | -3,8247644 | 0,0001309 | 0,00144774 | |
| ENSGALG00000006882 | 2236,27031 | -0,700351611 | 0,11195432 | -6,255691 | 3,96E-10 | 1,22E-08 | ST8SIA2 |
| ENSGALG00000002952 | 1456,76104 | -0,700899465 | 0,13291552 | -5,2732701 | 1,34E-07 | 2,78E-06 | NRG4 |
| ENSGALG00000012226 | 644,458201 | -0,707828644 | 0,18746681 | -3,7757545 | 0,00015952 | 0,00171729 | |
| ENSGALG00000012775 | 643,488801 | -0,709215839 | 0,20574547 | -3,4470545 | 0,00056673 | 0,00503143 | TFAP2A |
| ENSGALG00000004676 | 1415,38407 | -0,710261932 | 0,14188675 | -5,0058368 | 5,56E-07 | 1,05E-05 | |
| ENSGALG00000029140 | 9887,31246 | -0,710672238 | 0,11105305 | -6,3993944 | 1,56E-10 | 5,08E-09 | CRABP2 |
| ENSGALG00000017405 | 1348,63296 | -0,713912912 | 0,17423648 | -4,0973791 | 4,18E-05 | 0,00052797 | NPR3 |
| ENSGALG00000015926 | 932,376679 | -0,714450321 | 0,16657966 | -4,2889409 | 1,80E-05 | 0,00024869 | |
| ENSGALG00000019719 | 1763,03906 | -0,714724163 | 0,18211375 | -3,924603 | 8,69E-05 | 0,00101926 | KRT19 |
| ENSGALG00000025919 | 2203,75306 | -0,717706563 | 0,17787583 | -4,0348739 | 5,46E-05 | 0,00066999 | |
| ENSGALG00000024486 | 1640,6708 | -0,717823268 | 0,14268972 | -5,0306586 | 4,89E-07 | 9,31E-06 | |
| ENSGALG00000002016 | 716,489518 | -0,719060037 | 0,164965 | -4,3588642 | 1,31E-05 | 0,0001895 | PRKCD |
| ENSGALG00000000674 | 402,036081 | -0,719912044 | 0,20464736 | -3,5178175 | 0,00043511 | 0,00405273 | MYO1D |
| ENSGALG00000005608 | 3947,85993 | -0,721159959 | 0,11459976 | -6,2928576 | 3,12E-10 | 9,80E-09 | SLC9A3R2 |
| ENSGALG00000027198 | 9040,88356 | -0,721260873 | 0,09723673 | -7,4175766 | 1,19E-13 | 5,63E-12 | TMEM47 |
| ENSGALG00000011823 | 11238,0507 | -0,721999514 | 0,10012518 | -7,2109681 | 5,56E-13 | 2,43E-11 | FAT4 |
| ENSGALG00000003473 | 17669,5597 | -0,722769839 | 0,13594288 | -5,3167173 | 1,06E-07 | 2,23E-06 | SFRP1 |
| ENSGALG00000009248 | 4501,54609 | -0,722897324 | 0,13466182 | -5,3682427 | 7,95E-08 | 1,71E-06 | HSPA12A |
| ENSGALG00000002749 | 1465,51039 | -0,723137568 | 0,13433788 | -5,3829757 | 7,33E-08 | 1,59E-06 | |
| ENSGALG00000001449 | 3438,32151 | -0,723393814 | 0,10318556 | -7,0106112 | 2,37E-12 | 9,57E-11 | STRA6 |
| ENSGALG00000006792 | 350,301605 | -0,726229187 | 0,22053524 | -3,2930301 | 0,00099114 | 0,00807697 | VSTM2L |
| ENSGALG00000011712 | 1179,53872 | -0,727241211 | 0,14005331 | -5,1926027 | 2,07E-07 | 4,14E-06 | |
| ENSGALG00000012094 | 284,066214 | -0,727903351 | 0,22026176 | -3,3047196 | 0,00095071 | 0,00780859 | SLC35F4 |
| ENSGALG00000003894 | 796,557275 | -0,728609826 | 0,15200269 | -4,7934008 | 1,64E-06 | 2,86E-05 | |
| ENSGALG00000027907 | 6182,31359 | -0,731289025 | 0,08932741 | -8,1866136 | 2,69E-16 | 1,66E-14 | NR2F1 |
| ENSGALG00000029004 | 2171,38417 | -0,731300396 | 0,17746098 | -4,120908 | 3,77E-05 | 0,00048269 | |
| ENSGALG00000015597 | 952,530813 | -0,731477034 | 0,1532441 | -4,7732801 | 1,81E-06 | 3,13E-05 | |
| ENSGALG00000015673 | 6092,107 | -0,738858521 | 0,12050003 | -6,1316045 | 8,70E-10 | 2,52E-08 | |
| ENSGALG00000026858 | 6446,94204 | -0,741127309 | 0,11137264 | -6,6544825 | 2,84E-11 | 9,91E-10 | BTG2 |
| ENSGALG00000009020 | 13288,8425 | -0,742019908 | 0,08130962 | -9,125856 | 7,12E-20 | 5,90E-18 | JAG1 |
| ENSGALG00000005319 | 564,975279 | -0,742306515 | 0,19352205 | -3,8357723 | 0,00012517 | 0,00139302 | CDH8 |

| | | | | | | | |
|---|---|---|---|---|---|---|---|
| ENSGALG00000006211 | 4538,06281 | -0,742344985 | 0,0986063 | -7,5283727 | 5,14E-14 | 2,54E-12 | BCL2L1 |
| ENSGALG00000009536 | 396,027671 | -0,742471798 | 0,20594764 | -3,6051484 | 0,00031197 | 0,00307525 | |
| ENSGALG00000000044 | 1571,14006 | -0,742898149 | 0,12135747 | -6,1215694 | 9,27E-10 | 2,66E-08 | WFIKKN1 |
| ENSGALG00000026119 | 1205,64066 | -0,744514312 | 0,18875953 | -3,9442475 | 8,01E-05 | 0,00094826 | |
| ENSGALG00000002645 | 1167,98665 | -0,746278475 | 0,19471396 | -3,8326911 | 0,00012675 | 0,00140434 | |
| ENSGALG00000011655 | 1680,80637 | -0,746810997 | 0,13582015 | -5,4985288 | 3,83E-08 | 8,71E-07 | IQCB1 |
| ENSGALG00000015075 | 456,601258 | -0,751299073 | 0,202365 | -3,7125939 | 0,00020515 | 0,00214026 | |
| ENSGALG00000012440 | 13707,6458 | -0,752102714 | 0,10042226 | -7,4894023 | 6,92E-14 | 3,35E-12 | |
| ENSGALG00000004370 | 1173,97528 | -0,752418484 | 0,16274623 | -4,6232621 | 3,78E-06 | 6,14E-05 | MAP2K6 |
| ENSGALG00000009048 | 718,531943 | -0,75410293 | 0,17428775 | -4,3267695 | 1,51E-05 | 0,00021509 | ABLIM1 |
| ENSGALG00000026179 | 302,231909 | -0,755339606 | 0,23357232 | -3,2338576 | 0,0012213 | 0,00962652 | FAM163B |
| ENSGALG00000015448 | 3790,52276 | -0,755416575 | 0,16376879 | -4,6127018 | 3,97E-06 | 6,42E-05 | |
| ENSGALG00000005626 | 527,341226 | -0,756044449 | 0,20533084 | -3,6820793 | 0,00023134 | 0,00237979 | TPST2 |
| ENSGALG00000011182 | 9658,13673 | -0,756680805 | 0,13954289 | -5,4225681 | 5,87E-08 | 1,30E-06 | DLL1 |
| ENSGALG00000000690 | 8376,38333 | -0,756893418 | 0,08326065 | -9,0906497 | 9,84E-20 | 8,05E-18 | GDPD4 |
| ENSGALG00000015768 | 2395,79768 | -0,757420183 | 0,12945293 | -5,8509311 | 4,89E-09 | 1,25E-07 | |
| ENSGALG00000006793 | 352,610169 | -0,758190034 | 0,20784148 | -3,6479245 | 0,00026437 | 0,00266691 | |
| ENSGALG00000009959 | 1998,29964 | -0,759714189 | 0,12312472 | -6,1702815 | 6,82E-10 | 2,02E-08 | |
| ENSGALG00000006715 | 4235,48084 | -0,760061887 | 0,0985729 | -7,7106575 | 1,25E-14 | 6,63E-13 | OAF |
| ENSGALG00000003434 | 13153,9326 | -0,761134193 | 0,10176159 | -7,4795823 | 7,46E-14 | 3,60E-12 | C17orf75 |
| ENSGALG00000003837 | 1171,25005 | -0,761557993 | 0,14071896 | -5,4119075 | 6,24E-08 | 1,37E-06 | ADCY7 |
| ENSGALG00000002909 | 7457,53402 | -0,762624106 | 0,09533386 | -7,999509 | 1,25E-15 | 7,23E-14 | |
| ENSGALG00000003193 | 8812,21661 | -0,764114517 | 0,14621053 | -5,2261251 | 1,73E-07 | 3,51E-06 | CRABP-I |
| ENSGALG00000002742 | 2459,13281 | -0,7644405 | 0,11429203 | -6,6884845 | 2,25E-11 | 7,99E-10 | TMEM132B |
| ENSGALG00000015297 | 1133,52891 | -0,764510224 | 0,13956384 | -5,477853 | 4,31E-08 | 9,76E-07 | FOXO3 |
| ENSGALG00000016543 | 3109,19527 | -0,766325509 | 0,11889026 | -6,4456544 | 1,15E-10 | 3,78E-09 | NHS |
| ENSGALG00000006192 | 500,889627 | -0,767192565 | 0,19007181 | -4,0363301 | 5,43E-05 | 0,0006665 | |
| ENSGALG00000015399 | 380,128404 | -0,76931559 | 0,20396936 | -3,7717213 | 0,00016213 | 0,00174157 | FUT10 |
| ENSGALG00000011742 | 234,220676 | -0,769619243 | 0,23242839 | -3,3112102 | 0,00092893 | 0,00766493 | ART4 |
| ENSGALG00000009732 | 5423,65869 | -0,771536411 | 0,14233295 | -5,420645 | 5,94E-08 | 1,31E-06 | PCDH18 |
| ENSGALG00000013194 | 2884,73548 | -0,772416578 | 0,1120601 | -6,8928778 | 5,47E-12 | 2,10E-10 | |
| ENSGALG00000004331 | 987,414427 | -0,775822966 | 0,14440744 | -5,3724583 | 7,77E-08 | 1,68E-06 | |
| ENSGALG00000012595 | 1196,34853 | -0,775902432 | 0,16489518 | -4,7054283 | 2,53E-06 | 4,24E-05 | AGTPBP1 |
| ENSGALG00000007623 | 3044,29553 | -0,778310378 | 0,11922769 | -6,5279333 | 6,67E-11 | 2,22E-09 | CACNA1G |
| ENSGALG00000014853 | 603,727095 | -0,778595081 | 0,1836687 | -4,2391277 | 2,24E-05 | 0,0003031 | CLVS2 |
| ENSGALG00000026042 | 740,41298 | -0,780763307 | 0,23587999 | -3,3100024 | 0,00093295 | 0,00768428 | |
| ENSGALG00000006341 | 275,593591 | -0,786972369 | 0,23377109 | -3,366423 | 0,0007615 | 0,00649324 | |
| ENSGALG00000012191 | 7238,91303 | -0,787037882 | 0,10851003 | -7,2531352 | 4,07E-13 | 1,83E-11 | |
| ENSGALG00000021593 | 580,680268 | -0,787646676 | 0,19140603 | -4,1150567 | 3,87E-05 | 0,00049258 | |
| ENSGALG00000004028 | 313,965547 | -0,787784238 | 0,22915888 | -3,4377208 | 0,00058663 | 0,00519701 | MATN4 |
| ENSGALG00000010854 | 1552,15616 | -0,790623838 | 0,14163311 | -5,5821962 | 2,38E-08 | 5,61E-07 | HDAC9 |
| ENSGALG00000011487 | 7037,53698 | -0,797547344 | 0,08867908 | -8,9936355 | 2,39E-19 | 1,85E-17 | |
| ENSGALG00000015476 | 710,998877 | -0,797713032 | 0,19508358 | -4,0890835 | 4,33E-05 | 0,00054556 | |
| ENSGALG00000005892 | 22205,1699 | -0,805793553 | 0,08367702 | -9,6298065 | 5,98E-22 | 5,94E-20 | |
| ENSGALG00000012652 | 512,848116 | -0,806982417 | 0,18892568 | -4,2714278 | 1,94E-05 | 0,0002664 | |
| ENSGALG00000016223 | 637,815955 | -0,80722926 | 0,16999597 | -4,7485201 | 2,05E-06 | 3,50E-05 | |
| ENSGALG00000004844 | 6351,78047 | -0,807613149 | 0,13771449 | -5,8644021 | 4,51E-09 | 1,17E-07 | |
| ENSGALG00000007929 | 5051,2907 | -0,807992906 | 0,1300334 | -6,2137338 | 5,17E-10 | 1,56E-08 | DIXDC1 |
| ENSGALG00000002228 | 3137,43096 | -0,808972118 | 0,16544265 | -4,8897435 | 1,01E-06 | 1,83E-05 | |
| ENSGALG00000004986 | 20927,5453 | -0,810825031 | 0,09474777 | -8,5577211 | 1,15E-17 | 7,87E-16 | CERCAM |
| ENSGALG00000027498 | 3868,25329 | -0,813503322 | 0,13291213 | -6,1206102 | 9,32E-10 | 2,67E-08 | FOXB1 |
| ENSGALG00000004472 | 11062,9838 | -0,814406115 | 0,10696645 | -7,6136592 | 2,66E-14 | 1,37E-12 | ASTN1 |
| ENSGALG00000010055 | 752,36105 | -0,814715149 | 0,17382196 | -4,6870669 | 2,77E-06 | 4,60E-05 | DAAM2 |
| ENSGALG00000017374 | 378,439025 | -0,814717413 | 0,2081861 | -3,9134092 | 9,10E-05 | 0,00106189 | |
| ENSGALG00000005903 | 1351,52 | -0,81577863 | 0,16378148 | -4,9808968 | 6,33E-07 | 1,19E-05 | NRIP3 |
| ENSGALG00000009514 | 9638,41366 | -0,816414808 | 0,09378509 | -8,7051667 | 3,17E-18 | 2,24E-16 | NRCAM |
| ENSGALG00000013297 | 1134,91265 | -0,820011934 | 0,15144755 | -5,4144946 | 6,15E-08 | 1,35E-06 | BBOX1 |
| ENSGALG00000012092 | 334,595506 | -0,821365542 | 0,23392101 | -3,5112944 | 0,00044593 | 0,0041289 | ESYT3 |
| ENSGALG00000017332 | 676,273003 | -0,827504109 | 0,22658295 | -3,6521022 | 0,0002601 | 0,00262813 | KLHL35 |
| ENSGALG00000027300 | 4224,617 | -0,83359345 | 0,12039523 | -6,9238082 | 4,40E-12 | 1,70E-10 | |
| ENSGALG00000026388 | 192,213198 | -0,83565579 | 0,23979309 | -3,4849036 | 0,00049231 | 0,00447555 | |
| ENSGALG00000011206 | 1241,95142 | -0,836880661 | 0,13265101 | -6,30889 | 2,81E-10 | 8,90E-09 | KITLG |
| ENSGALG00000002911 | 3538,18445 | -0,840005591 | 0,1545939 | -5,4336271 | 5,52E-08 | 1,24E-06 | MOXD1 |
| ENSGALG00000012715 | 336,22161 | -0,842621875 | 0,2238429 | -3,7643449 | 0,00016699 | 0,00178533 | CAP2 |

| | | | | | | | |
|---|---|---|---|---|---|---|---|
| ENSGALG00000022967 | 514,043952 | -0,84515578 | 0,20455601 | -4,1316596 | 3,60E-05 | 0,00046252 | |
| ENSGALG00000011117 | 340,372899 | -0,850337952 | 0,2419193 | -3,5149653 | 0,00043981 | 0,00408392 | AGT |
| ENSGALG00000003424 | 1326,91239 | -0,851626485 | 0,15683659 | -5,4300242 | 5,63E-08 | 1,26E-06 | LMX1A |
| ENSGALG00000015084 | 885,260142 | -0,852219609 | 0,20558018 | -4,1454367 | 3,39E-05 | 0,00043874 | IGSF11 |
| ENSGALG00000023359 | 636,48694 | -0,852612547 | 0,1928659 | -4,4207533 | 9,84E-06 | 0,00014665 | BTBD17 |
| ENSGALG00000006873 | 2180,73864 | -0,854385087 | 0,11802019 | -7,2393131 | 4,51E-13 | 2,01E-11 | PHRF1 |
| ENSGALG00000017195 | 132,512378 | -0,858697871 | 0,2579367 | -3,3291031 | 0,00087126 | 0,0072803 | PGR |
| ENSGALG00000009052 | 2878,13312 | -0,861077306 | 0,16724187 | -5,1486945 | 2,62E-07 | 5,19E-06 | LBH |
| ENSGALG00000026192 | 3872,6926 | -0,861152656 | 0,11660797 | -7,3850243 | 1,52E-13 | 7,11E-12 | ST8SIA4 |
| ENSGALG00000026214 | 847,888186 | -0,867136808 | 0,18566435 | -4,6704539 | 3,01E-06 | 4,94E-05 | LRTM2 |
| ENSGALG00000028946 | 302,158738 | -0,867278013 | 0,21980251 | -3,9457147 | 7,96E-05 | 0,00094425 | |
| ENSGALG00000009478 | 940,161846 | -0,868396515 | 0,21374409 | -4,0627861 | 4,85E-05 | 0,00060596 | |
| ENSGALG00000006662 | 604,284175 | -0,86909427 | 0,21853019 | -3,9769986 | 6,98E-05 | 0,00083863 | |
| ENSGALG00000027484 | 107,369685 | -0,870548896 | 0,26698629 | -3,2606502 | 0,00111157 | 0,00890653 | C20orf85 |
| ENSGALG00000021031 | 6097,64928 | -0,871546666 | 0,10804236 | -8,0667127 | 7,22E-16 | 4,30E-14 | ACKR3 |
| ENSGALG00000008297 | 5581,87806 | -0,875877387 | 0,0888238 | -9,8608413 | 6,15E-23 | 6,64E-21 | SEMA4B |
| ENSGALG00000011709 | 491,428075 | -0,876476185 | 0,23285575 | -3,7640307 | 0,0001672 | 0,00178605 | |
| ENSGALG00000003345 | 838,196489 | -0,878084006 | 0,15479837 | -5,6724371 | 1,41E-08 | 3,43E-07 | |
| ENSGALG00000023738 | 720,292835 | -0,882263773 | 0,18476554 | -4,7750451 | 1,80E-06 | 3,11E-05 | |
| ENSGALG00000007441 | 1451,19846 | -0,88468627 | 0,1740619 | -5,0825959 | 3,72E-07 | 7,24E-06 | |
| ENSGALG00000027501 | 978,829878 | -0,88631831 | 0,14664686 | -6,0438954 | 1,50E-09 | 4,17E-08 | ANPEP |
| ENSGALG00000004527 | 511,661724 | -0,887058508 | 0,20448368 | -4,3380405 | 1,44E-05 | 0,00020599 | |
| ENSGALG00000006235 | 1550,13485 | -0,888605951 | 0,18068598 | -4,9179574 | 8,75E-07 | 1,61E-05 | |
| ENSGALG00000003426 | 2655,93043 | -0,891859274 | 0,1255624 | -7,1029166 | 1,22E-12 | 5,14E-11 | RHBDL3 |
| ENSGALG00000028371 | 2318,19897 | -0,893107554 | 0,18274531 | -4,8871708 | 1,02E-06 | 1,85E-05 | |
| ENSGALG00000012006 | 1652,91118 | -0,89358513 | 0,13601021 | -6,5699857 | 5,03E-11 | 1,70E-09 | RTN1 |
| ENSGALG00000028931 | 2326,56897 | -0,894112314 | 0,12608288 | -7,091465 | 1,33E-12 | 5,53E-11 | |
| ENSGALG00000001129 | 1687,21268 | -0,899923251 | 0,15466843 | -5,818403 | 5,94E-09 | 1,51E-07 | |
| ENSGALG00000000678 | 793,449754 | -0,899943985 | 0,22197436 | -4,0542701 | 5,03E-05 | 0,00062534 | CITED4 |
| ENSGALG00000015186 | 836,758425 | -0,900331794 | 0,15959516 | -5,6413479 | 1,69E-08 | 4,04E-07 | |
| ENSGALG00000006245 | 281,138415 | -0,900397348 | 0,25179693 | -3,5758869 | 0,00034904 | 0,00338989 | |
| ENSGALG00000029057 | 576,095316 | -0,907467539 | 0,18536984 | -4,8954433 | 9,81E-07 | 1,79E-05 | |
| ENSGALG00000009603 | 326,307151 | -0,909672796 | 0,25065366 | -3,6292021 | 0,0002843 | 0,00283596 | |
| ENSGALG00000017074 | 110,235407 | -0,912794343 | 0,27273176 | -3,3468575 | 0,00081733 | 0,00691746 | |
| ENSGALG00000028234 | 573,481229 | -0,912907561 | 0,22530534 | -4,0518682 | 5,08E-05 | 0,00063117 | |
| ENSGALG00000015967 | 195,046703 | -0,91657646 | 0,23708171 | -3,8660784 | 0,0001106 | 0,00125543 | LRRTM1 |
| ENSGALG00000004530 | 1049,1338 | -0,916608336 | 0,21450742 | -4,2730846 | 1,93E-05 | 0,00026497 | MMD2 |
| ENSGALG00000028790 | 633,225912 | -0,919355636 | 0,19021109 | -4,833344 | 1,34E-06 | 2,38E-05 | DNASE2B |
| ENSGALG00000020688 | 639,432719 | -0,921010154 | 0,21245986 | -4,3349843 | 1,46E-05 | 0,00020792 | |
| ENSGALG00000020376 | 2296,58151 | -0,925271568 | 0,10950523 | -8,449565 | 2,92E-17 | 1,90E-15 | |
| ENSGALG00000007078 | 118,265561 | -0,925877896 | 0,27021704 | -3,426423 | 0,00061159 | 0,00537621 | |
| ENSGALG00000005539 | 516,027592 | -0,927364348 | 0,19120632 | -4,8500715 | 1,23E-06 | 2,21E-05 | |
| ENSGALG00000006346 | 1363,78378 | -0,927790987 | 0,17515018 | -5,2971169 | 1,18E-07 | 2,47E-06 | CXCL14 |
| ENSGALG00000027154 | 1132,69645 | -0,928050292 | 0,17313065 | -5,3604045 | 8,30E-08 | 1,77E-06 | |
| ENSGALG00000029106 | 1760,23678 | -0,929278314 | 0,13137058 | -7,0737171 | 1,51E-12 | 6,27E-11 | |
| ENSGALG00000012747 | 3481,16686 | -0,930889427 | 0,16898793 | -5,508615 | 3,62E-08 | 8,30E-07 | NEDD9 |
| ENSGALG00000020575 | 478,814354 | -0,930944308 | 0,20604338 | -4,5181957 | 6,24E-06 | 9,79E-05 | |
| ENSGALG00000028745 | 260,770335 | -0,940548991 | 0,23003741 | -4,0886784 | 4,34E-05 | 0,00054596 | |
| ENSGALG00000009237 | 245,620907 | -0,942598283 | 0,22912432 | -4,1139163 | 3,89E-05 | 0,00049452 | |
| ENSGALG00000009042 | 1781,95784 | -0,948736093 | 0,13944347 | -6,8037326 | 1,02E-11 | 3,75E-10 | ISM1 |
| ENSGALG00000026622 | 538,54299 | -0,952490076 | 0,18017774 | -5,2863915 | 1,25E-07 | 2,60E-06 | |
| ENSGALG00000004849 | 4449,83906 | -0,955100006 | 0,11584173 | -8,2448699 | 1,65E-16 | 1,03E-14 | WSCD2 |
| ENSGALG00000000280 | 585,892203 | -0,956447554 | 0,17580335 | -5,4404398 | 5,31E-08 | 1,20E-06 | EFNA5 |
| ENSGALG00000027514 | 44,3005488 | -0,958727122 | 0,26964144 | -3,555563 | 0,00037717 | 0,00360705 | |
| ENSGALG00000011200 | 585,083893 | -0,962880109 | 0,21843281 | -4,4081295 | 1,04E-05 | 0,00015417 | THBS2 |
| ENSGALG00000001063 | 2024,22024 | -0,96616322 | 0,18451124 | -5,2363381 | 1,64E-07 | 3,34E-06 | |
| ENSGALG00000006507 | 1219,79935 | -0,967854239 | 0,13223106 | -7,3194168 | 2,49E-13 | 1,13E-11 | CLMP |
| ENSGALG00000004790 | 24425,8053 | -0,971232729 | 0,24050391 | -4,0383241 | 5,38E-05 | 0,00066151 | |
| ENSGALG00000028005 | 1938,07994 | -0,972644101 | 0,16792696 | -5,7920664 | 6,95E-09 | 1,74E-07 | GADD45G |
| ENSGALG00000016459 | 278,752189 | -0,980053688 | 0,21673389 | -4,5219218 | 6,13E-06 | 9,65E-05 | FAM84A |
| ENSGALG00000016222 | 2409,35151 | -0,982900573 | 0,14501715 | -6,7778229 | 1,22E-11 | 4,45E-10 | NDP |
| ENSGALG00000026153 | 543,41516 | -0,984548638 | 0,18483355 | -5,326677 | 1,00E-07 | 2,11E-06 | |
| ENSGALG00000014829 | 780,481119 | -0,984611419 | 0,22187254 | -4,4377345 | 9,09E-06 | 0,00013652 | |
| ENSGALG00000026560 | 1741,85122 | -0,986320846 | 0,13866695 | -7,1128761 | 1,14E-12 | 4,80E-11 | |

| | | | | | | | |
|---|---|---|---|---|---|---|---|
| ENSGALG00000007419 | 131,80991 | -0,986369449 | 0,27049343 | -3,6465561 | 0,00026578 | 0,00267683 | MASP1 |
| ENSGALG00000017021 | 594,509791 | -0,994338044 | 0,18535281 | -5,3645694 | 8,11E-08 | 1,74E-06 | ATP7B |
| ENSGALG00000001795 | 808,510482 | -0,995172186 | 0,18954952 | -5,2501962 | 1,52E-07 | 3,12E-06 | |
| ENSGALG00000002832 | 341,099375 | -0,995334878 | 0,21073325 | -4,7231981 | 2,32E-06 | 3,92E-05 | IFI35 |
| ENSGALG00000002098 | 280,73409 | -0,997392471 | 0,25960887 | -3,8419045 | 0,00012208 | 0,00136718 | GRIK3 |
| ENSGALG00000006563 | 4758,12134 | -1,001967625 | 0,10653632 | -9,4049393 | 5,21E-21 | 4,82E-19 | SEMA3D |
| ENSGALG00000019280 | 829,693531 | -1,005672086 | 0,15079747 | -6,669025 | 2,58E-11 | 9,05E-10 | |
| ENSGALG00000020876 | 273,366838 | -1,006719844 | 0,26113425 | -3,8551811 | 0,00011564 | 0,00130677 | |
| ENSGALG00000003554 | 4815,21553 | -1,008700457 | 0,1286997 | -7,8376287 | 4,59E-15 | 2,53E-13 | OPTC |
| ENSGALG00000016570 | 243,048597 | -1,009438758 | 0,22517634 | -4,4828811 | 7,36E-06 | 0,00011317 | |
| ENSGALG00000010500 | 3048,93457 | -1,016915026 | 0,1141321 | -8,9099825 | 5,10E-19 | 3,85E-17 | NKX6-2 |
| ENSGALG00000020615 | 425,649069 | -1,022483514 | 0,20534443 | -4,9793584 | 6,38E-07 | 1,19E-05 | STK32C |
| ENSGALG00000027891 | 12053,1726 | -1,030686173 | 0,08559178 | -12,041883 | 2,14E-33 | 3,77E-31 | |
| ENSGALG00000005981 | 1473,77881 | -1,035267465 | 0,18464036 | -5,6069401 | 2,06E-08 | 4,90E-07 | GDF2 |
| ENSGALG00000003551 | 1571,2827 | -1,036323807 | 0,13255448 | -7,8180974 | 5,36E-15 | 2,94E-13 | |
| ENSGALG00000016810 | 5739,43629 | -1,036681311 | 0,12690987 | -8,1686423 | 3,12E-16 | 1,90E-14 | |
| ENSGALG00000002298 | 1870,01778 | -1,039424987 | 0,15003565 | -6,9278535 | 4,27E-12 | 1,67E-10 | COL8A2 |
| ENSGALG00000020788 | 2719,7038 | -1,041075867 | 0,11340313 | -9,1803095 | 4,30E-20 | 3,66E-18 | |
| ENSGALG00000008279 | 45522,988 | -1,052545903 | 0,11115526 | -9,469151 | 2,82E-21 | 2,65E-19 | |
| ENSGALG00000008455 | 5120,18929 | -1,05482755 | 0,15577951 | -6,7712854 | 1,28E-11 | 4,64E-10 | |
| ENSGALG00000027821 | 3043,02821 | -1,05591502 | 0,14824678 | -7,1226844 | 1,06E-12 | 4,52E-11 | |
| ENSGALG00000013363 | 1658,14825 | -1,059850407 | 0,15135845 | -7,0022548 | 2,52E-12 | 1,01E-10 | |
| ENSGALG00000006576 | 474,441333 | -1,060241233 | 0,24449814 | -4,336398 | 1,45E-05 | 0,00020682 | GRM3 |
| ENSGALG00000009601 | 8847,52543 | -1,061491773 | 0,12977069 | -8,1797498 | 2,84E-16 | 1,75E-14 | NELL2 |
| ENSGALG00000003588 | 1399,08969 | -1,081475528 | 0,13003924 | -8,3165325 | 9,06E-17 | 5,72E-15 | |
| ENSGALG00000009638 | 352,651719 | -1,085140036 | 0,22094352 | -4,9113911 | 9,04E-07 | 1,66E-05 | ANO6 |
| ENSGALG00000007908 | 401,240258 | -1,09305059 | 0,19841137 | -5,5090119 | 3,61E-08 | 8,30E-07 | |
| ENSGALG00000007781 | 567,69339 | -1,095474125 | 0,20462468 | -5,3535776 | 8,62E-08 | 1,84E-06 | GAL3ST1 |
| ENSGALG00000013948 | 833,750503 | -1,099632212 | 0,18641525 | -5,8988318 | 3,66E-09 | 9,54E-08 | RASL11B |
| ENSGALG00000028682 | 140,240123 | -1,101427497 | 0,25860917 | -4,2590428 | 2,05E-05 | 0,00028006 | CBLN2 |
| ENSGALG00000008440 | 326,768063 | -1,108083257 | 0,21643685 | -5,1196608 | 3,06E-07 | 6,02E-06 | CHST1 |
| ENSGALG00000007211 | 263,73208 | -1,108214376 | 0,2365099 | -4,6856998 | 2,79E-06 | 4,62E-05 | CDH22 |
| ENSGALG00000016636 | 272,318125 | -1,110176301 | 0,23729514 | -4,6784621 | 2,89E-06 | 4,78E-05 | |
| ENSGALG00000028506 | 1366,3389 | -1,110787668 | 0,13126336 | -8,4622829 | 2,62E-17 | 1,71E-15 | NHSL2 |
| ENSGALG00000023904 | 802,719075 | -1,115042621 | 0,15879335 | -7,0219729 | 2,19E-12 | 8,94E-11 | FIBIN |
| ENSGALG00000005985 | 8172,66492 | -1,116443935 | 0,11421091 | -9,7752824 | 1,44E-22 | 1,49E-20 | GDF10 |
| ENSGALG00000016511 | 275,151393 | -1,119703281 | 0,22403272 | -4,9979453 | 5,79E-07 | 1,10E-05 | ADGRG2 |
| ENSGALG00000012123 | 16654,8584 | -1,123025117 | 0,07863736 | -14,281063 | 2,87E-46 | 8,16E-44 | PAX6 |
| ENSGALG00000002028 | 1441,95498 | -1,145532965 | 0,19074043 | -6,0057167 | 1,90E-09 | 5,16E-08 | GPRC5B |
| ENSGALG00000003965 | 6511,37534 | -1,147515866 | 0,15881452 | -7,2255096 | 4,99E-13 | 2,21E-11 | DBX1 |
| ENSGALG00000004448 | 445,618082 | -1,158751146 | 0,22989436 | -5,0403635 | 4,65E-07 | 8,87E-06 | BRINP2 |
| ENSGALG00000014699 | 1041,3098 | -1,163328429 | 0,21482752 | -5,4151741 | 6,12E-08 | 1,35E-06 | MEGF10 |
| ENSGALG00000013900 | 1854,85305 | -1,17030528 | 0,14712084 | -7,9547211 | 1,80E-15 | 1,03E-13 | MAP7 |
| ENSGALG00000027633 | 302,366053 | -1,199702733 | 0,21814739 | -5,4995053 | 3,81E-08 | 8,68E-07 | RASSF9 |
| ENSGALG00000001264 | 15314,0129 | -1,21609258 | 0,1243485 | -9,7797129 | 1,38E-22 | 1,43E-20 | PLXNA2 |
| ENSGALG00000008215 | 2557,20905 | -1,222668874 | 0,12813223 | -9,5422426 | 1,40E-21 | 1,36E-19 | |
| ENSGALG00000006210 | 10737,2281 | -1,225951378 | 0,09874124 | -12,415799 | 2,15E-35 | 4,40E-33 | |
| ENSGALG00000005967 | 601,737643 | -1,230002366 | 0,21719963 | -5,663004 | 1,49E-08 | 3,61E-07 | |
| ENSGALG00000023114 | 289,478138 | -1,251095176 | 0,23426953 | -5,3404092 | 9,27E-08 | 1,97E-06 | TAL2 |
| ENSGALG00000026395 | 5473,90727 | -1,254721779 | 0,10114284 | -12,405443 | 2,44E-35 | 4,93E-33 | TRABD2A |
| ENSGALG00000014258 | 1644,91976 | -1,256488908 | 0,16302387 | -7,7073924 | 1,28E-14 | 6,78E-13 | LIMCH1 |
| ENSGALG00000023577 | 185,179234 | -1,257090663 | 0,26329484 | -4,77446 | 1,80E-06 | 3,11E-05 | |
| ENSGALG00000016411 | 2414,195 | -1,262862187 | 0,12175871 | -10,371843 | 3,33E-25 | 4,12E-23 | |
| ENSGALG00000016428 | 17978,0384 | -1,276203339 | 0,12837408 | -9,9412849 | 2,75E-23 | 3,07E-21 | |
| ENSGALG00000029012 | 451,347482 | -1,299502297 | 0,18638978 | -6,971961 | 3,13E-12 | 1,24E-10 | |
| ENSGALG00000007945 | 609,326872 | -1,299576949 | 0,18158766 | -7,156747 | 8,26E-13 | 3,54E-11 | CRYAB |
| ENSGALG00000009604 | 7273,89514 | -1,349943691 | 0,11617707 | -11,619708 | 3,27E-31 | 5,32E-29 | DBX2 |
| ENSGALG00000009948 | 2089,1825 | -1,364538663 | 0,11221803 | -12,15971 | 5,09E-34 | 9,80E-32 | HHIP |
| ENSGALG00000012184 | 1097,0824 | -1,376965225 | 0,21041222 | -6,5441314 | 5,98E-11 | 2,01E-09 | |
| ENSGALG00000005413 | 755,657663 | -1,418701803 | 0,22290236 | -6,3646784 | 1,96E-10 | 6,28E-09 | |
| ENSGALG00000005975 | 1102,4413 | -1,45581134 | 0,21067685 | -6,9101628 | 4,84E-12 | 1,86E-10 | |
| ENSGALG00000011205 | 1465,86067 | -1,462754207 | 0,13057425 | -11,20247 | 3,97E-29 | 5,98E-27 | SMOC2 |

**Supplementary Table 3:** Commonly differentially regulated genes between MEK1+HOXB8 vs MEK1ca and pCIG vs MEK1ca (False Discovery Rate (padj) <= 0.0001). Genes that are inversely regulated are highlighted.

| ID | pCIG vs MEK1ca, Log2FC | MEK1ca+HOXB8 vs MEK1ca, Log2FC | Symbol |
|---|---|---|---|
| ENSGALG00000016783 | 6,569509059 | -5,301446621 | IL1R1 |
| ENSGALG00000026862 | 4,954899999 | -0,764882325 | CLDN1 |
| ENSGALG00000013624 | 4,748606334 | -3,6799039 | FAM65B |
| ENSGALG00000007701 | 4,453082303 | -2,906313035 | B3GNT7 |
| ENSGALG00000023406 | 4,424329106 | -1,774894708 | |
| ENSGALG00000004221 | 4,334100391 | -5,338632777 | IL22RA1 |
| ENSGALG00000016431 | 4,191515764 | -4,37340061 | PAX5 |
| ENSGALG00000006421 | 4,186231336 | -3,742853665 | SLC16A12 |
| ENSGALG00000014994 | 4,160137851 | -5,043766073 | CRHBP |
| ENSGALG00000016145 | 3,924548833 | -2,754137035 | RIPK4 |
| ENSGALG00000016906 | 3,743251462 | -1,870006943 | SPRY2 |
| ENSGALG00000020359 | 3,572883143 | -3,821393345 | SIX6 |
| ENSGALG00000023379 | 3,494464883 | -3,949811074 | F2RL2 |
| ENSGALG00000010435 | 3,471549614 | -2,340611141 | RASGRP3 |
| ENSGALG00000007656 | 3,445291545 | -2,823200316 | |
| ENSGALG00000005519 | 3,391598976 | -1,767947956 | TLL2 |
| ENSGALG00000026663 | 3,38430006 | -2,478656591 | |
| ENSGALG00000002562 | 3,3452383 | -3,12939254 | ARAP3 |
| ENSGALG00000008905 | 3,342984543 | -2,456637668 | HABP2 |
| ENSGALG00000016784 | 3,229015878 | -2,607549769 | IL1RL2 |
| ENSGALG00000008581 | 3,209755496 | -2,566439645 | DUSP5 |
| ENSGALG00000011099 | 3,19842177 | -2,760215905 | FAP |
| ENSGALG00000003456 | 3,130034846 | -3,934712358 | CA12 |
| ENSGALG00000015358 | 3,121559149 | -3,063209848 | MYH15 |
| ENSGALG00000011940 | 2,983749675 | -1,496267605 | CCK |
| ENSGALG00000007963 | 2,963911322 | -1,84239576 | NFATC2 |
| ENSGALG00000023521 | 2,942428455 | -1,651708778 | ADRA2A |
| ENSGALG00000007171 | 2,91119837 | -1,518198388 | CIITA |
| ENSGALG00000005000 | 2,843605103 | -2,158647655 | COMTD1 |
| ENSGALG00000000600 | 2,828234018 | -2,27272638 | PTPN7 |
| ENSGALG00000012018 | 2,812280515 | -1,435600968 | UGT8 |
| ENSGALG00000000177 | 2,765911854 | -1,083701806 | KLF3 |
| ENSGALG00000007346 | 2,763402401 | -2,361764382 | KIAA1462 |
| ENSGALG00000001621 | 2,731716978 | -1,495017381 | CLDN5 |
| ENSGALG00000010224 | 2,688243072 | -2,486198557 | GLIPR1L |
| ENSGALG00000009704 | 2,653508455 | -2,934403789 | CHST15 |
| ENSGALG00000005209 | 2,631256548 | -2,791291062 | RP5-877J2.1 |
| ENSGALG00000014984 | 2,599579307 | -2,931476354 | F2RL1 |
| ENSGALG00000022962 | 2,592879602 | -3,362304357 | OVST |
| ENSGALG00000002183 | 2,571301428 | -2,438499895 | PRR16 |
| ENSGALG00000015388 | 2,539489467 | -1,507384513 | PRDM1 |
| ENSGALG00000016140 | 2,527534269 | -2,998033733 | FAM3B |
| ENSGALG00000013102 | 2,517151592 | -1,724325032 | COBL |
| ENSGALG00000005855 | 2,515154305 | -1,488520056 | CA5B |
| ENSGALG00000002816 | 2,498779812 | -2,170634895 | STK10 |
| ENSGALG00000026258 | 2,487122645 | -0,909500707 | |
| ENSGALG00000003126 | 2,475370264 | -1,615974863 | ERBB4 |
| ENSGALG00000002540 | 2,44573463 | -2,132167559 | RGS2 |
| ENSGALG00000015826 | 2,399148661 | -2,484029553 | HTR1E |
| ENSGALG00000007538 | 2,386772965 | -3,275953369 | IRG1L |
| ENSGALG00000019054 | 2,38528849 | -1,427749976 | NAALAD2 |
| ENSGALG00000026366 | 2,348234721 | -1,956428809 | TTN |
| ENSGALG00000004306 | 2,335846949 | -1,830041848 | SYT12 |
| ENSGALG00000009393 | 2,335695464 | -0,782303357 | CAV1 |
| ENSGALG00000007657 | 2,332413704 | -2,412172788 | CDX4 |
| ENSGALG00000017091 | 2,329587636 | -2,363903034 | FLT1 |
| ENSGALG00000011562 | 2,318757284 | -2,817896047 | PARK2 |
| ENSGALG00000012051 | 2,272220282 | -2,878544854 | STAC |

| | | | |
|---|---|---|---|
| ENSGALG00000011207 | 2,267210819 | -1,434035643 | DUSP6 |
| ENSGALG00000010036 | 2,253052375 | -0,717787007 | FOSL2 |
| ENSGALG00000000949 | 2,249487158 | -1,013784064 | HBEGF |
| ENSGALG00000004376 | 2,243095428 | -2,323750078 | KCNJ2 |
| ENSGALG00000014159 | 2,234266769 | -1,81424576 | PACSIN2 |
| ENSGALG00000017244 | 2,226234853 | -2,189436741 | PRSS23 |
| ENSGALG00000007786 | 2,190066734 | -1,895122662 | DOK5 |
| ENSGALG00000017242 | 2,189728881 | -1,579394776 | FZD4 |
| ENSGALG00000005061 | 2,1754071 | -2,114741441 | PPM1E |
| ENSGALG00000003437 | 2,148196892 | -1,896224168 | ADAM8 |
| ENSGALG00000005499 | 2,068323361 | -1,531899327 | IL17RD |
| ENSGALG00000006995 | 2,052973523 | -1,759714805 | APOD |
| ENSGALG00000004594 | 2,028493881 | -0,774792529 | CD74 |
| ENSGALG00000006112 | 2,027013529 | -2,692857061 | SCN5A |
| ENSGALG00000025941 | 1,970690155 | -2,032822244 | RGS20 |
| ENSGALG00000012732 | 1,970234077 | -2,298252682 | PHACTR1 |
| ENSGALG00000017179 | 1,969058193 | -1,548130217 | PDGFD |
| ENSGALG00000007940 | 1,939184933 | -2,542363859 | RAG2 |
| ENSGALG00000011833 | 1,93191024 | -0,971207285 | SPATA5 |
| ENSGALG00000003323 | 1,918109692 | -1,591926379 | OSGIN1 |
| ENSGALG00000012635 | 1,911001153 | -1,755097361 | BTBD11 |
| ENSGALG00000022490 | 1,907440028 | -0,638026939 | ARPC1B |
| ENSGALG00000008461 | 1,900360149 | 0,900341356 | HGF |
| ENSGALG00000007415 | 1,899486225 | 0,938743099 | SH3RF2 |
| ENSGALG00000000544 | 1,895252672 | 1,877518925 | |
| ENSGALG00000016785 | 1,892879929 | -2,405348378 | IL1RL1 |
| ENSGALG00000005498 | 1,873960213 | -1,707286081 | |
| ENSGALG00000008353 | 1,848707427 | -1,679225856 | MAGI2 |
| ENSGALG00000010597 | 1,8345583 | -0,973101915 | CRIM1 |
| ENSGALG00000012106 | 1,828065766 | -2,236831516 | SCTR |
| ENSGALG00000000141 | 1,818659764 | -1,223136562 | BLB1 |
| ENSGALG00000026262 | 1,788029436 | -1,4922838 | PMP22 |
| ENSGALG00000010468 | 1,7857319 | -1,6309453 | ISM2 |
| ENSGALG00000003706 | 1,784490953 | -1,390323026 | DUSP1 |
| ENSGALG00000011581 | 1,763989645 | -1,059360643 | KIF26A |
| ENSGALG00000023241 | 1,749836088 | -0,938819916 | |
| ENSGALG00000027521 | 1,741041239 | -0,803725098 | ANKRD33B |
| ENSGALG00000012623 | 1,738546704 | -2,482707417 | CFC1B |
| ENSGALG00000029175 | 1,731471147 | -1,685259625 | EFNB2 |
| ENSGALG00000001560 | 1,726930021 | -2,555527127 | KCNU1 |
| ENSGALG00000024083 | 1,719739442 | -2,175833689 | C8orf4 |
| ENSGALG00000005069 | 1,702946767 | 0,669583836 | PTGS2 |
| ENSGALG00000005689 | 1,702270541 | -2,271345118 | PAX2 |
| ENSGALG00000009693 | 1,692012083 | -1,908196694 | CPXM2 |
| ENSGALG00000015332 | 1,689640626 | -2,044938557 | CPLX1 |
| ENSGALG00000006084 | 1,684603389 | -1,531133293 | WNT8A |
| ENSGALG00000028250 | 1,684255106 | -2,884651456 | ANKRD22 |
| ENSGALG00000010258 | 1,674285408 | -1,528410178 | LTBP2 |
| ENSGALG00000008281 | 1,659542176 | -1,009962725 | NEURL1 |
| ENSGALG00000013033 | 1,649436663 | -2,410715686 | CMBL |
| ENSGALG00000009740 | 1,648366239 | -1,063900656 | RASGRP1 |
| ENSGALG00000010910 | 1,643397512 | -0,801671392 | RAPGEF5 |
| ENSGALG00000016602 | 1,632391617 | -1,339565689 | ARHGAP6 |
| ENSGALG00000026989 | 1,621656649 | -0,47543634 | |
| ENSGALG00000016377 | 1,612690665 | -1,984049771 | PXDN |
| ENSGALG00000027218 | 1,609222298 | 1,190798556 | |
| ENSGALG00000008118 | 1,603231134 | -2,340544818 | RFTN2 |
| ENSGALG00000009282 | 1,600131022 | -1,769054515 | ELMSAN1 |
| ENSGALG00000027707 | 1,597314136 | -1,082314729 | ESM1 |
| ENSGALG00000005347 | 1,58404156 | -2,102469906 | ADAMTS18 |
| ENSGALG00000001383 | 1,582310757 | -1,543511015 | HTR4 |
| ENSGALG00000008890 | 1,553993643 | -2,449553325 | ADGRL4 |
| ENSGALG00000017714 | 1,553191397 | -1,052113718 | FHAD1 |
| ENSGALG00000011148 | 1,54566024 | -1,754606037 | HHIPL1 |

| | | | |
|---|---|---|---|
| ENSGALG00000011962 | 1,540068054 | -0,494326456 | NDNF |
| ENSGALG00000008875 | 1,539400497 | -1,156831558 | PLCB1 |
| ENSGALG00000001189 | 1,519846307 | 0,851541498 | NCF1 |
| ENSGALG00000016754 | 1,508945346 | -1,034769759 | MGAT4A |
| ENSGALG00000006997 | 1,507910301 | -1,684979264 | BRINP1 |
| ENSGALG00000019426 | 1,491625247 | -2,280006749 | TMEM71 |
| ENSGALG00000011236 | 1,484958654 | -1,070413941 | BEGAIN |
| ENSGALG00000002594 | 1,484249601 | -3,661637709 | TFPI |
| ENSGALG00000014930 | 1,479606742 | -1,696897635 | ENC1 |
| ENSGALG00000010683 | 1,466981653 | -0,919156764 | S1PR3 |
| ENSGALG00000007660 | 1,456310727 | -1,228207524 | CHIC1 |
| ENSGALG00000028130 | 1,44989129 | 1,140619832 | KIAA1644 |
| ENSGALG00000015865 | 1,445378264 | -0,393610955 | FAM46A |
| ENSGALG00000009667 | 1,442248321 | -1,219318734 | MFAP3L |
| ENSGALG00000005442 | 1,437319869 | -1,327990169 | PALMD |
| ENSGALG00000007771 | 1,43708626 | -1,353742442 | FAM210B |
| ENSGALG00000026790 | 1,433302155 | -0,903869706 | DMRT2 |
| ENSGALG00000027210 | 1,422887936 | -1,142415301 | |
| ENSGALG00000007218 | 1,41095016 | -1,732428983 | MB21D2 |
| ENSGALG00000012702 | 1,407925441 | -0,483830731 | JARID2 |
| ENSGALG00000009546 | 1,402581457 | -0,832251183 | HTRA1 |
| ENSGALG00000011468 | 1,39801296 | -1,424650779 | IGFBP5 |
| ENSGALG00000015665 | 1,394518864 | -1,188036426 | SH3BP2 |
| ENSGALG00000011680 | 1,389021273 | -0,709602993 | EPHA5 |
| ENSGALG00000011419 | 1,382981881 | -0,529859697 | DUSP4 |
| ENSGALG00000014852 | 1,380781721 | -1,119019515 | |
| ENSGALG00000010049 | 1,375590195 | -0,6748978 | PTPRB |
| ENSGALG00000006401 | 1,372021166 | -1,299022844 | OSBPL5 |
| ENSGALG00000002326 | 1,369220123 | -0,593338649 | |
| ENSGALG00000007846 | 1,3613354 | -0,863598394 | |
| ENSGALG00000011120 | 1,356774191 | -0,88227849 | SLC4A10 |
| ENSGALG00000004569 | 1,348974691 | -0,82424563 | UNC5B |
| ENSGALG00000012911 | 1,345449733 | -0,929872331 | SYT10 |
| ENSGALG00000015028 | 1,342064123 | -0,946739996 | RLN1 |
| ENSGALG00000005476 | 1,338834069 | -1,752170936 | PTH1R |
| ENSGALG00000000107 | 1,334266505 | 1,227291307 | TRIM7.1 |
| ENSGALG00000006258 | 1,330333906 | -0,778635815 | SPOCK1 |
| ENSGALG00000015089 | 1,327665029 | -1,867848339 | GAP43 |
| ENSGALG00000011110 | 1,322132253 | -1,431359085 | DPP4 |
| ENSGALG00000008893 | 1,318238187 | -2,444595373 | SYT4 |
| ENSGALG00000005805 | 1,313014625 | -0,954718819 | PLCD1 |
| ENSGALG00000010311 | 1,29508602 | -0,865329105 | NAV3 |
| ENSGALG00000002993 | 1,293663972 | -1,397301236 | CA10 |
| ENSGALG00000015256 | 1,284509548 | -1,397562325 | PAM |
| ENSGALG00000015908 | 1,276423655 | -0,970529728 | COL12A1 |
| ENSGALG00000014727 | 1,275557809 | -2,02826974 | PDE4D |
| ENSGALG00000016817 | 1,2720896 | -0,658902643 | RASA3 |
| ENSGALG00000007253 | 1,27114813 | 1,036667095 | IL1RAP |
| ENSGALG00000009920 | 1,271106358 | -1,80046145 | COCH |
| ENSGALG00000002632 | 1,269452785 | -1,220611399 | CALCRL |
| ENSGALG00000016102 | 1,263113327 | -1,630912312 | CYP26B1 |
| ENSGALG00000001608 | 1,262511696 | -2,081100043 | UNC5D |
| ENSGALG00000016998 | 1,256730923 | -1,177970615 | |
| ENSGALG00000011831 | 1,24831638 | -0,347478626 | ANKRD50 |
| ENSGALG00000028047 | 1,243898501 | -1,369008166 | RHOV |
| ENSGALG00000010010 | 1,237630354 | -0,869227214 | RHOQ |
| ENSGALG00000009947 | 1,236965236 | -1,358616298 | PLEKHH2 |
| ENSGALG00000014983 | 1,23637443 | -1,39278689 | F2R |
| ENSGALG00000005619 | 1,235271948 | -1,189698364 | F3 |
| ENSGALG00000015653 | 1,234972794 | -0,655959247 | JPH1 |
| ENSGALG00000003114 | 1,234804889 | -1,398035927 | NOG |
| ENSGALG00000019716 | 1,234170903 | -1,618712644 | KRT20 |
| ENSGALG00000019261 | 1,230099163 | -1,655584971 | ARHGAP8 |
| ENSGALG00000028821 | 1,225133338 | -1,46102831 | GLIPR2 |

| | | | |
|---|---|---|---|
| ENSGALG00000008999 | 1,220108565 | 0,583129446 | AFAP1L2 |
| ENSGALG00000014823 | 1,217853522 | -1,422401118 | C6orf58 |
| ENSGALG00000000307 | 1,215832976 | -0,894951246 | CERS4 |
| ENSGALG00000007268 | 1,214609385 | -1,168892699 | CADPS |
| ENSGALG00000000721 | 1,214440274 | -0,47173717 | SLC41A1 |
| ENSGALG00000026557 | 1,203253046 | -1,412974627 | PGF |
| ENSGALG00000002555 | 1,202502668 | -0,892902927 | RET |
| ENSGALG00000027133 | 1,199491369 | -1,405319409 | CD83 |
| ENSGALG00000009360 | 1,196193521 | -0,890830846 | CAPN2 |
| ENSGALG00000014864 | 1,190480858 | -1,150010609 | SMPDL3A |
| ENSGALG00000012927 | 1,187339616 | -2,419967236 | CDH10 |
| ENSGALG00000016455 | 1,186649639 | -2,034647249 | GREB1 |
| ENSGALG00000011867 | 1,179474298 | -1,242989678 | SERHL2 |
| ENSGALG00000002294 | 1,174006641 | -0,482168758 | SEMA6A |
| ENSGALG00000002543 | 1,169335598 | -0,840502021 | CSGALNACT2 |
| ENSGALG00000013892 | 1,167370699 | -1,122648268 | MAP3K5 |
| ENSGALG00000011439 | 1,165074777 | -1,116853678 | RBMS3 |
| ENSGALG00000009450 | 1,162765074 | -1,016064164 | DUSP10 |
| ENSGALG00000023768 | 1,155472868 | 0,58136874 | EPHA2 |
| ENSGALG00000016927 | 1,145535192 | -0,715628819 | KLF5 |
| ENSGALG00000006988 | 1,141001037 | 0,944077449 | RNF213 |
| ENSGALG00000012998 | 1,132683468 | -0,849149352 | WNT5B |
| ENSGALG00000028046 | 1,129369719 | -1,098357134 | INSIG2 |
| ENSGALG00000014866 | 1,123162841 | -1,312013425 | FABP7 |
| ENSGALG00000009567 | 1,119009905 | -1,113206952 | TLL1 |
| ENSGALG00000026970 | 1,11847741 | -0,785311616 | |
| ENSGALG00000006376 | 1,116147137 | -1,138122247 | TMC3 |
| ENSGALG00000008469 | 1,115125474 | -0,83875822 | SPINT1 |
| ENSGALG00000010692 | 1,111701613 | -0,996294098 | LRP8 |
| ENSGALG00000010081 | 1,110146926 | -0,563235953 | SH3D19 |
| ENSGALG00000006337 | 1,109144959 | -1,128489411 | K123 |
| ENSGALG00000026763 | 1,104647316 | -1,177313255 | |
| ENSGALG00000009170 | 1,101783321 | -1,581054934 | NCEH1 |
| ENSGALG00000006797 | 1,10056236 | -2,093089949 | ADARB2 |
| ENSGALG00000015980 | 1,100313329 | -0,841274637 | |
| ENSGALG00000012439 | 1,098736639 | -1,028638845 | IGFBP3 |
| ENSGALG00000026518 | 1,097099741 | -0,773283921 | RUNDC3A |
| ENSGALG00000028582 | 1,093959706 | 0,832821984 | LMO2 |
| ENSGALG00000002714 | 1,089564948 | -1,839377527 | TMEM132C |
| ENSGALG00000016875 | 1,086190922 | -1,825016077 | CLYBL |
| ENSGALG00000005492 | 1,08468259 | -0,609713298 | NBEAL2 |
| ENSGALG00000007113 | 1,078758406 | -1,77145981 | TNC |
| ENSGALG00000006822 | 1,077915026 | -2,117548167 | PCDH19 |
| ENSGALG00000023581 | 1,073595233 | -0,876769713 | |
| ENSGALG00000007056 | 1,069890641 | -0,638790551 | KLF6 |
| ENSGALG00000014907 | 1,069885451 | -0,80019622 | DCBLD1 |
| ENSGALG00000006808 | 1,068278734 | -0,948868926 | TSPAN6 |
| ENSGALG00000022871 | 1,067659168 | -1,176605905 | HHLA2 |
| ENSGALG00000028708 | 1,054408367 | -0,938854547 | |
| ENSGALG00000025773 | 1,052013322 | -1,506718569 | |
| ENSGALG00000001020 | 1,049473012 | -0,780112538 | KIFC3 |
| ENSGALG00000028602 | 1,04212846 | -0,881510014 | |
| ENSGALG00000012594 | 1,03128421 | -2,283939186 | NTRK2 |
| ENSGALG00000017411 | 1,030664685 | -1,17758282 | RXFP3 |
| ENSGALG00000011609 | 1,027774555 | -1,61436812 | ABTB2 |
| ENSGALG00000009129 | 1,026339944 | -0,878296092 | DLX5 |
| ENSGALG00000004393 | 1,024579421 | -0,962441655 | |
| ENSGALG00000007646 | 1,023532832 | -1,999886692 | |
| ENSGALG00000016841 | 1,021854986 | -0,576008454 | COL4A1 |
| ENSGALG00000006252 | 1,021389981 | -0,49037706 | LMO4 |
| ENSGALG00000008491 | 1,018906247 | -1,354708919 | XIAP |
| ENSGALG00000006638 | 1,014405382 | -0,469621749 | GRIK4 |
| ENSGALG00000028294 | 1,012130398 | -0,727698928 | VWA5B2 |
| ENSGALG00000002532 | 1,010182876 | -0,997428466 | GJA4 |

| | | | |
|---|---|---|---|
| ENSGALG00000007069 | 1,009530133 | -0,738101541 | ADAMTS17 |
| ENSGALG00000006908 | 1,006453251 | -0,789724754 | DLG1 |
| ENSGALG00000015440 | 1,001448719 | -0,996123586 | RCSD1 |
| ENSGALG00000001419 | 0,999950253 | -0,758492411 | DAB2IP |
| ENSGALG00000015208 | 0,998078994 | -0,778489641 | ROR2 |
| ENSGALG00000008918 | 0,995278864 | -1,685987772 | LAMP5 |
| ENSGALG00000003149 | 0,985359473 | -0,749443102 | ITPR3 |
| ENSGALG00000020723 | 0,981929794 | -0,913543779 | NOG2 |
| ENSGALG00000016537 | 0,979050265 | -0,739933749 | SCML2 |
| ENSGALG00000027497 | 0,976598532 | -0,973179721 | CFD |
| ENSGALG00000004930 | 0,975284752 | -1,782009807 | HRH1 |
| ENSGALG00000016143 | 0,975263193 | 0,607480677 | RHPN1 |
| ENSGALG00000010926 | 0,975212806 | -1,152009571 | SPP1 |
| ENSGALG00000012808 | 0,972939427 | -0,789210521 | CDYL |
| ENSGALG00000003739 | 0,97006222 | -1,237172502 | SALL1 |
| ENSGALG00000014343 | 0,963080504 | -1,412639292 | ARAP2 |
| ENSGALG00000002509 | 0,962423205 | -1,185807324 | COL5A2 |
| ENSGALG00000021879 | 0,950789217 | -1,34129965 | |
| ENSGALG00000006614 | 0,950175613 | 2,721769719 | TECTA |
| ENSGALG00000016843 | 0,946578268 | -0,926939677 | COL4A2 |
| ENSGALG00000017135 | 0,945725278 | -1,071209509 | CRYL1 |
| ENSGALG00000012224 | 0,94448356 | -1,20528636 | PDE1C |
| ENSGALG00000008014 | 0,943508119 | -1,126068911 | CEBPB |
| ENSGALG00000026892 | 0,943046494 | -1,6734567 | |
| ENSGALG00000010163 | 0,938977707 | -0,998496131 | LGR5 |
| ENSGALG00000004662 | 0,937959127 | -0,792400882 | DISP3 |
| ENSGALG00000010836 | 0,937907565 | -0,688393097 | AHR |
| ENSGALG00000002397 | 0,927215999 | -1,291814757 | PSPH |
| ENSGALG00000010412 | 0,925287079 | -0,756929566 | WWTR1 |
| ENSGALG00000007039 | 0,92399781 | -0,823801989 | IGF1R |
| ENSGALG00000008778 | 0,923325024 | -0,51039033 | SPRED2 |
| ENSGALG00000010785 | 0,922406238 | -0,818155142 | ACOT11 |
| ENSGALG00000015164 | 0,920081875 | -0,982829392 | PCSK5 |
| ENSGALG00000014855 | 0,916891226 | -0,627700184 | GHR |
| ENSGALG00000007667 | 0,910790115 | -0,757882718 | GPR158 |
| ENSGALG00000012533 | 0,909053879 | -0,619977689 | MYH9 |
| ENSGALG00000011618 | 0,908730543 | -0,609057453 | ADSSL1 |
| ENSGALG00000015624 | 0,906952704 | -0,97910183 | VCAN |
| ENSGALG00000001645 | 0,906502233 | -1,608468631 | KCNT1 |
| ENSGALG00000011258 | 0,899386948 | -0,946927699 | ATP2B1 |
| ENSGALG00000016308 | 0,888326042 | -0,690314419 | MYC |
| ENSGALG00000007952 | 0,887725324 | -0,932963194 | AMOT |
| ENSGALG00000001281 | 0,886212495 | 1,101845994 | PHOSPHO1 |
| ENSGALG00000010148 | 0,885936489 | -1,379685752 | PTPRR |
| ENSGALG00000001276 | 0,885240877 | -1,144775826 | HOXB9 |
| ENSGALG00000015896 | 0,883690573 | -1,023057651 | MMP16 |
| ENSGALG00000005256 | 0,880582454 | -0,63393871 | EPHA4 |
| ENSGALG00000009179 | 0,880187457 | -1,119360692 | TNFSF10 |
| ENSGALG00000011172 | 0,879786034 | -1,121518184 | MYO10L |
| ENSGALG00000027885 | 0,876800098 | -1,151437329 | C4orf19 |
| ENSGALG00000014783 | 0,875888949 | -0,572122303 | PTPRK |
| ENSGALG00000009405 | 0,875608773 | -1,391691484 | GRIA2 |
| ENSGALG00000006008 | 0,867963452 | 0,882382705 | HOMER2 |
| ENSGALG00000029023 | 0,865902724 | -1,242039059 | DMRTB1 |
| ENSGALG00000016909 | 0,864393738 | -0,488526858 | NDFIP2 |
| ENSGALG00000026582 | 0,863598751 | -0,997100015 | SOX18 |
| ENSGALG00000001617 | 0,855845667 | 1,898502237 | CAMKK1 |
| ENSGALG00000015180 | 0,851097646 | -0,540189844 | PSAT1 |
| ENSGALG00000016807 | 0,848101313 | -0,574724846 | LIMS1 |
| ENSGALG00000000977 | 0,842188766 | -1,032464943 | MAPKAP1 |
| ENSGALG00000009530 | 0,841159622 | -1,411853584 | PDZRN4 |
| ENSGALG00000004155 | 0,841077447 | -1,147734088 | MYOM3 |
| ENSGALG00000015409 | 0,837042297 | -0,599227418 | PROS1 |
| ENSGALG00000012278 | 0,835859144 | -2,383840075 | EMCN |

| | | | |
|---|---|---|---|
| ENSGALG00000010156 | 0,833499217 | -1,025083823 | DOCK8 |
| ENSGALG00000001220 | 0,831042878 | -0,818155857 | HERPUD1 |
| ENSGALG00000013085 | 0,830893947 | -0,535478159 | PTPRO |
| ENSGALG00000025770 | 0,830841194 | -1,089960517 | |
| ENSGALG00000012354 | 0,829092837 | -0,521764337 | VOPP1 |
| ENSGALG00000000639 | 0,827648848 | -0,79001541 | LRRN2 |
| ENSGALG00000011232 | 0,825691237 | -0,469477808 | STK11IP |
| ENSGALG00000020899 | 0,823141433 | -0,585391733 | |
| ENSGALG00000010664 | 0,817672769 | 0,852579615 | KIF26B |
| ENSGALG00000008670 | 0,815447938 | -0,486758564 | C1orf52 |
| ENSGALG00000013683 | 0,810200928 | -0,780228434 | ARID1B |
| ENSGALG00000012848 | 0,809997189 | -0,711139877 | KDM7A |
| ENSGALG00000014872 | 0,806209646 | -2,137267919 | FGF10 |
| ENSGALG00000005430 | 0,79963845 | -0,72235333 | ALS2CL |
| ENSGALG00000012849 | 0,79670503 | -0,57279907 | SLC37A3 |
| ENSGALG00000002443 | 0,789654091 | -1,353430658 | FBXL16 |
| ENSGALG00000009252 | 0,7810557 | -0,788959505 | PLD1 |
| ENSGALG00000002598 | 0,778822215 | -0,82047281 | TPST1 |
| ENSGALG00000004158 | 0,769428918 | -0,467330867 | GPT2 |
| ENSGALG00000001346 | 0,76776324 | -1,362109597 | |
| ENSGALG00000013124 | 0,765332021 | -1,026819624 | FHOD3 |
| ENSGALG00000011376 | 0,758366172 | -0,870171475 | ANKRD9 |
| ENSGALG00000001000 | 0,756534693 | -0,924720639 | HSPA5 |
| ENSGALG00000012012 | 0,743046962 | -0,895703636 | CMIP |
| ENSGALG00000011319 | 0,737544915 | -1,42017882 | LPHN3 |
| ENSGALG00000005512 | 0,736922168 | -0,667653797 | KIAA1671 |
| ENSGALG00000006138 | 0,736205494 | -1,026936244 | HELZ2 |
| ENSGALG00000003043 | 0,734167184 | -0,395518744 | BBS2 |
| ENSGALG00000005653 | 0,729875771 | -0,515161656 | NFKB2 |
| ENSGALG00000007501 | 0,729453053 | -0,702573548 | P2RX3 |
| ENSGALG00000015184 | 0,727266294 | -0,431383947 | TLE4 |
| ENSGALG00000009381 | 0,723727203 | -0,972226453 | SIPA1L1 |
| ENSGALG00000001370 | 0,722550699 | -1,073396904 | ADAMTS8 |
| ENSGALG00000015947 | 0,718924866 | -2,438977585 | CDH17 |
| ENSGALG00000008163 | 0,713636421 | -0,439350797 | PSME4 |
| ENSGALG00000015975 | 0,705892008 | -1,056406602 | DPY19L4 |
| ENSGALG00000015310 | 0,701805821 | -0,587354563 | SOBP |
| ENSGALG00000002198 | 0,701337674 | -0,485078535 | FHOD1 |
| ENSGALG00000010522 | 0,699893806 | -0,747786716 | ALDH18A1 |
| ENSGALG00000010204 | 0,699072042 | -0,660708753 | KCNC2 |
| ENSGALG00000012956 | 0,692712504 | 1,372804533 | TMTC1 |
| ENSGALG00000016835 | 0,685225343 | -0,539714594 | ATP11A |
| ENSGALG00000012696 | 0,682603698 | -0,443665071 | RANBP9 |
| ENSGALG00000016281 | 0,681257671 | -0,569360911 | DMD |
| ENSGALG00000003640 | 0,678918743 | 1,071788322 | FAM20C |
| ENSGALG00000027342 | 0,672830693 | -0,968330352 | METRN |
| ENSGALG00000027389 | 0,672548469 | -0,766788012 | RGS7BP |
| ENSGALG00000002646 | 0,671803119 | -0,515104787 | FAM171B |
| ENSGALG00000004612 | 0,671680213 | -0,579433401 | MTHFR |
| ENSGALG00000007704 | 0,666805577 | -0,445645488 | NPM3 |
| ENSGALG00000023876 | 0,666027019 | 0,598475769 | PRKCQ |
| ENSGALG00000015887 | 0,665885713 | -0,776552733 | URB1 |
| ENSGALG00000003957 | 0,665113393 | -1,565814231 | APOH |
| ENSGALG00000006872 | 0,664035354 | -0,823650254 | PISD |
| ENSGALG00000014957 | 0,661960443 | -0,81285735 | WDR1 |
| ENSGALG00000002655 | 0,657486198 | -0,468798244 | ITGAV |
| ENSGALG00000004231 | 0,654722209 | -0,911405982 | IFNLR1 |
| ENSGALG00000011770 | 0,65450966 | -0,525890593 | UMPS |
| ENSGALG00000016803 | 0,652622211 | -0,451555414 | ST6GAL2 |
| ENSGALG00000013031 | 0,641018743 | -0,574014118 | CECR1 |
| ENSGALG00000027835 | 0,637590923 | -0,430117051 | PCBD1 |
| ENSGALG00000024346 | 0,637233355 | 0,525612903 | FAM46B |
| ENSGALG00000008318 | 0,633974687 | -0,756789619 | SLK |
| ENSGALG00000004181 | 0,628709896 | -0,425938272 | GAA |

| | | | |
|---|---|---|---|
| ENSGALG00000014922 | 0,627329029 | -0,504425665 | ROCK1 |
| ENSGALG00000012726 | 0,625221924 | -0,941672551 | HSP90B1 |
| ENSGALG00000011435 | 0,614694006 | 0,35897775 | ELK3 |
| ENSGALG00000019397 | 0,608262914 | -0,975439675 | CERK |
| ENSGALG00000026468 | 0,607177306 | -0,514962132 | CHSY1 |
| ENSGALG00000000795 | 0,605895026 | -0,495778483 | MFSD12 |
| ENSGALG00000006939 | 0,603844144 | -0,728311968 | PFKP |
| ENSGALG00000016834 | 0,598292773 | -0,698351437 | MCF2L |
| ENSGALG00000011548 | 0,593504022 | -0,434202614 | PPP1R13B |
| ENSGALG00000002709 | 0,59270728 | -0,511853662 | MYO1C |
| ENSGALG00000008615 | 0,588449543 | -1,233824562 | LTK |
| ENSGALG00000011295 | 0,585889038 | -1,007421144 | SOCS2 |
| ENSGALG00000012979 | 0,583725131 | -0,528975634 | TRIO |
| ENSGALG00000007242 | 0,576491214 | -0,724284111 | SLITRK4 |
| ENSGALG00000011045 | 0,574987094 | -0,799075807 | AFF1 |
| ENSGALG00000020479 | 0,574556644 | -0,731996794 | RP11-872D17.8 |
| ENSGALG00000012429 | 0,572869136 | 1,680823476 | BMP4 |
| ENSGALG00000019961 | 0,57282511 | -0,7532835 | SLC18B1 |
| ENSGALG00000009415 | 0,570748764 | -0,939656238 | SMOC1 |
| ENSGALG00000011708 | 0,569570289 | -1,833902608 | MYLK |
| ENSGALG00000009156 | 0,56553435 | -0,682556058 | CDAN1 |
| ENSGALG00000012394 | 0,564570466 | -0,855781498 | NID2 |
| ENSGALG00000008293 | 0,562981441 | -0,66560113 | SH3PXD2A |
| ENSGALG00000008621 | 0,561672993 | -0,76612038 | NRP2 |
| ENSGALG00000003122 | 0,561038511 | 0,68839827 | ZFAND2A |
| ENSGALG00000015862 | 0,559415098 | -0,756350316 | CA2 |
| ENSGALG00000027757 | 0,559044043 | -1,076478944 | ARL4A |
| ENSGALG00000017462 | 0,558530219 | -0,763892042 | FNIP1 |
| ENSGALG00000000388 | 0,558245728 | -1,555950834 | LIN28A |
| ENSGALG00000007238 | 0,558183051 | 0,466556765 | PCSK6 |
| ENSGALG00000012852 | 0,557720187 | -0,42431457 | MKRN1 |
| ENSGALG00000016480 | 0,557041155 | -0,647091328 | SDC1 |
| ENSGALG00000028291 | 0,555590355 | -0,672824764 | BCAR1 |
| ENSGALG00000009433 | 0,551628222 | 0,528259009 | BAG3 |
| ENSGALG00000000680 | 0,550999971 | -1,411570576 | NUAK2 |
| ENSGALG00000002461 | 0,549280944 | -0,808165073 | MMP17 |
| ENSGALG00000015215 | 0,543882262 | -0,397390834 | MAPRE2 |
| ENSGALG00000017365 | 0,541410874 | -0,712049377 | REEP2 |
| ENSGALG00000014860 | 0,540418516 | -0,625239897 | YES1 |
| ENSGALG00000016781 | 0,539823119 | -0,504088725 | MAP4K4 |
| ENSGALG00000016883 | 0,537559856 | -0,555002491 | DOCK9 |
| ENSGALG00000011449 | 0,537091274 | -0,594721435 | |
| ENSGALG00000009855 | 0,536813312 | -0,567662024 | TBC1D9 |
| ENSGALG00000000891 | 0,531658469 | -0,533422492 | NSUN5 |
| ENSGALG00000015112 | 0,528998124 | -1,013051509 | ZNF521 |
| ENSGALG00000006002 | 0,517539205 | -0,422978689 | WHDC1 |
| ENSGALG00000011162 | 0,516837911 | -0,640538648 | CYP46A1 |
| ENSGALG00000000761 | 0,511636934 | -0,649687901 | TSKU |
| ENSGALG00000008811 | 0,510104856 | -0,931412058 | SLC1A4 |
| ENSGALG00000003668 | 0,508101808 | -0,85921128 | PITPNC1 |
| ENSGALG00000015458 | 0,50234495 | -0,453606012 | ILDR2 |
| ENSGALG00000017388 | 0,49802444 | -0,619582917 | YY1 |
| ENSGALG00000009503 | 0,49017284 | -0,547842813 | RAB3GAP2 |
| ENSGALG00000000936 | 0,490067343 | -1,193444435 | LMX1B |
| ENSGALG00000016768 | 0,489550679 | -0,54750013 | LONRF2 |
| ENSGALG00000001962 | 0,488180846 | -0,330389801 | PTBP1 |
| ENSGALG00000002316 | 0,486190455 | -0,700708149 | NUBP2 |
| ENSGALG00000005727 | 0,485528575 | -0,548590838 | SLC7A3 |
| ENSGALG00000013661 | 0,485373842 | -0,40778269 | MICU3 |
| ENSGALG00000006237 | 0,48464383 | -0,616678741 | PKN2 |
| ENSGALG00000006121 | 0,483017989 | -0,468808823 | PIK3C2A |
| ENSGALG00000003835 | 0,482616746 | -0,552761335 | ATP2A2 |
| ENSGALG00000028128 | 0,475145623 | 0,378077792 | HOXD8 |

| | | | |
|---|---|---|---|
| ENSGALG00000014953 | 0,474792932 | -0,429472364 | RWDD1 |
| ENSGALG00000014873 | 0,474224225 | -0,6514166 | GJA1 |
| ENSGALG00000009532 | 0,469457498 | -1,242374232 | GXYLT1 |
| ENSGALG00000016808 | 0,46860233 | -0,615970468 | RANBP2 |
| ENSGALG00000002726 | 0,464112357 | -0,708755608 | KIF1B |
| ENSGALG00000004144 | 0,459129046 | -0,588131949 | DDX50 |
| ENSGALG00000009850 | 0,458569963 | -0,63459607 | NOVA1 |
| ENSGALG00000014688 | 0,458090231 | -0,396059354 | ZYX |
| ENSGALG00000013480 | 0,457972294 | -0,615680556 | MTHFD2 |
| ENSGALG00000015940 | 0,457737572 | -0,972506334 | TMEM67 |
| ENSGALG00000005134 | 0,45766228 | -0,483437706 | ZSWIM8 |
| ENSGALG00000003295 | 0,455390197 | -1,063705711 | ADAMTS12 |
| ENSGALG00000008521 | 0,447794364 | -0,395949661 | TP53BP1 |
| ENSGALG00000006947 | 0,446651103 | 0,64187784 | BDH1 |
| ENSGALG00000001596 | 0,445911724 | -0,383722949 | MLLT1 |
| ENSGALG00000009300 | 0,444814977 | -0,413699825 | NUMB |
| ENSGALG00000005785 | 0,442398124 | -0,791416097 | UBAP2 |
| ENSGALG00000004169 | 0,441293938 | -0,613558741 | |
| ENSGALG00000001957 | 0,438219772 | -0,541703645 | ITCH |
| ENSGALG00000013861 | 0,437973226 | -0,913164571 | TNFAIP3 |
| ENSGALG00000014419 | 0,437804845 | -0,730417334 | SLIT2 |
| ENSGALG00000011464 | 0,435819807 | -0,582311331 | EIF5 |
| ENSGALG00000011643 | 0,435706974 | -0,816325927 | UTP20 |
| ENSGALG00000011707 | 0,43414341 | -0,516018775 | UBA5 |
| ENSGALG00000014686 | 0,43250268 | -0,691021625 | FBN2 |
| ENSGALG00000009534 | 0,430698068 | -0,408065762 | PLEKHA1 |
| ENSGALG00000001376 | 0,43069552 | 0,45012042 | ARID3B |
| ENSGALG00000007659 | 0,429907921 | -0,45246564 | HYOU1 |
| ENSGALG00000014784 | 0,427987842 | -0,520975246 | MAST4 |
| ENSGALG00000004638 | 0,427666392 | -0,48870476 | HEPH |
| ENSGALG00000010975 | 0,425801788 | -0,624280901 | TMTC2 |
| ENSGALG00000011246 | 0,419525625 | -0,471179653 | CHPF |
| ENSGALG00000004531 | 0,419189684 | -0,458962037 | PLOD1 |
| ENSGALG00000020805 | 0,416086722 | -0,719835862 | LRRC8D |
| ENSGALG00000005723 | 0,412949734 | -0,890647194 | RP11-96O20.4 |
| ENSGALG00000007141 | 0,412800257 | -0,70907976 | LRRK1 |
| ENSGALG00000005796 | 0,407045166 | -0,664735985 | XBP1 |
| ENSGALG00000023451 | 0,405630741 | -0,638494594 | TRPM7 |
| ENSGALG00000012814 | 0,404889777 | -0,898069575 | PXDC1 |
| ENSGALG00000028267 | 0,403950335 | -0,473386218 | GOLPH3 |
| ENSGALG00000017085 | 0,403507696 | -0,511832688 | SLC7A1 |
| ENSGALG00000009748 | 0,403044512 | -0,762363604 | ASNS |
| ENSGALG00000011101 | 0,40109735 | -0,47005983 | URB2 |
| ENSGALG00000008195 | 0,394418985 | -0,378910892 | NPHP1 |
| ENSGALG00000002763 | 0,38996775 | -0,423193935 | FRZB |
| ENSGALG00000014902 | 0,387988657 | -0,650325285 | RP1-179P9.3 |
| ENSGALG00000008708 | 0,384060223 | -0,423296325 | ESF1 |
| ENSGALG00000004200 | 0,383702067 | -0,347970886 | DPP9 |
| ENSGALG00000019077 | 0,383662275 | -1,030078317 | |
| ENSGALG00000015765 | 0,383191974 | -0,587270267 | POLR1A |
| ENSGALG00000000464 | 0,378042232 | 0,587838297 | RHOBTB2 |
| ENSGALG00000008240 | 0,376593038 | -0,670953542 | CFLAR |
| ENSGALG00000014213 | 0,372421493 | -0,596053951 | PRR5 |
| ENSGALG00000009428 | 0,369028406 | -0,792261942 | SUSD6 |
| ENSGALG00000003180 | 0,367625634 | -0,573999724 | TSR1 |
| ENSGALG00000006336 | 0,365567709 | -0,452052929 | PAXIP1 |
| ENSGALG00000005845 | 0,364474801 | -0,655210424 | SLC7A5 |
| ENSGALG00000004696 | 0,360778491 | -0,381141883 | LEO1 |
| ENSGALG00000009562 | 0,359101036 | -0,714412903 | PLEK2 |
| ENSGALG00000007145 | 0,358001987 | -0,440432711 | ITGB1 |
| ENSGALG00000010603 | 0,356862546 | -0,604746743 | CEBPZ |
| ENSGALG00000003563 | 0,354826794 | -0,364389392 | YARS |
| ENSGALG00000005160 | 0,351320099 | -0,454934871 | VMP1 |
| ENSGALG00000002954 | 0,350618821 | -0,457794297 | MBTD1 |

| | | | |
|---|---|---|---|
| ENSGALG00000001749 | 0,34679817 | -0,88875485 | ACSBG2 |
| ENSGALG00000003427 | 0,346212827 | -0,813323099 | USP3 |
| ENSGALG00000009540 | 0,344720194 | -0,9229711 | YAF2 |
| ENSGALG00000006437 | 0,344612756 | -0,682364759 | NOM1 |
| ENSGALG00000001003 | 0,344161191 | -0,600081977 | SEPN1 |
| ENSGALG00000007234 | 0,343143822 | -0,505755971 | CLCN5 |
| ENSGALG00000025884 | 0,339208538 | -0,543082298 | NABP1 |
| ENSGALG00000026349 | 0,337656858 | -0,736634172 | HSD17B12 |
| ENSGALG00000012202 | 0,337252452 | -0,341191282 | KBTBD2 |
| ENSGALG00000010976 | 0,337203087 | -0,434636867 | CCDC126 |
| ENSGALG00000006975 | 0,33439022 | -0,659116429 | TBC1D22A |
| ENSGALG00000003767 | 0,331888441 | -0,636798593 | NKD1 |
| ENSGALG00000006256 | 0,33180927 | -0,600800071 | MAP7D3 |
| ENSGALG00000015049 | 0,323309666 | -0,441268065 | AMD1 |
| ENSGALG00000004132 | 0,317911072 | -0,797075856 | UHRF1 |
| ENSGALG00000009448 | 0,31659087 | -0,43880669 | IFRD1 |
| ENSGALG00000000258 | 0,314352619 | -0,649481888 | HEATR1 |
| ENSGALG00000009992 | 0,311546697 | -0,521017168 | ARHGAP5 |
| ENSGALG00000008837 | 0,311145069 | -0,471745405 | PELI1 |
| ENSGALG00000008265 | 0,310830618 | -0,619236444 | PDCD11 |
| ENSGALG00000011228 | 0,310544896 | -0,565699287 | WARS |
| ENSGALG00000017111 | 0,309121441 | -0,428517877 | NUP58 |
| ENSGALG00000006971 | 0,308524903 | -0,549657911 | CDK5RAP2 |
| ENSGALG00000003039 | 0,308092774 | -0,372512284 | SRM |
| ENSGALG00000000346 | 0,306247189 | -0,429865287 | ZNF606 |
| ENSGALG00000002863 | 0,306238468 | -0,447764784 | ERGIC1 |
| ENSGALG00000010349 | 0,300633215 | -0,358393493 | POLR1C |
| ENSGALG00000007387 | 0,298075088 | -0,330955786 | GTF3C1 |
| ENSGALG00000011357 | 0,296426831 | -0,387039211 | VEZT |
| ENSGALG00000010536 | 0,29615271 | -0,646034394 | FAF1 |
| ENSGALG00000011400 | 0,295706421 | -0,550294288 | USP44 |
| ENSGALG00000012138 | 0,295412949 | -0,402996238 | sept-07 |
| ENSGALG00000002985 | 0,293105421 | -0,378678888 | UTP18 |
| ENSGALG00000003391 | 0,293097669 | -0,827807721 | EIF2AK1 |
| ENSGALG00000006856 | 0,290795459 | -0,537151634 | CD151 |
| ENSGALG00000006895 | 0,290027475 | -0,543915173 | CEP131 |
| ENSGALG00000016910 | 0,283615388 | -0,572230903 | RBM26 |
| ENSGALG00000009325 | 0,276324484 | -0,363294006 | OLA1 |
| ENSGALG00000003339 | 0,27482717 | -0,366040447 | MTOR |
| ENSGALG00000010481 | 0,274549325 | -0,330642085 | CMPK1 |
| ENSGALG00000005243 | 0,272933583 | -0,597738338 | PPP3CB |
| ENSGALG00000009782 | 0,272414043 | -0,507579083 | NAA15 |
| ENSGALG00000011596 | 0,272230864 | -0,519655791 | PTPN4 |
| ENSGALG00000006935 | 0,270244925 | -0,506724591 | BTAF1 |
| ENSGALG00000003078 | 0,264245742 | -0,483195985 | MMD |
| ENSGALG00000026300 | -0,257041042 | 0,345400821 | |
| ENSGALG00000020003 | -0,275409017 | -0,59891041 | MAP3K4 |
| ENSGALG00000027963 | -0,279887477 | 0,451771641 | COX7C |
| ENSGALG00000027710 | -0,296769112 | 0,421872733 | DAG1 |
| ENSGALG00000015326 | -0,301560094 | 0,524363909 | CLTA |
| ENSGALG00000006094 | -0,306148177 | -0,454736184 | PHF6 |
| ENSGALG00000012121 | -0,306219207 | -0,541445831 | RCN1 |
| ENSGALG00000012546 | -0,312592383 | 0,318249627 | MCM5 |
| ENSGALG00000026437 | -0,314002706 | 0,703536871 | |
| ENSGALG00000025764 | -0,316576106 | 0,675250138 | FKBP10 |
| ENSGALG00000006483 | -0,318521408 | -0,399267586 | AMOTL2 |
| ENSGALG00000013784 | -0,32170776 | 0,339967639 | CDT1 |
| ENSGALG00000016870 | -0,322919481 | 0,454001604 | TMTC4 |
| ENSGALG00000020528 | -0,323800436 | 0,424819785 | INIP |
| ENSGALG00000004621 | -0,328680536 | -0,512109985 | PREX1 |
| ENSGALG00000028470 | -0,330867645 | 0,39443368 | GSR |
| ENSGALG00000021365 | -0,33098251 | 0,432965219 | DCTN3 |
| ENSGALG00000025818 | -0,334464872 | 0,571475138 | KCNK5 |
| ENSGALG00000004581 | -0,341091866 | 0,466855623 | NDST1 |

| | | | |
|---|---|---|---|
| ENSGALG00000000684 | -0,343510226 | -0,552570682 | KLHDC8A |
| ENSGALG00000008758 | -0,348320506 | -0,696392948 | B3GNT5 |
| ENSGALG00000010674 | -0,348433039 | -0,399166032 | DCTD |
| ENSGALG00000016034 | -0,349640251 | -0,4051791 | ATRN |
| ENSGALG00000014970 | -0,352181226 | 0,397421157 | FSTL1 |
| ENSGALG00000000921 | -0,356222635 | 0,454646053 | SLC9A1 |
| ENSGALG00000017329 | -0,359975454 | 0,44555284 | FCHSD2 |
| ENSGALG00000026081 | -0,379250879 | 0,588777417 | RBM38 |
| ENSGALG00000015469 | -0,382238702 | 1,115704157 | IGSF3 |
| ENSGALG00000006702 | -0,38370743 | 0,471728545 | MFGE8 |
| ENSGALG00000024047 | -0,387381439 | -0,380223912 | MYCL |
| ENSGALG00000006606 | -0,392475054 | 1,610976489 | KIAA1324L |
| ENSGALG00000003457 | -0,397746039 | 0,662964808 | CNP |
| ENSGALG00000013067 | -0,398595631 | 0,494753671 | AKR1B1 |
| ENSGALG00000028567 | -0,405726705 | 0,515072598 | MYL9 |
| ENSGALG00000026870 | -0,410373931 | 0,966911139 | TMEM200C |
| ENSGALG00000016152 | -0,412571324 | 0,558712272 | LY6E |
| ENSGALG00000011778 | -0,415058611 | 0,494085012 | ITGB5 |
| ENSGALG00000011238 | -0,417063936 | 0,316616268 | WLS |
| ENSGALG00000013002 | -0,420522444 | 0,734684533 | DAP |
| ENSGALG00000005237 | -0,426583983 | -0,342909039 | FGD3 |
| ENSGALG00000005330 | -0,431988153 | 1,328511787 | SNCAIP |
| ENSGALG00000016328 | -0,434072431 | 1,146952103 | GSTA |
| ENSGALG00000026113 | -0,434078932 | 0,523056553 | CARHSP1 |
| ENSGALG00000029174 | -0,450699904 | 1,589321167 | |
| ENSGALG00000006036 | -0,461900797 | 0,644947298 | |
| ENSGALG00000022926 | -0,463163963 | 0,576442566 | GPR162 |
| ENSGALG00000017119 | -0,467363832 | -0,420708881 | TNFRSF19 |
| ENSGALG00000006885 | -0,469804206 | 0,574664846 | HRAS |
| ENSGALG00000010422 | -0,470203966 | 0,646628908 | FEZ1 |
| ENSGALG00000007710 | -0,471105348 | -0,637223247 | |
| ENSGALG00000027373 | -0,47216803 | -0,413207621 | |
| ENSGALG00000015107 | -0,4734568 | 0,542439326 | PIP5K1B |
| ENSGALG00000012658 | -0,475116315 | 1,214376265 | TCP11L2 |
| ENSGALG00000015660 | -0,475673447 | 0,831570906 | SNX30 |
| ENSGALG00000000316 | -0,476826949 | 0,549437781 | PKNOX2 |
| ENSGALG00000029011 | -0,483499319 | -0,444941401 | SLC35F1 |
| ENSGALG00000010316 | -0,484012085 | 0,662660688 | FRAS1 |
| ENSGALG00000001966 | -0,490465312 | 0,545673399 | RP11-106M3.5 |
| ENSGALG00000001816 | -0,495397442 | 0,414935997 | MAGI3 |
| ENSGALG00000009207 | -0,499904522 | -0,518081559 | TRIM2 |
| ENSGALG00000026505 | -0,503934058 | 0,837091894 | RDH10 |
| ENSGALG00000002777 | -0,509947463 | 0,524514209 | ACAA2 |
| ENSGALG00000025977 | -0,513209442 | 0,502103804 | GADD45A |
| ENSGALG00000008728 | -0,517913117 | 0,625207283 | PTER |
| ENSGALG00000022713 | -0,525946923 | 0,539793648 | FAM181B |
| ENSGALG00000005246 | -0,527306483 | -0,7272373 | PAX3 |
| ENSGALG00000010982 | -0,53129631 | 0,55035484 | NID1 |
| ENSGALG00000006480 | -0,534658874 | 0,810690895 | TCF7 |
| ENSGALG00000016551 | -0,548003277 | 1,321564075 | AC025048.1 |
| ENSGALG00000008433 | -0,549541331 | 0,732487 | FZD2 |
| ENSGALG00000013964 | -0,55034782 | 0,602864404 | SPATA18 |
| ENSGALG00000004323 | -0,5545441 | 0,7637615 | RASAL2 |
| ENSGALG00000015987 | -0,554680878 | 0,445391941 | TP53INP1 |
| ENSGALG00000008871 | -0,555142332 | -0,492812674 | LPHN2 |
| ENSGALG00000004593 | -0,555306466 | 0,839985137 | SULF2 |
| ENSGALG00000014242 | -0,557782152 | 0,583190799 | BEND4 |
| ENSGALG00000012710 | -0,557841328 | 0,660750331 | NCS1 |
| ENSGALG00000004537 | -0,559387674 | 0,454616982 | EFNB1 |
| ENSGALG00000028795 | -0,566961474 | 1,118039364 | CADM3 |
| ENSGALG00000004741 | -0,573167729 | 0,509578858 | EPHB2 |
| ENSGALG00000004256 | -0,575998874 | -0,396589984 | DHRS3 |
| ENSGALG00000003404 | -0,585273519 | 1,24748878 | TFEB |
| ENSGALG00000002707 | -0,585668905 | 0,58133458 | CHRNB2 |

| | | | |
|---|---|---|---|
| ENSGALG00000008961 | -0,593936394 | -0,780468343 | PTPRZ1 |
| ENSGALG00000006409 | -0,594856883 | -0,436770571 | PODXL |
| ENSGALG00000026180 | -0,59981586 | 0,591162305 | MSX2 |
| ENSGALG00000012712 | -0,607252424 | 0,742584862 | RBM24 |
| ENSGALG00000016251 | -0,611728855 | 1,000647408 | TSPAN7 |
| ENSGALG00000002974 | -0,611833543 | -0,529999669 | SPHKAP |
| ENSGALG00000008367 | -0,614602858 | 0,522250337 | MDK |
| ENSGALG00000010551 | -0,614933753 | 0,911984879 | CDC42EP3 |
| ENSGALG00000002652 | -0,621511478 | -0,509052755 | FZD10 |
| ENSGALG00000003582 | -0,622442826 | -1,218636528 | SLC1A3 |
| ENSGALG00000007076 | -0,624816157 | 0,617550884 | CCNY |
| ENSGALG00000016391 | -0,625110592 | -0,569812042 | CNKSR2 |
| ENSGALG00000027377 | -0,626128444 | 0,469161575 | NKAIN3 |
| ENSGALG00000015729 | -0,630253804 | 0,747405326 | LPAR1 |
| ENSGALG00000007748 | -0,63248212 | 0,758936347 | SLC16A2 |
| ENSGALG00000000040 | -0,633128633 | 0,669045681 | FAM195A |
| ENSGALG00000010320 | -0,636363776 | 0,704383823 | PAWR |
| ENSGALG00000006152 | -0,63795304 | 0,657637784 | |
| ENSGALG00000002437 | -0,638363537 | 0,993124174 | FAM69B |
| ENSGALG00000010269 | -0,639329258 | 0,886301871 | KCNAB1 |
| ENSGALG00000006826 | -0,641184602 | -0,530661584 | KLHL25 |
| ENSGALG00000028543 | -0,64250805 | 0,614319385 | FGFR4 |
| ENSGALG00000012577 | -0,644333952 | 0,956309243 | FRMD3 |
| ENSGALG00000015241 | -0,649652118 | -0,655397927 | SNAI2 |
| ENSGALG00000017036 | -0,649834759 | 1,353726483 | LHFP |
| ENSGALG00000009439 | -0,662995235 | 0,751125582 | GPR85 |
| ENSGALG00000005628 | -0,664092027 | 0,601333554 | COL9A3 |
| ENSGALG00000005657 | -0,664586696 | 0,780089243 | CRHR2 |
| ENSGALG00000007732 | -0,671216647 | 1,306274335 | RTN4R |
| ENSGALG00000010119 | -0,672877793 | 0,900097678 | CDKL2 |
| ENSGALG00000004309 | -0,673847887 | 0,913060877 | GDAP1L1 |
| ENSGALG00000005774 | -0,674007323 | -0,555555373 | NKAIN4 |
| ENSGALG00000017106 | -0,674065701 | 1,082115941 | ATP8A2 |
| ENSGALG00000016945 | -0,674331354 | 0,953551228 | LECT1 |
| ENSGALG00000006555 | -0,681158051 | -0,729109536 | IGF2 |
| ENSGALG00000001727 | -0,689838866 | 1,445847057 | LZTS1 |
| ENSGALG00000023073 | -0,692447843 | 0,353608475 | AKAP12 |
| ENSGALG00000009626 | -0,694120539 | -0,375704266 | THBS1 |
| ENSGALG00000003580 | -0,694931491 | 1,284312011 | MMP2 |
| ENSGALG00000002818 | -0,699438203 | 0,595523293 | VAT1 |
| ENSGALG00000003589 | -0,699453771 | 0,652278144 | VTN |
| ENSGALG00000006882 | -0,700351611 | 0,976472658 | ST8SIA2 |
| ENSGALG00000004676 | -0,710261932 | 0,910530695 | BTG3 |
| ENSGALG00000015926 | -0,714450321 | 0,934566237 | RUNX1T1 |
| ENSGALG00000025919 | -0,717706563 | 0,857090747 | BAMBI |
| ENSGALG00000002016 | -0,719060037 | 1,156233761 | PRKCD |
| ENSGALG00000000674 | -0,719912044 | 0,67836471 | MYO1D |
| ENSGALG00000005608 | -0,721159959 | 0,44662731 | SLC9A3R2 |
| ENSGALG00000002749 | -0,723137568 | 0,687223145 | RP11-565P22.6 |
| ENSGALG00000001449 | -0,723393814 | 0,740070536 | STRA6 |
| ENSGALG00000006792 | -0,726229187 | 1,109047707 | VSTM2L |
| ENSGALG00000009020 | -0,742019908 | -0,446477137 | JAG1 |
| ENSGALG00000009536 | -0,742471798 | 0,851130737 | |
| ENSGALG00000015448 | -0,755416575 | -0,533672096 | TOX |
| ENSGALG00000005626 | -0,756044449 | 0,92496767 | TPST2 |
| ENSGALG00000000690 | -0,756893418 | -0,657633862 | GDPD4 |
| ENSGALG00000015768 | -0,757420183 | 0,855142896 | ANKRD6 |
| ENSGALG00000006793 | -0,758190034 | 0,898163749 | NTRK3 |
| ENSGALG00000002742 | -0,7644405 | 0,847416181 | TMEM132B |
| ENSGALG00000015297 | -0,764510224 | 0,81224071 | FOXO3 |
| ENSGALG00000016543 | -0,766325509 | 0,654951402 | NHS |
| ENSGALG00000006192 | -0,767192565 | 0,958606154 | USH1C |
| ENSGALG00000007623 | -0,778310378 | 0,788173273 | CACNA1G |
| ENSGALG00000026042 | -0,780763307 | 0,987626868 | SHISA3 |

| | | | |
|---|---|---|---|
| ENSGALG00000012191 | -0,787037882 | 0,72968259 | LGR4 |
| ENSGALG00000012652 | -0,806982417 | 1,438231617 | KCNG2 |
| ENSGALG00000016223 | -0,80722926 | 1,013131958 | MAOB |
| ENSGALG00000004844 | -0,807613149 | 0,581688828 | SEMA6D |
| ENSGALG00000004986 | -0,810825031 | -0,430376609 | CERCAM |
| ENSGALG00000004472 | -0,814406115 | 0,672299012 | ASTN1 |
| ENSGALG00000017374 | -0,814717413 | 1,149174538 | SLITRK3 |
| ENSGALG00000009514 | -0,816414808 | -0,779501664 | NRCAM |
| ENSGALG00000012715 | -0,842621875 | 0,978917656 | CAP2 |
| ENSGALG00000015084 | -0,852219609 | 1,480304896 | IGSF11 |
| ENSGALG00000006873 | -0,854385087 | 0,565449117 | RASSF7 |
| ENSGALG00000009052 | -0,861077306 | 0,987910707 | LBH |
| ENSGALG00000026192 | -0,861152656 | -0,473341595 | ST8SIA4 |
| ENSGALG00000028946 | -0,867278013 | 1,217039733 | CSRP2 |
| ENSGALG00000008297 | -0,875877387 | 0,379216929 | SEMA4B |
| ENSGALG00000011709 | -0,876476185 | 0,918333355 | TMEM108 |
| ENSGALG00000003345 | -0,878084006 | 1,211546204 | PLCD4 |
| ENSGALG00000023738 | -0,882263773 | 0,704601132 | |
| ENSGALG00000027501 | -0,88631831 | 1,108782799 | ANPEP |
| ENSGALG00000003426 | -0,891859274 | -0,587203746 | RHBDL3 |
| ENSGALG00000001129 | -0,899923251 | 0,871868135 | KIRREL3 |
| ENSGALG00000015186 | -0,900331794 | 1,345677898 | KLHL14 |
| ENSGALG00000006245 | -0,900397348 | 1,589094556 | KLF15 |
| ENSGALG00000009603 | -0,909672796 | 0,844955114 | |
| ENSGALG00000028234 | -0,912907561 | 1,03829388 | SYNDIG1 |
| ENSGALG00000015967 | -0,91657646 | 0,936813576 | LRRTM1 |
| ENSGALG00000020688 | -0,921010154 | 0,678030575 | CYP4B1L |
| ENSGALG00000020376 | -0,925271568 | 0,781084057 | |
| ENSGALG00000007078 | -0,925877896 | 1,270778013 | ENTPD1 |
| ENSGALG00000005539 | -0,927364348 | 0,77704495 | ITM2C |
| ENSGALG00000006346 | -0,927790987 | 0,756405715 | CXCL14 |
| ENSGALG00000020575 | -0,930944308 | 0,690014438 | ERMN |
| ENSGALG00000009237 | -0,942598283 | 1,124183808 | TLR2A |
| ENSGALG00000009042 | -0,948736093 | 0,719468223 | ISM1 |
| ENSGALG00000004849 | -0,955100006 | 0,553445556 | WSCD2 |
| ENSGALG00000011200 | -0,962880109 | 1,609360966 | THBS2 |
| ENSGALG00000006507 | -0,967854239 | 0,793097237 | CLMP |
| ENSGALG00000026153 | -0,984548638 | 0,755983234 | FOXO6 |
| ENSGALG00000001795 | -0,995172186 | 1,261084777 | GFRA2 |
| ENSGALG00000002832 | -0,995334878 | 1,610990532 | IFI35 |
| ENSGALG00000019280 | -1,005672086 | 1,199070585 | LMO3 |
| ENSGALG00000010500 | -1,016915026 | 0,828235743 | NKX6-2 |
| ENSGALG00000027891 | -1,030686173 | 0,504324315 | NREP |
| ENSGALG00000020788 | -1,041075867 | -0,986044977 | CCDC141 |
| ENSGALG00000008455 | -1,05482755 | 0,695894902 | SEMA3C |
| ENSGALG00000009601 | -1,061491773 | 0,878730197 | NELL2 |
| ENSGALG00000003588 | -1,081475528 | 0,911216657 | |
| ENSGALG00000009638 | -1,085140036 | 0,988233263 | ANO6 |
| ENSGALG00000007908 | -1,09305059 | 1,477721596 | EFEMP1 |
| ENSGALG00000013948 | -1,099632212 | 0,760675086 | RASL11B |
| ENSGALG00000028682 | -1,101427497 | 0,990483134 | |
| ENSGALG00000008440 | -1,108083257 | 1,543692993 | CHST1 |
| ENSGALG00000007211 | -1,108214376 | 1,44654086 | CDH22 |
| ENSGALG00000028506 | -1,110787668 | 0,637581477 | |
| ENSGALG00000002028 | -1,145532965 | 0,675138088 | GPRC5B |
| ENSGALG00000004448 | -1,158751146 | 1,504790559 | BRINP2 |
| ENSGALG00000005967 | -1,230002366 | 1,545091384 | LMO1 |
| ENSGALG00000014258 | -1,256488908 | 1,040436704 | LIMCH1 |
| ENSGALG00000023577 | -1,257090663 | 2,257832359 | TBX2 |
| ENSGALG00000016411 | -1,262862187 | 0,657856223 | COL14A1 |
| ENSGALG00000009604 | -1,349943691 | 0,581479374 | DBX2 |
| ENSGALG00000011205 | -1,462754207 | 0,836341019 | SMOC2 |

**Supplementary Table 4:** MEK1ca+HOXB8 versus MEK1ca differentially regulated genes (False Discovery Rate (padj) <= 0.005)

| ID | baseMean | log2FoldChange | lfcSE | stat | pvalue | padj | Symbol |
|---|---|---|---|---|---|---|---|
| ENSGALG00000026853 | 1756,7605 | 5,751530736 | 0,1962667 | 29,304673 | 9,04E-189 | 6,16E-185 | FGF3 |
| ENSGALG00000011433 | 901,86118 | 4,353428748 | 0,1979937 | 21,987719 | 3,77E-107 | 1,28E-103 | VIL1 |
| ENSGALG00000009778 | 267,92271 | 4,29765073 | 0,2973351 | 14,453898 | 2,37E-47 | 9,77E-45 | HCLS1 |
| ENSGALG00000011582 | 175,17021 | 4,101846269 | 0,3182796 | 12,887557 | 5,29E-38 | 1,53E-35 | SLC17A8 |
| ENSGALG00000027742 | 312,3427 | 3,96517198 | 0,2663087 | 14,889379 | 3,86E-50 | 1,81E-47 | KLHL33 |
| ENSGALG00000028044 | 194,51466 | 3,90733608 | 0,3041023 | 12,848755 | 8,74E-38 | 2,48E-35 | ADMP |
| ENSGALG00000002108 | 813,37463 | 3,799830272 | 0,1965883 | 19,328872 | 3,07E-83 | 5,23E-80 | EVPL |
| ENSGALG00000021395 | 138,40693 | 3,681673719 | 0,3287873 | 11,197736 | 4,18E-29 | 6,70E-27 | |
| ENSGALG00000001080 | 589,09501 | 3,620989798 | 0,2231106 | 16,229571 | 3,12E-59 | 2,02E-56 | NR5A1 |
| ENSGALG00000005292 | 233,99511 | 3,593969871 | 0,2819274 | 12,747854 | 3,20E-37 | 8,90E-35 | KLHL40 |
| ENSGALG00000028376 | 1521,8358 | 3,47510163 | 0,1982878 | 17,525546 | 9,15E-69 | 7,78E-66 | FGF19 |
| ENSGALG00000023843 | 120,84193 | 3,393126537 | 0,3307759 | 10,258084 | 1,09E-24 | 1,35E-22 | FRMD7 |
| ENSGALG00000006471 | 279,29896 | 3,309144864 | 0,2889251 | 11,453298 | 2,26E-30 | 4,04E-28 | SLCO2A1 |
| ENSGALG00000028304 | 118,7931 | 3,251722149 | 0,3347604 | 9,7135804 | 2,64E-22 | 2,58E-20 | |
| ENSGALG00000004526 | 421,69106 | 3,217263436 | 0,2437389 | 13,199628 | 8,82E-40 | 2,93E-37 | TNR |
| ENSGALG00000008069 | 109,41167 | 3,182915984 | 0,3413839 | 9,3235675 | 1,12E-20 | 9,51E-19 | CYP2J6L3 |
| ENSGALG00000004134 | 133,43428 | 3,164728278 | 0,3267271 | 9,6861515 | 3,45E-22 | 3,31E-20 | GLRA1 |
| ENSGALG00000012254 | 87,731989 | 3,162704218 | 0,3459847 | 9,1411679 | 6,18E-20 | 4,98E-18 | KCNJ4 |
| ENSGALG00000006944 | 469,14467 | 3,116432937 | 0,2398381 | 12,993902 | 1,32E-38 | 4,01E-36 | |
| ENSGALG00000028612 | 103,60478 | 3,070056483 | 0,3464928 | 8,8603749 | 7,97E-19 | 5,81E-17 | MYH1G |
| ENSGALG00000005285 | 100,23743 | 2,996752626 | 0,3402533 | 8,8074179 | 1,28E-18 | 8,94E-17 | TBX4 |
| ENSGALG00000007335 | 107,29888 | 2,985281892 | 0,3405577 | 8,7658615 | 1,85E-18 | 1,28E-16 | KBTBD13 |
| ENSGALG00000001256 | 266,94499 | 2,963677659 | 0,2979331 | 9,9474608 | 2,59E-23 | 2,71E-21 | NLRC5 |
| ENSGALG00000016685 | 93,447456 | 2,945393761 | 0,3383685 | 8,7046936 | 3,18E-18 | 2,13E-16 | ASMT |
| ENSGALG00000016721 | 160,3282 | 2,940591405 | 0,3042022 | 9,6665674 | 4,18E-22 | 3,93E-20 | PLCXD1 |
| ENSGALG00000026167 | 171,61409 | 2,925839411 | 0,3177833 | 9,2070258 | 3,35E-20 | 2,78E-18 | PIK3R5 |
| ENSGALG00000002926 | 76,22895 | 2,900739748 | 0,3500461 | 8,2867354 | 1,16E-16 | 6,86E-15 | DBH |
| ENSGALG00000004700 | 97,840654 | 2,88173675 | 0,3343593 | 8,6186822 | 6,77E-18 | 4,45E-16 | NCF2 |
| ENSGALG00000016722 | 73,43112 | 2,829128945 | 0,3527315 | 8,0206314 | 1,05E-15 | 5,52E-14 | RGNL |
| ENSGALG00000012830 | 227,95073 | 2,760094911 | 0,2851044 | 9,6809972 | 3,63E-22 | 3,44E-20 | IRF4 |
| ENSGALG00000027247 | 97,436329 | 2,75413056 | 0,3327446 | 8,2770098 | 1,26E-16 | 7,41E-15 | EOMES |
| ENSGALG00000002823 | 276,03395 | 2,746001119 | 0,2565217 | 10,70475 | 9,67E-27 | 1,39E-24 | |
| ENSGALG00000006614 | 540,92661 | 2,721769719 | 0,2601033 | 10,464189 | 1,26E-25 | 1,70E-23 | TECTA |
| ENSGALG00000006744 | 446,95073 | 2,709854582 | 0,2639156 | 10,267881 | 9,83E-25 | 1,23E-22 | MUC2 |
| ENSGALG00000020485 | 352,82533 | 2,707785881 | 0,2864956 | 9,4514033 | 3,34E-21 | 2,90E-19 | |
| ENSGALG00000015322 | 401,95748 | 2,694878773 | 0,2468716 | 10,916117 | 9,65E-28 | 1,43E-25 | IMPG2 |
| ENSGALG00000025886 | 234,93273 | 2,681244585 | 0,2769653 | 9,6807963 | 3,64E-22 | 3,44E-20 | SUSD3 |
| ENSGALG00000013922 | 145,69265 | 2,651462599 | 0,3144764 | 8,4313551 | 3,42E-17 | 2,11E-15 | ZP1 |
| ENSGALG00000026590 | 256,39028 | 2,649598672 | 0,2655285 | 9,9785844 | 1,89E-23 | 2,01E-21 | KLF13 |
| ENSGALG00000016325 | 669,81495 | 2,632246576 | 0,2229691 | 11,805432 | 3,66E-32 | 7,66E-30 | GSTA3 |
| ENSGALG00000026627 | 138,55984 | 2,620280418 | 0,325291 | 8,0551894 | 7,94E-16 | 4,29E-14 | |
| ENSGALG00000001780 | 98,3971 | 2,613843166 | 0,3358627 | 7,7824758 | 7,11E-15 | 3,32E-13 | PROC |
| ENSGALG00000002987 | 118,45636 | 2,607861451 | 0,3206775 | 8,1323505 | 4,21E-16 | 2,34E-14 | SERPINF2 |
| ENSGALG00000027059 | 200,39104 | 2,606173672 | 0,2890015 | 9,0178542 | 1,92E-19 | 1,48E-17 | |
| ENSGALG00000023821 | 140,66943 | 2,605691323 | 0,3149662 | 8,2729246 | 1,31E-16 | 7,64E-15 | DHX58 |
| ENSGALG00000006032 | 296,77177 | 2,599165832 | 0,2722516 | 9,5469253 | 1,34E-21 | 1,19E-19 | IL13RA1 |
| ENSGALG00000002960 | 103,93249 | 2,574242623 | 0,3355369 | 7,6720094 | 1,69E-14 | 7,76E-13 | HMGCS2 |
| ENSGALG00000010841 | 110,20084 | 2,5587632 | 0,3321477 | 7,7036919 | 1,32E-14 | 6,10E-13 | |
| ENSGALG00000002829 | 649,13345 | 2,553310959 | 0,2049577 | 12,457744 | 1,27E-35 | 3,20E-33 | SCARF1 |
| ENSGALG00000021198 | 61,503229 | 2,552636185 | 0,3511007 | 7,2703825 | 3,58E-13 | 1,39E-11 | |
| ENSGALG00000015141 | 58,962649 | 2,548558749 | 0,3520748 | 7,2386848 | 4,53E-13 | 1,74E-11 | GDA |
| ENSGALG00000005122 | 338,26914 | 2,528381619 | 0,242755 | 10,415364 | 2,11E-25 | 2,79E-23 | MYO1H |
| ENSGALG00000010145 | 278,24749 | 2,508391386 | 0,3014197 | 8,3219221 | 8,65E-17 | 5,15E-15 | MDGA1 |
| ENSGALG00000014372 | 67,978413 | 2,495880293 | 0,3492839 | 7,1457073 | 8,95E-13 | 3,29E-11 | SLC34A2 |
| ENSGALG00000029144 | 72,328613 | 2,46980754 | 0,3488262 | 7,0803381 | 1,44E-12 | 5,17E-11 | |
| ENSGALG00000021399 | 89,329511 | 2,464358792 | 0,336861 | 7,3156547 | 2,56E-13 | 1,01E-11 | ABCA8 |
| ENSGALG00000001111 | 752,30516 | 2,451755117 | 0,2113059 | 11,602872 | 3,98E-31 | 7,64E-29 | DNAH9 |
| ENSGALG00000012985 | 261,61267 | 2,44205884 | 0,2658681 | 9,185228 | 4,11E-20 | 3,39E-18 | SLC6A13 |
| ENSGALG00000008105 | 47,87761 | 2,404806479 | 0,3539579 | 6,7940461 | 1,09E-11 | 3,44E-10 | CYP2J6L2 |
| ENSGALG00000012812 | 176,06145 | 2,39422427 | 0,2959701 | 8,089412 | 6,00E-16 | 3,29E-14 | SVOPL |

| | | | | | | | |
|---|---|---|---|---|---|---|---|
| ENSGALG00000002638 | 117,91406 | 2,388124012 | 0,3283139 | 7,2739052 | 3,49E-13 | 1,35E-11 | SRRL |
| ENSGALG00000002318 | 63,409663 | 2,376044719 | 0,3494777 | 6,7988449 | 1,05E-11 | 3,33E-10 | LVRN |
| ENSGALG00000002015 | 214,32321 | 2,374226325 | 0,2864093 | 8,2896265 | 1,14E-16 | 6,72E-15 | SNCG |
| ENSGALG00000026256 | 199,83767 | 2,357924834 | 0,3332057 | 7,0764832 | 1,48E-12 | 5,30E-11 | |
| ENSGALG00000007395 | 402,67751 | 2,356587415 | 0,2261671 | 10,419672 | 2,02E-25 | 2,69E-23 | ABCC2 |
| ENSGALG00000002046 | 463,72169 | 2,341658671 | 0,2422889 | 9,6647372 | 4,26E-22 | 3,97E-20 | SYT15 |
| ENSGALG00000007340 | 53,630639 | 2,338672246 | 0,3529687 | 6,6257213 | 3,46E-11 | 1,01E-09 | TMPRSS5 |
| ENSGALG00000023934 | 52,57595 | 2,337998666 | 0,3538351 | 6,6075938 | 3,91E-11 | 1,12E-09 | |
| ENSGALG00000028341 | 324,38787 | 2,327426751 | 0,2360199 | 9,8611477 | 6,13E-23 | 6,23E-21 | MADCAM1 |
| ENSGALG00000002260 | 279,41384 | 2,326075861 | 0,254913 | 9,1249794 | 7,17E-20 | 5,71E-18 | CISH |
| ENSGALG00000006402 | 199,28187 | 2,315345967 | 0,2776938 | 8,3377677 | 7,57E-17 | 4,56E-15 | NRROS |
| ENSGALG00000008709 | 98,437291 | 2,291854243 | 0,3268395 | 7,0121701 | 2,35E-12 | 8,17E-11 | MCOLN3 |
| ENSGALG00000013006 | 357,98483 | 2,275189997 | 0,2579484 | 8,8203283 | 1,14E-18 | 8,05E-17 | ROPN1L |
| ENSGALG00000006992 | 182,22358 | 2,270300547 | 0,2984455 | 7,6070847 | 2,80E-14 | 1,27E-12 | MMP9 |
| ENSGALG00000011113 | 527,00037 | 2,26759493 | 0,2713629 | 8,356319 | 6,47E-17 | 3,93E-15 | SGIP1 |
| ENSGALG00000023577 | 321,7197 | 2,257832359 | 0,2669922 | 8,4565486 | 2,75E-17 | 1,74E-15 | TBX2 |
| ENSGALG00000016852 | 56,030921 | 2,256531595 | 0,3516922 | 6,416211 | 1,40E-10 | 3,74E-09 | TNFSF13B |
| ENSGALG00000027617 | 219,45804 | 2,255993416 | 0,2851339 | 7,9120503 | 2,53E-15 | 1,25E-13 | RP11-385J1.3 |
| ENSGALG00000003677 | 113,60782 | 2,250263529 | 0,3449597 | 6,5232654 | 6,88E-11 | 1,92E-09 | NTNG2 |
| ENSGALG00000023742 | 249,60413 | 2,24719921 | 0,2820489 | 7,9674098 | 1,62E-15 | 8,26E-14 | |
| ENSGALG00000001589 | 111,25767 | 2,246033051 | 0,3441573 | 6,526182 | 6,75E-11 | 1,89E-09 | INPP5D |
| ENSGALG00000016369 | 952,12228 | 2,242110682 | 0,1770401 | 12,664423 | 9,31E-37 | 2,54E-34 | FBXO32 |
| ENSGALG00000027703 | 404,86881 | 2,238234147 | 0,2535546 | 8,8274262 | 1,07E-18 | 7,64E-17 | |
| ENSGALG00000003389 | 248,60083 | 2,230534118 | 0,2705236 | 8,2452465 | 1,65E-16 | 9,51E-15 | IFI30 |
| ENSGALG00000015244 | 123,74828 | 2,228224412 | 0,3204639 | 6,9531211 | 3,57E-12 | 1,20E-10 | TBX19 |
| ENSGALG00000001857 | 255,91037 | 2,203449137 | 0,2595043 | 8,4909939 | 2,05E-17 | 1,31E-15 | FAM132A |
| ENSGALG00000019861 | 47,604267 | 2,196772492 | 0,3539452 | 6,2065321 | 5,42E-10 | 1,33E-08 | |
| ENSGALG00000008514 | 959,21197 | 2,19329695 | 0,1769465 | 12,395254 | 2,77E-35 | 6,86E-33 | DLL4 |
| ENSGALG00000021714 | 88,405928 | 2,185372855 | 0,3343395 | 6,5363894 | 6,30E-11 | 1,78E-09 | PADI1 |
| ENSGALG00000009499 | 136,98531 | 2,185164296 | 0,3152046 | 6,932526 | 4,13E-12 | 1,37E-10 | NPY5R |
| ENSGALG00000026467 | 67,174686 | 2,184133106 | 0,3474873 | 6,2855052 | 3,27E-10 | 8,22E-09 | MESP1 |
| ENSGALG00000027532 | 181,56516 | 2,181261094 | 0,3353331 | 6,5047583 | 7,78E-11 | 2,16E-09 | COLEC11 |
| ENSGALG00000002817 | 43,721989 | 2,174112777 | 0,3531294 | 6,1567025 | 7,43E-10 | 1,79E-08 | CPLX4 |
| ENSGALG00000028339 | 572,22079 | 2,143746023 | 0,2535634 | 8,4544787 | 2,80E-17 | 1,76E-15 | AMIGO3 |
| ENSGALG00000003548 | 717,97931 | 2,131798574 | 0,1846233 | 11,546747 | 7,67E-31 | 1,43E-28 | BAIAP2L1 |
| ENSGALG00000029168 | 574,12283 | 2,125452005 | 0,2633801 | 8,069905 | 7,04E-16 | 3,85E-14 | TMEM27 |
| ENSGALG00000005062 | 695,37715 | 2,112402161 | 0,1891794 | 11,166133 | 5,97E-29 | 9,35E-27 | FOXN4 |
| ENSGALG00000001879 | 730,68274 | 2,106018407 | 0,1838916 | 11,4525 | 2,28E-30 | 4,04E-28 | GRID1 |
| ENSGALG00000028551 | 2508,945 | 2,105237568 | 0,1772334 | 11,878334 | 1,53E-32 | 3,32E-30 | |
| ENSGALG00000001696 | 267,69848 | 2,102610472 | 0,2564004 | 8,2004945 | 2,39E-16 | 1,36E-14 | SAG |
| ENSGALG00000008275 | 48,571485 | 2,098791425 | 0,3539549 | 5,9295439 | 3,04E-09 | 6,67E-08 | MESP2 |
| ENSGALG00000021567 | 284,86115 | 2,081191963 | 0,246615 | 8,4390326 | 3,20E-17 | 1,99E-15 | EPHA8 |
| ENSGALG00000001874 | 67,244987 | 2,080511409 | 0,3538919 | 5,8789453 | 4,13E-09 | 8,88E-08 | |
| ENSGALG00000028654 | 92,92864 | 2,074935686 | 0,3457662 | 6,0009788 | 1,96E-09 | 4,45E-08 | |
| ENSGALG00000006260 | 56,956461 | 2,062296513 | 0,3527945 | 5,8456027 | 5,05E-09 | 1,06E-07 | KLHL30 |
| ENSGALG00000007570 | 256,24389 | 2,055518081 | 0,2571315 | 7,9940356 | 1,31E-15 | 6,73E-14 | KLF8 |
| ENSGALG00000019527 | 38,897028 | 2,052781357 | 0,3521947 | 5,8285416 | 5,59E-09 | 1,16E-07 | SLC6A19 |
| ENSGALG00000000656 | 95,76214 | 2,036711968 | 0,3414768 | 5,964423 | 2,46E-09 | 5,51E-08 | TMEM81 |
| ENSGALG00000007104 | 125,61776 | 2,032418742 | 0,3200563 | 6,3501909 | 2,15E-10 | 5,59E-09 | LRRC15 |
| ENSGALG00000026541 | 175,99093 | 2,028653787 | 0,2976967 | 6,8144995 | 9,46E-12 | 3,00E-10 | |
| ENSGALG00000014203 | 138,12328 | 2,025939839 | 0,3005882 | 6,7399175 | 1,58E-11 | 4,87E-10 | PARVG |
| ENSGALG00000020195 | 143,5922 | 2,021370929 | 0,3033718 | 6,6630159 | 2,68E-11 | 7,92E-10 | DMP1 |
| ENSGALG00000012204 | 102,97928 | 2,019423041 | 0,32821 | 6,1528374 | 7,61E-10 | 1,83E-08 | FAM20CL |
| ENSGALG00000005797 | 87,794264 | 2,014484064 | 0,3341899 | 6,0279615 | 1,66E-09 | 3,83E-08 | COL20A1 |
| ENSGALG00000000437 | 37,093294 | 2,013731373 | 0,3518291 | 5,7236071 | 1,04E-08 | 2,04E-07 | |
| ENSGALG00000002931 | 53,692699 | 2,013598707 | 0,3534025 | 5,6977488 | 1,21E-08 | 2,37E-07 | REG4 |
| ENSGALG00000006273 | 107,39879 | 2,011386223 | 0,3229782 | 6,2276225 | 4,74E-10 | 1,16E-08 | MYLK2 |
| ENSGALG00000000722 | 46,564331 | 2,006522785 | 0,3537549 | 5,6720703 | 1,41E-08 | 2,72E-07 | |
| ENSGALG00000028147 | 229,82466 | 2,001096858 | 0,2664367 | 7,5105891 | 5,89E-14 | 2,49E-12 | CTSK |
| ENSGALG00000004248 | 56,901253 | 1,998749091 | 0,352065 | 5,6772168 | 1,37E-08 | 2,65E-07 | TACR2 |
| ENSGALG00000003075 | 93,078833 | 1,997443704 | 0,3325728 | 6,0060344 | 1,90E-09 | 4,34E-08 | SLC34A1 |
| ENSGALG00000001571 | 57,11691 | 1,994083312 | 0,35177 | 5,6687129 | 1,44E-08 | 2,77E-07 | MYO1F |
| ENSGALG00000010458 | 184,3782 | 1,990178775 | 0,2802188 | 7,1022313 | 1,23E-12 | 4,45E-11 | NOXRED1 |
| ENSGALG00000011889 | 230,92302 | 1,988400853 | 0,2676563 | 7,4289338 | 1,09E-13 | 4,48E-12 | RP11-47I22.4 |

| | | | | | | | |
|---|---|---|---|---|---|---|---|
| ENSGALG00000010924 | 209,96874 | 1,986898996 | 0,2874232 | 6,9127999 | 4,75E-12 | 1,56E-10 | NFIA |
| ENSGALG00000015846 | 313,73062 | 1,982793598 | 0,2465855 | 8,0409991 | 8,91E-16 | 4,78E-14 | SNAP91 |
| ENSGALG00000006835 | 112,62877 | 1,974805269 | 0,3171624 | 6,2264797 | 4,77E-10 | 1,17E-08 | TNNC2 |
| ENSGALG00000013253 | 157,86205 | 1,972349113 | 0,3198156 | 6,1671455 | 6,95E-10 | 1,68E-08 | INSRR |
| ENSGALG00000009302 | 367,16268 | 1,971912015 | 0,2879884 | 6,8471929 | 7,53E-12 | 2,42E-10 | EMX2 |
| ENSGALG00000004414 | 101,95784 | 1,959564216 | 0,3212322 | 6,1001479 | 1,06E-09 | 2,50E-08 | LUZP2 |
| ENSGALG00000009489 | 127,74049 | 1,956908773 | 0,320476 | 6,1062578 | 1,02E-09 | 2,42E-08 | FSTL5 |
| ENSGALG00000005822 | 147,05646 | 1,951526015 | 0,3027027 | 6,447005 | 1,14E-10 | 3,07E-09 | KCNQ2 |
| ENSGALG00000027147 | 228,05825 | 1,950896881 | 0,2801413 | 6,9639743 | 3,31E-12 | 1,13E-10 | SUB1L1 |
| ENSGALG00000021869 | 645,94199 | 1,950468731 | 0,2435545 | 8,0083473 | 1,16E-15 | 6,04E-14 | PODN |
| ENSGALG00000028975 | 532,93063 | 1,945491073 | 0,2058434 | 9,4513148 | 3,35E-21 | 2,90E-19 | FRMPD3 |
| ENSGALG00000007666 | 61,970313 | 1,945055405 | 0,3514319 | 5,5346576 | 3,12E-08 | 5,56E-07 | KCNIP2 |
| ENSGALG00000021656 | 37,511821 | 1,943986445 | 0,3507973 | 5,5416231 | 3,00E-08 | 5,37E-07 | PTPN22 |
| ENSGALG00000008188 | 164,94281 | 1,939910796 | 0,2907086 | 6,6730415 | 2,51E-11 | 7,46E-10 | TRIM36 |
| ENSGALG00000010362 | 56,130567 | 1,930559856 | 0,3538155 | 5,456402 | 4,86E-08 | 8,33E-07 | SUCNR1 |
| ENSGALG00000000560 | 58,758167 | 1,925570368 | 0,353335 | 5,4497022 | 5,05E-08 | 8,61E-07 | PI16 |
| ENSGALG00000015021 | 44,681844 | 1,923323022 | 0,3539578 | 5,4337643 | 5,52E-08 | 9,36E-07 | TMEM174 |
| ENSGALG00000015551 | 1931,1282 | 1,921428632 | 0,1720637 | 11,16696 | 5,92E-29 | 9,35E-27 | GRIN3A |
| ENSGALG00000010867 | 80,266993 | 1,919026004 | 0,3407804 | 5,6312691 | 1,79E-08 | 3,37E-07 | TACSTD2 |
| ENSGALG00000001004 | 271,27145 | 1,914258624 | 0,2596069 | 7,3736827 | 1,66E-13 | 6,69E-12 | MPO |
| ENSGALG00000011347 | 48,911798 | 1,909599526 | 0,3539061 | 5,3957801 | 6,82E-08 | 1,14E-06 | IHH |
| ENSGALG00000006413 | 117,16579 | 1,90767902 | 0,3112895 | 6,1283111 | 8,88E-10 | 2,11E-08 | CEMIP |
| ENSGALG00000003836 | 138,24842 | 1,907173269 | 0,3450544 | 5,5271675 | 3,25E-08 | 5,78E-07 | RBM44 |
| ENSGALG00000007345 | 111,97256 | 1,905774604 | 0,3245679 | 5,8717287 | 4,31E-09 | 9,20E-08 | TMEM114 |
| ENSGALG00000008701 | 97,136553 | 1,905747917 | 0,3400514 | 5,6042938 | 2,09E-08 | 3,87E-07 | XDH |
| ENSGALG00000010853 | 484,76878 | 1,904256394 | 0,2172677 | 8,7645617 | 1,88E-18 | 1,29E-16 | C8B |
| ENSGALG00000000221 | 51,676802 | 1,900065815 | 0,35396 | 5,3680237 | 7,96E-08 | 1,31E-06 | PROM2 |
| ENSGALG00000001617 | 1078,3293 | 1,898502237 | 0,2123138 | 8,9419639 | 3,82E-19 | 2,86E-17 | CAMKK1 |
| ENSGALG00000019290 | 78,123837 | 1,892925943 | 0,3496984 | 5,4130245 | 6,20E-08 | 1,04E-06 | SLC6A12 |
| ENSGALG00000009265 | 88,174325 | 1,891791729 | 0,3459146 | 5,4689565 | 4,53E-08 | 7,80E-07 | KCNK18 |
| ENSGALG00000003923 | 2335,3387 | 1,889033194 | 0,1503705 | 12,562527 | 3,39E-36 | 8,88E-34 | COL6A3 |
| ENSGALG00000008995 | 104,54637 | 1,882652598 | 0,3282959 | 5,7346211 | 9,77E-09 | 1,93E-07 | MAMDC4 |
| ENSGALG00000000544 | 347,72624 | 1,877518925 | 0,2328757 | 8,0623238 | 7,49E-16 | 4,08E-14 | |
| ENSGALG00000007706 | 86,703491 | 1,877381913 | 0,3347682 | 5,6080061 | 2,05E-08 | 3,82E-07 | FGF8 |
| ENSGALG00000015347 | 73,173206 | 1,876981559 | 0,3445279 | 5,4479816 | 5,09E-08 | 8,66E-07 | ZPLD1 |
| ENSGALG00000009078 | 124,06611 | 1,868595491 | 0,3338294 | 5,5974556 | 2,18E-08 | 4,01E-07 | GALNT14 |
| ENSGALG00000000899 | 303,59428 | 1,867380532 | 0,252928 | 7,3830526 | 1,55E-13 | 6,29E-12 | PSD2 |
| ENSGALG00000023459 | 62,523363 | 1,863475253 | 0,3506341 | 5,314586 | 1,07E-07 | 1,71E-06 | MC1R |
| ENSGALG00000013953 | 153,51081 | 1,853929495 | 0,316158 | 5,8639338 | 4,52E-09 | 9,60E-08 | TACR1 |
| ENSGALG00000020316 | 66,987938 | 1,852228829 | 0,3437054 | 5,3890016 | 7,09E-08 | 1,17E-06 | IL13RA2 |
| ENSGALG00000014770 | 4689,6836 | 1,850573228 | 0,1917177 | 9,6525964 | 4,79E-22 | 4,41E-20 | EPB41L3 |
| ENSGALG00000015406 | 32,824277 | 1,850052547 | 0,3495416 | 5,2927962 | 1,20E-07 | 1,90E-06 | MOS |
| ENSGALG00000007047 | 78,987356 | 1,844593336 | 0,3380656 | 5,4563178 | 4,86E-08 | 8,33E-07 | GAL |
| ENSGALG00000011421 | 155,78727 | 1,840566258 | 0,2941943 | 6,2562953 | 3,94E-10 | 9,81E-09 | HAL |
| ENSGALG00000003751 | 52,948026 | 1,835104614 | 0,3538037 | 5,186787 | 2,14E-07 | 3,22E-06 | ASAH2 |
| ENSGALG00000021693 | 69,312923 | 1,830329106 | 0,3430516 | 5,3354333 | 9,53E-08 | 1,54E-06 | |
| ENSGALG00000005634 | 92,232146 | 1,826798031 | 0,3274114 | 5,5795183 | 2,41E-08 | 4,40E-07 | CRYBA4 |
| ENSGALG00000005644 | 54,639892 | 1,819377424 | 0,3504577 | 5,1914323 | 2,09E-07 | 3,15E-06 | MYO1G |
| ENSGALG00000029006 | 89,819678 | 1,803532169 | 0,3296621 | 5,4708508 | 4,48E-08 | 7,73E-07 | SYPL1 |
| ENSGALG00000010126 | 116,19587 | 1,799221778 | 0,3298899 | 5,4540074 | 4,92E-08 | 8,42E-07 | BEST4 |
| ENSGALG00000019361 | 44,228613 | 1,792121861 | 0,3538597 | 5,0644988 | 4,09E-07 | 5,78E-06 | FAM19A2 |
| ENSGALG00000020982 | 53,872261 | 1,789263062 | 0,3513204 | 5,092966 | 3,53E-07 | 5,05E-06 | BPIFB3 |
| ENSGALG00000004841 | 236,70913 | 1,788099107 | 0,2763588 | 6,4702084 | 9,79E-11 | 2,66E-09 | APLNR |
| ENSGALG00000008845 | 211,0907 | 1,784489653 | 0,2697983 | 6,6141623 | 3,74E-11 | 1,08E-09 | HAO1 |
| ENSGALG00000023354 | 87,321322 | 1,782015396 | 0,3447223 | 5,1694224 | 2,35E-07 | 3,49E-06 | FSCN2 |
| ENSGALG00000010364 | 75,712359 | 1,777216343 | 0,3496501 | 5,0828426 | 3,72E-07 | 5,29E-06 | AADACL2 |
| ENSGALG00000027090 | 122,23539 | 1,772101833 | 0,3238631 | 5,4717629 | 4,46E-08 | 7,70E-07 | |
| ENSGALG00000005043 | 168,0345 | 1,771501032 | 0,2886982 | 6,13617 | 8,45E-10 | 2,02E-08 | ACACB |
| ENSGALG00000005747 | 39,000743 | 1,767653479 | 0,3536042 | 4,998961 | 5,76E-07 | 7,89E-06 | |
| ENSGALG00000014252 | 107,13553 | 1,765576737 | 0,346339 | 5,0978281 | 3,44E-07 | 4,94E-06 | |
| ENSGALG00000006007 | 80,52031 | 1,763247242 | 0,3431655 | 5,1381839 | 2,77E-07 | 4,06E-06 | SAMD10 |
| ENSGALG00000000630 | 44,986023 | 1,762139369 | 0,352772 | 4,9951231 | 5,88E-07 | 8,02E-06 | HAS3 |
| ENSGALG00000013971 | 623,6268 | 1,754983652 | 0,2217786 | 7,9132227 | 2,51E-15 | 1,24E-13 | SGK1 |
| ENSGALG00000025804 | 57,829823 | 1,753854275 | 0,3486801 | 5,0299803 | 4,91E-07 | 6,84E-06 | CLCN2 |

| | | | | | | | |
|---|---|---|---|---|---|---|---|
| ENSGALG00000016233 | 282,39076 | 1,751740847 | 0,2591609 | 6,7592786 | 1,39E-11 | 4,28E-10 | TG |
| ENSGALG00000000299 | 146,62252 | 1,750555222 | 0,3173752 | 5,5157274 | 3,47E-08 | 6,13E-07 | PKP1 |
| ENSGALG00000015154 | 107,54458 | 1,749162012 | 0,3192627 | 5,4787551 | 4,28E-08 | 7,43E-07 | TRPM6 |
| ENSGALG00000028801 | 326,04496 | 1,749152851 | 0,2317152 | 7,548717 | 4,40E-14 | 1,90E-12 | YPEL2 |
| ENSGALG00000003103 | 147,01296 | 1,745350116 | 0,2934725 | 5,9472356 | 2,73E-09 | 6,09E-08 | MST1R |
| ENSGALG00000010408 | 48,693762 | 1,74244975 | 0,3525174 | 4,9428754 | 7,70E-07 | 1,02E-05 | OPN4-1 |
| ENSGALG00000016580 | 156,41685 | 1,740943756 | 0,3137577 | 5,5486878 | 2,88E-08 | 5,20E-07 | SCARA5 |
| ENSGALG00000020779 | 287,43405 | 1,737619419 | 0,2447032 | 7,1009277 | 1,24E-12 | 4,48E-11 | |
| ENSGALG00000015544 | 64,576519 | 1,719200832 | 0,3511965 | 4,895268 | 9,82E-07 | 1,28E-05 | ALDOB |
| ENSGALG00000015305 | 121,8353 | 1,715527315 | 0,3092461 | 5,547451 | 2,90E-08 | 5,21E-07 | NR2E1 |
| ENSGALG00000008926 | 1594,7541 | 1,713961304 | 0,1563378 | 10,963191 | 5,74E-28 | 8,69E-26 | CADPS2 |
| ENSGALG00000002957 | 403,40008 | 1,713056332 | 0,2361807 | 7,2531612 | 4,07E-13 | 1,57E-11 | SLC12A3 |
| ENSGALG00000017212 | 76,54735 | 1,710911672 | 0,3424068 | 4,9967228 | 5,83E-07 | 7,97E-06 | GPR83 |
| ENSGALG00000004827 | 60,764569 | 1,70800582 | 0,3466264 | 4,9275126 | 8,33E-07 | 1,10E-05 | PAK6 |
| ENSGALG00000003242 | 570,22011 | 1,707949225 | 0,2101315 | 8,1280036 | 4,36E-16 | 2,42E-14 | STAR |
| ENSGALG00000001912 | 146,49438 | 1,699528733 | 0,3012842 | 5,6409491 | 1,69E-08 | 3,21E-07 | EPHA10 |
| ENSGALG00000003003 | 42,669769 | 1,694427776 | 0,3520315 | 4,8132841 | 1,48E-06 | 1,86E-05 | CCL20 |
| ENSGALG00000016261 | 158,70666 | 1,693178751 | 0,3145024 | 5,3836747 | 7,30E-08 | 1,20E-06 | CYBB |
| ENSGALG00000011007 | 1150,329 | 1,688733669 | 0,1907765 | 8,8518972 | 8,60E-19 | 6,20E-17 | OSBPL3 |
| ENSGALG00000015117 | 149,51437 | 1,688648354 | 0,2954854 | 5,7148283 | 1,10E-08 | 2,15E-07 | DRD3 |
| ENSGALG00000012429 | 6013,8279 | 1,680823476 | 0,1520104 | 11,05729 | 2,02E-28 | 3,09E-26 | BMP4 |
| ENSGALG00000000458 | 427,66514 | 1,679795969 | 0,2174489 | 7,7250163 | 1,12E-14 | 5,20E-13 | TMPRSS9 |
| ENSGALG00000011622 | 28,65156 | 1,67904306 | 0,3460309 | 4,8522925 | 1,22E-06 | 1,56E-05 | MAS1 |
| ENSGALG00000016572 | 77,200694 | 1,678816771 | 0,3434432 | 4,8881939 | 1,02E-06 | 1,32E-05 | C3H8ORF80 |
| ENSGALG00000001084 | 353,56487 | 1,677708871 | 0,2418689 | 6,9364397 | 4,02E-12 | 1,34E-10 | ATP8B3 |
| ENSGALG00000026473 | 1522,1511 | 1,674638021 | 0,2112768 | 7,9262761 | 2,26E-15 | 1,13E-13 | CHST11 |
| ENSGALG00000008178 | 111,71167 | 1,67186128 | 0,3225826 | 5,1827379 | 2,19E-07 | 3,28E-06 | CYP2J2L5 |
| ENSGALG00000005140 | 240,21489 | 1,671709253 | 0,257977 | 6,48007 | 9,17E-11 | 2,52E-09 | RRAD |
| ENSGALG00000001234 | 40,862653 | 1,668621796 | 0,3539185 | 4,7147069 | 2,42E-06 | 2,87E-05 | CETP |
| ENSGALG00000006514 | 1126,0055 | 1,665636549 | 0,1751467 | 9,5099504 | 1,91E-21 | 1,69E-19 | FSTL4 |
| ENSGALG00000004936 | 57,472904 | 1,658043102 | 0,3494689 | 4,744466 | 2,09E-06 | 2,51E-05 | GLRA4 |
| ENSGALG00000028174 | 599,32727 | 1,656109672 | 0,2408103 | 6,877239 | 6,10E-12 | 1,98E-10 | SDPR |
| ENSGALG00000011882 | 446,5677 | 1,655648124 | 0,2116439 | 7,8228016 | 5,17E-15 | 2,44E-13 | CDCP1 |
| ENSGALG00000020143 | 148,7819 | 1,654526791 | 0,3218985 | 5,1399029 | 2,75E-07 | 4,02E-06 | GABRG1 |
| ENSGALG00000027314 | 27,97681 | 1,648583002 | 0,3452618 | 4,7748783 | 1,80E-06 | 2,19E-05 | SAXO1 |
| ENSGALG00000019019 | 89,671454 | 1,647447024 | 0,3439031 | 4,7904393 | 1,66E-06 | 2,05E-05 | NTRK1 |
| ENSGALG00000026161 | 123,20926 | 1,647025641 | 0,3125278 | 5,2700132 | 1,36E-07 | 2,13E-06 | |
| ENSGALG00000024298 | 32,030697 | 1,646495761 | 0,3488596 | 4,7196511 | 2,36E-06 | 2,81E-05 | ADAMTSL5 |
| ENSGALG00000004394 | 167,65918 | 1,640660466 | 0,2837407 | 5,7822521 | 7,37E-09 | 1,49E-07 | PCBP3 |
| ENSGALG00000016262 | 396,40337 | 1,640510999 | 0,2287869 | 7,1704758 | 7,47E-13 | 2,77E-11 | XK |
| ENSGALG00000015905 | 84,004407 | 1,632452898 | 0,3358609 | 4,860503 | 1,17E-06 | 1,51E-05 | SLC7A7 |
| ENSGALG00000005293 | 115,92754 | 1,631397185 | 0,3272119 | 4,9857512 | 6,17E-07 | 8,36E-06 | GNG13 |
| ENSGALG00000025946 | 137,58478 | 1,630278266 | 0,3118772 | 5,2273088 | 1,72E-07 | 2,64E-06 | MOB3C |
| ENSGALG00000011806 | 33,35332 | 1,625032877 | 0,3475057 | 4,6762767 | 2,92E-06 | 3,39E-05 | MUC13 |
| ENSGALG00000010721 | 516,48932 | 1,61713725 | 0,1982525 | 8,1569561 | 3,44E-16 | 1,92E-14 | GLIS1 |
| ENSGALG00000008133 | 122,89128 | 1,61551597 | 0,3252043 | 4,9676957 | 6,78E-07 | 9,10E-06 | AS3MT |
| ENSGALG00000000820 | 127,58617 | 1,614544959 | 0,3185699 | 5,068102 | 4,02E-07 | 5,68E-06 | HTR1D |
| ENSGALG00000002832 | 501,68162 | 1,610990532 | 0,2127694 | 7,5715338 | 3,69E-14 | 1,62E-12 | IFI35 |
| ENSGALG00000006606 | 3620,8517 | 1,610976489 | 0,1224776 | 13,153237 | 1,63E-39 | 5,29E-37 | KIAA1324L |
| ENSGALG00000011200 | 907,82877 | 1,609360966 | 0,246667 | 6,5244282 | 6,83E-11 | 1,91E-09 | THBS2 |
| ENSGALG00000007980 | 258,34488 | 1,607363167 | 0,2746243 | 5,8529528 | 4,83E-09 | 1,02E-07 | KCNG1 |
| ENSGALG00000001062 | 680,57905 | 1,606326071 | 0,191836 | 8,3734359 | 5,60E-17 | 3,43E-15 | GGT7 |
| ENSGALG00000003106 | 661,98127 | 1,604428011 | 0,2222828 | 7,217957 | 5,28E-13 | 2,01E-11 | TOX2 |
| ENSGALG00000007815 | 333,86581 | 1,603027021 | 0,2797129 | 5,7309725 | 9,99E-09 | 1,96E-07 | ANKK1 |
| ENSGALG00000015366 | 80,07277 | 1,602600172 | 0,3432645 | 4,6687031 | 3,03E-06 | 3,52E-05 | |
| ENSGALG00000005725 | 91,108572 | 1,601523904 | 0,352027 | 4,5494347 | 5,38E-06 | 5,90E-05 | CSF1R |
| ENSGALG00000022644 | 26,874228 | 1,595235245 | 0,3437941 | 4,6400892 | 3,48E-06 | 3,98E-05 | CCDC36 |
| ENSGALG00000007759 | 856,86635 | 1,594985498 | 0,1989057 | 8,0188035 | 1,07E-15 | 5,57E-14 | TMEFF2 |
| ENSGALG00000008830 | 781,04938 | 1,591806155 | 0,2114024 | 7,5297432 | 5,08E-14 | 2,17E-12 | BMP2 |
| ENSGALG00000006412 | 53,058714 | 1,591456503 | 0,3527925 | 4,5110267 | 6,45E-06 | 6,90E-05 | VGLL1 |
| ENSGALG00000029174 | 5348,1501 | 1,589321167 | 0,1290901 | 12,311717 | 7,83E-35 | 1,87E-32 | |
| ENSGALG00000006245 | 402,97003 | 1,589094556 | 0,227751 | 6,9773328 | 3,01E-12 | 1,04E-10 | KLF15 |
| ENSGALG00000026733 | 77,5006 | 1,588089103 | 0,3406644 | 4,6617403 | 3,14E-06 | 3,62E-05 | FFAR4 |
| ENSGALG00000016138 | 93,251278 | 1,581494384 | 0,3296626 | 4,7973118 | 1,61E-06 | 2,00E-05 | DSCAM |

| | | | | | | | |
|---|---|---|---|---|---|---|---|
| ENSGALG00000016498 | 94,791705 | 1,580375213 | 0,3294934 | 4,7963786 | 1,62E-06 | 2,01E-05 | MFSD2B |
| ENSGALG00000011343 | 80,565753 | 1,579403764 | 0,3348001 | 4,7174527 | 2,39E-06 | 2,83E-05 | SLC23A3 |
| ENSGALG00000000733 | 1063,131 | 1,578595816 | 0,1987694 | 7,941845 | 1,99E-15 | 1,00E-13 | MYO7A |
| ENSGALG00000028175 | 72,934835 | 1,575180638 | 0,35396 | 4,4501653 | 8,58E-06 | 9,01E-05 | GJA9 |
| ENSGALG00000015293 | 333,00466 | 1,574970296 | 0,2444462 | 6,4430147 | 1,17E-10 | 3,15E-09 | TMEM45A |
| ENSGALG00000012115 | 228,29439 | 1,572479232 | 0,2734467 | 5,7505884 | 8,89E-09 | 1,77E-07 | |
| ENSGALG00000004692 | 57,025385 | 1,559793633 | 0,3537015 | 4,4099148 | 1,03E-05 | 0,0001064 | FDX1L |
| ENSGALG00000005958 | 558,63577 | 1,558583818 | 0,2329918 | 6,6894354 | 2,24E-11 | 6,70E-10 | TRIM66 |
| ENSGALG00000000826 | 97,953466 | 1,551558986 | 0,3236865 | 4,7933999 | 1,64E-06 | 2,03E-05 | MAPK13 |
| ENSGALG00000007690 | 2070,9973 | 1,54950396 | 0,1755538 | 8,8263752 | 1,08E-18 | 7,67E-17 | TFAP2C |
| ENSGALG00000006964 | 225,19101 | 1,546629935 | 0,2930271 | 5,2781128 | 1,31E-07 | 2,05E-06 | SDR42E2 |
| ENSGALG00000005967 | 735,63667 | 1,545091384 | 0,2216792 | 6,9699438 | 3,17E-12 | 1,09E-10 | LMO1 |
| ENSGALG00000008440 | 429,09527 | 1,543692993 | 0,21647 | 7,1312088 | 9,95E-13 | 3,64E-11 | CHST1 |
| ENSGALG00000001882 | 38,709827 | 1,540579881 | 0,352114 | 4,3752309 | 1,21E-05 | 0,0001224 | TNFRSF18 |
| ENSGALG00000015373 | 27,601769 | 1,538791481 | 0,3445115 | 4,4665892 | 7,95E-06 | 8,37E-05 | PDE6B |
| ENSGALG00000011577 | 399,56293 | 1,531134988 | 0,217286 | 7,0466337 | 1,83E-12 | 6,48E-11 | CNTNAP5 |
| ENSGALG00000025774 | 5724,9696 | 1,527733684 | 0,1270402 | 12,025592 | 2,61E-33 | 5,82E-31 | HOXB2 |
| ENSGALG00000005473 | 107,68664 | 1,527501434 | 0,3207303 | 4,7625734 | 1,91E-06 | 2,31E-05 | |
| ENSGALG00000000479 | 244,69562 | 1,527264642 | 0,2552088 | 5,984374 | 2,17E-09 | 4,90E-08 | PADI3 |
| ENSGALG00000011071 | 944,33324 | 1,527069671 | 0,169472 | 9,0107512 | 2,05E-19 | 1,57E-17 | CLMN |
| ENSGALG00000020459 | 25,515422 | 1,526419374 | 0,3416733 | 4,4674817 | 7,91E-06 | 8,34E-05 | |
| ENSGALG00000014267 | 99,335217 | 1,525149591 | 0,3197285 | 4,7701403 | 1,84E-06 | 2,24E-05 | RBM47 |
| ENSGALG00000007803 | 165,80729 | 1,522432144 | 0,2956669 | 5,1491467 | 2,62E-07 | 3,86E-06 | |
| ENSGALG00000001569 | 88,715373 | 1,518412601 | 0,3422278 | 4,4368471 | 9,13E-06 | 9,51E-05 | RHOC |
| ENSGALG00000020745 | 195,27404 | 1,515524715 | 0,2743093 | 5,5248748 | 3,30E-08 | 5,85E-07 | RHO |
| ENSGALG00000011227 | 778,97981 | 1,515331444 | 0,1818025 | 8,3350423 | 7,75E-17 | 4,65E-15 | CDS1 |
| ENSGALG00000007594 | 126,92297 | 1,512071087 | 0,3020168 | 5,0065791 | 5,54E-07 | 7,63E-06 | ASB12 |
| ENSGALG00000002314 | 85,131488 | 1,511687395 | 0,3333097 | 4,5353841 | 5,75E-06 | 6,23E-05 | CDHR1 |
| ENSGALG00000002353 | 221,89831 | 1,511078773 | 0,2623724 | 5,7592906 | 8,45E-09 | 1,69E-07 | C1orf216 |
| ENSGALG00000024397 | 46,37702 | 1,510326295 | 0,3539532 | 4,2670226 | 1,98E-05 | 0,000186 | |
| ENSGALG00000023866 | 254,33653 | 1,510055693 | 0,2802587 | 5,3880776 | 7,12E-08 | 1,18E-06 | ALPI |
| ENSGALG00000026089 | 41,811297 | 1,509966763 | 0,3504419 | 4,3087509 | 1,64E-05 | 0,0001581 | SLC26A6 |
| ENSGALG00000028386 | 982,7252 | 1,507238343 | 0,1699771 | 8,8673018 | 7,49E-19 | 5,49E-17 | SPEG |
| ENSGALG00000006479 | 34,346853 | 1,50588664 | 0,3481979 | 4,3248009 | 1,53E-05 | 0,0001484 | CCM2L |
| ENSGALG00000005297 | 304,6597 | 1,505251197 | 0,2455725 | 6,1295583 | 8,81E-10 | 2,10E-08 | HHATL |
| ENSGALG00000004448 | 547,11485 | 1,504790559 | 0,2323026 | 6,4777179 | 9,31E-11 | 2,55E-09 | BRINP2 |
| ENSGALG00000026983 | 590,10355 | 1,50209184 | 0,2137064 | 7,0287654 | 2,08E-12 | 7,31E-11 | CHRD |
| ENSGALG00000024264 | 40,681213 | 1,501303133 | 0,349559 | 4,2948485 | 1,75E-05 | 0,0001668 | CTD-2207O23.12 |
| ENSGALG00000028347 | 76,972548 | 1,501152953 | 0,3426288 | 4,381281 | 1,18E-05 | 0,0001192 | SDCBP2 |
| ENSGALG00000016556 | 93,532731 | 1,500831481 | 0,3253057 | 4,6136038 | 3,96E-06 | 4,47E-05 | GVIN1 |
| ENSGALG00000008424 | 31,555997 | 1,500581439 | 0,3495073 | 4,2934189 | 1,76E-05 | 0,0001675 | CALML3 |
| ENSGALG00000016287 | 253,69324 | 1,497802744 | 0,250661 | 5,9754121 | 2,30E-09 | 5,17E-08 | NR0B1 |
| ENSGALG00000028069 | 276,23402 | 1,497716991 | 0,2556436 | 5,858613 | 4,67E-09 | 9,87E-08 | WNT16 |
| ENSGALG00000015410 | 210,47767 | 1,497100835 | 0,2682786 | 5,5803964 | 2,40E-08 | 4,38E-07 | BVES |
| ENSGALG00000014074 | 676,43326 | 1,49566297 | 0,191936 | 7,7925091 | 6,57E-15 | 3,08E-13 | CPEB4 |
| ENSGALG00000003713 | 62,007135 | 1,494159857 | 0,3495106 | 4,2750063 | 1,91E-05 | 0,0001803 | CLCNKB |
| ENSGALG00000007409 | 87,08015 | 1,493981037 | 0,3289914 | 4,5410949 | 5,60E-06 | 6,09E-05 | SCN4B |
| ENSGALG00000028823 | 172,11814 | 1,491796906 | 0,2989787 | 4,9896427 | 6,05E-07 | 8,21E-06 | C9orf172 |
| ENSGALG00000026585 | 53,571101 | 1,487448109 | 0,3507492 | 4,240774 | 2,23E-05 | 0,0002057 | FAM19A3 |
| ENSGALG00000027548 | 35,763933 | 1,48608799 | 0,3490327 | 4,2577329 | 2,07E-05 | 0,0001927 | PROK1 |
| ENSGALG00000028770 | 169,21444 | 1,483894729 | 0,3097948 | 4,7899285 | 1,67E-06 | 2,06E-05 | INHBB |
| ENSGALG00000015237 | 27,293769 | 1,481694783 | 0,3444717 | 4,3013545 | 1,70E-05 | 0,0001626 | PPIL6 |
| ENSGALG00000015084 | 1284,7693 | 1,480304896 | 0,1853415 | 7,9869029 | 1,38E-15 | 7,11E-14 | IGSF11 |
| ENSGALG00000027357 | 83,363535 | 1,478068348 | 0,3342619 | 4,4218869 | 9,78E-06 | 0,000101 | PTGES |
| ENSGALG00000007908 | 523,87573 | 1,477721596 | 0,2119438 | 6,9722343 | 3,12E-12 | 1,08E-10 | EFEMP1 |
| ENSGALG00000008888 | 216,09563 | 1,475198435 | 0,2663438 | 5,5386996 | 3,05E-08 | 5,45E-07 | IL1RAPL2 |
| ENSGALG00000000984 | 31,631294 | 1,47391651 | 0,3504148 | 4,2062051 | 2,60E-05 | 0,0002346 | AGPAT1L |
| ENSGALG00000007862 | 431,76183 | 1,47265082 | 0,2334046 | 6,3094329 | 2,80E-10 | 7,11E-09 | CNTN3 |
| ENSGALG00000016538 | 1671,2767 | 1,472170836 | 0,1444523 | 10,191396 | 2,17E-24 | 2,59E-22 | RAI2 |
| ENSGALG00000002442 | 183,2009 | 1,464263461 | 0,2851722 | 5,1346638 | 2,83E-07 | 4,12E-06 | ANKS4B |
| ENSGALG00000010107 | 1860,6289 | 1,462273804 | 0,1417872 | 10,313158 | 6,14E-25 | 7,75E-23 | FHDC1 |
| ENSGALG00000026677 | 510,78911 | 1,461550873 | 0,2721661 | 5,3700699 | 7,87E-08 | 1,29E-06 | F10 |
| ENSGALG00000000703 | 748,40789 | 1,459568013 | 0,2288692 | 6,3773027 | 1,80E-10 | 4,77E-09 | SLC45A3 |
| ENSGALG00000026069 | 33,975719 | 1,459502097 | 0,3469009 | 4,207259 | 2,58E-05 | 0,0002339 | |

| | | | | | | | |
|---|---|---|---|---|---|---|---|
| ENSGALG00000001319 | 188,5387 | 1,458388464 | 0,2745911 | 5,3111274 | 1,09E-07 | 1,73E-06 | CAMK1G |
| ENSGALG00000014297 | 511,06078 | 1,457150717 | 0,2085114 | 6,9883507 | 2,78E-12 | 9,64E-11 | IRF7 |
| ENSGALG00000004565 | 123,01022 | 1,456247274 | 0,3389164 | 4,2967747 | 1,73E-05 | 0,0001657 | PHYHD1 |
| ENSGALG00000010024 | 105,10355 | 1,453185654 | 0,3195715 | 4,5472947 | 5,43E-06 | 5,94E-05 | |
| ENSGALG00000014941 | 729,89146 | 1,451697765 | 0,1864124 | 7,7875602 | 6,83E-15 | 3,20E-13 | STXBP5L |
| ENSGALG00000018878 | 40,669529 | 1,44967545 | 0,3538085 | 4,0973448 | 4,18E-05 | 0,000353 | |
| ENSGALG00000015128 | 44,346554 | 1,44940013 | 0,3539598 | 4,0948156 | 4,23E-05 | 0,0003559 | AQP4 |
| ENSGALG00000006751 | 459,34864 | 1,449311054 | 0,2141744 | 6,7669659 | 1,32E-11 | 4,09E-10 | THY1 |
| ENSGALG00000017041 | 876,05201 | 1,44806939 | 0,2013684 | 7,1911442 | 6,43E-13 | 2,41E-11 | FREM2 |
| ENSGALG00000007211 | 323,24032 | 1,44654086 | 0,2459257 | 5,8820245 | 4,05E-09 | 8,74E-08 | CDH22 |
| ENSGALG00000001727 | 728,89366 | 1,445847057 | 0,1802658 | 8,0206403 | 1,05E-15 | 5,52E-14 | LZTS1 |
| ENSGALG00000006239 | 29,398702 | 1,443346989 | 0,3491735 | 4,1336095 | 3,57E-05 | 0,0003093 | |
| ENSGALG00000023395 | 151,59782 | 1,441830671 | 0,2999274 | 4,807265 | 1,53E-06 | 1,91E-05 | PLIN1 |
| ENSGALG00000013640 | 91,822966 | 1,440450208 | 0,3243745 | 4,4407017 | 8,97E-06 | 9,38E-05 | MBP |
| ENSGALG00000014962 | 26,464506 | 1,440003001 | 0,3431954 | 4,1958697 | 2,72E-05 | 0,0002436 | FAM26F |
| ENSGALG00000008267 | 52,964686 | 1,439464257 | 0,3533811 | 4,0734046 | 4,63E-05 | 0,0003844 | IL5RA |
| ENSGALG00000011202 | 37,140825 | 1,438867662 | 0,3522352 | 4,0849627 | 4,41E-05 | 0,0003675 | COLQ |
| ENSGALG00000012652 | 778,18023 | 1,438231617 | 0,2169215 | 6,6301938 | 3,35E-11 | 9,79E-10 | KCNG2 |
| ENSGALG00000001710 | 36,697213 | 1,437606263 | 0,3472289 | 4,1402264 | 3,47E-05 | 0,0003013 | |
| ENSGALG00000014695 | 237,14245 | 1,436054042 | 0,2846492 | 5,0449961 | 4,54E-07 | 6,35E-06 | CLCN1 |
| ENSGALG00000024368 | 80,858185 | 1,435967902 | 0,3383077 | 4,2445619 | 2,19E-05 | 0,0002029 | AMH |
| ENSGALG00000003678 | 3218,3918 | 1,429238494 | 0,1247553 | 11,456331 | 2,19E-30 | 3,97E-28 | GLUL |
| ENSGALG00000012313 | 84,383689 | 1,428353126 | 0,34636 | 4,1238977 | 3,73E-05 | 0,0003194 | H1F0 |
| ENSGALG00000000914 | 24,818166 | 1,42659313 | 0,3378824 | 4,2221587 | 2,42E-05 | 0,0002211 | |
| ENSGALG00000006565 | 175,49957 | 1,424795357 | 0,3261375 | 4,3686951 | 1,25E-05 | 0,0001254 | GGT1 |
| ENSGALG00000011888 | 40,65783 | 1,420709802 | 0,353895 | 4,0144953 | 5,96E-05 | 0,0004785 | TGM4 |
| ENSGALG00000010213 | 97,780516 | 1,420151903 | 0,3251042 | 4,3682976 | 1,25E-05 | 0,0001255 | VSX2 |
| ENSGALG00000008901 | 121,60703 | 1,417468594 | 0,3085387 | 4,594136 | 4,35E-06 | 4,87E-05 | PTGFR |
| ENSGALG00000009596 | 108,17405 | 1,416089625 | 0,3214369 | 4,4054986 | 1,06E-05 | 0,0001081 | TMEM117 |
| ENSGALG00000014412 | 49,062969 | 1,415817045 | 0,3518535 | 4,0238822 | 5,72E-05 | 0,0004628 | CSTA |
| ENSGALG00000000589 | 81,385408 | 1,415474788 | 0,3386211 | 4,1801144 | 2,91E-05 | 0,0002588 | BTN1A1 |
| ENSGALG00000023834 | 58,026469 | 1,413727582 | 0,350805 | 4,0299523 | 5,58E-05 | 0,0004527 | |
| ENSGALG00000018998 | 220,94684 | 1,412120959 | 0,2810636 | 5,0242041 | 5,06E-07 | 7,03E-06 | UNC5A |
| ENSGALG00000013081 | 81,619637 | 1,411618895 | 0,3443628 | 4,0992204 | 4,15E-05 | 0,0003508 | VWC2 |
| ENSGALG00000027520 | 176,34242 | 1,409426755 | 0,294554 | 4,7849521 | 1,71E-06 | 2,10E-05 | |
| ENSGALG00000020423 | 91,673069 | 1,409349068 | 0,344453 | 4,0915566 | 4,28E-05 | 0,0003598 | NKX2-1 |
| ENSGALG00000026957 | 482,71907 | 1,407844107 | 0,20929 | 6,726761 | 1,73E-11 | 5,28E-10 | SEMA4G |
| ENSGALG00000010486 | 221,46033 | 1,406315246 | 0,2602657 | 5,4033833 | 6,54E-08 | 1,10E-06 | JAKMIP3 |
| ENSGALG00000028936 | 318,09117 | 1,405167114 | 0,2341016 | 6,0023824 | 1,94E-09 | 4,43E-08 | |
| ENSGALG00000002637 | 48,952453 | 1,404353552 | 0,3533799 | 3,9740621 | 7,07E-05 | 0,0005557 | RBP7 |
| ENSGALG00000026948 | 265,89811 | 1,40406892 | 0,2839426 | 4,9449037 | 7,62E-07 | 1,01E-05 | ADD2 |
| ENSGALG00000016782 | 106,88039 | 1,400781577 | 0,3284889 | 4,2643192 | 2,01E-05 | 0,000188 | IL1R2 |
| ENSGALG00000015918 | 313,69588 | 1,399804901 | 0,2380455 | 5,8804082 | 4,09E-09 | 8,82E-08 | NECAB1 |
| ENSGALG00000011594 | 42,210813 | 1,39976735 | 0,3539056 | 3,9552003 | 7,65E-05 | 0,0005929 | NR1H4 |
| ENSGALG00000009390 | 538,73431 | 1,398258988 | 0,2055508 | 6,8024979 | 1,03E-11 | 3,26E-10 | MET |
| ENSGALG00000004252 | 60,119703 | 1,3977567 | 0,3472961 | 4,0246834 | 5,71E-05 | 0,0004615 | |
| ENSGALG00000019864 | 58,892366 | 1,392548079 | 0,3487729 | 3,9927067 | 6,53E-05 | 0,0005177 | |
| ENSGALG00000029083 | 179,60714 | 1,390642479 | 0,2814687 | 4,9406649 | 7,79E-07 | 1,03E-05 | NXPH2 |
| ENSGALG00000012875 | 68,467203 | 1,389711553 | 0,3440597 | 4,0391582 | 5,36E-05 | 0,0004376 | WEE2 |
| ENSGALG00000023689 | 74,977508 | 1,388126911 | 0,3405714 | 4,0758757 | 4,58E-05 | 0,0003808 | ASS1 |
| ENSGALG00000026536 | 52,51215 | 1,387507758 | 0,3536902 | 3,9229466 | 8,75E-05 | 0,0006706 | IP6K3 |
| ENSGALG00000014857 | 696,58457 | 1,387229623 | 0,1828391 | 7,587161 | 3,27E-14 | 1,45E-12 | SEPP1 |
| ENSGALG00000002319 | 151,6038 | 1,386034084 | 0,2938554 | 4,7167213 | 2,40E-06 | 2,84E-05 | TEKT2 |
| ENSGALG00000015776 | 28,48922 | 1,382386109 | 0,3437254 | 4,0217745 | 5,78E-05 | 0,0004664 | GABRR2 |
| ENSGALG00000007636 | 33,478978 | 1,381489214 | 0,3508105 | 3,9379927 | 8,22E-05 | 0,0006327 | PCK1 |
| ENSGALG00000007014 | 150,69691 | 1,378668928 | 0,2898641 | 4,7562591 | 1,97E-06 | 2,38E-05 | PYROXD2 |
| ENSGALG00000027973 | 59,731595 | 1,378158223 | 0,3525539 | 3,909071 | 9,27E-05 | 0,0007038 | CATH1 |
| ENSGALG00000017099 | 99,501102 | 1,377440865 | 0,3228202 | 4,266898 | 1,98E-05 | 0,000186 | RASL11A |
| ENSGALG00000005493 | 91,862545 | 1,375327372 | 0,3276333 | 4,1977643 | 2,70E-05 | 0,0002424 | ITGB1BP2 |
| ENSGALG00000006522 | 45,353339 | 1,373172 | 0,3533367 | 3,8862984 | 0,0001018 | 0,0007623 | HCK |
| ENSGALG00000012956 | 1426,5049 | 1,372804533 | 0,1863192 | 7,3680244 | 1,73E-13 | 6,93E-12 | TMTC1 |
| ENSGALG00000002105 | 44,541184 | 1,371897103 | 0,3536727 | 3,8790025 | 0,0001049 | 0,0007838 | FOXI1 |
| ENSGALG00000016584 | 157,58752 | 1,371284929 | 0,2995639 | 4,5776048 | 4,70E-06 | 5,25E-05 | EGFL6 |
| ENSGALG00000000730 | 181,14132 | 1,369179792 | 0,2801686 | 4,8869848 | 1,02E-06 | 1,33E-05 | CACNA1S |

| | | | | | | | |
|---|---|---|---|---|---|---|---|
| ENSGALG00000009151 | 104,26311 | 1,368816021 | 0,3214356 | 4,2584453 | 2,06E-05 | 0,0001922 | FCN2 |
| ENSGALG00000027420 | 141,8792 | 1,367365407 | 0,2942071 | 4,6476294 | 3,36E-06 | 3,86E-05 | POPDC3 |
| ENSGALG00000012971 | 180,16128 | 1,365874431 | 0,2750242 | 4,9663794 | 6,82E-07 | 9,15E-06 | CCDC170 |
| ENSGALG00000005852 | 42,716737 | 1,364112477 | 0,3537079 | 3,8566077 | 0,000115 | 0,0008438 | SRMS |
| ENSGALG00000020458 | 27,129951 | 1,361629377 | 0,3397843 | 4,0073347 | 6,14E-05 | 0,0004906 | PLA2G4E |
| ENSGALG00000013221 | 46,584811 | 1,36089575 | 0,3526259 | 3,8593185 | 0,0001137 | 0,0008363 | ARHGEF33 |
| ENSGALG00000016155 | 67,519598 | 1,359832396 | 0,3425225 | 3,9700528 | 7,19E-05 | 0,0005636 | COL19A1 |
| ENSGALG00000004840 | 833,06506 | 1,358453061 | 0,1689566 | 8,0402477 | 8,97E-16 | 4,78E-14 | RPH3A |
| ENSGALG00000014913 | 89,586553 | 1,358196979 | 0,3332154 | 4,0760324 | 4,58E-05 | 0,0003808 | ROS1 |
| ENSGALG00000028822 | 179,37935 | 1,357789969 | 0,2792664 | 4,861989 | 1,16E-06 | 1,50E-05 | RNF152 |
| ENSGALG00000001932 | 150,41219 | 1,357009459 | 0,310456 | 4,3710208 | 1,24E-05 | 0,0001245 | SRRM3 |
| ENSGALG00000006104 | 74,225594 | 1,356707559 | 0,3368845 | 4,027219 | 5,64E-05 | 0,0004571 | ADAMTSL3 |
| ENSGALG00000013914 | 741,46351 | 1,355434386 | 0,2231994 | 6,0727508 | 1,26E-09 | 2,95E-08 | SERINC4 |
| ENSGALG00000001863 | 95,573016 | 1,354146203 | 0,3495798 | 3,8736396 | 0,0001072 | 0,0007966 | VTG2 |
| ENSGALG00000024064 | 33,370003 | 1,353791502 | 0,3529207 | 3,835965 | 0,0001251 | 0,0009082 | BMP8B |
| ENSGALG00000017036 | 3151,4224 | 1,353726483 | 0,1434983 | 9,4337437 | 3,96E-21 | 3,41E-19 | LHFP |
| ENSGALG00000016342 | 64,688956 | 1,353678127 | 0,3480736 | 3,8890574 | 0,0001006 | 0,0007541 | MYOM2 |
| ENSGALG00000028162 | 25,784822 | 1,353451729 | 0,3407708 | 3,9717367 | 7,14E-05 | 0,0005606 | |
| ENSGALG00000025857 | 83,462878 | 1,352908608 | 0,3353198 | 4,0346816 | 5,47E-05 | 0,0004451 | |
| ENSGALG00000005898 | 30,859755 | 1,351366295 | 0,3483263 | 3,8795984 | 0,0001046 | 0,0007823 | DND1 |
| ENSGALG00000026817 | 190,25856 | 1,350039834 | 0,3072699 | 4,3936618 | 1,11E-05 | 0,0001138 | |
| ENSGALG00000011259 | 318,18554 | 1,346953088 | 0,237873 | 5,6624895 | 1,49E-08 | 2,86E-07 | RPE65 |
| ENSGALG00000015186 | 1129,1529 | 1,345677898 | 0,1669938 | 8,0582487 | 7,74E-16 | 4,20E-14 | KLHL14 |
| ENSGALG00000027568 | 132,01086 | 1,344832291 | 0,3108069 | 4,3269066 | 1,51E-05 | 0,0001475 | |
| ENSGALG00000003671 | 47,547146 | 1,341551712 | 0,3538875 | 3,7908993 | 0,0001501 | 0,0010672 | |
| ENSGALG00000007106 | 977,16123 | 1,340640592 | 0,2071983 | 6,4703268 | 9,78E-11 | 2,66E-09 | DNAH17 |
| ENSGALG00000002466 | 67,363583 | 1,337320513 | 0,3459423 | 3,8657324 | 0,0001108 | 0,0008178 | SLC2A5 |
| ENSGALG00000008283 | 62,956159 | 1,336619904 | 0,3480958 | 3,8398045 | 0,0001231 | 0,0008951 | CYP2J2L6 |
| ENSGALG00000009058 | 854,57904 | 1,336327117 | 0,176152 | 7,5862168 | 3,29E-14 | 1,46E-12 | ENTPD2 |
| ENSGALG00000014523 | 712,2168 | 1,335395904 | 0,1808886 | 7,3824225 | 1,55E-13 | 6,30E-12 | MATN2 |
| ENSGALG00000027624 | 90,7092 | 1,332865286 | 0,324427 | 4,1083672 | 3,98E-05 | 0,0003397 | |
| ENSGALG00000020380 | 160,77545 | 1,332253662 | 0,2975792 | 4,476972 | 7,57E-06 | 8,01E-05 | |
| ENSGALG00000014117 | 36,1409 | 1,330198207 | 0,3537602 | 3,7601691 | 0,0001698 | 0,0011923 | AVP |
| ENSGALG00000005330 | 2519,7711 | 1,328511787 | 0,1342416 | 9,8964213 | 4,31E-23 | 4,45E-21 | SNCAIP |
| ENSGALG00000013481 | 89,334053 | 1,327640637 | 0,3315342 | 4,0045364 | 6,21E-05 | 0,0004953 | IKZF3 |
| ENSGALG00000008434 | 453,34748 | 1,327566577 | 0,2234773 | 5,9404989 | 2,84E-09 | 6,30E-08 | SORCS3 |
| ENSGALG00000006134 | 56,091168 | 1,326807835 | 0,348696 | 3,8050556 | 0,0001418 | 0,0010148 | HOGA1 |
| ENSGALG00000004903 | 45,997304 | 1,326402261 | 0,3537178 | 3,7498888 | 0,0001769 | 0,0012333 | CHST8 |
| ENSGALG00000015309 | 102,61836 | 1,321942011 | 0,3301857 | 4,0036322 | 6,24E-05 | 0,0004969 | SCML4 |
| ENSGALG00000016551 | 5221,0551 | 1,321564075 | 0,1177455 | 11,223902 | 3,11E-29 | 5,11E-27 | AC025048.1 |
| ENSGALG00000013302 | 789,25879 | 1,321288665 | 0,1757375 | 7,5185339 | 5,54E-14 | 2,35E-12 | SYT7 |
| ENSGALG00000009315 | 90,474554 | 1,318764229 | 0,3335429 | 3,9538073 | 7,69E-05 | 0,000596 | PAPLN |
| ENSGALG00000027575 | 153,60071 | 1,316495304 | 0,2966273 | 4,4382135 | 9,07E-06 | 9,46E-05 | RAMP3 |
| ENSGALG00000003050 | 83,782773 | 1,314582894 | 0,3351464 | 3,9224135 | 8,77E-05 | 0,0006717 | ADRA2B |
| ENSGALG00000002792 | 26,932942 | 1,31306542 | 0,34508 | 3,8051042 | 0,0001417 | 0,0010148 | SPDEF |
| ENSGALG00000016368 | 37,747118 | 1,308737431 | 0,3539388 | 3,6976373 | 0,0002176 | 0,0014711 | KLHL38 |
| ENSGALG00000002389 | 532,14602 | 1,308458099 | 0,1976049 | 6,6215867 | 3,55E-11 | 1,03E-09 | ITGB4 |
| ENSGALG00000000164 | 213,8267 | 1,307977843 | 0,2786458 | 4,6940518 | 2,68E-06 | 3,15E-05 | MYBPH |
| ENSGALG00000007732 | 613,17499 | 1,306274335 | 0,1852621 | 7,0509534 | 1,78E-12 | 6,32E-11 | RTN4R |
| ENSGALG00000013838 | 384,54265 | 1,305966304 | 0,2158246 | 6,0510536 | 1,44E-09 | 3,34E-08 | |
| ENSGALG00000003316 | 80,926571 | 1,303670454 | 0,3335441 | 3,9085405 | 9,29E-05 | 0,0007043 | FMO3 |
| ENSGALG00000005802 | 409,59537 | 1,302158184 | 0,2262217 | 5,7561153 | 8,61E-09 | 1,72E-07 | FLT4 |
| ENSGALG00000016109 | 151,6721 | 1,299400996 | 0,288909 | 4,4976137 | 6,87E-06 | 7,31E-05 | KCNV1 |
| ENSGALG00000009700 | 477,10962 | 1,299326338 | 0,2326682 | 5,5844594 | 2,34E-08 | 4,28E-07 | PDK4 |
| ENSGALG00000006045 | 136,04518 | 1,297721233 | 0,3015022 | 4,3041847 | 1,68E-05 | 0,0001609 | SH3GL3 |
| ENSGALG00000006165 | 67,169846 | 1,296804529 | 0,3443385 | 3,7660745 | 0,0001658 | 0,0011656 | WDFY4 |
| ENSGALG00000002234 | 407,0013 | 1,296274176 | 0,217295 | 5,9655032 | 2,44E-09 | 5,48E-08 | UNC13D |
| ENSGALG00000002051 | 21,039089 | 1,294841211 | 0,3323973 | 3,895462 | 9,80E-05 | 0,0007377 | DNASE1 |
| ENSGALG00000008022 | 294,02931 | 1,293915137 | 0,239607 | 5,4001556 | 6,66E-08 | 1,11E-06 | TAPBP |
| ENSGALG00000000775 | 136,7775 | 1,291374316 | 0,3199881 | 4,0356942 | 5,44E-05 | 0,0004436 | CTSS |
| ENSGALG00000010540 | 264,70457 | 1,289706596 | 0,2432527 | 5,3019215 | 1,15E-07 | 1,81E-06 | TTC39A |
| ENSGALG00000005790 | 146,07742 | 1,289660896 | 0,3149604 | 4,0946765 | 4,23E-05 | 0,0003559 | DEPTOR |
| ENSGALG00000004859 | 354,90162 | 1,288425333 | 0,2477818 | 5,1998389 | 1,99E-07 | 3,02E-06 | ZNFX1 |
| ENSGALG00000003118 | 754,37839 | 1,286216079 | 0,1877588 | 6,8503656 | 7,37E-12 | 2,38E-10 | GPR124 |

| | | | | | | | |
|---|---|---|---|---|---|---|---|
| ENSGALG00000006779 | 194,18454 | 1,285794374 | 0,2904687 | 4,4266201 | 9,57E-06 | 9,92E-05 | PDZD3 |
| ENSGALG00000012362 | 940,67168 | 1,285048698 | 0,1725255 | 7,4484556 | 9,44E-14 | 3,88E-12 | THSD7B |
| ENSGALG00000004044 | 637,18108 | 1,285008651 | 0,1978538 | 6,4947372 | 8,32E-11 | 2,30E-09 | CSRP3 |
| ENSGALG00000003580 | 1991,2953 | 1,284312011 | 0,1942227 | 6,6125738 | 3,78E-11 | 1,09E-09 | MMP2 |
| ENSGALG00000002855 | 1183,6448 | 1,278753967 | 0,1621147 | 7,8879566 | 3,07E-15 | 1,50E-13 | SARDH |
| ENSGALG00000016564 | 80,407544 | 1,275019755 | 0,3413328 | 3,7354158 | 0,0001874 | 0,0012926 | PTK2B |
| ENSGALG00000001028 | 434,61935 | 1,272121671 | 0,2282446 | 5,5735011 | 2,50E-08 | 4,54E-07 | RNF43 |
| ENSGALG00000007078 | 141,27907 | 1,270778013 | 0,3066415 | 4,1441819 | 3,41E-05 | 0,0002963 | ENTPD1 |
| ENSGALG00000009689 | 798,33378 | 1,269454305 | 0,1789844 | 7,0925427 | 1,32E-12 | 4,74E-11 | PON2 |
| ENSGALG00000012740 | 435,21639 | 1,269186578 | 0,2159823 | 5,8763447 | 4,19E-09 | 8,99E-08 | ADAM12 |
| ENSGALG00000000909 | 237,54325 | 1,268775969 | 0,2625461 | 4,832584 | 1,35E-06 | 1,70E-05 | GFAP |
| ENSGALG00000016967 | 712,9034 | 1,267335006 | 0,2153024 | 5,8863033 | 3,95E-09 | 8,55E-08 | ENOX1 |
| ENSGALG00000019797 | 403,10583 | 1,266041829 | 0,2858329 | 4,429307 | 9,45E-06 | 9,81E-05 | |
| ENSGALG00000007562 | 30,687084 | 1,263491411 | 0,3496081 | 3,6140222 | 0,0003015 | 0,00195 | FGF4 |
| ENSGALG00000008953 | 507,74233 | 1,263384491 | 0,2035775 | 6,2059148 | 5,44E-10 | 1,33E-08 | AASS |
| ENSGALG00000001795 | 978,86532 | 1,261084777 | 0,1873221 | 6,7321726 | 1,67E-11 | 5,13E-10 | GFRA2 |
| ENSGALG00000012978 | 566,09757 | 1,261073108 | 0,1899643 | 6,6384742 | 3,17E-11 | 9,30E-10 | CDYL2 |
| ENSGALG00000006725 | 183,57715 | 1,259943343 | 0,3248929 | 3,8780263 | 0,0001053 | 0,0007865 | ACAN |
| ENSGALG00000014398 | 578,99454 | 1,259804122 | 0,2168754 | 5,8088844 | 6,29E-09 | 1,29E-07 | PPARGC1A |
| ENSGALG00000015598 | 190,82086 | 1,259472248 | 0,277866 | 4,5326611 | 5,82E-06 | 6,30E-05 | RASGRF2 |
| ENSGALG00000002647 | 518,74694 | 1,258418808 | 0,2451254 | 5,1337755 | 2,84E-07 | 4,13E-06 | PLS1 |
| ENSGALG00000003490 | 147,80461 | 1,25835771 | 0,2972254 | 4,2336817 | 2,30E-05 | 0,0002112 | ALDH3B1 |
| ENSGALG00000005458 | 78,515702 | 1,258264066 | 0,3397845 | 3,7031237 | 0,000213 | 0,0014432 | RPL3L |
| ENSGALG00000005843 | 363,33688 | 1,25767053 | 0,2724846 | 4,6155656 | 3,92E-06 | 4,43E-05 | EEF1A2 |
| ENSGALG00000006809 | 89,492709 | 1,25736391 | 0,3250463 | 3,8682605 | 0,0001096 | 0,0008106 | POU2AF1 |
| ENSGALG00000014014 | 690,53146 | 1,255177749 | 0,1779605 | 7,0531265 | 1,75E-12 | 6,24E-11 | LAMA2 |
| ENSGALG00000001348 | 34,354329 | 1,253857774 | 0,350669 | 3,5756163 | 0,0003494 | 0,0022157 | TEKT3 |
| ENSGALG00000021855 | 271,81428 | 1,251787198 | 0,252542 | 4,9567492 | 7,17E-07 | 9,57E-06 | CCDC39 |
| ENSGALG00000007193 | 236,11491 | 1,251769992 | 0,2743537 | 4,5626132 | 5,05E-06 | 5,59E-05 | HRASLS |
| ENSGALG00000018557 | 36,565332 | 1,251460959 | 0,3537306 | 3,537893 | 0,0004033 | 0,0025029 | SOD3 |
| ENSGALG00000025896 | 117,28876 | 1,251281202 | 0,3211612 | 3,8961159 | 9,77E-05 | 0,0007361 | PLPPR4 |
| ENSGALG00000012547 | 219,75607 | 1,25117298 | 0,2908698 | 4,3014887 | 1,70E-05 | 0,0001626 | ACVR1C |
| ENSGALG00000013356 | 85,223066 | 1,250931573 | 0,339616 | 3,6833703 | 0,0002302 | 0,001543 | IKBKE |
| ENSGALG00000003404 | 2332,0677 | 1,24748878 | 0,1390419 | 8,972036 | 2,91E-19 | 2,21E-17 | TFEB |
| ENSGALG00000001206 | 49,037007 | 1,247009104 | 0,3530765 | 3,5318378 | 0,0004127 | 0,0025563 | |
| ENSGALG00000028496 | 22,12018 | 1,24376929 | 0,3359139 | 3,7026432 | 0,0002134 | 0,0014453 | |
| ENSGALG00000007065 | 27,838516 | 1,239888336 | 0,3478406 | 3,5645304 | 0,0003645 | 0,0022955 | GJD4 |
| ENSGALG00000015345 | 537,38522 | 1,237723953 | 0,2000428 | 6,187296 | 6,12E-10 | 1,49E-08 | |
| ENSGALG00000028448 | 68,098086 | 1,236909029 | 0,3465147 | 3,5695715 | 0,0003576 | 0,002257 | SH2D1B |
| ENSGALG00000001564 | 906,15479 | 1,234344815 | 0,196986 | 6,2661565 | 3,70E-10 | 9,23E-09 | ATP2A3 |
| ENSGALG00000023036 | 562,73194 | 1,234170444 | 0,2058267 | 5,9961639 | 2,02E-09 | 4,57E-08 | FRMPD1 |
| ENSGALG00000010927 | 22,212585 | 1,233723332 | 0,3353328 | 3,6791008 | 0,0002341 | 0,0015644 | MEPE |
| ENSGALG00000028681 | 168,61801 | 1,232657979 | 0,2905515 | 4,2424766 | 2,21E-05 | 0,0002043 | |
| ENSGALG00000015573 | 43,775019 | 1,232214926 | 0,3535094 | 3,4856638 | 0,0004909 | 0,0029548 | FHL5 |
| ENSGALG00000005298 | 800,81856 | 1,231219861 | 0,1718969 | 7,1625504 | 7,92E-13 | 2,92E-11 | ST6GALNAC6 |
| ENSGALG00000016112 | 539,87209 | 1,228482119 | 0,2093942 | 5,8668384 | 4,44E-09 | 9,46E-08 | NOV |
| ENSGALG00000000107 | 237,72923 | 1,227291307 | 0,277099 | 4,4290711 | 9,46E-06 | 9,81E-05 | TRIM7.1 |
| ENSGALG00000002086 | 83,650156 | 1,225708617 | 0,3336205 | 3,6739612 | 0,0002388 | 0,0015908 | RASL10B |
| ENSGALG00000002667 | 699,19963 | 1,225466735 | 0,1768396 | 6,9298206 | 4,21E-12 | 1,40E-10 | OLFML2B |
| ENSGALG00000013431 | 35,449073 | 1,223926576 | 0,3535105 | 3,4622067 | 0,0005358 | 0,0031756 | RAX2 |
| ENSGALG00000028031 | 210,36093 | 1,22342747 | 0,2837755 | 4,3112512 | 1,62E-05 | 0,0001564 | C1QL1 |
| ENSGALG00000012019 | 150,4401 | 1,222353012 | 0,2981526 | 4,0997566 | 4,14E-05 | 0,0003505 | ARSJ |
| ENSGALG00000000048 | 677,12707 | 1,221920851 | 0,1990761 | 6,1379597 | 8,36E-10 | 2,00E-08 | |
| ENSGALG00000028946 | 391,05664 | 1,217039733 | 0,2510887 | 4,8470509 | 1,25E-06 | 1,59E-05 | CSRP2 |
| ENSGALG00000014407 | 118,55781 | 1,216198172 | 0,3199077 | 3,801716 | 0,0001437 | 0,0010259 | CASR |
| ENSGALG00000012658 | 2563,9636 | 1,214376265 | 0,1523893 | 7,9689058 | 1,60E-15 | 8,19E-14 | TCP11L2 |
| ENSGALG00000028402 | 33,811586 | 1,211737057 | 0,3495685 | 3,466379 | 0,0005275 | 0,0031377 | |
| ENSGALG00000003345 | 1065,5168 | 1,211546204 | 0,160937 | 7,5280764 | 5,15E-14 | 2,19E-12 | PLCD4 |
| ENSGALG00000005083 | 47,868272 | 1,210240985 | 0,3512641 | 3,4453881 | 0,0005702 | 0,0033455 | SLC5A10 |
| ENSGALG00000003803 | 110,85757 | 1,208011558 | 0,3397797 | 3,555279 | 0,0003776 | 0,0023603 | DAB2 |
| ENSGALG00000026578 | 55,333978 | 1,20748664 | 0,3538872 | 3,4120664 | 0,0006447 | 0,0037053 | |
| ENSGALG00000014126 | 18,992333 | 1,207239433 | 0,3279747 | 3,6808919 | 0,0002324 | 0,0015558 | ENDOUL |
| ENSGALG00000015764 | 5967,7523 | 1,20416113 | 0,1908692 | 6,3088274 | 2,81E-10 | 7,13E-09 | FABP5 |
| ENSGALG00000015528 | 170,23305 | 1,204136401 | 0,3006436 | 4,0051957 | 6,20E-05 | 0,0004943 | JAKMIP1 |

| | | | | | | | |
|---|---|---|---|---|---|---|---|
| ENSGALG00000002355 | 234,00639 | 1,203134529 | 0,2548917 | 4,7201799 | 2,36E-06 | 2,80E-05 | RBPMS2 |
| ENSGALG00000004641 | 146,28084 | 1,20257315 | 0,2965454 | 4,0552753 | 5,01E-05 | 0,0004122 | MYO5C |
| ENSGALG00000026903 | 111,36196 | 1,200671289 | 0,3283141 | 3,6570806 | 0,0002551 | 0,0016869 | CALB2 |
| ENSGALG00000004745 | 1802,3575 | 1,199713581 | 0,141877 | 8,4560121 | 2,77E-17 | 1,74E-15 | NATD1 |
| ENSGALG00000013890 | 33,560749 | 1,1996045 | 0,3520141 | 3,4078307 | 0,0006548 | 0,0037491 | MC5R |
| ENSGALG00000019280 | 990,72458 | 1,199070585 | 0,1733998 | 6,915063 | 4,68E-12 | 1,54E-10 | LMO3 |
| ENSGALG00000004066 | 55,61946 | 1,199060923 | 0,3539016 | 3,3881191 | 0,0007037 | 0,0039939 | HTR6 |
| ENSGALG00000002082 | 57,1028 | 1,198569946 | 0,3484813 | 3,4394093 | 0,000583 | 0,0034066 | AMPD1 |
| ENSGALG00000013090 | 44,714809 | 1,197838739 | 0,3538143 | 3,3855007 | 0,0007105 | 0,0040265 | LOXL4 |
| ENSGALG00000014696 | 222,59602 | 1,197793931 | 0,2597544 | 4,6112558 | 4,00E-06 | 4,51E-05 | ZBTB7C |
| ENSGALG00000006922 | 19,384458 | 1,194740346 | 0,3271281 | 3,6522095 | 0,00026 | 0,0017142 | KCNJ14 |
| ENSGALG00000011489 | 4656,6813 | 1,193784841 | 0,1216657 | 9,8120096 | 1,00E-22 | 9,93E-21 | |
| ENSGALG00000016590 | 18,890623 | 1,192009768 | 0,3270811 | 3,6443865 | 0,000268 | 0,001757 | TLR7 |
| ENSGALG00000002407 | 144,98036 | 1,191296249 | 0,3041418 | 3,9169112 | 8,97E-05 | 0,0006837 | HYDIN |
| ENSGALG00000012890 | 273,96599 | 1,191168201 | 0,2640739 | 4,5107378 | 6,46E-06 | 6,90E-05 | DGKI |
| ENSGALG00000027218 | 239,44089 | 1,190798556 | 0,2757781 | 4,317959 | 1,57E-05 | 0,0001524 | |
| ENSGALG00000017040 | 33,080589 | 1,190245244 | 0,3523261 | 3,3782489 | 0,0007295 | 0,0041177 | |
| ENSGALG00000008362 | 175,02358 | 1,190011885 | 0,2754196 | 4,3207232 | 1,56E-05 | 0,000151 | SSTR4 |
| ENSGALG00000004521 | 401,35326 | 1,187877806 | 0,2197321 | 5,4060281 | 6,44E-08 | 1,08E-06 | GPX3 |
| ENSGALG00000006074 | 272,24078 | 1,186152058 | 0,28359 | 4,1826293 | 2,88E-05 | 0,0002563 | SOX6 |
| ENSGALG00000003842 | 39,781188 | 1,185854219 | 0,3538378 | 3,3514066 | 0,000804 | 0,0044662 | GHRH |
| ENSGALG00000021882 | 58,697437 | 1,185518376 | 0,3468224 | 3,4182285 | 0,0006303 | 0,0036347 | |
| ENSGALG00000002744 | 252,46116 | 1,184999187 | 0,2518537 | 4,705109 | 2,54E-06 | 3,00E-05 | PCDH15 |
| ENSGALG00000006768 | 37,969199 | 1,18499895 | 0,3536582 | 3,3506896 | 0,0008061 | 0,0044728 | |
| ENSGALG00000014812 | 711,71818 | 1,183271769 | 0,1870163 | 6,3271055 | 2,50E-10 | 6,39E-09 | SIDT1 |
| ENSGALG00000017331 | 630,73395 | 1,182854332 | 0,188652 | 6,2700317 | 3,61E-10 | 9,05E-09 | GDPD5 |
| ENSGALG00000005547 | 144,89722 | 1,180323703 | 0,2989787 | 3,9478524 | 7,89E-05 | 0,0006093 | PIK3AP1 |
| ENSGALG00000017291 | 662,46181 | 1,180080414 | 0,209044 | 5,6451284 | 1,65E-08 | 3,14E-07 | FAR2 |
| ENSGALG00000006308 | 28,078472 | 1,179461013 | 0,3370947 | 3,4989008 | 0,0004672 | 0,0028338 | |
| ENSGALG00000013112 | 320,80404 | 1,179282262 | 0,2330597 | 5,0600015 | 4,19E-07 | 5,90E-06 | RERGL |
| ENSGALG00000007333 | 325,51856 | 1,178440858 | 0,2584425 | 4,5597804 | 5,12E-06 | 5,65E-05 | UBAP1L |
| ENSGALG00000001021 | 191,27134 | 1,17756894 | 0,2838137 | 4,1490906 | 3,34E-05 | 0,0002911 | MYOCD |
| ENSGALG00000003821 | 322,04402 | 1,177115475 | 0,2347575 | 5,0141754 | 5,33E-07 | 7,38E-06 | RAMP1 |
| ENSGALG00000003164 | 243,86317 | 1,17697714 | 0,2590714 | 4,5430608 | 5,54E-06 | 6,04E-05 | DNM3 |
| ENSGALG00000016651 | 946,90096 | 1,174964335 | 0,1848632 | 6,35586 | 2,07E-10 | 5,41E-09 | TDH |
| ENSGALG00000002435 | 70,191489 | 1,174947125 | 0,34691 | 3,3868933 | 0,0007069 | 0,0040085 | DUOX2 |
| ENSGALG00000001331 | 1782,6492 | 1,174179888 | 0,1437468 | 8,16839 | 3,13E-16 | 1,75E-14 | ST14 |
| ENSGALG00000009114 | 216,59264 | 1,170782693 | 0,2608026 | 4,4891525 | 7,15E-06 | 7,60E-05 | DGAT2 |
| ENSGALG00000028580 | 59,186277 | 1,170297276 | 0,3494653 | 3,3488229 | 0,0008116 | 0,0044934 | C15orf48 |
| ENSGALG00000006099 | 3185,4261 | 1,169037408 | 0,1372144 | 8,5197869 | 1,60E-17 | 1,03E-15 | ZFPM1 |
| ENSGALG00000014261 | 23192,424 | 1,166708788 | 0,1480767 | 7,8790822 | 3,30E-15 | 1,59E-13 | UCHL1 |
| ENSGALG00000006627 | 44,723529 | 1,164076921 | 0,352923 | 3,2983882 | 0,0009724 | 0,0052103 | UPB1 |
| ENSGALG00000012668 | 622,84546 | 1,1628752 | 0,2448035 | 4,7502395 | 2,03E-06 | 2,44E-05 | DCDC2 |
| ENSGALG00000011446 | 368,84385 | 1,161895934 | 0,2215158 | 5,2452066 | 1,56E-07 | 2,42E-06 | TNFAIP2 |
| ENSGALG00000005630 | 38,097087 | 1,161740711 | 0,3428362 | 3,3886172 | 0,0007025 | 0,0039883 | CRYBB1 |
| ENSGALG00000023879 | 439,21059 | 1,160823609 | 0,2116765 | 5,4839511 | 4,16E-08 | 7,24E-07 | |
| ENSGALG00000007596 | 99,216367 | 1,160341621 | 0,3237884 | 3,5836422 | 0,0003388 | 0,0021608 | |
| ENSGALG00000005721 | 236,12764 | 1,159323873 | 0,251452 | 4,6105181 | 4,02E-06 | 4,52E-05 | DGKG |
| ENSGALG00000013911 | 546,72587 | 1,159209839 | 0,2024827 | 5,7249821 | 1,03E-08 | 2,03E-07 | ZC3HAV1 |
| ENSGALG00000011418 | 18,242274 | 1,158885244 | 0,3251653 | 3,5639875 | 0,0003653 | 0,002297 | CCR6 |
| ENSGALG00000028489 | 20,536962 | 1,158414063 | 0,3317704 | 3,4916132 | 0,0004801 | 0,0029001 | |
| ENSGALG00000001031 | 327,10314 | 1,156740289 | 0,2440319 | 4,7401197 | 2,14E-06 | 2,56E-05 | HSF5 |
| ENSGALG00000002016 | 953,70998 | 1,156233761 | 0,1649483 | 7,0096752 | 2,39E-12 | 8,30E-11 | PRKCD |
| ENSGALG00000001095 | 18,255475 | 1,156162616 | 0,3249866 | 3,5575704 | 0,0003743 | 0,0023463 | CCDC183 |
| ENSGALG00000013980 | 288,26492 | 1,155390063 | 0,2983628 | 3,8724333 | 0,0001078 | 0,0007995 | SLC2A12 |
| ENSGALG00000026241 | 639,21895 | 1,154849081 | 0,2302822 | 5,0149306 | 5,31E-07 | 7,36E-06 | GRIN3B |
| ENSGALG00000006208 | 100,69013 | 1,154488936 | 0,3219519 | 3,5859053 | 0,0003359 | 0,0021441 | OTOG |
| ENSGALG00000009641 | 18352,412 | 1,153845642 | 0,1458991 | 7,9085187 | 2,60E-15 | 1,28E-13 | COL1A2 |
| ENSGALG00000012288 | 52,934002 | 1,152071153 | 0,3501204 | 3,2905002 | 0,0010001 | 0,0053293 | SLC16A8 |
| ENSGALG00000015932 | 523,70552 | 1,152042382 | 0,2240419 | 5,1420853 | 2,72E-07 | 3,98E-06 | KCNQ5 |
| ENSGALG00000003789 | 56,043715 | 1,151405842 | 0,3500328 | 3,2894227 | 0,0010039 | 0,0053476 | FAM78A |
| ENSGALG00000010365 | 313,07869 | 1,151056068 | 0,2620814 | 4,39198 | 1,12E-05 | 0,0001143 | ESRRB |
| ENSGALG00000017691 | 74,633315 | 1,149429932 | 0,3427848 | 3,3532112 | 0,0007988 | 0,0044427 | |
| ENSGALG00000017374 | 478,61541 | 1,149174538 | 0,2222568 | 5,1704808 | 2,33E-07 | 3,48E-06 | SLITRK3 |

| | | | | | | | |
|---|---|---|---|---|---|---|---|
| ENSGALG00000028710 | 21,101173 | 1,148265888 | 0,3321095 | 3,4574919 | 0,0005452 | 0,0032233 | |
| ENSGALG00000012374 | 515,33143 | 1,147327656 | 0,1949041 | 5,8866278 | 3,94E-09 | 8,55E-08 | FRMD6 |
| ENSGALG00000016328 | 5259,3238 | 1,146952103 | 0,1404012 | 8,1691038 | 3,11E-16 | 1,75E-14 | GSTA |
| ENSGALG00000025721 | 19,243146 | 1,146921853 | 0,324147 | 3,5382766 | 0,0004027 | 0,0025004 | |
| ENSGALG00000012365 | 9658,5394 | 1,146353139 | 0,1101377 | 10,408358 | 2,27E-25 | 2,97E-23 | PYGL |
| ENSGALG00000005540 | 392,97544 | 1,145405784 | 0,219512 | 5,2179637 | 1,81E-07 | 2,76E-06 | MICAL2 |
| ENSGALG00000028702 | 48,721731 | 1,145044679 | 0,3522221 | 3,2509165 | 0,0011503 | 0,0060053 | EVX1 |
| ENSGALG00000022819 | 93,418767 | 1,14482978 | 0,3365037 | 3,4021308 | 0,0006686 | 0,0038249 | PCP4 |
| ENSGALG00000001968 | 55,570176 | 1,14394352 | 0,3522778 | 3,2472771 | 0,0011651 | 0,006064 | SYT6 |
| ENSGALG00000012505 | 1173,094 | 1,142096363 | 0,156235 | 7,3101186 | 2,67E-13 | 1,05E-11 | LRFN5 |
| ENSGALG00000015758 | 1198,984 | 1,141437782 | 0,170779 | 6,6837117 | 2,33E-11 | 6,96E-10 | PAG1 |
| ENSGALG00000003876 | 41,544916 | 1,140755556 | 0,3447861 | 3,3085895 | 0,0009377 | 0,0050721 | TIMD4 |
| ENSGALG00000009791 | 992,57388 | 1,140657464 | 0,2126878 | 5,3630614 | 8,18E-08 | 1,34E-06 | PROX1 |
| ENSGALG00000028130 | 1683,1758 | 1,140619832 | 0,1589917 | 7,1740827 | 7,28E-13 | 2,72E-11 | KIAA1644 |
| ENSGALG00000010673 | 22,777798 | 1,139845847 | 0,3403007 | 3,3495255 | 0,0008095 | 0,0044839 | SLC1A7 |
| ENSGALG00000006436 | 58,776846 | 1,138634151 | 0,3530266 | 3,2253491 | 0,0012582 | 0,0064643 | AQP12 |
| ENSGALG00000000620 | 143,16144 | 1,137872957 | 0,293674 | 3,8746119 | 0,0001068 | 0,0007954 | FABP3 |
| ENSGALG00000013818 | 524,97865 | 1,136933045 | 0,2108978 | 5,3909206 | 7,01E-08 | 1,17E-06 | CITED2 |
| ENSGALG00000007236 | 26,585317 | 1,136243459 | 0,3440899 | 3,30217 | 0,0009594 | 0,005167 | PLA2G1BL |
| ENSGALG00000023643 | 36,522348 | 1,136155947 | 0,3539078 | 3,2103166 | 0,0013259 | 0,0067509 | |
| ENSGALG00000007129 | 549,58573 | 1,134513156 | 0,2759889 | 4,1107204 | 3,94E-05 | 0,0003371 | ALDH1A3 |
| ENSGALG00000001442 | 382,78952 | 1,131898366 | 0,2156493 | 5,248791 | 1,53E-07 | 2,37E-06 | OPCML |
| ENSGALG00000000217 | 558,86585 | 1,131400306 | 0,2196843 | 5,1501182 | 2,60E-07 | 3,84E-06 | PPFIA4 |
| ENSGALG00000000911 | 18,802113 | 1,131377385 | 0,3232235 | 3,5002948 | 0,0004647 | 0,002821 | IL19 |
| ENSGALG00000003028 | 252,86338 | 1,131167716 | 0,2536539 | 4,4594925 | 8,22E-06 | 8,64E-05 | ADAMTS13 |
| ENSGALG00000027626 | 26,263554 | 1,129092221 | 0,3472331 | 3,251684 | 0,0011472 | 0,0059985 | |
| ENSGALG00000008063 | 112,83661 | 1,129089778 | 0,3199799 | 3,5286266 | 0,0004177 | 0,0025816 | NOX5 |
| ENSGALG00000016592 | 75,509326 | 1,127895523 | 0,3458232 | 3,2614806 | 0,0011083 | 0,0058217 | PRPS2 |
| ENSGALG00000021529 | 183,02929 | 1,127864143 | 0,273066 | 4,1303714 | 3,62E-05 | 0,0003123 | GP1BB |
| ENSGALG00000006503 | 1322,5302 | 1,127315674 | 0,1739831 | 6,4794533 | 9,21E-11 | 2,52E-09 | VDAC1 |
| ENSGALG00000026333 | 170,5148 | 1,126889713 | 0,280963 | 4,0108121 | 6,05E-05 | 0,0004843 | GNGT2 |
| ENSGALG00000015348 | 3709,6553 | 1,126202791 | 0,1601094 | 7,03396 | 2,01E-12 | 7,08E-11 | ALCAM |
| ENSGALG00000027872 | 562,41556 | 1,125834202 | 0,2065498 | 5,4506672 | 5,02E-08 | 8,57E-07 | |
| ENSGALG00000007964 | 712,00528 | 1,125506336 | 0,1939073 | 5,8043524 | 6,46E-09 | 1,32E-07 | GATSL3 |
| ENSGALG00000008940 | 122,07312 | 1,125469701 | 0,3044547 | 3,6966742 | 0,0002184 | 0,0014752 | SPTBN5 |
| ENSGALG00000012252 | 31,066473 | 1,124245419 | 0,3463855 | 3,2456478 | 0,0011718 | 0,0060895 | |
| ENSGALG00000009237 | 283,64685 | 1,124183808 | 0,2526532 | 4,4495132 | 8,61E-06 | 9,03E-05 | TLR2A |
| ENSGALG00000008629 | 362,4306 | 1,122527446 | 0,2874644 | 3,9049269 | 9,43E-05 | 0,0007125 | |
| ENSGALG00000004649 | 115,31891 | 1,122312882 | 0,3390875 | 3,3098029 | 0,0009336 | 0,0050602 | TRPV1 |
| ENSGALG00000016821 | 91,586414 | 1,121951764 | 0,3275569 | 3,4252114 | 0,0006143 | 0,0035683 | TMEM255B |
| ENSGALG00000000994 | 36,33755 | 1,120119947 | 0,3533528 | 3,1699761 | 0,0015245 | 0,0075368 | RAB42 |
| ENSGALG00000009943 | 1812,2354 | 1,119992507 | 0,1420982 | 7,8818219 | 3,23E-15 | 1,56E-13 | FAM117A |
| ENSGALG00000028795 | 3146,8331 | 1,118039364 | 0,160426 | 6,9691886 | 3,19E-12 | 1,09E-10 | CADM3 |
| ENSGALG00000026758 | 74,554613 | 1,11741437 | 0,3353836 | 3,33175 | 0,000863 | 0,0047277 | |
| ENSGALG00000001204 | 1443,1255 | 1,117076992 | 0,1466887 | 7,6152892 | 2,63E-14 | 1,19E-12 | ENKD1 |
| ENSGALG00000012055 | 30,631938 | 1,115864912 | 0,3495485 | 3,1923038 | 0,0014114 | 0,0070989 | TGFBR2L |
| ENSGALG00000015469 | 3626,2078 | 1,115704157 | 0,1321025 | 8,4457457 | 3,02E-17 | 1,89E-15 | IGSF3 |
| ENSGALG00000009963 | 76,780039 | 1,114244364 | 0,3375856 | 3,3006277 | 0,0009647 | 0,0051852 | LYZ |
| ENSGALG00000011414 | 712,8987 | 1,112714675 | 0,1796621 | 6,1933742 | 5,89E-10 | 1,44E-08 | AMDHD1 |
| ENSGALG00000015454 | 87,061362 | 1,111985108 | 0,3376828 | 3,2929875 | 0,0009913 | 0,0052948 | CA8 |
| ENSGALG00000010233 | 149,38519 | 1,111330421 | 0,289789 | 3,8349641 | 0,0001256 | 0,0009114 | SYNDIG1L |
| ENSGALG00000024411 | 95,571509 | 1,110994905 | 0,322591 | 3,4439741 | 0,0005732 | 0,0033582 | C2orf82 |
| ENSGALG00000003016 | 111,9759 | 1,109545556 | 0,3100714 | 3,5783554 | 0,0003458 | 0,0021977 | MASP2 |
| ENSGALG00000006792 | 447,85614 | 1,109047707 | 0,21888 | 5,0669216 | 4,04E-07 | 5,71E-06 | VSTM2L |
| ENSGALG00000009823 | 207,86307 | 1,108978484 | 0,2670183 | 4,1531933 | 3,28E-05 | 0,0002863 | PTPRE |
| ENSGALG00000027501 | 1178,2304 | 1,108782799 | 0,1555553 | 7,1279016 | 1,02E-12 | 3,72E-11 | ANPEP |
| ENSGALG00000004017 | 335,24395 | 1,108435951 | 0,2275944 | 4,8702245 | 1,11E-06 | 1,44E-05 | TOR1AIP2 |
| ENSGALG00000025912 | 62,633971 | 1,108425226 | 0,3437693 | 3,2243285 | 0,0012627 | 0,00648 | |
| ENSGALG00000000158 | 209,10615 | 1,107475494 | 0,2643096 | 4,1900691 | 2,79E-05 | 0,0002495 | DMA |
| ENSGALG00000002610 | 604,59063 | 1,105862871 | 0,189678 | 5,8302128 | 5,54E-09 | 1,15E-07 | |
| ENSGALG00000023621 | 702,55001 | 1,105773477 | 0,197195 | 5,6075141 | 2,05E-08 | 3,82E-07 | SFXN3 |
| ENSGALG00000009574 | 24,975698 | 1,105678443 | 0,3438055 | 3,216 | 0,0012999 | 0,0066483 | IL12A |
| ENSGALG00000006000 | 170,90987 | 1,105166021 | 0,292418 | 3,7794044 | 0,0001572 | 0,0011107 | ADAMTS2 |
| ENSGALG00000001011 | 623,47251 | 1,104831537 | 0,2206424 | 5,00734 | 5,52E-07 | 7,61E-06 | CNGB1 |

| | | | | | | | |
|---|---|---|---|---|---|---|---|
| ENSGALG00000011473 | 347,60043 | 1,104703027 | 0,2275411 | 4,8549595 | 1,20E-06 | 1,54E-05 | RPS6KA2 |
| ENSGALG00000026834 | 233,70852 | 1,10373431 | 0,2598301 | 4,2479073 | 2,16E-05 | 0,0002001 | PPAP2C |
| ENSGALG00000006003 | 58,995483 | 1,10329195 | 0,3504724 | 3,1480138 | 0,0016438 | 0,0080223 | C10orf10 |
| ENSGALG00000001281 | 270,65777 | 1,101845994 | 0,2489516 | 4,4259451 | 9,60E-06 | 9,94E-05 | PHOSPHO1 |
| ENSGALG00000006407 | 926,88526 | 1,100961832 | 0,1782262 | 6,1773296 | 6,52E-10 | 1,58E-08 | |
| ENSGALG00000011003 | 180,05065 | 1,099169303 | 0,2856147 | 3,8484334 | 0,0001189 | 0,0008669 | SLC35F3 |
| ENSGALG00000000168 | 20,237218 | 1,099055711 | 0,3363625 | 3,2674737 | 0,0010851 | 0,0057152 | ADORA1 |
| ENSGALG00000014798 | 1444,6552 | 1,098754437 | 0,1853247 | 5,9288075 | 3,05E-09 | 6,69E-08 | RP11-296A16.1 |
| ENSGALG00000015076 | 118,81874 | 1,094512369 | 0,3047536 | 3,5914667 | 0,0003288 | 0,0021068 | |
| ENSGALG00000002579 | 438,86248 | 1,09231668 | 0,2407976 | 4,5362436 | 5,73E-06 | 6,21E-05 | RIMBP2 |
| ENSGALG00000027102 | 27,931361 | 1,090659115 | 0,3485566 | 3,129073 | 0,0017536 | 0,0084616 | |
| ENSGALG00000007740 | 159,04973 | 1,089704597 | 0,297943 | 3,6574259 | 0,0002548 | 0,0016854 | |
| ENSGALG00000004972 | 200,69424 | 1,089536909 | 0,3028577 | 3,5975215 | 0,0003213 | 0,0020613 | ALKBH5 |
| ENSGALG00000004538 | 131,20236 | 1,088596778 | 0,3003085 | 3,6249277 | 0,000289 | 0,001882 | TNN |
| ENSGALG00000005470 | 915,92774 | 1,087631823 | 0,1770332 | 6,1436603 | 8,06E-10 | 1,93E-08 | LPPR5 |
| ENSGALG00000006189 | 81,047665 | 1,08750661 | 0,3396623 | 3,2017288 | 0,0013661 | 0,0069166 | CRYGN |
| ENSGALG00000001716 | 219,08602 | 1,084481493 | 0,276627 | 3,9203751 | 8,84E-05 | 0,0006759 | C15orf59 |
| ENSGALG00000017106 | 609,78404 | 1,082115941 | 0,1927819 | 5,6131603 | 1,99E-08 | 3,72E-07 | ATP8A2 |
| ENSGALG00000014264 | 24,119483 | 1,081789708 | 0,3421863 | 3,1614055 | 0,0015701 | 0,0077368 | NSUN7 |
| ENSGALG00000001470 | 804,49858 | 1,081645785 | 0,1818986 | 5,9464233 | 2,74E-09 | 6,10E-08 | STOML1 |
| ENSGALG00000023636 | 80,78387 | 1,079731402 | 0,3357478 | 3,2159003 | 0,0013004 | 0,0066483 | REM1 |
| ENSGALG00000021303 | 70,179062 | 1,078464744 | 0,3387078 | 3,1840559 | 0,0014523 | 0,0072578 | NPY7R |
| ENSGALG00000011353 | 167,02407 | 1,077984355 | 0,2948088 | 3,6565536 | 0,0002556 | 0,0016887 | TNNI3K |
| ENSGALG00000027354 | 35,225101 | 1,077448801 | 0,3534209 | 3,048628 | 0,0022989 | 0,0105741 | MPND |
| ENSGALG00000026651 | 469,2115 | 1,076820994 | 0,2627359 | 4,0984919 | 4,16E-05 | 0,0003517 | SCARA3 |
| ENSGALG00000005031 | 222,86675 | 1,076697936 | 0,2761103 | 3,8995209 | 9,64E-05 | 0,000727 | DRP2 |
| ENSGALG00000005076 | 658,30851 | 1,075685444 | 0,2013532 | 5,3422809 | 9,18E-08 | 1,49E-06 | GUCY1B4 |
| ENSGALG00000013180 | 23,783592 | 1,075255786 | 0,3379303 | 3,1818859 | 0,0014632 | 0,0072892 | SLC6A18 |
| ENSGALG00000004043 | 1743,1221 | 1,074636101 | 0,1810984 | 5,9339914 | 2,96E-09 | 6,52E-08 | NBL1 |
| ENSGALG00000008822 | 37,095962 | 1,073866227 | 0,3512806 | 3,0570039 | 0,0022356 | 0,0103284 | |
| ENSGALG00000000420 | 275,78606 | 1,072751202 | 0,2433406 | 4,4084345 | 1,04E-05 | 0,000107 | KLHDC7A |
| ENSGALG00000021559 | 51,236888 | 1,072736007 | 0,3500464 | 3,0645537 | 0,00218 | 0,0101332 | CD14 |
| ENSGALG00000003640 | 1881,5268 | 1,071788322 | 0,1472843 | 7,2770018 | 3,41E-13 | 1,33E-11 | FAM20C |
| ENSGALG00000009355 | 164,05056 | 1,070935457 | 0,2815733 | 3,8033985 | 0,0001427 | 0,0010204 | CLDN11 |
| ENSGALG00000026748 | 19,49545 | 1,069944148 | 0,3278792 | 3,2632271 | 0,0011015 | 0,0057881 | MYH1A |
| ENSGALG00000020492 | 36,098876 | 1,069561713 | 0,3538691 | 3,0224786 | 0,0025071 | 0,0113329 | MUC6 |
| ENSGALG00000004406 | 128,12145 | 1,066534863 | 0,3115962 | 3,4228111 | 0,0006198 | 0,0035892 | LRRC43 |
| ENSGALG00000006370 | 31,168172 | 1,066397762 | 0,3455845 | 3,0857802 | 0,0020302 | 0,0095644 | TULP1 |
| ENSGALG00000012377 | 68,881814 | 1,065846022 | 0,3384914 | 3,148813 | 0,0016394 | 0,0080143 | HNMT |
| ENSGALG00000006344 | 48,374655 | 1,06541292 | 0,3514839 | 3,0311857 | 0,002436 | 0,0110921 | GSTT1 |
| ENSGALG00000010752 | 133,53254 | 1,063752645 | 0,3170147 | 3,3555312 | 0,0007921 | 0,0044138 | CDCP2 |
| ENSGALG00000021049 | 25,160532 | 1,063749147 | 0,3349924 | 3,175443 | 0,0014961 | 0,0074177 | FNDC11 |
| ENSGALG00000006374 | 154,22163 | 1,06372742 | 0,3461141 | 3,0733436 | 0,0021167 | 0,0099138 | TBX6 |
| ENSGALG00000007780 | 1769,7678 | 1,063721106 | 0,1641554 | 6,4799655 | 9,17E-11 | 2,52E-09 | EIF4E3 |
| ENSGALG00000004149 | 50,119553 | 1,06296288 | 0,353007 | 3,0111663 | 0,0026025 | 0,01169 | KCNK15 |
| ENSGALG00000021542 | 348,90792 | 1,061917275 | 0,2447028 | 4,3396212 | 1,43E-05 | 0,000141 | SNCB |
| ENSGALG00000002055 | 5059,126 | 1,061032342 | 0,1289624 | 8,2274527 | 1,91E-16 | 1,10E-14 | HES4 |
| ENSGALG00000028742 | 318,00325 | 1,06018982 | 0,2447624 | 4,3315054 | 1,48E-05 | 0,0001453 | |
| ENSGALG00000001040 | 379,37277 | 1,060173693 | 0,2265161 | 4,6803463 | 2,86E-06 | 3,34E-05 | MEGF6 |
| ENSGALG00000008734 | 1277,1497 | 1,060092794 | 0,1784849 | 5,9393992 | 2,86E-09 | 6,33E-08 | PCSK2 |
| ENSGALG00000025771 | 418,4548 | 1,059868939 | 0,2130592 | 4,9745277 | 6,54E-07 | 8,82E-06 | PLCD3 |
| ENSGALG00000017311 | 141,94595 | 1,059578058 | 0,2931335 | 3,6146605 | 0,0003007 | 0,0019465 | P4HA3 |
| ENSGALG00000026017 | 67,59259 | 1,059475908 | 0,3510549 | 3,0179783 | 0,0025447 | 0,0114759 | DRGX |
| ENSGALG00000002554 | 211,29924 | 1,058725863 | 0,2726542 | 3,8830359 | 0,0001032 | 0,0007721 | TSPAN2 |
| ENSGALG00000011898 | 1941,7841 | 1,056447793 | 0,1400944 | 7,5409707 | 4,66E-14 | 2,00E-12 | FAM109B |
| ENSGALG00000012545 | 20,395626 | 1,051934631 | 0,3331292 | 3,1577373 | 0,00159 | 0,0078066 | CYTIP |
| ENSGALG00000016620 | 37,122352 | 1,049979269 | 0,351653 | 2,9858389 | 0,002828 | 0,0125541 | PNOC |
| ENSGALG00000010497 | 795,79291 | 1,048579632 | 0,1787889 | 5,8649021 | 4,49E-09 | 9,56E-08 | SLC29A4 |
| ENSGALG00000000503 | 90,521684 | 1,0483641 | 0,3288307 | 3,1881578 | 0,0014318 | 0,0071802 | PADI2 |
| ENSGALG00000007959 | 24,992818 | 1,04699816 | 0,3339903 | 3,134816 | 0,0017196 | 0,0083253 | SLC26A3 |
| ENSGALG00000012045 | 94,161025 | 1,046487546 | 0,3346828 | 3,1268044 | 0,0017672 | 0,0085078 | SLC12A8 |
| ENSGALG00000009340 | 441,50856 | 1,044750001 | 0,2505951 | 4,1690755 | 3,06E-05 | 0,0002702 | TRIM63 |
| ENSGALG00000007617 | 161,13247 | 1,044420437 | 0,3032396 | 3,4442081 | 0,0005727 | 0,0033568 | ANO1 |
| ENSGALG00000006854 | 35,896481 | 1,044373279 | 0,3538993 | 2,9510467 | 0,003167 | 0,0137583 | NEURL2 |

| | | | | | | | |
|---|---|---|---|---|---|---|---|
| ENSGALG00000007837 | 153,44217 | 1,044277791 | 0,317273 | 3,2914167 | 0,0009968 | 0,0053161 | KAZALD1 |
| ENSGALG00000001252 | 75,259479 | 1,044188086 | 0,3423423 | 3,0501291 | 0,0022874 | 0,0105321 | CREB3L3 |
| ENSGALG00000009588 | 47,706596 | 1,044149084 | 0,3525375 | 2,9618103 | 0,0030584 | 0,0133846 | |
| ENSGALG00000016020 | 82,686873 | 1,041815502 | 0,331022 | 3,1472698 | 0,001648 | 0,0080365 | CLIC6 |
| ENSGALG00000016608 | 1071,038 | 1,041630113 | 0,1614881 | 6,4501978 | 1,12E-10 | 3,01E-09 | ADCY3 |
| ENSGALG00000014258 | 1564,4528 | 1,040436704 | 0,1704968 | 6,1023832 | 1,04E-09 | 2,47E-08 | LIMCH1 |
| ENSGALG00000027096 | 1851,9247 | 1,040113834 | 0,1377861 | 7,5487579 | 4,39E-14 | 1,90E-12 | LGALSL |
| ENSGALG00000024369 | 197,40751 | 1,038738473 | 0,2651565 | 3,917454 | 8,95E-05 | 0,0006826 | |
| ENSGALG00000028234 | 617,7355 | 1,03829388 | 0,1910987 | 5,4332854 | 5,53E-08 | 9,37E-07 | SYNDIG1 |
| ENSGALG00000007219 | 1265,6281 | 1,038287687 | 0,1548697 | 6,7042673 | 2,02E-11 | 6,12E-10 | FGF12 |
| ENSGALG00000007253 | 1268,2995 | 1,036667095 | 0,1580585 | 6,5587573 | 5,43E-11 | 1,54E-09 | IL1RAP |
| ENSGALG00000005460 | 763,14713 | 1,036332157 | 0,1841326 | 5,6281837 | 1,82E-08 | 3,43E-07 | SLCO4A1 |
| ENSGALG00000005951 | 50,333342 | 1,035951706 | 0,3500228 | 2,9596694 | 0,0030797 | 0,0134575 | VPS29L |
| ENSGALG00000026714 | 2323,4921 | 1,035206481 | 0,1567075 | 6,6059781 | 3,95E-11 | 1,13E-09 | SHISA2 |
| ENSGALG00000024122 | 36,985836 | 1,03453695 | 0,3344358 | 3,0933796 | 0,0019789 | 0,0093558 | TNFRSF13B |
| ENSGALG00000023779 | 766,12884 | 1,033222085 | 0,198913 | 5,1943408 | 2,05E-07 | 3,11E-06 | TMEM82 |
| ENSGALG00000028318 | 1679,8373 | 1,033139386 | 0,1849882 | 5,584894 | 2,34E-08 | 4,28E-07 | CDKN1A |
| ENSGALG00000005114 | 245,74674 | 1,032666035 | 0,2545785 | 4,0563754 | 4,98E-05 | 0,0004108 | CES1 |
| ENSGALG00000028628 | 176,65893 | 1,032210406 | 0,27931 | 3,6955723 | 0,0002194 | 0,0014809 | LAPTM4B |
| ENSGALG00000011875 | 520,49686 | 1,032103241 | 0,2093207 | 4,9307268 | 8,19E-07 | 1,08E-05 | TRPC3 |
| ENSGALG00000001304 | 107,91968 | 1,031984172 | 0,3164648 | 3,2609758 | 0,0011103 | 0,0058298 | TMEM52 |
| ENSGALG00000002459 | 72,761835 | 1,030692731 | 0,3400622 | 3,0308944 | 0,0024383 | 0,0110991 | |
| ENSGALG00000004860 | 237,1229 | 1,03038994 | 0,2518626 | 4,0910789 | 4,29E-05 | 0,00036 | RASD1 |
| ENSGALG00000010566 | 1230,6364 | 1,029946325 | 0,1643356 | 6,2673336 | 3,67E-10 | 9,18E-09 | NPNT |
| ENSGALG00000012791 | 100,57094 | 1,028853172 | 0,3257471 | 3,1584418 | 0,0015861 | 0,0077906 | TBXAS1 |
| ENSGALG00000024024 | 333,88236 | 1,028182094 | 0,2283532 | 4,5025956 | 6,71E-06 | 7,16E-05 | TGFA |
| ENSGALG00000009739 | 1255,1864 | 1,027863615 | 0,1833266 | 5,6067343 | 2,06E-08 | 3,83E-07 | AMIGO2 |
| ENSGALG00000010078 | 138,04602 | 1,027109659 | 0,2951454 | 3,4800125 | 0,0005014 | 0,0030072 | GLP1R |
| ENSGALG00000003349 | 284,81496 | 1,025286613 | 0,2360846 | 4,3428788 | 1,41E-05 | 0,0001392 | PLEKHA2 |
| ENSGALG00000009722 | 2700,6877 | 1,024969081 | 0,1274848 | 8,0399312 | 8,99E-16 | 4,78E-14 | DYNC1I1 |
| ENSGALG00000013200 | 141,95986 | 1,024952745 | 0,3004034 | 3,4119214 | 0,0006451 | 0,0037057 | GABBR2 |
| ENSGALG00000029169 | 143,48628 | 1,024627176 | 0,2979791 | 3,4385878 | 0,0005848 | 0,003414 | HHEX |
| ENSGALG00000026420 | 49,217095 | 1,024145559 | 0,3514259 | 2,9142577 | 0,0035654 | 0,0152027 | CSF3 |
| ENSGALG00000013934 | 77,966051 | 1,022547067 | 0,3347704 | 3,0544729 | 0,0022546 | 0,0103983 | FAM196A |
| ENSGALG00000002370 | 25,819164 | 1,022537057 | 0,3456971 | 2,957899 | 0,0030974 | 0,0135165 | SH2D4B |
| ENSGALG00000028153 | 28,525881 | 1,022401628 | 0,3430351 | 2,9804582 | 0,0028782 | 0,012727 | FOXE1 |
| ENSGALG00000012470 | 43,321678 | 1,020606107 | 0,3536746 | 2,8857204 | 0,0039052 | 0,0163839 | LYPD6 |
| ENSGALG00000026070 | 69,472965 | 1,019730322 | 0,3407891 | 2,9922619 | 0,0027692 | 0,0123558 | |
| ENSGALG00000025906 | 86,105418 | 1,019071961 | 0,3299575 | 3,088495 | 0,0020117 | 0,0094905 | |
| ENSGALG00000012564 | 122,7387 | 1,018929434 | 0,3020158 | 3,3737619 | 0,0007415 | 0,0041682 | |
| ENSGALG00000009686 | 819,70472 | 1,017336386 | 0,2030924 | 5,0092304 | 5,46E-07 | 7,54E-06 | PPP1R9A |
| ENSGALG00000001698 | 101,57773 | 1,017252154 | 0,3196233 | 3,1826597 | 0,0014593 | 0,0072777 | GABRA1 |
| ENSGALG00000002991 | 1547,8463 | 1,016425478 | 0,1518876 | 6,6919594 | 2,20E-11 | 6,60E-10 | ARID5B |
| ENSGALG00000001191 | 726,03044 | 1,014738695 | 0,1760749 | 5,7631095 | 8,26E-09 | 1,66E-07 | MEOX1 |
| ENSGALG00000009066 | 83,156898 | 1,013658719 | 0,3282182 | 3,0883687 | 0,0020126 | 0,0094913 | KCND2 |
| ENSGALG00000003571 | 35,860323 | 1,013482972 | 0,3515721 | 2,882717 | 0,0039426 | 0,0164962 | KIF12 |
| ENSGALG00000024381 | 18,525889 | 1,013267738 | 0,3321669 | 3,050478 | 0,0022848 | 0,0105234 | LARP6L |
| ENSGALG00000016223 | 759,19748 | 1,013131958 | 0,1825727 | 5,5491961 | 2,87E-08 | 5,19E-07 | MAOB |
| ENSGALG00000012307 | 29,303894 | 1,012697653 | 0,3486352 | 2,904749 | 0,0036755 | 0,015599 | GALR3 |
| ENSGALG00000010203 | 639,91361 | 1,012100046 | 0,1964193 | 5,1527526 | 2,57E-07 | 3,79E-06 | |
| ENSGALG00000017362 | 22,179359 | 1,011846131 | 0,3421939 | 2,9569383 | 0,0031071 | 0,0135504 | |
| ENSGALG00000008874 | 19,446543 | 1,010524458 | 0,323522 | 3,1235106 | 0,0017871 | 0,0085793 | SLC13A1 |
| ENSGALG00000022718 | 221,71371 | 1,010197798 | 0,2618613 | 3,8577596 | 0,0001144 | 0,0008403 | FUT4 |
| ENSGALG00000014455 | 15,37556 | 1,009369671 | 0,3151604 | 3,2027176 | 0,0013614 | 0,0068955 | LPAR5 |
| ENSGALG00000008080 | 251,32912 | 1,008867324 | 0,2511963 | 4,01625 | 5,91E-05 | 0,0004755 | C1QTNF4 |
| ENSGALG00000021655 | 77,574154 | 1,008428495 | 0,3474014 | 2,902776 | 0,0036987 | 0,015683 | BCL2L15 |
| ENSGALG00000001951 | 315,95633 | 1,007959576 | 0,2412905 | 4,1773687 | 2,95E-05 | 0,0002612 | FAM102B |
| ENSGALG00000025876 | 51,163718 | 1,007886002 | 0,3524372 | 2,8597604 | 0,0042396 | 0,0175022 | THPO |
| ENSGALG00000027712 | 289,19222 | 1,006836187 | 0,246269 | 4,0883601 | 4,34E-05 | 0,0003633 | |
| ENSGALG00000005094 | 32,433793 | 1,005780474 | 0,350919 | 2,8661325 | 0,0041552 | 0,0172007 | GRAP |
| ENSGALG00000028186 | 897,73377 | 1,005437673 | 0,1860189 | 5,4050306 | 6,48E-08 | 1,09E-06 | RERG |
| ENSGALG00000003816 | 91,378339 | 1,005135037 | 0,3271198 | 3,0726821 | 0,0021214 | 0,009926 | NT5C1A |
| ENSGALG00000004653 | 66,385404 | 1,003752617 | 0,3500053 | 2,8678215 | 0,0041331 | 0,0171352 | CDH23 |
| ENSGALG00000007713 | 153,5491 | 1,002069085 | 0,297438 | 3,3690019 | 0,0007544 | 0,0042333 | CD300A |

| | | | | | | | |
|---|---|---|---|---|---|---|---|
| ENSGALG00000026584 | 85,189769 | 1,001767633 | 0,3522109 | 2,844227 | 0,0044519 | 0,0181912 | |
| ENSGALG00000016251 | 6835,3703 | 1,000647408 | 0,1546994 | 6,4683351 | 9,91E-11 | 2,69E-09 | TSPAN7 |
| ENSGALG00000028962 | 99,714152 | 1,000589641 | 0,3302214 | 3,030057 | 0,0024451 | 0,0111225 | |
| ENSGALG00000023953 | 637,17536 | 0,998916427 | 0,1866162 | 5,3527844 | 8,66E-08 | 1,41E-06 | |
| ENSGALG00000005887 | 163,6325 | 0,998359123 | 0,3075104 | 3,2465868 | 0,001168 | 0,0060741 | PXDNL |
| ENSGALG00000009511 | 29,357161 | 0,996402413 | 0,3503011 | 2,8444172 | 0,0044493 | 0,0181858 | SI |
| ENSGALG00000012480 | 541,53262 | 0,99597856 | 0,2064035 | 4,8253958 | 1,40E-06 | 1,76E-05 | NMI |
| ENSGALG00000014681 | 398,45148 | 0,995892967 | 0,2311425 | 4,3085667 | 1,64E-05 | 0,0001581 | PCSK1 |
| ENSGALG00000000172 | 20,017715 | 0,995836198 | 0,3363958 | 2,9603109 | 0,0030733 | 0,013437 | MYOG |
| ENSGALG00000017316 | 218,98095 | 0,995598244 | 0,3012626 | 3,3047519 | 0,0009506 | 0,0051298 | UCP3 |
| ENSGALG00000007045 | 32,468926 | 0,994891941 | 0,3526069 | 2,8215327 | 0,0047795 | 0,0193208 | GPR123 |
| ENSGALG00000005467 | 75,167039 | 0,994871662 | 0,3408309 | 2,9189596 | 0,003512 | 0,0150035 | HSD17B2 |
| ENSGALG00000001453 | 365,61787 | 0,994068241 | 0,2454962 | 4,0492198 | 5,14E-05 | 0,0004207 | FAM110D |
| ENSGALG00000002437 | 722,53208 | 0,993124174 | 0,1885536 | 5,2670664 | 1,39E-07 | 2,17E-06 | FAM69B |
| ENSGALG00000012613 | 121,97261 | 0,992227725 | 0,3163097 | 3,1368866 | 0,0017075 | 0,0082733 | FBP1 |
| ENSGALG00000015094 | 84,375027 | 0,9904996 | 0,3373138 | 2,9364333 | 0,0033201 | 0,0142958 | ZBTB20 |
| ENSGALG00000028682 | 130,82591 | 0,990483134 | 0,3063312 | 3,2333734 | 0,0012234 | 0,006314 | |
| ENSGALG00000013183 | 389,46671 | 0,988585254 | 0,2164446 | 4,5673816 | 4,94E-06 | 5,48E-05 | TERT |
| ENSGALG00000002766 | 25,140518 | 0,98837641 | 0,3444585 | 2,8693625 | 0,004113 | 0,0170883 | MLKL |
| ENSGALG00000009638 | 342,69847 | 0,988233263 | 0,2260661 | 4,3714351 | 1,23E-05 | 0,0001244 | ANO6 |
| ENSGALG00000016224 | 196,09798 | 0,988198095 | 0,2772614 | 3,5641386 | 0,0003651 | 0,0022968 | MAOA |
| ENSGALG00000009052 | 3269,1548 | 0,987910707 | 0,1624129 | 6,0827127 | 1,18E-09 | 2,78E-08 | LBH |
| ENSGALG00000026042 | 849,9138 | 0,987626868 | 0,2371785 | 4,1640662 | 3,13E-05 | 0,0002751 | SHISA3 |
| ENSGALG00000012536 | 763,86305 | 0,987580114 | 0,2116261 | 4,6666263 | 3,06E-06 | 3,54E-05 | GALNT13 |
| ENSGALG00000016599 | 16,001403 | 0,987073109 | 0,3131543 | 3,1520339 | 0,0016214 | 0,0079321 | MFAP4 |
| ENSGALG00000008638 | 22,301546 | 0,987044947 | 0,3280366 | 3,008948 | 0,0026215 | 0,011768 | |
| ENSGALG00000009463 | 2838,7047 | 0,986691121 | 0,1403118 | 7,0321343 | 2,03E-12 | 7,16E-11 | DCAF5 |
| ENSGALG00000006812 | 66,925337 | 0,985329432 | 0,3466414 | 2,8425032 | 0,0044761 | 0,0182625 | TTC36 |
| ENSGALG00000001764 | 93,207252 | 0,985279854 | 0,3248167 | 3,0333416 | 0,0024186 | 0,0110206 | HCN4 |
| ENSGALG00000005030 | 1142,7214 | 0,984706358 | 0,1778942 | 5,5353473 | 3,11E-08 | 5,55E-07 | DOCK10 |
| ENSGALG00000009488 | 49,450774 | 0,98346472 | 0,3539444 | 2,7785852 | 0,0054596 | 0,0216083 | LRRN3 |
| ENSGALG00000027429 | 108,06142 | 0,983232181 | 0,3123358 | 3,1479968 | 0,0016439 | 0,0080223 | HSD17B1 |
| ENSGALG00000011203 | 97,875883 | 0,983177315 | 0,3203781 | 3,0688031 | 0,0021492 | 0,0100141 | HPSE |
| ENSGALG00000005419 | 103,38309 | 0,983150481 | 0,3221858 | 3,0515019 | 0,002277 | 0,0104911 | ERC2 |
| ENSGALG00000015172 | 24,235396 | 0,982809248 | 0,3441667 | 2,85562 | 0,0042953 | 0,0176945 | |
| ENSGALG00000023847 | 205,80734 | 0,981888346 | 0,2682741 | 3,6600198 | 0,0002522 | 0,0016692 | |
| ENSGALG00000012552 | 632,63312 | 0,981819434 | 0,1820243 | 5,3938935 | 6,89E-08 | 1,15E-06 | TOM1 |
| ENSGALG00000027739 | 223,97962 | 0,981670308 | 0,2808826 | 3,4949492 | 0,0004742 | 0,0028704 | CPNE9 |
| ENSGALG00000000394 | 39,052306 | 0,981204777 | 0,3535388 | 2,7753805 | 0,0055137 | 0,0217718 | SLC6A17 |
| ENSGALG00000005883 | 37,607236 | 0,980233834 | 0,3534383 | 2,7734226 | 0,005547 | 0,0218779 | BRICD5 |
| ENSGALG00000012715 | 380,22403 | 0,978917656 | 0,2453996 | 3,9890758 | 6,63E-05 | 0,0005244 | CAP2 |
| ENSGALG00000012709 | 399,32692 | 0,978132572 | 0,259146 | 3,774446 | 0,0001604 | 0,0011301 | ATXN1 |
| ENSGALG00000027489 | 17,903941 | 0,978014836 | 0,3301784 | 2,9620804 | 0,0030557 | 0,0133772 | |
| ENSGALG00000026754 | 127,90019 | 0,977513049 | 0,3209323 | 3,0458545 | 0,0023202 | 0,0106577 | |
| ENSGALG00000028301 | 585,85613 | 0,976931086 | 0,2053037 | 4,7584672 | 1,95E-06 | 2,36E-05 | TRIM47 |
| ENSGALG00000006882 | 2766,7066 | 0,976472658 | 0,130813 | 7,4646479 | 8,35E-14 | 3,47E-12 | ST8SIA2 |
| ENSGALG00000026027 | 52,610545 | 0,976263156 | 0,3534975 | 2,7617256 | 0,0057497 | 0,0224496 | CLIC3 |
| ENSGALG00000024059 | 15,053333 | 0,976199065 | 0,3125125 | 3,1237123 | 0,0017858 | 0,0085765 | PGA |
| ENSGALG00000010797 | 412,00234 | 0,975073157 | 0,2100324 | 4,6424888 | 3,44E-06 | 3,95E-05 | SOSTDC1 |
| ENSGALG00000000902 | 273,8179 | 0,974915117 | 0,2442269 | 3,9918423 | 6,56E-05 | 0,0005187 | KDF1 |
| ENSGALG00000004591 | 574,2787 | 0,974905454 | 0,1969693 | 4,949529 | 7,44E-07 | 9,89E-06 | SERPINC1 |
| ENSGALG00000016715 | 46,103475 | 0,974213716 | 0,3521064 | 2,7668161 | 0,0056607 | 0,0221976 | MEP1A |
| ENSGALG00000016944 | 4957,4542 | 0,973864805 | 0,3289279 | 2,9607247 | 0,0030692 | 0,0134233 | PCDH8 |
| ENSGALG00000018875 | 14,740412 | 0,973272591 | 0,3123576 | 3,1158922 | 0,0018339 | 0,0087609 | ACSM4 |
| ENSGALG00000004721 | 51,579363 | 0,972599162 | 0,3523855 | 2,7600432 | 0,0057794 | 0,0225397 | KCNJ12 |
| ENSGALG00000003886 | 1023,8773 | 0,971801068 | 0,1773944 | 5,4781962 | 4,30E-08 | 7,44E-07 | SGCD |
| ENSGALG00000003069 | 21,65475 | 0,971652009 | 0,3390802 | 2,8655523 | 0,0041628 | 0,017227 | TNFSF4 |
| ENSGALG00000023869 | 23,260449 | 0,971412526 | 0,3437175 | 2,8261945 | 0,0047105 | 0,0190805 | |
| ENSGALG00000019231 | 26,295329 | 0,970164482 | 0,3464496 | 2,8003049 | 0,0051054 | 0,0203963 | CD200R1 |
| ENSGALG00000023439 | 59,847743 | 0,969553596 | 0,3512997 | 2,7599043 | 0,0057818 | 0,0225428 | |
| ENSGALG00000019325 | 54,610886 | 0,96951089 | 0,3507875 | 2,7638126 | 0,005713 | 0,0223258 | |
| ENSGALG00000005844 | 1501,3882 | 0,969445519 | 0,1528693 | 6,3416622 | 2,27E-10 | 5,87E-09 | CASKIN1 |
| ENSGALG00000016820 | 787,30983 | 0,968970008 | 0,1783249 | 5,4337332 | 5,52E-08 | 9,36E-07 | GAS6 |
| ENSGALG00000027287 | 736,98606 | 0,968636988 | 0,1893148 | 5,11654 | 3,11E-07 | 4,50E-06 | HK3 |

| | | | | | | | |
|---|---|---|---|---|---|---|---|
| ENSGALG00000003170 | 1633,7315 | 0,967045053 | 0,1741681 | 5,552367 | 2,82E-08 | 5,10E-07 | DNAH10 |
| ENSGALG00000026870 | 2578,9703 | 0,966911139 | 0,1632907 | 5,9214111 | 3,19E-09 | 6,99E-08 | TMEM200C |
| ENSGALG00000027511 | 3914,707 | 0,966828969 | 0,1525889 | 6,3361698 | 2,36E-10 | 6,06E-09 | SP5 |
| ENSGALG00000021275 | 22,663889 | 0,966605206 | 0,3386389 | 2,8543832 | 0,004312 | 0,0177474 | LTC4S |
| ENSGALG00000004734 | 21,747155 | 0,966247375 | 0,3385892 | 2,8537457 | 0,0043207 | 0,0177582 | |
| ENSGALG00000015418 | 42,123895 | 0,966243505 | 0,353786 | 2,7311521 | 0,0063113 | 0,0242874 | EPHA6 |
| ENSGALG00000001209 | 151,27678 | 0,963839438 | 0,2885245 | 3,3405805 | 0,000836 | 0,0046009 | TP53INP2 |
| ENSGALG00000008932 | 24,822651 | 0,962818941 | 0,3469054 | 2,7754514 | 0,0055125 | 0,0217718 | ENTPD8 |
| ENSGALG00000012681 | 687,54966 | 0,96179459 | 0,1933738 | 4,9737578 | 6,57E-07 | 8,84E-06 | MBOAT1 |
| ENSGALG00000026914 | 303,14643 | 0,960831377 | 0,2341299 | 4,1038388 | 4,06E-05 | 0,0003453 | DOK3 |
| ENSGALG00000016597 | 546,5809 | 0,96044418 | 0,2115615 | 4,5397871 | 5,63E-06 | 6,12E-05 | STMN4 |
| ENSGALG00000006911 | 92,463176 | 0,960163411 | 0,3331134 | 2,8823917 | 0,0039467 | 0,0165082 | TMC5 |
| ENSGALG00000028041 | 172,71276 | 0,96010795 | 0,292817 | 3,2788668 | 0,0010422 | 0,0055242 | DKK2 |
| ENSGALG00000008317 | 155,9685 | 0,958791686 | 0,2982524 | 3,2146994 | 0,0013058 | 0,0066662 | SDSL |
| ENSGALG00000006192 | 588,22456 | 0,958606154 | 0,1986953 | 4,824504 | 1,40E-06 | 1,77E-05 | USH1C |
| ENSGALG00000012285 | 88,550664 | 0,958415036 | 0,3255253 | 2,9442104 | 0,0032378 | 0,0140066 | BAIAP2L2 |
| ENSGALG00000017274 | 44,802442 | 0,957455073 | 0,3539461 | 2,7050872 | 0,0068286 | 0,0258936 | CD9 |
| ENSGALG00000009896 | 1725,9341 | 0,956710434 | 0,138947 | 6,8854343 | 5,76E-12 | 1,88E-10 | TRERF1 |
| ENSGALG00000008936 | 346,61065 | 0,956545148 | 0,2393518 | 3,9963977 | 6,43E-05 | 0,0005109 | ENTPD8L |
| ENSGALG00000012577 | 1716,8746 | 0,956309243 | 0,1378913 | 6,9352415 | 4,06E-12 | 1,35E-10 | FRMD3 |
| ENSGALG00000006375 | 24,972242 | 0,956253302 | 0,3445218 | 2,7755957 | 0,0055101 | 0,0217718 | TM4SF19 |
| ENSGALG00000012271 | 67,173186 | 0,955959348 | 0,3478063 | 2,7485396 | 0,0059861 | 0,0232197 | MTTP |
| ENSGALG00000023271 | 71,780111 | 0,955084764 | 0,3439036 | 2,7771873 | 0,0054832 | 0,0216826 | |
| ENSGALG00000016945 | 3627,6803 | 0,953551228 | 0,1270066 | 7,5078871 | 6,01E-14 | 2,53E-12 | LECT1 |
| ENSGALG00000003209 | 25,968303 | 0,953438931 | 0,3318207 | 2,8733556 | 0,0040614 | 0,0169048 | NODAL |
| ENSGALG00000014843 | 607,09399 | 0,953307805 | 0,2111335 | 4,5151905 | 6,33E-06 | 6,80E-05 | TPD52L1 |
| ENSGALG00000029150 | 2513,115 | 0,953288378 | 0,1486082 | 6,414778 | 1,41E-10 | 3,76E-09 | AK1 |
| ENSGALG00000005118 | 32,326575 | 0,952825585 | 0,3518222 | 2,7082585 | 0,0067637 | 0,0257069 | |
| ENSGALG00000000295 | 158,36113 | 0,951838119 | 0,2874721 | 3,3110626 | 0,0009294 | 0,0050415 | IGFN1 |
| ENSGALG00000002008 | 770,02151 | 0,951550674 | 0,2636914 | 3,6085762 | 0,0003079 | 0,0019866 | RNF157 |
| ENSGALG00000009216 | 17,250683 | 0,951287914 | 0,3202238 | 2,970697 | 0,0029712 | 0,0130664 | |
| ENSGALG00000005472 | 26,534994 | 0,951255745 | 0,3483401 | 2,7308243 | 0,0063176 | 0,024288 | NAT |
| ENSGALG00000017926 | 117,34516 | 0,950771836 | 0,3049521 | 3,1177741 | 0,0018222 | 0,0087174 | |
| ENSGALG00000009123 | 138,33508 | 0,94996493 | 0,2943779 | 3,227025 | 0,0012508 | 0,006434 | |
| ENSGALG00000017068 | 94,861308 | 0,947287623 | 0,3314816 | 2,8577385 | 0,0042667 | 0,0175875 | KL |
| ENSGALG00000021696 | 69,405082 | 0,947251279 | 0,3378068 | 2,8041212 | 0,0050454 | 0,0202098 | TRIM50 |
| ENSGALG00000026793 | 20,910942 | 0,946537074 | 0,3217404 | 2,9419277 | 0,0032618 | 0,0140891 | CYP2C8 |
| ENSGALG00000011046 | 918,66681 | 0,946061534 | 0,2245408 | 4,2133163 | 2,52E-05 | 0,0002287 | SNX10 |
| ENSGALG00000004792 | 282,6657 | 0,945718297 | 0,2609466 | 3,6241839 | 0,0002899 | 0,0018856 | PLCB2 |
| ENSGALG00000001091 | 62,437953 | 0,945610586 | 0,3444618 | 2,7451826 | 0,0060477 | 0,0234186 | C1orf116 |
| ENSGALG00000005857 | 90,998254 | 0,944585535 | 0,322613 | 2,9279211 | 0,0034124 | 0,0146283 | ALOX5 |
| ENSGALG00000006988 | 545,9632 | 0,944077449 | 0,2380928 | 3,9651665 | 7,33E-05 | 0,0005736 | RNF213 |
| ENSGALG00000009002 | 180,8252 | 0,943061124 | 0,2745333 | 3,4351427 | 0,0005922 | 0,0034533 | C7ORF58 |
| ENSGALG00000003370 | 6860,8975 | 0,941055859 | 0,1158875 | 8,120424 | 4,65E-16 | 2,56E-14 | KIAA1191 |
| ENSGALG00000015183 | 541,00251 | 0,940741895 | 0,2469295 | 3,8097593 | 0,0001391 | 0,0009994 | |
| ENSGALG00000010977 | 629,76264 | 0,94045823 | 0,1853813 | 5,0731011 | 3,91E-07 | 5,55E-06 | FAM221A |
| ENSGALG00000017169 | 2527,4044 | 0,94013642 | 0,1550564 | 6,0631902 | 1,33E-09 | 3,11E-08 | ELMOD1 |
| ENSGALG00000006351 | 260,93159 | 0,940035694 | 0,256606 | 3,6633421 | 0,0002489 | 0,0016526 | FAS |
| ENSGALG00000016699 | 24,25844 | 0,939513809 | 0,3453851 | 2,7201919 | 0,0065244 | 0,0249662 | OPN5L2 |
| ENSGALG00000007415 | 599,16556 | 0,938743099 | 0,2370594 | 3,9599488 | 7,50E-05 | 0,0005829 | SH3RF2 |
| ENSGALG00000002678 | 150,0779 | 0,938682309 | 0,293615 | 3,1969839 | 0,0013887 | 0,007008 | CSPG4 |
| ENSGALG00000003832 | 1422,7636 | 0,938218604 | 0,1475404 | 6,3590641 | 2,03E-10 | 5,31E-09 | TEX264 |
| ENSGALG00000009725 | 763,67798 | 0,938023358 | 0,224257 | 4,182805 | 2,88E-05 | 0,0002563 | SCG5 |
| ENSGALG00000015967 | 204,24999 | 0,936813576 | 0,2634911 | 3,5553898 | 0,0003774 | 0,0023603 | LRRTM1 |
| ENSGALG00000003435 | 45,697878 | 0,936462652 | 0,3537379 | 2,6473351 | 0,0081129 | 0,0297327 | PGC |
| ENSGALG00000002473 | 480,3177 | 0,93617222 | 0,2026166 | 4,6204127 | 3,83E-06 | 4,35E-05 | GPR157 |
| ENSGALG00000015926 | 1110,0554 | 0,934566237 | 0,1598776 | 5,84551 | 5,05E-09 | 1,06E-07 | RUNX1T1 |
| ENSGALG00000019312 | 16,25864 | 0,934023396 | 0,3177972 | 2,939055 | 0,0032921 | 0,0141844 | |
| ENSGALG00000001398 | 801,87151 | 0,933503925 | 0,1756569 | 5,3143584 | 1,07E-07 | 1,71E-06 | |
| ENSGALG00000010337 | 96,826556 | 0,93241733 | 0,319691 | 2,9166205 | 0,0035385 | 0,0151022 | GPR149 |
| ENSGALG00000028292 | 453,15086 | 0,931661788 | 0,2502644 | 3,7227094 | 0,0001971 | 0,0013471 | FAM19A4 |
| ENSGALG00000008735 | 158,65028 | 0,931477748 | 0,2951381 | 3,1560747 | 0,0015991 | 0,0078456 | BFSP1 |
| ENSGALG00000010139 | 381,67431 | 0,931031468 | 0,2175137 | 4,2803349 | 1,87E-05 | 0,0001763 | SSTR1 |
| ENSGALG00000002843 | 586,86661 | 0,930974563 | 0,1932115 | 4,8184228 | 1,45E-06 | 1,82E-05 | RILP |

| | | | | | | | |
|---|---|---|---|---|---|---|---|
| ENSGALG00000001694 | 115,671 | 0,929649369 | 0,3250641 | 2,8598955 | 0,0042378 | 0,0175001 | FER1L4 |
| ENSGALG00000001287 | 117,66472 | 0,929543301 | 0,3091097 | 3,0071632 | 0,002637 | 0,0118295 | ABI3 |
| ENSGALG00000023626 | 1834,269 | 0,929476271 | 0,1654852 | 5,6166736 | 1,95E-08 | 3,65E-07 | NTN1 |
| ENSGALG00000002083 | 14,444586 | 0,928533788 | 0,3085021 | 3,0098139 | 0,0026141 | 0,0117383 | PLIN3 |
| ENSGALG00000027053 | 71,270094 | 0,927920334 | 0,3515108 | 2,6398059 | 0,0082954 | 0,0302791 | C18orf42 |
| ENSGALG00000028091 | 182,30387 | 0,927849672 | 0,2899177 | 3,2003896 | 0,0013724 | 0,0069433 | MYF5 |
| ENSGALG00000025801 | 28,789593 | 0,927018359 | 0,349612 | 2,6515632 | 0,008012 | 0,0294645 | |
| ENSGALG00000000866 | 124,56244 | 0,926965653 | 0,3251101 | 2,8512364 | 0,004355 | 0,0178646 | SELENBP1 |
| ENSGALG00000006388 | 73,461217 | 0,926517942 | 0,3489684 | 2,6550197 | 0,0079304 | 0,029237 | IL16 |
| ENSGALG00000000919 | 14,595204 | 0,925422799 | 0,3081645 | 3,0030152 | 0,0026732 | 0,0119683 | PIGR |
| ENSGALG00000023517 | 259,64579 | 0,925400404 | 0,2568731 | 3,602558 | 0,0003151 | 0,0020246 | AGPAT2 |
| ENSGALG00000005626 | 609,64517 | 0,92496767 | 0,2159097 | 4,2840488 | 1,84E-05 | 0,0001735 | TPST2 |
| ENSGALG00000001035 | 102,60165 | 0,924878686 | 0,3465453 | 2,6688536 | 0,0076111 | 0,028305 | NR1D1 |
| ENSGALG00000020233 | 23,832939 | 0,924086016 | 0,3386279 | 2,7289127 | 0,0063544 | 0,0243978 | |
| ENSGALG00000000311 | 558,40847 | 0,92295118 | 0,2153587 | 4,2856468 | 1,82E-05 | 0,0001725 | DOCK5 |
| ENSGALG00000014988 | 88,836245 | 0,920904288 | 0,3383615 | 2,7216578 | 0,0064955 | 0,0248884 | POPDC2 |
| ENSGALG00000015207 | 1291,1294 | 0,92039826 | 0,1472006 | 6,2526786 | 4,03E-10 | 1,00E-08 | F5 |
| ENSGALG00000010417 | 18,919028 | 0,920255863 | 0,3320234 | 2,77166 | 0,0055771 | 0,0219585 | VDHAP |
| ENSGALG00000010288 | 20,644022 | 0,918819724 | 0,3351049 | 2,7418866 | 0,0061087 | 0,0236079 | |
| ENSGALG00000023936 | 805,04939 | 0,918539777 | 0,2012641 | 4,5638532 | 5,02E-06 | 5,56E-05 | |
| ENSGALG00000009509 | 42,581938 | 0,91836029 | 0,3539579 | 2,5945464 | 0,0094716 | 0,0336435 | CALCR |
| ENSGALG00000011709 | 513,36272 | 0,918333355 | 0,2317844 | 3,9620158 | 7,43E-05 | 0,0005792 | TMEM108 |
| ENSGALG00000027622 | 398,87687 | 0,916634803 | 0,2658647 | 3,4477494 | 0,0005653 | 0,0033245 | ADAM11 |
| ENSGALG00000015795 | 539,93954 | 0,91628777 | 0,2153216 | 4,2554383 | 2,09E-05 | 0,0001946 | ADAMTS5 |
| ENSGALG00000005120 | 2531,22 | 0,916224882 | 0,1549653 | 5,9124511 | 3,37E-09 | 7,37E-08 | FBLN2 |
| ENSGALG00000010736 | 93,908737 | 0,914854903 | 0,3255588 | 2,8101064 | 0,0049525 | 0,0199374 | DIO1 |
| ENSGALG00000013071 | 1399,4771 | 0,914850391 | 0,17423 | 5,2508198 | 1,51E-07 | 2,35E-06 | CALD1 |
| ENSGALG00000010382 | 346,13588 | 0,914779385 | 0,2280121 | 4,0119779 | 6,02E-05 | 0,0004822 | LRRC73 |
| ENSGALG00000004309 | 938,33548 | 0,913060877 | 0,1855348 | 4,9212369 | 8,60E-07 | 1,13E-05 | GDAP1L1 |
| ENSGALG00000028622 | 281,67527 | 0,912974481 | 0,2559896 | 3,566452 | 0,0003618 | 0,0022808 | PEAR1 |
| ENSGALG00000017214 | 131,47371 | 0,912890352 | 0,3103783 | 2,9412185 | 0,0032692 | 0,0141048 | HEPHL1 |
| ENSGALG00000016013 | 42,116835 | 0,912662466 | 0,3534877 | 2,5818789 | 0,0098264 | 0,0347137 | HTR7L |
| ENSGALG00000026687 | 69,587609 | 0,912634606 | 0,347276 | 2,6279803 | 0,0085893 | 0,0311435 | PLCD4 |
| ENSGALG00000005331 | 677,48087 | 0,912514017 | 0,1798939 | 5,0725131 | 3,93E-07 | 5,56E-06 | GJC2 |
| ENSGALG00000003529 | 116,76012 | 0,912498344 | 0,3196723 | 2,8544806 | 0,0043107 | 0,0177474 | SYNC |
| ENSGALG00000027462 | 1468,6722 | 0,912376278 | 0,1432394 | 6,369589 | 1,90E-10 | 4,98E-09 | LEPREL4 |
| ENSGALG00000015789 | 559,92994 | 0,912066241 | 0,2123869 | 4,2943618 | 1,75E-05 | 0,000167 | CNR1 |
| ENSGALG00000026563 | 13,690632 | 0,912028647 | 0,3072132 | 2,9687161 | 0,0029905 | 0,0131255 | HAPLN4 |
| ENSGALG00000010551 | 919,31695 | 0,911984879 | 0,1832286 | 4,9773072 | 6,45E-07 | 8,70E-06 | CDC42EP3 |
| ENSGALG00000012298 | 550,04797 | 0,911513817 | 0,2224948 | 4,0967868 | 4,19E-05 | 0,0003536 | SLC39A8 |
| ENSGALG00000003588 | 1398,6038 | 0,911216657 | 0,1584852 | 5,7495378 | 8,95E-09 | 1,78E-07 | |
| ENSGALG00000005814 | 212,00295 | 0,910795359 | 0,271855 | 3,3502982 | 0,0008072 | 0,0044752 | |
| ENSGALG00000002581 | 512,90179 | 0,910753526 | 0,1951976 | 4,6658018 | 3,07E-06 | 3,56E-05 | HTATIP2 |
| ENSGALG00000004676 | 1678,5831 | 0,910530695 | 0,1433097 | 6,3535879 | 2,10E-10 | 5,48E-09 | BTG3 |
| ENSGALG00000008727 | 92,320134 | 0,910292988 | 0,3286188 | 2,7700573 | 0,0056046 | 0,022035 | |
| ENSGALG00000023195 | 260,80986 | 0,909179266 | 0,2471713 | 3,6783362 | 0,0002348 | 0,0015683 | DLX6 |
| ENSGALG00000027013 | 1552,1008 | 0,908413942 | 0,3505249 | 2,5915816 | 0,0095536 | 0,0338377 | HOX1 |
| ENSGALG00000015725 | 901,98347 | 0,907979483 | 0,1701411 | 5,3366278 | 9,47E-08 | 1,53E-06 | FGFRL1 |
| ENSGALG00000007114 | 305,81852 | 0,907573399 | 0,2767971 | 3,2788398 | 0,0010423 | 0,0055242 | APOA1 |
| ENSGALG00000010635 | 38,448493 | 0,906970906 | 0,3515336 | 2,5800407 | 0,0098789 | 0,0348087 | |
| ENSGALG00000011213 | 25,560047 | 0,906880966 | 0,3462592 | 2,619081 | 0,0088167 | 0,0317985 | DEGS2 |
| ENSGALG00000019635 | 32,562895 | 0,90677452 | 0,3521651 | 2,5748562 | 0,0100282 | 0,0352345 | PP2D1 |
| ENSGALG00000006202 | 116,81237 | 0,906421176 | 0,31598 | 2,8686028 | 0,0041229 | 0,0171086 | SCNN1B |
| ENSGALG00000003018 | 309,19026 | 0,906093174 | 0,2488521 | 3,6410907 | 0,0002715 | 0,0017771 | SCARB1 |
| ENSGALG00000016076 | 120,93641 | 0,906052944 | 0,3046837 | 2,9737494 | 0,0029419 | 0,0129706 | DPYS |
| ENSGALG00000015495 | 24,536119 | 0,904690337 | 0,3465615 | 2,6104756 | 0,0090416 | 0,0324294 | POU1F1 |
| ENSGALG00000013886 | 841,1366 | 0,904236576 | 0,1804703 | 5,0104453 | 5,43E-07 | 7,51E-06 | LDLRAD4 |
| ENSGALG00000005795 | 32,624744 | 0,904049889 | 0,343537 | 2,6315941 | 0,0084985 | 0,0308718 | CYP2H1 |
| ENSGALG00000021442 | 473,2158 | 0,903637173 | 0,225822 | 4,0015457 | 6,29E-05 | 0,000501 | RLTPR |
| ENSGALG00000005014 | 159,55971 | 0,903243848 | 0,2894313 | 3,1207539 | 0,0018039 | 0,0086448 | AIPL1 |
| ENSGALG00000002963 | 189,40667 | 0,902926335 | 0,2699116 | 3,3452673 | 0,000822 | 0,0045354 | PID1 |
| ENSGALG00000000836 | 482,38155 | 0,902335572 | 0,2085245 | 4,3272398 | 1,51E-05 | 0,0001475 | RFX5 |
| ENSGALG00000011124 | 436,07123 | 0,902068067 | 0,2097494 | 4,3006934 | 1,70E-05 | 0,000163 | ARHGAP24 |
| ENSGALG00000010827 | 4357,4834 | 0,901311467 | 0,1146698 | 7,8600622 | 3,84E-15 | 1,84E-13 | ITPK1 |

| | | | | | | | |
|---|---|---|---|---|---|---|---|
| ENSGALG00000006756 | 144,39 | 0,900987054 | 0,2996417 | 3,0068817 | 0,0026394 | 0,0118366 | BPI |
| ENSGALG00000006740 | 150,08447 | 0,900492229 | 0,3014742 | 2,9869628 | 0,0028176 | 0,0125162 | |
| ENSGALG00000008461 | 387,65672 | 0,900341356 | 0,2481083 | 3,6288245 | 0,0002847 | 0,0018574 | HGF |
| ENSGALG00000016761 | 58,000294 | 0,90015721 | 0,3467952 | 2,5956447 | 0,0094414 | 0,0335537 | LYG2 |
| ENSGALG00000010119 | 1290,7566 | 0,900097678 | 0,1879287 | 4,78957 | 1,67E-06 | 2,06E-05 | CDKL2 |
| ENSGALG00000020905 | 199,73795 | 0,899806675 | 0,2823411 | 3,1869491 | 0,0014378 | 0,007205 | KSR2 |
| ENSGALG00000009737 | 97,351691 | 0,899099989 | 0,342334 | 2,6263823 | 0,0086298 | 0,0312485 | TAC1 |
| ENSGALG00000006793 | 399,96964 | 0,898163749 | 0,2122528 | 4,2315762 | 2,32E-05 | 0,0002129 | NTRK3 |
| ENSGALG00000008632 | 774,98804 | 0,897411708 | 0,1727746 | 5,1941169 | 2,06E-07 | 3,11E-06 | ACSS1 |
| ENSGALG00000001896 | 633,5555 | 0,897398544 | 0,1829807 | 4,9043349 | 9,37E-07 | 1,23E-05 | NTNG1 |
| ENSGALG00000006083 | 883,4583 | 0,896818662 | 0,1847643 | 4,8538537 | 1,21E-06 | 1,55E-05 | |
| ENSGALG00000023772 | 19,974194 | 0,896459603 | 0,3370772 | 2,6595087 | 0,0078255 | 0,02896 | HSPB7 |
| ENSGALG00000005853 | 55,709692 | 0,89635133 | 0,3538655 | 2,5330285 | 0,0113082 | 0,0387543 | HTR2C |
| ENSGALG00000007052 | 507,96535 | 0,895651499 | 0,203478 | 4,4017107 | 1,07E-05 | 0,0001099 | PXK |
| ENSGALG00000015977 | 788,48732 | 0,89563673 | 0,2070525 | 4,3256514 | 1,52E-05 | 0,0001481 | SMOX |
| ENSGALG00000005349 | 125,03741 | 0,895405672 | 0,3020632 | 2,9642993 | 0,0030337 | 0,0132897 | MSLN |
| ENSGALG00000005085 | 110,02771 | 0,895178522 | 0,3179722 | 2,8152725 | 0,0048736 | 0,0196662 | DRD4 |
| ENSGALG00000008262 | 171,78981 | 0,894529697 | 0,2811864 | 3,1812692 | 0,0014663 | 0,0072994 | RASGRF1 |
| ENSGALG00000000374 | 1105,8431 | 0,894239416 | 0,1877046 | 4,7640791 | 1,90E-06 | 2,30E-05 | RNPEP |
| ENSGALG00000017204 | 419,66256 | 0,891856915 | 0,2224702 | 4,0088824 | 6,10E-05 | 0,000488 | SESN3 |
| ENSGALG00000002854 | 440,26532 | 0,890941536 | 0,2420509 | 3,6808025 | 0,0002325 | 0,0015558 | PDZD2 |
| ENSGALG00000008599 | 20,776579 | 0,889603293 | 0,3339725 | 2,6637025 | 0,0077286 | 0,0286811 | |
| ENSGALG00000005203 | 773,68467 | 0,889220424 | 0,2133641 | 4,1676197 | 3,08E-05 | 0,0002718 | OLFM3 |
| ENSGALG00000027603 | 933,53527 | 0,889190298 | 0,1666499 | 5,335677 | 9,52E-08 | 1,54E-06 | ZHX3 |
| ENSGALG00000003932 | 494,66101 | 0,888085177 | 0,2291723 | 3,8751853 | 0,0001065 | 0,000794 | SDC4 |
| ENSGALG00000007388 | 193,87974 | 0,887520344 | 0,3042223 | 2,9173416 | 0,0035303 | 0,0150721 | SRRM4 |
| ENSGALG00000013265 | 26,768688 | 0,887507398 | 0,3491977 | 2,5415617 | 0,0110358 | 0,0380484 | GYS2 |
| ENSGALG00000001917 | 36,330226 | 0,88709327 | 0,3537861 | 2,5074281 | 0,0121613 | 0,0410137 | SSC4D |
| ENSGALG00000021666 | 75,893072 | 0,886771767 | 0,3536689 | 2,5073499 | 0,012164 | 0,0410137 | TMEM240 |
| ENSGALG00000010269 | 1110,1934 | 0,886301871 | 0,1651233 | 5,3675161 | 7,98E-08 | 1,31E-06 | KCNAB1 |
| ENSGALG00000004093 | 635,62125 | 0,885654514 | 0,2097536 | 4,2223558 | 2,42E-05 | 0,0002211 | TBX22 |
| ENSGALG00000009256 | 32,540437 | 0,884987978 | 0,3436981 | 2,5748992 | 0,0100269 | 0,0352345 | LEFTY2 |
| ENSGALG00000004729 | 91,741024 | 0,88490464 | 0,3219745 | 2,7483686 | 0,0059893 | 0,0232252 | SLC7A10 |
| ENSGALG00000008722 | 81,303501 | 0,884773994 | 0,3362693 | 2,6311471 | 0,0085097 | 0,030896 | MCOLN2 |
| ENSGALG00000015788 | 630,45937 | 0,884593035 | 0,2015792 | 4,3883151 | 1,14E-05 | 0,000116 | CHMP4C |
| ENSGALG00000004970 | 368,25658 | 0,884420332 | 0,2518256 | 3,5120356 | 0,0004447 | 0,0027174 | SVOP |
| ENSGALG00000026152 | 376,28373 | 0,884418189 | 0,2320545 | 3,8112524 | 0,0001383 | 0,0009939 | GBP |
| ENSGALG00000002252 | 31,22735 | 0,883701655 | 0,3473707 | 2,5439732 | 0,0109599 | 0,0378538 | |
| ENSGALG00000010862 | 27,986731 | 0,883307692 | 0,3482432 | 2,5364677 | 0,0111977 | 0,0384768 | FERD3L |
| ENSGALG00000004854 | 90,659457 | 0,882798305 | 0,3265019 | 2,7038079 | 0,006855 | 0,0259757 | |
| ENSGALG00000025733 | 660,61374 | 0,882587352 | 0,1955092 | 4,5143008 | 6,35E-06 | 6,83E-05 | |
| ENSGALG00000006008 | 1270,6534 | 0,882382705 | 0,1743852 | 5,0599632 | 4,19E-07 | 5,90E-06 | HOMER2 |
| ENSGALG00000027027 | 1969,0809 | 0,882215122 | 0,163127 | 5,4081483 | 6,37E-08 | 1,07E-06 | FBXL7 |
| ENSGALG00000011235 | 63,312192 | 0,882157118 | 0,3457444 | 2,5514721 | 0,0107269 | 0,0372093 | GALNT15 |
| ENSGALG00000003270 | 387,81088 | 0,881994372 | 0,2135989 | 4,1292087 | 3,64E-05 | 0,0003135 | DRD1 |
| ENSGALG00000020160 | 119,73819 | 0,881263086 | 0,3087394 | 2,8543913 | 0,0043119 | 0,0177474 | ADH6 |
| ENSGALG00000011234 | 31,35804 | 0,880790396 | 0,3529516 | 2,4954988 | 0,012578 | 0,0422005 | INHA |
| ENSGALG00000001519 | 27,987195 | 0,879960975 | 0,3507261 | 2,5089692 | 0,0121084 | 0,0408565 | NEU2 |
| ENSGALG00000009580 | 19,504731 | 0,879798414 | 0,32149 | 2,7366276 | 0,0062073 | 0,0239207 | IQCJ |
| ENSGALG00000016986 | 175,40028 | 0,879609126 | 0,2982944 | 2,9487952 | 0,0031902 | 0,0138545 | LCP1 |
| ENSGALG00000008349 | 209,56657 | 0,879571782 | 0,2609378 | 3,3708107 | 0,0007495 | 0,0042079 | LMCD1 |
| ENSGALG00000009601 | 8743,3872 | 0,878730197 | 0,1376458 | 6,3839954 | 1,73E-10 | 4,59E-09 | NELL2 |
| ENSGALG00000008121 | 42,835354 | 0,876566467 | 0,3539561 | 2,4764833 | 0,0132684 | 0,0439535 | CYP17A1 |
| ENSGALG00000008364 | 36,778976 | 0,876075442 | 0,3534635 | 2,4785458 | 0,0131919 | 0,0437898 | |
| ENSGALG00000023415 | 29,787572 | 0,875042862 | 0,3509632 | 2,4932611 | 0,0126576 | 0,0424047 | HMX3 |
| ENSGALG00000027920 | 386,75217 | 0,874011674 | 0,2284444 | 3,8259279 | 0,0001303 | 0,0009425 | CPLX2 |
| ENSGALG00000026059 | 270,41633 | 0,873870501 | 0,2397099 | 3,6455337 | 0,0002668 | 0,0017508 | TWIST1 |
| ENSGALG00000004788 | 68,83418 | 0,873507388 | 0,3452821 | 2,5298369 | 0,0114116 | 0,0389981 | TNFAIP8L3 |
| ENSGALG00000011219 | 83,302927 | 0,873388378 | 0,3403042 | 2,5664933 | 0,0102733 | 0,0359195 | GCGR |
| ENSGALG00000016017 | 590,76905 | 0,873175292 | 0,2154495 | 4,0528075 | 5,06E-05 | 0,000416 | SLC4A11 |
| ENSGALG00000005974 | 256,91419 | 0,872920311 | 0,2521004 | 3,4625904 | 0,000535 | 0,0031739 | COL6A1 |
| ENSGALG00000012422 | 27,355404 | 0,872315124 | 0,3483618 | 2,5040496 | 0,0122781 | 0,0413266 | SH3BP1 |
| ENSGALG00000026939 | 41,153772 | 0,872255445 | 0,3538073 | 2,4653407 | 0,0136883 | 0,044964 | |
| ENSGALG00000009640 | 27,779879 | 0,872186333 | 0,3505999 | 2,487697 | 0,0128573 | 0,0429155 | CHRM5 |

| | | | | | | | |
|---|---|---|---|---|---|---|---|
| ENSGALG00000001840 | 243,72168 | 0,872138912 | 0,2890387 | 3,0173773 | 0,0025497 | 0,0114911 | TEK |
| ENSGALG00000001129 | 1785,0406 | 0,871868135 | 0,1530444 | 5,69683 | 1,22E-08 | 2,38E-07 | KIRREL3 |
| ENSGALG00000028928 | 43,951223 | 0,871728708 | 0,3534532 | 2,4663203 | 0,0136509 | 0,0448931 | LCAT |
| ENSGALG00000001532 | 54,980298 | 0,870029294 | 0,3496522 | 2,4882708 | 0,0128366 | 0,0428667 | WNT2B |
| ENSGALG00000009867 | 212,19054 | 0,869748432 | 0,2675059 | 3,2513241 | 0,0011487 | 0,0059997 | WIF1 |
| ENSGALG00000027364 | 6291,5882 | 0,867638443 | 0,1162491 | 7,4636145 | 8,42E-14 | 3,48E-12 | HEY1 |
| ENSGALG00000026246 | 29,222799 | 0,867468639 | 0,3517182 | 2,4663739 | 0,0136489 | 0,0448931 | GIF |
| ENSGALG00000020876 | 244,82737 | 0,867421463 | 0,2948139 | 2,942268 | 0,0032582 | 0,0140826 | AOX2 |
| ENSGALG00000011624 | 479,03518 | 0,867273579 | 0,2018671 | 4,2962601 | 1,74E-05 | 0,000166 | PNLDC1 |
| ENSGALG00000016384 | 268,59226 | 0,867172927 | 0,2685681 | 3,2288758 | 0,0012428 | 0,0064011 | MYT1L |
| ENSGALG00000013515 | 39,808741 | 0,866546513 | 0,3539601 | 2,4481471 | 0,0143593 | 0,0467149 | RP11-287D1.3 |
| ENSGALG00000000225 | 112,7664 | 0,866321273 | 0,3372745 | 2,5685943 | 0,0102112 | 0,0357668 | MGAT4C |
| ENSGALG00000011354 | 21,346176 | 0,866050474 | 0,3402355 | 2,5454444 | 0,0109139 | 0,0377138 | CRYBA2 |
| ENSGALG00000027965 | 101,43615 | 0,865747962 | 0,3205874 | 2,700505 | 0,0069234 | 0,0261695 | OPRL1 |
| ENSGALG00000026172 | 493,17496 | 0,865691508 | 0,2111797 | 4,0993117 | 4,14E-05 | 0,0003508 | CYBA |
| ENSGALG00000016115 | 186,82276 | 0,864642259 | 0,2830702 | 3,0545156 | 0,0022542 | 0,0103983 | SAMD12 |
| ENSGALG00000027380 | 43,976414 | 0,862762922 | 0,3539532 | 2,4375054 | 0,014789 | 0,0478271 | |
| ENSGALG00000002206 | 95,268458 | 0,862737257 | 0,3285875 | 2,6255938 | 0,0086498 | 0,031296 | SNX22 |
| ENSGALG00000019804 | 21,501251 | 0,862231301 | 0,3371774 | 2,5572037 | 0,0105517 | 0,0367422 | |
| ENSGALG00000003895 | 788,02112 | 0,861640305 | 0,2589823 | 3,3270233 | 0,0008778 | 0,0047996 | PRDM12 |
| ENSGALG00000012072 | 201,72423 | 0,861021311 | 0,2753299 | 3,1272347 | 0,0017646 | 0,0084984 | PARP14 |
| ENSGALG00000007838 | 40,646821 | 0,860354034 | 0,3539557 | 2,4306829 | 0,0150704 | 0,0485568 | NLRC3 |
| ENSGALG00000000953 | 1344,7125 | 0,860062109 | 0,1616375 | 5,3209304 | 1,03E-07 | 1,66E-06 | TEAD3 |
| ENSGALG00000009902 | 340,49684 | 0,859561208 | 0,2490936 | 3,4507553 | 0,000559 | 0,0032934 | CFAP57 |
| ENSGALG00000005655 | 299,93592 | 0,858231064 | 0,2557112 | 3,3562518 | 0,0007901 | 0,0044067 | TSPEAR |
| ENSGALG00000017072 | 1099,8408 | 0,857400203 | 0,1810022 | 4,7369608 | 2,17E-06 | 2,59E-05 | N4BP2L1 |
| ENSGALG00000025919 | 2494,4885 | 0,857090747 | 0,142817 | 6,0013216 | 1,96E-09 | 4,45E-08 | BAMBI |
| ENSGALG00000016473 | 44,778577 | 0,856849135 | 0,3459792 | 2,4765913 | 0,0132644 | 0,0439509 | OSR1 |
| ENSGALG00000017228 | 49,296654 | 0,856739492 | 0,3532728 | 2,4251498 | 0,0153021 | 0,0491478 | MTNR1B |
| ENSGALG00000016755 | 172,26247 | 0,856689481 | 0,2873424 | 2,9814236 | 0,0028691 | 0,0126954 | KIAA1211L |
| ENSGALG00000007376 | 87,397516 | 0,856687427 | 0,3445883 | 2,4861189 | 0,0129145 | 0,0430009 | CCDC60 |
| ENSGALG00000015768 | 2710,7019 | 0,855142896 | 0,1297486 | 6,5907703 | 4,38E-11 | 1,25E-09 | ANKRD6 |
| ENSGALG00000026896 | 77,09018 | 0,854217305 | 0,3380109 | 2,527189 | 0,011498 | 0,0392048 | |
| ENSGALG00000012515 | 43,853432 | 0,854026842 | 0,3512742 | 2,4312255 | 0,0150478 | 0,0485029 | TPPP |
| ENSGALG00000010664 | 712,1771 | 0,852579615 | 0,2288179 | 3,7260174 | 0,0001945 | 0,0013328 | KIF26B |
| ENSGALG00000001189 | 452,19238 | 0,851541498 | 0,2063764 | 4,126158 | 3,69E-05 | 0,0003169 | NCF1 |
| ENSGALG00000009536 | 446,62772 | 0,851130737 | 0,2275785 | 3,7399437 | 0,0001841 | 0,0012727 | |
| ENSGALG00000003612 | 268,62308 | 0,850682606 | 0,2669792 | 3,1863255 | 0,0014409 | 0,0072179 | RNLS |
| ENSGALG00000023172 | 1255,7616 | 0,849498162 | 0,1537527 | 5,5250928 | 3,29E-08 | 5,85E-07 | |
| ENSGALG00000004511 | 28,137324 | 0,848004478 | 0,350316 | 2,4206844 | 0,0154913 | 0,0496386 | MTTPL |
| ENSGALG00000026671 | 2116,5692 | 0,847898773 | 0,1425868 | 5,9465465 | 2,74E-09 | 6,10E-08 | |
| ENSGALG00000002742 | 2787,3387 | 0,847416181 | 0,1449021 | 5,8481963 | 4,97E-09 | 1,04E-07 | TMEM132B |
| ENSGALG00000015670 | 217,38931 | 0,846232332 | 0,2802832 | 3,0192043 | 0,0025344 | 0,0114409 | HNF4G |
| ENSGALG00000005791 | 520,60885 | 0,84592391 | 0,2039846 | 4,146998 | 3,37E-05 | 0,0002934 | JPH3 |
| ENSGALG00000009603 | 307,91481 | 0,844955114 | 0,2387657 | 3,5388467 | 0,0004019 | 0,0024962 | |
| ENSGALG00000006379 | 1670,5263 | 0,844378171 | 0,1827468 | 4,6204804 | 3,83E-06 | 4,35E-05 | SHH |
| ENSGALG00000005895 | 481,78824 | 0,844174027 | 0,2049844 | 4,1182354 | 3,82E-05 | 0,0003271 | FYTTD1L |
| ENSGALG00000026103 | 166,74522 | 0,842618716 | 0,3162093 | 2,6647499 | 0,0077046 | 0,0286293 | CACNG3 |
| ENSGALG00000016677 | 26,013707 | 0,842209203 | 0,3459083 | 2,4347759 | 0,014901 | 0,0481322 | IL17F |
| ENSGALG00000003952 | 60,952026 | 0,841003956 | 0,3435108 | 2,4482605 | 0,0143548 | 0,0467114 | RBPJL |
| ENSGALG00000007043 | 22,708865 | 0,840648204 | 0,3395839 | 2,4755244 | 0,0133041 | 0,0440503 | |
| ENSGALG00000007775 | 18,86724 | 0,840390709 | 0,3327596 | 2,525519 | 0,0115528 | 0,0393522 | USH1G |
| ENSGALG00000000731 | 28,360648 | 0,840347147 | 0,3458767 | 2,4296148 | 0,0151149 | 0,0486392 | THEMIS2 |
| ENSGALG00000004593 | 2963,7081 | 0,839985137 | 0,1559067 | 5,3877418 | 7,13E-08 | 1,18E-06 | SULF2 |
| ENSGALG00000010857 | 11562,252 | 0,839106931 | 0,1136809 | 7,381249 | 1,57E-13 | 6,34E-12 | DAB1 |
| ENSGALG00000004946 | 111,2048 | 0,83835973 | 0,3100886 | 2,7036139 | 0,006859 | 0,0259771 | COL4A4 |
| ENSGALG00000010762 | 102,40709 | 0,837861719 | 0,3182395 | 2,6328025 | 0,0084684 | 0,0307881 | OPN3 |
| ENSGALG00000005204 | 101,3211 | 0,837525509 | 0,330602 | 2,5333346 | 0,0112983 | 0,0387374 | GSTT1L |
| ENSGALG00000026505 | 1579,9758 | 0,837091894 | 0,1526832 | 5,48254 | 4,19E-08 | 7,29E-07 | RDH10 |
| ENSGALG00000003354 | 276,83001 | 0,837083394 | 0,2477788 | 3,3783489 | 0,0007292 | 0,0041177 | KCNH4 |
| ENSGALG00000011205 | 1172,7422 | 0,836341019 | 0,2011925 | 4,1569189 | 3,23E-05 | 0,0002826 | SMOC2 |
| ENSGALG00000005161 | 2738,7124 | 0,836184904 | 0,1385814 | 6,0338884 | 1,60E-09 | 3,70E-08 | PHF2 |
| ENSGALG00000006014 | 961,40069 | 0,836085862 | 0,1594344 | 5,2440759 | 1,57E-07 | 2,43E-06 | PRKCB |
| ENSGALG00000007770 | 193,51785 | 0,835610241 | 0,2684966 | 3,1121814 | 0,0018571 | 0,00885 | FADS6 |

| | | | | | | | |
|---|---|---|---|---|---|---|---|
| ENSGALG00000020515 | 13,200692 | 0,834854635 | 0,2996969 | 2,7856631 | 0,0053418 | 0,0211799 | |
| ENSGALG00000010044 | 118,46337 | 0,834748583 | 0,304154 | 2,7444935 | 0,0060604 | 0,0234611 | KCNMB4 |
| ENSGALG00000001071 | 194,63937 | 0,834483858 | 0,2807285 | 2,9725654 | 0,0029532 | 0,0130039 | ELN |
| ENSGALG00000026098 | 20,591793 | 0,833498696 | 0,3202723 | 2,6024691 | 0,0092555 | 0,0330238 | IL8L2 |
| ENSGALG00000028582 | 1565,1403 | 0,832821984 | 0,1507685 | 5,5238471 | 3,32E-08 | 5,87E-07 | LMO2 |
| ENSGALG00000005481 | 1783,1099 | 0,83266607 | 0,1448108 | 5,7500286 | 8,92E-09 | 1,77E-07 | ARHGEF3 |
| ENSGALG00000015660 | 2889,8193 | 0,831570906 | 0,1228249 | 6,7703746 | 1,28E-11 | 4,00E-10 | SNX30 |
| ENSGALG00000028125 | 921,39565 | 0,831550629 | 0,1803628 | 4,6104334 | 4,02E-06 | 4,52E-05 | CACNG4 |
| ENSGALG00000011883 | 233,92984 | 0,831245154 | 0,2791409 | 2,9778696 | 0,0029026 | 0,0128183 | CLEC3B |
| ENSGALG00000027201 | 25,956081 | 0,831158121 | 0,3370736 | 2,4658058 | 0,0136705 | 0,0449359 | |
| ENSGALG00000011808 | 15,036213 | 0,830943596 | 0,3107832 | 2,6737086 | 0,0075018 | 0,0279749 | CCR9 |
| ENSGALG00000002771 | 197,13737 | 0,83021733 | 0,2733379 | 3,0373302 | 0,0023868 | 0,01091 | AL928654.7 |
| ENSGALG00000025776 | 414,67908 | 0,83014723 | 0,214629 | 3,8678243 | 0,0001098 | 0,0008117 | EMID1 |
| ENSGALG00000027903 | 10650,644 | 0,829170411 | 0,110533 | 7,501567 | 6,31E-14 | 2,64E-12 | HOXB5 |
| ENSGALG00000010500 | 3026,4125 | 0,828235743 | 0,1300266 | 6,3697426 | 1,89E-10 | 4,98E-09 | NKX6-2 |
| ENSGALG00000024487 | 160,03616 | 0,827463833 | 0,3258506 | 2,5393961 | 0,0111044 | 0,0382267 | C8orf22 |
| ENSGALG00000020002 | 21,954447 | 0,826702298 | 0,3346434 | 2,4703975 | 0,0134963 | 0,0445036 | SLC22A3 |
| ENSGALG00000007015 | 336,01016 | 0,825119263 | 0,2296496 | 3,5929492 | 0,000327 | 0,0020958 | CD40 |
| ENSGALG00000015720 | 662,9716 | 0,824601233 | 0,2077728 | 3,9687648 | 7,22E-05 | 0,000566 | CHODL |
| ENSGALG00000011744 | 1298,602 | 0,824149013 | 0,1729843 | 4,7642989 | 1,90E-06 | 2,30E-05 | KALRN |
| ENSGALG00000015575 | 115,532 | 0,823970797 | 0,3263661 | 2,5246826 | 0,0115803 | 0,0394262 | HTRA3 |
| ENSGALG00000017155 | 286,34498 | 0,823340146 | 0,2794831 | 2,9459393 | 0,0032198 | 0,0139435 | C11orf87 |
| ENSGALG00000016558 | 512,36729 | 0,822973317 | 0,2009905 | 4,0945888 | 4,23E-05 | 0,0003559 | FIGF |
| ENSGALG00000002383 | 12,21302 | 0,822929288 | 0,2987538 | 2,75454 | 0,0058775 | 0,0228764 | CD72AG |
| ENSGALG00000006833 | 142,93183 | 0,822822635 | 0,2953281 | 2,7861304 | 0,0053341 | 0,0211609 | ITIH2 |
| ENSGALG00000005662 | 148,91563 | 0,822087333 | 0,2869197 | 2,865217 | 0,0041672 | 0,01724 | AMPD3 |
| ENSGALG00000001688 | 9554,3654 | 0,822044682 | 0,1265194 | 6,497383 | 8,17E-11 | 2,27E-09 | sept-05 |
| ENSGALG00000015941 | 330,15649 | 0,819153887 | 0,2336045 | 3,5065841 | 0,0004539 | 0,0027625 | AP000295.9 |
| ENSGALG00000019768 | 3223,3224 | 0,819142269 | 0,1222158 | 6,7024262 | 2,05E-11 | 6,19E-10 | ACSF2 |
| ENSGALG00000003360 | 122,83324 | 0,818748241 | 0,302344 | 2,7080025 | 0,006769 | 0,0257069 | ARL10 |
| ENSGALG00000026276 | 151,57437 | 0,818604795 | 0,2887563 | 2,834933 | 0,0045835 | 0,0186339 | TCF15 |
| ENSGALG00000003014 | 83,658638 | 0,818530331 | 0,3298981 | 2,4811612 | 0,0130955 | 0,0435185 | CHRNA3 |
| ENSGALG00000005175 | 201,26271 | 0,818212824 | 0,2767102 | 2,9569306 | 0,0031072 | 0,0135504 | BAAT |
| ENSGALG00000017089 | 188,71558 | 0,817752266 | 0,2686445 | 3,0439946 | 0,0023346 | 0,0107165 | SLC46A3 |
| ENSGALG00000016774 | 140,17126 | 0,817235755 | 0,294664 | 2,7734498 | 0,0055465 | 0,0218779 | NPAS2 |
| ENSGALG00000010698 | 1578,3511 | 0,816916741 | 0,1404823 | 5,8150867 | 6,06E-09 | 1,25E-07 | RPS6KA5 |
| ENSGALG00000012628 | 1115,1061 | 0,816609796 | 0,1882236 | 4,3385098 | 1,43E-05 | 0,0001415 | HABP4 |
| ENSGALG00000027830 | 138,16537 | 0,816343449 | 0,2952222 | 2,7651828 | 0,0056891 | 0,0222705 | CACNG2 |
| ENSGALG00000027534 | 844,14704 | 0,815389343 | 0,1740275 | 4,6854054 | 2,79E-06 | 3,27E-05 | GM2A |
| ENSGALG00000023601 | 110,08849 | 0,814252411 | 0,3257765 | 2,4994207 | 0,0124397 | 0,0418084 | |
| ENSGALG00000010397 | 282,5332 | 0,813527008 | 0,2534717 | 3,2095382 | 0,0013295 | 0,0067667 | GRID2 |
| ENSGALG00000027338 | 409,81098 | 0,813282814 | 0,2198892 | 3,6986034 | 0,0002168 | 0,0014663 | |
| ENSGALG00000015297 | 1260,5465 | 0,81224071 | 0,1674371 | 4,8510192 | 1,23E-06 | 1,57E-05 | FOXO3 |
| ENSGALG00000000954 | 405,07056 | 0,812029159 | 0,2420413 | 3,3549193 | 0,0007939 | 0,0044207 | SMPD2 |
| ENSGALG00000028570 | 89,253148 | 0,811507626 | 0,3263252 | 2,4868067 | 0,0128895 | 0,0429494 | C6orf132 |
| ENSGALG00000006480 | 1820,4374 | 0,810690895 | 0,1479509 | 5,4794584 | 4,27E-08 | 7,41E-07 | TCF7 |
| ENSGALG00000011861 | 430,59758 | 0,810665245 | 0,2125716 | 3,8136096 | 0,000137 | 0,0009866 | SYT16 |
| ENSGALG00000011615 | 12,005704 | 0,809627409 | 0,2973783 | 2,7225505 | 0,006478 | 0,0248312 | SLC22A2 |
| ENSGALG00000009680 | 349,5215 | 0,809243098 | 0,2208092 | 3,6648968 | 0,0002474 | 0,0016447 | PAQR7 |
| ENSGALG00000001605 | 101,47062 | 0,807232306 | 0,3186097 | 2,5336084 | 0,0112895 | 0,0387248 | PPM1J |
| ENSGALG00000015051 | 777,65205 | 0,807158582 | 0,1825122 | 4,4224914 | 9,76E-06 | 0,0001009 | CDK19 |
| ENSGALG00000013051 | 1092,6973 | 0,806546944 | 0,1922229 | 4,1958947 | 2,72E-05 | 0,0002436 | SEMA5A |
| ENSGALG00000024090 | 246,87805 | 0,806200799 | 0,263668 | 3,0576358 | 0,0022309 | 0,0103145 | UNC93B1 |
| ENSGALG00000005147 | 486,92842 | 0,80606613 | 0,2005832 | 4,0186116 | 5,85E-05 | 0,0004713 | AMY1A |
| ENSGALG00000013600 | 13,810453 | 0,805946686 | 0,307319 | 2,6225087 | 0,0087285 | 0,0315473 | |
| ENSGALG00000004260 | 114,05372 | 0,805411915 | 0,3114119 | 2,5863235 | 0,0097006 | 0,0343137 | PKP3 |
| ENSGALG00000008980 | 97,942464 | 0,805130642 | 0,3170949 | 2,5390838 | 0,0111143 | 0,0382415 | VWA2 |
| ENSGALG00000017016 | 265,33781 | 0,804980828 | 0,247719 | 3,2495724 | 0,0011558 | 0,0060245 | FAM124A |
| ENSGALG00000015608 | 746,79897 | 0,802634493 | 0,1733654 | 4,629727 | 3,66E-06 | 4,17E-05 | ADRA2C |
| ENSGALG00000011394 | 463,3825 | 0,801850887 | 0,2051329 | 3,9089335 | 9,27E-05 | 0,0007038 | CYP2J2L3 |
| ENSGALG00000003261 | 114,4772 | 0,800174677 | 0,3173455 | 2,5214623 | 0,0116868 | 0,0397492 | PTRF |
| ENSGALG00000007186 | 16,312433 | 0,798835823 | 0,3250305 | 2,457726 | 0,013982 | 0,0457372 | C9orf91 |
| ENSGALG00000007565 | 677,7029 | 0,798729286 | 0,2213224 | 3,6088951 | 0,0003075 | 0,0019851 | PPP2R2B |
| ENSGALG00000027043 | 126,47192 | 0,798613446 | 0,3175009 | 2,515311 | 0,0118927 | 0,0403187 | KCNH2 |

| | | | | | | | |
|---|---|---|---|---|---|---|---|
| ENSGALG00000012073 | 470,07631 | 0,797159786 | 0,2039204 | 3,9091719 | 9,26E-05 | 0,0007038 | SFRP4 |
| ENSGALG00000008408 | 1323,0351 | 0,796075417 | 0,1521355 | 5,2326742 | 1,67E-07 | 2,57E-06 | GAB3 |
| ENSGALG00000014122 | 10545,592 | 0,795943411 | 0,1349861 | 5,8964838 | 3,71E-09 | 8,08E-08 | CYB5R3 |
| ENSGALG00000000865 | 119,7276 | 0,795679669 | 0,3113005 | 2,5559862 | 0,0105887 | 0,0368427 | |
| ENSGALG00000014313 | 191,48762 | 0,795282607 | 0,2722671 | 2,9209643 | 0,0034895 | 0,0149206 | |
| ENSGALG00000013489 | 630,05884 | 0,794845244 | 0,1872111 | 4,2457157 | 2,18E-05 | 0,000202 | CCDC82 |
| ENSGALG00000015233 | 5666,2368 | 0,794464087 | 0,1575131 | 5,0437964 | 4,56E-07 | 6,39E-06 | ATP1B1 |
| ENSGALG00000006507 | 1217,4322 | 0,793097237 | 0,1534206 | 5,169432 | 2,35E-07 | 3,49E-06 | CLMP |
| ENSGALG00000017267 | 12,565544 | 0,792427514 | 0,2952723 | 2,6837179 | 0,0072809 | 0,0272932 | THRSP |
| ENSGALG00000026605 | 1028,743 | 0,792369586 | 0,1581582 | 5,0099806 | 5,44E-07 | 7,52E-06 | BAG2 |
| ENSGALG00000007720 | 244,44298 | 0,792327844 | 0,2555746 | 3,1001818 | 0,001934 | 0,009178 | AIFM3 |
| ENSGALG00000006319 | 831,26714 | 0,791303644 | 0,1737644 | 4,5538886 | 5,27E-06 | 5,80E-05 | TGFBI |
| ENSGALG00000028576 | 735,47031 | 0,790565069 | 0,2313869 | 3,4166365 | 0,000634 | 0,0036529 | PAX1 |
| ENSGALG00000010708 | 2290,4638 | 0,790532368 | 0,1502852 | 5,2602149 | 1,44E-07 | 2,24E-06 | ICA1 |
| ENSGALG00000021589 | 1830,7464 | 0,790216265 | 0,177386 | 4,4547837 | 8,40E-06 | 8,83E-05 | RTN4RL1 |
| ENSGALG00000027458 | 914,19579 | 0,790135397 | 0,1634873 | 4,8330066 | 1,34E-06 | 1,70E-05 | |
| ENSGALG00000014300 | 310,95646 | 0,790002156 | 0,2504565 | 3,1542493 | 0,0016091 | 0,0078863 | KLB |
| ENSGALG00000013862 | 6295,0756 | 0,789845326 | 0,1439094 | 5,4884886 | 4,05E-08 | 7,08E-07 | OLIG3 |
| ENSGALG00000026899 | 678,26051 | 0,789824953 | 0,18654 | 4,2340774 | 2,29E-05 | 0,000211 | CHSY3 |
| ENSGALG00000020522 | 2265,9648 | 0,78974705 | 0,152638 | 5,1739888 | 2,29E-07 | 3,42E-06 | MYRF |
| ENSGALG00000009039 | 1670,7449 | 0,789311851 | 0,1468245 | 5,375886 | 7,62E-08 | 1,25E-06 | NPDC1 |
| ENSGALG00000003933 | 4489,5312 | 0,788744436 | 0,1311022 | 6,0162586 | 1,78E-09 | 4,11E-08 | QSOX1 |
| ENSGALG00000013642 | 1029,7112 | 0,788560165 | 0,1619937 | 4,8678436 | 1,13E-06 | 1,46E-05 | KAZN |
| ENSGALG00000006453 | 2069,6286 | 0,788464487 | 0,1533354 | 5,1420896 | 2,72E-07 | 3,98E-06 | TF |
| ENSGALG00000007623 | 3327,2226 | 0,788173273 | 0,1396726 | 5,6430046 | 1,67E-08 | 3,18E-07 | CACNA1G |
| ENSGALG00000024097 | 498,63238 | 0,787003033 | 0,253676 | 3,102395 | 0,0019196 | 0,0091192 | AGBL2 |
| ENSGALG00000011913 | 446,23727 | 0,78594155 | 0,2729807 | 2,8791099 | 0,003988 | 0,0166514 | sept-03 |
| ENSGALG00000005849 | 1033,3817 | 0,784505411 | 0,1637307 | 4,7914382 | 1,66E-06 | 2,05E-05 | PPDPF |
| ENSGALG00000000076 | 922,87992 | 0,7843985 | 0,1622181 | 4,8354571 | 1,33E-06 | 1,68E-05 | NRBP2 |
| ENSGALG00000016507 | 201,43925 | 0,784023754 | 0,2659211 | 2,9483321 | 0,0031949 | 0,0138709 | FNDC4 |
| ENSGALG00000012116 | 675,67694 | 0,783832372 | 0,1875924 | 4,1783795 | 2,94E-05 | 0,0002606 | EEPD1 |
| ENSGALG00000006508 | 175,86669 | 0,783784604 | 0,2797588 | 2,8016444 | 0,0050843 | 0,0203416 | FGF13 |
| ENSGALG00000011635 | 890,9248 | 0,783625313 | 0,1778317 | 4,4065551 | 1,05E-05 | 0,0001078 | TFCP2L1 |
| ENSGALG00000000313 | 165,09493 | 0,78307364 | 0,2881217 | 2,7178574 | 0,0065706 | 0,0251007 | TNNI1 |
| ENSGALG00000003900 | 14,168362 | 0,782866932 | 0,3173181 | 2,467136 | 0,0136199 | 0,0448342 | CACNG1 |
| ENSGALG00000027788 | 2064,8116 | 0,782858625 | 0,1847884 | 4,2365144 | 2,27E-05 | 0,0002093 | CPE |
| ENSGALG00000009866 | 599,48537 | 0,782677583 | 0,2085508 | 3,7529357 | 0,0001748 | 0,0012203 | CAMK1D |
| ENSGALG00000023069 | 225,10329 | 0,782582264 | 0,2592407 | 3,0187478 | 0,0025382 | 0,0114506 | MYRIP |
| ENSGALG00000020376 | 2337,0409 | 0,781084057 | 0,129917 | 6,0121758 | 1,83E-09 | 4,20E-08 | |
| ENSGALG00000024447 | 255,3388 | 0,780783451 | 0,2581275 | 3,0247981 | 0,002488 | 0,0112763 | NMB |
| ENSGALG00000007357 | 1727,6304 | 0,780757803 | 0,1788255 | 4,3660311 | 1,27E-05 | 0,0001264 | BCL6 |
| ENSGALG00000021240 | 149,64373 | 0,780471598 | 0,2990547 | 2,6097954 | 0,0090596 | 0,0324549 | |
| ENSGALG00000009407 | 420,64448 | 0,780277328 | 0,2147109 | 3,6340831 | 0,000279 | 0,0018234 | |
| ENSGALG00000021620 | 400,19342 | 0,780112352 | 0,215822 | 3,6146106 | 0,0003008 | 0,0019465 | TMEM59L |
| ENSGALG00000005657 | 1485,5044 | 0,780089243 | 0,2099353 | 3,7158564 | 0,0002025 | 0,0013786 | CRHR2 |
| ENSGALG00000017327 | 228,75567 | 0,779838018 | 0,2748161 | 2,8376719 | 0,0045444 | 0,0184913 | P2RY6 |
| ENSGALG00000017065 | 605,99073 | 0,779643959 | 0,2212955 | 3,5230893 | 0,0004265 | 0,0026302 | STARD13 |
| ENSGALG00000015379 | 3926,961 | 0,779239634 | 0,1720773 | 4,5284285 | 5,94E-06 | 6,41E-05 | TAGLN3 |
| ENSGALG00000010898 | 121,86089 | 0,778406503 | 0,3067735 | 2,5373979 | 0,011168 | 0,0383873 | BMP3 |
| ENSGALG00000001735 | 332,00127 | 0,778269785 | 0,3162815 | 2,4606869 | 0,0138671 | 0,0454285 | LHX4 |
| ENSGALG00000005418 | 474,48986 | 0,777790922 | 0,2764455 | 2,8135412 | 0,0048999 | 0,019755 | FRRS1 |
| ENSGALG00000015615 | 136,43246 | 0,777196481 | 0,3023569 | 2,5704607 | 0,0101563 | 0,0355929 | DOK7 |
| ENSGALG00000005539 | 507,50867 | 0,77704495 | 0,1983441 | 3,9176614 | 8,94E-05 | 0,0006824 | ITM2C |
| ENSGALG00000009612 | 1284,262 | 0,776962988 | 0,1567277 | 4,9574072 | 7,14E-07 | 9,55E-06 | TGFB2 |
| ENSGALG00000005421 | 527,43362 | 0,776882668 | 0,193304 | 4,0189685 | 5,85E-05 | 0,0004709 | TEX2L |
| ENSGALG00000017086 | 244,06613 | 0,776712467 | 0,2590522 | 2,9982852 | 0,002715 | 0,0121396 | MTUS2 |
| ENSGALG00000026326 | 2889,7968 | 0,77526969 | 0,1465876 | 5,2887801 | 1,23E-07 | 1,94E-06 | WBP1L |
| ENSGALG00000003747 | 210,55966 | 0,773681017 | 0,268223 | 2,8844693 | 0,0039207 | 0,0164299 | OSMR |
| ENSGALG00000002708 | 2271,2308 | 0,772455755 | 0,1693238 | 4,5620036 | 5,07E-06 | 5,60E-05 | LINGO1 |
| ENSGALG00000011986 | 13,457929 | 0,772453397 | 0,3038739 | 2,5420199 | 0,0110214 | 0,0380183 | FABP2 |
| ENSGALG00000004183 | 169,51264 | 0,772146407 | 0,2847172 | 2,7119762 | 0,0066883 | 0,0255075 | FAM20A |
| ENSGALG00000013404 | 465,07043 | 0,770980594 | 0,2045333 | 3,7694617 | 0,0001636 | 0,0011517 | AK8 |
| ENSGALG00000002283 | 1172,584 | 0,770174106 | 0,1718025 | 4,482905 | 7,36E-06 | 7,80E-05 | MAPKAPK3 |
| ENSGALG00000017208 | 521,56067 | 0,769802089 | 0,1959747 | 3,9280679 | 8,56E-05 | 0,0006572 | AMOTL1 |

| | | | | | | | |
|---|---|---|---|---|---|---|---|
| ENSGALG00000013350 | 163,60204 | 0,766573209 | 0,2847601 | 2,6919969 | 0,0071026 | 0,0267426 | MTMR11 |
| ENSGALG00000027592 | 1174,3515 | 0,765780883 | 0,1667631 | 4,5920291 | 4,39E-06 | 4,91E-05 | ERN1 |
| ENSGALG00000005779 | 2301,589 | 0,765244932 | 0,1290039 | 5,9319529 | 2,99E-09 | 6,58E-08 | FBXO31 |
| ENSGALG00000006900 | 414,72698 | 0,764663976 | 0,2420639 | 3,158934 | 0,0015835 | 0,0077802 | ST6GALNAC2L |
| ENSGALG00000004323 | 3702,9555 | 0,7637615 | 0,1520557 | 5,0229053 | 5,09E-07 | 7,07E-06 | RASAL2 |
| ENSGALG00000007744 | 1224,4032 | 0,76351713 | 0,1645392 | 4,6403371 | 3,48E-06 | 3,98E-05 | CASS4 |
| ENSGALG00000000897 | 268,90562 | 0,763363816 | 0,249632 | 3,0579566 | 0,0022285 | 0,0103145 | LHFPL5 |
| ENSGALG00000004491 | 190,54596 | 0,761794314 | 0,2733192 | 2,7871964 | 0,0053166 | 0,0211161 | DMGDH |
| ENSGALG00000002069 | 610,16885 | 0,761037525 | 0,2096031 | 3,6308501 | 0,0002825 | 0,0018438 | FAM214B |
| ENSGALG00000013948 | 736,87288 | 0,760675086 | 0,1727814 | 4,402528 | 1,07E-05 | 0,0001095 | RASL11B |
| ENSGALG00000006642 | 162,08607 | 0,760653303 | 0,2972229 | 2,5592016 | 0,0104913 | 0,0365597 | ADORA2A |
| ENSGALG00000015115 | 155,99801 | 0,760473682 | 0,2860137 | 2,6588715 | 0,0078403 | 0,028999 | MAMDC2 |
| ENSGALG00000002804 | 155,62632 | 0,759679339 | 0,2858837 | 2,6573024 | 0,0078769 | 0,0290949 | RND2 |
| ENSGALG00000007956 | 2266,7266 | 0,759404394 | 0,1455052 | 5,2190893 | 1,80E-07 | 2,75E-06 | PLXDC2 |
| ENSGALG00000007748 | 2468,6975 | 0,758936347 | 0,126557 | 5,9967954 | 2,01E-09 | 4,56E-08 | SLC16A2 |
| ENSGALG00000004170 | 385,32134 | 0,75875998 | 0,2343812 | 3,23729 | 0,0012067 | 0,0062493 | ADA |
| ENSGALG00000015688 | 1188,5305 | 0,758515207 | 0,1730216 | 4,3839346 | 1,17E-05 | 0,0001181 | MXD4 |
| ENSGALG00000006576 | 399,34369 | 0,758241661 | 0,2615129 | 2,8994424 | 0,0037383 | 0,015831 | GRM3 |
| ENSGALG00000001933 | 120,41471 | 0,757286355 | 0,3102287 | 2,4410587 | 0,0146443 | 0,0474719 | CORO2A |
| ENSGALG00000003115 | 505,20482 | 0,756684773 | 0,2022143 | 3,7419946 | 0,0001826 | 0,0012662 | COL4A3 |
| ENSGALG00000007889 | 769,26198 | 0,756666335 | 0,1846622 | 4,0975693 | 4,18E-05 | 0,0003529 | CCDC85A |
| ENSGALG00000006346 | 1339,3203 | 0,756405715 | 0,1832878 | 4,1268742 | 3,68E-05 | 0,0003163 | CXCL14 |
| ENSGALG00000026153 | 524,42092 | 0,755983234 | 0,2314691 | 3,2660225 | 0,0010907 | 0,0057424 | FOXO6 |
| ENSGALG00000026091 | 117,64664 | 0,755485898 | 0,3112696 | 2,4271108 | 0,0152196 | 0,0489092 | |
| ENSGALG00000002958 | 2112,9504 | 0,755313619 | 0,167448 | 4,510735 | 6,46E-06 | 6,90E-05 | DNER |
| ENSGALG00000001311 | 1301,9009 | 0,752137925 | 0,1518518 | 4,9531061 | 7,30E-07 | 9,72E-06 | ARHGAP44 |
| ENSGALG00000027905 | 3422,0143 | 0,751625166 | 0,132987 | 5,6518692 | 1,59E-08 | 3,03E-07 | CSRNP1 |
| ENSGALG00000009439 | 631,53261 | 0,751125582 | 0,2024985 | 3,7092895 | 0,0002078 | 0,0014128 | GPR85 |
| ENSGALG00000006795 | 336,85781 | 0,75042189 | 0,2480756 | 3,0249723 | 0,0024866 | 0,0112763 | ABCG4 |
| ENSGALG00000029047 | 339,90745 | 0,749178885 | 0,226666 | 3,30521 | 0,0009491 | 0,0051255 | TMEM204 |
| ENSGALG00000015729 | 1439,0922 | 0,747405326 | 0,14976 | 4,9906871 | 6,02E-07 | 8,18E-06 | LPAR1 |
| ENSGALG00000019030 | 11,696679 | 0,747320322 | 0,2902826 | 2,5744581 | 0,0100397 | 0,035266 | |
| ENSGALG00000006203 | 182,30527 | 0,747208785 | 0,2854721 | 2,6174493 | 0,008859 | 0,0319171 | |
| ENSGALG00000001969 | 1252,395 | 0,746864271 | 0,1550045 | 4,8183402 | 1,45E-06 | 1,82E-05 | LPPR3 |
| ENSGALG00000006050 | 280,4499 | 0,74642112 | 0,2472463 | 3,0189372 | 0,0025366 | 0,0114472 | CYP2R1 |
| ENSGALG00000004420 | 2554,1446 | 0,745430944 | 0,1535099 | 4,8559134 | 1,20E-06 | 1,54E-05 | SDK1 |
| ENSGALG00000018733 | 341,10493 | 0,744919582 | 0,2235871 | 3,3316754 | 0,0008632 | 0,0047277 | |
| ENSGALG00000023828 | 228,11537 | 0,744788155 | 0,2585381 | 2,8807671 | 0,0039671 | 0,0165833 | LDHD |
| ENSGALG00000007849 | 195,07279 | 0,744715772 | 0,2780479 | 2,6783724 | 0,0073981 | 0,0276565 | CD44 |
| ENSGALG00000005063 | 630,43033 | 0,74466275 | 0,1956389 | 3,8063128 | 0,0001411 | 0,0010113 | ST6GALNAC4 |
| ENSGALG00000017050 | 201,15206 | 0,743800542 | 0,2755212 | 2,6996126 | 0,006942 | 0,0262325 | SMAD9 |
| ENSGALG00000026919 | 497,57567 | 0,743491927 | 0,2001741 | 3,7142261 | 0,0002038 | 0,0013869 | |
| ENSGALG00000025937 | 424,74226 | 0,743354667 | 0,2192559 | 3,3903524 | 0,000698 | 0,0039714 | FXYD6 |
| ENSGALG00000012712 | 2097,8069 | 0,742584862 | 0,1498005 | 4,9571602 | 7,15E-07 | 9,56E-06 | RBM24 |
| ENSGALG00000015994 | 1103,9909 | 0,741778145 | 0,1550114 | 4,7853126 | 1,71E-06 | 2,10E-05 | MAVS |
| ENSGALG00000016622 | 136,85244 | 0,741316835 | 0,3044477 | 2,4349564 | 0,0148936 | 0,0481221 | STS |
| ENSGALG00000001449 | 3789,5374 | 0,740070536 | 0,1451567 | 5,0984252 | 3,42E-07 | 4,93E-06 | STRA6 |
| ENSGALG00000006467 | 165,38559 | 0,739825587 | 0,2835935 | 2,6087534 | 0,0090873 | 0,0325074 | XKR7 |
| ENSGALG00000003022 | 149,11768 | 0,738879702 | 0,2881931 | 2,5638358 | 0,0103522 | 0,0361678 | |
| ENSGALG00000026449 | 3878,5798 | 0,737442369 | 0,1336353 | 5,5183189 | 3,42E-08 | 6,04E-07 | |
| ENSGALG00000011524 | 227,70396 | 0,737317525 | 0,2799 | 2,6342174 | 0,0084331 | 0,0306998 | PPEF2 |
| ENSGALG00000002539 | 465,9328 | 0,737264559 | 0,2112723 | 3,4896412 | 0,0004837 | 0,002919 | KCNN2 |
| ENSGALG00000022001 | 220,24358 | 0,737013016 | 0,2592948 | 2,8423753 | 0,0044779 | 0,0182643 | CAMKV |
| ENSGALG00000016749 | 204,45908 | 0,736312128 | 0,2623793 | 2,8062894 | 0,0050116 | 0,0200861 | CNGA3 |
| ENSGALG00000008339 | 615,63573 | 0,736061451 | 0,1948925 | 3,776756 | 0,0001589 | 0,0011208 | GRM7 |
| ENSGALG00000013002 | 3217,4534 | 0,734684533 | 0,1469698 | 4,9988824 | 5,77E-07 | 7,89E-06 | DAP |
| ENSGALG00000019087 | 2799,7159 | 0,734240753 | 0,1295389 | 5,6681115 | 1,44E-08 | 2,78E-07 | RFXAP |
| ENSGALG00000011059 | 526,88294 | 0,733913318 | 0,20212 | 3,6310777 | 0,0002822 | 0,001843 | HOX1B |
| ENSGALG00000019291 | 131,18453 | 0,733807509 | 0,2877581 | 2,5500843 | 0,0107697 | 0,0373103 | |
| ENSGALG00000010326 | 286,24182 | 0,733495449 | 0,2372268 | 3,0919588 | 0,0019884 | 0,0093882 | FLVCR2 |
| ENSGALG00000008433 | 2910,7777 | 0,732487 | 0,1303149 | 5,6209017 | 1,90E-08 | 3,57E-07 | FZD2 |
| ENSGALG00000008076 | 5047,2961 | 0,731953468 | 0,1274268 | 5,7441112 | 9,24E-09 | 1,83E-07 | TMEM164 |
| ENSGALG00000006897 | 1904,4846 | 0,73176707 | 0,1370838 | 5,3381014 | 9,39E-08 | 1,52E-06 | |
| ENSGALG00000025916 | 1511,635 | 0,730226914 | 0,1971166 | 3,7045423 | 0,0002118 | 0,0014359 | EMILIN3 |

| | | | | | | | |
|---|---|---|---|---|---|---|---|
| ENSGALG00000002651 | 2712,1143 | 0,730025866 | 0,128861 | 5,6652212 | 1,47E-08 | 2,82E-07 | ST3GAL2 |
| ENSGALG00000012191 | 7671,5827 | 0,72968259 | 0,1256369 | 5,8078663 | 6,33E-09 | 1,30E-07 | LGR4 |
| ENSGALG00000002451 | 1225,1572 | 0,728956851 | 0,1533766 | 4,7527269 | 2,01E-06 | 2,41E-05 | KCNT2 |
| ENSGALG00000019317 | 6925,165 | 0,728083244 | 0,1200043 | 6,0671438 | 1,30E-09 | 3,05E-08 | JOSD1 |
| ENSGALG00000001561 | 154,80244 | 0,72739532 | 0,2918515 | 2,4923472 | 0,0126902 | 0,0424526 | MXRA8 |
| ENSGALG00000013925 | 194,37903 | 0,727322252 | 0,27832 | 2,6132593 | 0,0089683 | 0,0321919 | KIT |
| ENSGALG00000000400 | 352,09264 | 0,727285624 | 0,2194005 | 3,3148775 | 0,0009168 | 0,0049811 | KCNC4 |
| ENSGALG00000013143 | 266,7402 | 0,72680287 | 0,2471882 | 2,9402819 | 0,0032791 | 0,0141417 | PDE3A |
| ENSGALG00000014910 | 3784,5308 | 0,72588259 | 0,1623113 | 4,4721633 | 7,74E-06 | 8,18E-05 | COLEC12 |
| ENSGALG00000023083 | 215,10004 | 0,725643274 | 0,2997631 | 2,4207225 | 0,0154897 | 0,0496386 | KIAA1958 |
| ENSGALG00000004875 | 3335,2739 | 0,724796191 | 0,1381226 | 5,2474846 | 1,54E-07 | 2,39E-06 | PEMT |
| ENSGALG00000015123 | 798,45006 | 0,722653298 | 0,1722336 | 4,1957734 | 2,72E-05 | 0,0002436 | ZDHHC23 |
| ENSGALG00000026864 | 531,42097 | 0,721498621 | 0,203036 | 3,5535495 | 0,0003801 | 0,0023748 | ZCCHC24 |
| ENSGALG00000011426 | 2063,2289 | 0,721230387 | 0,1613379 | 4,4703085 | 7,81E-06 | 8,24E-05 | RP11-514O12.4 |
| ENSGALG00000000723 | 8396,3268 | 0,72079059 | 0,1405098 | 5,1298237 | 2,90E-07 | 4,21E-06 | STX12 |
| ENSGALG00000008914 | 183,84781 | 0,720697485 | 0,2822547 | 2,5533588 | 0,010669 | 0,0370556 | NRAP |
| ENSGALG00000002558 | 389,45186 | 0,720642154 | 0,2290837 | 3,1457591 | 0,0016566 | 0,0080666 | ASL1 |
| ENSGALG00000001459 | 1037,3333 | 0,719793089 | 0,1787902 | 4,0259094 | 5,68E-05 | 0,0004594 | TNNC1 |
| ENSGALG00000007981 | 2067,6116 | 0,719546729 | 0,1428586 | 5,0367766 | 4,73E-07 | 6,61E-06 | CLN6 |
| ENSGALG00000009042 | 1726,6169 | 0,719468223 | 0,1493757 | 4,8165026 | 1,46E-06 | 1,83E-05 | ISM1 |
| ENSGALG00000017296 | 332,74685 | 0,719276435 | 0,2726134 | 2,6384485 | 0,0083286 | 0,0303762 | KLHL42 |
| ENSGALG00000003855 | 2094,2466 | 0,719100031 | 0,1368455 | 5,2548333 | 1,48E-07 | 2,31E-06 | SRC |
| ENSGALG00000017031 | 283,53237 | 0,718524914 | 0,2888562 | 2,4874835 | 0,012865 | 0,0429181 | TPTE2 |
| ENSGALG00000015403 | 10307,491 | 0,718054416 | 0,1317872 | 5,4485916 | 5,08E-08 | 8,65E-07 | EPHA3 |
| ENSGALG00000007383 | 515,08856 | 0,717575013 | 0,2017592 | 3,5565912 | 0,0003757 | 0,0023529 | HSPB8 |
| ENSGALG00000012792 | 5661,7981 | 0,717520705 | 0,1251036 | 5,7354139 | 9,73E-09 | 1,92E-07 | HIPK2 |
| ENSGALG00000013583 | 619,65104 | 0,717484093 | 0,2169479 | 3,3071723 | 0,0009424 | 0,0050958 | FAM114A1 |
| ENSGALG00000024012 | 315,05149 | 0,716701131 | 0,2440127 | 2,937147 | 0,0033125 | 0,0142674 | TDRKH |
| ENSGALG00000023607 | 192,83095 | 0,714467677 | 0,2785112 | 2,5653106 | 0,0103083 | 0,0360237 | ASCL2 |
| ENSGALG00000004812 | 420,51393 | 0,714148656 | 0,2276151 | 3,1375277 | 0,0017038 | 0,0082611 | FAM129A |
| ENSGALG00000003340 | 440,9962 | 0,714011033 | 0,2132487 | 3,3482552 | 0,0008132 | 0,0044953 | GBGT1 |
| ENSGALG00000001233 | 1022,3151 | 0,713516886 | 0,1751304 | 4,0742031 | 4,62E-05 | 0,0003833 | RHCE |
| ENSGALG00000026230 | 11,282047 | 0,713226116 | 0,2861696 | 2,49232 | 0,0126912 | 0,0424526 | |
| ENSGALG00000007244 | 345,61048 | 0,711922899 | 0,2206692 | 3,2261997 | 0,0012545 | 0,0064477 | SIRT4 |
| ENSGALG00000009628 | 330,24566 | 0,711918795 | 0,2307632 | 3,085062 | 0,0020351 | 0,0095809 | |
| ENSGALG00000003282 | 5419,4179 | 0,711820753 | 0,1111661 | 6,4032191 | 1,52E-10 | 4,05E-09 | STAT5B |
| ENSGALG00000007196 | 4102,6967 | 0,710478048 | 0,1280928 | 5,5465879 | 2,91E-08 | 5,23E-07 | STXBP1 |
| ENSGALG00000003805 | 6589,8039 | 0,710406685 | 0,1297333 | 5,4759004 | 4,35E-08 | 7,53E-07 | PRKG1 |
| ENSGALG00000001130 | 1241,2514 | 0,709264893 | 0,1624051 | 4,3672563 | 1,26E-05 | 0,0001259 | WBSCR17 |
| ENSGALG00000020794 | 10,814613 | 0,707249449 | 0,285562 | 2,4766932 | 0,0132606 | 0,0439509 | |
| ENSGALG00000002038 | 769,11793 | 0,706334316 | 0,1755894 | 4,0226475 | 5,75E-05 | 0,000465 | DENND2C |
| ENSGALG00000007618 | 10,281175 | 0,705775117 | 0,2855705 | 2,4714563 | 0,0134564 | 0,0444413 | BTG4 |
| ENSGALG00000028443 | 5716,4355 | 0,705249503 | 0,169055 | 4,1717175 | 3,02E-05 | 0,0002673 | CSPG5 |
| ENSGALG00000023738 | 705,32888 | 0,704601132 | 0,2001565 | 3,5202511 | 0,0004311 | 0,0026513 | |
| ENSGALG00000010320 | 1626,0227 | 0,704383823 | 0,138468 | 5,0869789 | 3,64E-07 | 5,19E-06 | PAWR |
| ENSGALG00000028685 | 205,34816 | 0,704382223 | 0,2842057 | 2,4784238 | 0,0131964 | 0,0437898 | |
| ENSGALG00000001640 | 312,33982 | 0,704326284 | 0,2358711 | 2,9860645 | 0,0028259 | 0,0125489 | RNF122 |
| ENSGALG00000000599 | 300,39451 | 0,703788264 | 0,2449165 | 2,8735847 | 0,0040584 | 0,0168977 | mars-02 |
| ENSGALG00000026437 | 5573,4884 | 0,703536871 | 0,1156883 | 6,0813142 | 1,19E-09 | 2,80E-08 | |
| ENSGALG00000004735 | 2180,5981 | 0,703327732 | 0,1488288 | 4,7257505 | 2,29E-06 | 2,73E-05 | MAP2K3 |
| ENSGALG00000026973 | 1340,8175 | 0,702510997 | 0,1463553 | 4,8000384 | 1,59E-06 | 1,97E-05 | KIF3C |
| ENSGALG00000026873 | 606,99857 | 0,701267796 | 0,1971895 | 3,5563142 | 0,0003761 | 0,0023543 | RSPO2 |
| ENSGALG00000017269 | 189,52991 | 0,70061916 | 0,2816451 | 2,4875958 | 0,012861 | 0,0429172 | USP35 |
| ENSGALG00000023266 | 1017,6993 | 0,69953934 | 0,177551 | 3,9399339 | 8,15E-05 | 0,0006287 | INPP5J |
| ENSGALG00000027058 | 1118,3528 | 0,699526976 | 0,185719 | 3,7665882 | 0,0001655 | 0,001164 | EMP2 |
| ENSGALG00000028018 | 221,75017 | 0,69904366 | 0,2851436 | 2,4515496 | 0,0142243 | 0,0463753 | RAB38 |
| ENSGALG00000020497 | 375,96471 | 0,698148301 | 0,21726 | 3,2134229 | 0,0013116 | 0,0066923 | IFITM10 |
| ENSGALG00000008902 | 238,20892 | 0,6980432 | 0,2801236 | 2,4919118 | 0,0127058 | 0,0424825 | GIPC2 |
| ENSGALG00000004647 | 2581,3028 | 0,697164089 | 0,1609537 | 4,3314572 | 1,48E-05 | 0,0001453 | C5H15ORF57 |
| ENSGALG00000015181 | 1358,5983 | 0,696928823 | 0,1980876 | 3,518286 | 0,0004343 | 0,0026634 | GAREM1 |
| ENSGALG00000016299 | 188,53968 | 0,696461514 | 0,2699353 | 2,5801055 | 0,009877 | 0,0348087 | FAM83B |
| ENSGALG00000028945 | 1325,8832 | 0,696399862 | 0,1972938 | 3,5297611 | 0,0004159 | 0,0025729 | LGI2 |
| ENSGALG00000007349 | 388,2147 | 0,696333265 | 0,2142206 | 3,2505423 | 0,0011519 | 0,0060109 | RASL12 |
| ENSGALG00000008455 | 4612,7445 | 0,695894902 | 0,1608753 | 4,3256789 | 1,52E-05 | 0,0001481 | SEMA3C |

| | | | | | | | |
|---|---|---|---|---|---|---|---|
| ENSGALG00000010094 | 8338,3892 | 0,695566561 | 0,1102371 | 6,3097346 | 2,80E-10 | 7,11E-09 | B4GALT2 |
| ENSGALG00000008250 | 259,75115 | 0,695347261 | 0,2674448 | 2,5999651 | 0,0093233 | 0,0332035 | TBX3 |
| ENSGALG00000010331 | 1143,0104 | 0,693784562 | 0,1997471 | 3,4733156 | 0,0005141 | 0,0030671 | MME |
| ENSGALG00000010406 | 684,71072 | 0,693540735 | 0,1775952 | 3,9051762 | 9,42E-05 | 0,0007122 | TMEM63C |
| ENSGALG00000001889 | 934,57066 | 0,693259058 | 0,1726852 | 4,0145828 | 5,96E-05 | 0,0004785 | CARD9 |
| ENSGALG00000008651 | 2483,2004 | 0,693028969 | 0,1577851 | 4,3922326 | 1,12E-05 | 0,0001142 | APMAP |
| ENSGALG00000005103 | 1726,7826 | 0,692645693 | 0,1383584 | 5,0061716 | 5,55E-07 | 7,63E-06 | GDPD1 |
| ENSGALG00000023188 | 975,83818 | 0,692540652 | 0,1600052 | 4,3282373 | 1,50E-05 | 0,0001471 | NANP |
| ENSGALG00000001697 | 10,047458 | 0,69224725 | 0,2838775 | 2,4385418 | 0,0147467 | 0,0477354 | ITIH3 |
| ENSGALG00000002192 | 251,47548 | 0,692054061 | 0,2782223 | 2,4874138 | 0,0128676 | 0,0429181 | PTPRC |
| ENSGALG00000003636 | 432,59045 | 0,690941222 | 0,2090885 | 3,3045388 | 0,0009513 | 0,0051317 | FBLIM1 |
| ENSGALG00000020575 | 447,18026 | 0,690014438 | 0,2210762 | 3,1211618 | 0,0018014 | 0,0086359 | ERMN |
| ENSGALG00000023655 | 726,63007 | 0,689103212 | 0,1781324 | 3,8684889 | 0,0001095 | 0,0008103 | |
| ENSGALG00000019436 | 255,33631 | 0,68869647 | 0,2721433 | 2,5306391 | 0,0113855 | 0,0389481 | MAL2 |
| ENSGALG00000011356 | 243,58893 | 0,688404159 | 0,2584311 | 2,6637818 | 0,0077268 | 0,0286811 | |
| ENSGALG00000003122 | 2359,0002 | 0,68839827 | 0,1635612 | 4,2088128 | 2,57E-05 | 0,0002327 | ZFAND2A |
| ENSGALG00000026361 | 265,71557 | 0,687466561 | 0,2597715 | 2,6464277 | 0,0081347 | 0,0297885 | SLITRK5 |
| ENSGALG00000002749 | 1561,2531 | 0,687223145 | 0,1398964 | 4,9123728 | 9,00E-07 | 1,18E-05 | RP11-565P22.6 |
| ENSGALG00000001342 | 285,4866 | 0,686942007 | 0,2446203 | 2,8081975 | 0,004982 | 0,0200146 | HCN2 |
| ENSGALG00000016844 | 1336,9691 | 0,685906484 | 0,1561619 | 4,3922785 | 1,12E-05 | 0,0001142 | RAB20 |
| ENSGALG00000016123 | 816,29325 | 0,685692574 | 0,2096562 | 3,2705578 | 0,0010734 | 0,0056686 | TRPS1 |
| ENSGALG00000002204 | 670,70196 | 0,684953864 | 0,196026 | 3,4941984 | 0,0004755 | 0,0028759 | SLC7A4 |
| ENSGALG00000009444 | 276,54744 | 0,684544481 | 0,2525059 | 2,7110044 | 0,006708 | 0,0255466 | HHIPL2 |
| ENSGALG00000014559 | 1808,9059 | 0,684328525 | 0,1507966 | 4,5380898 | 5,68E-06 | 6,17E-05 | MEF2D |
| ENSGALG00000016570 | 213,7613 | 0,684262758 | 0,2599004 | 2,6327884 | 0,0084687 | 0,0307881 | |
| ENSGALG00000009075 | 1175,3212 | 0,683878573 | 0,1540118 | 4,44043 | 8,98E-06 | 9,38E-05 | CD99L2 |
| ENSGALG00000010770 | 1614,3878 | 0,683740122 | 0,1643769 | 4,159587 | 3,19E-05 | 0,0002801 | SCIN |
| ENSGALG00000026269 | 172,06551 | 0,683345796 | 0,2809077 | 2,432635 | 0,0149894 | 0,0483604 | TAP1 |
| ENSGALG00000009523 | 834,57418 | 0,681144754 | 0,1860744 | 3,6606041 | 0,0002516 | 0,0016663 | CNTN1 |
| ENSGALG00000007251 | 3608,0883 | 0,681060329 | 0,121444 | 5,6080193 | 2,05E-08 | 3,82E-07 | PCYT2 |
| ENSGALG00000004341 | 261,34286 | 0,680949124 | 0,262149 | 2,5975655 | 0,0093887 | 0,0333927 | |
| ENSGALG00000019702 | 11617,301 | 0,678512546 | 0,132661 | 5,1146335 | 3,14E-07 | 4,54E-06 | RP4-613B23.5 |
| ENSGALG00000000674 | 418,54848 | 0,67836471 | 0,2130129 | 3,1846173 | 0,0014495 | 0,00725 | MYO1D |
| ENSGALG00000020688 | 589,72193 | 0,678030575 | 0,2132646 | 3,1792929 | 0,0014763 | 0,0073359 | CYP4B1L |
| ENSGALG00000026986 | 1312,7721 | 0,67793682 | 0,1686465 | 4,0198699 | 5,82E-05 | 0,0004694 | KIAA1324 |
| ENSGALG00000004240 | 977,7105 | 0,676367039 | 0,1738198 | 3,8911964 | 9,98E-05 | 0,0007483 | ORAI1 |
| ENSGALG00000025764 | 5164,9467 | 0,675250138 | 0,1225181 | 5,5114325 | 3,56E-08 | 6,27E-07 | FKBP10 |
| ENSGALG00000002028 | 1196,673 | 0,675138088 | 0,1985373 | 3,4005596 | 0,0006725 | 0,0038438 | GPRC5B |
| ENSGALG00000015425 | 550,05237 | 0,674783987 | 0,2346154 | 2,8761286 | 0,0040259 | 0,0167724 | LPL |
| ENSGALG00000012583 | 448,70952 | 0,674342421 | 0,2079713 | 3,242479 | 0,0011849 | 0,0061506 | IDNK |
| ENSGALG00000016471 | 262,57308 | 0,674081782 | 0,2520085 | 2,6748379 | 0,0074765 | 0,0278962 | NT5C1B |
| ENSGALG00000021345 | 271,10964 | 0,673475888 | 0,2582429 | 2,6079164 | 0,0091095 | 0,0325699 | B3GNT9 |
| ENSGALG00000016399 | 1323,2505 | 0,673456474 | 0,1609458 | 4,1843685 | 2,86E-05 | 0,000255 | SNTB1 |
| ENSGALG00000004472 | 11283,073 | 0,672299012 | 0,1222317 | 5,5002025 | 3,79E-08 | 6,65E-07 | ASTN1 |
| ENSGALG00000005041 | 1267,2443 | 0,67175632 | 0,1650486 | 4,0700516 | 4,70E-05 | 0,0003893 | TMEM35 |
| ENSGALG00000011896 | 1990,3879 | 0,671754963 | 0,1334515 | 5,0337007 | 4,81E-07 | 6,71E-06 | SIDT2 |
| ENSGALG00000007038 | 2302,528 | 0,670533206 | 0,1352237 | 4,9586965 | 7,10E-07 | 9,50E-06 | ASTN2 |
| ENSGALG00000005069 | 1045,9472 | 0,669583836 | 0,1580685 | 4,2360364 | 2,27E-05 | 0,0002096 | PTGS2 |
| ENSGALG00000006700 | 425,17548 | 0,669478691 | 0,2185126 | 3,0637986 | 0,0021855 | 0,0101553 | PROSER2 |
| ENSGALG00000000040 | 692,57517 | 0,669045681 | 0,1867751 | 3,5820926 | 0,0003409 | 0,0021706 | FAM195A |
| ENSGALG00000000920 | 727,43745 | 0,668851002 | 0,1978996 | 3,3797499 | 0,0007255 | 0,0041021 | CGN |
| ENSGALG00000016415 | 1383,1065 | 0,668618298 | 0,1866966 | 3,581309 | 0,0003419 | 0,0021744 | MAP7D2 |
| ENSGALG00000002551 | 236,59495 | 0,667886193 | 0,2539141 | 2,6303623 | 0,0085294 | 0,0309591 | TTC18 |
| ENSGALG00000016617 | 4902,5535 | 0,667497538 | 0,1287787 | 5,1832934 | 2,18E-07 | 3,27E-06 | NCOA1 |
| ENSGALG00000008970 | 735,02263 | 0,666872911 | 0,1806417 | 3,6916889 | 0,0002228 | 0,0015 | NRK |
| ENSGALG00000006449 | 910,14644 | 0,666661271 | 0,1961576 | 3,3985997 | 0,0006773 | 0,0038698 | GRIP2 |
| ENSGALG00000003736 | 312,42816 | 0,665332911 | 0,230427 | 2,8873912 | 0,0038845 | 0,0163183 | ABCB9 |
| ENSGALG00000007324 | 392,14542 | 0,66463294 | 0,2296182 | 2,8945136 | 0,0037975 | 0,0160169 | TP63 |
| ENSGALG00000003457 | 3064,2612 | 0,662964808 | 0,1431321 | 4,6318371 | 3,62E-06 | 4,14E-05 | CNP |
| ENSGALG00000013766 | 661,355 | 0,662853472 | 0,1976254 | 3,35409 | 0,0007963 | 0,0044322 | DSEL |
| ENSGALG00000008135 | 345,6592 | 0,662771473 | 0,2243246 | 2,9545197 | 0,0031316 | 0,0136305 | SATB2 |
| ENSGALG00000010316 | 7099,3915 | 0,662660688 | 0,1194525 | 5,5474833 | 2,90E-08 | 5,21E-07 | FRAS1 |
| ENSGALG00000016800 | 213,64071 | 0,662047446 | 0,2628994 | 2,5182536 | 0,0117938 | 0,0400432 | C2orf40 |
| ENSGALG00000006775 | 290,55417 | 0,661623797 | 0,267005 | 2,4779452 | 0,0132141 | 0,0438272 | TGM2 |

| | | | | | | | |
|---|---|---|---|---|---|---|---|
| ENSGALG00000002775 | 509,11695 | 0,661278251 | 0,2056466 | 3,2156052 | 0,0013017 | 0,0066526 | FA2H |
| ENSGALG00000012710 | 1279,2972 | 0,660750331 | 0,1586684 | 4,1643484 | 3,12E-05 | 0,000275 | NCS1 |
| ENSGALG00000023708 | 346,0463 | 0,660417389 | 0,226761 | 2,9123934 | 0,0035867 | 0,0152699 | UBXN10 |
| ENSGALG00000018815 | 410,19043 | 0,659828915 | 0,2173041 | 3,0364305 | 0,002394 | 0,0109266 | PCGF2 |
| ENSGALG00000006236 | 481,20913 | 0,65916962 | 0,2166125 | 3,0430824 | 0,0023417 | 0,0107419 | TPH1 |
| ENSGALG00000016411 | 1960,7449 | 0,657856223 | 0,1571317 | 4,1866556 | 2,83E-05 | 0,0002527 | COL14A1 |
| ENSGALG00000006152 | 1208,2078 | 0,657637784 | 0,1593989 | 4,125736 | 3,70E-05 | 0,0003172 | |
| ENSGALG00000025698 | 1218,1933 | 0,657354664 | 0,227163 | 2,8937581 | 0,0038066 | 0,0160455 | |
| ENSGALG00000018967 | 1911,6852 | 0,656123394 | 0,1849185 | 3,5481766 | 0,0003879 | 0,0024149 | LYN |
| ENSGALG00000016543 | 3236,9117 | 0,654951402 | 0,1701332 | 3,8496396 | 0,0001183 | 0,000864 | NHS |
| ENSGALG00000010732 | 284,29764 | 0,654838117 | 0,2441236 | 2,6824038 | 0,0073095 | 0,027378 | GALNTL6 |
| ENSGALG00000009473 | 257,7395 | 0,654742804 | 0,2591149 | 2,5268436 | 0,0115093 | 0,0392335 | EFCAB1 |
| ENSGALG00000012092 | 321,27125 | 0,654654529 | 0,2674895 | 2,4474025 | 0,014389 | 0,046778 | ESYT3 |
| ENSGALG00000012023 | 986,86948 | 0,654468757 | 0,1614892 | 4,0527095 | 5,06E-05 | 0,000416 | DACT1 |
| ENSGALG00000003845 | 2106,1901 | 0,653837192 | 0,1371393 | 4,7676855 | 1,86E-06 | 2,26E-05 | MKNK2 |
| ENSGALG00000006676 | 521,68608 | 0,652541968 | 0,2289326 | 2,8503676 | 0,0043669 | 0,0179081 | RLBP1 |
| ENSGALG00000003589 | 5654,7984 | 0,652278144 | 0,1207315 | 5,4027172 | 6,56E-08 | 1,10E-06 | VTN |
| ENSGALG00000001426 | 1341,2118 | 0,651807691 | 0,1484727 | 4,3900856 | 1,13E-05 | 0,0001152 | MECR |
| ENSGALG00000002403 | 2862,6989 | 0,651643976 | 0,1530067 | 4,2589243 | 2,05E-05 | 0,0001919 | ULK1 |
| ENSGALG00000001956 | 2538,005 | 0,650817237 | 0,1761305 | 3,6950845 | 0,0002198 | 0,001483 | OLFML3 |
| ENSGALG00000014225 | 320,24798 | 0,65009587 | 0,2560628 | 2,5388138 | 0,0111229 | 0,0382613 | KCTD8 |
| ENSGALG00000008120 | 6095,8052 | 0,649872522 | 0,1262195 | 5,1487478 | 2,62E-07 | 3,86E-06 | TLE3 |
| ENSGALG00000024104 | 1479,6002 | 0,64969951 | 0,1504929 | 4,3171453 | 1,58E-05 | 0,0001528 | |
| ENSGALG00000005145 | 354,77662 | 0,64926713 | 0,2184796 | 2,9717518 | 0,0029611 | 0,0130342 | SS18L1 |
| ENSGALG00000024488 | 3112,8373 | 0,649250333 | 0,1855727 | 3,4986306 | 0,0004677 | 0,0028349 | CD99 |
| ENSGALG00000008763 | 548,60866 | 0,64905793 | 0,2030908 | 3,1959002 | 0,001394 | 0,0070213 | SSX2IP |
| ENSGALG00000007090 | 354,84899 | 0,648812032 | 0,2646101 | 2,4519552 | 0,0142082 | 0,0463564 | CPN1 |
| ENSGALG00000008780 | 291,63944 | 0,648139707 | 0,2519098 | 2,5729041 | 0,0100849 | 0,0354064 | CTBS |
| ENSGALG00000007848 | 1247,6356 | 0,647083546 | 0,1839624 | 3,5174772 | 0,0004357 | 0,0026671 | PTS |
| ENSGALG00000010422 | 5513,2698 | 0,646628908 | 0,1246783 | 5,1863789 | 2,14E-07 | 3,22E-06 | FEZ1 |
| ENSGALG00000010719 | 1138,2128 | 0,646215963 | 0,1598376 | 4,0429522 | 5,28E-05 | 0,0004319 | NEIL3 |
| ENSGALG00000006036 | 6926,7297 | 0,644947298 | 0,1418361 | 4,547131 | 5,44E-06 | 5,94E-05 | |
| ENSGALG00000003655 | 1118,2527 | 0,643488267 | 0,1848793 | 3,4805866 | 0,0005003 | 0,0030021 | GAS2 |
| ENSGALG00000000402 | 743,97016 | 0,642116084 | 0,180432 | 3,5587699 | 0,0003726 | 0,0023388 | LOXL2 |
| ENSGALG00000006947 | 23413,053 | 0,64187784 | 0,1108573 | 5,7901257 | 7,03E-09 | 1,43E-07 | BDH1 |
| ENSGALG00000003568 | 441,24282 | 0,640936632 | 0,2094653 | 3,0598708 | 0,0022143 | 0,0102684 | PPP1R16B |
| ENSGALG00000011335 | 567,87108 | 0,64086738 | 0,205863 | 3,113077 | 0,0018515 | 0,0088263 | NHEJ1 |
| ENSGALG00000008479 | 572,78152 | 0,640276298 | 0,2217437 | 2,8874605 | 0,0038837 | 0,0163183 | PLEKHM3 |
| ENSGALG00000026159 | 2624,117 | 0,639768169 | 0,1366904 | 4,6804192 | 2,86E-06 | 3,34E-05 | PDLIM7 |
| ENSGALG00000004074 | 504,98036 | 0,63919781 | 0,2089342 | 3,0593254 | 0,0022184 | 0,0102792 | KCNS1 |
| ENSGALG00000026295 | 3751,1713 | 0,638993968 | 0,1256122 | 5,0870391 | 3,64E-07 | 5,19E-06 | CDR2L |
| ENSGALG00000000935 | 1038,8243 | 0,638135684 | 0,1951826 | 3,2694289 | 0,0010776 | 0,0056891 | TUFT1 |
| ENSGALG00000009478 | 877,18337 | 0,637933371 | 0,2202594 | 2,8962816 | 0,0037761 | 0,0159566 | BCHE |
| ENSGALG00000028506 | 1185,606 | 0,637581477 | 0,166917 | 3,8197523 | 0,0001336 | 0,0009649 | |
| ENSGALG00000003527 | 1639,1106 | 0,636215476 | 0,1470878 | 4,3254142 | 1,52E-05 | 0,0001482 | ARRDC2 |
| ENSGALG00000028756 | 1081,2256 | 0,636133999 | 0,1663302 | 3,8245244 | 0,000131 | 0,0009474 | SLC4A2 |
| ENSGALG00000004585 | 1609,3421 | 0,634633954 | 0,1652199 | 3,841147 | 0,0001225 | 0,0008907 | SLC29A3 |
| ENSGALG00000020726 | 246,91021 | 0,63266925 | 0,2559148 | 2,4721867 | 0,0134289 | 0,0443774 | PDZD7 |
| ENSGALG00000008057 | 2849,9643 | 0,63255389 | 0,1469667 | 4,3040624 | 1,68E-05 | 0,0001609 | LLGL2 |
| ENSGALG00000011053 | 10065,657 | 0,630484727 | 0,107695 | 5,8543543 | 4,79E-09 | 1,01E-07 | HOXA2 |
| ENSGALG00000008151 | 5862,8164 | 0,629127593 | 0,1186504 | 5,3023621 | 1,14E-07 | 1,81E-06 | WSB2 |
| ENSGALG00000027658 | 594,25398 | 0,628365542 | 0,1906089 | 3,2966228 | 0,0009785 | 0,005237 | TESK1 |
| ENSGALG00000002664 | 1165,5467 | 0,627944771 | 0,1531652 | 4,0997864 | 4,14E-05 | 0,0003505 | ANXA11 |
| ENSGALG00000029000 | 1142,4251 | 0,627547484 | 0,158422 | 3,9612397 | 7,46E-05 | 0,0005808 | HYAL3 |
| ENSGALG00000028204 | 8948,3531 | 0,626531339 | 0,1218206 | 5,1430676 | 2,70E-07 | 3,97E-06 | GPX1 |
| ENSGALG00000001100 | 796,40837 | 0,626475621 | 0,1802096 | 3,4763714 | 0,0005082 | 0,0030393 | ST3GAL4 |
| ENSGALG00000002313 | 2718,417 | 0,626374122 | 0,1346423 | 4,6521335 | 3,29E-06 | 3,79E-05 | SLC38A7 |
| ENSGALG00000016745 | 455,542 | 0,625941712 | 0,2478958 | 2,5250189 | 0,0115692 | 0,0393984 | GABRB3 |
| ENSGALG00000026762 | 1809,4015 | 0,625670937 | 0,1367941 | 4,5738152 | 4,79E-06 | 5,34E-05 | WBSCR27 |
| ENSGALG00000008728 | 894,54762 | 0,625207283 | 0,1707065 | 3,6624697 | 0,0002498 | 0,0016558 | PTER |
| ENSGALG00000011856 | 1033,6518 | 0,624422163 | 0,1575632 | 3,9629959 | 7,40E-05 | 0,0005778 | C7orf55 |
| ENSGALG00000007267 | 4451,8628 | 0,623851268 | 0,1170233 | 5,3309986 | 9,77E-08 | 1,57E-06 | MVD |
| ENSGALG00000014890 | 1413,3331 | 0,620763059 | 0,1462687 | 4,2439922 | 2,20E-05 | 0,0002031 | TNFRSF1A |
| ENSGALG00000000426 | 554,75724 | 0,620684473 | 0,1991349 | 3,1169053 | 0,0018276 | 0,0087369 | IGSF21 |

| | | | | | | | |
|---|---|---|---|---|---|---|---|
| ENSGALG00000006114 | 909,8822 | 0,61972927 | 0,1710742 | 3,6225768 | 0,0002917 | 0,0018965 | |
| ENSGALG00000016595 | 534,98984 | 0,618865991 | 0,2042 | 3,0306858 | 0,00244 | 0,0111031 | TRIM35 |
| ENSGALG00000011900 | 1365,0771 | 0,61861917 | 0,1798125 | 3,440356 | 0,0005809 | 0,0033976 | NAGA |
| ENSGALG00000007076 | 2564,3514 | 0,617550884 | 0,1437626 | 4,2956288 | 1,74E-05 | 0,0001663 | CCNY |
| ENSGALG00000016934 | 244,09661 | 0,616949085 | 0,2531836 | 2,4367659 | 0,0148193 | 0,0479038 | PCDH20 |
| ENSGALG00000014561 | 290,72868 | 0,616417234 | 0,2372422 | 2,5982612 | 0,0093697 | 0,0333426 | PTPN6 |
| ENSGALG00000007647 | 329,01065 | 0,615818159 | 0,2520071 | 2,443654 | 0,0145394 | 0,0471767 | MYO3A |
| ENSGALG00000007381 | 1220,4425 | 0,615500074 | 0,1553201 | 3,9627846 | 7,41E-05 | 0,000578 | SERPING1 |
| ENSGALG00000028543 | 1301,4945 | 0,614319385 | 0,1647996 | 3,7276752 | 0,0001933 | 0,0013256 | FGFR4 |
| ENSGALG00000015734 | 903,95178 | 0,61429478 | 0,1983985 | 3,0962669 | 0,0019597 | 0,0092839 | ZC2HC1A |
| ENSGALG00000016728 | 5650,4017 | 0,613362339 | 0,115156 | 5,3263591 | 1,00E-07 | 1,61E-06 | RP2 |
| ENSGALG00000023881 | 330,14663 | 0,612305207 | 0,2457805 | 2,4912681 | 0,0127288 | 0,0425491 | PLXDC1 |
| ENSGALG00000016092 | 954,71804 | 0,611645343 | 0,1643983 | 3,720508 | 0,0001988 | 0,0013569 | RAB11FIP5 |
| ENSGALG00000011975 | 2830,8254 | 0,61116717 | 0,1227563 | 4,9787019 | 6,40E-07 | 8,65E-06 | FBXL2 |
| ENSGALG00000014699 | 806,27033 | 0,610005378 | 0,2334147 | 2,6133976 | 0,0089647 | 0,0321873 | MEGF10 |
| ENSGALG00000012495 | 822,21166 | 0,608707981 | 0,2014447 | 3,0217131 | 0,0025135 | 0,0113541 | Q9DEH4_CHICK |
| ENSGALG00000016143 | 5447,091 | 0,607480677 | 0,1713253 | 3,5457739 | 0,0003915 | 0,0024359 | RHPN1 |
| ENSGALG00000004103 | 351,09074 | 0,607102605 | 0,2488829 | 2,4393098 | 0,0147153 | 0,0476455 | FAM81A |
| ENSGALG00000008512 | 665,49717 | 0,606896507 | 0,1931945 | 3,141375 | 0,0016816 | 0,0081679 | GRIA3 |
| ENSGALG00000008272 | 1496,8242 | 0,6063144 | 0,1570984 | 3,8594554 | 0,0001136 | 0,0008363 | TMED3 |
| ENSGALG00000012934 | 686,11794 | 0,606004522 | 0,2092753 | 2,8957294 | 0,0037828 | 0,0159748 | BICD1 |
| ENSGALG00000016071 | 698,48269 | 0,605474788 | 0,1980237 | 3,0575882 | 0,0022313 | 0,0103145 | CTHRC1 |
| ENSGALG00000015414 | 1285,1197 | 0,605396806 | 0,1804212 | 3,3554643 | 0,0007923 | 0,0044138 | NSUN3 |
| ENSGALG00000002930 | 5292,4924 | 0,603605896 | 0,1376656 | 4,3845818 | 1,16E-05 | 0,0001178 | STX7 |
| ENSGALG00000005303 | 729,47439 | 0,603106981 | 0,1827363 | 3,3004217 | 0,0009654 | 0,005187 | CCDC13 |
| ENSGALG00000006742 | 8579,7495 | 0,60288418 | 0,1075096 | 5,6077227 | 2,05E-08 | 3,82E-07 | NECTIN1 |
| ENSGALG00000013964 | 1158,3447 | 0,602864404 | 0,163189 | 3,6942711 | 0,0002205 | 0,0014863 | SPATA18 |
| ENSGALG00000015075 | 452,11233 | 0,602809512 | 0,2158206 | 2,7931043 | 0,0052205 | 0,0207767 | |
| ENSGALG00000000608 | 482,47931 | 0,60269407 | 0,1990068 | 3,0285101 | 0,0024576 | 0,0111684 | CDH1 |
| ENSGALG00000000739 | 566,84692 | 0,602546305 | 0,188602 | 3,1948037 | 0,0013993 | 0,0070455 | XKR8 |
| ENSGALG00000009047 | 3574,1219 | 0,601534497 | 0,1392573 | 4,3195906 | 1,56E-05 | 0,0001517 | YPEL5 |
| ENSGALG00000005628 | 1589,6542 | 0,601333554 | 0,1810639 | 3,3211128 | 0,0008966 | 0,0048849 | COL9A3 |
| ENSGALG00000029021 | 4810,2675 | 0,600451025 | 0,1243158 | 4,830046 | 1,37E-06 | 1,72E-05 | CACNA2D2 |
| ENSGALG00000023435 | 807,96294 | 0,600270173 | 0,1899681 | 3,1598469 | 0,0015785 | 0,0077671 | GATM |
| ENSGALG00000028023 | 391,94362 | 0,598578703 | 0,2158662 | 2,7729159 | 0,0055556 | 0,0219007 | C14ORF166B |
| ENSGALG00000023876 | 5804,1712 | 0,598475769 | 0,1258285 | 4,7562796 | 1,97E-06 | 2,38E-05 | PRKCQ |
| ENSGALG00000016330 | 507,71021 | 0,598164421 | 0,2008484 | 2,9781891 | 0,0028996 | 0,0128091 | ANGPT2 |
| ENSGALG00000008475 | 1732,9827 | 0,598095279 | 0,145561 | 4,1088979 | 3,98E-05 | 0,0003394 | ARRDC1 |
| ENSGALG00000013167 | 14635,204 | 0,597899753 | 0,1179107 | 5,0707829 | 3,96E-07 | 5,61E-06 | SDHA |
| ENSGALG00000007801 | 1966,8829 | 0,597201094 | 0,1460776 | 4,0882463 | 4,35E-05 | 0,0003633 | CORO7 |
| ENSGALG00000027450 | 1958,1012 | 0,596865465 | 0,1360214 | 4,3880272 | 1,14E-05 | 0,000116 | |
| ENSGALG00000005839 | 281,11607 | 0,595553706 | 0,2359627 | 2,5239319 | 0,011605 | 0,0394908 | ARID3C |
| ENSGALG00000002818 | 12498,951 | 0,595523293 | 0,1252741 | 4,7537617 | 2,00E-06 | 2,41E-05 | VAT1 |
| ENSGALG00000015399 | 371,45534 | 0,595444245 | 0,2198201 | 2,7087794 | 0,0067531 | 0,0256826 | FUT10 |
| ENSGALG00000008285 | 2205,3966 | 0,594659875 | 0,1335709 | 4,4520161 | 8,51E-06 | 8,94E-05 | NAPEPLD |
| ENSGALG00000026756 | 608,70147 | 0,593058479 | 0,1859613 | 3,1891498 | 0,0014269 | 0,0071635 | |
| ENSGALG00000022661 | 267,36767 | 0,593055191 | 0,2440949 | 2,429609 | 0,0151151 | 0,0486392 | |
| ENSGALG00000011145 | 3513,1587 | 0,5922185 | 0,1195159 | 4,9551454 | 7,23E-07 | 9,63E-06 | TRIL |
| ENSGALG00000026180 | 4122,3044 | 0,591162305 | 0,1425421 | 4,1472816 | 3,36E-05 | 0,0002933 | MSX2 |
| ENSGALG00000005999 | 5083,9087 | 0,591144605 | 0,1218787 | 4,8502694 | 1,23E-06 | 1,57E-05 | ABTB1 |
| ENSGALG00000004596 | 540,83556 | 0,59090413 | 0,1964513 | 3,0078909 | 0,0026307 | 0,0118051 | AR |
| ENSGALG00000013436 | 9895,6737 | 0,590008237 | 0,1280495 | 4,6076575 | 4,07E-06 | 4,58E-05 | TET3 |
| ENSGALG00000013742 | 6102,599 | 0,589905166 | 0,1472857 | 4,0051753 | 6,20E-05 | 0,0004943 | EZR |
| ENSGALG00000025745 | 2020,2184 | 0,589609597 | 0,1655708 | 3,5610734 | 0,0003693 | 0,0023216 | ARHGAP27 |
| ENSGALG00000026081 | 3861,3662 | 0,588777417 | 0,1223416 | 4,8125694 | 1,49E-06 | 1,86E-05 | RBM38 |
| ENSGALG00000013882 | 324,83067 | 0,587886739 | 0,2342948 | 2,5091749 | 0,0121014 | 0,0408428 | MXD1 |
| ENSGALG00000000464 | 6973,1204 | 0,587838297 | 0,1100043 | 5,3437778 | 9,10E-08 | 1,48E-06 | RHOBTB2 |
| ENSGALG00000004900 | 814,58968 | 0,587812156 | 0,240363 | 2,4455181 | 0,0144644 | 0,0469671 | FAM101B |
| ENSGALG00000005102 | 2254,4611 | 0,58732124 | 0,1676607 | 3,503035 | 0,00046 | 0,0027934 | CDH4 |
| ENSGALG00000016594 | 1080,5684 | 0,58674351 | 0,1703086 | 3,4451783 | 0,0005707 | 0,0033462 | FRMPD4 |
| ENSGALG00000010934 | 813,60176 | 0,586239182 | 0,1706421 | 3,4354907 | 0,0005915 | 0,0034503 | INADL |
| ENSGALG00000025822 | 1186,6239 | 0,586217768 | 0,1900116 | 3,0851675 | 0,0020344 | 0,0095808 | CYP1B1 |
| ENSGALG00000017558 | 835,37967 | 0,585776262 | 0,1728652 | 3,3886308 | 0,0007024 | 0,0039883 | |
| ENSGALG00000000407 | 552,43307 | 0,58498369 | 0,2033571 | 2,876632 | 0,0040194 | 0,0167507 | |

| | | | | | | | |
|---|---|---|---|---|---|---|---|
| ENSGALG00000002169 | 5307,0492 | 0,58477373 | 0,1393007 | 4,1979236 | 2,69E-05 | 0,0002424 | PLEKHB2 |
| ENSGALG00000016108 | 377,17299 | 0,58441259 | 0,2150799 | 2,7171887 | 0,0065839 | 0,0251303 | SYBU |
| ENSGALG00000025738 | 2313,9355 | 0,584187633 | 0,1378228 | 4,2386855 | 2,25E-05 | 0,0002075 | RHOU |
| ENSGALG00000028942 | 551,36062 | 0,584183813 | 0,2020534 | 2,8912352 | 0,0038373 | 0,0161649 | ZBTB25 |
| ENSGALG00000015938 | 2888,9851 | 0,583396297 | 0,1447547 | 4,0302413 | 5,57E-05 | 0,0004524 | IFNAR2 |
| ENSGALG00000001574 | 3769,6924 | 0,583342794 | 0,1296384 | 4,499767 | 6,80E-06 | 7,24E-05 | MLLT6 |
| ENSGALG00000014242 | 1647,3047 | 0,583190799 | 0,1556527 | 3,7467429 | 0,0001791 | 0,0012457 | BEND4 |
| ENSGALG00000008999 | 3276,8472 | 0,583129446 | 0,1551657 | 3,7581073 | 0,0001712 | 0,001199 | AFAP1L2 |
| ENSGALG00000010706 | 6618,9625 | 0,582943162 | 0,1129494 | 5,161101 | 2,46E-07 | 3,64E-06 | TENM3 |
| ENSGALG00000015952 | 611,25698 | 0,582757747 | 0,2345109 | 2,4849925 | 0,0129554 | 0,0431161 | B3GAT2 |
| ENSGALG00000004844 | 6201,28 | 0,581688828 | 0,1476751 | 3,9389777 | 8,18E-05 | 0,0006305 | SEMA6D |
| ENSGALG00000009604 | 5450,2438 | 0,581479374 | 0,1323792 | 4,392529 | 1,12E-05 | 0,0001142 | DBX2 |
| ENSGALG00000001446 | 6585,1604 | 0,581431635 | 0,1501068 | 3,8734532 | 0,0001073 | 0,0007966 | GSN |
| ENSGALG00000023768 | 4728,0275 | 0,58136874 | 0,1164642 | 4,9918245 | 5,98E-07 | 8,14E-06 | EPHA2 |
| ENSGALG00000002707 | 937,59842 | 0,58133458 | 0,1802757 | 3,2246966 | 0,0012611 | 0,0064766 | CHRNB2 |
| ENSGALG00000029020 | 1577,3992 | 0,581217481 | 0,1719893 | 3,3793821 | 0,0007265 | 0,0041059 | PPAPDC1A |
| ENSGALG00000028478 | 1030,9127 | 0,580829913 | 0,1824278 | 3,1838898 | 0,0014531 | 0,0072578 | CELF3 |
| ENSGALG00000007531 | 904,34237 | 0,57752222 | 0,1731776 | 3,3348561 | 0,0008534 | 0,0046815 | ZFYVE27 |
| ENSGALG00000006054 | 759,82179 | 0,576882549 | 0,1820676 | 3,1685082 | 0,0015322 | 0,0075694 | CALCA |
| ENSGALG00000022926 | 4686,6657 | 0,576442566 | 0,1261533 | 4,569381 | 4,89E-06 | 5,45E-05 | GPR162 |
| ENSGALG00000013185 | 3965,0445 | 0,575558846 | 0,1485635 | 3,8741602 | 0,000107 | 0,0007965 | PITPNM1 |
| ENSGALG00000000869 | 855,33376 | 0,574690811 | 0,1735124 | 3,3121031 | 0,000926 | 0,0050248 | ZDHHC18 |
| ENSGALG00000006885 | 6383,5426 | 0,574664846 | 0,1264083 | 4,5461019 | 5,46E-06 | 5,96E-05 | HRAS |
| ENSGALG00000017150 | 518,06641 | 0,574401936 | 0,1974777 | 2,9086925 | 0,0036294 | 0,0154277 | ARHGAP20 |
| ENSGALG00000015110 | 438,79213 | 0,573807714 | 0,2198442 | 2,6100658 | 0,0090525 | 0,0324512 | FAM189A2 |
| ENSGALG00000015136 | 2417,8066 | 0,573614313 | 0,1609296 | 3,5643815 | 0,0003647 | 0,0022957 | ILDR1 |
| ENSGALG00000015744 | 3529,1332 | 0,573305804 | 0,1255432 | 4,5666009 | 4,96E-06 | 5,50E-05 | RP11-26J3.4 |
| ENSGALG00000001489 | 8334,3262 | 0,572451499 | 0,129456 | 4,4219762 | 9,78E-06 | 0,000101 | SSU72 |
| ENSGALG00000014911 | 1071,9721 | 0,572338387 | 0,1572272 | 3,6402005 | 0,0002724 | 0,0017824 | ARL15 |
| ENSGALG00000015351 | 2719,2778 | 0,572206068 | 0,1704231 | 3,357561 | 0,0007863 | 0,0043931 | CBLB |
| ENSGALG00000025818 | 8546,9056 | 0,571475138 | 0,109953 | 5,1974483 | 2,02E-07 | 3,06E-06 | KCNK5 |
| ENSGALG00000005549 | 1034,7545 | 0,569387138 | 0,1561531 | 3,6463379 | 0,000266 | 0,0017479 | CUEDC1 |
| ENSGALG00000024153 | 1062,2502 | 0,56872295 | 0,170972 | 3,326409 | 0,0008797 | 0,0048083 | NLRX1 |
| ENSGALG00000000186 | 787,09326 | 0,568415238 | 0,1771285 | 3,2090555 | 0,0013317 | 0,006773 | MTCH1 |
| ENSGALG00000001146 | 1103,9797 | 0,568231367 | 0,1686686 | 3,3689227 | 0,0007546 | 0,0042333 | UBXN6 |
| ENSGALG00000026471 | 418,38308 | 0,568003242 | 0,2176763 | 2,6093938 | 0,0090703 | 0,0324722 | CRIP2 |
| ENSGALG00000009403 | 580,12466 | 0,567910527 | 0,2061904 | 2,7543016 | 0,0058818 | 0,02288 | GRK5 |
| ENSGALG00000012788 | 1309,5248 | 0,567748724 | 0,1613363 | 3,5190387 | 0,0004331 | 0,0026598 | PARP12 |
| ENSGALG00000026703 | 524,51341 | 0,567716959 | 0,1939681 | 2,926857 | 0,0034241 | 0,0146738 | SLC2A4RG |
| ENSGALG00000027275 | 1081,3036 | 0,567504466 | 0,2139281 | 2,6527819 | 0,0079831 | 0,0293917 | PPP1R3B |
| ENSGALG00000008682 | 380,85967 | 0,566238069 | 0,2325764 | 2,4346325 | 0,0149069 | 0,0481399 | SLC25A43 |
| ENSGALG00000009880 | 899,08782 | 0,565731325 | 0,2142452 | 2,6405788 | 0,0082765 | 0,0302345 | INPP4B |
| ENSGALG00000006873 | 2079,2766 | 0,565449117 | 0,1310751 | 4,3139312 | 1,60E-05 | 0,0001546 | RASSF7 |
| ENSGALG00000007358 | 319,01574 | 0,565211229 | 0,2295584 | 2,462168 | 0,01381 | 0,0452631 | CILP |
| ENSGALG00000007093 | 1910,9478 | 0,563647311 | 0,1531316 | 3,6808026 | 0,0002325 | 0,0015558 | CYTH1 |
| ENSGALG00000002778 | 10757,974 | 0,563240809 | 0,1099995 | 5,1203932 | 3,05E-07 | 4,42E-06 | C6orf106 |
| ENSGALG00000028605 | 2035,8396 | 0,561167945 | 0,1578097 | 3,5559782 | 0,0003766 | 0,0023562 | CRELD1 |
| ENSGALG00000001478 | 436,63763 | 0,560376671 | 0,2038738 | 2,7486445 | 0,0059842 | 0,0232189 | |
| ENSGALG00000002681 | 1896,5953 | 0,559637304 | 0,1492264 | 3,750257 | 0,0001767 | 0,0012328 | HSD17B7 |
| ENSGALG00000008416 | 491,34771 | 0,559354461 | 0,2224807 | 2,5141709 | 0,0119313 | 0,040409 | CFAP61 |
| ENSGALG00000016152 | 7208,3385 | 0,558712272 | 0,1259225 | 4,4369523 | 9,12E-06 | 9,51E-05 | LY6E |
| ENSGALG00000021079 | 356,52842 | 0,558275751 | 0,2206598 | 2,5300293 | 0,0114053 | 0,0389865 | RGS19 |
| ENSGALG00000010307 | 1309,92 | 0,558082008 | 0,160225 | 3,4831137 | 0,0004956 | 0,0029792 | ACYP1 |
| ENSGALG00000007997 | 607,00115 | 0,557958539 | 0,1934786 | 2,883826 | 0,0039288 | 0,0164585 | ACCS |
| ENSGALG00000011604 | 1003,2775 | 0,557581352 | 0,1841576 | 3,0277409 | 0,0024639 | 0,0111932 | SLC4A4 |
| ENSGALG00000010614 | 5022,3744 | 0,557460992 | 0,1255639 | 4,4396588 | 9,01E-06 | 9,41E-05 | SLC25A4 |
| ENSGALG00000001682 | 1516,3508 | 0,557286636 | 0,150105 | 3,712645 | 0,0002051 | 0,0013949 | FBXL20 |
| ENSGALG00000001238 | 3616,0144 | 0,556401234 | 0,1285764 | 4,3273971 | 1,51E-05 | 0,0001475 | STX8 |
| ENSGALG00000002286 | 73998,022 | 0,555324016 | 0,1110271 | 5,0016995 | 5,68E-07 | 7,79E-06 | H3F3C |
| ENSGALG00000025787 | 2442,8908 | 0,554206894 | 0,1300356 | 4,2619625 | 2,03E-05 | 0,0001895 | CDK5RAP3 |
| ENSGALG00000004849 | 4017,8273 | 0,553445556 | 0,1159402 | 4,7735421 | 1,81E-06 | 2,21E-05 | WSCD2 |
| ENSGALG00000013929 | 1086,8441 | 0,552883419 | 0,2064792 | 2,6776709 | 0,0074136 | 0,0277069 | PDGFRA |
| ENSGALG00000001766 | 1421,2057 | 0,552758547 | 0,1737186 | 3,1819197 | 0,001463 | 0,0072892 | RNF165 |
| ENSGALG00000005377 | 707,88046 | 0,551995057 | 0,1867603 | 2,9556343 | 0,0031203 | 0,0135944 | WWOX |

| | | | | | | | |
|---|---|---|---|---|---|---|---|
| ENSGALG00000023025 | 647,83568 | 0,551042888 | 0,1904818 | 2,8928903 | 0,0038171 | 0,0160849 | ZBTB42 |
| ENSGALG00000010982 | 4422,5508 | 0,55035484 | 0,1393577 | 3,9492248 | 7,84E-05 | 0,0006062 | NID1 |
| ENSGALG00000000316 | 1699,5534 | 0,549437781 | 0,1376171 | 3,9925111 | 6,54E-05 | 0,0005177 | PKNOX2 |
| ENSGALG00000027194 | 1293,5716 | 0,548094206 | 0,1517129 | 3,6127065 | 0,000303 | 0,001959 | PMVK |
| ENSGALG00000016166 | 3894,3455 | 0,547970517 | 0,1393864 | 3,9313064 | 8,45E-05 | 0,0006489 | SLC45A4 |
| ENSGALG00000006663 | 9717,8185 | 0,547489923 | 0,1171093 | 4,6750332 | 2,94E-06 | 3,41E-05 | ARHGEF12 |
| ENSGALG00000009092 | 736,78214 | 0,546285921 | 0,1858557 | 2,9393012 | 0,0032895 | 0,0141776 | GTF2A1L |
| ENSGALG00000001966 | 2313,195 | 0,545673399 | 0,1456394 | 3,7467418 | 0,0001791 | 0,0012457 | RP11-106M3.5 |
| ENSGALG00000001864 | 1163,993 | 0,545343287 | 0,1686132 | 3,2342865 | 0,0012195 | 0,006301 | C9orf72 |
| ENSGALG00000003201 | 647,35709 | 0,545148693 | 0,199247 | 2,7360447 | 0,0062183 | 0,0239563 | RP11-5A19.5 |
| ENSGALG00000006598 | 2327,6663 | 0,544504824 | 0,1375901 | 3,9574412 | 7,58E-05 | 0,0005879 | SORL1 |
| ENSGALG00000004272 | 762,39975 | 0,543009602 | 0,1799766 | 3,017113 | 0,0025519 | 0,0114973 | ASB1 |
| ENSGALG00000015107 | 1737,1601 | 0,542439326 | 0,135653 | 3,9987266 | 6,37E-05 | 0,0005065 | PIP5K1B |
| ENSGALG00000008860 | 2251,9145 | 0,541261109 | 0,1571898 | 3,4433603 | 0,0005745 | 0,0033644 | LONRF3 |
| ENSGALG00000004438 | 432,7024 | 0,540948656 | 0,2158033 | 2,5066749 | 0,0121873 | 0,0410717 | ARSB |
| ENSGALG00000005315 | 1118,2152 | 0,540760838 | 0,1598616 | 3,3826823 | 0,0007178 | 0,0040636 | RPUSD1 |
| ENSGALG00000004033 | 1001,8846 | 0,540669975 | 0,1647104 | 3,2825495 | 0,0010287 | 0,0054647 | GALNT10 |
| ENSGALG00000016176 | 404,69822 | 0,540238641 | 0,2176628 | 2,4819979 | 0,0130648 | 0,0434589 | BAI3 |
| ENSGALG00000022713 | 4245,7427 | 0,539793648 | 0,1498998 | 3,6010297 | 0,000317 | 0,0020356 | FAM181B |
| ENSGALG00000025800 | 976,78159 | 0,538353026 | 0,1626794 | 3,3092876 | 0,0009353 | 0,0050615 | |
| ENSGALG00000028513 | 5510,2985 | 0,53711532 | 0,1123093 | 4,7824632 | 1,73E-06 | 2,12E-05 | TMUB1 |
| ENSGALG00000026466 | 809,72654 | 0,537047161 | 0,1854082 | 2,8965663 | 0,0037727 | 0,015947 | |
| ENSGALG00000015437 | 3953,661 | 0,536831128 | 0,1328102 | 4,0420929 | 5,30E-05 | 0,0004329 | MPC2 |
| ENSGALG00000024049 | 660,99489 | 0,536773384 | 0,1891289 | 2,8381347 | 0,0045378 | 0,0184744 | RAB33A |
| ENSGALG00000029027 | 627,86616 | 0,53597017 | 0,182674 | 2,9340251 | 0,003346 | 0,0143844 | |
| ENSGALG00000007389 | 2084,8413 | 0,53455468 | 0,1327207 | 4,0276668 | 5,63E-05 | 0,0004565 | ANKRD40 |
| ENSGALG00000021313 | 982,25761 | 0,534412472 | 0,180088 | 2,9675073 | 0,0030023 | 0,0131729 | PDGFRB |
| ENSGALG00000016037 | 768,10735 | 0,532870331 | 0,1779289 | 2,9948499 | 0,0027458 | 0,0122611 | SPAG1 |
| ENSGALG00000009805 | 563,53778 | 0,53211751 | 0,1982743 | 2,6837437 | 0,0072803 | 0,0272932 | PPM1H |
| ENSGALG00000026039 | 442,77163 | 0,531809088 | 0,2097706 | 2,5351933 | 0,0112385 | 0,0385811 | |
| ENSGALG00000016167 | 2827,4394 | 0,530005093 | 0,1278741 | 4,1447408 | 3,40E-05 | 0,0002958 | DENND3 |
| ENSGALG00000027498 | 3720,6236 | 0,529700401 | 0,1865582 | 2,8393312 | 0,0045208 | 0,0184174 | FOXB1 |
| ENSGALG00000023189 | 809,88073 | 0,529094048 | 0,1674238 | 3,1602079 | 0,0015766 | 0,0077603 | UMAD1 |
| ENSGALG00000009433 | 12972,3 | 0,528259009 | 0,1367266 | 3,8636145 | 0,0001117 | 0,0008235 | BAG3 |
| ENSGALG00000011350 | 565,5219 | 0,52709173 | 0,1912783 | 2,7556272 | 0,005858 | 0,0228071 | NEGR1 |
| ENSGALG00000015372 | 6118,1225 | 0,526646329 | 0,1098598 | 4,7938042 | 1,64E-06 | 2,03E-05 | ATP5I |
| ENSGALG00000024346 | 1302,1617 | 0,525612903 | 0,1486838 | 3,5351054 | 0,0004076 | 0,0025272 | FAM46B |
| ENSGALG00000013839 | 706,87354 | 0,524875259 | 0,1772725 | 2,9608384 | 0,003068 | 0,0134226 | |
| ENSGALG00000015306 | 3107,5418 | 0,524586766 | 0,1467809 | 3,5739432 | 0,0003516 | 0,0022279 | OSTM1 |
| ENSGALG00000026545 | 5534,2449 | 0,524540794 | 0,126285 | 4,1536273 | 3,27E-05 | 0,000286 | |
| ENSGALG00000002777 | 12410,374 | 0,524514209 | 0,1049816 | 4,996249 | 5,85E-07 | 7,98E-06 | ACAA2 |
| ENSGALG00000015326 | 10708,875 | 0,524363909 | 0,1179079 | 4,4472344 | 8,70E-06 | 9,12E-05 | CLTA |
| ENSGALG00000014846 | 1408,263 | 0,524315684 | 0,1475731 | 3,5529214 | 0,000381 | 0,0023794 | OXCT1 |
| ENSGALG00000011945 | 1101,0533 | 0,524157211 | 0,1675307 | 3,1287225 | 0,0017557 | 0,0084616 | SCRN2 |
| ENSGALG00000011349 | 1290,6252 | 0,523508946 | 0,2028294 | 2,5810314 | 0,0098506 | 0,0347594 | MNR2 |
| ENSGALG00000006901 | 2767,4012 | 0,523326668 | 0,1242839 | 4,2107361 | 2,55E-05 | 0,0002309 | AATK |
| ENSGALG00000026113 | 4499,0711 | 0,523056553 | 0,1465805 | 3,5683905 | 0,0003592 | 0,0022661 | CARHSP1 |
| ENSGALG00000000433 | 145504,42 | 0,523011809 | 0,1027427 | 5,0905032 | 3,57E-07 | 5,11E-06 | TUBA4A |
| ENSGALG00000016886 | 6214,4284 | 0,522545779 | 0,1102341 | 4,7403267 | 2,13E-06 | 2,56E-05 | FARP1 |
| ENSGALG00000014321 | 925,70397 | 0,522295082 | 0,1727289 | 3,0237853 | 0,0024963 | 0,0113033 | SMIM14 |
| ENSGALG00000008367 | 12605,915 | 0,522250337 | 0,1198719 | 4,3567381 | 1,32E-05 | 0,0001314 | MDK |
| ENSGALG00000023812 | 573,55559 | 0,520869219 | 0,1910734 | 2,7260163 | 0,0064104 | 0,024599 | CCNI2 |
| ENSGALG00000008911 | 1167,8871 | 0,520705327 | 0,1637362 | 3,1801469 | 0,001472 | 0,0073224 | RNF208 |
| ENSGALG00000023338 | 5372,4298 | 0,520605081 | 0,1279451 | 4,0689734 | 4,72E-05 | 0,0003908 | CBX2 |
| ENSGALG00000016266 | 844,50002 | 0,519693116 | 0,1718383 | 3,0243148 | 0,002492 | 0,0112906 | PRRG1 |
| ENSGALG00000027069 | 549,10808 | 0,519660528 | 0,2040614 | 2,5465893 | 0,0108781 | 0,037619 | TSC22D3 |
| ENSGALG00000001944 | 1567,5072 | 0,519218499 | 0,1502243 | 3,4562878 | 0,0005477 | 0,0032349 | TBC1D2 |
| ENSGALG00000000059 | 125389,82 | 0,519152064 | 0,1205209 | 4,307569 | 1,65E-05 | 0,0001587 | TUBB3 |
| ENSGALG00000004775 | 978,22061 | 0,519049003 | 0,1721556 | 3,0149984 | 0,0025698 | 0,0115624 | COLGALT2 |
| ENSGALG00000005346 | 2351,8167 | 0,518505546 | 0,1295966 | 4,0009207 | 6,31E-05 | 0,0005021 | PPIC |
| ENSGALG00000026560 | 1526,6551 | 0,518030334 | 0,1900719 | 2,7254446 | 0,0064215 | 0,0246347 | PRDM8 |
| ENSGALG00000028033 | 2016,6076 | 0,517359519 | 0,1337773 | 3,8673194 | 0,00011 | 0,0008129 | PURA |
| ENSGALG00000026891 | 1109,4806 | 0,516642503 | 0,1835376 | 2,814914 | 0,004879 | 0,0196824 | NINJ1 |
| ENSGALG00000013024 | 1376,2985 | 0,516485359 | 0,1557483 | 3,3161536 | 0,0009127 | 0,0049604 | IL17RA |

| | | | | | | | |
|---|---|---|---|---|---|---|---|
| ENSGALG00000028567 | 2485,7677 | 0,515072598 | 0,1278731 | 4,027999 | 5,63E-05 | 0,0004562 | MYL9 |
| ENSGALG00000001998 | 1765,5757 | 0,514536405 | 0,138305 | 3,7203024 | 0,000199 | 0,0013569 | R3HDM4 |
| ENSGALG00000027073 | 784,17281 | 0,514122556 | 0,1915064 | 2,684623 | 0,0072612 | 0,0272551 | AAMDC |
| ENSGALG00000012089 | 5360,5907 | 0,513995299 | 0,1209653 | 4,2491144 | 2,15E-05 | 0,0001992 | C14orf37 |
| ENSGALG00000013708 | 5397,0941 | 0,513478256 | 0,1315099 | 3,9044846 | 9,44E-05 | 0,0007134 | CYB5A |
| ENSGALG00000029159 | 2974,1732 | 0,51305649 | 0,129165 | 3,9721007 | 7,12E-05 | 0,00056 | APBB1 |
| ENSGALG00000005901 | 2799,2351 | 0,510684045 | 0,1324352 | 3,8561047 | 0,0001152 | 0,0008451 | PCMTD2 |
| ENSGALG00000023786 | 1008,229 | 0,510469642 | 0,1622026 | 3,1471121 | 0,0016489 | 0,008038 | PLEKHH3 |
| ENSGALG00000017053 | 1433,2665 | 0,50974214 | 0,1465107 | 3,4792149 | 0,0005029 | 0,0030135 | SPG20 |
| ENSGALG00000004741 | 5108,3909 | 0,509578858 | 0,1446763 | 3,5221998 | 0,000428 | 0,0026378 | EPHB2 |
| ENSGALG00000012644 | 1843,5171 | 0,509005651 | 0,1372404 | 3,7088609 | 0,0002082 | 0,0014145 | FOXD3_CHICK |
| ENSGALG00000012697 | 713,61608 | 0,508988728 | 0,1895408 | 2,6853778 | 0,0072448 | 0,0272029 | SLC41A2 |
| ENSGALG00000016014 | 587,48188 | 0,508625141 | 0,2007004 | 2,534251 | 0,0112688 | 0,0386655 | RCAN1 |
| ENSGALG00000029036 | 2011,0671 | 0,506773529 | 0,1625524 | 3,1176018 | 0,0018233 | 0,0087194 | ZNF687 |
| ENSGALG00000016708 | 1004,6127 | 0,506256389 | 0,1774268 | 2,8533257 | 0,0043264 | 0,0177744 | ENPP4 |
| ENSGALG00000011324 | 77589,574 | 0,50624037 | 0,1039388 | 4,8705612 | 1,11E-06 | 1,44E-05 | TUBA4B |
| ENSGALG00000015597 | 932,24281 | 0,505967585 | 0,1755067 | 2,8828969 | 0,0039404 | 0,0164962 | CPZ |
| ENSGALG00000005757 | 2287,1727 | 0,505955901 | 0,148039 | 3,4177214 | 0,0006315 | 0,0036399 | IRF8 |
| ENSGALG00000011942 | 693,1373 | 0,505923034 | 0,1765129 | 2,8662097 | 0,0041542 | 0,0172007 | |
| ENSGALG00000003175 | 433,6143 | 0,505815122 | 0,2067003 | 2,4470941 | 0,0144013 | 0,0467957 | STKLD1 |
| ENSGALG00000012670 | 2710,2763 | 0,505329169 | 0,1395627 | 3,6208026 | 0,0002937 | 0,0019059 | NRSN1 |
| ENSGALG00000028145 | 4237,9607 | 0,505320547 | 0,1554994 | 3,2496615 | 0,0011554 | 0,0060245 | |
| ENSGALG00000027891 | 10377,445 | 0,504324315 | 0,1084665 | 4,649585 | 3,33E-06 | 3,83E-05 | NREP |
| ENSGALG00000024295 | 3217,4381 | 0,504249612 | 0,1324209 | 3,8079298 | 0,0001401 | 0,0010058 | |
| ENSGALG00000008631 | 3955,1009 | 0,503418102 | 0,1318317 | 3,8186431 | 0,0001342 | 0,0009682 | TYRO3 |
| ENSGALG00000015026 | 714,61563 | 0,503288049 | 0,182056 | 2,7644677 | 0,0057016 | 0,0223002 | TRAF3IP2 |
| ENSGALG00000005811 | 1644,0505 | 0,503226947 | 0,1577089 | 3,1908596 | 0,0014185 | 0,0071292 | COL7A1 |
| ENSGALG00000006802 | 2159,6578 | 0,502556738 | 0,1878266 | 2,6756417 | 0,0074586 | 0,0278522 | LAMB2 |
| ENSGALG00000025977 | 2572,6109 | 0,502103804 | 0,1296104 | 3,8739483 | 0,0001071 | 0,0007966 | GADD45A |
| ENSGALG00000023554 | 790,06032 | 0,500814372 | 0,1776938 | 2,8184128 | 0,0048262 | 0,0194807 | PIGW |
| ENSGALG00000024398 | 21096,229 | 0,500531072 | 0,1162284 | 4,3064435 | 1,66E-05 | 0,0001594 | RPS28 |
| ENSGALG00000004851 | 3943,1572 | 0,500191453 | 0,1444617 | 3,4624504 | 0,0005353 | 0,0031741 | MED9 |
| ENSGALG00000008341 | 3732,2674 | 0,499807137 | 0,1274369 | 3,9219961 | 8,78E-05 | 0,0006725 | FURIN |
| ENSGALG00000012487 | 1459,0736 | 0,499098415 | 0,1787383 | 2,7923417 | 0,0052328 | 0,0208196 | MPST |
| ENSGALG00000011326 | 2703,9221 | 0,4983963 | 0,1426307 | 3,4943128 | 0,0004753 | 0,0028759 | STK16 |
| ENSGALG00000000842 | 2625,6895 | 0,49824929 | 0,1497155 | 3,3279751 | 0,0008748 | 0,0047852 | NDUFA2 |
| ENSGALG00000011181 | 876,58708 | 0,497942212 | 0,1756687 | 2,834553 | 0,004589 | 0,0186403 | FKBP14 |
| ENSGALG00000018819 | 651,0396 | 0,497763645 | 0,1880595 | 2,6468407 | 0,0081248 | 0,0297602 | TBCEL |
| ENSGALG00000007370 | 4047,939 | 0,4969378 | 0,1207586 | 4,115134 | 3,87E-05 | 0,0003312 | CLP1 |
| ENSGALG00000012947 | 2224,1495 | 0,495800698 | 0,1377049 | 3,6004578 | 0,0003177 | 0,0020391 | CAPRIN2 |
| ENSGALG00000013067 | 8815,6345 | 0,494753671 | 0,1163549 | 4,2521102 | 2,12E-05 | 0,0001972 | AKR1B1 |
| ENSGALG00000007441 | 1294,7426 | 0,494713822 | 0,1878271 | 2,633879 | 0,0084416 | 0,030714 | MAP6 |
| ENSGALG00000000527 | 6471,7809 | 0,494652161 | 0,1195904 | 4,1362185 | 3,53E-05 | 0,0003062 | SH3BGRL3 |
| ENSGALG00000012540 | 1788,9218 | 0,494557337 | 0,1380803 | 3,5816639 | 0,0003414 | 0,0021731 | RBFOX2 |
| ENSGALG00000011778 | 2592,1932 | 0,494085012 | 0,1468186 | 3,3652744 | 0,0007647 | 0,0042862 | ITGB5 |
| ENSGALG00000009090 | 635,87673 | 0,493735019 | 0,1952486 | 2,5287506 | 0,0114469 | 0,0390797 | UAP1L1 |
| ENSGALG00000001242 | 570,55086 | 0,493511493 | 0,1867093 | 2,6432083 | 0,0082124 | 0,030041 | |
| ENSGALG00000027824 | 1654,6946 | 0,493225579 | 0,1467768 | 3,360379 | 0,0007784 | 0,0043539 | MRPL55 |
| ENSGALG00000005897 | 4766,7142 | 0,493064333 | 0,1262096 | 3,9067089 | 9,36E-05 | 0,0007093 | HMCES |
| ENSGALG00000012816 | 2261,4061 | 0,492973169 | 0,1565978 | 3,1480205 | 0,0016438 | 0,0080223 | SLC22A23 |
| ENSGALG00000005071 | 6694,0161 | 0,492854493 | 0,1137172 | 4,3340352 | 1,46E-05 | 0,000144 | MPP1 |
| ENSGALG00000000336 | 3779,8129 | 0,492268059 | 0,1394428 | 3,5302496 | 0,0004152 | 0,0025705 | PPP1R12B |
| ENSGALG00000004351 | 891,83433 | 0,49204731 | 0,1689419 | 2,9125234 | 0,0035852 | 0,0152683 | |
| ENSGALG00000004902 | 4281,0079 | 0,491426093 | 0,1175409 | 4,1808935 | 2,90E-05 | 0,0002581 | TOM1L2 |
| ENSGALG00000026195 | 759,73378 | 0,491395512 | 0,1717565 | 2,8610013 | 0,0042231 | 0,017455 | GALT |
| ENSGALG00000011894 | 1400,2218 | 0,491394089 | 0,1454582 | 3,3782503 | 0,0007295 | 0,0041177 | CYP2D6 |
| ENSGALG00000017076 | 6428,3314 | 0,490620681 | 0,1467776 | 3,3426126 | 0,0008299 | 0,0045704 | B3GLCT |
| ENSGALG00000008409 | 5336,4487 | 0,490103959 | 0,1431132 | 3,4245884 | 0,0006157 | 0,0035734 | GSTO1 |
| ENSGALG00000005751 | 599,07782 | 0,487576383 | 0,1863158 | 2,6169354 | 0,0088723 | 0,0319489 | RAB26 |
| ENSGALG00000005299 | 758,00774 | 0,487367317 | 0,1823876 | 2,6721512 | 0,0075367 | 0,028082 | ANXA7 |
| ENSGALG00000010230 | 677,39962 | 0,486896744 | 0,1774474 | 2,7438932 | 0,0060715 | 0,0234907 | TESK2 |
| ENSGALG00000011531 | 2214,443 | 0,486736013 | 0,1514726 | 3,2133593 | 0,0013119 | 0,0066923 | APAF1 |
| ENSGALG00000006915 | 1392,1391 | 0,485309324 | 0,1555538 | 3,1198817 | 0,0018092 | 0,0086661 | BAIAP2 |
| ENSGALG00000011798 | 1079,8245 | 0,484782273 | 0,1756184 | 2,7604306 | 0,0057725 | 0,0225205 | HEBP1 |

| | | | | | | | |
|---|---|---|---|---|---|---|---|
| ENSGALG00000017419 | 644,62948 | 0,484006649 | 0,1950269 | 2,4817427 | 0,0130742 | 0,0434688 | HPS1 |
| ENSGALG00000000947 | 2256,1097 | 0,482847185 | 0,1494215 | 3,2314447 | 0,0012317 | 0,0063519 | FKBP5 |
| ENSGALG00000027173 | 1663,1472 | 0,482690141 | 0,1744895 | 2,7662985 | 0,0056697 | 0,0222137 | TMEM159 |
| ENSGALG00000008604 | 829,00435 | 0,482676253 | 0,1667486 | 2,8946345 | 0,003796 | 0,0160157 | TMEM255A |
| ENSGALG00000014581 | 2314,88 | 0,482286067 | 0,1382204 | 3,4892527 | 0,0004844 | 0,0029219 | MEF2BNB |
| ENSGALG00000011738 | 6621,7844 | 0,482221251 | 0,1182329 | 4,0785695 | 4,53E-05 | 0,0003771 | ARHGDIB |
| ENSGALG00000027688 | 6057,2057 | 0,481915396 | 0,1181641 | 4,0783585 | 4,54E-05 | 0,0003772 | NDUFB4 |
| ENSGALG00000004476 | 498,99618 | 0,48187718 | 0,1958501 | 2,4604391 | 0,0138767 | 0,045449 | SEMA3G |
| ENSGALG00000000625 | 1809,062 | 0,481753107 | 0,1576512 | 3,0558166 | 0,0022445 | 0,0103589 | MAPT |
| ENSGALG00000015774 | 1090,9886 | 0,481727224 | 0,159212 | 3,0256962 | 0,0024806 | 0,0112579 | CYYR1 |
| ENSGALG00000001763 | 1095,0503 | 0,481398992 | 0,1634996 | 2,9443437 | 0,0032364 | 0,0140062 | CZH18ORF25 |
| ENSGALG00000005713 | 4508,0892 | 0,480930735 | 0,1143118 | 4,2071834 | 2,59E-05 | 0,0002339 | NCKIPSD |
| ENSGALG00000022838 | 522,00396 | 0,480740174 | 0,1959359 | 2,4535587 | 0,014145 | 0,0461834 | GAREM2 |
| ENSGALG00000003405 | 5095,9892 | 0,480657067 | 0,1124415 | 4,2747319 | 1,91E-05 | 0,0001804 | BSDC1 |
| ENSGALG00000002033 | 6624,4207 | 0,480537464 | 0,1111006 | 4,3252479 | 1,52E-05 | 0,0001482 | NDUFB6 |
| ENSGALG00000001531 | 815,50813 | 0,480440457 | 0,1690567 | 2,8418887 | 0,0044847 | 0,0182867 | FN3K |
| ENSGALG00000029135 | 7403,5283 | 0,479980361 | 0,1296505 | 3,7021087 | 0,0002138 | 0,0014476 | YEATS4 |
| ENSGALG00000011436 | 3916,7711 | 0,477975314 | 0,1192676 | 4,0075871 | 6,13E-05 | 0,0004904 | TMBIM1 |
| ENSGALG00000003466 | 4818,2759 | 0,477695903 | 0,1342072 | 3,5593901 | 0,0003717 | 0,0023355 | AKR1B1L |
| ENSGALG00000007661 | 1313,7584 | 0,477627101 | 0,1756565 | 2,7190975 | 0,006546 | 0,0250419 | MYCBPAP |
| ENSGALG00000029123 | 831,21662 | 0,477624117 | 0,1743995 | 2,7386782 | 0,0061687 | 0,0238124 | RBFOX3 |
| ENSGALG00000001606 | 1726,3623 | 0,477443775 | 0,1469062 | 3,2499912 | 0,0011541 | 0,0060203 | INPP5B |
| ENSGALG00000000785 | 1523,6714 | 0,476932316 | 0,1468983 | 3,2466828 | 0,0011676 | 0,0060741 | HYAL1 |
| ENSGALG00000028573 | 1369,3549 | 0,476874115 | 0,1566656 | 3,0438978 | 0,0023353 | 0,0107165 | SAYSD1 |
| ENSGALG00000014765 | 2914,6817 | 0,476665637 | 0,1245755 | 3,8263205 | 0,0001301 | 0,0009415 | TRIM23 |
| ENSGALG00000016994 | 1571,3108 | 0,474313428 | 0,1588828 | 2,9853031 | 0,002833 | 0,012572 | NUDT15 |
| ENSGALG00000000943 | 4846,5059 | 0,473716041 | 0,112782 | 4,2002794 | 2,67E-05 | 0,0002402 | WDTC1 |
| ENSGALG00000006588 | 1949,8657 | 0,473632802 | 0,1386756 | 3,4154005 | 0,0006369 | 0,0036653 | LRRC75B |
| ENSGALG00000012704 | 1775,4396 | 0,473322212 | 0,1740897 | 2,7188406 | 0,0065511 | 0,0250543 | MYLIP |
| ENSGALG00000002615 | 1139,5869 | 0,472751564 | 0,1639644 | 2,883258 | 0,0039358 | 0,0164831 | DEF6 |
| ENSGALG00000023157 | 3625,8738 | 0,472554642 | 0,1422354 | 3,3223424 | 0,0008927 | 0,0048731 | REEP5 |
| ENSGALG00000001849 | 1893,9454 | 0,47187756 | 0,1796068 | 2,6272817 | 0,008607 | 0,0311909 | BSN |
| ENSGALG00000006702 | 2572,8058 | 0,471728545 | 0,1314835 | 3,5877394 | 0,0003336 | 0,0021305 | MFGE8 |
| ENSGALG00000003359 | 6429,5457 | 0,469661247 | 0,1200374 | 3,9126251 | 9,13E-05 | 0,0006948 | RAB5C |
| ENSGALG00000027377 | 1588,8633 | 0,469161575 | 0,1429069 | 3,2829883 | 0,0010271 | 0,0054584 | NKAIN3 |
| ENSGALG00000013511 | 6097,6262 | 0,469095656 | 0,1205264 | 3,892056 | 9,94E-05 | 0,000746 | ANKRA2 |
| ENSGALG00000014963 | 978,12348 | 0,468684934 | 0,1590044 | 2,9476227 | 0,0032023 | 0,0138939 | DSE |
| ENSGALG00000016988 | 780,62352 | 0,468570179 | 0,173966 | 2,6934591 | 0,0070715 | 0,026633 | KIAA0226L |
| ENSGALG00000003348 | 1706,407 | 0,468370054 | 0,138122 | 3,3909876 | 0,0006964 | 0,0039639 | HIGD2A |
| ENSGALG00000027993 | 718,86063 | 0,467989481 | 0,184762 | 2,5329319 | 0,0113113 | 0,0387543 | MXD3 |
| ENSGALG00000028342 | 3684,9232 | 0,4672884 | 0,1405267 | 3,3252632 | 0,0008834 | 0,0048262 | CHRAC1 |
| ENSGALG00000004581 | 7502,7339 | 0,466855623 | 0,1169353 | 3,9924255 | 6,54E-05 | 0,0005177 | NDST1 |
| ENSGALG00000013795 | 1427,0027 | 0,466841888 | 0,1691704 | 2,759595 | 0,0057873 | 0,0225577 | HIVEP2 |
| ENSGALG00000007238 | 3294,4647 | 0,466556765 | 0,1396865 | 3,3400276 | 0,0008377 | 0,0046082 | PCSK6 |
| ENSGALG00000001556 | 1997,4095 | 0,466292104 | 0,1362595 | 3,4220882 | 0,0006214 | 0,0035957 | ROMO1 |
| ENSGALG00000006942 | 9942,2696 | 0,465540173 | 0,155783 | 2,988388 | 0,0028045 | 0,0124702 | PIK3IP1 |
| ENSGALG00000009655 | 1343,4112 | 0,465196113 | 0,1463089 | 3,1795467 | 0,0014751 | 0,0073322 | AADAT |
| ENSGALG00000005074 | 969,33864 | 0,464823262 | 0,1663096 | 2,7949274 | 0,0051911 | 0,020684 | FAM102A |
| ENSGALG00000002485 | 4126,4279 | 0,464623616 | 0,1275886 | 3,6415767 | 0,000271 | 0,0017746 | PHKG1 |
| ENSGALG00000011474 | 2398,5188 | 0,463345487 | 0,1386209 | 3,3425371 | 0,0008302 | 0,0045704 | OSBPL10 |
| ENSGALG00000013698 | 1914,3877 | 0,462883571 | 0,1399784 | 3,3068211 | 0,0009436 | 0,0051001 | EFHD2 |
| ENSGALG00000007582 | 9089,3986 | 0,462276159 | 0,111854 | 4,1328546 | 3,58E-05 | 0,0003098 | DPYSL3 |
| ENSGALG00000003976 | 1632,2576 | 0,461042882 | 0,1442377 | 3,1964111 | 0,0013915 | 0,0070167 | BCORL1 |
| ENSGALG00000024391 | 9415,8207 | 0,460947942 | 0,1147209 | 4,0179932 | 5,87E-05 | 0,0004723 | CACTIN |
| ENSGALG00000027418 | 872,62308 | 0,460841146 | 0,1699387 | 2,7118078 | 0,0066917 | 0,0255122 | HINT2 |
| ENSGALG00000003406 | 1016,1622 | 0,460645128 | 0,185859 | 2,4784659 | 0,0131949 | 0,0437898 | RXRG |
| ENSGALG00000014818 | 715,41879 | 0,460609845 | 0,1805705 | 2,5508589 | 0,0107458 | 0,0372559 | TRIM45 |
| ENSGALG00000024034 | 3571,4505 | 0,460548751 | 0,1231211 | 3,740617 | 0,0001836 | 0,0012713 | PI4KB |
| ENSGALG00000014226 | 4001,8851 | 0,460133597 | 0,1394922 | 3,2986321 | 0,0009716 | 0,0052095 | KIAA0930 |
| ENSGALG00000022980 | 7902,7397 | 0,459059074 | 0,1123607 | 4,0855838 | 4,40E-05 | 0,0003668 | CDKN1B |
| ENSGALG00000001728 | 1860,8126 | 0,458348865 | 0,1498189 | 3,0593533 | 0,0022182 | 0,0102792 | NACC2 |
| ENSGALG00000014324 | 2616,4144 | 0,458208088 | 0,1484479 | 3,0866594 | 0,0020242 | 0,0095394 | SLC2A3 |
| ENSGALG00000026871 | 2483,7701 | 0,457953468 | 0,1821233 | 2,5145249 | 0,0119193 | 0,0403885 | SC5D |
| ENSGALG00000015972 | 3109,9302 | 0,45786464 | 0,1525893 | 3,0006332 | 0,0026942 | 0,0120504 | IFNGR2 |

| | | | | | | | |
|---|---|---|---|---|---|---|---|
| ENSGALG00000001881 | 2571,7226 | 0,456046301 | 0,1474962 | 3,0919195 | 0,0019887 | 0,0093882 | DNLZ |
| ENSGALG00000001779 | 903,85433 | 0,455324974 | 0,1721096 | 2,6455517 | 0,0081558 | 0,0298497 | KIAA0895L |
| ENSGALG00000000921 | 4858,4012 | 0,454646053 | 0,1160077 | 3,9191038 | 8,89E-05 | 0,0006791 | SLC9A1 |
| ENSGALG00000004537 | 5387,8208 | 0,454616982 | 0,1154772 | 3,9368562 | 8,26E-05 | 0,0006354 | EFNB1 |
| ENSGALG00000016870 | 5615,9964 | 0,454001604 | 0,1190199 | 3,8145028 | 0,0001365 | 0,000984 | TMTC4 |
| ENSGALG00000011655 | 1599,7524 | 0,452530841 | 0,1507993 | 3,0008809 | 0,002692 | 0,0120445 | IQCB1 |
| ENSGALG00000011592 | 753,887 | 0,452054353 | 0,1756234 | 2,5739982 | 0,0100531 | 0,0353038 | MRAS |
| ENSGALG00000027963 | 10511,195 | 0,451771641 | 0,1053151 | 4,2897137 | 1,79E-05 | 0,0001697 | COX7C |
| ENSGALG00000012110 | 938,42508 | 0,451377511 | 0,1665857 | 2,7095817 | 0,0067368 | 0,0256277 | KIAA0895 |
| ENSGALG00000009054 | 1742,5243 | 0,451368563 | 0,1444522 | 3,1246911 | 0,0017799 | 0,008551 | TMEM141 |
| ENSGALG00000017351 | 2392,614 | 0,45115395 | 0,1297902 | 3,4760237 | 0,0005089 | 0,0030416 | STIM1 |
| ENSGALG00000007286 | 2211,1348 | 0,450759437 | 0,1433837 | 3,1437292 | 0,0016681 | 0,0081111 | SIRT7 |
| ENSGALG00000004623 | 967,41624 | 0,450139696 | 0,1793951 | 2,5092091 | 0,0121002 | 0,0408428 | AGTRAP |
| ENSGALG00000001376 | 9938,3188 | 0,45012042 | 0,1054385 | 4,2690347 | 1,96E-05 | 0,0001846 | ARID3B |
| ENSGALG00000017126 | 2105,4507 | 0,449836372 | 0,1445661 | 3,111632 | 0,0018606 | 0,0088634 | ZDHHC20 |
| ENSGALG00000004353 | 9460,5472 | 0,448918917 | 0,1126661 | 3,9845069 | 6,76E-05 | 0,0005331 | TBCA |
| ENSGALG00000015780 | 22777,794 | 0,447991765 | 0,1105939 | 4,0507816 | 5,10E-05 | 0,0004182 | PNRC1 |
| ENSGALG00000026200 | 2407,7782 | 0,447943252 | 0,1654706 | 2,7070867 | 0,0067877 | 0,0257707 | FAM134B |
| ENSGALG00000012218 | 3810,3697 | 0,44776649 | 0,1283039 | 3,4898898 | 0,0004832 | 0,0029175 | CBY1 |
| ENSGALG00000008274 | 1660,0276 | 0,446767639 | 0,1420336 | 3,1455061 | 0,001658 | 0,0080707 | CRBN |
| ENSGALG00000005608 | 3810,2468 | 0,44662731 | 0,1300871 | 3,4332951 | 0,0005963 | 0,0034739 | SLC9A3R2 |
| ENSGALG00000007296 | 1317,9251 | 0,446130433 | 0,1486148 | 3,0019248 | 0,0026828 | 0,0120073 | MAFG |
| ENSGALG00000016545 | 1336,374 | 0,445723593 | 0,1531647 | 2,910094 | 0,0036132 | 0,0153682 | REPS2 |
| ENSGALG00000017329 | 3706,9673 | 0,44555284 | 0,1363133 | 3,2685934 | 0,0010808 | 0,0057037 | FCHSD2 |
| ENSGALG00000016210 | 979,19029 | 0,445450514 | 0,1608438 | 2,7694608 | 0,0056149 | 0,0220587 | ST3GAL1 |
| ENSGALG00000015987 | 4807,1926 | 0,445391941 | 0,1245569 | 3,5758102 | 0,0003491 | 0,0022157 | TP53INP1 |
| ENSGALG00000008681 | 1024,5364 | 0,444903231 | 0,167862 | 2,65041 | 0,0080394 | 0,029535 | NSMF |
| ENSGALG00000000067 | 2534,6314 | 0,444870546 | 0,1286038 | 3,4592336 | 0,0005417 | 0,0032081 | SPR |
| ENSGALG00000011921 | 3797,7786 | 0,444716768 | 0,1414811 | 3,1432948 | 0,0016706 | 0,0081203 | DHRS7 |
| ENSGALG00000000233 | 2504,2434 | 0,444272958 | 0,1492937 | 2,9758312 | 0,002922 | 0,0128913 | SP2 |
| ENSGALG00000015344 | 1754,4897 | 0,442789405 | 0,1532928 | 2,88852 | 0,0038706 | 0,0162799 | NXPE3 |
| ENSGALG00000027337 | 6395,1608 | 0,441351335 | 0,1148234 | 3,8437395 | 0,0001212 | 0,0008818 | CLDND1 |
| ENSGALG00000004907 | 2377,2693 | 0,441232207 | 0,1337197 | 3,2996803 | 0,000968 | 0,0051966 | KCTD15 |
| ENSGALG00000002626 | 1744,7946 | 0,440405851 | 0,1788023 | 2,4630889 | 0,0137746 | 0,0451855 | RXRA |
| ENSGALG00000017096 | 2040,4546 | 0,440278375 | 0,1337931 | 3,29074 | 0,0009992 | 0,0053268 | LNX2 |
| ENSGALG00000012043 | 13548,036 | 0,439717314 | 0,1042663 | 4,2172536 | 2,47E-05 | 0,0002255 | PDCD6IP |
| ENSGALG00000012265 | 3656,8809 | 0,439419657 | 0,1289098 | 3,4087374 | 0,0006526 | 0,0037398 | MICALL1 |
| ENSGALG00000015988 | 2456,5859 | 0,439214581 | 0,1576704 | 2,785651 | 0,005342 | 0,0211799 | PLEKHF2 |
| ENSGALG00000002986 | 2686,672 | 0,439006087 | 0,125125 | 3,5085396 | 0,0004506 | 0,0027448 | WNK4 |
| ENSGALG00000010811 | 7190,0128 | 0,438424216 | 0,1472899 | 2,9766068 | 0,0029146 | 0,0128629 | LGMN |
| ENSGALG00000005820 | 8527,7428 | 0,438136271 | 0,1128485 | 3,8825187 | 0,0001034 | 0,0007734 | RNF130 |
| ENSGALG00000012913 | 1327,8126 | 0,437865562 | 0,1766936 | 2,478107 | 0,0132082 | 0,043818 | PKP2 |
| ENSGALG00000006204 | 1464,0386 | 0,437842966 | 0,1451037 | 3,0174484 | 0,0025491 | 0,0114911 | CHST13 |
| ENSGALG00000012136 | 1595,9568 | 0,437388472 | 0,1538246 | 2,8434236 | 0,0044632 | 0,0182316 | MPPED2 |
| ENSGALG00000026116 | 1987,2825 | 0,437155594 | 0,135619 | 3,2234086 | 0,0012667 | 0,006496 | MAPRE3 |
| ENSGALG00000013091 | 2461,6299 | 0,436802578 | 0,1334788 | 3,272449 | 0,0010662 | 0,0056418 | EPS8 |
| ENSGALG00000008444 | 16562,743 | 0,436671101 | 0,1126641 | 3,8758677 | 0,0001062 | 0,0007926 | TUBAL3 |
| ENSGALG00000012821 | 60743,332 | 0,436526282 | 0,1102259 | 3,9602861 | 7,49E-05 | 0,0005824 | TUBB2B |
| ENSGALG00000016322 | 1469,7377 | 0,435481617 | 0,165226 | 2,6356732 | 0,0083971 | 0,030593 | |
| ENSGALG00000006399 | 2888,1679 | 0,435019992 | 0,1286562 | 3,38126 | 0,0007215 | 0,0040828 | MESDC1 |
| ENSGALG00000001289 | 1028,449 | 0,434897808 | 0,1641592 | 2,6492441 | 0,0080672 | 0,0296038 | SHC2 |
| ENSGALG00000010134 | 2529,1958 | 0,434710794 | 0,126122 | 3,4467496 | 0,0005674 | 0,003334 | ZFAND3 |
| ENSGALG00000000270 | 895,80796 | 0,434589704 | 0,1737942 | 2,500599 | 0,0123983 | 0,0416902 | THRA |
| ENSGALG00000016059 | 2071,2967 | 0,434358409 | 0,1376843 | 3,1547427 | 0,0016064 | 0,0078758 | ETS2 |
| ENSGALG00000000049 | 1247,3305 | 0,433824072 | 0,1496385 | 2,8991468 | 0,0037418 | 0,015841 | CSDC2 |
| ENSGALG00000006655 | 1295,5112 | 0,433042079 | 0,15427 | 2,8070402 | 0,0049999 | 0,0200672 | MAP3K13 |
| ENSGALG00000021365 | 7390,78 | 0,432965219 | 0,1172033 | 3,6941392 | 0,0002206 | 0,0014864 | DCTN3 |
| ENSGALG00000003447 | 2587,3652 | 0,432921053 | 0,1381602 | 3,1334707 | 0,0017275 | 0,0083553 | FRS3 |
| ENSGALG00000015607 | 3625,3575 | 0,432451982 | 0,1684544 | 2,5671749 | 0,0102531 | 0,035895 | NCOA2 |
| ENSGALG00000004725 | 7054,5494 | 0,431307897 | 0,129893 | 3,3204852 | 0,0008986 | 0,0048938 | RP11-162P23.2 |
| ENSGALG00000014410 | 1635,3072 | 0,430076039 | 0,1754863 | 2,450767 | 0,0142552 | 0,0464477 | CSTB |
| ENSGALG00000029157 | 4821,9983 | 0,430051945 | 0,1512627 | 2,8430801 | 0,004468 | 0,0182458 | MBNL3 |
| ENSGALG00000000551 | 4770,6021 | 0,428841133 | 0,1175738 | 3,6474204 | 0,0002649 | 0,0017414 | C6orf89 |
| ENSGALG00000014804 | 1784,8867 | 0,427778224 | 0,1683627 | 2,5408135 | 0,0110595 | 0,0381106 | THBS4 |

| | | | | | | | |
|---|---|---|---|---|---|---|---|
| ENSGALG00000005295 | 1837,9552 | 0,4276113 | 0,1611793 | 2,653017 | 0,0079776 | 0,029382 | BCAS3 |
| ENSGALG00000023612 | 4284,8687 | 0,427551312 | 0,1222371 | 3,4977227 | 0,0004692 | 0,0028433 | LZTS2 |
| ENSGALG00000010838 | 5580,7171 | 0,427079258 | 0,1310404 | 3,2591413 | 0,0011175 | 0,0058586 | ERO1LB |
| ENSGALG00000016605 | 1894,8074 | 0,426964808 | 0,1498455 | 2,8493661 | 0,0043806 | 0,0179484 | EFR3B |
| ENSGALG00000007630 | 1371,8459 | 0,426278895 | 0,1475832 | 2,8883966 | 0,0038721 | 0,0162813 | CDIP1 |
| ENSGALG00000026022 | 1208,8026 | 0,425857109 | 0,1749797 | 2,4337512 | 0,0149433 | 0,0482458 | CREG2 |
| ENSGALG00000027822 | 849,07694 | 0,425453865 | 0,1646142 | 2,5845511 | 0,0097506 | 0,0344727 | |
| ENSGALG00000015304 | 5360,7582 | 0,425237788 | 0,1168587 | 3,6389042 | 0,0002738 | 0,0017905 | SNX3 |
| ENSGALG00000020528 | 5739,52 | 0,424819785 | 0,1109172 | 3,8300637 | 0,0001281 | 0,0009293 | INIP |
| ENSGALG00000014569 | 787,05235 | 0,42466715 | 0,1733979 | 2,4490901 | 0,0143218 | 0,0466262 | PNPLA2 |
| ENSGALG00000015603 | 3809,7497 | 0,424497436 | 0,1338532 | 3,171366 | 0,0015172 | 0,0075035 | ZCCHC9 |
| ENSGALG00000021688 | 803,0907 | 0,424387775 | 0,170819 | 2,4844292 | 0,0129759 | 0,0431738 | CD320 |
| ENSGALG00000001975 | 1467,2899 | 0,42427092 | 0,1628237 | 2,6057072 | 0,0091685 | 0,0327635 | TMEM120A |
| ENSGALG00000026692 | 7703,2554 | 0,424103985 | 0,1361891 | 3,1140823 | 0,0018452 | 0,0088024 | NRAS |
| ENSGALG00000013194 | 2699,5463 | 0,423818786 | 0,1417822 | 2,9892249 | 0,0027969 | 0,0124483 | IRX1 |
| ENSGALG00000015903 | 1395,6597 | 0,423426358 | 0,1458209 | 2,9037432 | 0,0036873 | 0,0156395 | FILIP1 |
| ENSGALG00000012009 | 3507,1232 | 0,422977808 | 0,1235894 | 3,4224446 | 0,0006206 | 0,0035925 | JKAMP |
| ENSGALG00000009107 | 1525,7253 | 0,422108392 | 0,149748 | 2,8187922 | 0,0048205 | 0,0194634 | NRXN1 |
| ENSGALG00000008257 | 1616,4159 | 0,421896886 | 0,1646167 | 2,5629047 | 0,0103801 | 0,036237 | MERTK |
| ENSGALG00000027710 | 12308,995 | 0,421872733 | 0,1075864 | 3,9212473 | 8,81E-05 | 0,0006742 | DAG1 |
| ENSGALG00000000436 | 3616,9217 | 0,421563067 | 0,1280766 | 3,2914916 | 0,0009966 | 0,0053161 | RABIF |
| ENSGALG00000008613 | 2544,3778 | 0,420884082 | 0,1331062 | 3,1620172 | 0,0015668 | 0,007729 | NDUFA1 |
| ENSGALG00000028302 | 1437,8585 | 0,420702301 | 0,1488301 | 2,826728 | 0,0047026 | 0,0190554 | |
| ENSGALG00000012131 | 1906,2436 | 0,419815141 | 0,1493221 | 2,8114727 | 0,0049315 | 0,0198588 | ELOVL6 |
| ENSGALG00000009595 | 1381,8135 | 0,419771594 | 0,1544649 | 2,7175851 | 0,006576 | 0,0251107 | TWF1 |
| ENSGALG00000007619 | 4223,0444 | 0,419444023 | 0,1439729 | 2,9133548 | 0,0035757 | 0,015242 | UBALD1 |
| ENSGALG00000007025 | 838,27153 | 0,41933553 | 0,165861 | 2,5282343 | 0,0114638 | 0,0391176 | CPNE8 |
| ENSGALG00000007054 | 5550,7308 | 0,419262331 | 0,1227261 | 3,4162453 | 0,0006349 | 0,0036566 | TBC1D16 |
| ENSGALG00000017675 | 6606,9674 | 0,417757976 | 0,11207 | 3,7276537 | 0,0001933 | 0,0013256 | NDUFS8 |
| ENSGALG00000008818 | 5178,1883 | 0,417517144 | 0,1293144 | 3,2286974 | 0,0012436 | 0,0064011 | IDH1 |
| ENSGALG00000004038 | 1559,4978 | 0,417388311 | 0,1404734 | 2,9712977 | 0,0029654 | 0,0130493 | ADAL |
| ENSGALG00000026394 | 6922,0955 | 0,417110916 | 0,1102725 | 3,782546 | 0,0001552 | 0,0010991 | CORO1B |
| ENSGALG00000002840 | 4242,1731 | 0,415750464 | 0,1518147 | 2,7385381 | 0,0061713 | 0,0238158 | GRM4 |
| ENSGALG00000007220 | 48744,176 | 0,415111704 | 0,1048964 | 3,9573492 | 7,58E-05 | 0,0005879 | FTH1 |
| ENSGALG00000001816 | 2247,2353 | 0,414935997 | 0,1327278 | 3,1262169 | 0,0017707 | 0,0085128 | MAGI3 |
| ENSGALG00000028812 | 2137,7927 | 0,414162814 | 0,1436705 | 2,8827261 | 0,0039425 | 0,0164962 | CREBL2 |
| ENSGALG00000015352 | 2437,8785 | 0,413775598 | 0,1391873 | 2,9727977 | 0,002951 | 0,0130039 | GAK |
| ENSGALG00000026802 | 39202,809 | 0,413746284 | 0,1146157 | 3,6098563 | 0,0003064 | 0,0019787 | TUBB2A |
| ENSGALG00000015915 | 2924,5989 | 0,413737551 | 0,1308763 | 3,1612868 | 0,0015707 | 0,0077372 | TMEM64 |
| ENSGALG00000020626 | 1251,1665 | 0,41305269 | 0,1546192 | 2,6714191 | 0,0075531 | 0,0281203 | C3ORF55 |
| ENSGALG00000012426 | 3567,3724 | 0,412512543 | 0,125055 | 3,2986494 | 0,0009715 | 0,0052095 | DDHD1 |
| ENSGALG00000004918 | 890,67566 | 0,411822203 | 0,1629138 | 2,5278534 | 0,0114762 | 0,0391454 | RAB9B |
| ENSGALG00000001201 | 7498,136 | 0,411468635 | 0,10775 | 3,8187341 | 0,0001341 | 0,0009682 | GFOD2 |
| ENSGALG00000024081 | 3055,1077 | 0,410810718 | 0,129763 | 3,1658532 | 0,0015463 | 0,0076361 | |
| ENSGALG00000004873 | 2191,1914 | 0,410670756 | 0,1393688 | 2,9466478 | 0,0032124 | 0,0139255 | PHACTR3 |
| ENSGALG00000017405 | 1260,4536 | 0,410522272 | 0,1629615 | 2,5191367 | 0,0117643 | 0,0399927 | NPR3 |
| ENSGALG00000003750 | 7887,3706 | 0,409049332 | 0,1094937 | 3,7358254 | 0,0001871 | 0,0012918 | PLCG1 |
| ENSGALG00000016828 | 4406,9031 | 0,407053917 | 0,1467975 | 2,7728942 | 0,005556 | 0,0219007 | GRTP1 |
| ENSGALG00000000478 | 3159,4613 | 0,406777826 | 0,1532189 | 2,6548799 | 0,0079337 | 0,0292412 | TANC2 |
| ENSGALG00000015004 | 7430,1607 | 0,406416416 | 0,1365123 | 2,9771421 | 0,0029095 | 0,0128446 | RIOK3 |
| ENSGALG00000011488 | 1838,711 | 0,405575212 | 0,1659995 | 2,443232 | 0,0145564 | 0,0472207 | CMTM7 |
| ENSGALG00000008597 | 7184,8669 | 0,404910294 | 0,1198878 | 3,37741 | 0,0007317 | 0,0041252 | MRPL2 |
| ENSGALG00000006505 | 3147,8767 | 0,40439474 | 0,1221546 | 3,310517 | 0,0009312 | 0,0050493 | ZFAND6 |
| ENSGALG00000014981 | 1640,2876 | 0,404127478 | 0,1524452 | 2,6509683 | 0,0080261 | 0,0295009 | COX17 |
| ENSGALG00000008143 | 14658,924 | 0,402789538 | 0,1063945 | 3,7858132 | 0,0001532 | 0,0010858 | UACA |
| ENSGALG00000003924 | 3485,2604 | 0,402077112 | 0,1262253 | 3,1853918 | 0,0014456 | 0,0072359 | ACBD6 |
| ENSGALG00000027768 | 2765,0207 | 0,401592495 | 0,1379224 | 2,9117282 | 0,0035944 | 0,0152976 | CTDSP1 |
| ENSGALG00000028931 | 2027,2809 | 0,401212621 | 0,1470133 | 2,7290911 | 0,0063509 | 0,0243915 | IGFBP4 |
| ENSGALG00000008508 | 3096,6728 | 0,401158456 | 0,1203234 | 3,3340018 | 0,0008561 | 0,004694 | PYGB |
| ENSGALG00000010704 | 15057,888 | 0,401049242 | 0,1064765 | 3,7665529 | 0,0001655 | 0,001164 | GLCCI1 |
| ENSGALG00000003371 | 6971,3556 | 0,400787786 | 0,1174349 | 3,4128513 | 0,0006429 | 0,0036962 | ELL |
| ENSGALG00000003575 | 5543,8738 | 0,400668854 | 0,1362202 | 2,9413311 | 0,003268 | 0,0141048 | DNTTIP1 |
| ENSGALG00000026316 | 1590,1424 | 0,400010985 | 0,1398143 | 2,8610159 | 0,0042229 | 0,017455 | TMEM136 |
| ENSGALG00000001473 | 2324,7953 | 0,399774883 | 0,1339201 | 2,9851752 | 0,0028342 | 0,0125732 | MIB2 |

| | | | | | | | |
|---|---|---|---|---|---|---|---|
| ENSGALG00000000253 | 2196,4708 | 0,399568995 | 0,1345932 | 2,9687154 | 0,0029905 | 0,0131255 | SLC25A46 |
| ENSGALG00000001096 | 5395,305 | 0,399476875 | 0,1210243 | 3,3007994 | 0,0009641 | 0,0051841 | AC005943.2 |
| ENSGALG00000002492 | 1100,2646 | 0,398681223 | 0,1527657 | 2,6097562 | 0,0090607 | 0,0324549 | H6PD |
| ENSGALG00000016191 | 3940,4435 | 0,398485008 | 0,1411098 | 2,8239358 | 0,0047438 | 0,0191937 | PKNOX1 |
| ENSGALG00000000972 | 1494,0966 | 0,398341588 | 0,1611855 | 2,4713244 | 0,0134614 | 0,0444414 | PBX3 |
| ENSGALG00000015961 | 15590,16 | 0,398310373 | 0,130734 | 3,0467236 | 0,0023135 | 0,0106334 | SUCLG1 |
| ENSGALG00000003412 | 4305,4895 | 0,398169298 | 0,118859 | 3,34993 | 0,0008083 | 0,0044792 | RALGDS |
| ENSGALG00000016549 | 1882,0348 | 0,397970824 | 0,1562578 | 2,5468866 | 0,0108689 | 0,037606 | CTPS2 |
| ENSGALG00000004176 | 2961,1415 | 0,397874037 | 0,1416462 | 2,8089278 | 0,0049707 | 0,0199869 | ATOX1 |
| ENSGALG00000014970 | 13935,559 | 0,397421157 | 0,1074971 | 3,6970395 | 0,0002181 | 0,0014739 | FSTL1 |
| ENSGALG00000003422 | 9411,1693 | 0,397187449 | 0,1102449 | 3,6027755 | 0,0003148 | 0,0020238 | TRIM3 |
| ENSGALG00000017262 | 1782,0171 | 0,39670681 | 0,1365862 | 2,9044423 | 0,0036791 | 0,0156094 | NARS2 |
| ENSGALG00000006736 | 1555,2322 | 0,396360498 | 0,1559292 | 2,5419267 | 0,0110243 | 0,0380183 | AEN |
| ENSGALG00000006889 | 2493,8401 | 0,396136143 | 0,1473921 | 2,6876356 | 0,007196 | 0,0270495 | TMC7 |
| ENSGALG00000014233 | 13243,506 | 0,394951392 | 0,1481986 | 2,6650135 | 0,0076985 | 0,0286146 | FBLN1 |
| ENSGALG00000005046 | 2539,1359 | 0,394497591 | 0,1359729 | 2,9012952 | 0,0037162 | 0,0157524 | SHMT1 |
| ENSGALG00000001572 | 4170,7383 | 0,394480266 | 0,1178093 | 3,3484648 | 0,0008126 | 0,0044938 | COMMD4 |
| ENSGALG00000028470 | 5093,1917 | 0,39443368 | 0,1241813 | 3,1762725 | 0,0014918 | 0,007402 | GSR |
| ENSGALG00000005675 | 3085,819 | 0,394073439 | 0,1342631 | 2,9350829 | 0,0033346 | 0,014349 | PTTG1IP |
| ENSGALG00000021047 | 9190,8891 | 0,393972396 | 0,122897 | 3,2057124 | 0,0013473 | 0,0068343 | KIFAP3 |
| ENSGALG00000008039 | 2411,3069 | 0,392392533 | 0,1551271 | 2,5294903 | 0,0114228 | 0,0390268 | MFSD13A |
| ENSGALG00000008050 | 1623,9365 | 0,391124824 | 0,1411515 | 2,7709581 | 0,0055892 | 0,0219805 | HBP1 |
| ENSGALG00000013348 | 2256,8618 | 0,390928939 | 0,128295 | 3,0471095 | 0,0023105 | 0,0106241 | OTUD7B |
| ENSGALG00000010490 | 10248,566 | 0,389494511 | 0,1117875 | 3,4842389 | 0,0004935 | 0,002968 | DPYSL4 |
| ENSGALG00000013000 | 7034,8497 | 0,388948819 | 0,1248778 | 3,1146355 | 0,0018417 | 0,008789 | ADIPOR2 |
| ENSGALG00000026095 | 1420,7135 | 0,388637494 | 0,1454616 | 2,6717534 | 0,0075456 | 0,0281076 | CDC26 |
| ENSGALG00000000955 | 5256,5616 | 0,388613492 | 0,1145617 | 3,3921753 | 0,0006934 | 0,0039501 | TMEM222 |
| ENSGALG00000011035 | 1343,9931 | 0,388589204 | 0,1474226 | 2,6358867 | 0,0083918 | 0,0305819 | AK4 |
| ENSGALG00000008386 | 1936,8714 | 0,388150893 | 0,1387904 | 2,7966699 | 0,0051632 | 0,0205937 | KIZ |
| ENSGALG00000015311 | 2435,4616 | 0,387705603 | 0,1261741 | 3,0727833 | 0,0021207 | 0,009926 | RNF38 |
| ENSGALG00000014496 | 6279,2131 | 0,385917845 | 0,134402 | 2,8713694 | 0,004087 | 0,0169958 | PROM1 |
| ENSGALG00000004011 | 4239,1769 | 0,385892717 | 0,1224058 | 3,1525685 | 0,0016184 | 0,0079204 | CAMKK2 |
| ENSGALG00000005337 | 6236,1037 | 0,385644449 | 0,1096926 | 3,5156828 | 0,0004386 | 0,0026828 | SNX2 |
| ENSGALG00000004086 | 6417,7852 | 0,385423377 | 0,1245819 | 3,0937355 | 0,0019765 | 0,0093504 | CHMP1B |
| ENSGALG00000026102 | 19802,423 | 0,385336727 | 0,121726 | 3,1656082 | 0,0015476 | 0,0076398 | CHTOP |
| ENSGALG00000015898 | 1319,6724 | 0,384642657 | 0,1580612 | 2,4335051 | 0,0149534 | 0,0482633 | MYO6 |
| ENSGALG00000016639 | 3515,2869 | 0,384196385 | 0,1227968 | 3,1287174 | 0,0017557 | 0,0084616 | KIF13B |
| ENSGALG00000026188 | 5641,8964 | 0,384179427 | 0,1152566 | 3,3332525 | 0,0008584 | 0,0047048 | TMEM8A |
| ENSGALG00000008736 | 6339,8487 | 0,384117634 | 0,1174006 | 3,271854 | 0,0010684 | 0,0056493 | PNPLA7 |
| ENSGALG00000020005 | 6706,2183 | 0,383841955 | 0,130019 | 2,9521995 | 0,0031552 | 0,0137158 | SFT2D1 |
| ENSGALG00000014808 | 2216,3488 | 0,383214364 | 0,1540134 | 2,488189 | 0,0128395 | 0,0428667 | USF3 |
| ENSGALG00000001536 | 1337,5687 | 0,382405736 | 0,149925 | 2,5506477 | 0,0107523 | 0,0372615 | ARHGAP39 |
| ENSGALG00000028635 | 2927,7161 | 0,382285105 | 0,1328162 | 2,8783008 | 0,0039982 | 0,0166777 | C9ORF58 |
| ENSGALG00000000456 | 4419,0221 | 0,381123553 | 0,1256788 | 3,0325206 | 0,0024252 | 0,0110469 | MRPS21 |
| ENSGALG00000011325 | 5273,0258 | 0,380378399 | 0,1366737 | 2,7831135 | 0,005384 | 0,0213339 | NDUFA12 |
| ENSGALG00000008297 | 4909,6132 | 0,379216929 | 0,1148855 | 3,3008262 | 0,000964 | 0,0051841 | SEMA4B |
| ENSGALG00000001166 | 2588,454 | 0,378691011 | 0,1266846 | 2,9892423 | 0,0027967 | 0,0124483 | GATSL2 |
| ENSGALG00000007205 | 14498,189 | 0,37857906 | 0,1049589 | 3,6069259 | 0,0003098 | 0,0019961 | COX6A1 |
| ENSGALG00000015958 | 3885,4269 | 0,378120956 | 0,1313335 | 2,8790889 | 0,0039883 | 0,0166514 | EIF2AK3 |
| ENSGALG00000028128 | 3864,6629 | 0,378077792 | 0,1172702 | 3,2239889 | 0,0012642 | 0,0064852 | HOXD8 |
| ENSGALG00000015436 | 20607,243 | 0,377556464 | 0,1126811 | 3,3506644 | 0,0008062 | 0,0044728 | SDCBP |
| ENSGALG00000002053 | 9601,2632 | 0,376194926 | 0,1115908 | 3,3711997 | 0,0007484 | 0,0042037 | POR |
| ENSGALG00000027416 | 1624,7836 | 0,375994995 | 0,1449326 | 2,5942749 | 0,0094791 | 0,0336613 | ROMO1L |
| ENSGALG00000002959 | 3279,2685 | 0,375394519 | 0,1495524 | 2,5101204 | 0,012069 | 0,0407741 | NRBF2 |
| ENSGALG00000009948 | 1430,5348 | 0,375138393 | 0,1485226 | 2,5258008 | 0,0115435 | 0,0393305 | HHIP |
| ENSGALG00000023180 | 1278,6895 | 0,374920803 | 0,1483185 | 2,527809 | 0,0114777 | 0,0391454 | |
| ENSGALG00000004191 | 5196,6988 | 0,374652449 | 0,1167651 | 3,2086007 | 0,0013338 | 0,0067774 | RIC8A |
| ENSGALG00000009959 | 1836,2989 | 0,374451686 | 0,1402648 | 2,6696054 | 0,007594 | 0,0282649 | AC025263.3 |
| ENSGALG00000003717 | 5316,7007 | 0,373543889 | 0,1130969 | 3,3028656 | 0,000957 | 0,0051565 | MYO9B |
| ENSGALG00000004324 | 3684,0767 | 0,373249102 | 0,1339589 | 2,7862949 | 0,0053314 | 0,0211564 | SETD1B |
| ENSGALG00000008438 | 6164,0602 | 0,373204358 | 0,1216092 | 3,0688816 | 0,0021486 | 0,0100141 | SLC35C1 |
| ENSGALG00000001032 | 4578,5026 | 0,373005757 | 0,1227121 | 3,039681 | 0,0023683 | 0,010842 | STUB1 |
| ENSGALG00000026291 | 5117,1597 | 0,37266418 | 0,1170472 | 3,1838789 | 0,0014532 | 0,0072578 | CDH3 |
| ENSGALG00000026421 | 20179,815 | 0,372144283 | 0,1169454 | 3,1822057 | 0,0014616 | 0,0072865 | PSMB2 |

| | | | | | | | |
|---|---|---|---|---|---|---|---|
| ENSGALG00000002990 | 14374,489 | 0,371847834 | 0,1302887 | 2,8540301 | 0,0043168 | 0,017756 | PRELID1 |
| ENSGALG00000002605 | 7879,732 | 0,371275034 | 0,1234991 | 3,006297 | 0,0026445 | 0,0118554 | MRPL17 |
| ENSGALG00000009073 | 2474,3956 | 0,370739827 | 0,1479952 | 2,5050805 | 0,0122424 | 0,0412267 | PSEN2 |
| ENSGALG00000004357 | 2772,1991 | 0,370669228 | 0,1314113 | 2,8206804 | 0,0047922 | 0,019355 | ANXA6 |
| ENSGALG00000016850 | 6183,7618 | 0,369402589 | 0,1221012 | 3,0253801 | 0,0024832 | 0,0112659 | ANKRD10 |
| ENSGALG00000015543 | 2204,9702 | 0,369304606 | 0,1292142 | 2,8580804 | 0,0042621 | 0,0175766 | MRPL50 |
| ENSGALG00000003185 | 3436,5941 | 0,368988195 | 0,1514261 | 2,4367543 | 0,0148197 | 0,0479038 | SCPEP1 |
| ENSGALG00000015537 | 3286,5424 | 0,368709498 | 0,1440858 | 2,5589581 | 0,0104986 | 0,036576 | MAN2B2 |
| ENSGALG00000006857 | 3516,2609 | 0,36847671 | 0,1191186 | 3,0933594 | 0,001979 | 0,0093558 | SPATA20 |
| ENSGALG00000026716 | 2067,7158 | 0,368315116 | 0,1476133 | 2,4951349 | 0,0125909 | 0,042223 | CELF5 |
| ENSGALG00000000747 | 4334,9927 | 0,36652476 | 0,1212386 | 3,0231688 | 0,0025014 | 0,0113184 | CHD5 |
| ENSGALG00000028249 | 8201,3974 | 0,366515679 | 0,1075014 | 3,409405 | 0,000651 | 0,0037354 | MCL1 |
| ENSGALG00000006149 | 3876,2934 | 0,366158451 | 0,1322855 | 2,7679406 | 0,0056412 | 0,0221403 | MORN4 |
| ENSGALG00000012952 | 17818,525 | 0,365044211 | 0,1286631 | 2,8372099 | 0,004551 | 0,0185125 | BASP1 |
| ENSGALG00000013848 | 3142,0736 | 0,364984572 | 0,1264318 | 2,8868109 | 0,0038917 | 0,0163434 | MVK |
| ENSGALG00000010621 | 3304,8585 | 0,364976716 | 0,1292617 | 2,8235486 | 0,0047495 | 0,0192112 | DPY30 |
| ENSGALG00000015663 | 10480,988 | 0,363842291 | 0,1220245 | 2,9817148 | 0,0028664 | 0,0126913 | HSDL2 |
| ENSGALG00000001597 | 12155,403 | 0,36336647 | 0,135585 | 2,6799894 | 0,0073624 | 0,027546 | PSMB3 |
| ENSGALG00000013876 | 3474,876 | 0,361738997 | 0,1264822 | 2,8599989 | 0,0042364 | 0,0174997 | FAM136A |
| ENSGALG00000021692 | 6544,9246 | 0,361385734 | 0,1165284 | 3,1012667 | 0,0019269 | 0,0091476 | AC024592.12 |
| ENSGALG00000016333 | 1977,9922 | 0,361282215 | 0,1426527 | 2,5326003 | 0,011322 | 0,0387796 | MTSS1 |
| ENSGALG00000005704 | 10751,887 | 0,361085404 | 0,1196869 | 3,0169164 | 0,0025536 | 0,0115009 | RP11-411B6.6 |
| ENSGALG00000006630 | 3807,9216 | 0,360923749 | 0,1347522 | 2,6784253 | 0,0073969 | 0,0276565 | COMMD7 |
| ENSGALG00000014589 | 16537,437 | 0,360345797 | 0,1177437 | 3,0604243 | 0,0022102 | 0,0102529 | |
| ENSGALG00000007589 | 8938,4163 | 0,360072922 | 0,1085811 | 3,3161649 | 0,0009126 | 0,0049604 | DEDD |
| ENSGALG00000011435 | 8069,4783 | 0,35897775 | 0,1123046 | 3,1964666 | 0,0013912 | 0,0070167 | ELK3 |
| ENSGALG00000010171 | 1427,5927 | 0,358902977 | 0,1475799 | 2,4319235 | 0,0150189 | 0,048421 | NFKBIE |
| ENSGALG00000001330 | 11236,984 | 0,358718136 | 0,1146464 | 3,128908 | 0,0017546 | 0,0084616 | ATP5G1 |
| ENSGALG00000006143 | 4041,5216 | 0,358630949 | 0,1180875 | 3,0369922 | 0,0023895 | 0,0109172 | RP11-548K23.11 |
| ENSGALG00000002159 | 5312,4581 | 0,35820654 | 0,1168583 | 3,0653076 | 0,0021745 | 0,0101146 | ACOX1 |
| ENSGALG00000005478 | 5105,9851 | 0,35705086 | 0,1183988 | 3,015662 | 0,0025642 | 0,011541 | SNX7 |
| ENSGALG00000026531 | 11386,746 | 0,356957338 | 0,1206613 | 2,9583418 | 0,003093 | 0,0135058 | CHST14 |
| ENSGALG00000026517 | 6681,3961 | 0,355711572 | 0,1112537 | 3,1973011 | 0,0013872 | 0,0070029 | RFC5 |
| ENSGALG00000011176 | 5734,5779 | 0,354690121 | 0,1402295 | 2,5293538 | 0,0114273 | 0,0390322 | SCRN1 |
| ENSGALG00000003096 | 2680,9001 | 0,354610608 | 0,1443928 | 2,455874 | 0,0140542 | 0,045942 | CBFA2T2 |
| ENSGALG00000001425 | 2462,9176 | 0,354258416 | 0,1359645 | 2,6055206 | 0,0091735 | 0,0327727 | SNX19 |
| ENSGALG00000023073 | 19350,483 | 0,353608475 | 0,1057356 | 3,3442711 | 0,000825 | 0,004546 | AKAP12 |
| ENSGALG00000013036 | 6599,1108 | 0,353370495 | 0,1362318 | 2,5938909 | 0,0094897 | 0,0336858 | ATP6V1E1 |
| ENSGALG00000014891 | 4747,3286 | 0,353275878 | 0,1368587 | 2,5813185 | 0,0098424 | 0,034752 | PELO |
| ENSGALG00000026600 | 7491,3546 | 0,352554713 | 0,1374236 | 2,5654603 | 0,0103039 | 0,0360174 | |
| ENSGALG00000026565 | 5799,2794 | 0,352044819 | 0,1113845 | 3,1606258 | 0,0015743 | 0,007752 | ENSA |
| ENSGALG00000007396 | 5182,898 | 0,351138505 | 0,1196447 | 2,9348426 | 0,0033372 | 0,0143556 | TAOK3 |
| ENSGALG00000012229 | 22179,4 | 0,350504617 | 0,1021787 | 3,430311 | 0,0006029 | 0,0035093 | RPS29 |
| ENSGALG00000007305 | 2042,815 | 0,350349865 | 0,1310633 | 2,6731358 | 0,0075146 | 0,0280151 | NME1 |
| ENSGALG00000000461 | 6717,843 | 0,348147161 | 0,1224667 | 2,8427896 | 0,0044721 | 0,0182515 | MRC2 |
| ENSGALG00000016241 | 5823,7161 | 0,347015145 | 0,1359999 | 2,5515842 | 0,0107234 | 0,0372068 | ATP6AP2 |
| ENSGALG00000001381 | 43057,795 | 0,34689365 | 0,1163519 | 2,9814171 | 0,0028692 | 0,0126954 | ACTG1 |
| ENSGALG00000010347 | 11028,081 | 0,346318448 | 0,1080262 | 3,2058747 | 0,0013465 | 0,0068343 | CCNG2 |
| ENSGALG00000005135 | 11151,897 | 0,345569481 | 0,1285407 | 2,688405 | 0,0071794 | 0,0270058 | SERPINE2 |
| ENSGALG00000026300 | 11402,689 | 0,345400821 | 0,1081778 | 3,1928993 | 0,0014085 | 0,0070895 | |
| ENSGALG00000020922 | 2968,9637 | 0,345299483 | 0,1246827 | 2,769425 | 0,0056155 | 0,0220587 | C1orf21 |
| ENSGALG00000023447 | 10483,165 | 0,344836567 | 0,1046431 | 3,2953598 | 0,000983 | 0,0052585 | RP11-343C2.11 |
| ENSGALG00000013563 | 2902,2623 | 0,344320197 | 0,1287254 | 2,6748425 | 0,0074764 | 0,0278962 | CCDC102A |
| ENSGALG00000002484 | 4545,8313 | 0,343583814 | 0,1190666 | 2,8856437 | 0,0039061 | 0,0163839 | CRYM |
| ENSGALG00000003215 | 4360,1328 | 0,343135123 | 0,1166654 | 2,9411903 | 0,0032695 | 0,0141048 | PEF1 |
| ENSGALG00000016665 | 8909,5125 | 0,342839663 | 0,1328584 | 2,5804883 | 0,0098661 | 0,0347816 | FDFT1 |
| ENSGALG00000026203 | 4631,9513 | 0,341949081 | 0,1195864 | 2,8594311 | 0,004244 | 0,0175151 | FAM174A |
| ENSGALG00000012586 | 2037,1791 | 0,341920905 | 0,1360582 | 2,5130482 | 0,0119693 | 0,0405176 | GKAP1 |
| ENSGALG00000003293 | 12978,607 | 0,341652813 | 0,1149964 | 2,9709875 | 0,0029684 | 0,0130583 | MBTPS1 |
| ENSGALG00000012770 | 2998,9246 | 0,340485203 | 0,1279524 | 2,6610294 | 0,0077902 | 0,028853 | MAK |
| ENSGALG00000013784 | 13631,601 | 0,339967639 | 0,1079829 | 3,1483476 | 0,001642 | 0,0080213 | CDT1 |
| ENSGALG00000006837 | 2354,847 | 0,339965168 | 0,1390009 | 2,4457766 | 0,0144541 | 0,0469446 | TSPAN4 |
| ENSGALG00000006463 | 6778,1634 | 0,33979504 | 0,1341936 | 2,5321248 | 0,0113374 | 0,0388225 | TMA7 |
| ENSGALG00000000833 | 15942,313 | 0,339708342 | 0,1136284 | 2,9896436 | 0,002793 | 0,0124394 | IK |

| | | | | | | | |
|---|---|---|---|---|---|---|---|
| ENSGALG00000015998 | 9692,1223 | 0,339150714 | 0,1058855 | 3,2029939 | 0,0013601 | 0,0068915 | PTDSS1 |
| ENSGALG00000006056 | 5221,8645 | 0,338801705 | 0,1296 | 2,6142105 | 0,0089434 | 0,0321363 | CHCHD6 |
| ENSGALG00000027609 | 5715,7167 | 0,338322071 | 0,1138079 | 2,9727469 | 0,0029515 | 0,0130039 | RAB40C |
| ENSGALG00000024261 | 6604,4448 | 0,336614329 | 0,1086574 | 3,0979431 | 0,0019487 | 0,0092412 | NHP2 |
| ENSGALG00000001476 | 3822,0948 | 0,335505795 | 0,1349976 | 2,485273 | 0,0129452 | 0,0430927 | RAP1A |
| ENSGALG00000001329 | 7967,7759 | 0,33542437 | 0,1280369 | 2,6197481 | 0,0087995 | 0,0317532 | EPB41 |
| ENSGALG00000012093 | 2809,4561 | 0,335251509 | 0,1300305 | 2,5782533 | 0,0099301 | 0,0349712 | ELMO1 |
| ENSGALG00000002157 | 23751,205 | 0,335162629 | 0,1132861 | 2,958549 | 0,0030909 | 0,013501 | RPS17 |
| ENSGALG00000011476 | 10900,501 | 0,334810497 | 0,1121934 | 2,9842264 | 0,002843 | 0,0125999 | sept-11 |
| ENSGALG00000012277 | 5198,3118 | 0,33449799 | 0,122213 | 2,7370074 | 0,0062001 | 0,0239052 | MAFF |
| ENSGALG00000024033 | 5551,4467 | 0,334154631 | 0,1192958 | 2,8010587 | 0,0050935 | 0,0203689 | PHLDB1 |
| ENSGALG00000012312 | 5618,5193 | 0,333752483 | 0,1164047 | 2,8671731 | 0,0041416 | 0,017158 | GCAT |
| ENSGALG00000007519 | 2801,5468 | 0,333421083 | 0,1296588 | 2,5715271 | 0,0101251 | 0,0355292 | ACBD5 |
| ENSGALG00000009234 | 2846,7839 | 0,332951253 | 0,1239542 | 2,6860825 | 0,0072295 | 0,0271531 | RP11-20I23.3 |
| ENSGALG00000006546 | 6021,0715 | 0,332225604 | 0,1201276 | 2,765605 | 0,0056817 | 0,0222545 | CD81 |
| ENSGALG00000003934 | 3446,0691 | 0,332093035 | 0,1296938 | 2,5605934 | 0,0104494 | 0,0364416 | WIPF2 |
| ENSGALG00000013529 | 4334,8333 | 0,331833816 | 0,1204748 | 2,7543838 | 0,0058803 | 0,02288 | NUP210L |
| ENSGALG00000014236 | 4088,5792 | 0,331720827 | 0,1185166 | 2,7989389 | 0,0051271 | 0,0204647 | ATP8A1 |
| ENSGALG00000001761 | 3252,5626 | 0,33061823 | 0,1263585 | 2,6165091 | 0,0088834 | 0,0319713 | HAUS1 |
| ENSGALG00000000905 | 28204,583 | 0,330518761 | 0,1072649 | 3,0813327 | 0,0020608 | 0,0096783 | PSMB4 |
| ENSGALG00000000055 | 4657,6107 | 0,330211238 | 0,1218412 | 2,7101775 | 0,0067247 | 0,0256032 | LAMTOR3 |
| ENSGALG00000002967 | 2956,1697 | 0,329951632 | 0,1271164 | 2,5956654 | 0,0094408 | 0,0335537 | ZNF365 |
| ENSGALG00000008457 | 3707,8992 | 0,329256556 | 0,1280479 | 2,5713549 | 0,0101301 | 0,0355377 | RIN2 |
| ENSGALG00000015844 | 18562,8 | 0,329214954 | 0,1087925 | 3,0260806 | 0,0024775 | 0,0112473 | SOD1 |
| ENSGALG00000000629 | 2184,8298 | 0,329174801 | 0,1291076 | 2,5496155 | 0,0107842 | 0,0373494 | |
| ENSGALG00000010109 | 8235,1722 | 0,328812909 | 0,1134692 | 2,8978173 | 0,0037577 | 0,0158984 | RNF220 |
| ENSGALG00000029052 | 4904,723 | 0,328574855 | 0,1134688 | 2,8957295 | 0,0037828 | 0,0159748 | CERS2L1 |
| ENSGALG00000003331 | 9717,1633 | 0,328559948 | 0,1145187 | 2,8690497 | 0,0041171 | 0,0171 | FAF2 |
| ENSGALG00000002919 | 3330,9646 | 0,328149978 | 0,1337566 | 2,4533361 | 0,0141538 | 0,0462009 | MON1A |
| ENSGALG00000007272 | 8940,9306 | 0,327852784 | 0,1056422 | 3,1034251 | 0,0019129 | 0,0090938 | RNF166 |
| ENSGALG00000001604 | 7206,5279 | 0,327620067 | 0,1148758 | 2,8519514 | 0,0043452 | 0,0178353 | |
| ENSGALG00000010441 | 3362,4409 | 0,327510367 | 0,1300487 | 2,5183664 | 0,0117901 | 0,0400403 | GYG1 |
| ENSGALG00000008966 | 9784,0345 | 0,327459196 | 0,1078303 | 3,0368019 | 0,002391 | 0,0109204 | FBXW5 |
| ENSGALG00000006439 | 2469,3771 | 0,325297261 | 0,1285361 | 2,5307849 | 0,0113808 | 0,0389417 | ARHGEF6 |
| ENSGALG00000004000 | 3131,3983 | 0,325159334 | 0,1227269 | 2,6494547 | 0,0080622 | 0,0295947 | FLOT2 |
| ENSGALG00000008925 | 2847,1933 | 0,32505073 | 0,1271866 | 2,555699 | 0,0105975 | 0,0368637 | TMEM203 |
| ENSGALG00000006161 | 2625,1209 | 0,324498322 | 0,1340451 | 2,4208145 | 0,0154858 | 0,0496386 | CHMP1A |
| ENSGALG00000004899 | 7704,948 | 0,323966145 | 0,1311985 | 2,469283 | 0,0135384 | 0,0446201 | PEPD |
| ENSGALG00000013039 | 15959,11 | 0,323619 | 0,1177645 | 2,7480188 | 0,0059957 | 0,0232434 | BID |
| ENSGALG00000014555 | 6185,863 | 0,32349139 | 0,119778 | 2,7007582 | 0,0069182 | 0,0261637 | C12orf57 |
| ENSGALG00000015259 | 6306,083 | 0,322565394 | 0,1124418 | 2,8687313 | 0,0041212 | 0,0171086 | RB1CC1 |
| ENSGALG00000015828 | 4399,085 | 0,322397999 | 0,1225293 | 2,6311901 | 0,0085086 | 0,030896 | BACH1 |
| ENSGALG00000010985 | 3705,9823 | 0,321700662 | 0,1306157 | 2,462956 | 0,0137797 | 0,0451855 | ATG4C |
| ENSGALG00000028499 | 5696,2423 | 0,321645303 | 0,1148381 | 2,8008589 | 0,0050967 | 0,0203732 | FAM63A |
| ENSGALG00000013738 | 7162,7044 | 0,321523306 | 0,1318588 | 2,4383909 | 0,0147528 | 0,0477354 | OPTN |
| ENSGALG00000025781 | 5021,0191 | 0,321200342 | 0,1135034 | 2,8298746 | 0,0046566 | 0,0188803 | CERS2L2 |
| ENSGALG00000028937 | 3780,2165 | 0,321152786 | 0,1241263 | 2,5873067 | 0,0096729 | 0,0342248 | TIMM13 |
| ENSGALG00000008443 | 36760,133 | 0,321038212 | 0,1010416 | 3,1772886 | 0,0014866 | 0,0073788 | ARHGDIA |
| ENSGALG00000003513 | 5800,0504 | 0,320426255 | 0,1140656 | 2,8091408 | 0,0049674 | 0,0199796 | KLHL20 |
| ENSGALG00000014682 | 2661,1178 | 0,320335943 | 0,1298503 | 2,4669635 | 0,0136264 | 0,044845 | CAST |
| ENSGALG00000005701 | 4780,9727 | 0,319914204 | 0,1181292 | 2,708173 | 0,0067655 | 0,0257069 | IP6K2 |
| ENSGALG00000016172 | 54051,231 | 0,319415143 | 0,1086593 | 2,9396034 | 0,0032863 | 0,0141682 | RPLP1 |
| ENSGALG00000012843 | 3927,8364 | 0,319142472 | 0,131197 | 2,4325431 | 0,0149932 | 0,0483612 | GMDS |
| ENSGALG00000015664 | 24898,273 | 0,319058165 | 0,1003222 | 3,1803344 | 0,0014711 | 0,0073203 | ADD1 |
| ENSGALG00000012546 | 35449,688 | 0,318249627 | 0,099977 | 3,1832278 | 0,0014564 | 0,0072688 | MCM5 |
| ENSGALG00000011192 | 4976,6178 | 0,317451368 | 0,1192137 | 2,6628766 | 0,0077476 | 0,0287264 | PHF10 |
| ENSGALG00000002595 | 4724,8428 | 0,316793207 | 0,1219683 | 2,5973403 | 0,0093949 | 0,0334059 | |
| ENSGALG00000011238 | 46142,463 | 0,316616268 | 0,1021796 | 3,0986245 | 0,0019442 | 0,0092232 | WLS |
| ENSGALG00000013242 | 4048,7621 | 0,316427862 | 0,1207163 | 2,621252 | 0,0087607 | 0,0316302 | MRPL24 |
| ENSGALG00000002891 | 7363,8293 | 0,315737267 | 0,1107929 | 2,8497977 | 0,0043747 | 0,0179348 | ACADL |
| ENSGALG00000023308 | 5464,7909 | 0,314498813 | 0,1190501 | 2,6417343 | 0,0082483 | 0,0301558 | NPB |
| ENSGALG00000020078 | 34717,171 | 0,313921783 | 0,0988798 | 3,174781 | 0,0014995 | 0,007432 | |
| ENSGALG00000028600 | 142500,09 | 0,313870865 | 0,0985585 | 3,1846146 | 0,0014495 | 0,00725 | |
| ENSGALG00000011106 | 5264,4735 | 0,312738357 | 0,1159112 | 2,698085 | 0,006974 | 0,0263167 | GALNT2 |

| | | | | | | | |
|---|---|---|---|---|---|---|---|
| ENSGALG00000006880 | 22127,188 | 0,310525591 | 0,1109413 | 2,7990071 | 0,005126 | 0,0204647 | SLC38A10 |
| ENSGALG00000011958 | 2755,4733 | 0,30674184 | 0,1239537 | 2,4746489 | 0,0133367 | 0,0441477 | TEF |
| ENSGALG00000008586 | 62912,793 | 0,306372777 | 0,1047411 | 2,9250486 | 0,003444 | 0,0147454 | TUBB4B |
| ENSGALG00000003691 | 21366,038 | 0,306291187 | 0,1008897 | 3,0359009 | 0,0023982 | 0,0109384 | RNF145 |
| ENSGALG00000014190 | 4272,095 | 0,305976217 | 0,1248086 | 2,4515639 | 0,0142237 | 0,0463753 | PNPLA3 |
| ENSGALG00000003428 | 8918,8411 | 0,30596567 | 0,1074998 | 2,8461972 | 0,0044245 | 0,0181062 | PIK3R2 |
| ENSGALG00000010997 | 7028,5315 | 0,305349661 | 0,119889 | 2,5469371 | 0,0108673 | 0,037606 | DFNA5 |
| ENSGALG00000001024 | 11719,39 | 0,304587459 | 0,1095728 | 2,779773 | 0,0054397 | 0,0215357 | TPRG1L |
| ENSGALG00000004898 | 4936,2049 | 0,304436049 | 0,1215502 | 2,5046118 | 0,0122586 | 0,0412712 | |
| ENSGALG00000015273 | 5611,3118 | 0,304040992 | 0,1190879 | 2,5530798 | 0,0106775 | 0,0370663 | TBC1D23 |
| ENSGALG00000000606 | 4262,8792 | 0,302883764 | 0,1188851 | 2,5477019 | 0,0108435 | 0,0375373 | ARL8A |
| ENSGALG00000002712 | 8556,0623 | 0,302868054 | 0,1070979 | 2,8279558 | 0,0046846 | 0,0189881 | LIPG |
| ENSGALG00000014118 | 12062,787 | 0,302595823 | 0,1052318 | 2,8755167 | 0,0040337 | 0,0167998 | MRPS26 |
| ENSGALG00000000094 | 5913,4414 | 0,30200193 | 0,1190933 | 2,5358428 | 0,0112177 | 0,0385193 | ADIPOR1 |
| ENSGALG00000000970 | 7143,9739 | 0,300853659 | 0,1087984 | 2,7652385 | 0,0056881 | 0,0222705 | CHEK1 |
| ENSGALG00000006876 | 3198,9684 | 0,300817961 | 0,1240621 | 2,4247362 | 0,0153195 | 0,0491923 | CTSA |
| ENSGALG00000015211 | 3091,6006 | 0,300524293 | 0,1216505 | 2,4703913 | 0,0134965 | 0,0445036 | DTNA |
| ENSGALG00000004156 | 5252,3729 | 0,300005854 | 0,114727 | 2,6149543 | 0,0089239 | 0,0320918 | KIF1BP |
| ENSGALG00000003973 | 3666,163 | 0,297790777 | 0,1193872 | 2,4943284 | 0,0126196 | 0,0422982 | TLCD1 |
| ENSGALG00000027838 | 4595,5664 | 0,29706177 | 0,1173947 | 2,5304523 | 0,0113916 | 0,0389591 | HMG20B |
| ENSGALG00000002279 | 6077,2003 | 0,297008819 | 0,1136905 | 2,6124335 | 0,00899 | 0,0322612 | CTD-2116N17.1 |
| ENSGALG00000017249 | 12265,651 | 0,295667982 | 0,1084027 | 2,7274952 | 0,0063817 | 0,0244959 | PCF11 |
| ENSGALG00000009570 | 6991,744 | 0,29562858 | 0,1160829 | 2,5467029 | 0,0108746 | 0,0376163 | ATP6V1D |
| ENSGALG00000001913 | 13013,108 | 0,29356182 | 0,1209326 | 2,4274828 | 0,015204 | 0,048879 | DYNLRB1 |
| ENSGALG00000000944 | 3984,6591 | 0,29296383 | 0,1166894 | 2,5106287 | 0,0120516 | 0,0407356 | SUPT4H1 |
| ENSGALG00000014408 | 16252,764 | 0,292920829 | 0,1032929 | 2,8358262 | 0,0045707 | 0,0185874 | LLGL1 |
| ENSGALG00000023975 | 11745,641 | 0,291584025 | 0,1045871 | 2,7879551 | 0,0053042 | 0,0210728 | SLC25A39 |
| ENSGALG00000028356 | 9980,9657 | 0,288424278 | 0,1104192 | 2,6120845 | 0,0089992 | 0,0322857 | LMAN2 |
| ENSGALG00000006904 | 13921,155 | 0,287723415 | 0,1047262 | 2,7473884 | 0,0060072 | 0,0232815 | RNH1 |
| ENSGALG00000008865 | 4561,3827 | 0,286783741 | 0,1175361 | 2,4399621 | 0,0146888 | 0,0475709 | HDHD3 |
| ENSGALG00000013341 | 5963,8498 | 0,285851588 | 0,116546 | 2,4526925 | 0,0141791 | 0,0462726 | AGAP3 |
| ENSGALG00000007250 | 25520,219 | 0,285601509 | 0,1086627 | 2,6283308 | 0,0085805 | 0,0311198 | SNRPB |
| ENSGALG00000021242 | 6224,5764 | 0,284935514 | 0,1141595 | 2,4959426 | 0,0125623 | 0,0421581 | PHPT1 |
| ENSGALG00000004410 | 6738,2931 | 0,28304622 | 0,1082646 | 2,6143925 | 0,0089386 | 0,0321276 | ANAPC16 |
| ENSGALG00000006073 | 7325,56 | 0,281547021 | 0,1095092 | 2,5709901 | 0,0101408 | 0,0355602 | NDUFAB1 |
| ENSGALG00000009940 | 12012,29 | 0,281084158 | 0,1035397 | 2,7147474 | 0,0066326 | 0,0253092 | SLC35E3 |
| ENSGALG00000014800 | 8363,7154 | 0,280973212 | 0,1136279 | 2,4727492 | 0,0134078 | 0,0443399 | ATP6V1A |
| ENSGALG00000014467 | 5734,7231 | 0,280861046 | 0,1140326 | 2,462989 | 0,0137784 | 0,0451855 | COPS7A |
| ENSGALG00000006917 | 6209,9134 | 0,280478545 | 0,1125356 | 2,4923538 | 0,01269 | 0,0424526 | CHKA |
| ENSGALG00000001026 | 9235,9335 | 0,279930511 | 0,1055048 | 2,65325 | 0,0079721 | 0,0293748 | EPB41L1 |
| ENSGALG00000007260 | 8065,1993 | 0,279432916 | 0,1094636 | 2,5527474 | 0,0106877 | 0,0370923 | ZEB1 |
| ENSGALG00000014004 | 8156,8334 | 0,27938572 | 0,1132363 | 2,4672797 | 0,0136144 | 0,0448342 | EPB41L2 |
| ENSGALG00000002546 | 9375,5325 | 0,275701764 | 0,1107787 | 2,4887613 | 0,0128189 | 0,0428188 | COL5A1 |
| ENSGALG00000005138 | 8773,7935 | 0,274876887 | 0,1065601 | 2,5795489 | 0,0098929 | 0,0348493 | KCTD10 |
| ENSGALG00000003625 | 7388,448 | 0,273140927 | 0,1127098 | 2,4233993 | 0,015376 | 0,0493388 | NDUFS5 |
| ENSGALG00000006611 | 7163,1018 | 0,272646564 | 0,1103641 | 2,4704288 | 0,0134951 | 0,0445036 | KIF7 |
| ENSGALG00000011916 | 19570,61 | 0,271939056 | 0,112004 | 2,4279397 | 0,0151849 | 0,048829 | SREBF2 |
| ENSGALG00000008149 | 20254,48 | 0,269265878 | 0,1016307 | 2,6494538 | 0,0080622 | 0,0295947 | EWSR1 |
| ENSGALG00000004639 | 22048,421 | 0,266827236 | 0,1006911 | 2,6499598 | 0,0080501 | 0,0295664 | GNAI2 |
| ENSGALG00000008609 | 46235,247 | 0,259800517 | 0,1027436 | 2,5286292 | 0,0114509 | 0,0390835 | PTK7 |
| ENSGALG00000015617 | 50870,723 | 0,25500812 | 0,1021977 | 2,4952431 | 0,0125871 | 0,0422205 | RPS23 |
| ENSGALG00000017299 | 53500,096 | 0,254780179 | 0,1021944 | 2,4930939 | 0,0126635 | 0,0424143 | RPS11 |
| ENSGALG00000011951 | 179564,91 | 0,254345179 | 0,1001273 | 2,5402178 | 0,0110783 | 0,0381466 | RPSA |
| ENSGALG00000008812 | 67809,432 | -0,244632027 | 0,100856 | -2,425557 | 0,0152849 | 0,0491044 | CCT4 |
| ENSGALG00000008038 | 49840,281 | -0,24989979 | 0,1024864 | -2,438371 | 0,0147536 | 0,0477354 | SF3B1 |
| ENSGALG00000010027 | 20404,953 | -0,251221533 | 0,1016636 | -2,471106 | 0,0134696 | 0,0444515 | CFL2 |
| ENSGALG00000009228 | 73654,992 | -0,251350219 | 0,0988688 | -2,54226 | 0,0110138 | 0,0380109 | PARP1 |
| ENSGALG00000001053 | 21507,706 | -0,251351424 | 0,101079 | -2,486682 | 0,0128941 | 0,0429539 | SRPRA |
| ENSGALG00000009309 | 84581,571 | -0,253473054 | 0,1003203 | -2,526638 | 0,011516 | 0,0392467 | EIF5A2 |
| ENSGALG00000028484 | 18420,667 | -0,257706043 | 0,100811 | -2,556328 | 0,0105783 | 0,036816 | |
| ENSGALG00000029167 | 20948,612 | -0,258969024 | 0,102918 | -2,516265 | 0,0118606 | 0,0402497 | THOP1 |
| ENSGALG00000010842 | 26982,926 | -0,259509696 | 0,1041078 | -2,492703 | 0,0126775 | 0,0424402 | UBR7 |
| ENSGALG00000008944 | 11174,776 | -0,259905745 | 0,1056639 | -2,459741 | 0,0139037 | 0,0455156 | FBXO11 |
| ENSGALG00000004040 | 10377,603 | -0,259935536 | 0,107237 | -2,423935 | 0,0153534 | 0,0492777 | MFAP3 |

| | | | | | | | |
|---|---|---|---|---|---|---|---|
| ENSGALG00000015706 | 7834,195 | -0,260107652 | 0,1070306 | -2,430217 | 0,0150898 | 0,0485921 | LETM1 |
| ENSGALG00000008962 | 22921,574 | -0,260994621 | 0,1007267 | -2,591116 | 0,0095665 | 0,0338747 | MSH2 |
| ENSGALG00000004965 | 10149,59 | -0,261538698 | 0,1058067 | -2,471854 | 0,0134414 | 0,044408 | SLC27A4 |
| ENSGALG00000002569 | 85602,751 | -0,262007268 | 0,1060299 | -2,47107 | 0,013471 | 0,0444515 | RAN |
| ENSGALG00000015596 | 26927,118 | -0,262154887 | 0,1027805 | -2,550629 | 0,0107529 | 0,0372615 | MAP3K7 |
| ENSGALG00000015821 | 61741,915 | -0,262392566 | 0,1001176 | -2,620842 | 0,0087713 | 0,0316599 | CCT8 |
| ENSGALG00000002488 | 22309,721 | -0,262864461 | 0,100189 | -2,623685 | 0,0086984 | 0,0314552 | PAIP2 |
| ENSGALG00000008180 | 11420,002 | -0,263317561 | 0,1088104 | -2,419968 | 0,0155219 | 0,0497248 | SPG11 |
| ENSGALG00000008500 | 8629,7439 | -0,263668812 | 0,1056933 | -2,494659 | 0,0126078 | 0,0422692 | RBSN |
| ENSGALG00000012405 | 16329,496 | -0,26384951 | 0,1069456 | -2,467137 | 0,0136198 | 0,0448342 | EZH2 |
| ENSGALG00000001350 | 6586,6747 | -0,264940499 | 0,1094289 | -2,421119 | 0,0154728 | 0,0496142 | RSPRY1 |
| ENSGALG00000001515 | 15194,726 | -0,265107192 | 0,1041998 | -2,54422 | 0,0109522 | 0,0378366 | ATAD3A |
| ENSGALG00000016042 | 8800,9648 | -0,265273818 | 0,1060345 | -2,501769 | 0,0123574 | 0,0415732 | CHAF1B |
| ENSGALG00000000581 | 14682,192 | -0,265993327 | 0,1094668 | -2,4299 | 0,015103 | 0,0486232 | DDX42 |
| ENSGALG00000014330 | 13035,527 | -0,266560386 | 0,1080798 | -2,466329 | 0,0136506 | 0,0448931 | NECAP1 |
| ENSGALG00000011504 | 43957,582 | -0,266735415 | 0,0983735 | -2,711456 | 0,0066988 | 0,0255261 | TMPO |
| ENSGALG00000016226 | 14374,267 | -0,266804523 | 0,10248 | -2,603479 | 0,0092283 | 0,0329512 | CASK |
| ENSGALG00000009781 | 15826,657 | -0,267258812 | 0,1074538 | -2,487198 | 0,0128754 | 0,0429337 | DOCK1 |
| ENSGALG00000009941 | 12598,143 | -0,268249202 | 0,1049288 | -2,556488 | 0,0105735 | 0,0368085 | SLC35B1 |
| ENSGALG00000017118 | 10000,178 | -0,268386575 | 0,1059335 | -2,533537 | 0,0112918 | 0,0387248 | MIPEP |
| ENSGALG00000011553 | 8178,0483 | -0,268405007 | 0,1061309 | -2,528999 | 0,0114389 | 0,0390619 | UTP3 |
| ENSGALG00000016417 | 8419,6417 | -0,268534668 | 0,1066163 | -2,518702 | 0,0117788 | 0,0400122 | DSCC1 |
| ENSGALG00000016894 | 12688,39 | -0,268921468 | 0,1105656 | -2,432233 | 0,015006 | 0,0483911 | DNAJC3 |
| ENSGALG00000005587 | 297190,08 | -0,269468112 | 0,0978414 | -2,754133 | 0,0058848 | 0,0228853 | EIF4G2 |
| ENSGALG00000026111 | 15113,012 | -0,269770034 | 0,1052633 | -2,562812 | 0,0103828 | 0,0362374 | SAAL1 |
| ENSGALG00000006113 | 19999,679 | -0,272100604 | 0,1073343 | -2,535075 | 0,0112423 | 0,0385844 | ZNF326 |
| ENSGALG00000028900 | 8664,7981 | -0,272551766 | 0,1086758 | -2,507935 | 0,0121439 | 0,0409661 | |
| ENSGALG00000010284 | 17075,328 | -0,272708241 | 0,1031857 | -2,642887 | 0,0082202 | 0,0300614 | GMPS |
| ENSGALG00000016953 | 7971,8105 | -0,272857363 | 0,1064399 | -2,563488 | 0,0103626 | 0,0361948 | NAA16 |
| ENSGALG00000012402 | 11643,369 | -0,273460141 | 0,1041025 | -2,626836 | 0,0086183 | 0,0312235 | CUL1 |
| ENSGALG00000005870 | 10634,139 | -0,27364395 | 0,1062314 | -2,575923 | 0,0099973 | 0,0351622 | MCM10 |
| ENSGALG00000014331 | 49106,781 | -0,273875014 | 0,1046006 | -2,618292 | 0,0088371 | 0,0318637 | FKBP4 |
| ENSGALG00000005654 | 13436,846 | -0,27412231 | 0,1031825 | -2,656674 | 0,0078916 | 0,0291333 | PITPNB |
| ENSGALG00000000907 | 34606,894 | -0,274397863 | 0,1051305 | -2,610069 | 0,0090524 | 0,0324512 | KARS |
| ENSGALG00000006974 | 21987,582 | -0,275795791 | 0,1053463 | -2,617992 | 0,0088449 | 0,0318788 | DDX27 |
| ENSGALG00000014439 | 23731,455 | -0,275802443 | 0,1013775 | -2,72055 | 0,0065173 | 0,024948 | NCAPD2 |
| ENSGALG00000012244 | 14183,266 | -0,27619833 | 0,1043027 | -2,648045 | 0,0080959 | 0,0296783 | KLHDC2 |
| ENSGALG00000011130 | 11036,561 | -0,276443798 | 0,1084288 | -2,549541 | 0,0107865 | 0,0373494 | COPS4 |
| ENSGALG00000007491 | 11685,784 | -0,276864687 | 0,1090804 | -2,538171 | 0,0111433 | 0,0383219 | VWA9 |
| ENSGALG00000000856 | 7768,1765 | -0,277333067 | 0,1078018 | -2,57262 | 0,0100932 | 0,0354264 | EIF2D |
| ENSGALG00000017128 | 6812,3227 | -0,278120188 | 0,1105176 | -2,516523 | 0,0118519 | 0,0402303 | SKA3 |
| ENSGALG00000002179 | 5553,9071 | -0,278228201 | 0,1141171 | -2,438094 | 0,0147649 | 0,0477607 | RP11-552F3.12 |
| ENSGALG00000008394 | 17588,746 | -0,278483321 | 0,1028235 | -2,708363 | 0,0067616 | 0,0257069 | THUMPD3 |
| ENSGALG00000007712 | 6385,5881 | -0,27852982 | 0,1114303 | -2,499589 | 0,0124338 | 0,0417989 | MYO1B |
| ENSGALG00000003532 | 126353,26 | -0,278703141 | 0,1005665 | -2,771332 | 0,0055828 | 0,0219743 | DDX5 |
| ENSGALG00000009774 | 8409,19 | -0,279620047 | 0,1118848 | -2,499177 | 0,0124482 | 0,0418268 | ELF2 |
| ENSGALG00000014425 | 11773,73 | -0,280179605 | 0,1054976 | -2,655791 | 0,0079123 | 0,0291781 | NCAPG |
| ENSGALG00000016340 | 46321,016 | -0,280763167 | 0,1135687 | -2,472187 | 0,0134289 | 0,0443774 | EIF2S3L |
| ENSGALG00000012798 | 32319,553 | -0,280818753 | 0,1118437 | -2,510814 | 0,0120453 | 0,0407243 | SSR1 |
| ENSGALG00000016447 | 53614,037 | -0,281195284 | 0,1001808 | -2,806878 | 0,0050024 | 0,0200672 | PDIA6 |
| ENSGALG00000023682 | 6598,1149 | -0,28149273 | 0,1105533 | -2,546216 | 0,0108898 | 0,0376475 | GTF2H3 |
| ENSGALG00000026390 | 10622,058 | -0,281499002 | 0,1082118 | -2,60137 | 0,0092852 | 0,0330938 | BAHD1 |
| ENSGALG00000011762 | 7368,3608 | -0,281679323 | 0,1102252 | -2,55549 | 0,0106038 | 0,03687 | UBA6 |
| ENSGALG00000000726 | 16204,608 | -0,284722338 | 0,1131864 | -2,515517 | 0,0118858 | 0,0403051 | ELAVL1 |
| ENSGALG00000003248 | 40887,68 | -0,284935192 | 0,0989747 | -2,878868 | 0,003991 | 0,0166579 | SUB1 |
| ENSGALG00000011171 | 7789,5801 | -0,285220301 | 0,1097597 | -2,59859 | 0,0093608 | 0,0333194 | TBP |
| ENSGALG00000015285 | 36051,228 | -0,286253589 | 0,1140443 | -2,510021 | 0,0120724 | 0,0407754 | TOMM70 |
| ENSGALG00000014773 | 11518,187 | -0,287175691 | 0,1109758 | -2,587732 | 0,009661 | 0,0342004 | ERBB2IP |
| ENSGALG00000008086 | 20675,956 | -0,2873629 | 0,1034267 | -2,778421 | 0,0054624 | 0,021613 | ACTR1A |
| ENSGALG00000026761 | 29796,866 | -0,287544813 | 0,1163484 | -2,471412 | 0,0134581 | 0,0444413 | LIN28B |
| ENSGALG00000007101 | 5189,3303 | -0,287687091 | 0,1145161 | -2,512197 | 0,0119982 | 0,0406054 | ZPR1 |
| ENSGALG00000000823 | 24312,395 | -0,287692196 | 0,1169353 | -2,460268 | 0,0138833 | 0,0454597 | BAZ1B |
| ENSGALG00000001219 | 4265,8107 | -0,287809408 | 0,1164176 | -2,472216 | 0,0134278 | 0,0443774 | PIGU |
| ENSGALG00000010925 | 7979,9749 | -0,288036245 | 0,1081614 | -2,663022 | 0,0077442 | 0,0287218 | FAM126A |

| | | | | | | | |
|---|---|---|---|---|---|---|---|
| ENSGALG00000005302 | 10087,37 | -0,288116362 | 0,105057 | -2,742476 | 0,0060978 | 0,0235755 | SASS6 |
| ENSGALG00000016238 | 8227,2702 | -0,288804451 | 0,1172255 | -2,463666 | 0,0137524 | 0,0451396 | MED14 |
| ENSGALG00000001277 | 6953,8461 | -0,28888441 | 0,1159131 | -2,492249 | 0,0126937 | 0,0424526 | STAG1 |
| ENSGALG00000003086 | 11675,474 | -0,289022537 | 0,1042361 | -2,772768 | 0,0055582 | 0,0219029 | EXOSC10 |
| ENSGALG00000010829 | 17151,788 | -0,289288545 | 0,1076078 | -2,68836 | 0,0071804 | 0,0270058 | CDCA7 |
| ENSGALG00000014570 | 6323,6704 | -0,289387372 | 0,1174534 | -2,463847 | 0,0137455 | 0,0451277 | LPCAT3 |
| ENSGALG00000010318 | 35384,519 | -0,289579204 | 0,1116994 | -2,592487 | 0,0095285 | 0,0337839 | TMED10 |
| ENSGALG00000015804 | 7311,6424 | -0,289600409 | 0,1113075 | -2,601805 | 0,0092734 | 0,0330778 | LTN1 |
| ENSGALG00000003355 | 8990,1272 | -0,289601526 | 0,1161151 | -2,49409 | 0,0126281 | 0,0423163 | NOP16 |
| ENSGALG00000008095 | 17234,199 | -0,289644341 | 0,113924 | -2,542434 | 0,0110083 | 0,0380016 | KIF23 |
| ENSGALG00000005769 | 9850,6081 | -0,289680709 | 0,1121661 | -2,582606 | 0,0098057 | 0,0346586 | WEE1 |
| ENSGALG00000010167 | 23334,849 | -0,289818191 | 0,1045631 | -2,771707 | 0,0055763 | 0,0219585 | PNN |
| ENSGALG00000003194 | 7179,1586 | -0,290381921 | 0,1074323 | -2,702929 | 0,0068731 | 0,0260208 | ATP6V0A2 |
| ENSGALG00000011304 | 6781,2258 | -0,290613007 | 0,1160055 | -2,505165 | 0,0122394 | 0,0412267 | NGLY1 |
| ENSGALG00000005858 | 11910,812 | -0,290886069 | 0,1040132 | -2,796626 | 0,0051639 | 0,0205937 | DR1 |
| ENSGALG00000006706 | 5131,464 | -0,290887314 | 0,1158194 | -2,511558 | 0,0120199 | 0,0406486 | USP6NL |
| ENSGALG00000013369 | 5721,3826 | -0,290969648 | 0,1118474 | -2,601488 | 0,009282 | 0,0330933 | NCLN |
| ENSGALG00000001443 | 24266,064 | -0,291025391 | 0,1058054 | -2,750573 | 0,0059491 | 0,0231222 | PTPRU |
| ENSGALG00000014854 | 16295,817 | -0,291513975 | 0,1167727 | -2,496423 | 0,0125453 | 0,0421114 | MYL12A |
| ENSGALG00000006934 | 7206,0363 | -0,291528022 | 0,1149418 | -2,536309 | 0,0112028 | 0,0384778 | PATZ1 |
| ENSGALG00000007254 | 3754,7945 | -0,291530428 | 0,1205128 | -2,419082 | 0,0155597 | 0,0498226 | ARHGAP12 |
| ENSGALG00000006884 | 8596,5657 | -0,291624168 | 0,113545 | -2,568359 | 0,0102181 | 0,0357819 | mars-05 |
| ENSGALG00000015120 | 9551,3619 | -0,291739099 | 0,116855 | -2,49659 | 0,0125394 | 0,042102 | QTRT2 |
| ENSGALG00000001438 | 13509,485 | -0,292098905 | 0,1078016 | -2,709598 | 0,0067365 | 0,0256277 | WDR77 |
| ENSGALG00000015483 | 16683,55 | -0,292201948 | 0,1099352 | -2,657946 | 0,0078618 | 0,029063 | PNISR |
| ENSGALG00000002341 | 9767,8137 | -0,292212762 | 0,1068419 | -2,735002 | 0,006238 | 0,0240255 | AP3S1 |
| ENSGALG00000001733 | 10762,095 | -0,292618206 | 0,1060051 | -2,760415 | 0,0057728 | 0,0225205 | LRIG2 |
| ENSGALG00000015933 | 13856,149 | -0,292842928 | 0,1142604 | -2,562944 | 0,0103789 | 0,036237 | PAXBP1 |
| ENSGALG00000007516 | 13969,274 | -0,293041961 | 0,1052871 | -2,783265 | 0,0053815 | 0,0213301 | RP11-449H3.3 |
| ENSGALG00000002136 | 3591,7129 | -0,293341559 | 0,1202128 | -2,440185 | 0,0146797 | 0,0475641 | AP3B2 |
| ENSGALG00000016106 | 14612,023 | -0,293596343 | 0,1087124 | -2,700671 | 0,00692 | 0,0261637 | NUDCD1 |
| ENSGALG00000013424 | 3938,6768 | -0,293681861 | 0,1183319 | -2,481848 | 0,0130703 | 0,0434666 | TULP3 |
| ENSGALG00000008529 | 7562,5902 | -0,293829308 | 0,1095259 | -2,682737 | 0,0073022 | 0,0273583 | FASTKD2 |
| ENSGALG00000001175 | 5051,078 | -0,294210197 | 0,1151299 | -2,555462 | 0,0106047 | 0,03687 | C4BPA |
| ENSGALG00000000890 | 14838,887 | -0,29430607 | 0,1074987 | -2,737765 | 0,0061858 | 0,0238651 | SRPK1 |
| ENSGALG00000001503 | 12542,871 | -0,294569539 | 0,1129054 | -2,608993 | 0,0090809 | 0,0325017 | TBCD |
| ENSGALG00000005927 | 7896,7145 | -0,295114532 | 0,1102438 | -2,676927 | 0,0074301 | 0,0277608 | ST5 |
| ENSGALG00000016006 | 15900,752 | -0,295193317 | 0,10687 | -2,762171 | 0,0057418 | 0,0224255 | ITSN1 |
| ENSGALG00000011392 | 3156,1633 | -0,295432379 | 0,1200153 | -2,461624 | 0,013831 | 0,045321 | MSH4 |
| ENSGALG00000001301 | 23007,802 | -0,295608227 | 0,1064113 | -2,777977 | 0,0054699 | 0,0216363 | CLTCL1 |
| ENSGALG00000003262 | 4105,2449 | -0,296160558 | 0,1196838 | -2,474525 | 0,0133413 | 0,0441513 | VAMP4 |
| ENSGALG00000002134 | 7387,1514 | -0,296314687 | 0,1196419 | -2,47668 | 0,0132611 | 0,0439509 | THUMPD1 |
| ENSGALG00000016270 | 5777,0167 | -0,296663159 | 0,109934 | -2,698557 | 0,0069641 | 0,0262954 | EFR3A |
| ENSGALG00000002670 | 23872,138 | -0,297199956 | 0,1006311 | -2,953362 | 0,0031433 | 0,0136686 | ZC3H15 |
| ENSGALG00000008094 | 100007,59 | -0,297620991 | 0,1010893 | -2,94414 | 0,0032385 | 0,0140066 | HSPD1 |
| ENSGALG00000010787 | 8044,4352 | -0,297642358 | 0,1129413 | -2,635371 | 0,0084045 | 0,030612 | TTC4 |
| ENSGALG00000016203 | 5413,6009 | -0,297788719 | 0,1132631 | -2,629177 | 0,0085592 | 0,0310507 | KHDRBS3 |
| ENSGALG00000017134 | 4848,4656 | -0,298805127 | 0,1142181 | -2,616093 | 0,0088942 | 0,0320019 | IFT88 |
| ENSGALG00000023703 | 6517,2936 | -0,298876557 | 0,1228202 | -2,433448 | 0,0149558 | 0,0482633 | GIT1 |
| ENSGALG00000006646 | 12074,844 | -0,299111546 | 0,108755 | -2,750324 | 0,0059536 | 0,0231332 | ABCC1 |
| ENSGALG00000008325 | 37371,278 | -0,299659865 | 0,1093847 | -2,739505 | 0,0061532 | 0,0237661 | CKAP5 |
| ENSGALG00000011445 | 6835,5845 | -0,299863462 | 0,1227479 | -2,442921 | 0,0145689 | 0,0472501 | AAMP |
| ENSGALG00000013382 | 5584,8468 | -0,299878268 | 0,1168395 | -2,566582 | 0,0102706 | 0,0359195 | SCO1 |
| ENSGALG00000012561 | 14201,494 | -0,299998657 | 0,1061635 | -2,825817 | 0,004716 | 0,0190927 | PKP4 |
| ENSGALG00000010120 | 15856,807 | -0,300036521 | 0,1036618 | -2,894379 | 0,0037991 | 0,0160188 | KIF2C |
| ENSGALG00000010818 | 4623,916 | -0,300127816 | 0,1207061 | -2,486435 | 0,012903 | 0,0429732 | PLPP3 |
| ENSGALG00000010619 | 3449,0695 | -0,300249899 | 0,1235308 | -2,430566 | 0,0150752 | 0,0485568 | CC2D1B |
| ENSGALG00000001964 | 6474,3266 | -0,300692826 | 0,114042 | -2,636685 | 0,0083721 | 0,0305182 | INPP5E |
| ENSGALG00000014370 | 5230,6919 | -0,300941728 | 0,1119644 | -2,687833 | 0,0071917 | 0,027041 | SEL1L3 |
| ENSGALG00000010275 | 8613,432 | -0,301228493 | 0,1133946 | -2,656462 | 0,0078965 | 0,0291438 | FCF1 |
| ENSGALG00000015611 | 5412,5223 | -0,301903565 | 0,1149449 | -2,626507 | 0,0086266 | 0,0312454 | CASP8AP2 |
| ENSGALG00000007351 | 8585,3707 | -0,301983 | 0,1078117 | -2,801023 | 0,0050941 | 0,0203689 | MTPAP |
| ENSGALG00000014248 | 12329,033 | -0,30224682 | 0,1037621 | -2,912883 | 0,0035811 | 0,0152603 | TMEM33 |
| ENSGALG00000011301 | 5632,1725 | -0,302449789 | 0,1176772 | -2,570164 | 0,010165 | 0,0356142 | LRRC40 |

| | | | | | | | |
|---|---|---|---|---|---|---|---|
| ENSGALG00000005786 | 13742,557 | -0,302474038 | 0,1235613 | -2,447967 | 0,0143665 | 0,046727 | DNTTIP2 |
| ENSGALG00000016872 | 5074,8717 | -0,30261365 | 0,1222955 | -2,474446 | 0,0133443 | 0,0441513 | PCCA |
| ENSGALG00000009990 | 5003,3757 | -0,302651446 | 0,112748 | -2,684317 | 0,0072678 | 0,0272594 | SRBD1 |
| ENSGALG00000005054 | 41673,988 | -0,303560897 | 0,1093589 | -2,775822 | 0,0055062 | 0,0217675 | DKC1 |
| ENSGALG00000005525 | 43269,502 | -0,303682785 | 0,1060484 | -2,863624 | 0,0041882 | 0,0173217 | SRSF1 |
| ENSGALG00000013573 | 4217,1955 | -0,303884724 | 0,1221979 | -2,486825 | 0,0128889 | 0,0429494 | ALG2 |
| ENSGALG00000020454 | 8238,8869 | -0,304067404 | 0,1241364 | -2,449462 | 0,014307 | 0,0465892 | C5H14ORF169 |
| ENSGALG00000001726 | 4862,9398 | -0,304268782 | 0,1143657 | -2,660491 | 0,0078027 | 0,0288914 | NUDCD2 |
| ENSGALG00000007102 | 6521,5758 | -0,304336302 | 0,1232551 | -2,469158 | 0,0135432 | 0,0446249 | MLEC |
| ENSGALG00000000627 | 7600,7154 | -0,305117543 | 0,1084461 | -2,81354 | 0,0048999 | 0,019755 | DNAJC11 |
| ENSGALG00000026838 | 4227,5246 | -0,305207931 | 0,1214042 | -2,513982 | 0,0119377 | 0,0404206 | MDFIC |
| ENSGALG00000003052 | 7714,0905 | -0,30523634 | 0,1156088 | -2,640252 | 0,0082844 | 0,0302474 | AGFG1 |
| ENSGALG00000006418 | 20583,346 | -0,305254291 | 0,1100474 | -2,773843 | 0,0055398 | 0,0218623 | PDPK1 |
| ENSGALG00000015013 | 17595,102 | -0,305329384 | 0,1094445 | -2,789809 | 0,0052739 | 0,0209647 | MSX1 |
| ENSGALG00000006488 | 10099,124 | -0,305368261 | 0,111059 | -2,749605 | 0,0059667 | 0,0231774 | RYK |
| ENSGALG00000009856 | 9272,2302 | -0,305500873 | 0,1237141 | -2,46941 | 0,0135336 | 0,044615 | GNS |
| ENSGALG00000001007 | 27355,205 | -0,305959563 | 0,1072059 | -2,853942 | 0,004318 | 0,017756 | MMP15 |
| ENSGALG00000015465 | 4968,7172 | -0,305990751 | 0,1148791 | -2,66359 | 0,0077312 | 0,0286811 | ASCC3 |
| ENSGALG00000001267 | 9421,3215 | -0,306647481 | 0,1081863 | -2,83444 | 0,0045906 | 0,0186403 | MAP2K2 |
| ENSGALG00000013064 | 12429,63 | -0,306789618 | 0,1032068 | -2,972572 | 0,0029532 | 0,0130039 | NSUN2 |
| ENSGALG00000010042 | 7954,106 | -0,30721043 | 0,1081752 | -2,839935 | 0,0045123 | 0,0183881 | CNOT2 |
| ENSGALG00000007307 | 30782,066 | -0,307584455 | 0,1030681 | -2,984284 | 0,0028424 | 0,0125999 | USP7 |
| ENSGALG00000015331 | 7117,5258 | -0,308171718 | 0,1237599 | -2,490077 | 0,0127715 | 0,0426815 | GNE |
| ENSGALG00000012109 | 33178,661 | -0,308358257 | 0,1122444 | -2,747204 | 0,0060106 | 0,023288 | EIF3M |
| ENSGALG00000017073 | 7452,5062 | -0,308364987 | 0,1160232 | -2,657786 | 0,0078656 | 0,0290689 | BRCA2 |
| ENSGALG00000006445 | 15242,685 | -0,308553523 | 0,1035516 | -2,979708 | 0,0028852 | 0,012754 | ARNT2 |
| ENSGALG00000003373 | 12183,365 | -0,309311012 | 0,1178882 | -2,623766 | 0,0086964 | 0,0314552 | BRIX1 |
| ENSGALG00000000990 | 39372,44 | -0,309348416 | 0,12727 | -2,430646 | 0,0150719 | 0,0485568 | SRSF6 |
| ENSGALG00000012936 | 8859,5776 | -0,309500805 | 0,1175868 | -2,632105 | 0,0084858 | 0,0308336 | KIAA1551 |
| ENSGALG00000020164 | 4566,1787 | -0,31008137 | 0,1239655 | -2,501353 | 0,012372 | 0,0416118 | GAR1 |
| ENSGALG00000001504 | 8296,3292 | -0,31053349 | 0,1111444 | -2,793964 | 0,0052066 | 0,0207336 | DDX20 |
| ENSGALG00000010961 | 89733,227 | -0,310634298 | 0,100957 | -3,076896 | 0,0020917 | 0,0098032 | IGF2BP3 |
| ENSGALG00000008211 | 5098,0335 | -0,311240889 | 0,1149304 | -2,708081 | 0,0067673 | 0,0257069 | ACP2 |
| ENSGALG00000003815 | 34063,921 | -0,31144656 | 0,1009753 | -3,084385 | 0,0020397 | 0,0095994 | RPN2 |
| ENSGALG00000007905 | 37631,106 | -0,311643664 | 0,1126417 | -2,766681 | 0,005663 | 0,0222004 | LAMB1 |
| ENSGALG00000016267 | 13457,673 | -0,312264026 | 0,115716 | -2,698538 | 0,0069645 | 0,0262954 | PHF3 |
| ENSGALG00000010344 | 30762,544 | -0,312514856 | 0,1018116 | -3,069541 | 0,0021439 | 0,0100027 | DHX36 |
| ENSGALG00000001848 | 13173,32 | -0,312572754 | 0,1053696 | -2,96644 | 0,0030127 | 0,0132102 | UBE2J2 |
| ENSGALG00000002106 | 4264,2615 | -0,312942638 | 0,1236027 | -2,531844 | 0,0113465 | 0,0388341 | CLCC1 |
| ENSGALG00000013462 | 2483,9812 | -0,3130983 | 0,1273973 | -2,457652 | 0,0139849 | 0,0457372 | STX17 |
| ENSGALG00000008714 | 4292,5321 | -0,313343892 | 0,1254822 | -2,497119 | 0,0125207 | 0,04206 | ABI2 |
| ENSGALG00000011797 | 18609,134 | -0,313395866 | 0,1178867 | -2,65845 | 0,0078501 | 0,0290274 | MTHFD1 |
| ENSGALG00000007089 | 17474,95 | -0,313684689 | 0,1041053 | -3,013149 | 0,0025855 | 0,0116254 | KDELR2 |
| ENSGALG00000005647 | 9129,0785 | -0,313707238 | 0,1081392 | -2,900958 | 0,0037202 | 0,0157595 | ABCD3 |
| ENSGALG00000016736 | 23809,889 | -0,314343681 | 0,1027519 | -3,05925 | 0,0022189 | 0,0102792 | CYFIP1 |
| ENSGALG00000001973 | 14919,644 | -0,314418391 | 0,103095 | -3,049793 | 0,00229 | 0,0105372 | GNL2 |
| ENSGALG00000007410 | 10084,856 | -0,314498411 | 0,1237962 | -2,540453 | 0,0110709 | 0,0381306 | IGDCC4 |
| ENSGALG00000012827 | 3045,2609 | -0,314591513 | 0,12464 | -2,524001 | 0,0116028 | 0,0394908 | RIPK1 |
| ENSGALG00000016072 | 4693,3951 | -0,314891598 | 0,1172976 | -2,684553 | 0,0072627 | 0,0272551 | SLC25A32 |
| ENSGALG00000003419 | 3186,2022 | -0,314999563 | 0,1269361 | -2,481561 | 0,0130808 | 0,0434804 | MPHOSPH9 |
| ENSGALG00000016893 | 6244,0698 | -0,315224782 | 0,1174264 | -2,684447 | 0,007265 | 0,0272563 | UGGT2 |
| ENSGALG00000006356 | 8494,9234 | -0,315726233 | 0,1090464 | -2,895338 | 0,0037875 | 0,0159897 | MKLN1 |
| ENSGALG00000013174 | 3418,9295 | -0,315789129 | 0,1225684 | -2,576431 | 0,0099826 | 0,0351244 | RECQL |
| ENSGALG00000003948 | 5405,0125 | -0,315917308 | 0,1189483 | -2,65592 | 0,0079092 | 0,0291748 | ALAS1 |
| ENSGALG00000010115 | 2409,6668 | -0,316246717 | 0,1295797 | -2,440558 | 0,0146646 | 0,0475264 | SCLT1 |
| ENSGALG00000009305 | 39225,132 | -0,31625936 | 0,1073859 | -2,945072 | 0,0032288 | 0,0139777 | LBR |
| ENSGALG00000014378 | 6758,1006 | -0,3165666 | 0,1112178 | -2,846367 | 0,0044221 | 0,018102 | ANAPC4 |
| ENSGALG00000025996 | 45890,797 | -0,316767046 | 0,1146038 | -2,76402 | 0,0057094 | 0,0223244 | MKI67 |
| ENSGALG00000009438 | 13859,101 | -0,316803968 | 0,1103871 | -2,869936 | 0,0041056 | 0,0170626 | SLC39A9 |
| ENSGALG00000002158 | 4001,6826 | -0,31684048 | 0,1201134 | -2,637844 | 0,0083435 | 0,0304223 | GOLGA3 |
| ENSGALG00000004375 | 15903,904 | -0,317187722 | 0,1029166 | -3,081989 | 0,0020562 | 0,0096636 | VPS35 |
| ENSGALG00000005518 | 5338,317 | -0,317368673 | 0,1224356 | -2,592127 | 0,0095385 | 0,0338105 | ADRBK2 |
| ENSGALG00000005567 | 4701,7613 | -0,317423466 | 0,1173805 | -2,704227 | 0,0068464 | 0,0259502 | |
| ENSGALG00000016056 | 26843,51 | -0,317775278 | 0,1063633 | -2,98764 | 0,0028114 | 0,0124926 | AZIN1 |

| | | | | | | | |
|---|---|---|---|---|---|---|---|
| ENSGALG00000003753 | 2677,8795 | -0,317987251 | 0,1281857 | -2,480677 | 0,0131133 | 0,0435671 | CYLD |
| ENSGALG00000028935 | 5415,5214 | -0,318166766 | 0,1109111 | -2,868666 | 0,0041221 | 0,0171086 | BCL10 |
| ENSGALG00000016243 | 13413,478 | -0,318335491 | 0,1120682 | -2,840551 | 0,0045036 | 0,0183581 | PHF20L1 |
| ENSGALG00000006525 | 5626,0083 | -0,318337533 | 0,1236191 | -2,575147 | 0,0100197 | 0,035223 | NCAPG2 |
| ENSGALG00000008045 | 3399,549 | -0,318998276 | 0,1242709 | -2,566959 | 0,0102595 | 0,0359082 | SLC9A8 |
| ENSGALG00000010748 | 12682,351 | -0,319116199 | 0,1031995 | -3,092226 | 0,0019866 | 0,0093851 | EXO1 |
| ENSGALG00000006861 | 3243,9283 | -0,319421528 | 0,1239662 | -2,576682 | 0,0099754 | 0,0351124 | COQ7 |
| ENSGALG00000006400 | 5333,2312 | -0,319435266 | 0,1204467 | -2,652088 | 0,0079996 | 0,0294363 | LMBR1 |
| ENSGALG00000004283 | 4377,1001 | -0,319446156 | 0,1200243 | -2,661512 | 0,007779 | 0,0288274 | GCOM1 |
| ENSGALG00000009854 | 6064,675 | -0,319466559 | 0,1106787 | -2,886432 | 0,0038964 | 0,016358 | NEK2 |
| ENSGALG00000003294 | 7834,3924 | -0,31979622 | 0,1079493 | -2,962467 | 0,0030518 | 0,0133647 | ATAD5 |
| ENSGALG00000010312 | 8288,8613 | -0,319830197 | 0,1182081 | -2,705655 | 0,006817 | 0,0258605 | PLCH1 |
| ENSGALG00000000847 | 5954,7792 | -0,320140743 | 0,1166802 | -2,743746 | 0,0060743 | 0,0234946 | GARNL3 |
| ENSGALG00000015983 | 15093,436 | -0,320176555 | 0,1272555 | -2,516013 | 0,0118691 | 0,0402685 | GART |
| ENSGALG00000012160 | 7671,7979 | -0,320265774 | 0,1148045 | -2,789663 | 0,0052763 | 0,0209681 | KIF18A |
| ENSGALG00000005139 | 53456,399 | -0,320485908 | 0,1129243 | -2,838059 | 0,0045389 | 0,0184744 | CLTC |
| ENSGALG00000011568 | 15058,649 | -0,320702144 | 0,1071903 | -2,991896 | 0,0027725 | 0,0123601 | GRSF1 |
| ENSGALG00000010677 | 4086,7271 | -0,320970598 | 0,1263152 | -2,54103 | 0,0110526 | 0,0380966 | NRDE2 |
| ENSGALG00000005367 | 6338,6685 | -0,320977764 | 0,116103 | -2,764594 | 0,0056994 | 0,022298 | MRPS22 |
| ENSGALG00000004197 | 7479,7717 | -0,321070935 | 0,1089802 | -2,94614 | 0,0032177 | 0,0139429 | SUPV3L1 |
| ENSGALG00000005668 | 17427,249 | -0,321274618 | 0,102142 | -3,145373 | 0,0016588 | 0,0080714 | WSB1 |
| ENSGALG00000001817 | 20809,427 | -0,321828859 | 0,1040048 | -3,094366 | 0,0019723 | 0,0093338 | CTCF |
| ENSGALG00000000235 | 25074,326 | -0,321994098 | 0,1026312 | -3,13739 | 0,0017046 | 0,008262 | PPP2R2A |
| ENSGALG00000016496 | 7694,8086 | -0,322028168 | 0,1163281 | -2,768276 | 0,0056354 | 0,0221302 | ATAD2B |
| ENSGALG00000007021 | 10708,327 | -0,322236639 | 0,122335 | -2,634052 | 0,0084373 | 0,0307066 | PCNT |
| ENSGALG00000010178 | 15181,256 | -0,322485727 | 0,103283 | -3,122351 | 0,0017941 | 0,0086102 | PUM3 |
| ENSGALG00000012415 | 30254,104 | -0,322526711 | 0,1060731 | -3,040608 | 0,002361 | 0,010816 | PDIA4 |
| ENSGALG00000010629 | 6077,087 | -0,322714728 | 0,1236984 | -2,608883 | 0,0090838 | 0,0325036 | ZCCHC11 |
| ENSGALG00000015420 | 5618,1384 | -0,322913162 | 0,1200773 | -2,689211 | 0,0071621 | 0,0269519 | MINA |
| ENSGALG00000016312 | 24555,619 | -0,323062498 | 0,1115176 | -2,896963 | 0,0037679 | 0,0159318 | POLA1 |
| ENSGALG00000027524 | 18598,963 | -0,323112776 | 0,1191737 | -2,711276 | 0,0067025 | 0,0255328 | TNKS1BP1 |
| ENSGALG00000004905 | 15958,906 | -0,323232118 | 0,1254595 | -2,576385 | 0,0099839 | 0,0351244 | MYEF2 |
| ENSGALG00000026825 | 4552,9877 | -0,323431285 | 0,117936 | -2,742431 | 0,0060986 | 0,0235755 | CENPL |
| ENSGALG00000005758 | 19288,831 | -0,323694502 | 0,1027713 | -3,149659 | 0,0016346 | 0,007994 | SPPL2A |
| ENSGALG00000012795 | 9238,0512 | -0,32399695 | 0,115695 | -2,80044 | 0,0051033 | 0,0203938 | RIOK1 |
| ENSGALG00000006477 | 10679,273 | -0,324083678 | 0,1040257 | -3,11542 | 0,0018368 | 0,0087718 | DNAJB6 |
| ENSGALG00000001225 | 6585,9552 | -0,324115689 | 0,1122468 | -2,887527 | 0,0038828 | 0,0163183 | RABGAP1 |
| ENSGALG00000002248 | 6999,8386 | -0,324167266 | 0,1095291 | -2,959644 | 0,0030799 | 0,0134575 | NAB1 |
| ENSGALG00000004468 | 4954,5803 | -0,324300015 | 0,1240229 | -2,61484 | 0,0089269 | 0,0320941 | RMDN3 |
| ENSGALG00000001774 | 27937,325 | -0,324482091 | 0,1102622 | -2,942822 | 0,0032524 | 0,0140619 | NEO1 |
| ENSGALG00000014740 | 9788,9965 | -0,324585149 | 0,1200564 | -2,703606 | 0,0068592 | 0,0259771 | IPO11 |
| ENSGALG00000002222 | 13238,295 | -0,324594377 | 0,102741 | -3,159348 | 0,0015812 | 0,0077775 | NOC2L |
| ENSGALG00000001691 | 7796,647 | -0,324650287 | 0,1188991 | -2,73047 | 0,0063244 | 0,0243034 | CPSF3L |
| ENSGALG00000016630 | 14201,696 | -0,324650816 | 0,1156607 | -2,806924 | 0,0050017 | 0,0200672 | EXTL3 |
| ENSGALG00000008963 | 9612,6755 | -0,324767208 | 0,1203182 | -2,699236 | 0,0069499 | 0,0262549 | DBF4 |
| ENSGALG00000016383 | 4080,3919 | -0,324853698 | 0,1238239 | -2,623513 | 0,0087028 | 0,0314627 | TBC1D31 |
| ENSGALG00000000814 | 43324,255 | -0,325022823 | 0,0992306 | -3,275429 | 0,001055 | 0,0055891 | HNRNPR |
| ENSGALG00000014575 | 10529,411 | -0,325103377 | 0,1055361 | -3,080496 | 0,0020666 | 0,0096988 | SEPHS1 |
| ENSGALG00000010228 | 4674,7787 | -0,325552019 | 0,1143948 | -2,845863 | 0,0044291 | 0,0181143 | TOE1 |
| ENSGALG00000017098 | 6011,3801 | -0,325598928 | 0,1104524 | -2,947867 | 0,0031997 | 0,0138873 | GTF3A |
| ENSGALG00000013626 | 6229,0015 | -0,325639127 | 0,1131349 | -2,878325 | 0,0039979 | 0,0166777 | GMNN |
| ENSGALG00000011251 | 4075,2982 | -0,326088701 | 0,1204829 | -2,706514 | 0,0067994 | 0,025808 | TBC1D5 |
| ENSGALG00000004780 | 17002,116 | -0,326184617 | 0,1040438 | -3,135071 | 0,0017181 | 0,0083217 | NAA25 |
| ENSGALG00000006512 | 270063,95 | -0,326293391 | 0,1225651 | -2,662205 | 0,0077631 | 0,028776 | HSPA8 |
| ENSGALG00000017200 | 4010,3886 | -0,326558761 | 0,134864 | -2,421394 | 0,0154611 | 0,0495885 | MTMR2 |
| ENSGALG00000003303 | 8501,0081 | -0,326575058 | 0,1195862 | -2,730875 | 0,0063166 | 0,024288 | IDH3A |
| ENSGALG00000012986 | 12589,688 | -0,326734663 | 0,1169574 | -2,793621 | 0,0052122 | 0,0207496 | KDM5A |
| ENSGALG00000011091 | 12877,91 | -0,326829517 | 0,1036675 | -3,152671 | 0,0016178 | 0,0079204 | NUP133 |
| ENSGALG00000010185 | 21490,623 | -0,326946921 | 0,1096738 | -2,981085 | 0,0028723 | 0,0127051 | HSPA4L |
| ENSGALG00000004263 | 8041,9782 | -0,32716572 | 0,1124664 | -2,909008 | 0,0036258 | 0,0154169 | MYDGF |
| ENSGALG00000001386 | 15479,004 | -0,32719689 | 0,1250442 | -2,61665 | 0,0088797 | 0,0319666 | PI4KA |
| ENSGALG00000015118 | 7147,4862 | -0,327224219 | 0,1258447 | -2,600222 | 0,0093163 | 0,0331874 | SMC5 |
| ENSGALG00000004691 | 6193,4106 | -0,327404992 | 0,1145398 | -2,85844 | 0,0042573 | 0,0175646 | NLE1 |
| ENSGALG00000001262 | 15946,322 | -0,32743844 | 0,1065971 | -3,07174 | 0,0021282 | 0,0099471 | CLIC4 |

| | | | | | | | |
|---|---|---|---|---|---|---|---|
| ENSGALG00000003624 | 5974,0185 | -0,32749199 | 0,132372 | -2,474028 | 0,0133599 | 0,0441923 | KLHL10 |
| ENSGALG00000021128 | 30791,423 | -0,32759269 | 0,1148555 | -2,852217 | 0,0043415 | 0,0178311 | FSCN1 |
| ENSGALG00000004161 | 24484,078 | -0,327725215 | 0,1031907 | -3,175919 | 0,0014936 | 0,0074083 | CCNB2 |
| ENSGALG00000012192 | 7395,3976 | -0,328017116 | 0,1315753 | -2,492999 | 0,0126669 | 0,0424152 | NT5C3A |
| ENSGALG00000015367 | 2777,8919 | -0,328321788 | 0,124308 | -2,641195 | 0,0082614 | 0,0301957 | PVRL3 |
| ENSGALG00000009944 | 17813,767 | -0,328405469 | 0,1120225 | -2,931602 | 0,0033722 | 0,0144834 | MYST2 |
| ENSGALG00000006829 | 14523,345 | -0,328581471 | 0,1344576 | -2,443756 | 0,0145352 | 0,0471746 | SMG1 |
| ENSGALG00000014286 | 15937,559 | -0,329230926 | 0,1063546 | -3,095595 | 0,0019642 | 0,0093017 | PDS5A |
| ENSGALG00000015334 | 2718,7701 | -0,329290061 | 0,1296871 | -2,539112 | 0,0111134 | 0,0382415 | ZBTB11 |
| ENSGALG00000027091 | 4014,8809 | -0,329390279 | 0,1266481 | -2,60083 | 0,0092998 | 0,0331373 | GPR27 |
| ENSGALG00000013962 | 7389,8352 | -0,329646116 | 0,1086383 | -3,034345 | 0,0024106 | 0,0109913 | HBS1L |
| ENSGALG00000007603 | 19301,424 | -0,329941075 | 0,1204714 | -2,738749 | 0,0061673 | 0,0238124 | RRP12 |
| ENSGALG00000002062 | 16195,779 | -0,329944458 | 0,105102 | -3,139279 | 0,0016936 | 0,0082195 | SRP68 |
| ENSGALG00000001962 | 54240,973 | -0,330389801 | 0,103799 | -3,182976 | 0,0014577 | 0,0072725 | PTBP1 |
| ENSGALG00000001283 | 38879,748 | -0,330483919 | 0,1158085 | -2,853711 | 0,0043212 | 0,0177582 | SRRM1 |
| ENSGALG00000010576 | 13964,899 | -0,330632554 | 0,1026092 | -3,22225 | 0,0012719 | 0,0065198 | GTF2A1 |
| ENSGALG00000010481 | 12612,575 | -0,330642085 | 0,1059113 | -3,121879 | 0,001797 | 0,008621 | CMPK1 |
| ENSGALG00000028637 | 4067,4699 | -0,330733659 | 0,135202 | -2,446219 | 0,0144363 | 0,0468982 | PTPN23 |
| ENSGALG00000010341 | 71633,337 | -0,33081711 | 0,1033707 | -3,2003 | 0,0013728 | 0,0069433 | XPO5 |
| ENSGALG00000015540 | 16673,279 | -0,330835071 | 0,1205533 | -2,744306 | 0,0060639 | 0,0234679 | RAD23B |
| ENSGALG00000007387 | 14223,805 | -0,330955786 | 0,1065147 | -3,107138 | 0,0018891 | 0,0089867 | GTF3C1 |
| ENSGALG00000021683 | 8310,9409 | -0,330974467 | 0,1292911 | -2,559917 | 0,0104697 | 0,0364939 | RER1 |
| ENSGALG00000003547 | 13605,07 | -0,331189406 | 0,1062178 | -3,118021 | 0,0018207 | 0,0087131 | SKP2 |
| ENSGALG00000002889 | 12683,471 | -0,331591263 | 0,1090874 | -3,039685 | 0,0023683 | 0,010842 | ASPSCR1 |
| ENSGALG00000001788 | 3234,7706 | -0,331738591 | 0,1195268 | -2,775432 | 0,0055128 | 0,0217718 | ADPGK |
| ENSGALG00000014445 | 19258,2 | -0,331954776 | 0,1074559 | -3,08922 | 0,0020068 | 0,0094707 | NOP2 |
| ENSGALG00000011904 | 3328,8747 | -0,332013213 | 0,1288249 | -2,577244 | 0,0099592 | 0,0350644 | TRMT5 |
| ENSGALG00000007590 | 10666,564 | -0,332146464 | 0,1081338 | -3,071625 | 0,002129 | 0,0099471 | MTMR8 |
| ENSGALG00000004806 | 7000,4013 | -0,332607085 | 0,1094342 | -3,039334 | 0,002371 | 0,0108489 | EDEM3 |
| ENSGALG00000004108 | 32339,09 | -0,332634184 | 0,1003676 | -3,314158 | 0,0009192 | 0,004992 | CCAR1 |
| ENSGALG00000015813 | 10809,928 | -0,332684119 | 0,1173697 | -2,834497 | 0,0045898 | 0,0186403 | USP16 |
| ENSGALG00000010967 | 21405,06 | -0,332907911 | 0,1160936 | -2,867581 | 0,0041362 | 0,017143 | DOCK7 |
| ENSGALG00000007528 | 28367,48 | -0,33311917 | 0,1060953 | -3,139811 | 0,0016906 | 0,0082087 | TCERG1 |
| ENSGALG00000016766 | 4441,5098 | -0,333177403 | 0,118864 | -2,803014 | 0,0050627 | 0,0202733 | REV1 |
| ENSGALG00000006051 | 6105,2991 | -0,333204336 | 0,1106146 | -3,0123 | 0,0025928 | 0,0116542 | CDC7 |
| ENSGALG00000005177 | 21314,802 | -0,333333735 | 0,1014573 | -3,285458 | 0,0010182 | 0,0054171 | FAM120A |
| ENSGALG00000001976 | 9937,8689 | -0,333600989 | 0,1086579 | -3,070194 | 0,0021392 | 0,0099866 | TRIM33 |
| ENSGALG00000011704 | 13971,193 | -0,333952521 | 0,1117683 | -2,987901 | 0,002809 | 0,012486 | PTPLB |
| ENSGALG00000010282 | 7159,3971 | -0,334004467 | 0,1089729 | -3,065022 | 0,0021765 | 0,0101207 | TMEM63B |
| ENSGALG00000026315 | 3315,4842 | -0,334081105 | 0,119333 | -2,799571 | 0,0051171 | 0,0204367 | RAB34 |
| ENSGALG00000015505 | 6307,2082 | -0,334198908 | 0,1151962 | -2,901129 | 0,0037182 | 0,0157558 | YTHDF3 |
| ENSGALG00000009332 | 24425,521 | -0,334253024 | 0,1022919 | -3,26764 | 0,0010845 | 0,0057141 | RBM25 |
| ENSGALG00000005042 | 3847,7663 | -0,334275581 | 0,1289111 | -2,593071 | 0,0095123 | 0,0337354 | IQSEC1 |
| ENSGALG00000003169 | 2023,2915 | -0,334437759 | 0,1334861 | -2,505413 | 0,0122309 | 0,0412084 | CBFB |
| ENSGALG00000012130 | 1792,2693 | -0,334703976 | 0,1357589 | -2,46543 | 0,0136849 | 0,044964 | DNAJC24 |
| ENSGALG00000004325 | 13667,765 | -0,334704266 | 0,1162502 | -2,879172 | 0,0039872 | 0,0166514 | P4HA1 |
| ENSGALG00000004088 | 4344,9033 | -0,335048738 | 0,1281962 | -2,613561 | 0,0089604 | 0,0321873 | FNBP1 |
| ENSGALG00000009500 | 5544,5023 | -0,33525036 | 0,1244385 | -2,694104 | 0,0070578 | 0,0265889 | VPS50 |
| ENSGALG00000010733 | 14379,472 | -0,335293656 | 0,1230465 | -2,724935 | 0,0064314 | 0,0246658 | CEP170 |
| ENSGALG00000015858 | 4190,1025 | -0,335665691 | 0,1163166 | -2,885795 | 0,0039043 | 0,0163839 | DOPEY1 |
| ENSGALG00000004523 | 6662,5854 | -0,335874075 | 0,1147961 | -2,925833 | 0,0034354 | 0,0147175 | MFN2 |
| ENSGALG00000011533 | 4440,3034 | -0,335898283 | 0,1139943 | -2,946625 | 0,0032126 | 0,0139255 | XRCC3 |
| ENSGALG00000014298 | 14009,413 | -0,335955725 | 0,1061838 | -3,163909 | 0,0015567 | 0,0076817 | RFC1 |
| ENSGALG00000028471 | 3553,6714 | -0,335976223 | 0,1281632 | -2,621473 | 0,0087551 | 0,0316265 | CNPY3 |
| ENSGALG00000014822 | 2960,1308 | -0,336894676 | 0,1318948 | -2,554268 | 0,0106411 | 0,0369778 | CTC1 |
| ENSGALG00000003231 | 28849,937 | -0,336908133 | 0,1031629 | -3,265788 | 0,0010916 | 0,0057449 | KIF20A |
| ENSGALG00000018800 | 6576,7401 | -0,337277222 | 0,1183588 | -2,849617 | 0,0043772 | 0,0179396 | POP1 |
| ENSGALG00000016898 | 2197,9018 | -0,33731956 | 0,1297596 | -2,599573 | 0,009334 | 0,0332327 | TGDS |
| ENSGALG00000006617 | 7019,2341 | -0,337429872 | 0,117642 | -2,868278 | 0,0041271 | 0,0171209 | BCCIP |
| ENSGALG00000009494 | 6742,6053 | -0,337457352 | 0,1118976 | -3,015769 | 0,0025633 | 0,0115407 | THAP5 |
| ENSGALG00000009410 | 4067,5004 | -0,337671951 | 0,1159276 | -2,912783 | 0,0035822 | 0,0152603 | PEX1 |
| ENSGALG00000009097 | 3815,7028 | -0,338568689 | 0,1171869 | -2,889135 | 0,003863 | 0,0162676 | SNAP23 |
| ENSGALG00000009464 | 6021,3203 | -0,338614502 | 0,1119841 | -3,023773 | 0,0024964 | 0,0113033 | GOLIM4 |
| ENSGALG00000014327 | 7067,9269 | -0,338948318 | 0,1081812 | -3,133152 | 0,0017294 | 0,0083614 | ARID4B |

| | | | | | | | |
|---|---|---|---|---|---|---|---|
| ENSGALG00000002302 | 11281,327 | -0,339317581 | 0,1087619 | -3,119821 | 0,0018096 | 0,0086661 | YEATS2 |
| ENSGALG00000020569 | 5039,2721 | -0,339770586 | 0,1193898 | -2,845892 | 0,0044287 | 0,0181143 | DIMT1 |
| ENSGALG00000012617 | 6367,5718 | -0,339828609 | 0,1237855 | -2,745303 | 0,0060455 | 0,0234167 | TGFBR1 |
| ENSGALG00000000408 | 12091,853 | -0,339880371 | 0,121973 | -2,786521 | 0,0053277 | 0,0211518 | LSM7 |
| ENSGALG00000013403 | 2742,2413 | -0,340019904 | 0,1302877 | -2,609761 | 0,0090605 | 0,0324549 | MSANTD3 |
| ENSGALG00000004089 | 8925,1812 | -0,340215935 | 0,1167229 | -2,914732 | 0,0035599 | 0,0151891 | PRDM2 |
| ENSGALG00000005966 | 13999,743 | -0,340533213 | 0,1101239 | -3,092273 | 0,0019863 | 0,0093851 | SEC61A1 |
| ENSGALG00000016255 | 2201,4032 | -0,340652705 | 0,131986 | -2,580977 | 0,0098521 | 0,0347594 | RPGR |
| ENSGALG00000006601 | 4208,8165 | -0,340895714 | 0,1177608 | -2,894815 | 0,0037938 | 0,0160114 | MTMR14 |
| ENSGALG00000012202 | 8130,087 | -0,341191282 | 0,1080608 | -3,157402 | 0,0015918 | 0,0078127 | KBTBD2 |
| ENSGALG00000004759 | 3321,8502 | -0,341574811 | 0,1390989 | -2,455626 | 0,0140639 | 0,0459627 | DHRS7B |
| ENSGALG00000007989 | 28672,589 | -0,341827011 | 0,1092587 | -3,128603 | 0,0017564 | 0,0084619 | ADNP |
| ENSGALG00000003263 | 5162,4302 | -0,341906085 | 0,1243755 | -2,748982 | 0,0059781 | 0,0232054 | SFXN1 |
| ENSGALG00000010361 | 9329,7103 | -0,342127455 | 0,1233327 | -2,774021 | 0,0055368 | 0,0218567 | MBNL1 |
| ENSGALG00000004515 | 10691,241 | -0,342166688 | 0,1169935 | -2,924664 | 0,0034483 | 0,014759 | KNTC1 |
| ENSGALG00000010541 | 14976,655 | -0,3427229 | 0,1277901 | -2,681921 | 0,0073201 | 0,02741 | BIRC6 |
| ENSGALG00000004628 | 2015,8556 | -0,342763414 | 0,1400684 | -2,447115 | 0,0144005 | 0,0467957 | CTNS |
| ENSGALG00000005237 | 11277,397 | -0,342909039 | 0,1056363 | -3,24613 | 0,0011699 | 0,0060815 | FGD3 |
| ENSGALG00000005420 | 7995,347 | -0,343276467 | 0,1261306 | -2,721596 | 0,0064967 | 0,0248884 | AATF |
| ENSGALG00000002648 | 3882,3909 | -0,343349251 | 0,1203014 | -2,854076 | 0,0043162 | 0,017756 | FUK |
| ENSGALG00000016314 | 22102,711 | -0,343364339 | 0,1069912 | -3,209276 | 0,0013307 | 0,0067704 | ELOVL5 |
| ENSGALG00000006455 | 4218,2466 | -0,343579981 | 0,1158475 | -2,965794 | 0,003019 | 0,0132318 | SMPD4 |
| ENSGALG00000012239 | 9046,4232 | -0,343596083 | 0,1058983 | -3,244586 | 0,0011762 | 0,0061076 | METAP1 |
| ENSGALG00000004624 | 11632,549 | -0,34416889 | 0,1347829 | -2,553506 | 0,0106645 | 0,0370494 | MYO5A |
| ENSGALG00000003365 | 1822,7969 | -0,344340057 | 0,1360837 | -2,530355 | 0,0113947 | 0,0389601 | RAD1 |
| ENSGALG00000002527 | 4444,3372 | -0,344545241 | 0,1227619 | -2,806613 | 0,0050065 | 0,0200778 | TMEM201 |
| ENSGALG00000009813 | 1922,5946 | -0,344705696 | 0,1334833 | -2,582388 | 0,0098119 | 0,0346715 | B0LHV3_CHICK |
| ENSGALG00000013053 | 3803,4898 | -0,345339591 | 0,1218833 | -2,833364 | 0,0046061 | 0,0186977 | MTRR |
| ENSGALG00000001494 | 9000,8803 | -0,345374318 | 0,1246336 | -2,771118 | 0,0055864 | 0,021976 | NISCH |
| ENSGALG00000011884 | 7925,1907 | -0,345461445 | 0,1150003 | -3,004005 | 0,0026645 | 0,0119412 | EXOSC9 |
| ENSGALG00000001236 | 5792,0595 | -0,345759524 | 0,1103803 | -3,132439 | 0,0017336 | 0,0083777 | ZBTB26 |
| ENSGALG00000005261 | 15832,957 | -0,345768981 | 0,1275962 | -2,70987 | 0,006731 | 0,0256198 | RP11-403P17.5 |
| ENSGALG00000012837 | 7840,9496 | -0,345878883 | 0,1096612 | -3,154068 | 0,0016101 | 0,0078884 | EXOC2 |
| ENSGALG00000006307 | 2423,8788 | -0,346032557 | 0,1338086 | -2,586026 | 0,009709 | 0,0343344 | NSMCE1 |
| ENSGALG00000003434 | 9368,6549 | -0,346127339 | 0,1194628 | -2,897364 | 0,0037631 | 0,0159164 | C17orf75 |
| ENSGALG00000006153 | 2844,7772 | -0,346474519 | 0,1286035 | -2,69413 | 0,0070573 | 0,0265889 | EXOG |
| ENSGALG00000006475 | 6864,1942 | -0,347016477 | 0,1079405 | -3,214886 | 0,001305 | 0,0066643 | KIAA0430 |
| ENSGALG00000002276 | 1478,127 | -0,34721896 | 0,1416798 | -2,45073 | 0,0142567 | 0,0464477 | EME2 |
| ENSGALG00000015553 | 10445,976 | -0,347410904 | 0,1127 | -3,082616 | 0,0020519 | 0,0096499 | ARFGEF1 |
| ENSGALG00000011831 | 9083,6449 | -0,347478626 | 0,1108523 | -3,13461 | 0,0017208 | 0,0083259 | ANKRD50 |
| ENSGALG00000004269 | 64393,653 | -0,347526191 | 0,103785 | -3,348521 | 0,0008124 | 0,0044938 | EIF3B |
| ENSGALG00000004548 | 15606,046 | -0,347529204 | 0,1215975 | -2,85803 | 0,0042628 | 0,0175766 | RP11-101E3.5 |
| ENSGALG00000005477 | 8038,2726 | -0,347589933 | 0,1308189 | -2,657031 | 0,0078832 | 0,0291104 | MPHOSPH6 |
| ENSGALG00000004200 | 11378,582 | -0,347970886 | 0,1075116 | -3,236588 | 0,0012097 | 0,0062623 | DPP9 |
| ENSGALG00000012364 | 4848,5693 | -0,348109095 | 0,1264902 | -2,752064 | 0,0059221 | 0,0230238 | PPIL4 |
| ENSGALG00000016887 | 64639,106 | -0,348222743 | 0,0978749 | -3,557836 | 0,0003739 | 0,002345 | IPO5 |
| ENSGALG00000013649 | 18631,187 | -0,348311839 | 0,1020152 | -3,414312 | 0,0006394 | 0,003678 | CNOT7 |
| ENSGALG00000017241 | 2897,41 | -0,348538577 | 0,1300034 | -2,680995 | 0,0073404 | 0,0274784 | TMEM135 |
| ENSGALG00000016931 | 5043,2864 | -0,348656857 | 0,1247665 | -2,794476 | 0,0051984 | 0,0207069 | BORA |
| ENSGALG00000002469 | 8118,1027 | -0,348923543 | 0,1164279 | -2,996907 | 0,0027273 | 0,0121867 | CDC73 |
| ENSGALG00000004168 | 7087,525 | -0,349365069 | 0,1171636 | -2,981857 | 0,0028651 | 0,0126896 | FEM1A |
| ENSGALG00000027444 | 3608,9781 | -0,349517124 | 0,1245455 | -2,806341 | 0,0050108 | 0,0200861 | |
| ENSGALG00000017139 | 10651,91 | -0,349554954 | 0,107047 | -3,265433 | 0,001093 | 0,0057499 | ZMYM2 |
| ENSGALG00000002719 | 17659,446 | -0,349571816 | 0,1018084 | -3,433625 | 0,0005956 | 0,0034712 | CSNK1D |
| ENSGALG00000008645 | 8917,9646 | -0,349922505 | 0,1156037 | -3,026915 | 0,0024706 | 0,0112201 | NKRF |
| ENSGALG00000004042 | 4817,4388 | -0,349988244 | 0,1306697 | -2,67842 | 0,007397 | 0,0276565 | NUFIP2 |
| ENSGALG00000009633 | 4605,1827 | -0,350007911 | 0,1141085 | -3,067326 | 0,0021598 | 0,0100534 | KATNBL1 |
| ENSGALG00000007500 | 4924,5265 | -0,350118918 | 0,1432067 | -2,44485 | 0,0144912 | 0,047043 | DENND4A |
| ENSGALG00000009249 | 5232,7054 | -0,350123017 | 0,1287908 | -2,71854 | 0,0065571 | 0,0250618 | UBR1 |
| ENSGALG00000008454 | 36878,682 | -0,351028215 | 0,1038189 | -3,38116 | 0,0007218 | 0,0040828 | NOP58 |
| ENSGALG00000016727 | 9641,1293 | -0,351861521 | 0,1225244 | -2,871766 | 0,0040818 | 0,0169796 | JADE3 |
| ENSGALG00000001023 | 2133,9525 | -0,352229585 | 0,1347749 | -2,613466 | 0,0089629 | 0,0321873 | BZRAP1 |
| ENSGALG00000017026 | 9366,3877 | -0,352330056 | 0,1086684 | -3,24225 | 0,0011859 | 0,0061532 | VPS36 |
| ENSGALG00000007861 | 11084,886 | -0,352469383 | 0,1082462 | -3,256183 | 0,0011292 | 0,0059177 | MAGT1 |

| | | | | | | | |
|---|---|---|---|---|---|---|---|
| ENSGALG00000005569 | 11340,438 | -0,352585966 | 0,10935 | -3,224379 | 0,0012625 | 0,00648 | USP47 |
| ENSGALG00000009283 | 5220,1674 | -0,352608684 | 0,1245251 | -2,831627 | 0,0046312 | 0,0187848 | KIAA1715 |
| ENSGALG00000006612 | 7533,7656 | -0,353107966 | 0,1223847 | -2,885229 | 0,0039113 | 0,0164004 | ASXL1 |
| ENSGALG00000009913 | 62516,864 | -0,353547892 | 0,0988051 | -3,578237 | 0,0003459 | 0,0021977 | SMARCA5 |
| ENSGALG00000011620 | 9603,3836 | -0,353615473 | 0,1055478 | -3,350286 | 0,0008073 | 0,0044752 | AKT1 |
| ENSGALG00000010177 | 12046,319 | -0,353917428 | 0,1080741 | -3,274766 | 0,0010575 | 0,0056001 | ZFC3H1 |
| ENSGALG00000015043 | 11145,365 | -0,35411601 | 0,1099631 | -3,220318 | 0,0012805 | 0,006559 | RPF2 |
| ENSGALG00000011368 | 2716,7046 | -0,354344153 | 0,1315134 | -2,694357 | 0,0070525 | 0,0265834 | ZNF839 |
| ENSGALG00000015024 | 3631,7153 | -0,355056011 | 0,138118 | -2,570671 | 0,0101502 | 0,0355805 | RCL1 |
| ENSGALG00000013863 | 3597,6259 | -0,35519316 | 0,1205708 | -2,94593 | 0,0032199 | 0,0139435 | PSMG2 |
| ENSGALG00000009380 | 1987,1722 | -0,355339175 | 0,1456281 | -2,440045 | 0,0146854 | 0,0475709 | ST7 |
| ENSGALG00000008036 | 5782,6274 | -0,355406874 | 0,1116416 | -3,183463 | 0,0014552 | 0,0072656 | TMEM209 |
| ENSGALG00000014383 | 9301,5506 | -0,355665092 | 0,1163485 | -3,056896 | 0,0022364 | 0,0103287 | PI4K2B |
| ENSGALG00000010805 | 16069,419 | -0,355813089 | 0,1260649 | -2,822459 | 0,0047657 | 0,0192708 | USP24 |
| ENSGALG00000021031 | 4102,9147 | -0,355958904 | 0,1319256 | -2,698179 | 0,006972 | 0,0263165 | ACKR3 |
| ENSGALG00000005125 | 7959,3271 | -0,356020767 | 0,1067406 | -3,335383 | 0,0008518 | 0,0046764 | LSM14B |
| ENSGALG00000012717 | 4761,9391 | -0,356301296 | 0,1163989 | -3,061036 | 0,0022057 | 0,0102355 | NFYB |
| ENSGALG00000006123 | 7411,1798 | -0,356657256 | 0,13028 | -2,73762 | 0,0061885 | 0,0238689 | RTEL1 |
| ENSGALG00000006254 | 6379,7603 | -0,357023226 | 0,1173769 | -3,041681 | 0,0023526 | 0,0107811 | ERCC6 |
| ENSGALG00000001811 | 14773,974 | -0,357025162 | 0,1141916 | -3,126546 | 0,0017687 | 0,0085114 | RP4-614O4.11 |
| ENSGALG00000003441 | 4610,7331 | -0,357140672 | 0,1182742 | -3,0196 | 0,0025311 | 0,0114298 | GTF3C5 |
| ENSGALG00000008957 | 22456,525 | -0,357191503 | 0,1156693 | -3,088042 | 0,0020148 | 0,0094985 | MSH6 |
| ENSGALG00000012782 | 4940,896 | -0,35754473 | 0,1182911 | -3,022584 | 0,0025063 | 0,0113327 | BLOC1S5 |
| ENSGALG00000013657 | 2568,382 | -0,35785969 | 0,136842 | -2,615131 | 0,0089193 | 0,0320837 | ZNF236 |
| ENSGALG00000008218 | 3293,8403 | -0,357888042 | 0,1386724 | -2,580816 | 0,0098567 | 0,0347614 | DDB2 |
| ENSGALG00000009330 | 6639,4947 | -0,358006963 | 0,1142065 | -3,134734 | 0,0017201 | 0,0083253 | DEGS1 |
| ENSGALG00000010763 | 6453,8983 | -0,358059956 | 0,1126815 | -3,177629 | 0,0014848 | 0,0073728 | TRIP11 |
| ENSGALG00000008507 | 23518,146 | -0,358201653 | 0,1023769 | -3,498854 | 0,0004673 | 0,0028338 | THOC2 |
| ENSGALG00000009839 | 30979,576 | -0,358257804 | 0,1012165 | -3,539518 | 0,0004009 | 0,002491 | PPIG |
| ENSGALG00000010349 | 7048,1543 | -0,358393493 | 0,1129044 | -3,174309 | 0,0015019 | 0,0074414 | POLR1C |
| ENSGALG00000007721 | 11622,16 | -0,358457512 | 0,1098179 | -3,264109 | 0,0010981 | 0,0057724 | ARHGAP21 |
| ENSGALG00000012414 | 1870,7753 | -0,358664924 | 0,1359051 | -2,639084 | 0,0083131 | 0,0303275 | GNPNAT1 |
| ENSGALG00000016419 | 4650,1008 | -0,358846606 | 0,1220043 | -2,941261 | 0,0032688 | 0,0141048 | ASAP2 |
| ENSGALG00000002399 | 4037,3816 | -0,359044332 | 0,1192358 | -3,011213 | 0,0026021 | 0,01169 | GTF2H2 |
| ENSGALG00000014795 | 5578,5367 | -0,359083857 | 0,1276747 | -2,812491 | 0,0049159 | 0,0198137 | TGIF1 |
| ENSGALG00000005694 | 30867,661 | -0,359162398 | 0,1021823 | -3,514918 | 0,0004399 | 0,0026893 | GARS |
| ENSGALG00000016598 | 5686,2877 | -0,359376196 | 0,1202475 | -2,988638 | 0,0028022 | 0,0124641 | MSL3 |
| ENSGALG00000008254 | 17757,266 | -0,35964551 | 0,1079509 | -3,331565 | 0,0008636 | 0,0047277 | ANAPC1 |
| ENSGALG00000007683 | 31803,136 | -0,359759566 | 0,1106419 | -3,251567 | 0,0011477 | 0,0059985 | |
| ENSGALG00000005374 | 3276,9339 | -0,359759753 | 0,1426205 | -2,522497 | 0,0116525 | 0,0396423 | MYO19 |
| ENSGALG00000004670 | 7571,3318 | -0,360055666 | 0,109156 | -3,298544 | 0,0009719 | 0,0052095 | ATRIP |
| ENSGALG00000024039 | 12181,866 | -0,36099515 | 0,1049573 | -3,439449 | 0,0005829 | 0,0034066 | UTP14A |
| ENSGALG00000004379 | 20460,522 | -0,361272684 | 0,1033693 | -3,494972 | 0,0004741 | 0,0028704 | BCL7A |
| ENSGALG00000000463 | 2471,2656 | -0,361708293 | 0,131949 | -2,741273 | 0,0061202 | 0,0236453 | DDX25 |
| ENSGALG00000006955 | 3003,9828 | -0,36242424 | 0,1397632 | -2,59313 | 0,0095107 | 0,0337354 | CCNJ |
| ENSGALG00000012763 | 12297,883 | -0,362574608 | 0,1084037 | -3,344671 | 0,0008238 | 0,0045427 | GNPTAB |
| ENSGALG00000001942 | 6053,3632 | -0,362623052 | 0,1158608 | -3,129817 | 0,0017492 | 0,0084479 | SPDL1 |
| ENSGALG00000012638 | 9149,4343 | -0,362691803 | 0,1227328 | -2,955134 | 0,0031253 | 0,0136121 | CRY1 |
| ENSGALG00000012916 | 8111,1057 | -0,362729901 | 0,1091942 | -3,321879 | 0,0008941 | 0,0048792 | DROSHA |
| ENSGALG00000002139 | 98917,168 | -0,362887161 | 0,1016046 | -3,571563 | 0,0003549 | 0,0022423 | CSDE1 |
| ENSGALG00000009325 | 21860,643 | -0,363294006 | 0,1110082 | -3,272678 | 0,0010653 | 0,0056394 | OLA1 |
| ENSGALG00000015993 | 4919,2212 | -0,363761464 | 0,123068 | -2,955776 | 0,0031188 | 0,0135925 | MTERF3 |
| ENSGALG00000015363 | 3478,9826 | -0,363849789 | 0,1186073 | -3,067684 | 0,0021572 | 0,0100482 | HJURP |
| ENSGALG00000003826 | 8394,886 | -0,363869123 | 0,1071172 | -3,396925 | 0,0006815 | 0,0038903 | RP11-478C19.2 |
| ENSGALG00000001790 | 14734,645 | -0,363943316 | 0,1104661 | -3,294616 | 0,0009856 | 0,0052683 | RP11-318A15.7 |
| ENSGALG00000004553 | 9745,0828 | -0,363967467 | 0,1049477 | -3,468085 | 0,0005242 | 0,0031206 | SGPL1 |
| ENSGALG00000002471 | 4521,5673 | -0,363984132 | 0,1261711 | -2,884846 | 0,0039161 | 0,0164153 | ZNF592 |
| ENSGALG00000001663 | 10259,573 | -0,364202029 | 0,1099047 | -3,3138 | 0,0009204 | 0,0049964 | MAK16 |
| ENSGALG00000007199 | 3668,9433 | -0,36428019 | 0,1273389 | -2,860714 | 0,0042269 | 0,0174655 | TRIAP1 |
| ENSGALG00000010900 | 12043,923 | -0,364291872 | 0,1128172 | -3,229045 | 0,001242 | 0,0064006 | SPC25 |
| ENSGALG00000010618 | 2510,6274 | -0,364367021 | 0,127904 | -2,848755 | 0,0043891 | 0,0179775 | SLC30A6 |
| ENSGALG00000003563 | 13287,766 | -0,364389392 | 0,1101047 | -3,309482 | 0,0009347 | 0,0050615 | YARS |
| ENSGALG00000000638 | 12789,677 | -0,364411808 | 0,1251634 | -2,911488 | 0,0035971 | 0,0153046 | UTP4 |
| ENSGALG00000010006 | 4182,5012 | -0,364534028 | 0,1219632 | -2,988885 | 0,0028 | 0,0124581 | SLC10A7 |

| | | | | | | | |
|---|---|---|---|---|---|---|---|
| ENSGALG00000002137 | 15220,893 | -0,364569255 | 0,1047688 | -3,479749 | 0,0005019 | 0,0030089 | PGAM5 |
| ENSGALG00000004485 | 1975,7552 | -0,364730971 | 0,1340663 | -2,720526 | 0,0065178 | 0,024948 | DOLPP1 |
| ENSGALG00000004467 | 5426,4673 | -0,365351749 | 0,1191675 | -3,065867 | 0,0021704 | 0,0100991 | CLIP1 |
| ENSGALG00000001126 | 13435,921 | -0,365650894 | 0,1039475 | -3,51765 | 0,0004354 | 0,0026666 | CHAF1A |
| ENSGALG00000003674 | 4753,1822 | -0,365887371 | 0,1502656 | -2,434938 | 0,0148943 | 0,0481221 | DHTKD1 |
| ENSGALG00000015874 | 13228,099 | -0,365980992 | 0,1079617 | -3,389915 | 0,0006991 | 0,0039761 | TTK |
| ENSGALG00000003339 | 10767,061 | -0,366040447 | 0,1177244 | -3,109301 | 0,0018753 | 0,0089243 | MTOR |
| ENSGALG00000010400 | 7431,8392 | -0,366106792 | 0,1070902 | -3,418676 | 0,0006293 | 0,0036317 | SMARCAD1 |
| ENSGALG00000012205 | 2497,2514 | -0,366370844 | 0,1293829 | -2,831679 | 0,0046304 | 0,0187848 | AVL9 |
| ENSGALG00000014867 | 5553,4309 | -0,366420731 | 0,1173966 | -3,12122 | 0,001801 | 0,0086359 | PAIP1 |
| ENSGALG00000006193 | 20925,466 | -0,366551091 | 0,1152835 | -3,179563 | 0,001475 | 0,0073322 | TXNRD3 |
| ENSGALG00000012807 | 3029,2492 | -0,366699033 | 0,1239604 | -2,958195 | 0,0030945 | 0,0135079 | RPP40 |
| ENSGALG00000009587 | 5817,192 | -0,366858159 | 0,1201432 | -3,053508 | 0,0022618 | 0,0104283 | GPHN |
| ENSGALG00000011242 | 2355,7326 | -0,367017354 | 0,1308827 | -2,80417 | 0,0050446 | 0,0202098 | OBSL1 |
| ENSGALG00000005353 | 4596,566 | -0,367159893 | 0,1195448 | -3,071317 | 0,0021312 | 0,009954 | FAR1 |
| ENSGALG00000001872 | 9491,7801 | -0,367383699 | 0,1243406 | -2,954657 | 0,0031302 | 0,0136288 | SDF4 |
| ENSGALG00000015683 | 11433,193 | -0,367498221 | 0,1101651 | -3,335886 | 0,0008503 | 0,0046699 | GNG10 |
| ENSGALG00000003171 | 20304,055 | -0,367614474 | 0,1019213 | -3,606847 | 0,0003099 | 0,0019961 | IREB2 |
| ENSGALG00000028828 | 22676,693 | -0,367651199 | 0,1028075 | -3,576112 | 0,0003487 | 0,0022146 | EIF4E |
| ENSGALG00000009637 | 4483,6182 | -0,367846509 | 0,1376888 | -2,671578 | 0,0075495 | 0,0281146 | EMC7 |
| ENSGALG00000013100 | 3042,9761 | -0,368006107 | 0,1414611 | -2,601465 | 0,0092826 | 0,0330933 | GRB10 |
| ENSGALG00000015009 | 1981,8828 | -0,368164733 | 0,1354772 | -2,71754 | 0,0065769 | 0,0251107 | ZBTB49 |
| ENSGALG00000024159 | 24561,804 | -0,368265263 | 0,1065192 | -3,457267 | 0,0005457 | 0,0032246 | |
| ENSGALG00000003881 | 9972,6997 | -0,368271023 | 0,1180506 | -3,119604 | 0,0018109 | 0,0086695 | ERCC6L |
| ENSGALG00000015957 | 2851,4747 | -0,368440661 | 0,1383223 | -2,663638 | 0,0077301 | 0,0286811 | RNF151 |
| ENSGALG00000001619 | 9403,9871 | -0,368532524 | 0,1142023 | -3,227015 | 0,0012509 | 0,006434 | ATG16L1 |
| ENSGALG00000004115 | 26356,095 | -0,369026879 | 0,1077431 | -3,425063 | 0,0006147 | 0,0035687 | DNAJA2 |
| ENSGALG00000012266 | 5845,0519 | -0,369135559 | 0,1123441 | -3,285757 | 0,0010171 | 0,0054135 | LRRFIP2 |
| ENSGALG00000015355 | 10608,533 | -0,369224786 | 0,1221453 | -3,022833 | 0,0025042 | 0,0113271 | CD47 |
| ENSGALG00000008526 | 11139,841 | -0,369313202 | 0,1088311 | -3,393452 | 0,0006902 | 0,003935 | BRD1 |
| ENSGALG00000001618 | 4391,5637 | -0,369471319 | 0,1261006 | -2,929973 | 0,0033899 | 0,0145504 | OGFOD3 |
| ENSGALG00000001485 | 7897,8046 | -0,369653264 | 0,1079491 | -3,424331 | 0,0006163 | 0,0035753 | TOP3B |
| ENSGALG00000012263 | 9464,4086 | -0,36996383 | 0,1064406 | -3,475778 | 0,0005094 | 0,0030417 | NEMF |
| ENSGALG00000007509 | 10510,151 | -0,37005709 | 0,1074005 | -3,44558 | 0,0005698 | 0,0033455 | GNAS |
| ENSGALG00000008807 | 7903,608 | -0,370129883 | 0,107044 | -3,457735 | 0,0005447 | 0,0032218 | DCUN1D1 |
| ENSGALG00000009547 | 3006,9462 | -0,370182446 | 0,1224834 | -3,022307 | 0,0025086 | 0,0113355 | PPHLN1 |
| ENSGALG00000014582 | 21606,21 | -0,370326496 | 0,1092612 | -3,389368 | 0,0007005 | 0,0039824 | CADM1 |
| ENSGALG00000010052 | 1789,2611 | -0,370466621 | 0,138855 | -2,668012 | 0,0076302 | 0,0283683 | PPP2R3C |
| ENSGALG00000010296 | 6483,9405 | -0,370634079 | 0,1153594 | -3,212864 | 0,0013142 | 0,0067014 | MRPS18A |
| ENSGALG00000011215 | 4111,7689 | -0,370718656 | 0,1158847 | -3,199029 | 0,0013789 | 0,0069662 | POC1B |
| ENSGALG00000004254 | 3591,4431 | -0,370755667 | 0,1184788 | -3,1293 | 0,0017522 | 0,0084598 | TMEM120B |
| ENSGALG00000015649 | 13211,542 | -0,370890117 | 0,1067035 | -3,475894 | 0,0005092 | 0,0030417 | NOP14 |
| ENSGALG00000008557 | 15377,869 | -0,371010327 | 0,1075723 | -3,44894 | 0,0005628 | 0,0033142 | ARL5B |
| ENSGALG00000004224 | 5462,5737 | -0,371126705 | 0,1408849 | -2,634254 | 0,0084322 | 0,0306998 | MAD1L1 |
| ENSGALG00000008795 | 3264,8024 | -0,371277159 | 0,1212162 | -3,062934 | 0,0021918 | 0,0101812 | GPAM |
| ENSGALG00000006586 | 3512,8975 | -0,371428946 | 0,1391432 | -2,6694 | 0,0075987 | 0,0282744 | PLAGL2 |
| ENSGALG00000016614 | 8922,873 | -0,371522735 | 0,1401003 | -2,651835 | 0,0080056 | 0,0294504 | SHROOM2 |
| ENSGALG00000015847 | 1545,0428 | -0,371535052 | 0,1402786 | -2,64855 | 0,0080838 | 0,02965 | PRSS35 |
| ENSGALG00000009817 | 1241,5525 | -0,371617458 | 0,1484632 | -2,503095 | 0,0123113 | 0,041428 | ZNF770 |
| ENSGALG00000005935 | 6194,9397 | -0,371684146 | 0,110711 | -3,357247 | 0,0007872 | 0,0043945 | EVI5 |
| ENSGALG00000009807 | 6561,2068 | -0,371941676 | 0,1238286 | -3,003683 | 0,0026673 | 0,011946 | FLVCR1 |
| ENSGALG00000004834 | 12710,539 | -0,372460145 | 0,1085125 | -3,432418 | 0,0005982 | 0,0034837 | COPS3 |
| ENSGALG00000016578 | 7057,3465 | -0,372483757 | 0,11837 | -3,146774 | 0,0016508 | 0,0080444 | |
| ENSGALG00000003039 | 8776,92 | -0,372512284 | 0,1093684 | -3,406031 | 0,0006591 | 0,0037723 | SRM |
| ENSGALG00000008787 | 2268,5743 | -0,372527529 | 0,1464722 | -2,543332 | 0,0109801 | 0,0379137 | CEP68 |
| ENSGALG00000015705 | 13014,296 | -0,372615903 | 0,1210163 | -3,079056 | 0,0020766 | 0,0097424 | WHSC1 |
| ENSGALG00000015800 | 3887,9053 | -0,37263738 | 0,128126 | -2,908367 | 0,0036332 | 0,0154341 | ORC3 |
| ENSGALG00000011713 | 7918,9457 | -0,372947939 | 0,1156001 | -3,226189 | 0,0012545 | 0,0064477 | CDV3 |
| ENSGALG00000001513 | 6151,0051 | -0,373009041 | 0,1249789 | -2,984577 | 0,0028397 | 0,0125937 | CNTRL |
| ENSGALG00000023536 | 26304,92 | -0,373188455 | 0,1263177 | -2,954363 | 0,0031332 | 0,0136331 | FZD7 |
| ENSGALG00000000412 | 5015,4199 | -0,373361237 | 0,1191944 | -3,132372 | 0,001734 | 0,0083777 | CCDC15 |
| ENSGALG00000003942 | 9873,2639 | -0,373719716 | 0,1078722 | -3,464467 | 0,0005313 | 0,0031559 | LONP2 |
| ENSGALG00000016236 | 30922,987 | -0,374432078 | 0,1135095 | -3,298685 | 0,0009714 | 0,0052095 | USP9X |
| ENSGALG00000005618 | 12381,341 | -0,374714579 | 0,1064892 | -3,518803 | 0,0004335 | 0,002661 | CTR9 |

| | | | | | | | |
|---|---|---|---|---|---|---|---|
| ENSGALG00000010744 | 5078,2186 | -0,375014992 | 0,1175988 | -3,188937 | 0,001428 | 0,0071662 | GALNT7 |
| ENSGALG00000012689 | 4208,649 | -0,375087396 | 0,1416897 | -2,647245 | 0,0081151 | 0,0297327 | KIF13A |
| ENSGALG00000001939 | 11757,482 | -0,375413327 | 0,1227756 | -3,057719 | 0,0022303 | 0,0103145 | HIPK1 |
| ENSGALG00000009626 | 5852,8685 | -0,375704266 | 0,1201786 | -3,126215 | 0,0017707 | 0,0085128 | THBS1 |
| ENSGALG00000021251 | 12519,129 | -0,375765974 | 0,1228993 | -3,057511 | 0,0022318 | 0,0103145 | DDX6 |
| ENSGALG00000003325 | 1577,6189 | -0,375811622 | 0,1416562 | -2,652984 | 0,0079784 | 0,029382 | KMT5A |
| ENSGALG00000013034 | 8837,0774 | -0,375844048 | 0,1530686 | -2,455396 | 0,0140729 | 0,045981 | CECR2 |
| ENSGALG00000009677 | 24959,786 | -0,376298569 | 0,10418 | -3,612004 | 0,0003038 | 0,0019633 | SSB |
| ENSGALG00000003776 | 26231,392 | -0,376541512 | 0,1408091 | -2,674127 | 0,0074924 | 0,0279477 | PRRC2B |
| ENSGALG00000004867 | 10358,587 | -0,376546151 | 0,1068635 | -3,523619 | 0,0004257 | 0,0026261 | CEBPG |
| ENSGALG00000000085 | 18552,019 | -0,37666785 | 0,1073497 | -3,508793 | 0,0004501 | 0,0027434 | GFPT1 |
| ENSGALG00000007929 | 3448,5443 | -0,376720223 | 0,1459515 | -2,581133 | 0,0098477 | 0,0347594 | DIXDC1 |
| ENSGALG00000011281 | 6523,2043 | -0,37690737 | 0,1178655 | -3,197776 | 0,0013849 | 0,0069939 | SGOL1 |
| ENSGALG00000003740 | 3014,38 | -0,377276945 | 0,1362236 | -2,769542 | 0,0056135 | 0,0220587 | IKBKB |
| ENSGALG00000016337 | 3166,2527 | -0,377573201 | 0,1356955 | -2,782504 | 0,0053941 | 0,0213616 | TATDN1 |
| ENSGALG00000012251 | 2154,1949 | -0,377607005 | 0,1483053 | -2,546147 | 0,0108919 | 0,0376475 | KDELR3 |
| ENSGALG00000008247 | 7600,6846 | -0,377692943 | 0,109538 | -3,448054 | 0,0005646 | 0,0033222 | ARFGAP2 |
| ENSGALG00000005006 | 11070,754 | -0,377708377 | 0,1075988 | -3,510339 | 0,0004475 | 0,0027311 | UBA2 |
| ENSGALG00000000978 | 4782,6943 | -0,377763607 | 0,1179131 | -3,203745 | 0,0013565 | 0,0068761 | CEP104 |
| ENSGALG00000013146 | 5329,2902 | -0,377879987 | 0,1326893 | -2,847857 | 0,0044015 | 0,0180229 | PLXNB2 |
| ENSGALG00000006558 | 1973,6885 | -0,378031821 | 0,1426055 | -2,650891 | 0,008028 | 0,0295009 | WDR60 |
| ENSGALG00000005468 | 6023,6864 | -0,378125454 | 0,1402851 | -2,695408 | 0,0070303 | 0,026507 | SYNRG |
| ENSGALG00000009093 | 6655,7671 | -0,378156036 | 0,1279163 | -2,956277 | 0,0031138 | 0,0135748 | PNO1 |
| ENSGALG00000002985 | 14604,542 | -0,378678888 | 0,1056758 | -3,583404 | 0,0003391 | 0,0021617 | UTP18 |
| ENSGALG00000016200 | 6807,9958 | -0,378720539 | 0,1505954 | -2,514821 | 0,0119093 | 0,0403647 | SIK1 |
| ENSGALG00000016851 | 8342,7517 | -0,378897954 | 0,1201029 | -3,154778 | 0,0016062 | 0,0078758 | ARHGEF7 |
| ENSGALG00000008195 | 5727,1148 | -0,378910892 | 0,1131533 | -3,348651 | 0,0008121 | 0,0044938 | NPHP1 |
| ENSGALG00000003326 | 2369,3349 | -0,379089886 | 0,1276328 | -2,97016 | 0,0029764 | 0,0130808 | AMACR |
| ENSGALG00000011015 | 2757,9302 | -0,379357377 | 0,1543896 | -2,457143 | 0,0140047 | 0,045791 | NTPCR |
| ENSGALG00000014085 | 1445,5651 | -0,379926489 | 0,1417435 | -2,68038 | 0,0073539 | 0,0275214 | TM7SF3 |
| ENSGALG00000006011 | 4210,2643 | -0,380179632 | 0,1512099 | -2,51425 | 0,0119286 | 0,040409 | KIAA0753 |
| ENSGALG00000024047 | 6900,307 | -0,380223912 | 0,115759 | -3,284616 | 0,0010212 | 0,0054312 | MYCL |
| ENSGALG00000003322 | 2664,8777 | -0,380407199 | 0,1320573 | -2,880623 | 0,0039689 | 0,0165858 | TBC1D2B |
| ENSGALG00000007036 | 8649,5744 | -0,380541953 | 0,1060007 | -3,589994 | 0,0003307 | 0,0021167 | PITRM1 |
| ENSGALG00000005573 | 47811,321 | -0,380592325 | 0,1004918 | -3,787297 | 0,0001523 | 0,0010811 | NOLC1 |
| ENSGALG00000008187 | 22960,701 | -0,380821067 | 0,1023642 | -3,720256 | 0,000199 | 0,0013569 | EIF3J |
| ENSGALG00000000864 | 4682,0682 | -0,380939683 | 0,115984 | -3,284416 | 0,0010219 | 0,0054329 | FBXW2 |
| ENSGALG00000004696 | 6529,7846 | -0,381141883 | 0,109965 | -3,46603 | 0,0005282 | 0,0031404 | LEO1 |
| ENSGALG00000011330 | 86092,558 | -0,381207584 | 0,1173583 | -3,248237 | 0,0011612 | 0,0060506 | DYNC1H1 |
| ENSGALG00000000543 | 2560,0751 | -0,381260503 | 0,1306196 | -2,91886 | 0,0035131 | 0,0150036 | TNFRSF9 |
| ENSGALG00000005776 | 1699,2203 | -0,38126111 | 0,1437588 | -2,65209 | 0,0079995 | 0,0294363 | TECR |
| ENSGALG00000007354 | 7140,2177 | -0,381532857 | 0,1451021 | -2,629409 | 0,0085533 | 0,0310378 | CIT |
| ENSGALG00000011484 | 5458,5585 | -0,381581605 | 0,1495733 | -2,551135 | 0,0107373 | 0,0372359 | CMTM8 |
| ENSGALG00000012953 | 7564,6193 | -0,381744469 | 0,1101496 | -3,465692 | 0,0005289 | 0,003143 | IPO8 |
| ENSGALG00000007559 | 4279,2164 | -0,381904484 | 0,1214524 | -3,144479 | 0,0016638 | 0,0080933 | PDSS1 |
| ENSGALG00000017333 | 5175,4799 | -0,382465165 | 0,1273235 | -3,003886 | 0,0026656 | 0,0119419 | LCMT2 |
| ENSGALG00000010680 | 1744,2067 | -0,38261968 | 0,1387981 | -2,756664 | 0,0058394 | 0,0227414 | TTC7B |
| ENSGALG00000015226 | 1410,5634 | -0,382624366 | 0,1475175 | -2,593756 | 0,0094934 | 0,0336858 | SPIDR |
| ENSGALG00000008587 | 12026,073 | -0,382953864 | 0,1040771 | -3,679521 | 0,0002337 | 0,0015626 | NDUFAF1 |
| ENSGALG00000008493 | 1237,0726 | -0,38325533 | 0,1561662 | -2,45415 | 0,0141218 | 0,0461297 | METTL21A |
| ENSGALG00000005301 | 7189,628 | -0,383471192 | 0,1309696 | -2,92794 | 0,0034122 | 0,0146283 | CYB5R2 |
| ENSGALG00000027355 | 3798,5712 | -0,383473155 | 0,1422345 | -2,696064 | 0,0070164 | 0,0264622 | TPBG |
| ENSGALG00000001596 | 12885,592 | -0,383722949 | 0,1090252 | -3,51958 | 0,0004322 | 0,0026566 | MLLT1 |
| ENSGALG00000000054 | 13558,561 | -0,383859531 | 0,1440809 | -2,664195 | 0,0077173 | 0,0286687 | OAT |
| ENSGALG00000004839 | 5677,6109 | -0,383869776 | 0,1125796 | -3,409763 | 0,0006502 | 0,003732 | GPATCH1 |
| ENSGALG00000002346 | 7982,3843 | -0,383971209 | 0,1121749 | -3,422968 | 0,0006194 | 0,0035886 | CLSPN |
| ENSGALG00000007556 | 1766,8481 | -0,384355829 | 0,1442884 | -2,663803 | 0,0077263 | 0,0286811 | ORAOV1 |
| ENSGALG00000016040 | 7227,3379 | -0,384424184 | 0,1139331 | -3,374123 | 0,0007405 | 0,0041657 | MORC3 |
| ENSGALG00000011881 | 17920,009 | -0,384788496 | 0,101237 | -3,800869 | 0,0001442 | 0,0010289 | CCNA2 |
| ENSGALG00000016751 | 19933,902 | -0,385163412 | 0,1317503 | -2,923435 | 0,0034619 | 0,0148127 | TMEM131 |
| ENSGALG00000017105 | 3524,6327 | -0,385190009 | 0,1190655 | -3,235111 | 0,001216 | 0,0062852 | RNF6 |
| ENSGALG00000015010 | 9588,3759 | -0,38521292 | 0,1153698 | -3,338939 | 0,000841 | 0,0046244 | NSG1 |
| ENSGALG00000010061 | 5963,3093 | -0,385385873 | 0,159264 | -2,419792 | 0,0155294 | 0,0497371 | LRBA |
| ENSGALG00000008542 | 24254,362 | -0,385688801 | 0,137665 | -2,801647 | 0,0050842 | 0,0203416 | ADD3 |

| | | | | | | | |
|---|---|---|---|---|---|---|---|
| ENSGALG00000015040 | 4593,8104 | -0,38608617 | 0,1172115 | -3,293928 | 0,000988 | 0,0052792 | SLC16A10 |
| ENSGALG00000001876 | 1186,4565 | -0,386144808 | 0,1524506 | -2,532918 | 0,0113118 | 0,0387543 | RASA4 |
| ENSGALG00000013061 | 7957,2214 | -0,38617661 | 0,1127911 | -3,423821 | 0,0006175 | 0,0035805 | PAPD7 |
| ENSGALG00000011357 | 6554,7154 | -0,387039211 | 0,1110437 | -3,485467 | 0,0004913 | 0,0029557 | VEZT |
| ENSGALG00000006259 | 19676,996 | -0,387217866 | 0,1021071 | -3,792272 | 0,0001493 | 0,0010619 | KIRREL |
| ENSGALG00000027774 | 12094,125 | -0,387231267 | 0,1136139 | -3,40831 | 0,0006537 | 0,0037441 | ALMS1 |
| ENSGALG00000002143 | 8359,2157 | -0,387272727 | 0,1268467 | -3,053076 | 0,0022651 | 0,0104398 | ANKLE2 |
| ENSGALG00000008280 | 7969,0775 | -0,387606949 | 0,111093 | -3,48903 | 0,0004848 | 0,002923 | PMPCB |
| ENSGALG00000012724 | 10174,958 | -0,387874274 | 0,1052864 | -3,683992 | 0,0002296 | 0,001541 | PAK1IP1 |
| ENSGALG00000015247 | 7003,5436 | -0,387982807 | 0,1080781 | -3,589838 | 0,0003309 | 0,002117 | GPR161 |
| ENSGALG00000027262 | 21153,65 | -0,388074983 | 0,1013462 | -3,829203 | 0,0001286 | 0,0009315 | CAB39 |
| ENSGALG00000005575 | 18548,779 | -0,388395592 | 0,1080882 | -3,593321 | 0,0003265 | 0,0020938 | AKAP1 |
| ENSGALG00000010609 | 3709,2893 | -0,388677075 | 0,1243181 | -3,126472 | 0,0017692 | 0,0085114 | NDUFAF7 |
| ENSGALG00000005012 | 6065,171 | -0,388993034 | 0,1215819 | -3,199432 | 0,001377 | 0,006959 | ISY1 |
| ENSGALG00000005904 | 32126,543 | -0,389017981 | 0,1104816 | -3,521111 | 0,0004297 | 0,0026439 | TMED5 |
| ENSGALG00000011103 | 30281,636 | -0,389199345 | 0,1020702 | -3,813057 | 0,0001373 | 0,0009872 | PAPOLA |
| ENSGALG00000007788 | 4541,3087 | -0,389226557 | 0,117185 | -3,321472 | 0,0008954 | 0,0048844 | ABCB7 |
| ENSGALG00000001073 | 7165,4084 | -0,389421011 | 0,147194 | -2,645631 | 0,0081539 | 0,0298497 | NR6A1 |
| ENSGALG00000003166 | 4550,9206 | -0,38962163 | 0,1147713 | -3,394765 | 0,0006869 | 0,0039178 | REXO4 |
| ENSGALG00000002129 | 6971,2134 | -0,390332447 | 0,1113848 | -3,504359 | 0,0004577 | 0,0027845 | WDR47 |
| ENSGALG00000027696 | 2856,8564 | -0,390616277 | 0,1467859 | -2,661129 | 0,0077879 | 0,0288524 | |
| ENSGALG00000007485 | 13617,236 | -0,391583137 | 0,1247391 | -3,139217 | 0,001694 | 0,0082195 | TFRC |
| ENSGALG00000008983 | 6819,4343 | -0,391807233 | 0,114877 | -3,410666 | 0,000648 | 0,0037213 | GNB4 |
| ENSGALG00000009364 | 15815,387 | -0,391850394 | 0,1069886 | -3,662544 | 0,0002497 | 0,0016558 | PRKCI |
| ENSGALG00000015396 | 993,36818 | -0,391937266 | 0,1599566 | -2,450272 | 0,0142748 | 0,0464956 | CD200 |
| ENSGALG00000028035 | 5338,9185 | -0,391942986 | 0,1230917 | -3,184154 | 0,0014518 | 0,0072578 | FBXO22 |
| ENSGALG00000025858 | 3504,3476 | -0,392172869 | 0,1322344 | -2,96574 | 0,0030196 | 0,0132318 | LUZP1 |
| ENSGALG00000017064 | 15165,119 | -0,392266132 | 0,1075067 | -3,648759 | 0,0002635 | 0,001734 | RFC3 |
| ENSGALG00000013410 | 8555,6915 | -0,392331865 | 0,1073488 | -3,654738 | 0,0002574 | 0,0016999 | UBA3 |
| ENSGALG00000003277 | 6528,6806 | -0,393093094 | 0,1163756 | -3,377797 | 0,0007307 | 0,0041228 | SUZ12 |
| ENSGALG00000007312 | 1591,4187 | -0,393109181 | 0,1537105 | -2,557464 | 0,0105438 | 0,0367241 | GNPDA1 |
| ENSGALG00000002682 | 5643,0549 | -0,393160876 | 0,1228268 | -3,200938 | 0,0013698 | 0,0069331 | BRD3 |
| ENSGALG00000004481 | 5167,7274 | -0,39336646 | 0,121645 | -3,233724 | 0,0012219 | 0,0063086 | EDA |
| ENSGALG00000015865 | 4560,3514 | -0,393610955 | 0,1160346 | -3,392187 | 0,0006934 | 0,0039501 | FAM46A |
| ENSGALG00000009569 | 11508,232 | -0,393918558 | 0,1069286 | -3,683939 | 0,0002297 | 0,001541 | TLK1 |
| ENSGALG00000009201 | 11168,289 | -0,39393497 | 0,1118548 | -3,521842 | 0,0004286 | 0,0026399 | CCNF |
| ENSGALG00000028133 | 7664,2653 | -0,394703045 | 0,1165319 | -3,387082 | 0,0007064 | 0,0040074 | UBE2V2 |
| ENSGALG00000028883 | 9007,8407 | -0,394818784 | 0,1358341 | -2,906626 | 0,0036535 | 0,0155154 | NOL11 |
| ENSGALG00000028898 | 2507,0991 | -0,395098922 | 0,1286111 | -3,072045 | 0,002126 | 0,0099434 | GLTPD1 |
| ENSGALG00000014166 | 1449,8688 | -0,395137478 | 0,1437153 | -2,749445 | 0,0059696 | 0,0231821 | TSPO |
| ENSGALG00000016467 | 8711,5563 | -0,395365049 | 0,1194699 | -3,309327 | 0,0009352 | 0,0050615 | SMC6 |
| ENSGALG00000003043 | 3489,7523 | -0,395518744 | 0,1229266 | -3,217519 | 0,001293 | 0,0066184 | BBS2 |
| ENSGALG00000017024 | 1501,8199 | -0,395600818 | 0,1535883 | -2,575722 | 0,0100031 | 0,0351736 | NEK3 |
| ENSGALG00000008738 | 1562,619 | -0,395613631 | 0,1395655 | -2,83461 | 0,0045882 | 0,0186403 | |
| ENSGALG00000002076 | 11878,124 | -0,395918706 | 0,1140373 | -3,471836 | 0,0005169 | 0,0030827 | DGCR8 |
| ENSGALG00000008521 | 16274,178 | -0,395949661 | 0,1276174 | -3,10263 | 0,0019181 | 0,0091151 | TP53BP1 |
| ENSGALG00000014688 | 9710,7864 | -0,396059354 | 0,1249828 | -3,16891 | 0,0015301 | 0,0075617 | ZYX |
| ENSGALG00000003273 | 1825,9769 | -0,396166645 | 0,1597575 | -2,4798 | 0,0131456 | 0,0436637 | HSDL1 |
| ENSGALG00000004256 | 8236,2208 | -0,396589984 | 0,1140041 | -3,478736 | 0,0005038 | 0,003015 | DHRS3 |
| ENSGALG00000010643 | 4501,8566 | -0,396627508 | 0,117516 | -3,375094 | 0,0007379 | 0,004154 | ZYG11B |
| ENSGALG00000012759 | 9679,7293 | -0,396656849 | 0,1130301 | -3,509303 | 0,0004493 | 0,0027403 | NUP37 |
| ENSGALG00000004435 | 2804,793 | -0,396797105 | 0,1243897 | -3,189952 | 0,001423 | 0,0071489 | RAB27A |
| ENSGALG00000009608 | 5107,6787 | -0,396825966 | 0,1273293 | -3,116534 | 0,0018299 | 0,0087449 | UBR3 |
| ENSGALG00000009838 | 16688,775 | -0,397053502 | 0,1016373 | -3,906571 | 9,36E-05 | 0,0007093 | AQR |
| ENSGALG00000009533 | 7127,7941 | -0,397081701 | 0,1281809 | -3,097822 | 0,0019495 | 0,0092417 | NMD3 |
| ENSGALG00000006563 | 2968,8404 | -0,397183147 | 0,1427397 | -2,78257 | 0,005393 | 0,0213616 | SEMA3D |
| ENSGALG00000015435 | 3490,2323 | -0,397319834 | 0,1216986 | -3,264786 | 0,0010955 | 0,0057608 | DCAF6 |
| ENSGALG00000010886 | 2623,1957 | -0,397321501 | 0,1327902 | -2,992101 | 0,0027706 | 0,0123558 | PAQR3 |
| ENSGALG00000015215 | 6704,2917 | -0,397390834 | 0,1204862 | -3,298228 | 0,000973 | 0,0052112 | MAPRE2 |
| ENSGALG00000013776 | 3256,8373 | -0,397455529 | 0,1277422 | -3,111389 | 0,0018621 | 0,0088676 | CEP135 |
| ENSGALG00000002675 | 3557,3289 | -0,397531672 | 0,1252523 | -3,173848 | 0,0015043 | 0,0074478 | SCUBE3 |
| ENSGALG00000009322 | 5962,2024 | -0,397973054 | 0,1135762 | -3,504017 | 0,0004583 | 0,0027868 | NVL |
| ENSGALG00000001576 | 13166,504 | -0,398019961 | 0,106825 | -3,725905 | 0,0001946 | 0,0013328 | HIRA |
| ENSGALG00000003675 | 2269,9539 | -0,398220954 | 0,1307081 | -3,046644 | 0,0023141 | 0,0106334 | PRKAR1B |

| | | | | | | | |
|---|---|---|---|---|---|---|---|
| ENSGALG00000010256 | 2730,229 | -0,398269176 | 0,1444457 | -2,757224 | 0,0058294 | 0,0227155 | TIPARP |
| ENSGALG00000000838 | 3236,5634 | -0,398388838 | 0,1429731 | -2,786459 | 0,0053287 | 0,0211518 | THAP12 |
| ENSGALG00000008889 | 7462,9462 | -0,398667809 | 0,107895 | -3,694959 | 0,0002199 | 0,001483 | TTC14 |
| ENSGALG00000010674 | 9662,4596 | -0,399166032 | 0,1113474 | -3,58487 | 0,0003372 | 0,0021516 | DCTD |
| ENSGALG00000011186 | 1570,9959 | -0,399261707 | 0,1516141 | -2,633408 | 0,0084533 | 0,0307484 | ERMARD |
| ENSGALG00000006483 | 5623,947 | -0,399267586 | 0,1210637 | -3,297995 | 0,0009738 | 0,0052135 | AMOTL2 |
| ENSGALG00000004368 | 3768,914 | -0,39990946 | 0,1253746 | -3,189717 | 0,0014241 | 0,0071521 | GNA12 |
| ENSGALG00000001377 | 9703,5691 | -0,400093128 | 0,1073106 | -3,728364 | 0,0001927 | 0,0013239 | POLRMT |
| ENSGALG00000023725 | 1256,3767 | -0,400126756 | 0,1588481 | -2,518927 | 0,0117713 | 0,0399966 | IER5 |
| ENSGALG00000020251 | 2658,2357 | -0,40015479 | 0,1247153 | -3,208547 | 0,0013341 | 0,0067774 | FNIP2 |
| ENSGALG00000007663 | 14617,833 | -0,400171791 | 0,1147124 | -3,488479 | 0,0004858 | 0,0029278 | KDM3B |
| ENSGALG00000016437 | 3164,1402 | -0,40025509 | 0,1368097 | -2,925634 | 0,0034375 | 0,0147223 | TAF1B |
| ENSGALG00000008289 | 17537,453 | -0,401142997 | 0,1117051 | -3,59109 | 0,0003293 | 0,0021088 | RBM19 |
| ENSGALG00000000332 | 20852,835 | -0,401294712 | 0,1053884 | -3,80777 | 0,0001402 | 0,0010059 | FTSJ3 |
| ENSGALG00000016747 | 15836,06 | -0,401466819 | 0,10405 | -3,858403 | 0,0001141 | 0,0008386 | UBE3A |
| ENSGALG00000011542 | 3771,8431 | -0,401482328 | 0,1540301 | -2,606518 | 0,0091468 | 0,0326946 | METTL6 |
| ENSGALG00000001094 | 14665,623 | -0,40156915 | 0,1240977 | -3,235912 | 0,0012125 | 0,0062731 | ADGRD2 |
| ENSGALG00000011493 | 4936,2961 | -0,401631833 | 0,1189649 | -3,376053 | 0,0007353 | 0,0041422 | TMEM169 |
| ENSGALG00000010304 | 4399,9164 | -0,401743756 | 0,1198974 | -3,350729 | 0,000806 | 0,0044728 | MLH3 |
| ENSGALG00000001422 | 12037,379 | -0,402405365 | 0,1260368 | -3,192761 | 0,0014092 | 0,0070902 | RRP9 |
| ENSGALG00000005516 | 8167,2279 | -0,402406965 | 0,1079699 | -3,72703 | 0,0001937 | 0,0013282 | HEATR6 |
| ENSGALG00000010903 | 1862,2125 | -0,40247318 | 0,14557 | -2,764808 | 0,0056956 | 0,0222898 | RASGEF1B |
| ENSGALG00000012138 | 29874,231 | -0,402996238 | 0,1040941 | -3,871462 | 0,0001082 | 0,0008023 | sept-07 |
| ENSGALG00000011379 | 3502,2993 | -0,403264726 | 0,1218523 | -3,309456 | 0,0009348 | 0,0050615 | SLC44A5 |
| ENSGALG00000004486 | 7663,2736 | -0,403537703 | 0,1189928 | -3,391278 | 0,0006957 | 0,0039614 | ZCCHC8 |
| ENSGALG00000011505 | 19437,736 | -0,403647787 | 0,1041549 | -3,875456 | 0,0001064 | 0,0007935 | MARK3 |
| ENSGALG00000007610 | 8753,2545 | -0,403710794 | 0,1080893 | -3,734976 | 0,0001877 | 0,0012942 | CDC23 |
| ENSGALG00000003219 | 2817,8802 | -0,403748967 | 0,1250219 | -3,229426 | 0,0012404 | 0,0063945 | RP11-77K12.7 |
| ENSGALG00000002602 | 3162,965 | -0,403772578 | 0,1577228 | -2,560015 | 0,0104668 | 0,036493 | CSMD2 |
| ENSGALG00000012919 | 4205,6363 | -0,40384181 | 0,1377531 | -2,931636 | 0,0033718 | 0,0144834 | YARS2 |
| ENSGALG00000015033 | 6515,5396 | -0,403855123 | 0,1645709 | -2,453988 | 0,0141282 | 0,0461393 | REV3L |
| ENSGALG00000003033 | 4208,0329 | -0,4040823 | 0,1239451 | -3,260173 | 0,0011134 | 0,0058396 | STXBP4 |
| ENSGALG00000009887 | 7499,0415 | -0,404219345 | 0,1116137 | -3,621592 | 0,0002928 | 0,0019019 | DIEXF |
| ENSGALG00000016070 | 5235,9258 | -0,404284672 | 0,1139162 | -3,548966 | 0,0003867 | 0,0024088 | WRB |
| ENSGALG00000028679 | 2739,4279 | -0,404433944 | 0,1560396 | -2,591867 | 0,0095457 | 0,0338272 | RRP8 |
| ENSGALG00000005558 | 18603,537 | -0,404537586 | 0,1021007 | -3,962142 | 7,43E-05 | 0,0005792 | TBL3 |
| ENSGALG00000003358 | 9790,289 | -0,404595333 | 0,1105122 | -3,661092 | 0,0002511 | 0,0016639 | UBIAD1 |
| ENSGALG00000010428 | 7926,1945 | -0,404704607 | 0,1082049 | -3,74017 | 0,0001839 | 0,0012722 | FAM98A |
| ENSGALG00000012509 | 6088,1178 | -0,40489721 | 0,1172775 | -3,452472 | 0,0005555 | 0,0032739 | ARL5A |
| ENSGALG00000016034 | 11941,122 | -0,4051791 | 0,11639 | -3,481218 | 0,0004991 | 0,0029977 | ATRN |
| ENSGALG00000016397 | 10814,151 | -0,405217096 | 0,1464803 | -2,766359 | 0,0056686 | 0,0222137 | SOX11 |
| ENSGALG00000009015 | 5602,2699 | -0,40522305 | 0,1351963 | -2,997294 | 0,0027239 | 0,0121752 | PIGK |
| ENSGALG00000011405 | 5639,0186 | -0,405273633 | 0,1171817 | -3,458507 | 0,0005432 | 0,0032154 | SLC4A7 |
| ENSGALG00000002042 | 9110,0202 | -0,405517273 | 0,1100944 | -3,683359 | 0,0002302 | 0,001543 | KNOP1 |
| ENSGALG00000015313 | 1727,1874 | -0,405609451 | 0,1673222 | -2,424122 | 0,0153455 | 0,049264 | PDSS2 |
| ENSGALG00000011118 | 5098,9857 | -0,405615226 | 0,1135418 | -3,572385 | 0,0003537 | 0,0022401 | MIER1 |
| ENSGALG00000012756 | 1600,2653 | -0,405783919 | 0,1444512 | -2,809142 | 0,0049674 | 0,0199796 | PARPBP |
| ENSGALG00000004300 | 11426,337 | -0,405827457 | 0,1082534 | -3,748866 | 0,0001776 | 0,0012377 | PTDSS2 |
| ENSGALG00000002107 | 25522,201 | -0,406162719 | 0,1018979 | -3,985977 | 6,72E-05 | 0,0005304 | RANBP1 |
| ENSGALG00000011571 | 3230,4791 | -0,406488263 | 0,1210977 | -3,356697 | 0,0007888 | 0,0044014 | AGPAT4 |
| ENSGALG00000015439 | 3241,7503 | -0,406514431 | 0,1539648 | -2,640308 | 0,0082831 | 0,0302474 | SLC44A1 |
| ENSGALG00000009800 | 11257,087 | -0,406824322 | 0,137832 | -2,951596 | 0,0031614 | 0,0137382 | SETD7 |
| ENSGALG00000014404 | 9644,7108 | -0,407214942 | 0,1124218 | -3,622207 | 0,0002921 | 0,0018983 | GPR125 |
| ENSGALG00000004001 | 29564,347 | -0,407260268 | 0,1220375 | -3,337173 | 0,0008464 | 0,0046501 | LARP1 |
| ENSGALG00000014165 | 2563,4692 | -0,407468645 | 0,1414422 | -2,880813 | 0,0039665 | 0,0165833 | MCAT |
| ENSGALG00000000275 | 5245,7551 | -0,407767805 | 0,1196134 | -3,409049 | 0,0006519 | 0,0037377 | KCTD9 |
| ENSGALG00000013661 | 4102,8132 | -0,40778269 | 0,1293022 | -3,153718 | 0,001612 | 0,007895 | MICU3 |
| ENSGALG00000009534 | 2852,2534 | -0,408065762 | 0,1282621 | -3,1815 | 0,0014651 | 0,0072963 | PLEKHA1 |
| ENSGALG00000013715 | 2414,831 | -0,408079285 | 0,1364191 | -2,991364 | 0,0027773 | 0,0123776 | DLC1 |
| ENSGALG00000003143 | 15625,053 | -0,408123735 | 0,1067545 | -3,823012 | 0,0001318 | 0,0009527 | ZC3H7A |
| ENSGALG00000001616 | 13431,828 | -0,408530697 | 0,1073575 | -3,805328 | 0,0001416 | 0,0010148 | CDC45 |
| ENSGALG00000003995 | 3806,3629 | -0,408610387 | 0,1221472 | -3,34523 | 0,0008221 | 0,0045354 | ERAL1 |
| ENSGALG00000028536 | 1522,6438 | -0,40868801 | 0,1548439 | -2,639355 | 0,0083064 | 0,0303114 | MPEG1 |
| ENSGALG00000011100 | 3743,986 | -0,408960809 | 0,1243724 | -3,288197 | 0,0010083 | 0,0053689 | HIBADH |

| | | | | | | | |
|---|---|---|---|---|---|---|---|
| ENSGALG00000008523 | 12531,511 | -0,409027447 | 0,1050057 | -3,895289 | 9,81E-05 | 0,0007378 | XPNPEP1 |
| ENSGALG00000012607 | 5903,9235 | -0,409282973 | 0,1276687 | -3,20582 | 0,0013468 | 0,0068343 | ZCCHC6 |
| ENSGALG00000009483 | 9325,7747 | -0,409303762 | 0,1092985 | -3,744826 | 0,0001805 | 0,0012537 | MARK1 |
| ENSGALG00000008658 | 2008,7076 | -0,409694565 | 0,1367947 | -2,994959 | 0,0027448 | 0,0122607 | ZNHIT6 |
| ENSGALG00000013858 | 2841,1323 | -0,410414198 | 0,1233001 | -3,328579 | 0,0008729 | 0,0047767 | CEP76 |
| ENSGALG00000002673 | 4173,3236 | -0,41052388 | 0,114577 | -3,582951 | 0,0003397 | 0,0021645 | GRPEL2 |
| ENSGALG00000016345 | 4179,8127 | -0,410542245 | 0,1255035 | -3,271162 | 0,0010711 | 0,0056609 | KLHL15 |
| ENSGALG00000006404 | 23302,915 | -0,411265326 | 0,1022948 | -4,020395 | 5,81E-05 | 0,0004686 | DDX46 |
| ENSGALG00000002830 | 2218,8919 | -0,411355669 | 0,1470401 | -2,797575 | 0,0051488 | 0,0205453 | NUDT3 |
| ENSGALG00000011662 | 25762,115 | -0,412100057 | 0,1171723 | -3,517043 | 0,0004364 | 0,0026703 | NUP205 |
| ENSGALG00000006157 | 6346,9334 | -0,412112134 | 0,1105255 | -3,728661 | 0,0001925 | 0,0013231 | DDX26B |
| ENSGALG00000015944 | 3056,9164 | -0,412472548 | 0,1405652 | -2,934387 | 0,0033421 | 0,0143722 | RIMS1 |
| ENSGALG00000000780 | 20148,91 | -0,413093901 | 0,1091845 | -3,783448 | 0,0001547 | 0,0010956 | SMPD3 |
| ENSGALG00000010734 | 12330,712 | -0,413140268 | 0,115155 | -3,587689 | 0,0003336 | 0,0021305 | PPP4R3A |
| ENSGALG00000027373 | 11590,467 | -0,413207621 | 0,1275875 | -3,238621 | 0,0012011 | 0,0062229 | |
| ENSGALG00000014811 | 9624,081 | -0,413279775 | 0,1068315 | -3,868519 | 0,0001095 | 0,0008103 | SMCHD1 |
| ENSGALG00000000860 | 3991,9401 | -0,413426982 | 0,141168 | -2,928617 | 0,0034047 | 0,0146048 | SLC7A6 |
| ENSGALG00000014464 | 1467,51 | -0,413666995 | 0,1677373 | -2,466159 | 0,0136571 | 0,0449024 | MTR |
| ENSGALG00000027864 | 1064,9642 | -0,413669777 | 0,1682884 | -2,458101 | 0,0139674 | 0,045702 | NFKBIA |
| ENSGALG00000009300 | 6311,4725 | -0,413699825 | 0,1174909 | -3,521122 | 0,0004297 | 0,0026439 | NUMB |
| ENSGALG00000014978 | 5258,7361 | -0,41400107 | 0,1181716 | -3,503389 | 0,0004594 | 0,0027915 | IQGAP2 |
| ENSGALG00000004947 | 6549,1555 | -0,414737624 | 0,1229203 | -3,374037 | 0,0007407 | 0,0041657 | POLR3A |
| ENSGALG00000001613 | 28424,625 | -0,414922322 | 0,1050352 | -3,950318 | 7,80E-05 | 0,0006041 | GNL3 |
| ENSGALG00000013943 | 3049,8006 | -0,414996341 | 0,1265367 | -3,279652 | 0,0010394 | 0,0055126 | FIP1L1 |
| ENSGALG00000012241 | 2635,2581 | -0,415005332 | 0,1298425 | -3,196222 | 0,0013924 | 0,0070187 | KLHDC1 |
| ENSGALG00000014708 | 1768,2325 | -0,415209722 | 0,1395692 | -2,974938 | 0,0029305 | 0,0129247 | GSTK1 |
| ENSGALG00000027593 | 7745,809 | -0,415214085 | 0,1233358 | -3,366535 | 0,0007612 | 0,0042684 | ZNF292 |
| ENSGALG00000000800 | 3898,7504 | -0,415275292 | 0,1200235 | -3,459951 | 0,0005403 | 0,003201 | PRMT7 |
| ENSGALG00000027666 | 1353,3857 | -0,415292257 | 0,1696173 | -2,448407 | 0,014349 | 0,0467036 | TMEM80 |
| ENSGALG00000012608 | 2590,1311 | -0,415354931 | 0,1398222 | -2,970593 | 0,0029723 | 0,0130666 | DAPK1 |
| ENSGALG00000009634 | 10679,835 | -0,415471196 | 0,1072605 | -3,873478 | 0,0001073 | 0,0007966 | GFM1 |
| ENSGALG00000012030 | 9331,9586 | -0,415761326 | 0,140767 | -2,953543 | 0,0031415 | 0,013665 | QSER1 |
| ENSGALG00000016972 | 5736,7284 | -0,415815188 | 0,1184924 | -3,509213 | 0,0004494 | 0,0027403 | NUFIP1 |
| ENSGALG00000026470 | 13532,223 | -0,41655555 | 0,1120502 | -3,71758 | 0,0002011 | 0,0013699 | BOD1L1 |
| ENSGALG00000010996 | 3758,0298 | -0,416979888 | 0,1506983 | -2,766985 | 0,0056577 | 0,0221925 | IRF2BP2 |
| ENSGALG00000003737 | 3492,3176 | -0,417242803 | 0,1329577 | -3,138163 | 0,0017001 | 0,0082461 | TOX3 |
| ENSGALG00000006818 | 7817,9637 | -0,417614565 | 0,1586281 | -2,632664 | 0,0084718 | 0,0307911 | KMT2A |
| ENSGALG00000014386 | 4214,1295 | -0,417751127 | 0,1215158 | -3,437832 | 0,0005864 | 0,0034221 | SEPSECS |
| ENSGALG00000014013 | 11695,533 | -0,418753457 | 0,1054585 | -3,970791 | 7,16E-05 | 0,0005625 | ARHGAP18 |
| ENSGALG00000015132 | 49891,397 | -0,418923213 | 0,1007384 | -4,158527 | 3,20E-05 | 0,000281 | CDH2 |
| ENSGALG00000004121 | 4190,9905 | -0,419252057 | 0,1222891 | -3,428369 | 0,0006072 | 0,003533 | LPAR4 |
| ENSGALG00000013681 | 3809,2674 | -0,419314087 | 0,1641248 | -2,55485 | 0,0106234 | 0,0369255 | ZNF407 |
| ENSGALG00000009961 | 27537,675 | -0,419402034 | 0,1063815 | -3,942434 | 8,07E-05 | 0,0006229 | ABCE1 |
| ENSGALG00000010201 | 4830,0616 | -0,41947829 | 0,1360243 | -3,083848 | 0,0020434 | 0,0096134 | ABHD18 |
| ENSGALG00000006526 | 6573,3308 | -0,419710882 | 0,1536968 | -2,730772 | 0,0063186 | 0,024288 | PLXNB1 |
| ENSGALG00000004716 | 1607,3564 | -0,41985683 | 0,1470278 | -2,855629 | 0,0042952 | 0,0176945 | ZC4H2 |
| ENSGALG00000019228 | 2351,715 | -0,419863222 | 0,1356378 | -3,095474 | 0,001965 | 0,0093023 | SFT2D2 |
| ENSGALG00000007227 | 17010,789 | -0,420130207 | 0,125082 | -3,358839 | 0,0007827 | 0,0043764 | MSI1 |
| ENSGALG00000000975 | 4091,6956 | -0,420207875 | 0,1199453 | -3,50333 | 0,0004595 | 0,0027915 | |
| ENSGALG00000003985 | 6692,4044 | -0,42022626 | 0,1152659 | -3,645711 | 0,0002667 | 0,0017505 | TARSL2 |
| ENSGALG00000007736 | 8209,2388 | -0,420323444 | 0,1102306 | -3,813127 | 0,0001372 | 0,0009872 | ZWILCH |
| ENSGALG00000014827 | 2639,1918 | -0,420567115 | 0,1657257 | -2,537731 | 0,0111574 | 0,0383605 | KIAA0408 |
| ENSGALG00000012188 | 1607,0265 | -0,420636341 | 0,1572383 | -2,675152 | 0,0074695 | 0,0278853 | BBS9 |
| ENSGALG00000008989 | 27837,925 | -0,420708851 | 0,1092049 | -3,852471 | 0,0001169 | 0,0008555 | WDR43 |
| ENSGALG00000017119 | 1864,3003 | -0,420708881 | 0,1365688 | -3,080563 | 0,0020661 | 0,0096988 | TNFRSF19 |
| ENSGALG00000023650 | 764,17678 | -0,420788842 | 0,1697354 | -2,479087 | 0,0131719 | 0,0437404 | |
| ENSGALG00000010630 | 14099,6 | -0,420914921 | 0,1126811 | -3,735454 | 0,0001874 | 0,0012926 | AHCTF1 |
| ENSGALG00000004310 | 3386,8687 | -0,4211147 | 0,1211839 | -3,475005 | 0,0005108 | 0,0030492 | TTYH3 |
| ENSGALG00000010671 | 83381,23 | -0,421121888 | 0,1031007 | -4,084569 | 4,42E-05 | 0,0003679 | HNRNPU |
| ENSGALG00000008466 | 2872,7321 | -0,421406429 | 0,1332906 | -3,16156 | 0,0015693 | 0,0077355 | FAM117B |
| ENSGALG00000027311 | 10812,972 | -0,421551178 | 0,1501476 | -2,807579 | 0,0049915 | 0,0200413 | NGFR |
| ENSGALG00000014937 | 3129,4872 | -0,421603891 | 0,1255788 | -3,357285 | 0,0007871 | 0,0043945 | KPNA5 |
| ENSGALG00000018803 | 3405,6338 | -0,421885923 | 0,1263968 | -3,33779 | 0,0008445 | 0,0046417 | TMEM198 |
| ENSGALG00000011965 | 3023,1564 | -0,422009879 | 0,1389961 | -3,036128 | 0,0023964 | 0,0109339 | ZC3H7B |

| | | | | | | | |
|---|---|---|---|---|---|---|---|
| ENSGALG00000015692 | 1953,5689 | -0,422243248 | 0,1336531 | -3,159247 | 0,0015818 | 0,0077775 | POLN |
| ENSGALG00000012579 | 13996,553 | -0,422359298 | 0,1452155 | -2,908501 | 0,0036317 | 0,0154323 | BAZ2B |
| ENSGALG00000013607 | 7250,9812 | -0,422561681 | 0,11201 | -3,772535 | 0,0001616 | 0,0011382 | FBXO5 |
| ENSGALG00000016276 | 1028,3831 | -0,422833677 | 0,1624303 | -2,60317 | 0,0092366 | 0,0329724 | KHDRBS2 |
| ENSGALG00000015589 | 4158,3548 | -0,422922749 | 0,1597028 | -2,648186 | 0,0080925 | 0,029674 | MSH3 |
| ENSGALG00000006002 | 2601,7776 | -0,422978689 | 0,1302324 | -3,247876 | 0,0011627 | 0,0060536 | WHDC1 |
| ENSGALG00000006652 | 5529,9226 | -0,423060365 | 0,1599744 | -2,644551 | 0,0081799 | 0,0299301 | DIP2C |
| ENSGALG00000027334 | 5378,2134 | -0,423190484 | 0,1394998 | -3,033628 | 0,0024163 | 0,0110138 | |
| ENSGALG00000002763 | 8455,8856 | -0,423193935 | 0,11583 | -3,653579 | 0,0002586 | 0,0017067 | FRZB |
| ENSGALG00000000962 | 2645,4224 | -0,423212428 | 0,1264405 | -3,347126 | 0,0008165 | 0,00451 | PUS3 |
| ENSGALG00000008708 | 9031,5502 | -0,423296325 | 0,1063024 | -3,982003 | 6,83E-05 | 0,0005384 | ESF1 |
| ENSGALG00000004297 | 1889,6269 | -0,42334794 | 0,1444708 | -2,930335 | 0,003386 | 0,014538 | DNAH1 |
| ENSGALG00000014568 | 6675,2521 | -0,423460012 | 0,1137337 | -3,723259 | 0,0001967 | 0,0013449 | RP3-461F17.3 |
| ENSGALG00000012216 | 1240,2281 | -0,423752018 | 0,1593085 | -2,659946 | 0,0078153 | 0,0289303 | BMPR1B |
| ENSGALG00000001850 | 22801,494 | -0,423798382 | 0,1070003 | -3,960721 | 7,47E-05 | 0,0005817 | RARS |
| ENSGALG00000016701 | 2038,8685 | -0,423807866 | 0,1319527 | -3,211817 | 0,001319 | 0,0067183 | |
| ENSGALG00000003288 | 22428,698 | -0,423843105 | 0,1239826 | -3,418569 | 0,0006295 | 0,0036317 | TARS |
| ENSGALG00000009177 | 7539,3588 | -0,423896895 | 0,110067 | -3,851261 | 0,0001175 | 0,0008592 | FMR1 |
| ENSGALG00000012852 | 33011,495 | -0,42431457 | 0,1189985 | -3,565714 | 0,0003629 | 0,0022862 | MKRN1 |
| ENSGALG00000002288 | 13379,292 | -0,424413539 | 0,1057585 | -4,013044 | 5,99E-05 | 0,0004809 | LIG3 |
| ENSGALG00000000644 | 5332,9382 | -0,424448701 | 0,1720142 | -2,467522 | 0,0136052 | 0,0448185 | NFASC |
| ENSGALG00000011177 | 6032,2306 | -0,424449663 | 0,1147226 | -3,699791 | 0,0002158 | 0,0014601 | CEP290 |
| ENSGALG00000005945 | 41976,029 | -0,425016747 | 0,1048948 | -4,051839 | 5,08E-05 | 0,0004173 | CANX |
| ENSGALG00000002119 | 6930,5852 | -0,425364527 | 0,1209038 | -3,518206 | 0,0004345 | 0,0026634 | C16orf62 |
| ENSGALG00000000033 | 3918,6949 | -0,425866952 | 0,1266041 | -3,363768 | 0,0007689 | 0,0043061 | NFU1 |
| ENSGALG00000004181 | 1761,4875 | -0,425938272 | 0,1375166 | -3,09736 | 0,0019525 | 0,0092529 | GAA |
| ENSGALG00000016911 | 5511,4642 | -0,426013919 | 0,1155064 | -3,688227 | 0,0002258 | 0,0015183 | RNF219 |
| ENSGALG00000001229 | 3132,3073 | -0,426291988 | 0,1628978 | -2,616929 | 0,0088725 | 0,0319489 | SKI |
| ENSGALG00000005205 | 5076,6373 | -0,426364463 | 0,1335815 | -3,191793 | 0,0014139 | 0,0071088 | RNFT1 |
| ENSGALG00000016955 | 4576,7081 | -0,426960619 | 0,1306354 | -3,268337 | 0,0010818 | 0,0057067 | VWA8 |
| ENSGALG00000008616 | 22133,522 | -0,427012968 | 0,114208 | -3,738907 | 0,0001848 | 0,0012773 | GTF2H4 |
| ENSGALG00000001181 | 1351,4114 | -0,427049071 | 0,1574822 | -2,711729 | 0,0066933 | 0,0255122 | KCNJ5 |
| ENSGALG00000004104 | 16560,152 | -0,427862285 | 0,1288312 | -3,321108 | 0,0008966 | 0,0048849 | INTS1 |
| ENSGALG00000002262 | 28425,435 | -0,428265663 | 0,1046681 | -4,091655 | 4,28E-05 | 0,0003598 | ETF1 |
| ENSGALG00000009145 | 1321,8829 | -0,428291708 | 0,1635071 | -2,619407 | 0,0088083 | 0,0317766 | STARD9 |
| ENSGALG00000012068 | 1109,9265 | -0,428298706 | 0,1525081 | -2,808367 | 0,0049793 | 0,0200099 | ZGRF1 |
| ENSGALG00000005164 | 1906,2497 | -0,428309171 | 0,1642287 | -2,608005 | 0,0091072 | 0,0325699 | MTG2 |
| ENSGALG00000004776 | 7278,2903 | -0,428490556 | 0,1147583 | -3,733852 | 0,0001886 | 0,0012986 | AP4E1 |
| ENSGALG00000012227 | 10485,474 | -0,428492425 | 0,1091678 | -3,92508 | 8,67E-05 | 0,000665 | TSPAN5 |
| ENSGALG00000017111 | 14958,655 | -0,428517877 | 0,1112294 | -3,852559 | 0,0001169 | 0,0008555 | NUP58 |
| ENSGALG00000008844 | 4547,7203 | -0,428880605 | 0,1351184 | -3,17411 | 0,001503 | 0,0074438 | SLC31A1 |
| ENSGALG00000023571 | 6639,2137 | -0,428973189 | 0,1232423 | -3,480731 | 0,0005 | 0,0030018 | TRIM59 |
| ENSGALG00000014953 | 10323,626 | -0,429472364 | 0,1343816 | -3,195916 | 0,0013939 | 0,0070213 | RWDD1 |
| ENSGALG00000016454 | 3850,1908 | -0,429504414 | 0,1220362 | -3,519484 | 0,0004324 | 0,0026566 | E2F6 |
| ENSGALG00000000346 | 4943,5232 | -0,429865287 | 0,1169676 | -3,675082 | 0,0002378 | 0,0015854 | ZNF606 |
| ENSGALG00000004133 | 28687,584 | -0,430003398 | 0,1149324 | -3,741359 | 0,000183 | 0,0012682 | SRSF10 |
| ENSGALG00000006730 | 17262,845 | -0,430049755 | 0,1029557 | -4,177038 | 2,95E-05 | 0,0002614 | PM5 |
| ENSGALG00000027835 | 5051,5612 | -0,430117051 | 0,1151335 | -3,735813 | 0,0001871 | 0,0012918 | PCBD1 |
| ENSGALG00000001501 | 5273,6643 | -0,430190725 | 0,1139513 | -3,775216 | 0,0001599 | 0,0011272 | MAPK1 |
| ENSGALG00000006500 | 13286,475 | -0,430344251 | 0,1186807 | -3,626068 | 0,0002878 | 0,0018755 | NDE1 |
| ENSGALG00000004986 | 14224,048 | -0,430376609 | 0,1189456 | -3,618265 | 0,0002966 | 0,0019238 | CERCAM |
| ENSGALG00000004756 | 2781,9182 | -0,43117271 | 0,1246958 | -3,457796 | 0,0005446 | 0,0032218 | CEP89 |
| ENSGALG00000015184 | 7792,5962 | -0,431383947 | 0,1228874 | -3,510399 | 0,0004474 | 0,0027311 | TLE4 |
| ENSGALG00000007705 | 251805,66 | -0,431935045 | 0,0988918 | -4,367752 | 1,26E-05 | 0,0001257 | NCL |
| ENSGALG00000011555 | 11516,332 | -0,431962973 | 0,1142249 | -3,781689 | 0,0001558 | 0,0011017 | QKI |
| ENSGALG00000014222 | 26624,515 | -0,431998929 | 0,0998402 | -4,326904 | 1,51E-05 | 0,0001475 | NUP50 |
| ENSGALG00000011013 | 3992,5007 | -0,43257266 | 0,1184225 | -3,65279 | 0,0002594 | 0,0017112 | PCNX2 |
| ENSGALG00000015635 | 5954,0554 | -0,432598021 | 0,1516977 | -2,851712 | 0,0043485 | 0,0178433 | HTT |
| ENSGALG00000008413 | 12232,739 | -0,432788919 | 0,1151054 | -3,759937 | 0,00017 | 0,0011928 | PPP4R1 |
| ENSGALG00000007481 | 6576,2725 | -0,43296248 | 0,1091555 | -3,966473 | 7,29E-05 | 0,0005708 | HACD3 |
| ENSGALG00000006378 | 6236,1912 | -0,43312035 | 0,1311772 | -3,301795 | 0,0009607 | 0,0051719 | LIPA |
| ENSGALG00000013920 | 7232,4329 | -0,433479079 | 0,1210408 | -3,581264 | 0,0003419 | 0,0021744 | SUV39H2 |
| ENSGALG00000002877 | 3840,7613 | -0,433897954 | 0,1361261 | -3,187471 | 0,0014352 | 0,0071946 | LRRC45 |
| ENSGALG00000011548 | 3329,6306 | -0,434202614 | 0,128113 | -3,389215 | 0,0007009 | 0,003983 | PPP1R13B |

| | | | | | | | |
|---|---|---|---|---|---|---|---|
| ENSGALG00000016279 | 1166,2207 | -0,434455337 | 0,1634837 | -2,657485 | 0,0078726 | 0,029087 | RAB23 |
| ENSGALG00000010976 | 5423,3364 | -0,434636867 | 0,1246154 | -3,487825 | 0,000487 | 0,0029336 | CCDC126 |
| ENSGALG00000009930 | 17651,917 | -0,434701609 | 0,1010801 | -4,300564 | 1,70E-05 | 0,000163 | STRN3 |
| ENSGALG00000006221 | 21172,656 | -0,434757063 | 0,1020087 | -4,261961 | 2,03E-05 | 0,0001895 | KIF1A |
| ENSGALG00000028462 | 7479,587 | -0,434862527 | 0,1269541 | -3,425353 | 0,000614 | 0,0035679 | USP36 |
| ENSGALG00000008355 | 3052,8201 | -0,434887931 | 0,1335972 | -3,255217 | 0,0011331 | 0,0059333 | CASP8 |
| ENSGALG00000011543 | 9465,1404 | -0,435034048 | 0,1235278 | -3,521751 | 0,0004287 | 0,0026399 | SDAD1 |
| ENSGALG00000009519 | 11233,674 | -0,43521067 | 0,1059425 | -4,107988 | 3,99E-05 | 0,0003401 | ARG2 |
| ENSGALG00000009598 | 50606,56 | -0,435652779 | 0,1207717 | -3,607242 | 0,0003095 | 0,0019954 | EPRS |
| ENSGALG00000006409 | 11343,942 | -0,436770571 | 0,1258794 | -3,469755 | 0,0005209 | 0,0031026 | PODXL |
| ENSGALG00000028675 | 1287,5833 | -0,436847257 | 0,1749597 | -2,496845 | 0,0125304 | 0,0420822 | |
| ENSGALG00000001717 | 40955,931 | -0,437024576 | 0,1153453 | -3,788837 | 0,0001514 | 0,0010755 | SLC16A1 |
| ENSGALG00000012680 | 2735,4526 | -0,437301833 | 0,1231056 | -3,552249 | 0,000382 | 0,0023844 | E2F3 |
| ENSGALG00000026093 | 981,75895 | -0,437495765 | 0,1708286 | -2,561021 | 0,0104365 | 0,0364061 | PTPRN |
| ENSGALG00000011475 | 4517,3627 | -0,437819648 | 0,1241287 | -3,527143 | 0,0004201 | 0,0025938 | SMARCAL1 |
| ENSGALG00000014375 | 10299,481 | -0,437825336 | 0,1079876 | -4,054405 | 5,03E-05 | 0,0004135 | |
| ENSGALG00000017038 | 6592,4492 | -0,437833021 | 0,1366122 | -3,204934 | 0,0013509 | 0,0068503 | PROSER1 |
| ENSGALG00000012642 | 14397,962 | -0,437930988 | 0,1176937 | -3,720938 | 0,0001985 | 0,0013557 | RIC8B |
| ENSGALG00000008796 | 7379,0553 | -0,438293488 | 0,1157441 | -3,786746 | 0,0001526 | 0,0010829 | RPF1 |
| ENSGALG00000007999 | 6686,6947 | -0,438325223 | 0,1112507 | -3,939978 | 8,15E-05 | 0,0006287 | MRPS7 |
| ENSGALG00000009448 | 19270,632 | -0,43880669 | 0,1070346 | -4,099673 | 4,14E-05 | 0,0003505 | IFRD1 |
| ENSGALG00000008163 | 17633,406 | -0,439350797 | 0,1262915 | -3,478863 | 0,0005035 | 0,0030149 | PSME4 |
| ENSGALG00000004522 | 17438,777 | -0,439397212 | 0,1129665 | -3,889624 | 0,0001004 | 0,0007527 | |
| ENSGALG00000005854 | 8969,9116 | -0,439557852 | 0,1134785 | -3,87349 | 0,0001073 | 0,0007966 | CLUH |
| ENSGALG00000014870 | 12682,587 | -0,439767118 | 0,1169979 | -3,75876 | 0,0001708 | 0,0011965 | HSF2 |
| ENSGALG00000009731 | 11305,313 | -0,439893222 | 0,106114 | -4,145476 | 3,39E-05 | 0,000295 | ARHGAP11A |
| ENSGALG00000006427 | 15012,164 | -0,439960428 | 0,10348 | -4,251647 | 2,12E-05 | 0,0001975 | CARS |
| ENSGALG00000012497 | 3402,6563 | -0,440007974 | 0,1212931 | -3,627643 | 0,000286 | 0,001865 | SLC12A7 |
| ENSGALG00000022802 | 18215,83 | -0,44018485 | 0,113024 | -3,894614 | 9,84E-05 | 0,0007387 | PWP2 |
| ENSGALG00000000430 | 10996,849 | -0,440297246 | 0,1104513 | -3,986348 | 6,71E-05 | 0,0005299 | SPPL2B |
| ENSGALG00000007145 | 45254,09 | -0,440432711 | 0,0991199 | -4,443434 | 8,85E-06 | 9,27E-05 | ITGB1 |
| ENSGALG00000002732 | 1764,9109 | -0,440734535 | 0,1504889 | -2,928684 | 0,003404 | 0,0146048 | AKAP7 |
| ENSGALG00000003071 | 6465,8929 | -0,441229907 | 0,1183654 | -3,727693 | 0,0001932 | 0,0013256 | OGFOD1 |
| ENSGALG00000015049 | 15802,948 | -0,441268065 | 0,1109947 | -3,975577 | 7,02E-05 | 0,0005529 | AMD1 |
| ENSGALG00000008988 | 11231,556 | -0,442984401 | 0,1126159 | -3,933587 | 8,37E-05 | 0,0006437 | ZZZ3 |
| ENSGALG00000006403 | 20700,703 | -0,442987653 | 0,1077974 | -4,109445 | 3,97E-05 | 0,0003388 | MESDC2 |
| ENSGALG00000010957 | 23588,305 | -0,443064247 | 0,1020626 | -4,341102 | 1,42E-05 | 0,0001402 | USP1 |
| ENSGALG00000004820 | 1103,092 | -0,443483479 | 0,1622612 | -2,733145 | 0,0062733 | 0,0241546 | FAM57A |
| ENSGALG00000020879 | 1971,7439 | -0,443502872 | 0,137058 | -3,235877 | 0,0012127 | 0,0062731 | SLC30A7 |
| ENSGALG00000012696 | 9654,978 | -0,443665071 | 0,1168806 | -3,795883 | 0,0001471 | 0,0010476 | RANBP9 |
| ENSGALG00000012000 | 7157,9161 | -0,444031788 | 0,1096112 | -4,05097 | 5,10E-05 | 0,0004182 | METTL14 |
| ENSGALG00000005206 | 7037,81 | -0,444288079 | 0,1130121 | -3,931331 | 8,45E-05 | 0,0006489 | ACSL3 |
| ENSGALG00000003861 | 8032,1428 | -0,444434747 | 0,1081924 | -4,107819 | 3,99E-05 | 0,0003401 | HERC4 |
| ENSGALG00000005241 | 3696,959 | -0,444802963 | 0,1260848 | -3,527809 | 0,000419 | 0,0025884 | DYNC1LI2 |
| ENSGALG00000029011 | 22651,885 | -0,444941401 | 0,1003035 | -4,435953 | 9,17E-06 | 9,53E-05 | SLC35F1 |
| ENSGALG00000026971 | 4341,3764 | -0,445130127 | 0,1193093 | -3,730892 | 0,0001908 | 0,0013127 | RRAGD |
| ENSGALG00000026753 | 1244,1641 | -0,445460857 | 0,1753388 | -2,540573 | 0,0110671 | 0,0381272 | FSBP |
| ENSGALG00000007704 | 19378,595 | -0,445645488 | 0,1426064 | -3,125003 | 0,001778 | 0,008545 | NPM3 |
| ENSGALG00000010786 | 1210,4793 | -0,44600197 | 0,1746834 | -2,553203 | 0,0106737 | 0,0370627 | DGKB |
| ENSGALG00000003400 | 9822,1857 | -0,446090625 | 0,1073373 | -4,155972 | 3,24E-05 | 0,0002836 | AIMP2 |
| ENSGALG00000009020 | 9304,4549 | -0,446477137 | 0,1140153 | -3,91594 | 9,01E-05 | 0,0006861 | JAG1 |
| ENSGALG00000000619 | 1429,9309 | -0,446602215 | 0,1730494 | -2,580779 | 0,0098578 | 0,0347614 | ANGPTL4 |
| ENSGALG00000013131 | 1864,5844 | -0,447051346 | 0,1333424 | -3,352657 | 0,0008004 | 0,0044479 | C18orf21 |
| ENSGALG00000002536 | 10005,2 | -0,447408092 | 0,1148794 | -3,894588 | 9,84E-05 | 0,0007387 | STX2 |
| ENSGALG00000002863 | 21477,092 | -0,447764784 | 0,1264663 | -3,540586 | 0,0003992 | 0,002482 | ERGIC1 |
| ENSGALG00000009645 | 4180,3141 | -0,447866483 | 0,1704432 | -2,627659 | 0,0085975 | 0,0311647 | ESRRG |
| ENSGALG00000007437 | 4980,3271 | -0,447866955 | 0,1129748 | -3,96431 | 7,36E-05 | 0,0005753 | DPP8 |
| ENSGALG00000012914 | 8462,3227 | -0,447972492 | 0,15259 | -2,935792 | 0,003327 | 0,0143208 | PRKDC |
| ENSGALG00000010612 | 2716,3259 | -0,44797699 | 0,144448 | -3,101302 | 0,0019267 | 0,0091476 | PRKD3 |
| ENSGALG00000013568 | 1779,8432 | -0,448058074 | 0,1709275 | -2,621333 | 0,0087587 | 0,0316302 | NR4A3 |
| ENSGALG00000016385 | 1906,4631 | -0,448510168 | 0,1379255 | -3,251829 | 0,0011467 | 0,0059985 | TSSC1 |
| ENSGALG00000000424 | 6758,6507 | -0,448587625 | 0,1462417 | -3,067441 | 0,002159 | 0,0100529 | CMTM3 |
| ENSGALG00000006656 | 4293,7935 | -0,448668509 | 0,1620852 | -2,768103 | 0,0056384 | 0,0221356 | FRMD4A |
| ENSGALG00000011642 | 18424,345 | -0,448893503 | 0,1458916 | -3,076897 | 0,0020917 | 0,0098032 | ANKRD17 |

| | | | | | | | |
|---|---|---|---|---|---|---|---|
| ENSGALG00000001845 | 10040,892 | -0,449444654 | 0,1070784 | -4,197342 | 2,70E-05 | 0,0002425 | PHTF1 |
| ENSGALG00000012568 | 2033,2155 | -0,449796718 | 0,1384814 | -3,248067 | 0,0011619 | 0,0060519 | TIMP3 |
| ENSGALG00000006324 | 5540,5991 | -0,450353768 | 0,1447725 | -3,110768 | 0,001866 | 0,0088831 | PFKFB3 |
| ENSGALG00000012598 | 6595,5132 | -0,450640697 | 0,1225301 | -3,677795 | 0,0002353 | 0,0015709 | NAA35 |
| ENSGALG00000002519 | 3426,1294 | -0,450983038 | 0,1241584 | -3,632319 | 0,0002809 | 0,0018351 | SLC25A33 |
| ENSGALG00000007652 | 11015,319 | -0,45121126 | 0,1103319 | -4,089582 | 4,32E-05 | 0,0003619 | SUCLG2 |
| ENSGALG00000005959 | 3317,973 | -0,451515508 | 0,1209724 | -3,732383 | 0,0001897 | 0,0013056 | GLMN |
| ENSGALG00000016803 | 4775,6071 | -0,451555414 | 0,1329918 | -3,395363 | 0,0006854 | 0,0039109 | ST6GAL2 |
| ENSGALG00000002663 | 8380,3175 | -0,451591009 | 0,1168807 | -3,863693 | 0,0001117 | 0,0008235 | ATR |
| ENSGALG00000007075 | 11514,123 | -0,451627608 | 0,1269002 | -3,558919 | 0,0003724 | 0,0023386 | ATP13A3 |
| ENSGALG00000004049 | 11640,773 | -0,45169714 | 0,107806 | -4,189907 | 2,79E-05 | 0,0002495 | ITFG1 |
| ENSGALG00000013579 | 24869,718 | -0,451701092 | 0,1629664 | -2,771743 | 0,0055757 | 0,0219585 | FAT1 |
| ENSGALG00000013233 | 9834,1286 | -0,4517759 | 0,1141463 | -3,957866 | 7,56E-05 | 0,0005873 | PPP1R9B |
| ENSGALG00000006336 | 7051,0846 | -0,452052929 | 0,1247016 | -3,625078 | 0,0002889 | 0,0018818 | PAXIP1 |
| ENSGALG00000007659 | 35189,399 | -0,45246564 | 0,1027053 | -4,405474 | 1,06E-05 | 0,0001081 | HYOU1 |
| ENSGALG00000008088 | 3813,1419 | -0,452924609 | 0,1170567 | -3,869275 | 0,0001092 | 0,0008088 | ACSL4 |
| ENSGALG00000004291 | 6511,6259 | -0,453004371 | 0,1491475 | -3,037292 | 0,0023871 | 0,01091 | VPS13D |
| ENSGALG00000008848 | 8858,4601 | -0,453382128 | 0,1083977 | -4,182581 | 2,88E-05 | 0,0002563 | SOX2 |
| ENSGALG00000009835 | 14808,429 | -0,453500739 | 0,1117239 | -4,059121 | 4,93E-05 | 0,0004065 | XPOT |
| ENSGALG00000023480 | 1643,2534 | -0,453589407 | 0,1642924 | -2,760867 | 0,0057648 | 0,0225023 | FBXO39 |
| ENSGALG00000015458 | 9174,0129 | -0,453606012 | 0,1307234 | -3,469967 | 0,0005205 | 0,0031015 | ILDR2 |
| ENSGALG00000005716 | 3689,9521 | -0,453921562 | 0,1547747 | -2,932789 | 0,0033593 | 0,0144372 | SBF2 |
| ENSGALG00000004971 | 14464,929 | -0,454228028 | 0,152979 | -2,969218 | 0,0029856 | 0,0131129 | URM1 |
| ENSGALG00000005840 | 6869,8394 | -0,454270613 | 0,1158533 | -3,921086 | 8,82E-05 | 0,0006743 | DENND5A |
| ENSGALG00000016074 | 1091,7628 | -0,454409142 | 0,1579184 | -2,877493 | 0,0040085 | 0,0167153 | RIMS2 |
| ENSGALG00000006094 | 10694,318 | -0,454736184 | 0,1116856 | -4,071574 | 4,67E-05 | 0,000387 | PHF6 |
| ENSGALG00000004713 | 12581,373 | -0,454819325 | 0,1044249 | -4,355468 | 1,33E-05 | 0,0001321 | LAS1L |
| ENSGALG00000005160 | 9338,8368 | -0,454934871 | 0,1255105 | -3,624675 | 0,0002893 | 0,001883 | VMP1 |
| ENSGALG00000021883 | 1337,969 | -0,455471836 | 0,1467354 | -3,104035 | 0,001909 | 0,0090783 | |
| ENSGALG00000005489 | 5406,3342 | -0,455551792 | 0,1294829 | -3,518238 | 0,0004344 | 0,0026634 | DDX52 |
| ENSGALG00000016196 | 5946,1092 | -0,455745009 | 0,1321926 | -3,447582 | 0,0005656 | 0,0033251 | CBS |
| ENSGALG00000011191 | 11563,883 | -0,455971006 | 0,1072313 | -4,252219 | 2,12E-05 | 0,0001972 | EML1 |
| ENSGALG00000008839 | 9872,7485 | -0,456849267 | 0,1286669 | -3,550635 | 0,0003843 | 0,0023968 | ATP11B |
| ENSGALG00000008415 | 6540,8044 | -0,456904989 | 0,1186805 | -3,849874 | 0,0001182 | 0,0008637 | ALS2 |
| ENSGALG00000000884 | 7194,478 | -0,456928071 | 0,1125409 | -4,060106 | 4,91E-05 | 0,000405 | CXXC5 |
| ENSGALG00000021592 | 966,74998 | -0,457097931 | 0,1634651 | -2,796303 | 0,0051691 | 0,0206083 | TLCD2 |
| ENSGALG00000002338 | 29369,786 | -0,457290151 | 0,1429103 | -3,199841 | 0,001375 | 0,0069517 | ASPM |
| ENSGALG00000001179 | 4301,5313 | -0,45735712 | 0,1154017 | -3,963175 | 7,40E-05 | 0,0005777 | ARMC8 |
| ENSGALG00000023193 | 1113,686 | -0,457481355 | 0,1527665 | -2,994644 | 0,0027477 | 0,0122653 | SOCS1B |
| ENSGALG00000002954 | 6734,124 | -0,457794297 | 0,1205807 | -3,79658 | 0,0001467 | 0,0010458 | MBTD1 |
| ENSGALG00000008497 | 15581,475 | -0,457962816 | 0,1156674 | -3,959307 | 7,52E-05 | 0,0005841 | SLC38A2 |
| ENSGALG00000007777 | 3900,1787 | -0,45814234 | 0,1287478 | -3,558449 | 0,0003731 | 0,0023406 | SLC39A10 |
| ENSGALG00000026322 | 5686,9342 | -0,458344915 | 0,1182055 | -3,877526 | 0,0001055 | 0,0007877 | ARMT1 |
| ENSGALG00000013647 | 1849,9954 | -0,458404091 | 0,1358218 | -3,375041 | 0,000738 | 0,004154 | TFB1M |
| ENSGALG00000026639 | 1529,3689 | -0,458562471 | 0,139721 | -3,281988 | 0,0010308 | 0,0054735 | SMCO4 |
| ENSGALG00000004531 | 7125,0235 | -0,458962037 | 0,1117417 | -4,107348 | 4,00E-05 | 0,0003406 | PLOD1 |
| ENSGALG00000007842 | 5019,27 | -0,459124954 | 0,1174579 | -3,908847 | 9,27E-05 | 0,0007038 | PDHX |
| ENSGALG00000016930 | 13372,763 | -0,459461549 | 0,1064549 | -4,316019 | 1,59E-05 | 0,0001534 | DIS3 |
| ENSGALG00000008705 | 14944,012 | -0,4597037 | 0,1096313 | -4,193181 | 2,75E-05 | 0,0002462 | TRABD |
| ENSGALG00000015773 | 9016,2735 | -0,459782065 | 0,1064627 | -4,318717 | 1,57E-05 | 0,0001521 | PTCD3 |
| ENSGALG00000008482 | 22772,22 | -0,459855147 | 0,1105258 | -4,160612 | 3,17E-05 | 0,000279 | STAG2 |
| ENSGALG00000013406 | 738,83887 | -0,460037256 | 0,1721459 | -2,672369 | 0,0075318 | 0,0280715 | EOGT |
| ENSGALG00000010411 | 14781,22 | -0,460452211 | 0,1181671 | -3,896619 | 9,75E-05 | 0,000735 | RNF13 |
| ENSGALG00000002501 | 3377,4292 | -0,46100497 | 0,1227582 | -3,755392 | 0,0001731 | 0,0012109 | RP11-196G11.1 |
| ENSGALG00000006281 | 5170,2004 | -0,461106682 | 0,1148963 | -4,013241 | 5,99E-05 | 0,0004808 | GTF2H1 |
| ENSGALG00000006078 | 2849,4853 | -0,461126659 | 0,1296477 | -3,556767 | 0,0003754 | 0,0023524 | PALB2 |
| ENSGALG00000016216 | 6550,317 | -0,461519219 | 0,1642494 | -2,809869 | 0,0049562 | 0,0199462 | NDRG1 |
| ENSGALG00000013893 | 1508,8689 | -0,461829974 | 0,1439203 | -3,208929 | 0,0013323 | 0,0067735 | MNS1 |
| ENSGALG00000015633 | 9547,4016 | -0,462141421 | 0,1057703 | -4,369294 | 1,25E-05 | 0,0001252 | TERF1 |
| ENSGALG00000016980 | 3122,8167 | -0,462357479 | 0,1258227 | -3,674674 | 0,0002382 | 0,0015871 | COG3 |
| ENSGALG00000013136 | 2283,1481 | -0,462408191 | 0,131854 | -3,506971 | 0,0004532 | 0,0027598 | AEBP2 |
| ENSGALG00000000687 | 3686,837 | -0,462411333 | 0,1264523 | -3,656806 | 0,0002554 | 0,0016878 | KIAA0556 |
| ENSGALG00000013306 | 1859,3469 | -0,463147904 | 0,1696975 | -2,729256 | 0,0063477 | 0,0243862 | DAGLA |
| ENSGALG00000016445 | 7095,5905 | -0,463408352 | 0,1140354 | -4,063725 | 4,83E-05 | 0,000399 | NOL10 |

| | | | | | | | |
|---|---|---|---|---|---|---|---|
| ENSGALG00000009899 | 35232,794 | -0,46345514 | 0,1085537 | -4,269364 | 1,96E-05 | 0,0001844 | CAND1 |
| ENSGALG00000009974 | 634,62537 | -0,463813865 | 0,1807038 | -2,566708 | 0,0102669 | 0,0359195 | MMAA |
| ENSGALG00000011794 | 10368,91 | -0,463824908 | 0,1147491 | -4,042078 | 5,30E-05 | 0,0004329 | DDX47 |
| ENSGALG00000007057 | 3861,2618 | -0,46473909 | 0,1208896 | -3,844327 | 0,0001209 | 0,0008806 | LRRC28 |
| ENSGALG00000010234 | 635,90171 | -0,464990416 | 0,1851223 | -2,511801 | 0,0120117 | 0,0406308 | MMACHC |
| ENSGALG00000012655 | 11727,465 | -0,4651119 | 0,1097394 | -4,23833 | 2,25E-05 | 0,0002077 | POLR3B |
| ENSGALG00000011630 | 13721,826 | -0,465130144 | 0,1774121 | -2,62175 | 0,008748 | 0,0316092 | GLI2 |
| ENSGALG00000015138 | 14137,444 | -0,465157078 | 0,1241857 | -3,745659 | 0,0001799 | 0,0012504 | TMEM2 |
| ENSGALG00000009507 | 1700,6819 | -0,465226456 | 0,1787644 | -2,602456 | 0,0092559 | 0,0330238 | PLEKHH1 |
| ENSGALG00000011677 | 985,1206 | -0,465468876 | 0,1657897 | -2,807586 | 0,0049914 | 0,0200413 | HSPBAP1 |
| ENSGALG00000002942 | 14672,633 | -0,465828236 | 0,1343592 | -3,467036 | 0,0005262 | 0,0031314 | JMJD1C |
| ENSGALG00000014217 | 2291,1249 | -0,46602463 | 0,1389942 | -3,352834 | 0,0007999 | 0,0044469 | GUF1 |
| ENSGALG00000010054 | 3266,3331 | -0,466429533 | 0,1270851 | -3,670215 | 0,0002423 | 0,0016127 | RP11-561B11.2 |
| ENSGALG00000004158 | 2874,2398 | -0,467330867 | 0,1229214 | -3,801867 | 0,0001436 | 0,0010258 | GPT2 |
| ENSGALG00000028849 | 26828,63 | -0,467375912 | 0,1168899 | -3,99843 | 6,38E-05 | 0,0005068 | SCAF11 |
| ENSGALG00000011309 | 5857,3376 | -0,467663025 | 0,1226263 | -3,813725 | 0,0001369 | 0,0009866 | PLXNC1 |
| ENSGALG00000013728 | 12921,155 | -0,467853342 | 0,1080614 | -4,329513 | 1,49E-05 | 0,0001465 | PPAT |
| ENSGALG00000013765 | 8392,681 | -0,468026469 | 0,1155365 | -4,050895 | 5,10E-05 | 0,0004182 | RP3-468K18.7 |
| ENSGALG00000002282 | 2493,1225 | -0,468163907 | 0,1304601 | -3,588561 | 0,0003325 | 0,0021254 | TMEM200A |
| ENSGALG00000000895 | 27062,291 | -0,468166268 | 0,1101353 | -4,250829 | 2,13E-05 | 0,0001979 | NUDC |
| ENSGALG00000017309 | 912,73532 | -0,468669212 | 0,1805876 | -2,595246 | 0,0094523 | 0,0335839 | RNF169 |
| ENSGALG00000002655 | 6693,9795 | -0,468798244 | 0,1325586 | -3,536536 | 0,0004054 | 0,0025147 | ITGAV |
| ENSGALG00000006121 | 11011,198 | -0,468808823 | 0,116573 | -4,021589 | 5,78E-05 | 0,0004665 | PIK3C2A |
| ENSGALG00000011232 | 2855,7622 | -0,469477808 | 0,1239226 | -3,788475 | 0,0001516 | 0,0010765 | STK11IP |
| ENSGALG00000006638 | 2375,8984 | -0,469621749 | 0,1392735 | -3,37194 | 0,0007464 | 0,0041941 | GRIK4 |
| ENSGALG00000009954 | 1375,372 | -0,469791805 | 0,1450112 | -3,239694 | 0,0011966 | 0,0062039 | DYNC2LI1 |
| ENSGALG00000007058 | 9479,7297 | -0,469857249 | 0,1077974 | -4,358706 | 1,31E-05 | 0,0001303 | LSG1 |
| ENSGALG00000011101 | 5741,1035 | -0,47005983 | 0,1114966 | -4,215912 | 2,49E-05 | 0,0002267 | URB2 |
| ENSGALG00000001424 | 896,527 | -0,470247 | 0,1654153 | -2,842827 | 0,0044715 | 0,0182515 | YDJC |
| ENSGALG00000010434 | 945,24487 | -0,470849898 | 0,1876053 | -2,509791 | 0,0120803 | 0,0407919 | CP |
| ENSGALG00000000843 | 7211,6886 | -0,471124949 | 0,1430863 | -3,292592 | 0,0009927 | 0,0053002 | DOT1L |
| ENSGALG00000011246 | 5026,827 | -0,471179653 | 0,1165719 | -4,041966 | 5,30E-05 | 0,0004329 | CHPF |
| ENSGALG00000007564 | 1255,8688 | -0,471313446 | 0,1490623 | -3,161856 | 0,0015677 | 0,0077305 | |
| ENSGALG00000011537 | 2857,0805 | -0,471339277 | 0,1817158 | -2,593827 | 0,0094914 | 0,0336858 | PDE10A |
| ENSGALG00000016836 | 6379,6617 | -0,471449962 | 0,1187504 | -3,970091 | 7,18E-05 | 0,0005636 | TUBGCP3 |
| ENSGALG00000002439 | 1784,311 | -0,471657703 | 0,1533679 | -3,075335 | 0,0021027 | 0,0098513 | BDP1 |
| ENSGALG00000000721 | 4145,8062 | -0,47173717 | 0,1427965 | -3,303562 | 0,0009546 | 0,0051475 | SLC41A1 |
| ENSGALG00000008837 | 13730,848 | -0,471745405 | 0,1080446 | -4,366209 | 1,26E-05 | 0,0001264 | PELI1 |
| ENSGALG00000026412 | 3599,375 | -0,471881516 | 0,1379941 | -3,419578 | 0,0006272 | 0,0036213 | |
| ENSGALG00000016195 | 2664,1039 | -0,472549861 | 0,1405226 | -3,362804 | 0,0007716 | 0,0043194 | COL22A1 |
| ENSGALG00000004228 | 2475,7819 | -0,472593441 | 0,1261151 | -3,747317 | 0,0001787 | 0,0012448 | TRAF3IP1 |
| ENSGALG00000012647 | 9993,6453 | -0,472701385 | 0,125933 | -3,753594 | 0,0001743 | 0,0012177 | RFX4 |
| ENSGALG00000001627 | 4772,9003 | -0,472962819 | 0,1154681 | -4,096047 | 4,20E-05 | 0,0003545 | CPNE1 |
| ENSGALG00000026192 | 2515,1802 | -0,473341595 | 0,1470736 | -3,218399 | 0,0012891 | 0,0066005 | ST8SIA4 |
| ENSGALG00000028267 | 15487,152 | -0,473386218 | 0,114594 | -4,130988 | 3,61E-05 | 0,0003117 | GOLPH3 |
| ENSGALG00000006296 | 11149,265 | -0,47366212 | 0,1093498 | -4,331622 | 1,48E-05 | 0,0001453 | SON |
| ENSGALG00000014903 | 1822,8102 | -0,474039661 | 0,1650108 | -2,87278 | 0,0040688 | 0,0169304 | ITGA2 |
| ENSGALG00000004055 | 3994,9616 | -0,474043343 | 0,1194933 | -3,967113 | 7,27E-05 | 0,0005696 | C7orf50 |
| ENSGALG00000027678 | 2192,33 | -0,474178987 | 0,1641393 | -2,888881 | 0,0038662 | 0,0162692 | SLC35G1 |
| ENSGALG00000027440 | 2934,027 | -0,474361212 | 0,127321 | -3,725711 | 0,0001948 | 0,0013332 | COX20 |
| ENSGALG00000004889 | 8112,9525 | -0,474641509 | 0,1093142 | -4,341993 | 1,41E-05 | 0,0001397 | SLC2A1 |
| ENSGALG00000005211 | 6813,7998 | -0,474686708 | 0,1155357 | -4,108572 | 3,98E-05 | 0,0003396 | NAE1 |
| ENSGALG00000026034 | 1070,3788 | -0,47474594 | 0,1567205 | -3,029253 | 0,0024516 | 0,0111484 | RGS17 |
| ENSGALG00000010182 | 7219,0566 | -0,474781283 | 0,1123939 | -4,224264 | 2,40E-05 | 0,0002195 | SLC29A1 |
| ENSGALG00000015925 | 3585,1162 | -0,475263897 | 0,1282784 | -3,704941 | 0,0002114 | 0,0014344 | MB21D1 |
| ENSGALG00000026989 | 16347,278 | -0,47543634 | 0,1239145 | -3,83681 | 0,0001246 | 0,0009056 | |
| ENSGALG00000006698 | 967,37858 | -0,475588456 | 0,1653212 | -2,876753 | 0,0040179 | 0,0167494 | ABCC6 |
| ENSGALG00000002783 | 1895,8368 | -0,475653855 | 0,1505722 | -3,158976 | 0,0015832 | 0,0077802 | SLC43A2 |
| ENSGALG00000016409 | 21993,092 | -0,475709899 | 0,1119047 | -4,251027 | 2,13E-05 | 0,0001979 | KIDINS220 |
| ENSGALG00000004957 | 935,39904 | -0,476296229 | 0,1744085 | -2,730924 | 0,0063157 | 0,024288 | RHBDD1 |
| ENSGALG00000008686 | 2315,3907 | -0,476496754 | 0,1292591 | -3,686368 | 0,0002275 | 0,0015287 | TRDMT1 |
| ENSGALG00000006906 | 7509,2168 | -0,477103418 | 0,1079356 | -4,420261 | 9,86E-06 | 0,0001016 | CHM |
| ENSGALG00000009841 | 1397,5061 | -0,477667905 | 0,1552355 | -3,077053 | 0,0020906 | 0,0098032 | FASTKD1 |
| ENSGALG00000011342 | 3208,6738 | -0,478233256 | 0,1425056 | -3,355892 | 0,0007911 | 0,0044106 | FGD6 |

| | | | | | | | |
|---|---|---|---|---|---|---|---|
| ENSGALG00000002045 | 30343,402 | -0,478318973 | 0,0998188 | -4,791874 | 1,65E-06 | 2,05E-05 | EIF2S2 |
| ENSGALG00000002460 | 17010 | -0,480154381 | 0,105142 | -4,566722 | 4,95E-06 | 5,50E-05 | WDR75 |
| ENSGALG00000002862 | 5434,2601 | -0,480739106 | 0,1110641 | -4,328483 | 1,50E-05 | 0,0001471 | RRN3 |
| ENSGALG00000012484 | 32855,96 | -0,4809575 | 0,1211628 | -3,969513 | 7,20E-05 | 0,0005645 | RIF1 |
| ENSGALG00000010884 | 11252,816 | -0,481115067 | 0,1148894 | -4,187635 | 2,82E-05 | 0,0002518 | DDX24 |
| ENSGALG00000015901 | 15729,886 | -0,481643321 | 0,1054274 | -4,568482 | 4,91E-06 | 5,46E-05 | SENP6 |
| ENSGALG00000023516 | 18924,681 | -0,48179177 | 0,1052391 | -4,578068 | 4,69E-06 | 5,25E-05 | PHRF1 |
| ENSGALG00000002294 | 3404,1182 | -0,482168758 | 0,1282462 | -3,759711 | 0,0001701 | 0,0011932 | SEMA6A |
| ENSGALG00000007948 | 1369,1179 | -0,482500979 | 0,1814408 | -2,659276 | 0,0078309 | 0,0289722 | LRRC4C |
| ENSGALG00000001612 | 1053,9675 | -0,482944134 | 0,1589083 | -3,039137 | 0,0023726 | 0,0108507 | HEXDC |
| ENSGALG00000003078 | 20488,019 | -0,483195985 | 0,106301 | -4,545546 | 5,48E-06 | 5,97E-05 | MMD |
| ENSGALG00000008765 | 5344,7574 | -0,483352119 | 0,1278807 | -3,779711 | 0,000157 | 0,0011099 | C1D |
| ENSGALG00000005134 | 3683,9209 | -0,483437706 | 0,1457126 | -3,317748 | 0,0009075 | 0,0049361 | ZSWIM8 |
| ENSGALG00000000569 | 10192,548 | -0,483580181 | 0,1749621 | -2,763914 | 0,0057113 | 0,0223253 | SDC3 |
| ENSGALG00000009967 | 51604,136 | -0,483675694 | 0,1030754 | -4,692447 | 2,70E-06 | 3,17E-05 | LRPPRC |
| ENSGALG00000012702 | 17854,641 | -0,483830731 | 0,1194258 | -4,051309 | 5,09E-05 | 0,000418 | JARID2 |
| ENSGALG00000015830 | 24016,122 | -0,484314141 | 0,1030076 | -4,70173 | 2,58E-06 | 3,04E-05 | SYNCRIP |
| ENSGALG00000004887 | 17300,511 | -0,484420703 | 0,1068743 | -4,532622 | 5,83E-06 | 6,30E-05 | SART3 |
| ENSGALG00000002781 | 17608,172 | -0,484707648 | 0,1272601 | -3,808796 | 0,0001396 | 0,0010028 | BRCA1 |
| ENSGALG00000001950 | 14346,845 | -0,484721287 | 0,11288 | -4,294128 | 1,75E-05 | 0,0001671 | PMPCA |
| ENSGALG00000001777 | 2307,3102 | -0,484938734 | 0,1469529 | -3,299961 | 0,000967 | 0,0051935 | MFSD11 |
| ENSGALG00000002198 | 1337,3607 | -0,485078535 | 0,1489905 | -3,255768 | 0,0011309 | 0,0059241 | FHOD1 |
| ENSGALG00000004416 | 1683,2285 | -0,485858255 | 0,141996 | -3,421634 | 0,0006225 | 0,0035994 | TTC19 |
| ENSGALG00000012258 | 6299,7876 | -0,485912726 | 0,1153331 | -4,213123 | 2,52E-05 | 0,0002288 | GOLGA4 |
| ENSGALG00000013745 | 3956,926 | -0,486562498 | 0,1295758 | -3,755041 | 0,0001733 | 0,0012113 | RTTN |
| ENSGALG00000008862 | 12336,53 | -0,486712741 | 0,1063335 | -4,57723 | 4,71E-06 | 5,26E-05 | DNAJC10 |
| ENSGALG00000008670 | 2394,2538 | -0,486758564 | 0,1345509 | -3,617654 | 0,0002973 | 0,0019265 | C1orf52 |
| ENSGALG00000009157 | 10495,932 | -0,48696748 | 0,111678 | -4,36046 | 1,30E-05 | 0,0001295 | ECT2 |
| ENSGALG00000002906 | 23481,039 | -0,487686151 | 0,1087191 | -4,485745 | 7,27E-06 | 7,71E-05 | TARDBP |
| ENSGALG00000008155 | 684,45488 | -0,487898949 | 0,1882495 | -2,591767 | 0,0095484 | 0,0338282 | KCTD18 |
| ENSGALG00000005198 | 7882,4671 | -0,488262054 | 0,1163467 | -4,196612 | 2,71E-05 | 0,0002432 | RPS6KB1 |
| ENSGALG00000004750 | 7577,7666 | -0,488295007 | 0,1092829 | -4,468172 | 7,89E-06 | 8,32E-05 | NOL8 |
| ENSGALG00000010570 | 52092,719 | -0,488515596 | 0,1005153 | -4,860111 | 1,17E-06 | 1,51E-05 | BTF3L4 |
| ENSGALG00000016909 | 4324,6991 | -0,488526858 | 0,1162572 | -4,202121 | 2,64E-05 | 0,0002386 | NDFIP2 |
| ENSGALG00000004638 | 3370,7614 | -0,48870476 | 0,1414816 | -3,454194 | 0,0005519 | 0,0032587 | HEPH |
| ENSGALG00000001036 | 9684,7005 | -0,488799588 | 0,1105545 | -4,421345 | 9,81E-06 | 0,0001012 | ELAC2 |
| ENSGALG00000009981 | 1405,745 | -0,49003231 | 0,1465298 | -3,34425 | 0,0008251 | 0,004546 | PREPL |
| ENSGALG00000010638 | 13156,313 | -0,49033693 | 0,1208756 | -4,056541 | 4,98E-05 | 0,0004107 | CASP3 |
| ENSGALG00000006252 | 14647,749 | -0,49037706 | 0,1393987 | -3,517803 | 0,0004351 | 0,0026662 | LMO4 |
| ENSGALG00000000272 | 5267,8402 | -0,490419343 | 0,1354436 | -3,620838 | 0,0002936 | 0,0019059 | CDCA2 |
| ENSGALG00000008553 | 5356,6273 | -0,490452503 | 0,1346372 | -3,642771 | 0,0002697 | 0,0017672 | INO80 |
| ENSGALG00000010013 | 1321,1558 | -0,490665695 | 0,1680223 | -2,920241 | 0,0034976 | 0,0149466 | EDNRA |
| ENSGALG00000000367 | 793,50452 | -0,492073862 | 0,2010375 | -2,447672 | 0,0143783 | 0,0467543 | ASIC2 |
| ENSGALG00000016404 | 5806,1629 | -0,492286998 | 0,1192985 | -4,126516 | 3,68E-05 | 0,0003166 | MTBP |
| ENSGALG00000010335 | 1726,889 | -0,492481433 | 0,1437148 | -3,426798 | 0,0006107 | 0,0035505 | PIK3R3 |
| ENSGALG00000002141 | 14007,058 | -0,492600599 | 0,1062141 | -4,637807 | 3,52E-06 | 4,03E-05 | UGGT1 |
| ENSGALG00000008871 | 16063,59 | -0,492812674 | 0,1349508 | -3,651796 | 0,0002604 | 0,0017161 | LPHN2 |
| ENSGALG00000015059 | 7575,55 | -0,49315511 | 0,1131114 | -4,359906 | 1,30E-05 | 0,0001297 | |
| ENSGALG00000005652 | 5954,5779 | -0,493244478 | 0,113265 | -4,354781 | 1,33E-05 | 0,0001323 | DIDO1 |
| ENSGALG00000014218 | 876,2756 | -0,493463947 | 0,1862713 | -2,649168 | 0,008069 | 0,0296038 | PHF21B |
| ENSGALG00000023576 | 3299,2992 | -0,493564294 | 0,1207816 | -4,086419 | 4,38E-05 | 0,0003657 | PIM3 |
| ENSGALG00000008798 | 4574,7649 | -0,493864419 | 0,1182978 | -4,174757 | 2,98E-05 | 0,0002639 | ZMYND19 |
| ENSGALG00000020742 | 13114,145 | -0,493871262 | 0,1393963 | -3,54293 | 0,0003957 | 0,0024612 | PPRC1 |
| ENSGALG00000011962 | 7036,6211 | -0,494326456 | 0,1348846 | -3,664809 | 0,0002475 | 0,0016447 | NDNF |
| ENSGALG00000015478 | 5554,5683 | -0,494614804 | 0,1302844 | -3,796423 | 0,0001468 | 0,0010459 | USP45 |
| ENSGALG00000007326 | 7076,3591 | -0,494698301 | 0,1094474 | -4,519964 | 6,19E-06 | 6,66E-05 | PMM2 |
| ENSGALG00000005683 | 5447,0767 | -0,494902817 | 0,117901 | -4,197614 | 2,70E-05 | 0,0002424 | ARHGAP29 |
| ENSGALG00000010960 | 1065,9668 | -0,495730596 | 0,183425 | -2,702634 | 0,0068792 | 0,0260314 | GALNT3 |
| ENSGALG00000000795 | 1504,9365 | -0,495778483 | 0,1425043 | -3,479042 | 0,0005032 | 0,0030142 | MFSD12 |
| ENSGALG00000004273 | 11441,37 | -0,495936282 | 0,1511882 | -3,280258 | 0,0010371 | 0,005505 | RALGPS2 |
| ENSGALG00000014998 | 1987,961 | -0,495971981 | 0,1542046 | -3,216325 | 0,0012984 | 0,0066435 | CABLES1 |
| ENSGALG00000015684 | 4408,6888 | -0,496145153 | 0,1139834 | -4,352783 | 1,34E-05 | 0,0001334 | DNAJC25 |
| ENSGALG00000004570 | 5026,7973 | -0,496270556 | 0,1351131 | -3,673 | 0,0002397 | 0,001596 | NCOA3 |
| ENSGALG00000013257 | 123007,22 | -0,496798558 | 0,1026267 | -4,840832 | 1,29E-06 | 1,64E-05 | LDHB |

| | | | | | | | |
|---|---|---|---|---|---|---|---|
| ENSGALG00000027635 | 9027,1813 | -0,49702612 | 0,1189634 | -4,177976 | 2,94E-05 | 0,0002609 | NME4 |
| ENSGALG00000005583 | 1029,2928 | -0,497171251 | 0,1674424 | -2,969208 | 0,0029857 | 0,0131129 | ALG14 |
| ENSGALG00000003596 | 25796,912 | -0,497856616 | 0,162082 | -3,071634 | 0,0021289 | 0,0099471 | TRRAP |
| ENSGALG00000010624 | 9648,3759 | -0,498764877 | 0,1086145 | -4,592066 | 4,39E-06 | 4,91E-05 | MEMO1 |
| ENSGALG00000026865 | 1245,9917 | -0,49909443 | 0,1619603 | -3,081585 | 0,002059 | 0,0096734 | XRCC2 |
| ENSGALG00000007691 | 3206,9135 | -0,499115113 | 0,1294529 | -3,855572 | 0,0001155 | 0,0008465 | ARMC9 |
| ENSGALG00000007155 | 2000,5013 | -0,499370494 | 0,1491539 | -3,348022 | 0,0008139 | 0,0044973 | RMI2 |
| ENSGALG00000003874 | 8943,2552 | -0,499635075 | 0,1097884 | -4,550893 | 5,34E-06 | 5,87E-05 | HEATR3 |
| ENSGALG00000005695 | 3266,3031 | -0,500082186 | 0,1314863 | -3,803302 | 0,0001428 | 0,0010204 | CHEK2 |
| ENSGALG00000025792 | 3188,1064 | -0,500292681 | 0,1218494 | -4,105829 | 4,03E-05 | 0,0003426 | FRRS1L |
| ENSGALG00000011264 | 1925,3077 | -0,500518992 | 0,1479576 | -3,382854 | 0,0007174 | 0,0040628 | SULF1 |
| ENSGALG00000016758 | 1465,7896 | -0,5005603 | 0,1465611 | -3,415369 | 0,000637 | 0,0036653 | LIPT1 |
| ENSGALG00000010716 | 5501,7296 | -0,50085931 | 0,1188557 | -4,214011 | 2,51E-05 | 0,0002282 | AGA |
| ENSGALG00000009269 | 3597,5456 | -0,501724842 | 0,1302073 | -3,853278 | 0,0001165 | 0,000854 | RP5-1021I20.4 |
| ENSGALG00000011857 | 24467,231 | -0,502240381 | 0,105172 | -4,775418 | 1,79E-06 | 2,19E-05 | LUC7L2 |
| ENSGALG00000003719 | 6864,0435 | -0,502933526 | 0,135236 | -3,718932 | 0,0002001 | 0,0013633 | CHD9 |
| ENSGALG00000024035 | 1950,9384 | -0,503656541 | 0,1918019 | -2,625921 | 0,0086415 | 0,0312743 | EPS8L2 |
| ENSGALG00000006524 | 12378,285 | -0,503792354 | 0,1077015 | -4,677671 | 2,90E-06 | 3,38E-05 | NOC3L |
| ENSGALG00000016781 | 17546,507 | -0,504088725 | 0,1248435 | -4,037765 | 5,40E-05 | 0,0004399 | MAP4K4 |
| ENSGALG00000014922 | 15076,997 | -0,504425665 | 0,118689 | -4,249979 | 2,14E-05 | 0,0001986 | ROCK1 |
| ENSGALG00000004690 | 52340,48 | -0,504461763 | 0,1007635 | -5,006396 | 5,55E-07 | 7,63E-06 | MAPK6 |
| ENSGALG00000002728 | 6868,7758 | -0,504726866 | 0,1233642 | -4,091357 | 4,29E-05 | 0,0003598 | SLC16A3 |
| ENSGALG00000011877 | 4219,7935 | -0,504776099 | 0,1219876 | -4,137928 | 3,50E-05 | 0,0003041 | LARS2 |
| ENSGALG00000005533 | 4875,7726 | -0,504976403 | 0,1222357 | -4,131171 | 3,61E-05 | 0,0003116 | ZMYM3 |
| ENSGALG00000013209 | 5984,6391 | -0,505418476 | 0,1115613 | -4,530409 | 5,89E-06 | 6,36E-05 | SLC39A6 |
| ENSGALG00000015252 | 4947,9442 | -0,505578034 | 0,1423502 | -3,551649 | 0,0003828 | 0,0023887 | ST3GAL6 |
| ENSGALG00000007234 | 4564,3675 | -0,505755971 | 0,117953 | -4,287774 | 1,80E-05 | 0,000171 | CLCN5 |
| ENSGALG00000010034 | 2578,0952 | -0,50624459 | 0,1322184 | -3,828852 | 0,0001287 | 0,0009324 | |
| ENSGALG00000009556 | 2369,7227 | -0,506316289 | 0,2035819 | -2,487039 | 0,0128811 | 0,0429423 | PRICKLE1 |
| ENSGALG00000003540 | 2816,5136 | -0,506324474 | 0,1426366 | -3,549751 | 0,0003856 | 0,0024027 | CEP95 |
| ENSGALG00000005992 | 1391,4533 | -0,506625581 | 0,1532979 | -3,304844 | 0,0009503 | 0,0051298 | PDE8A |
| ENSGALG00000000271 | 3107,1088 | -0,506687982 | 0,1663878 | -3,045222 | 0,0023251 | 0,0106766 | C2CD2L |
| ENSGALG00000006935 | 12264,183 | -0,506724591 | 0,1094246 | -4,630812 | 3,64E-06 | 4,15E-05 | BTAF1 |
| ENSGALG00000019867 | 6166,9755 | -0,506841982 | 0,1119268 | -4,528335 | 5,95E-06 | 6,41E-05 | WDCP |
| ENSGALG00000008420 | 3322,248 | -0,507225725 | 0,1221905 | -4,151106 | 3,31E-05 | 0,0002888 | CAND2 |
| ENSGALG00000012020 | 2468,0174 | -0,507245855 | 0,184003 | -2,756726 | 0,0058383 | 0,0227414 | KIAA1549L |
| ENSGALG00000009782 | 45976,854 | -0,507579083 | 0,1062698 | -4,776324 | 1,79E-06 | 2,18E-05 | NAA15 |
| ENSGALG00000004118 | 1623,6627 | -0,508229667 | 0,1686061 | -3,014301 | 0,0025757 | 0,0115852 | |
| ENSGALG00000002424 | 2020,6569 | -0,508419582 | 0,160293 | -3,171813 | 0,0015149 | 0,0074947 | RECQL5 |
| ENSGALG00000002652 | 8163,3179 | -0,509052755 | 0,1365252 | -3,728635 | 0,0001925 | 0,0013231 | FZD10 |
| ENSGALG00000005509 | 6460,5427 | -0,509428544 | 0,1427708 | -3,568157 | 0,0003595 | 0,0022671 | DPYD |
| ENSGALG00000004990 | 1688,4348 | -0,509590179 | 0,1471207 | -3,463756 | 0,0005327 | 0,0031615 | C10orf11 |
| ENSGALG00000002505 | 458,85789 | -0,509844408 | 0,2099041 | -2,42894 | 0,015143 | 0,0487175 | |
| ENSGALG00000010735 | 16013,937 | -0,510060882 | 0,1046006 | -4,876271 | 1,08E-06 | 1,40E-05 | NDC1 |
| ENSGALG00000003628 | 14552,598 | -0,510073233 | 0,1414317 | -3,6065 | 0,0003104 | 0,0019979 | TLN2 |
| ENSGALG00000015787 | 3935,8112 | -0,510253054 | 0,1410404 | -3,617779 | 0,0002971 | 0,0019265 | RNGTT |
| ENSGALG00000008778 | 6210,2201 | -0,51039033 | 0,1491907 | -3,421059 | 0,0006238 | 0,0036047 | SPRED2 |
| ENSGALG00000012135 | 48183,581 | -0,511735314 | 0,1079748 | -4,739395 | 2,14E-06 | 2,56E-05 | ATF4 |
| ENSGALG00000017085 | 4116,9494 | -0,511832688 | 0,125074 | -4,092239 | 4,27E-05 | 0,0003593 | SLC7A1 |
| ENSGALG00000002709 | 2938,4248 | -0,511853662 | 0,1402453 | -3,649703 | 0,0002625 | 0,0017285 | MYO1C |
| ENSGALG00000004621 | 8629,8848 | -0,512109985 | 0,1230296 | -4,162493 | 3,15E-05 | 0,0002769 | PREX1 |
| ENSGALG00000005278 | 9525,2437 | -0,512823817 | 0,1583471 | -3,238605 | 0,0012012 | 0,0062229 | CDH11 |
| ENSGALG00000014791 | 22071,934 | -0,513191503 | 0,1237594 | -4,146686 | 3,37E-05 | 0,0002936 | WDR3 |
| ENSGALG00000009825 | 16041,569 | -0,51401966 | 0,1105113 | -4,651284 | 3,30E-06 | 3,80E-05 | SRGAP1 |
| ENSGALG00000001441 | 10679 | -0,514329303 | 0,1052351 | -4,88743 | 1,02E-06 | 1,33E-05 | UBE2G1 |
| ENSGALG00000026468 | 7333,857 | -0,514962132 | 0,1096475 | -4,696525 | 2,65E-06 | 3,11E-05 | CHSY1 |
| ENSGALG00000002646 | 11173,638 | -0,515104787 | 0,1477459 | -3,486422 | 0,0004895 | 0,0029478 | FAM171B |
| ENSGALG00000005653 | 3018,7755 | -0,515161656 | 0,1231259 | -4,184022 | 2,86E-05 | 0,0002552 | NFKB2 |
| ENSGALG00000011707 | 2975,9589 | -0,516018775 | 0,1353146 | -3,813473 | 0,000137 | 0,0009866 | UBA5 |
| ENSGALG00000003566 | 2456,1483 | -0,516245691 | 0,1356954 | -3,804444 | 0,0001421 | 0,0010168 | SMURF2 |
| ENSGALG00000003047 | 12574,653 | -0,516252465 | 0,132068 | -3,908991 | 9,27E-05 | 0,0007038 | SUGP2 |
| ENSGALG00000009784 | 1512,522 | -0,516604655 | 0,1915407 | -2,697101 | 0,0069946 | 0,0263872 | PTPN14 |
| ENSGALG00000006830 | 714,61116 | -0,51666647 | 0,1768872 | -2,920882 | 0,0034904 | 0,0149206 | CHID1 |
| ENSGALG00000007527 | 4564,3635 | -0,516973735 | 0,1183481 | -4,368248 | 1,25E-05 | 0,0001255 | ADARB1 |

| | | | | | | | |
|---|---|---|---|---|---|---|---|
| ENSGALG00000012550 | 2418,3256 | -0,517745286 | 0,1390423 | -3,723653 | 0,0001964 | 0,0013434 | HMOX1 |
| ENSGALG00000009207 | 1636,7589 | -0,518081559 | 0,1511771 | -3,426985 | 0,0006103 | 0,0035496 | TRIM2 |
| ENSGALG00000012207 | 4193,9067 | -0,518404651 | 0,1155257 | -4,487352 | 7,21E-06 | 7,66E-05 | CCNT2 |
| ENSGALG00000009451 | 10542,681 | -0,518444547 | 0,1054026 | -4,918707 | 8,71E-07 | 1,14E-05 | PPID |
| ENSGALG00000010189 | 6061,8235 | -0,518843565 | 0,1108188 | -4,681909 | 2,84E-06 | 3,32E-05 | RP11-407N17.3 |
| ENSGALG00000011596 | 8210,1595 | -0,519655791 | 0,1247256 | -4,166392 | 3,09E-05 | 0,0002731 | PTPN4 |
| ENSGALG00000006622 | 2389,7433 | -0,520113735 | 0,1331747 | -3,9055 | 9,40E-05 | 0,0007116 | FRA10AC1 |
| ENSGALG00000014784 | 4045,4061 | -0,520975246 | 0,1405154 | -3,707602 | 0,0002092 | 0,0014201 | MAST4 |
| ENSGALG00000009992 | 11680,473 | -0,521017168 | 0,1135606 | -4,588008 | 4,47E-06 | 5,01E-05 | ARHGAP5 |
| ENSGALG00000015889 | 13285,71 | -0,52134607 | 0,1484689 | -3,511482 | 0,0004456 | 0,0027219 | PHIP |
| ENSGALG00000007752 | 6790,4614 | -0,521363763 | 0,1083691 | -4,811 | 1,50E-06 | 1,88E-05 | COPS7B |
| ENSGALG00000005850 | 15672,107 | -0,521743975 | 0,1094822 | -4,765559 | 1,88E-06 | 2,29E-05 | FNBP1L |
| ENSGALG00000012354 | 2395,6585 | -0,521764337 | 0,1590822 | -3,27984 | 0,0010387 | 0,005511 | VOPP1 |
| ENSGALG00000015314 | 12934,444 | -0,521997148 | 0,1092952 | -4,77603 | 1,79E-06 | 2,19E-05 | BEND3 |
| ENSGALG00000001109 | 1011,8383 | -0,522206198 | 0,1701303 | -3,069449 | 0,0021445 | 0,0100027 | GTSF1 |
| ENSGALG00000004397 | 3452,7405 | -0,523035736 | 0,1275508 | -4,100607 | 4,12E-05 | 0,0003498 | MLXIP |
| ENSGALG00000012592 | 3142,2738 | -0,523397807 | 0,1406662 | -3,720849 | 0,0001986 | 0,0013557 | RMI1 |
| ENSGALG00000006811 | 8467,8095 | -0,523451296 | 0,1844356 | -2,838125 | 0,0045379 | 0,0184744 | ZIC1 |
| ENSGALG00000002487 | 10163,363 | -0,523537895 | 0,1313011 | -3,987308 | 6,68E-05 | 0,0005281 | SFSWAP |
| ENSGALG00000003260 | 9438,4125 | -0,523953501 | 0,1105482 | -4,739594 | 2,14E-06 | 2,56E-05 | UTP6 |
| ENSGALG00000012720 | 24474,404 | -0,524679114 | 0,1115414 | -4,703897 | 2,55E-06 | 3,01E-05 | NUP153 |
| ENSGALG00000005425 | 38874,994 | -0,524696803 | 0,1183507 | -4,433407 | 9,28E-06 | 9,63E-05 | OGT |
| ENSGALG00000010861 | 2392,2986 | -0,524736464 | 0,1328216 | -3,950687 | 7,79E-05 | 0,0006035 | OMA1 |
| ENSGALG00000004489 | 1483,1995 | -0,524803254 | 0,1455768 | -3,604991 | 0,0003122 | 0,0020076 | |
| ENSGALG00000011770 | 14710,919 | -0,525890593 | 0,1188458 | -4,424984 | 9,64E-06 | 9,98E-05 | UMPS |
| ENSGALG00000002850 | 6570,4074 | -0,526275555 | 0,1347444 | -3,905733 | 9,39E-05 | 0,0007113 | RASA2 |
| ENSGALG00000023562 | 1492,3921 | -0,526424205 | 0,1608921 | -3,271908 | 0,0010682 | 0,0056493 | TEX9 |
| ENSGALG00000015836 | 3297,8542 | -0,526441345 | 0,132466 | -3,974163 | 7,06E-05 | 0,0005557 | CEP162 |
| ENSGALG00000014426 | 14003,05 | -0,526522361 | 0,1207296 | -4,361172 | 1,29E-05 | 0,0001291 | KPNA1 |
| ENSGALG00000009172 | 566,03742 | -0,527120334 | 0,2081817 | -2,532021 | 0,0113407 | 0,0388242 | OSBPL6 |
| ENSGALG00000011982 | 5512,8717 | -0,527666168 | 0,1403594 | -3,759392 | 0,0001703 | 0,0011941 | PDE5A |
| ENSGALG00000011149 | 879,0742 | -0,527681437 | 0,1626198 | -3,244879 | 0,001175 | 0,0061037 | PLA2R1 |
| ENSGALG00000008096 | 6070,9895 | -0,528190931 | 0,1247684 | -4,233371 | 2,30E-05 | 0,0002114 | SUFU |
| ENSGALG00000007140 | 3712,2899 | -0,528407251 | 0,1869729 | -2,826116 | 0,0047116 | 0,0190805 | NRP1 |
| ENSGALG00000008847 | 3493,1884 | -0,528756478 | 0,1263169 | -4,185952 | 2,84E-05 | 0,0002534 | VPS54 |
| ENSGALG00000013177 | 2545,0183 | -0,528877548 | 0,1412318 | -3,744749 | 0,0001806 | 0,0012537 | BCAT1 |
| ENSGALG00000017392 | 1036,5696 | -0,528892969 | 0,1585879 | -3,335014 | 0,000853 | 0,0046807 | SPDYA |
| ENSGALG00000012979 | 10526,261 | -0,528975634 | 0,1480989 | -3,571772 | 0,0003546 | 0,0022422 | TRIO |
| ENSGALG00000000208 | 3421,1766 | -0,529383362 | 0,1352035 | -3,915457 | 9,02E-05 | 0,0006871 | MCC |
| ENSGALG00000004474 | 1133,2959 | -0,529777367 | 0,1534298 | -3,452897 | 0,0005546 | 0,0032709 | PLEKHF1 |
| ENSGALG00000013203 | 7478,1862 | -0,52983334 | 0,1163226 | -4,554862 | 5,24E-06 | 5,78E-05 | ELP2 |
| ENSGALG00000011419 | 3053,0195 | -0,529859697 | 0,123662 | -4,284741 | 1,83E-05 | 0,0001731 | DUSP4 |
| ENSGALG00000010159 | 4422,5959 | -0,529906457 | 0,122307 | -4,332595 | 1,47E-05 | 0,0001449 | SLC20A2 |
| ENSGALG00000002974 | 1015,7762 | -0,529999669 | 0,168327 | -3,148631 | 0,0016404 | 0,0080164 | SPHKAP |
| ENSGALG00000002624 | 31172,832 | -0,530323004 | 0,1123984 | -4,718243 | 2,38E-06 | 2,82E-05 | BMS1 |
| ENSGALG00000006826 | 4092,1467 | -0,530661584 | 0,1454636 | -3,648071 | 0,0002642 | 0,0017378 | KLHL25 |
| ENSGALG00000002253 | 1425,4871 | -0,530784268 | 0,1456031 | -3,64542 | 0,000267 | 0,0017508 | HEMK1 |
| ENSGALG00000011569 | 5815,2909 | -0,530885029 | 0,12043 | -4,408246 | 1,04E-05 | 0,000107 | SCYL2 |
| ENSGALG00000003944 | 1246,484 | -0,531232705 | 0,1600895 | -3,318348 | 0,0009055 | 0,0049275 | FAM189A1 |
| ENSGALG00000004347 | 6495,1605 | -0,533321296 | 0,1103628 | -4,832435 | 1,35E-06 | 1,70E-05 | NEDD4 |
| ENSGALG00000006710 | 714,0564 | -0,533378082 | 0,1783936 | -2,989895 | 0,0027907 | 0,0124332 | RAB36 |
| ENSGALG00000000891 | 3716,0687 | -0,533422492 | 0,1230689 | -4,33434 | 1,46E-05 | 0,000144 | NSUN5 |
| ENSGALG00000015818 | 5198,5394 | -0,53353462 | 0,1266077 | -4,214076 | 2,51E-05 | 0,0002282 | E2F5 |
| ENSGALG00000015448 | 2486,2853 | -0,533672096 | 0,1639578 | -3,254936 | 0,0011342 | 0,0059369 | TOX |
| ENSGALG00000008312 | 1844,0991 | -0,534378374 | 0,1384766 | -3,858978 | 0,0001139 | 0,0008371 | GSAP |
| ENSGALG00000004506 | 11897,642 | -0,534405871 | 0,1101239 | -4,852769 | 1,22E-06 | 1,55E-05 | URI1 |
| ENSGALG00000013085 | 2436,8268 | -0,535478159 | 0,1366821 | -3,917689 | 8,94E-05 | 0,0006824 | PTPRO |
| ENSGALG00000008098 | 5171,8512 | -0,536763264 | 0,1152605 | -4,656959 | 3,21E-06 | 3,70E-05 | NAMPT |
| ENSGALG00000016024 | 730,05733 | -0,537143611 | 0,1775788 | -3,024818 | 0,0024878 | 0,0112763 | SETD4 |
| ENSGALG00000006856 | 8733,4208 | -0,537151634 | 0,1162633 | -4,62013 | 3,83E-06 | 4,35E-05 | CD151 |
| ENSGALG00000011024 | 13428,049 | -0,537183599 | 0,1048649 | -5,122625 | 3,01E-07 | 4,37E-06 | CACHD1 |
| ENSGALG00000010564 | 3014,7883 | -0,537373895 | 0,1233997 | -4,354741 | 1,33E-05 | 0,0001323 | TBCK |
| ENSGALG00000015360 | 9022,7064 | -0,537653684 | 0,1055635 | -5,093178 | 3,52E-07 | 5,05E-06 | KIAA1524 |
| ENSGALG00000026253 | 1652,9426 | -0,5379573 | 0,1506081 | -3,571902 | 0,0003544 | 0,0022422 | TMEM144 |

| | | | | | | | |
|---|---|---|---|---|---|---|---|
| ENSGALG00000003705 | 3639,3367 | -0,538250704 | 0,1231851 | -4,369445 | 1,25E-05 | 0,0001252 | VPS13C |
| ENSGALG00000012468 | 13779,95 | -0,538390713 | 0,1366618 | -3,939585 | 8,16E-05 | 0,0006293 | PRPF39 |
| ENSGALG00000016440 | 2000,29 | -0,538612864 | 0,1606177 | -3,353384 | 0,0007983 | 0,0044417 | KLF11 |
| ENSGALG00000016835 | 6008,9103 | -0,539714594 | 0,1468334 | -3,675694 | 0,0002372 | 0,0015823 | ATP11A |
| ENSGALG00000009476 | 2384,795 | -0,539812489 | 0,1852255 | -2,914353 | 0,0035643 | 0,0152027 | CDK6 |
| ENSGALG00000002873 | 15190,766 | -0,539979366 | 0,10839 | -4,981821 | 6,30E-07 | 8,52E-06 | RSL1D1 |
| ENSGALG00000015701 | 9337,5788 | -0,540112853 | 0,1299709 | -4,155644 | 3,24E-05 | 0,0002838 | CXADR |
| ENSGALG00000015180 | 15097,101 | -0,540189844 | 0,1129211 | -4,78378 | 1,72E-06 | 2,11E-05 | PSAT1 |
| ENSGALG00000001389 | 3052,2811 | -0,540945617 | 0,1260291 | -4,292227 | 1,77E-05 | 0,0001682 | TTLL11 |
| ENSGALG00000029022 | 1420,5998 | -0,540984923 | 0,2146447 | -2,520374 | 0,011723 | 0,0398623 | NPAS3 |
| ENSGALG00000015491 | 4074,1339 | -0,541085652 | 0,1219675 | -4,436312 | 9,15E-06 | 9,53E-05 | GPR89B |
| ENSGALG00000012121 | 12053,922 | -0,541445831 | 0,1045431 | -5,179165 | 2,23E-07 | 3,34E-06 | RCN1 |
| ENSGALG00000023089 | 788,71726 | -0,541614618 | 0,1982755 | -2,731627 | 0,0063022 | 0,0242593 | SSBP2 |
| ENSGALG00000013438 | 21891,879 | -0,541667727 | 0,1120475 | -4,834271 | 1,34E-06 | 1,69E-05 | TEX10 |
| ENSGALG00000001957 | 8870,632 | -0,541703645 | 0,1341035 | -4,039444 | 5,36E-05 | 0,0004373 | ITCH |
| ENSGALG00000016460 | 6190,6626 | -0,541890883 | 0,14842 | -3,651065 | 0,0002612 | 0,0017202 | NBAS |
| ENSGALG00000014573 | 17671,905 | -0,541989088 | 0,1103445 | -4,911792 | 9,02E-07 | 1,18E-05 | NAT10 |
| ENSGALG00000025884 | 6669,8373 | -0,543082298 | 0,1134242 | -4,788063 | 1,68E-06 | 2,07E-05 | NABP1 |
| ENSGALG00000007835 | 1738,059 | -0,543517994 | 0,1483373 | -3,664068 | 0,0002482 | 0,0016487 | PUS10 |
| ENSGALG00000006895 | 13372,527 | -0,543915173 | 0,1058888 | -5,136663 | 2,80E-07 | 4,09E-06 | CEP131 |
| ENSGALG00000004364 | 7914,9764 | -0,544307466 | 0,1156323 | -4,707225 | 2,51E-06 | 2,97E-05 | ABR |
| ENSGALG00000014769 | 613,98427 | -0,544618003 | 0,20267 | -2,687216 | 0,007205 | 0,0270761 | NLN |
| ENSGALG00000026452 | 7763,7201 | -0,545277299 | 0,1319298 | -4,133086 | 3,58E-05 | 0,0003098 | MURC |
| ENSGALG00000008623 | 615,02509 | -0,545289509 | 0,1967721 | -2,771173 | 0,0055855 | 0,021976 | TOR4A |
| ENSGALG00000012223 | 1554,6199 | -0,545359079 | 0,1399232 | -3,89756 | 9,72E-05 | 0,0007325 | ZRANB3 |
| ENSGALG00000004879 | 617,17052 | -0,545838008 | 0,1927673 | -2,831591 | 0,0046317 | 0,0187848 | SLC6A11 |
| ENSGALG00000004744 | 525,6252 | -0,546332008 | 0,1957015 | -2,791659 | 0,0052439 | 0,0208575 | DISP2 |
| ENSGALG00000003440 | 1101,0238 | -0,546469024 | 0,1603017 | -3,409004 | 0,000652 | 0,0037377 | |
| ENSGALG00000011351 | 157820,47 | -0,547374715 | 0,1580098 | -3,464182 | 0,0005318 | 0,0031579 | HSP90AA1 |
| ENSGALG00000016768 | 7527,2287 | -0,54750013 | 0,11987 | -4,56745 | 4,94E-06 | 5,48E-05 | LONRF2 |
| ENSGALG00000012894 | 568,24499 | -0,54779071 | 0,2155009 | -2,541941 | 0,0110239 | 0,0380183 | CHRM2 |
| ENSGALG00000009503 | 5606,8039 | -0,547842813 | 0,1186519 | -4,617227 | 3,89E-06 | 4,41E-05 | RAB3GAP2 |
| ENSGALG00000015746 | 3303,5119 | -0,547863301 | 0,1403611 | -3,903241 | 9,49E-05 | 0,0007163 | JAM2 |
| ENSGALG00000003937 | 13547,502 | -0,548529686 | 0,130015 | -4,218972 | 2,45E-05 | 0,0002241 | GEMIN5 |
| ENSGALG00000005727 | 12274,551 | -0,548590838 | 0,1154072 | -4,753525 | 2,00E-06 | 2,41E-05 | SLC7A3 |
| ENSGALG00000017083 | 1252,4876 | -0,54907192 | 0,1596795 | -3,438588 | 0,0005848 | 0,003414 | KATNAL1 |
| ENSGALG00000006971 | 10409,679 | -0,549657911 | 0,1186418 | -4,63292 | 3,61E-06 | 4,12E-05 | CDK5RAP2 |
| ENSGALG00000003026 | 1987,5595 | -0,550107166 | 0,1687281 | -3,260317 | 0,0011129 | 0,0058389 | MKL2 |
| ENSGALG00000029127 | 622,20536 | -0,550279038 | 0,2169523 | -2,536406 | 0,0111997 | 0,0384768 | |
| ENSGALG00000011400 | 12123,593 | -0,550294288 | 0,103122 | -5,336343 | 9,48E-08 | 1,53E-06 | USP44 |
| ENSGALG00000028501 | 7019,464 | -0,550742687 | 0,1179763 | -4,668248 | 3,04E-06 | 3,52E-05 | DCK |
| ENSGALG00000013503 | 838,79916 | -0,551514976 | 0,1874586 | -2,942063 | 0,0032603 | 0,0140874 | FAM149A |
| ENSGALG00000007097 | 2035,7636 | -0,551520351 | 0,1696313 | -3,25129 | 0,0011488 | 0,0059997 | RPS6KA6 |
| ENSGALG00000017006 | 4626,7368 | -0,551986159 | 0,1179626 | -4,679334 | 2,88E-06 | 3,35E-05 | SETDB2 |
| ENSGALG00000004071 | 3249,4429 | -0,552437495 | 0,123172 | -4,485092 | 7,29E-06 | 7,73E-05 | NETO2 |
| ENSGALG00000000684 | 3181,0431 | -0,552570682 | 0,1435454 | -3,849448 | 0,0001184 | 0,0008642 | KLHDC8A |
| ENSGALG00000003835 | 27823,208 | -0,552761335 | 0,1061578 | -5,20698 | 1,92E-07 | 2,92E-06 | ATP2A2 |
| ENSGALG00000011052 | 797,4232 | -0,553217179 | 0,2034892 | -2,718657 | 0,0065548 | 0,0250612 | SLC38A11 |
| ENSGALG00000010309 | 346,20571 | -0,554430553 | 0,2211705 | -2,506801 | 0,0121829 | 0,0410672 | ZC2HC1C |
| ENSGALG00000016883 | 3231,0954 | -0,555002491 | 0,1279098 | -4,339014 | 1,43E-05 | 0,0001413 | DOCK9 |
| ENSGALG00000004813 | 26912,189 | -0,555221474 | 0,117319 | -4,732578 | 2,22E-06 | 2,65E-05 | IARS |
| ENSGALG00000005774 | 2418,9054 | -0,555555373 | 0,133361 | -4,165801 | 3,10E-05 | 0,0002736 | NKAIN4 |
| ENSGALG00000013635 | 10619,161 | -0,555662229 | 0,1221599 | -4,548648 | 5,40E-06 | 5,91E-05 | SCAF8 |
| ENSGALG00000016468 | 6793,0911 | -0,555855719 | 0,108947 | -5,102077 | 3,36E-07 | 4,84E-06 | GEN1 |
| ENSGALG00000026379 | 480,26073 | -0,557143449 | 0,1981326 | -2,811973 | 0,0049239 | 0,0198356 | OPRD1 |
| ENSGALG00000013430 | 2775,6809 | -0,557212283 | 0,1237629 | -4,502256 | 6,72E-06 | 7,16E-05 | |
| ENSGALG00000021487 | 6790,2413 | -0,55890103 | 0,1279072 | -4,369583 | 1,24E-05 | 0,0001252 | |
| ENSGALG00000016231 | 123947,5 | -0,55907395 | 0,1098338 | -5,09018 | 3,58E-07 | 5,11E-06 | DDX3X |
| ENSGALG00000019178 | 3990,1334 | -0,559313492 | 0,1249859 | -4,475011 | 7,64E-06 | 8,08E-05 | PRDM15 |
| ENSGALG00000010022 | 5530,328 | -0,559381952 | 0,1189802 | -4,701473 | 2,58E-06 | 3,04E-05 | PRMT9 |
| ENSGALG00000011692 | 9047,4774 | -0,559417827 | 0,1160828 | -4,819127 | 1,44E-06 | 1,81E-05 | CDCA4 |
| ENSGALG00000017021 | 332,60369 | -0,55959631 | 0,2270786 | -2,464329 | 0,013727 | 0,045078 | ATP7B |
| ENSGALG00000000447 | 760,9204 | -0,559669779 | 0,171256 | -3,268029 | 0,001083 | 0,0057107 | KCNA3 |
| ENSGALG00000011541 | 6975,1988 | -0,559795067 | 0,1404905 | -3,984576 | 6,76E-05 | 0,0005331 | BIN1 |

| | | | | | | | |
|---|---|---|---|---|---|---|---|
| ENSGALG00000003510 | 5135,2375 | -0,562160497 | 0,117815 | -4,771553 | 1,83E-06 | 2,23E-05 | UCK2 |
| ENSGALG00000012508 | 4886,2726 | -0,562203872 | 0,114843 | -4,895414 | 9,81E-07 | 1,28E-05 | TRIP13 |
| ENSGALG00000027231 | 9354,1487 | -0,562988244 | 0,1537161 | -3,66252 | 0,0002497 | 0,0016558 | NRARP |
| ENSGALG00000016897 | 2920,7661 | -0,563110321 | 0,1234746 | -4,560535 | 5,10E-06 | 5,64E-05 | GPR180 |
| ENSGALG00000010081 | 2535,9184 | -0,563235953 | 0,1656002 | -3,401179 | 0,000671 | 0,0038367 | SH3D19 |
| ENSGALG00000003074 | 3831,5986 | -0,563796907 | 0,1286586 | -4,382116 | 1,18E-05 | 0,0001188 | HOMER3 |
| ENSGALG00000012256 | 15406,505 | -0,563809375 | 0,163288 | -3,452852 | 0,0005547 | 0,0032709 | UTRN |
| ENSGALG00000026438 | 2075,6069 | -0,563826313 | 0,1305634 | -4,318409 | 1,57E-05 | 0,0001522 | |
| ENSGALG00000008132 | 7657,4934 | -0,564518166 | 0,1163041 | -4,853812 | 1,21E-06 | 1,55E-05 | PUS7 |
| ENSGALG00000013641 | 530,0064 | -0,565215365 | 0,2025498 | -2,790501 | 0,0052627 | 0,0209261 | MTMR7 |
| ENSGALG00000026583 | 481,9172 | -0,565402739 | 0,219019 | -2,581523 | 0,0098365 | 0,0347405 | |
| ENSGALG00000022808 | 24156,793 | -0,565497585 | 0,1052406 | -5,373379 | 7,73E-08 | 1,27E-06 | RRP1B |
| ENSGALG00000011228 | 7018,8875 | -0,565699287 | 0,1248478 | -4,53111 | 5,87E-06 | 6,35E-05 | WARS |
| ENSGALG00000002623 | 550,49256 | -0,566538314 | 0,1922408 | -2,947024 | 0,0032085 | 0,0139164 | PCOLCE2 |
| ENSGALG00000014940 | 2607,3053 | -0,566659105 | 0,1584762 | -3,575674 | 0,0003493 | 0,0022157 | ZUFSP |
| ENSGALG00000016917 | 1018,5023 | -0,567495374 | 0,1648915 | -3,441629 | 0,0005782 | 0,0033831 | CLN5 |
| ENSGALG00000015431 | 2610,4581 | -0,567537408 | 0,1279963 | -4,434014 | 9,25E-06 | 9,61E-05 | UBXN2B |
| ENSGALG00000009855 | 7273,6535 | -0,567662024 | 0,1223057 | -4,641336 | 3,46E-06 | 3,97E-05 | TBC1D9 |
| ENSGALG00000017079 | 12990,655 | -0,56791639 | 0,124822 | -4,549812 | 5,37E-06 | 5,89E-05 | USPL1 |
| ENSGALG00000000195 | 1824,66 | -0,568283448 | 0,1600789 | -3,55002 | 0,0003852 | 0,0024013 | SLC23A2 |
| ENSGALG00000003906 | 12475,038 | -0,568387401 | 0,109475 | -5,191937 | 2,08E-07 | 3,14E-06 | MPHOSPH10 |
| ENSGALG00000016281 | 11421,229 | -0,569360911 | 0,1081146 | -5,266272 | 1,39E-07 | 2,17E-06 | DMD |
| ENSGALG00000016391 | 2197,6068 | -0,569812042 | 0,1354173 | -4,207823 | 2,58E-05 | 0,0002336 | CNKSR2 |
| ENSGALG00000015142 | 3202,1058 | -0,570285644 | 0,1334226 | -4,27428 | 1,92E-05 | 0,0001807 | DSG2 |
| ENSGALG00000009714 | 1150,5594 | -0,570305895 | 0,1555825 | -3,665616 | 0,0002467 | 0,0016411 | CBR4 |
| ENSGALG00000014783 | 10636,1 | -0,572122303 | 0,1261051 | -4,53687 | 5,71E-06 | 6,20E-05 | PTPRK |
| ENSGALG00000016910 | 23108,199 | -0,572230903 | 0,1116391 | -5,125723 | 2,96E-07 | 4,30E-06 | RBM26 |
| ENSGALG00000012849 | 5113,4695 | -0,57279907 | 0,1488538 | -3,848064 | 0,0001191 | 0,0008677 | SLC37A3 |
| ENSGALG00000001363 | 21664,352 | -0,573517858 | 0,1098878 | -5,219123 | 1,80E-07 | 2,75E-06 | MYBBP1A |
| ENSGALG00000003180 | 15412,658 | -0,573999724 | 0,1081856 | -5,305695 | 1,12E-07 | 1,78E-06 | TSR1 |
| ENSGALG00000013031 | 1094,9404 | -0,574014118 | 0,173605 | -3,306438 | 0,0009449 | 0,0051051 | CECR1 |
| ENSGALG00000010035 | 2145,4649 | -0,574430705 | 0,1965144 | -2,923098 | 0,0034657 | 0,0148241 | NR3C2 |
| ENSGALG00000016807 | 10081,673 | -0,574724846 | 0,1138634 | -5,047493 | 4,48E-07 | 6,28E-06 | LIMS1 |
| ENSGALG00000016841 | 10674,737 | -0,576008454 | 0,1276781 | -4,511411 | 6,44E-06 | 6,90E-05 | COL4A1 |
| ENSGALG00000020057 | 1737,427 | -0,576621955 | 0,1365122 | -4,223958 | 2,40E-05 | 0,0002196 | PIGF |
| ENSGALG00000007992 | 6680,6643 | -0,5769061 | 0,1093351 | -5,276494 | 1,32E-07 | 2,06E-06 | GGA3 |
| ENSGALG00000003591 | 1041,8529 | -0,577610116 | 0,1793022 | -3,221433 | 0,0012755 | 0,006536 | FTO |
| ENSGALG00000014939 | 2493,3187 | -0,579384653 | 0,1743511 | -3,323091 | 0,0008903 | 0,004862 | GFM2 |
| ENSGALG00000004612 | 6633,1212 | -0,579433401 | 0,1094771 | -5,292734 | 1,21E-07 | 1,90E-06 | MTHFR |
| ENSGALG00000007008 | 8124,263 | -0,579649516 | 0,117537 | -4,931634 | 8,15E-07 | 1,08E-05 | POLR3E |
| ENSGALG00000004140 | 1503,5608 | -0,579940212 | 0,1679604 | -3,452839 | 0,0005547 | 0,0032709 | SH3BP4 |
| ENSGALG00000015629 | 16787,85 | -0,581629702 | 0,1013849 | -5,736848 | 9,65E-09 | 1,91E-07 | TRAM1 |
| ENSGALG00000010202 | 1508,6027 | -0,581722546 | 0,1688428 | -3,445349 | 0,0005703 | 0,0033455 | ENTPD5 |
| ENSGALG00000006713 | 2292,8924 | -0,58181715 | 0,1450076 | -4,012322 | 6,01E-05 | 0,0004821 | CELF2 |
| ENSGALG00000011464 | 79055,793 | -0,582311331 | 0,1058239 | -5,502646 | 3,74E-08 | 6,56E-07 | EIF5 |
| ENSGALG00000017130 | 3473,8029 | -0,582570563 | 0,1291633 | -4,510341 | 6,47E-06 | 6,91E-05 | LATS2 |
| ENSGALG00000008288 | 656,98572 | -0,582969807 | 0,1906368 | -3,058013 | 0,0022281 | 0,0103145 | ARMC10 |
| ENSGALG00000015000 | 15510,253 | -0,584898632 | 0,104418 | -5,601509 | 2,12E-08 | 3,93E-07 | LYAR |
| ENSGALG00000020899 | 644,11569 | -0,585391733 | 0,1791404 | -3,267782 | 0,0010839 | 0,0057134 | |
| ENSGALG00000027517 | 979,67929 | -0,585667855 | 0,1639516 | -3,5722 | 0,000354 | 0,0022407 | gga-mir-3064 |
| ENSGALG00000007677 | 1856,8714 | -0,585818002 | 0,1452003 | -4,03455 | 5,47E-05 | 0,0004451 | THNSL1 |
| ENSGALG00000003426 | 1636,1649 | -0,587203746 | 0,1565835 | -3,750099 | 0,0001768 | 0,0012329 | RHBDL3 |
| ENSGALG00000015765 | 12998,254 | -0,587270267 | 0,1455632 | -4,034469 | 5,47E-05 | 0,0004451 | POLR1A |
| ENSGALG00000015310 | 4463,7435 | -0,587354563 | 0,1636253 | -3,589632 | 0,0003311 | 0,0021177 | SOBP |
| ENSGALG00000008263 | 616,84886 | -0,587476931 | 0,2027246 | -2,897907 | 0,0037566 | 0,0158984 | CNTN4 |
| ENSGALG00000004144 | 27007,859 | -0,588131949 | 0,109754 | -5,358638 | 8,39E-08 | 1,37E-06 | DDX50 |
| ENSGALG00000012946 | 11580,052 | -0,588849778 | 0,1146976 | -5,133933 | 2,84E-07 | 4,13E-06 | FAM60A |
| ENSGALG00000016187 | 8513,8325 | -0,588903063 | 0,112954 | -5,213653 | 1,85E-07 | 2,82E-06 | WDR4 |
| ENSGALG00000000284 | 7651,6591 | -0,589684267 | 0,1227001 | -4,8059 | 1,54E-06 | 1,92E-05 | HOXB4 |
| ENSGALG00000025668 | 302,68906 | -0,590006716 | 0,2416858 | -2,441214 | 0,014638 | 0,0474627 | |
| ENSGALG00000015556 | 4802,1991 | -0,591646833 | 0,1279382 | -4,624475 | 3,76E-06 | 4,27E-05 | AFAP1 |
| ENSGALG00000012147 | 18627,042 | -0,591734529 | 0,112588 | -5,255749 | 1,47E-07 | 2,30E-06 | DDX18 |
| ENSGALG00000008376 | 8424,8255 | -0,592197559 | 0,1214407 | -4,876433 | 1,08E-06 | 1,40E-05 | EIF2B5 |
| ENSGALG00000005803 | 3271,7652 | -0,592808251 | 0,1582098 | -3,746976 | 0,000179 | 0,0012457 | ZNRF3 |

| | | | | | | | |
|---|---|---|---|---|---|---|---|
| ENSGALG00000002326 | 890,06973 | -0,593338649 | 0,1837643 | -3,228802 | 0,0012431 | 0,0064011 | |
| ENSGALG00000028863 | 286,98407 | -0,594135525 | 0,2453357 | -2,421725 | 0,015447 | 0,049555 | |
| ENSGALG00000011449 | 1934,5212 | -0,594721435 | 0,1428135 | -4,164321 | 3,12E-05 | 0,000275 | |
| ENSGALG00000007536 | 1357,1535 | -0,59551277 | 0,156731 | -3,799585 | 0,0001449 | 0,0010337 | |
| ENSGALG00000007048 | 584,57505 | -0,595618933 | 0,2088367 | -2,85208 | 0,0043434 | 0,0178334 | SYNM |
| ENSGALG00000005692 | 514,92687 | -0,595884023 | 0,2256362 | -2,640906 | 0,0082685 | 0,0302134 | CRISPLD2 |
| ENSGALG00000014213 | 4954,5042 | -0,596053951 | 0,129599 | -4,599216 | 4,24E-06 | 4,76E-05 | PRR5 |
| ENSGALG00000009458 | 5058,5683 | -0,596085417 | 0,1247109 | -4,779738 | 1,76E-06 | 2,15E-05 | |
| ENSGALG00000005171 | 1357,2271 | -0,597406416 | 0,14458 | -4,132013 | 3,60E-05 | 0,0003107 | AP1S3 |
| ENSGALG00000005243 | 13547,504 | -0,597738338 | 0,1028436 | -5,812113 | 6,17E-09 | 1,27E-07 | PPP3CB |
| ENSGALG00000020438 | 1854,2018 | -0,597817303 | 0,1402495 | -4,262527 | 2,02E-05 | 0,0001893 | |
| ENSGALG00000027159 | 3433,7581 | -0,598718251 | 0,1676492 | -3,571256 | 0,0003553 | 0,0022435 | CNKSR3 |
| ENSGALG00000014615 | 13865,336 | -0,598774766 | 0,1450012 | -4,129446 | 3,64E-05 | 0,0003134 | LAMA1 |
| ENSGALG00000011210 | 4763,2532 | -0,598839459 | 0,1129355 | -5,302492 | 1,14E-07 | 1,81E-06 | FAM175A |
| ENSGALG00000020003 | 15165,061 | -0,59891041 | 0,1111234 | -5,389599 | 7,06E-08 | 1,17E-06 | MAP3K4 |
| ENSGALG00000015409 | 17031,867 | -0,599227418 | 0,1473812 | -4,065834 | 4,79E-05 | 0,0003956 | PROS1 |
| ENSGALG00000011710 | 2961,6307 | -0,599591029 | 0,1881948 | -3,186013 | 0,0014425 | 0,007223 | BTBD6 |
| ENSGALG00000009437 | 2415,945 | -0,59959234 | 0,2125339 | -2,821161 | 0,004785 | 0,0193375 | MECOM |
| ENSGALG00000001003 | 4385,7574 | -0,600081977 | 0,1214895 | -4,939372 | 7,84E-07 | 1,04E-05 | SEPN1 |
| ENSGALG00000006256 | 7733,3335 | -0,600800071 | 0,1126462 | -5,333512 | 9,63E-08 | 1,55E-06 | MAP7D3 |
| ENSGALG00000003110 | 375,89024 | -0,601903021 | 0,2441478 | -2,465322 | 0,013689 | 0,044964 | |
| ENSGALG00000011280 | 7822,1495 | -0,603497491 | 0,1227685 | -4,915737 | 8,84E-07 | 1,16E-05 | EEA1 |
| ENSGALG00000010603 | 12309,752 | -0,604746743 | 0,1082631 | -5,585897 | 2,32E-08 | 4,26E-07 | CEBPZ |
| ENSGALG00000005807 | 96623,567 | -0,604834873 | 0,1046771 | -5,778103 | 7,55E-09 | 1,52E-07 | IPO7 |
| ENSGALG00000007030 | 1042,0347 | -0,605544059 | 0,1602843 | -3,777937 | 0,0001581 | 0,0011161 | |
| ENSGALG00000010623 | 6764,8612 | -0,606701235 | 0,111882 | -5,422691 | 5,87E-08 | 9,92E-07 | ORC1 |
| ENSGALG00000006147 | 7732,277 | -0,607224855 | 0,1099367 | -5,523405 | 3,32E-08 | 5,88E-07 | NUCB2 |
| ENSGALG00000009348 | 14314,788 | -0,607303929 | 0,150627 | -4,031839 | 5,53E-05 | 0,0004496 | AKAP9 |
| ENSGALG00000011522 | 2524,7796 | -0,607850232 | 0,1342636 | -4,527291 | 5,97E-06 | 6,44E-05 | TRMT61A |
| ENSGALG00000012407 | 335,57386 | -0,607964974 | 0,2234167 | -2,721215 | 0,0065042 | 0,02491 | LRP1B |
| ENSGALG00000010538 | 11960,175 | -0,60879058 | 0,1100325 | -5,532827 | 3,15E-08 | 5,61E-07 | TTC27 |
| ENSGALG00000011618 | 1230,2077 | -0,609057453 | 0,1895822 | -3,21263 | 0,0013153 | 0,0067043 | ADSSL1 |
| ENSGALG00000015689 | 13523,347 | -0,609077567 | 0,14383 | -4,234705 | 2,29E-05 | 0,0002105 | KIAA0368 |
| ENSGALG00000005492 | 6703,9725 | -0,609713298 | 0,1420841 | -4,291213 | 1,78E-05 | 0,0001688 | NBEAL2 |
| ENSGALG00000009698 | 6354,4127 | -0,610790424 | 0,1589015 | -3,843831 | 0,0001211 | 0,0008818 | SLC38A1 |
| ENSGALG00000016302 | 7538,4329 | -0,61104581 | 0,1423529 | -4,292472 | 1,77E-05 | 0,0001681 | ASAP1 |
| ENSGALG00000000769 | 395,02523 | -0,611277614 | 0,2259769 | -2,705045 | 0,0068295 | 0,0258936 | RAB7B |
| ENSGALG00000000438 | 3178,9721 | -0,61235395 | 0,1445774 | -4,235475 | 2,28E-05 | 0,0002099 | ERMP1 |
| ENSGALG00000010607 | 463,53756 | -0,613528032 | 0,2050464 | -2,992142 | 0,0027703 | 0,0123558 | SPATA7 |
| ENSGALG00000004169 | 13748,93 | -0,613558741 | 0,1070672 | -5,730594 | 1,00E-08 | 1,97E-07 | |
| ENSGALG00000013760 | 3088,0215 | -0,614432788 | 0,1210707 | -5,07499 | 3,88E-07 | 5,50E-06 | GFPT2 |
| ENSGALG00000016574 | 1715,0637 | -0,614768808 | 0,142039 | -4,328168 | 1,50E-05 | 0,0001471 | GEMIN8 |
| ENSGALG00000007703 | 1248,9929 | -0,614958252 | 0,1535298 | -4,005464 | 6,19E-05 | 0,0004943 | |
| ENSGALG00000013480 | 4896,2242 | -0,615680556 | 0,1242349 | -4,955777 | 7,20E-07 | 9,61E-06 | MTHFD2 |
| ENSGALG00000016808 | 40422,889 | -0,615970468 | 0,1230783 | -5,004705 | 5,59E-07 | 7,68E-06 | RANBP2 |
| ENSGALG00000006237 | 12767,815 | -0,616678741 | 0,1272849 | -4,844871 | 1,27E-06 | 1,61E-05 | PKN2 |
| ENSGALG00000016951 | 1808,556 | -0,617422601 | 0,1357822 | -4,547156 | 5,44E-06 | 5,94E-05 | MTRF1 |
| ENSGALG00000007631 | 328,24576 | -0,617881783 | 0,2272908 | -2,718464 | 0,0065586 | 0,0250618 | MEGF11 |
| ENSGALG00000028279 | 367,51739 | -0,618647245 | 0,2374707 | -2,605152 | 0,0091834 | 0,0327994 | |
| ENSGALG00000008265 | 23793,57 | -0,619236444 | 0,1097973 | -5,639813 | 1,70E-08 | 3,22E-07 | PDCD11 |
| ENSGALG00000017388 | 6991,3984 | -0,619582917 | 0,1159517 | -5,343459 | 9,12E-08 | 1,48E-06 | YY1 |
| ENSGALG00000005815 | 11029,748 | -0,619938317 | 0,1099084 | -5,640501 | 1,70E-08 | 3,22E-07 | TMEM41B |
| ENSGALG00000012533 | 37098,602 | -0,619977689 | 0,143535 | -4,31935 | 1,56E-05 | 0,0001518 | MYH9 |
| ENSGALG00000001192 | 22737,825 | -0,620466491 | 0,1546499 | -4,012072 | 6,02E-05 | 0,0004822 | ARHGAP32 |
| ENSGALG00000011058 | 317,84832 | -0,620682204 | 0,234088 | -2,651491 | 0,0080137 | 0,0294645 | LEPR |
| ENSGALG00000016653 | 1523,2789 | -0,621295808 | 0,1474254 | -4,214308 | 2,51E-05 | 0,0002282 | FAM167A |
| ENSGALG00000012447 | 5839,1863 | -0,62156312 | 0,1278297 | -4,862432 | 1,16E-06 | 1,49E-05 | TNS3 |
| ENSGALG00000005319 | 350,2389 | -0,621682491 | 0,2536246 | -2,451192 | 0,0142384 | 0,0464103 | CDH8 |
| ENSGALG00000008539 | 7630,1458 | -0,622137491 | 0,1211519 | -5,135185 | 2,82E-07 | 4,11E-06 | ALG12 |
| ENSGALG00000023419 | 9218,5287 | -0,622730836 | 0,1310534 | -4,751733 | 2,02E-06 | 2,42E-05 | HOXD4 |
| ENSGALG00000015271 | 532,65573 | -0,623845538 | 0,2308973 | -2,701831 | 0,0068959 | 0,0260871 | FILIP1L |
| ENSGALG00000000234 | 2924,3601 | -0,623968223 | 0,1484686 | -4,202696 | 2,64E-05 | 0,0002381 | EPB41L4A |
| ENSGALG00000005438 | 938,76871 | -0,624083264 | 0,1616022 | -3,861849 | 0,0001125 | 0,0008286 | PARVA |
| ENSGALG00000010975 | 6618,9998 | -0,624280901 | 0,1157101 | -5,395213 | 6,84E-08 | 1,14E-06 | TMTC2 |

| | | | | | | | |
|---|---|---|---|---|---|---|---|
| ENSGALG00000005504 | 1081,6574 | -0,624552568 | 0,1731409 | -3,607192 | 0,0003095 | 0,0019954 | HNF1B |
| ENSGALG00000006609 | 9375,2554 | -0,624834648 | 0,1373671 | -4,548649 | 5,40E-06 | 5,91E-05 | EPHB1 |
| ENSGALG00000014860 | 10686,581 | -0,625239897 | 0,107074 | -5,839324 | 5,24E-09 | 1,09E-07 | YES1 |
| ENSGALG00000017159 | 5806,6713 | -0,626252438 | 0,142901 | -4,382421 | 1,17E-05 | 0,0001187 | ATM |
| ENSGALG00000004046 | 1117,4696 | -0,627070394 | 0,169937 | -3,690016 | 0,0002242 | 0,0015092 | LARP6 |
| ENSGALG00000014855 | 4504,1501 | -0,627700184 | 0,1611556 | -3,894995 | 9,82E-05 | 0,0007383 | GHR |
| ENSGALG00000010702 | 11951,045 | -0,628390454 | 0,1083577 | -5,799225 | 6,66E-09 | 1,36E-07 | ADSS |
| ENSGALG00000015472 | 37144,636 | -0,629536006 | 0,1475373 | -4,266963 | 1,98E-05 | 0,000186 | CHD7 |
| ENSGALG00000012966 | 5358,7576 | -0,62976328 | 0,1160975 | -5,424435 | 5,81E-08 | 9,83E-07 | ZBTB2 |
| ENSGALG00000005450 | 5500,7304 | -0,632586778 | 0,1208381 | -5,234993 | 1,65E-07 | 2,54E-06 | CCDC66 |
| ENSGALG00000005256 | 11273,337 | -0,63393871 | 0,1641492 | -3,861966 | 0,0001125 | 0,0008286 | EPHA4 |
| ENSGALG00000014337 | 1369,7017 | -0,634337238 | 0,1462286 | -4,337984 | 1,44E-05 | 0,0001418 | NWD2 |
| ENSGALG00000009850 | 11171,857 | -0,63459607 | 0,108041 | -5,87366 | 4,26E-09 | 9,11E-08 | NOVA1 |
| ENSGALG00000004937 | 10776,497 | -0,634904509 | 0,1095156 | -5,797386 | 6,74E-09 | 1,37E-07 | VGLL4 |
| ENSGALG00000017368 | 7609,7868 | -0,635358308 | 0,1095099 | -5,801832 | 6,56E-09 | 1,34E-07 | TRNT1 |
| ENSGALG00000007392 | 3781,9294 | -0,635704603 | 0,1408897 | -4,512074 | 6,42E-06 | 6,88E-05 | ADAMTS9 |
| ENSGALG00000003767 | 4407,983 | -0,636798593 | 0,1177151 | -5,409657 | 6,31E-08 | 1,06E-06 | NKD1 |
| ENSGALG00000007710 | 1737,5833 | -0,637223247 | 0,1525396 | -4,177427 | 2,95E-05 | 0,0002612 | |
| ENSGALG00000012813 | 15846,526 | -0,637913408 | 0,1220106 | -5,228345 | 1,71E-07 | 2,63E-06 | PRPF4B |
| ENSGALG00000005521 | 7067,2749 | -0,637995859 | 0,1317048 | -4,844135 | 1,27E-06 | 1,62E-05 | PER2 |
| ENSGALG00000022490 | 838,24845 | -0,638026939 | 0,2048258 | -3,114973 | 0,0018396 | 0,008782 | ARPC1B |
| ENSGALG00000023451 | 10451,614 | -0,638494594 | 0,1052822 | -6,064604 | 1,32E-09 | 3,09E-08 | TRPM7 |
| ENSGALG00000007056 | 3476,5095 | -0,638790551 | 0,1810296 | -3,528652 | 0,0004177 | 0,0025816 | KLF6 |
| ENSGALG00000003891 | 23052,983 | -0,638963678 | 0,1001377 | -6,380853 | 1,76E-10 | 4,67E-09 | ZNF423 |
| ENSGALG00000008304 | 1332,9115 | -0,639371492 | 0,2087659 | -3,062623 | 0,0021941 | 0,0101883 | OBFC1 |
| ENSGALG00000012662 | 3593,3785 | -0,639523486 | 0,1240921 | -5,153618 | 2,56E-07 | 3,78E-06 | NUAK1 |
| ENSGALG00000014138 | 4048,403 | -0,640267154 | 0,1406798 | -4,551239 | 5,33E-06 | 5,86E-05 | NFXL1 |
| ENSGALG00000011162 | 1836,7364 | -0,640538648 | 0,1515623 | -4,226239 | 2,38E-05 | 0,0002177 | CYP46A1 |
| ENSGALG00000021238 | 337,09341 | -0,640934901 | 0,2252878 | -2,84496 | 0,0044417 | 0,0181603 | CYP2W1 |
| ENSGALG00000009240 | 3768,4908 | -0,641416186 | 0,1167026 | -5,49616 | 3,88E-08 | 6,79E-07 | NFE2L2 |
| ENSGALG00000007352 | 21915,639 | -0,641667347 | 0,1092352 | -5,874181 | 4,25E-09 | 9,10E-08 | SPAG9 |
| ENSGALG00000009445 | 3221,2949 | -0,642383978 | 0,1226543 | -5,237353 | 1,63E-07 | 2,51E-06 | INPP5F |
| ENSGALG00000010642 | 1388,2464 | -0,643267076 | 0,1679801 | -3,829424 | 0,0001284 | 0,0009312 | IRF2 |
| ENSGALG00000027704 | 758,7446 | -0,643979597 | 0,2120706 | -3,036628 | 0,0023924 | 0,0109231 | |
| ENSGALG00000005321 | 14326,683 | -0,645095033 | 0,1462871 | -4,409787 | 1,03E-05 | 0,0001064 | LAMA5 |
| ENSGALG00000005024 | 4274,063 | -0,645541534 | 0,1373988 | -4,698306 | 2,62E-06 | 3,09E-05 | EFCC1 |
| ENSGALG00000026181 | 1587,8742 | -0,645581251 | 0,1576569 | -4,094848 | 4,22E-05 | 0,0003559 | STRIP2 |
| ENSGALG00000010536 | 11449,034 | -0,646034394 | 0,1106982 | -5,835997 | 5,35E-09 | 1,11E-07 | FAF1 |
| ENSGALG00000016480 | 10063,583 | -0,647091328 | 0,1222876 | -5,291552 | 1,21E-07 | 1,91E-06 | SDC1 |
| ENSGALG00000000258 | 23273,697 | -0,649481888 | 0,1229147 | -5,284004 | 1,26E-07 | 1,98E-06 | HEATR1 |
| ENSGALG00000000761 | 2678,9011 | -0,649687901 | 0,1320076 | -4,921596 | 8,58E-07 | 1,13E-05 | TSKU |
| ENSGALG00000014902 | 5870,3239 | -0,650325285 | 0,1293417 | -5,027965 | 4,96E-07 | 6,90E-06 | RP1-179P9.3 |
| ENSGALG00000012847 | 300,13784 | -0,651077536 | 0,259355 | -2,510372 | 0,0120604 | 0,0407551 | |
| ENSGALG00000006864 | 655,13638 | -0,651213169 | 0,1997356 | -3,260377 | 0,0011126 | 0,0058389 | COL24A1 |
| ENSGALG00000014873 | 10860,594 | -0,6514166 | 0,1060541 | -6,142302 | 8,13E-10 | 1,95E-08 | GJA1 |
| ENSGALG00000012232 | 769,02109 | -0,652989979 | 0,197777 | -3,301647 | 0,0009612 | 0,0051726 | DNAAF2 |
| ENSGALG00000003154 | 3509,1767 | -0,653496705 | 0,122188 | -5,348291 | 8,88E-08 | 1,45E-06 | TXNDC11 |
| ENSGALG00000005845 | 8323,5942 | -0,655210424 | 0,124744 | -5,252442 | 1,50E-07 | 2,33E-06 | SLC7A5 |
| ENSGALG00000015241 | 1466,0595 | -0,655397927 | 0,141762 | -4,623228 | 3,78E-06 | 4,29E-05 | SNAI2 |
| ENSGALG00000015653 | 733,01948 | -0,655959247 | 0,2027357 | -3,235538 | 0,0012141 | 0,0062782 | JPH1 |
| ENSGALG00000016681 | 817,92363 | -0,656368052 | 0,1696389 | -3,869207 | 0,0001092 | 0,0008088 | DHRSX |
| ENSGALG00000000690 | 5466,0012 | -0,657633862 | 0,1415168 | -4,647039 | 3,37E-06 | 3,87E-05 | GDPD4 |
| ENSGALG00000016817 | 6729,2541 | -0,658902643 | 0,1108429 | -5,944473 | 2,77E-09 | 6,16E-08 | RASA3 |
| ENSGALG00000006975 | 7158,0796 | -0,659116429 | 0,117355 | -5,616432 | 1,95E-08 | 3,65E-07 | TBC1D22A |
| ENSGALG00000006840 | 2751,6848 | -0,659559733 | 0,1711898 | -3,852798 | 0,0001168 | 0,0008552 | AKAP13 |
| ENSGALG00000006681 | 619,27869 | -0,660452265 | 0,2149441 | -3,07267 | 0,0021215 | 0,009926 | BRSK2 |
| ENSGALG00000010204 | 972,3601 | -0,660708753 | 0,1680654 | -3,93126 | 8,45E-05 | 0,0006489 | KCNC2 |
| ENSGALG00000008551 | 7057,1759 | -0,661341847 | 0,1249566 | -5,292574 | 1,21E-07 | 1,90E-06 | CRELD2 |
| ENSGALG00000007870 | 12818,558 | -0,66197661 | 0,1166502 | -5,674884 | 1,39E-08 | 2,68E-07 | SMAD3 |
| ENSGALG00000001405 | 334,6643 | -0,663151012 | 0,2350856 | -2,820891 | 0,004789 | 0,019348 | IRF6 |
| ENSGALG00000005796 | 13227,627 | -0,664735985 | 0,1063503 | -6,25044 | 4,09E-10 | 1,01E-08 | XBP1 |
| ENSGALG00000008293 | 829,33594 | -0,66560113 | 0,2005473 | -3,318924 | 0,0009036 | 0,0049193 | SH3PXD2A |
| ENSGALG00000016246 | 560,38544 | -0,667393695 | 0,2438472 | -2,736934 | 0,0062015 | 0,0239052 | KCNQ3 |
| ENSGALG00000009915 | 13723,099 | -0,66741216 | 0,1151588 | -5,79558 | 6,81E-09 | 1,39E-07 | EML4 |

| | | | | | | | |
|---|---|---|---|---|---|---|---|
| ENSGALG00000005512 | 8645,0111 | -0,667653797 | 0,1184896 | -5,634703 | 1,75E-08 | 3,31E-07 | KIAA1671 |
| ENSGALG00000008418 | 2809,2517 | -0,668553187 | 0,1227062 | -5,448405 | 5,08E-08 | 8,65E-07 | GYLTL1B |
| ENSGALG00000011139 | 4725,8286 | -0,66929694 | 0,1340022 | -4,99467 | 5,89E-07 | 8,03E-06 | CCDC85C |
| ENSGALG00000011686 | 12938,733 | -0,670528172 | 0,1470322 | -4,560417 | 5,11E-06 | 5,64E-05 | SEMA5B |
| ENSGALG00000007278 | 350,13209 | -0,670809216 | 0,2308902 | -2,905317 | 0,0036688 | 0,0155756 | GRIN2A |
| ENSGALG00000008240 | 2340,2977 | -0,670953542 | 0,1295685 | -5,178368 | 2,24E-07 | 3,35E-06 | CFLAR |
| ENSGALG00000006080 | 30589,303 | -0,672288685 | 0,1018389 | -6,601491 | 4,07E-11 | 1,17E-09 | GPC4 |
| ENSGALG00000028291 | 15611,072 | -0,672824764 | 0,1341958 | -5,013756 | 5,34E-07 | 7,39E-06 | BCAR1 |
| ENSGALG00000004715 | 1867,9414 | -0,672888292 | 0,1525035 | -4,41228 | 1,02E-05 | 0,0001053 | ZC3H12B |
| ENSGALG00000015912 | 3388,0589 | -0,674238434 | 0,1460392 | -4,616833 | 3,90E-06 | 4,41E-05 | NBN |
| ENSGALG00000015708 | 22628,562 | -0,674375402 | 0,1784895 | -3,778235 | 0,0001579 | 0,0011154 | FGFR3 |
| ENSGALG00000008684 | 70609,009 | -0,674536207 | 0,1288068 | -5,236804 | 1,63E-07 | 2,52E-06 | EIF4A2 |
| ENSGALG00000010049 | 749,198 | -0,6748978 | 0,2082509 | -3,240792 | 0,001192 | 0,0061824 | PTPRB |
| ENSGALG00000003249 | 7468,8212 | -0,675105103 | 0,1116673 | -6,045684 | 1,49E-09 | 3,45E-08 | CWF19L1 |
| ENSGALG00000006035 | 6729,7716 | -0,676290578 | 0,1083357 | -6,242548 | 4,30E-10 | 1,06E-08 | PDE3B |
| ENSGALG00000004494 | 4756,9197 | -0,676838081 | 0,11958 | -5,66013 | 1,51E-08 | 2,89E-07 | CCNE1 |
| ENSGALG00000016451 | 7945,9007 | -0,679900325 | 0,139439 | -4,87597 | 1,08E-06 | 1,40E-05 | ROCK2 |
| ENSGALG00000025674 | 227,33639 | -0,681326883 | 0,2807179 | -2,427088 | 0,0152206 | 0,0489092 | |
| ENSGALG00000006437 | 3771,9905 | -0,682364759 | 0,1217534 | -5,60448 | 2,09E-08 | 3,87E-07 | NOM1 |
| ENSGALG00000009156 | 5048,9849 | -0,682556058 | 0,1676146 | -4,072176 | 4,66E-05 | 0,0003862 | CDAN1 |
| ENSGALG00000014274 | 3304,6353 | -0,683648491 | 0,1394936 | -4,900932 | 9,54E-07 | 1,25E-05 | RIMKLB |
| ENSGALG00000012390 | 2074,4615 | -0,684899295 | 0,1473975 | -4,646615 | 3,37E-06 | 3,87E-05 | PPP1R14C |
| ENSGALG00000009152 | 8451,9252 | -0,685418448 | 0,1346381 | -5,09082 | 3,57E-07 | 5,10E-06 | ATRNL1 |
| ENSGALG00000004631 | 4926,2821 | -0,686947368 | 0,1966631 | -3,493016 | 0,0004776 | 0,0028862 | DRAXIN |
| ENSGALG00000010836 | 1260,2339 | -0,688393097 | 0,166876 | -4,125178 | 3,70E-05 | 0,0003178 | AHR |
| ENSGALG00000005990 | 327,32624 | -0,688776212 | 0,2796142 | -2,463309 | 0,0137661 | 0,0451737 | MGLL |
| ENSGALG00000003726 | 1048,3673 | -0,689550024 | 0,2280538 | -3,023628 | 0,0024976 | 0,0113049 | EGFLAM |
| ENSGALG00000015879 | 11968,416 | -0,689623134 | 0,1480044 | -4,659477 | 3,17E-06 | 3,66E-05 | HUNK |
| ENSGALG00000008234 | 5250,562 | -0,689849994 | 0,1273096 | -5,41868 | 6,00E-08 | 1,01E-06 | ORC2 |
| ENSGALG00000016308 | 5133,4014 | -0,690314419 | 0,1281438 | -5,387031 | 7,16E-08 | 1,18E-06 | MYC |
| ENSGALG00000014686 | 1912,6046 | -0,691021625 | 0,1730464 | -3,993274 | 6,52E-05 | 0,0005167 | FBN2 |
| ENSGALG00000025627 | 543,3063 | -0,691903941 | 0,1957541 | -3,534557 | 0,0004085 | 0,0025313 | |
| ENSGALG00000017866 | 156,05125 | -0,692156313 | 0,285038 | -2,428295 | 0,01517 | 0,0487927 | |
| ENSGALG00000008292 | 9417,4765 | -0,69263042 | 0,1222675 | -5,664879 | 1,47E-08 | 2,83E-07 | LRP4 |
| ENSGALG00000017857 | 160,75501 | -0,693298969 | 0,2819397 | -2,459032 | 0,0139312 | 0,0455946 | |
| ENSGALG00000011679 | 1487,1899 | -0,695574487 | 0,1740708 | -3,995929 | 6,44E-05 | 0,0005116 | DIRC2 |
| ENSGALG00000007371 | 25835,451 | -0,695805113 | 0,1165608 | -5,969459 | 2,38E-09 | 5,36E-08 | IGDCC3 |
| ENSGALG00000017220 | 8610,4222 | -0,695886193 | 0,1257779 | -5,532661 | 3,15E-08 | 5,61E-07 | |
| ENSGALG00000008758 | 3528,4952 | -0,696392948 | 0,1183183 | -5,88576 | 3,96E-09 | 8,56E-08 | B3GNT5 |
| ENSGALG00000028415 | 2970,1144 | -0,698216592 | 0,2041115 | -3,42076 | 0,0006245 | 0,0036072 | |
| ENSGALG00000016834 | 1336,2251 | -0,698351437 | 0,1627936 | -4,289798 | 1,79E-05 | 0,0001697 | MCF2L |
| ENSGALG00000017812 | 214,0597 | -0,698433493 | 0,2584052 | -2,702862 | 0,0068745 | 0,0260208 | |
| ENSGALG00000002316 | 6016,7291 | -0,700708149 | 0,1098757 | -6,377279 | 1,80E-10 | 4,77E-09 | NUBP2 |
| ENSGALG00000002689 | 3193,9897 | -0,701352786 | 0,134772 | -5,203994 | 1,95E-07 | 2,96E-06 | NUP35 |
| ENSGALG00000012654 | 165,03411 | -0,702415833 | 0,2836155 | -2,476648 | 0,0132623 | 0,0439509 | NFATC1 |
| ENSGALG00000007501 | 495,82592 | -0,702573548 | 0,2210424 | -3,178456 | 0,0014806 | 0,0073545 | P2RX3 |
| ENSGALG00000006592 | 563,46793 | -0,703616422 | 0,1908922 | -3,685937 | 0,0002279 | 0,0015305 | MEIKIN |
| ENSGALG00000021316 | 17378,378 | -0,704586437 | 0,1023184 | -6,886217 | 5,73E-12 | 1,88E-10 | SLC12A4 |
| ENSGALG00000012153 | 842,90994 | -0,706186244 | 0,1736471 | -4,066791 | 4,77E-05 | 0,0003943 | METTL15 |
| ENSGALG00000002762 | 6873,8438 | -0,70662137 | 0,1320656 | -5,350533 | 8,77E-08 | 1,43E-06 | UHRF1BP1 |
| ENSGALG00000002726 | 8639,3567 | -0,708755608 | 0,1395335 | -5,079466 | 3,78E-07 | 5,38E-06 | KIF1B |
| ENSGALG00000007141 | 2245,3684 | -0,70907976 | 0,1459375 | -4,858791 | 1,18E-06 | 1,52E-05 | LRRK1 |
| ENSGALG00000011680 | 3554,5974 | -0,709602993 | 0,1836532 | -3,86382 | 0,0001116 | 0,0008235 | EPHA5 |
| ENSGALG00000012848 | 3745,194 | -0,711139877 | 0,1738156 | -4,091346 | 4,29E-05 | 0,0003598 | KDM7A |
| ENSGALG00000022857 | 997,97159 | -0,711204976 | 0,1675693 | -4,244244 | 2,19E-05 | 0,000203 | |
| ENSGALG00000015733 | 14565,427 | -0,711632203 | 0,1740069 | -4,089677 | 4,32E-05 | 0,0003619 | MDN1 |
| ENSGALG00000017365 | 2855,5664 | -0,712049377 | 0,1517404 | -4,692551 | 2,70E-06 | 3,17E-05 | REEP2 |
| ENSGALG00000009562 | 5126,9731 | -0,714412903 | 0,1287612 | -5,548356 | 2,88E-08 | 5,20E-07 | PLEK2 |
| ENSGALG00000003970 | 25720,818 | -0,714513793 | 0,1240731 | -5,758812 | 8,47E-09 | 1,70E-07 | TJP1 |
| ENSGALG00000017871 | 177,34037 | -0,715510132 | 0,2848578 | -2,511815 | 0,0120112 | 0,0406308 | |
| ENSGALG00000003161 | 1569,9242 | -0,715518291 | 0,1658137 | -4,315193 | 1,59E-05 | 0,0001539 | GDF3 |
| ENSGALG00000016927 | 1190,189 | -0,715628819 | 0,1764386 | -4,055965 | 4,99E-05 | 0,0004112 | KLF5 |
| ENSGALG00000013615 | 780,84937 | -0,715808204 | 0,1833387 | -3,904293 | 9,45E-05 | 0,0007136 | MTUS1 |
| ENSGALG00000011076 | 244,64999 | -0,715907242 | 0,2845675 | -2,515773 | 0,0118772 | 0,0402859 | SYNE3 |

| | | | | | | | |
|---|---|---|---|---|---|---|---|
| ENSGALG00000010036 | 359,24032 | -0,717787007 | 0,2219662 | -3,233767 | 0,0012217 | 0,0063086 | FOSL2 |
| ENSGALG00000015122 | 2528,5766 | -0,719672556 | 0,1390865 | -5,174279 | 2,29E-07 | 3,42E-06 | TAF4B |
| ENSGALG00000020805 | 5789,8017 | -0,719835862 | 0,1139176 | -6,318916 | 2,63E-10 | 6,72E-09 | LRRC8D |
| ENSGALG00000005430 | 950,09062 | -0,72235333 | 0,2022537 | -3,571522 | 0,0003549 | 0,0022423 | ALS2CL |
| ENSGALG00000002553 | 2782,5411 | -0,723490416 | 0,1606217 | -4,504313 | 6,66E-06 | 7,10E-05 | PCDH1 |
| ENSGALG00000007242 | 1612,8307 | -0,724284111 | 0,1436296 | -5,04272 | 4,59E-07 | 6,42E-06 | SLITRK4 |
| ENSGALG00000008078 | 7758,4981 | -0,725207523 | 0,1252194 | -5,791493 | 6,98E-09 | 1,42E-07 | FUT8 |
| ENSGALG00000006978 | 1693,5845 | -0,725851222 | 0,16966 | -4,27827 | 1,88E-05 | 0,0001778 | APOOL |
| ENSGALG00000004004 | 8836,3289 | -0,725989536 | 0,1219894 | -5,951253 | 2,66E-09 | 5,95E-08 | PHKB |
| ENSGALG00000005246 | 9658,48 | -0,7272373 | 0,1233553 | -5,895469 | 3,74E-09 | 8,11E-08 | PAX3 |
| ENSGALG00000028294 | 3609,4147 | -0,727698928 | 0,1304345 | -5,579039 | 2,42E-08 | 4,40E-07 | VWA5B2 |
| ENSGALG00000006939 | 26463,229 | -0,728311968 | 0,1092521 | -6,666343 | 2,62E-11 | 7,76E-10 | PFKP |
| ENSGALG00000004519 | 237,39511 | -0,728806195 | 0,2713012 | -2,686336 | 0,007224 | 0,02714 | TRPV2 |
| ENSGALG00000006555 | 1381,6916 | -0,729109536 | 0,1554307 | -4,690898 | 2,72E-06 | 3,19E-05 | IGF2 |
| ENSGALG00000025743 | 3141,8277 | -0,729896626 | 0,1215881 | -6,003027 | 1,94E-09 | 4,42E-08 | CDR2 |
| ENSGALG00000016078 | 860,37098 | -0,73004938 | 0,1938639 | -3,765783 | 0,000166 | 0,0011664 | ZFPM2 |
| ENSGALG00000014419 | 9837,2387 | -0,730417334 | 0,1131065 | -6,457784 | 1,06E-10 | 2,87E-09 | SLIT2 |
| ENSGALG00000020479 | 3560,7787 | -0,731996794 | 0,1299087 | -5,634701 | 1,75E-08 | 3,31E-07 | RP11-872D17.8 |
| ENSGALG00000003378 | 7303,9139 | -0,732735156 | 0,1242316 | -5,89814 | 3,68E-09 | 8,01E-08 | USP42 |
| ENSGALG00000015627 | 1040,2204 | -0,73285779 | 0,2129145 | -3,442028 | 0,0005774 | 0,0033796 | HAPLN1 |
| ENSGALG00000026349 | 29196,624 | -0,736634172 | 0,0999371 | -7,370975 | 1,69E-13 | 6,80E-12 | HSD17B12 |
| ENSGALG00000007069 | 592,20837 | -0,738101541 | 0,2325984 | -3,173288 | 0,0015072 | 0,0074595 | ADAMTS17 |
| ENSGALG00000016537 | 11629,834 | -0,739933749 | 0,1619397 | -4,569193 | 4,90E-06 | 5,45E-05 | SCML2 |
| ENSGALG00000010448 | 40598,94 | -0,742255902 | 0,1099244 | -6,752421 | 1,45E-11 | 4,48E-10 | LTBP1 |
| ENSGALG00000002679 | 1869,603 | -0,743662338 | 0,1515547 | -4,90689 | 9,25E-07 | 1,21E-05 | GLT1D1 |
| ENSGALG00000007331 | 18072,434 | -0,744880346 | 0,1084385 | -6,869149 | 6,46E-12 | 2,09E-10 | SVIL |
| ENSGALG00000027639 | 27019,601 | -0,74689172 | 0,1081151 | -6,908304 | 4,90E-12 | 1,61E-10 | RGMA |
| ENSGALG00000015077 | 367,6495 | -0,747092368 | 0,2656853 | -2,811945 | 0,0049243 | 0,0198356 | ARHGAP31 |
| ENSGALG00000010522 | 11132,1 | -0,747786716 | 0,1199212 | -6,23565 | 4,50E-10 | 1,11E-08 | ALDH18A1 |
| ENSGALG00000008741 | 273,37173 | -0,747824755 | 0,2458046 | -3,042355 | 0,0023474 | 0,0107607 | |
| ENSGALG00000003975 | 3354,1135 | -0,748226558 | 0,1450875 | -5,15707 | 2,51E-07 | 3,71E-06 | CEP112 |
| ENSGALG00000008910 | 360,62302 | -0,749041615 | 0,2212545 | -3,385431 | 0,0007107 | 0,0040265 | CERKL |
| ENSGALG00000003149 | 1878,4222 | -0,749443102 | 0,1529534 | -4,899812 | 9,59E-07 | 1,25E-05 | ITPR3 |
| ENSGALG00000012932 | 193,43191 | -0,749547645 | 0,2920342 | -2,566643 | 0,0102688 | 0,0359195 | FGD4 |
| ENSGALG00000016144 | 227,03636 | -0,75041933 | 0,282532 | -2,656051 | 0,0079062 | 0,0291714 | TMPRSS2 |
| ENSGALG00000007011 | 11491,263 | -0,750635431 | 0,1125238 | -6,670901 | 2,54E-11 | 7,56E-10 | ZBTB16 |
| ENSGALG00000023464 | 2176,7949 | -0,751478603 | 0,162842 | -4,614771 | 3,94E-06 | 4,45E-05 | |
| ENSGALG00000018479 | 498,05044 | -0,753094112 | 0,2693867 | -2,795587 | 0,0051806 | 0,0206479 | |
| ENSGALG00000019961 | 8882,8923 | -0,7532835 | 0,1292302 | -5,829007 | 5,58E-09 | 1,16E-07 | SLC18B1 |
| ENSGALG00000003430 | 9094,4226 | -0,754857044 | 0,1082845 | -6,971053 | 3,15E-12 | 1,08E-10 | PMS2 |
| ENSGALG00000015862 | 12207,703 | -0,756350316 | 0,1499013 | -5,045657 | 4,52E-07 | 6,34E-06 | CA2 |
| ENSGALG00000008318 | 18620,526 | -0,756789619 | 0,1179267 | -6,41746 | 1,39E-10 | 3,71E-09 | SLK |
| ENSGALG00000010412 | 643,75824 | -0,756929566 | 0,1797201 | -4,211714 | 2,53E-05 | 0,00023 | WWTR1 |
| ENSGALG00000004981 | 3521,9026 | -0,757070103 | 0,1329306 | -5,695226 | 1,23E-08 | 2,40E-07 | TSEN2 |
| ENSGALG00000006981 | 1290,8961 | -0,757579596 | 0,1692071 | -4,477234 | 7,56E-06 | 8,01E-05 | MCTP2 |
| ENSGALG00000007667 | 1024,279 | -0,757882718 | 0,1665229 | -4,551224 | 5,33E-06 | 5,86E-05 | GPR158 |
| ENSGALG00000000742 | 7852,9776 | -0,757900463 | 0,1311808 | -5,777528 | 7,58E-09 | 1,53E-07 | PIM1 |
| ENSGALG00000001419 | 2086,701 | -0,758492411 | 0,1407402 | -5,389307 | 7,07E-08 | 1,17E-06 | DAB2IP |
| ENSGALG00000016294 | 651,26015 | -0,761040192 | 0,2034016 | -3,741564 | 0,0001829 | 0,0012678 | HMGCLL1 |
| ENSGALG00000009748 | 12417,603 | -0,762363604 | 0,1219472 | -6,251585 | 4,06E-10 | 1,01E-08 | ASNS |
| ENSGALG00000006167 | 4665,1269 | -0,763429455 | 0,1613684 | -4,730971 | 2,23E-06 | 2,66E-05 | LRRC8B |
| ENSGALG00000017462 | 3834,7093 | -0,763892042 | 0,1177124 | -6,489476 | 8,61E-11 | 2,38E-09 | FNIP1 |
| ENSGALG00000028254 | 428,45012 | -0,764100303 | 0,2118876 | -3,606159 | 0,0003108 | 0,0019995 | |
| ENSGALG00000007300 | 11025,546 | -0,764508198 | 0,1238668 | -6,172016 | 6,74E-10 | 1,63E-08 | PYCR2 |
| ENSGALG00000026862 | 22574,745 | -0,764882325 | 0,1624148 | -4,709438 | 2,48E-06 | 2,94E-05 | CLDN1 |
| ENSGALG00000008621 | 9873,262 | -0,76612038 | 0,131649 | -5,819416 | 5,91E-09 | 1,22E-07 | NRP2 |
| ENSGALG00000027389 | 3852,8154 | -0,766788012 | 0,1308552 | -5,85982 | 4,63E-09 | 9,83E-08 | RGS7BP |
| ENSGALG00000010598 | 220,78888 | -0,767088888 | 0,2736996 | -2,802667 | 0,0050682 | 0,0202892 | KCNK10 |
| ENSGALG00000013782 | 3281,2809 | -0,769533335 | 0,1760353 | -4,371472 | 1,23E-05 | 0,0001244 | CDH7 |
| ENSGALG00000026897 | 591,30007 | -0,770711311 | 0,2031153 | -3,794453 | 0,000148 | 0,0010531 | |
| ENSGALG00000028737 | 5045,3512 | -0,771568179 | 0,1454081 | -5,306224 | 1,12E-07 | 1,78E-06 | |
| ENSGALG00000000317 | 6552,3317 | -0,771905791 | 0,3101223 | -2,489037 | 0,012809 | 0,0427961 | NEFM |
| ENSGALG00000026518 | 766,81567 | -0,773283921 | 0,178411 | -4,334283 | 1,46E-05 | 0,000144 | RUNDC3A |
| ENSGALG00000005610 | 640,25249 | -0,773621314 | 0,242648 | -3,188245 | 0,0014314 | 0,0071802 | SLC44A3 |

| | | | | | | | |
|---|---|---|---|---|---|---|---|
| ENSGALG00000004594 | 1038,5627 | -0,774792529 | 0,2089107 | -3,708727 | 0,0002083 | 0,0014145 | CD74 |
| ENSGALG00000013627 | 453,51839 | -0,775913381 | 0,2108029 | -3,680753 | 0,0002325 | 0,0015558 | SLC7A2 |
| ENSGALG00000015887 | 12708,103 | -0,776552733 | 0,129241 | -6,008564 | 1,87E-09 | 4,28E-08 | URB1 |
| ENSGALG00000015208 | 6752,9307 | -0,778489641 | 0,1611505 | -4,830824 | 1,36E-06 | 1,72E-05 | ROR2 |
| ENSGALG00000006258 | 9759,6638 | -0,778635815 | 0,1224981 | -6,356312 | 2,07E-10 | 5,40E-09 | SPOCK1 |
| ENSGALG00000009514 | 5928,6059 | -0,779501664 | 0,1147864 | -6,790888 | 1,11E-11 | 3,50E-10 | NRCAM |
| ENSGALG00000001020 | 7177,3339 | -0,780112538 | 0,1084839 | -7,191044 | 6,43E-13 | 2,41E-11 | KIFC3 |
| ENSGALG00000013683 | 18236,905 | -0,780228434 | 0,1531512 | -5,094499 | 3,50E-07 | 5,02E-06 | ARID1B |
| ENSGALG00000008961 | 32098,143 | -0,780468343 | 0,1668861 | -4,676652 | 2,92E-06 | 3,39E-05 | PTPRZ1 |
| ENSGALG00000012834 | 693,3013 | -0,78090617 | 0,243104 | -3,212231 | 0,0013171 | 0,0067111 | AKR1D1 |
| ENSGALG00000024964 | 133,81844 | -0,781593119 | 0,3102875 | -2,518932 | 0,0117711 | 0,0399966 | |
| ENSGALG00000009393 | 806,58478 | -0,782303357 | 0,1913491 | -4,088356 | 4,34E-05 | 0,0003633 | CAV1 |
| ENSGALG00000012329 | 10571,297 | -0,782765701 | 0,1633862 | -4,790892 | 1,66E-06 | 2,05E-05 | GLI3 |
| ENSGALG00000017864 | 140,93672 | -0,783664392 | 0,2920232 | -2,683569 | 0,0072841 | 0,0272978 | |
| ENSGALG00000026970 | 1108,6536 | -0,785311616 | 0,1563721 | -5,022071 | 5,11E-07 | 7,09E-06 | |
| ENSGALG00000009252 | 7419,3312 | -0,788959505 | 0,1242797 | -6,348257 | 2,18E-10 | 5,65E-09 | PLD1 |
| ENSGALG00000012808 | 5957,06 | -0,789210521 | 0,1360454 | -5,801083 | 6,59E-09 | 1,35E-07 | CDYL |
| ENSGALG00000006908 | 14243,327 | -0,789724754 | 0,1486312 | -5,313318 | 1,08E-07 | 1,72E-06 | DLG1 |
| ENSGALG00000000639 | 611,51653 | -0,79001541 | 0,2261059 | -3,494006 | 0,0004758 | 0,0028767 | LRRN2 |
| ENSGALG00000008411 | 91,4625 | -0,790605178 | 0,3210824 | -2,462313 | 0,0138044 | 0,0452558 | GHRL |
| ENSGALG00000016412 | 2161,4314 | -0,791360693 | 0,1346388 | -5,877656 | 4,16E-09 | 8,94E-08 | MBOAT2 |
| ENSGALG00000005785 | 2650,4252 | -0,791416097 | 0,1851664 | -4,274081 | 1,92E-05 | 0,0001807 | UBAP2 |
| ENSGALG00000009428 | 6528,6197 | -0,792261942 | 0,1416583 | -5,592767 | 2,23E-08 | 4,11E-07 | SUSD6 |
| ENSGALG00000004662 | 3631,9487 | -0,792400882 | 0,1664941 | -4,759334 | 1,94E-06 | 2,35E-05 | DISP3 |
| ENSGALG00000001995 | 9089,1491 | -0,794307619 | 0,1100185 | -7,219767 | 5,21E-13 | 1,99E-11 | LIMS2 |
| ENSGALG00000012317 | 261,45531 | -0,794799886 | 0,2552487 | -3,113826 | 0,0018468 | 0,008807 | ADGB |
| ENSGALG00000003470 | 221,25413 | -0,795180239 | 0,2878091 | -2,762873 | 0,0057295 | 0,0223837 | ZMAT4 |
| ENSGALG00000025464 | 234,06627 | -0,795229903 | 0,2531202 | -3,141708 | 0,0016797 | 0,0081615 | |
| ENSGALG00000009808 | 209,51024 | -0,795296038 | 0,2773049 | -2,867948 | 0,0041314 | 0,0171336 | AVPR1A |
| ENSGALG00000012178 | 117,35655 | -0,79634844 | 0,3097478 | -2,570958 | 0,0101418 | 0,0355602 | PDGFB |
| ENSGALG00000004132 | 38489,958 | -0,797075856 | 0,1145646 | -6,957436 | 3,47E-12 | 1,17E-10 | UHRF1 |
| ENSGALG00000026486 | 578,46167 | -0,797913674 | 0,2123431 | -3,757662 | 0,0001715 | 0,0012006 | |
| ENSGALG00000008378 | 25019,971 | -0,798096236 | 0,1224811 | -6,516075 | 7,22E-11 | 2,01E-09 | SRGAP3 |
| ENSGALG00000009932 | 6037,7104 | -0,798314026 | 0,1391751 | -5,736039 | 9,69E-09 | 1,92E-07 | THADA |
| ENSGALG00000011045 | 4900,9878 | -0,799075807 | 0,1382254 | -5,780962 | 7,43E-09 | 1,50E-07 | AFF1 |
| ENSGALG00000014907 | 2917,0076 | -0,80019622 | 0,1504507 | -5,318662 | 1,05E-07 | 1,68E-06 | DCBLD1 |
| ENSGALG00000010910 | 3629,9835 | -0,801671392 | 0,1211047 | -6,619657 | 3,60E-11 | 1,04E-09 | RAPGEF5 |
| ENSGALG00000012576 | 1359,5745 | -0,801916632 | 0,1553997 | -5,160349 | 2,46E-07 | 3,65E-06 | RASEF |
| ENSGALG00000000583 | 9340,9356 | -0,802342978 | 0,1195873 | -6,709264 | 1,96E-11 | 5,94E-10 | SOX13 |
| ENSGALG00000017044 | 315,56866 | -0,803518305 | 0,2643771 | -3,039289 | 0,0023714 | 0,0108489 | TRPC4 |
| ENSGALG00000002900 | 2509,7754 | -0,803716995 | 0,1458872 | -5,509169 | 3,61E-08 | 6,34E-07 | NOTUM |
| ENSGALG00000027521 | 307,65741 | -0,803725098 | 0,23936 | -3,357809 | 0,0007856 | 0,0043909 | ANKRD33B |
| ENSGALG00000013212 | 21085,124 | -0,805085082 | 0,1156431 | -6,961806 | 3,36E-12 | 1,14E-10 | TMEFF1 |
| ENSGALG00000001883 | 2479,1556 | -0,805297687 | 0,1269406 | -6,343891 | 2,24E-10 | 5,80E-09 | ST6GALNAC2 |
| ENSGALG00000003105 | 820,42554 | -0,805784417 | 0,1904502 | -4,230946 | 2,33E-05 | 0,0002134 | ANKFN1 |
| ENSGALG00000002461 | 9290,0239 | -0,808165073 | 0,1405519 | -5,749941 | 8,93E-09 | 1,77E-07 | MMP17 |
| ENSGALG00000002242 | 344,64399 | -0,808545932 | 0,229428 | -3,524182 | 0,0004248 | 0,0026217 | GALNT9 |
| ENSGALG00000017103 | 2337,4348 | -0,811501513 | 0,1279991 | -6,339902 | 2,30E-10 | 5,93E-09 | WASF3 |
| ENSGALG00000014957 | 24709,132 | -0,81285735 | 0,1124851 | -7,226354 | 4,96E-13 | 1,90E-11 | WDR1 |
| ENSGALG00000003427 | 8550,6609 | -0,813323099 | 0,1234103 | -6,590399 | 4,39E-11 | 1,25E-09 | USP3 |
| ENSGALG00000001136 | 15067,474 | -0,813551169 | 0,3048155 | -2,668996 | 0,0076078 | 0,0283008 | |
| ENSGALG00000011089 | 415,15739 | -0,814043658 | 0,2251011 | -3,616347 | 0,0002988 | 0,0019353 | IFIH1 |
| ENSGALG00000011643 | 20679,582 | -0,816325927 | 0,1468695 | -5,558172 | 2,73E-08 | 4,94E-07 | UTP20 |
| ENSGALG00000010785 | 498,93848 | -0,818155142 | 0,1993699 | -4,103705 | 4,07E-05 | 0,0003453 | ACOT11 |
| ENSGALG00000001220 | 3875,7891 | -0,818155857 | 0,1299079 | -6,297969 | 3,02E-10 | 7,62E-09 | HERPUD1 |
| ENSGALG00000002598 | 7963,9328 | -0,82047281 | 0,1085944 | -7,555386 | 4,18E-14 | 1,82E-12 | TPST1 |
| ENSGALG00000011511 | 101055,81 | -0,820640197 | 0,126541 | -6,485172 | 8,86E-11 | 2,44E-09 | CKB |
| ENSGALG00000003160 | 1417,9918 | -0,823423786 | 0,1758829 | -4,68166 | 2,85E-06 | 3,32E-05 | CERS1 |
| ENSGALG00000026370 | 123,7781 | -0,823508807 | 0,3145625 | -2,61795 | 0,008846 | 0,0318788 | |
| ENSGALG00000006872 | 8998,3245 | -0,823650254 | 0,1148886 | -7,169119 | 7,55E-13 | 2,79E-11 | PISD |
| ENSGALG00000007039 | 11318,955 | -0,823801989 | 0,1338606 | -6,154178 | 7,55E-10 | 1,82E-08 | IGF1R |
| ENSGALG00000004569 | 2222,4465 | -0,82424563 | 0,1654621 | -4,981477 | 6,31E-07 | 8,53E-06 | UNC5B |
| ENSGALG00000010507 | 1443,24 | -0,824303692 | 0,1501991 | -5,488074 | 4,06E-08 | 7,08E-07 | FCRL2 |
| ENSGALG00000011623 | 9810,9245 | -0,826439103 | 0,3226307 | -2,561564 | 0,0104202 | 0,0363586 | ADAMTS3 |

| | | | | | | | |
|---|---|---|---|---|---|---|---|
| ENSGALG00000003391 | 10590,452 | -0,827807721 | 0,1094916 | -7,56047 | 4,02E-14 | 1,76E-12 | EIF2AK1 |
| ENSGALG00000017855 | 167,53221 | -0,830680639 | 0,3021834 | -2,748929 | 0,005979 | 0,0232054 | |
| ENSGALG00000009546 | 1643,0136 | -0,832251183 | 0,1524822 | -5,45802 | 4,81E-08 | 8,27E-07 | HTRA1 |
| ENSGALG00000007717 | 242,86019 | -0,834765713 | 0,2552035 | -3,270981 | 0,0010718 | 0,0056624 | GCNT7 |
| ENSGALG00000014954 | 7661,4102 | -0,835213371 | 0,1158293 | -7,210726 | 5,57E-13 | 2,10E-11 | ZNF518B |
| ENSGALG00000008469 | 4008,4217 | -0,83875822 | 0,1497579 | -5,600763 | 2,13E-08 | 3,94E-07 | SPINT1 |
| ENSGALG00000002543 | 5887,0903 | -0,840502021 | 0,1240047 | -6,777985 | 1,22E-11 | 3,81E-10 | CSGALNACT2 |
| ENSGALG00000015980 | 410,78574 | -0,841274637 | 0,2500328 | -3,364658 | 0,0007664 | 0,004294 | |
| ENSGALG00000011388 | 12926,485 | -0,842306194 | 0,1210688 | -6,957254 | 3,47E-12 | 1,17E-10 | RABGGTB |
| ENSGALG00000004627 | 119,19825 | -0,843337638 | 0,3097612 | -2,722542 | 0,0064782 | 0,0248312 | LAMC2 |
| ENSGALG00000015519 | 4471,9334 | -0,844394446 | 0,1245073 | -6,781886 | 1,19E-11 | 3,71E-10 | ROBO2 |
| ENSGALG00000015656 | 12813,534 | -0,849003497 | 0,1245832 | -6,814749 | 9,44E-12 | 3,00E-10 | AKAP2 |
| ENSGALG00000012998 | 2617,3459 | -0,849149352 | 0,1768308 | -4,802043 | 1,57E-06 | 1,96E-05 | WNT5B |
| ENSGALG00000024984 | 247,00772 | -0,849437052 | 0,2463411 | -3,448214 | 0,0005643 | 0,0033217 | |
| ENSGALG00000023609 | 5841,3037 | -0,855145185 | 0,1351075 | -6,329368 | 2,46E-10 | 6,31E-09 | |
| ENSGALG00000014178 | 3963,5155 | -0,855544377 | 0,1502564 | -5,693895 | 1,24E-08 | 2,41E-07 | SCUBE1 |
| ENSGALG00000012394 | 13948,867 | -0,855781498 | 0,1036166 | -8,259118 | 1,47E-16 | 8,54E-15 | NID2 |
| ENSGALG00000012873 | 33,50313 | -0,858492519 | 0,3501442 | -2,451826 | 0,0142133 | 0,0463619 | SERPINB5 |
| ENSGALG00000003668 | 1594,4405 | -0,85921128 | 0,1422579 | -6,039814 | 1,54E-09 | 3,57E-08 | PITPNC1 |
| ENSGALG00000007846 | 1019,2888 | -0,863598394 | 0,2308913 | -3,740281 | 0,0001838 | 0,0012722 | |
| ENSGALG00000010311 | 430,97249 | -0,865329105 | 0,2059571 | -4,201502 | 2,65E-05 | 0,0002391 | NAV3 |
| ENSGALG00000015681 | 7286,4362 | -0,869123123 | 0,1385988 | -6,270786 | 3,59E-10 | 9,02E-09 | NRIP1 |
| ENSGALG00000010010 | 1250,7492 | -0,869227214 | 0,1534764 | -5,66359 | 1,48E-08 | 2,84E-07 | RHOQ |
| ENSGALG00000008815 | 1450,4816 | -0,86979829 | 0,1663609 | -5,22838 | 1,71E-07 | 2,63E-06 | LRRN4 |
| ENSGALG00000011376 | 402,72686 | -0,870171475 | 0,2115735 | -4,112858 | 3,91E-05 | 0,0003342 | ANKRD9 |
| ENSGALG00000020703 | 83,715675 | -0,87414394 | 0,3266079 | -2,676432 | 0,0074411 | 0,0277943 | GRB14 |
| ENSGALG00000016896 | 2175,6547 | -0,874558686 | 0,1602235 | -5,458368 | 4,81E-08 | 8,26E-07 | ABCC4 |
| ENSGALG00000016981 | 36,264584 | -0,876405769 | 0,3539545 | -2,476041 | 0,0132848 | 0,0439973 | SPERT |
| ENSGALG00000023581 | 218,88299 | -0,876769713 | 0,2786993 | -3,145934 | 0,0016556 | 0,0080646 | |
| ENSGALG00000009129 | 1209,8208 | -0,878296092 | 0,1715571 | -5,119557 | 3,06E-07 | 4,43E-06 | DLX5 |
| ENSGALG00000004542 | 7447,8359 | -0,879244031 | 0,1445917 | -6,080875 | 1,20E-09 | 2,81E-08 | ZMYND8 |
| ENSGALG00000016462 | 9552,6988 | -0,880157248 | 0,1399598 | -6,288642 | 3,20E-10 | 8,07E-09 | MYCN |
| ENSGALG00000028602 | 5443,1954 | -0,881510014 | 0,1576328 | -5,592172 | 2,24E-08 | 4,12E-07 | |
| ENSGALG00000011120 | 494,96982 | -0,88227849 | 0,2357487 | -3,742453 | 0,0001822 | 0,0012646 | SLC4A10 |
| ENSGALG00000003657 | 764,547 | -0,88270712 | 0,1987799 | -4,440625 | 8,97E-06 | 9,38E-05 | UNC119 |
| ENSGALG00000027633 | 124,92778 | -0,88394595 | 0,3059642 | -2,88905 | 0,0038641 | 0,0162676 | RASSF9 |
| ENSGALG00000028320 | 417,90377 | -0,88451445 | 0,2720255 | -3,251586 | 0,0011476 | 0,0059985 | |
| ENSGALG00000001749 | 4549,2907 | -0,88875485 | 0,1275791 | -6,966303 | 3,25E-12 | 1,11E-10 | ACSBG2 |
| ENSGALG00000005723 | 2016,6496 | -0,890647194 | 0,1398661 | -6,367856 | 1,92E-10 | 5,03E-09 | RP11-96O20.4 |
| ENSGALG00000009360 | 602,38543 | -0,890830846 | 0,1924613 | -4,628624 | 3,68E-06 | 4,19E-05 | CAPN2 |
| ENSGALG00000002555 | 1358,1498 | -0,892902927 | 0,2116768 | -4,218237 | 2,46E-05 | 0,0002247 | RET |
| ENSGALG00000000307 | 8913,0835 | -0,894951246 | 0,1134627 | -7,887626 | 3,08E-15 | 1,50E-13 | CERS4 |
| ENSGALG00000012012 | 5383,1455 | -0,895703636 | 0,1725569 | -5,190772 | 2,09E-07 | 3,15E-06 | CMIP |
| ENSGALG00000017870 | 493,11834 | -0,896459218 | 0,2052027 | -4,368652 | 1,25E-05 | 0,0001254 | |
| ENSGALG00000008547 | 11178,2 | -0,896850751 | 0,1262491 | -7,103817 | 1,21E-12 | 4,41E-11 | ZBED4 |
| ENSGALG00000008509 | 336,23193 | -0,897122538 | 0,2669373 | -3,360799 | 0,0007772 | 0,004349 | SEMA3E |
| ENSGALG00000012814 | 2520,1319 | -0,898069575 | 0,1279271 | -7,020165 | 2,22E-12 | 7,76E-11 | PXDC1 |
| ENSGALG00000017838 | 224,75144 | -0,898451215 | 0,2607674 | -3,445413 | 0,0005702 | 0,0033455 | |
| ENSGALG00000027269 | 123,982 | -0,898861923 | 0,3460342 | -2,59761 | 0,0093875 | 0,0333927 | GUCA2B |
| ENSGALG00000009687 | 2155,9656 | -0,89897016 | 0,1572674 | -5,71619 | 1,09E-08 | 2,13E-07 | KCNK2 |
| ENSGALG00000014645 | 469,37544 | -0,89901657 | 0,2175335 | -4,132773 | 3,58E-05 | 0,0003098 | MEF2C |
| ENSGALG00000006498 | 20254,306 | -0,899416586 | 0,1691498 | -5,317278 | 1,05E-07 | 1,69E-06 | PTPRN2 |
| ENSGALG00000013149 | 142,93297 | -0,899640208 | 0,2949853 | -3,04978 | 0,0022901 | 0,0105372 | MOCOS |
| ENSGALG00000025723 | 123,40432 | -0,903219239 | 0,3196649 | -2,825519 | 0,0047204 | 0,0191048 | NKX1-1 |
| ENSGALG00000026790 | 419,69129 | -0,903869706 | 0,2560648 | -3,529848 | 0,0004158 | 0,0025729 | DMRT2 |
| ENSGALG00000002704 | 377,64333 | -0,906514113 | 0,2414033 | -3,755185 | 0,0001732 | 0,0012112 | AFAP1L1 |
| ENSGALG00000027960 | 55,099779 | -0,908090032 | 0,3479917 | -2,609516 | 0,009067 | 0,0324691 | GRPR |
| ENSGALG00000017876 | 51,084697 | -0,908586905 | 0,3511323 | -2,587592 | 0,0096649 | 0,0342054 | |
| ENSGALG00000026258 | 289,1614 | -0,909500707 | 0,2404705 | -3,782171 | 0,0001555 | 0,0011001 | |
| ENSGALG00000004231 | 640,94875 | -0,911405982 | 0,183472 | -4,967547 | 6,78E-07 | 9,10E-06 | IFNLR1 |
| ENSGALG00000013861 | 1063,5397 | -0,913164571 | 0,1795586 | -5,085609 | 3,66E-07 | 5,22E-06 | TNFAIP3 |
| ENSGALG00000020723 | 828,74872 | -0,913543779 | 0,1755447 | -5,204053 | 1,95E-07 | 2,96E-06 | NOG2 |
| ENSGALG00000019622 | 6843,9055 | -0,918336915 | 0,3199295 | -2,870435 | 0,0040991 | 0,0170409 | TRIM71 |
| ENSGALG00000005994 | 969,59836 | -0,91862167 | 0,1776308 | -5,171523 | 2,32E-07 | 3,46E-06 | DERL3 |

| | | | | | | | |
|---|---|---|---|---|---|---|---|
| ENSGALG00000010683 | 599,9885 | -0,919156764 | 0,2019322 | -4,551809 | 5,32E-06 | 5,85E-05 | S1PR3 |
| ENSGALG00000001656 | 26148,058 | -0,922800056 | 0,1104664 | -8,35367 | 6,62E-17 | 4,00E-15 | sept-09 |
| ENSGALG00000009540 | 5865,4242 | -0,9229711 | 0,1165261 | -7,920723 | 2,36E-15 | 1,17E-13 | YAF2 |
| ENSGALG00000025166 | 165,3461 | -0,924288426 | 0,2874597 | -3,215367 | 0,0013028 | 0,0066557 | |
| ENSGALG00000001000 | 110996,8 | -0,924720639 | 0,1374659 | -6,726907 | 1,73E-11 | 5,28E-10 | HSPA5 |
| ENSGALG00000008305 | 1229,5865 | -0,925580955 | 0,1854564 | -4,990827 | 6,01E-07 | 8,18E-06 | CASP10 |
| ENSGALG00000016843 | 7409,4478 | -0,926939677 | 0,1285541 | -7,210501 | 5,57E-13 | 2,10E-11 | COL4A2 |
| ENSGALG00000023100 | 28,255969 | -0,928527718 | 0,3431483 | -2,705908 | 0,0068118 | 0,025848 | |
| ENSGALG00000014961 | 70,382782 | -0,928667888 | 0,3536337 | -2,626073 | 0,0086376 | 0,0312686 | FAM26E |
| ENSGALG00000009262 | 127,08834 | -0,928869092 | 0,3039249 | -3,056245 | 0,0022413 | 0,0103476 | FGB |
| ENSGALG00000012911 | 418,92558 | -0,929872331 | 0,2601161 | -3,574836 | 0,0003504 | 0,0022213 | SYT10 |
| ENSGALG00000008811 | 6177,8009 | -0,931412058 | 0,1489053 | -6,255065 | 3,97E-10 | 9,87E-09 | SLC1A4 |
| ENSGALG00000007952 | 17655,501 | -0,932963194 | 0,1344416 | -6,939543 | 3,93E-12 | 1,32E-10 | AMOT |
| ENSGALG00000023241 | 1184,6759 | -0,938819916 | 0,1882587 | -4,986861 | 6,14E-07 | 8,32E-06 | |
| ENSGALG00000028708 | 6182,0112 | -0,938854547 | 0,1258631 | -7,459329 | 8,70E-14 | 3,59E-12 | |
| ENSGALG00000009415 | 20765,099 | -0,939656238 | 0,1402127 | -6,701647 | 2,06E-11 | 6,21E-10 | SMOC1 |
| ENSGALG00000012726 | 95084,968 | -0,941672551 | 0,1261141 | -7,466831 | 8,21E-14 | 3,42E-12 | HSP90B1 |
| ENSGALG00000015622 | 657,0594 | -0,944352456 | 0,2173775 | -4,344298 | 1,40E-05 | 0,0001385 | EYA1 |
| ENSGALG00000015028 | 404,3337 | -0,946739996 | 0,2500725 | -3,785863 | 0,0001532 | 0,0010858 | RLN1 |
| ENSGALG00000011258 | 44996,798 | -0,946927699 | 0,1449344 | -6,533494 | 6,43E-11 | 1,81E-09 | ATP2B1 |
| ENSGALG00000011068 | 1649,8566 | -0,947751489 | 0,1445198 | -6,557937 | 5,46E-11 | 1,54E-09 | COBLL1 |
| ENSGALG00000005601 | 522,73024 | -0,948098308 | 0,2254007 | -4,206279 | 2,60E-05 | 0,0002346 | DNAH12 |
| ENSGALG00000006808 | 9236,5969 | -0,948868926 | 0,1071422 | -8,856165 | 8,28E-19 | 6,00E-17 | TSPAN6 |
| ENSGALG00000013193 | 667,61334 | -0,948945842 | 0,2164896 | -4,383332 | 1,17E-05 | 0,0001183 | IRX2 |
| ENSGALG00000023608 | 3676,9857 | -0,950348621 | 0,134551 | -7,06311 | 1,63E-12 | 5,82E-11 | |
| ENSGALG00000027260 | 44,084585 | -0,951699131 | 0,3538862 | -2,68928 | 0,0071606 | 0,0269519 | MC3R |
| ENSGALG00000005805 | 4566,7573 | -0,954718819 | 0,1455167 | -6,560889 | 5,35E-11 | 1,52E-09 | PLCD1 |
| ENSGALG00000023463 | 1598,1146 | -0,956997326 | 0,1712907 | -5,586978 | 2,31E-08 | 4,24E-07 | |
| ENSGALG00000017875 | 78,294933 | -0,958367374 | 0,3317498 | -2,888825 | 0,0038668 | 0,0162692 | |
| ENSGALG00000002577 | 53,909401 | -0,959612138 | 0,3537853 | -2,712414 | 0,0066795 | 0,025481 | STARD10 |
| ENSGALG00000004393 | 1366,8686 | -0,962441655 | 0,2126961 | -4,524962 | 6,04E-06 | 6,51E-05 | |
| ENSGALG00000004284 | 19106,512 | -0,967926178 | 0,1544238 | -6,267988 | 3,66E-10 | 9,15E-09 | LFNG |
| ENSGALG00000027342 | 11330,786 | -0,968330352 | 0,1154603 | -8,386697 | 5,00E-17 | 3,08E-15 | METRN |
| ENSGALG00000015908 | 1062,9623 | -0,970529728 | 0,2034554 | -4,770232 | 1,84E-06 | 2,24E-05 | COL12A1 |
| ENSGALG00000011833 | 13327,666 | -0,971207285 | 0,1137465 | -8,538346 | 1,36E-17 | 8,83E-16 | SPATA5 |
| ENSGALG00000009381 | 15338,368 | -0,972226453 | 0,1330298 | -7,308336 | 2,70E-13 | 1,06E-11 | SIPA1L1 |
| ENSGALG00000015940 | 2397,4582 | -0,972506334 | 0,1424398 | -6,827491 | 8,64E-12 | 2,76E-10 | TMEM67 |
| ENSGALG00000007367 | 230,09823 | -0,973082329 | 0,2646976 | -3,676204 | 0,0002367 | 0,0015799 | WFIKKN2 |
| ENSGALG00000010597 | 2446,5354 | -0,973101915 | 0,127224 | -7,64873 | 2,03E-14 | 9,27E-13 | CRIM1 |
| ENSGALG00000027497 | 292,35552 | -0,973179721 | 0,2340886 | -4,157313 | 3,22E-05 | 0,0002823 | CFD |
| ENSGALG00000019397 | 12676,413 | -0,975439675 | 0,1148854 | -8,490543 | 2,06E-17 | 1,31E-15 | CERK |
| ENSGALG00000026995 | 97,429549 | -0,97623355 | 0,3272256 | -2,983365 | 0,002851 | 0,0126313 | RHOG |
| ENSGALG00000002103 | 289,28254 | -0,979001464 | 0,2861261 | -3,421574 | 0,0006226 | 0,0035994 | TPPP3 |
| ENSGALG00000015624 | 34457,429 | -0,97910183 | 0,1758979 | -5,566309 | 2,60E-08 | 4,72E-07 | VCAN |
| ENSGALG00000012418 | 35,255982 | -0,979401863 | 0,3538633 | -2,767741 | 0,0056446 | 0,0221474 | KYNU |
| ENSGALG00000015164 | 3228,2862 | -0,982829392 | 0,1480222 | -6,639743 | 3,14E-11 | 9,24E-10 | PCSK5 |
| ENSGALG00000020788 | 1411,7356 | -0,986044977 | 0,148882 | -6,622998 | 3,52E-11 | 1,02E-09 | CCDC141 |
| ENSGALG00000001371 | 2323,3283 | -0,990350626 | 0,1647412 | -6,011554 | 1,84E-09 | 4,21E-08 | |
| ENSGALG00000009758 | 793,28474 | -0,991410492 | 0,1927041 | -5,144731 | 2,68E-07 | 3,94E-06 | SLC7A11 |
| ENSGALG00000015914 | 35,887211 | -0,993778818 | 0,3538558 | -2,808429 | 0,0049784 | 0,0200099 | CALB1 |
| ENSGALG00000015440 | 170,65631 | -0,996123586 | 0,2860343 | -3,482532 | 0,0004967 | 0,0029843 | RCSD1 |
| ENSGALG00000010692 | 6955,6689 | -0,996294098 | 0,1389198 | -7,171722 | 7,41E-13 | 2,76E-11 | LRP8 |
| ENSGALG00000026582 | 1194,487 | -0,997100015 | 0,1907735 | -5,226618 | 1,73E-07 | 2,64E-06 | SOX18 |
| ENSGALG00000002532 | 116,30592 | -0,997428466 | 0,3168923 | -3,147531 | 0,0016466 | 0,0080322 | GJA4 |
| ENSGALG00000010163 | 10928,524 | -0,998496131 | 0,1269077 | -7,867891 | 3,61E-15 | 1,74E-13 | LGR5 |
| ENSGALG00000007483 | 18280,16 | -0,999963194 | 0,1499281 | -6,669618 | 2,56E-11 | 7,61E-10 | LRIG1 |
| ENSGALG00000005483 | 1190,7719 | -1,005472997 | 0,1552955 | -6,474578 | 9,51E-11 | 2,59E-09 | CDH13 |
| ENSGALG00000012454 | 37,219552 | -1,005696393 | 0,3507698 | -2,867112 | 0,0041424 | 0,017158 | CYTH4 |
| ENSGALG00000011295 | 1389,1449 | -1,007421144 | 0,1580798 | -6,372863 | 1,86E-10 | 4,90E-09 | SOCS2 |
| ENSGALG00000016627 | 15863,827 | -1,008110823 | 0,1082287 | -9,314637 | 1,22E-20 | 1,03E-18 | FZD3 |
| ENSGALG00000008281 | 355,49596 | -1,009962725 | 0,2355243 | -4,288147 | 1,80E-05 | 0,0001708 | NEURL1 |
| ENSGALG00000015078 | 5440,7576 | -1,010765666 | 0,1823319 | -5,543548 | 2,96E-08 | 5,32E-07 | MLLT3 |
| ENSGALG00000015112 | 8576,8627 | -1,013051509 | 0,1332664 | -7,601703 | 2,92E-14 | 1,31E-12 | ZNF521 |
| ENSGALG00000000949 | 297,0835 | -1,013784064 | 0,2329319 | -4,352276 | 1,35E-05 | 0,0001336 | HBEGF |

| | | | | | | | |
|---|---|---|---|---|---|---|---|
| ENSGALG00000016616 | 1435,9249 | -1,01551489 | 0,1675974 | -6,059252 | 1,37E-09 | 3,18E-08 | ANOS1 |
| ENSGALG00000009450 | 2394,0442 | -1,016064164 | 0,1296479 | -7,837106 | 4,61E-15 | 2,19E-13 | DUSP10 |
| ENSGALG00000001009 | 527,81732 | -1,018620159 | 0,2467658 | -4,127883 | 3,66E-05 | 0,0003151 | |
| ENSGALG00000015896 | 2100,3865 | -1,023057651 | 0,1679504 | -6,091426 | 1,12E-09 | 2,64E-08 | MMP16 |
| ENSGALG00000010156 | 274,63907 | -1,025083823 | 0,2479625 | -4,134028 | 3,56E-05 | 0,0003089 | DOCK8 |
| ENSGALG00000013124 | 3974,9034 | -1,026819624 | 0,1623011 | -6,326635 | 2,51E-10 | 6,40E-09 | FHOD3 |
| ENSGALG00000006138 | 2396,0525 | -1,026936244 | 0,150452 | -6,825674 | 8,75E-12 | 2,79E-10 | HELZ2 |
| ENSGALG00000010579 | 4573,0234 | -1,028169899 | 0,1257094 | -8,178939 | 2,86E-16 | 1,62E-14 | STON2 |
| ENSGALG00000012439 | 1314,5759 | -1,028638845 | 0,2279249 | -4,513061 | 6,39E-06 | 6,86E-05 | IGFBP3 |
| ENSGALG00000019077 | 11786,668 | -1,030078317 | 0,1385218 | -7,436218 | 1,04E-13 | 4,25E-12 | |
| ENSGALG00000012522 | 26,038382 | -1,03063871 | 0,345927 | -2,979353 | 0,0028886 | 0,0127647 | PVALB |
| ENSGALG00000000977 | 7324,7206 | -1,032464943 | 0,1097312 | -9,409039 | 5,01E-21 | 4,29E-19 | MAPKAP1 |
| ENSGALG00000016754 | 1717,8597 | -1,034769759 | 0,1770332 | -5,845059 | 5,06E-09 | 1,06E-07 | MGAT4A |
| ENSGALG00000012412 | 7272,9925 | -1,036824012 | 0,1452529 | -7,138059 | 9,47E-13 | 3,47E-11 | MTHFD1L |
| ENSGALG00000027213 | 84,217699 | -1,037130822 | 0,3364624 | -3,082456 | 0,002053 | 0,0096518 | ALX1 |
| ENSGALG00000007555 | 13407,29 | -1,038877738 | 0,1147853 | -9,050617 | 1,42E-19 | 1,11E-17 | CCND1 |
| ENSGALG00000025284 | 64,080075 | -1,04409147 | 0,3409884 | -3,061956 | 0,002199 | 0,0102076 | |
| ENSGALG00000008465 | 8001,1338 | -1,045042375 | 0,1298809 | -8,046158 | 8,54E-16 | 4,60E-14 | SORCS1 |
| ENSGALG00000022800 | 1489,665 | -1,045072381 | 0,145669 | -7,174293 | 7,27E-13 | 2,72E-11 | KCNB2 |
| ENSGALG00000011336 | 49,782944 | -1,048887527 | 0,3535455 | -2,966768 | 0,0030095 | 0,0132004 | TECRL |
| ENSGALG00000015008 | 37,265424 | -1,049053854 | 0,3418003 | -3,069201 | 0,0021463 | 0,0100076 | ZNF366 |
| ENSGALG00000023355 | 72,955518 | -1,050070076 | 0,34512 | -3,042623 | 0,0023453 | 0,0107547 | TMEM154 |
| ENSGALG00000017714 | 665,00414 | -1,052113718 | 0,2083331 | -5,050151 | 4,41E-07 | 6,20E-06 | FHAD1 |
| ENSGALG00000015975 | 5069,4191 | -1,056406602 | 0,1284565 | -8,223848 | 1,97E-16 | 1,13E-14 | DPY19L4 |
| ENSGALG00000011581 | 10286,837 | -1,059360643 | 0,338584 | -3,128797 | 0,0017552 | 0,0084616 | KIF26A |
| ENSGALG00000015736 | 279,62808 | -1,060236168 | 0,3192341 | -3,321187 | 0,0008964 | 0,0048849 | STMN2 |
| ENSGALG00000003295 | 2261,7027 | -1,063705711 | 0,1585827 | -6,707576 | 1,98E-11 | 6,00E-10 | ADAMTS12 |
| ENSGALG00000009740 | 783,87139 | -1,063900656 | 0,1815827 | -5,859043 | 4,66E-09 | 9,86E-08 | RASGRP1 |
| ENSGALG00000013604 | 164,1582 | -1,065244459 | 0,2852669 | -3,734202 | 0,0001883 | 0,0012975 | VIP |
| ENSGALG00000016470 | 856,35527 | -1,066533888 | 0,3521055 | -3,029018 | 0,0024535 | 0,0111534 | KCNS3 |
| ENSGALG00000011236 | 190,29168 | -1,070413941 | 0,315055 | -3,397546 | 0,0006799 | 0,0038831 | BEGAIN |
| ENSGALG00000017135 | 1406,4237 | -1,071209509 | 0,1612008 | -6,645186 | 3,03E-11 | 8,92E-10 | CRYL1 |
| ENSGALG00000002988 | 7561,0439 | -1,072357906 | 0,152099 | -7,050395 | 1,78E-12 | 6,33E-11 | PHGDH |
| ENSGALG00000001370 | 1171,2311 | -1,073396904 | 0,1915947 | -5,602436 | 2,11E-08 | 3,91E-07 | ADAMTS8 |
| ENSGALG00000010707 | 29,836376 | -1,074169362 | 0,351316 | -3,057559 | 0,0022315 | 0,0103145 | GPR68 |
| ENSGALG00000027757 | 10827,926 | -1,076478944 | 0,1080166 | -9,965866 | 2,15E-23 | 2,27E-21 | ARL4A |
| ENSGALG00000010891 | 1573,4126 | -1,079203445 | 0,1845381 | -5,848133 | 4,97E-09 | 1,04E-07 | ABCB11 |
| ENSGALG00000027707 | 112,77294 | -1,082314729 | 0,3118106 | -3,471064 | 0,0005184 | 0,0030902 | ESM1 |
| ENSGALG00000016899 | 32,641313 | -1,08275528 | 0,3526736 | -3,070135 | 0,0021396 | 0,0099866 | DCT |
| ENSGALG00000000177 | 4145,0213 | -1,083701806 | 0,1219668 | -8,885217 | 6,38E-19 | 4,70E-17 | KLF3 |
| ENSGALG00000025770 | 345,96122 | -1,089960517 | 0,2525089 | -4,316523 | 1,59E-05 | 0,0001532 | |
| ENSGALG00000028046 | 6843,8734 | -1,098357134 | 0,1139847 | -9,636006 | 5,63E-22 | 5,11E-20 | INSIG2 |
| ENSGALG00000012917 | 4548,5783 | -1,102419391 | 0,1907106 | -5,780587 | 7,44E-09 | 1,50E-07 | CDH6 |
| ENSGALG00000005842 | 443,93623 | -1,104947048 | 0,2118757 | -5,215072 | 1,84E-07 | 2,80E-06 | GDPD2 |
| ENSGALG00000012095 | 202,10968 | -1,108243378 | 0,2716377 | -4,079858 | 4,51E-05 | 0,0003753 | C14orf105 |
| ENSGALG00000002518 | 128,6744 | -1,109129449 | 0,3369249 | -3,291919 | 0,0009951 | 0,0053108 | |
| ENSGALG00000009567 | 5248,5052 | -1,113206952 | 0,1654659 | -6,727712 | 1,72E-11 | 5,27E-10 | TLL1 |
| ENSGALG00000011439 | 1241,3905 | -1,116853678 | 0,1622406 | -6,883936 | 5,82E-12 | 1,90E-10 | RBMS3 |
| ENSGALG00000013548 | 35,231247 | -1,118469658 | 0,3386376 | -3,302851 | 0,0009571 | 0,0051565 | GZMA |
| ENSGALG00000014852 | 15743,642 | -1,119019515 | 0,1113638 | -10,04833 | 9,34E-24 | 1,05E-21 | |
| ENSGALG00000009179 | 4347,973 | -1,119360692 | 0,1337923 | -8,366405 | 5,94E-17 | 3,63E-15 | TNFSF10 |
| ENSGALG00000011172 | 2541,5225 | -1,121518184 | 0,1274182 | -8,801871 | 1,35E-18 | 9,35E-17 | MYO10L |
| ENSGALG00000013892 | 1340,4062 | -1,122648268 | 0,1612643 | -6,961542 | 3,37E-12 | 1,14E-10 | MAP3K5 |
| ENSGALG00000001079 | 117,89095 | -1,122757301 | 0,3356009 | -3,345513 | 0,0008213 | 0,0045345 | WNT3 |
| ENSGALG00000016680 | 793,76813 | -1,12428894 | 0,234202 | -4,80051 | 1,58E-06 | 1,97E-05 | TFAP2B |
| ENSGALG00000008014 | 284,58059 | -1,126068911 | 0,2641663 | -4,262727 | 2,02E-05 | 0,0001892 | CEBPB |
| ENSGALG00000006337 | 35,98151 | -1,128489411 | 0,3491935 | -3,231702 | 0,0012306 | 0,0063486 | K123 |
| ENSGALG00000020151 | 65,103733 | -1,129234483 | 0,3522369 | -3,205895 | 0,0013464 | 0,0068343 | NMU |
| ENSGALG00000006376 | 107,4182 | -1,138122247 | 0,3273909 | -3,476341 | 0,0005083 | 0,0030393 | TMC3 |
| ENSGALG00000027210 | 288,32435 | -1,142415301 | 0,2380637 | -4,798779 | 1,60E-06 | 1,98E-05 | |
| ENSGALG00000001085 | 2988,7865 | -1,143093103 | 0,1311931 | -8,71306 | 2,96E-18 | 1,98E-16 | ADGRG1 |
| ENSGALG00000001276 | 18773,518 | -1,144775826 | 0,131613 | -8,698048 | 3,38E-18 | 2,24E-16 | HOXB9 |
| ENSGALG00000004155 | 218,30979 | -1,147734088 | 0,2762334 | -4,154943 | 3,25E-05 | 0,0002845 | MYOM3 |
| ENSGALG00000027107 | 1206,9544 | -1,149521147 | 0,1567788 | -7,332123 | 2,27E-13 | 9,02E-12 | PROCA1 |

| | | | | | | | |
|---|---|---|---|---|---|---|---|
| ENSGALG00000014864 | 769,88057 | -1,150010609 | 0,170127 | -6,759718 | 1,38E-11 | 4,28E-10 | SMPDL3A |
| ENSGALG00000027885 | 31,983561 | -1,151437329 | 0,3494378 | -3,295114 | 0,0009838 | 0,0052611 | C4orf19 |
| ENSGALG00000010926 | 145,65 | -1,152009571 | 0,3159768 | -3,645868 | 0,0002665 | 0,0017503 | SPP1 |
| ENSGALG00000008875 | 1752,669 | -1,156831558 | 0,1589922 | -7,276029 | 3,44E-13 | 1,34E-11 | PLCB1 |
| ENSGALG00000007268 | 3702,3293 | -1,168892699 | 0,1417063 | -8,248697 | 1,60E-16 | 9,28E-15 | CADPS |
| ENSGALG00000006584 | 535,75964 | -1,169466644 | 0,2313724 | -5,054478 | 4,32E-07 | 6,07E-06 | |
| ENSGALG00000016114 | 1343,2019 | -1,174926553 | 0,198016 | -5,933492 | 2,97E-09 | 6,53E-08 | TNFRSF11B |
| ENSGALG00000022871 | 818,02057 | -1,176605905 | 0,1867706 | -6,299737 | 2,98E-10 | 7,55E-09 | HHLA2 |
| ENSGALG00000026763 | 724,90274 | -1,177313255 | 0,2069483 | -5,688924 | 1,28E-08 | 2,48E-07 | |
| ENSGALG00000017411 | 73,326206 | -1,17758282 | 0,3486458 | -3,377591 | 0,0007312 | 0,0041242 | RXFP3 |
| ENSGALG00000016998 | 449,62073 | -1,177970615 | 0,2144018 | -5,494221 | 3,92E-08 | 6,86E-07 | |
| ENSGALG00000008941 | 29,808839 | -1,184874519 | 0,3508426 | -3,377225 | 0,0007322 | 0,0041263 | FEZF1 |
| ENSGALG00000002509 | 14719,435 | -1,185807324 | 0,1326958 | -8,936281 | 4,02E-19 | 2,98E-17 | COL5A2 |
| ENSGALG00000012357 | 5240,5441 | -1,187472008 | 0,1225271 | -9,691503 | 3,28E-22 | 3,16E-20 | CXCR4 |
| ENSGALG00000015665 | 1052,1425 | -1,188036426 | 0,1715856 | -6,923872 | 4,39E-12 | 1,45E-10 | SH3BP2 |
| ENSGALG00000005619 | 863,25589 | -1,189698364 | 0,2239703 | -5,311858 | 1,09E-07 | 1,73E-06 | F3 |
| ENSGALG00000005685 | 3302,1261 | -1,189953127 | 0,1570606 | -7,576394 | 3,55E-14 | 1,57E-12 | KSR1 |
| ENSGALG00000016083 | 216,72251 | -1,190269979 | 0,2711704 | -4,38938 | 1,14E-05 | 0,0001155 | ANGPT1 |
| ENSGALG00000000936 | 1054,1306 | -1,193444435 | 0,1573416 | -7,585055 | 3,32E-14 | 1,47E-12 | LMX1B |
| ENSGALG00000006170 | 3710,8245 | -1,194070483 | 0,1488786 | -8,020428 | 1,05E-15 | 5,52E-14 | LRRC8C |
| ENSGALG00000015433 | 4115,112 | -1,199115509 | 0,1254233 | -9,560547 | 1,17E-21 | 1,05E-19 | ABCA1 |
| ENSGALG00000002923 | 1146,0005 | -1,199213505 | 0,1576913 | -7,60482 | 2,85E-14 | 1,29E-12 | |
| ENSGALG00000011104 | 29,588884 | -1,203717219 | 0,3516022 | -3,42352 | 0,0006182 | 0,0035829 | GCG |
| ENSGALG00000012224 | 1928,9682 | -1,20528636 | 0,1513933 | -7,961295 | 1,70E-15 | 8,65E-14 | PDE1C |
| ENSGALG00000003582 | 6407,4212 | -1,218636528 | 0,1784672 | -6,828349 | 8,59E-12 | 2,75E-10 | SLC1A3 |
| ENSGALG00000009667 | 660,39309 | -1,219318734 | 0,2495248 | -4,886564 | 1,03E-06 | 1,33E-05 | MFAP3L |
| ENSGALG00000002632 | 1307,7165 | -1,220611399 | 0,1739574 | -7,016727 | 2,27E-12 | 7,93E-11 | CALCRL |
| ENSGALG00000009215 | 122,78745 | -1,220936874 | 0,3057355 | -3,993442 | 6,51E-05 | 0,0005167 | PDE11A |
| ENSGALG00000000141 | 479,97527 | -1,223136562 | 0,2587692 | -4,726747 | 2,28E-06 | 2,72E-05 | BLB1 |
| ENSGALG00000006473 | 340,04191 | -1,223226887 | 0,2654847 | -4,607524 | 4,07E-06 | 4,58E-05 | PLXNA4B |
| ENSGALG00000007660 | 3604,7946 | -1,228207524 | 0,1637881 | -7,49876 | 6,44E-14 | 2,69E-12 | CHIC1 |
| ENSGALG00000013239 | 67578,529 | -1,231185989 | 0,1344215 | -9,159144 | 5,23E-20 | 4,24E-18 | NES |
| ENSGALG00000008615 | 1056,6531 | -1,233824562 | 0,1860614 | -6,631275 | 3,33E-11 | 9,74E-10 | LTK |
| ENSGALG00000003739 | 39854,675 | -1,237172502 | 0,1212391 | -10,2044 | 1,89E-24 | 2,28E-22 | SALL1 |
| ENSGALG00000029023 | 187,05762 | -1,242039059 | 0,2692492 | -4,612972 | 3,97E-06 | 4,48E-05 | DMRTB1 |
| ENSGALG00000009532 | 9821,7952 | -1,242374232 | 0,113576 | -10,9387 | 7,53E-28 | 1,13E-25 | GXYLT1 |
| ENSGALG00000011867 | 2059,3195 | -1,242989678 | 0,1421856 | -8,742025 | 2,29E-18 | 1,56E-16 | SERHL2 |
| ENSGALG00000005427 | 736,09619 | -1,25622691 | 0,1850037 | -6,790278 | 1,12E-11 | 3,51E-10 | |
| ENSGALG00000013237 | 693,51796 | -1,260226776 | 0,2152859 | -5,853736 | 4,81E-09 | 1,01E-07 | BCAN |
| ENSGALG00000010858 | 12929,131 | -1,268608971 | 0,3211828 | -3,949803 | 7,82E-05 | 0,0006051 | LRP2 |
| ENSGALG00000003353 | 12512,084 | -1,269046359 | 0,175098 | -7,247634 | 4,24E-13 | 1,63E-11 | RAI14 |
| ENSGALG00000019552 | 24,982545 | -1,269295425 | 0,3440776 | -3,688981 | 0,0002252 | 0,0015146 | SERPINB2 |
| ENSGALG00000004361 | 87800,454 | -1,276561341 | 0,1268377 | -10,06452 | 7,93E-24 | 8,99E-22 | PRTG |
| ENSGALG00000008352 | 151,36281 | -1,283975751 | 0,2919953 | -4,397248 | 1,10E-05 | 0,0001121 | CKMT1A |
| ENSGALG00000002397 | 837,30231 | -1,291814757 | 0,2176946 | -5,934069 | 2,96E-09 | 6,52E-08 | PSPH |
| ENSGALG00000006401 | 4187,8898 | -1,299022844 | 0,1294305 | -10,03645 | 1,05E-23 | 1,16E-21 | OSBPL5 |
| ENSGALG00000014866 | 52452,789 | -1,312013425 | 0,1313405 | -9,989404 | 1,70E-23 | 1,83E-21 | FABP7 |
| ENSGALG00000005442 | 270,98777 | -1,327990169 | 0,2831728 | -4,689682 | 2,74E-06 | 3,20E-05 | PALMD |
| ENSGALG00000016602 | 1659,682 | -1,339565689 | 0,1637218 | -8,181961 | 2,79E-16 | 1,58E-14 | ARHGAP6 |
| ENSGALG00000021879 | 40,043275 | -1,34129965 | 0,3484949 | -3,848836 | 0,0001187 | 0,0008659 | |
| ENSGALG00000002443 | 492,98343 | -1,353430658 | 0,2272211 | -5,956448 | 2,58E-09 | 5,77E-08 | FBXL16 |
| ENSGALG00000007771 | 2716,4461 | -1,353742442 | 0,1514834 | -8,936574 | 4,01E-19 | 2,98E-17 | FAM210B |
| ENSGALG00000008491 | 3635,9278 | -1,354708919 | 0,1393335 | -9,72278 | 2,41E-22 | 2,38E-20 | XIAP |
| ENSGALG00000009947 | 4729,0815 | -1,358616298 | 0,1191908 | -11,39867 | 4,25E-30 | 7,32E-28 | PLEKHH2 |
| ENSGALG00000001346 | 421,97082 | -1,362109597 | 0,2357782 | -5,77708 | 7,60E-09 | 1,53E-07 | |
| ENSGALG00000028048 | 17805,974 | -1,362450241 | 0,1038968 | -13,1135 | 2,76E-39 | 8,73E-37 | PTN |
| ENSGALG00000028047 | 613,76602 | -1,369008166 | 0,2271663 | -6,026458 | 1,68E-09 | 3,86E-08 | RHOV |
| ENSGALG00000010148 | 96,068007 | -1,379685752 | 0,3285947 | -4,198746 | 2,68E-05 | 0,0002417 | PTPRR |
| ENSGALG00000003706 | 2880,2185 | -1,390323026 | 0,1522007 | -9,134799 | 6,55E-20 | 5,25E-18 | DUSP1 |
| ENSGALG00000009405 | 351,46777 | -1,391691484 | 0,2252283 | -6,179026 | 6,45E-10 | 1,57E-08 | GRIA2 |
| ENSGALG00000014983 | 1427,7077 | -1,39278689 | 0,1752564 | -7,947137 | 1,91E-15 | 9,66E-14 | F2R |
| ENSGALG00000002993 | 1938,9589 | -1,397301236 | 0,1581691 | -8,834225 | 1,01E-18 | 7,22E-17 | CA10 |
| ENSGALG00000015256 | 9909,979 | -1,397562325 | 0,1233891 | -11,32647 | 9,70E-30 | 1,61E-27 | PAM |
| ENSGALG00000003114 | 426,74171 | -1,398035927 | 0,2162026 | -6,466322 | 1,00E-10 | 2,72E-09 | NOG |

| | | | | | | | |
|---|---|---|---|---|---|---|---|
| ENSGALG00000011306 | 2909,7094 | -1,403496478 | 0,1324006 | -10,60038 | 2,97E-26 | 4,17E-24 | DES |
| ENSGALG00000027133 | 162,08932 | -1,405319409 | 0,31973 | -4,395332 | 1,11E-05 | 0,000113 | CD83 |
| ENSGALG00000001768 | 6648,8124 | -1,406520646 | 0,156826 | -8,968671 | 3,00E-19 | 2,27E-17 | TENM2 |
| ENSGALG00000005123 | 593,9961 | -1,408198856 | 0,2224863 | -6,329374 | 2,46E-10 | 6,31E-09 | WNT7A |
| ENSGALG00000000680 | 6447,7598 | -1,411570576 | 0,1125835 | -12,53799 | 4,63E-36 | 1,19E-33 | NUAK2 |
| ENSGALG00000009530 | 697,31067 | -1,411853584 | 0,1982451 | -7,121758 | 1,07E-12 | 3,88E-11 | PDZRN4 |
| ENSGALG00000014343 | 783,22971 | -1,412639292 | 0,1787261 | -7,903934 | 2,70E-15 | 1,32E-13 | ARAP2 |
| ENSGALG00000026557 | 168,82638 | -1,412974627 | 0,2911389 | -4,853266 | 1,21E-06 | 1,55E-05 | PGF |
| ENSGALG00000011319 | 10478,782 | -1,42017882 | 0,126589 | -11,21882 | 3,30E-29 | 5,34E-27 | LPHN3 |
| ENSGALG00000014823 | 40,51625 | -1,422401118 | 0,3456571 | -4,115064 | 3,87E-05 | 0,0003312 | C6orf58 |
| ENSGALG00000011468 | 1760,2982 | -1,424650779 | 0,1875286 | -7,596979 | 3,03E-14 | 1,36E-12 | IGFBP5 |
| ENSGALG00000016297 | 52,464778 | -1,427639697 | 0,3539064 | -4,033947 | 5,48E-05 | 0,0004458 | HCRTR2 |
| ENSGALG00000019054 | 529,71662 | -1,427749976 | 0,2415069 | -5,911839 | 3,38E-09 | 7,38E-08 | NAALAD2 |
| ENSGALG00000011110 | 296,25723 | -1,431359085 | 0,2488071 | -5,752887 | 8,77E-09 | 1,75E-07 | DPP4 |
| ENSGALG00000011207 | 13680,733 | -1,434035643 | 0,1207094 | -11,88007 | 1,50E-32 | 3,30E-30 | DUSP6 |
| ENSGALG00000012018 | 2349,8518 | -1,435600968 | 0,1869145 | -7,680524 | 1,58E-14 | 7,29E-13 | UGT8 |
| ENSGALG00000013402 | 51,692715 | -1,446510354 | 0,3538534 | -4,087881 | 4,35E-05 | 0,0003636 | AHRR |
| ENSGALG00000011164 | 2326,4168 | -1,446841804 | 0,2083124 | -6,945539 | 3,77E-12 | 1,26E-10 | CHN2 |
| ENSGALG00000028821 | 11178,852 | -1,46102831 | 0,1488545 | -9,815142 | 9,69E-23 | 9,70E-21 | GLIPR2 |
| ENSGALG00000001169 | 20569,632 | -1,469628578 | 0,1453759 | -10,10916 | 5,03E-24 | 5,85E-22 | CRB2 |
| ENSGALG00000006072 | 43,451296 | -1,471939488 | 0,3539382 | -4,158747 | 3,20E-05 | 0,0002809 | SCN10A |
| ENSGALG00000005855 | 6149,8313 | -1,488520056 | 0,1624534 | -9,162749 | 5,06E-20 | 4,12E-18 | CA5B |
| ENSGALG00000026262 | 5433,5657 | -1,4922838 | 0,1545273 | -9,657086 | 4,59E-22 | 4,25E-20 | PMP22 |
| ENSGALG00000001621 | 458,45489 | -1,495017381 | 0,2209806 | -6,765379 | 1,33E-11 | 4,12E-10 | CLDN5 |
| ENSGALG00000011940 | 614,50292 | -1,496267605 | 0,2191218 | -6,828475 | 8,58E-12 | 2,75E-10 | CCK |
| ENSGALG00000008978 | 15989,385 | -1,505709648 | 0,1318417 | -11,42059 | 3,30E-30 | 5,76E-28 | ITGA4 |
| ENSGALG00000025773 | 391,15903 | -1,506718569 | 0,2193821 | -6,868009 | 6,51E-12 | 2,11E-10 | |
| ENSGALG00000015388 | 3557,5816 | -1,507384513 | 0,1325751 | -11,37004 | 5,90E-30 | 1,00E-27 | PRDM1 |
| ENSGALG00000007171 | 156,08231 | -1,518198388 | 0,2937383 | -5,168541 | 2,36E-07 | 3,50E-06 | CIITA |
| ENSGALG00000010258 | 1087,7077 | -1,528410178 | 0,2320488 | -6,58659 | 4,50E-11 | 1,28E-09 | LTBP2 |
| ENSGALG00000006084 | 149,65329 | -1,531133293 | 0,3082185 | -4,967689 | 6,78E-07 | 9,10E-06 | WNT8A |
| ENSGALG00000005499 | 52125,792 | -1,531899327 | 0,1548752 | -9,891183 | 4,55E-23 | 4,65E-21 | IL17RD |
| ENSGALG00000001383 | 88,961213 | -1,543511015 | 0,3420022 | -4,513161 | 6,39E-06 | 6,86E-05 | HTR4 |
| ENSGALG00000017179 | 397,23584 | -1,548130217 | 0,2756174 | -5,616954 | 1,94E-08 | 3,65E-07 | PDGFD |
| ENSGALG00000000388 | 10384,966 | -1,555950834 | 0,1505748 | -10,33341 | 4,98E-25 | 6,33E-23 | LIN28A |
| ENSGALG00000016293 | 1486,8847 | -1,561397543 | 0,1523922 | -10,24591 | 1,23E-24 | 1,51E-22 | BMP5 |
| ENSGALG00000014657 | 1240,6807 | -1,561923173 | 0,2131383 | -7,328215 | 2,33E-13 | 9,26E-12 | ADGRV1 |
| ENSGALG00000003957 | 1745,618 | -1,565814231 | 0,1495869 | -10,46759 | 1,22E-25 | 1,66E-23 | APOH |
| ENSGALG00000017242 | 1988,3309 | -1,579394776 | 0,1359327 | -11,61895 | 3,30E-31 | 6,42E-29 | FZD4 |
| ENSGALG00000009170 | 1955,3023 | -1,581054934 | 0,1355952 | -11,66011 | 2,04E-31 | 4,02E-29 | NCEH1 |
| ENSGALG00000003323 | 100,58975 | -1,591926379 | 0,3209027 | -4,960776 | 7,02E-07 | 9,41E-06 | OSGIN1 |
| ENSGALG00000001645 | 733,71496 | -1,608468631 | 0,2050944 | -7,842576 | 4,41E-15 | 2,10E-13 | KCNT1 |
| ENSGALG00000011609 | 3466,2373 | -1,61436812 | 0,3265448 | -4,943788 | 7,66E-07 | 1,02E-05 | ABTB2 |
| ENSGALG00000003126 | 1209,6364 | -1,615974863 | 0,1617448 | -9,990893 | 1,67E-23 | 1,82E-21 | ERBB4 |
| ENSGALG00000019716 | 86,528621 | -1,618712644 | 0,3285879 | -4,926269 | 8,38E-07 | 1,10E-05 | KRT20 |
| ENSGALG00000016102 | 256,08529 | -1,630912312 | 0,2481459 | -6,572392 | 4,95E-11 | 1,41E-09 | CYP26B1 |
| ENSGALG00000010468 | 1107,4297 | -1,6309453 | 0,1573602 | -10,36441 | 3,60E-25 | 4,67E-23 | ISM2 |
| ENSGALG00000023521 | 1102,9061 | -1,651708778 | 0,1982919 | -8,329684 | 8,11E-17 | 4,84E-15 | ADRA2A |
| ENSGALG00000019261 | 1018,7346 | -1,655584971 | 0,2240687 | -7,388737 | 1,48E-13 | 6,04E-12 | ARHGAP8 |
| ENSGALG00000026892 | 80,007612 | -1,6734567 | 0,3494002 | -4,789512 | 1,67E-06 | 2,06E-05 | |
| ENSGALG00000008353 | 2780,0954 | -1,679225856 | 0,1922691 | -8,733726 | 2,46E-18 | 1,67E-16 | MAGI2 |
| ENSGALG00000006997 | 745,7891 | -1,684979264 | 0,1736039 | -9,705885 | 2,85E-22 | 2,77E-20 | BRINP1 |
| ENSGALG00000029175 | 12384,251 | -1,685259625 | 0,1660405 | -10,14969 | 3,32E-24 | 3,90E-22 | EFNB2 |
| ENSGALG00000008918 | 724,31518 | -1,685987772 | 0,1865069 | -9,039814 | 1,57E-19 | 1,22E-17 | LAMP5 |
| ENSGALG00000014930 | 4308,5867 | -1,696897635 | 0,1564042 | -10,84944 | 2,01E-27 | 2,94E-25 | ENC1 |
| ENSGALG00000005498 | 1151,8463 | -1,707286081 | 0,2153293 | -7,928721 | 2,21E-15 | 1,11E-13 | |
| ENSGALG00000014235 | 549,44433 | -1,721709963 | 0,2641679 | -6,517485 | 7,15E-11 | 1,99E-09 | WNT7B |
| ENSGALG00000013102 | 1811,9189 | -1,724325032 | 0,1396681 | -12,34587 | 5,13E-35 | 1,25E-32 | COBL |
| ENSGALG00000007218 | 6209,2721 | -1,732428983 | 0,1735379 | -9,982999 | 1,81E-23 | 1,94E-21 | MB21D2 |
| ENSGALG00000014908 | 2604,0347 | -1,732825245 | 0,1633068 | -10,61086 | 2,65E-26 | 3,76E-24 | FST |
| ENSGALG00000005476 | 213,71162 | -1,752170936 | 0,3180675 | -5,508801 | 3,61E-08 | 6,35E-07 | PTH1R |
| ENSGALG00000011148 | 2471,4965 | -1,754606037 | 0,1499315 | -11,70272 | 1,23E-31 | 2,51E-29 | HHIPL1 |
| ENSGALG00000027874 | 793,36454 | -1,754956403 | 0,232655 | -7,54317 | 4,59E-14 | 1,97E-12 | CHAC1 |
| ENSGALG00000012635 | 1110,8535 | -1,755097361 | 0,1664549 | -10,54398 | 5,42E-26 | 7,45E-24 | BTBD11 |

| | | | | | | | |
|---|---|---|---|---|---|---|---|
| ENSGALG00000006995 | 175,82725 | -1,759714805 | 0,3023936 | -5,819285 | 5,91E-09 | 1,22E-07 | APOD |
| ENSGALG00000027316 | 126,11379 | -1,764476803 | 0,3121114 | -5,653356 | 1,57E-08 | 3,00E-07 | CRNN |
| ENSGALG00000005519 | 2075,7509 | -1,767947956 | 0,3335103 | -5,30103 | 1,15E-07 | 1,82E-06 | TLL2 |
| ENSGALG00000009282 | 5108,519 | -1,769054515 | 0,16415 | -10,77706 | 4,42E-27 | 6,40E-25 | ELMSAN1 |
| ENSGALG00000007113 | 2619,5663 | -1,77145981 | 0,1909494 | -9,277115 | 1,74E-20 | 1,45E-18 | TNC |
| ENSGALG00000015148 | 101,3242 | -1,772924708 | 0,3468771 | -5,111103 | 3,20E-07 | 4,62E-06 | ANXA1 |
| ENSGALG00000023406 | 3748,5665 | -1,774894708 | 0,1767754 | -10,04039 | 1,01E-23 | 1,12E-21 | |
| ENSGALG00000004930 | 259,83629 | -1,782009807 | 0,3260345 | -5,465709 | 4,61E-08 | 7,94E-07 | HRH1 |
| ENSGALG00000009920 | 1810,7316 | -1,80046145 | 0,2059481 | -8,742306 | 2,28E-18 | 1,56E-16 | COCH |
| ENSGALG00000014159 | 18203,792 | -1,81424576 | 0,1061679 | -17,08846 | 1,81E-65 | 1,30E-62 | PACSIN2 |
| ENSGALG00000016875 | 1624,3852 | -1,825016077 | 0,1855323 | -9,836647 | 7,83E-23 | 7,89E-21 | CLYBL |
| ENSGALG00000004306 | 1596,7644 | -1,830041848 | 0,1566925 | -11,67919 | 1,63E-31 | 3,26E-29 | SYT12 |
| ENSGALG00000011708 | 13494,005 | -1,833902608 | 0,1212478 | -15,12524 | 1,10E-51 | 6,01E-49 | MYLK |
| ENSGALG00000002714 | 18305,731 | -1,839377527 | 0,1231358 | -14,93779 | 1,87E-50 | 9,10E-48 | TMEM132C |
| ENSGALG00000007963 | 269,37035 | -1,84239576 | 0,2523089 | -7,302143 | 2,83E-13 | 1,11E-11 | NFATC2 |
| ENSGALG00000014935 | 24043,958 | -1,864724119 | 0,1640609 | -11,36605 | 6,17E-30 | 1,04E-27 | GREB1L |
| ENSGALG00000015089 | 1244,3881 | -1,867848339 | 0,2166645 | -8,620924 | 6,64E-18 | 4,39E-16 | GAP43 |
| ENSGALG00000016906 | 16521,342 | -1,870006943 | 0,1023605 | -18,26883 | 1,47E-74 | 2,00E-71 | SPRY2 |
| ENSGALG00000012392 | 124,6248 | -1,879813042 | 0,3053078 | -6,157108 | 7,41E-10 | 1,79E-08 | IYD |
| ENSGALG00000007786 | 809,87356 | -1,895122662 | 0,2583339 | -7,335942 | 2,20E-13 | 8,79E-12 | DOK5 |
| ENSGALG00000003437 | 453,52356 | -1,896224168 | 0,2236049 | -8,480244 | 2,25E-17 | 1,43E-15 | ADAM8 |
| ENSGALG00000009693 | 162,98636 | -1,908196694 | 0,2849148 | -6,69743 | 2,12E-11 | 6,38E-10 | CPXM2 |
| ENSGALG00000026366 | 659,49696 | -1,956428809 | 0,2559307 | -7,644371 | 2,10E-14 | 9,56E-13 | TTN |
| ENSGALG00000016377 | 39318,721 | -1,984049771 | 0,1516015 | -13,08727 | 3,89E-39 | 1,20E-36 | PXDN |
| ENSGALG00000007646 | 1162,4308 | -1,999886692 | 0,2341543 | -8,540893 | 1,33E-17 | 8,72E-16 | |
| ENSGALG00000014727 | 707,4954 | -2,02826974 | 0,2257943 | -8,982819 | 2,64E-19 | 2,02E-17 | PDE4D |
| ENSGALG00000025941 | 508,27465 | -2,032822244 | 0,2108225 | -9,642339 | 5,30E-22 | 4,84E-20 | RGS20 |
| ENSGALG00000016455 | 36148,301 | -2,034647249 | 0,1415614 | -14,37289 | 7,66E-47 | 3,07E-44 | GREB1 |
| ENSGALG00000015332 | 1189,9865 | -2,044938557 | 0,2318737 | -8,819191 | 1,15E-18 | 8,09E-17 | CPLX1 |
| ENSGALG00000001608 | 473,98158 | -2,081100043 | 0,2297131 | -9,059563 | 1,31E-19 | 1,03E-17 | UNC5D |
| ENSGALG00000006797 | 719,73345 | -2,093089949 | 0,1825483 | -11,46595 | 1,96E-30 | 3,60E-28 | ADARB2 |
| ENSGALG00000005347 | 2547,968 | -2,102469906 | 0,1728132 | -12,16614 | 4,71E-34 | 1,11E-31 | ADAMTS18 |
| ENSGALG00000005061 | 4294,1757 | -2,114741441 | 0,1575348 | -13,42396 | 4,38E-41 | 1,53E-38 | PPM1E |
| ENSGALG00000006822 | 6099,6969 | -2,117548167 | 0,190863 | -11,0946 | 1,33E-28 | 2,06E-26 | PCDH19 |
| ENSGALG00000002540 | 1690,5022 | -2,132167559 | 0,1444055 | -14,76514 | 2,46E-49 | 1,08E-46 | RGS2 |
| ENSGALG00000014872 | 706,84998 | -2,137267919 | 0,2117349 | -10,09407 | 5,87E-24 | 6,77E-22 | FGF10 |
| ENSGALG00000005000 | 10982,035 | -2,158647655 | 0,1512305 | -14,27389 | 3,18E-46 | 1,24E-43 | COMTD1 |
| ENSGALG00000002816 | 1397,7493 | -2,170634895 | 0,1725497 | -12,57976 | 2,73E-36 | 7,28E-34 | STK10 |
| ENSGALG00000024083 | 2014,3842 | -2,175833689 | 0,1641192 | -13,25765 | 4,07E-40 | 1,39E-37 | C8orf4 |
| ENSGALG00000017244 | 1712,4698 | -2,189436741 | 0,1475328 | -14,84034 | 8,03E-50 | 3,65E-47 | PRSS23 |
| ENSGALG00000012106 | 326,65016 | -2,236831516 | 0,2846604 | -7,857895 | 3,91E-15 | 1,87E-13 | SCTR |
| ENSGALG00000005689 | 443,46806 | -2,271345118 | 0,2372034 | -9,575515 | 1,01E-21 | 9,14E-20 | PAX2 |
| ENSGALG00000000600 | 217,11331 | -2,27272638 | 0,2827556 | -8,037777 | 9,15E-16 | 4,85E-14 | PTPN7 |
| ENSGALG00000019426 | 136,01564 | -2,280006749 | 0,3003394 | -7,591434 | 3,16E-14 | 1,41E-12 | TMEM71 |
| ENSGALG00000012594 | 465,97876 | -2,283939186 | 0,2248463 | -10,15778 | 3,06E-24 | 3,62E-22 | NTRK2 |
| ENSGALG00000012732 | 358,70918 | -2,298252682 | 0,2277106 | -10,09287 | 5,94E-24 | 6,80E-22 | PHACTR1 |
| ENSGALG00000004376 | 1102,6462 | -2,323750078 | 0,1982175 | -11,72323 | 9,69E-32 | 2,00E-29 | KCNJ2 |
| ENSGALG00000008118 | 2184,0618 | -2,340544818 | 0,1326409 | -17,64572 | 1,10E-69 | 1,15E-66 | RFTN2 |
| ENSGALG00000010435 | 1040,6262 | -2,340611141 | 0,2214259 | -10,57063 | 4,08E-26 | 5,66E-24 | RASGRP3 |
| ENSGALG00000007346 | 9971,3513 | -2,361764382 | 0,1565921 | -15,08227 | 2,12E-51 | 1,11E-48 | KIAA1462 |
| ENSGALG00000017091 | 828,29835 | -2,363903034 | 0,1707454 | -13,84461 | 1,37E-43 | 5,05E-41 | FLT1 |
| ENSGALG00000012278 | 11240,166 | -2,383840075 | 0,163892 | -14,54519 | 6,27E-48 | 2,67E-45 | EMCN |
| ENSGALG00000016785 | 93,113308 | -2,405348378 | 0,3296339 | -7,297029 | 2,94E-13 | 1,15E-11 | IL1RL1 |
| ENSGALG00000013033 | 107,44587 | -2,410715686 | 0,3195602 | -7,543854 | 4,56E-14 | 1,97E-12 | CMBL |
| ENSGALG00000007657 | 4917,8686 | -2,412172788 | 0,3343808 | -7,213849 | 5,44E-13 | 2,06E-11 | CDX4 |
| ENSGALG00000012927 | 231,31802 | -2,419967236 | 0,2657891 | -9,104839 | 8,64E-20 | 6,84E-18 | CDH10 |
| ENSGALG00000002183 | 431,19833 | -2,438499895 | 0,2660795 | -9,164552 | 4,98E-20 | 4,08E-18 | PRR16 |
| ENSGALG00000015947 | 963,91091 | -2,438977585 | 0,1891685 | -12,89315 | 4,92E-38 | 1,46E-35 | CDH17 |
| ENSGALG00000008893 | 301,74621 | -2,444595373 | 0,2469193 | -9,900382 | 4,15E-23 | 4,31E-21 | SYT4 |
| ENSGALG00000008890 | 114,85977 | -2,449553325 | 0,3261648 | -7,51017 | 5,91E-14 | 2,49E-12 | ADGRL4 |
| ENSGALG00000008905 | 317,21852 | -2,456637668 | 0,2373167 | -10,35173 | 4,11E-25 | 5,28E-23 | HABP2 |
| ENSGALG00000026663 | 524,42245 | -2,478656591 | 0,2092266 | -11,84675 | 2,24E-32 | 4,76E-30 | |
| ENSGALG00000012623 | 104,28468 | -2,482707417 | 0,3217829 | -7,715473 | 1,21E-14 | 5,58E-13 | CFC1B |
| ENSGALG00000015826 | 274,53091 | -2,484029553 | 0,3182796 | -7,804553 | 5,97E-15 | 2,81E-13 | HTR1E |

| | | | | | | | |
|---|---|---|---|---|---|---|---|
| ENSGALG00000010224 | 221,98807 | -2,486198557 | 0,2911733 | -8,538554 | 1,36E-17 | 8,83E-16 | GLIPR1L |
| ENSGALG00000007940 | 128,05228 | -2,542363859 | 0,32048 | -7,932988 | 2,14E-15 | 1,07E-13 | RAG2 |
| ENSGALG00000001560 | 195,37323 | -2,555527127 | 0,3382967 | -7,554101 | 4,22E-14 | 1,83E-12 | KCNU1 |
| ENSGALG00000008581 | 16731,944 | -2,566439645 | 0,1474527 | -17,40517 | 7,54E-68 | 5,70E-65 | DUSP5 |
| ENSGALG00000016784 | 1039,8515 | -2,607549769 | 0,1741423 | -14,97367 | 1,09E-50 | 5,50E-48 | IL1RL2 |
| ENSGALG00000006112 | 988,65304 | -2,692857061 | 0,1899063 | -14,17993 | 1,22E-45 | 4,61E-43 | SCN5A |
| ENSGALG00000016154 | 140,89016 | -2,742713554 | 0,306496 | -8,948611 | 3,60E-19 | 2,71E-17 | ARC |
| ENSGALG00000016145 | 1265,1012 | -2,754137035 | 0,1745953 | -15,7744 | 4,67E-56 | 2,76E-53 | RIPK4 |
| ENSGALG00000011099 | 581,91676 | -2,760215905 | 0,269967 | -10,22427 | 1,54E-24 | 1,88E-22 | FAP |
| ENSGALG00000005209 | 4151,2431 | -2,791291062 | 0,3487778 | -8,003063 | 1,21E-15 | 6,28E-14 | RP5-877J2.1 |
| ENSGALG00000011562 | 1031,64 | -2,817896047 | 0,1809122 | -15,57604 | 1,06E-54 | 6,01E-52 | PARK2 |
| ENSGALG00000007656 | 17151,759 | -2,823200316 | 0,3012681 | -9,371057 | 7,18E-21 | 6,11E-19 | |
| ENSGALG00000012051 | 829,36725 | -2,878544854 | 0,2482483 | -11,59542 | 4,35E-31 | 8,22E-29 | STAC |
| ENSGALG00000028250 | 102,79679 | -2,884651456 | 0,3308091 | -8,719988 | 2,78E-18 | 1,88E-16 | ANKRD22 |
| ENSGALG00000007701 | 2491,5718 | -2,906313035 | 0,1654067 | -17,57071 | 4,13E-69 | 3,75E-66 | B3GNT7 |
| ENSGALG00000014984 | 5902,5896 | -2,931476354 | 0,1495694 | -19,59944 | 1,56E-85 | 3,04E-82 | F2RL1 |
| ENSGALG00000009704 | 18346,867 | -2,934403789 | 0,1676164 | -17,50666 | 1,27E-68 | 1,02E-65 | CHST15 |
| ENSGALG00000016140 | 355,67412 | -2,998033733 | 0,2492573 | -12,02787 | 2,54E-33 | 5,76E-31 | FAM3B |
| ENSGALG00000015358 | 3246,1808 | -3,063209848 | 0,145761 | -21,0153 | 4,75E-98 | 1,29E-94 | MYH15 |
| ENSGALG00000002562 | 2027,1289 | -3,12939254 | 0,1764516 | -17,73513 | 2,25E-70 | 2,55E-67 | ARAP3 |
| ENSGALG00000007538 | 106,89217 | -3,275953369 | 0,3461412 | -9,464212 | 2,96E-21 | 2,60E-19 | IRG1L |
| ENSGALG00000022962 | 106,15694 | -3,362304357 | 0,3346563 | -10,04704 | 9,47E-24 | 1,06E-21 | OVST |
| ENSGALG00000002594 | 1003,2406 | -3,661637709 | 0,2050909 | -17,85373 | 2,70E-71 | 3,35E-68 | TFPI |
| ENSGALG00000013624 | 4838,4306 | -3,6799039 | 0,1462758 | -25,15731 | 1,18E-139 | 5,34E-136 | FAM65B |
| ENSGALG00000006421 | 916,75433 | -3,742853665 | 0,2370467 | -15,78952 | 3,67E-56 | 2,27E-53 | SLC16A12 |
| ENSGALG00000020359 | 246,79437 | -3,821393345 | 0,2845023 | -13,43185 | 3,93E-41 | 1,41E-38 | SIX6 |
| ENSGALG00000003456 | 187,12467 | -3,934712358 | 0,3247803 | -12,115 | 8,80E-34 | 2,03E-31 | CA12 |
| ENSGALG00000023379 | 453,27145 | -3,949811074 | 0,2313834 | -17,07042 | 2,46E-65 | 1,68E-62 | F2RL2 |
| ENSGALG00000016431 | 670,26005 | -4,37340061 | 0,2358196 | -18,54553 | 8,86E-77 | 1,34E-73 | PAX5 |
| ENSGALG00000014994 | 310,70765 | -5,043766073 | 0,2868267 | -17,58472 | 3,23E-69 | 3,14E-66 | CRHBP |
| ENSGALG00000016783 | 9121,1085 | -5,301446621 | 0,1450351 | -36,55285 | 1,61E-292 | 2,19E-288 | IL1R1 |
| ENSGALG00000004221 | 898,01485 | -5,338632777 | 0,2699166 | -19,77882 | 4,53E-87 | 1,03E-83 | IL22RA1 |

**Table 5:** Classification of the 2316 genes deregulated in the “MEK1ca versus pCIG” condition (False Discovery Rate (padj) <= 0.005) clustered using “kmeans” with k=12, with the mean TPM (transcripts per kilobase million) of each gene for the two replicates of pCIG, HOXB8, MEK1ca, and MEK1ca+HOXB8 in each condition.

| ID | Symbol | pCIG | HOXB8 | MEK1ca | MEK1_HOXB8 | Cluster |
|---|---|---|---|---|---|---|
| ENSGALG00000000141 | BLB1 | 4,346461397 | 6,936083838 | 25,83463981 | 9,813152958 | 1 |
| ENSGALG00000000307 | | 59,23843877 | 89,71588529 | 143,6917892 | 78,57624532 | 1 |
| ENSGALG00000000600 | | 0,16716819 | 0,219662917 | 4,032909459 | 0,585105026 | 1 |
| ENSGALG00000000639 | LRRN2 | 4,627904717 | 4,790564693 | 9,521707125 | 5,306668428 | 1 |
| ENSGALG00000000761 | TSKU | 53,35356817 | 54,75872543 | 78,02304089 | 50,7231382 | 1 |
| ENSGALG00000000795 | MFSD12 | 15,27116365 | 19,51741298 | 24,13081695 | 17,43156709 | 1 |
| ENSGALG00000000977 | MAPKAP1 | 76,21721102 | 75,42044153 | 139,8918526 | 69,33283402 | 1 |
| ENSGALG00000001094 | ADGRD2 | 102,7073676 | 111,4654329 | 121,4385963 | 94,7567043 | 1 |
| ENSGALG00000001220 | HERPUD1 | 52,87970256 | 43,77611515 | 97,02489846 | 55,98900064 | 1 |
| ENSGALG00000001346 | | 12,45034341 | 13,79821487 | 25,17364864 | 8,821835481 | 1 |
| ENSGALG00000001370 | ADAMTS8 | 10,00645846 | 11,44367995 | 17,37313634 | 8,098120229 | 1 |
| ENSGALG00000001383 | HTR4 | 0,223346685 | 0,462240728 | 2,654657367 | 0,480455141 | 1 |
| ENSGALG00000001405 | IRF6 | 2,701639368 | 2,9520233 | 6,01110282 | 3,704203216 | 1 |
| ENSGALG00000001419 | DAB2IP | 8,238485904 | 12,58819262 | 17,26179821 | 10,32238732 | 1 |
| ENSGALG00000001560 | | 0,150607399 | 0,295573757 | 2,997594118 | 0,175011822 | 1 |
| ENSGALG00000001596 | MLLT1 | 104,5141917 | 111,7769614 | 144,5899475 | 114,1039573 | 1 |
| ENSGALG00000001621 | CLDN5 | 1,779507208 | 3,559103071 | 26,91827318 | 8,738372257 | 1 |
| ENSGALG00000002183 | PRR16 | 1,802939556 | 2,415784551 | 21,12574473 | 2,851415468 | 1 |
| ENSGALG00000002242 | | 3,30906785 | 4,319855305 | 5,828198288 | 3,158054392 | 1 |
| ENSGALG00000002509 | | 30,0871481 | 32,18952881 | 61,1232203 | 26,93140662 | 1 |
| ENSGALG00000002532 | GJA4 | 0,544399491 | 0,638100325 | 1,969673798 | 0,784806618 | 1 |
| ENSGALG00000002540 | | 6,446932133 | 5,308050588 | 40,08364447 | 8,825438475 | 1 |
| ENSGALG00000002543 | CSGALNACT2 | 32,12663012 | 34,93337178 | 76,29195476 | 43,26033436 | 1 |
| ENSGALG00000002555 | RET | 2,88163933 | 4,228435821 | 7,895631031 | 4,06619279 | 1 |
| ENSGALG00000002562 | ARAP3 | 1,48713749 | 1,131854243 | 19,88581577 | 2,017191883 | 1 |
| ENSGALG00000002594 | TFPI | 1,818415284 | 2,178414033 | 30,95014095 | 1,96144887 | 1 |
| ENSGALG00000002598 | TPST1 | 110,6957744 | 124,8080538 | 194,3906783 | 111,9974286 | 1 |
| ENSGALG00000002623 | PCOLCE2 | 6,900390576 | 7,625754792 | 13,68975598 | 9,209524817 | 1 |
| ENSGALG00000002632 | CALCRL | 12,01510153 | 10,78717477 | 32,20239461 | 13,53387588 | 1 |
| ENSGALG00000002646 | FAM171B | 70,39669635 | 76,78219835 | 117,3361401 | 83,48810086 | 1 |
| ENSGALG00000002655 | ITGAV | 34,70042214 | 37,88033757 | 56,07206538 | 41,54256787 | 1 |
| ENSGALG00000002704 | | 1,560758451 | 1,645911429 | 3,272911068 | 1,63570101 | 1 |
| ENSGALG00000002763 | FRZB | 88,08590473 | 101,8551818 | 118,1762783 | 90,87514056 | 1 |
| ENSGALG00000002816 | STK10 | 1,302638213 | 1,070607568 | 9,306934994 | 1,934117709 | 1 |
| ENSGALG00000002830 | NUDT3 | 82,47138192 | 83,39720868 | 119,0949223 | 90,87056175 | 1 |
| ENSGALG00000002863 | ERGIC1 | 200,1815036 | 199,8499674 | 251,621888 | 188,1263931 | 1 |
| ENSGALG00000002954 | MBTD1 | 61,91506849 | 61,41366817 | 80,11960497 | 59,8379799 | 1 |
| ENSGALG00000002993 | CA10 | 4,907630575 | 7,330232074 | 13,05830949 | 4,830057404 | 1 |
| ENSGALG00000003114 | NOG | 8,459709423 | 11,93287775 | 23,8221868 | 8,350925143 | 1 |
| ENSGALG00000003149 | | 3,188704926 | 3,828601125 | 6,728823346 | 4,067555236 | 1 |
| ENSGALG00000003323 | OSGIN1 | 0,150652789 | 0,306431517 | 2,76792484 | 0,600147405 | 1 |
| ENSGALG00000003339 | MTOR | 32,72614893 | 33,27392871 | 40,05816094 | 32,03402147 | 1 |
| ENSGALG00000003353 | RAI14 | 72,22040829 | 61,3924759 | 156,0160059 | 63,07779452 | 1 |
| ENSGALG00000003437 | | 1,140967301 | 0,975600437 | 7,404177579 | 1,755276738 | 1 |
| ENSGALG00000003456 | CA12 | 0,043477625 | 0,052506397 | 4,443271062 | 0,056186982 | 1 |
| ENSGALG00000003668 | | 0,107991316 | 0,101421289 | 0,499834388 | 0,201779775 | 1 |
| ENSGALG00000003706 | | 7,669767671 | 8,241795488 | 28,6640262 | 10,83001181 | 1 |
| ENSGALG00000004155 | MYOM3 | 0,565715899 | 0,738462272 | 1,297153697 | 0,505525313 | 1 |
| ENSGALG00000004221 | | 0,270203292 | 0,285792101 | 18,79414137 | 0,190172822 | 1 |
| ENSGALG00000004273 | RALGPS2 | 68,26603769 | 74,39473921 | 93,44497048 | 67,30976616 | 1 |
| ENSGALG00000004306 | SYT12 | 2,605589645 | 4,144014971 | 15,45617568 | 4,186305684 | 1 |
| ENSGALG00000004376 | KCNJ2 | 3,77289362 | 4,959356504 | 22,31025244 | 3,944582 | 1 |
| ENSGALG00000004393 | | 4,430270701 | 6,514512501 | 10,5028138 | 5,156954248 | 1 |
| ENSGALG00000004531 | PLOD1 | 39,43375984 | 39,16887142 | 53,49951129 | 39,85679447 | 1 |
| ENSGALG00000004569 | UNC5B | 7,868192231 | 10,03498033 | 22,37566356 | 12,64304576 | 1 |
| ENSGALG00000004612 | MTHFR | 29,88510462 | 33,82285398 | 48,61122399 | 33,33543084 | 1 |
| ENSGALG00000004624 | | 38,51562207 | 38,74443918 | 46,92510617 | 37,97246389 | 1 |
| ENSGALG00000004638 | | 12,39324598 | 12,9301946 | 17,06424419 | 12,4188215 | 1 |

| | | | | | | |
|---|---|---|---|---|---|---|
| ENSGALG00000004696 | LEO1 | 51,73562652 | 54,64381268 | 67,49391493 | 53,30887079 | 1 |
| ENSGALG00000004879 | SLC6A11 | 6,044456134 | 7,986360168 | 9,907299162 | 6,80721088 | 1 |
| ENSGALG00000004930 | HRH1 | 2,252570121 | 1,790119278 | 7,87682517 | 1,349179431 | 1 |
| ENSGALG00000005000 | COMTD1 | 48,07598474 | 44,35755518 | 397,3367537 | 85,33221268 | 1 |
| ENSGALG00000005061 | PPM1E | 15,5883157 | 12,4158043 | 76,17698494 | 16,84323915 | 1 |
| ENSGALG00000005134 | ZSWIM8 | 11,30147215 | 11,70309029 | 15,86549961 | 11,59078122 | 1 |
| ENSGALG00000005160 | VMP1 | 97,80171612 | 94,91460927 | 126,6233829 | 94,36577445 | 1 |
| ENSGALG00000005209 | | 0,978757944 | 1,181001848 | 116,1518425 | 3,250210874 | 1 |
| ENSGALG00000005347 | ADAMTS18 | 7,626809484 | 5,838020535 | 25,8863696 | 5,61477326 | 1 |
| ENSGALG00000005430 | ALS2CL | 1,955531218 | 2,01749045 | 3,820475596 | 2,280218129 | 1 |
| ENSGALG00000005442 | PALMD | 1,233769732 | 2,376997769 | 5,258268724 | 1,702027405 | 1 |
| ENSGALG00000005468 | SYNRG | 31,68189573 | 33,65304843 | 40,16103869 | 31,72045473 | 1 |
| ENSGALG00000005476 | PTH1R | 0,874654422 | 0,830470831 | 5,138549733 | 0,976037129 | 1 |
| ENSGALG00000005498 | | 6,346864963 | 6,410134604 | 28,72716752 | 8,222793763 | 1 |
| ENSGALG00000005499 | IL17RD | 42,29096857 | 36,47768079 | 270,3943487 | 91,41665589 | 1 |
| ENSGALG00000005519 | TLL2 | 1,410283264 | 2,247551296 | 22,60288502 | 3,669028574 | 1 |
| ENSGALG00000005610 | SLC44A3 | 5,452687132 | 6,470508823 | 10,61156478 | 5,917968895 | 1 |
| ENSGALG00000005619 | | 12,91699497 | 13,14521332 | 36,29311148 | 14,78644626 | 1 |
| ENSGALG00000005785 | | 6,273844084 | 6,527878689 | 9,293607737 | 4,778628942 | 1 |
| ENSGALG00000005805 | PLCD1 | 8,133157801 | 12,3694805 | 21,47493828 | 11,05897104 | 1 |
| ENSGALG00000005855 | | 17,59991732 | 15,64179009 | 118,2991947 | 40,94877771 | 1 |
| ENSGALG00000006002 | WHAMM | 15,4050577 | 16,68613201 | 22,65836529 | 17,34219926 | 1 |
| ENSGALG00000006072 | SCN10A | 0,071733051 | 0,072511186 | 0,322233693 | 0,040260369 | 1 |
| ENSGALG00000006084 | | 0,415711399 | 1,339960031 | 3,460295102 | 0,856058604 | 1 |
| ENSGALG00000006112 | SCN5A | 0,916499676 | 0,661017755 | 4,51117604 | 0,605407018 | 1 |
| ENSGALG00000006121 | PIK3C2A | 42,74947084 | 49,54970032 | 60,78554868 | 45,17039169 | 1 |
| ENSGALG00000006337 | | 0,002439521 | 0,023624003 | 0,312529764 | 0,041882001 | 1 |
| ENSGALG00000006376 | TMC3 | 0,192169524 | 0,233676578 | 1,09689024 | 0,353408492 | 1 |
| ENSGALG00000006401 | OSBPL5 | 11,36431471 | 10,26220066 | 31,66422311 | 12,85811602 | 1 |
| ENSGALG00000006421 | SLC16A12 | 0,786470857 | 0,789157454 | 28,31054622 | 1,468096746 | 1 |
| ENSGALG00000006609 | EPHB1 | 70,32545848 | 78,56087564 | 89,8486204 | 58,97301751 | 1 |
| ENSGALG00000006652 | DIP2C | 13,37555911 | 12,67551415 | 17,61300034 | 13,378876 | 1 |
| ENSGALG00000006808 | TSPAN6 | 84,8160072 | 92,4891866 | 182,1056518 | 96,16407149 | 1 |
| ENSGALG00000006975 | | 35,48790893 | 37,00507441 | 45,63493758 | 29,37760443 | 1 |
| ENSGALG00000006995 | APOD | 0,696051145 | 1,390188703 | 7,733888215 | 1,572610716 | 1 |
| ENSGALG00000006997 | BRINP1 | 3,499835162 | 4,356493158 | 11,27517561 | 3,348538768 | 1 |
| ENSGALG00000007039 | | 40,28481666 | 45,9836234 | 79,80599717 | 45,70745825 | 1 |
| ENSGALG00000007069 | | 1,998973396 | 3,269505285 | 4,779320231 | 2,74022749 | 1 |
| ENSGALG00000007113 | | 4,572530679 | 5,629890394 | 15,02562976 | 4,089934281 | 1 |
| ENSGALG00000007171 | CIITA | 0,022403266 | 0,204546083 | 2,706469378 | 0,728094044 | 1 |
| ENSGALG00000007206 | | 0 | 0,022465315 | 0,940527994 | 0,230962067 | 1 |
| ENSGALG00000007218 | MB21D2 | 21,50353374 | 22,18131924 | 63,39535546 | 18,17572811 | 1 |
| ENSGALG00000007242 | SLITRK4 | 11,61340101 | 13,68905429 | 17,8729469 | 10,94148732 | 1 |
| ENSGALG00000007268 | | 12,02218487 | 17,33922043 | 29,52538581 | 13,1036655 | 1 |
| ENSGALG00000007346 | KIAA1462 | 9,556019949 | 13,87679845 | 75,69402273 | 13,92754889 | 1 |
| ENSGALG00000007501 | | 6,923403837 | 6,775032581 | 12,8684032 | 7,692808633 | 1 |
| ENSGALG00000007538 | | 0 | 0,011572698 | 2,680467049 | 0,023686378 | 1 |
| ENSGALG00000007646 | | 1,828426892 | 2,407614248 | 12,04323697 | 2,574591371 | 1 |
| ENSGALG00000007656 | | 9,354467191 | 12,73350311 | 306,4981787 | 24,13865507 | 1 |
| ENSGALG00000007657 | CDX4 | 10,43751312 | 5,656329149 | 135,1902583 | 10,65993568 | 1 |
| ENSGALG00000007660 | | 18,53508552 | 20,01450682 | 57,59481836 | 23,79729358 | 1 |
| ENSGALG00000007667 | GPR158 | 2,017006699 | 2,351089848 | 4,019170374 | 2,390691361 | 1 |
| ENSGALG00000007701 | B3GNT7 | 1,965758713 | 2,885006525 | 58,70594437 | 7,123195196 | 1 |
| ENSGALG00000007771 | FAM210B | 48,42301262 | 61,59675671 | 141,9653945 | 54,87237005 | 1 |
| ENSGALG00000007786 | DOK5 | 1,639190441 | 2,831374722 | 12,89140399 | 2,824710311 | 1 |
| ENSGALG00000007940 | | 0,298085716 | 0,405740152 | 3,03794633 | 0,245682446 | 1 |
| ENSGALG00000007952 | AMOT | 42,59469668 | 43,129628 | 81,92458191 | 43,27930325 | 1 |
| ENSGALG00000007963 | NFATC2 | 0,197351415 | 0,379445389 | 3,998563902 | 0,912021662 | 1 |
| ENSGALG00000008118 | | 11,21822805 | 8,858651443 | 37,96056253 | 7,261176994 | 1 |
| ENSGALG00000008195 | NPHP1 | 52,32037043 | 55,15367663 | 70,06251176 | 55,17279702 | 1 |
| ENSGALG00000008293 | SH3PXD2A | 4,963643686 | 5,106021399 | 7,660740968 | 4,819388759 | 1 |
| ENSGALG00000008318 | SLK | 85,01647415 | 80,15010998 | 134,3520566 | 81,18806127 | 1 |
| ENSGALG00000008353 | | 4,025184036 | 3,680815167 | 16,66369921 | 4,874571991 | 1 |
| ENSGALG00000008469 | SPINT1 | 14,24665038 | 16,90480956 | 32,8484745 | 18,45751953 | 1 |

| | | | | | | |
|---|---|---|---|---|---|---|
| ENSGALG00000008542 | ADD3 | 118,4151596 | 124,28577 | 157,6607813 | 123,3415266 | 1 |
| ENSGALG00000008581 | | 10,16528829 | 6,061157018 | 256,373084 | 41,13464142 | 1 |
| ENSGALG00000008621 | NRP2 | 78,31928008 | 89,14932983 | 118,8571698 | 70,77451755 | 1 |
| ENSGALG00000008739 | FAM188A | 46,04693561 | 50,96110843 | 56,92147735 | 48,6759362 | 1 |
| ENSGALG00000008875 | PLCB1 | 4,345271238 | 4,743918313 | 14,04603032 | 6,225376121 | 1 |
| ENSGALG00000008890 | ADGRL4 | 0,129602088 | 0,092978733 | 0,996601503 | 0,083175824 | 1 |
| ENSGALG00000008905 | HABP2 | 0,266437294 | 0,48347075 | 6,734457842 | 0,981954515 | 1 |
| ENSGALG00000009170 | NCEH1 | 26,8945161 | 27,63392523 | 60,16087129 | 19,95188217 | 1 |
| ENSGALG00000009179 | TNFSF10 | 50,54952664 | 51,97607136 | 96,43379558 | 44,37200827 | 1 |
| ENSGALG00000009222 | FNDC3B | 20,05392267 | 20,78387393 | 24,33913641 | 20,24031624 | 1 |
| ENSGALG00000009252 | | 17,05340609 | 20,22552019 | 30,26838438 | 17,79052756 | 1 |
| ENSGALG00000009282 | ELMSAN1 | 12,79207245 | 10,87027045 | 43,26764769 | 12,23866375 | 1 |
| ENSGALG00000009300 | NUMB | 37,95931361 | 39,65737919 | 52,66309106 | 40,50552035 | 1 |
| ENSGALG00000009360 | CAPN2 | 3,497797582 | 4,055116565 | 9,127908666 | 4,85175942 | 1 |
| ENSGALG00000009415 | SMOC1 | 117,4087959 | 134,7914352 | 180,6575257 | 94,49588202 | 1 |
| ENSGALG00000009450 | DUSP10 | 9,354175589 | 11,43364627 | 21,919628 | 10,97128745 | 1 |
| ENSGALG00000009562 | | 71,61834554 | 77,27651814 | 93,73781016 | 58,03501792 | 1 |
| ENSGALG00000009567 | TLL1 | 12,75490022 | 14,9239491 | 30,19675472 | 13,74189903 | 1 |
| ENSGALG00000009667 | MFAP3L | 4,900753944 | 6,755229067 | 18,92590259 | 7,235188798 | 1 |
| ENSGALG00000009693 | CPXM2 | 0,297057574 | 0,224259756 | 1,916995035 | 0,375685436 | 1 |
| ENSGALG00000009704 | CHST15 | 26,42019968 | 31,60061585 | 300,8079085 | 35,72264945 | 1 |
| ENSGALG00000009740 | RASGRP1 | 2,238442875 | 3,221407126 | 8,304577968 | 3,876544109 | 1 |
| ENSGALG00000009752 | C1GALT1 | 73,87944835 | 76,00031211 | 88,11179224 | 77,6483376 | 1 |
| ENSGALG00000009850 | NOVA1 | 65,62071803 | 72,5812928 | 91,8483665 | 60,49260852 | 1 |
| ENSGALG00000009855 | TBC1D9 | 35,32107704 | 35,40997725 | 52,56579067 | 36,25236136 | 1 |
| ENSGALG00000009920 | COCH | 13,26044931 | 11,28208934 | 38,09312731 | 9,926593491 | 1 |
| ENSGALG00000009947 | PLEKHH2 | 8,548346442 | 7,655850005 | 20,90561351 | 8,219735522 | 1 |
| ENSGALG00000010010 | RHOQ | 22,36249765 | 32,86035501 | 57,1305441 | 31,25736751 | 1 |
| ENSGALG00000010061 | LRBA | 13,57799133 | 15,84454033 | 19,19400383 | 15,01651499 | 1 |
| ENSGALG00000010148 | | 0,376485353 | 0,303044356 | 1,090711174 | 0,276420272 | 1 |
| ENSGALG00000010156 | | 0,484158088 | 0,407906519 | 1,056535994 | 0,478788157 | 1 |
| ENSGALG00000010163 | LGR5 | 29,22401237 | 42,84673911 | 58,12291645 | 29,33576971 | 1 |
| ENSGALG00000010204 | KCNC2 | 8,315391818 | 9,598554626 | 14,43598051 | 9,193014079 | 1 |
| ENSGALG00000010224 | | 0,29053295 | 0,779126106 | 7,883361459 | 0,88877587 | 1 |
| ENSGALG00000010258 | LTBP2 | 1,700141471 | 1,689599283 | 7,343564576 | 2,281390244 | 1 |
| ENSGALG00000010282 | TMEM63B | 53,53998682 | 52,35379756 | 67,85386895 | 55,3585158 | 1 |
| ENSGALG00000010318 | TMED10 | 380,0260003 | 378,4420008 | 456,2386739 | 383,0242366 | 1 |
| ENSGALG00000010412 | WWTR1 | 7,372640458 | 8,212018771 | 15,65426723 | 9,207581713 | 1 |
| ENSGALG00000010435 | RASGRP3 | 0,951722698 | 1,288332265 | 16,04572445 | 2,667849388 | 1 |
| ENSGALG00000010468 | ISM2 | 6,961603369 | 5,451403991 | 28,88288749 | 9,016585179 | 1 |
| ENSGALG00000010522 | | 52,19125685 | 45,4437284 | 88,95246599 | 53,90163476 | 1 |
| ENSGALG00000010642 | IRF2 | 14,12465929 | 15,27261891 | 20,26270283 | 12,94760591 | 1 |
| ENSGALG00000010683 | | 1,286887376 | 2,082891848 | 4,318730895 | 2,217138302 | 1 |
| ENSGALG00000010688 | SHC3 | 0,377086245 | 0,473231752 | 1,28457644 | 0,656462583 | 1 |
| ENSGALG00000010692 | LRP8 | 17,33417611 | 14,1459726 | 39,14255891 | 19,74069674 | 1 |
| ENSGALG00000010707 | GPR68 | 0 | 0,159372632 | 1,201854665 | 0,204226743 | 1 |
| ENSGALG00000010785 | ACOT11 | 3,3120391 | 3,449588553 | 7,445190708 | 4,169283534 | 1 |
| ENSGALG00000010926 | | 1,182613457 | 1,250155615 | 4,861891125 | 1,667707956 | 1 |
| ENSGALG00000011099 | FAP | 0,524872973 | 0,627940647 | 8,568804109 | 0,852104076 | 1 |
| ENSGALG00000011110 | | 1,049280017 | 1,070578153 | 3,569578228 | 1,155450113 | 1 |
| ENSGALG00000011120 | | 0,903887543 | 1,717288396 | 2,968494683 | 1,501450921 | 1 |
| ENSGALG00000011148 | | 10,04557421 | 13,49083348 | 31,35368024 | 9,031119595 | 1 |
| ENSGALG00000011172 | | 7,443800272 | 6,657256083 | 14,36492915 | 6,634505832 | 1 |
| ENSGALG00000011207 | DUSP6 | 38,54581524 | 33,84315846 | 198,3618888 | 73,6726962 | 1 |
| ENSGALG00000011236 | BEGAIN | 0,363396332 | 1,174831529 | 2,353729205 | 0,884245626 | 1 |
| ENSGALG00000011246 | | 34,4951174 | 37,48548023 | 47,02695891 | 34,71936867 | 1 |
| ENSGALG00000011258 | | 115,2808442 | 105,6819299 | 225,5885638 | 117,6950123 | 1 |
| ENSGALG00000011336 | | 0,100777952 | 0,154515943 | 0,745110987 | 0,176260961 | 1 |
| ENSGALG00000011376 | ANKRD9 | 5,699927441 | 6,47582585 | 10,76381249 | 5,70463788 | 1 |
| ENSGALG00000011439 | | 11,48455465 | 11,97057423 | 28,13859794 | 12,9402475 | 1 |
| ENSGALG00000011468 | IGFBP5 | 22,18372906 | 31,8043152 | 67,63240972 | 24,25782653 | 1 |
| ENSGALG00000011548 | PPP1R13B | 17,59096224 | 18,42868364 | 27,31810324 | 20,71203786 | 1 |
| ENSGALG00000011562 | PARK2 | 2,307166008 | 2,209921812 | 14,10410652 | 1,800517372 | 1 |
| ENSGALG00000011581 | KIF26A | 4,38085147 | 8,865650561 | 49,95121099 | 16,48311585 | 1 |

| | | | | | | |
|---|---|---|---|---|---|---|
| ENSGALG00000011609 | | 9,843500616 | 8,417625726 | 40,27301602 | 8,233732841 | 1 |
| ENSGALG00000011618 | ADSSL1 | 8,786398246 | 10,75524319 | 18,1647197 | 11,87110001 | 1 |
| ENSGALG00000011623 | ADAMTS3 | 24,331848 | 27,29215191 | 78,31157876 | 35,78867135 | 1 |
| ENSGALG00000011707 | | 53,24117172 | 52,60615779 | 73,60572574 | 52,46461184 | 1 |
| ENSGALG00000011762 | UBA6 | 45,15145647 | 45,49374679 | 54,61323703 | 46,15736387 | 1 |
| ENSGALG00000011770 | UMPS | 129,7844633 | 135,257901 | 207,8820378 | 147,8994428 | 1 |
| ENSGALG00000011867 | SERHL2 | 29,43272669 | 33,34712098 | 71,24407539 | 29,95176995 | 1 |
| ENSGALG00000011940 | | 2,58677773 | 5,2650269 | 34,93790074 | 11,23222133 | 1 |
| ENSGALG00000012018 | UGT8 | 3,339674913 | 5,868867271 | 29,87840982 | 10,482522 | 1 |
| ENSGALG00000012051 | | 4,041058882 | 2,27262952 | 31,35683641 | 3,147836465 | 1 |
| ENSGALG00000012106 | SCTR | 0,647796476 | 1,062662762 | 4,974582478 | 0,728529092 | 1 |
| ENSGALG00000012439 | | 5,987859862 | 8,057194256 | 15,71567866 | 7,213637245 | 1 |
| ENSGALG00000012454 | CYTH4 | 0,040933627 | 0,148295635 | 0,646186586 | 0,122442754 | 1 |
| ENSGALG00000012522 | PVALB | 0,10415029 | 0,018320357 | 1,845669062 | 0,264205542 | 1 |
| ENSGALG00000012533 | | 82,14235353 | 89,9220224 | 161,7487773 | 106,8315371 | 1 |
| ENSGALG00000012608 | DAPK1 | 18,63601716 | 17,25242431 | 24,89907156 | 19,04902754 | 1 |
| ENSGALG00000012623 | | 0,308723784 | 0,17685494 | 2,783031501 | 0,236507182 | 1 |
| ENSGALG00000012635 | | 2,956578448 | 4,062704068 | 12,89063246 | 3,64967825 | 1 |
| ENSGALG00000012732 | PHACTR1 | 1,002723787 | 0,92758825 | 5,310014441 | 0,874633616 | 1 |
| ENSGALG00000012808 | CDYL | 18,02096582 | 21,34276206 | 37,17353321 | 21,75666775 | 1 |
| ENSGALG00000012848 | | 20,64815843 | 19,22874436 | 38,64533737 | 23,70889337 | 1 |
| ENSGALG00000012849 | SLC37A3 | 25,5841736 | 29,25564121 | 46,57867282 | 31,70186837 | 1 |
| ENSGALG00000012911 | SYT10 | 1,659882445 | 2,809814019 | 5,964646057 | 2,833486186 | 1 |
| ENSGALG00000012979 | | 22,57549197 | 22,92713118 | 34,79249726 | 24,59961237 | 1 |
| ENSGALG00000012998 | WNT5B | 32,51568668 | 42,44877134 | 79,13605981 | 43,44051816 | 1 |
| ENSGALG00000013033 | | 0,623034671 | 0,523981207 | 6,611512329 | 0,624842166 | 1 |
| ENSGALG00000013102 | COBL | 1,89987265 | 2,53428906 | 12,26309687 | 3,628837952 | 1 |
| ENSGALG00000013149 | MOCOS | 0,31264351 | 0,69552403 | 1,148089973 | 0,526605797 | 1 |
| ENSGALG00000013406 | EOGT | 8,979738398 | 9,102324177 | 13,64010251 | 10,0250873 | 1 |
| ENSGALG00000013548 | GZMA | 0,244325018 | 0,143724945 | 1,250994806 | 0,095859771 | 1 |
| ENSGALG00000013624 | | 2,042343298 | 4,425049521 | 70,07747071 | 5,032989524 | 1 |
| ENSGALG00000013661 | MICU3 | 16,86990908 | 18,35601379 | 24,39035699 | 18,72105289 | 1 |
| ENSGALG00000013683 | ARID1B | 50,21288267 | 47,56895592 | 91,55652092 | 53,76612448 | 1 |
| ENSGALG00000013892 | MAP3K5 | 3,106959802 | 3,891469812 | 7,477861463 | 3,385152426 | 1 |
| ENSGALG00000014159 | PACSIN2 | 46,93642045 | 52,69096243 | 231,699989 | 66,05762406 | 1 |
| ENSGALG00000014165 | | 34,4949156 | 36,21634446 | 46,38287449 | 35,66651524 | 1 |
| ENSGALG00000014213 | PRR5 | 45,61960191 | 49,63409703 | 60,32943821 | 40,69527126 | 1 |
| ENSGALG00000014337 | NWD2 | 4,063259347 | 3,87414171 | 6,001585157 | 3,902325985 | 1 |
| ENSGALG00000014370 | | 25,6946956 | 26,45741685 | 32,9477035 | 27,52616136 | 1 |
| ENSGALG00000014688 | ZYX | 73,08300226 | 79,10890291 | 102,6158311 | 80,01024875 | 1 |
| ENSGALG00000014783 | | 22,3498316 | 24,871629 | 42,37641705 | 29,08458804 | 1 |
| ENSGALG00000014823 | | 0 | 0,035377845 | 0,977269445 | 0,046829328 | 1 |
| ENSGALG00000014852 | | 58,83953299 | 69,54541567 | 178,8283916 | 76,07766834 | 1 |
| ENSGALG00000014855 | | 27,82040525 | 38,69420984 | 56,11193291 | 36,63572912 | 1 |
| ENSGALG00000014860 | YES1 | 130,3824873 | 130,6854913 | 192,983834 | 128,1750395 | 1 |
| ENSGALG00000014864 | | 3,731602592 | 5,046367263 | 9,359766476 | 4,111091192 | 1 |
| ENSGALG00000014866 | | 1161,822033 | 1014,002441 | 2650,648004 | 1063,272464 | 1 |
| ENSGALG00000014872 | FGF10 | 1,077530612 | 0,935843353 | 31,13159693 | 6,221341059 | 1 |
| ENSGALG00000014877 | EMB | 13,70340987 | 18,5752455 | 42,8108232 | 19,61514892 | 1 |
| ENSGALG00000014907 | DCBLD1 | 14,02744527 | 19,4714576 | 31,49340589 | 18,13536669 | 1 |
| ENSGALG00000014930 | ENC1 | 32,6260464 | 23,96935622 | 95,15822903 | 28,5198255 | 1 |
| ENSGALG00000014953 | RWDD1 | 34,3211322 | 35,70428677 | 48,85056151 | 36,89366799 | 1 |
| ENSGALG00000014957 | WDR1 | 149,3387297 | 141,564327 | 241,426839 | 140,1462197 | 1 |
| ENSGALG00000014961 | | 0,030063775 | 0,212281951 | 0,53262582 | 0,159553166 | 1 |
| ENSGALG00000014983 | F2R | 11,99732554 | 10,51644411 | 30,61968372 | 11,21982229 | 1 |
| ENSGALG00000014984 | F2RL1 | 9,191228695 | 7,297000019 | 60,18915078 | 7,334530663 | 1 |
| ENSGALG00000014994 | CRHBP | 0,094206422 | 0,123018797 | 8,155312142 | 0,069432825 | 1 |
| ENSGALG00000015008 | ZNF366 | 0,025612818 | 0,007984965 | 0,218726746 | 0,02805032 | 1 |
| ENSGALG00000015028 | | 5,215749923 | 6,301235219 | 16,40492276 | 7,825223421 | 1 |
| ENSGALG00000015033 | REV3L | 14,90469289 | 16,2960558 | 20,42678903 | 15,73038983 | 1 |
| ENSGALG00000015089 | GAP43 | 10,73590963 | 9,303884931 | 33,04526973 | 8,030676006 | 1 |
| ENSGALG00000015208 | ROR2 | 17,52597629 | 20,97201143 | 36,98791655 | 21,6549491 | 1 |
| ENSGALG00000015252 | ST3GAL6 | 50,15692337 | 52,54417622 | 62,67006181 | 44,72893 | 1 |
| ENSGALG00000015256 | PAM | 37,6976856 | 29,97280604 | 96,16475159 | 36,54825142 | 1 |

| | | | | | | |
|---|---|---|---|---|---|---|
| ENSGALG00000015332 | CPLX1 | 1,265996124 | 4,716193333 | 92,44153266 | 18,97429343 | 1 |
| ENSGALG00000015358 | MYH15 | 2,502155814 | 2,562264429 | 24,52524418 | 2,753567038 | 1 |
| ENSGALG00000015388 | | 0,998528448 | 2,80672025 | 22,77458166 | 7,945924483 | 1 |
| ENSGALG00000015439 | SLC44A1 | 37,1490751 | 42,22953839 | 47,91821194 | 36,92039081 | 1 |
| ENSGALG00000015440 | | 0,649690061 | 0,632601394 | 1,775464964 | 0,785617529 | 1 |
| ENSGALG00000015458 | ILDR2 | 95,9535283 | 99,09522051 | 139,284134 | 104,0358316 | 1 |
| ENSGALG00000015624 | VCAN | 33,86551135 | 34,75318733 | 94,61693612 | 47,34830969 | 1 |
| ENSGALG00000015634 | | 0,190327277 | 0,320177513 | 1,214767371 | 0,43293815 | 1 |
| ENSGALG00000015665 | SH3BP2 | 3,286250973 | 2,895344261 | 9,524027287 | 4,083976177 | 1 |
| ENSGALG00000015826 | HTR1E | 0,33449219 | 0,44224444 | 11,45369572 | 1,027953514 | 1 |
| ENSGALG00000015896 | | 6,388186822 | 7,197999605 | 12,51939522 | 6,10029628 | 1 |
| ENSGALG00000015947 | | 0,048261531 | 0,051324674 | 8,627025826 | 1,412173952 | 1 |
| ENSGALG00000015975 | | 46,39913828 | 46,69943177 | 77,97746804 | 37,87795667 | 1 |
| ENSGALG00000016074 | | 2,965663767 | 3,518415951 | 4,36951983 | 3,22716103 | 1 |
| ENSGALG00000016102 | | 1,474592182 | 2,407557081 | 8,858325248 | 2,452322765 | 1 |
| ENSGALG00000016140 | FAM3B | 1,345511674 | 1,561374658 | 14,90793732 | 1,339776747 | 1 |
| ENSGALG00000016145 | | 0,594265981 | 1,380102509 | 16,360371 | 2,19880009 | 1 |
| ENSGALG00000016226 | | 82,42631993 | 87,8153227 | 97,22152217 | 82,97244346 | 1 |
| ENSGALG00000016236 | | 79,83962807 | 81,6981472 | 94,32375099 | 74,79813666 | 1 |
| ENSGALG00000016297 | HCRTR2 | 0,172470075 | 0,189519273 | 0,91709899 | 0,109463532 | 1 |
| ENSGALG00000016377 | PXDN | 48,44717284 | 47,31975564 | 221,1364094 | 53,85481487 | 1 |
| ENSGALG00000016431 | Pax5B | 0,759670645 | 1,184984006 | 24,88772514 | 0,76701086 | 1 |
| ENSGALG00000016470 | KCNS3 | 0,820548787 | 1,835471231 | 21,82469461 | 5,812307253 | 1 |
| ENSGALG00000016537 | SCML2 | 23,96103489 | 29,8195471 | 50,09057341 | 29,94459128 | 1 |
| ENSGALG00000016602 | | 3,664177334 | 3,917626255 | 12,58470849 | 4,88155337 | 1 |
| ENSGALG00000016630 | EXTL3 | 78,945634 | 78,16521266 | 100,6801361 | 82,53544774 | 1 |
| ENSGALG00000016736 | CYFIP1 | 115,4849451 | 117,5058947 | 151,3780148 | 125,0449365 | 1 |
| ENSGALG00000016751 | | 74,54427214 | 79,14506937 | 102,3489998 | 80,62212588 | 1 |
| ENSGALG00000016754 | | 6,519633751 | 9,835468627 | 20,77499853 | 9,953787953 | 1 |
| ENSGALG00000016768 | LONRF2 | 44,67796838 | 42,46064416 | 63,91927301 | 44,69491354 | 1 |
| ENSGALG00000016781 | MAP4K4 | 80,99823221 | 79,95663486 | 120,0802219 | 86,66172865 | 1 |
| ENSGALG00000016783 | | 0,807775429 | 0,850085852 | 114,9806394 | 2,530944487 | 1 |
| ENSGALG00000016784 | | 0,558045454 | 0,472553521 | 7,610637786 | 1,133090803 | 1 |
| ENSGALG00000016785 | IL1RL1 | 0,070474547 | 0 | 1,971779618 | 0,162809382 | 1 |
| ENSGALG00000016834 | MCF2L | 4,225420748 | 4,647949468 | 6,579407696 | 4,100185155 | 1 |
| ENSGALG00000016835 | | 19,91459988 | 25,58593002 | 33,27889918 | 23,30715898 | 1 |
| ENSGALG00000016883 | DOCK9 | 10,45434421 | 10,83646735 | 15,52539363 | 10,77548298 | 1 |
| ENSGALG00000016893 | | 30,0481437 | 30,75098399 | 36,9917239 | 30,50750363 | 1 |
| ENSGALG00000016899 | DCT | 0,082019435 | 0,067986713 | 0,424223435 | 0,074974082 | 1 |
| ENSGALG00000016906 | SPRY2 | 22,79491815 | 23,80423911 | 369,1836197 | 101,4167028 | 1 |
| ENSGALG00000016911 | RNF219 | 35,29870957 | 34,99094267 | 44,02485713 | 33,4703359 | 1 |
| ENSGALG00000016998 | LPAR6 | 5,959033195 | 5,246978391 | 17,44700786 | 7,250724714 | 1 |
| ENSGALG00000017044 | TRPC4 | 0,533180653 | 0,692407894 | 1,395530609 | 0,748381254 | 1 |
| ENSGALG00000017091 | FLT1 | 1,390041035 | 0,693832705 | 8,294413787 | 1,492735256 | 1 |
| ENSGALG00000017135 | CRYL1 | 4,9261885 | 4,709864357 | 10,17588307 | 4,813391563 | 1 |
| ENSGALG00000017179 | PDGFD | 2,173029014 | 3,505795831 | 14,26690549 | 3,894835223 | 1 |
| ENSGALG00000017186 | | 69,04285483 | 68,94924482 | 88,45648678 | 73,22242114 | 1 |
| ENSGALG00000017242 | FZD4 | 4,01523892 | 5,003599185 | 20,09846171 | 6,618181415 | 1 |
| ENSGALG00000017244 | PRSS23 | 12,28501908 | 11,45283797 | 64,02403712 | 13,46351005 | 1 |
| ENSGALG00000017714 | | 2,458678092 | 3,270301188 | 9,373616504 | 4,3477783 | 1 |
| ENSGALG00000018479 | | 19,30607593 | 27,19022439 | 34,32889124 | 17,26565929 | 1 |
| ENSGALG00000019261 | | 6,423634206 | 5,898459466 | 17,86083615 | 5,032292518 | 1 |
| ENSGALG00000019397 | CERK | 65,45684989 | 75,63715367 | 102,1461158 | 52,57771362 | 1 |
| ENSGALG00000019426 | | 1,895961953 | 1,778982812 | 8,767179842 | 1,135546894 | 1 |
| ENSGALG00000019552 | SERPINB2 | 0,03276452 | 0,012366785 | 0,305019782 | 0,020558652 | 1 |
| ENSGALG00000019716 | KRT20 | 0,427331711 | 0,54228415 | 2,142286477 | 0,421447442 | 1 |
| ENSGALG00000020057 | PIGF | 47,49119487 | 53,61856839 | 63,6504531 | 43,06229201 | 1 |
| ENSGALG00000020359 | SIX6 | 0,147864328 | 0,28328832 | 7,275442564 | 0,245244412 | 1 |
| ENSGALG00000020723 | | 23,95654484 | 27,95652952 | 51,89848439 | 27,20320324 | 1 |
| ENSGALG00000021238 | | 3,377343555 | 3,609177279 | 6,58959715 | 4,12451232 | 1 |
| ENSGALG00000021879 | | 0,060680129 | 0,02783627 | 4,532687997 | 0,361707989 | 1 |
| ENSGALG00000022871 | | 9,149877167 | 11,59069382 | 20,94301968 | 8,951752944 | 1 |
| ENSGALG00000022962 | | 0,005547593 | 0 | 1,110790954 | 0,021210214 | 1 |
| ENSGALG00000023355 | TMEM154 | 0,274198999 | 0,242777653 | 1,595469388 | 0,503883204 | 1 |

| | | | | | | |
|---|---|---|---|---|---|---|
| ENSGALG00000023379 | F2RL2 | 0,699357483 | 0,206509379 | 17,20987384 | 0,770639649 | 1 |
| ENSGALG00000023406 | | 7,189218224 | 27,88560585 | 237,0573377 | 66,09046748 | 1 |
| ENSGALG00000023430 | | 0,204871896 | 0,107896168 | 5,766235294 | 0,103839874 | 1 |
| ENSGALG00000023510 | ERP29 | 244,8276365 | 243,7579874 | 286,6547438 | 250,6109926 | 1 |
| ENSGALG00000023521 | ADRA2A | 2,990826686 | 6,4207327 | 33,46692465 | 9,885743021 | 1 |
| ENSGALG00000023581 | | 2,293508573 | 3,632286312 | 6,833722653 | 3,344298943 | 1 |
| ENSGALG00000024083 | C8orf4 | 24,35008397 | 20,56522548 | 90,61964264 | 18,98861546 | 1 |
| ENSGALG00000025723 | | 0,558060232 | 0,679196686 | 0,810932422 | 0,573634828 | 1 |
| ENSGALG00000025773 | | 12,25494928 | 11,44694026 | 30,15620585 | 9,759339323 | 1 |
| ENSGALG00000025941 | RGS20 | 0,992958806 | 1,120198002 | 4,961879868 | 1,076166451 | 1 |
| ENSGALG00000026262 | | 23,02651413 | 19,44179408 | 89,23902687 | 31,06609278 | 1 |
| ENSGALG00000026366 | | 0,032450796 | 0,03560783 | 0,297910888 | 0,062257621 | 1 |
| ENSGALG00000026518 | RUNDC3A | 10,57114969 | 16,46754486 | 25,12660637 | 14,66077777 | 1 |
| ENSGALG00000026557 | PGF | 1,365378945 | 1,285576654 | 6,812213801 | 1,989316648 | 1 |
| ENSGALG00000026582 | | 6,114977984 | 5,267516926 | 12,31174473 | 6,045225829 | 1 |
| ENSGALG00000026663 | CX3CL1 | 0,28585624 | 0,517672331 | 5,224016511 | 0,810407917 | 1 |
| ENSGALG00000026753 | | 4,373338809 | 4,733385198 | 6,354129053 | 4,718191959 | 1 |
| ENSGALG00000026763 | | 5,105466174 | 4,675186082 | 12,43728303 | 5,145242205 | 1 |
| ENSGALG00000026790 | DMRT2 | 2,44415194 | 3,013071402 | 9,651534911 | 4,741219823 | 1 |
| ENSGALG00000026892 | | 0,630709919 | 0,49426779 | 2,669713204 | 0,343635403 | 1 |
| ENSGALG00000026970 | | 19,9958411 | 29,56495965 | 47,72673087 | 27,76473883 | 1 |
| ENSGALG00000026995 | RHOG | 2,125612682 | 1,771546724 | 5,966320997 | 2,254397519 | 1 |
| ENSGALG00000027133 | | 0,276818664 | 0,334539577 | 1,403416102 | 0,364180584 | 1 |
| ENSGALG00000027210 | | 0,626776305 | 0,767218066 | 2,251672836 | 0,943311678 | 1 |
| ENSGALG00000027497 | | 4,239074936 | 4,349860411 | 10,12003694 | 4,862962086 | 1 |
| ENSGALG00000027864 | NFKBIA | 5,145580181 | 5,317887462 | 7,594774216 | 5,76461762 | 1 |
| ENSGALG00000027885 | C4orf19 | 0,006715072 | 0,052136259 | 0,371881758 | 0,049775385 | 1 |
| ENSGALG00000028046 | INSIG2 | 47,56660897 | 56,64100654 | 107,3362206 | 50,7535001 | 1 |
| ENSGALG00000028047 | RHOV | 2,271837176 | 2,028232459 | 7,468783549 | 2,647132976 | 1 |
| ENSGALG00000028250 | ANKRD22 | 0,112085424 | 0,162263291 | 0,899150004 | 0,041047018 | 1 |
| ENSGALG00000028267 | | 76,90546701 | 75,83970622 | 103,3050563 | 76,29588931 | 1 |
| ENSGALG00000028291 | | 96,11071203 | 93,78119127 | 144,7372696 | 92,58940493 | 1 |
| ENSGALG00000028294 | VWA5B2 | 14,51886882 | 15,87519135 | 30,51008951 | 18,72172895 | 1 |
| ENSGALG00000028637 | | 37,10791538 | 39,94856328 | 50,36322451 | 41,0166067 | 1 |
| ENSGALG00000028708 | | 24,99318737 | 36,73989626 | 54,90494637 | 28,8846325 | 1 |
| ENSGALG00000028938 | | 0,513837926 | 0,624776649 | 0,839919222 | 0,435033808 | 1 |
| ENSGALG00000029175 | EFNB2 | 30,56530828 | 27,21275717 | 113,8041773 | 33,97932717 | 1 |
| ENSGALG00000000195 | SVCT2 | 19,10462588 | 17,48116214 | 25,91142744 | 17,63462036 | 2 |
| ENSGALG00000000208 | MCC | 25,83310361 | 20,09476945 | 33,82276296 | 23,91037382 | 2 |
| ENSGALG00000000258 | | 75,85689269 | 54,92413434 | 95,52174596 | 62,32390544 | 2 |
| ENSGALG00000000275 | KCTD9 | 37,75610815 | 32,89373192 | 45,75008795 | 35,24739442 | 2 |
| ENSGALG00000000332 | | 72,56124041 | 58,37784284 | 85,08368566 | 66,53036384 | 2 |
| ENSGALG00000000346 | | 75,70929419 | 72,18519081 | 95,30895247 | 71,18874501 | 2 |
| ENSGALG00000000388 | | 173,5309683 | 83,88403469 | 265,5077681 | 88,686351 | 2 |
| ENSGALG00000000680 | | 65,55002426 | 35,12675538 | 98,37170682 | 37,34008803 | 2 |
| ENSGALG00000000936 | | 18,6039239 | 13,94226071 | 27,03075663 | 11,68456073 | 2 |
| ENSGALG00000001000 | | 903,1481891 | 710,3094654 | 1580,638121 | 839,8439333 | 2 |
| ENSGALG00000001003 | SEPN1 | 54,28402873 | 50,00244607 | 70,19927743 | 47,23413232 | 2 |
| ENSGALG00000001169 | CRB2 | 129,1232018 | 81,31040842 | 173,364492 | 61,90998035 | 2 |
| ENSGALG00000001192 | | 56,80326116 | 47,24213542 | 66,68178678 | 43,96666792 | 2 |
| ENSGALG00000001276 | HOXB9 | 440,9241521 | 286,4552462 | 833,8511125 | 381,4973378 | 2 |
| ENSGALG00000001363 | MYBBP1A | 108,2631392 | 74,57137633 | 129,5761289 | 87,68932285 | 2 |
| ENSGALG00000001608 | UNC5D | 1,806421005 | 1,336857976 | 5,395223323 | 1,082125149 | 2 |
| ENSGALG00000001645 | KCNT1 | 3,342845418 | 2,419654306 | 6,945138399 | 2,085754068 | 2 |
| ENSGALG00000001656 | | 187,6401815 | 129,4403167 | 226,396159 | 121,0670172 | 2 |
| ENSGALG00000001749 | ACSBG2 | 25,77749834 | 18,04184176 | 33,32440771 | 18,2584482 | 2 |
| ENSGALG00000001768 | TENM2 | 13,34961314 | 7,569539699 | 19,32663023 | 7,163569584 | 2 |
| ENSGALG00000001957 | | 30,17853095 | 28,11283549 | 41,98257618 | 29,43069139 | 2 |
| ENSGALG00000002119 | C16orf62 | 53,57445126 | 51,96391471 | 66,2690369 | 50,53612293 | 2 |
| ENSGALG00000002316 | NUBP2 | 79,87979134 | 66,76426362 | 113,9079487 | 71,91374142 | 2 |
| ENSGALG00000002397 | PSPH | 16,90500002 | 11,13674451 | 38,37209435 | 14,53627012 | 2 |
| ENSGALG00000002443 | FBXL16 | 5,482262774 | 3,393627985 | 10,2747505 | 3,721226607 | 2 |
| ENSGALG00000002460 | | 169,5656604 | 134,6549735 | 199,9940187 | 147,1796955 | 2 |
| ENSGALG00000002461 | | 85,54594955 | 74,49743937 | 129,1652031 | 74,457145 | 2 |

| | | | | | | |
|---|---|---|---|---|---|---|
| ENSGALG00000002714 | TMEM132C | 81,17568898 | 53,4953987 | 178,5879937 | 49,72398397 | 2 |
| ENSGALG00000002726 | KIF1B | 37,34066731 | 35,19418972 | 52,6551427 | 32,73497754 | 2 |
| ENSGALG00000002923 | | 42,50041281 | 25,78454685 | 58,32063392 | 24,5275742 | 2 |
| ENSGALG00000002985 | UTP18 | 59,80283029 | 50,99843338 | 74,38386445 | 58,7986555 | 2 |
| ENSGALG00000003052 | AGFG1 | 53,98350241 | 52,12101214 | 63,9794555 | 53,31865967 | 2 |
| ENSGALG00000003078 | MMD | 166,6104847 | 155,0883677 | 202,3376044 | 148,475486 | 2 |
| ENSGALG00000003180 | | 107,8427517 | 81,65197474 | 140,4725058 | 96,80702055 | 2 |
| ENSGALG00000003277 | SUZ12 | 67,78173578 | 55,86030796 | 80,4002124 | 62,8663404 | 2 |
| ENSGALG00000003295 | | 11,95332143 | 9,200781695 | 16,90039539 | 8,047388377 | 2 |
| ENSGALG00000003375 | | 0,555521759 | 0,504517259 | 0,641921993 | 0,451150123 | 2 |
| ENSGALG00000003391 | EIF2AK1 | 120,3525644 | 98,69879665 | 149,048697 | 85,56996995 | 2 |
| ENSGALG00000003427 | USP3 | 144,6869467 | 126,2563974 | 186,8105836 | 108,1291213 | 2 |
| ENSGALG00000003430 | PMS2 | 83,93336261 | 66,63608403 | 99,6035854 | 60,47076563 | 2 |
| ENSGALG00000003566 | SMURF2 | 22,13122327 | 21,05696082 | 28,29312846 | 20,18762732 | 2 |
| ENSGALG00000003739 | | 107,7186679 | 79,57676443 | 217,3446885 | 92,92358891 | 2 |
| ENSGALG00000003767 | NKD1 | 64,58229372 | 56,04338743 | 82,66219062 | 54,31127427 | 2 |
| ENSGALG00000003835 | | 95,26245909 | 80,97210783 | 135,4218017 | 94,66924992 | 2 |
| ENSGALG00000003957 | | 7,497005362 | 7,280862747 | 12,32775655 | 4,077169463 | 2 |
| ENSGALG00000004049 | ITFG1 | 143,3658699 | 143,1696704 | 169,0452251 | 126,3388766 | 2 |
| ENSGALG00000004121 | LPAR4 | 57,41190777 | 51,21366709 | 73,35902376 | 56,01058859 | 2 |
| ENSGALG00000004132 | | 313,0280743 | 222,5345954 | 396,1457648 | 232,4516201 | 2 |
| ENSGALG00000004144 | | 230,9295811 | 165,493935 | 322,5375425 | 219,6317991 | 2 |
| ENSGALG00000004169 | | 52,0041871 | 45,80321214 | 71,82848643 | 48,04815439 | 2 |
| ENSGALG00000004231 | IFNLR1 | 13,20283795 | 12,15973039 | 22,07165879 | 11,62321498 | 2 |
| ENSGALG00000004416 | TTC19 | 35,18587556 | 33,81562749 | 46,08392978 | 33,45574377 | 2 |
| ENSGALG00000004435 | RAB27A | 37,01217138 | 30,70100511 | 46,40397428 | 36,08556376 | 2 |
| ENSGALG00000004474 | PLEKHF1 | 16,35912847 | 14,33700972 | 22,89893142 | 16,04362022 | 2 |
| ENSGALG00000004813 | IARS | 121,2613046 | 91,77804842 | 145,089873 | 101,0949477 | 2 |
| ENSGALG00000005123 | WNT7A | 12,40158553 | 7,623339638 | 21,37318743 | 7,362852482 | 2 |
| ENSGALG00000005177 | | 102,6782147 | 98,37619348 | 120,4338593 | 98,08612871 | 2 |
| ENSGALG00000005243 | PPP3CB | 90,43009538 | 80,11505572 | 110,9801319 | 75,05008011 | 2 |
| ENSGALG00000005425 | OGT | 218,3232715 | 170,7793164 | 269,4024715 | 191,9762176 | 2 |
| ENSGALG00000005427 | | 2,231810001 | 1,699165441 | 2,481654149 | 1,214604019 | 2 |
| ENSGALG00000005601 | | 0,874226757 | 0,788269723 | 1,394307851 | 0,68739779 | 2 |
| ENSGALG00000005685 | KSR1 | 28,99559087 | 18,12035724 | 39,2301589 | 17,04233962 | 2 |
| ENSGALG00000005689 | Pax-2 | 2,427170557 | 1,41090904 | 9,598393861 | 1,637819447 | 2 |
| ENSGALG00000005723 | | 20,77746675 | 15,4473937 | 28,34501666 | 15,42480292 | 2 |
| ENSGALG00000005796 | XBP1 | 178,2861977 | 149,6817159 | 240,7969177 | 155,1233149 | 2 |
| ENSGALG00000005815 | TMEM41B | 103,3957725 | 81,41002257 | 126,0115268 | 83,74442495 | 2 |
| ENSGALG00000005845 | SLC7A5 | 122,4599338 | 88,23427206 | 160,3761058 | 103,8001208 | 2 |
| ENSGALG00000005854 | CLUH | 53,12207763 | 46,43056255 | 64,52154409 | 48,66611182 | 2 |
| ENSGALG00000005945 | | 312,5580735 | 266,4501752 | 377,6849137 | 289,237717 | 2 |
| ENSGALG00000005966 | SEC61A1 | 146,4192137 | 135,8133269 | 172,2380178 | 140,0010865 | 2 |
| ENSGALG00000006080 | GPC4 | 336,1866881 | 277,2326888 | 398,7191432 | 255,9354884 | 2 |
| ENSGALG00000006138 | | 5,3558423 | 4,222816208 | 9,559136283 | 4,692914386 | 2 |
| ENSGALG00000006237 | PKN2 | 94,14155442 | 80,04499622 | 133,7485665 | 89,13597808 | 2 |
| ENSGALG00000006254 | ERCC6 | 20,09113345 | 17,29890431 | 24,82822974 | 19,87584242 | 2 |
| ENSGALG00000006256 | | 81,30960437 | 70,77595812 | 104,1957411 | 69,99809839 | 2 |
| ENSGALG00000006336 | | 24,78170301 | 21,0855019 | 32,50165931 | 24,28317002 | 2 |
| ENSGALG00000006378 | | 46,64016235 | 43,92374135 | 60,27231414 | 45,47990755 | 2 |
| ENSGALG00000006437 | | 21,70370986 | 15,95524381 | 28,04172555 | 17,79255113 | 2 |
| ENSGALG00000006797 | ADARB2 | 2,386525484 | 1,723097368 | 5,609955946 | 1,217344116 | 2 |
| ENSGALG00000006822 | PCDH19 | 27,11782022 | 17,17899723 | 78,54913552 | 16,55189982 | 2 |
| ENSGALG00000006856 | CD151 | 66,08781237 | 56,83548982 | 81,90386525 | 57,64667081 | 2 |
| ENSGALG00000006872 | PISD | 43,47551418 | 38,78352232 | 70,45275748 | 40,42254893 | 2 |
| ENSGALG00000006895 | CEP131 | 90,4838243 | 72,05921603 | 111,7343496 | 78,83214694 | 2 |
| ENSGALG00000006935 | | 41,19849281 | 36,85474382 | 50,28356686 | 36,31503506 | 2 |
| ENSGALG00000006939 | PFKP | 159,5195891 | 114,2592094 | 248,2482576 | 153,3129457 | 2 |
| ENSGALG00000006971 | | 39,6698917 | 31,40577651 | 49,95181554 | 34,88964426 | 2 |
| ENSGALG00000007141 | LRRK1 | 8,471065321 | 7,23115594 | 11,52135294 | 7,124193969 | 2 |
| ENSGALG00000007145 | ITGB1 | 273,7290161 | 235,6454154 | 355,8809593 | 269,2892165 | 2 |
| ENSGALG00000007234 | CLCN5 | 26,22166298 | 22,62208897 | 33,75980899 | 24,31186251 | 2 |
| ENSGALG00000007555 | CCND1 | 306,4125901 | 154,2072849 | 365,1007316 | 180,3456931 | 2 |
| ENSGALG00000007703 | | 9,24001434 | 8,395210751 | 12,59696711 | 8,329025499 | 2 |

| | | | | | | |
|---|---|---|---|---|---|---|
| ENSGALG00000007870 | | 87,32940421 | 79,94947931 | 110,2506191 | 70,91164374 | 2 |
| ENSGALG00000007992 | GGA3 | 36,45435135 | 33,68958392 | 46,79638773 | 32,15023613 | 2 |
| ENSGALG00000008014 | CEBPB | 4,510096151 | 2,905838226 | 10,42101228 | 4,244775839 | 2 |
| ENSGALG00000008088 | ACSL4 | 42,77845399 | 39,53431922 | 53,52228442 | 40,19984773 | 2 |
| ENSGALG00000008132 | | 65,00574864 | 54,88816517 | 78,94598152 | 54,53613107 | 2 |
| ENSGALG00000008240 | CFLAR | 18,56362706 | 16,85330063 | 24,63252397 | 15,72359014 | 2 |
| ENSGALG00000008265 | | 99,00016091 | 72,4970655 | 124,8646432 | 83,07442811 | 2 |
| ENSGALG00000008352 | | 2,250088005 | 1,0956429 | 4,791149948 | 1,598463212 | 2 |
| ENSGALG00000008376 | EIF2B5 | 94,27576083 | 72,47419873 | 115,9006191 | 78,67487542 | 2 |
| ENSGALG00000008420 | CAND2 | 23,37386731 | 21,50334329 | 30,87205611 | 22,23847698 | 2 |
| ENSGALG00000008491 | | 17,15532317 | 13,9477056 | 36,83972093 | 14,33742773 | 2 |
| ENSGALG00000008547 | | 64,24771593 | 51,30719206 | 82,19229622 | 44,61001661 | 2 |
| ENSGALG00000008615 | | 5,06223098 | 3,526819344 | 8,116738936 | 3,342214797 | 2 |
| ENSGALG00000008705 | | 70,28714953 | 55,76149858 | 84,361465 | 62,76428579 | 2 |
| ENSGALG00000008811 | SLC1A4 | 58,52949193 | 43,75140023 | 86,41435492 | 45,37141837 | 2 |
| ENSGALG00000008837 | PELI1 | 94,23917029 | 88,93657106 | 118,7910083 | 87,71875282 | 2 |
| ENSGALG00000008848 | | 228,2464595 | 192,5521067 | 285,6860563 | 202,9251398 | 2 |
| ENSGALG00000008893 | | 0,788890271 | 0,471371295 | 2,736222886 | 0,392092612 | 2 |
| ENSGALG00000008918 | LAMP5 | 14,93803939 | 12,88300339 | 33,54629897 | 9,821656942 | 2 |
| ENSGALG00000009156 | CDAN1 | 24,72426506 | 21,37975812 | 38,04864069 | 23,8875669 | 2 |
| ENSGALG00000009157 | | 87,96461416 | 76,74072672 | 108,7893047 | 79,4696583 | 2 |
| ENSGALG00000009325 | | 294,7081113 | 263,852235 | 362,0594636 | 288,1555786 | 2 |
| ENSGALG00000009381 | | 32,68110455 | 28,74970946 | 55,76103582 | 28,68002235 | 2 |
| ENSGALG00000009405 | GRIA2 | 1,407370493 | 1,108824914 | 2,875596002 | 1,012364018 | 2 |
| ENSGALG00000009428 | SUSD6 | 57,11791895 | 42,67061228 | 75,42226656 | 43,93842172 | 2 |
| ENSGALG00000009448 | IFRD1 | 202,5579686 | 184,5575928 | 256,5170811 | 193,8512693 | 2 |
| ENSGALG00000009503 | RAB3GAP2 | 27,58247884 | 24,32314054 | 39,58475158 | 27,79781709 | 2 |
| ENSGALG00000009530 | PDZRN4 | 4,501893374 | 4,232231688 | 8,897453929 | 3,14255931 | 2 |
| ENSGALG00000009532 | GXYLT1 | 80,28520746 | 50,71712672 | 113,1318587 | 48,13296544 | 2 |
| ENSGALG00000009540 | YAF2 | 59,24038791 | 44,15994271 | 76,99125086 | 41,14854067 | 2 |
| ENSGALG00000009598 | EPRS | 197,6861501 | 158,8121803 | 232,2356108 | 176,423736 | 2 |
| ENSGALG00000009748 | ASNS | 97,82806709 | 83,16578594 | 131,8122178 | 78,83074589 | 2 |
| ENSGALG00000009782 | NAA15 | 266,9481403 | 207,4047491 | 327,0135056 | 235,6097075 | 2 |
| ENSGALG00000009899 | | 236,5885982 | 203,9108633 | 284,3763821 | 211,3515889 | 2 |
| ENSGALG00000009932 | THADA | 22,14429481 | 19,57605019 | 29,02730205 | 16,89609973 | 2 |
| ENSGALG00000009992 | ARHGAP5 | 58,77117318 | 59,03006064 | 73,92712 | 52,73589474 | 2 |
| ENSGALG00000010185 | HSPA4L | 163,6669681 | 145,5240912 | 193,6623799 | 158,5908649 | 2 |
| ENSGALG00000010344 | | 156,9519341 | 147,9541769 | 185,7383997 | 153,6414155 | 2 |
| ENSGALG00000010481 | CMPK1 | 87,69769168 | 80,49136199 | 107,5540033 | 88,01504488 | 2 |
| ENSGALG00000010536 | FAF1 | 88,30370611 | 72,66469157 | 110,2520086 | 71,68965841 | 2 |
| ENSGALG00000010603 | | 84,1905846 | 68,02532034 | 109,3218398 | 73,68783444 | 2 |
| ENSGALG00000010609 | NDUFAF7 | 58,29146836 | 51,96812842 | 72,7689523 | 57,11232085 | 2 |
| ENSGALG00000010975 | TMTC2 | 23,82759321 | 19,13077263 | 32,54483622 | 21,60811044 | 2 |
| ENSGALG00000010976 | CCDC126 | 54,6027561 | 47,2667848 | 70,21267556 | 52,96789525 | 2 |
| ENSGALG00000011045 | | 26,46809802 | 23,19298715 | 40,20404704 | 23,42656319 | 2 |
| ENSGALG00000011101 | URB2 | 25,42984582 | 20,93888932 | 34,16483045 | 25,23812695 | 2 |
| ENSGALG00000011162 | CYP46A1 | 13,41648813 | 11,27861549 | 19,78883987 | 12,85367535 | 2 |
| ENSGALG00000011228 | WARS | 106,1162631 | 99,35563125 | 133,5656722 | 91,79714635 | 2 |
| ENSGALG00000011280 | EEA1 | 45,16187273 | 37,63511378 | 56,03426363 | 37,5455685 | 2 |
| ENSGALG00000011295 | SOCS2 | 35,2007449 | 27,48311534 | 54,87659139 | 27,05575114 | 2 |
| ENSGALG00000011319 | ADGRL3 | 45,1852269 | 32,57841533 | 77,39214523 | 28,90106474 | 2 |
| ENSGALG00000011351 | | 1685,984803 | 1051,80178 | 2090,592418 | 1455,900582 | 2 |
| ENSGALG00000011357 | VEZT | 46,17530033 | 43,8453212 | 57,83328907 | 44,85175026 | 2 |
| ENSGALG00000011388 | RABGGTB | 110,4609778 | 75,06399545 | 135,3567109 | 76,64938861 | 2 |
| ENSGALG00000011400 | USP44 | 105,190674 | 89,81710894 | 130,7144947 | 91,51130572 | 2 |
| ENSGALG00000011449 | | 22,58118535 | 18,20250459 | 33,94025319 | 22,81520682 | 2 |
| ENSGALG00000011464 | EIF5 | 917,8237588 | 741,5110635 | 1254,948905 | 864,0505476 | 2 |
| ENSGALG00000011541 | BIN1 | 42,772136 | 38,21688698 | 55,05676669 | 37,86617878 | 2 |
| ENSGALG00000011596 | | 57,99322619 | 44,45105626 | 70,97049646 | 50,67792225 | 2 |
| ENSGALG00000011630 | GLI2 | 38,42268045 | 36,6914924 | 48,8882817 | 35,92337252 | 2 |
| ENSGALG00000011643 | UTP20 | 49,30168208 | 34,35447527 | 68,15857187 | 39,08059154 | 2 |
| ENSGALG00000011708 | MYLK | 42,12023537 | 23,42722651 | 64,28850589 | 17,96485165 | 2 |
| ENSGALG00000012012 | | 37,18646392 | 31,88793184 | 65,7915782 | 35,088643 | 2 |
| ENSGALG00000012135 | ATF4 | 797,1581709 | 658,7569115 | 982,7650311 | 706,8909729 | 2 |

| | | | | | | |
|---|---|---|---|---|---|---|
| ENSGALG00000012138 | | 509,5484881 | 456,9957633 | 634,8041273 | 493,2502814 | 2 |
| ENSGALG00000012147 | DDX18 | 65,05833895 | 45,55953755 | 80,75087614 | 54,70066068 | 2 |
| ENSGALG00000012224 | | 12,79412136 | 10,47045165 | 26,42294332 | 11,28060466 | 2 |
| ENSGALG00000012278 | | 69,38557328 | 35,10562561 | 151,1486425 | 26,90469823 | 2 |
| ENSGALG00000012394 | NID2 | 68,69887592 | 50,72697074 | 103,5972114 | 58,40743145 | 2 |
| ENSGALG00000012412 | MTHFD1L | 68,14080099 | 40,88211832 | 86,10529775 | 42,0382293 | 2 |
| ENSGALG00000012447 | | 30,18390029 | 29,50018616 | 37,62561165 | 24,8779874 | 2 |
| ENSGALG00000012594 | NTRK2 | 2,037686782 | 1,161148992 | 4,803911879 | 0,826826974 | 2 |
| ENSGALG00000012726 | | 731,3536309 | 562,0189803 | 1152,278498 | 610,6379714 | 2 |
| ENSGALG00000012807 | | 41,90128094 | 38,20539641 | 52,18993579 | 41,4992427 | 2 |
| ENSGALG00000012814 | PXDC1 | 18,15321703 | 15,98836675 | 24,69297093 | 13,47136744 | 2 |
| ENSGALG00000012834 | AKR1D1 | 11,22123762 | 9,012546744 | 17,6164965 | 9,658495181 | 2 |
| ENSGALG00000012916 | DROSHA | 22,62795344 | 20,36463746 | 27,36524709 | 21,85837675 | 2 |
| ENSGALG00000012927 | | 0,883855398 | 0,584154188 | 2,63363293 | 0,355129019 | 2 |
| ENSGALG00000013124 | FHOD3 | 15,42123161 | 11,36420449 | 27,34521932 | 13,3362274 | 2 |
| ENSGALG00000013203 | | 24,74101095 | 18,16871784 | 30,60325872 | 21,74233169 | 2 |
| ENSGALG00000013480 | | 63,83136272 | 43,0080344 | 89,59781528 | 60,13279899 | 2 |
| ENSGALG00000013861 | TNFAIP3 | 10,67839855 | 9,456256342 | 14,89948611 | 7,83500739 | 2 |
| ENSGALG00000014343 | | 2,461528107 | 1,903728583 | 5,19294882 | 1,881861539 | 2 |
| ENSGALG00000014419 | | 28,48257578 | 26,61598867 | 39,61862765 | 24,2985378 | 2 |
| ENSGALG00000014426 | KPNA1 | 75,53504815 | 58,7970706 | 92,09409276 | 65,46068843 | 2 |
| ENSGALG00000014686 | FBN2 | 3,824183353 | 3,617210217 | 5,312013174 | 3,28868843 | 2 |
| ENSGALG00000014727 | PDE4D | 3,095516476 | 2,120314733 | 9,004244594 | 1,906866262 | 2 |
| ENSGALG00000014784 | MAST4 | 7,644433311 | 6,829537967 | 10,49021956 | 7,439951494 | 2 |
| ENSGALG00000014873 | GJA1 | 151,5200503 | 125,8535441 | 213,8081133 | 139,1697079 | 2 |
| ENSGALG00000014902 | | 36,12871703 | 36,02307188 | 48,16761796 | 31,1982202 | 2 |
| ENSGALG00000015049 | | 107,1047214 | 88,92587671 | 135,8651695 | 102,4480149 | 2 |
| ENSGALG00000015112 | | 52,46535646 | 48,47995695 | 77,60078191 | 38,76419637 | 2 |
| ENSGALG00000015164 | | 8,124039618 | 6,253772744 | 15,92231741 | 8,079666445 | 2 |
| ENSGALG00000015472 | | 103,7455883 | 84,8260084 | 125,5639489 | 82,46251332 | 2 |
| ENSGALG00000015765 | | 57,24314739 | 44,13731219 | 76,00962273 | 51,54508175 | 2 |
| ENSGALG00000015862 | | 174,4258517 | 150,9505279 | 265,4828154 | 159,1555857 | 2 |
| ENSGALG00000015887 | URB1 | 25,51642297 | 20,47718015 | 41,57109807 | 25,13000945 | 2 |
| ENSGALG00000015912 | | 32,16222318 | 29,58832844 | 40,98127953 | 26,04493072 | 2 |
| ENSGALG00000015940 | | 20,73939007 | 19,85946192 | 29,04560533 | 14,93146904 | 2 |
| ENSGALG00000016412 | MBOAT2 | 20,66710203 | 20,80239859 | 26,79137108 | 15,68219706 | 2 |
| ENSGALG00000016455 | GREB1 | 71,30258407 | 44,76556849 | 171,8842869 | 40,7988828 | 2 |
| ENSGALG00000016480 | SDC1 | 67,79712381 | 46,12962808 | 101,6542603 | 65,96909209 | 2 |
| ENSGALG00000016808 | RANBP2 | 88,68408088 | 76,82428287 | 126,4454672 | 83,37223599 | 2 |
| ENSGALG00000016875 | | 35,81511404 | 23,39580851 | 81,48141489 | 21,24280635 | 2 |
| ENSGALG00000016896 | ABCC4 | 9,804323199 | 7,015685475 | 14,63344086 | 7,987146561 | 2 |
| ENSGALG00000016897 | GPR180 | 25,35505379 | 21,30681925 | 31,82543062 | 22,00243895 | 2 |
| ENSGALG00000016910 | RBM26 | 141,5608071 | 114,5852063 | 174,753134 | 120,2734436 | 2 |
| ENSGALG00000017085 | SLC7A1 | 40,05394599 | 35,00050584 | 54,14362264 | 38,81530443 | 2 |
| ENSGALG00000017111 | NUP58 | 63,43269188 | 54,90711803 | 79,68018034 | 60,76739727 | 2 |
| ENSGALG00000017365 | | 22,49317964 | 17,18921742 | 33,78900494 | 20,82699363 | 2 |
| ENSGALG00000017388 | | 186,1814132 | 172,046714 | 268,9548589 | 178,5764003 | 2 |
| ENSGALG00000017462 | FNIP1 | 23,16041033 | 20,29852756 | 34,80619268 | 20,97040204 | 2 |
| ENSGALG00000019077 | | 75,80953752 | 63,82999065 | 101,13045 | 49,73846401 | 2 |
| ENSGALG00000019622 | TRIM71 | 63,53466997 | 45,08282347 | 138,6372475 | 59,01429912 | 2 |
| ENSGALG00000019961 | SLC18B1 | 94,7023836 | 82,41511543 | 144,636966 | 86,89618913 | 2 |
| ENSGALG00000020479 | | 23,15151259 | 19,80281907 | 35,36804246 | 21,72259355 | 2 |
| ENSGALG00000020805 | LRRC8D | 52,23662484 | 47,43227859 | 70,9545852 | 43,87669915 | 2 |
| ENSGALG00000021316 | | 95,97879893 | 77,14187045 | 118,2758942 | 74,31764513 | 2 |
| ENSGALG00000023451 | TRPM7 | 25,75064391 | 21,92560329 | 34,73493332 | 22,8366183 | 2 |
| ENSGALG00000023576 | PIM3 | 38,39240852 | 36,6823612 | 50,24199049 | 36,47779083 | 2 |
| ENSGALG00000024039 | | 104,7252252 | 86,46498974 | 126,7964154 | 101,8972811 | 2 |
| ENSGALG00000025770 | | 5,489257369 | 3,00959702 | 12,02421921 | 5,106431368 | 2 |
| ENSGALG00000025884 | NABP1 | 53,416184 | 46,44733288 | 68,89014188 | 48,27093752 | 2 |
| ENSGALG00000026349 | | 276,236352 | 197,8659478 | 354,8447655 | 217,6076426 | 2 |
| ENSGALG00000027316 | | 2,126519018 | 1,150096401 | 4,599057408 | 0,899918777 | 2 |
| ENSGALG00000027342 | | 54,55996923 | 43,14565226 | 88,63782029 | 46,00922333 | 2 |
| ENSGALG00000027355 | TPBG | 77,04597047 | 65,11404966 | 96,88531936 | 76,00938707 | 2 |
| ENSGALG00000027389 | RGS7BP | 16,07367378 | 12,09558999 | 26,33652515 | 15,7130332 | 2 |

| | | | | | | |
|---|---|---|---|---|---|---|
| ENSGALG00000027757 | ARL4A | 92,36112237 | 89,07729168 | 139,8864616 | 67,39567749 | 2 |
| ENSGALG00000028462 | | 49,20550825 | 40,59815974 | 63,79574475 | 48,48730827 | 2 |
| ENSGALG00000028821 | GLIPR2 | 73,84703902 | 51,69185499 | 185,78976 | 66,36412003 | 2 |
| ENSGALG00000029023 | DMRTB1 | 0,460922202 | 0,311878344 | 1,17310912 | 0,418308344 | 2 |
| ENSGALG00000000054 | | 40,61041039 | 77,77927106 | 101,7475053 | 79,46683091 | 3 |
| ENSGALG00000000162 | | 0,462119365 | 0,938839881 | 1,64917459 | 0,88469459 | 3 |
| ENSGALG00000000385 | | 31,22124771 | 34,99404406 | 41,45690236 | 38,58983564 | 3 |
| ENSGALG00000000409 | CHMP7 | 79,5962766 | 87,42800269 | 94,16850493 | 90,97109387 | 3 |
| ENSGALG00000000546 | | 13,1839576 | 15,33846461 | 17,00256672 | 16,22873609 | 3 |
| ENSGALG00000000721 | SLC41A1 | 10,56410509 | 14,67315988 | 25,86101352 | 19,0181628 | 3 |
| ENSGALG00000001007 | | 249,8579466 | 266,9325144 | 356,1755387 | 297,3682242 | 3 |
| ENSGALG00000001482 | | 39,02660104 | 42,05429417 | 46,83086881 | 45,01081844 | 3 |
| ENSGALG00000001789 | | 5,660481882 | 5,998946978 | 8,128074203 | 6,887258106 | 3 |
| ENSGALG00000001962 | PTBP1 | 232,0447099 | 243,7695722 | 332,3908518 | 271,6208432 | 3 |
| ENSGALG00000002232 | | 38,27382102 | 41,1639609 | 46,66833375 | 42,91091967 | 3 |
| ENSGALG00000002248 | | 31,30319367 | 32,96247972 | 49,67058945 | 40,65598095 | 3 |
| ENSGALG00000002294 | SEMA6A | 10,49550142 | 13,58348745 | 24,74662011 | 18,10914393 | 3 |
| ENSGALG00000002326 | | 1,47250781 | 2,63934996 | 4,3086282 | 2,825363581 | 3 |
| ENSGALG00000002335 | DOCK3 | 1,86448435 | 2,175929741 | 3,65664367 | 2,786015113 | 3 |
| ENSGALG00000002336 | | 31,11744712 | 33,42002163 | 37,20470924 | 36,29167307 | 3 |
| ENSGALG00000002521 | | 18,85761021 | 20,11966762 | 24,33563548 | 21,95444037 | 3 |
| ENSGALG00000002604 | | 56,3890493 | 58,74501369 | 73,02098131 | 67,097638 | 3 |
| ENSGALG00000002745 | | 150,7994546 | 155,5390263 | 180,6431172 | 172,3193816 | 3 |
| ENSGALG00000003113 | RGS14 | 32,70926878 | 38,37080551 | 45,95420135 | 39,64318938 | 3 |
| ENSGALG00000003488 | | 1,062824324 | 2,489829504 | 4,41900121 | 3,4655683 | 3 |
| ENSGALG00000003634 | PTEN | 109,6705257 | 112,8873555 | 130,5075766 | 116,9977413 | 3 |
| ENSGALG00000003701 | SGMS1 | 9,258518873 | 11,10229042 | 15,94820162 | 13,26938247 | 3 |
| ENSGALG00000003948 | ALAS1 | 49,59705625 | 55,47941093 | 71,59274829 | 59,09107457 | 3 |
| ENSGALG00000004088 | FNBP1 | 24,99237284 | 26,63166615 | 41,69591151 | 33,84795116 | 3 |
| ENSGALG00000004158 | | 10,55351282 | 11,6407991 | 18,6163799 | 13,95560906 | 3 |
| ENSGALG00000004181 | | 10,65424926 | 12,03452833 | 16,95695499 | 13,00094636 | 3 |
| ENSGALG00000004384 | | 69,61318883 | 72,98146284 | 91,46967536 | 84,65324587 | 3 |
| ENSGALG00000004509 | | 973,1657851 | 1201,316866 | 1361,543225 | 1311,496279 | 3 |
| ENSGALG00000004517 | PARP3 | 6,608038224 | 7,4623483 | 10,339109 | 8,753186004 | 3 |
| ENSGALG00000004573 | LRRC8A | 0,06745578 | 0,071659756 | 0,107430357 | 0,088247902 | 3 |
| ENSGALG00000004594 | CD74 | 4,406632912 | 8,750656652 | 24,51781075 | 14,099464 | 3 |
| ENSGALG00000004661 | | 55,81884913 | 71,36370724 | 89,93526626 | 84,78952034 | 3 |
| ENSGALG00000004730 | | 2,547946628 | 4,415127922 | 9,113776156 | 6,91193394 | 3 |
| ENSGALG00000004917 | | 2,896199012 | 3,859495387 | 6,644848881 | 4,974527218 | 3 |
| ENSGALG00000004974 | PPARG | 1,096913427 | 1,387820745 | 3,376628365 | 2,480137643 | 3 |
| ENSGALG00000005310 | | 4,865705062 | 6,154572974 | 9,226338957 | 6,706461454 | 3 |
| ENSGALG00000005492 | | 10,69479144 | 14,25428111 | 24,12257422 | 16,1669692 | 3 |
| ENSGALG00000005678 | FLNB | 60,41912172 | 63,75180733 | 77,98540804 | 71,20030379 | 3 |
| ENSGALG00000005692 | | 3,071738847 | 4,652874224 | 7,830445668 | 5,084013812 | 3 |
| ENSGALG00000005990 | | 0,702479687 | 0,971890089 | 2,092257816 | 1,181810364 | 3 |
| ENSGALG00000006097 | PLXNA1 | 90,9441087 | 93,67817119 | 112,3883703 | 102,2276242 | 3 |
| ENSGALG00000006252 | LMO4 | 39,86339327 | 44,78129011 | 86,70760634 | 62,88011361 | 3 |
| ENSGALG00000006258 | SPOCK1 | 35,25706314 | 52,38093264 | 93,08564779 | 55,08217348 | 3 |
| ENSGALG00000006314 | | 2,490008645 | 4,28813785 | 7,168503958 | 6,122938793 | 3 |
| ENSGALG00000006417 | SEC24A | 58,74346813 | 64,91074036 | 75,01570455 | 66,74698384 | 3 |
| ENSGALG00000006638 | GRIK4 | 10,90795756 | 15,65522755 | 23,06474952 | 17,01050771 | 3 |
| ENSGALG00000006717 | | 0,341376117 | 3,858277819 | 12,25143393 | 5,62632686 | 3 |
| ENSGALG00000006786 | | 9,374384568 | 13,921691 | 19,00994587 | 16,68642534 | 3 |
| ENSGALG00000006845 | | 2,670363578 | 3,667942238 | 4,464215057 | 3,687958638 | 3 |
| ENSGALG00000006928 | | 28,56711841 | 31,59803573 | 37,31258837 | 32,05630269 | 3 |
| ENSGALG00000007140 | NRP1 | 6,786804779 | 9,937661813 | 29,18904165 | 20,3705717 | 3 |
| ENSGALG00000007608 | ARHGEF9 | 19,95586878 | 33,81382649 | 49,630279 | 46,62145769 | 3 |
| ENSGALG00000007707 | TRAPPC4 | 89,44703604 | 93,8442147 | 110,2275432 | 97,87185693 | 3 |
| ENSGALG00000007712 | MYO1B | 15,61469896 | 20,14680824 | 26,22598793 | 22,24150912 | 3 |
| ENSGALG00000007920 | PRR5L | 4,129042771 | 4,972105698 | 7,776312534 | 6,401104872 | 3 |
| ENSGALG00000008018 | SNAI1 | 4,838614652 | 6,837750638 | 10,0616924 | 8,716434475 | 3 |
| ENSGALG00000008128 | | 75,28877718 | 82,09362626 | 103,3431878 | 93,65315249 | 3 |
| ENSGALG00000008206 | TESC | 3,628035071 | 6,771511577 | 8,739719175 | 7,526555407 | 3 |
| ENSGALG00000008321 | RNPEPL1 | 22,85721363 | 26,24567137 | 33,01671824 | 30,40006594 | 3 |

| | | | | | | |
|---|---|---|---|---|---|---|
| ENSGALG00000008359 | CD93 | 1,29957602 | 1,606526404 | 2,626230481 | 1,994368504 | 3 |
| ENSGALG00000008445 | SLC24A3 | 7,111549609 | 8,166183777 | 13,26231133 | 11,30603962 | 3 |
| ENSGALG00000008623 | TOR4A | 2,885819993 | 3,8654735 | 7,453674406 | 5,093186098 | 3 |
| ENSGALG00000009145 | | 1,464711516 | 1,626807682 | 2,784818386 | 2,107621671 | 3 |
| ENSGALG00000009293 | CHN1 | 67,90118322 | 74,31595236 | 82,89622495 | 76,74118835 | 3 |
| ENSGALG00000009393 | | 6,562399308 | 15,29420916 | 43,36617594 | 24,96798456 | 3 |
| ENSGALG00000009808 | | 1,416304393 | 2,421872238 | 5,557291922 | 2,900189483 | 3 |
| ENSGALG00000010005 | EPAS1 | 1,665911612 | 3,648797813 | 8,298542118 | 6,247343231 | 3 |
| ENSGALG00000010035 | NR3C2 | 6,203251658 | 9,632638993 | 13,68730791 | 9,212635965 | 3 |
| ENSGALG00000010081 | SH3D19 | 9,151663444 | 12,27542336 | 21,57827913 | 14,7478447 | 3 |
| ENSGALG00000010090 | FAM160A1 | 2,972923275 | 4,146500246 | 6,55742234 | 5,035528514 | 3 |
| ENSGALG00000010186 | | 26,1218641 | 34,11416012 | 41,89387746 | 40,73222649 | 3 |
| ENSGALG00000010518 | | 0,144840281 | 0,417602932 | 0,617619694 | 0,379300378 | 3 |
| ENSGALG00000010597 | | 4,611954486 | 9,57844156 | 19,361201 | 10,00647113 | 3 |
| ENSGALG00000010780 | | 14,90096854 | 16,74023252 | 20,06717072 | 17,00357201 | 3 |
| ENSGALG00000010910 | | 5,56043356 | 11,09159137 | 18,79138106 | 10,94754966 | 3 |
| ENSGALG00000010996 | IRF2BP2 | 74,78507418 | 89,5496145 | 142,7037833 | 109,2878448 | 3 |
| ENSGALG00000011061 | | 0 | 0,089561338 | 0,194908996 | 0,087418642 | 3 |
| ENSGALG00000011076 | SYNE3 | 0,555939022 | 0,853639426 | 2,736597554 | 1,544551691 | 3 |
| ENSGALG00000011220 | SLC25A29 | 13,82485106 | 20,4490233 | 37,69585477 | 30,16838999 | 3 |
| ENSGALG00000011232 | | 8,944661229 | 10,28962659 | 16,59653933 | 12,22287995 | 3 |
| ENSGALG00000011241 | RFTN1 | 4,68880177 | 5,893184053 | 7,360556398 | 6,707841258 | 3 |
| ENSGALG00000011550 | | 40,28733517 | 55,05082442 | 67,35622718 | 54,65736176 | 3 |
| ENSGALG00000011613 | | 4,739203641 | 5,310547233 | 9,592133476 | 7,686518403 | 3 |
| ENSGALG00000011831 | ANKRD50 | 13,06309751 | 25,09232183 | 32,26143246 | 25,9987912 | 3 |
| ENSGALG00000011963 | PRDM5 | 2,076763979 | 3,308086443 | 4,532414832 | 3,751882733 | 3 |
| ENSGALG00000012033 | SGSM3 | 12,06891416 | 14,05699702 | 17,1981993 | 14,96637778 | 3 |
| ENSGALG00000012111 | TMEM37 | 3,030173772 | 3,743044977 | 7,331132284 | 5,289018921 | 3 |
| ENSGALG00000012407 | LRP1B | 0,189428611 | 0,330664455 | 0,574724991 | 0,367586865 | 3 |
| ENSGALG00000012449 | MBD5 | 36,89274386 | 42,54116552 | 53,76802751 | 44,82551839 | 3 |
| ENSGALG00000012462 | KIF5C | 65,21507153 | 73,27025091 | 90,38041221 | 78,71948215 | 3 |
| ENSGALG00000012739 | HIVEP1 | 15,82020811 | 17,77656476 | 21,4826725 | 19,34743987 | 3 |
| ENSGALG00000012850 | RAB19 | 3,884293705 | 4,594800469 | 7,321110934 | 5,597645823 | 3 |
| ENSGALG00000013060 | ADCY2 | 1,064982925 | 1,936215345 | 2,96245522 | 2,691019878 | 3 |
| ENSGALG00000013085 | PTPRO | 8,814882072 | 9,635541753 | 16,31598922 | 11,48544595 | 3 |
| ENSGALG00000013204 | | 11,67890458 | 12,83559276 | 15,73940433 | 14,21451059 | 3 |
| ENSGALG00000013605 | LRRC16A | 24,93445177 | 27,46094865 | 32,7533139 | 29,62259302 | 3 |
| ENSGALG00000014139 | | 48,56470481 | 50,53915714 | 62,44067282 | 56,14372316 | 3 |
| ENSGALG00000014371 | LGALS8 | 42,99482411 | 51,47866425 | 59,99952802 | 56,0859926 | 3 |
| ENSGALG00000014441 | | 10,1718582 | 12,15414806 | 16,15178081 | 12,91195223 | 3 |
| ENSGALG00000014545 | | 5,504651212 | 7,224808336 | 8,631274642 | 6,873714829 | 3 |
| ENSGALG00000014834 | NCOA7 | 17,62360916 | 20,65742055 | 26,25090959 | 22,94517424 | 3 |
| ENSGALG00000014841 | HDDC2 | 155,8070969 | 174,4976494 | 189,514832 | 184,7270869 | 3 |
| ENSGALG00000015196 | ASXL3 | 12,54053734 | 13,3023841 | 16,82687164 | 15,16050633 | 3 |
| ENSGALG00000015355 | CD47 | 76,26899784 | 118,0330955 | 173,0471949 | 137,0561941 | 3 |
| ENSGALG00000015653 | | 3,796339821 | 6,553253791 | 10,50484987 | 6,581133531 | 3 |
| ENSGALG00000015667 | | 15,19349969 | 17,21199916 | 19,65556076 | 17,55600721 | 3 |
| ENSGALG00000015721 | SVEP1 | 1,737960293 | 2,245209146 | 3,02298257 | 2,299009295 | 3 |
| ENSGALG00000015858 | DOPEY1 | 10,26362443 | 12,06032829 | 15,43817177 | 12,56861101 | 3 |
| ENSGALG00000015865 | | 8,149500097 | 14,67315572 | 23,51658618 | 18,36456973 | 3 |
| ENSGALG00000015995 | PTPRA | 111,1055577 | 127,974971 | 137,2289686 | 127,4894494 | 3 |
| ENSGALG00000016285 | | 18,08199823 | 20,16991605 | 23,48414703 | 21,6556274 | 3 |
| ENSGALG00000016329 | AGPAT5 | 54,88045679 | 57,69984224 | 65,85606335 | 59,46993298 | 3 |
| ENSGALG00000016350 | ARHGEF10 | 13,20943184 | 16,47206961 | 19,97855358 | 16,74087613 | 3 |
| ENSGALG00000016356 | TDRP | 19,61514474 | 24,85234346 | 31,91658421 | 26,97607924 | 3 |
| ENSGALG00000016443 | HPCAL1 | 45,49523459 | 60,94224423 | 91,38350236 | 82,37034179 | 3 |
| ENSGALG00000016719 | TNFRSF21 | 32,04606952 | 36,62371331 | 51,90970368 | 46,80574718 | 3 |
| ENSGALG00000016746 | | 6,325273962 | 6,601627683 | 8,839401918 | 7,699088309 | 3 |
| ENSGALG00000016779 | RNF149 | 40,32057066 | 43,89003588 | 53,7136991 | 48,40680769 | 3 |
| ENSGALG00000016909 | NDFIP2 | 79,60051734 | 96,09155119 | 149,2317403 | 108,9565955 | 3 |
| ENSGALG00000016927 | | 4,423680336 | 5,818898304 | 11,01889315 | 6,732269887 | 3 |
| ENSGALG00000017112 | MTMR6 | 27,31233071 | 28,93025574 | 35,58260609 | 30,94460943 | 3 |
| ENSGALG00000017283 | CCND2 | 58,83812867 | 71,53308904 | 78,60163688 | 74,96269338 | 3 |
| ENSGALG00000017335 | | 49,94809765 | 58,41748149 | 65,31328378 | 61,44158464 | 3 |

| | | | | | | |
|---|---|---|---|---|---|---|
| ENSGALG00000017715 | TINAGL1 | 0,390432351 | 0,886129706 | 1,981143569 | 1,219554501 | 3 |
| ENSGALG00000021812 | | 1,33055689 | 2,218290992 | 4,819468653 | 3,797137631 | 3 |
| ENSGALG00000022887 | KBTBD11 | 9,128381161 | 10,94614726 | 24,8936814 | 16,96298322 | 3 |
| ENSGALG00000022909 | PTPN3 | 1,705551313 | 2,69191956 | 5,75680131 | 4,605522677 | 3 |
| ENSGALG00000023027 | | 5,819536568 | 8,734844706 | 12,14989903 | 8,702812739 | 3 |
| ENSGALG00000023193 | | 22,82628116 | 26,83374341 | 46,03186329 | 34,00628372 | 3 |
| ENSGALG00000023857 | | 23,87214226 | 28,64172799 | 31,048651 | 28,83857214 | 3 |
| ENSGALG00000026093 | | 5,630092855 | 7,023865753 | 9,045481093 | 6,813892067 | 3 |
| ENSGALG00000026484 | | 0,135024738 | 0,263566954 | 0,838178415 | 0,546375363 | 3 |
| ENSGALG00000026728 | AMMECR1L | 29,84210004 | 31,09949395 | 35,82038624 | 32,81834419 | 3 |
| ENSGALG00000026776 | TIGAR | 13,56828479 | 16,77103564 | 21,89957869 | 19,88878977 | 3 |
| ENSGALG00000026795 | | 0,217228639 | 0,495115948 | 2,256733718 | 1,191236832 | 3 |
| ENSGALG00000027070 | TIMP2 | 63,034473 | 97,3863476 | 154,0452617 | 134,7888119 | 3 |
| ENSGALG00000027241 | | 0,952560408 | 1,466719985 | 3,039748701 | 2,207159329 | 3 |
| ENSGALG00000027255 | NRXN3 | 0,181357946 | 0,830745305 | 2,361320482 | 1,14505164 | 3 |
| ENSGALG00000027521 | | 0,404163847 | 0,994883411 | 2,02843061 | 1,126800761 | 3 |
| ENSGALG00000027835 | PCBD1 | 94,80491215 | 113,5855851 | 151,1573142 | 114,9552317 | 3 |
| ENSGALG00000027983 | TNK2 | 4,828489899 | 7,027495256 | 8,351294461 | 7,398414931 | 3 |
| ENSGALG00000028242 | | 6,695143255 | 8,395783618 | 9,169385188 | 8,751478338 | 3 |
| ENSGALG00000028935 | BCL10 | 31,45990136 | 34,35834737 | 53,25704847 | 43,70890102 | 3 |
| ENSGALG00000000177 | | 3,672925592 | 4,7133494 | 28,69442229 | 13,64228432 | 4 |
| ENSGALG00000000581 | DDX42 | 79,71475033 | 75,27436341 | 94,20135848 | 80,66368967 | 4 |
| ENSGALG00000000638 | UTP4 | 130,0676455 | 96,70332836 | 165,3136722 | 132,8315835 | 4 |
| ENSGALG00000000652 | ACOT7 | 91,0618764 | 84,44688017 | 110,5059683 | 102,0297592 | 4 |
| ENSGALG00000000875 | PHLPP2 | 20,54132997 | 19,47427289 | 25,59885106 | 23,08633198 | 4 |
| ENSGALG00000000890 | | 115,4827014 | 99,93575405 | 161,284949 | 135,4581557 | 4 |
| ENSGALG00000000891 | NSUN5 | 53,01503719 | 44,67338132 | 77,96581988 | 55,35290206 | 4 |
| ENSGALG00000000949 | HBEGF | 1,41847004 | 1,85041107 | 12,65482264 | 5,806391814 | 4 |
| ENSGALG00000001020 | | 29,88405432 | 26,56154792 | 64,98663159 | 38,60596779 | 4 |
| ENSGALG00000001025 | GOLGA1 | 22,45573384 | 18,51150657 | 29,42118809 | 24,79417751 | 4 |
| ENSGALG00000001194 | | 212,5442816 | 205,1105713 | 287,0990995 | 265,262679 | 4 |
| ENSGALG00000001364 | CSNK1A1 | 415,1207983 | 409,9539609 | 501,5029834 | 437,3747681 | 4 |
| ENSGALG00000001382 | CDC2L1 | 72,94889565 | 67,80678106 | 88,43393376 | 81,89331757 | 4 |
| ENSGALG00000001546 | ZZEF1 | 12,49878851 | 12,56771979 | 17,15985806 | 15,54466993 | 4 |
| ENSGALG00000001551 | CCNL2 | 122,413604 | 108,8267176 | 162,3532986 | 139,7824135 | 4 |
| ENSGALG00000001876 | | 9,90311991 | 9,249057674 | 14,02958487 | 10,97007453 | 4 |
| ENSGALG00000002198 | FHOD1 | 5,26743915 | 4,954885979 | 9,38561863 | 6,836917746 | 4 |
| ENSGALG00000002222 | NOC2L | 118,3584902 | 95,02050883 | 142,0720819 | 116,3738506 | 4 |
| ENSGALG00000002362 | MANF | 364,1212991 | 353,3282859 | 462,1312606 | 409,2625306 | 4 |
| ENSGALG00000002364 | | 32,94239485 | 32,71450827 | 39,97668091 | 35,39664528 | 4 |
| ENSGALG00000002399 | | 52,36416197 | 46,22509547 | 70,78493069 | 56,5784979 | 4 |
| ENSGALG00000002471 | | 16,17379426 | 15,90253266 | 23,9774057 | 19,02253072 | 4 |
| ENSGALG00000002709 | MYO1C | 16,40141681 | 14,00693265 | 25,42132948 | 18,29325483 | 4 |
| ENSGALG00000002759 | PEAK1 | 12,11070695 | 12,37294828 | 18,07755992 | 15,14928933 | 4 |
| ENSGALG00000003039 | SRM | 110,2374056 | 95,18663787 | 138,4535202 | 109,8554395 | 4 |
| ENSGALG00000003043 | | 22,75792127 | 21,97305286 | 39,26576174 | 30,68440091 | 4 |
| ENSGALG00000003082 | DPH1 | 43,35222965 | 39,90415548 | 56,22303159 | 46,64836959 | 4 |
| ENSGALG00000003130 | DDX49 | 65,08104231 | 64,76205282 | 78,85615956 | 71,53496308 | 4 |
| ENSGALG00000003181 | SURF2 | 68,66649596 | 63,22622727 | 93,24040328 | 77,05363503 | 4 |
| ENSGALG00000003401 | FBXL22 | 1,842586497 | 2,069041413 | 5,714050546 | 4,460181323 | 4 |
| ENSGALG00000003563 | YARS | 124,5878605 | 100,806762 | 161,5804147 | 129,3590756 | 4 |
| ENSGALG00000003726 | EGFLAM | 3,127640917 | 2,882156629 | 6,50447413 | 3,899600608 | 4 |
| ENSGALG00000003830 | | 56,83320796 | 53,87981451 | 71,17442725 | 65,73104839 | 4 |
| ENSGALG00000004042 | | 38,46767509 | 32,97068481 | 61,87801155 | 49,83556181 | 4 |
| ENSGALG00000004063 | SLC25A14 | 52,88533686 | 51,42823816 | 66,67629574 | 61,35330486 | 4 |
| ENSGALG00000004089 | | 17,77256296 | 16,78271592 | 24,42500908 | 19,86038229 | 4 |
| ENSGALG00000004098 | STK4 | 78,3035915 | 77,0974588 | 96,93702951 | 88,80516834 | 4 |
| ENSGALG00000004200 | DPP9 | 53,75067211 | 45,69989026 | 71,29460307 | 57,55657601 | 4 |
| ENSGALG00000004295 | CPD | 62,31416544 | 60,36967779 | 77,78486944 | 67,0127332 | 4 |
| ENSGALG00000004437 | LHFPL2 | 20,63775236 | 19,68831498 | 34,32158717 | 26,52799986 | 4 |
| ENSGALG00000004625 | ATXN2 | 88,67526317 | 82,01836417 | 109,5403049 | 102,5728716 | 4 |
| ENSGALG00000004662 | DISP3 | 14,38521556 | 11,55502683 | 29,05261461 | 16,89063595 | 4 |
| ENSGALG00000004691 | | 56,33686982 | 40,4318144 | 71,73587757 | 59,00776865 | 4 |
| ENSGALG00000004820 | FAM57A | 12,31601366 | 10,07982253 | 19,55856559 | 14,55945516 | 4 |

| | | | | | | |
|---|---|---|---|---|---|---|
| ENSGALG00000004938 | | 54,1479251 | 51,08009956 | 67,80947377 | 62,11469064 | 4 |
| ENSGALG00000005139 | CLTC | 200,1181658 | 191,6793455 | 253,6981165 | 209,8034908 | 4 |
| ENSGALG00000005318 | | 3,152263541 | 2,662414727 | 5,120954069 | 3,784663378 | 4 |
| ENSGALG00000005433 | | 36,83290793 | 32,95618637 | 55,24825615 | 43,10703235 | 4 |
| ENSGALG00000005464 | | 85,83962449 | 85,8443126 | 105,9021193 | 93,37178801 | 4 |
| ENSGALG00000005582 | KLHL18 | 31,94479309 | 31,30348133 | 42,78533543 | 37,78377686 | 4 |
| ENSGALG00000005653 | NFKB2 | 19,40013661 | 19,08669938 | 32,80863272 | 23,57889165 | 4 |
| ENSGALG00000005660 | NF1 | 25,50711294 | 23,71387031 | 34,53169071 | 28,24626635 | 4 |
| ENSGALG00000005668 | WSB1 | 179,678984 | 146,1632157 | 214,4435918 | 176,4064239 | 4 |
| ENSGALG00000005679 | CDX1 | 1,304631914 | 0,905701649 | 2,563014719 | 1,644766779 | 4 |
| ENSGALG00000005691 | | 0,879450332 | 1,016907284 | 4,287680194 | 2,372582554 | 4 |
| ENSGALG00000005727 | | 87,5175031 | 54,87972446 | 124,3665649 | 87,29261047 | 4 |
| ENSGALG00000005962 | | 96,05596249 | 97,74823491 | 129,0244635 | 115,5419182 | 4 |
| ENSGALG00000006225 | EFL1 | 25,89461358 | 24,34066277 | 33,72870887 | 27,27818148 | 4 |
| ENSGALG00000006430 | | 34,42578206 | 34,74030494 | 42,66924729 | 38,00467817 | 4 |
| ENSGALG00000006512 | HSPA8 | 1476,479912 | 1243,966272 | 1904,025233 | 1566,091754 | 4 |
| ENSGALG00000006564 | | 135,1349308 | 137,694922 | 164,941611 | 147,7888853 | 4 |
| ENSGALG00000006569 | | 30,28902795 | 30,34830504 | 44,14456419 | 34,89953162 | 4 |
| ENSGALG00000006703 | GTPBP4 | 263,023663 | 226,6648498 | 306,5407089 | 275,5959813 | 4 |
| ENSGALG00000006908 | DLG1 | 79,94395383 | 69,9741556 | 167,5563942 | 97,80401695 | 4 |
| ENSGALG00000007056 | | 14,79571572 | 13,46626993 | 33,62262451 | 21,76104972 | 4 |
| ENSGALG00000007174 | TNFSF15 | 1,102995748 | 0,448114415 | 2,915856114 | 1,855636786 | 4 |
| ENSGALG00000007241 | NUBP1 | 109,3181308 | 94,68377161 | 138,0537018 | 121,3146554 | 4 |
| ENSGALG00000007278 | GRIN2A | 1,410215689 | 1,099864815 | 2,433976143 | 1,483307418 | 4 |
| ENSGALG00000007312 | | 0,515818805 | 0,324808533 | 0,640115412 | 0,516898894 | 4 |
| ENSGALG00000007387 | GTF3C1 | 49,4431244 | 44,10989653 | 61,73807855 | 50,46690808 | 4 |
| ENSGALG00000007434 | RHBDF1 | 8,971085939 | 8,883515652 | 14,18157043 | 11,77887017 | 4 |
| ENSGALG00000007516 | | 62,43238732 | 57,39706836 | 74,52591155 | 62,64255662 | 4 |
| ENSGALG00000007603 | | 107,5989261 | 79,43933012 | 130,7533918 | 107,5767645 | 4 |
| ENSGALG00000007659 | HYOU1 | 247,2830843 | 202,3775908 | 337,8470085 | 253,6943761 | 4 |
| ENSGALG00000007704 | | 344,7122616 | 259,8186212 | 567,3248203 | 425,2898747 | 4 |
| ENSGALG00000007793 | UPRT | 63,74779044 | 56,0337098 | 86,10974443 | 72,54448635 | 4 |
| ENSGALG00000007845 | REL | 11,42236566 | 10,4053628 | 17,28277767 | 14,80106617 | 4 |
| ENSGALG00000007846 | | 0,386598336 | 0,938692576 | 50,96425002 | 26,68289287 | 4 |
| ENSGALG00000007932 | TRAF6 | 10,42206821 | 10,02548702 | 20,10355496 | 15,77060665 | 4 |
| ENSGALG00000007957 | | 27,06104057 | 24,98161486 | 33,66292504 | 31,22532025 | 4 |
| ENSGALG00000008097 | CELF1 | 26,96388959 | 26,53468446 | 34,24988615 | 30,0909987 | 4 |
| ENSGALG00000008163 | PSME4 | 46,80933972 | 45,11907125 | 79,1287345 | 59,67413509 | 4 |
| ENSGALG00000008182 | CHAC2 | 6,548181465 | 7,336110999 | 17,67917062 | 12,66985204 | 4 |
| ENSGALG00000008211 | ACP2 | 64,70328045 | 58,45883411 | 80,09941154 | 66,54844454 | 4 |
| ENSGALG00000008263 | CNTN4 | 1,106200523 | 1,176160283 | 3,185854517 | 2,100686659 | 4 |
| ENSGALG00000008348 | PDIA3 | 746,0442099 | 742,0928002 | 909,6299808 | 852,0733662 | 4 |
| ENSGALG00000008414 | DVL3 | 135,6013387 | 130,367946 | 176,8576759 | 158,0362434 | 4 |
| ENSGALG00000008520 | EIF4G1 | 229,5376092 | 185,6983546 | 301,8025317 | 265,7430302 | 4 |
| ENSGALG00000008521 | | 31,18069235 | 29,21280914 | 43,31917635 | 33,76297669 | 4 |
| ENSGALG00000008680 | | 3,676371429 | 3,695854361 | 6,469859596 | 5,35658988 | 4 |
| ENSGALG00000008708 | | 37,14584777 | 32,77792521 | 49,18084278 | 37,57520453 | 4 |
| ENSGALG00000008778 | SPRED2 | 34,65788535 | 37,41244323 | 69,179714 | 49,64306392 | 4 |
| ENSGALG00000008942 | FOXN2 | 31,91191136 | 31,77313307 | 38,7914746 | 33,70013914 | 4 |
| ENSGALG00000009129 | DLX5 | 10,01865983 | 5,040758016 | 22,8485431 | 12,34243947 | 4 |
| ENSGALG00000009172 | OSBPL6 | 2,650564571 | 2,756249804 | 5,544541276 | 3,837527585 | 4 |
| ENSGALG00000009534 | | 25,76733875 | 24,62278561 | 35,59283804 | 27,46733311 | 4 |
| ENSGALG00000009546 | | 10,46067246 | 7,836081786 | 30,46916994 | 17,16886901 | 4 |
| ENSGALG00000009594 | RARRES1 | 9,3163689 | 9,894726709 | 19,208027 | 15,83960019 | 4 |
| ENSGALG00000009699 | CCNL1 | 91,8351549 | 82,06759115 | 115,5386418 | 102,8629539 | 4 |
| ENSGALG00000009906 | UBR2 | 22,46888792 | 22,12414082 | 27,23693376 | 24,46789564 | 4 |
| ENSGALG00000009946 | HECTD1 | 48,79520303 | 46,18145293 | 63,02540497 | 54,54830376 | 4 |
| ENSGALG00000010036 | FOSL2 | 1,42569897 | 0,793171681 | 12,63940938 | 7,470718426 | 4 |
| ENSGALG00000010049 | PTPRB | 0,961664682 | 1,128851705 | 3,443635365 | 2,118202631 | 4 |
| ENSGALG00000010084 | EIF4G3 | 114,1687657 | 103,5506191 | 143,5390186 | 117,7492928 | 4 |
| ENSGALG00000010098 | SLC6A9 | 21,12603755 | 15,71173611 | 30,01896704 | 26,093241 | 4 |
| ENSGALG00000010167 | PNN | 265,0067262 | 220,0057882 | 322,9605906 | 271,8747079 | 4 |
| ENSGALG00000010349 | POLR1C | 117,943623 | 93,52322591 | 145,73653 | 117,5515932 | 4 |
| ENSGALG00000010361 | | 48,97012579 | 44,43975772 | 68,27866824 | 55,34965362 | 4 |

| | | | | | | |
|---|---|---|---|---|---|---|
| ENSGALG00000010786 | | 4,097877003 | 2,931870248 | 5,490876839 | 4,103642012 | 4 |
| ENSGALG00000010836 | | 3,785106733 | 3,728133151 | 7,708117179 | 4,823935284 | 4 |
| ENSGALG00000010967 | | 58,3636706 | 56,63549114 | 79,71286329 | 65,06813368 | 4 |
| ENSGALG00000011055 | | 3,528425682 | 2,14343308 | 3,539716441 | 3,81794966 | 4 |
| ENSGALG00000011058 | LEPR | 0,561037727 | 0,528085813 | 2,462715898 | 1,549528016 | 4 |
| ENSGALG00000011331 | CTH | 248,0035867 | 236,6542483 | 303,6507875 | 275,389819 | 4 |
| ENSGALG00000011419 | DUSP4 | 44,01115652 | 33,26358857 | 121,1807682 | 85,68812834 | 4 |
| ENSGALG00000011537 | PDE10A | 17,52900263 | 13,79619641 | 27,86596367 | 20,34592748 | 4 |
| ENSGALG00000011833 | | 29,27658949 | 25,74330458 | 117,5407031 | 60,95401121 | 4 |
| ENSGALG00000011962 | | 24,37861772 | 19,78836623 | 76,1380067 | 54,90216003 | 4 |
| ENSGALG00000012137 | KTN1 | 52,45719129 | 52,70647636 | 63,46102016 | 56,68222967 | 4 |
| ENSGALG00000012202 | | 52,23663855 | 47,13644927 | 66,99015096 | 54,26052413 | 4 |
| ENSGALG00000012342 | SASH1 | 23,67583218 | 24,00657812 | 31,8459396 | 26,74112268 | 4 |
| ENSGALG00000012354 | | 38,36252665 | 31,9693923 | 72,19373709 | 50,92066877 | 4 |
| ENSGALG00000012404 | ERO1A | 22,83857753 | 22,01192185 | 30,88099896 | 28,27942365 | 4 |
| ENSGALG00000012410 | | 67,06196987 | 58,3818171 | 81,80427215 | 74,10521197 | 4 |
| ENSGALG00000012414 | GNPNAT1 | 66,78273296 | 56,41713595 | 92,31540052 | 73,43548792 | 4 |
| ENSGALG00000012626 | | 14,77954066 | 13,73791725 | 18,79074056 | 15,62077944 | 4 |
| ENSGALG00000012696 | | 60,55715685 | 57,86504 | 99,67821995 | 75,06643816 | 4 |
| ENSGALG00000012702 | JARID2 | 37,33195628 | 36,20655994 | 105,618813 | 76,98869311 | 4 |
| ENSGALG00000012847 | | 0,986713969 | 0,845118709 | 4,302990379 | 2,59612048 | 4 |
| ENSGALG00000012852 | MKRN1 | 230,2783715 | 221,0113476 | 347,0213055 | 264,4287667 | 4 |
| ENSGALG00000013454 | | 182,0155039 | 123,4948983 | 229,2326813 | 190,0588303 | 4 |
| ENSGALG00000013616 | OPRM1 | 8,498418048 | 8,602322585 | 13,37881711 | 10,98826365 | 4 |
| ENSGALG00000013677 | | 30,75467791 | 29,88810558 | 39,45604034 | 37,02318823 | 4 |
| ENSGALG00000014080 | | 0,21595968 | 0,263148688 | 1,065482871 | 0,727785613 | 4 |
| ENSGALG00000014445 | | 231,7075249 | 162,1297126 | 273,7779191 | 224,85464 | 4 |
| ENSGALG00000014795 | TGIF1 | 90,42983611 | 80,36474464 | 117,3826664 | 93,7149106 | 4 |
| ENSGALG00000014880 | | 7,049861195 | 6,527543364 | 11,03857212 | 8,462532694 | 4 |
| ENSGALG00000014922 | ROCK1 | 58,13294478 | 55,12527106 | 91,44699065 | 66,17491261 | 4 |
| ENSGALG00000015180 | PSAT1 | 39,87210776 | 38,25469818 | 74,32407226 | 52,26746661 | 4 |
| ENSGALG00000015184 | | 18,40942927 | 17,96441873 | 31,32066199 | 23,65139942 | 4 |
| ENSGALG00000015271 | FILIP1L | 1,507114952 | 0,978526739 | 2,740725006 | 1,723861266 | 4 |
| ENSGALG00000015284 | RGMB | 11,87565903 | 10,63508532 | 17,84314469 | 15,52968056 | 4 |
| ENSGALG00000015310 | | 26,26727406 | 23,40911182 | 43,92421285 | 29,60330699 | 4 |
| ENSGALG00000015820 | | 5,795989287 | 4,835828284 | 10,12157143 | 6,826990388 | 4 |
| ENSGALG00000015908 | COL12A1 | 1,280525315 | 0,982791397 | 3,642005312 | 1,798732148 | 4 |
| ENSGALG00000015930 | | 89,77271382 | 89,18055945 | 105,8631291 | 96,52936382 | 4 |
| ENSGALG00000015942 | | 25,13739734 | 24,29186435 | 34,37777216 | 29,63188224 | 4 |
| ENSGALG00000015980 | | 0,596196735 | 0,41874926 | 1,674898262 | 0,878230845 | 4 |
| ENSGALG00000016308 | | 26,44250919 | 19,32816829 | 50,33957199 | 31,74534839 | 4 |
| ENSGALG00000016387 | TRAPPC12 | 41,96858601 | 38,40737819 | 51,57440402 | 44,44505587 | 4 |
| ENSGALG00000016506 | ZNF512 | 88,06015215 | 79,02693958 | 104,7277824 | 96,40793662 | 4 |
| ENSGALG00000016513 | PPM1G | 230,2916367 | 196,6821079 | 271,5136768 | 260,5494092 | 4 |
| ENSGALG00000016764 | EIF5B | 216,2000449 | 194,9595544 | 249,9786739 | 241,9844265 | 4 |
| ENSGALG00000016803 | ST6GAL2 | 35,94066123 | 32,07201411 | 58,08741372 | 43,47710939 | 4 |
| ENSGALG00000016807 | | 34,55977522 | 25,87689197 | 63,00840571 | 44,52494993 | 4 |
| ENSGALG00000016816 | CDC16 | 62,16393798 | 54,16495998 | 74,3131724 | 66,19654442 | 4 |
| ENSGALG00000016817 | RASA3 | 18,72143367 | 21,1947238 | 46,6794009 | 30,14801484 | 4 |
| ENSGALG00000016841 | | 23,36476602 | 15,1644545 | 49,52648097 | 33,88640549 | 4 |
| ENSGALG00000016843 | COL4A2 | 23,50684554 | 16,00417492 | 47,20989436 | 25,07421643 | 4 |
| ENSGALG00000017185 | TMEM123 | 82,3272294 | 82,74278569 | 123,1509651 | 97,26527034 | 4 |
| ENSGALG00000017320 | | 172,8178042 | 158,3012361 | 208,2067185 | 188,3508122 | 4 |
| ENSGALG00000017418 | NPTX1 | 2,015625875 | 1,372808327 | 4,33776454 | 3,060994409 | 4 |
| ENSGALG00000018879 | | 6,563062255 | 5,094820232 | 11,25672365 | 8,341936654 | 4 |
| ENSGALG00000019054 | | 0,728717128 | 0,463400019 | 6,735112481 | 2,212402684 | 4 |
| ENSGALG00000019470 | | 1,444283154 | 1,413792107 | 3,796706053 | 2,057755864 | 4 |
| ENSGALG00000020899 | | 6,083043676 | 4,511874861 | 13,31624301 | 8,922200915 | 4 |
| ENSGALG00000022490 | | 3,542294964 | 4,40309386 | 17,43885586 | 11,12767909 | 4 |
| ENSGALG00000023241 | | 14,69724223 | 12,08407893 | 57,94569675 | 29,73935332 | 4 |
| ENSGALG00000023725 | | 74,22983509 | 68,94311731 | 125,647953 | 97,24504122 | 4 |
| ENSGALG00000026034 | RGS17 | 3,024836883 | 2,866775508 | 5,550569778 | 4,08214868 | 4 |
| ENSGALG00000026251 | | 0,057739324 | 0 | 0,072028586 | 0,0498639 | 4 |
| ENSGALG00000026379 | | 0,919332817 | 0,655580292 | 2,303728052 | 1,597097731 | 4 |

| | | | | | | |
|---|---|---|---|---|---|---|
| ENSGALG00000026468 | CHSY1 | 71,12020109 | 60,49110129 | 111,0111548 | 79,65498057 | 4 |
| ENSGALG00000026631 | HOXA10 | 0,174683257 | 0,06671589 | 0,623564049 | 0,246427429 | 4 |
| ENSGALG00000026862 | CLDN1 | 14,1101416 | 17,88491877 | 498,851265 | 292,4470901 | 4 |
| ENSGALG00000027057 | | 45,65853393 | 41,50345279 | 55,55225266 | 47,46219036 | 4 |
| ENSGALG00000027132 | | 21,35779753 | 20,57984953 | 27,44116286 | 24,14699039 | 4 |
| ENSGALG00000027526 | COL9A2 | 18,60130136 | 18,58810977 | 27,51237863 | 22,80853569 | 4 |
| ENSGALG00000027696 | | 17,22300656 | 13,40010504 | 27,54529579 | 21,46414333 | 4 |
| ENSGALG00000027707 | ESM1 | 0,150067881 | 0,073206173 | 1,347708981 | 0,510169001 | 4 |
| ENSGALG00000028279 | SNORD14 | 59,56148275 | 36,73539954 | 85,15192697 | 70,25858401 | 4 |
| ENSGALG00000028675 | | 43,07663481 | 36,65018625 | 58,259897 | 43,676594 | 4 |
| ENSGALG00000028773 | | 88,44681977 | 88,70929071 | 105,8823606 | 98,70306764 | 4 |
| ENSGALG00000000166 | | 4,929206982 | 8,868613541 | 10,16203898 | 7,489518013 | 5 |
| ENSGALG00000000874 | | 27,5257408 | 42,0642166 | 38,71313331 | 35,44203257 | 5 |
| ENSGALG00000001443 | | 61,90702775 | 100,220611 | 84,17995064 | 70,86083476 | 5 |
| ENSGALG00000001741 | NPTN | 112,9186561 | 136,9981308 | 138,6382684 | 124,8191766 | 5 |
| ENSGALG00000001926 | HSPB1 | 26,87936471 | 51,36562943 | 52,8540106 | 47,19220813 | 5 |
| ENSGALG00000002602 | | 3,588473784 | 4,875391746 | 5,77341269 | 4,458144988 | 5 |
| ENSGALG00000003126 | ERBB4 | 1,696937945 | 7,085918708 | 11,05966427 | 3,505512894 | 5 |
| ENSGALG00000003263 | | 76,37963484 | 99,04824669 | 105,8462646 | 85,42780449 | 5 |
| ENSGALG00000003578 | FN1 | 62,38375818 | 70,31597704 | 75,66792654 | 66,94626169 | 5 |
| ENSGALG00000003642 | PDGF-A | 28,92734021 | 56,03765372 | 46,12301374 | 39,27720096 | 5 |
| ENSGALG00000003748 | ANO5 | 25,09400369 | 31,77740288 | 32,82517973 | 30,40964521 | 5 |
| ENSGALG00000004071 | NETO2 | 43,78776373 | 55,53083264 | 54,37099849 | 37,75449027 | 5 |
| ENSGALG00000004118 | | 7,47667819 | 26,22733823 | 20,20577066 | 14,30878756 | 5 |
| ENSGALG00000004184 | SPARC | 155,0540441 | 232,7699504 | 211,3314171 | 199,4811275 | 5 |
| ENSGALG00000004215 | WIPI1 | 33,13808616 | 43,77015696 | 42,74430902 | 40,4803601 | 5 |
| ENSGALG00000004519 | TRPV2 | 0,822447882 | 1,60414895 | 2,273876542 | 1,268485872 | 5 |
| ENSGALG00000004686 | | 30,2279262 | 36,35087632 | 37,69253986 | 35,20409282 | 5 |
| ENSGALG00000004710 | TMOD3 | 46,23733435 | 62,98353115 | 66,8079787 | 60,43811176 | 5 |
| ENSGALG00000004724 | | 14,31585256 | 20,5405451 | 19,66998379 | 18,70531832 | 5 |
| ENSGALG00000005223 | PAC1 | 12,65150175 | 17,99234789 | 18,81912824 | 15,2360441 | 5 |
| ENSGALG00000005230 | MED13 | 32,91669497 | 36,87653151 | 39,04153735 | 36,48895089 | 5 |
| ENSGALG00000005256 | EPHA4 | 42,66846892 | 78,56615799 | 96,36961707 | 62,46596208 | 5 |
| ENSGALG00000005512 | | 47,88593846 | 69,10907971 | 81,90943527 | 52,75638588 | 5 |
| ENSGALG00000006472 | | 1,419681564 | 4,536593254 | 3,288727942 | 2,539433944 | 5 |
| ENSGALG00000006705 | MYOF | 2,618645578 | 7,53302833 | 6,251568851 | 4,810717487 | 5 |
| ENSGALG00000007343 | | 0,49771636 | 3,066447888 | 2,335443105 | 1,682163718 | 5 |
| ENSGALG00000007766 | KIAA1217 | 5,142338869 | 9,945849569 | 7,737757011 | 6,6003621 | 5 |
| ENSGALG00000007809 | TSHZ2 | 5,283631458 | 12,9087792 | 13,412855 | 11,27929818 | 5 |
| ENSGALG00000008053 | PRDM11 | 21,30498282 | 24,55027354 | 26,50984007 | 23,21540024 | 5 |
| ENSGALG00000008222 | SLC26A5 | 29,61861728 | 40,31425403 | 46,54677335 | 38,47187747 | 5 |
| ENSGALG00000008281 | NEURL1 | 0,643796003 | 2,406738252 | 2,878119044 | 1,315326778 | 5 |
| ENSGALG00000008670 | | 60,9188267 | 92,15789368 | 112,4269423 | 81,96474528 | 5 |
| ENSGALG00000008903 | SSFA2 | 9,137506036 | 12,52608672 | 14,56587518 | 11,95743887 | 5 |
| ENSGALG00000009163 | | 8,759302894 | 14,89860879 | 16,52575933 | 11,8340751 | 5 |
| ENSGALG00000009378 | PDGFC | 12,77846886 | 24,19271297 | 24,15556174 | 22,07089313 | 5 |
| ENSGALG00000009556 | PRICKLE1 | 10,09396138 | 19,24833012 | 25,35963008 | 17,81681241 | 5 |
| ENSGALG00000009645 | ESRRG | 13,80191386 | 17,39972079 | 19,42718421 | 14,40652117 | 5 |
| ENSGALG00000010311 | NAV3 | 0,615932694 | 1,312817613 | 1,821822919 | 0,975422452 | 5 |
| ENSGALG00000010470 | TCERG1L | 2,421080738 | 7,105845544 | 7,441496756 | 5,889451234 | 5 |
| ENSGALG00000010588 | | 0,618826163 | 7,910358704 | 6,674385318 | 4,860719678 | 5 |
| ENSGALG00000011047 | LEPROT | 49,88870182 | 65,03197595 | 64,41076564 | 61,76206291 | 5 |
| ENSGALG00000011254 | SATB1 | 11,10630754 | 13,32672382 | 15,03624216 | 12,84351007 | 5 |
| ENSGALG00000011288 | DACT2 | 43,3233096 | 77,67540397 | 56,71321695 | 50,47908136 | 5 |
| ENSGALG00000011373 | TECPR2 | 14,79142682 | 18,36494399 | 18,18702754 | 17,49834935 | 5 |
| ENSGALG00000011539 | | 57,67498946 | 85,42119822 | 84,42143077 | 72,20633676 | 5 |
| ENSGALG00000011542 | | 58,95969276 | 74,88826879 | 78,93145405 | 60,78404348 | 5 |
| ENSGALG00000011680 | EPHA5 | 8,20621494 | 24,36580423 | 24,51946644 | 14,89187836 | 5 |
| ENSGALG00000012251 | | 20,12025104 | 26,36888417 | 27,50762152 | 21,56503412 | 5 |
| ENSGALG00000013031 | CECR1 | 5,057170782 | 7,128107118 | 8,312127528 | 5,595783665 | 5 |
| ENSGALG00000013134 | | 22,82793318 | 29,22780923 | 31,32115935 | 27,27461137 | 5 |
| ENSGALG00000013503 | FAM149A | 1,752925892 | 4,848988383 | 4,853083076 | 3,295521827 | 5 |
| ENSGALG00000013710 | SNX9 | 28,51304311 | 33,65160133 | 35,14415843 | 30,51214921 | 5 |
| ENSGALG00000013768 | PHACTR2 | 16,97204936 | 22,53311454 | 26,1533975 | 21,45971135 | 5 |

| | | | | | | |
|---|---|---|---|---|---|---|
| ENSGALG00000015013 | MSX1 | 272,3083235 | 372,1131576 | 353,787571 | 293,7007336 | 5 |
| ENSGALG00000015096 | SH3GL2 | 6,752855595 | 9,972778277 | 10,62531102 | 9,697118369 | 5 |
| ENSGALG00000015167 | PRUNE2 | 0,376231707 | 0,978364886 | 0,754818696 | 0,73083871 | 5 |
| ENSGALG00000015193 | CCDC80 | 14,67870793 | 25,22532121 | 21,73900557 | 21,1053184 | 5 |
| ENSGALG00000015215 | MAPRE2 | 117,8139702 | 164,251409 | 175,9162941 | 136,884432 | 5 |
| ENSGALG00000015409 | PROS1 | 45,426329 | 66,08672514 | 83,84526057 | 55,91392118 | 5 |
| ENSGALG00000015794 | | 5,438251718 | 7,227879991 | 7,741266482 | 6,719009922 | 5 |
| ENSGALG00000015797 | | 0,53814002 | 7,856916105 | 2,807378812 | 2,732711798 | 5 |
| ENSGALG00000016160 | | 8,263899107 | 10,02238742 | 10,67328648 | 9,407543829 | 5 |
| ENSGALG00000016258 | | 0,878895793 | 3,431767393 | 3,190781752 | 1,828922771 | 5 |
| ENSGALG00000016276 | KHDRBS2 | 17,88353402 | 32,13830872 | 27,94112843 | 21,08977544 | 5 |
| ENSGALG00000016281 | DMD | 17,81837254 | 29,46320013 | 29,11375678 | 20,07434969 | 5 |
| ENSGALG00000016495 | KLHL29 | 7,027776784 | 9,00783271 | 9,910837711 | 8,941313628 | 5 |
| ENSGALG00000016776 | TBC1D8 | 8,308297301 | 10,63215235 | 11,46728711 | 10,62476118 | 5 |
| ENSGALG00000016839 | | 23,55110961 | 30,45895529 | 34,20721695 | 28,99322678 | 5 |
| ENSGALG00000016947 | ELF1 | 16,00351494 | 19,80439985 | 20,80439973 | 18,68259934 | 5 |
| ENSGALG00000016981 | | 0,057692799 | 0,773832888 | 1,091983863 | 0,323032301 | 5 |
| ENSGALG00000017246 | ME3 | 13,03979429 | 22,01093333 | 24,71107053 | 22,21530556 | 5 |
| ENSGALG00000017411 | RXFP3 | 0,51168238 | 1,422949404 | 2,090496742 | 0,518778426 | 5 |
| ENSGALG00000024035 | | 5,499637289 | 10,63733364 | 13,88164669 | 9,819516571 | 5 |
| ENSGALG00000025747 | NQO2 | 8,133537598 | 12,93516452 | 11,59383961 | 10,23613308 | 5 |
| ENSGALG00000026258 | MAF | 0,778820903 | 9,324146442 | 13,1495057 | 6,561829291 | 5 |
| ENSGALG00000026370 | | 0,891124883 | 2,751588481 | 2,510061088 | 1,175691721 | 5 |
| ENSGALG00000026486 | | 4,237996386 | 7,869569204 | 6,981784134 | 3,839481651 | 5 |
| ENSGALG00000026991 | | 1,07803858 | 1,847127388 | 1,949984514 | 1,262349842 | 5 |
| ENSGALG00000027627 | | 30,89543317 | 50,46301542 | 50,55295596 | 43,60585579 | 5 |
| ENSGALG00000027947 | GHSR | 0,104688776 | 0,849436519 | 0,862659404 | 0,749153227 | 5 |
| ENSGALG00000028602 | | 13,19864069 | 32,69987166 | 40,67321367 | 22,05665612 | 5 |
| ENSGALG00000000151 | ADAMTS19 | 11,80611965 | 12,46514745 | 7,204855475 | 9,17871014 | 6 |
| ENSGALG00000000227 | DPYSL2 | 270,6999858 | 268,6183531 | 219,938641 | 254,0376962 | 6 |
| ENSGALG00000000280 | EFNA5 | 24,32055926 | 20,52988368 | 11,64998519 | 17,31698677 | 6 |
| ENSGALG00000000362 | | 37,35064361 | 32,02718335 | 27,44376501 | 33,1655256 | 6 |
| ENSGALG00000000449 | | 49,2665107 | 49,10237718 | 33,20160451 | 41,94914256 | 6 |
| ENSGALG00000000529 | LIMD2 | 93,67951598 | 94,89616479 | 75,32979772 | 80,66727455 | 6 |
| ENSGALG00000000547 | | 3,018242875 | 2,863860974 | 1,595558221 | 2,347638467 | 6 |
| ENSGALG00000000584 | | 3,261584505 | 2,895004253 | 2,038105409 | 2,118177535 | 6 |
| ENSGALG00000000607 | GPR37L1 | 5,416118593 | 4,00865424 | 2,523565581 | 3,30135529 | 6 |
| ENSGALG00000000678 | CITED4 | 15,62473381 | 11,43958359 | 7,239375596 | 9,048783186 | 6 |
| ENSGALG00000000692 | CDK18 | 22,23095371 | 21,04019437 | 14,37658239 | 15,37207022 | 6 |
| ENSGALG00000000812 | OAZ1 | 489,9056014 | 482,0065382 | 407,200401 | 467,4568226 | 6 |
| ENSGALG00000000839 | WNT11 | 2,75489007 | 2,199440661 | 1,264090823 | 1,917561529 | 6 |
| ENSGALG00000000912 | | 46,3292383 | 41,75530213 | 31,67896574 | 36,03517849 | 6 |
| ENSGALG00000001264 | PLXNA2 | 58,22770217 | 53,91327501 | 24,13638965 | 32,04174249 | 6 |
| ENSGALG00000001374 | | 104,8241518 | 112,9489393 | 90,04261095 | 91,50863097 | 6 |
| ENSGALG00000001472 | JAM3 | 100,011492 | 93,74068448 | 82,19173947 | 92,91975166 | 6 |
| ENSGALG00000001506 | | 368,3526839 | 372,9183268 | 296,0295963 | 339,4166627 | 6 |
| ENSGALG00000001567 | | 126,5813492 | 129,435459 | 100,5746864 | 111,6667323 | 6 |
| ENSGALG00000001611 | B3GAT1 | 7,43006463 | 8,500157291 | 3,962658545 | 5,321053943 | 6 |
| ENSGALG00000001916 | SLIT3 | 25,76301015 | 25,02868434 | 16,99253818 | 21,07931886 | 6 |
| ENSGALG00000002005 | | 5,413669282 | 5,77801867 | 2,751506276 | 4,550706014 | 6 |
| ENSGALG00000002024 | COMT | 27,25339768 | 29,45942369 | 19,71910445 | 20,44636948 | 6 |
| ENSGALG00000002028 | GPRC5B | 14,34185212 | 14,07342152 | 5,786874678 | 9,906480205 | 6 |
| ENSGALG00000002040 | | 79,93552599 | 81,85898438 | 64,79086608 | 61,98429748 | 6 |
| ENSGALG00000002041 | | 59,40242733 | 57,27740971 | 48,38350615 | 52,07872637 | 6 |
| ENSGALG00000002056 | TANGO2 | 45,88582661 | 46,90045634 | 35,03961486 | 37,17107403 | 6 |
| ENSGALG00000002116 | TEN1 | 20,56756935 | 19,76690486 | 13,13442349 | 13,44002686 | 6 |
| ENSGALG00000002155 | | 76,89660127 | 81,52844767 | 49,12868241 | 64,57841702 | 6 |
| ENSGALG00000002377 | ENO1 | 5212,537489 | 5074,505023 | 4322,535611 | 4651,610527 | 6 |
| ENSGALG00000002552 | | 10,14344218 | 8,496682359 | 6,743722308 | 7,370594553 | 6 |
| ENSGALG00000002561 | | 27,30204849 | 27,24602203 | 20,69160555 | 25,05439519 | 6 |
| ENSGALG00000002645 | | 15,71672487 | 13,70701973 | 8,631292786 | 8,68302629 | 6 |
| ENSGALG00000002897 | SLC16A14 | 27,20069002 | 23,63489477 | 17,04976801 | 16,06268463 | 6 |
| ENSGALG00000002909 | | 111,6429653 | 108,8364841 | 65,45516928 | 74,21179258 | 6 |
| ENSGALG00000002911 | MOXD1 | 4,443996829 | 3,766709566 | 2,890068858 | 2,950915419 | 6 |

| | | | | | | |
|---|---|---|---|---|---|---|
| ENSGALG00000002952 | NRG4 | 28,16643542 | 25,91401013 | 16,97766028 | 18,81496187 | 6 |
| ENSGALG00000003193 | CRABP-I | 372,078802 | 401,8286749 | 212,1153567 | 250,9542932 | 6 |
| ENSGALG00000003212 | TSPO2 | 9,375586046 | 7,497418907 | 5,455285302 | 5,632090299 | 6 |
| ENSGALG00000003322 | TBC1D2B | 23,0364406 | 23,04385432 | 18,28372397 | 14,42807756 | 6 |
| ENSGALG00000003374 | ESRP2 | 4,290851004 | 4,01526902 | 2,786837195 | 2,82526351 | 6 |
| ENSGALG00000003473 | SFRP1 | 198,6174832 | 161,8765433 | 117,5653707 | 121,237755 | 6 |
| ENSGALG00000003539 | KIAA1522 | 48,54035458 | 41,79698536 | 34,13776991 | 42,25080563 | 6 |
| ENSGALG00000003550 | ARHGAP40 | 3,317838701 | 2,653438966 | 1,952460676 | 2,348843771 | 6 |
| ENSGALG00000003602 | | 17,85228161 | 18,32582058 | 13,19044577 | 14,56512893 | 6 |
| ENSGALG00000003732 | UCK1 | 74,42929071 | 77,91477397 | 61,41002009 | 65,22123954 | 6 |
| ENSGALG00000003742 | CPAMD8 | 22,90988764 | 20,34187827 | 16,93949378 | 18,38217809 | 6 |
| ENSGALG00000003829 | HPCAL4 | 9,672599215 | 8,323189404 | 5,220820337 | 7,313009587 | 6 |
| ENSGALG00000003894 | | 39,49235103 | 48,65174612 | 23,31361969 | 26,19853057 | 6 |
| ENSGALG00000003965 | DBX1 | 120,9552444 | 117,7217027 | 51,14908628 | 70,42580158 | 6 |
| ENSGALG00000003973 | TLCD1 | 112,8068814 | 117,1801993 | 73,74619124 | 94,39823275 | 6 |
| ENSGALG00000004048 | PTPRS | 155,7119771 | 161,4097176 | 128,6427922 | 143,3686471 | 6 |
| ENSGALG00000004249 | GRHL3 | 6,349924787 | 5,24388785 | 3,935762529 | 4,71839621 | 6 |
| ENSGALG00000004270 | ALDH1A2 | 27,69749738 | 24,65884251 | 18,46023442 | 21,00172637 | 6 |
| ENSGALG00000004370 | MAP2K6 | 29,31711165 | 33,73015033 | 16,64289398 | 21,44172167 | 6 |
| ENSGALG00000004527 | | 2,134308872 | 2,240087571 | 1,027730715 | 1,26194997 | 6 |
| ENSGALG00000004598 | CUX2 | 10,91100525 | 10,38753268 | 7,769537303 | 10,06444635 | 6 |
| ENSGALG00000004849 | WSCD2 | 45,34791706 | 50,70050864 | 22,81269623 | 34,91261466 | 6 |
| ENSGALG00000004888 | RAI1 | 40,93525626 | 33,1953892 | 27,30738815 | 31,72051679 | 6 |
| ENSGALG00000005036 | SH2D3C | 19,05266339 | 20,32939106 | 12,13835847 | 15,76846777 | 6 |
| ENSGALG00000005064 | PRPSAP2 | 180,1816199 | 171,8333543 | 152,3830656 | 151,8777115 | 6 |
| ENSGALG00000005107 | | 70,58508054 | 63,43166224 | 49,22045308 | 51,47493334 | 6 |
| ENSGALG00000005396 | | 24,53292592 | 23,39277847 | 16,09342546 | 13,16614788 | 6 |
| ENSGALG00000005409 | | 9,602911865 | 11,56291556 | 4,665267822 | 5,939605674 | 6 |
| ENSGALG00000005608 | SLC9A3R2 | 41,76346794 | 36,24125491 | 25,01105653 | 35,56871411 | 6 |
| ENSGALG00000005616 | ARSI | 10,31329519 | 8,403075571 | 5,912173057 | 7,138945964 | 6 |
| ENSGALG00000005632 | MRVI1 | 1,035430484 | 0,888938675 | 0,506588986 | 0,613816863 | 6 |
| ENSGALG00000005886 | WSCD1 | 42,42581136 | 36,95926631 | 26,36940368 | 25,78087186 | 6 |
| ENSGALG00000005976 | TCEA2 | 151,1204022 | 158,0299347 | 129,1939558 | 140,5406725 | 6 |
| ENSGALG00000005981 | GDF2 | 35,21583503 | 34,95950736 | 15,63020753 | 18,34501736 | 6 |
| ENSGALG00000005985 | GDF10 | 151,8179242 | 133,3393762 | 68,22657308 | 67,90626041 | 6 |
| ENSGALG00000006211 | BCL2L1 | 77,71688202 | 69,24101045 | 46,01690957 | 57,52750738 | 6 |
| ENSGALG00000006277 | | 43,37142919 | 41,65643624 | 34,95474443 | 37,42762996 | 6 |
| ENSGALG00000006341 | | 2,477772914 | 1,863998893 | 1,22355805 | 1,629215459 | 6 |
| ENSGALG00000006360 | | 116,2259025 | 108,0171956 | 99,60835707 | 100,7079967 | 6 |
| ENSGALG00000006420 | ABHD17C | 43,96192153 | 44,25815693 | 36,5118052 | 39,97659664 | 6 |
| ENSGALG00000006456 | C16orf45 | 126,2885268 | 123,5755496 | 103,0104563 | 113,0436349 | 6 |
| ENSGALG00000006539 | C21orf2 | 91,92359624 | 96,93066537 | 72,63220666 | 80,75071846 | 6 |
| ENSGALG00000006576 | GRM3 | 5,093834571 | 4,402339701 | 1,89484859 | 3,657846954 | 6 |
| ENSGALG00000006583 | LSP1 | 130,1339628 | 114,8168042 | 87,05056318 | 96,05195296 | 6 |
| ENSGALG00000006662 | | 4,073009343 | 3,076860613 | 1,949960342 | 2,615131782 | 6 |
| ENSGALG00000006715 | OAF | 67,7069681 | 52,68685915 | 39,6578323 | 46,72294434 | 6 |
| ENSGALG00000006886 | DACH2 | 97,17562698 | 96,3669717 | 74,86391205 | 74,85847215 | 6 |
| ENSGALG00000006887 | FAM174B | 113,7792578 | 123,4632073 | 89,0301308 | 92,30488335 | 6 |
| ENSGALG00000006950 | LIMK2 | 39,24605928 | 44,80038999 | 28,89988718 | 31,66912322 | 6 |
| ENSGALG00000006982 | | 28,22770649 | 26,3955686 | 20,28855166 | 21,73180767 | 6 |
| ENSGALG00000007007 | | 3,672988192 | 3,675424255 | 2,403461423 | 2,636506344 | 6 |
| ENSGALG00000007127 | | 72,79567328 | 70,87318067 | 59,36657504 | 64,29580864 | 6 |
| ENSGALG00000007131 | | 1741,940063 | 1718,967686 | 1405,400638 | 1577,061404 | 6 |
| ENSGALG00000007132 | ACOX2 | 3,920015175 | 3,800327799 | 2,227110374 | 3,369924911 | 6 |
| ENSGALG00000007364 | TOB1 | 25,92250443 | 23,41006218 | 15,10205731 | 18,83966369 | 6 |
| ENSGALG00000007441 | | 37,78990923 | 39,61953583 | 19,20223432 | 28,70034349 | 6 |
| ENSGALG00000007719 | | 8,587739597 | 8,673743587 | 4,950497049 | 4,790200635 | 6 |
| ENSGALG00000007819 | PDZRN3 | 7,790604113 | 6,689038204 | 4,884979375 | 6,326245839 | 6 |
| ENSGALG00000007839 | NCAM1 | 224,900862 | 248,3471659 | 164,3211414 | 191,3588384 | 6 |
| ENSGALG00000007912 | ATP9A | 14,30496017 | 14,40885784 | 11,23211708 | 12,79682857 | 6 |
| ENSGALG00000007945 | CRYAB | 28,66809483 | 20,95669537 | 10,44303707 | 14,14521444 | 6 |
| ENSGALG00000007993 | DCX | 35,56507544 | 35,44942575 | 22,95234521 | 19,99682824 | 6 |
| ENSGALG00000008034 | CASKIN2 | 15,6733548 | 15,28382385 | 10,1352996 | 13,19245044 | 6 |
| ENSGALG00000008060 | PRKAR2B | 56,53549837 | 55,35469483 | 39,37569127 | 48,11630919 | 6 |

| | | | | | | |
|---|---|---|---|---|---|---|
| ENSGALG00000008136 | C2orf69 | 23,07666382 | 22,96082979 | 16,74740759 | 19,66350146 | 6 |
| ENSGALG00000008152 | | 44,0565263 | 40,40259712 | 32,97020753 | 34,00914523 | 6 |
| ENSGALG00000008164 | | 16,82913916 | 16,88116082 | 13,13788045 | 11,98403826 | 6 |
| ENSGALG00000008297 | SEMA4B | 63,40218087 | 60,59213273 | 34,32024933 | 46,40412182 | 6 |
| ENSGALG00000008555 | MXI1 | 62,54223051 | 63,65172867 | 39,52038549 | 50,53829158 | 6 |
| ENSGALG00000008579 | OIP5 | 80,39985195 | 71,04664556 | 62,04889257 | 65,37711843 | 6 |
| ENSGALG00000008677 | VIM | 1934,220794 | 2091,395034 | 1528,513774 | 1633,778736 | 6 |
| ENSGALG00000008774 | | 240,3536889 | 235,8958052 | 155,5629281 | 192,5922241 | 6 |
| ENSGALG00000008850 | TTLL7 | 9,855630847 | 9,638036441 | 6,465694185 | 7,831903916 | 6 |
| ENSGALG00000009004 | ZNF804B | 0,528433071 | 0,560499823 | 0,242320416 | 0,44430822 | 6 |
| ENSGALG00000009048 | ABLIM1 | 10,71327222 | 8,768191528 | 6,002604343 | 7,651491713 | 6 |
| ENSGALG00000009064 | FZD1 | 94,20126626 | 80,37022502 | 67,72298945 | 75,05676907 | 6 |
| ENSGALG00000009248 | HSPA12A | 32,76381377 | 29,41540416 | 19,35339196 | 24,0607483 | 6 |
| ENSGALG00000009280 | NPY2R | 7,52643434 | 6,490731964 | 4,241120616 | 4,842042787 | 6 |
| ENSGALG00000009291 | EPHX1 | 24,12810147 | 21,68268279 | 16,94897373 | 14,68925954 | 6 |
| ENSGALG00000009362 | | 91,11750451 | 103,1144581 | 72,59595314 | 68,03634497 | 6 |
| ENSGALG00000009475 | | 366,4300954 | 326,2865393 | 265,7944967 | 289,0206255 | 6 |
| ENSGALG00000009559 | | 62,46382877 | 63,35737381 | 50,60514824 | 51,23070493 | 6 |
| ENSGALG00000009604 | DBX2 | 188,02469 | 164,9576559 | 71,47102484 | 112,4678314 | 6 |
| ENSGALG00000009650 | | 40,35619744 | 41,46793477 | 31,51222177 | 31,26515047 | 6 |
| ENSGALG00000009799 | | 131,9263511 | 123,4148587 | 88,50089928 | 102,2744474 | 6 |
| ENSGALG00000009905 | MDM1 | 34,94608598 | 38,49309929 | 27,93640968 | 26,68582306 | 6 |
| ENSGALG00000009948 | HHIP | 34,10489404 | 29,81672032 | 12,8659564 | 17,40556709 | 6 |
| ENSGALG00000010055 | DAAM2 | 7,571786163 | 6,688168824 | 4,066430756 | 4,819714522 | 6 |
| ENSGALG00000010158 | KANK1 | 34,42266386 | 33,10703465 | 25,60147101 | 25,08690202 | 6 |
| ENSGALG00000010243 | PRDX1 | 1920,279994 | 1901,867506 | 1624,165646 | 1809,468067 | 6 |
| ENSGALG00000010711 | NXPH1 | 1,336267919 | 1,08347215 | 0,412641959 | 0,793884376 | 6 |
| ENSGALG00000010745 | HMGB2 | 960,0569077 | 950,3575761 | 759,8576275 | 846,3521199 | 6 |
| ENSGALG00000010758 | | 80,09122628 | 85,39793585 | 63,98441519 | 61,48005141 | 6 |
| ENSGALG00000010761 | MRPL37 | 104,7510396 | 105,0379991 | 87,00391806 | 96,37058418 | 6 |
| ENSGALG00000010854 | HDAC9 | 15,52765661 | 13,90606664 | 8,67588231 | 11,10862945 | 6 |
| ENSGALG00000010892 | | 94,54684796 | 97,06101349 | 74,78496364 | 77,87489575 | 6 |
| ENSGALG00000010940 | ACSS3 | 17,0859282 | 19,73733137 | 11,03537862 | 13,6172326 | 6 |
| ENSGALG00000011019 | ROR1 | 31,89158503 | 33,92332362 | 19,77815514 | 25,04993277 | 6 |
| ENSGALG00000011083 | | 31,53400696 | 29,83533586 | 21,70010114 | 21,42515155 | 6 |
| ENSGALG00000011117 | AGT | 7,011848699 | 6,40595233 | 3,225035171 | 4,179528638 | 6 |
| ENSGALG00000011205 | SMOC2 | 28,04565303 | 26,98470908 | 9,630259431 | 18,70884659 | 6 |
| ENSGALG00000011283 | ZNF385D | 2,605818014 | 2,669322669 | 1,67305957 | 2,163474606 | 6 |
| ENSGALG00000011296 | FRMD1 | 70,2232087 | 67,45990844 | 55,38004143 | 52,65139895 | 6 |
| ENSGALG00000011298 | RARB | 55,69535963 | 47,68028638 | 38,04727303 | 37,68492663 | 6 |
| ENSGALG00000011442 | TGFBR2 | 2,861068196 | 2,68853117 | 1,838107115 | 2,275891693 | 6 |
| ENSGALG00000011487 | | 37,86246542 | 33,98107732 | 21,65815303 | 19,53005182 | 6 |
| ENSGALG00000011518 | POMGNT2 | 15,30539131 | 14,01183671 | 10,32877903 | 9,911142688 | 6 |
| ENSGALG00000011532 | | 55,69108076 | 52,76270126 | 42,60190613 | 42,40119097 | 6 |
| ENSGALG00000011558 | | 23,50941161 | 20,49599556 | 17,43700181 | 18,64310313 | 6 |
| ENSGALG00000011580 | | 116,9362646 | 111,1956453 | 101,032554 | 102,3871869 | 6 |
| ENSGALG00000011592 | MRAS | 26,78163289 | 27,15769346 | 16,17293361 | 23,32241226 | 6 |
| ENSGALG00000011657 | EAF2 | 88,86209982 | 78,74142964 | 59,13328659 | 70,88662684 | 6 |
| ENSGALG00000011712 | | 27,43874052 | 26,29278103 | 16,13911667 | 16,52237231 | 6 |
| ENSGALG00000011742 | ART4 | 2,842510701 | 2,786801503 | 1,453508753 | 1,821990845 | 6 |
| ENSGALG00000011826 | | 24,11633262 | 24,94919055 | 16,13711683 | 18,0713928 | 6 |
| ENSGALG00000011848 | | 61,83068956 | 50,7503725 | 40,99509459 | 46,82618647 | 6 |
| ENSGALG00000012006 | RTN1 | 15,5752017 | 15,69038812 | 8,141532846 | 10,07584334 | 6 |
| ENSGALG00000012010 | | 12,13552249 | 10,81139678 | 8,109147776 | 7,96587428 | 6 |
| ENSGALG00000012082 | FAIM | 44,71906582 | 40,82570202 | 33,31693495 | 29,35247649 | 6 |
| ENSGALG00000012094 | SLC35F4 | 3,504266079 | 2,479954589 | 1,868276354 | 2,82870976 | 6 |
| ENSGALG00000012158 | | 144,978586 | 127,8854779 | 106,8748489 | 109,7215444 | 6 |
| ENSGALG00000012219 | UNC5C | 21,57499882 | 22,95837716 | 14,58740088 | 16,32433091 | 6 |
| ENSGALG00000012282 | L2HGDH | 20,93313563 | 22,32547931 | 15,22212186 | 15,47060553 | 6 |
| ENSGALG00000012308 | MANBA | 11,89786869 | 12,7530861 | 8,200203397 | 9,308437527 | 6 |
| ENSGALG00000012346 | | 7,862982605 | 8,864901197 | 5,102603107 | 6,396826485 | 6 |
| ENSGALG00000012569 | FBXO7 | 73,79359843 | 76,51114178 | 60,47045786 | 56,31379388 | 6 |
| ENSGALG00000012575 | TLE1 | 42,64744251 | 40,35590358 | 26,26275038 | 30,7525678 | 6 |
| ENSGALG00000012621 | | 23,12197697 | 21,50098662 | 17,51990666 | 20,73745298 | 6 |

| | | | | | | |
|---|---|---|---|---|---|---|
| ENSGALG00000012747 | NEDD9 | 51,16943075 | 46,55544661 | 25,2270541 | 28,40372824 | 6 |
| ENSGALG00000012748 | ELOVL2 | 6,703664016 | 5,77345016 | 4,279673705 | 3,893667093 | 6 |
| ENSGALG00000012796 | KIAA1549 | 51,09108109 | 44,95257214 | 38,63565476 | 39,44896655 | 6 |
| ENSGALG00000013182 | | 64,54611055 | 57,87116822 | 44,54062532 | 41,58771622 | 6 |
| ENSGALG00000013194 | | 37,97153678 | 35,58529342 | 22,21115632 | 30,79003586 | 6 |
| ENSGALG00000013207 | CTNNAL1 | 96,43762659 | 88,20907865 | 78,48817168 | 78,03326849 | 6 |
| ENSGALG00000013297 | BBOX1 | 23,95421511 | 20,28591096 | 13,84454117 | 17,01177863 | 6 |
| ENSGALG00000013363 | | 59,76145701 | 53,34909219 | 27,20240346 | 30,97263445 | 6 |
| ENSGALG00000013505 | | 3,889722018 | 3,478768697 | 2,452222386 | 2,376785773 | 6 |
| ENSGALG00000013869 | IL20RA | 6,972034921 | 7,7365372 | 3,434256512 | 4,89649334 | 6 |
| ENSGALG00000013948 | RASL11B | 44,13163783 | 48,97517759 | 18,53083963 | 33,62688077 | 6 |
| ENSGALG00000014033 | | 104,7477162 | 94,29244852 | 81,03494353 | 95,32221883 | 6 |
| ENSGALG00000014094 | STK38L | 63,92061824 | 56,75448411 | 49,03019712 | 53,65761291 | 6 |
| ENSGALG00000014262 | APBB2 | 50,41157639 | 49,52225086 | 39,71176444 | 41,3726837 | 6 |
| ENSGALG00000014627 | | 6,186249674 | 6,570860566 | 3,973407227 | 5,195659962 | 6 |
| ENSGALG00000014648 | CETN3 | 120,6378615 | 124,7051507 | 91,07219377 | 107,9915591 | 6 |
| ENSGALG00000014749 | EPHB6 | 39,83982973 | 37,84869471 | 30,92879956 | 29,01876058 | 6 |
| ENSGALG00000014853 | CLVS2 | 20,14220368 | 18,4145602 | 10,94311348 | 13,28443107 | 6 |
| ENSGALG00000014971 | SLC2A9 | 5,990387675 | 5,899169872 | 4,692471627 | 4,901640879 | 6 |
| ENSGALG00000015037 | EVC | 23,92704947 | 26,1129777 | 17,06719452 | 14,17492822 | 6 |
| ENSGALG00000015109 | | 47,46943411 | 46,36864366 | 35,65399625 | 39,11989952 | 6 |
| ENSGALG00000015349 | PIGG | 15,28388747 | 14,19631757 | 12,44050293 | 12,25488892 | 6 |
| ENSGALG00000015368 | TMEM175 | 40,97091402 | 42,09879659 | 30,18566566 | 35,6655243 | 6 |
| ENSGALG00000015399 | FUT10 | 5,766769243 | 5,749814663 | 3,113288123 | 5,094696357 | 6 |
| ENSGALG00000015512 | | 12,75994149 | 13,90362938 | 8,228539011 | 10,70286882 | 6 |
| ENSGALG00000015542 | PLPPR1 | 116,2967487 | 112,3139725 | 75,3404969 | 69,59821363 | 6 |
| ENSGALG00000015593 | EPHA7 | 23,76270266 | 25,30203349 | 15,93621129 | 16,69520231 | 6 |
| ENSGALG00000015628 | MSANTD1 | 27,98934213 | 27,89487479 | 19,00355994 | 16,65250297 | 6 |
| ENSGALG00000015661 | | 25,53972785 | 27,31771949 | 16,46254751 | 21,18719894 | 6 |
| ENSGALG00000015702 | | 165,4779657 | 153,2689949 | 126,1182243 | 131,223407 | 6 |
| ENSGALG00000015713 | FAM53A | 97,02921321 | 94,00902169 | 72,8675515 | 69,6370911 | 6 |
| ENSGALG00000015749 | | 21,2120221 | 17,80838104 | 14,06053735 | 17,19977004 | 6 |
| ENSGALG00000015812 | SMIM8 | 22,70170198 | 23,41415321 | 15,5950842 | 19,1637852 | 6 |
| ENSGALG00000015880 | HMGN3 | 377,9392173 | 394,1557704 | 311,1143085 | 339,3443829 | 6 |
| ENSGALG00000016027 | | 94,40142812 | 100,8328093 | 71,31951897 | 75,50602922 | 6 |
| ENSGALG00000016067 | HMG14 | 441,5841588 | 451,8120476 | 376,4578579 | 378,3157428 | 6 |
| ENSGALG00000016100 | | 16,68836719 | 14,10822856 | 9,670667046 | 14,44480754 | 6 |
| ENSGALG00000016161 | | 20,22006123 | 21,56436875 | 14,06276308 | 15,46768923 | 6 |
| ENSGALG00000016222 | NDP | 75,17879418 | 59,52253055 | 36,29361092 | 33,58711701 | 6 |
| ENSGALG00000016318 | | 27,27843983 | 23,46508248 | 18,58560473 | 20,47415343 | 6 |
| ENSGALG00000016390 | | 37,86603951 | 38,28740842 | 29,30950252 | 35,44055736 | 6 |
| ENSGALG00000016403 | | 266,8591886 | 263,757375 | 206,1924561 | 172,5882396 | 6 |
| ENSGALG00000016411 | | 14,25340121 | 13,22735653 | 5,731016335 | 9,570264498 | 6 |
| ENSGALG00000016477 | | 32,59657778 | 30,23039623 | 26,7611813 | 26,75620295 | 6 |
| ENSGALG00000016511 | ADGRG2 | 2,386466849 | 2,957863341 | 0,902910309 | 1,512144222 | 6 |
| ENSGALG00000016570 | | 5,798752701 | 5,090455098 | 2,401852442 | 4,386433754 | 6 |
| ENSGALG00000016636 | | 5,617816157 | 5,062765544 | 2,035430911 | 2,908022952 | 6 |
| ENSGALG00000016731 | | 239,7537815 | 224,2247214 | 208,7586328 | 215,0388526 | 6 |
| ENSGALG00000016895 | DZIP1 | 27,04772767 | 25,34766727 | 21,42357243 | 21,88134556 | 6 |
| ENSGALG00000017015 | RNASEH2B | 56,39861841 | 51,8363735 | 44,83361055 | 48,79351186 | 6 |
| ENSGALG00000017032 | | 58,5366442 | 56,1349687 | 48,61797983 | 48,88052127 | 6 |
| ENSGALG00000017074 | | 4,074357168 | 3,691118729 | 1,412239719 | 2,952723128 | 6 |
| ENSGALG00000017119 | TNFRSF19 | 58,66491746 | 57,05921994 | 42,25493843 | 32,20227457 | 6 |
| ENSGALG00000017133 | EEF1AKMT1 | 26,99940931 | 29,0525506 | 20,74451252 | 21,42944072 | 6 |
| ENSGALG00000017195 | PGR | 3,608790529 | 2,632589836 | 1,517382064 | 1,94031908 | 6 |
| ENSGALG00000017229 | FAT3 | 8,89802285 | 8,313903578 | 6,609729949 | 6,008519085 | 6 |
| ENSGALG00000017247 | | 35,90192669 | 33,73272811 | 24,29212924 | 26,07653344 | 6 |
| ENSGALG00000017302 | SLCO2B1 | 1,429813469 | 1,146573087 | 0,673898438 | 0,668516203 | 6 |
| ENSGALG00000017371 | | 51,38692014 | 50,91225956 | 38,09521913 | 45,83153263 | 6 |
| ENSGALG00000017405 | NPR3 | 23,00399694 | 22,53783009 | 13,33544434 | 18,73145328 | 6 |
| ENSGALG00000019198 | | 17,59843277 | 15,08692745 | 10,51068089 | 8,621485841 | 6 |
| ENSGALG00000019514 | CNDP2 | 182,9891291 | 180,1386682 | 158,0964217 | 167,2312883 | 6 |
| ENSGALG00000019719 | KRT19 | 37,68434018 | 33,76138595 | 21,75346913 | 23,72609315 | 6 |
| ENSGALG00000020575 | | 8,11802815 | 7,979917996 | 3,72206627 | 6,630432556 | 6 |

| | | | | | | |
|---|---|---|---|---|---|---|
| ENSGALG00000020615 | STK32C | 11,47277599 | 10,0537539 | 4,973755997 | 7,984705212 | 6 |
| ENSGALG00000020688 | | 7,755595976 | 7,74751763 | 3,581525145 | 6,288357343 | 6 |
| ENSGALG00000020876 | | 2,382805079 | 2,166538193 | 0,848276969 | 1,89263607 | 6 |
| ENSGALG00000020956 | | 38,20475834 | 33,66535769 | 30,02776671 | 33,98390736 | 6 |
| ENSGALG00000020984 | FAM163A | 20,23037736 | 16,5164079 | 11,68786047 | 14,02896446 | 6 |
| ENSGALG00000021285 | | 186,2854475 | 165,9217969 | 136,042098 | 155,4656808 | 6 |
| ENSGALG00000021653 | IFT22 | 101,3073949 | 105,5647369 | 78,19787743 | 75,492151 | 6 |
| ENSGALG00000022891 | | 63,01409828 | 62,3114166 | 49,31573941 | 51,86061496 | 6 |
| ENSGALG00000022967 | | 19,86053476 | 14,53555584 | 9,909871032 | 11,74545585 | 6 |
| ENSGALG00000023073 | | 75,25252995 | 70,70073732 | 46,37176598 | 61,58963078 | 6 |
| ENSGALG00000023279 | EPN3 | 18,11156599 | 18,44465948 | 11,60965438 | 15,0198188 | 6 |
| ENSGALG00000023542 | | 44,21934953 | 41,11173749 | 33,22451866 | 36,77576397 | 6 |
| ENSGALG00000023683 | TCTN2 | 77,20303141 | 74,18428746 | 64,19070348 | 58,30899806 | 6 |
| ENSGALG00000023904 | FIBIN | 19,51600353 | 16,39135199 | 8,430598252 | 11,5313733 | 6 |
| ENSGALG00000024294 | | 3,558273438 | 3,080519379 | 1,266138207 | 1,949493615 | 6 |
| ENSGALG00000024481 | | 133,9861745 | 137,3212927 | 96,10190154 | 120,9686545 | 6 |
| ENSGALG00000024486 | | 127,4593345 | 112,050928 | 75,08512045 | 74,30239813 | 6 |
| ENSGALG00000025915 | ARRDC3 | 26,46755421 | 22,42622815 | 19,05714962 | 23,18559721 | 6 |
| ENSGALG00000026055 | PALM | 62,96970401 | 58,46982113 | 37,62754481 | 51,55881293 | 6 |
| ENSGALG00000026119 | | 19,50030797 | 15,03597578 | 10,89450307 | 13,56917513 | 6 |
| ENSGALG00000026153 | | 13,21809934 | 13,27180391 | 6,109358729 | 11,47472691 | 6 |
| ENSGALG00000026351 | | 7,714996646 | 7,132791831 | 4,733453719 | 4,737550727 | 6 |
| ENSGALG00000026388 | | 7,176755468 | 5,70158915 | 3,356921795 | 5,377783467 | 6 |
| ENSGALG00000026424 | TOR2A | 49,43334651 | 51,09796496 | 40,22928877 | 42,49356237 | 6 |
| ENSGALG00000026460 | | 35,89548571 | 39,92113818 | 22,86822683 | 27,96586662 | 6 |
| ENSGALG00000026544 | | 58,65981865 | 57,10160799 | 41,59272235 | 50,81686876 | 6 |
| ENSGALG00000026560 | | 36,96829613 | 26,69799391 | 17,90105267 | 27,48806165 | 6 |
| ENSGALG00000026622 | | 3,176811988 | 2,665491808 | 1,529135988 | 2,006855906 | 6 |
| ENSGALG00000026704 | GPX8 | 14,6970356 | 14,21488529 | 8,762242734 | 6,869731027 | 6 |
| ENSGALG00000026722 | UNC119B | 143,9059999 | 141,8749309 | 120,3644467 | 119,8965562 | 6 |
| ENSGALG00000026941 | | 11,45825275 | 12,65274753 | 7,360856213 | 7,871403016 | 6 |
| ENSGALG00000027154 | | 32,85298841 | 26,87435096 | 15,98536481 | 21,99669163 | 6 |
| ENSGALG00000027300 | | 255,9415831 | 208,7324314 | 140,1583855 | 136,9067811 | 6 |
| ENSGALG00000027454 | CENPW | 133,7719182 | 117,0934874 | 105,1892275 | 115,178855 | 6 |
| ENSGALG00000027498 | FOXB1 | 141,2191776 | 137,5736463 | 78,19797441 | 120,6686286 | 6 |
| ENSGALG00000027648 | | 297,8558779 | 297,8555526 | 235,0735679 | 265,6448529 | 6 |
| ENSGALG00000027655 | TRABD2B | 3,732000135 | 2,929243457 | 1,949007079 | 2,75895498 | 6 |
| ENSGALG00000027714 | ATG10 | 14,65174394 | 14,12262445 | 10,09229515 | 11,13826046 | 6 |
| ENSGALG00000027891 | | 245,592978 | 266,7752548 | 119,3002211 | 176,3890275 | 6 |
| ENSGALG00000027907 | NR2F1 | 167,8585793 | 148,627158 | 100,6374936 | 126,2471647 | 6 |
| ENSGALG00000028005 | GADD45G | 26,61578991 | 21,03730188 | 12,66743452 | 14,80395902 | 6 |
| ENSGALG00000028015 | | 40,97729593 | 39,21000061 | 27,73413561 | 27,479133 | 6 |
| ENSGALG00000028142 | HACD4 | 1,371297566 | 1,176318136 | 0,666595527 | 0,732081177 | 6 |
| ENSGALG00000028168 | | 21,67520772 | 22,20929179 | 18,06713907 | 16,59662597 | 6 |
| ENSGALG00000028221 | | 9,866386893 | 9,886670244 | 5,735567879 | 7,645874692 | 6 |
| ENSGALG00000028506 | NHSL2 | 18,71016948 | 15,75409848 | 8,290834081 | 13,78916943 | 6 |
| ENSGALG00000028508 | | 19,24169204 | 18,49325928 | 11,74503381 | 15,41263549 | 6 |
| ENSGALG00000028745 | | 8,953044339 | 8,076698766 | 3,902548389 | 4,23747302 | 6 |
| ENSGALG00000028931 | | 107,4437216 | 87,27433427 | 55,7926251 | 77,21406238 | 6 |
| ENSGALG00000029012 | | 14,73441004 | 15,72458847 | 5,28285304 | 8,311412343 | 6 |
| ENSGALG00000000040 | | 23,44475508 | 28,21280039 | 14,22402533 | 24,43401832 | 7 |
| ENSGALG00000000209 | PRNP | 10,74066283 | 12,46930325 | 6,402314152 | 9,988641721 | 7 |
| ENSGALG00000000231 | | 149,0870697 | 185,9234478 | 117,8750199 | 142,1973043 | 7 |
| ENSGALG00000000316 | | 33,64295455 | 43,6406022 | 24,05590723 | 36,87840136 | 7 |
| ENSGALG00000000426 | | 10,27991254 | 12,41662544 | 6,880846351 | 11,34770947 | 7 |
| ENSGALG00000000674 | MYO1D | 4,406809187 | 6,316304931 | 2,455472407 | 4,256521683 | 7 |
| ENSGALG00000000713 | ZFHX3 | 39,06233202 | 43,14699726 | 29,26088997 | 38,05997761 | 7 |
| ENSGALG00000000897 | | 4,978147826 | 7,378389371 | 2,837088455 | 5,487379033 | 7 |
| ENSGALG00000000899 | | 0,963046208 | 5,946365074 | 0,475413557 | 2,252274664 | 7 |
| ENSGALG00000001024 | TPRG1L | 120,9971135 | 152,3547079 | 103,4903126 | 132,788969 | 7 |
| ENSGALG00000001100 | | 13,95505436 | 17,65108414 | 9,356507162 | 15,50884815 | 7 |
| ENSGALG00000001129 | | 23,86423909 | 32,84739588 | 12,21283125 | 23,798218 | 7 |
| ENSGALG00000001329 | EPB41 | 65,34880758 | 78,02186828 | 51,24432241 | 67,40825669 | 7 |
| ENSGALG00000001446 | | 64,62473612 | 87,21072718 | 48,6331509 | 76,52155764 | 7 |

| | | | | | | |
|---|---|---|---|---|---|---|
| ENSGALG00000001449 | STRA6 | 45,37016661 | 58,55812578 | 27,01253734 | 47,99719354 | 7 |
| ENSGALG00000001475 | | 3350,976054 | 3963,682871 | 2691,829167 | 3366,524715 | 7 |
| ENSGALG00000001492 | NDRG3 | 65,58399327 | 69,55303732 | 54,78233205 | 62,46719773 | 7 |
| ENSGALG00000001604 | | 52,26872952 | 56,56561231 | 40,78881554 | 53,29231133 | 7 |
| ENSGALG00000001727 | LZTS1 | 8,163488473 | 25,58923197 | 4,764125596 | 14,44205782 | 7 |
| ENSGALG00000001795 | | 10,08663077 | 20,53677 | 4,664121167 | 12,34777599 | 7 |
| ENSGALG00000001816 | MAGI3 | 22,47607338 | 29,09793529 | 15,71484571 | 21,83158076 | 7 |
| ENSGALG00000001966 | | 25,7785394 | 36,09230687 | 17,73218323 | 27,60168099 | 7 |
| ENSGALG00000001996 | DNALI1 | 33,7845802 | 40,3539152 | 23,78692763 | 32,34237044 | 7 |
| ENSGALG00000002016 | PRKCD | 9,534583077 | 25,41833693 | 5,509761135 | 13,37710012 | 7 |
| ENSGALG00000002055 | hairy1 | 100,9426832 | 191,0596931 | 71,9414387 | 159,5868237 | 7 |
| ENSGALG00000002098 | GRIK3 | 3,884532135 | 6,948087025 | 1,414247851 | 2,595929492 | 7 |
| ENSGALG00000002125 | TMOD1 | 19,78933687 | 23,97315207 | 12,05769249 | 16,67488273 | 7 |
| ENSGALG00000002169 | PLEKHB2 | 32,6969416 | 49,0144176 | 25,10240291 | 39,31342986 | 7 |
| ENSGALG00000002225 | RFFL | 28,82366062 | 33,4994994 | 22,79136152 | 26,46140023 | 7 |
| ENSGALG00000002437 | FAM69B | 10,34052265 | 18,5207246 | 6,353529848 | 13,77821286 | 7 |
| ENSGALG00000002484 | CRYM | 52,30050528 | 59,7062453 | 38,85736704 | 51,40699758 | 7 |
| ENSGALG00000002610 | | 13,09856503 | 27,92032271 | 8,551552499 | 20,41307335 | 7 |
| ENSGALG00000002692 | | 7,601162153 | 8,489273999 | 4,867865763 | 6,523571836 | 7 |
| ENSGALG00000002707 | CHRNB2 | 22,21560311 | 33,9138735 | 14,13296126 | 22,61297767 | 7 |
| ENSGALG00000002712 | | 98,06557273 | 124,0459701 | 60,94436313 | 78,26408645 | 7 |
| ENSGALG00000002742 | TMEM132B | 21,18839394 | 28,46994537 | 12,23650943 | 23,57680408 | 7 |
| ENSGALG00000002749 | | 15,4477782 | 18,83801502 | 9,210681708 | 15,54271584 | 7 |
| ENSGALG00000002777 | | 206,4893527 | 235,0657741 | 145,3623973 | 217,0270845 | 7 |
| ENSGALG00000002804 | RND2 | 7,564698815 | 11,37308686 | 3,691136555 | 7,382372138 | 7 |
| ENSGALG00000002818 | | 76,01900762 | 92,05204683 | 46,3235432 | 73,16432864 | 7 |
| ENSGALG00000002840 | GRM4 | 38,07446155 | 45,47238554 | 23,80739886 | 33,41537259 | 7 |
| ENSGALG00000002891 | ACADL | 101,9928444 | 126,3938118 | 85,60119687 | 110,7373997 | 7 |
| ENSGALG00000002940 | | 63,67353296 | 73,60062167 | 49,11114405 | 52,63840174 | 7 |
| ENSGALG00000003099 | PCTP | 55,94909963 | 68,55104571 | 39,17056499 | 50,10135455 | 7 |
| ENSGALG00000003135 | ANK3 | 29,82803871 | 36,54263 | 22,66405296 | 23,3794697 | 7 |
| ENSGALG00000003185 | SCPEP1 | 39,41581989 | 47,08505296 | 26,77264651 | 36,20660242 | 7 |
| ENSGALG00000003345 | | 8,011075734 | 13,2054126 | 4,141316066 | 10,41345166 | 7 |
| ENSGALG00000003445 | MGST3 | 164,0311496 | 177,1818225 | 134,4152099 | 167,0547988 | 7 |
| ENSGALG00000003474 | MED20 | 110,444728 | 116,7026163 | 88,99338596 | 109,2480186 | 7 |
| ENSGALG00000003580 | MMP2 | 7,325674272 | 32,10481709 | 4,372959973 | 11,83675111 | 7 |
| ENSGALG00000003588 | | 41,80910808 | 61,28137773 | 18,99649734 | 38,34816043 | 7 |
| ENSGALG00000003589 | VTN | 114,2225217 | 150,0745088 | 69,50664764 | 114,9001653 | 7 |
| ENSGALG00000003595 | SARM1 | 30,68252668 | 34,98549632 | 21,8893545 | 27,60935273 | 7 |
| ENSGALG00000003648 | SVIP | 22,22982974 | 29,46021773 | 15,12642168 | 17,95720385 | 7 |
| ENSGALG00000003832 | TEX264 | 15,86809856 | 27,52547475 | 11,7554828 | 23,97522593 | 7 |
| ENSGALG00000003999 | | 13,72215726 | 16,36238679 | 11,0995887 | 12,90633719 | 7 |
| ENSGALG00000004038 | ADAL | 25,17562849 | 34,56044312 | 19,02897515 | 26,71696561 | 7 |
| ENSGALG00000004236 | TRAF3IP1 | 33,67265581 | 37,7057156 | 28,80217302 | 29,82775189 | 7 |
| ENSGALG00000004323 | RASAL2 | 17,46458791 | 24,05795653 | 11,68718553 | 21,19348944 | 7 |
| ENSGALG00000004448 | BRINP2 | 7,422301548 | 13,10562437 | 2,669166415 | 9,017834641 | 7 |
| ENSGALG00000004472 | ASTN1 | 45,27241295 | 56,98089896 | 25,40185133 | 42,4550224 | 7 |
| ENSGALG00000004581 | NDST1 | 36,77466572 | 44,71190372 | 29,08485213 | 41,89895888 | 7 |
| ENSGALG00000004593 | SULF2 | 30,06159727 | 56,14934678 | 19,86175058 | 37,91786843 | 7 |
| ENSGALG00000004623 | AGTRAP | 23,55850976 | 32,14710496 | 15,71775217 | 22,70328234 | 7 |
| ENSGALG00000004676 | | 44,69765223 | 67,37462526 | 26,5567982 | 52,68656003 | 7 |
| ENSGALG00000004705 | BUD31 | 190,1039203 | 210,7250422 | 159,5584737 | 195,6353874 | 7 |
| ENSGALG00000004769 | | 231,0703098 | 265,9633181 | 188,117676 | 223,1112578 | 7 |
| ENSGALG00000004812 | FAM129A | 3,011515596 | 4,419696057 | 1,922761698 | 3,474926272 | 7 |
| ENSGALG00000004844 | | 38,76266742 | 46,85070174 | 21,59843211 | 33,82333789 | 7 |
| ENSGALG00000004851 | | 102,8684263 | 153,4847972 | 80,99247061 | 119,9569497 | 7 |
| ENSGALG00000004873 | | 21,88445189 | 28,72943819 | 17,20757599 | 23,79151605 | 7 |
| ENSGALG00000005062 | FoxN4 | 4,362960232 | 23,41120468 | 2,672268849 | 13,15759672 | 7 |
| ENSGALG00000005076 | | 2,97258771 | 5,738604455 | 1,868991974 | 4,403336028 | 7 |
| ENSGALG00000005203 | OLFM3 | 9,917656348 | 19,51400136 | 6,107981318 | 12,50996987 | 7 |
| ENSGALG00000005446 | | 433,5296312 | 485,9292058 | 357,6286444 | 378,096764 | 7 |
| ENSGALG00000005502 | KCNG4 | 0,912952006 | 1,203243361 | 0,297105278 | 0,60110466 | 7 |
| ENSGALG00000005539 | | 10,06496779 | 11,46804537 | 4,811243877 | 8,97883854 | 7 |
| ENSGALG00000005549 | CUEDC1 | 24,9585639 | 31,73680379 | 17,96776786 | 28,07856242 | 7 |

| | | | | | | |
|---|---|---|---|---|---|---|
| ENSGALG00000005626 | TPST2 | 11,80433122 | 16,81885809 | 6,363075005 | 13,39419824 | 7 |
| ENSGALG00000005657 | CRHR2 | 16,41937448 | 21,14097462 | 9,617236635 | 17,9916876 | 7 |
| ENSGALG00000005901 | PCMTD2 | 15,03826855 | 22,94366732 | 12,20442379 | 18,1485811 | 7 |
| ENSGALG00000005967 | | 16,34607915 | 27,95004368 | 5,734444886 | 19,79268951 | 7 |
| ENSGALG00000006014 | | 11,83239403 | 17,79556194 | 8,524616365 | 16,16215962 | 7 |
| ENSGALG00000006036 | | 51,78288032 | 72,39218737 | 37,65758569 | 61,49569637 | 7 |
| ENSGALG00000006083 | | 4,451611687 | 9,146359521 | 3,028486674 | 6,106851928 | 7 |
| ENSGALG00000006172 | ABCC8 | 10,46414413 | 12,87547866 | 7,792903566 | 8,971268996 | 7 |
| ENSGALG00000006192 | | 6,56151079 | 13,70633898 | 3,575297057 | 7,644373335 | 7 |
| ENSGALG00000006233 | | 73,88654127 | 86,5339275 | 62,67969355 | 65,81661353 | 7 |
| ENSGALG00000006490 | SCN3B | 48,56309698 | 61,96270518 | 30,47526127 | 36,08774643 | 7 |
| ENSGALG00000006588 | LRRC75B | 20,28960188 | 26,05813662 | 15,10431218 | 21,90900245 | 7 |
| ENSGALG00000006606 | | 6,809091308 | 32,17770953 | 5,138243105 | 16,8368564 | 7 |
| ENSGALG00000006780 | | 245,3841891 | 304,5147127 | 196,4819313 | 254,7782702 | 7 |
| ENSGALG00000006792 | VSTM2L | 14,91896564 | 24,34060472 | 8,041765026 | 19,66450542 | 7 |
| ENSGALG00000006857 | SPATA20 | 13,000609 | 15,94295579 | 10,59989797 | 14,33927151 | 7 |
| ENSGALG00000006873 | PHRF1 | 51,794473 | 56,56115769 | 28,13525923 | 43,64214396 | 7 |
| ENSGALG00000006882 | ST8SIA2 | 43,06730195 | 67,38681182 | 26,22541309 | 54,35238193 | 7 |
| ENSGALG00000006942 | | 132,4198867 | 227,8606087 | 86,86056301 | 125,7845777 | 7 |
| ENSGALG00000007076 | CCNY | 18,51574898 | 24,71732273 | 11,69363679 | 18,88976684 | 7 |
| ENSGALG00000007078 | | 2,87252386 | 4,383770523 | 1,005692341 | 3,383060819 | 7 |
| ENSGALG00000007128 | SH3BGRL | 214,2530874 | 229,1837125 | 167,2589166 | 211,0428624 | 7 |
| ENSGALG00000007211 | CDH22 | 1,485142793 | 2,494244579 | 0,548146547 | 1,816562264 | 7 |
| ENSGALG00000007262 | SPG21 | 71,40763929 | 85,58144568 | 58,23479154 | 64,58167991 | 7 |
| ENSGALG00000007732 | RTN4R | 3,699735966 | 6,809590742 | 2,178228147 | 5,983400116 | 7 |
| ENSGALG00000007748 | SLC16A2 | 20,32729004 | 25,80043621 | 12,93321002 | 22,99796604 | 7 |
| ENSGALG00000007780 | EIF4E3 | 18,03876093 | 38,22663339 | 13,64586903 | 30,5542396 | 7 |
| ENSGALG00000007908 | | 5,792965987 | 13,97554053 | 2,389386077 | 7,707065248 | 7 |
| ENSGALG00000007943 | | 19,29727515 | 23,99833693 | 15,17755759 | 15,86135712 | 7 |
| ENSGALG00000007965 | HN1 | 411,0251371 | 434,100802 | 329,5995071 | 408,922843 | 7 |
| ENSGALG00000008438 | SLC35C1 | 124,3184986 | 149,9093654 | 98,43575195 | 132,7600199 | 7 |
| ENSGALG00000008455 | | 32,31963069 | 57,54084189 | 14,64828513 | 25,24367492 | 7 |
| ENSGALG00000008697 | RBM20 | 3,846768505 | 4,926267748 | 2,456023592 | 3,647659329 | 7 |
| ENSGALG00000008700 | PDCD4 | 325,6337309 | 399,5164392 | 238,9464322 | 293,0805554 | 7 |
| ENSGALG00000008728 | PTER | 6,3399657 | 8,156910559 | 4,337055382 | 7,10541263 | 7 |
| ENSGALG00000008755 | | 67,34340554 | 73,50049392 | 52,70850781 | 64,66558339 | 7 |
| ENSGALG00000008818 | IDH1 | 71,66766176 | 91,61503146 | 53,6340664 | 74,86640267 | 7 |
| ENSGALG00000008865 | ALAD | 139,8444823 | 169,0120545 | 111,5900089 | 141,221191 | 7 |
| ENSGALG00000009002 | | 1,28675173 | 1,674918274 | 0,613968423 | 1,412361787 | 7 |
| ENSGALG00000009042 | ISM1 | 19,65944442 | 27,17765292 | 9,838603957 | 17,08408347 | 7 |
| ENSGALG00000009052 | LBH | 37,94812104 | 96,15335335 | 19,71154839 | 42,02747293 | 7 |
| ENSGALG00000009134 | CTTNBP2 | 11,40771871 | 15,88432695 | 8,530144038 | 10,85876386 | 7 |
| ENSGALG00000009212 | MND1 | 66,79405001 | 75,25435741 | 49,15985084 | 63,46334427 | 7 |
| ENSGALG00000009354 | | 11,65567463 | 14,51617495 | 8,805310484 | 9,634955967 | 7 |
| ENSGALG00000009439 | GPR85 | 17,44009003 | 22,44330748 | 10,32374078 | 18,89710187 | 7 |
| ENSGALG00000009478 | | 12,99711452 | 17,0630503 | 6,327520434 | 10,68724831 | 7 |
| ENSGALG00000009536 | | 8,930747832 | 11,26382701 | 4,868019414 | 9,734234384 | 7 |
| ENSGALG00000009537 | PPM1L | 15,19652092 | 19,19973765 | 9,532288845 | 11,64296507 | 7 |
| ENSGALG00000009601 | NELL2 | 110,2961344 | 154,2484566 | 51,22048418 | 99,21561222 | 7 |
| ENSGALG00000009603 | | 11,49975548 | 14,99438031 | 4,75072532 | 9,623024967 | 7 |
| ENSGALG00000009670 | SGCE | 59,43794591 | 65,07796911 | 46,83087493 | 55,09067955 | 7 |
| ENSGALG00000009959 | | 35,62728541 | 41,80367214 | 20,63974566 | 27,91189617 | 7 |
| ENSGALG00000009977 | SMAD1 | 78,48081668 | 94,47814686 | 67,52132303 | 77,34497377 | 7 |
| ENSGALG00000010119 | CDKL2 | 8,919919874 | 17,83271101 | 5,399685531 | 10,94164418 | 7 |
| ENSGALG00000010127 | GLO1 | 231,8522951 | 279,4946021 | 191,8389983 | 204,8112253 | 7 |
| ENSGALG00000010137 | TTC6 | 1,287993124 | 1,948479866 | 0,779674705 | 0,991764128 | 7 |
| ENSGALG00000010139 | SSTR1 | 7,978023763 | 12,61992979 | 4,912549304 | 10,41199792 | 7 |
| ENSGALG00000010269 | KCNAB1 | 10,86190361 | 16,59727193 | 6,653570316 | 13,2184581 | 7 |
| ENSGALG00000010320 | PAWR | 34,60903172 | 62,08502203 | 21,78625211 | 37,50489825 | 7 |
| ENSGALG00000010422 | | 103,3041586 | 158,2231145 | 73,72776386 | 121,1270632 | 7 |
| ENSGALG00000010441 | | 56,2447908 | 64,77433617 | 43,50838618 | 56,84118443 | 7 |
| ENSGALG00000010444 | | 30,2192376 | 32,99747957 | 23,77039516 | 26,95022033 | 7 |
| ENSGALG00000010533 | ELAVL4 | 101,3698774 | 123,7671942 | 80,28112842 | 100,1644983 | 7 |
| ENSGALG00000010611 | CFAP97 | 27,26735677 | 31,60703776 | 21,00245601 | 25,6223608 | 7 |

| | | | | | | |
|---|---|---|---|---|---|---|
| ENSGALG00000010797 | | 2,204962564 | 3,597171829 | 1,242295373 | 2,737107629 | 7 |
| ENSGALG00000010982 | | 34,9067668 | 47,34640946 | 23,85218963 | 36,57797015 | 7 |
| ENSGALG00000010997 | DFNA5 | 30,98330411 | 36,05207494 | 25,73539441 | 33,02230238 | 7 |
| ENSGALG00000011078 | | 29,00849301 | 32,4201808 | 22,47842078 | 27,65933043 | 7 |
| ENSGALG00000011176 | SCRN1 | 73,33672636 | 113,3873336 | 57,54804702 | 76,21746157 | 7 |
| ENSGALG00000011200 | THBS2 | 4,729378665 | 17,96316127 | 2,101963158 | 7,934820548 | 7 |
| ENSGALG00000011238 | WLS | 398,6085265 | 452,0423133 | 298,9173487 | 386,8213626 | 7 |
| ENSGALG00000011276 | BTG1 | 502,7455821 | 558,6321082 | 412,3661019 | 452,7358963 | 7 |
| ENSGALG00000011350 | NEGR1 | 20,33143431 | 23,86084881 | 12,71729034 | 19,59581703 | 7 |
| ENSGALG00000011444 | MFHAS1 | 58,17054063 | 70,52843469 | 40,96561842 | 49,56226035 | 7 |
| ENSGALG00000011655 | IQCB1 | 23,64106498 | 26,10604734 | 13,7260757 | 19,72872665 | 7 |
| ENSGALG00000011656 | | 0,973892515 | 2,15913225 | 0,52062462 | 0,800954632 | 7 |
| ENSGALG00000011709 | | 10,59850516 | 15,02453029 | 4,939474354 | 10,42674333 | 7 |
| ENSGALG00000011778 | | 23,66787702 | 28,8235186 | 17,91116836 | 26,20801542 | 7 |
| ENSGALG00000011798 | HEBP1 | 12,49320521 | 25,56914364 | 7,850759982 | 11,62061774 | 7 |
| ENSGALG00000011810 | | 65,9521746 | 82,87507381 | 51,45272236 | 58,71022908 | 7 |
| ENSGALG00000012044 | | 9,229584329 | 12,71447637 | 7,5350979 | 9,063435059 | 7 |
| ENSGALG00000012057 | | 11,18597673 | 12,61517469 | 8,164876186 | 10,80367383 | 7 |
| ENSGALG00000012092 | ESYT3 | 4,680130309 | 6,111516607 | 2,253806553 | 3,990397078 | 7 |
| ENSGALG00000012116 | | 9,646935596 | 13,32089976 | 6,14467466 | 11,39954133 | 7 |
| ENSGALG00000012185 | PLA2G12A | 102,7974734 | 112,7403772 | 84,43323048 | 105,0643144 | 7 |
| ENSGALG00000012191 | | 32,82020057 | 46,37361014 | 18,84566996 | 32,6990559 | 7 |
| ENSGALG00000012362 | THSD7B | 3,775092996 | 10,36024079 | 2,69688283 | 7,22396186 | 7 |
| ENSGALG00000012377 | HNMT | 1,717135166 | 6,060908343 | 0,573693554 | 1,878896194 | 7 |
| ENSGALG00000012487 | | 24,24348782 | 31,4734152 | 16,11839467 | 23,96854961 | 7 |
| ENSGALG00000012652 | | 10,7529658 | 25,54973014 | 5,663246629 | 17,81444342 | 7 |
| ENSGALG00000012658 | TCP11L2 | 15,92414007 | 33,05220242 | 11,27908373 | 28,17571503 | 7 |
| ENSGALG00000012692 | SIRT5 | 45,36256371 | 63,38567531 | 35,0481337 | 44,42038027 | 7 |
| ENSGALG00000012714 | | 43,94791511 | 52,13807433 | 33,7764496 | 43,47609852 | 7 |
| ENSGALG00000012715 | CAP2 | 8,59191649 | 11,46991422 | 4,148180559 | 9,301130175 | 7 |
| ENSGALG00000012791 | TBXAS1 | 1,864983737 | 3,103081301 | 0,74879117 | 2,141225714 | 7 |
| ENSGALG00000013002 | DAP1 | 32,2061213 | 57,91577834 | 24,03389605 | 42,05183833 | 7 |
| ENSGALG00000013067 | | 137,5096802 | 191,3369487 | 110,2279785 | 152,0190566 | 7 |
| ENSGALG00000013837 | | 59,04391774 | 71,3460663 | 48,21803842 | 49,19810217 | 7 |
| ENSGALG00000013876 | | 202,1069193 | 227,9456838 | 159,7823013 | 213,2983315 | 7 |
| ENSGALG00000013964 | SPATA18 | 5,231802216 | 6,450354247 | 3,531114535 | 5,627475195 | 7 |
| ENSGALG00000013980 | SLC2A12 | 1,779452239 | 4,860254934 | 0,730603455 | 2,144183869 | 7 |
| ENSGALG00000014004 | | 38,19788449 | 43,26576373 | 28,71001642 | 36,16158832 | 7 |
| ENSGALG00000014233 | FBLN1 | 100,2616639 | 153,9996924 | 85,13157946 | 116,6781323 | 7 |
| ENSGALG00000014242 | | 24,47453838 | 32,12228945 | 16,30724171 | 25,76182082 | 7 |
| ENSGALG00000014258 | LIMCH1 | 11,23039493 | 16,58171204 | 4,316562472 | 9,590761217 | 7 |
| ENSGALG00000014300 | | 2,585478889 | 4,334338666 | 1,496149583 | 2,982896803 | 7 |
| ENSGALG00000014496 | Prom1 | 38,69686144 | 62,3200769 | 30,55765133 | 41,6577711 | 7 |
| ENSGALG00000014537 | BMF | 66,59173617 | 87,81261663 | 47,00373868 | 44,31366941 | 7 |
| ENSGALG00000014661 | FAM172A | 53,62830077 | 59,84785505 | 44,237488 | 54,90827811 | 7 |
| ENSGALG00000014662 | RBP5 | 94,12229359 | 110,0266074 | 71,71186194 | 92,49555572 | 7 |
| ENSGALG00000014699 | MEGF10 | 5,113117701 | 7,375190092 | 1,931874889 | 3,210163737 | 7 |
| ENSGALG00000014812 | SIDT1 | 6,795260519 | 16,41334784 | 4,233919069 | 10,62192737 | 7 |
| ENSGALG00000014833 | RPL37 | 1596,687135 | 1720,735968 | 1387,064356 | 1578,22352 | 7 |
| ENSGALG00000014844 | | 19,41962485 | 24,18777381 | 13,76691142 | 17,52298328 | 7 |
| ENSGALG00000014941 | | 4,499019336 | 9,516121976 | 2,714813327 | 8,305086794 | 7 |
| ENSGALG00000015075 | | 14,78424772 | 16,24793751 | 7,992995674 | 13,23798293 | 7 |
| ENSGALG00000015183 | RAB13 | 27,51711853 | 44,6852246 | 16,72602217 | 36,46505352 | 7 |
| ENSGALG00000015297 | FOXO3 | 24,37599027 | 33,24133814 | 13,94064542 | 26,23468786 | 7 |
| ENSGALG00000015537 | MAN2B2 | 30,3066011 | 45,5128118 | 21,78564425 | 29,42547877 | 7 |
| ENSGALG00000015543 | MRPL50 | 97,51826513 | 108,3654216 | 76,17159113 | 102,4663452 | 7 |
| ENSGALG00000015551 | GRIN3A | 5,679152207 | 17,73134218 | 3,841307699 | 16,43553314 | 7 |
| ENSGALG00000015660 | SNX30 | 42,56777807 | 79,83420568 | 30,02031755 | 56,72491395 | 7 |
| ENSGALG00000015762 | LYRM2 | 37,69692925 | 47,81440333 | 26,81605255 | 32,66012008 | 7 |
| ENSGALG00000015768 | | 28,37209383 | 38,55451722 | 16,41325642 | 31,2891459 | 7 |
| ENSGALG00000015774 | CYYR1 | 73,07270964 | 93,2325043 | 46,68077123 | 68,44045186 | 7 |
| ENSGALG00000015780 | PNRC1 | 344,5438061 | 552,0456497 | 280,6093647 | 398,9770095 | 7 |
| ENSGALG00000015903 | | 11,13980945 | 17,28591052 | 7,220689109 | 10,09139434 | 7 |
| ENSGALG00000015967 | LRRTM1 | 3,686312959 | 5,438787671 | 1,609278461 | 3,628427823 | 7 |

| | | | | | | |
|---|---|---|---|---|---|---|
| ENSGALG00000015987 | | 23,3708759 | 36,39569956 | 15,7535009 | 22,36869225 | 7 |
| ENSGALG00000016223 | | 3,343218403 | 5,947487934 | 1,855740174 | 4,018942369 | 7 |
| ENSGALG00000016251 | | 93,77894807 | 167,1461162 | 59,86796921 | 128,2348016 | 7 |
| ENSGALG00000016289 | DST | 15,90950987 | 19,20924281 | 11,20078851 | 13,75441035 | 7 |
| ENSGALG00000016322 | | 31,14927638 | 36,17694833 | 22,78652856 | 32,10595086 | 7 |
| ENSGALG00000016463 | FAM49A | 89,87725864 | 97,84634246 | 78,24594873 | 86,88895017 | 7 |
| ENSGALG00000016518 | PHKA2 | 22,70891315 | 25,28803726 | 16,10212794 | 20,50180035 | 7 |
| ENSGALG00000016543 | NHS | 23,2035252 | 27,27258538 | 13,39269629 | 22,59847883 | 7 |
| ENSGALG00000016549 | CTPS2 | 7,332248006 | 10,148353 | 5,828515996 | 8,094284 | 7 |
| ENSGALG00000016551 | AP1S2 | 150,331608 | 388,8047531 | 100,7222479 | 269,0706899 | 7 |
| ENSGALG00000016697 | MUT | 33,31728166 | 36,06625525 | 27,10395955 | 31,40871266 | 7 |
| ENSGALG00000016870 | TMTC4 | 37,55391684 | 53,03696736 | 30,09752082 | 43,11923341 | 7 |
| ENSGALG00000016945 | LECT1 | 36,93378975 | 51,37890081 | 22,55103978 | 46,08609363 | 7 |
| ENSGALG00000017036 | LHFP | 57,15964101 | 140,0778702 | 35,52548196 | 97,5065896 | 7 |
| ENSGALG00000017072 | N4BP2L1 | 13,74550565 | 22,46637191 | 10,03427041 | 19,42210549 | 7 |
| ENSGALG00000017106 | ATP8A2 | 4,502878382 | 10,93890475 | 2,617722948 | 6,161186837 | 7 |
| ENSGALG00000017219 | C11orf54 | 30,51387908 | 33,47914202 | 23,10946363 | 29,3091597 | 7 |
| ENSGALG00000017321 | PLEKHB1 | 21,68327263 | 24,29861118 | 15,96522023 | 20,23831437 | 7 |
| ENSGALG00000017326 | | 51,59859703 | 55,00738274 | 41,05442188 | 47,2276208 | 7 |
| ENSGALG00000017329 | FCHSD2 | 14,45697367 | 17,25365137 | 11,28085158 | 16,13712426 | 7 |
| ENSGALG00000017331 | | 3,159907146 | 5,517577508 | 1,862654243 | 4,660927839 | 7 |
| ENSGALG00000017332 | KLHL35 | 12,22287679 | 16,1880175 | 5,959667234 | 8,934592728 | 7 |
| ENSGALG00000017334 | | 8,363048271 | 10,26995253 | 6,118966477 | 8,105085971 | 7 |
| ENSGALG00000017342 | | 3,582341309 | 4,647630554 | 1,063461596 | 2,894726481 | 7 |
| ENSGALG00000017374 | | 7,866867941 | 11,57898563 | 3,96740595 | 10,0765853 | 7 |
| ENSGALG00000017926 | 7SK | 13,43570614 | 17,43926497 | 5,12370302 | 13,13564085 | 7 |
| ENSGALG00000018624 | | 211,612459 | 247,6923528 | 121,693404 | 178,5967543 | 7 |
| ENSGALG00000019280 | | 31,88812177 | 75,53173467 | 15,04830691 | 37,69512136 | 7 |
| ENSGALG00000020376 | | 138,5609605 | 183,9703067 | 71,81940611 | 129,9222041 | 7 |
| ENSGALG00000020528 | INIP | 136,2608495 | 170,1147196 | 109,1941668 | 152,6630137 | 7 |
| ENSGALG00000020626 | | 21,17069503 | 23,62411008 | 14,87384185 | 20,507544 | 7 |
| ENSGALG00000021021 | ARL4C | 275,3523921 | 337,3144485 | 220,9487353 | 242,8119461 | 7 |
| ENSGALG00000021344 | | 9,857224916 | 10,97152481 | 7,024591987 | 9,213278791 | 7 |
| ENSGALG00000021972 | | 101,8915641 | 119,2500243 | 83,48016628 | 106,4157224 | 7 |
| ENSGALG00000022713 | | 139,0657773 | 202,1677216 | 94,76044509 | 146,646949 | 7 |
| ENSGALG00000022718 | | 6,91125738 | 10,33059383 | 3,599725653 | 8,616716382 | 7 |
| ENSGALG00000022838 | | 6,670263015 | 8,568090621 | 4,322962094 | 6,400590802 | 7 |
| ENSGALG00000022896 | IRX4 | 0,349425233 | 0,593038773 | 0,239827947 | 0,525323997 | 7 |
| ENSGALG00000023042 | | 11,1327311 | 13,31359085 | 7,176717915 | 11,03847192 | 7 |
| ENSGALG00000023131 | PDXP | 56,87831586 | 59,99147542 | 45,64640553 | 52,21241518 | 7 |
| ENSGALG00000023189 | UMAD1 | 64,4654313 | 78,99153297 | 45,33959426 | 68,96410803 | 7 |
| ENSGALG00000023621 | SFXN3 | 7,518241748 | 11,9378823 | 4,437892273 | 10,8074635 | 7 |
| ENSGALG00000023655 | | 27,08945009 | 36,20494428 | 18,57596443 | 31,99177494 | 7 |
| ENSGALG00000023738 | | 48,2607002 | 63,37886353 | 23,3954474 | 42,14401685 | 7 |
| ENSGALG00000023751 | | 2,121704326 | 39,53753646 | 8,474198433 | 21,85058761 | 7 |
| ENSGALG00000023812 | | 15,06664138 | 19,18104403 | 10,25652845 | 15,75468054 | 7 |
| ENSGALG00000023881 | PLXDC1 | 4,82633085 | 5,58847381 | 2,844462158 | 4,814461859 | 7 |
| ENSGALG00000025163 | gga-mir-1627 | 110,9384893 | 123,3023999 | 64,38404565 | 94,51602649 | 7 |
| ENSGALG00000025698 | 5_8S_rRNA | 214,2147206 | 274,3953045 | 109,3746589 | 240,8884729 | 7 |
| ENSGALG00000025818 | | 34,66056389 | 52,36778343 | 27,68112335 | 42,84794983 | 7 |
| ENSGALG00000025977 | GADD45A | 65,82047393 | 81,42861239 | 46,15966381 | 67,89681053 | 7 |
| ENSGALG00000026042 | | 16,59308302 | 28,8879479 | 8,277397044 | 18,77618411 | 7 |
| ENSGALG00000026103 | | 9,300012163 | 12,28619669 | 3,508789346 | 8,103143537 | 7 |
| ENSGALG00000026113 | | 55,31332896 | 69,44178259 | 41,36543103 | 62,22853953 | 7 |
| ENSGALG00000026180 | | 43,38860094 | 74,08650761 | 27,9624619 | 44,23281888 | 7 |
| ENSGALG00000026437 | | 53,32978665 | 106,3553563 | 42,99932252 | 73,23714682 | 7 |
| ENSGALG00000026505 | | 6,840702248 | 11,78765652 | 4,728224671 | 8,990305294 | 7 |
| ENSGALG00000026600 | C1orf198 | 62,51064096 | 97,06115423 | 52,49014482 | 69,67854538 | 7 |
| ENSGALG00000026870 | | 22,81793746 | 48,40836989 | 17,04944091 | 35,73639395 | 7 |
| ENSGALG00000027027 | | 12,17687155 | 19,36749454 | 8,327209902 | 16,49890464 | 7 |
| ENSGALG00000027058 | EMP2 | 9,670752435 | 14,47252177 | 6,676394562 | 11,60111876 | 7 |
| ENSGALG00000027173 | TMEM159 | 101,3603219 | 133,3366653 | 66,720424 | 98,09898988 | 7 |
| ENSGALG00000027337 | | 35,30559561 | 46,3009554 | 28,71685084 | 40,66049371 | 7 |
| ENSGALG00000027351 | LRRC71 | 12,41099266 | 13,81465571 | 7,63665904 | 11,62692947 | 7 |

| | | | | | | |
|---|---|---|---|---|---|---|
| ENSGALG00000027377 | | 13,23118924 | 17,08047841 | 8,393624566 | 12,22092941 | 7 |
| ENSGALG00000027418 | HINT2 | 54,35709786 | 65,97192429 | 37,50431959 | 54,53621881 | 7 |
| ENSGALG00000027511 | | 94,02464102 | 203,238416 | 68,70151952 | 146,2623945 | 7 |
| ENSGALG00000027534 | | 22,63698 | 42,16451109 | 16,01154358 | 30,54121339 | 7 |
| ENSGALG00000027695 | PCBP4 | 162,956744 | 178,0507957 | 132,4616368 | 163,8638337 | 7 |
| ENSGALG00000027778 | | 0,760117162 | 7,843427702 | 2,238665183 | 3,422613312 | 7 |
| ENSGALG00000027793 | SCN9A | 2,260801542 | 2,687945527 | 1,301123732 | 1,749485357 | 7 |
| ENSGALG00000027904 | | 130,5879679 | 138,1008328 | 105,1857775 | 127,8946055 | 7 |
| ENSGALG00000027920 | | 8,224921324 | 11,67541101 | 4,941372418 | 10,16266736 | 7 |
| ENSGALG00000028031 | C1QL1 | 3,855114072 | 6,583157269 | 1,783653583 | 5,460140979 | 7 |
| ENSGALG00000028187 | FUNDC1 | 50,26900328 | 65,59407968 | 42,88306331 | 53,15622678 | 7 |
| ENSGALG00000028234 | | 4,065747386 | 5,588221597 | 1,848844705 | 4,176917872 | 7 |
| ENSGALG00000028281 | | 0,539773136 | 0,627847197 | 0,388756816 | 0,499963427 | 7 |
| ENSGALG00000028284 | PTX3 | 1,360269657 | 1,613746408 | 0,640931131 | 1,214844992 | 7 |
| ENSGALG00000028371 | | 1205,986207 | 2113,34695 | 600,0776823 | 1309,782147 | 7 |
| ENSGALG00000028443 | | 106,3466323 | 117,1726174 | 52,65285501 | 90,77932913 | 7 |
| ENSGALG00000028567 | MYL9 | 25,71491939 | 30,84447999 | 19,34651722 | 28,9008983 | 7 |
| ENSGALG00000028605 | | 33,48063752 | 44,87000567 | 24,5313162 | 38,26157979 | 7 |
| ENSGALG00000028635 | | 36,25602462 | 40,97336372 | 25,19241388 | 34,21441251 | 7 |
| ENSGALG00000028946 | | 2,681879198 | 4,752514748 | 1,279230013 | 3,51800798 | 7 |
| ENSGALG00000029037 | | 150,1495751 | 180,3499614 | 124,9709847 | 155,381485 | 7 |
| ENSGALG00000029102 | PXYLP1 | 43,95109752 | 57,83768431 | 29,96371923 | 35,84248398 | 7 |
| ENSGALG00000029150 | AK1 | 74,20126426 | 117,6906446 | 51,63919399 | 106,9208251 | 7 |
| ENSGALG00000000402 | LOXL2 | 5,954483122 | 5,824945864 | 3,968242539 | 6,612512106 | 8 |
| ENSGALG00000000433 | | 1272,018653 | 1440,431589 | 928,8310457 | 1560,586316 | 8 |
| ENSGALG00000000921 | | 20,02878103 | 21,81551147 | 15,53568376 | 22,32711039 | 8 |
| ENSGALG00000000966 | | 20,47256091 | 19,52267998 | 15,14294559 | 19,21668586 | 8 |
| ENSGALG00000000970 | | 121,5454735 | 107,7587292 | 88,7857163 | 113,9503321 | 8 |
| ENSGALG00000001026 | | 36,75836285 | 36,46182397 | 30,03854447 | 37,77669558 | 8 |
| ENSGALG00000001040 | | 2,135610314 | 2,701891274 | 1,274424793 | 3,016335863 | 8 |
| ENSGALG00000001096 | | 305,8554674 | 349,0409519 | 253,9138438 | 350,2592113 | 8 |
| ENSGALG00000001144 | ZNF346 | 38,22164066 | 33,87809183 | 30,12408592 | 37,38262219 | 8 |
| ENSGALG00000001191 | MEOX1 | 7,04229087 | 7,660896452 | 4,783886826 | 10,65337693 | 8 |
| ENSGALG00000001342 | HCN2 | 3,566827073 | 3,913196094 | 2,003233469 | 3,567430691 | 8 |
| ENSGALG00000001668 | | 65,31424271 | 61,00669811 | 56,15496715 | 68,17862012 | 8 |
| ENSGALG00000001879 | | 1,337810461 | 2,692005897 | 0,827824211 | 4,113696738 | 8 |
| ENSGALG00000002308 | COMMD10 | 64,65931335 | 63,73727871 | 52,43554159 | 65,08224 | 8 |
| ENSGALG00000002451 | | 7,892336133 | 8,433004452 | 5,368002869 | 9,378353576 | 8 |
| ENSGALG00000002832 | IFI35 | 13,41565324 | 19,92428156 | 5,840906287 | 20,84894169 | 8 |
| ENSGALG00000003282 | STAT5B | 43,47011723 | 59,21868599 | 36,01354452 | 61,71799679 | 8 |
| ENSGALG00000003404 | | 16,38506125 | 22,47425288 | 10,42923248 | 26,49809005 | 8 |
| ENSGALG00000003457 | CNP | 27,91472575 | 24,76733122 | 21,15447525 | 35,31883756 | 8 |
| ENSGALG00000004309 | GDAP1L1 | 24,07708294 | 28,27887319 | 14,31439188 | 29,41749102 | 8 |
| ENSGALG00000004518 | | 5,456576175 | 5,358687351 | 3,636599085 | 5,328686101 | 8 |
| ENSGALG00000004537 | | 180,6757873 | 161,8033161 | 121,9790746 | 175,016988 | 8 |
| ENSGALG00000004725 | | 82,94245202 | 95,93644244 | 69,23109947 | 97,51285509 | 8 |
| ENSGALG00000004741 | EPHB2 | 47,80916319 | 49,45118607 | 31,7185343 | 48,00633611 | 8 |
| ENSGALG00000004775 | | 9,40967762 | 9,784964463 | 6,248181276 | 9,515999548 | 8 |
| ENSGALG00000004875 | PEMT | 36,58111755 | 49,2637599 | 28,18151635 | 49,03587614 | 8 |
| ENSGALG00000004902 | TOM1L2 | 47,90888274 | 54,14283223 | 38,4291727 | 56,34056348 | 8 |
| ENSGALG00000004936 | GLRA4 | 0,79751784 | 0,927900085 | 0,1832244 | 1,510310992 | 8 |
| ENSGALG00000005297 | HHATL | 3,387077303 | 2,668515935 | 1,79964287 | 6,359255378 | 8 |
| ENSGALG00000005330 | SNCAIP | 8,488103316 | 16,17894196 | 6,249057345 | 16,83568789 | 8 |
| ENSGALG00000005628 | COL9A3 | 17,96213617 | 15,94826923 | 10,69613986 | 17,40578323 | 8 |
| ENSGALG00000005751 | | 5,36854481 | 5,781142754 | 3,794119758 | 5,564424762 | 8 |
| ENSGALG00000005820 | RNF130 | 115,9755276 | 132,7019958 | 96,44159114 | 135,6127204 | 8 |
| ENSGALG00000005930 | PLS3 | 207,3820465 | 213,2667153 | 176,3834995 | 205,374161 | 8 |
| ENSGALG00000006152 | | 21,62598292 | 24,37801658 | 13,1869189 | 22,40566092 | 8 |
| ENSGALG00000006245 | | 7,594383946 | 6,44434218 | 3,155446769 | 11,38846586 | 8 |
| ENSGALG00000006346 | CXCL14 | 20,0059794 | 16,52261414 | 9,684803529 | 17,74246896 | 8 |
| ENSGALG00000006453 | | 23,05683828 | 28,21009978 | 15,71140953 | 28,93303717 | 8 |
| ENSGALG00000006480 | TCF7 | 40,96204917 | 40,44824339 | 28,05334558 | 52,25914443 | 8 |
| ENSGALG00000006507 | CLMP | 11,08247747 | 10,29291948 | 5,480177048 | 10,05172185 | 8 |
| ENSGALG00000006702 | MFGE8 | 21,72913025 | 18,25785553 | 16,69202306 | 24,14199294 | 8 |

| | | | | | | |
|---|---|---|---|---|---|---|
| ENSGALG00000006793 | | 4,520154861 | 4,229306859 | 2,448533311 | 5,022349685 | 8 |
| ENSGALG00000006885 | | 193,6654886 | 228,2465968 | 140,0387935 | 217,4857151 | 8 |
| ENSGALG00000007388 | | 3,861549661 | 3,783674214 | 1,646524718 | 3,861815162 | 8 |
| ENSGALG00000007447 | TUBB1 | 7,115252041 | 6,858661343 | 4,858476059 | 6,983463694 | 8 |
| ENSGALG00000007623 | CACNA1G | 9,626950958 | 10,78838576 | 5,504792009 | 10,1006864 | 8 |
| ENSGALG00000007981 | CLN6 | 46,70691407 | 61,89784382 | 34,14706975 | 59,56872405 | 8 |
| ENSGALG00000008188 | TRIM36 | 1,365256301 | 2,320330846 | 0,507058176 | 2,908026625 | 8 |
| ENSGALG00000008250 | | 2,334382918 | 2,222336548 | 1,211757734 | 2,093247638 | 8 |
| ENSGALG00000008349 | LMCD1 | 6,381606237 | 5,950498424 | 3,175987053 | 6,800362349 | 8 |
| ENSGALG00000008367 | MDK | 538,002912 | 538,5537481 | 348,2539918 | 523,7018112 | 8 |
| ENSGALG00000008433 | | 49,98160633 | 63,0842024 | 33,53067368 | 59,10964698 | 8 |
| ENSGALG00000008440 | CHST1 | 6,557087603 | 7,782559043 | 2,575428704 | 8,796610773 | 8 |
| ENSGALG00000008479 | PLEKHM3 | 7,425884037 | 8,009813156 | 4,951758076 | 8,469638599 | 8 |
| ENSGALG00000008888 | | 2,717591914 | 3,726185576 | 1,209465832 | 4,278582904 | 8 |
| ENSGALG00000008953 | AASS | 2,104289207 | 3,485597766 | 1,298976223 | 3,507044234 | 8 |
| ENSGALG00000009060 | | 55,52198203 | 50,99564319 | 43,7247881 | 55,32060319 | 8 |
| ENSGALG00000009237 | | 3,961438611 | 0,772111634 | 1,60979255 | 4,327297542 | 8 |
| ENSGALG00000009638 | ANO6 | 1,715961455 | 1,740638076 | 0,68045043 | 1,500474705 | 8 |
| ENSGALG00000010083 | ST3GAL3 | 50,26691798 | 46,55220626 | 40,90820647 | 47,83758969 | 8 |
| ENSGALG00000010316 | FRAS1 | 14,37701744 | 14,44809891 | 10,14312586 | 16,86422595 | 8 |
| ENSGALG00000010500 | NKX6-2 | 82,02997841 | 72,56241153 | 39,76739828 | 74,26592223 | 8 |
| ENSGALG00000010551 | CDC42EP3 | 32,94479902 | 38,50554976 | 20,60412631 | 41,86968279 | 8 |
| ENSGALG00000010706 | | 15,94984957 | 17,60812199 | 13,24497666 | 20,76705437 | 8 |
| ENSGALG00000010750 | | 90,15991016 | 92,3598817 | 69,55665579 | 93,56553739 | 8 |
| ENSGALG00000010811 | LGMN | 104,6799372 | 108,867442 | 72,91205227 | 103,5052655 | 8 |
| ENSGALG00000011324 | | 645,0588375 | 753,4902301 | 447,1398409 | 797,1195948 | 8 |
| ENSGALG00000011409 | SNRPF | 537,0729519 | 537,2486928 | 446,0636939 | 526,1919388 | 8 |
| ENSGALG00000011951 | | 4348,85487 | 4582,468074 | 3827,371123 | 4750,322822 | 8 |
| ENSGALG00000012165 | DLGAP5 | 180,1309956 | 166,0315001 | 151,8503163 | 181,618057 | 8 |
| ENSGALG00000012546 | MCM5 | 377,5873693 | 349,3150659 | 303,0517423 | 394,3894071 | 8 |
| ENSGALG00000012577 | FRMD3 | 8,135128562 | 10,76858863 | 5,124147526 | 10,54811856 | 8 |
| ENSGALG00000012586 | GKAP1 | 49,38065987 | 48,31048476 | 35,18915075 | 46,61458231 | 8 |
| ENSGALG00000012644 | | 118,2208301 | 107,161874 | 82,58924356 | 124,3352882 | 8 |
| ENSGALG00000012710 | NCS1 | 53,38605119 | 59,47857108 | 35,31604687 | 59,30108349 | 8 |
| ENSGALG00000012712 | RBM24 | 11,69804863 | 12,65506283 | 7,511658153 | 13,38509159 | 8 |
| ENSGALG00000012792 | HIPK2 | 34,10544651 | 47,51606445 | 28,0810563 | 48,57650721 | 8 |
| ENSGALG00000013370 | | 19,96508379 | 17,82198179 | 14,85078035 | 19,24758448 | 8 |
| ENSGALG00000013784 | CDT1 | 165,2005614 | 151,8783838 | 132,5244372 | 174,9558592 | 8 |
| ENSGALG00000013848 | MVK | 80,28802642 | 88,74775882 | 66,89244986 | 85,71101038 | 8 |
| ENSGALG00000014408 | LLGL1 | 136,7318583 | 150,9541375 | 116,2058785 | 148,1708454 | 8 |
| ENSGALG00000014818 | TRIM45 | 13,13116048 | 11,86036513 | 9,402249388 | 13,7018197 | 8 |
| ENSGALG00000014910 | COLEC12 | 36,98106275 | 49,28334328 | 26,80255957 | 46,75071046 | 8 |
| ENSGALG00000014970 | FSTL1 | 400,311328 | 408,6417616 | 314,2104498 | 430,8721221 | 8 |
| ENSGALG00000015082 | RPS6 | 3517,307527 | 3686,176035 | 3027,546214 | 3739,458637 | 8 |
| ENSGALG00000015084 | IGSF11 | 7,264231584 | 9,578005382 | 3,591960257 | 11,29906875 | 8 |
| ENSGALG00000015107 | PIP5K1B | 18,92175169 | 18,57005395 | 13,47044353 | 20,52336847 | 8 |
| ENSGALG00000015186 | | 14,74535991 | 21,55944147 | 7,489188851 | 20,8815943 | 8 |
| ENSGALG00000015326 | CLTA | 267,2978304 | 294,1857064 | 219,3695547 | 329,8482612 | 8 |
| ENSGALG00000015372 | ATP5I | 403,7583558 | 509,6131573 | 333,1327727 | 501,0696761 | 8 |
| ENSGALG00000015379 | TAGLN3 | 87,81368477 | 100,9944499 | 62,06751842 | 113,9094292 | 8 |
| ENSGALG00000015469 | | 20,37996053 | 36,86257283 | 15,57629849 | 36,01578911 | 8 |
| ENSGALG00000015597 | | 5,227904206 | 3,833284117 | 3,037151249 | 4,594564421 | 8 |
| ENSGALG00000015617 | RPS23 | 2617,899457 | 2862,618867 | 2286,280255 | 2782,266287 | 8 |
| ENSGALG00000015663 | HSDL2 | 133,5386111 | 139,1086821 | 101,7937975 | 136,1974338 | 8 |
| ENSGALG00000015729 | LPAR1 | 21,41474649 | 23,35821948 | 13,55389449 | 24,06974178 | 8 |
| ENSGALG00000015895 | HTR1B | 25,76646903 | 23,82650311 | 19,0394431 | 23,92584621 | 8 |
| ENSGALG00000015926 | | 5,567947829 | 7,134824852 | 3,268581878 | 6,673131012 | 8 |
| ENSGALG00000016152 | | 183,9160675 | 192,1318333 | 138,2517808 | 212,5560354 | 8 |
| ENSGALG00000016328 | | 83,28263556 | 147,5069832 | 61,86336813 | 143,313037 | 8 |
| ENSGALG00000016665 | FDFT1 | 96,95097122 | 82,48027441 | 75,2834769 | 99,1418902 | 8 |
| ENSGALG00000016692 | | 85,88865234 | 83,54926437 | 67,04372699 | 83,5908187 | 8 |
| ENSGALG00000021174 | | 57,3355086 | 57,82250149 | 43,91995666 | 55,54606164 | 8 |
| ENSGALG00000021365 | DCTN3 | 212,9718058 | 222,1596474 | 170,2155642 | 239,7819731 | 8 |
| ENSGALG00000022926 | GPR162 | 43,58694006 | 47,89104575 | 31,47238754 | 49,14485034 | 8 |

| | | | | | | |
|---|---|---|---|---|---|---|
| ENSGALG00000023435 | GATM | 10,90557171 | 11,56749101 | 7,424217313 | 11,9559467 | 8 |
| ENSGALG00000023577 | | 3,448939012 | 3,331537676 | 0,899477294 | 6,243389678 | 8 |
| ENSGALG00000023626 | NTN1 | 24,02953412 | 32,21432394 | 16,58916643 | 33,96160684 | 8 |
| ENSGALG00000023689 | ASS1 | 1,249320069 | 1,92940501 | 0,417616289 | 1,950364347 | 8 |
| ENSGALG00000023801 | | 3,651253916 | 2,716726028 | 0,815991185 | 2,845431632 | 8 |
| ENSGALG00000023936 | | 14,3827097 | 16,65315064 | 9,644678815 | 19,86677939 | 8 |
| ENSGALG00000025764 | FKBP10 | 34,79818805 | 41,73571023 | 28,05073896 | 46,92463723 | 8 |
| ENSGALG00000025919 | | 20,97538621 | 16,70085792 | 12,13546828 | 23,20345206 | 8 |
| ENSGALG00000026081 | RBM38 | 44,21346634 | 48,37930983 | 34,01608481 | 53,55787387 | 8 |
| ENSGALG00000026203 | FAM174A | 27,05573691 | 28,32165205 | 20,54054072 | 27,04979678 | 8 |
| ENSGALG00000026291 | | 44,51892603 | 46,84798811 | 36,7820635 | 50,21922775 | 8 |
| ENSGALG00000026300 | | 417,310051 | 435,3420646 | 350,8924467 | 463,9459697 | 8 |
| ENSGALG00000026449 | | 234,0396809 | 311,3452264 | 183,6563344 | 320,9495089 | 8 |
| ENSGALG00000026517 | RFC5 | 156,0885731 | 158,5361803 | 126,8196552 | 169,2035955 | 8 |
| ENSGALG00000026919 | | 2,441650333 | 2,953248286 | 1,625810597 | 2,961202745 | 8 |
| ENSGALG00000027462 | P3H4 | 17,15905149 | 24,46370473 | 12,9024833 | 25,92857528 | 8 |
| ENSGALG00000027501 | ANPEP | 12,01379711 | 10,03763968 | 6,221923555 | 14,47536557 | 8 |
| ENSGALG00000027710 | | 118,024832 | 134,7918283 | 96,52690233 | 134,4618088 | 8 |
| ENSGALG00000027824 | | 76,49717615 | 91,19420938 | 59,38902312 | 87,936403 | 8 |
| ENSGALG00000027917 | | 29,58120555 | 28,64651933 | 24,83745695 | 28,98249309 | 8 |
| ENSGALG00000027963 | COX7C | 177,2982167 | 215,1096727 | 146,7488079 | 208,9018198 | 8 |
| ENSGALG00000028470 | | 29,51653543 | 32,02852181 | 23,47533065 | 32,21145656 | 8 |
| ENSGALG00000028543 | FGFR4 | 15,68663076 | 14,79137651 | 9,453274085 | 15,46542694 | 8 |
| ENSGALG00000028682 | CBLN2 | 14,71375594 | 13,14190177 | 4,554452616 | 12,43147608 | 8 |
| ENSGALG00000028716 | | 67,55341979 | 61,61072026 | 54,87200105 | 66,7873307 | 8 |
| ENSGALG00000028795 | | 18,82719941 | 21,71314579 | 12,37729026 | 28,90667954 | 8 |
| ENSGALG00000029019 | | 0,071552757 | 0,02569526 | 0,013204207 | 0,059434845 | 8 |
| ENSGALG00000029174 | | 27,89925869 | 54,48511253 | 20,36078384 | 65,69049819 | 8 |
| ENSGALG00000000044 | WFIKKN1 | 31,25400692 | 17,05051776 | 18,3375194 | 16,23160749 | 9 |
| ENSGALG00000000161 | ISOC1 | 46,60655276 | 39,59532572 | 35,69279471 | 37,31603953 | 9 |
| ENSGALG00000000528 | | 57,8074608 | 48,44024498 | 44,75009669 | 37,2413949 | 9 |
| ENSGALG00000000587 | | 10,87385763 | 7,033016405 | 7,125083511 | 6,717576626 | 9 |
| ENSGALG00000000619 | ANGPTL4 | 19,0867957 | 11,9712835 | 14,11823462 | 10,50099385 | 9 |
| ENSGALG00000000623 | | 123,5225802 | 85,31962828 | 80,7882465 | 78,48784073 | 9 |
| ENSGALG00000000628 | | 30,5489741 | 20,87801046 | 22,15847088 | 20,43363063 | 9 |
| ENSGALG00000000644 | | 27,34191038 | 15,24672665 | 17,77134235 | 13,4679886 | 9 |
| ENSGALG00000000681 | PAK1 | 19,8427259 | 17,46535245 | 16,20668324 | 17,09279966 | 9 |
| ENSGALG00000000684 | KLHDC8A | 52,21088274 | 37,62714869 | 41,16059659 | 28,61108362 | 9 |
| ENSGALG00000000690 | GDPD4 | 108,6457182 | 52,11130181 | 64,04614044 | 41,17074748 | 9 |
| ENSGALG00000000750 | | 5,480945592 | 4,028561748 | 3,354812383 | 4,175923743 | 9 |
| ENSGALG00000000780 | | 100,949447 | 93,14132877 | 82,50393208 | 63,461697 | 9 |
| ENSGALG00000000927 | ANGPTL2 | 12,96678898 | 9,307183466 | 8,306381085 | 6,636659284 | 9 |
| ENSGALG00000000939 | | 88,7332569 | 65,62779965 | 64,03714459 | 58,49544371 | 9 |
| ENSGALG00000001063 | | 9,161939454 | 6,091395269 | 4,295525218 | 4,422690334 | 9 |
| ENSGALG00000001141 | HES5 | 1418,60207 | 1112,884661 | 1038,241962 | 898,373404 | 9 |
| ENSGALG00000001153 | AUTS2 | 115,680856 | 92,03951073 | 73,84336585 | 81,27424337 | 9 |
| ENSGALG00000001516 | | 36,66443788 | 25,78277742 | 22,91121743 | 22,30922256 | 9 |
| ENSGALG00000001733 | LRIG2 | 69,99915275 | 58,54048669 | 55,02885328 | 46,10602716 | 9 |
| ENSGALG00000001774 | NEO1 | 144,6407555 | 114,9123423 | 104,6053004 | 85,78682404 | 9 |
| ENSGALG00000001954 | APTX | 80,10762546 | 60,11975832 | 63,03128633 | 55,10270093 | 9 |
| ENSGALG00000002090 | GPSM2 | 133,1906718 | 117,2282647 | 113,4776413 | 103,8677502 | 9 |
| ENSGALG00000002091 | ZDHHC1 | 26,3461381 | 22,81614683 | 19,80544615 | 19,55804929 | 9 |
| ENSGALG00000002228 | | 44,10956358 | 26,8456593 | 23,99308478 | 24,50200802 | 9 |
| ENSGALG00000002249 | AGO1 | 128,6081271 | 93,61556828 | 98,265346 | 85,67338208 | 9 |
| ENSGALG00000002268 | | 57,88933765 | 48,13784859 | 48,5939597 | 43,31254971 | 9 |
| ENSGALG00000002298 | COL8A2 | 33,36976189 | 19,29568543 | 15,36725962 | 21,08774073 | 9 |
| ENSGALG00000002317 | | 28,69380943 | 24,97653503 | 23,05559118 | 21,56051721 | 9 |
| ENSGALG00000002375 | NOTCH1 | 137,3911419 | 112,1394882 | 91,48126487 | 99,84782244 | 9 |
| ENSGALG00000002599 | ARHGEF39 | 41,36660965 | 30,08437158 | 31,67263412 | 32,09717563 | 9 |
| ENSGALG00000002652 | | 215,5577691 | 158,3864206 | 138,1374797 | 98,78094971 | 9 |
| ENSGALG00000002774 | TSPAN3 | 599,9764973 | 447,3646823 | 467,922637 | 392,2362777 | 9 |
| ENSGALG00000002831 | SH3PXD2B | 43,1122237 | 31,76322262 | 32,27401658 | 27,94680373 | 9 |
| ENSGALG00000002974 | SPHKAP | 6,235148986 | 3,427073538 | 3,961823218 | 2,770629109 | 9 |
| ENSGALG00000003085 | CDK1 | 835,642788 | 711,6170086 | 717,3170313 | 734,33598 | 9 |

| | | | | | | |
|---|---|---|---|---|---|---|
| ENSGALG00000003163 | GNAO1 | 161,9917225 | 108,4121059 | 105,2394616 | 85,1330615 | 9 |
| ENSGALG00000003320 | | 2,681105339 | 1,925580202 | 1,577834384 | 1,719761201 | 9 |
| ENSGALG00000003424 | LMX1A | 14,62572453 | 6,771825817 | 7,698712962 | 5,993144908 | 9 |
| ENSGALG00000003426 | RHBDL3 | 84,52214462 | 44,61134271 | 44,5094291 | 29,9201131 | 9 |
| ENSGALG00000003429 | PBX1 | 90,75741527 | 72,39295828 | 72,36913039 | 65,41501567 | 9 |
| ENSGALG00000003434 | C17orf75 | 116,8784541 | 77,66935733 | 68,15217644 | 55,35753847 | 9 |
| ENSGALG00000003503 | MYBL2 | 103,3422234 | 89,91141222 | 86,47418819 | 88,49615621 | 9 |
| ENSGALG00000003551 | | 16,08587485 | 9,180770702 | 7,561269976 | 10,02134429 | 9 |
| ENSGALG00000003554 | OPTC | 119,1239593 | 71,84161518 | 56,89099392 | 54,86320698 | 9 |
| ENSGALG00000003582 | SLC1A3 | 84,64090633 | 32,60614203 | 53,95299351 | 22,44357991 | 9 |
| ENSGALG00000003601 | ATP2B4 | 84,10436696 | 68,83661031 | 57,81129784 | 65,5936423 | 9 |
| ENSGALG00000003763 | | 30,39802539 | 26,39791988 | 26,01635594 | 25,84006884 | 9 |
| ENSGALG00000003782 | | 67,52962601 | 48,8949534 | 48,05084689 | 44,1293765 | 9 |
| ENSGALG00000003837 | ADCY7 | 11,25806886 | 7,141289587 | 6,436046755 | 7,162858002 | 9 |
| ENSGALG00000003930 | APBA2 | 56,79764343 | 48,00506316 | 44,99572286 | 37,56595504 | 9 |
| ENSGALG00000003987 | TRAF4 | 24,93184967 | 18,42870213 | 17,6207474 | 15,76983074 | 9 |
| ENSGALG00000004028 | MATN4 | 4,142818748 | 1,759184805 | 2,092418693 | 1,787010196 | 9 |
| ENSGALG00000004256 | | 117,1187154 | 94,6556023 | 78,78379073 | 61,2475351 | 9 |
| ENSGALG00000004331 | | 78,43473809 | 47,01093113 | 44,17985984 | 35,90771284 | 9 |
| ENSGALG00000004338 | | 184,3647772 | 126,1585532 | 128,6882618 | 115,000824 | 9 |
| ENSGALG00000004364 | ABR | 63,93113798 | 46,56275273 | 54,95450035 | 38,63387769 | 9 |
| ENSGALG00000004400 | | 22,88103183 | 17,29827581 | 15,39979111 | 11,65059965 | 9 |
| ENSGALG00000004464 | | 29,2120941 | 22,14419799 | 23,17263832 | 21,18993522 | 9 |
| ENSGALG00000004516 | | 159,5821658 | 133,8086625 | 134,3461022 | 133,3681191 | 9 |
| ENSGALG00000004530 | MMD2 | 36,22728323 | 25,45444714 | 16,61274281 | 19,35254048 | 9 |
| ENSGALG00000004621 | PREX1 | 57,87358886 | 41,24143534 | 46,0542225 | 33,12573231 | 9 |
| ENSGALG00000004790 | | 654,3246722 | 352,1800754 | 269,3322898 | 224,3252022 | 9 |
| ENSGALG00000004954 | | 335,5094457 | 269,4578124 | 262,8551051 | 262,2329529 | 9 |
| ENSGALG00000004986 | CERCAM | 289,2108064 | 153,4223911 | 163,1756543 | 124,3823144 | 9 |
| ENSGALG00000005214 | | 22,97422669 | 17,61003498 | 17,43784342 | 14,70240257 | 9 |
| ENSGALG00000005237 | FGD3 | 100,5472145 | 86,36549191 | 75,01190805 | 60,70387368 | 9 |
| ENSGALG00000005246 | | 90,58806014 | 51,35115641 | 62,64847331 | 38,43466117 | 9 |
| ENSGALG00000005319 | CDH8 | 5,445262477 | 4,173315769 | 3,017728175 | 1,866804765 | 9 |
| ENSGALG00000005403 | DHRS11 | 99,94667899 | 73,04572469 | 64,1697335 | 58,53117191 | 9 |
| ENSGALG00000005405 | | 31,49344217 | 26,42825252 | 25,35628462 | 25,1695081 | 9 |
| ENSGALG00000005413 | | 17,63752439 | 7,770628589 | 5,025165513 | 4,739852004 | 9 |
| ENSGALG00000005426 | FREM1 | 4,04595993 | 2,468827971 | 2,431482695 | 2,135553458 | 9 |
| ENSGALG00000005680 | | 9,640144343 | 7,265188978 | 6,687105253 | 7,850074723 | 9 |
| ENSGALG00000005718 | BHLHE23 | 7,660886537 | 4,527062152 | 3,141076562 | 4,207532061 | 9 |
| ENSGALG00000005739 | SCD | 120,233915 | 71,49058858 | 95,28129954 | 82,94998739 | 9 |
| ENSGALG00000005774 | | 107,6754727 | 75,61866044 | 66,91043187 | 46,31621048 | 9 |
| ENSGALG00000005806 | KIF24 | 14,08523354 | 11,47064874 | 9,930660996 | 9,661822978 | 9 |
| ENSGALG00000005817 | | 40,19436852 | 33,50907622 | 33,14220262 | 32,53433765 | 9 |
| ENSGALG00000005892 | | 231,8060061 | 154,3941218 | 131,8842244 | 122,7009004 | 9 |
| ENSGALG00000005903 | NRIP3 | 13,90935106 | 8,083012884 | 7,461017706 | 7,936189903 | 9 |
| ENSGALG00000005956 | | 2,030478214 | 0,939634647 | 0,819765053 | 0,774947504 | 9 |
| ENSGALG00000005961 | dAK | 237,4982121 | 209,651884 | 186,032781 | 188,3048286 | 9 |
| ENSGALG00000005975 | | 25,36720101 | 9,858054738 | 7,549317126 | 8,015243217 | 9 |
| ENSGALG00000005989 | | 67,30399299 | 61,24008237 | 57,66738697 | 52,15578407 | 9 |
| ENSGALG00000006037 | MCM2 | 331,6274576 | 265,1735327 | 287,4206462 | 284,4425533 | 9 |
| ENSGALG00000006094 | PHF6 | 205,6581455 | 163,1742698 | 166,1087468 | 124,9981669 | 9 |
| ENSGALG00000006158 | ACVR2B | 187,5109516 | 161,5663473 | 141,2383772 | 119,6603309 | 9 |
| ENSGALG00000006210 | | 681,9171879 | 337,8310196 | 285,5834543 | 265,8640383 | 9 |
| ENSGALG00000006219 | PRKAG2 | 116,4614015 | 99,89346909 | 100,664404 | 99,31929353 | 9 |
| ENSGALG00000006235 | | 25,72391513 | 18,24632753 | 12,9330952 | 11,31001119 | 9 |
| ENSGALG00000006267 | TPX2 | 285,3239263 | 234,4828814 | 242,4073029 | 244,9347288 | 9 |
| ENSGALG00000006409 | | 98,96573337 | 59,31551837 | 64,86590911 | 48,98443263 | 9 |
| ENSGALG00000006483 | | 66,59212784 | 47,861138 | 53,79593876 | 41,7959226 | 9 |
| ENSGALG00000006488 | RYK | 123,9224556 | 101,6109651 | 102,7321424 | 85,2358062 | 9 |
| ENSGALG00000006555 | | 36,25083342 | 15,25441221 | 21,94626456 | 13,42875792 | 9 |
| ENSGALG00000006563 | SEMA3D | 64,15874594 | 42,35039981 | 31,42366476 | 24,29139394 | 9 |
| ENSGALG00000006623 | ATP11C | 84,40040171 | 75,82040472 | 72,31961566 | 76,48580465 | 9 |
| ENSGALG00000006634 | | 312,6569359 | 230,7046187 | 244,5534232 | 217,9314324 | 9 |
| ENSGALG00000006666 | BPIFB2 | 12,13866269 | 7,437423024 | 6,49942857 | 7,202762558 | 9 |

| | | | | | | |
|---|---|---|---|---|---|---|
| ENSGALG00000006826 | KLHL25 | 110,0451345 | 65,01099472 | 69,38138855 | 48,75260528 | 9 |
| ENSGALG00000006859 | | 8,765240974 | 7,31641822 | 6,12271801 | 5,389653833 | 9 |
| ENSGALG00000007245 | | 27,97164142 | 18,0114389 | 18,55377799 | 13,50467657 | 9 |
| ENSGALG00000007254 | ARHGAP12 | 45,44941843 | 33,57720882 | 36,21730402 | 30,37036253 | 9 |
| ENSGALG00000007269 | GABRA3 | 13,96411172 | 10,86879485 | 9,501500127 | 8,542584012 | 9 |
| ENSGALG00000007304 | | 34,23812047 | 29,61794147 | 25,721381 | 25,45594431 | 9 |
| ENSGALG00000007419 | MASP1 | 1,57539922 | 0,938984074 | 0,512908118 | 0,711741724 | 9 |
| ENSGALG00000007668 | BMP7 | 76,36611621 | 53,6951202 | 61,40584973 | 54,13905456 | 9 |
| ENSGALG00000007710 | | 99,30828406 | 71,94323354 | 71,27438268 | 46,25151534 | 9 |
| ENSGALG00000007781 | GAL3ST1 | 14,13111353 | 9,3883292 | 5,772312125 | 6,986217646 | 9 |
| ENSGALG00000007892 | SPAG6 | 61,11626497 | 47,97350969 | 39,2464153 | 36,4836125 | 9 |
| ENSGALG00000007929 | DIXDC1 | 68,58037619 | 48,0580845 | 38,09909643 | 29,97386629 | 9 |
| ENSGALG00000008174 | | 124,348004 | 107,8743784 | 104,2337788 | 99,6087797 | 9 |
| ENSGALG00000008215 | | 50,38257469 | 31,52731813 | 20,60181027 | 23,6834746 | 9 |
| ENSGALG00000008279 | | 577,5801838 | 394,4296542 | 272,5352455 | 249,8305959 | 9 |
| ENSGALG00000008425 | | 14,69561107 | 12,47619676 | 11,11587652 | 9,119602956 | 9 |
| ENSGALG00000008525 | ABHD12 | 51,3904353 | 43,30549983 | 40,64464425 | 38,93553863 | 9 |
| ENSGALG00000008616 | | 156,5243897 | 126,1365769 | 136,68892 | 104,2885426 | 9 |
| ENSGALG00000008698 | MGME1 | 71,41290823 | 58,06460253 | 56,79294211 | 53,48784783 | 9 |
| ENSGALG00000008758 | | 124,4228967 | 82,10905734 | 98,0079813 | 61,64666093 | 9 |
| ENSGALG00000008871 | | 80,84342079 | 47,41819894 | 54,93095143 | 39,863161 | 9 |
| ENSGALG00000008873 | POLE3 | 321,2902245 | 262,7556375 | 240,5475354 | 218,3913542 | 9 |
| ENSGALG00000008887 | | 33,84974642 | 27,03979324 | 26,04572531 | 22,59057103 | 9 |
| ENSGALG00000008896 | | 6,022737193 | 3,499565077 | 2,398429886 | 2,115469582 | 9 |
| ENSGALG00000008961 | | 138,0057818 | 67,99395964 | 89,6981597 | 52,304204 | 9 |
| ENSGALG00000009020 | JAG1 | 124,7023905 | 75,41664337 | 74,13724728 | 55,83821036 | 9 |
| ENSGALG00000009027 | ST6GALNAC3 | 29,42036569 | 25,27116058 | 20,59626824 | 18,22933473 | 9 |
| ENSGALG00000009207 | TRIM2 | 30,67711925 | 25,11356027 | 21,42653595 | 15,16568337 | 9 |
| ENSGALG00000009279 | SFMBT2 | 10,91446674 | 6,537015591 | 7,691053349 | 7,838415429 | 9 |
| ENSGALG00000009474 | | 165,3825058 | 136,4697521 | 134,2634588 | 138,1460328 | 9 |
| ENSGALG00000009514 | NRCAM | 42,45191179 | 20,00427649 | 23,8798494 | 14,18976528 | 9 |
| ENSGALG00000009516 | | 41,86000548 | 32,23669772 | 30,81092781 | 28,62779004 | 9 |
| ENSGALG00000009619 | | 40,1118097 | 32,22391492 | 31,93240207 | 32,85425266 | 9 |
| ENSGALG00000009626 | THBS1 | 45,34052685 | 25,28042099 | 27,54267112 | 21,88710977 | 9 |
| ENSGALG00000009732 | PCDH18 | 41,0860473 | 31,86811586 | 23,36024215 | 21,9012931 | 9 |
| ENSGALG00000010097 | GATB | 32,46219309 | 27,21579359 | 23,9674918 | 19,30756619 | 9 |
| ENSGALG00000010170 | INTU | 36,14029054 | 26,89626511 | 28,52273715 | 23,97967786 | 9 |
| ENSGALG00000010312 | PLCH1 | 44,07417619 | 31,12405937 | 36,06839209 | 29,68185375 | 9 |
| ENSGALG00000010346 | TGFB3 | 93,98664535 | 76,02551444 | 69,58015827 | 79,88767103 | 9 |
| ENSGALG00000010448 | LTBP1 | 245,7170762 | 174,8131199 | 213,6709569 | 130,4004299 | 9 |
| ENSGALG00000010598 | KCNK10 | 7,855388034 | 3,90410236 | 4,553758606 | 2,460899286 | 9 |
| ENSGALG00000010674 | DCTD | 192,9283112 | 144,6806116 | 152,1865547 | 117,8680635 | 9 |
| ENSGALG00000010700 | RPA3 | 260,5499153 | 214,8768156 | 223,3044233 | 200,2168515 | 9 |
| ENSGALG00000010705 | ZBTB18 | 44,10376849 | 24,51748535 | 31,58377038 | 25,0157285 | 9 |
| ENSGALG00000010775 | | 12,70190269 | 9,002694914 | 7,808452185 | 6,029336459 | 9 |
| ENSGALG00000010798 | DHCR24 | 54,13550003 | 46,53369275 | 42,79643706 | 47,16973778 | 9 |
| ENSGALG00000010818 | PLPP3 | 69,29865779 | 48,32497394 | 49,3012161 | 41,06142891 | 9 |
| ENSGALG00000010825 | AGR2 | 2,225899698 | 0,437850605 | 0,260687102 | 0,445933444 | 9 |
| ENSGALG00000010829 | | 167,2465489 | 136,4200703 | 138,2033727 | 116,3988713 | 9 |
| ENSGALG00000010835 | DTX4 | 1195,686573 | 840,6890347 | 831,069909 | 723,909037 | 9 |
| ENSGALG00000010903 | | 30,51325584 | 19,6495363 | 18,42950404 | 14,34456426 | 9 |
| ENSGALG00000010960 | GALNT3 | 20,3476324 | 13,46078214 | 12,92983377 | 9,283859309 | 9 |
| ENSGALG00000010971 | CSRNP3 | 17,1320569 | 10,94668327 | 12,37192759 | 9,463981496 | 9 |
| ENSGALG00000011109 | MAPK10 | 10,6059035 | 8,201546462 | 6,224593472 | 6,370787412 | 9 |
| ENSGALG00000011182 | DLL1 | 115,6739078 | 83,57962434 | 66,45209234 | 63,60645287 | 9 |
| ENSGALG00000011206 | KITLG | 22,29167397 | 16,26019056 | 12,14019114 | 12,29957305 | 9 |
| ENSGALG00000011571 | AGPAT4 | 93,67204267 | 68,94234947 | 78,24764951 | 60,3321412 | 9 |
| ENSGALG00000011823 | FAT4 | 25,5685505 | 20,27150072 | 15,35641303 | 12,56335526 | 9 |
| ENSGALG00000012017 | DAAM1 | 66,51638573 | 51,6956635 | 46,97005806 | 42,50632291 | 9 |
| ENSGALG00000012121 | RCN1 | 128,2812544 | 85,34230795 | 104,339948 | 73,50530684 | 9 |
| ENSGALG00000012123 | PAX6 | 207,3310704 | 125,6192176 | 94,53567349 | 104,5564319 | 9 |
| ENSGALG00000012124 | PELI2 | 97,01672853 | 72,38390338 | 68,07241714 | 72,99638735 | 9 |
| ENSGALG00000012130 | DNAJC24 | 87,42750265 | 67,82813268 | 67,26018354 | 54,6134636 | 9 |
| ENSGALG00000012184 | | 17,53434364 | 10,51607147 | 5,571243627 | 4,897927265 | 9 |

| | | | | | | |
|---|---|---|---|---|---|---|
| ENSGALG00000012216 | BMPR1B | 19,73470606 | 17,41660439 | 14,638607 | 11,03804069 | 9 |
| ENSGALG00000012226 | | 10,14170208 | 6,091780586 | 5,829669854 | 4,876167191 | 9 |
| ENSGALG00000012332 | | 1,75499313 | 0,334938857 | 0,049686386 | 0,405453413 | 9 |
| ENSGALG00000012361 | NIN | 38,54121987 | 30,6965853 | 28,85722382 | 29,36252216 | 9 |
| ENSGALG00000012388 | PCMT1 | 114,9803287 | 97,79226505 | 96,7908776 | 85,73985935 | 9 |
| ENSGALG00000012395 | PLEKHG1 | 40,97672463 | 34,45473513 | 31,88510956 | 26,43661125 | 9 |
| ENSGALG00000012440 | | 126,942135 | 81,54303435 | 74,69736259 | 83,67449226 | 9 |
| ENSGALG00000012595 | AGTPBP1 | 5,843425877 | 3,87973109 | 3,244881839 | 3,714060964 | 9 |
| ENSGALG00000012618 | FANCC | 77,15501466 | 61,95447317 | 63,86958208 | 59,5116733 | 9 |
| ENSGALG00000012620 | PTCH1 | 128,0187265 | 79,01327975 | 78,98591949 | 63,93344831 | 9 |
| ENSGALG00000012662 | NUAK1 | 65,54806121 | 48,25369742 | 54,49515428 | 35,50923797 | 9 |
| ENSGALG00000012684 | DEK | 213,1310822 | 191,1534846 | 182,5897587 | 191,0382858 | 9 |
| ENSGALG00000012697 | SLC41A2 | 0,058029502 | 0,037541976 | 0,025719414 | 0,034391684 | 9 |
| ENSGALG00000012703 | DTNBP1 | 65,11808287 | 60,06352135 | 53,570554 | 45,8855677 | 9 |
| ENSGALG00000012756 | PARPBP | 8,741901058 | 7,493838313 | 6,914878548 | 5,358083966 | 9 |
| ENSGALG00000012775 | TFAP2A | 8,004715347 | 5,508538021 | 5,589140758 | 6,133219794 | 9 |
| ENSGALG00000012936 | | 43,2525541 | 33,16536089 | 32,41777008 | 26,8771372 | 9 |
| ENSGALG00000013155 | | 60,96549085 | 50,10471915 | 41,405435 | 39,39397896 | 9 |
| ENSGALG00000013211 | Sial-T2 | 53,06870305 | 39,2868121 | 36,22341479 | 29,76988665 | 9 |
| ENSGALG00000013212 | | 352,0737733 | 218,2231586 | 304,4999677 | 177,1428711 | 9 |
| ENSGALG00000013357 | | 14,5238095 | 9,676299999 | 9,468806965 | 11,33051766 | 9 |
| ENSGALG00000013672 | TSHZ1 | 47,63040303 | 40,01248352 | 37,12051956 | 40,35485231 | 9 |
| ENSGALG00000013722 | SEMA3B | 67,72409615 | 45,67603936 | 47,22283227 | 43,11405926 | 9 |
| ENSGALG00000013754 | | 99,90845036 | 74,32231981 | 73,38471802 | 63,38796719 | 9 |
| ENSGALG00000013756 | | 9,551795017 | 8,012150023 | 7,152967635 | 6,59881996 | 9 |
| ENSGALG00000013782 | CDH7 | 44,73753961 | 24,83144713 | 33,13366661 | 19,36298515 | 9 |
| ENSGALG00000013900 | MAP7 | 18,05284523 | 9,552077542 | 7,566128142 | 7,59934912 | 9 |
| ENSGALG00000013961 | | 56,10973329 | 47,74639425 | 42,56173827 | 38,2782928 | 9 |
| ENSGALG00000014069 | TP53I11 | 462,0420259 | 303,2892015 | 318,9636775 | 276,2729485 | 9 |
| ENSGALG00000014100 | | 7,763915083 | 3,510610015 | 2,846977556 | 4,067062932 | 9 |
| ENSGALG00000014218 | PHF21B | 44,41267673 | 32,88106133 | 33,30902396 | 23,77376394 | 9 |
| ENSGALG00000014284 | | 25,24429229 | 20,35052461 | 20,85799387 | 23,81657043 | 9 |
| ENSGALG00000014347 | | 21,79724499 | 18,61697194 | 16,21325842 | 13,48321788 | 9 |
| ENSGALG00000014485 | LDB2 | 58,65859245 | 44,03980556 | 38,92048977 | 32,99839391 | 9 |
| ENSGALG00000014508 | CD38 | 34,52009233 | 22,68968774 | 20,56010187 | 19,01534557 | 9 |
| ENSGALG00000014620 | | 1,956048892 | 0,782889319 | 0,862399115 | 0,580997363 | 9 |
| ENSGALG00000014646 | SYT11 | 171,4109919 | 151,5937547 | 140,9231322 | 134,8023289 | 9 |
| ENSGALG00000014672 | ARSK | 14,70009358 | 11,78884508 | 10,76404571 | 9,602709952 | 9 |
| ENSGALG00000014680 | | 24,0520188 | 19,83747893 | 17,85066188 | 16,91394554 | 9 |
| ENSGALG00000014753 | CENPK | 106,4544342 | 83,51630756 | 81,07856229 | 90,52213786 | 9 |
| ENSGALG00000014756 | PPWD1 | 129,0762799 | 102,4153578 | 107,5831259 | 108,0635267 | 9 |
| ENSGALG00000014829 | | 15,02253293 | 10,43374439 | 6,545915045 | 6,828450455 | 9 |
| ENSGALG00000014897 | TYMS | 276,8568161 | 246,1181255 | 233,2358706 | 234,7946898 | 9 |
| ENSGALG00000014903 | ITGA2 | 19,28102901 | 12,04416315 | 14,06402156 | 10,26187687 | 9 |
| ENSGALG00000014934 | HS3ST1 | 5,540861535 | 3,210938472 | 2,425738011 | 1,67621552 | 9 |
| ENSGALG00000015053 | GLDC | 65,20466566 | 54,00282438 | 55,14865591 | 47,938418 | 9 |
| ENSGALG00000015091 | HAUS6 | 107,8740638 | 77,47610692 | 90,80409523 | 80,42181649 | 9 |
| ENSGALG00000015103 | CCDC171 | 9,278746716 | 7,442349244 | 6,81158066 | 5,688912777 | 9 |
| ENSGALG00000015152 | BOC | 61,37451521 | 50,47041324 | 41,88963775 | 37,19576843 | 9 |
| ENSGALG00000015241 | | 34,52994734 | 20,74355677 | 21,80562309 | 14,0162192 | 9 |
| ENSGALG00000015386 | RNF170 | 48,11695973 | 40,96561413 | 36,04033923 | 29,04558532 | 9 |
| ENSGALG00000015444 | | 17,77233078 | 14,64411238 | 13,67274692 | 12,36829449 | 9 |
| ENSGALG00000015448 | | 26,99122737 | 19,26894575 | 15,38003926 | 10,64663709 | 9 |
| ENSGALG00000015476 | | 14,64646726 | 7,865669198 | 7,720149158 | 5,783165827 | 9 |
| ENSGALG00000015564 | | 7,156970347 | 5,334313058 | 5,500962035 | 4,919475757 | 9 |
| ENSGALG00000015673 | | 19,31944035 | 13,00690578 | 11,37988935 | 9,833987567 | 9 |
| ENSGALG00000015685 | CFAP99 | 9,786223813 | 7,029241391 | 6,950135982 | 6,216729289 | 9 |
| ENSGALG00000015712 | SLBP | 102,5760884 | 74,18453744 | 83,0184383 | 76,31708087 | 9 |
| ENSGALG00000015786 | REEP1 | 53,13915006 | 42,50403724 | 36,81544162 | 36,11332154 | 9 |
| ENSGALG00000015799 | ELAVL2 | 6,037654055 | 4,325382874 | 3,767789255 | 2,819033678 | 9 |
| ENSGALG00000015847 | PRSS35 | 35,11704576 | 25,66370192 | 24,07123267 | 19,02952064 | 9 |
| ENSGALG00000016034 | ATRN | 111,1123874 | 81,26651019 | 87,63945569 | 67,82246842 | 9 |
| ENSGALG00000016148 | C2CD2 | 39,27360457 | 32,23641902 | 29,98843351 | 28,60296797 | 9 |
| ENSGALG00000016171 | | 218,2787932 | 205,7354159 | 192,6760184 | 172,0707128 | 9 |

| | | | | | | |
|---|---|---|---|---|---|---|
| ENSGALG00000016200 | | 51,81097754 | 39,08952357 | 37,81925937 | 29,69260203 | 9 |
| ENSGALG00000016244 | | 34,80907173 | 29,60541455 | 24,53665949 | 19,37430659 | 9 |
| ENSGALG00000016326 | | 10,59842146 | 8,398086434 | 7,852987771 | 6,850555245 | 9 |
| ENSGALG00000016391 | CNKSR2 | 26,29502855 | 16,46476384 | 16,96231718 | 11,5736257 | 9 |
| ENSGALG00000016394 | | 32,19475368 | 17,39524143 | 21,9320163 | 17,49313991 | 9 |
| ENSGALG00000016417 | | 131,3461383 | 103,1768005 | 104,7836855 | 88,93333367 | 9 |
| ENSGALG00000016428 | | 98,90427495 | 50,27843955 | 39,10462421 | 33,95468388 | 9 |
| ENSGALG00000016442 | RRM2 | 593,9920997 | 471,2620483 | 469,5176719 | 463,1466191 | 9 |
| ENSGALG00000016456 | Lpin1 | 11,86024943 | 9,523941198 | 9,270708107 | 9,810098096 | 9 |
| ENSGALG00000016459 | FAM84A | 7,093676028 | 4,179695873 | 3,107376993 | 3,325425099 | 9 |
| ENSGALG00000016487 | LDAH | 39,21408017 | 36,90700532 | 33,03840175 | 29,58276228 | 9 |
| ENSGALG00000016497 | UBXN2A | 68,16553674 | 58,34933795 | 51,49336847 | 45,51619727 | 9 |
| ENSGALG00000016575 | GPM6B | 267,7110691 | 174,3280247 | 163,2139649 | 132,8150489 | 9 |
| ENSGALG00000016576 | OFD1 | 80,96250322 | 62,01808879 | 54,92287394 | 47,92994163 | 9 |
| ENSGALG00000016676 | MCM3 | 220,3958455 | 163,3016763 | 186,0877349 | 184,8337011 | 9 |
| ENSGALG00000016810 | | 34,46474494 | 22,03432788 | 16,29129687 | 17,02120699 | 9 |
| ENSGALG00000016811 | | 20,87242426 | 16,56591395 | 15,39201879 | 12,9699234 | 9 |
| ENSGALG00000017021 | ATP7B | 3,9869324 | 2,371265345 | 1,829014636 | 1,212619235 | 9 |
| ENSGALG00000017154 | ZC3H12C | 16,69008154 | 11,1294025 | 10,93196073 | 9,942631937 | 9 |
| ENSGALG00000017393 | | 14,97893571 | 10,68802307 | 10,58952889 | 9,672886802 | 9 |
| ENSGALG00000019987 | LRP11 | 101,4977697 | 84,74657711 | 74,00932276 | 62,64430136 | 9 |
| ENSGALG00000020003 | MAP3K4 | 101,7199951 | 71,21199104 | 84,45899897 | 57,14632163 | 9 |
| ENSGALG00000020554 | | 2,745534391 | 1,931406303 | 1,752178514 | 1,782013487 | 9 |
| ENSGALG00000020788 | | 16,08950004 | 7,37692355 | 7,655663237 | 3,861511503 | 9 |
| ENSGALG00000020813 | | 171,290155 | 148,1329936 | 144,4510861 | 138,3803558 | 9 |
| ENSGALG00000021031 | ACKR3 | 197,7454394 | 96,43341863 | 106,3679352 | 85,09027757 | 9 |
| ENSGALG00000021340 | CA9 | 246,544707 | 203,2768771 | 190,5602386 | 192,1909568 | 9 |
| ENSGALG00000021592 | | 24,90805272 | 18,51353834 | 18,06631656 | 13,3222017 | 9 |
| ENSGALG00000021593 | | 21,6290003 | 14,87215611 | 11,48507743 | 16,43767163 | 9 |
| ENSGALG00000021616 | | 660,7892 | 560,4597192 | 492,802757 | 474,9610111 | 9 |
| ENSGALG00000022603 | | 127,7440137 | 94,25649465 | 98,2989799 | 83,9412793 | 9 |
| ENSGALG00000023114 | TAL2 | 10,83090997 | 4,516618379 | 3,527657045 | 4,165674735 | 9 |
| ENSGALG00000023359 | BTBD17 | 12,37917296 | 9,215387972 | 6,210062578 | 6,211710478 | 9 |
| ENSGALG00000023546 | | 103,3250361 | 82,98091285 | 85,32961236 | 77,35742371 | 9 |
| ENSGALG00000023647 | | 16,32978445 | 10,43731524 | 9,865902217 | 8,46856396 | 9 |
| ENSGALG00000023898 | | 25,15757129 | 20,28376564 | 21,16857225 | 21,71796482 | 9 |
| ENSGALG00000024047 | MYCL | 130,6449284 | 91,27348025 | 100,1887482 | 78,95088543 | 9 |
| ENSGALG00000024120 | | 251,9186613 | 186,3133705 | 184,633668 | 156,1919821 | 9 |
| ENSGALG00000025796 | ADI1 | 151,9056229 | 128,8288136 | 130,0669627 | 115,3487407 | 9 |
| ENSGALG00000026085 | | 37,55971233 | 29,79205273 | 28,08976508 | 26,0061611 | 9 |
| ENSGALG00000026179 | FAM163B | 12,70392319 | 7,406176596 | 6,530189162 | 10,54638186 | 9 |
| ENSGALG00000026192 | ST8SIA4 | 26,69578621 | 14,31843914 | 14,38997567 | 10,57310427 | 9 |
| ENSGALG00000026214 | LRTM2 | 5,889632729 | 3,621162228 | 2,979514896 | 2,858887818 | 9 |
| ENSGALG00000026313 | RND3 | 54,52252904 | 42,60080306 | 41,54066922 | 34,55755523 | 9 |
| ENSGALG00000026395 | TRABD2A | 34,96858896 | 22,03852277 | 14,4142307 | 13,7208985 | 9 |
| ENSGALG00000026452 | MURC | 68,5514329 | 42,65798561 | 57,55999236 | 40,22180647 | 9 |
| ENSGALG00000026741 | | 92,52230306 | 78,97977067 | 79,76273285 | 68,87406566 | 9 |
| ENSGALG00000026757 | | 115,1492891 | 83,10995002 | 78,42339249 | 85,39515762 | 9 |
| ENSGALG00000026858 | BTG2 | 73,25524576 | 56,978272 | 43,18762707 | 46,83714731 | 9 |
| ENSGALG00000026981 | NHSL1 | 90,0052068 | 79,25555686 | 72,87488074 | 70,3460249 | 9 |
| ENSGALG00000027002 | | 335,0560184 | 189,0165812 | 223,361849 | 151,3843853 | 9 |
| ENSGALG00000027183 | | 19,96308058 | 12,60167171 | 12,49826777 | 10,81410515 | 9 |
| ENSGALG00000027198 | TMEM47 | 68,18436645 | 46,36338174 | 41,15361629 | 45,81100202 | 9 |
| ENSGALG00000027240 | TSPAN13 | 60,13229624 | 43,7309013 | 38,32508629 | 35,23013388 | 9 |
| ENSGALG00000027373 | | 255,2616537 | 165,0099254 | 173,1189979 | 141,526405 | 9 |
| ENSGALG00000027472 | TMEM38B | 18,53265539 | 16,23438527 | 14,16492945 | 12,41744795 | 9 |
| ENSGALG00000027484 | C20orf85 | 5,62155177 | 3,44560581 | 2,174506144 | 2,913151077 | 9 |
| ENSGALG00000027514 | | 0,528787008 | 0,166761415 | 0,080304322 | 0,124803265 | 9 |
| ENSGALG00000027633 | RASSF9 | 2,035731325 | 0,854912883 | 0,737308999 | 0,339184915 | 9 |
| ENSGALG00000027821 | | 23,11507984 | 14,67663083 | 10,50775245 | 14,49954578 | 9 |
| ENSGALG00000027897 | DTX1 | 44,84876496 | 37,26651436 | 33,21350155 | 34,29440557 | 9 |
| ENSGALG00000028043 | | 35,35893431 | 28,05893819 | 22,40021823 | 20,30301901 | 9 |
| ENSGALG00000028140 | CNTFR | 21,29176694 | 15,68480074 | 14,7877547 | 16,3046985 | 9 |
| ENSGALG00000028327 | TMEM178A | 15,25643824 | 12,99141162 | 11,17599999 | 10,775708 | 9 |

| | | | | | | |
|---|---|---|---|---|---|---|
| ENSGALG00000028415 | | 51,81555289 | 16,42028113 | 20,4065691 | 12,36370773 | 9 |
| ENSGALG00000028536 | MPEG1 | 20,5427871 | 13,37260379 | 15,08320974 | 11,52346016 | 9 |
| ENSGALG00000028790 | DNASE2B | 19,09091612 | 11,85162585 | 9,263964302 | 10,09521056 | 9 |
| ENSGALG00000029004 | | 0,626344243 | 0,267590302 | 0,357524634 | 0,25546319 | 9 |
| ENSGALG00000029011 | SLC35F1 | 147,2321386 | 101,3673948 | 105,6335088 | 79,54888119 | 9 |
| ENSGALG00000029057 | | 6,612061759 | 3,565809227 | 3,192304914 | 3,978423806 | 9 |
| ENSGALG00000029070 | FAM181A | 140,8743623 | 93,20761779 | 88,5842806 | 63,71077724 | 9 |
| ENSGALG00000029072 | NTN3 | 23,04277493 | 13,47398348 | 14,06728438 | 13,23933001 | 9 |
| ENSGALG00000029106 | | 11,86577158 | 8,120777874 | 6,029059075 | 6,275995647 | 9 |
| ENSGALG00000029140 | CRABP2 | 943,46397 | 516,5320484 | 569,9989128 | 473,9399887 | 9 |
| ENSGALG00000029151 | | 3,237130342 | 2,159102861 | 1,570247908 | 2,377856659 | 9 |
| ENSGALG00000000067 | SPR | 25,19151507 | 38,09496599 | 32,36154561 | 46,01001796 | 10 |
| ENSGALG00000000243 | RDM1 | 64,16894169 | 84,57366491 | 88,19253189 | 87,07892618 | 10 |
| ENSGALG00000000505 | KCNH6 | 1,806346197 | 3,772203834 | 3,349604234 | 3,427118714 | 10 |
| ENSGALG00000000836 | RFX5 | 0,220047592 | 0,798370098 | 0,387008719 | 0,842809634 | 10 |
| ENSGALG00000000894 | APCDD1 | 12,17953658 | 20,37298423 | 18,67259013 | 21,14941456 | 10 |
| ENSGALG00000001376 | ARID3B | 81,77846269 | 128,5884065 | 111,6210537 | 158,6263766 | 10 |
| ENSGALG00000001664 | PTPN9 | 27,32342552 | 45,33442697 | 41,5208593 | 49,78702526 | 10 |
| ENSGALG00000002054 | | 4,463691977 | 10,11551737 | 8,122956552 | 9,768543779 | 10 |
| ENSGALG00000002210 | PPIB | 615,7965084 | 812,398235 | 771,3773995 | 889,5451731 | 10 |
| ENSGALG00000002342 | | 76,97404676 | 104,3309613 | 95,54319676 | 118,6337645 | 10 |
| ENSGALG00000002922 | NOTCH2 | 34,87569574 | 45,95827585 | 44,26184738 | 50,20078048 | 10 |
| ENSGALG00000002969 | MAU2 | 41,46144033 | 47,17099845 | 48,73910042 | 47,72736011 | 10 |
| ENSGALG00000003011 | TOM1L1 | 0,671138223 | 2,313862165 | 2,230320103 | 3,177873963 | 10 |
| ENSGALG00000003019 | | 7,412203933 | 9,138713853 | 9,422655346 | 10,02764078 | 10 |
| ENSGALG00000003340 | GBGT1 | 1,301060604 | 4,566529257 | 2,549542908 | 4,535141421 | 10 |
| ENSGALG00000003562 | NPTX2 | 1,276374613 | 2,747257273 | 2,562428679 | 3,583310899 | 10 |
| ENSGALG00000003640 | | 9,84244551 | 40,12648844 | 16,73702806 | 37,56346879 | 10 |
| ENSGALG00000003841 | DGKK | 1,170538651 | 1,987308984 | 2,249687426 | 2,15756725 | 10 |
| ENSGALG00000004083 | GRIA1 | 6,30072857 | 18,26990907 | 15,82341279 | 17,5097179 | 10 |
| ENSGALG00000004719 | SPTAN1 | 93,79037656 | 111,3665527 | 109,9098282 | 108,9079703 | 10 |
| ENSGALG00000004740 | TMOD2 | 7,782204442 | 14,88555351 | 10,95510486 | 12,75531378 | 10 |
| ENSGALG00000004805 | | 5,914997516 | 7,271558567 | 7,670585037 | 7,762114467 | 10 |
| ENSGALG00000004909 | | 59,70087412 | 104,0709532 | 86,70697475 | 99,82170433 | 10 |
| ENSGALG00000005065 | PLA2G4A | 4,27041438 | 8,286182082 | 7,268353555 | 8,503620832 | 10 |
| ENSGALG00000005389 | | 0,141856332 | 3,826903443 | 1,782989285 | 2,582117723 | 10 |
| ENSGALG00000005861 | | 2,00701187 | 3,862514989 | 3,827339633 | 5,164782157 | 10 |
| ENSGALG00000005906 | | 25,34069165 | 46,7681257 | 40,66967147 | 52,53393838 | 10 |
| ENSGALG00000006313 | | 7,796058512 | 11,54622893 | 10,90957967 | 13,40926496 | 10 |
| ENSGALG00000006641 | | 15,64772345 | 21,61228415 | 20,72433764 | 23,81644591 | 10 |
| ENSGALG00000006659 | | 40,08388889 | 51,59649186 | 52,50217807 | 58,09914445 | 10 |
| ENSGALG00000006729 | | 14,16368852 | 150,8222642 | 64,1838708 | 77,30692972 | 10 |
| ENSGALG00000007412 | MPZL2 | 1,426673901 | 2,603616958 | 3,116406355 | 3,436684581 | 10 |
| ENSGALG00000007543 | APCDD1L | 1,285779838 | 10,76373217 | 10,1632961 | 15,43303975 | 10 |
| ENSGALG00000007565 | PPP2R2B | 4,498606815 | 16,55729926 | 7,013796757 | 13,48424013 | 10 |
| ENSGALG00000007833 | | 22,46417563 | 30,2369186 | 29,28165289 | 32,02014786 | 10 |
| ENSGALG00000008009 | BCAP29 | 54,09129251 | 74,79823611 | 73,83395516 | 82,24575148 | 10 |
| ENSGALG00000008181 | MADD | 15,2172411 | 21,6887672 | 21,73612765 | 25,35030935 | 10 |
| ENSGALG00000008999 | AFAP1L2 | 5,746909958 | 16,02416497 | 14,60767657 | 23,10791907 | 10 |
| ENSGALG00000009155 | | 6,468639196 | 11,71015107 | 12,0614327 | 12,97982335 | 10 |
| ENSGALG00000009433 | BAG3 | 43,32330391 | 89,33785815 | 65,96819578 | 99,83819404 | 10 |
| ENSGALG00000009505 | PNPLA8 | 33,57778621 | 40,98407666 | 40,30681413 | 42,47879678 | 10 |
| ENSGALG00000009570 | ATP6V1D | 74,62845941 | 128,6859866 | 102,1283936 | 130,2908971 | 10 |
| ENSGALG00000009718 | FAM53B | 5,787080902 | 16,4568785 | 11,11024098 | 15,08477821 | 10 |
| ENSGALG00000009759 | SLC16A7 | 0,371037879 | 0,893895773 | 0,86989287 | 1,026405516 | 10 |
| ENSGALG00000009783 | | 10,2405462 | 14,28065595 | 13,0313556 | 14,17793631 | 10 |
| ENSGALG00000009792 | | 10,84341484 | 21,73021356 | 16,86750659 | 17,88482058 | 10 |
| ENSGALG00000009896 | TRERF1 | 4,808170267 | 13,5104085 | 7,397743277 | 15,27344774 | 10 |
| ENSGALG00000009972 | | 8,233254639 | 14,77085833 | 10,96112463 | 12,48984776 | 10 |
| ENSGALG00000009988 | ELOVL1 | 24,54970046 | 40,25627092 | 37,95280868 | 40,63830581 | 10 |
| ENSGALG00000010164 | SMARCA2 | 31,48210827 | 48,72221196 | 46,7844494 | 57,39705613 | 10 |
| ENSGALG00000010268 | | 4,921325317 | 13,01196961 | 15,00161538 | 16,21986662 | 10 |
| ENSGALG00000010664 | KIF26B | 0,871077545 | 3,300649183 | 1,910395118 | 3,87997435 | 10 |
| ENSGALG00000010911 | | 1,617219387 | 3,748364438 | 2,982354432 | 4,754746898 | 10 |

| | | | | | | |
|---|---|---|---|---|---|---|
| ENSGALG00000010934 | | 1,8031112 | 3,709899605 | 2,91645905 | 4,653025085 | 10 |
| ENSGALG00000011016 | PGM1 | 23,44574688 | 32,53423526 | 32,62243154 | 31,85192134 | 10 |
| ENSGALG00000011286 | UBE2E1 | 62,79994395 | 81,23871675 | 90,03965196 | 96,89006504 | 10 |
| ENSGALG00000011488 | CMTM7 | 7,277427951 | 20,1323349 | 11,74053362 | 16,23963969 | 10 |
| ENSGALG00000011690 | | 0,103946776 | 0,939637581 | 0,88149688 | 0,834198832 | 10 |
| ENSGALG00000011972 | | 21,41005163 | 26,662867 | 28,93241264 | 30,75647977 | 10 |
| ENSGALG00000012093 | | 12,1904691 | 23,2396876 | 16,99874812 | 22,36987224 | 10 |
| ENSGALG00000012118 | TMEM260 | 10,53321393 | 13,22765331 | 14,21926447 | 15,08071395 | 10 |
| ENSGALG00000012211 | PDLIM5 | 16,67664874 | 23,83760349 | 27,59486714 | 28,08174448 | 10 |
| ENSGALG00000012280 | | 32,6377987 | 47,09813408 | 43,51845385 | 46,57456552 | 10 |
| ENSGALG00000012304 | NFKB1 | 18,92353309 | 36,72187679 | 23,71285028 | 29,81253681 | 10 |
| ENSGALG00000012345 | UST | 25,98919869 | 34,81550817 | 36,11724943 | 34,83622704 | 10 |
| ENSGALG00000012877 | CREB3L2 | 14,15122188 | 29,46359767 | 22,89406436 | 24,70796898 | 10 |
| ENSGALG00000013738 | OPTN | 31,20424125 | 63,0962501 | 64,58752569 | 83,85842847 | 10 |
| ENSGALG00000013830 | PRELID3A | 39,10486792 | 55,02903372 | 52,71471236 | 54,50673512 | 10 |
| ENSGALG00000013902 | | 1,573316302 | 4,401902139 | 4,571135381 | 4,45045502 | 10 |
| ENSGALG00000014014 | | 0,595599345 | 2,558484407 | 0,964881493 | 2,547986889 | 10 |
| ENSGALG00000014106 | PPFIBP1 | 48,33798078 | 79,49747287 | 78,82503111 | 87,90269502 | 10 |
| ENSGALG00000014933 | HEXB | 13,82875151 | 27,24707297 | 21,86614475 | 28,18919125 | 10 |
| ENSGALG00000015250 | | 6,860711489 | 17,44011316 | 12,3963785 | 12,88410587 | 10 |
| ENSGALG00000015424 | | 48,88941571 | 72,42774461 | 65,23032901 | 67,1628436 | 10 |
| ENSGALG00000015459 | | 19,21668801 | 24,21055023 | 25,90194667 | 25,85794579 | 10 |
| ENSGALG00000015652 | MFSD10 | 12,11072038 | 16,15790442 | 16,48891498 | 16,37173519 | 10 |
| ENSGALG00000015677 | SUSD1 | 5,974385986 | 10,6464791 | 9,370624852 | 13,62756519 | 10 |
| ENSGALG00000015770 | APP | 380,9874972 | 492,5151558 | 458,3835411 | 482,5133678 | 10 |
| ENSGALG00000015792 | | 8,931724434 | 14,36302132 | 13,26278114 | 14,78442869 | 10 |
| ENSGALG00000015958 | EIF2AK3 | 9,629444865 | 17,72798233 | 12,0730299 | 16,39705179 | 10 |
| ENSGALG00000016050 | TTC3 | 21,38230929 | 25,43226191 | 26,64498514 | 26,08190222 | 10 |
| ENSGALG00000016069 | | 14,59709436 | 23,86183631 | 25,39502668 | 24,64718079 | 10 |
| ENSGALG00000016090 | | 2,529020993 | 11,65505479 | 5,834035244 | 6,766647026 | 10 |
| ENSGALG00000016311 | | 9,237809102 | 13,39156196 | 14,74306104 | 15,77492644 | 10 |
| ENSGALG00000016341 | | 0,202104856 | 0,525241112 | 0,549551156 | 0,636899768 | 10 |
| ENSGALG00000017199 | MAML2 | 16,24202136 | 24,17136146 | 25,99411058 | 26,45082516 | 10 |
| ENSGALG00000017379 | | 1,368912243 | 3,891543067 | 3,862137749 | 5,074133424 | 10 |
| ENSGALG00000017414 | JUP | 51,88278313 | 71,84240343 | 75,80087103 | 85,42821 | 10 |
| ENSGALG00000017652 | TTC9 | 0,468001686 | 2,538977862 | 2,85619406 | 2,815917147 | 10 |
| ENSGALG00000018634 | | 3,028664762 | 6,076484682 | 4,960409416 | 5,873704945 | 10 |
| ENSGALG00000019835 | | 5,591831465 | 7,516668319 | 8,481902626 | 8,879886504 | 10 |
| ENSGALG00000023541 | SPHK1 | 22,91275764 | 31,05187359 | 30,31393316 | 31,95169083 | 10 |
| ENSGALG00000023806 | | 73,79335109 | 95,32276617 | 100,6008133 | 101,5648268 | 10 |
| ENSGALG00000023886 | ZNF804A | 1,685790356 | 4,57926496 | 3,192521495 | 4,573801224 | 10 |
| ENSGALG00000025780 | ITPRIP | 5,385578786 | 9,094153341 | 10,15330177 | 10,84817331 | 10 |
| ENSGALG00000026184 | | 20,45088044 | 35,45119107 | 28,32020272 | 36,40521622 | 10 |
| ENSGALG00000026694 | | 1,351457644 | 3,212622506 | 2,533295638 | 2,645072591 | 10 |
| ENSGALG00000026727 | SLC30A10 | 1,391821025 | 2,828916703 | 2,502986912 | 3,564811485 | 10 |
| ENSGALG00000026822 | WDR81 | 18,46040207 | 24,87090381 | 24,50072815 | 25,59442237 | 10 |
| ENSGALG00000027764 | | 2,512489522 | 9,103387834 | 10,99845138 | 12,01146352 | 10 |
| ENSGALG00000027772 | | 0,004892914 | 0,62961338 | 0,353233983 | 0,624609016 | 10 |
| ENSGALG00000028155 | | 5,782098605 | 11,47490998 | 10,05991248 | 10,8361757 | 10 |
| ENSGALG00000028354 | METRNL | 37,35106289 | 58,93445961 | 46,00048458 | 56,99942589 | 10 |
| ENSGALG00000028582 | | 3,241190702 | 10,61160058 | 7,55174664 | 14,2492626 | 10 |
| ENSGALG00000000107 | TRIM7 | 0,223879993 | 3,047486999 | 1,757372561 | 5,176952685 | 11 |
| ENSGALG00000000117 | | 0,040480999 | 0,220063553 | 0,353819011 | 0,526729322 | 11 |
| ENSGALG00000000464 | | 34,05612158 | 42,48913859 | 44,95313416 | 70,7967403 | 11 |
| ENSGALG00000000544 | RAB44 | 0,061671392 | 0,998421135 | 3,862756333 | 17,67716506 | 11 |
| ENSGALG00000000935 | | 5,314625544 | 7,277338245 | 8,425064177 | 13,7096056 | 11 |
| ENSGALG00000001163 | NDEL1 | 8,521796698 | 10,78416631 | 12,97721211 | 15,40550268 | 11 |
| ENSGALG00000001189 | NCF1 | 1,08189946 | 3,436075078 | 4,646876547 | 9,23946829 | 11 |
| ENSGALG00000001281 | | 0,846894805 | 2,258112365 | 2,370684136 | 5,949678488 | 11 |
| ENSGALG00000001617 | CAMKK1 | 2,604019637 | 7,289324417 | 5,664646749 | 25,44035569 | 11 |
| ENSGALG00000001728 | NACC2 | 11,80186684 | 15,10233978 | 18,75610144 | 27,14939763 | 11 |
| ENSGALG00000002221 | | 17,07683677 | 20,30245478 | 22,63798635 | 24,90324994 | 11 |
| ENSGALG00000002260 | CISH | 0,663424147 | 2,139171593 | 2,487720205 | 17,431701 | 11 |
| ENSGALG00000002548 | TLN1 | 20,31627411 | 22,61612195 | 25,81496882 | 28,39885397 | 11 |

| | | | | | | |
|---|---|---|---|---|---|---|
| ENSGALG00000002583 | PIK3CD | 2,842773013 | 3,679276554 | 4,250538255 | 5,394793108 | 11 |
| ENSGALG00000002651 | ST3GAL2 | 40,21612852 | 57,64615583 | 53,02907343 | 92,8557766 | 11 |
| ENSGALG00000003122 | ZFAND2A | 38,01972748 | 57,81426075 | 57,94506614 | 99,08508814 | 11 |
| ENSGALG00000003439 | PLEKHM2 | 48,97856485 | 55,11774842 | 59,44696304 | 66,65072501 | 11 |
| ENSGALG00000003939 | | 0,73347476 | 1,054393959 | 2,139119382 | 3,112879397 | 11 |
| ENSGALG00000004599 | | 14,05222685 | 16,58556716 | 26,26019986 | 36,29605918 | 11 |
| ENSGALG00000004626 | | 160,6674727 | 192,0383383 | 196,9247106 | 241,3004908 | 11 |
| ENSGALG00000004735 | | 18,71576319 | 24,95209333 | 27,715552 | 47,74079099 | 11 |
| ENSGALG00000005069 | | 0,925868522 | 2,967476206 | 3,749577092 | 6,294077837 | 11 |
| ENSGALG00000005422 | PDLIM1 | 22,92658872 | 27,45113237 | 32,72226862 | 44,12324064 | 11 |
| ENSGALG00000006008 | HOMER2 | 9,984368442 | 15,75188365 | 20,62165034 | 40,94770081 | 11 |
| ENSGALG00000006407 | | 11,54087345 | 29,36740869 | 20,30551587 | 48,42948482 | 11 |
| ENSGALG00000006614 | | 0,081869495 | 0,459517562 | 0,360325202 | 3,561928827 | 11 |
| ENSGALG00000006655 | MAP3K13 | 5,192744917 | 7,000137701 | 8,021460066 | 11,36909123 | 11 |
| ENSGALG00000006947 | BDH1 | 134,1451069 | 150,8578862 | 185,9764126 | 304,1197319 | 11 |
| ENSGALG00000006958 | COL27A1 | 1,237521755 | 2,205820699 | 2,92438058 | 3,742640145 | 11 |
| ENSGALG00000006988 | | 0,188436541 | 0,498050832 | 0,566069098 | 1,241061957 | 11 |
| ENSGALG00000007238 | | 7,877525855 | 11,00497278 | 12,15849698 | 17,65303471 | 11 |
| ENSGALG00000007253 | IL1RAP | 5,654414984 | 5,558750136 | 15,57106007 | 34,03095536 | 11 |
| ENSGALG00000007271 | | 14,84086276 | 18,48294427 | 21,14237742 | 25,377985 | 11 |
| ENSGALG00000007394 | | 8,540622935 | 12,18654051 | 12,89609102 | 16,54507416 | 11 |
| ENSGALG00000007415 | | 0,625774296 | 1,462296833 | 4,567836451 | 9,979634759 | 11 |
| ENSGALG00000007700 | CTTN | 128,2039185 | 140,4859116 | 150,5105378 | 160,6028241 | 11 |
| ENSGALG00000008143 | | 54,04131097 | 43,71598201 | 70,22117139 | 96,62123273 | 11 |
| ENSGALG00000008461 | | 0,10085251 | 0,358449616 | 0,976354066 | 2,087420033 | 11 |
| ENSGALG00000008855 | UGP2 | 111,7329891 | 128,4121345 | 150,4721612 | 177,2730406 | 11 |
| ENSGALG00000008860 | LONRF3 | 13,08961567 | 11,56252738 | 17,86767603 | 27,59267882 | 11 |
| ENSGALG00000008914 | NRAP | 0,060907983 | 0,269803827 | 0,554252687 | 1,06803981 | 11 |
| ENSGALG00000009298 | TNIK | 63,46441582 | 78,28507302 | 87,33775935 | 95,12356324 | 11 |
| ENSGALG00000009755 | LRIG3 | 8,560302311 | 9,689196018 | 13,21297799 | 16,14258758 | 11 |
| ENSGALG00000011054 | | 0,288321762 | 0,262980056 | 0,523082511 | 0,94837143 | 11 |
| ENSGALG00000011435 | | 39,02276939 | 40,19212106 | 61,11491938 | 81,52693241 | 11 |
| ENSGALG00000011489 | | 14,63826567 | 23,22066191 | 21,51808684 | 52,80909388 | 11 |
| ENSGALG00000011682 | CNOT4 | 22,61054622 | 27,70663704 | 31,04588956 | 34,4550159 | 11 |
| ENSGALG00000012425 | FERMT2 | 83,37649219 | 100,1967554 | 100,4088724 | 114,1270352 | 11 |
| ENSGALG00000012429 | BMP4 | 17,82855398 | 36,14532957 | 28,25726137 | 99,3060206 | 11 |
| ENSGALG00000012799 | RREB1 | 11,30126957 | 13,32117937 | 15,67245479 | 20,56566068 | 11 |
| ENSGALG00000012956 | TMTC1 | 3,209890424 | 8,381486759 | 5,824287986 | 16,89837709 | 11 |
| ENSGALG00000013971 | SGK1 | 1,199458409 | 4,40482548 | 2,542307669 | 10,62932801 | 11 |
| ENSGALG00000014261 | | 242,5207982 | 377,9510577 | 322,2554339 | 780,0694354 | 11 |
| ENSGALG00000015351 | CBLB | 12,07404883 | 15,15682028 | 17,17091245 | 27,17633895 | 11 |
| ENSGALG00000015764 | | 44,0304829 | 84,94006302 | 66,45816223 | 168,7490346 | 11 |
| ENSGALG00000015842 | TIAM1 | 20,70041796 | 28,78174708 | 32,87000523 | 38,20472121 | 11 |
| ENSGALG00000015952 | | 4,032607318 | 5,861909098 | 8,43091945 | 13,8938431 | 11 |
| ENSGALG00000016143 | | 17,59022168 | 32,08840457 | 37,51357006 | 60,99757206 | 11 |
| ENSGALG00000016245 | BCOR | 28,64786295 | 33,51699852 | 36,2266639 | 41,65368829 | 11 |
| ENSGALG00000016608 | ADCY3 | 2,853041335 | 6,651742728 | 4,591714727 | 10,19414025 | 11 |
| ENSGALG00000016728 | RP2 | 25,23490471 | 36,91489398 | 30,38401635 | 48,54519915 | 11 |
| ENSGALG00000016782 | | 0,10442844 | 0,304036925 | 0,846006738 | 3,678282371 | 11 |
| ENSGALG00000017053 | SPG20 | 11,56761385 | 17,96862498 | 16,31923789 | 24,44057475 | 11 |
| ENSGALG00000023768 | | 12,58827437 | 14,79960637 | 29,53884984 | 46,46901449 | 11 |
| ENSGALG00000023876 | | 18,91142705 | 31,12230827 | 30,65086759 | 49,13801227 | 11 |
| ENSGALG00000023950 | | 16,76930744 | 21,1204594 | 24,25813505 | 28,37848226 | 11 |
| ENSGALG00000024094 | UBAP2L | 121,4816481 | 122,1153283 | 148,8144652 | 174,3988709 | 11 |
| ENSGALG00000024346 | | 7,457246685 | 8,304577678 | 12,23979195 | 18,63106951 | 11 |
| ENSGALG00000024357 | | 4,028752212 | 4,219547662 | 5,072418923 | 7,277278899 | 11 |
| ENSGALG00000026998 | | 3,550274189 | 4,733865265 | 5,160014923 | 6,176494911 | 11 |
| ENSGALG00000027168 | | 0,136502676 | 0,267739429 | 0,637942204 | 1,071922405 | 11 |
| ENSGALG00000027218 | | 0,155738515 | 0,513268647 | 2,446272203 | 7,015905066 | 11 |
| ENSGALG00000027275 | | 0,679644543 | 3,953449831 | 4,492768067 | 7,153248591 | 11 |
| ENSGALG00000028130 | | 1,446421878 | 5,232635436 | 4,495461427 | 10,72481048 | 11 |
| ENSGALG00000028318 | CDKN1A | 11,85233268 | 16,60453562 | 17,57147394 | 39,65416547 | 11 |
| ENSGALG00000028326 | HSD11B2 | 17,52552612 | 21,01950351 | 22,5837827 | 24,73250896 | 11 |
| ENSGALG00000028478 | | 8,846493717 | 10,6899465 | 14,16438457 | 22,92265049 | 11 |

| | | | | | | |
|---|---|---|---|---|---|---|
| ENSGALG00000028908 | HOXA6 | 0,080342382 | 0,199969978 | 0,088329259 | 0,326037306 | 11 |
| ENSGALG00000029104 | gga-mir-1732-3 | 0 | 0 | 0 | 0,104480823 | 11 |
| ENSGALG00000000146 | SUPT6H | 75,73757331 | 73,2737215 | 93,33629554 | 93,02462799 | 12 |
| ENSGALG00000000156 | BRD2 | 395,5412624 | 375,8987072 | 462,9625348 | 448,7455005 | 12 |
| ENSGALG00000000239 | | 0,126231267 | 0,270426788 | 1,459421001 | 1,749971744 | 12 |
| ENSGALG00000000363 | | 17,72172408 | 18,08275976 | 21,91750525 | 24,64111121 | 12 |
| ENSGALG00000000452 | | 13,88409826 | 12,9025315 | 23,92675951 | 31,4506522 | 12 |
| ENSGALG00000000669 | ETV3 | 55,2844349 | 51,98736662 | 82,87935161 | 85,15029012 | 12 |
| ENSGALG00000001076 | LIMK1 | 27,37079683 | 33,09902324 | 41,69368553 | 48,09803052 | 12 |
| ENSGALG00000001108 | | 36,7691568 | 37,71543648 | 46,04875359 | 44,15642446 | 12 |
| ENSGALG00000001160 | | 15,70235161 | 23,82939143 | 39,42507425 | 37,6126867 | 12 |
| ENSGALG00000001188 | | 26,63368303 | 16,27228527 | 87,3108226 | 78,12265485 | 12 |
| ENSGALG00000001258 | NUMA1 | 75,21228323 | 80,47740842 | 88,36224786 | 93,51829499 | 12 |
| ENSGALG00000001434 | STOM | 1,111021049 | 1,193655921 | 3,310272534 | 3,967409139 | 12 |
| ENSGALG00000001439 | RANBP10 | 20,85370319 | 20,3325829 | 24,978668 | 24,63196551 | 12 |
| ENSGALG00000001629 | | 0,1320548 | 0,348616407 | 1,058011551 | 1,157701708 | 12 |
| ENSGALG00000001786 | | 0,066749272 | 0,014283645 | 0,159044053 | 0,214445459 | 12 |
| ENSGALG00000002203 | FGF-18 | 1,02113805 | 2,989854627 | 7,968073498 | 11,18804202 | 12 |
| ENSGALG00000002219 | WBP2 | 14,30140436 | 15,18900359 | 20,5204241 | 18,87317075 | 12 |
| ENSGALG00000002672 | | 12,23172351 | 13,31637203 | 17,11505231 | 17,08473791 | 12 |
| ENSGALG00000002710 | IPMK | 25,59409512 | 25,81557827 | 33,58394728 | 37,18293113 | 12 |
| ENSGALG00000002892 | | 3,768927028 | 3,881473842 | 6,678895746 | 8,225997381 | 12 |
| ENSGALG00000002941 | | 31,89980035 | 29,96014663 | 38,7163801 | 38,24721579 | 12 |
| ENSGALG00000002955 | ADAMTSL2 | 7,228745086 | 6,663532384 | 13,21950783 | 14,13514694 | 12 |
| ENSGALG00000003329 | GORAB | 29,21089981 | 30,59281993 | 35,78325922 | 36,27201232 | 12 |
| ENSGALG00000003669 | | 30,87085295 | 31,84817216 | 37,92728777 | 40,73316328 | 12 |
| ENSGALG00000003770 | ANXA2 | 34,30091985 | 36,59000096 | 49,56068963 | 57,6372842 | 12 |
| ENSGALG00000003774 | ATP13A2 | 32,84399915 | 32,52396046 | 56,0962193 | 53,02322702 | 12 |
| ENSGALG00000003862 | | 14,85029412 | 13,21434675 | 20,79018931 | 19,72952996 | 12 |
| ENSGALG00000003977 | TWF2 | 111,4566955 | 113,279942 | 130,8659209 | 133,7761085 | 12 |
| ENSGALG00000004002 | LYPLA2 | 150,7495569 | 166,7823137 | 199,4046703 | 212,3326077 | 12 |
| ENSGALG00000004189 | MAFK | 18,5742469 | 15,59354155 | 38,4494806 | 41,1022777 | 12 |
| ENSGALG00000004209 | ELFN1 | 0,814582164 | 1,927779755 | 2,943360603 | 2,951185081 | 12 |
| ENSGALG00000004365 | WDR66 | 2,811763915 | 2,218322715 | 7,695524749 | 7,52136835 | 12 |
| ENSGALG00000004620 | LAMC1 | 67,46199036 | 66,61106568 | 80,89251769 | 77,47594626 | 12 |
| ENSGALG00000004644 | | 29,84829422 | 33,04333611 | 40,59987383 | 41,34144336 | 12 |
| ENSGALG00000004853 | LRP3 | 14,18967878 | 14,79468174 | 18,33567747 | 17,50060833 | 12 |
| ENSGALG00000005027 | | 72,94509595 | 75,84276563 | 86,87569122 | 95,21946926 | 12 |
| ENSGALG00000005095 | SLC25A25 | 35,12452075 | 31,73824572 | 50,94724862 | 51,41349297 | 12 |
| ENSGALG00000005176 | | 3,413908368 | 6,654325261 | 18,68400718 | 17,79864417 | 12 |
| ENSGALG00000005181 | SEC24C | 112,2572831 | 109,0976774 | 135,2911855 | 130,1713029 | 12 |
| ENSGALG00000005249 | BICD2 | 41,4406151 | 47,18645604 | 56,32728041 | 59,8048339 | 12 |
| ENSGALG00000005263 | SOX8 | 9,020400549 | 10,58055905 | 34,04858434 | 38,05051596 | 12 |
| ENSGALG00000005397 | GGNBP2 | 38,09740839 | 43,89356976 | 47,73002156 | 49,9842953 | 12 |
| ENSGALG00000005410 | | 30,14761432 | 40,13307191 | 57,20488775 | 54,56007134 | 12 |
| ENSGALG00000005423 | GRHPR | 33,87507342 | 41,46612067 | 57,59587439 | 60,8918674 | 12 |
| ENSGALG00000005475 | | 42,00791977 | 40,17065565 | 51,63447746 | 53,70763465 | 12 |
| ENSGALG00000005505 | PIK3CB | 5,618150407 | 6,581557383 | 7,805316892 | 8,557807703 | 12 |
| ENSGALG00000005624 | | 40,16275662 | 42,1965956 | 48,86600313 | 48,24907824 | 12 |
| ENSGALG00000005768 | ETV5 | 17,84320142 | 19,27243279 | 32,56938156 | 36,72354266 | 12 |
| ENSGALG00000006310 | | 2,754882711 | 5,047741943 | 6,407070844 | 6,761563236 | 12 |
| ENSGALG00000006340 | | 7,323548685 | 5,786364119 | 11,16780943 | 11,8700516 | 12 |
| ENSGALG00000006372 | RBM33 | 60,01467349 | 56,67563861 | 72,27843028 | 72,37077428 | 12 |
| ENSGALG00000006447 | | 12,1905467 | 16,08270401 | 60,23850836 | 48,57012952 | 12 |
| ENSGALG00000006757 | XYLT1 | 21,69948906 | 21,78358564 | 28,28419731 | 28,13276454 | 12 |
| ENSGALG00000006855 | MELTF | 7,135006571 | 6,352915053 | 18,88027596 | 17,81802431 | 12 |
| ENSGALG00000006863 | | 42,12487657 | 42,61257462 | 56,69086954 | 58,62079625 | 12 |
| ENSGALG00000006919 | POF1B | 10,72105321 | 9,803658308 | 14,95587342 | 15,84499082 | 12 |
| ENSGALG00000007016 | FAM169B | 16,76630825 | 19,67869242 | 24,92697293 | 23,82466955 | 12 |
| ENSGALG00000007144 | SIK3 | 2,474106812 | 3,27210838 | 3,763083294 | 4,031320651 | 12 |
| ENSGALG00000007416 | CD3E | 0,796692068 | 0,963660773 | 2,18018488 | 1,892526906 | 12 |
| ENSGALG00000007512 | | 62,47306751 | 60,26951594 | 76,45241873 | 83,74261589 | 12 |
| ENSGALG00000007514 | | 40,05564722 | 43,21170517 | 53,37200909 | 50,85259172 | 12 |
| ENSGALG00000007530 | | 2,185558313 | 3,093638992 | 7,22955912 | 8,390199431 | 12 |

| | | | | | | |
|---|---|---|---|---|---|---|
| ENSGALG00000007669 | EGR1 | 20,96798972 | 15,10254827 | 112,2799795 | 93,85732019 | 12 |
| ENSGALG00000008099 | | 38,73018399 | 38,99872595 | 49,856415 | 56,42754101 | 12 |
| ENSGALG00000008315 | | 48,98731667 | 49,55807408 | 61,57033181 | 59,50346104 | 12 |
| ENSGALG00000008369 | ABCC5 | 21,72606872 | 30,01042058 | 54,12213459 | 53,90325716 | 12 |
| ENSGALG00000008403 | PHF21A | 43,18235274 | 47,88824436 | 51,58618832 | 51,77574719 | 12 |
| ENSGALG00000008676 | RRBP1 | 23,1998561 | 23,99502204 | 31,01580542 | 29,92130683 | 12 |
| ENSGALG00000008856 | | 1,031931903 | 1,608558804 | 2,634592306 | 3,16631245 | 12 |
| ENSGALG00000008883 | TCF7L2 | 20,496599 | 30,99622973 | 37,62172759 | 37,77014196 | 12 |
| ENSGALG00000009034 | | 0,801126927 | 0,932946754 | 1,753738946 | 1,779517801 | 12 |
| ENSGALG00000009086 | EHD3 | 146,0272989 | 157,6078485 | 205,2381547 | 195,3390011 | 12 |
| ENSGALG00000009396 | | 13,79079951 | 19,66608855 | 26,51338215 | 25,68230408 | 12 |
| ENSGALG00000010030 | | 41,45059966 | 41,25182098 | 57,13689599 | 55,53405584 | 12 |
| ENSGALG00000010166 | VLDLR | 4,067429632 | 6,335447679 | 9,796757208 | 10,20517655 | 12 |
| ENSGALG00000010291 | RBPMS | 40,68094996 | 40,8529908 | 75,23070837 | 65,91476169 | 12 |
| ENSGALG00000010385 | TJAP1 | 28,82460773 | 27,21958624 | 36,81273204 | 35,93680702 | 12 |
| ENSGALG00000010778 | CHRM3 | 2,273839966 | 2,32517469 | 6,105235943 | 5,391486719 | 12 |
| ENSGALG00000011025 | | 11,49510331 | 13,44047249 | 19,03749752 | 17,51789112 | 12 |
| ENSGALG00000011137 | | 0,621854178 | 0,557885914 | 2,441826917 | 1,948944133 | 12 |
| ENSGALG00000011266 | PPM1K | 3,953648629 | 4,152677018 | 6,846909686 | 7,380787396 | 12 |
| ENSGALG00000011608 | INF2 | 1,633620915 | 6,49652447 | 13,65966714 | 14,33444624 | 12 |
| ENSGALG00000011944 | | 29,7277375 | 25,20155428 | 47,10159373 | 48,8566892 | 12 |
| ENSGALG00000012277 | | 35,33038415 | 33,48644634 | 63,08082454 | 82,91803991 | 12 |
| ENSGALG00000012380 | SPOPL | 26,64799127 | 26,90568857 | 37,32642684 | 35,31642033 | 12 |
| ENSGALG00000012538 | NR4A2 | 0,505244825 | 0,625081889 | 1,1713555 | 1,07035435 | 12 |
| ENSGALG00000012885 | BCL2 | 12,34404469 | 14,24953187 | 16,80990968 | 17,44414904 | 12 |
| ENSGALG00000013021 | | 30,98315435 | 33,14000451 | 66,60766145 | 57,42790105 | 12 |
| ENSGALG00000013342 | GBX2 | 104,3637185 | 76,23496549 | 163,838919 | 164,7512996 | 12 |
| ENSGALG00000013628 | C6orf62 | 110,0447131 | 111,9104576 | 145,7924007 | 158,3265634 | 12 |
| ENSGALG00000013757 | TMX3 | 82,04092518 | 84,73616801 | 98,56482332 | 96,04531272 | 12 |
| ENSGALG00000014084 | FGFR1OP2 | 38,22441741 | 44,06957863 | 58,20825246 | 56,45097665 | 12 |
| ENSGALG00000014349 | | 8,560237913 | 9,973650087 | 16,79967362 | 17,03527571 | 12 |
| ENSGALG00000014355 | STIM2 | 27,91184823 | 26,64561514 | 35,21479281 | 40,31897857 | 12 |
| ENSGALG00000014516 | CPEB2 | 3,813359717 | 7,524646883 | 11,31353119 | 13,35774029 | 12 |
| ENSGALG00000014598 | | 121,596947 | 119,4705335 | 150,3934093 | 156,4723329 | 12 |
| ENSGALG00000014687 | EPHA1 | 0,875109773 | 0,887791079 | 2,439656862 | 2,902792716 | 12 |
| ENSGALG00000014719 | | 53,36926679 | 56,13568368 | 75,95786608 | 70,89585202 | 12 |
| ENSGALG00000014815 | | 7,514303562 | 7,928521182 | 16,27836163 | 14,27131615 | 12 |
| ENSGALG00000014854 | | 235,2101919 | 242,2048178 | 262,4097502 | 261,6360768 | 12 |
| ENSGALG00000015025 | VANGL1 | 14,75235588 | 14,95356303 | 18,5151432 | 19,32445822 | 12 |
| ENSGALG00000015036 | RIC1 | 24,6564351 | 26,54672688 | 34,77569889 | 32,31957583 | 12 |
| ENSGALG00000015943 | | 22,32640939 | 23,59775922 | 28,84956292 | 31,67339581 | 12 |
| ENSGALG00000016086 | EGR4 | 0,312754609 | 0,349739454 | 3,626256956 | 2,868130472 | 12 |
| ENSGALG00000016105 | DYSF | 11,86244012 | 10,67649701 | 17,62292607 | 18,03089 | 12 |
| ENSGALG00000016283 | BEND6 | 8,65472961 | 10,21102149 | 12,41413372 | 13,84260597 | 12 |
| ENSGALG00000016348 | SAT1 | 65,846725 | 146,5356664 | 203,5314021 | 222,2269098 | 12 |
| ENSGALG00000016457 | TRIB2 | 147,3950287 | 132,4539352 | 194,1354664 | 187,1424656 | 12 |
| ENSGALG00000016500 | FKBP12.6 | 12,53712907 | 24,62077173 | 47,98704792 | 47,94808325 | 12 |
| ENSGALG00000016508 | | 27,46728733 | 30,92244173 | 37,55597573 | 37,58110828 | 12 |
| ENSGALG00000017117 | | 0,247769645 | 0,503287821 | 1,181898236 | 1,186818356 | 12 |
| ENSGALG00000020719 | | 0,7762351 | 0,748726958 | 3,053419651 | 2,604518111 | 12 |
| ENSGALG00000021041 | | 48,79130614 | 47,81392988 | 60,79348483 | 57,94430863 | 12 |
| ENSGALG00000023347 | PTCHD3 | 1,787758741 | 2,844053111 | 5,289343591 | 6,207946544 | 12 |
| ENSGALG00000023468 | VPS9D1 | 20,7533336 | 23,64061043 | 26,5205822 | 26,21200221 | 12 |
| ENSGALG00000023648 | PPP3R1 | 108,4834821 | 119,5709887 | 131,3437155 | 137,8492022 | 12 |
| ENSGALG00000025887 | ATF3 | 6,584528653 | 6,144744457 | 11,98779346 | 16,71462268 | 12 |
| ENSGALG00000025911 | | 7,173430124 | 8,328187345 | 12,41146067 | 14,53810568 | 12 |
| ENSGALG00000026077 | ALDOC | 1287,720681 | 1303,453672 | 1587,752058 | 1585,907858 | 12 |
| ENSGALG00000026154 | CARD19 | 2,519303597 | 5,718212359 | 10,62772597 | 11,77837126 | 12 |
| ENSGALG00000026231 | ARID3A | 55,59951305 | 68,67400855 | 79,54534838 | 84,31937056 | 12 |
| ENSGALG00000026237 | | 197,6339803 | 214,683724 | 292,3753153 | 315,5343838 | 12 |
| ENSGALG00000026263 | RGS8 | 0,38280482 | 0,973247805 | 10,59393568 | 8,731063711 | 12 |
| ENSGALG00000026355 | | 2,797729238 | 3,795263667 | 5,576978031 | 6,019303511 | 12 |
| ENSGALG00000026534 | | 0,434765116 | 0,435780888 | 1,090824905 | 1,486910738 | 12 |
| ENSGALG00000026850 | | 19,3448547 | 23,33560488 | 26,08136183 | 27,49169005 | 12 |

| | | | | | | |
|---|---|---|---|---|---|---|
| ENSGALG00000026989 | HOXA7 | 1,842634302 | 1,750162784 | 3,503069673 | 3,914427 | 12 |
| ENSGALG00000027208 | | 33,6710471 | 32,97550046 | 43,12346055 | 42,66001317 | 12 |
| ENSGALG00000027331 | | 1,065988859 | 1,550790838 | 11,17131653 | 17,68458576 | 12 |
| ENSGALG00000027352 | | 283,8386064 | 243,9179572 | 481,0308996 | 471,5712627 | 12 |
| ENSGALG00000027365 | HIVEP3 | 1,475692748 | 1,810205476 | 5,644935787 | 4,967885552 | 12 |
| ENSGALG00000027554 | | 0,179438786 | 0,053342062 | 1,868827591 | 2,239675166 | 12 |
| ENSGALG00000027684 | | 17,74702399 | 21,59636073 | 35,22185222 | 45,64200281 | 12 |
| ENSGALG00000027761 | TNFAIP8 | 5,255419926 | 6,047134406 | 7,671477467 | 7,823425868 | 12 |
| ENSGALG00000027805 | | 3,22591528 | 3,316534643 | 8,789132761 | 9,027754662 | 12 |
| ENSGALG00000027925 | | 2,246110271 | 1,656033894 | 2,090585784 | 2,40260606 | 12 |
| ENSGALG00000028095 | | 116,6593576 | 116,1033043 | 153,7263599 | 150,47256 | 12 |
| ENSGALG00000028128 | | 48,54899189 | 31,62799235 | 69,21356455 | 93,53834461 | 12 |
| ENSGALG00000028203 | SPRED1 | 21,61093344 | 21,70393773 | 28,89856099 | 28,51271586 | 12 |
| ENSGALG00000028726 | | 128,7914409 | 85,78500371 | 167,5124021 | 189,4637389 | 12 |
| ENSGALG00000028749 | | 2923,755457 | 3158,474332 | 3440,767081 | 3520,806683 | 12 |
| ENSGALG00000028759 | LYPD6B | 3,251758824 | 4,924290781 | 7,287330896 | 9,009994746 | 12 |
| ENSGALG00000028785 | | 0,606392138 | 1,210251308 | 2,188601903 | 2,472445717 | 12 |
| ENSGALG00000029090 | | 169,0725833 | 156,4931705 | 386,9724027 | 460,4292989 | 12 |
